# Dynamic cell fate plasticity and tissue integration drive functional synovial joint regeneration

**DOI:** 10.1101/2024.12.12.628180

**Authors:** Maria Blumenkrantz, Felicia Woron, Ernesto Gagarin, Everett Weinstein, Maryam H. Kamel, Leonardo Campos, Agnieszka Geras, Troy Anderson, Julia Mo, Desmarie Sherwood, Maya Gwin, Bianca Dumitrascu, Nadeen O. Chahine, Joanna Smeeton

## Abstract

Adult mammalian synovial joints have limited regenerative capacity, where injuries heal with mechanically inferior fibrotic tissues. Here we developed a unilateral whole-joint resection model in adult zebrafish to advance our understanding of how to stimulate regrowth of native synovial joint tissues. Using a combination of microCT, histological, live imaging, and single-cell RNA sequencing (scRNAseq) approaches after complete removal of all joint tissues, we find de novo regeneration of articular cartilage, ligament, and synovium into a functional joint. Clonal lineage tracing and scRNAseq implicate a multipotent, neural crest-derived population in the adult skeleton as a cell source for these regenerating tissues. Together, our findings reveal latent molecular and cellular programs within the adult skeleton that are deployed to regenerate a complex joint with lubricated articular cartilage.

## Introduction

Synovial joints are complex organs consisting of articular cartilage that minimizes joint friction, synovium that secretes lubricating fluid, and ligaments that stabilize the articulating joint. Injury or lifelong wear of joint tissues can lead to osteoarthritis (OA), a degenerative joint disease that is a leading cause of disability (*1*). Damaged articular cartilage and ligaments exhibit poor regenerative capacity in mammals (*2*), and regeneration of functional hyaline cartilage with lubricating properties has remained an elusive target. Current disease-modifying strategies, such as surgical joint replacement and autologous connective tissue grafting, have limitations such as prosthetic joint loosening, high donor site morbidity, graft failure, and repair with fibrous tissue that has inferior biomechanical properties (*3–5*). There is an urgent clinical need to develop more effective treatments for joint injury; however, the fundamental mechanisms driving ligament and articular cartilage cell fate specification and integration into functional tissues are still largely unknown.

Success in activating regenerative programs in mammals has been largely limited to neonates (*6, 7*), and even the highly regenerative axolotl fails to repair a critical size joint defect (*8*). However, the highly regenerative zebrafish possesses unique regenerative properties distinct from the mammalian fibrotic response, and the study of these differences has led to new tools for improving regenerative outcomes in mammalian cardiac injury models (*9*). Zebrafish have over 70% genetic conservation with humans, including important similarities in the composition of their synovial joints, such as the presence of synovium and multilayered articular cartilage that secrete lubricating proteins (*10, 11*). While adult zebrafish retain high capacity to regenerate individual skeletal tissues (*12–15*), currently, there are no published adult models for the regeneration of a complex synovial joint with lubricating articular cartilage. Existing surgical injury models for articular cartilage regeneration in adult zebrafish only indirectly damage the cartilage, a major limitation as less than half of injured fish undergo subsequent joint degeneration (*13*). Thus, we devised a novel whole-joint injury model to probe the zebrafish’s ability to rebuild multiple skeletal cell and tissue types de novo and reintegrate them into a functional joint after catastrophic injury.

Here we establish adult zebrafish whole-joint resection injury as a new model for understanding the coordinated regeneration of an integrated synovial joint organ. This surgery completely removes all pre-existing joint tissues, including articular cartilage, synovium, all intra- and extra-synovial ligaments, and some subchondral bone. Through histological analysis and 3D fluorescence imaging, we find that proliferation of endogenous mesenchymal cells leads to functional synovial joint regeneration by 2 months post-injury. Strikingly, we observe that regenerated articular cartilage expresses *prg4b*, the hallmark of superficial hyaline articular cartilage (*11, 16*). From single-cell RNA sequencing (scRNAseq) analysis of the first 70 days after resection injury, we describe the timeline and transcriptional signatures of four phases of joint regeneration. Through pseudotime analysis and lineage tracing, we identify a population of multipotent neural crest-derived progenitors as a cell source for regenerated joint tissues. Taken together, our findings establish adult zebrafish whole-joint resection as a new model for de novo regeneration of a functional synovial joint with lubricating articular cartilage.

## Results

### Adult joint resection injury and functional regeneration

To test whether zebrafish have the capacity to regenerate multiple synovial joint cell types after removal of pre-existing mature joint tissues, we devised a new joint resection injury to the jaw joint. The resection surgery consists of three cuts to the anguloarticular, interopercular, and quadrate bones (Fig. 1A). Gross imaging immediately after surgery shows complete removal of the jaw joint, including the articular cartilage, synovial membrane, and interopercular-mandibular ligament, from one side of the zebrafish’s jaw (Fig. 1B). Morphometric analysis at 0 days post-joint resection (dpjr) shows an initial surgical resection length of 1mm generated by three independent surgeons, demonstrating the consistency of the surgery (Fig. 1C). Repeated gross measurements of individual fish until 56 dpjr showed no change in weight of resected vs uninjured (n=21) (Fig. S1A-B). μCT visualization of the craniofacial skeleton confirmed that joint resection fully removes the articulation present in the uninjured jaw (Fig. 1D). By 28 dpjr, we observed bone growth in the form of protrusions from the cut sites (green arrowhead) and islands of bone (green asterisk). At 1 year post-joint resection (ypjr), the gap was largely bridged by new bone, and a structure resembling an articulating joint surface was present (green arrow). Histological analysis showed the regeneration of a joint structure capped with tissue resembling multilayered articular cartilage, in both morphology and location, by 56 dpjr (Fig. 1E). Both the uninjured and regenerated joints contained cavities lined with articular chondrocytes, including cells with the rounded morphology of deeper articular chondrocytes (black arrowhead) and cells with the flatter morphology of superficial articular chondrocytes (black arrow).

**Fig. 1:**
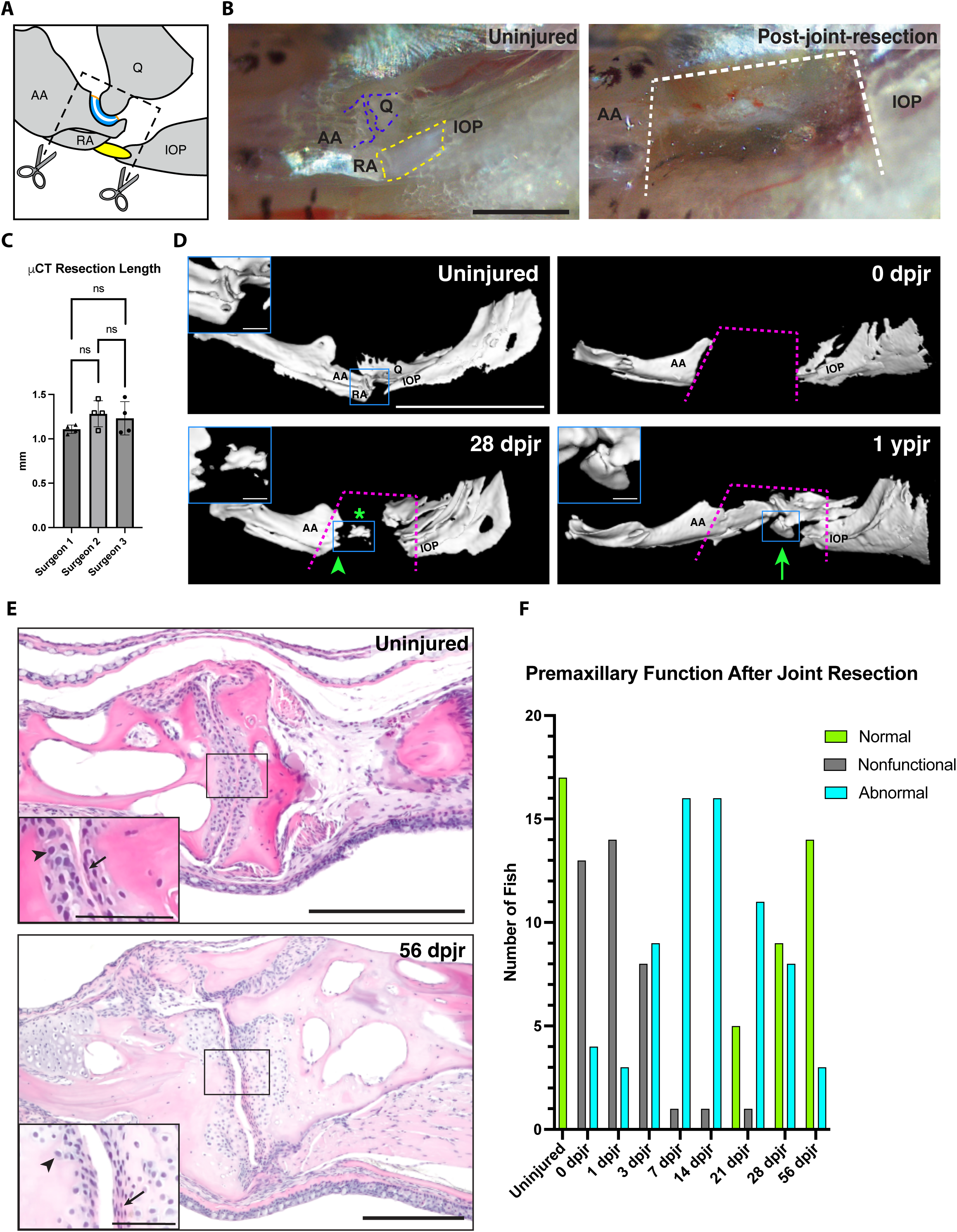
Zebrafish whole-joint resection injury and regeneration. **(A)** Schematic of the articulating bones at the zebrafish jaw joint that are cut to surgically remove all jaw joint tissues including the articular cartilage (blue) and the interopercular-mandibular (IOM) ligament (yellow). **(B)** Reflected light images of the zebrafish jaw in ventral view show the zebrafish jaw joint (blue dashed line) and IOM ligament (yellow dashed line) completely removed after joint resection (cut sites indicated by white dashed line). **(C)** Quantification of length of bone removed by 3 independent surgeons in millimeters (n=4 per surgeon), error bar represents SEM. **(D)** Micro-computed tomography (µCT)-based skeletal reconstructions of the lateral view of the uninjured zebrafish jaw joint as well as the bones in the injury site at 0 dpjr, 28 dpjr, and 1 ypjr. Blue insets show the articulating bones of the uninjured jaw joint, bone growth at 28 dpjr (green arrow and asterisk), and a regenerated joint-like structure (green arrow) at 1 ypjr. Magenta dashed lines indicate cut sites. **(E)** H&E staining of an uninjured joint and a regenerated joint at 56 dpjr. Insets show deep (black arrowhead) and superficial (black arrow) articular chondrocyte layers. **(F)** Quantification of premaxillary function over the time course of joint regeneration. Premaxillary function in each fish was scored as normal, nonfunctional, or abnormal. AA, anguloarticular bone; IOP, interopercular bone; Q, quadrate bone; RA, retroarticular bone. Scale bars = 200µm (B), 2.5mm (D), 200µm (D insets), 100µm (E), 25µm (E insets).

Next, we sought to determine whether joint function was restored after resection. To do so, we quantified footage of jaw activity during swimming and feeding. Slow-motion analysis of jaw movements in individual live fish did not show any major disruption in lower jaw rotational movement following injury (n=21), likely due to the intact jaw joint on the contralateral side driving lower jaw movement. However, qualitative scoring of jaw movement recordings showed that immediately post-surgery and out to 21 dpjr, premaxillary protrusion was disrupted (abnormal or nonfunctional) (n=17/17) (Fig. S1C). We observed restored craniofacial coordination by 56 dpjr in the form of normal premaxillary protrusion (n=14/17) (Fig. 1F). Additionally, premaxilla retraction was disrupted following injury, resulting in larger intervals of time between consecutive suction feeding motions (n=6; p=0.0056), and was restored by 56 dpjr (Fig. S1D). Overall, these observations point to the regeneration of a functional joint-like structure covered by multilayered articular cartilage following complete resection.

### Epithelial and mesenchymal cells proliferate in response to injury and bridge the resection site

After observing the regeneration of an articulating surface, we assessed the early injury response to whole-joint resection. Histological analysis showed that at 0 dpjr, the resected bone stubs were exposed (Fig. 2A). By 1 dpjr, the resection site was enclosed by epithelia directly covering the cut bones. At 3 dpjr, bands of mesenchymal cells were present under the cap of epithelial tissue at the anterior, posterior, and dorsal cut sites. By 7 dpjr, epithelial and mesenchymal tissues formed a bridge spanning the regenerating zone. Morphological hallmarks of cellular lineage commitment were visible by 14 dpjr, with chondrocytes and islands of bone arising within the bridge of mesenchymal tissue (Fig. 2A’).

**Fig. 2:**
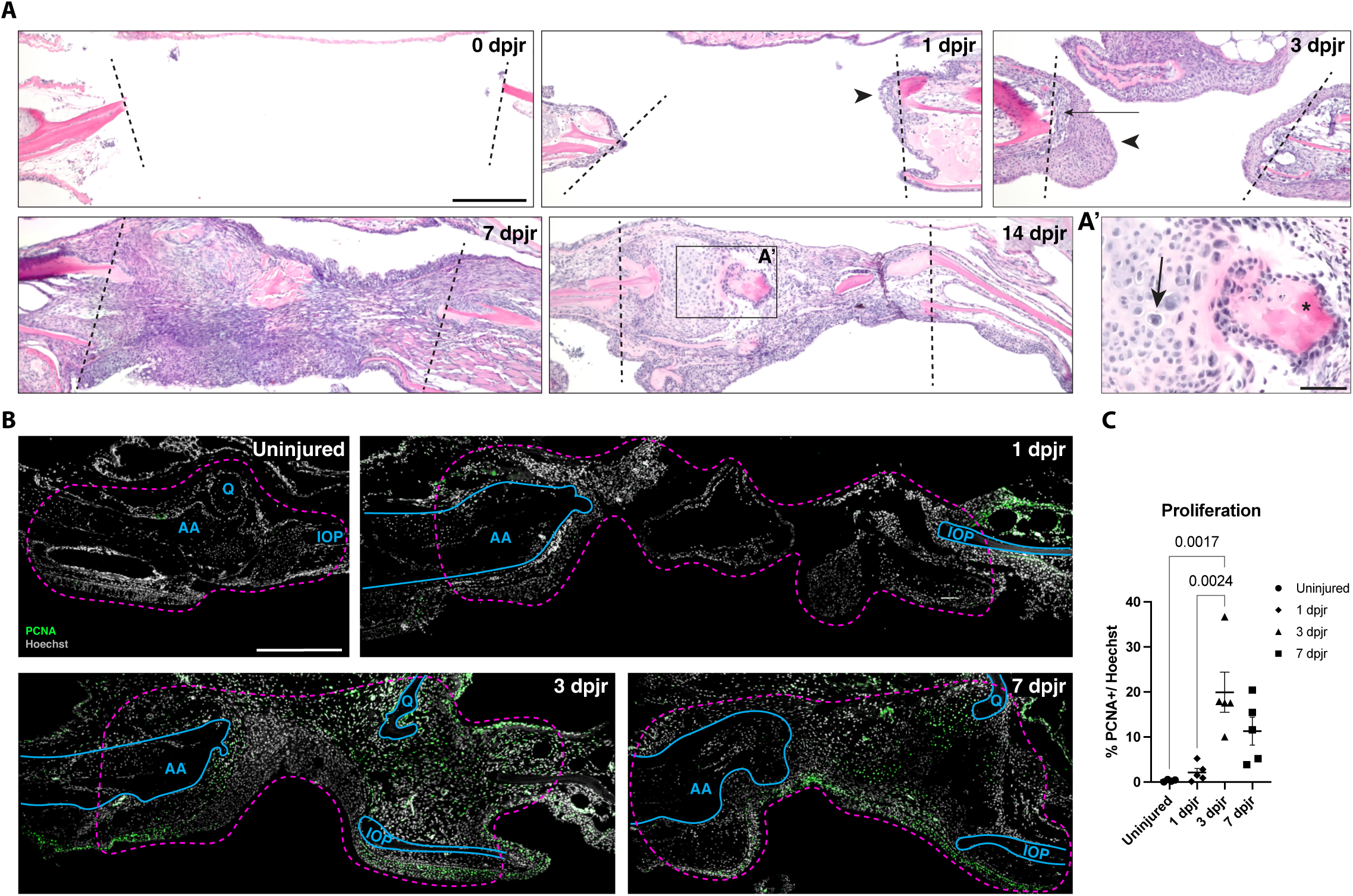
Early epithelial and mesenchymal cell response to joint resection. **(A)** H&E staining of early injury response in 0 dpjr (n=5), 1 dpjr (n=4), 3 dpjr (n=4), 7 dpjr (n=6), and 14 dpjr (n=8). Black arrowhead denotes epithelium and black arrow denotes mesenchyme. **(A’)** Inset shows regenerating chondrocytes (arrow) and bone (asterisk) at 14 dpjr. Black dashed lines indicate cut sites. **(B)** Representative images of PCNA immunofluorescence in uninjured joint tissue and at 1 dpjr, 3 dpjr, and 7 dpjr. Region of interest for quantification is outlined in magenta dashed lines and resected bone stubs are outlined in solid blue lines. **(C)** Quantification of percentage of PCNA-positive nuclei showing significant increase at 3 dpjr (n=4-5 per time point); error bar represents SEM. AA, anguloarticular bone; IOP, interopercular bone; Q, quadrate bone; RA, retroarticular bone. Scale bars = 100µm (A), 25µm (A’), 300µm (B).

To characterize the proliferative response of the injured and regenerating tissues, we performed immunofluorescence staining for Proliferating Cell Nuclear Antigen (PCNA). While we observed low levels of proliferation prior to injury or at 1 dpjr, there was a significant increase in PCNA+ cells by 3 dpjr (Fig. 2B & 2C). Proliferating cells were found surrounding the resected bone stubs (up to 650 µm from each resection site) and throughout the mesenchyme. By 7 dpjr, the proliferative response significantly decreased and was largely limited to epithelial cells and chondrocytes. Proliferating chondrocytes (PCNA+, Sox9a+) were observed surrounding the resected bone stubs and in chondrocyte islands within the bridging mesenchyme domain (Fig. S2A). Compared to articular chondrocytes in the uninjured joint, proliferation was significantly increased in regenerating chondrocytes at 7 dpjr (Fig. S2B). Proliferation in the skin at 3 dpjr was mainly localized to injury-adjacent epithelium (IAE) surrounding the resected bone stubs, with few proliferating cells within the injury epithelium (IE) bridging the wound site (Fig. S2C). By 7 dpjr, the IE had significantly higher PCNA+ cells compared to the 3 dpjr IE (Fig. S2D), indicating that the cell cycle dynamics of the IE are distinct from that of the IAE. Overall, these data reveal that the identity of proliferating cell populations in the regenerating joint undergoes major shifts over time.

### Mature joint tissue types regenerate following resection

To characterize the timing and pattern of bone regrowth, we conducted repeated live imaging of the resection site using early osteoblast reporter line *sp7:eGFP* (Fig. 3A). Immediately after joint resection, there was a gap in GFP fluorescence consistent with the area of the resected bone. By 14 dpjr, we saw the re-emergence of GFP fluorescence in the form of both bony protrusions from the resected stubs and bony islands within the injury mesenchyme. We then sought to determine whether mature joint cell types regenerate via assessment of gene expression as well as cell and tissue morphology. Imaging uninjured jaw joints of *thbs4a_p1:eGFP;scxa:mCherry* transgenic fish with calcein-stained bones showed that the articular chondrocytes lining the joint were *thbs4a_p1:eGFP*+;*scxa:mCherry*-and show a rounded cell morphology (Fig. 3B), whereas the IOM (interopercular-mandibular) ligamentocytes were *thbs4a_p1:eGFP*+;*scxa:mCherry*+ with an elongated cell morphology (Fig. 3C). Whole-joint resection completely removed the articular cartilage and IOM ligament (Fig. 3D). At 56 dpjr, regenerated articular chondrocytes were *thbs4a_p1:eGFP*+;*scxa:mCherry*- and lined a cavity between two regenerated bones (Fig. 3E). Regenerated ligamentocytes were *thbs4a_p1:eGFP*+;*scxa:mCherry*+, had an elongated morphology, and bridged two regenerated bones (Fig. 3F). Live imaging of *scxa:mCherry;sp7:eGFP* double-transgenics confirmed new ligament tissue bundles connecting newly formed joint-associated jaw bones by 57 dpjr (Fig. 3G). These data show that after removal of pre-existing mature joint tissues, zebrafish can re-establish ligamentocyte and articular chondrocyte cell fate.

**Fig. 3:**
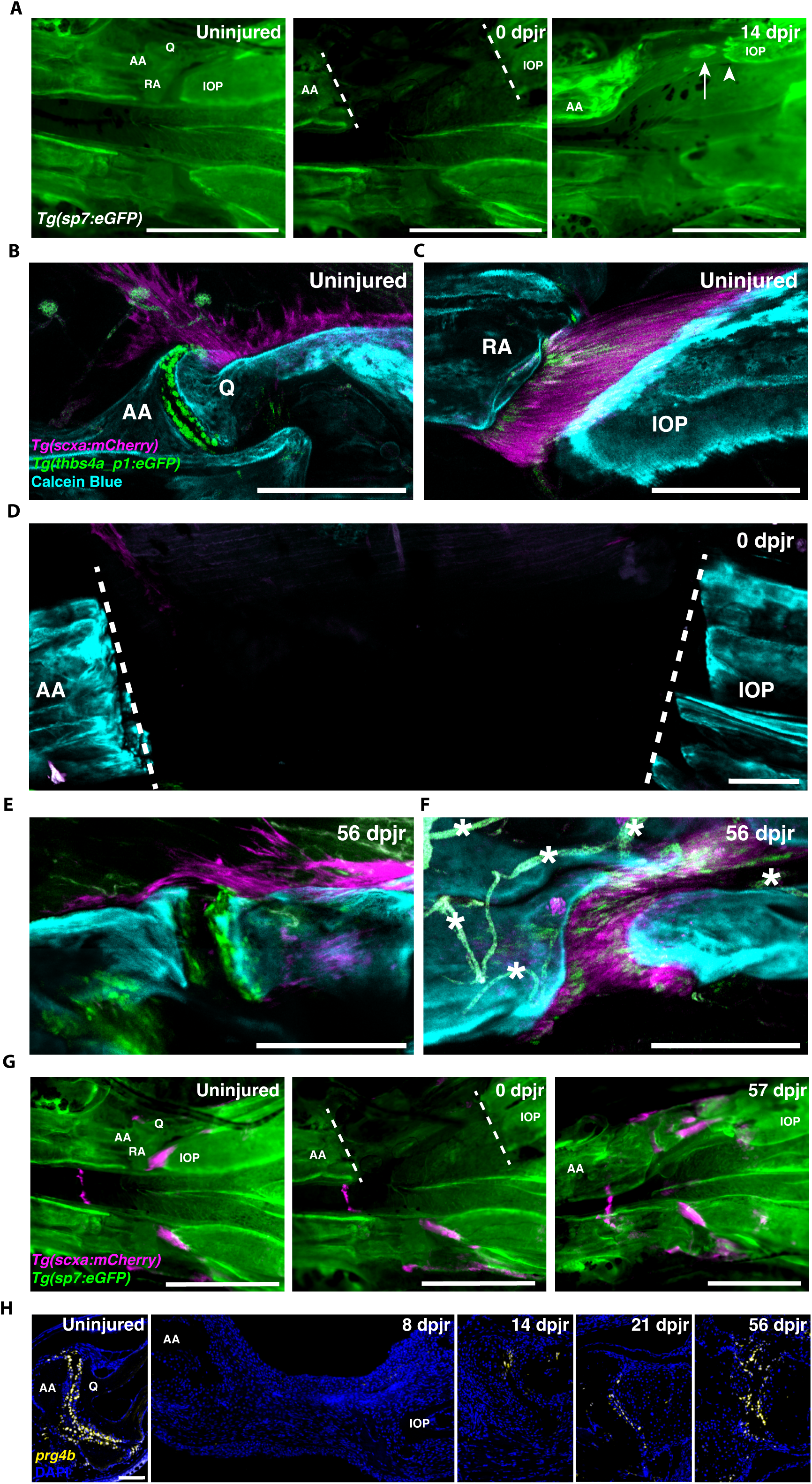
Regeneration of all mature joint tissues after joint resection. **(A)** Repeated live imaging of the ventral view of *sp7:eGFP* expression before injury (n=28), at 0 dpjr (n=25), and at 14 dpjr (n=19). Bony protrusions denoted by white arrowhead and bony islands indicated by white arrow. **(B** to **F)** Confocal microscopy imaging of tissue-cleared *thbs4a_p1:eGFP* and *scxa:mCherry* expression in the uninjured joint (B) and uninjured ligament (C) (n=4), at 0 dpjr (n=3) (D), and in a regenerated joint (E) and regenerated ligament (F) at 56 dpjr (n=7). Bone visualized with calcein blue staining. White asterisks mark autofluorescent blood vessels. **(G)** Repeated live imaging of the ventral view of *scxa:mCherry* and *sp7:eGFP* expression before injury (n=28), at 0 dpjr (n=25), and at 57 dpjr (n=8). **(H)** *prg4b* (yellow) smFISH in the uninjured jaw joint and at 8 dpjr, 14 dpjr, 21 dpjr, and 56 dpjr (n=3 per time point). White dashed lines indicate cut sites. AA, anguloarticular bone; IOP, interopercular bone; Q, quadrate bone; RA, retroarticular bone. Scale bars = 1mm (A and G), 200µm (B-F), 100µm (H).

To determine whether regenerated joints are true lubricated synovial joints, we performed single-molecule fluorescent *in situ* hybridization (smFISH) for *prg4b,* which encodes for a lubricating glycoprotein secreted by articular chondrocytes and synoviocytes. Uninjured animals had robust *prg4b* expression in cells lining the joint cavity. At 8 dpjr, we observed minimal *prg4b* expression within the mesenchymal bridge. At 14 dpjr, *prg4b* was re-expressed by cells at the edges of a regenerating joint cavity. Interestingly, *prg4b* was robustly expressed along both surfaces of a new articulating joint structure by 21 dpjr and maintained at 56 dpjr (Fig. 3H). These data show that after complete removal of the joint, regenerated articular chondrocytes and synoviocytes re-establish the capacity to secrete the lubricating proteins necessary for healthy joint function and movement.

### The regenerative mesenchyme undergoes dynamic ECM changes and reactivation of skeletal development programs

To define the cellular and transcriptional mechanisms driving whole-joint regeneration, we performed scRNAseq on live-sorted cells from uninjured joints and regenerated joint tissues at 6 timepoints post-resection (1, 3, 7, 14, 28, and 70 dpjr) (Fig. 4A). Unbiased clustering at a low resolution yielded 10 broad superclusters of cells (Fig. 4B). Skeletal cells were marked by either the mesenchymal marker *prrx1b* (Skeletal 1) or osteoblast marker *bglap* (Skeletal 2) (Fig. S3A). The Immune 1 supercluster included neutrophils (*mpx+*) and macrophages (*mpeg1.1+*), while the Immune 2 supercluster primarily included lymphocytes, such as B cells and T cells (*ikzf1+*). Elevated numbers of neutrophils and macrophages at 1 dpjr indicated an influx of immune cells into the injury site as part of the wound healing response (Fig. S4). Other superclusters included epithelial cells (*epcam+*), endothelial cells (*pecam1+*), perivascular cells (*myh11a+*), muscle cells (*myod1+*), neural cells (*mpz+*), and odontoblasts (*scpp1+*).

**Fig. 4:**
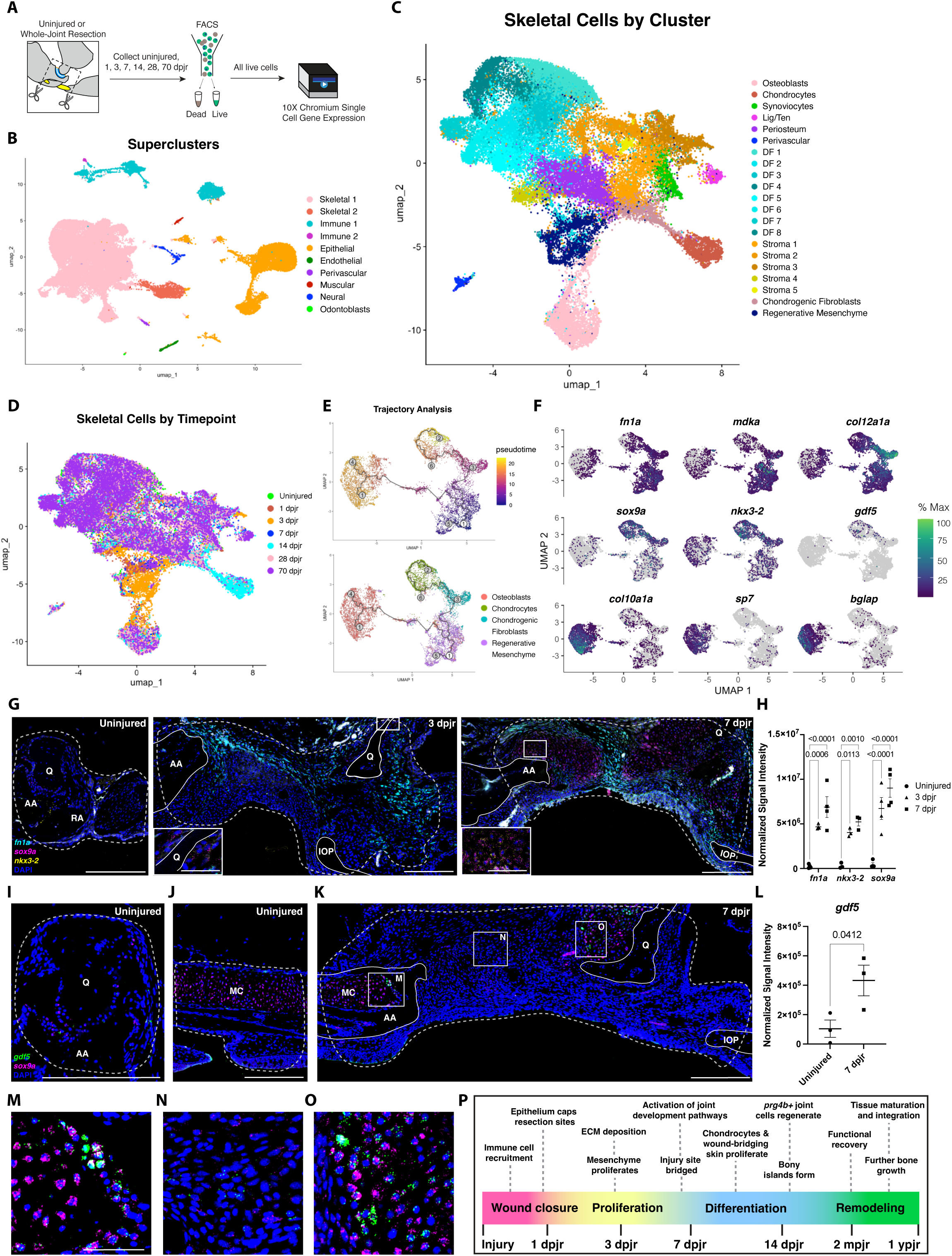
Transcriptional dynamics of skeletal cells throughout regeneration. (**A**) Schematic detailing the generation of single-cell RNA sequencing (scRNAseq) dataset. Jaw joint tissue was collected from uninjured joints and regenerated joints at 1 dpjr, 3 dpjr, 7 dpjr, 14 dpjr, 28 dpjr, and 70 dpjr (n=2-3 per time point). Samples were dissociated and sorted for all live cells prior to scRNAseq library generation. (**B**) UMAP of low-resolution clusters (r=0.02) for integrated samples. Superclusters include 2 skeletal clusters, 2 immune clusters, and individual epithelial, endothelial, perivascular, muscular, neural, and odontoblast clusters. (**C** and **D**) UMAP of skeletal cells clustered at a high resolution (r=0.95) and colored by cluster (C) or timepoint (D). (**E**) Single-cell trajectory analysis of the osteoblasts, chondrocytes, chondrogenic fibroblasts, and regenerative mesenchyme clusters using Monocle3 colored by pseudotime (top) or cluster (bottom). (**F**) Differential gene expression analysis plotted along the single-cell trajectory over time. (**G**) HCR smFISH for *fn1a* (cyan), *sox9a* (magenta), and *nkx3-2* (yellow) in uninjured, at 3 dpjr, and at 7 dpjr. 3 dpjr inset displays *sox9a* expression along the edges of the resected quadrate bone, and 7 dpjr inset shows *sox9a* and *nkx3-2* co-expression in regenerating chondrocyte islands. (**H**) Quantification of smFISH *fn1a*, *sox9a*, and *nkx3-2* transcript expression from uninjured, 3 dpjr, and 7 dpjr (n=3-4 per time point). (**I** to **O**) RNAscope smFISH for *gdf5* (green) and *sox9a* (magenta) transcripts in the uninjured joint (I), uninjured Meckel’s cartilage (J) and at 7 dpjr (K) (n=3 per time point). Quantification of *gdf5* transcript expression from uninjured and 7 dpjr (L). Insets illustrate increased *gdf5* transcript expression in the perichondrium surrounding Meckel’s cartilage (M), the regenerative mesenchyme (N), and in regenerating chondrocyte islands (O) at 7 dpjr. (**P**) Schematic of the major events along the whole joint regeneration time course. Regions of interest for quantification are outlined in light gray dashed lines; error bars represent SEM. Resected bone stubs are outlined in solid white lines. AA, anguloarticular bone; IOP, interopercular bone; Q, quadrate bone; RA, retroarticular bone; MC, Meckel’s cartilage. Scale bars = 200µm (G, I-K), 50µm (G insets and M-O).

To elucidate the transcriptional dynamics of regenerating skeletal cells, we subsetted the Skeletal 1, Skeletal 2, and Perivascular clusters and re-clustered at a higher resolution to yield 21 clusters (Fig. 4C, S3B-C, S5A-B, Table S1). Within the skeletal population, we captured the major cell types present in uninjured joints: osteoblasts (*bglap+*), chondrocytes (*col2a1a*+), synoviocytes (*prg4b+, col2a1a-*), and ligamentocytes/tenocytes (*scxa+, thbs4a+, tnmd+*) (Table S2). In addition to the perivascular (*myh11a+*) cluster, we also identified a periosteal (*mmel1*+) cluster. Clustering at this resolution revealed 8 clusters of dermal fibroblasts (DF; *pcdb1+, pah+, qdpra+*) and 5 clusters of stromal cells (*col5a3a+*, *coch+*, and/or *cxcl12a+*).

Additionally, we identified a cluster specific to early regeneration, comprised mainly of cells from 1 dpjr and 3 dpjr (82% of cells in this cluster) (Fig. 4D, S5C). This cluster is defined by expression of proliferation (*pcna+*, *mki67+*) and injury-responsive mesenchyme (*fn1a*+, *mdka*+) markers. We also noted a population of chondrogenic fibroblasts (71% from 14 dpjr and 28 dpjr) that co-expressed markers of fibroblasts (*col1a1a+, col15a1+*) and joint development programs (*sox9a+, nkx3-2+, trps1+*). We then used Monocle3 to measure transcriptional changes in “pseudotime” in the early regenerative fibroblasts, chondrogenic fibroblasts, chondrocytes, and osteoblasts. Monocle3 identified regenerative mesenchyme as the earliest developmental stage and showed that the regenerative mesenchyme cells diverged into two differentiation trajectories, terminating in the osteoblast and chondrocyte clusters (Fig. 4E). Visualization of the trajectories showed the chondrogenic fibroblasts at an intermediate differentiation state between the regenerative mesenchyme and chondrocytes. Regenerative mesenchyme genes (*fn1a*, *mdka*, *col12a1a*) were highly expressed both at early and intermediate phases along the differentiation trajectories, while joint development genes (*sox9a*, *nkx3-2*, *gdf5*) and osteoblast markers (*col10a1a*, *sp7*, *bglap*) were most highly expressed in the chondrocyte and osteoblast differentiation trajectories, respectively (Fig. 4F, Fig. S3C).

We then used Monocle3 to group genes with similar expression patterns along the trajectory into 70 modules (Fig. S6). Modules specific to the osteoblast cluster included Module 21 (enriched in pre-osteoblast genes *runx2a* and *spp1*), Module 26, (enriched in osteoblast differentiation gene *sp7*), and Module 51 (enriched in mature osteoblast genes *bglap* and *ifitm5*). (Fig. S7). Modules specific to the chondrocyte cluster included Module 8 (enriched in cartilage development genes *sox9a* and *gdf5*) and Module 32 (enriched in mature cartilage genes *matn1* and *col2a1a*). Modules specific to the chondrogenic fibroblast cluster include Module 3 (enriched in joint cell fate markers *trps1* and *clu*) and Module 1 (enriched in fibroblast markers *col12a1a* and *col5a1*). Among the modules specific to the regenerative mesenchyme cluster, we identified 3 modules (19, 29, and 38) associated with cell cycle re-entry and two modules (18 and 20) enriched for pro-regenerative fibroblast markers. Together, these gene module analyses revealed groups of co-regulated genes that were unique to various regenerative cell states and whose expression patterns shifted as mesenchymal cells differentiated into mature skeletal cell lineages.

Given the lineage relationships we inferred between the *fn1a+* early mesenchyme and *sox9a+;nkx3-2+* regenerating chondrocytes, next we assessed the spatiotemporal expression of these markers in vivo. At 3 dpjr, combinatorial smFISH analyses show broad expression of *fn1a* throughout the regenerative mesenchyme, while the majority of *sox9a* and *nkx3-2* transcripts were observed lining the resected bone stubs (Fig. 4G). At 7 dpjr, *fn1a+, sox9a+, nkx3-2+* mesenchymal cells were found within the regenerating zone, while chondrocyte islands were *fn1a-* and *sox9a+;nkx3-2+*. These data confirm the presence of injury-induced regenerative cell states transcriptionally similar to the chondrogenic fibroblasts found in the scRNAseq (Fig. S3C). Expression of *fn1a*, *sox9a*, and *nkx3-2* was significantly increased at 3 and 7 dpjr compared to the uninjured joint (Fig. 4H). While *fn1a* was broadly expressed throughout the regenerative mesenchyme at 3 dpjr, its expression at 7 dpjr was maintained in mesenchymal cells but largely excluded from nascent chondrocyte islands. This transcriptional shift indicates a change in cell populations in the regenerating zone by 7 dpjr as injury-responsive mesenchymal cells begin differentiating into mature skeletal cell types.

Next, we assessed the expression of *gdf5*, a TGF-Beta superfamily member which we found to be associated with the cartilage trajectory in pseudotime analysis. While it is expressed in low levels in the adult joint (Fig. 4I) and in perichondrium around Meckel’s cartilage (Fig. 4J), *gdf5* was significantly upregulated in the regenerating zone at 7 dpjr (Fig. 4K and 4L). Increased expression of *gdf5* was observed in three populations of regenerating cells: in perichondrium around Meckel’s cartilage (Fig. 4M), at low levels in subsets of the regenerative mesenchyme (Fig. 4N), and at high levels in a subpopulation of *sox9a*+ chondrocytes (Fig. 4O). These data suggest that after whole-joint resection, re-establishment of mature joint tissues is preceded by injury-responsive mesenchymal cells undergoing a phase of broad ECM deposition that becomes spatially restricted and coincides with a redeployment of skeletal developmental programs by 7 dpjr. Taken together, our analyses demonstrate that whole-joint regeneration progresses through 4 main stages: wound closure, mesenchymal cell proliferation, cell lineage differentiation, and tissue remodeling (Fig. 4P).

### Regenerating joint structures are derived from multipotent neural crest-lineage progenitors

During development, zebrafish craniofacial skeletal structures are derived from the cranial neural crest. To test whether new joint tissues similarly arise from a neural crest-derived cell source during regeneration, we performed joint resections on *Sox10:Cre;actb2:loxP-BFP-STOP-loxP-DsRed* fish, in which blue fluorescence (BFP) is converted to red fluorescence (DsRed) in all neural crest lineage-derived cells (*17–19*). Prior to injury, the IOM ligament and articular chondrocytes lining the jaw joint were DsRed+ (Fig. 5A). Immediately after joint resection, no DsRed fluorescence was detected in the injury site. At 56 dpjr, DsRed expression was observed in regenerated bone, ligament, and cartilage. These data show that similar to development, joint tissues are derived from the neural crest lineage during regeneration.

**Fig. 5:**
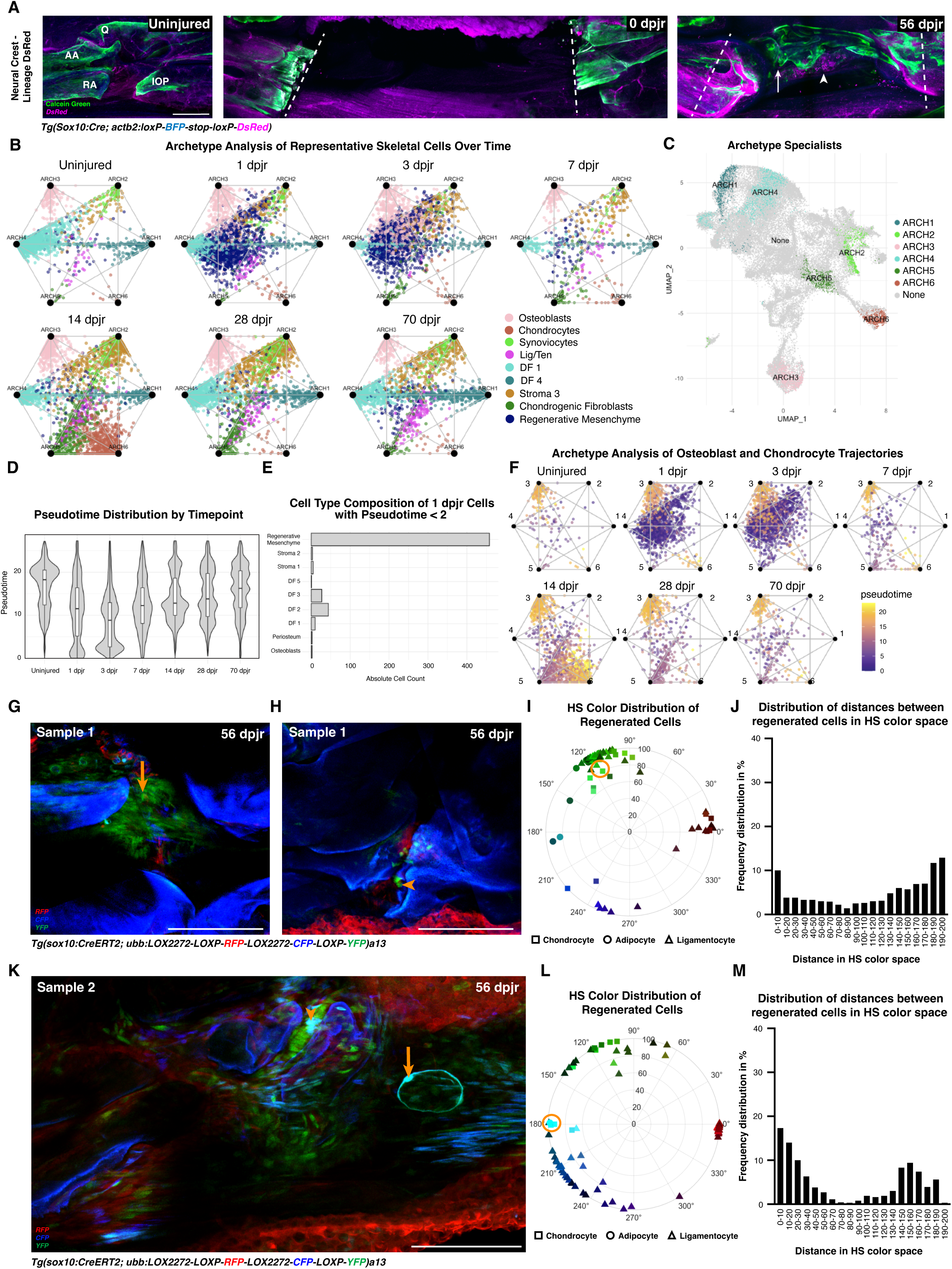
Neural crest-derived multipotent cells contribute to regenerated joint tissues. (**A**) Lineage trace of DsRed-labeled neural crest lineage-derived cells using *Sox10:Cre*;*actb2:loxP-BFP-STOP-loxP-DsRed*; bone visualized with calcein green staining. White arrow denotes regenerated joint surface and white arrowhead indicates regenerating chondrocytes. (**B**) Simplex plots of archetype analysis colored by cluster and split by time point. Clusters include DF 1, Regenerative Mesenchyme, DF 4, Stroma 3, Osteoblasts, Lig/Ten, Chondrogenic Fibroblasts, Chondrocytes, and Synoviocytes. (**C**) UMAP of skeletal cells colored by most similar archetypes. (**D**) Distribution of pseudotime of skeletal cell clusters across timepoints. (**E**) Cell type composition of cells in the 1 dpjr sample that have pseudotime between 0 and 2. Regenerative mesenchyme represents the majority of cells with the earliest pseudotime. (**F**) Simplex plots of archetype analysis with osteoblasts, chondrocytes, chondrogenic fibroblasts, and regenerative mesenchyme colored by pseudotime. (**G**) Single-clone lineage trace of regenerated Zebrabow ligament from Sample 1. Orange arrow denotes green ligamentocyte circled in the corresponding polar graph. (**H**) Single-clone lineage trace of regenerated Zebrabow joint from Sample 1. Orange arrowhead denotes green chondrocyte circled in the corresponding polar graph. (**I**) Polar graph of HS color distribution of regenerated cells from Sample 1. Orange circle indicates a green ligamentocyte and chondrocyte clone pair fewer than 10 units apart in color space. (**J**) Histogram displaying the frequency distribution of distances between regenerated cells from Sample 1 in HS color space. Approximately 10% of cell pairs in Sample 1 are predicted clones. (**K**) Single-clone lineage trace of regenerated joint tissues from Sample 2. Orange arrowhead denotes cyan chondrocyte circled in the corresponding polar graph. Orange arrow denotes cyan adipocyte circled in the corresponding polar graph. (**L**) Polar graph of HS color distribution of regenerated cells from Sample 2. Orange circle indicates a cyan adipocyte and chondrocyte clone pair fewer than 10 units apart in color space. (**M**) Histogram displaying the frequency distribution of distances between regenerated cells from Sample 2 in HS color space. Approximately 17.5% of cell pairs in Sample 2 are predicted clones. AA, anguloarticular bone; IOP, interopercular bone; Q, quadrate bone; RA, retroarticular bone. Scale bars = 250µm (A), 125µm (G, H, K).

Since our neural crest lineage tracing data revealed DsRed+ regenerated joint tissues of distinct cell types, we sought to delineate the degree of cellular plasticity in regenerating skeletal cells. To model plasticity, we applied archetype analysis (AA) to the scRNAseq dataset of zebrafish jaw joint regeneration. AA maps individual cells onto points within a simplex, offering insights into how cells balance multiple functions. The vertices of the simplex correspond to archetypes, which represent extreme gene expression profiles optimized for a functional task (*20–24*). Where each cluster falls within this simplex is indicative of how specifically it fits a single task (specialists close to archetypes) or how generalized it is to complete several tasks (generalists near the center of the simplex or along a continuum between two or more vertices). In the context of regeneration, we assume “generalist cells” correspond to multipotent progenitor-like states, while “specialists” reflect cells committed to particular lineages or functions. We calculated the variance explained by varying numbers of k vertices (k = 1-30) (Fig. S8A). As there was little biological variance explained by including additional archetypes, we chose a 6-vertex simplex to represent joint regeneration phenotypic space.

Next, we mapped the skeletal clusters from the scRNAseq dataset onto the simplex and performed Gene Ontology (GO) analysis to identify the genes and biological processes enriched at each vertex (Fig. S8B-C, S9, S10, Table S3). Throughout regeneration, mature skeletal cell populations were optimized for a single task (e.g., osteoblasts: ARCH3 (ossification and osteoblast differentiation) and chondrocytes: ARCH6 (cartilage development and positive regulation of developmental processes)), indicating a specialized function for each of these populations within the joint (Fig. 5B). Stromal and dermal fibroblast populations were also found among the specialists (i.e., DF 1: ARCH4 (L-phenylalanine metabolism); Stroma 3: ARCH2 (extracellular matrix organization)). In contrast, the regenerative mesenchyme mapped near the center of the simplex at all time points. Chondrogenic fibroblasts optimized a single task (ARCH5: cartilage development and negative regulation of developmental processes) in the uninjured and 70 dpjr samples. However, at 14 dpjr, chondrogenic fibroblasts plotted along a continuum between ARCH 5/ARCH 6 and between ARCH 5/ARCH 2, suggesting a degree of cell plasticity that enables them to shift between archetypes during joint regeneration. Highlighting cells classified as specialists on a UMAP of skeletal clusters further confirmed that none of the archetypes align with the regenerative mesenchyme cluster (Fig. 5C).

Additionally, we identified the cell type composition of cells close to each archetype and generated box plots of similarity scores for each cluster to each archetype (Fig. S11 & S12). While the majority of mature skeletal clusters had a high similarity score corresponding to one archetype (e.g., chondrocytes with a similarity score of >0.87 to ARCH6), the regenerative mesenchyme cluster did not have a similarity score above 0.40 when compared to any archetype. The large distance of the regenerative mesenchyme cluster from archetype vertices suggested that these cells are a generalist population with the ability to perform several functional tasks.

We then assessed the pseudotime distribution of all skeletal cells and found that the population with the earliest pseudotime (0 to 2) in the 1 dpjr sample largely consisted of regenerative mesenchyme cells, signifying their undifferentiated, progenitor-like state (Fig. 5D-E). Plotting osteoblast and chondrocyte differentiation trajectories onto simplex maps showed that mature clusters with the latest pseudotime (osteoblasts and chondrocytes) were optimized at archetype vertices, while the cell population with the earliest pseudotime (regenerative mesenchyme) was found near the center of the simplex (Fig. 5F). These data shed light on cell plasticity within the regenerating joint and suggest that neural crest-derived regenerative mesenchyme may be a multipotent population with the capacity to differentiate into multiple mature joint cell lineages with specialized functions.

To confirm in vivo whether neural crest-derived cells in the adult joint have multipotent potential during regeneration, we used inducible *sox10:CreERT2* (*25, 26*) in combination with *ubi:Zebrabow* (*27*). In the Zebrabow transgenic line, recombination results in permanent expression of either dTomato, mCerulean, or eYFP, enabling the tracking of both converted cells and their progeny. We treated *sox10:CreERT2;ubi:Zebrabow* fish with one dose of 5 μM 4-OHT from 24 to 36 hours post-fertilization (hpf) and performed clonal analysis in adulthood. In each joint, we identified chondrocytes, ligamentocytes, and adipocytes based on cell location and morphology. To determine distance in hue-saturation (HS) color space between cells in each sample, we developed a custom application in MATLAB 2024a (Fig. S13A-F). We assigned each cell a cell-type code (1: ligamentocyte, 2: adipocyte, 3: chondrocyte) (Fig. S14A-C); then, mean hue (H) and saturation (S) for each cell were calculated in the app, and these HS values were plotted on a polar graph. In uninjured controls, all 3 cell types were found dispersed across the RGB spectrum, (Fig. S15A-J). Previously, it has been shown that the mean distance between clonally derived cells is <40 units, with the frequency distribution peaking at ∼20 units apart in color space (*28, 29*). We employed a conservative binning approach of designating cell pairs between 0-10 color space units apart as predicted clones. Within this first bin of the histogram for each control sample, we identified cells of distinct lineages, highlighting the multipotent potential of the neural crest during development.

We then sought to discern whether these neural crest-derived tissues retain multipotency in adulthood. Single-clone lineage tracing analysis was performed on 56 dpjr regenerated samples (Fig. 5G-M, Fig. S16A-F). Similarly to control samples, we found that regenerated joints had a high proportion of cells 0-10 units apart in color space. Next, we assessed whether predicted clone pairs in regenerated samples included cells of different lineages. We found that regenerated samples contained predicted clone pairs of distinct lineages that were fewer than 10 units apart (i.e., green ligamentocyte and chondrocyte, Figure 5G-I; cyan chondrocyte and adipocyte, Fig. 5K-L). Additionally, we observed cells of the same lineage far apart in color space (i.e., >180 units), suggesting that multiple distinct clones contribute to each regenerated joint tissue type. As all preexisting joint tissues had been removed by joint resection, these single-clone lineage tracing data suggest that similarly to development, neural crest-derived cells responding to whole-joint injury in the adult zebrafish are not lineage restricted. Taken together, our computational and in vivo single-cell analyses implicate a multipotent, neural crest-derived cell population in retaining the ability to rebuild mature joint tissues in adulthood.

## Discussion

Here we establish a model of adult de novo synovial joint regeneration. While previous studies have explored mechanisms driving regeneration of individual joint tissues, single-tissue injuries do not provide a model for rebuilding joint tissues without a pre-existing source of the same cell type. Moreover, in the whole-limb regeneration models published to date, an ability to regenerate lubricating hyaline cartilage has not been described. In this study, we show that after complete removal of the synovial jaw joint in the adult zebrafish, all native joint tissues regenerate. The regenerated articular cartilage expresses *prg4b*, thereby restoring the self-lubricating function of the joint, a phenomenon which has not been demonstrated in previous joint regeneration models. While tissue patterning of the uninjured joint is not wholly recapitulated after complete resection, we used μCT reconstructions, testing of feeding coordination, and imaging of compatible transgenics to show that regenerated joint tissues assemble into an integrated 3D structure that generates functional movement.

Whether regenerating tissues arise from a progenitor pool or via cellular dedifferentiation has long been a topic of debate. Many published models, such as zebrafish fin, ligament, and heart regeneration, as well as newt muscle regeneration, rely primarily on mechanisms of dedifferentiation (*14, 30–32*). Our joint resection surgery differs from these injury models in that we fully remove the articular cartilage, ligaments, and synovium that could serve as a source for rebuilding these tissues. Some dedifferentiation of osteoblasts in the remaining bone stubs may occur, as we observe bony protrusions as early as 3 dpjr. However, we also see new bone growth in the form of bony islands within the mesenchyme, indicating that regenerating osteoblasts may be derived from multiple sources. The de novo regeneration of joint tissues after complete resection suggests either the presence of a resident progenitor pool in the uninjured joint or the occurrence of non-lineage-restricted cell dedifferentiation. This differs from existing zebrafish regeneration models where dedifferentiation does not result in the acquisition of multipotency (e.g., fin regeneration) (*30*). The archetype and clonality analyses we performed on regenerated cells in our model, however, support a multipotent neural crest-derived population giving rise to distinct synovial joint cell lineages. This observation parallels published work which found that axolotl limbs regenerate through dedifferentiation of connective tissue cells into a multipotent progenitor (*29*). Since regenerated axolotl limbs are mesoderm-derived and regenerated zebrafish jaw joints are neural crest-derived, activation of the multipotent state observed in both models is likely independent of cells’ embryonic lineage.

An emerging theme in the field of skeletal biology is the importance of tissue cross-talk in maintaining joint homeostasis (*33, 34*). Our whole-joint resection model enables the investigation of how cross-talk between different joint tissues (i.e., bone and articular cartilage) may facilitate the regeneration of the joint as an integrated organ. Our scRNAseq data also captured non-skeletal populations such as immune cells, neural cells, and endothelial cells. Immune cells in particular show major population shifts during early regeneration (Fig. S4), and future work will probe the functional role of such non-skeletal cells in rebuilding the joint. We also observed that the domains of proliferating epithelial cells were dynamic (Fig. S2C-D). Given the wound epithelium’s role in recruiting fibroblasts to form a blastema (*35*), our model provides a new paradigm to investigate how epithelial-mesenchymal cell cross-talk may contribute to zebrafish whole-joint regeneration.

The regenerative program induced after our surgery differs from the fracture healing response of other zebrafish craniofacial skeletal regeneration models (*12*). While lower jawbone healing in zebrafish relies on hybrid bone-cartilage cells derived from the periosteum, we did not identify any clusters co-expressing osteoblast and chondrocyte markers in our scRNAseq data nor any islands of hybrid bone-chondrocytes in our histological analysis. This indicates that after joint resection, zebrafish skeletal tissues undergo a mechanism of healing unique from the response to lower jawbone resection or fracture healing. In mouse models of osteoarthritis, *Sox9*-expressing progenitors contribute to osteophyte formation rather than joint tissue regeneration (*36*). Hence, although mammalian joint cells respond to acute injury, they fail to correctly re-deploy developmental pathways. In contrast, we have shown that adult zebrafish are able to engage resident skeletal cells to correctly activate developmental genes to rebuild multiple joint tissue lineages. Our finding of *sox9a* upregulation lining the resected bone stubs is consistent with a model wherein the initial re-expression of joint development genes occurs in periosteal cells. Future lineage tracing experiments are required to determine whether the periosteum serves as a source for regenerating joint cells. Unraveling the cues guiding resident skeletal cells to successfully regenerate joint tissues in response to resection injury in adult zebrafish lays the foundation for awakening a similar regenerative program in human joints.

## Materials & Methods

### Zebrafish Lines

All zebrafish experiments were approved by the Columbia University Institutional Animal Care and Use Committee. Published lines used in this study include *AB*, *Tg(scxa:mCherry*)*fb301* (*37*), *Tg(sp7:EGFP)b1212* (*38*), *Tg(thbs4a_p1:eGFP)el912* (*14*), *Tg(actab2:loxP-BFP-STOP-loxP-DsRed)sd27* (*39*), *Tg(Mmu.Sox10-Mmu.Fos:Cre)zf384* (*17*), Tg(*sox10*:CreERT2)*el777* (*25*), and Zebrabow - *Tg(ubb:LOX2272-LOXP-RFP-LOX2272-CFP-LOXP-YFP)a13* (*27*). Embryos were raised in a methylene blue salt solution at 28.5 °C, while juvenile and adult fish were maintained in groups of 15-30 animals. Joint resection surgeries were performed in size-matched 3-9 mpf adult zebrafish using standard body length (SL) measurements.

### Adult Joint Resection Surgery

Fish were anesthetized with Tricaine MS-222 at a concentration of 0.17mg/mL and placed into a damp sponge ventral side up. 3-mm Vannas spring scissors (Fine Science Tools, cat. #1500000) were then used to make 3 cuts to remove the entirety of the quadrate-articular joint on the left side. The surgery also removed the retroarticular bone between the IOM ligament and anguloarticular bone. The first cut was performed anterior to the quadrate through the anguloarticular bone, and the second cut was performed posterior to the quadrate through the interopercular bone (Fig. 1A). Student Dumont #5 Forceps (Fine Science Tools, cat. #9115020) were used to grasp the joint tissue before making the third cut dorsal to the joint along the quadrate bone. The forceps were then used to remove the joint as well as any residual tissue or bone fragments in the resection site. Fish were then revived by circulating clean system water over the fish’s gills, housed on-system, and monitored for three days to ensure normal swimming and behavior post-surgery. For euthanasia at experimental endpoints, fish were placed into a lethal dose of Tricaine MS-222 for 30 minutes on ice.

### MicroCT

For microCT (µCT), adult zebrafish were euthanized, fixed in 4% PFA overnight, and rinsed twice in 1x PBS. The scans were performed on the µCT system (MILabs, Netherlands) using a fixed 100µm aluminum primary filter and a standard 400µm aluminum secondary filter. The parameters were as follows: voltage of 20 kV, current of 0.07 mA, and exposure time of 38 ms, resulting in a 10 micron voxel volume. Raw data was reconstructed in 3D Slicer image computing platform.

### Functional Testing

Functional testing was conducted by quantifying slow motion footage of jaw activity during swimming and feeding. All fish used for functional testing were wild-type ABs ∼8-9 months old, with an equal number of males and females tested, and fish were selected to be a similar weight and length for their respective sex. Videos of the fish were collected one day before whole-joint resection, 3 hours after whole-joint resection, and then again for 1, 3, 7, 14, 21, 28, and 56 days after whole-joint resection, all at the same time of day. Footage was captured using a Canon EOS Rebel T6 DSLR Camera with an EF-S 18-55mm lens. Videos were taken using a resolution of 1280×720 at 60fps, then imported to Microsoft Clipchamp software where they are slowed down to one tenth speed for analysis.

### Histology

Adult zebrafish samples were prepared for histological analysis as previously described (*14*). Briefly, following euthanasia and overnight fixation in 4% PFA/1xPBS at 4 °C, samples were washed twice for 30 minutes in 1xPBS. The head was then separated from the trunk for embedding of desired structures. Samples were decalcified in 0.5 M EDTA solution for 10 days rocking at room temperature. Prior to embedding, the tissue was dehydrated with a series of ethanol/DepC-treated water dilutions (DepC, 30% EtOH, 50% EtOH, 70% EtOH, 95% EtOH, 100% EtOH) followed by a second series of ethanol to Hemo-De Xylene Substitute dilutions (50% Hemo-De, 75% Hemo-De, 100% Hemo-De) (Electron Microscopy Sciences, 23412-01). The samples were then incubated in a 1:1 Hemo-De:Paraffin solution at 65 °C for an hour. After an overnight incubation in 100% paraffin at 65 °C, samples were embedded in freshly melted paraffin. Paraffin sectioning was performed using Thermo HM355S automatic microtome to collect 5 μm sections.

For hematoxylin & eosin (H&E) staining, 5 μm paraffin sections were deparaffinized with Hemo-De Xylene Substitute (Electron Microscopy Sciences, 23412-01) and rehydrated through an ethanol series to distilled water. Sections were then stained in hematoxylin (Sigma-Aldrich, HHS16) for 2min followed by a brief rinse in acetic acid (Fisher Scientific, A38-212), 20x dip in water, 2min in Scott Tap Water Bluing Reagent (Mercedes Scientific, 11160), 2 x 15x dips in water, and 2 x 15x dips in 95% ethanol. The sections were then stained in Eosin (Sigma-Aldrich, E6003) for 30 s followed by 3 x 1 min washes in 95% ethanol and 2 x 1 min washes in 100% ethanol. After 2 x 2 min in Hemo-De, samples were mounted with cytoseal (Fisher Scientific, 23-244256) for imaging.

### Tissue Clearing

CUBIC tissue clearing protocol was adapted from Susaki *et al.,* 2014 (*40*) and used for zebrafish as described in Anderson & Mo *et al.,* 2023 (*14*). Whole adult zebrafish were fixed using 4% PFA/PBS at 4°C overnight. After samples were washed in PBS twice for 30 minutes each while rocking, heads were separated from the trunk of the body. At a ratio of 5-8 heads per 50mL, samples were incubated in a CUBIC-1 solution in the dark at 37°C with rocking for 3-4 days. Tissue was then transferred to fresh CUBIC-1 solution for another 2-4 days. After removing CUBIC-1, samples were washed twice in PBS for 30 minutes each at RT and transferred to a 20% sucrose/PBS solution for ∼2 hours at RT (all washes with rocking). Then, the samples were transferred to CUBIC-2 solution in the dark at RT, with gentle rocking for 2-3 days before imaging. Once sufficiently cleared, heads were mounted in a chambered coverglass with a drop of CUBIC-2 (Fisher Scientific, 12-565-335) and imaged using a Leica SP8 confocal microscope.

Both CUBIC-1 and CUBIC-2 solutions were prepared at 100°C while covered. CUBIC-1 was a mixture of 25 wt% urea, 25 wt% N,N,N’,N’-tetrakis(2-hydroxypropyl) ethylenediamine (Fisher Scientific, AAL16280AP) and 15 wt% Triton X-100. Deionized water was heated and added to N,N,N’,N’-Tetrakis(2-hydroxypropyl)ethylenediamine. After dissolving, urea was added to the solution and mixed until homogenous. Solution was then removed from heat, and after cooling to room temperature, Triton-X-100 was added at RT with gentle stirring.

CUBIC-2 was prepared as a mixture of 50 wt% sucrose, 25 wt% urea, 10 wt% 2,2′,2′’-nitrilotriethanol (Fisher Scientific, AAL044860E), and 0.1% (v/v) Triton X-100. Deionized water was heated and added to 2,2′,2′’-nitrilotriethanol. When the mixture was homogenous, urea was added. After the urea was sufficiently dissolved, sucrose was mixed into the solution. Once the solution was clear, it was removed from heat and allowed to cool to room temperature. Triton-X-100 was then added at RT with gentle stirring. Both CUBIC-1 and CUBIC-2 solutions were stored in the dark at RT after preparation and used within 1 week.

### Calcein Blue Bone Staining

Calcein blue bone staining solution was prepared by adding Calcein blue powder (Millipore Sigma, C1429) to 1X PBS to a final concentration of 1mg Calcein blue/2 mL 1X PBS. After fixation, samples were stained for 2-3 days at 4 °C in a Calcein blue solution, covered to minimize light exposure. Samples were washed twice in 1X PBS before proceeding to tissue clearing and imaging.

### Immunofluorescence, HCR, and RNAscope

Immunofluorescence and smFISH experiments were performed on 5μm formalin-fixed paraffin-embedded (FFPE) tissue sections. For immunofluorescence, sections were deparaffinized with 3 Hemo-De washes and rehydrated through an ethanol series (100%, 80%, 50%). Antigen retrieval was performed using sodium citrate buffer, and samples were blocked for 1 hour with GeneTex immunostain blocker solution (GTX73323). After an overnight 4 °C incubation with primary antibody (PCNA, 1:1000, Sigma-Aldrich, P8825; Sox9a, 1:200, GeneTex, GTX128370), samples were washed with 1X PBS-T and incubated with secondary antibody (1:500, Invitrogen, A-21245, A-11001) diluted in ImmunoStain blocker solution (GeneTex, GTX73323) and Hoechst 33342 nuclear stain (1:1000, Fisher Scientific, 51–17). Sections were mounted with Fluoromount-G (Southern Biotech, 0100-01).

Single-molecule fluorescence in situ hybridization was performed according to manufacturer guidelines using RNAScope Multiplex Fluorescent Reagent Kit v2 Assay (ACD Bio) or Hybridization Chain Reaction (HCR) RNA-Fish (Molecular Instruments), except that slides were covered in parafilm for the HCR overnight incubation step in probe solution. Target retrieval was performed for 4 min in a steamer (using 1x Target Retrieval Reagent for RNAscope and sodium citrate buffer for HCR).

RNAScope probes used were Dr-*sox9a*-C3 (543491-C3) and Dr-*gdf5*-C2 (1255581-C2) from ACD Bio. Opal 520 and 570 fluorophores were used for visualizing expression (1:1000; Akoya Biosciences, cat. #FP1487001KT, #FP1488001KT). HCR probes used were *sox9*a-B1 (0.4pmol, Molecular Instruments), *nkx3-2*-B3 (0.4pmol, Molecular Instruments), and *fn1a*-B5 (0.4pmol, IDT). The amplifiers used were B1-647, B3-546, and B5-488 (Molecular Instruments). Sections were mounted with Fluoromount-G+DAPI (Southern Biotech, 0100-02).

### Drug Treatments

One treatment with 5 µM (Z)-4-Hydroxytamoxifen, ≥98% Z isomer (Sigma-Aldrich, H7904) was used to convert *sox10:CreERT2* lines in the dark from 24 to 36 hpf. Following drug treatment, embryos were washed 2×15 min in methylene blue water and then screened for conversion using a Leica M165FC stereo microscope before being housed on-system.

### Imaging

Images of histological sections were captured using a Zeiss AxioLab 5 Microscope with a Axiocam 208 color camera (Zen v3.9.101.01000).

Live imaging was performed using a Leica M165FC stereo microscope with Leica K5 sCMOS camera and LASX software v3.7.4.23463. Fluorescence imaging was performed using a Leica SP8 confocal microscope with LASX software v3.5.7.23225. All images were processed in Fiji (*41*).

### Single-cell RNA sequencing

Whole-joint resection (n=18 joints for 1 dpjr, 10 joints for 3 dpjr, 7 dpjr, and 14 dpjr, 12 joints for 28 dpjr, and 13 joints for 70 dpjr) was performed on size-matched *Sox10Cre;actb2:loxP-BFP-loxP-DsRed; flk1:GFP* fish at 3-6 mpf. Regenerated jaw joint tissues from each injured fish and uninjured controls (32 joints) were microdissected and incubated in [0.25% trypsin (Life Technologies, 15090–046), 1 mM EDTA, and 400 mg/ml collagenase D (Sigma-Aldrich, 11088882001) in HBSS] for 1 hr at 28 °C. Mechanical dissociation of joints was achieved by nutation and agitation with a P1000 pipette every 5 min for the 1 hr incubation period. Regenerated joints are represented by two independent samples per time point (1 sample of pooled joints and 1 sample from a single animal), and uninjured joints are represented by 3 independent samples (1 sample of pooled joints and 2 samples from single animals). Live cells were sorted with FACS and immediately processed to create single-cell RNA Sequencing libraries using the Chromium Single Cell 3′ Library & Gel Bead Kit v2 (10X Genomics), following the manufacturer’s recommendations. Libraries were sequenced at the Columbia Single Cell Analysis Core on the NovaSeq 6000 (Illumina, San Diego, CA) with 101 bp sequencing for Read1, 10 bp sequencing for Index1, 101 bp sequencing for Read2, and 10 bp sequencing for Index2. Each sample was sequenced to a mean read depth of greater than 86,000 reads per cell. CellRanger v6.1.2 (10X Genomics) was used for alignment to GRCz11 with default parameters to generate cell-by-gene count matrices.

scRNAseq libraries were analyzed using R version 4.4.0 in RStudio with Seurat (v5.0.3.) CellRanger outputs for each sample were converted to Seurat objects and merged. Outputs were filtered for cells between 600 and 2,500 unique features per cell and less than 7% mitochondrial RNA detected per cell to remove low-quality cells and doublets. Next, data were processed with log normalization, selection of 2,500 variable features, and scaled using all features expressed in 3 or more cells. Samples were then integrated with RPCA-based integration, and principal component analysis was used to perform linear dimensional reduction to compute 150 principal components. Jackstraw permutation identified all 150 principal components as significant (p<0.05), and K-nearest-neighbor graph construction (KNN) was performed. After quality control filtering, a total of 63,305 high-quality cells with a median 600 genes per cell were obtained. Metadata for pooled and single-animal samples from the same injury time points were merged for downstream analysis. Broad superclusters were identified by performing clustering at a low resolution of 0.02. Clusters and gene expression plots were graphed using UMAP-non-linear dimensional reduction to maintain global variability.

To analyze scRNAseq data from skeletal cells, the ‘Skeletal 1’, ‘Skeletal 2’, and ‘Perivascular’ superclusters were subset and merged. Unprocessed gene expression data from these cells was re-processed with log-normalization, feature selection, and feature scaling. Principal components 1-150 were used for KNN and graphed using UMAP-non-linear dimensional reduction before clustering at a resolution of r=0.9 (Fig. S5A-B). Two clusters (cluster 18 and 22) expressed epithelial cell markers (*epcam*) and were excluded as a likely doublet cluster. Unprocessed data from cells in remaining clusters were then re-processed with log normalization, feature selection, and feature scaling. Principal components 1-150 were used for KNN and graphed with UMAP-non-linear dimensional reduction, and clustering was performed once more at a higher resolution of r=0.95. Cluster markers were identified with function FindAllMarkers using a Wilcoxon Rank Sum test for features expressed in at least 10% of the cluster and a minimum of 0.2 log-fold-change greater expression than the other cells in the skeletal clusters.

### Pseudotime Analysis

R package Monocle3 (v1.3.7) was used for pseudotime trajectory analysis. The pre-analyzed Seurat object containing all skeletal cells was converted to a monocle format. To limit bias, the root of the trajectory was specified programmatically rather than with manual selection. A helper function was used to group cells from the 1 dpjr sample according to which trajectory graph they are nearest to, calculate the fraction of cells at each node from the earliest time point, and designate the node that is most heavily occupied by early cells as the root (regenerative mesenchyme). The output was used to assess pseudotime distribution and cell type composition of 1 dpjr cells (Fig. 5D-E). To identify genes upregulated along osteoblast and chondrocyte trajectories, the osteoblast, chondrocyte, chondrogenic fibroblasts, and regenerative mesenchyme clusters were subset from the skeletal cell object and converted to a monocle format. Pre-processing, alignment, and UMAP dimensionality reduction were then performed. Cells were clustered using Monocle3’s algorithm and a principal graph within each partition was fit using the learn_graph function. The same helper function described above was then used to programmatically select regenerative mesenchyme as the root node. Cells were plotted and colored by both pseudotime and cluster (Fig. 4E). The output was used for further analyses, including finding genes that change as a function of pseudotime (Fig. 4F) and finding modules of co-regulated genes (Fig. S6 & S7).

### Quantification and Statistical Analysis

Proliferation was measured as the percentage PCNA+ cells relative to the total number of Hoechst+ cells within a region of interest (ROI) in the uninjured joint domain or regenerating joint injury domain between the resection sites. Additional ROIs were defined within the larger joint area, targeting specific regions such as the uninjured chondrocyte domain, regenerating chondrocyte domain, uninjured epithelium domain, injury epithelium domain, and injury-adjacent epithelium domain. ROIs were determined based on the regional nuclear morphology in the tissue section. Hoechst+ cells were quantified using FIJI’s StarDist nuclei detection and segmentation tool, while PCNA+ cells were manually scored by an observer blinded to both the time point and ROI identity.

Expression levels of *fn1a*, *nkx3.2*, *sox9a*, and *gdf5* were quantified from smFISH images using ImageJ. Uninjured and regenerating joint domains were delineated based on regional nuclear morphology within the tissue section. Regenerating joint domains were specifically outlined adjacent to the cut sites to encompass the entire joint injury area, capturing bone stubs, regenerating chondrocyte islands, injury epithelium, and other regenerating tissues located between the cut sites. The corrected total fluorescence was calculated by determining the integrated density within each ROI for each channel. Tissue autofluorescence and background signal was accounted for by calculating the average mean gray value from 3 distinct background regions within each image. The corrected total fluorescence (CTF) was determined using the following formula: CTF = Integrated Density – (Area of ROI × Average Mean Gray Value).

To quantify the output of regeneration after joint resection, slow motion footage of zebrafish suction feeding was analyzed and assigned a score based on premaxillary protrusion as a function of musculoskeletal coordination. A double-blind verification was used to confirm the assigned functional scores which maintained the observed trend. Additional recordings were taken of the time in between consecutive suction feeding actions, measured as the time in milliseconds between the closing of the mouth and the very beginning of the next opening movement. These values were normalized relative to the uninjured time between feeding actions for each individual zebrafish to attain a fold change in the time between consecutive feeding actions.

All statistical analyses were conducted using GraphPad Prism 10.3.0. Proliferation in the whole joint region and epithelial domains was evaluated using ordinary 1-way ANOVA with Tukey’s multiple comparison test, while proliferation in the chondrocyte domains was assessed with a 2-tailed unpaired t-test. Transcript expression of *fn1a*, *nkx3-2*, and *sox9a* across different post-injury timepoints was assessed using 2-way ANOVA with Tukey’s multiple comparison test. The expression of *gdf5* transcripts was evaluated with ordinary 1-way ANOVA with Tukey’s multiple comparison test. Comparisons of functional testing outputs across timepoints post-injury were evaluated using a chi-square test. Error bars in all statistical analyses represent standard error of mean (SEM).

### Archetype Analysis

To perform archetype analysis (AA), the gene expression encodings of cells within the Perivascular, Skeletal 1, and Skeletal 2 superclusters, derived from the dimensionality reduction procedure outlined in the Single-Cell RNA Sequencing section, were utilized. Each cell’s expression encoding was represented by *x̃_i_* ϵ ℝ^*m*^, where *m* = 150 principal components (PCs) were selected.

Archetype was defined as a convex combination of the PC encodings *x̃_i_* in the dataset, as follows:

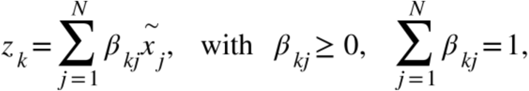

where *N* denoted the number of all cells in the scRNA-seq dataset.

Each gene expression encoding *x̃_i_* was subsequently modeled as a convex combination of the archetypes:

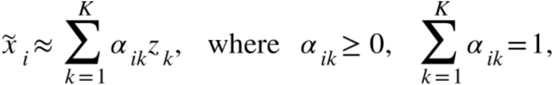

where α_*ik*_ corresponds to the coordinates of cell *i* with respect to archetype *k*, allowing the positioning within the simplex, and *K* is the total number of archetypes.

To find the archetype vectors *z_k_* and coefficients α, β the original data reconstruction error was minimized:

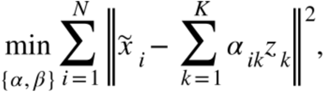

subject to the constraints:

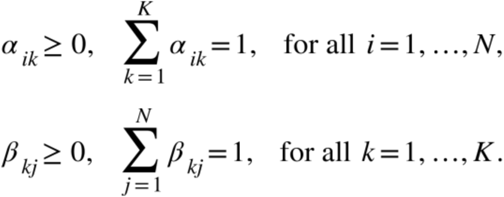

The Seurat object, containing the PC encodings, was converted to a Scanpy object using the R package MuDataSeurat (v0.0.1), enabling further analysis in Python. To optimize the resulting objective function (*21*), the Projected Gradient Descent algorithm (*42*), as implemented in the scikit-learn-compatible Python package archetypes, was utilized with default parameters. The analysis used a forked version of the package (October 19, 2024) available at https://github.com/wangzcl/archetypes.git (original repository at https://github.com/aleixalcacer/archetypes). This package provides the α coefficients, referred to as similarity scores, which quantified each data point’s association with each archetype. The scores ranged from 0 to 1, where 0 indicated maximal distance from an archetype, and 1 signified perfect alignment.

### Gene Ontology Enrichment at Archetype Locations

Raw gene expression counts were normalized using the NormalizeData function from the Seurat package with the default parameter normalization.method = ’LogNormalize’. For each archetype, cells were classified into three groups based on the value of similarity scores. The classification criteria were as follows: close cells with a similarity score greater than 0.9, medium cells with a similarity score between 0.8 and 0.9, and distant cells with a similarity score less than 0.8. Using the Seurat function FindMarkers, differentially expressed genes between close and distant cells were identified with a minimum expression threshold of 0.1 (min.pct = 0.1) and a log-fold change threshold of 0.2 (logfc.threshold = 0.2). The top 150 genes, ranked by average log-fold change and with the adjusted p-value significance less than 0.001, were selected for Gene Ontology (GO) Enrichment Analysis. The GO analysis was performed with the following parameters: ontology ont=’BP’ (Biological Process), adjustment method pAdjustMethod=’BH’ (Benjamini-Hochberg correction), pvalueCutoff=0.01, and qvalueCutoff=0.05.

### Zebrabow Clonal Lineage Tracing

For clonal analysis using the *ubi:Zebrabow* lines, images were captured on a Leica SP8 confocal microscope with LASX software v3.5.7.23225 and processed with Lightning. Using representative slices throughout the Z-stack, cells were categorized into cell type (1: ligamentocyte, 2: adipocyte, 3: chondrocyte) based on cell location and morphology (Fig. S14A-C). Cell-type codes were manually assigned using the Point Tool in Fiji. Confocal images were then analyzed using a custom application developed in MATLAB 2024a. The input to the application was a set of image pairs consisting of the raw, unlabeled confocal image (Fig. S13A) and its corresponding cell-coded image (Fig. S13B). For each pair, the difference between the two images was calculated to provide a ‘cell label’ only image (Fig. S13C). This was then scanned to sequentially identify labels indicating cell markers and cell-type code, providing the location of the cell and its cell type, respectively (Fig. S13D). RGB values were extracted from the raw images of the cells based on the position delineated by each cell marker and averaged across these pixels to provide a mean hue and saturation value for a given cell (Fig. S13E). The number associated with that marker was then recognized to identify cell type. The corresponding cell hue and saturation were then plotted in a polar plot as angle and radius, respectively (Fig. S13F). The polar coordinates were used to calculate distance in color space for every potential pairwise comparison of individual cells, allowing for determination of cell lineage, using the standardized equation:

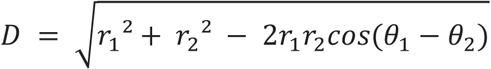

where *D* is distance, *r* is radius, and *θ* is the angle in radians. This application allows for tabulation of cells, quantification of distances, and polar plot visualization with interactive functionality that allows users to click on points in the graph to determine that corresponding cell information and visualize its position in the image.

To determine the distance in color space between known clones, the uninjured contralateral side of the joint was imaged. Hue and saturation were recorded for cells with similar color, location, and morphology in these control samples, as these cells were assumed to be clones. The distance in color space between known clones was calculated. Using this value of expected variance in color space for clonally related cells, cells in regenerated joints were then determined to be either predicted as clones or non-clones. Five contralateral uninjured control and five regenerated samples were included in the analysis.

## Acknowledgments

We thank Joshua Barber for fish care, Michael Kissner at the CSCI Flow Core, Dr. Andrei Molotkov and Dr. Akiva Mintz (supported by the Accelerate Resource of NCATS UL1TR001873 (Reilly)) at the CUIMC PET Imaging Core, and Erin Bush, Joseph Mullen, and Kwasi Osae-Kwapong at the CUIMC Single Cell Analysis Core.

## Author contributions

Conceptualization: JS, FW, MB

Methodology: MB, FW, EG, EW, AG, LC, MK, BD, NC, JS

Investigation: MB, FW, EG, EW, AG, LC, TA, JM, DS, MG, MK

Visualization: MB, FW, EG, EW, AG, LC, MK, MG

Funding acquisition: JS, MB, EW, LC, AG, NC

Project administration: JS

Supervision: JS

Writing – original draft: MB, EW, MK, AG, LC, JS

Writing – review & editing: MB, FW, EG, EW, MK, AG, LC, TA, JM, DS, BD, NC, JS

## Data and materials availability

Raw and processed scRNA sequencing files have been deposited in GEO and can be retrieved using accession number GSE283763. All other raw data, transgenic zebrafish lines, and/or materials from this study are available upon request from the corresponding author (JS).

