## Supplemental Figures and Legends for "Dynamic cell fate plasticity and tissue integration drive functional synovial joint regeneration"

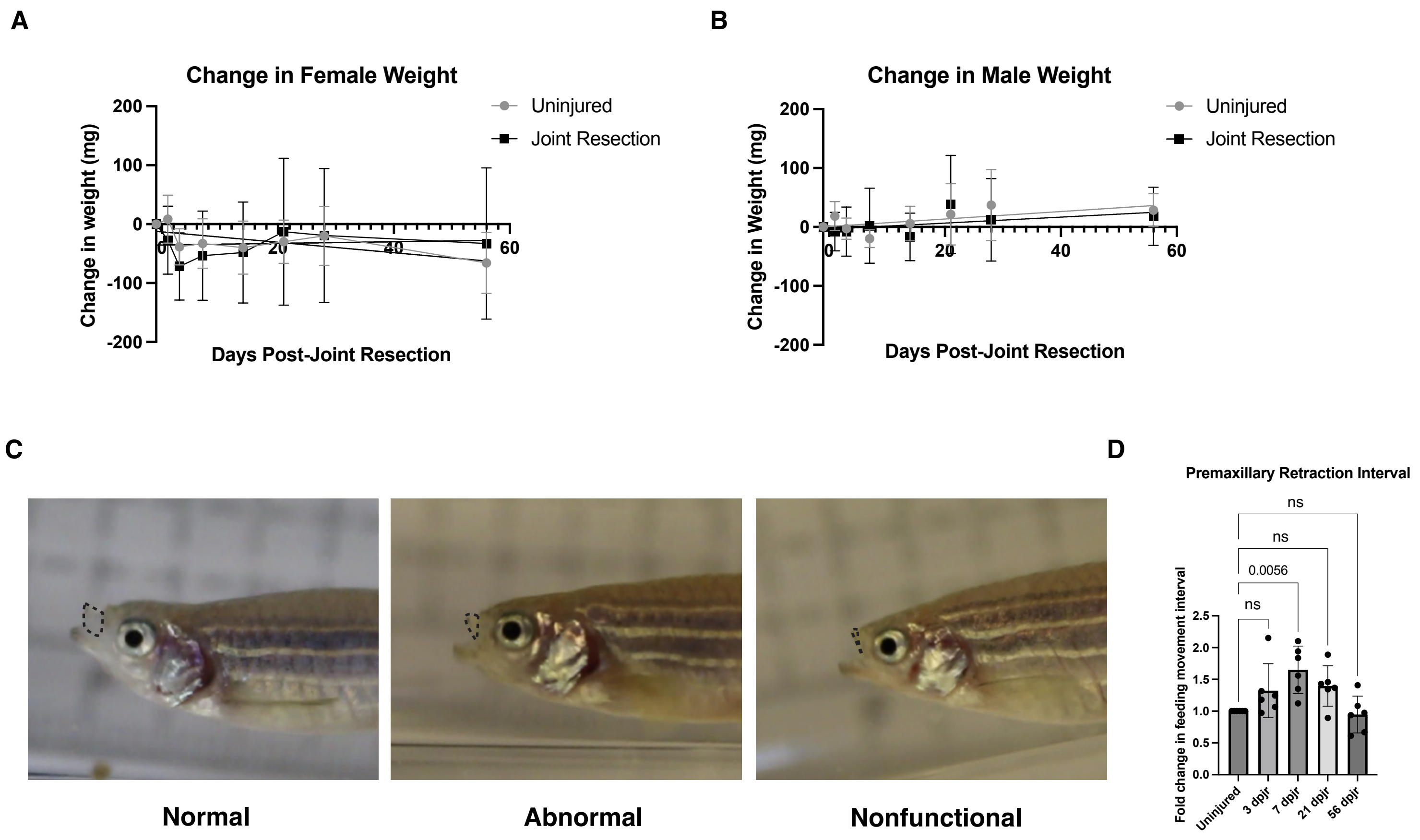

A
Change in Female Weight
Uninjured
Joint Resection
Change in weight (mg)
Days Post-Joint Resection
B
Change in Male Weight
Uninjured
Joint Resection
Change in Weight (mg)
Days Post-Joint Resection
C
Normal
Abnormal
Nonfunctional
D
Premaxillary Retraction Interval
Fold change in feeding movement interval
ns
0.0056
Uninjured
3 dpjr
7 dpjr
21 dpjr
56 dpjr

**Fig. S1: Restoration of feeding coordination during joint regeneration**. (**A** to **B**) Repeated measurements of weight of female (A) and male (B) fish over a period of 56 dpjr. There were no significant changes in weight between the injured and uninjured samples. (**C**) Representative images of normal, abnormal, and nonfunctional premaxillary protrusion as a measure of joint function. Premaxillary protrusions outlined in black dashed lines. (**D**) Quantification of the time interval between craniofacial suction feeding movements. Error bars represent SEM.

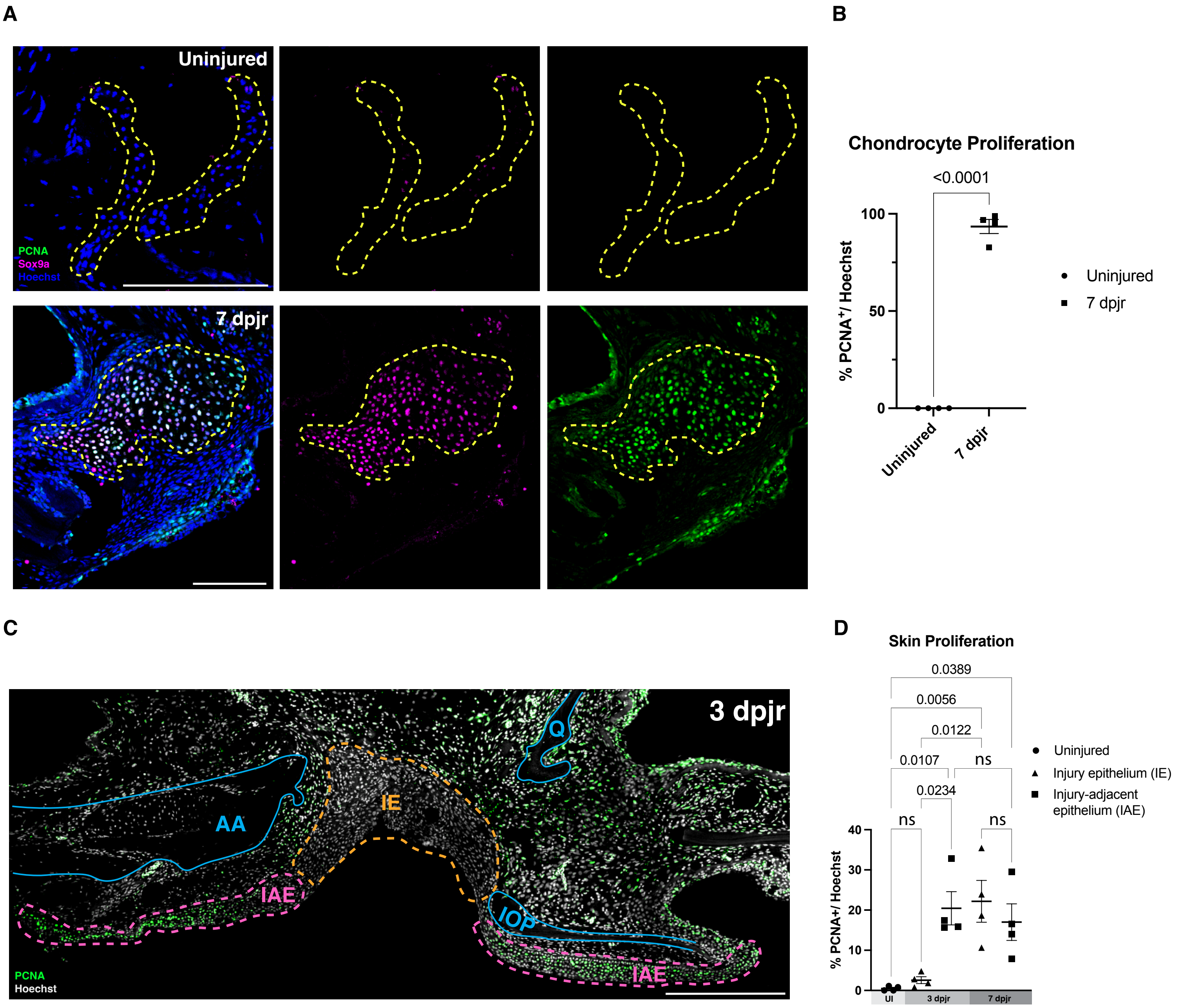
A
Uninjured
PCNA
Sox9a
Hoechst
7 dpjr
B
Chondrocyte Proliferation
<0.0001
100
50
0
% PCNA+/ Hoechst
Uninjured
7 dpjr
Uninjured
7 dpjr
C
3 dpjr
Q
AA
IE
IAE
IOP
IAE
PCNA
Hoechst
D
Skin Proliferation
0.0389
0.0056
0.0122
0.0107
ns
0.0234
ns
ns
40
30
20
10
0
% PCNA+/ Hoechst
UI
3 dpjr
7 dpjr
Uninjured
Injury epithelium (IE)
Injury-adjacent epithelium (IAE)

**Fig. S2: Dynamics of proliferating cells in early regeneration**. (**A**) PCNA and Sox9a immunofluorescence (IF) in the uninjured joint cavity and in chondrocyte islands at 7 dpjr. (**B**) Quantification of PCNA+ nuclei in articular chondrocytes of the uninjured joint and in regenerating chondrocyte islands at 7 dpjr, showing an increase in chondrocyte proliferation at 7 dpjr. Regions of interest for quantification outlined in yellow dashed lines; error bar represents SEM. (**C**) Representative image of regions of interest for quantification of epithelial proliferation at 3 dpjr. Pink dashed lines denote injury-adjacent epithelium (IAE) surrounding the resected bone stubs and orange dashed line indicates injury epithelium (IE) bridging the wound. (**D**) Quantification of PCNA+ IAE and IE at 3 dpjr and 7 dpjr compared to uninjured skin (n=4 per time point); error bars represent SEM. AA, anguloarticular bone; IOP, interopercular bone; Q, quadrate bone. Scale bars = 100µm (A), 300µm (C).

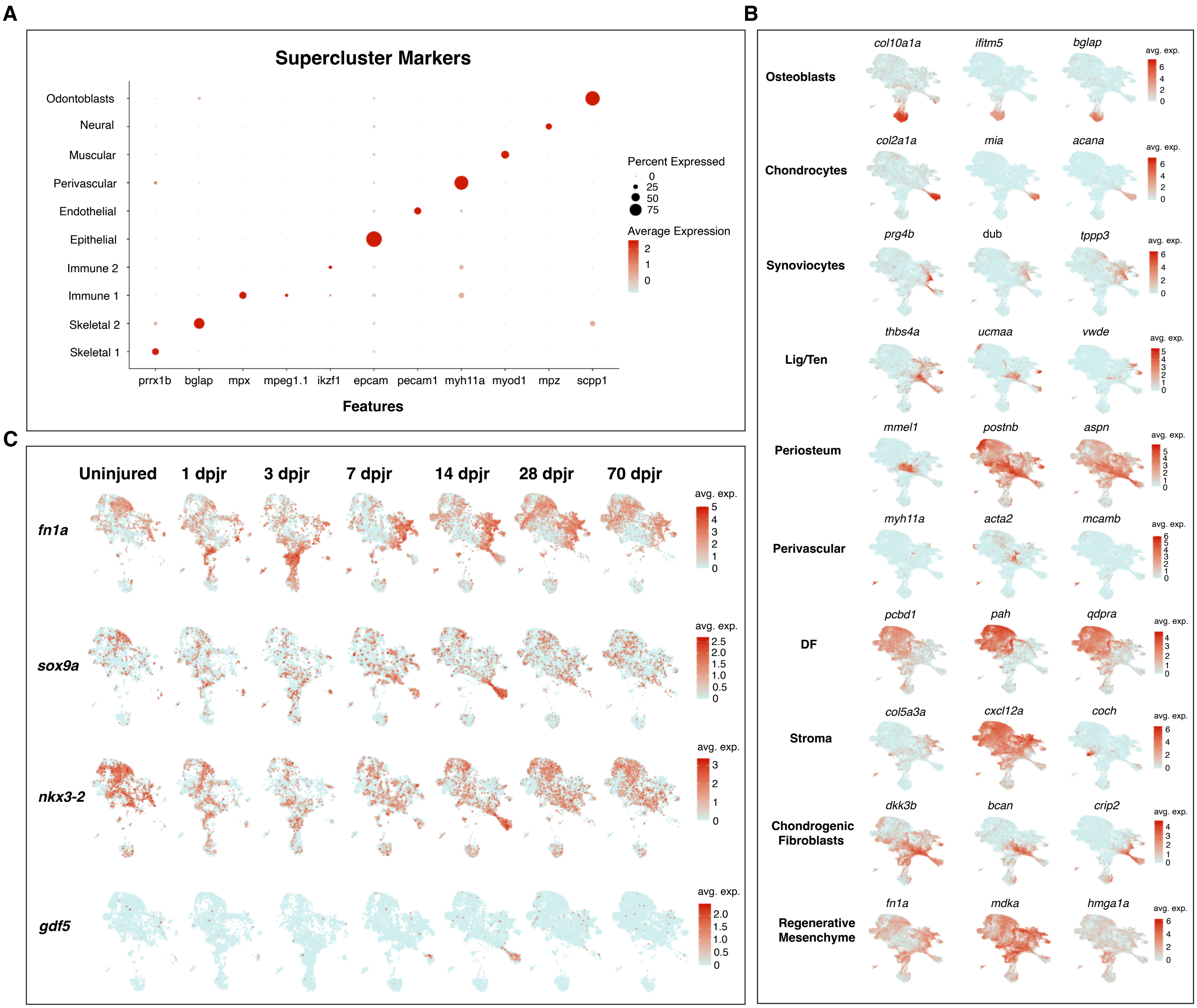

A
Supercluster Markers
Odontoblasts
Neural
Muscular
Perivascular
Endothelial
Epithelial
Immune 2
Immune 1
Skeletal 2
Skeletal 1
prrx1b
bglap
mpx
mpeg1.1
ikzf1
epcam
pecam1
myh11a
myod1
mpz
scpp1
Features
Percent Expressed
0
25
50
75
Average Expression
2
1
0
B
Osteoblasts
col10a1a
ifitm5
bglap
Chondrocytes
col2a1a
mia
acana
Synoviocytes
prg4b
dub
tppp3
Lig/Ten
thbs4a
ucmaa
vwde
Periosteum
mmel1
postnb
aspn
Perivascular
myh11a
acta2
mcamb
DF
pcbd1
pah
qdpra
Stroma
col5a3a
cxcl12a
coch
Chondrogenic Fibroblasts
dkk3b
bcan
crip2
Regenerative Mesenchyme
fn1a
mdka
hmga1a
avg. exp.
C
Uninjured
1 dpjr
3 dpjr
7 dpjr
14 dpjr
28 dpjr
70 dpjr
fn1a
sox9a
nkx3-2
gdf5
avg. exp.

**Fig. S3: Transcriptional heterogeneity of regenerating joint tissues via scRNAseq**. (**A**) DotPlot of marker genes used to identify each supercluster (r=0.02). (**B**) FeaturePlot expression of *fn1a*, *sox9a*, *nkx3-2*, and *gdf5* over time in the skeletal clusters. (**C**) FeaturePlots of differentially expressed marker genes used to define skeletal cell clusters at a resolution of 0.95.

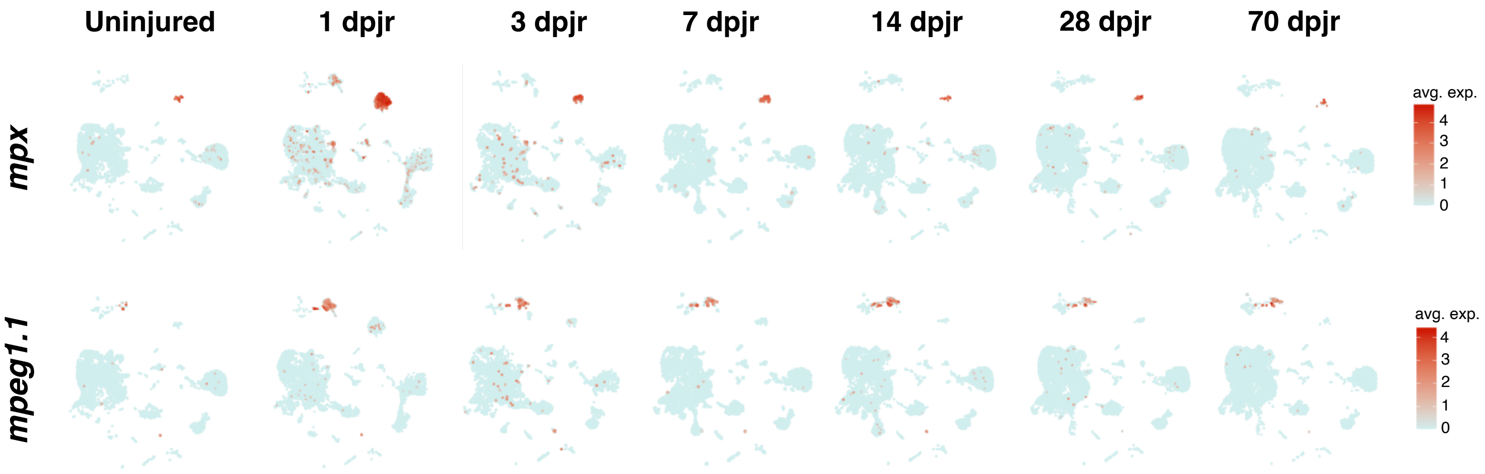
Uninjured
1 dpjr
3 dpjr
7 dpjr
14 dpjr
28 dpjr
70 dpjr
mpx
mpeg1.1
avg. exp.
4
3
2
1
0
avg. exp.
4
3
2
1
0

**Fig. S4: Dynamics of neutrophils and macrophages in joint regeneration.** FeaturePlot expression of *mpx* and *mpeg1.1* over time in superclusters.

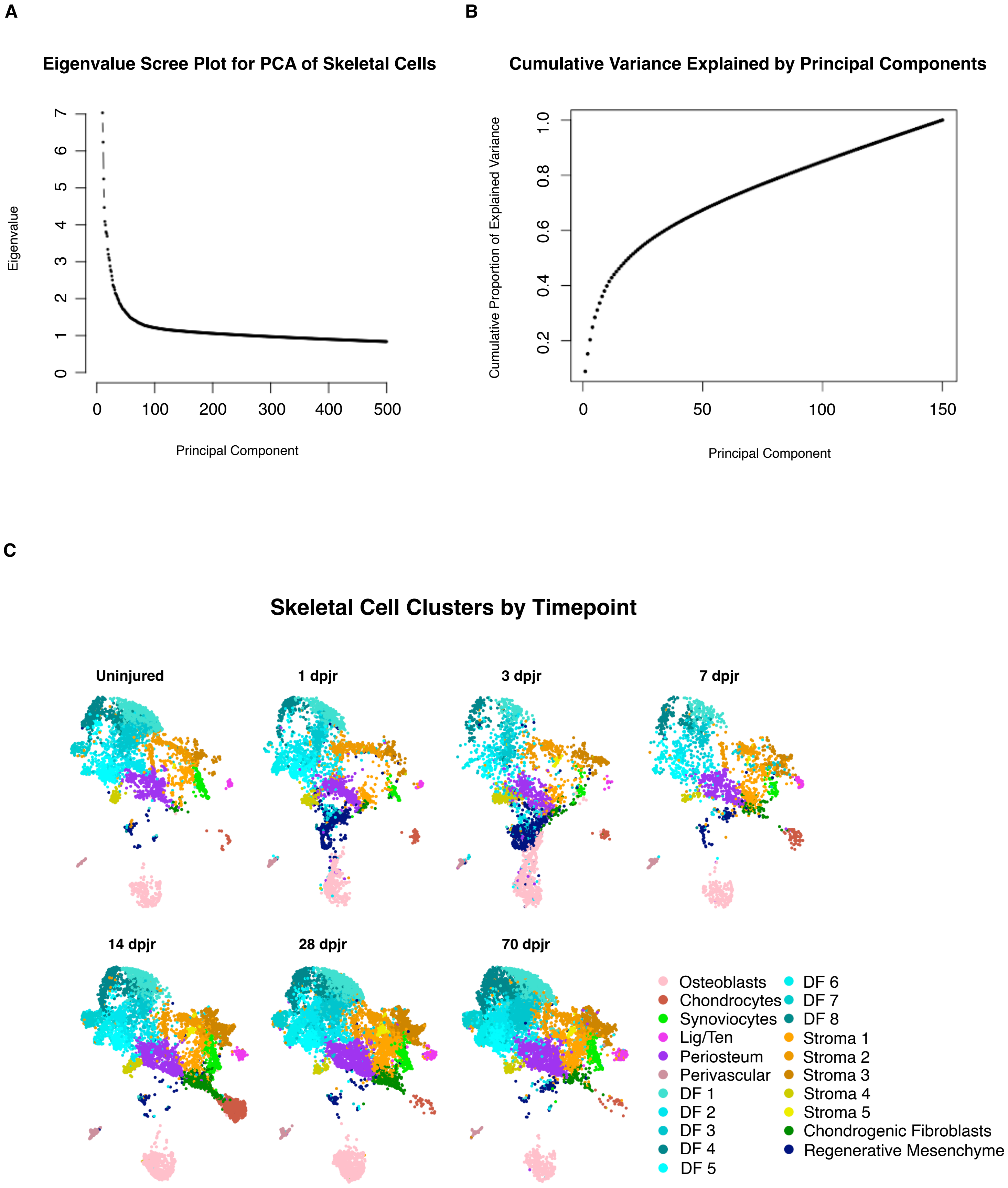
A
Eigenvalue Scree Plot for PCA of Skeletal Cells
Eigenvalue
0
1
2
3
4
5
6
7
0
100
200
300
400
500
Principal Component
B
Cumulative Variance Explained by Principal Components
Cumulative Proportion of Explained Variance
0.2
0.4
0.6
0.8
1.0
0
50
100
150
Principal Component
C
Skeletal Cell Clusters by Timepoint
Uninjured
1 dpjr
3 dpjr
7 dpjr
14 dpjr
28 dpjr
70 dpjr
Osteoblasts
Chondrocytes
Synoviocytes
Lig/Ten
Periosteum
Perivascular
DF 1
DF 2
DF 3
DF 4
DF 5
DF 6
DF 7
DF 8
Stroma 1
Stroma 2
Stroma 3
Stroma 4
Stroma 5
Chondrogenic Fibroblasts
Regenerative Mesenchyme

**Fig. S5: Generation of skeletal cell subclusters.** (**A**) Eigenvalue scree plot for principal component analysis of skeletal cells. (**B**) Cumulative variance explained by principal components. 150 principal components were used for clustering. (**C**) UMAPs of skeletal cells clustered at r=0.95 and split by time point.

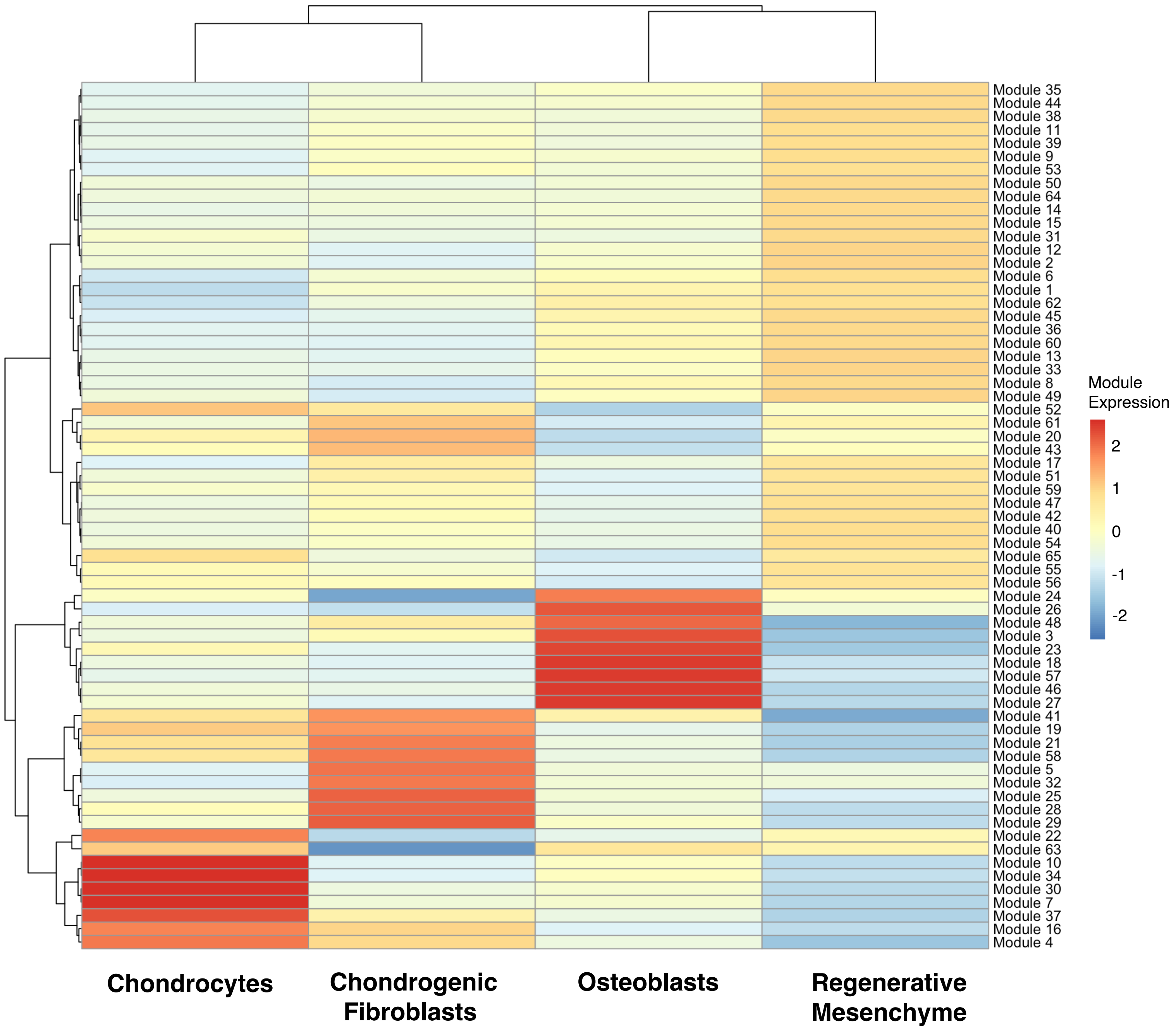
Module 35
Module 44
Module 38
Module 11
Module 39
Module 9
Module 53
Module 50
Module 64
Module 14
Module 15
Module 31
Module 12
Module 2
Module 6
Module 1
Module 62
Module 45
Module 36
Module 60
Module 13
Module 33
Module 8
Module 49
Module 52
Module 61
Module 20
Module 43
Module 17
Module 51
Module 59
Module 47
Module 42
Module 40
Module 54
Module 65
Module 55
Module 56
Module 24
Module 26
Module 48
Module 3
Module 23
Module 18
Module 57
Module 46
Module 27
Module 41
Module 19
Module 21
Module 58
Module 5
Module 32
Module 25
Module 28
Module 29
Module 22
Module 63
Module 10
Module 34
Module 30
Module 7
Module 37
Module 16
Module 4
Module Expression
2
1
0
-1
-2
Chondrocytes
Chondrogenic Fibroblasts
Osteoblasts
Regenerative Mesenchyme

**Fig. S6: Modules of co-regulated genes along the osteoblast and chondrocyte trajectories.** Heatmap of gene modules highly expressed in chondrocyte, chondrogenic fibroblast, osteoblast, and regenerative mesenchyme clusters.

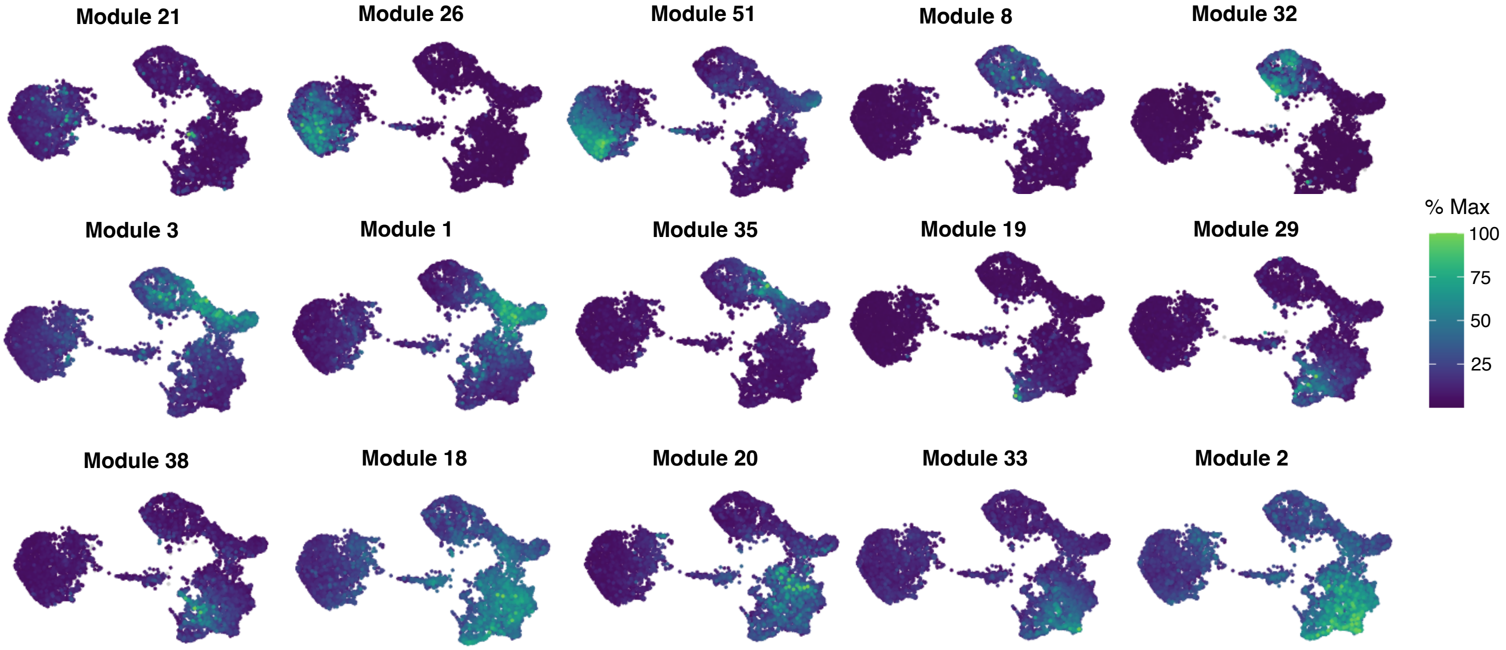
Module 21
Module 26
Module 51
Module 8
Module 32
Module 3
Module 1
Module 35
Module 19
Module 29
Module 38
Module 18
Module 20
Module 33
Module 2
% Max
100
75
50
25

**Fig. S7: Expression dynamics of co-regulated genes throughout joint regeneration**. Expression of modules of co-regulated genes along pseudotime, with some modules being specific to one cluster and other modules being shared by multiple cell states.

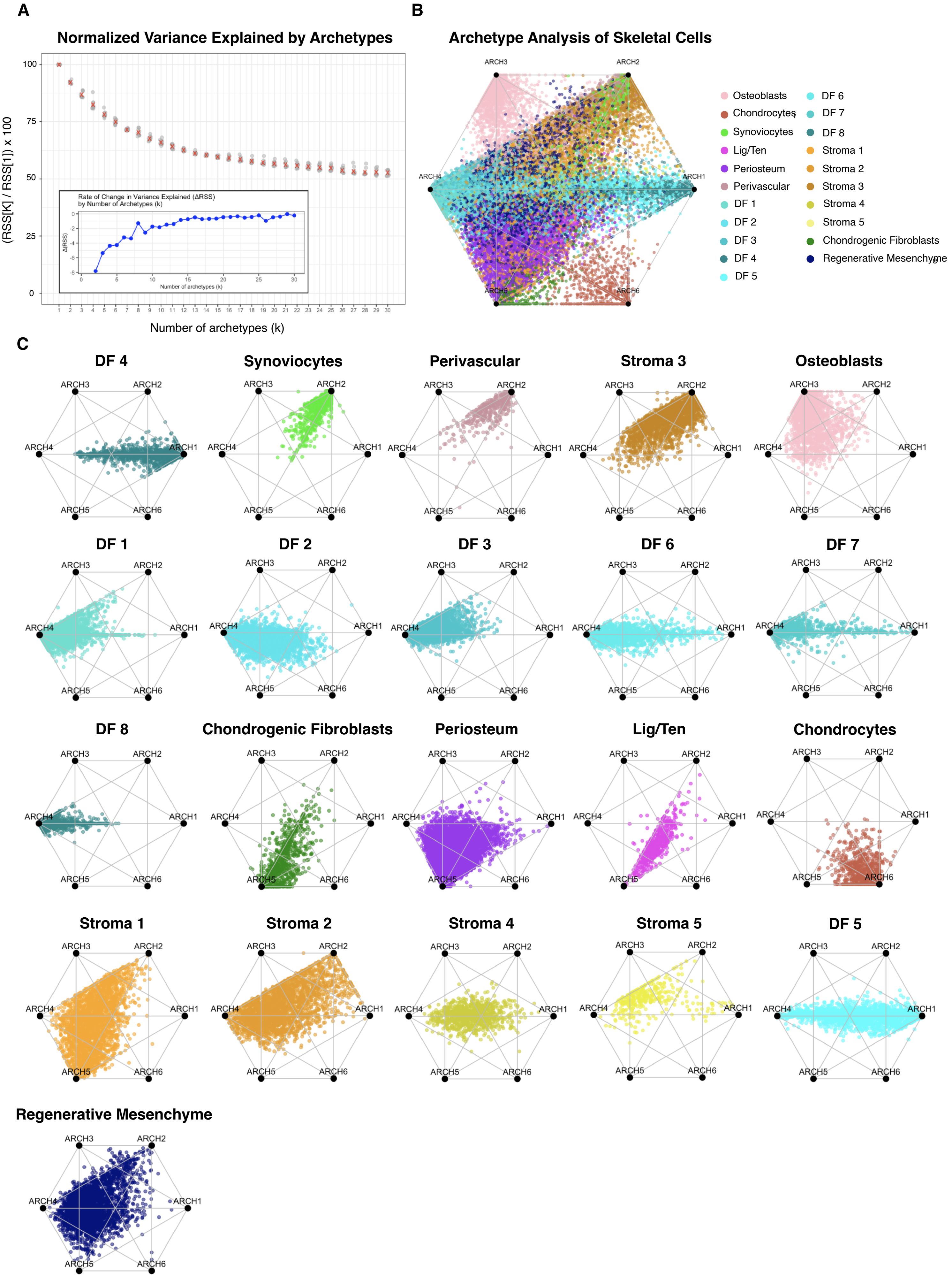
A
Normalized Variance Explained by Archetypes
(RSS[K] / RSS[1]) x 100
100
75
50
25
0
Number of archetypes (k)
Rate of Change in Variance Explained (ΔRSS)
by Number of Archetypes (k)
Δ(RSS)
Number of archetypes (k)
B
Archetype Analysis of Skeletal Cells
ARCH1
ARCH2
ARCH3
ARCH4
ARCH5
ARCH6
Osteoblasts
Chondrocytes
Synoviocytes
Lig/Ten
Periosteum
Perivascular
DF 1
DF 2
DF 3
DF 4
DF 5
DF 6
DF 7
DF 8
Stroma 1
Stroma 2
Stroma 3
Stroma 4
Stroma 5
Chondrogenic Fibroblasts
Regenerative Mesenchyme
C
DF 4
Synoviocytes
Perivascular
Stroma 3
Osteoblasts
DF 1
DF 2
DF 3
DF 6
DF 7
DF 8
Chondrogenic Fibroblasts
Periosteum
Lig/Ten
Chondrocytes
Stroma 1
Stroma 2
Stroma 4
Stroma 5
DF 5
Regenerative Mesenchyme

**Fig. S8: Archetype analysis of skeletal clusters**. (**A**) Variance explained by varying numbers of k archetypes as measured by Residual Sum of Squares (RSS). (**B**) Simplex plot of archetype analysis of skeletal clusters colored by cluster. (**C**) Individual skeletal cell clusters plotted onto archetypes.

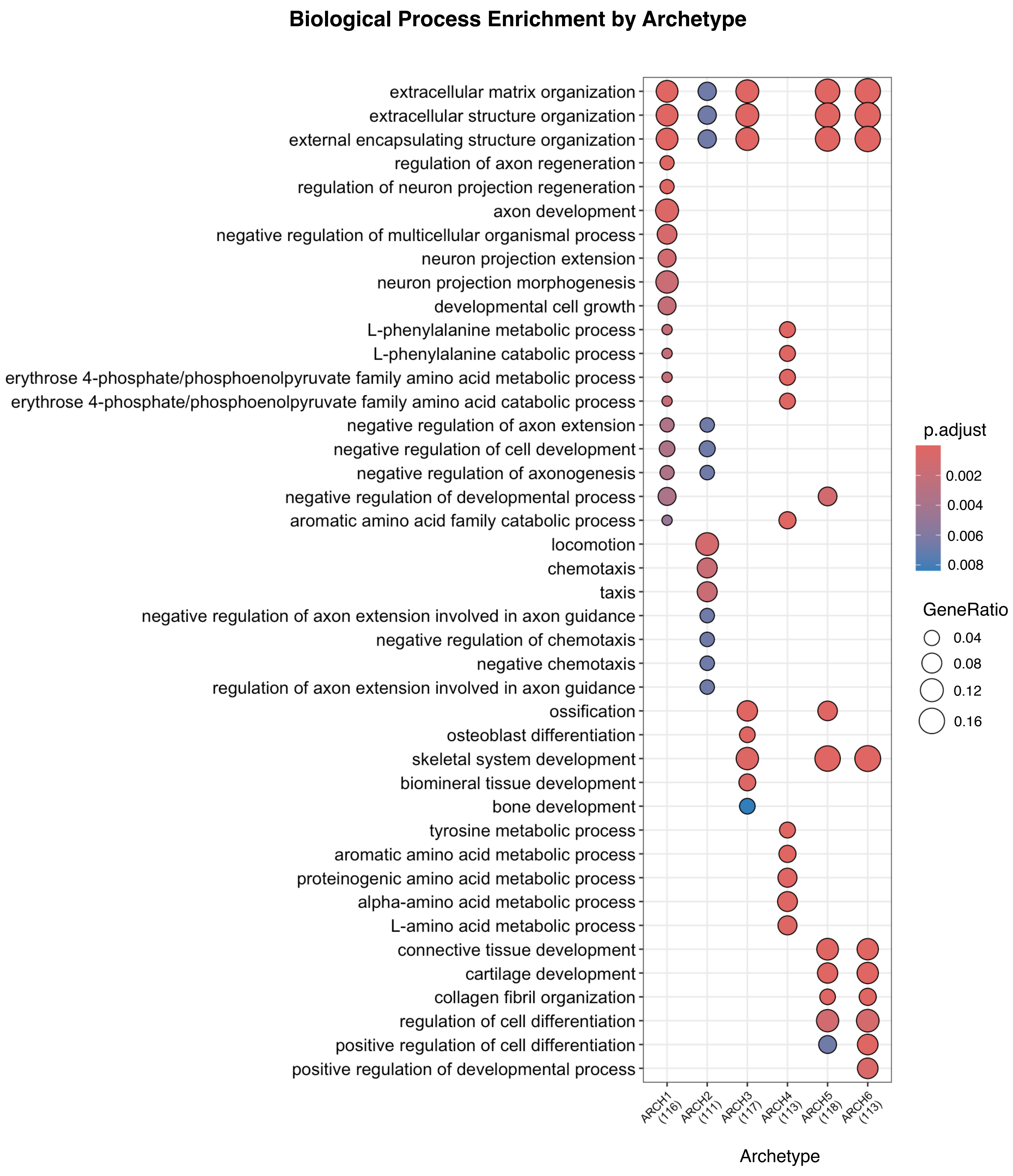
Biological Process Enrichment by Archetype
extracellular matrix organization
extracellular structure organization
external encapsulating structure organization
regulation of axon regeneration
regulation of neuron projection regeneration
axon development
negative regulation of multicellular organismal process
neuron projection extension
neuron projection morphogenesis
developmental cell growth
L-phenylalanine metabolic process
L-phenylalanine catabolic process
erythrose 4-phosphate/phosphoenolpyruvate family amino acid metabolic process
erythrose 4-phosphate/phosphoenolpyruvate family amino acid catabolic process
negative regulation of axon extension
negative regulation of cell development
negative regulation of axonogenesis
negative regulation of developmental process
aromatic amino acid family catabolic process
locomotion
chemotaxis
taxis
negative regulation of axon extension involved in axon guidance
negative regulation of chemotaxis
negative chemotaxis
regulation of axon extension involved in axon guidance
ossification
osteoblast differentiation
skeletal system development
biomineral tissue development
bone development
tyrosine metabolic process
aromatic amino acid metabolic process
proteinogenic amino acid metabolic process
alpha-amino acid metabolic process
L-amino acid metabolic process
connective tissue development
cartilage development
collagen fibril organization
regulation of cell differentiation
positive regulation of cell differentiation
positive regulation of developmental process
ARCH1 (116)
ARCH2 (111)
ARCH3 (117)
ARCH4 (113)
ARCH5 (118)
ARCH6 (113)
Archetype
p.adjust
0.002
0.004
0.006
0.008
GeneRatio
0.04
0.08
0.12
0.16

**Fig. S9: Gene ontology of biological process by archetype**. Biological processes enriched at each archetype as identified by gene ontology (GO) enrichment analysis.

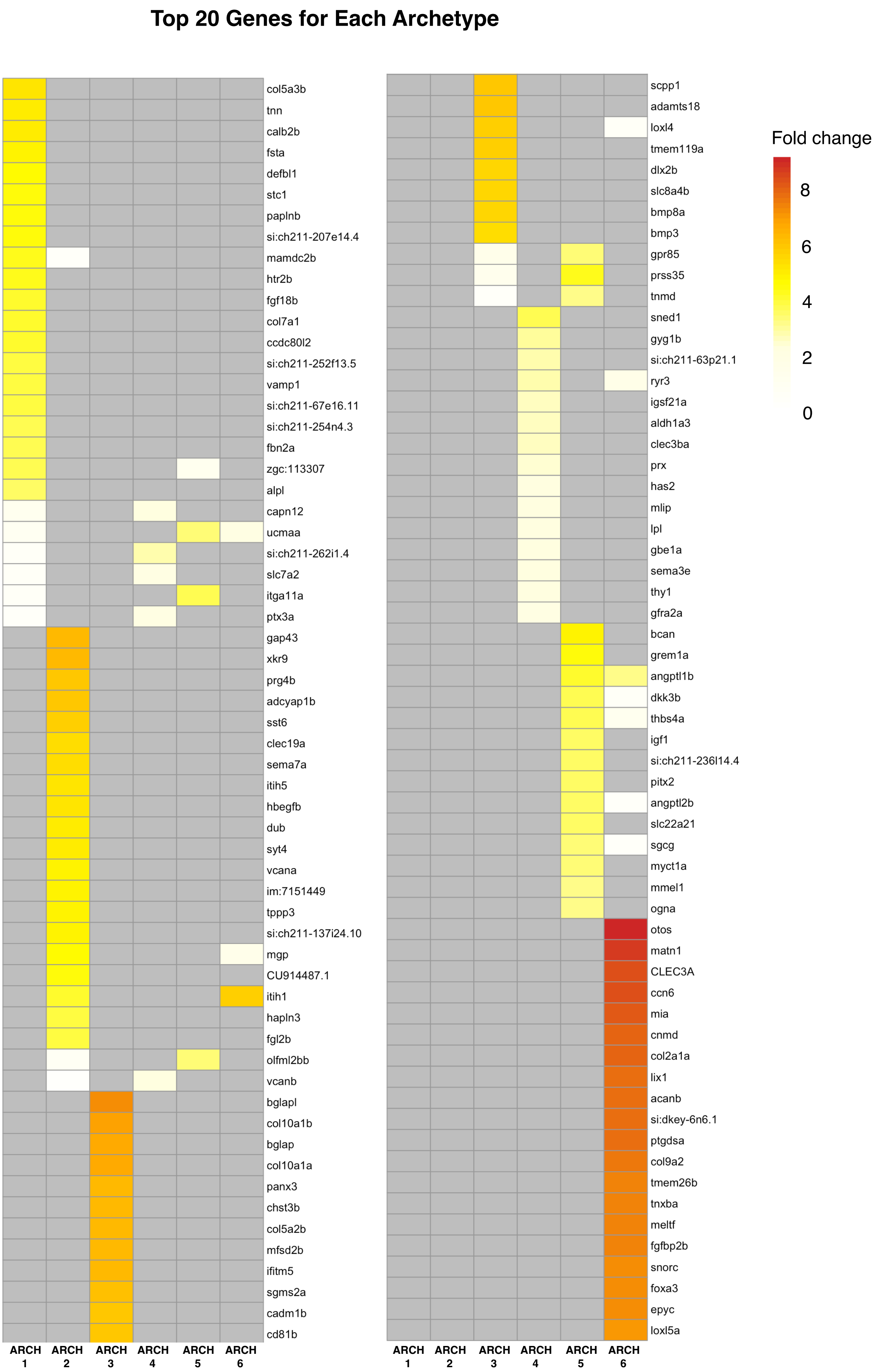

Top 20 Genes for Each Archetype
col5a3b
tnn
calb2b
fsta
defbl1
stc1
paplnb
si:ch211-207e14.4
mamdc2b
htr2b
fgf18b
col7a1
ccdc80l2
si:ch211-252f13.5
vamp1
si:ch211-67e16.11
si:ch211-254n4.3
fbn2a
zgc:113307
alpl
capn12
ucmaa
si:ch211-262i1.4
slc7a2
itga11a
ptx3a
gap43
xkr9
prg4b
adcyap1b
sst6
clec19a
sema7a
itih5
hbegfb
dub
syt4
vcana
im:7151449
tppp3
si:ch211-137i24.10
mgp
CU914487.1
itih1
hapln3
fgl2b
olfml2bb
vcanb
bglapl
col10a1b
bglap
col10a1a
panx3
chst3b
col5a2b
mfsd2b
ifitm5
sgms2a
cadm1b
cd81b
ARCH 1
ARCH 2
ARCH 3
ARCH 4
ARCH 5
ARCH 6
scpp1
adamts18
loxl4
tmem119a
dlx2b
slc8a4b
bmp8a
bmp3
gpr85
prss35
tnmd
sned1
gyg1b
si:ch211-63p21.1
ryr3
igsf21a
aldh1a3
clec3ba
prx
has2
mlip
lpl
gbe1a
sema3e
thy1
gfra2a
bcan
grem1a
angptl1b
dkk3b
thbs4a
igf1
si:ch211-236l14.4
pitx2
angptl2b
slc22a21
sgcg
myct1a
mmel1
ogna
otos
matn1
CLEC3A
ccn6
mia
cnmd
col2a1a
lix1
acanb
si:dkey-6n6.1
ptgdsa
col9a2
tmem26b
tnxba
meltf
fgfbp2b
snorc
foxa3
epyc
loxl5a
ARCH 1
ARCH 2
ARCH 3
ARCH 4
ARCH 5
ARCH 6
Fold change
8
6
4
2
0

**Fig. S10: Top genes defining each archetype**. Heatmap of top 20 genes for each archetype in terms of average log2 fold change.

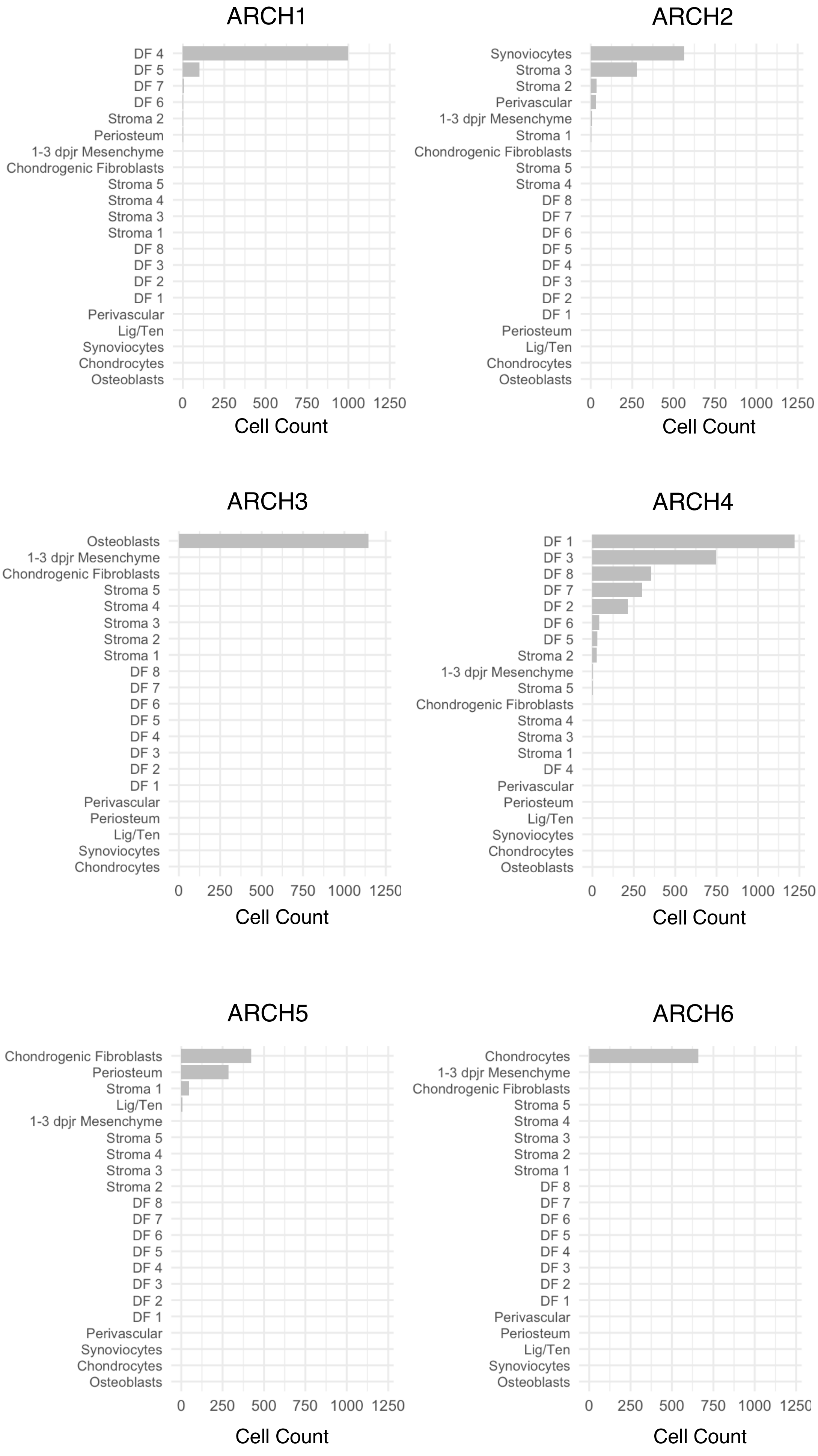
ARCH1
DF 4
DF 5
DF 7
DF 6
Stroma 2
Periosteum
1-3 dpjr Mesenchyme
Chondrogenic Fibroblasts
Stroma 5
Stroma 4
Stroma 3
Stroma 1
DF 8
DF 3
DF 2
DF 1
Perivascular
Lig/Ten
Synoviocytes
Chondrocytes
Osteoblasts
0 250 500 750 1000 1250
Cell Count
ARCH2
Synoviocytes
Stroma 3
Stroma 2
Perivascular
1-3 dpjr Mesenchyme
Stroma 1
Chondrogenic Fibroblasts
Stroma 5
Stroma 4
DF 8
DF 7
DF 6
DF 5
DF 4
DF 3
DF 2
DF 1
Periosteum
Lig/Ten
Chondrocytes
Osteoblasts
0 250 500 750 1000 1250
Cell Count
ARCH3
Osteoblasts
1-3 dpjr Mesenchyme
Chondrogenic Fibroblasts
Stroma 5
Stroma 4
Stroma 3
Stroma 2
Stroma 1
DF 8
DF 7
DF 6
DF 5
DF 4
DF 3
DF 2
DF 1
Perivascular
Periosteum
Lig/Ten
Synoviocytes
Chondrocytes
0 250 500 750 1000 1250
Cell Count
ARCH4
DF 1
DF 3
DF 8
DF 7
DF 2
DF 6
DF 5
Stroma 2
1-3 dpjr Mesenchyme
Stroma 5
Chondrogenic Fibroblasts
Stroma 4
Stroma 3
Stroma 1
DF 4
Perivascular
Periosteum
Lig/Ten
Synoviocytes
Chondrocytes
Osteoblasts
0 250 500 750 1000 1250
Cell Count
ARCH5
Chondrogenic Fibroblasts
Periosteum
Stroma 1
Lig/Ten
1-3 dpjr Mesenchyme
Stroma 5
Stroma 4
Stroma 3
Stroma 2
DF 8
DF 7
DF 6
DF 5
DF 4
DF 3
DF 2
DF 1
Perivascular
Synoviocytes
Chondrocytes
Osteoblasts
0 250 500 750 1000 1250
Cell Count
ARCH6
Chondrocytes
1-3 dpjr Mesenchyme
Chondrogenic Fibroblasts
Stroma 5
Stroma 4
Stroma 3
Stroma 2
Stroma 1
DF 8
DF 7
DF 6
DF 5
DF 4
DF 3
DF 2
DF 1
Perivascular
Periosteum
Lig/Ten
Synoviocytes
Osteoblasts
0 250 500 750 1000 1250
Cell Count

**Fig. S11: Cell type composition of cells close to archetypes**. Absolute count of skeletal cell types mapping close to each archetype.

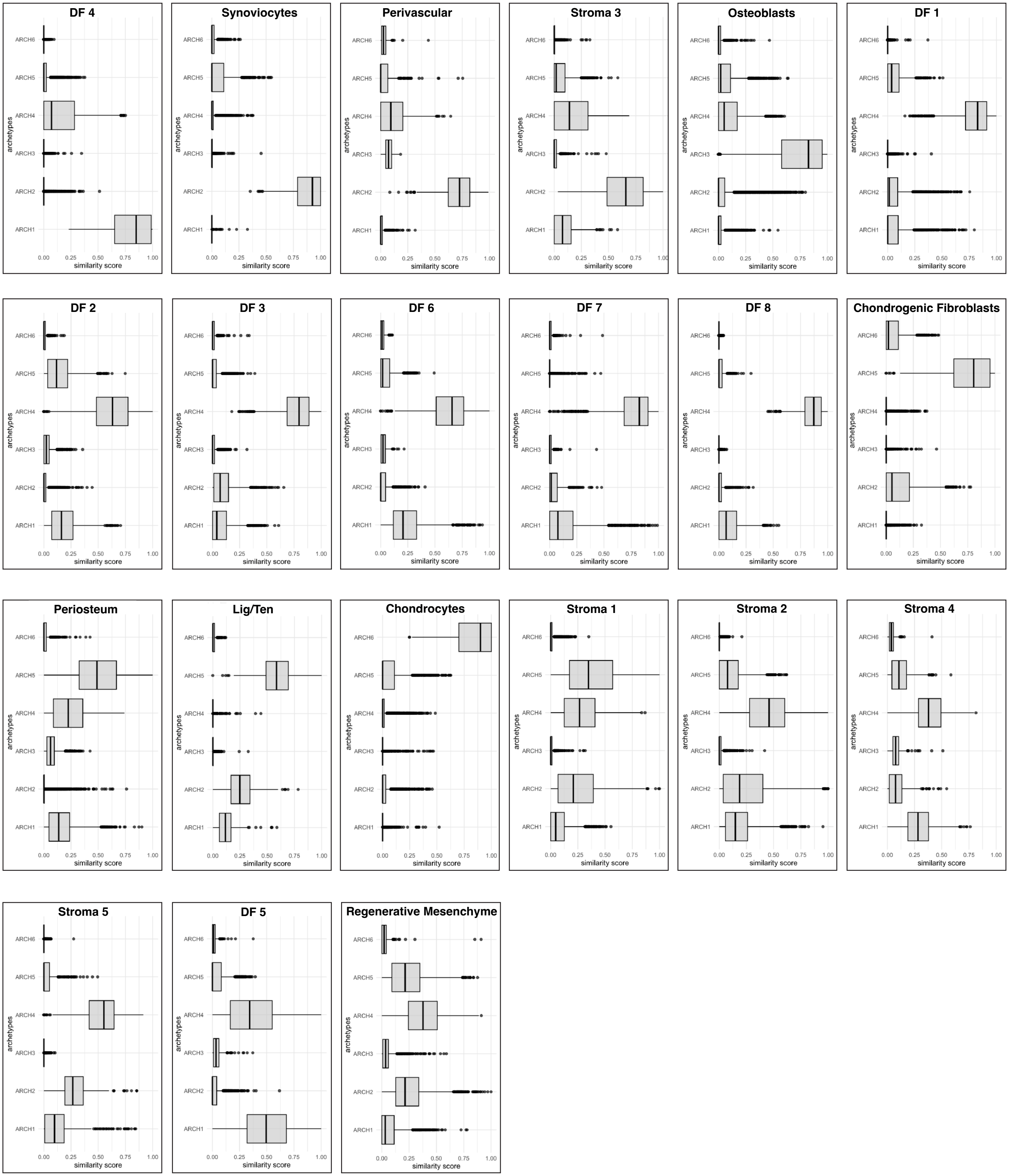
DF 4
Synoviocytes
Perivascular
Stroma 3
Osteoblasts
DF 1
DF 2
DF 3
DF 6
DF 7
DF 8
Chondrogenic Fibroblasts
Periosteum
Lig/Ten
Chondrocytes
Stroma 1
Stroma 2
Stroma 4
Stroma 5
DF 5
Regenerative Mesenchyme
ARCH6
ARCH5
ARCH4
ARCH3
ARCH2
ARCH1
archetypes
0.00
0.25
0.50
0.75
1.00
similarity score

**Fig. S12: Distribution of similarity scores of skeletal cell clusters to archetypes**. Distribution of similarity of each skeletal cluster to each archetype. Each cell has a similarity score to each archetype.

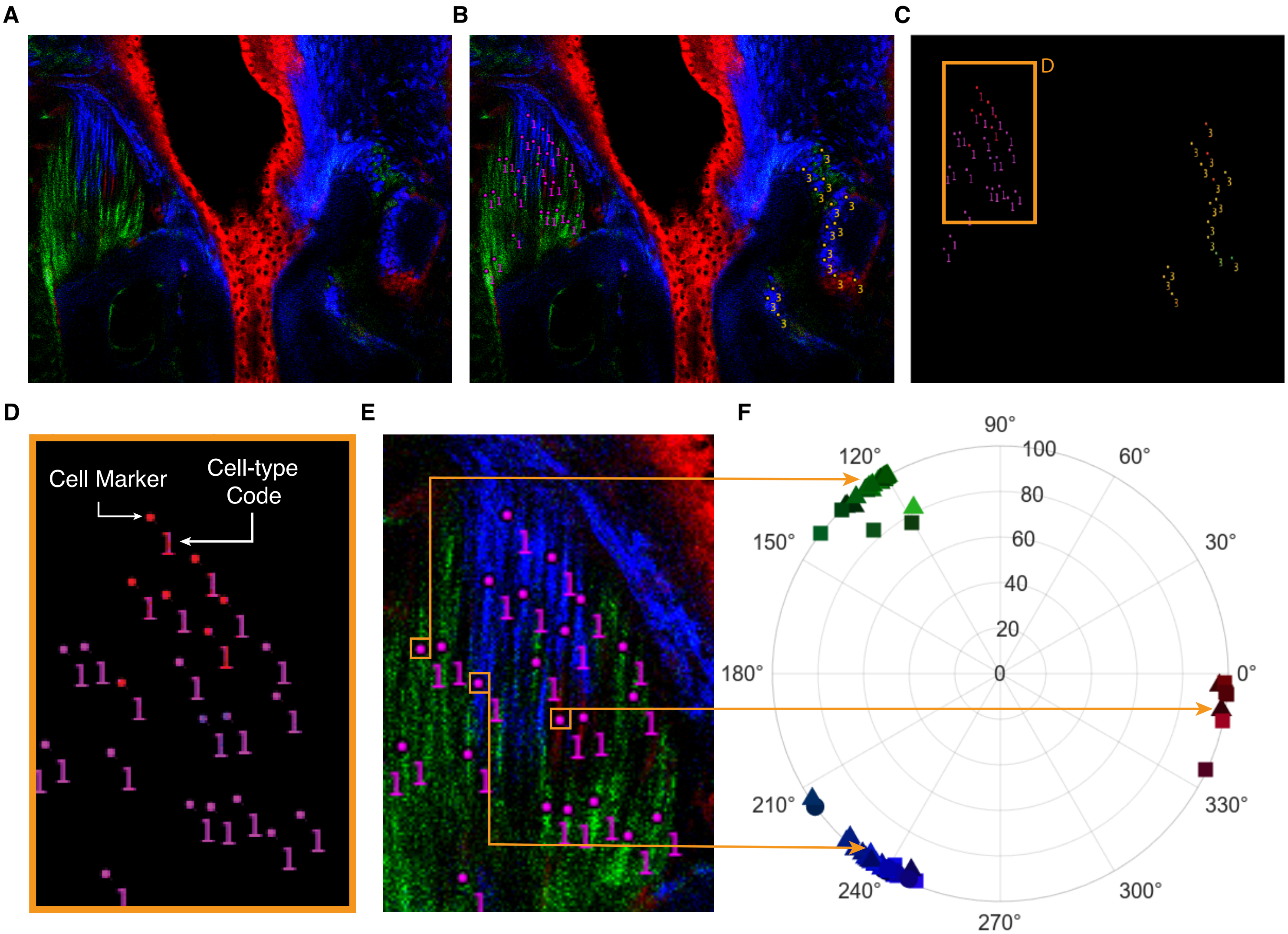
A
B
C
D
E
F
Cell Marker
Cell-type Code
90°
120°
60°
150°
30°
180°
0°
210°
330°
240°
300°
270°
100
80
60
40
20
0

**Fig. S13: Overview of analysis pipeline in developed clonality application.** (**A** to **C**) Raw confocal images of the tissue (A) and its corresponding labeled image (B) are used as inputs into the application. The difference between the two images yields the (C) 'cell label' only image. (**D**) This image is scanned to identify cell markers and their corresponding cell-type code. (**E** and **F**) RGB values are extracted from the pixels within each cell marker (E) and plotted on a polar plot as hue and saturation (F). 1 = Ligamentocytes, 2 = Adipocytes, 3 = Chondrocytes.

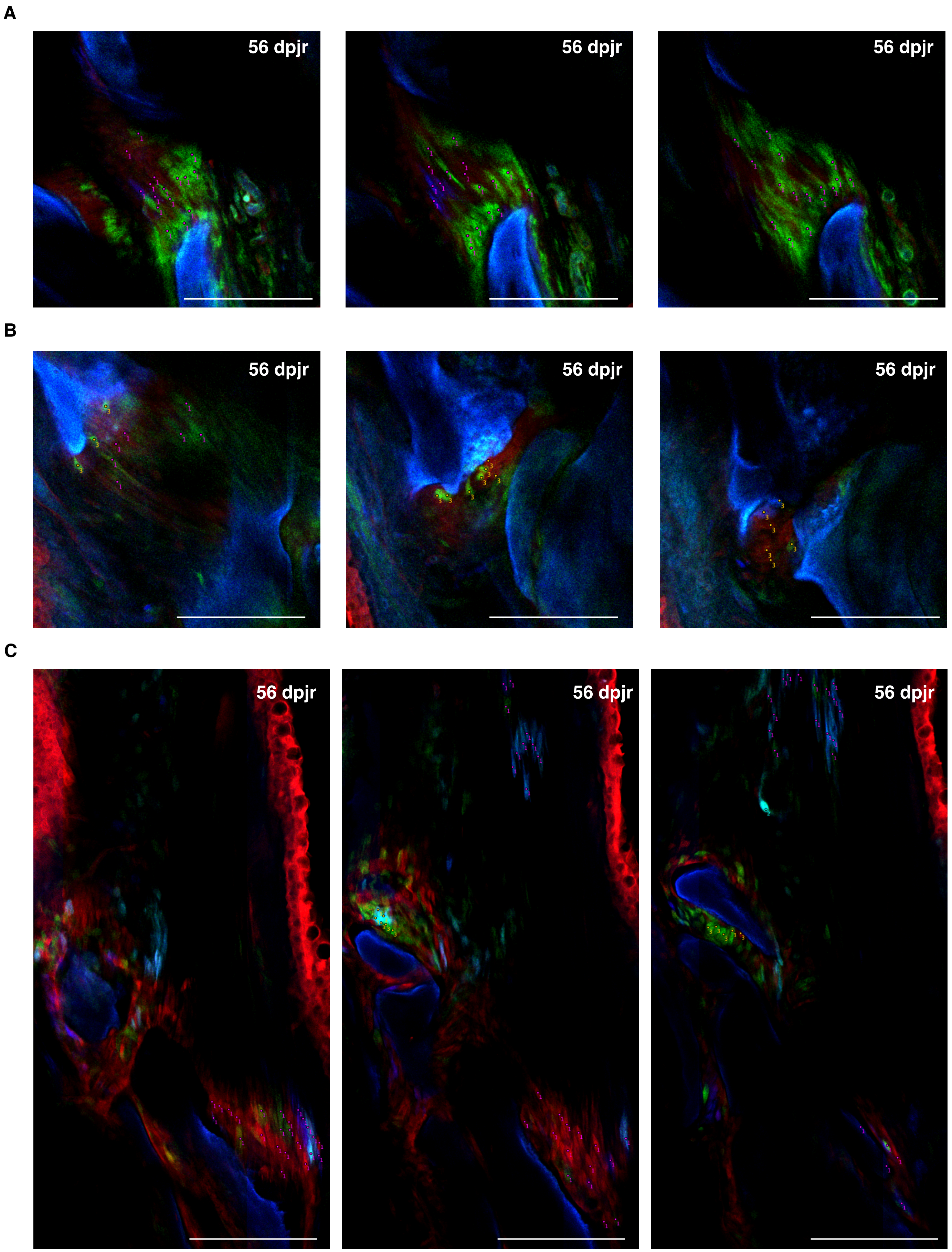
A
56 dpjr
56 dpjr
56 dpjr
B
56 dpjr
56 dpjr
56 dpjr
C
56 dpjr
56 dpjr
56 dpjr

**Fig. S14: Representative cell-type code and cell markers for single-clone lineage tracing analysis**. (**A** to **C**) Cell-type code and cell markers used to label cells in representative z-slices for the sample corresponding to Fig. 5G (A); the sample corresponding to Fig. 5H (B); and the sample corresponding to Fig. 5K (C). 1 = Ligamentocytes, 2 = Adipocytes, 3 = Chondrocytes. Scale bars = 125µm (A to C).

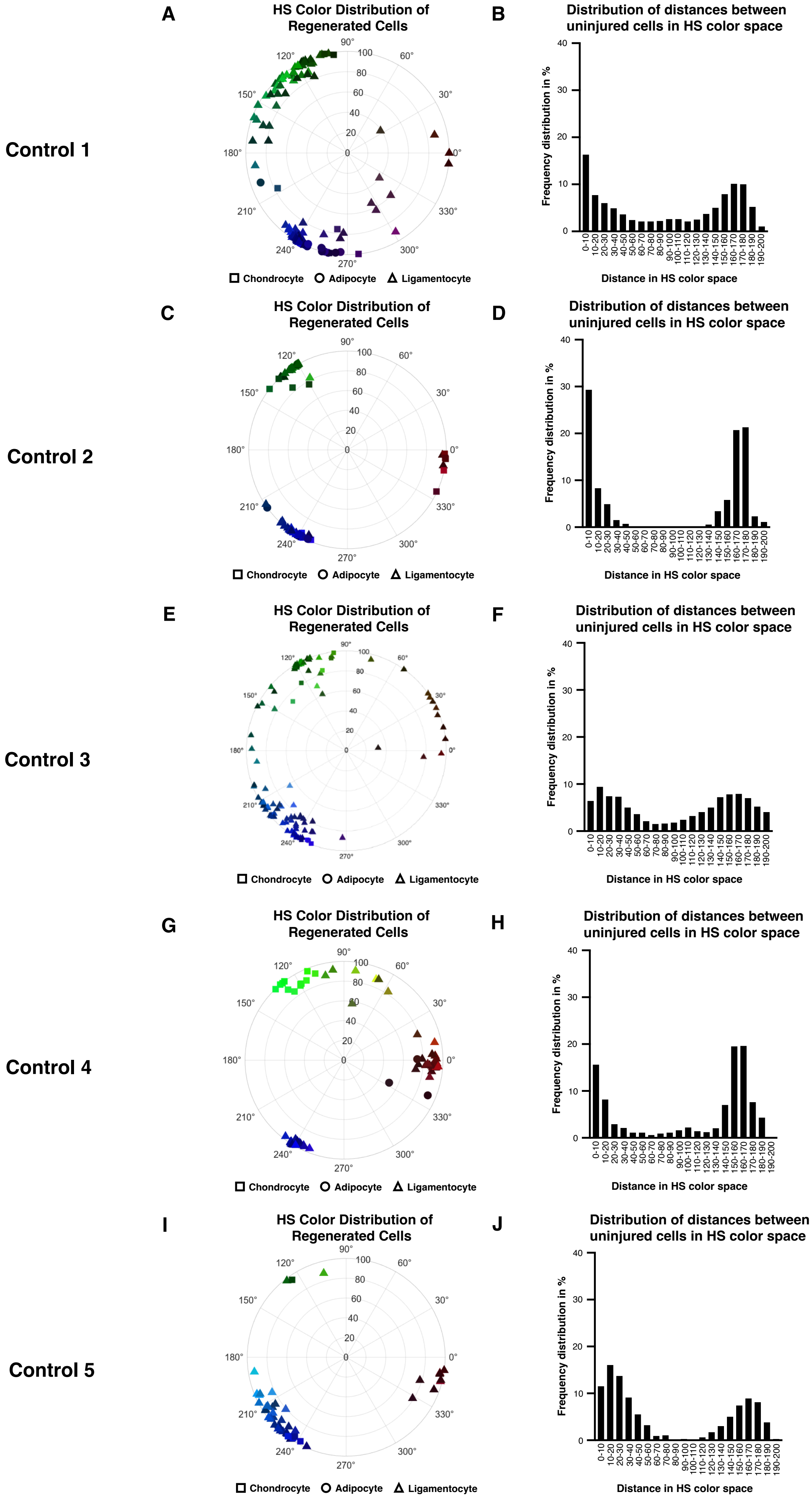
A
HS Color Distribution of Regenerated Cells
B
Distribution of distances between uninjured cells in HS color space
Control 1
C
HS Color Distribution of Regenerated Cells
D
Distribution of distances between uninjured cells in HS color space
Control 2
E
HS Color Distribution of Regenerated Cells
F
Distribution of distances between uninjured cells in HS color space
Control 3
G
HS Color Distribution of Regenerated Cells
H
Distribution of distances between uninjured cells in HS color space
Control 4
I
HS Color Distribution of Regenerated Cells
J
Distribution of distances between uninjured cells in HS color space
Control 5
Chondrocyte
Adipocyte
Ligamentocyte
Frequency distribution in %
Distance in HS color space
0-10
10-20
20-30
30-40
40-50
50-60
60-70
70-80
80-90
90-100
100-110
110-120
120-130
130-140
140-150
150-160
160-170
170-180
180-190
190-200

**Fig. S15: Clonality of cells in the uninjured joint**. (**A** to **J**) Representative graphs of HS color plots and frequency distribution of joint cells in 5 uninjured control samples.

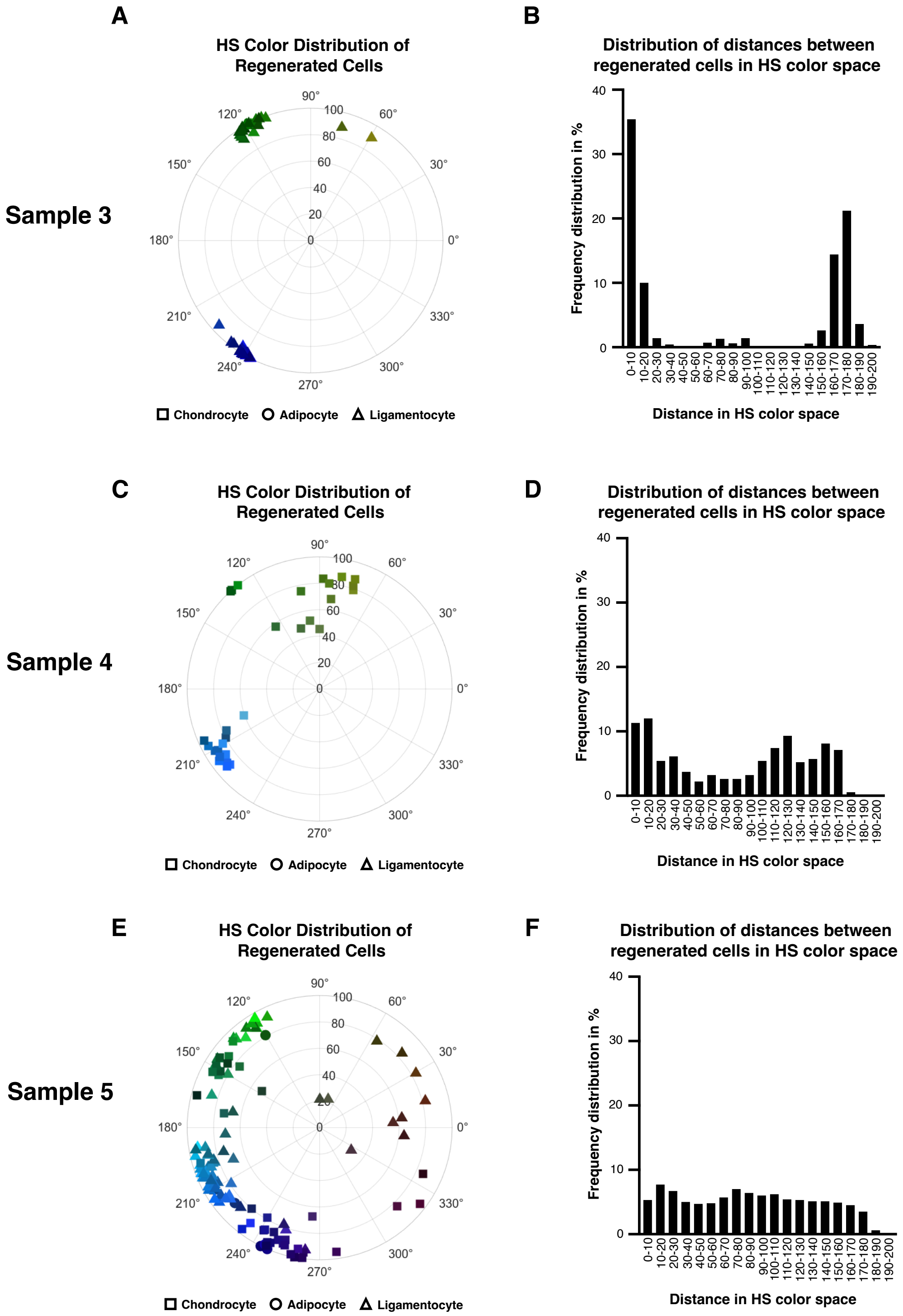

Sample 3
Sample 4
Sample 5
A
HS Color Distribution of Regenerated Cells
90°
60°
30°
0°
330°
300°
270°
240°
210°
180°
150°
120°
0
20
40
60
80
100
Chondrocyte
Adipocyte
Ligamentocyte
B
Distribution of distances between regenerated cells in HS color space
Frequency distribution in %
0
10
20
30
40
0-10
10-20
20-30
30-40
40-50
50-60
60-70
70-80
80-90
90-100
100-110
110-120
120-130
130-140
140-150
150-160
160-170
170-180
180-190
190-200
Distance in HS color space
C
HS Color Distribution of Regenerated Cells
D
Distribution of distances between regenerated cells in HS color space
E
HS Color Distribution of Regenerated Cells
F
Distribution of distances between regenerated cells in HS color space

**Fig. S16: Clonality of cells in the regenerated joint**. (**A** to **F**) Additional representative graphs of HS color plots and frequency distribution of joint cells in 3 regenerated samples at 56 dpjr.

**Table S1: scRNAseq cell counts by cluster and sample.**

**Table S2: Marker genes of skeletal scRNAseq clusters.**

**Table S3: Top genes and biological processes enriched at each archetype.**
