## Supplemental Table 1 for "Dynamic cell fate plasticity and tissue integration drive functional synovial joint regeneration"

| | Periosteum | DF 1 | DF 2 | DF 3 | Osteoblasts | Stroma 1 | Regenerative Mesenchyme | Stroma 2 | DF 4 | Stroma 3 | DF 5 | DF 6 | Chondrocytes | Chondrogenic Fibroblasts | DF 7 | Synoviocytes | Stroma 4 | DF 8 | Lig/Ten | Perivascular | Stroma 5 |
|---|---|---|---|---|---|---|---|---|---|---|---|---|---|---|---|---|---|---|---|---|---|
| **Uninjured** | 377 | 1104 | 327 | 547 | 237 | 158 | 49 | 315 | 281 | 68 | 350 | 60 | 19 | 18 | 83 | 119 | 79 | 250 | 25 | 13 | 8 |
| **1 dpjr** | 479 | 380 | 380 | 409 | 468 | 164 | 984 | 356 | 157 | 118 | 309 | 168 | 47 | 21 | 12 | 84 | 145 | 6 | 48 | 42 | 0 |
| **3 dpjr** | 429 | 137 | 406 | 274 | 752 | 156 | 1083 | 222 | 123 | 240 | 82 | 20 | 40 | 70 | 2 | 39 | 267 | 9 | 7 | 59 | 34 |
| **7 dpjr** | 327 | 129 | 159 | 176 | 177 | 254 | 90 | 150 | 109 | 128 | 90 | 48 | 54 | 97 | 2 | 53 | 43 | 43 | 31 | 49 | 9 |
| **14dpjr** | 675 | 616 | 558 | 298 | 623 | 352 | 92 | 384 | 409 | 498 | 252 | 203 | 1107 | 495 | 368 | 246 | 163 | 68 | 110 | 133 | 61 |
| **28 dpjr** | 1071 | 941 | 708 | 603 | 664 | 810 | 87 | 540 | 545 | 608 | 324 | 482 | 32 | 340 | 203 | 283 | 96 | 243 | 99 | 75 | 163 |
| **70 dpjr** | 931 | 823 | 886 | 1002 | 283 | 624 | 114 | 427 | 712 | 368 | 613 | 422 | 26 | 123 | 463 | 198 | 92 | 243 | 131 | 22 | 83 |
| Total | 4289 | 4130 | 3424 | 3309 | 3204 | 2518 | 2499 | 2394 | 2336 | 2028 | 2020 | 1403 | 1325 | 1164 | 1133 | 1022 | 885 | 862 | 451 | 393 | 358 |
