## Supplemental Table 2 for "Dynamic cell fate plasticity and tissue integration drive functional synovial joint regeneration"

| | p_val | avg_log2FC | pct.1 | pct.2 | p_val_adj | cluster | gene |
|---|---|---|---|---|---|---|---|
| aspn | 0 | 1.9357273 | 0.856 | 0.352 | 0 | Periosteum | aspn |
| si:ch211-76l23.4 | 0 | 3.5115432 | 0.606 | 0.14 | 0 | Periosteum | si:ch211-76l23.4 |
| clu | 0 | 1.6295652 | 0.782 | 0.338 | 0 | Periosteum | clu |
| postnb | 0 | 1.9100718 | 0.85 | 0.411 | 0 | Periosteum | postnb |
| tpm1 | 0 | 1.9546942 | 0.717 | 0.294 | 0 | Periosteum | tpm1 |
| mmel1 | 0 | 6.1146145 | 0.422 | 0.018 | 0 | Periosteum | mmel1 |
| ogna | 0 | 2.1489831 | 0.545 | 0.149 | 0 | Periosteum | ogna |
| fgfr3 | 0 | 1.4172478 | 0.607 | 0.232 | 0 | Periosteum | fgfr3 |
| si:busm1-57f23.1 | 0 | 1.1980899 | 0.906 | 0.537 | 0 | Periosteum | si:busm1-57f23.1 |
| si:dkey-12l12.1 | 0 | 1.4165112 | 0.871 | 0.547 | 0 | Periosteum | si:dkey-12l12.1 |
| tnc | 0 | 0.9879848 | 0.76 | 0.438 | 0 | Periosteum | tnc |
| col11a1b | 0 | 1.0611046 | 0.82 | 0.513 | 0 | Periosteum | col11a1b |
| cdh11 | 0 | 1.0854413 | 0.863 | 0.559 | 0 | Periosteum | cdh11 |
| mxra8b | 0 | 1.2389667 | 0.676 | 0.378 | 0 | Periosteum | mxra8b |
| itga11a | 0 | 2.3751738 | 0.373 | 0.081 | 0 | Periosteum | itga11a |
| angptl2b | 0 | 2.2700523 | 0.395 | 0.112 | 0 | Periosteum | angptl2b |
| col27a1b | 0 | 1.206103 | 0.592 | 0.309 | 0 | Periosteum | col27a1b |
| CABZ01080702.1 | 0 | 1.2696962 | 0.551 | 0.274 | 0 | Periosteum | CABZ01080702.1 |
| kctd12.2 | 0 | 1.3731351 | 0.464 | 0.2 | 0 | Periosteum | kctd12.2 |
| dkk3b | 0 | 1.184182 | 0.483 | 0.222 | 0 | Periosteum | dkk3b |
| col14a1a | 0 | 3.147858 | 0.308 | 0.048 | 0 | Periosteum | col14a1a |
| pcolceb | 0 | 0.8600029 | 0.403 | 0.147 | 0 | Periosteum | pcolceb |
| cpxm1a | 0 | 1.6394356 | 0.406 | 0.153 | 0 | Periosteum | cpxm1a |
| mmp2 | 0 | 1.5922678 | 0.971 | 0.719 | 0 | Periosteum | mmp2 |
| ANPEP | 0 | 1.3101332 | 0.435 | 0.185 | 0 | Periosteum | ANPEP |
| vasnb | 0 | 1.7504466 | 0.365 | 0.119 | 0 | Periosteum | vasnb |
| tnmd | 0 | 1.5334673 | 0.373 | 0.127 | 0 | Periosteum | tnmd |
| hyal4 | 0 | 1.9962184 | 0.324 | 0.081 | 0 | Periosteum | hyal4 |
| bcan | 0 | 2.4691115 | 0.315 | 0.078 | 0 | Periosteum | bcan |
| s1pr1 | 0 | 2.2575179 | 0.306 | 0.069 | 0 | Periosteum | s1pr1 |
| lmod1b | 0 | 2.1949069 | 0.28 | 0.076 | 0 | Periosteum | lmod1b |
| nog2 | 0 | 2.2346437 | 0.263 | 0.064 | 0 | Periosteum | nog2 |
| aqp3b | 0 | 1.8936588 | 0.252 | 0.063 | 0 | Periosteum | aqp3b |
| prss35 | 0 | 2.1359912 | 0.262 | 0.074 | 0 | Periosteum | prss35 |
| nog3 | 0 | 1.8277395 | 0.241 | 0.064 | 0 | Periosteum | nog3 |
| timp2a | 0 | 0.8731189 | 0.906 | 0.738 | 0 | Periosteum | timp2a |
| fgfr4 | 0 | 2.1283876 | 0.224 | 0.06 | 0 | Periosteum | fgfr4 |
| si:ch1073-459j12.1 | 0 | 0.8897974 | 0.958 | 0.795 | 0 | Periosteum | si:ch1073-459j12.1 |
| col8a2 | 0 | 1.8442963 | 0.206 | 0.052 | 0 | Periosteum | col8a2 |
| ca10a | 0 | 2.3432835 | 0.187 | 0.036 | 0 | Periosteum | ca10a |
| col1a1a | 0 | 0.8755514 | 0.998 | 0.978 | 0 | Periosteum | col1a1a |
| sparc | 0 | 0.6975532 | 0.997 | 0.978 | 0 | Periosteum | sparc |
| col1a1b | 0 | 0.8583797 | 1 | 0.985 | 0 | Periosteum | col1a1b |
| col1a2 | 0 | 0.8741373 | 0.999 | 0.99 | 0 | Periosteum | col1a2 |
| lum | 2E-306 | 0.7528954 | 0.923 | 0.74 | 6.8E-302 | Periosteum | lum |
| col5a2a | 2E-287 | 0.7490578 | 0.921 | 0.778 | 5.2E-283 | Periosteum | col5a2a |
| si:dkey-21e2.8 | 1E-278 | 3.1511892 | 0.1 | 0.014 | 4.4E-274 | Periosteum | si:dkey-21e2.8 |

| | | | | | | | |
|---|---|---|---|---|---|---|---|
| lrrc17 | 9E-272 | 0.9624026 | 0.637 | 0.37 | 2.8E-267 | Periosteum | lrrc17 |
| ecm2 | 7E-268 | 1.0800945 | 0.529 | 0.272 | 2.4E-263 | Periosteum | ecm2 |
| rps14 | 9E-267 | 0.3802095 | 0.999 | 0.993 | 3E-262 | Periosteum | rps14 |
| ccdc3b | 1E-266 | 1.5171447 | 0.292 | 0.106 | 4.8E-262 | Periosteum | ccdc3b |
| si:ch211-160j14.3 | 1E-259 | 1.1100812 | 0.542 | 0.289 | 4E-255 | Periosteum | si:ch211-160j14.3 |
| selenop | 4E-253 | 0.7282886 | 0.892 | 0.724 | 1.2E-248 | Periosteum | selenop |
| esr2a | 1E-252 | 2.5099731 | 0.103 | 0.017 | 3.6E-248 | Periosteum | esr2a |
| rspo3 | 1E-249 | 1.760745 | 0.186 | 0.051 | 3.3E-245 | Periosteum | rspo3 |
| krt18a.1 | 4E-243 | 0.6177986 | 0.961 | 0.828 | 1.4E-238 | Periosteum | krt18a.1 |
| rpl34 | 2E-240 | 0.3865049 | 0.996 | 0.984 | 5.5E-236 | Periosteum | rpl34 |
| selenon | 1E-237 | 1.4482412 | 0.326 | 0.138 | 3.5E-233 | Periosteum | selenon |
| rplp2 | 2E-236 | 0.3668941 | 0.996 | 0.986 | 6.1E-232 | Periosteum | rplp2 |
| zgc:92161 | 5E-235 | 1.2410001 | 0.382 | 0.173 | 1.6E-230 | Periosteum | zgc:92161 |
| ptgs2a | 2E-232 | 1.5022159 | 0.288 | 0.113 | 5.8E-228 | Periosteum | ptgs2a |
| acta2 | 9E-229 | 3.3091348 | 0.127 | 0.028 | 3.1E-224 | Periosteum | acta2 |
| tnfaip8l3 | 5E-226 | 2.1409909 | 0.144 | 0.036 | 1.6E-221 | Periosteum | tnfaip8l3 |
| eef1g | 2E-225 | 0.4605027 | 0.983 | 0.949 | 5.5E-221 | Periosteum | eef1g |
| rpl32 | 5E-223 | 0.3639379 | 0.997 | 0.988 | 1.8E-218 | Periosteum | rpl32 |
| ppiaa | 3E-222 | 0.4094057 | 0.987 | 0.957 | 8.6E-218 | Periosteum | ppiaa |
| rpsa | 2E-218 | 0.3574949 | 0.996 | 0.984 | 7.9E-214 | Periosteum | rpsa |
| fndc1 | 3E-218 | 1.1146891 | 0.34 | 0.148 | 9.7E-214 | Periosteum | fndc1 |
| uba52 | 1E-215 | 0.3212229 | 0.998 | 0.991 | 4.6E-211 | Periosteum | uba52 |
| hspb6 | 5E-215 | 2.5917466 | 0.1 | 0.019 | 1.6E-210 | Periosteum | hspb6 |
| lmo7a | 1E-214 | 1.4558247 | 0.314 | 0.136 | 3.6E-210 | Periosteum | lmo7a |
| rps2 | 7E-212 | 0.3737684 | 0.996 | 0.99 | 2.2E-207 | Periosteum | rps2 |
| angptl1b | 1E-211 | 1.7812379 | 0.176 | 0.053 | 3.4E-207 | Periosteum | angptl1b |
| cald1a | 2E-211 | 0.8102719 | 0.793 | 0.603 | 7.7E-207 | Periosteum | cald1a |
| dlx5a | 5E-209 | 1.0328424 | 0.498 | 0.276 | 1.6E-204 | Periosteum | dlx5a |
| si:dkey-151g10.6 | 5E-209 | 0.3321516 | 1 | 0.999 | 1.6E-204 | Periosteum | si:dkey-151g10.6 |
| id2b | 2E-208 | 1.1314806 | 0.367 | 0.171 | 8E-204 | Periosteum | id2b |
| lgals2a | 2E-207 | 0.3949743 | 0.845 | 0.618 | 6.8E-203 | Periosteum | lgals2a |
| rpl7a | 6E-207 | 0.3697313 | 0.997 | 0.99 | 2.1E-202 | Periosteum | rpl7a |
| rps27.1 | 4E-206 | 0.3555527 | 0.991 | 0.976 | 1.3E-201 | Periosteum | rps27.1 |
| plekha5 | 7E-206 | 1.5563251 | 0.187 | 0.059 | 2.1E-201 | Periosteum | plekha5 |
| col16a1 | 5E-204 | 0.8578951 | 0.669 | 0.432 | 1.5E-199 | Periosteum | col16a1 |
| rplp0 | 2E-202 | 0.4077774 | 0.994 | 0.981 | 5.2E-198 | Periosteum | rplp0 |
| mafa | 3E-202 | 2.0046886 | 0.144 | 0.038 | 8.4E-198 | Periosteum | mafa |
| col21a1 | 4E-202 | 1.592523 | 0.201 | 0.067 | 1.4E-197 | Periosteum | col21a1 |
| rpl35a | 1E-200 | 0.3300705 | 0.997 | 0.988 | 3.8E-196 | Periosteum | rpl35a |
| sgcg | 4E-200 | 1.4430653 | 0.219 | 0.077 | 1.2E-195 | Periosteum | sgcg |
| slc25a5 | 6E-200 | 0.4069711 | 0.988 | 0.96 | 2E-195 | Periosteum | slc25a5 |
| c1qtnf12 | 1E-196 | 1.3266635 | 0.234 | 0.086 | 4.2E-192 | Periosteum | c1qtnf12 |
| epha2b | 2E-195 | 1.7612047 | 0.154 | 0.044 | 5.2E-191 | Periosteum | epha2b |
| rpl28 | 2E-195 | 0.3126245 | 0.998 | 0.99 | 7.1E-191 | Periosteum | rpl28 |
| rps27a | 3E-195 | 0.3140108 | 0.997 | 0.992 | 8.7E-191 | Periosteum | rps27a |
| rpl12 | 8E-195 | 0.358351 | 0.994 | 0.981 | 2.7E-190 | Periosteum | rpl12 |
| fgfr2 | 2E-194 | 0.9548935 | 0.55 | 0.326 | 5.8E-190 | Periosteum | fgfr2 |
| p4ha3 | 7E-193 | 2.1393434 | 0.117 | 0.028 | 2.3E-188 | Periosteum | p4ha3 |

| | | | | | | | |
|---|---|---|---|---|---|---|---|
| rpl23 | 1E-192 | 0.3217994 | 0.997 | 0.992 | 3.5E-188 | Periosteum | rpl23 |
| sec61g | 3E-192 | 0.4618962 | 0.969 | 0.919 | 9.4E-188 | Periosteum | sec61g |
| rpl30 | 5E-192 | 0.3334175 | 0.994 | 0.981 | 1.5E-187 | Periosteum | rpl30 |
| fibina | 7E-191 | 1.263468 | 0.386 | 0.194 | 2.3E-186 | Periosteum | fibina |
| spam1 | 6E-190 | 2.1186453 | 0.1 | 0.021 | 1.8E-185 | Periosteum | spam1 |
| kank1b | 6E-190 | 1.3781951 | 0.212 | 0.076 | 2.1E-185 | Periosteum | kank1b |
| rps5 | 1E-188 | 0.3739183 | 0.989 | 0.971 | 3.3E-184 | Periosteum | rps5 |
| rps15 | 6E-187 | 0.3205553 | 0.995 | 0.988 | 1.9E-182 | Periosteum | rps15 |
| c1qtnf6b | 2E-186 | 1.5581397 | 0.189 | 0.064 | 7.6E-182 | Periosteum | c1qtnf6b |
| vaspb | 6E-185 | 1.4424542 | 0.225 | 0.086 | 1.9E-180 | Periosteum | vaspb |
| rpl36a | 2E-184 | 0.3145932 | 0.998 | 0.994 | 6.2E-180 | Periosteum | rpl36a |
| eef1a1l1 | 6E-184 | 0.3170998 | 0.999 | 0.996 | 1.9E-179 | Periosteum | eef1a1l1 |
| faua | 1E-183 | 0.3443779 | 0.992 | 0.972 | 3.3E-179 | Periosteum | faua |
| rps4x | 1E-181 | 0.3739487 | 0.988 | 0.964 | 4.9E-177 | Periosteum | rps4x |
| rpl13a | 1E-180 | 0.333241 | 0.994 | 0.984 | 4.4E-176 | Periosteum | rpl13a |
| egfl6 | 1E-180 | 1.7460057 | 0.151 | 0.045 | 4.6E-176 | Periosteum | egfl6 |
| rpl17 | 5E-180 | 0.3299378 | 0.997 | 0.988 | 1.7E-175 | Periosteum | rpl17 |
| nme2b.1 | 3E-179 | 0.4257738 | 0.97 | 0.913 | 9.7E-175 | Periosteum | nme2b.1 |
| rps6 | 4E-179 | 0.3610398 | 0.984 | 0.958 | 1.1E-174 | Periosteum | rps6 |
| eef1b2 | 6E-178 | 0.4099203 | 0.972 | 0.924 | 1.9E-173 | Periosteum | eef1b2 |
| FP102018.1 | 1E-177 | 0.7803189 | 0.595 | 0.369 | 4.2E-173 | Periosteum | FP102018.1 |
| rpl13 | 1E-177 | 0.3099435 | 0.997 | 0.993 | 4.6E-173 | Periosteum | rpl13 |
| fap | 7E-177 | 1.5806353 | 0.168 | 0.054 | 2.4E-172 | Periosteum | fap |
| c1qtnf5 | 1E-175 | 1.0510026 | 0.302 | 0.135 | 3.9E-171 | Periosteum | c1qtnf5 |
| rack1 | 1E-174 | 0.380186 | 0.98 | 0.945 | 4.6E-170 | Periosteum | rack1 |
| rpl9 | 4E-174 | 0.3145777 | 0.995 | 0.987 | 1.4E-169 | Periosteum | rpl9 |
| tfpia | 9E-174 | 0.9517267 | 0.339 | 0.162 | 3.1E-169 | Periosteum | tfpia |
| kng1 | 5E-173 | 1.46069 | 0.192 | 0.068 | 1.5E-168 | Periosteum | kng1 |
| rps23 | 9E-173 | 0.3181168 | 0.991 | 0.981 | 2.8E-168 | Periosteum | rps23 |
| ndnf | 5E-172 | 1.5459442 | 0.228 | 0.091 | 1.6E-167 | Periosteum | ndnf |
| stxbp5a | 4E-171 | 1.922727 | 0.118 | 0.031 | 1.3E-166 | Periosteum | stxbp5a |
| rps11 | 4E-170 | 0.3013779 | 0.996 | 0.988 | 1.2E-165 | Periosteum | rps11 |
| ncam1a | 4E-169 | 1.513077 | 0.176 | 0.06 | 1.3E-164 | Periosteum | ncam1a |
| rps3a | 2E-168 | 0.3470794 | 0.993 | 0.98 | 7.9E-164 | Periosteum | rps3a |
| rps24 | 3E-168 | 0.2775136 | 0.998 | 0.996 | 1.1E-163 | Periosteum | rps24 |
| rps16 | 4E-168 | 0.2984417 | 0.997 | 0.988 | 1.2E-163 | Periosteum | rps16 |
| rps3 | 9E-167 | 0.3744387 | 0.979 | 0.943 | 2.9E-162 | Periosteum | rps3 |
| CR381618.1 | 6E-164 | 0.9359163 | 0.394 | 0.209 | 1.9E-159 | Periosteum | CR381618.1 |
| rpl18 | 2E-162 | 0.3224719 | 0.994 | 0.98 | 5E-158 | Periosteum | rpl18 |
| rpl22 | 3E-162 | 0.3283227 | 0.99 | 0.971 | 1.1E-157 | Periosteum | rpl22 |
| rpl18a | 1E-161 | 0.3083687 | 0.994 | 0.983 | 4.4E-157 | Periosteum | rpl18a |
| rps10 | 2E-161 | 0.2862146 | 0.998 | 0.994 | 7.2E-157 | Periosteum | rps10 |
| nid1a | 3E-161 | 0.7572919 | 0.409 | 0.216 | 1.1E-156 | Periosteum | nid1a |
| rpl14 | 1E-160 | 0.2471881 | 0.999 | 0.995 | 3.2E-156 | Periosteum | rpl14 |
| mmp14b | 9E-160 | 0.7192267 | 0.732 | 0.53 | 3E-155 | Periosteum | mmp14b |
| fhl2a | 2E-159 | 1.4765342 | 0.267 | 0.121 | 7.6E-155 | Periosteum | fhl2a |
| lurap1 | 9E-159 | 1.6558227 | 0.137 | 0.041 | 2.8E-154 | Periosteum | lurap1 |
| rpl3 | 2E-158 | 0.3302053 | 0.988 | 0.977 | 7.6E-154 | Periosteum | rpl3 |

| | | | | | | | |
|---|---|---|---|---|---|---|---|
| igsf10 | 2E-158 | 0.74788 | 0.462 | 0.258 | 7.7E-154 | Periosteum | igsf10 |
| rpl10 | 2E-157 | 0.3389557 | 0.985 | 0.967 | 4.9E-153 | Periosteum | rpl10 |
| rpl35 | 2E-156 | 0.2899562 | 0.997 | 0.991 | 7.7E-152 | Periosteum | rpl35 |
| fhl1a | 3E-155 | 2.6328082 | 0.106 | 0.027 | 9.7E-151 | Periosteum | fhl1a |
| col2a1b | 2E-154 | 0.3214636 | 0.697 | 0.48 | 5.6E-150 | Periosteum | col2a1b |
| pear1 | 1E-153 | 1.1717157 | 0.262 | 0.116 | 3.8E-149 | Periosteum | pear1 |
| tpm4a | 1E-153 | 1.2339001 | 0.708 | 0.536 | 4E-149 | Periosteum | tpm4a |
| tpt1 | 2E-153 | 0.3803226 | 0.962 | 0.923 | 6E-149 | Periosteum | tpt1 |
| rps18 | 1E-151 | 0.339274 | 0.983 | 0.961 | 4.7E-147 | Periosteum | rps18 |
| rps19 | 4E-151 | 0.2616532 | 0.998 | 0.996 | 1.3E-146 | Periosteum | rps19 |
| mxra8a | 1E-150 | 0.8108666 | 0.424 | 0.235 | 4.3E-146 | Periosteum | mxra8a |
| rpl21 | 8E-149 | 0.3001984 | 0.996 | 0.984 | 2.6E-144 | Periosteum | rpl21 |
| c1qtnf6a | 1E-148 | 1.637651 | 0.152 | 0.052 | 4.5E-144 | Periosteum | c1qtnf6a |
| rpl15 | 2E-148 | 0.3417986 | 0.989 | 0.969 | 5.9E-144 | Periosteum | rpl15 |
| tspan4b | 4E-148 | 0.6927507 | 0.526 | 0.316 | 1.4E-143 | Periosteum | tspan4b |
| col5a1 | 5E-148 | 0.4313226 | 0.992 | 0.961 | 1.6E-143 | Periosteum | col5a1 |
| dcn | 8E-148 | 0.3372779 | 0.967 | 0.858 | 2.7E-143 | Periosteum | dcn |
| rpl8 | 4E-147 | 0.3238285 | 0.99 | 0.977 | 1.2E-142 | Periosteum | rpl8 |
| rpl31 | 6E-146 | 0.2781289 | 0.997 | 0.993 | 1.8E-141 | Periosteum | rpl31 |
| rpl5a | 8E-146 | 0.352824 | 0.964 | 0.914 | 2.5E-141 | Periosteum | rpl5a |
| rps20 | 9E-146 | 0.3122163 | 0.996 | 0.985 | 2.8E-141 | Periosteum | rps20 |
| cdo1 | 1E-144 | 0.7218929 | 0.505 | 0.31 | 3.4E-140 | Periosteum | cdo1 |
| rps15a | 2E-144 | 0.2833619 | 0.997 | 0.989 | 7.3E-140 | Periosteum | rps15a |
| rps25 | 8E-143 | 0.2442047 | 0.999 | 0.993 | 2.5E-138 | Periosteum | rps25 |
| rpl37.1 | 1E-142 | 0.303 | 0.997 | 0.99 | 3.2E-138 | Periosteum | rpl37.1 |
| f10 | 5E-142 | 0.7379536 | 0.577 | 0.38 | 1.8E-137 | Periosteum | f10 |
| rps9 | 2E-141 | 0.2881678 | 0.996 | 0.986 | 5.9E-137 | Periosteum | rps9 |
| aebp1 | 2E-141 | 0.7054173 | 0.721 | 0.52 | 6.6E-137 | Periosteum | aebp1 |
| jam3b | 3E-141 | 0.9228128 | 0.373 | 0.204 | 8.5E-137 | Periosteum | jam3b |
| rplp2l | 2E-139 | 0.2734902 | 0.995 | 0.986 | 5.4E-135 | Periosteum | rplp2l |
| rps12 | 7E-137 | 0.3207757 | 0.99 | 0.977 | 2.3E-132 | Periosteum | rps12 |
| rpl10a | 2E-136 | 0.3183479 | 0.99 | 0.977 | 4.9E-132 | Periosteum | rpl10a |
| glula | 9E-136 | 0.5484539 | 0.791 | 0.6 | 3E-131 | Periosteum | glula |
| rpl7 | 3E-134 | 0.3218749 | 0.988 | 0.971 | 1.1E-129 | Periosteum | rpl7 |
| rps26 | 4E-134 | 0.3160352 | 0.987 | 0.967 | 1.5E-129 | Periosteum | rps26 |
| rps8a | 3E-133 | 0.2760156 | 0.996 | 0.991 | 8.8E-129 | Periosteum | rps8a |
| rps7 | 2E-132 | 0.271268 | 0.997 | 0.991 | 8E-128 | Periosteum | rps7 |
| trib2 | 3E-132 | 1.339338 | 0.189 | 0.078 | 1.1E-127 | Periosteum | trib2 |
| rplp1 | 2E-131 | 0.2307395 | 1 | 0.999 | 5.5E-127 | Periosteum | rplp1 |
| rpl5b | 2E-131 | 0.3793975 | 0.922 | 0.842 | 6.4E-127 | Periosteum | rpl5b |
| rpl39 | 2E-131 | 0.2919101 | 0.993 | 0.983 | 6.9E-127 | Periosteum | rpl39 |
| rpl36 | 2E-131 | 0.2802455 | 0.992 | 0.985 | 6.9E-127 | Periosteum | rpl36 |
| gpm6ab | 7E-130 | 1.1501889 | 0.19 | 0.077 | 2.2E-125 | Periosteum | gpm6ab |
| ndnfl | 3E-129 | 1.3885901 | 0.102 | 0.029 | 9.4E-125 | Periosteum | ndnfl |
| cnn3a | 4E-129 | 0.6422605 | 0.654 | 0.457 | 1.3E-124 | Periosteum | cnn3a |
| srpx | 5E-129 | 1.0032403 | 0.299 | 0.153 | 1.5E-124 | Periosteum | srpx |
| eif4a1b | 1E-128 | 0.5457871 | 0.76 | 0.594 | 3.7E-124 | Periosteum | eif4a1b |
| RPS17 | 6E-128 | 0.2700859 | 0.994 | 0.986 | 1.9E-123 | Periosteum | RPS17 |

| | | | | | | | |
|---|---|---|---|---|---|---|---|
| apcdd1l | 8E-128 | 1.3078314 | 0.22 | 0.098 | 2.6E-123 | Periosteum | apcdd1l |
| rpl4 | 8E-127 | 0.347897 | 0.955 | 0.889 | 2.4E-122 | Periosteum | rpl4 |
| myl9a | 2E-126 | 1.0023241 | 0.54 | 0.364 | 5.1E-122 | Periosteum | myl9a |
| tpm4b | 7E-125 | 1.5422533 | 0.179 | 0.074 | 2.2E-120 | Periosteum | tpm4b |
| pabpc1b | 9E-125 | 0.5740155 | 0.736 | 0.57 | 2.9E-120 | Periosteum | pabpc1b |
| rpl19 | 1E-124 | 0.2875666 | 0.995 | 0.982 | 4.7E-120 | Periosteum | rpl19 |
| rpl24 | 4E-123 | 0.2509107 | 0.998 | 0.993 | 1.2E-118 | Periosteum | rpl24 |
| rpl23a | 5E-121 | 0.2753104 | 0.988 | 0.969 | 1.6E-116 | Periosteum | rpl23a |
| tmem47 | 9E-121 | 1.0529748 | 0.285 | 0.147 | 2.8E-116 | Periosteum | tmem47 |
| rpl11 | 5E-120 | 0.2560366 | 0.995 | 0.988 | 1.7E-115 | Periosteum | rpl11 |
| ghra | 6E-120 | 0.8022531 | 0.374 | 0.212 | 2E-115 | Periosteum | ghra |
| mafba | 8E-120 | 0.6959487 | 0.553 | 0.373 | 2.6E-115 | Periosteum | mafba |
| palld | 1E-119 | 1.1674136 | 0.266 | 0.134 | 4.6E-115 | Periosteum | palld |
| rps26l | 2E-119 | 0.2735403 | 0.99 | 0.973 | 7.6E-115 | Periosteum | rps26l |
| CABZ01068367.1 | 3E-119 | 0.5957871 | 0.567 | 0.374 | 9.9E-115 | Periosteum | CABZ01068367.1 |
| spon1b | 3E-118 | 1.2953448 | 0.135 | 0.048 | 9.8E-114 | Periosteum | spon1b |
| rpl27 | 1E-117 | 0.2651983 | 0.99 | 0.977 | 4.4E-113 | Periosteum | rpl27 |
| rspo1 | 2E-117 | 1.1499512 | 0.146 | 0.054 | 6E-113 | Periosteum | rspo1 |
| bcl2l10 | 5E-117 | 0.9695952 | 0.308 | 0.166 | 1.7E-112 | Periosteum | bcl2l10 |
| tnfrsf11a | 1E-116 | 0.8258314 | 0.317 | 0.169 | 4.6E-112 | Periosteum | tnfrsf11a |
| tanc2a | 5E-116 | 1.2549443 | 0.169 | 0.069 | 1.7E-111 | Periosteum | tanc2a |
| col12a1a | 8E-115 | 0.3794149 | 0.896 | 0.736 | 2.5E-110 | Periosteum | col12a1a |
| tent5bb | 5E-114 | 0.9795297 | 0.258 | 0.13 | 1.6E-109 | Periosteum | tent5bb |
| rsl24d1 | 5E-111 | 0.440794 | 0.848 | 0.738 | 1.6E-106 | Periosteum | rsl24d1 |
| serbp1a | 6E-111 | 0.3162374 | 0.958 | 0.903 | 2.1E-106 | Periosteum | serbp1a |
| cbfb | 7E-111 | 0.963447 | 0.233 | 0.112 | 2.3E-106 | Periosteum | cbfb |
| tagln | 2E-110 | 0.7645225 | 0.718 | 0.519 | 7.9E-106 | Periosteum | tagln |
| cx43 | 1E-109 | 0.7507341 | 0.295 | 0.156 | 4.5E-105 | Periosteum | cx43 |
| ENSDARG000001152 | 5E-108 | 0.2396033 | 0.997 | 0.993 | 1.6E-103 | Periosteum | ENSDARG00000115 |
| rpl6 | 1E-107 | 0.3078563 | 0.98 | 0.956 | 3.3E-103 | Periosteum | rpl6 |
| ppib | 4E-107 | 0.3163088 | 0.971 | 0.911 | 1.3E-102 | Periosteum | ppib |
| loxl2b | 3E-106 | 0.6298704 | 0.472 | 0.3 | 1.1E-101 | Periosteum | loxl2b |
| rps13 | 4E-106 | 0.2244953 | 0.996 | 0.99 | 1.2E-101 | Periosteum | rps13 |
| naca | 5E-104 | 0.2831569 | 0.981 | 0.959 | 1.8E-99 | Periosteum | naca |
| kazald3 | 3E-103 | 0.6679758 | 0.405 | 0.245 | 1E-98 | Periosteum | kazald3 |
| olfml3b | 8E-103 | 0.5673939 | 0.67 | 0.497 | 2.52E-98 | Periosteum | olfml3b |
| neo1b | 1E-102 | 1.0483859 | 0.168 | 0.072 | 3.36E-98 | Periosteum | neo1b |
| eif3ea | 2E-102 | 0.4960565 | 0.712 | 0.543 | 5.2E-98 | Periosteum | eif3ea |
| relch | 4E-102 | 1.2013354 | 0.15 | 0.061 | 1.16E-97 | Periosteum | relch |
| cyp1c1 | 2E-101 | 1.0502921 | 0.198 | 0.093 | 6.84E-97 | Periosteum | cyp1c1 |
| rgs5b | 1E-100 | 0.5510762 | 0.673 | 0.507 | 3.77E-96 | Periosteum | rgs5b |
| mmp14a | 1E-98 | 0.5244429 | 0.783 | 0.633 | 3.1E-94 | Periosteum | mmp14a |
| si:ch1073-184j22.1 | 4.1E-98 | 1.065888 | 0.165 | 0.071 | 1.32E-93 | Periosteum | si:ch1073-184j22.1 |
| loxa | 6.8E-98 | 0.7647972 | 0.292 | 0.161 | 2.22E-93 | Periosteum | loxa |
| romo1 | 7.3E-96 | 0.4201789 | 0.817 | 0.694 | 2.38E-91 | Periosteum | romo1 |
| stc2a | 2.8E-94 | 2.6500598 | 0.121 | 0.047 | 9.21E-90 | Periosteum | stc2a |
| gask1a | 4.4E-94 | 0.6694127 | 0.318 | 0.182 | 1.42E-89 | Periosteum | gask1a |
| ssr2 | 6.5E-94 | 0.3633805 | 0.825 | 0.676 | 2.1E-89 | Periosteum | ssr2 |

| | | | | | | | |
|---|---|---|---|---|---|---|---|
| rps21 | 6E-92 | 0.2410882 | 0.992 | 0.977 | 1.96E-87 | Periosteum | rps21 |
| rps28 | 6.8E-92 | 0.2745156 | 0.991 | 0.977 | 2.2E-87 | Periosteum | rps28 |
| nedd9 | 4.7E-90 | 1.2836449 | 0.149 | 0.065 | 1.53E-85 | Periosteum | nedd9 |
| nsa2 | 2.2E-89 | 0.4553866 | 0.755 | 0.619 | 7.04E-85 | Periosteum | nsa2 |
| smarcd3b | 8.1E-89 | 1.3663146 | 0.105 | 0.038 | 2.65E-84 | Periosteum | smarcd3b |
| nrp2b | 9.2E-89 | 0.934455 | 0.223 | 0.115 | 2.98E-84 | Periosteum | nrp2b |
| si:ch211-284f22.3 | 1E-87 | 0.84228 | 0.213 | 0.107 | 3.4E-83 | Periosteum | si:ch211-284f22.3 |
| luzp1 | 5.7E-87 | 0.8789789 | 0.284 | 0.163 | 1.85E-82 | Periosteum | luzp1 |
| flrt2 | 5.9E-87 | 1.1011254 | 0.152 | 0.067 | 1.92E-82 | Periosteum | flrt2 |
| niban2a | 1.2E-86 | 1.5078982 | 0.115 | 0.045 | 3.75E-82 | Periosteum | niban2a |
| slit3 | 2.5E-86 | 0.8176211 | 0.319 | 0.189 | 8.12E-82 | Periosteum | slit3 |
| ccn2a | 4.8E-86 | 0.950519 | 0.324 | 0.194 | 1.57E-81 | Periosteum | ccn2a |
| f3b | 3.1E-85 | 0.5880003 | 0.518 | 0.365 | 9.92E-81 | Periosteum | f3b |
| mxra5a | 3.8E-85 | 1.1232897 | 0.153 | 0.069 | 1.22E-80 | Periosteum | mxra5a |
| svep1 | 1.6E-84 | 0.8178735 | 0.246 | 0.134 | 5.34E-80 | Periosteum | svep1 |
| dchs1a | 6E-83 | 0.9339195 | 0.158 | 0.072 | 1.94E-78 | Periosteum | dchs1a |
| adamts2 | 1.1E-82 | 0.4181267 | 0.527 | 0.357 | 3.45E-78 | Periosteum | adamts2 |
| serpinh1a | 1.1E-82 | 0.6589962 | 0.361 | 0.225 | 3.48E-78 | Periosteum | serpinh1a |
| sema3fa | 1.3E-82 | 1.352839 | 0.132 | 0.056 | 4.09E-78 | Periosteum | sema3fa |
| steap4 | 3.5E-81 | 1.0039613 | 0.288 | 0.173 | 1.15E-76 | Periosteum | steap4 |
| csrp2 | 2.3E-79 | 0.7988396 | 0.21 | 0.109 | 7.59E-75 | Periosteum | csrp2 |
| snai1b | 4.5E-79 | 0.8402823 | 0.202 | 0.103 | 1.47E-74 | Periosteum | snai1b |
| crispld1a | 7.6E-79 | 1.2993536 | 0.104 | 0.04 | 2.46E-74 | Periosteum | crispld1a |
| tenm3 | 3.4E-78 | 0.6437629 | 0.323 | 0.195 | 1.12E-73 | Periosteum | tenm3 |
| rps17 | 1E-76 | 0.2252446 | 0.977 | 0.942 | 3.28E-72 | Periosteum | rps17 |
| fat3a | 2.5E-76 | 1.3422288 | 0.108 | 0.043 | 7.97E-72 | Periosteum | fat3a |
| serpinf1 | 3.4E-76 | 0.401055 | 0.704 | 0.538 | 1.11E-71 | Periosteum | serpinf1 |
| cmtm4 | 7.7E-76 | 1.0460722 | 0.202 | 0.107 | 2.51E-71 | Periosteum | cmtm4 |
| eif3f | 4.3E-75 | 0.338593 | 0.827 | 0.718 | 1.4E-70 | Periosteum | eif3f |
| gal3st2 | 1.1E-74 | 1.0879066 | 0.148 | 0.069 | 3.47E-70 | Periosteum | gal3st2 |
| BX927333.1 | 4.9E-74 | 1.2606334 | 0.106 | 0.043 | 1.6E-69 | Periosteum | BX927333.1 |
| phlda3 | 7.5E-74 | 0.7171136 | 0.284 | 0.169 | 2.45E-69 | Periosteum | phlda3 |
| eef2b | 2.2E-73 | 0.269923 | 0.958 | 0.904 | 7.02E-69 | Periosteum | eef2b |
| entpd5a | 2.5E-73 | 0.8770691 | 0.113 | 0.047 | 8.06E-69 | Periosteum | entpd5a |
| rpl29 | 3.2E-73 | 0.2796187 | 0.966 | 0.928 | 1.05E-68 | Periosteum | rpl29 |
| klf12a | 4.4E-73 | 0.6389814 | 0.233 | 0.128 | 1.44E-68 | Periosteum | klf12a |
| tacc1 | 1.2E-71 | 0.6524341 | 0.326 | 0.205 | 3.88E-67 | Periosteum | tacc1 |
| snai2 | 1.3E-71 | 0.7523789 | 0.188 | 0.096 | 4.38E-67 | Periosteum | snai2 |
| frzb | 2.6E-71 | 0.6603484 | 0.217 | 0.12 | 8.43E-67 | Periosteum | frzb |
| si:ch211-251b21.1 | 4.5E-71 | 0.6494118 | 0.562 | 0.44 | 1.46E-66 | Periosteum | si:ch211-251b21.1 |
| si:ch211-133n4.4 | 3.9E-69 | 0.5043881 | 0.577 | 0.434 | 1.25E-64 | Periosteum | si:ch211-133n4.4 |
| ptmaa | 5E-69 | 0.314825 | 0.916 | 0.827 | 1.62E-64 | Periosteum | ptmaa |
| id2a | 7.9E-69 | 0.4696196 | 0.634 | 0.489 | 2.57E-64 | Periosteum | id2a |
| si:ch211-126j24.1 | 1.8E-68 | 0.8424518 | 0.154 | 0.074 | 5.71E-64 | Periosteum | si:ch211-126j24.1 |
| emilin1a | 6.1E-68 | 0.8135648 | 0.17 | 0.086 | 1.99E-63 | Periosteum | emilin1a |
| fbln5 | 3.3E-67 | 0.9431589 | 0.155 | 0.076 | 1.07E-62 | Periosteum | fbln5 |
| c7a | 4.3E-66 | 0.5228752 | 0.276 | 0.164 | 1.4E-61 | Periosteum | c7a |
| serinc5 | 1.2E-65 | 0.5647324 | 0.351 | 0.228 | 4.02E-61 | Periosteum | serinc5 |

| | | | | | | | |
|---|---|---|---|---|---|---|---|
| robo1 | 2E-65 | 0.9123877 | 0.146 | 0.071 | 6.6E-61 | Periosteum | robo1 |
| dipk1c | 2.3E-65 | 0.6840275 | 0.197 | 0.106 | 7.52E-61 | Periosteum | dipk1c |
| gtpbp4 | 3.1E-65 | 0.4517501 | 0.541 | 0.397 | 1.02E-60 | Periosteum | gtpbp4 |
| ENSDARG000000762 | 6E-65 | 0.3928367 | 0.739 | 0.611 | 1.94E-60 | Periosteum | ENSDARG00000076 |
| ssr4 | 7.8E-65 | 0.3080808 | 0.746 | 0.593 | 2.54E-60 | Periosteum | ssr4 |
| asip1 | 1.9E-64 | 0.7065769 | 0.274 | 0.167 | 6.29E-60 | Periosteum | asip1 |
| ssr3 | 5.2E-64 | 0.2592166 | 0.723 | 0.562 | 1.7E-59 | Periosteum | ssr3 |
| sec61a1 | 3.3E-63 | 0.4303403 | 0.548 | 0.406 | 1.08E-58 | Periosteum | sec61a1 |
| pik3r3b | 8E-63 | 0.524946 | 0.388 | 0.262 | 2.59E-58 | Periosteum | pik3r3b |
| sox6 | 4.1E-62 | 0.7501224 | 0.178 | 0.095 | 1.35E-57 | Periosteum | sox6 |
| ddx21 | 6.5E-62 | 0.4629072 | 0.467 | 0.33 | 2.11E-57 | Periosteum | ddx21 |
| acvrl1 | 6.9E-62 | 0.5051448 | 0.358 | 0.234 | 2.26E-57 | Periosteum | acvrl1 |
| eif5a | 9.7E-62 | 0.2914312 | 0.824 | 0.709 | 3.15E-57 | Periosteum | eif5a |
| nfixb | 2.7E-61 | 0.4577477 | 0.415 | 0.281 | 8.92E-57 | Periosteum | nfixb |
| mylka | 2.4E-60 | 0.574075 | 0.186 | 0.1 | 7.88E-56 | Periosteum | mylka |
| vat1 | 3.8E-60 | 0.2933791 | 0.442 | 0.302 | 1.23E-55 | Periosteum | vat1 |
| rbfox1 | 8.4E-60 | 0.3797575 | 0.539 | 0.392 | 2.73E-55 | Periosteum | rbfox1 |
| wu:fj16a03 | 2.8E-58 | 0.592356 | 0.286 | 0.182 | 9.11E-54 | Periosteum | wu:fj16a03 |
| ccdc80 | 6.2E-58 | 0.2383093 | 0.517 | 0.373 | 2.01E-53 | Periosteum | ccdc80 |
| tdh | 8.8E-58 | 0.5989943 | 0.343 | 0.23 | 2.85E-53 | Periosteum | tdh |
| cirbpb | 9.2E-58 | 0.2889193 | 0.9 | 0.826 | 2.99E-53 | Periosteum | cirbpb |
| htra1a | 9.3E-58 | 0.4331118 | 0.333 | 0.216 | 3.04E-53 | Periosteum | htra1a |
| mych | 1.9E-57 | 0.5688158 | 0.491 | 0.368 | 6.1E-53 | Periosteum | mych |
| gstp1 | 2.3E-57 | 0.2775637 | 0.766 | 0.631 | 7.49E-53 | Periosteum | gstp1 |
| rpl38 | 2.8E-57 | 0.2563925 | 0.944 | 0.907 | 9.09E-53 | Periosteum | rpl38 |
| thbs2a | 3.1E-57 | 0.6372932 | 0.302 | 0.193 | 1.01E-52 | Periosteum | thbs2a |
| rpe65b | 5E-57 | 0.8087029 | 0.13 | 0.063 | 1.62E-52 | Periosteum | rpe65b |
| gnai2b | 1.3E-56 | 0.9858415 | 0.12 | 0.057 | 4.15E-52 | Periosteum | gnai2b |
| myl9b | 2.6E-56 | 0.5064631 | 0.542 | 0.413 | 8.61E-52 | Periosteum | myl9b |
| cx43.4 | 2.9E-56 | 0.5879397 | 0.32 | 0.212 | 9.35E-52 | Periosteum | cx43.4 |
| dlx4a | 3.6E-56 | 0.5203363 | 0.289 | 0.183 | 1.18E-51 | Periosteum | dlx4a |
| gsc | 4.4E-56 | 0.9381483 | 0.19 | 0.109 | 1.42E-51 | Periosteum | gsc |
| rai14 | 1E-55 | 0.9176889 | 0.151 | 0.08 | 3.28E-51 | Periosteum | rai14 |
| CR855311.1 | 7.7E-55 | 1.1041414 | 0.234 | 0.145 | 2.52E-50 | Periosteum | CR855311.1 |
| fthl27 | 1.6E-54 | 0.2485297 | 0.901 | 0.833 | 5.16E-50 | Periosteum | fthl27 |
| gse1 | 1.7E-54 | 0.5612621 | 0.204 | 0.117 | 5.39E-50 | Periosteum | gse1 |
| pno1 | 2.5E-54 | 0.3889931 | 0.527 | 0.395 | 7.98E-50 | Periosteum | pno1 |
| srpx2 | 5.2E-54 | 0.4477912 | 0.364 | 0.243 | 1.68E-49 | Periosteum | srpx2 |
| tgfb3 | 5.3E-54 | 0.7355464 | 0.144 | 0.074 | 1.71E-49 | Periosteum | tgfb3 |
| olfml3a | 5.5E-54 | 0.4482849 | 0.363 | 0.247 | 1.77E-49 | Periosteum | olfml3a |
| SCARF2 | 6.8E-54 | 0.5063334 | 0.372 | 0.257 | 2.22E-49 | Periosteum | SCARF2 |
| six1a | 7.6E-54 | 0.7219223 | 0.207 | 0.122 | 2.48E-49 | Periosteum | six1a |
| scxa | 9.1E-54 | 0.9691462 | 0.145 | 0.077 | 2.95E-49 | Periosteum | scxa |
| rbp5 | 1.7E-53 | 0.6036692 | 0.193 | 0.11 | 5.49E-49 | Periosteum | rbp5 |
| ola1 | 2.6E-53 | 0.3393471 | 0.629 | 0.493 | 8.37E-49 | Periosteum | ola1 |
| CU929070.5 | 3.1E-53 | 0.4977918 | 0.34 | 0.23 | 1.02E-48 | Periosteum | CU929070.5 |
| rhpn2 | 7.3E-53 | 0.5777565 | 0.225 | 0.134 | 2.37E-48 | Periosteum | rhpn2 |
| zgc:153372 | 2.9E-52 | 0.7423876 | 0.151 | 0.08 | 9.36E-48 | Periosteum | zgc:153372 |

| | | | | | | | |
|---|---|---|---|---|---|---|---|
| eif3g | 6E-52 | 0.3213999 | 0.652 | 0.513 | 1.94E-47 | Periosteum | eif3g |
| dap1b.1 | 6.9E-52 | 0.3573668 | 0.64 | 0.493 | 2.25E-47 | Periosteum | dap1b.1 |
| lrp5 | 1.1E-51 | 0.8141627 | 0.183 | 0.105 | 3.46E-47 | Periosteum | lrp5 |
| cox6b2 | 1.2E-51 | 0.2214838 | 0.894 | 0.806 | 4E-47 | Periosteum | cox6b2 |
| usp54a | 1.4E-51 | 0.7771674 | 0.111 | 0.053 | 4.49E-47 | Periosteum | usp54a |
| kank4 | 4E-51 | 0.6656477 | 0.173 | 0.096 | 1.3E-46 | Periosteum | kank4 |
| eef1da | 2.3E-50 | 0.2013437 | 0.925 | 0.87 | 7.47E-46 | Periosteum | eef1da |
| slc38a5b | 3.3E-50 | 0.3828048 | 0.467 | 0.341 | 1.06E-45 | Periosteum | slc38a5b |
| lpp | 5.7E-50 | 0.5425114 | 0.351 | 0.243 | 1.85E-45 | Periosteum | lpp |
| serf2 | 1.8E-49 | 0.256002 | 0.863 | 0.762 | 5.77E-45 | Periosteum | serf2 |
| il11ra | 3.3E-49 | 0.3254863 | 0.344 | 0.229 | 1.07E-44 | Periosteum | il11ra |
| rpl22l1 | 9.1E-49 | 0.270918 | 0.842 | 0.738 | 2.97E-44 | Periosteum | rpl22l1 |
| rps29 | 1.1E-48 | 0.2116625 | 0.987 | 0.968 | 3.7E-44 | Periosteum | rps29 |
| uacab | 1.3E-48 | 0.8236335 | 0.13 | 0.067 | 4.15E-44 | Periosteum | uacab |
| rtn4b | 1.4E-48 | 0.6862824 | 0.156 | 0.086 | 4.54E-44 | Periosteum | rtn4b |
| sox11b | 2.1E-48 | 0.494217 | 0.302 | 0.201 | 6.71E-44 | Periosteum | sox11b |
| ctsk | 2.3E-48 | 0.4912483 | 0.707 | 0.581 | 7.52E-44 | Periosteum | ctsk |
| zgc:110239 | 3.3E-48 | 0.4852865 | 0.328 | 0.222 | 1.07E-43 | Periosteum | zgc:110239 |
| ptpn11b | 4E-48 | 0.8049363 | 0.118 | 0.059 | 1.29E-43 | Periosteum | ptpn11b |
| ppp1r1c | 8E-48 | 0.7088575 | 0.204 | 0.124 | 2.61E-43 | Periosteum | ppp1r1c |
| lfng | 8.4E-48 | 0.5042444 | 0.354 | 0.244 | 2.72E-43 | Periosteum | lfng |
| si:ch211-288d18.1 | 2.5E-47 | 0.8578206 | 0.14 | 0.076 | 8.25E-43 | Periosteum | si:ch211-288d18.1 |
| gpx1a | 3.1E-47 | 0.530185 | 0.28 | 0.184 | 9.99E-43 | Periosteum | gpx1a |
| limch1a | 2.4E-46 | 0.7952258 | 0.135 | 0.072 | 7.75E-42 | Periosteum | limch1a |
| cd9b | 3.5E-46 | 0.57488 | 0.164 | 0.094 | 1.13E-41 | Periosteum | cd9b |
| pdia3 | 6.4E-46 | 0.2255978 | 0.788 | 0.675 | 2.09E-41 | Periosteum | pdia3 |
| vdac2 | 1E-45 | 0.2835532 | 0.64 | 0.506 | 3.25E-41 | Periosteum | vdac2 |
| marcksl1b | 1.4E-45 | 0.3557892 | 0.352 | 0.244 | 4.7E-41 | Periosteum | marcksl1b |
| mfge8a | 2.7E-45 | 0.7137124 | 0.187 | 0.113 | 8.83E-41 | Periosteum | mfge8a |
| bloc1s6 | 5.4E-45 | 0.5754307 | 0.261 | 0.171 | 1.76E-40 | Periosteum | bloc1s6 |
| btf3 | 1.1E-44 | 0.2066229 | 0.93 | 0.87 | 3.56E-40 | Periosteum | btf3 |
| rrbp1b | 1.2E-44 | 0.380333 | 0.334 | 0.229 | 4.06E-40 | Periosteum | rrbp1b |
| s1pr5a | 1.3E-44 | 0.8907916 | 0.111 | 0.056 | 4.21E-40 | Periosteum | s1pr5a |
| rnd1b | 5.5E-44 | 0.5187065 | 0.167 | 0.096 | 1.79E-39 | Periosteum | rnd1b |
| itgb1b.2 | 5.6E-44 | 0.5041308 | 0.178 | 0.104 | 1.81E-39 | Periosteum | itgb1b.2 |
| tgfb5 | 1E-43 | 0.5657646 | 0.18 | 0.107 | 3.28E-39 | Periosteum | tgfb5 |
| idh2 | 1.2E-43 | 0.3071041 | 0.454 | 0.332 | 3.87E-39 | Periosteum | idh2 |
| fkbp7 | 1.4E-43 | 0.3061774 | 0.723 | 0.601 | 4.46E-39 | Periosteum | fkbp7 |
| nme3 | 1.8E-43 | 0.3242404 | 0.564 | 0.437 | 5.72E-39 | Periosteum | nme3 |
| enpp1 | 4.8E-43 | 0.5491839 | 0.267 | 0.179 | 1.56E-38 | Periosteum | enpp1 |
| ptpn13 | 7.3E-43 | 0.3860236 | 0.401 | 0.291 | 2.38E-38 | Periosteum | ptpn13 |
| gstt1a | 4.9E-42 | 0.5455249 | 0.13 | 0.07 | 1.61E-37 | Periosteum | gstt1a |
| actn1 | 6.7E-42 | 0.5043257 | 0.414 | 0.307 | 2.18E-37 | Periosteum | actn1 |
| eif2s3 | 7.8E-42 | 0.3017618 | 0.609 | 0.484 | 2.53E-37 | Periosteum | eif2s3 |
| tbx2b | 2.1E-41 | 0.4906941 | 0.195 | 0.12 | 6.9E-37 | Periosteum | tbx2b |
| eif2s1b | 2.1E-41 | 0.3309957 | 0.495 | 0.376 | 6.94E-37 | Periosteum | eif2s1b |
| ak2 | 7.3E-41 | 0.3398224 | 0.483 | 0.369 | 2.39E-36 | Periosteum | ak2 |
| mob2a | 7.7E-41 | 0.5694144 | 0.207 | 0.13 | 2.5E-36 | Periosteum | mob2a |

| | | | | | | | |
|---|---|---|---|---|---|---|---|
| podn | 9.3E-41 | 0.4734914 | 0.25 | 0.164 | 3.03E-36 | Periosteum | podn |
| sash1a | 3.3E-40 | 0.4202036 | 0.294 | 0.201 | 1.08E-35 | Periosteum | sash1a |
| nop53 | 3.5E-40 | 0.3027011 | 0.572 | 0.446 | 1.13E-35 | Periosteum | nop53 |
| eif3m | 5.1E-40 | 0.2641383 | 0.729 | 0.616 | 1.67E-35 | Periosteum | eif3m |
| ahcy | 5.3E-40 | 0.5545334 | 0.261 | 0.177 | 1.71E-35 | Periosteum | ahcy |
| fkbp10b | 6.8E-40 | 0.4088357 | 0.345 | 0.245 | 2.22E-35 | Periosteum | fkbp10b |
| si:dkey-164f24.2 | 1.3E-39 | 0.6281363 | 0.246 | 0.162 | 4.27E-35 | Periosteum | si:dkey-164f24.2 |
| neo1a | 1.3E-39 | 0.4419638 | 0.261 | 0.174 | 4.36E-35 | Periosteum | neo1a |
| plekhg5a | 3.1E-39 | 0.8590297 | 0.122 | 0.067 | 1.02E-34 | Periosteum | plekhg5a |
| serpine1 | 3.4E-39 | 0.2216668 | 0.837 | 0.754 | 1.1E-34 | Periosteum | serpine1 |
| csrp1a | 4.5E-39 | 0.5210398 | 0.342 | 0.247 | 1.46E-34 | Periosteum | csrp1a |
| pcxa | 5.1E-39 | 0.5096961 | 0.18 | 0.11 | 1.64E-34 | Periosteum | pcxa |
| eif3s6ip | 1.1E-38 | 0.3267413 | 0.418 | 0.309 | 3.68E-34 | Periosteum | eif3s6ip |
| vdac3 | 1.8E-38 | 0.2485928 | 0.642 | 0.514 | 5.93E-34 | Periosteum | vdac3 |
| tomm5 | 1.9E-38 | 0.2733372 | 0.563 | 0.436 | 6.2E-34 | Periosteum | tomm5 |
| lpin1 | 2.7E-38 | 0.7105847 | 0.109 | 0.057 | 8.82E-34 | Periosteum | lpin1 |
| c1qtnf1 | 3.6E-38 | 0.7628276 | 0.126 | 0.07 | 1.18E-33 | Periosteum | c1qtnf1 |
| hmgn7 | 4.5E-38 | 0.285982 | 0.632 | 0.511 | 1.45E-33 | Periosteum | hmgn7 |
| zgc:113531 | 5.5E-38 | 0.4142899 | 0.371 | 0.269 | 1.79E-33 | Periosteum | zgc:113531 |
| eva1a | 1.1E-37 | 0.7054033 | 0.101 | 0.053 | 3.64E-33 | Periosteum | eva1a |
| sat1b | 1.5E-37 | 0.6177269 | 0.17 | 0.104 | 4.92E-33 | Periosteum | sat1b |
| si:dkey-56m19.5 | 2.6E-37 | 0.8906693 | 0.138 | 0.08 | 8.38E-33 | Periosteum | si:dkey-56m19.5 |
| fhl3b | 8.2E-37 | 0.4610119 | 0.172 | 0.106 | 2.68E-32 | Periosteum | fhl3b |
| arl4d | 8.5E-37 | 0.6649495 | 0.118 | 0.065 | 2.75E-32 | Periosteum | arl4d |
| camk1b | 1E-36 | 0.6310924 | 0.103 | 0.054 | 3.25E-32 | Periosteum | camk1b |
| slc25a48 | 1.3E-36 | 0.3983435 | 0.207 | 0.131 | 4.17E-32 | Periosteum | slc25a48 |
| cd82a | 1.8E-36 | 0.3118044 | 0.458 | 0.344 | 5.73E-32 | Periosteum | cd82a |
| rbpjb | 2.9E-36 | 0.6166975 | 0.153 | 0.092 | 9.47E-32 | Periosteum | rbpjb |
| rasgef1ba | 4.5E-36 | 0.3987377 | 0.232 | 0.153 | 1.46E-31 | Periosteum | rasgef1ba |
| mknk2a | 9.9E-36 | 0.3628292 | 0.314 | 0.22 | 3.22E-31 | Periosteum | mknk2a |
| slc43a3b | 1.3E-35 | 0.6902627 | 0.129 | 0.074 | 4.33E-31 | Periosteum | slc43a3b |
| myo10l3 | 2.6E-35 | 0.5524723 | 0.156 | 0.094 | 8.39E-31 | Periosteum | myo10l3 |
| frem1b | 4.2E-35 | 0.4379768 | 0.129 | 0.073 | 1.36E-30 | Periosteum | frem1b |
| cox4i2 | 4.6E-35 | 0.2705265 | 0.421 | 0.31 | 1.5E-30 | Periosteum | cox4i2 |
| cct2.1 | 5.7E-35 | 0.2586416 | 0.528 | 0.413 | 1.84E-30 | Periosteum | cct2.1 |
| fam120a | 4.1E-34 | 0.4037435 | 0.254 | 0.174 | 1.33E-29 | Periosteum | fam120a |
| aldh9a1a.1 | 1.7E-33 | 0.2109389 | 0.591 | 0.47 | 5.64E-29 | Periosteum | aldh9a1a.1 |
| zfhx4 | 1.8E-33 | 0.264535 | 0.553 | 0.439 | 5.94E-29 | Periosteum | zfhx4 |
| atox1 | 2.6E-33 | 0.3626592 | 0.341 | 0.249 | 8.37E-29 | Periosteum | atox1 |
| npc2 | 4.8E-33 | 0.3133222 | 0.442 | 0.336 | 1.55E-28 | Periosteum | npc2 |
| fkbp9 | 7.6E-33 | 0.2357853 | 0.487 | 0.373 | 2.48E-28 | Periosteum | fkbp9 |
| tfe3a | 1.2E-32 | 0.6131583 | 0.147 | 0.089 | 3.78E-28 | Periosteum | tfe3a |
| nedd4l | 2.7E-32 | 0.5986122 | 0.153 | 0.095 | 8.74E-28 | Periosteum | nedd4l |
| caprin1b | 2.8E-32 | 0.2955503 | 0.388 | 0.287 | 9.11E-28 | Periosteum | caprin1b |
| sort1a | 6.3E-32 | 0.460483 | 0.171 | 0.108 | 2.04E-27 | Periosteum | sort1a |
| idh1 | 1.1E-31 | 0.6481462 | 0.146 | 0.089 | 3.43E-27 | Periosteum | idh1 |
| eya1 | 1.1E-31 | 0.4126128 | 0.198 | 0.129 | 3.44E-27 | Periosteum | eya1 |
| fkbp14 | 1.8E-31 | 0.2988139 | 0.427 | 0.322 | 5.73E-27 | Periosteum | fkbp14 |

| | | | | | | | |
|---|---|---|---|---|---|---|---|
| pik3r2 | 1.8E-31 | 0.3409752 | 0.275 | 0.193 | 5.93E-27 | Periosteum | pik3r2 |
| irf1b | 2.4E-31 | 0.4837986 | 0.174 | 0.112 | 7.87E-27 | Periosteum | irf1b |
| serp1 | 3.1E-31 | 0.3088462 | 0.405 | 0.306 | 1.02E-26 | Periosteum | serp1 |
| rp9 | 3.4E-31 | 0.4901058 | 0.163 | 0.103 | 1.1E-26 | Periosteum | rp9 |
| bambib | 5.3E-31 | 0.6800522 | 0.111 | 0.063 | 1.71E-26 | Periosteum | bambib |
| nectin1b | 5.8E-31 | 0.4737046 | 0.12 | 0.069 | 1.9E-26 | Periosteum | nectin1b |
| smad6b | 6.1E-31 | 0.6004282 | 0.156 | 0.098 | 1.98E-26 | Periosteum | smad6b |
| cxcl8a | 6.8E-31 | 0.7556041 | 0.332 | 0.259 | 2.2E-26 | Periosteum | cxcl8a |
| eif3i | 7.6E-31 | 0.275741 | 0.437 | 0.335 | 2.46E-26 | Periosteum | eif3i |
| cpn1 | 9.4E-31 | 0.3289217 | 0.288 | 0.204 | 3.07E-26 | Periosteum | cpn1 |
| ivns1abpa | 9.9E-31 | 0.2368861 | 0.374 | 0.277 | 3.21E-26 | Periosteum | ivns1abpa |
| cacna1g | 1.1E-30 | 0.4328618 | 0.149 | 0.091 | 3.48E-26 | Periosteum | cacna1g |
| nanos1 | 1.2E-30 | 0.3040646 | 0.28 | 0.198 | 3.94E-26 | Periosteum | nanos1 |
| itgb1a | 1.8E-30 | 0.3664453 | 0.345 | 0.256 | 5.81E-26 | Periosteum | itgb1a |
| dhrs1 | 1.8E-30 | 0.308951 | 0.265 | 0.184 | 5.82E-26 | Periosteum | dhrs1 |
| smad6a | 2.2E-30 | 0.7962246 | 0.101 | 0.057 | 7.24E-26 | Periosteum | smad6a |
| pdia7 | 2.3E-30 | 0.3412026 | 0.208 | 0.138 | 7.56E-26 | Periosteum | pdia7 |
| gpm6bb | 5E-30 | 0.2922351 | 0.426 | 0.326 | 1.62E-25 | Periosteum | gpm6bb |
| ckap4 | 6.7E-30 | 0.270869 | 0.613 | 0.509 | 2.17E-25 | Periosteum | ckap4 |
| rgs12a | 1.8E-29 | 0.5961449 | 0.1 | 0.056 | 5.83E-25 | Periosteum | rgs12a |
| fscn1a | 2.9E-29 | 0.3622701 | 0.245 | 0.171 | 9.47E-25 | Periosteum | fscn1a |
| pld7 | 3.2E-29 | 0.3611823 | 0.215 | 0.146 | 1.04E-24 | Periosteum | pld7 |
| myo1ea | 6.1E-29 | 0.6802175 | 0.127 | 0.077 | 1.98E-24 | Periosteum | myo1ea |
| nuak1b | 6.6E-29 | 0.2595254 | 0.366 | 0.273 | 2.16E-24 | Periosteum | nuak1b |
| ppp1r14bb | 8.1E-29 | 0.3861138 | 0.211 | 0.143 | 2.65E-24 | Periosteum | ppp1r14bb |
| si:dkey-23a13.22 | 1.2E-28 | 0.6533539 | 0.112 | 0.066 | 3.89E-24 | Periosteum | si:dkey-23a13.22 |
| elmo1 | 1.2E-28 | 0.5583981 | 0.144 | 0.09 | 4.04E-24 | Periosteum | elmo1 |
| foxp4 | 3.3E-28 | 0.2966288 | 0.274 | 0.196 | 1.08E-23 | Periosteum | foxp4 |
| gas1b | 3.5E-28 | 0.3254875 | 0.266 | 0.189 | 1.12E-23 | Periosteum | gas1b |
| fkbp3 | 3.6E-28 | 0.2501845 | 0.448 | 0.348 | 1.18E-23 | Periosteum | fkbp3 |
| rcn3 | 5.2E-28 | 0.2838453 | 0.343 | 0.255 | 1.68E-23 | Periosteum | rcn3 |
| rrbp1a | 1E-27 | 0.2427694 | 0.525 | 0.421 | 3.41E-23 | Periosteum | rrbp1a |
| atp6v0e1 | 1.1E-27 | 0.2688186 | 0.465 | 0.366 | 3.55E-23 | Periosteum | atp6v0e1 |
| hsd11b2 | 1.4E-27 | 0.4066512 | 0.142 | 0.089 | 4.71E-23 | Periosteum | hsd11b2 |
| adh5 | 1.7E-27 | 0.2273651 | 0.615 | 0.507 | 5.67E-23 | Periosteum | adh5 |
| scpep1 | 2E-27 | 0.4198238 | 0.188 | 0.126 | 6.35E-23 | Periosteum | scpep1 |
| runx3 | 2.1E-27 | 0.2030031 | 0.354 | 0.263 | 6.82E-23 | Periosteum | runx3 |
| egr2a | 9.2E-27 | 0.5744486 | 0.142 | 0.09 | 2.99E-22 | Periosteum | egr2a |
| setbp1 | 1.1E-26 | 0.284304 | 0.324 | 0.241 | 3.7E-22 | Periosteum | setbp1 |
| znf503 | 2E-26 | 0.3957358 | 0.166 | 0.108 | 6.44E-22 | Periosteum | znf503 |
| eif4ebp3l | 2.9E-26 | 0.4929308 | 0.212 | 0.149 | 9.34E-22 | Periosteum | eif4ebp3l |
| eef1db | 4.2E-26 | 0.2174551 | 0.673 | 0.571 | 1.37E-21 | Periosteum | eef1db |
| si:dkey-238c7.16 | 6.3E-26 | 0.3672261 | 0.226 | 0.159 | 2.04E-21 | Periosteum | si:dkey-238c7.16 |
| atp5po | 7.2E-26 | 0.2085393 | 0.523 | 0.415 | 2.35E-21 | Periosteum | atp5po |
| eif3ha | 1.3E-25 | 0.218807 | 0.516 | 0.411 | 4.14E-21 | Periosteum | eif3ha |
| vdac1 | 1.3E-25 | 0.2538357 | 0.407 | 0.315 | 4.17E-21 | Periosteum | vdac1 |
| tomm20b | 1.8E-25 | 0.2359707 | 0.432 | 0.334 | 5.96E-21 | Periosteum | tomm20b |
| CR759885.1 | 3.2E-25 | 0.3243217 | 0.15 | 0.096 | 1.05E-20 | Periosteum | CR759885.1 |

| | | | | | | | |
|---|---|---|---|---|---|---|---|
| C20H1orf198 | 3.3E-25 | 0.3541455 | 0.247 | 0.178 | 1.08E-20 | Periosteum | C20H1orf198 |
| p3h3 | 4E-25 | 0.387429 | 0.19 | 0.13 | 1.31E-20 | Periosteum | p3h3 |
| tspan18b | 5E-25 | 0.2203377 | 0.201 | 0.136 | 1.63E-20 | Periosteum | tspan18b |
| pdk2b | 6.5E-25 | 0.2660592 | 0.266 | 0.192 | 2.11E-20 | Periosteum | pdk2b |
| bmp2b | 7E-25 | 0.4196425 | 0.149 | 0.097 | 2.27E-20 | Periosteum | bmp2b |
| net1 | 1.2E-24 | 0.2561471 | 0.46 | 0.364 | 3.84E-20 | Periosteum | net1 |
| rpl26 | 1.8E-24 | 0.308491 | 0.215 | 0.15 | 5.93E-20 | Periosteum | rpl26 |
| nfyba | 1.9E-24 | 0.2824189 | 0.246 | 0.176 | 6.05E-20 | Periosteum | nfyba |
| lzts2a | 2E-24 | 0.5650893 | 0.104 | 0.062 | 6.35E-20 | Periosteum | lzts2a |
| prdx4 | 2.6E-24 | 0.2065959 | 0.454 | 0.355 | 8.55E-20 | Periosteum | prdx4 |
| C7H16orf87 | 2.9E-24 | 0.3983948 | 0.178 | 0.122 | 9.55E-20 | Periosteum | C7H16orf87 |
| prss23 | 3.4E-24 | 0.2955511 | 0.266 | 0.194 | 1.1E-19 | Periosteum | prss23 |
| efna1a | 4.1E-24 | 0.304236 | 0.239 | 0.171 | 1.34E-19 | Periosteum | efna1a |
| creb3l1 | 8E-24 | 0.4112452 | 0.162 | 0.109 | 2.61E-19 | Periosteum | creb3l1 |
| ccnd1 | 9.6E-24 | 0.4600877 | 0.117 | 0.073 | 3.14E-19 | Periosteum | ccnd1 |
| aldob | 2E-23 | 0.2389201 | 0.39 | 0.303 | 6.55E-19 | Periosteum | aldob |
| prss16 | 2E-23 | 0.5254763 | 0.133 | 0.086 | 6.57E-19 | Periosteum | prss16 |
| arl5c | 3.7E-23 | 0.3391252 | 0.172 | 0.117 | 1.2E-18 | Periosteum | arl5c |
| epha4a | 3.8E-23 | 0.5503593 | 0.105 | 0.064 | 1.22E-18 | Periosteum | epha4a |
| arvcfb | 7.3E-23 | 0.4125646 | 0.149 | 0.098 | 2.37E-18 | Periosteum | arvcfb |
| coro1ca | 3.4E-22 | 0.2717396 | 0.305 | 0.231 | 1.11E-17 | Periosteum | coro1ca |
| pim2 | 6.7E-22 | 0.2868791 | 0.293 | 0.219 | 2.19E-17 | Periosteum | pim2 |
| tspan36 | 2.9E-21 | 0.3281927 | 0.216 | 0.157 | 9.46E-17 | Periosteum | tspan36 |
| si:dkey-199f5.8 | 3E-21 | 0.2731982 | 0.221 | 0.158 | 9.7E-17 | Periosteum | si:dkey-199f5.8 |
| bambia | 3E-21 | 0.2827254 | 0.2 | 0.143 | 9.72E-17 | Periosteum | bambia |
| SSBP4 | 8.7E-21 | 0.2957728 | 0.169 | 0.117 | 2.82E-16 | Periosteum | SSBP4 |
| fkbp10a | 8.9E-21 | 0.3435222 | 0.184 | 0.13 | 2.88E-16 | Periosteum | fkbp10a |
| dhrs13l1 | 1E-20 | 0.2515804 | 0.207 | 0.147 | 3.36E-16 | Periosteum | dhrs13l1 |
| ahr2 | 1.8E-20 | 0.2018194 | 0.467 | 0.375 | 6E-16 | Periosteum | ahr2 |
| sdr16c5b | 4E-20 | 0.3480123 | 0.192 | 0.137 | 1.3E-15 | Periosteum | sdr16c5b |
| dnpep | 1.8E-19 | 0.3668014 | 0.141 | 0.095 | 5.9E-15 | Periosteum | dnpep |
| zgc:103438 | 2E-19 | 0.3142886 | 0.128 | 0.085 | 6.62E-15 | Periosteum | zgc:103438 |
| deptor | 2.9E-19 | 0.4759684 | 0.106 | 0.068 | 9.54E-15 | Periosteum | deptor |
| pttg1ipb | 6.2E-19 | 0.4431916 | 0.101 | 0.064 | 2.01E-14 | Periosteum | pttg1ipb |
| FP074902.2 | 9.2E-19 | 0.4565923 | 0.108 | 0.07 | 2.98E-14 | Periosteum | FP074902.2 |
| ptges | 9.4E-19 | 0.3910872 | 0.151 | 0.105 | 3.06E-14 | Periosteum | ptges |
| slain2 | 9.6E-19 | 0.322246 | 0.152 | 0.105 | 3.12E-14 | Periosteum | slain2 |
| cav2 | 1.2E-18 | 0.2885494 | 0.225 | 0.167 | 3.89E-14 | Periosteum | cav2 |
| etv5b | 1.7E-18 | 0.347637 | 0.14 | 0.096 | 5.44E-14 | Periosteum | etv5b |
| rgs3a | 1.8E-18 | 0.3269602 | 0.133 | 0.09 | 5.85E-14 | Periosteum | rgs3a |
| si:ch211-213a13.1 | 1.9E-18 | 0.277659 | 0.175 | 0.124 | 6.21E-14 | Periosteum | si:ch211-213a13.1 |
| dpp7 | 1.2E-17 | 0.2019203 | 0.273 | 0.21 | 3.92E-13 | Periosteum | dpp7 |
| jcada | 1.6E-17 | 0.3669078 | 0.164 | 0.117 | 5.31E-13 | Periosteum | jcada |
| pbx1a | 1.7E-17 | 0.2684134 | 0.29 | 0.227 | 5.55E-13 | Periosteum | pbx1a |
| sestd1 | 2E-17 | 0.3209976 | 0.146 | 0.102 | 6.58E-13 | Periosteum | sestd1 |
| nupr1a | 2.4E-17 | 0.2263603 | 0.319 | 0.25 | 7.75E-13 | Periosteum | nupr1a |
| smad7 | 2.5E-17 | 0.2661186 | 0.202 | 0.149 | 8.27E-13 | Periosteum | smad7 |
| rin2 | 2.9E-17 | 0.3277061 | 0.154 | 0.109 | 9.48E-13 | Periosteum | rin2 |

| | | | | | | | |
|---|---|---|---|---|---|---|---|
| socs1a | 4.3E-17 | 0.426444 | 0.166 | 0.12 | 1.4E-12 | Periosteum | socs1a |
| jade1 | 4.7E-17 | 0.3222707 | 0.143 | 0.099 | 1.53E-12 | Periosteum | jade1 |
| hmgn3 | 5.8E-17 | 0.2337234 | 0.212 | 0.157 | 1.9E-12 | Periosteum | hmgn3 |
| afap1 | 6.1E-17 | 0.4852395 | 0.104 | 0.068 | 1.97E-12 | Periosteum | afap1 |
| efna5a | 6.4E-17 | 0.3960546 | 0.112 | 0.075 | 2.09E-12 | Periosteum | efna5a |
| sh3pxd2ab | 7.4E-17 | 0.3154653 | 0.113 | 0.075 | 2.4E-12 | Periosteum | sh3pxd2ab |
| pebp1 | 9.7E-17 | 0.4098343 | 0.106 | 0.07 | 3.17E-12 | Periosteum | pebp1 |
| sorl1 | 9.9E-17 | 0.2829391 | 0.105 | 0.069 | 3.22E-12 | Periosteum | sorl1 |
| bckdk | 1.1E-16 | 0.4072412 | 0.124 | 0.085 | 3.69E-12 | Periosteum | bckdk |
| smad1 | 1.4E-16 | 0.2758042 | 0.153 | 0.108 | 4.67E-12 | Periosteum | smad1 |
| selenom | 1.6E-16 | 0.2149238 | 0.221 | 0.167 | 5.36E-12 | Periosteum | selenom |
| aamp | 2E-16 | 0.2290351 | 0.202 | 0.15 | 6.53E-12 | Periosteum | aamp |
| phactr4b | 2.1E-16 | 0.3492098 | 0.106 | 0.07 | 6.84E-12 | Periosteum | phactr4b |
| ENSDARG00000908 | 2.2E-16 | 0.2692695 | 0.127 | 0.087 | 7.29E-12 | Periosteum | ENSDARG00000090 |
| mrc2 | 3E-16 | 0.3753889 | 0.124 | 0.085 | 9.85E-12 | Periosteum | mrc2 |
| sept7b | 3E-16 | 0.2123964 | 0.252 | 0.193 | 9.91E-12 | Periosteum | sept7b |
| CU984600.1 | 3.4E-16 | 0.210553 | 0.221 | 0.165 | 1.11E-11 | Periosteum | CU984600.1 |
| b3gnt5a | 4E-16 | 0.3250719 | 0.152 | 0.108 | 1.3E-11 | Periosteum | b3gnt5a |
| impdh1b | 4.2E-16 | 0.4803567 | 0.116 | 0.08 | 1.36E-11 | Periosteum | impdh1b |
| gmps | 4.8E-16 | 0.2858615 | 0.18 | 0.133 | 1.55E-11 | Periosteum | gmps |
| adam19a | 6.3E-16 | 0.2239034 | 0.119 | 0.081 | 2.05E-11 | Periosteum | adam19a |
| gsto2 | 6.4E-16 | 0.2121106 | 0.143 | 0.1 | 2.07E-11 | Periosteum | gsto2 |
| rnaset2 | 1E-15 | 0.3135014 | 0.172 | 0.126 | 3.26E-11 | Periosteum | rnaset2 |
| actb2 | 1.1E-15 | 0.2088498 | 0.994 | 0.991 | 3.64E-11 | Periosteum | actb2 |
| csrnp1b | 1.5E-15 | 0.2074053 | 0.238 | 0.182 | 5.03E-11 | Periosteum | csrnp1b |
| chid1 | 2.2E-15 | 0.3638077 | 0.119 | 0.082 | 7.27E-11 | Periosteum | chid1 |
| tnfsf12 | 2.8E-15 | 0.419219 | 0.189 | 0.142 | 8.95E-11 | Periosteum | tnfsf12 |
| smco4 | 3.4E-15 | 0.2414137 | 0.185 | 0.137 | 1.11E-10 | Periosteum | smco4 |
| fam3c | 3.9E-15 | 0.2437331 | 0.139 | 0.099 | 1.27E-10 | Periosteum | fam3c |
| ngdn | 4.3E-15 | 0.2404719 | 0.203 | 0.153 | 1.41E-10 | Periosteum | ngdn |
| rab34a | 4.5E-15 | 0.263854 | 0.158 | 0.115 | 1.48E-10 | Periosteum | rab34a |
| cald1b | 1.2E-14 | 0.502298 | 0.21 | 0.164 | 3.8E-10 | Periosteum | cald1b |
| dhrs3b | 1.6E-14 | 0.2931204 | 0.154 | 0.113 | 5.05E-10 | Periosteum | dhrs3b |
| farp2 | 1.9E-14 | 0.3442659 | 0.152 | 0.112 | 6.28E-10 | Periosteum | farp2 |
| zgc:136864 | 2.3E-14 | 0.3640747 | 0.106 | 0.072 | 7.51E-10 | Periosteum | zgc:136864 |
| kank1a | 3.6E-14 | 0.2263269 | 0.15 | 0.109 | 1.18E-09 | Periosteum | kank1a |
| fhod1 | 5E-14 | 0.4049547 | 0.113 | 0.079 | 1.62E-09 | Periosteum | fhod1 |
| mrpl41 | 7E-14 | 0.3227677 | 0.117 | 0.082 | 2.27E-09 | Periosteum | mrpl41 |
| zgc:153018 | 7.1E-14 | 0.2042431 | 0.286 | 0.227 | 2.29E-09 | Periosteum | zgc:153018 |
| mrpl32 | 7.1E-14 | 0.2155254 | 0.214 | 0.164 | 2.31E-09 | Periosteum | mrpl32 |
| fut8a | 7.9E-14 | 0.2997391 | 0.102 | 0.07 | 2.56E-09 | Periosteum | fut8a |
| palm2akap2.1 | 8E-14 | 0.4489716 | 0.116 | 0.082 | 2.59E-09 | Periosteum | palm2akap2.1 |
| hvcn1 | 8.6E-14 | 0.3050096 | 0.106 | 0.073 | 2.81E-09 | Periosteum | hvcn1 |
| ggcx | 1.1E-13 | 0.2278559 | 0.211 | 0.162 | 3.46E-09 | Periosteum | ggcx |
| slc7a10a | 1.1E-13 | 0.2708811 | 0.143 | 0.104 | 3.71E-09 | Periosteum | slc7a10a |
| ak1 | 1.4E-13 | 0.273161 | 0.164 | 0.123 | 4.63E-09 | Periosteum | ak1 |
| cnn2 | 1.5E-13 | 0.4580918 | 0.497 | 0.44 | 4.77E-09 | Periosteum | cnn2 |
| nos1apa | 1.5E-13 | 0.251407 | 0.145 | 0.105 | 4.86E-09 | Periosteum | nos1apa |

| | | | | | | | |
|---|---|---|---|---|---|---|---|
| aopep | 1.9E-13 | 0.4012165 | 0.105 | 0.073 | 6.13E-09 | Periosteum | aopep |
| zgc:100829 | 3.2E-13 | 0.206088 | 0.157 | 0.116 | 1.03E-08 | Periosteum | zgc:100829 |
| grwd1 | 3.3E-13 | 0.2042353 | 0.182 | 0.138 | 1.08E-08 | Periosteum | grwd1 |
| rab22a | 4.8E-13 | 0.2737861 | 0.136 | 0.099 | 1.55E-08 | Periosteum | rab22a |
| zgc:110796 | 5.7E-13 | 0.224358 | 0.111 | 0.078 | 1.87E-08 | Periosteum | zgc:110796 |
| pds5b | 6.2E-13 | 0.4217082 | 0.107 | 0.075 | 2.01E-08 | Periosteum | pds5b |
| pprc1 | 7.9E-13 | 0.2001934 | 0.153 | 0.113 | 2.57E-08 | Periosteum | pprc1 |
| TST | 9.8E-13 | 0.2030575 | 0.101 | 0.07 | 3.19E-08 | Periosteum | TST |
| cdc42ep4b | 1.7E-12 | 0.2665968 | 0.116 | 0.083 | 5.57E-08 | Periosteum | cdc42ep4b |
| si:ch73-335l21.1 | 2E-12 | 0.2301765 | 0.101 | 0.07 | 6.61E-08 | Periosteum | si:ch73-335l21.1 |
| slc39a13 | 2.3E-12 | 0.2182425 | 0.18 | 0.138 | 7.59E-08 | Periosteum | slc39a13 |
| ctsla | 4.4E-12 | 0.2104845 | 0.763 | 0.717 | 1.42E-07 | Periosteum | ctsla |
| plxnb2a.1 | 6E-12 | 0.248019 | 0.121 | 0.087 | 1.94E-07 | Periosteum | plxnb2a.1 |
| si:ch211-107o10.3 | 7.7E-12 | 0.2668299 | 0.125 | 0.091 | 2.5E-07 | Periosteum | si:ch211-107o10.3 |
| ino80c | 9E-12 | 0.2695245 | 0.104 | 0.074 | 2.91E-07 | Periosteum | ino80c |
| ppp1r9ba | 1.4E-11 | 0.3431828 | 0.102 | 0.072 | 4.46E-07 | Periosteum | ppp1r9ba |
| snx7 | 1.4E-11 | 0.3045688 | 0.103 | 0.074 | 4.68E-07 | Periosteum | snx7 |
| si:dkeyp-113d7.1 | 1.5E-11 | 0.2007726 | 0.182 | 0.141 | 4.76E-07 | Periosteum | si:dkeyp-113d7.1 |
| fbxo9 | 2E-11 | 0.2471776 | 0.109 | 0.078 | 6.4E-07 | Periosteum | fbxo9 |
| arhgap23a | 2.4E-11 | 0.3071647 | 0.116 | 0.085 | 7.76E-07 | Periosteum | arhgap23a |
| rflnb | 2.6E-11 | 0.2200636 | 0.144 | 0.109 | 8.55E-07 | Periosteum | rflnb |
| CU464124.1 | 3.5E-11 | 0.2165107 | 0.136 | 0.101 | 1.13E-06 | Periosteum | CU464124.1 |
| atp10a | 4.3E-11 | 0.243689 | 0.127 | 0.094 | 1.4E-06 | Periosteum | atp10a |
| pdcd4a | 7.1E-11 | 0.2738874 | 0.117 | 0.086 | 2.32E-06 | Periosteum | pdcd4a |
| ERC1 | 7.3E-11 | 0.238751 | 0.104 | 0.075 | 2.37E-06 | Periosteum | ERC1 |
| dusp2 | 9.9E-11 | 0.2190005 | 0.266 | 0.218 | 3.23E-06 | Periosteum | dusp2 |
| cdkn1ba | 2.3E-10 | 0.2966022 | 0.148 | 0.114 | 7.39E-06 | Periosteum | cdkn1ba |
| dohh | 2.3E-10 | 0.2074567 | 0.185 | 0.146 | 7.6E-06 | Periosteum | dohh |
| mrpl23 | 2.4E-10 | 0.2518907 | 0.105 | 0.077 | 7.68E-06 | Periosteum | mrpl23 |
| zgc:194981 | 2.4E-10 | 0.3065937 | 0.108 | 0.079 | 7.94E-06 | Periosteum | zgc:194981 |
| rgs12b | 4.4E-10 | 0.2683227 | 0.116 | 0.086 | 1.43E-05 | Periosteum | rgs12b |
| polr1d | 4.9E-10 | 0.2407731 | 0.112 | 0.083 | 1.6E-05 | Periosteum | polr1d |
| nenf | 1.3E-09 | 0.2581842 | 0.125 | 0.095 | 4.16E-05 | Periosteum | nenf |
| spry2 | 2E-09 | 0.2493327 | 0.104 | 0.076 | 6.38E-05 | Periosteum | spry2 |
| srsf10b | 5.3E-09 | 0.2202197 | 0.112 | 0.084 | 0.000172 | Periosteum | srsf10b |
| six2a | 7.7E-09 | 0.3467504 | 0.128 | 0.099 | 0.000251 | Periosteum | six2a |
| tmem18 | 8.3E-09 | 0.2064139 | 0.113 | 0.086 | 0.000269 | Periosteum | tmem18 |
| cygb1 | 1.3E-08 | 0.4590914 | 0.37 | 0.429 | 0.000408 | Periosteum | cygb1 |
| p4ha1b | 1.6E-08 | 0.2562441 | 0.171 | 0.138 | 0.00053 | Periosteum | p4ha1b |
| cpvl | 2E-08 | 0.2049141 | 0.118 | 0.091 | 0.000655 | Periosteum | cpvl |
| mkrn1 | 2.4E-08 | 0.2065567 | 0.111 | 0.085 | 0.00078 | Periosteum | mkrn1 |
| NDUFB1 | 4.4E-08 | 0.2013733 | 0.119 | 0.092 | 0.001418 | Periosteum | NDUFB1 |
| tpte | 6.2E-08 | 0.2729249 | 0.111 | 0.086 | 0.00201 | Periosteum | tpte |
| tmed3 | 1.9E-06 | 0.2389906 | 0.118 | 0.095 | 0.062448 | Periosteum | tmed3 |
| tpm2 | 2E-06 | 0.4218005 | 0.141 | 0.114 | 0.063628 | Periosteum | tpm2 |
| si:dkeyp-44a8.4 | 2.7E-05 | 0.2661426 | 0.191 | 0.164 | 0.880621 | Periosteum | si:dkeyp-44a8.4 |
| nfatc1 | 3.3E-05 | 0.2278734 | 0.104 | 0.085 | 1 | Periosteum | nfatc1 |
| fgfbp2a | 0.00074 | 0.2660595 | 0.498 | 0.541 | 1 | Periosteum | fgfbp2a |

| | | | | | | | |
|---|---|---|---|---|---|---|---|
| hapln1a | 0 | 1.8449028 | 0.905 | 0.43 | 0 | DF 1 | hapln1a |
| vcanb | 0 | 2.2417959 | 0.898 | 0.424 | 0 | DF 1 | vcanb |
| pah | 0 | 2.0139125 | 0.758 | 0.305 | 0 | DF 1 | pah |
| fgfbp2a1 | 0 | 1.0336189 | 0.94 | 0.491 | 0 | DF 1 | fgfbp2a |
| ptx3a | 0 | 1.5662045 | 0.888 | 0.442 | 0 | DF 1 | ptx3a |
| zgc:194398 | 0 | 1.9025621 | 0.744 | 0.343 | 0 | DF 1 | zgc:194398 |
| postna | 0 | 1.2633014 | 0.946 | 0.551 | 0 | DF 1 | postna |
| hpdb | 0 | 1.7109221 | 0.861 | 0.47 | 0 | DF 1 | hpdb |
| capn12 | 0 | 2.1158668 | 0.574 | 0.187 | 0 | DF 1 | capn12 |
| gyg1b | 0 | 1.6170937 | 0.588 | 0.231 | 0 | DF 1 | gyg1b |
| si:ch73-173h19.3 | 0 | 1.9467801 | 0.559 | 0.204 | 0 | DF 1 | si:ch73-173h19.3 |
| fam169aa | 0 | 1.7312365 | 0.72 | 0.369 | 0 | DF 1 | fam169aa |
| spock3 | 0 | 0.8810418 | 0.711 | 0.379 | 0 | DF 1 | spock3 |
| ptgdsb.1 | 0 | 1.1009058 | 0.755 | 0.431 | 0 | DF 1 | ptgdsb.1 |
| BX663503.3 | 0 | 1.536239 | 0.694 | 0.371 | 0 | DF 1 | BX663503.3 |
| ptgdsb.2 | 0 | 1.224821 | 0.95 | 0.636 | 0 | DF 1 | ptgdsb.2 |
| aqp3a | 0 | 1.3545415 | 0.597 | 0.289 | 0 | DF 1 | aqp3a |
| hgd | 0 | 1.5239493 | 0.705 | 0.399 | 0 | DF 1 | hgd |
| C18H15orf48 | 0 | 1.489173 | 0.583 | 0.28 | 0 | DF 1 | C18H15orf48 |
| plpp1a | 0 | 1.2132802 | 0.635 | 0.347 | 0 | DF 1 | plpp1a |
| qdpra | 0 | 1.3603299 | 0.681 | 0.398 | 0 | DF 1 | qdpra |
| thy1 | 0 | 1.8015945 | 0.479 | 0.204 | 0 | DF 1 | thy1 |
| alx4a | 0 | 1.3836338 | 0.615 | 0.343 | 0 | DF 1 | alx4a |
| cxcl12a | 0 | 0.7597374 | 0.924 | 0.656 | 0 | DF 1 | cxcl12a |
| syt11a | 0 | 2.0388505 | 0.401 | 0.139 | 0 | DF 1 | syt11a |
| vmo1a | 0 | 1.136034 | 0.934 | 0.675 | 0 | DF 1 | vmo1a |
| tnfaip6 | 0 | 1.090336 | 0.665 | 0.406 | 0 | DF 1 | tnfaip6 |
| mfap5 | 0 | 0.9949471 | 0.779 | 0.52 | 0 | DF 1 | mfap5 |
| tat | 0 | 1.5176322 | 0.475 | 0.22 | 0 | DF 1 | tat |
| olfml2ba | 0 | 2.2117167 | 0.345 | 0.101 | 0 | DF 1 | olfml2ba |
| bhmt | 0 | 1.25909 | 0.924 | 0.684 | 0 | DF 1 | bhmt |
| cavin1b | 0 | 1.2382143 | 0.857 | 0.623 | 0 | DF 1 | cavin1b |
| crybg1b | 0 | 1.3105071 | 0.691 | 0.457 | 0 | DF 1 | crybg1b |
| ecm1b | 0 | 1.1250623 | 0.678 | 0.449 | 0 | DF 1 | ecm1b |
| krt15 | 0 | 1.3835881 | 0.907 | 0.678 | 0 | DF 1 | krt15 |
| si:ch211-262i1.4 | 0 | 2.751679 | 0.286 | 0.058 | 0 | DF 1 | si:ch211-262i1.4 |
| gbe1a | 0 | 2.1384982 | 0.311 | 0.09 | 0 | DF 1 | gbe1a |
| prx | 0 | 2.8370671 | 0.261 | 0.044 | 0 | DF 1 | prx |
| slc51a | 0 | 1.7782802 | 0.354 | 0.137 | 0 | DF 1 | slc51a |
| cav1 | 0 | 1.3434831 | 0.82 | 0.612 | 0 | DF 1 | cav1 |
| lxn | 0 | 0.9548921 | 0.887 | 0.681 | 0 | DF 1 | lxn |
| aldh1a3 | 0 | 1.9604452 | 0.287 | 0.092 | 0 | DF 1 | aldh1a3 |
| ryr3 | 0 | 2.1871458 | 0.244 | 0.073 | 0 | DF 1 | ryr3 |
| krt4 | 0 | 1.1730818 | 0.91 | 0.742 | 0 | DF 1 | krt4 |
| elnb | 0 | 2.8850269 | 0.201 | 0.033 | 0 | DF 1 | elnb |
| mxra5b | 0 | 2.6153314 | 0.2 | 0.044 | 0 | DF 1 | mxra5b |
| sned1 | 0 | 2.2792566 | 0.199 | 0.048 | 0 | DF 1 | sned1 |
| lmna | 0 | 0.856911 | 0.897 | 0.768 | 0 | DF 1 | lmna |

| | | | | | | | |
|---|---|---|---|---|---|---|---|
| etnppl | 0 | 3.3940136 | 0.114 | 0.017 | 0 | DF 1 | etnppl |
| rbp4 | 0 | 0.9885594 | 0.994 | 0.9 | 0 | DF 1 | rbp4 |
| krt91 | 0 | 0.8059384 | 0.981 | 0.893 | 0 | DF 1 | krt91 |
| anxa1a | 0 | 1.0185726 | 0.963 | 0.899 | 0 | DF 1 | anxa1a |
| col6a3.1 | 0 | 0.7654894 | 0.992 | 0.932 | 0 | DF 1 | col6a3.1 |
| ahnak | 0 | 1.4712792 | 0.995 | 0.958 | 0 | DF 1 | ahnak |
| s100a10b | 0 | 0.5771346 | 1 | 0.993 | 0 | DF 1 | s100a10b |
| icn | 0 | 1.1475563 | 1 | 0.995 | 0 | DF 1 | icn |
| CR383676.2 | 0 | 0.6004326 | 1 | 1 | 0 | DF 1 | CR383676.2 |
| smoc1 | 3E-293 | 1.5555432 | 0.405 | 0.186 | 9.1E-289 | DF 1 | smoc1 |
| sfrp1b | 6E-285 | 1.0480066 | 0.588 | 0.337 | 1.9E-280 | DF 1 | sfrp1b |
| pawr | 6E-281 | 1.6542599 | 0.374 | 0.167 | 2E-276 | DF 1 | pawr |
| vim | 2E-277 | 1.266511 | 0.58 | 0.347 | 6.6E-273 | DF 1 | vim |
| aoc2 | 3E-275 | 1.449138 | 0.439 | 0.22 | 1E-270 | DF 1 | aoc2 |
| pcolcea | 6E-271 | 0.7824374 | 0.769 | 0.546 | 2.1E-266 | DF 1 | pcolcea |
| gstz1 | 4E-265 | 1.3437796 | 0.531 | 0.324 | 1.4E-260 | DF 1 | gstz1 |
| ccl25b | 3E-264 | 0.3442763 | 0.888 | 0.639 | 1E-259 | DF 1 | ccl25b |
| apoeb | 4E-264 | 1.1350023 | 0.452 | 0.226 | 1.4E-259 | DF 1 | apoeb |
| sema3e | 3E-261 | 1.9125421 | 0.239 | 0.079 | 9.7E-257 | DF 1 | sema3e |
| si:ch73-237c6.1 | 4E-248 | 1.1105499 | 0.56 | 0.344 | 1.2E-243 | DF 1 | si:ch73-237c6.1 |
| NC-002333.4 | 3E-243 | 0.4654956 | 1 | 1 | 1.1E-238 | DF 1 | NC-002333.4 |
| tmsb4x | 7E-240 | 0.3884207 | 0.998 | 0.991 | 2.3E-235 | DF 1 | tmsb4x |
| si:dkey-261h17.1 | 2E-237 | 0.6875344 | 0.5 | 0.268 | 6.7E-233 | DF 1 | si:dkey-261h17.1 |
| igfbp5b | 5E-230 | 0.8349166 | 0.737 | 0.553 | 1.6E-225 | DF 1 | igfbp5b |
| ppl | 3E-225 | 1.6916891 | 0.312 | 0.138 | 1E-220 | DF 1 | ppl |
| anxa4 | 2E-218 | 0.8093503 | 0.787 | 0.696 | 7.5E-214 | DF 1 | anxa4 |
| FO203432.1 | 2E-217 | 2.1290142 | 0.16 | 0.044 | 5.1E-213 | DF 1 | FO203432.1 |
| eppk1 | 8E-215 | 1.4270694 | 0.379 | 0.194 | 2.5E-210 | DF 1 | eppk1 |
| ifitm1 | 2E-214 | 0.4392138 | 0.969 | 0.77 | 6E-210 | DF 1 | ifitm1 |
| ppap2d | 5E-209 | 1.3161296 | 0.399 | 0.211 | 1.6E-204 | DF 1 | ppap2d |
| pleca | 5E-208 | 0.5076322 | 0.965 | 0.912 | 1.7E-203 | DF 1 | pleca |
| slc7a2 | 2E-205 | 1.4548654 | 0.313 | 0.142 | 4.9E-201 | DF 1 | slc7a2 |
| col6a2 | 1E-204 | 0.4795108 | 0.953 | 0.869 | 4E-200 | DF 1 | col6a2 |
| cast | 4E-202 | 0.8995288 | 0.705 | 0.579 | 1.2E-197 | DF 1 | cast |
| clec3ba | 7E-202 | 1.7887859 | 0.265 | 0.112 | 2.3E-197 | DF 1 | clec3ba |
| si:ch211-105c13.3 | 4E-198 | 0.6293194 | 0.735 | 0.551 | 1.4E-193 | DF 1 | si:ch211-105c13.3 |
| itga6b | 3E-197 | 1.1651292 | 0.464 | 0.278 | 9.5E-193 | DF 1 | itga6b |
| kazald31 | 1E-195 | 1.2834911 | 0.426 | 0.243 | 3.1E-191 | DF 1 | kazald3 |
| si:ch211-160j14.31 | 2E-194 | 0.9965491 | 0.495 | 0.295 | 5.7E-190 | DF 1 | si:ch211-160j14.3 |
| myh9b | 4E-189 | 1.2213242 | 0.463 | 0.285 | 1.4E-184 | DF 1 | myh9b |
| cfp | 2E-188 | 0.9234453 | 0.52 | 0.321 | 6.8E-184 | DF 1 | cfp |
| ca6 | 1E-187 | 0.3962222 | 0.567 | 0.323 | 3.8E-183 | DF 1 | ca6 |
| synm | 3E-186 | 1.3570674 | 0.389 | 0.213 | 1.1E-181 | DF 1 | synm |
| ppm1e | 1E-180 | 2.4932886 | 0.108 | 0.025 | 4.7E-176 | DF 1 | ppm1e |
| si:ch211-170d8.8 | 2E-180 | 0.843356 | 0.63 | 0.469 | 5.5E-176 | DF 1 | si:ch211-170d8.8 |
| si:ch73-380l3.2 | 1E-173 | 1.0007272 | 0.569 | 0.4 | 3.3E-169 | DF 1 | si:ch73-380l3.2 |
| ttc36 | 5E-172 | 0.8379472 | 0.53 | 0.335 | 1.5E-167 | DF 1 | ttc36 |
| pcbd1 | 3E-171 | 1.1209242 | 0.473 | 0.3 | 1.1E-166 | DF 1 | pcbd1 |

| | | | | | | | |
|---|---|---|---|---|---|---|---|
| zgc:56585 | 1E-164 | 0.9264705 | 0.493 | 0.31 | 3.2E-160 | DF 1 | zgc:56585 |
| ablim1b | 3E-164 | 1.0808272 | 0.409 | 0.24 | 9.1E-160 | DF 1 | ablim1b |
| neu3.2 | 9E-162 | 1.1893666 | 0.37 | 0.206 | 3.1E-157 | DF 1 | neu3.2 |
| enpp2 | 7E-159 | 0.7701744 | 0.603 | 0.428 | 2.2E-154 | DF 1 | enpp2 |
| agtr2 | 4E-158 | 1.5702677 | 0.236 | 0.103 | 1.4E-153 | DF 1 | agtr2 |
| plpp3 | 3E-149 | 0.8329075 | 0.598 | 0.439 | 8.9E-145 | DF 1 | plpp3 |
| itgb1b.21 | 5E-148 | 1.5339333 | 0.225 | 0.099 | 1.5E-143 | DF 1 | itgb1b.2 |
| si:dkey-85k7.7 | 3E-147 | 0.9371721 | 0.437 | 0.264 | 9.6E-143 | DF 1 | si:dkey-85k7.7 |
| lpl | 3E-145 | 1.5380117 | 0.226 | 0.101 | 1E-140 | DF 1 | lpl |
| b2ml | 2E-143 | 0.3929768 | 0.935 | 0.894 | 4.9E-139 | DF 1 | b2ml |
| aldob1 | 2E-142 | 1.0404737 | 0.448 | 0.297 | 6E-138 | DF 1 | aldob |
| add3a | 9E-142 | 0.7571896 | 0.584 | 0.434 | 3E-137 | DF 1 | add3a |
| ehd2b | 1E-139 | 0.7658625 | 0.667 | 0.549 | 3.2E-135 | DF 1 | ehd2b |
| cavin2b | 1E-139 | 0.7853028 | 0.642 | 0.519 | 4.5E-135 | DF 1 | cavin2b |
| myh10 | 2E-139 | 1.0436138 | 0.467 | 0.321 | 5.7E-135 | DF 1 | myh10 |
| NC-002333.17 | 4E-137 | 0.3491498 | 0.999 | 0.999 | 1.2E-132 | DF 1 | NC-002333.17 |
| prrx1a | 4E-135 | 0.6379086 | 0.695 | 0.575 | 1.3E-130 | DF 1 | prrx1a |
| crhr1 | 4E-134 | 1.7058016 | 0.173 | 0.068 | 1.3E-129 | DF 1 | crhr1 |
| si:ch211-1a19.3 | 5E-132 | 1.0915217 | 0.627 | 0.504 | 1.6E-127 | DF 1 | si:ch211-1a19.3 |
| si:ch211-194e15.5 | 3E-130 | 1.668697 | 0.205 | 0.092 | 8.5E-126 | DF 1 | si:ch211-194e15.5 |
| ly75 | 8E-130 | 1.872946 | 0.158 | 0.061 | 2.7E-125 | DF 1 | ly75 |
| krt18b | 1E-129 | 0.4746877 | 0.937 | 0.886 | 3.3E-125 | DF 1 | krt18b |
| tagln1 | 1E-129 | 0.3748741 | 0.687 | 0.523 | 4E-125 | DF 1 | tagln |
| gpm6aa | 1E-127 | 1.9047432 | 0.137 | 0.049 | 3.4E-123 | DF 1 | gpm6aa |
| has2 | 2E-125 | 1.5351115 | 0.218 | 0.102 | 5.9E-121 | DF 1 | has2 |
| metrnla | 1E-123 | 1.0361883 | 0.438 | 0.302 | 3.5E-119 | DF 1 | metrnla |
| krt8 | 2E-122 | 0.28277 | 0.991 | 0.984 | 5.8E-118 | DF 1 | krt8 |
| ctbp2a | 2E-122 | 1.224681 | 0.351 | 0.218 | 6E-118 | DF 1 | ctbp2a |
| si:dkey-11f4.20 | 1E-121 | 0.7265928 | 0.608 | 0.476 | 3.8E-117 | DF 1 | si:dkey-11f4.20 |
| s100w | 1E-117 | 0.8525174 | 0.55 | 0.421 | 3.9E-113 | DF 1 | s100w |
| slc1a5 | 1E-116 | 1.1312744 | 0.378 | 0.245 | 3.5E-112 | DF 1 | slc1a5 |
| selenow1 | 6E-115 | 0.4237876 | 0.912 | 0.896 | 2E-110 | DF 1 | selenow1 |
| herc1 | 1E-113 | 1.7189528 | 0.191 | 0.089 | 3.2E-109 | DF 1 | herc1 |
| tcf12 | 1E-112 | 0.7158916 | 0.626 | 0.512 | 3.5E-108 | DF 1 | tcf12 |
| notch3 | 3E-112 | 0.7618568 | 0.471 | 0.333 | 1.1E-107 | DF 1 | notch3 |
| sfrp1a | 1E-110 | 0.705923 | 0.564 | 0.424 | 4.2E-106 | DF 1 | sfrp1a |
| klf15 | 2E-110 | 1.2036481 | 0.322 | 0.196 | 8E-106 | DF 1 | klf15 |
| dio1 | 1E-107 | 1.4789216 | 0.221 | 0.112 | 3.1E-103 | DF 1 | dio1 |
| scara5 | 1E-105 | 1.2370761 | 0.258 | 0.14 | 3.3E-101 | DF 1 | scara5 |
| ppp1r9alb | 1E-104 | 1.9410935 | 0.105 | 0.036 | 3.6E-100 | DF 1 | ppp1r9alb |
| ucp2 | 2E-104 | 0.8014556 | 0.533 | 0.403 | 6.7E-100 | DF 1 | ucp2 |
| skia | 1E-103 | 0.7542986 | 0.55 | 0.439 | 3.2E-99 | DF 1 | skia |
| glud1a | 1E-103 | 0.8174839 | 0.535 | 0.431 | 4.5E-99 | DF 1 | glud1a |
| tril | 7E-102 | 1.3914729 | 0.199 | 0.097 | 2.27E-97 | DF 1 | tril |
| ITLN1 | 3E-100 | 1.0962788 | 0.322 | 0.199 | 1.05E-95 | DF 1 | ITLN1 |
| gig2d | 1.7E-98 | 1.7956056 | 0.12 | 0.046 | 5.57E-94 | DF 1 | gig2d |
| nfe2l2a | 2.7E-97 | 0.904068 | 0.449 | 0.333 | 8.66E-93 | DF 1 | nfe2l2a |
| spon2b | 8.8E-97 | 1.5622767 | 0.223 | 0.118 | 2.87E-92 | DF 1 | spon2b |

| | | | | | | | |
|---|---|---|---|---|---|---|---|
| fstb | 1.7E-96 | 0.8129781 | 0.485 | 0.349 | 5.59E-92 | DF 1 | fstb |
| serpinb14 | 1.1E-95 | 1.3388015 | 0.252 | 0.143 | 3.52E-91 | DF 1 | serpinb14 |
| cd81a | 1.2E-94 | 0.318551 | 0.921 | 0.901 | 3.77E-90 | DF 1 | cd81a |
| GNAZ | 1.5E-94 | 1.3024489 | 0.231 | 0.125 | 4.99E-90 | DF 1 | GNAZ |
| pamr1 | 2.8E-91 | 1.3323602 | 0.197 | 0.1 | 9.01E-87 | DF 1 | pamr1 |
| PAMR1 | 2.5E-90 | 1.5448388 | 0.207 | 0.111 | 8.13E-86 | DF 1 | PAMR1 |
| CR384075.1 | 8.5E-90 | 1.6801466 | 0.124 | 0.05 | 2.76E-85 | DF 1 | CR384075.1 |
| hspa12b | 9.7E-90 | 1.300857 | 0.213 | 0.113 | 3.16E-85 | DF 1 | hspa12b |
| soul5 | 2.7E-88 | 0.7991735 | 0.388 | 0.266 | 8.64E-84 | DF 1 | soul5 |
| twist2 | 2.5E-87 | 0.7580778 | 0.451 | 0.329 | 7.99E-83 | DF 1 | twist2 |
| itm2ba | 5.8E-87 | 0.4621072 | 0.772 | 0.739 | 1.88E-82 | DF 1 | itm2ba |
| itga9 | 2.6E-86 | 1.2972049 | 0.216 | 0.118 | 8.46E-82 | DF 1 | itga9 |
| thrap3a | 1.7E-85 | 0.793171 | 0.48 | 0.374 | 5.64E-81 | DF 1 | thrap3a |
| lsp1b | 2.4E-84 | 0.9302291 | 0.363 | 0.247 | 7.72E-80 | DF 1 | lsp1b |
| gstm.1 | 6.3E-84 | 0.5338088 | 0.67 | 0.589 | 2.06E-79 | DF 1 | gstm.1 |
| gch1 | 1.2E-83 | 1.1918833 | 0.227 | 0.126 | 4.01E-79 | DF 1 | gch1 |
| stmn1b | 2.2E-83 | 1.3074389 | 0.211 | 0.114 | 7.03E-79 | DF 1 | stmn1b |
| wnk1b | 1.7E-82 | 1.4406419 | 0.212 | 0.118 | 5.69E-78 | DF 1 | wnk1b |
| arhgap36 | 2.9E-82 | 1.5142782 | 0.129 | 0.056 | 9.46E-78 | DF 1 | arhgap36 |
| actb21 | 8.1E-81 | 0.2306893 | 0.994 | 0.991 | 2.63E-76 | DF 1 | actb2 |
| myo1ca | 6E-79 | 1.0348064 | 0.299 | 0.193 | 1.95E-74 | DF 1 | myo1ca |
| pcdh18a | 1.4E-78 | 1.3445349 | 0.156 | 0.075 | 4.45E-74 | DF 1 | pcdh18a |
| gpc6a | 2.2E-78 | 0.9885632 | 0.299 | 0.192 | 7.11E-74 | DF 1 | gpc6a |
| tmsb1 | 9E-78 | 0.444592 | 0.556 | 0.439 | 2.91E-73 | DF 1 | tmsb1 |
| slc12a2 | 6.9E-77 | 1.2263462 | 0.23 | 0.135 | 2.24E-72 | DF 1 | slc12a2 |
| tnc1 | 3.7E-76 | 0.5720636 | 0.583 | 0.458 | 1.2E-71 | DF 1 | tnc |
| zgc:153031 | 5.9E-76 | 1.6054712 | 0.125 | 0.056 | 1.92E-71 | DF 1 | zgc:153031 |
| fbp1a | 5.9E-76 | 0.9396421 | 0.369 | 0.268 | 1.92E-71 | DF 1 | fbp1a |
| loxl1 | 7.7E-75 | 0.8103262 | 0.301 | 0.19 | 2.51E-70 | DF 1 | loxl1 |
| ebf3a | 1.4E-73 | 1.0114414 | 0.308 | 0.206 | 4.4E-69 | DF 1 | ebf3a |
| tob1b | 1.7E-73 | 0.4394706 | 0.759 | 0.697 | 5.63E-69 | DF 1 | tob1b |
| angptl2a | 7.6E-72 | 0.5098362 | 0.431 | 0.307 | 2.47E-67 | DF 1 | angptl2a |
| slco2b1 | 8.6E-72 | 0.8760722 | 0.319 | 0.211 | 2.78E-67 | DF 1 | slco2b1 |
| slc12a7b | 1.4E-71 | 1.1131686 | 0.237 | 0.143 | 4.48E-67 | DF 1 | slc12a7b |
| ppdpfb | 3.6E-71 | 0.4008552 | 0.803 | 0.776 | 1.19E-66 | DF 1 | ppdpfb |
| tuft1a | 5.9E-71 | 0.5871258 | 0.655 | 0.585 | 1.91E-66 | DF 1 | tuft1a |
| si:dkey-40c11.2 | 1.1E-70 | 0.8295449 | 0.406 | 0.308 | 3.42E-66 | DF 1 | si:dkey-40c11.2 |
| slc48a1b | 1.2E-69 | 1.2268656 | 0.222 | 0.133 | 4.04E-65 | DF 1 | slc48a1b |
| si:ch211-183d21.1 | 1.4E-69 | 0.8011782 | 0.468 | 0.373 | 4.64E-65 | DF 1 | si:ch211-183d21.1 |
| aldh9a1a.11 | 8.7E-69 | 0.7309702 | 0.546 | 0.475 | 2.83E-64 | DF 1 | aldh9a1a.1 |
| mao | 8.8E-69 | 1.3395573 | 0.137 | 0.066 | 2.86E-64 | DF 1 | mao |
| ism2a | 1.5E-68 | 1.5399071 | 0.108 | 0.046 | 4.81E-64 | DF 1 | ism2a |
| slc22a2 | 1.5E-68 | 0.8847465 | 0.287 | 0.185 | 5.01E-64 | DF 1 | slc22a2 |
| myof | 1.6E-67 | 1.2371926 | 0.217 | 0.13 | 5.3E-63 | DF 1 | myof |
| pdlim5a | 2.6E-67 | 1.5825243 | 0.109 | 0.048 | 8.55E-63 | DF 1 | pdlim5a |
| vegfc | 9.4E-67 | 1.3409739 | 0.157 | 0.081 | 3.06E-62 | DF 1 | vegfc |
| mhc1zaa | 3.3E-66 | 1.2758885 | 0.163 | 0.086 | 1.07E-61 | DF 1 | mhc1zaa |
| dpysl2b | 7.4E-66 | 1.1078013 | 0.243 | 0.153 | 2.41E-61 | DF 1 | dpysl2b |

| | | | | | | | |
|---|---|---|---|---|---|---|---|
| pthlha | 1.4E-65 | 1.3444624 | 0.135 | 0.065 | 4.54E-61 | DF 1 | pthlha |
| sf3b1 | 2.4E-65 | 0.8066585 | 0.418 | 0.33 | 7.7E-61 | DF 1 | sf3b1 |
| gpt2l | 8.8E-65 | 1.2229856 | 0.205 | 0.122 | 2.86E-60 | DF 1 | gpt2l |
| gpc1b | 1.5E-63 | 0.7161574 | 0.481 | 0.394 | 4.9E-59 | DF 1 | gpc1b |
| mlip | 3.9E-63 | 1.4393948 | 0.135 | 0.067 | 1.28E-58 | DF 1 | mlip |
| abi3bpa | 7.6E-63 | 0.2213321 | 0.42 | 0.287 | 2.46E-58 | DF 1 | abi3bpa |
| si:dkey-240h12.4 | 5E-62 | 1.1143164 | 0.192 | 0.11 | 1.63E-57 | DF 1 | si:dkey-240h12.4 |
| si:ch211-63p21.1 | 1.2E-60 | 1.4935071 | 0.102 | 0.045 | 3.74E-56 | DF 1 | si:ch211-63p21.1 |
| CABZ01092746.1 | 7.3E-60 | 0.3024445 | 0.809 | 0.699 | 2.39E-55 | DF 1 | CABZ01092746.1 |
| cebpd | 1.6E-59 | 0.4601354 | 0.645 | 0.57 | 5.16E-55 | DF 1 | cebpd |
| itgb4 | 1.8E-59 | 0.6816555 | 0.401 | 0.303 | 5.69E-55 | DF 1 | itgb4 |
| KCTD15 | 2.4E-59 | 0.8077098 | 0.417 | 0.335 | 7.77E-55 | DF 1 | KCTD15 |
| gpatch8 | 1E-58 | 0.7586872 | 0.408 | 0.321 | 3.34E-54 | DF 1 | gpatch8 |
| pmp22b | 1.2E-58 | 0.7763592 | 0.44 | 0.354 | 3.76E-54 | DF 1 | pmp22b |
| prrx1b | 1.5E-58 | 0.6392135 | 0.465 | 0.371 | 5E-54 | DF 1 | prrx1b |
| plecb | 2.2E-58 | 0.4270981 | 0.704 | 0.653 | 7.05E-54 | DF 1 | plecb |
| axin2 | 3.3E-58 | 0.827699 | 0.305 | 0.211 | 1.06E-53 | DF 1 | axin2 |
| fgfr21 | 3.6E-58 | 0.7029146 | 0.43 | 0.34 | 1.18E-53 | DF 1 | fgfr2 |
| cdk6 | 3.9E-58 | 1.4349307 | 0.122 | 0.06 | 1.25E-53 | DF 1 | cdk6 |
| emilin2b | 3.6E-57 | 1.1213082 | 0.127 | 0.063 | 1.18E-52 | DF 1 | emilin2b |
| cdh2 | 5.9E-57 | 0.7478194 | 0.373 | 0.284 | 1.91E-52 | DF 1 | cdh2 |
| tmod2 | 9.6E-57 | 0.991427 | 0.248 | 0.164 | 3.11E-52 | DF 1 | tmod2 |
| b2m | 1.2E-56 | 0.5285288 | 0.716 | 0.683 | 3.75E-52 | DF 1 | b2m |
| zgc:195173 | 2.3E-56 | 0.60625 | 0.477 | 0.389 | 7.48E-52 | DF 1 | zgc:195173 |
| zcchc24 | 2.3E-56 | 0.79156 | 0.339 | 0.248 | 7.55E-52 | DF 1 | zcchc24 |
| rnf213a | 1.4E-55 | 0.8221186 | 0.349 | 0.261 | 4.4E-51 | DF 1 | rnf213a |
| arid5b | 1.5E-55 | 1.0826496 | 0.208 | 0.128 | 4.88E-51 | DF 1 | arid5b |
| actn11 | 3.6E-55 | 0.7449223 | 0.395 | 0.31 | 1.16E-50 | DF 1 | actn1 |
| emid1 | 5.5E-55 | 0.6594315 | 0.216 | 0.132 | 1.78E-50 | DF 1 | emid1 |
| igsf21a | 7.4E-55 | 1.4825589 | 0.111 | 0.053 | 2.41E-50 | DF 1 | igsf21a |
| top1 | 7.6E-55 | 1.2922048 | 0.142 | 0.076 | 2.46E-50 | DF 1 | top1 |
| tgfb1a | 8.8E-55 | 0.7188832 | 0.356 | 0.266 | 2.86E-50 | DF 1 | tgfb1a |
| lhfpl6 | 1.2E-53 | 0.6263135 | 0.484 | 0.411 | 3.86E-49 | DF 1 | lhfpl6 |
| cdon | 1.4E-53 | 0.9078078 | 0.265 | 0.18 | 4.49E-49 | DF 1 | cdon |
| cd248a | 2.3E-53 | 0.9127883 | 0.186 | 0.109 | 7.49E-49 | DF 1 | cd248a |
| scinlb | 2.4E-53 | 0.7867461 | 0.381 | 0.301 | 7.95E-49 | DF 1 | scinlb |
| arl6ip5b | 2.6E-53 | 0.6116649 | 0.554 | 0.506 | 8.56E-49 | DF 1 | arl6ip5b |
| rn7sk | 1.2E-50 | 0.5968476 | 0.523 | 0.449 | 3.98E-46 | DF 1 | rn7sk |
| itgb2 | 1.3E-50 | 1.176167 | 0.169 | 0.099 | 4.24E-46 | DF 1 | itgb2 |
| enah | 1.4E-50 | 1.1009909 | 0.18 | 0.109 | 4.6E-46 | DF 1 | enah |
| edil3a | 2E-50 | 0.4668028 | 0.515 | 0.427 | 6.66E-46 | DF 1 | edil3a |
| ARHGAP6 | 2.6E-50 | 1.2754108 | 0.116 | 0.059 | 8.4E-46 | DF 1 | ARHGAP6 |
| si:cabz01069012.2 | 8.2E-50 | 1.1209715 | 0.165 | 0.097 | 2.65E-45 | DF 1 | si:cabz01069012.2 |
| cd82a1 | 1E-49 | 0.5967956 | 0.432 | 0.347 | 3.34E-45 | DF 1 | cd82a |
| mt-co3 | 2.1E-49 | 0.2103087 | 0.998 | 0.996 | 6.99E-45 | DF 1 | mt-co3 |
| rspo11 | 2.6E-49 | 1.2229307 | 0.115 | 0.058 | 8.42E-45 | DF 1 | rspo1 |
| slc29a1a | 3.1E-49 | 0.9283303 | 0.27 | 0.191 | 1.01E-44 | DF 1 | slc29a1a |
| cgnl1 | 1.4E-48 | 1.2177532 | 0.126 | 0.067 | 4.48E-44 | DF 1 | cgnl1 |

| | | | | | | | |
|---|---|---|---|---|---|---|---|
| tnfsf10 | 4.1E-48 | 1.0873335 | 0.159 | 0.092 | 1.33E-43 | DF 1 | tnfsf10 |
| dchs1b | 4.9E-48 | 1.2140363 | 0.149 | 0.085 | 1.59E-43 | DF 1 | dchs1b |
| wnt11 | 7.3E-48 | 0.7486285 | 0.362 | 0.276 | 2.38E-43 | DF 1 | wnt11 |
| gpr143 | 9E-48 | 1.25064 | 0.123 | 0.065 | 2.94E-43 | DF 1 | gpr143 |
| PPM1K | 1.9E-47 | 1.3607046 | 0.113 | 0.058 | 6.29E-43 | DF 1 | PPM1K |
| dpyda.1 | 2E-47 | 1.119586 | 0.173 | 0.105 | 6.36E-43 | DF 1 | dpyda.1 |
| igf2a | 4.1E-47 | 1.1188164 | 0.151 | 0.086 | 1.34E-42 | DF 1 | igf2a |
| ktn1 | 5.3E-47 | 0.8356038 | 0.34 | 0.266 | 1.72E-42 | DF 1 | ktn1 |
| colec12 | 1.8E-46 | 0.5231592 | 0.501 | 0.432 | 5.71E-42 | DF 1 | colec12 |
| si:dkey-16p21.8 | 4.8E-45 | 0.7920868 | 0.281 | 0.202 | 1.57E-40 | DF 1 | si:dkey-16p21.8 |
| gpm6bb1 | 7.2E-45 | 0.6745571 | 0.4 | 0.329 | 2.33E-40 | DF 1 | gpm6bb |
| fosb | 5E-44 | 0.4784487 | 0.715 | 0.69 | 1.63E-39 | DF 1 | fosb |
| cd99 | 3E-43 | 0.3230395 | 0.751 | 0.724 | 9.74E-39 | DF 1 | cd99 |
| fibinb | 9.7E-43 | 0.5582595 | 0.351 | 0.267 | 3.16E-38 | DF 1 | fibinb |
| plcd4b | 9.7E-43 | 1.226002 | 0.133 | 0.076 | 3.16E-38 | DF 1 | plcd4b |
| tmeff1b | 1.3E-42 | 1.2446117 | 0.162 | 0.1 | 4.09E-38 | DF 1 | tmeff1b |
| mgst1.1 | 1.8E-42 | 0.7699676 | 0.317 | 0.244 | 5.85E-38 | DF 1 | mgst1.1 |
| srgap3 | 5.3E-42 | 0.865123 | 0.268 | 0.196 | 1.73E-37 | DF 1 | srgap3 |
| adam10a | 8.2E-42 | 0.4969658 | 0.445 | 0.375 | 2.67E-37 | DF 1 | adam10a |
| gsta.1 | 1.5E-41 | 0.6043769 | 0.42 | 0.351 | 4.75E-37 | DF 1 | gsta.1 |
| capn2b | 1.5E-41 | 0.6187194 | 0.391 | 0.315 | 4.86E-37 | DF 1 | capn2b |
| pcm1 | 4.6E-41 | 0.7085489 | 0.22 | 0.146 | 1.51E-36 | DF 1 | pcm1 |
| hsd3b7 | 1.2E-40 | 1.073935 | 0.222 | 0.153 | 3.94E-36 | DF 1 | hsd3b7 |
| FQ323156.1 | 1.9E-40 | 0.7850209 | 0.262 | 0.189 | 6.04E-36 | DF 1 | FQ323156.1 |
| ildr2 | 3.1E-40 | 1.3313806 | 0.106 | 0.057 | 1.02E-35 | DF 1 | ildr2 |
| svila | 3.3E-40 | 0.8867102 | 0.211 | 0.142 | 1.09E-35 | DF 1 | svila |
| MYADM.1 | 5.5E-40 | 0.7154891 | 0.3 | 0.223 | 1.78E-35 | DF 1 | MYADM.1 |
| ndfip2 | 9E-40 | 0.7502837 | 0.346 | 0.281 | 2.93E-35 | DF 1 | ndfip2 |
| kcne4 | 1.2E-39 | 0.9940745 | 0.107 | 0.057 | 3.78E-35 | DF 1 | kcne4 |
| tead1b | 1.2E-39 | 0.7907472 | 0.352 | 0.287 | 3.8E-35 | DF 1 | tead1b |
| tmem200b | 1.2E-39 | 1.020667 | 0.148 | 0.089 | 3.83E-35 | DF 1 | tmem200b |
| rtn1a | 2.3E-39 | 0.8026293 | 0.286 | 0.216 | 7.52E-35 | DF 1 | rtn1a |
| sfxn5b | 2.6E-39 | 1.0075823 | 0.197 | 0.132 | 8.45E-35 | DF 1 | sfxn5b |
| hyal2b | 3E-39 | 0.8188923 | 0.261 | 0.191 | 9.7E-35 | DF 1 | hyal2b |
| LO018188.1 | 3.3E-39 | 0.4205681 | 0.533 | 0.473 | 1.06E-34 | DF 1 | LO018188.1 |
| celf2 | 1E-38 | 0.6907553 | 0.266 | 0.191 | 3.41E-34 | DF 1 | celf2 |
| ak4 | 4.6E-38 | 0.8541399 | 0.246 | 0.179 | 1.49E-33 | DF 1 | ak4 |
| anxa11a | 1.1E-37 | 0.6056079 | 0.458 | 0.406 | 3.62E-33 | DF 1 | anxa11a |
| zeb2a | 6.4E-37 | 0.5531511 | 0.404 | 0.335 | 2.08E-32 | DF 1 | zeb2a |
| ccdc80l1 | 3.4E-36 | 1.0709879 | 0.127 | 0.074 | 1.12E-31 | DF 1 | ccdc80l1 |
| asip11 | 3.6E-36 | 0.570424 | 0.243 | 0.171 | 1.17E-31 | DF 1 | asip1 |
| itga5 | 7E-36 | 0.7353859 | 0.319 | 0.251 | 2.28E-31 | DF 1 | itga5 |
| mmp15a | 1.8E-35 | 1.1106058 | 0.131 | 0.079 | 5.88E-31 | DF 1 | mmp15a |
| tspan18b1 | 2.6E-35 | 0.9217811 | 0.198 | 0.136 | 8.5E-31 | DF 1 | tspan18b |
| tmem59l | 3.9E-35 | 1.0362407 | 0.158 | 0.101 | 1.27E-30 | DF 1 | tmem59l |
| pdcd6 | 6.4E-35 | 0.4242492 | 0.518 | 0.473 | 2.08E-30 | DF 1 | pdcd6 |
| si:ch73-1a9.3 | 9.5E-35 | 0.359586 | 0.725 | 0.72 | 3.1E-30 | DF 1 | si:ch73-1a9.3 |
| flna | 3.8E-34 | 0.3113507 | 0.593 | 0.539 | 1.23E-29 | DF 1 | flna |

| | | | | | | | |
|---|---|---|---|---|---|---|---|
| hmgb1a | 5.4E-34 | 0.4398562 | 0.592 | 0.562 | 1.75E-29 | DF 1 | hmgb1a |
| madd | 5.5E-34 | 1.307421 | 0.105 | 0.059 | 1.79E-29 | DF 1 | madd |
| dcbld2 | 6.3E-34 | 0.8678172 | 0.179 | 0.119 | 2.05E-29 | DF 1 | dcbld2 |
| CR383676.3 | 7.9E-34 | 0.5948574 | 0.505 | 0.462 | 2.56E-29 | DF 1 | CR383676.3 |
| psd2 | 1.3E-33 | 1.0925858 | 0.146 | 0.093 | 4.36E-29 | DF 1 | psd2 |
| tshz1 | 3E-33 | 1.0124451 | 0.189 | 0.131 | 9.71E-29 | DF 1 | tshz1 |
| fgl2a | 7.1E-33 | 0.5857114 | 0.219 | 0.152 | 2.29E-28 | DF 1 | fgl2a |
| cst3 | 1.8E-32 | 0.2629188 | 0.732 | 0.724 | 5.97E-28 | DF 1 | cst3 |
| tnrc18 | 4.7E-32 | 0.5257465 | 0.453 | 0.403 | 1.54E-27 | DF 1 | tnrc18 |
| lpar1 | 4.8E-32 | 0.8830716 | 0.236 | 0.177 | 1.56E-27 | DF 1 | lpar1 |
| tspan31 | 6.3E-32 | 0.7403032 | 0.282 | 0.221 | 2.05E-27 | DF 1 | tspan31 |
| ackr3b | 8.2E-32 | 0.709999 | 0.172 | 0.113 | 2.68E-27 | DF 1 | ackr3b |
| si:ch211-110e21.3 | 1E-31 | 1.2097136 | 0.12 | 0.072 | 3.26E-27 | DF 1 | si:ch211-110e21.3 |
| wsb1 | 1.7E-31 | 0.9148794 | 0.24 | 0.182 | 5.65E-27 | DF 1 | wsb1 |
| gchfr | 1.8E-31 | 0.8476187 | 0.217 | 0.158 | 5.76E-27 | DF 1 | gchfr |
| slc35c2 | 2.3E-31 | 0.9347396 | 0.204 | 0.146 | 7.36E-27 | DF 1 | slc35c2 |
| rbm5 | 9.3E-31 | 0.7746087 | 0.24 | 0.18 | 3.02E-26 | DF 1 | rbm5 |
| rtn3 | 1E-30 | 0.7829245 | 0.279 | 0.222 | 3.27E-26 | DF 1 | rtn3 |
| kitlga | 1.3E-30 | 1.0562288 | 0.113 | 0.067 | 4.08E-26 | DF 1 | kitlga |
| kctd1 | 1.3E-30 | 1.145796 | 0.122 | 0.075 | 4.09E-26 | DF 1 | kctd1 |
| arhgdia | 2.9E-30 | 0.5492194 | 0.447 | 0.409 | 9.51E-26 | DF 1 | arhgdia |
| cobll1b | 5E-30 | 1.0104728 | 0.103 | 0.059 | 1.62E-25 | DF 1 | cobll1b |
| snai21 | 2.6E-29 | 1.0051827 | 0.152 | 0.101 | 8.32E-25 | DF 1 | snai2 |
| si:ch73-86n18.1 | 3.5E-29 | 0.8063409 | 0.208 | 0.149 | 1.13E-24 | DF 1 | si:ch73-86n18.1 |
| nrp2a | 4.4E-29 | 0.984859 | 0.176 | 0.124 | 1.45E-24 | DF 1 | nrp2a |
| acap1 | 1.8E-28 | 1.129478 | 0.124 | 0.078 | 5.71E-24 | DF 1 | acap1 |
| il15l | 2.2E-28 | 0.8219202 | 0.167 | 0.114 | 7.29E-24 | DF 1 | il15l |
| emilin2a | 2.5E-28 | 0.8851336 | 0.168 | 0.115 | 8.15E-24 | DF 1 | emilin2a |
| nfil3-6 | 2.5E-28 | 0.4970307 | 0.368 | 0.3 | 8.24E-24 | DF 1 | nfil3-6 |
| adamts15a | 3.3E-28 | 0.8277117 | 0.195 | 0.138 | 1.06E-23 | DF 1 | adamts15a |
| heyl | 4.2E-28 | 0.9841357 | 0.112 | 0.068 | 1.35E-23 | DF 1 | heyl |
| si:dkey-164f24.21 | 9.7E-28 | 0.3950005 | 0.229 | 0.164 | 3.15E-23 | DF 1 | si:dkey-164f24.2 |
| rfx2 | 1.2E-27 | 1.1684974 | 0.109 | 0.067 | 4.04E-23 | DF 1 | rfx2 |
| rpz5 | 1.4E-27 | 0.4439379 | 0.368 | 0.301 | 4.66E-23 | DF 1 | rpz5 |
| slc16a1b | 2E-27 | 0.9898052 | 0.177 | 0.127 | 6.48E-23 | DF 1 | slc16a1b |
| CABZ01074397.1 | 2.2E-27 | 0.8723136 | 0.198 | 0.145 | 7.3E-23 | DF 1 | CABZ01074397.1 |
| ncor1 | 3.8E-27 | 0.6826415 | 0.328 | 0.278 | 1.24E-22 | DF 1 | ncor1 |
| crybg1a | 4.8E-27 | 0.7719906 | 0.269 | 0.215 | 1.56E-22 | DF 1 | crybg1a |
| ogt.1.1 | 6.5E-27 | 0.9285676 | 0.158 | 0.109 | 2.12E-22 | DF 1 | ogt.1.1 |
| asah1b | 7.5E-27 | 0.861168 | 0.234 | 0.18 | 2.43E-22 | DF 1 | asah1b |
| ccnd2a | 1.8E-26 | 0.7754248 | 0.267 | 0.213 | 5.75E-22 | DF 1 | ccnd2a |
| si:dkeyp-23e4.3 | 4.3E-26 | 0.8927056 | 0.127 | 0.082 | 1.39E-21 | DF 1 | si:dkeyp-23e4.3 |
| rnd3b | 9E-26 | 0.6070205 | 0.37 | 0.328 | 2.92E-21 | DF 1 | rnd3b |
| tln1 | 1.5E-25 | 0.3165143 | 0.62 | 0.611 | 4.75E-21 | DF 1 | tln1 |
| flii | 1.6E-25 | 1.0362549 | 0.152 | 0.105 | 5.23E-21 | DF 1 | flii |
| mdh1aa | 1.9E-25 | 0.7181234 | 0.401 | 0.372 | 6.06E-21 | DF 1 | mdh1aa |
| prrg1 | 2.5E-25 | 0.7837569 | 0.253 | 0.203 | 8.05E-21 | DF 1 | prrg1 |
| aff2 | 3.5E-25 | 0.8011797 | 0.227 | 0.174 | 1.14E-20 | DF 1 | aff2 |

| | | | | | | | |
|---|---|---|---|---|---|---|---|
| mt-nd4 | 6E-25 | 0.2124821 | 0.866 | 0.856 | 1.95E-20 | DF 1 | mt-nd4 |
| ndrg2 | 7.3E-25 | 0.5978932 | 0.363 | 0.319 | 2.38E-20 | DF 1 | ndrg2 |
| emilin1b | 9.3E-25 | 0.9655675 | 0.126 | 0.083 | 3.04E-20 | DF 1 | emilin1b |
| peli1a | 1.9E-24 | 0.8075232 | 0.175 | 0.124 | 6.07E-20 | DF 1 | peli1a |
| lrrc15 | 2.8E-24 | 0.8294929 | 0.18 | 0.13 | 9.16E-20 | DF 1 | lrrc15 |
| sept7a | 2.9E-24 | 0.6351792 | 0.391 | 0.359 | 9.27E-20 | DF 1 | sept7a |
| capgb | 3.5E-24 | 0.8353622 | 0.145 | 0.099 | 1.13E-19 | DF 1 | capgb |
| s100u | 4.9E-24 | 0.3383996 | 0.551 | 0.516 | 1.58E-19 | DF 1 | s100u |
| peli3 | 1.1E-23 | 1.1133968 | 0.102 | 0.063 | 3.49E-19 | DF 1 | peli3 |
| zfp36l1a | 1.5E-23 | 0.729625 | 0.253 | 0.202 | 4.72E-19 | DF 1 | zfp36l1a |
| myofl | 1.7E-23 | 0.7860519 | 0.205 | 0.155 | 5.38E-19 | DF 1 | myofl |
| zyx | 2E-23 | 0.5147775 | 0.363 | 0.314 | 6.36E-19 | DF 1 | zyx |
| fxyd1 | 2.5E-23 | 0.280099 | 0.595 | 0.571 | 8.07E-19 | DF 1 | fxyd1 |
| fbxo34 | 2.8E-23 | 0.9598833 | 0.129 | 0.086 | 8.99E-19 | DF 1 | fbxo34 |
| serpina10a | 3.6E-23 | 0.4571679 | 0.416 | 0.375 | 1.16E-18 | DF 1 | serpina10a |
| ldhba | 4.1E-23 | 0.461923 | 0.506 | 0.493 | 1.34E-18 | DF 1 | ldhba |
| ccdc125 | 4.4E-23 | 0.9141941 | 0.112 | 0.071 | 1.45E-18 | DF 1 | ccdc125 |
| agrn | 6.9E-23 | 0.6893372 | 0.231 | 0.18 | 2.26E-18 | DF 1 | agrn |
| mhc1zba | 9.6E-23 | 0.326529 | 0.595 | 0.584 | 3.11E-18 | DF 1 | mhc1zba |
| si:ch1073-291c23.2 | 9.7E-23 | 0.3244921 | 0.509 | 0.472 | 3.16E-18 | DF 1 | si:ch1073-291c23.2 |
| ENSDARG0000005970 | 1.3E-22 | 1.027121 | 0.122 | 0.081 | 4.26E-18 | DF 1 | ENSDARG00000059 |
| si:ch211-281l24.3 | 1.4E-22 | 0.7299923 | 0.265 | 0.218 | 4.41E-18 | DF 1 | si:ch211-281l24.3 |
| jund | 1.8E-22 | 0.3917772 | 0.584 | 0.569 | 5.89E-18 | DF 1 | jund |
| capns1b | 2.2E-22 | 0.4194849 | 0.509 | 0.495 | 7.27E-18 | DF 1 | capns1b |
| thbs2b | 5E-22 | 0.3807071 | 0.304 | 0.246 | 1.62E-17 | DF 1 | thbs2b |
| svild | 5.6E-22 | 0.6750958 | 0.299 | 0.252 | 1.83E-17 | DF 1 | svild |
| fh | 1.2E-21 | 1.0794536 | 0.104 | 0.067 | 3.96E-17 | DF 1 | fh |
| capn2a | 1.6E-21 | 0.6141776 | 0.389 | 0.36 | 5.08E-17 | DF 1 | capn2a |
| soga1 | 1.7E-21 | 0.9251397 | 0.167 | 0.123 | 5.68E-17 | DF 1 | soga1 |
| si:ch73-366l1.5 | 1.8E-21 | 1.0409418 | 0.118 | 0.079 | 5.77E-17 | DF 1 | si:ch73-366l1.5 |
| si:ch73-335l21.11 | 1.8E-21 | 1.0801474 | 0.107 | 0.069 | 5.84E-17 | DF 1 | si:ch73-335l21.1 |
| mfap2 | 2E-21 | 0.4071365 | 0.428 | 0.38 | 6.35E-17 | DF 1 | mfap2 |
| atp2b4 | 2E-21 | 0.5384664 | 0.388 | 0.353 | 6.5E-17 | DF 1 | atp2b4 |
| CPNE8 | 2E-20 | 0.5406785 | 0.328 | 0.283 | 6.38E-16 | DF 1 | CPNE8 |
| ddx6 | 2.4E-20 | 0.5293274 | 0.38 | 0.346 | 7.82E-16 | DF 1 | ddx6 |
| kank2 | 3.3E-20 | 0.6378766 | 0.255 | 0.208 | 1.06E-15 | DF 1 | kank2 |
| si:ch211-57n23.1 | 3.5E-20 | 0.6218653 | 0.262 | 0.214 | 1.15E-15 | DF 1 | si:ch211-57n23.1 |
| gstp11 | 6.1E-20 | 0.2321342 | 0.677 | 0.641 | 1.97E-15 | DF 1 | gstp1 |
| pigr | 7.2E-20 | 0.6982359 | 0.23 | 0.185 | 2.36E-15 | DF 1 | pigr |
| tsku | 8.8E-20 | 0.4647678 | 0.357 | 0.316 | 2.86E-15 | DF 1 | tsku |
| slc40a1 | 9.3E-20 | 0.6025576 | 0.246 | 0.198 | 3.03E-15 | DF 1 | slc40a1 |
| pdgfrb | 1.5E-19 | 0.3467477 | 0.442 | 0.409 | 4.72E-15 | DF 1 | pdgfrb |
| ephb3a | 2.1E-19 | 0.7687117 | 0.144 | 0.103 | 6.73E-15 | DF 1 | ephb3a |
| cbsb | 4.3E-19 | 0.6911953 | 0.179 | 0.135 | 1.41E-14 | DF 1 | cbsb |
| zbtb4 | 9.1E-19 | 0.5181087 | 0.358 | 0.322 | 2.95E-14 | DF 1 | zbtb4 |
| ash1l | 1.2E-18 | 0.4883138 | 0.426 | 0.406 | 3.78E-14 | DF 1 | ash1l |
| twsg1a | 2.1E-18 | 0.6068392 | 0.294 | 0.255 | 6.91E-14 | DF 1 | twsg1a |
| creb3l2 | 3.8E-18 | 0.4287963 | 0.446 | 0.425 | 1.23E-13 | DF 1 | creb3l2 |

| | | | | | | | |
|---|---|---|---|---|---|---|---|
| tnfsf10l | 4.2E-18 | 0.5788971 | 0.263 | 0.218 | 1.36E-13 | DF 1 | tnfsf10l |
| smarca2 | 7.2E-18 | 0.6880973 | 0.261 | 0.223 | 2.34E-13 | DF 1 | smarca2 |
| rbpms2a | 8.3E-18 | 0.4484495 | 0.174 | 0.13 | 2.69E-13 | DF 1 | rbpms2a |
| ppp1r3ca | 1.5E-17 | 0.531121 | 0.25 | 0.205 | 4.86E-13 | DF 1 | ppp1r3ca |
| enc1 | 1.6E-17 | 0.8697697 | 0.161 | 0.121 | 5.19E-13 | DF 1 | enc1 |
| col6a3 | 1.8E-17 | 0.5967236 | 0.275 | 0.235 | 5.81E-13 | DF 1 | col6a3 |
| lix1l | 2.5E-17 | 0.6022333 | 0.214 | 0.171 | 8.12E-13 | DF 1 | lix1l |
| sema3aa | 2.9E-17 | 0.6856784 | 0.145 | 0.105 | 9.34E-13 | DF 1 | sema3aa |
| bsg | 4.2E-17 | 0.3258083 | 0.567 | 0.576 | 1.36E-12 | DF 1 | bsg |
| lgals3b | 4.2E-17 | 0.5047466 | 0.424 | 0.407 | 1.37E-12 | DF 1 | lgals3b |
| map3k14a | 5.3E-17 | 0.923388 | 0.107 | 0.073 | 1.73E-12 | DF 1 | map3k14a |
| ptbp3 | 5.3E-17 | 0.8112854 | 0.17 | 0.131 | 1.73E-12 | DF 1 | ptbp3 |
| lamb1b | 7.2E-17 | 0.3825444 | 0.239 | 0.192 | 2.34E-12 | DF 1 | lamb1b |
| syne3 | 8.5E-17 | 0.8824867 | 0.145 | 0.108 | 2.78E-12 | DF 1 | syne3 |
| si:ch211-196l7.4 | 9.3E-17 | 0.391385 | 0.489 | 0.475 | 3.02E-12 | DF 1 | si:ch211-196l7.4 |
| chka | 1.4E-16 | 0.9052236 | 0.122 | 0.087 | 4.56E-12 | DF 1 | chka |
| sesn1 | 1.7E-16 | 0.5576566 | 0.256 | 0.214 | 5.43E-12 | DF 1 | sesn1 |
| rac3b | 1.8E-16 | 0.8643586 | 0.114 | 0.08 | 5.89E-12 | DF 1 | rac3b |
| si:ch211-212k18.8 | 3.9E-16 | 0.7087819 | 0.177 | 0.138 | 1.28E-11 | DF 1 | si:ch211-212k18.8 |
| dedd1 | 4.9E-16 | 0.3325153 | 0.558 | 0.552 | 1.59E-11 | DF 1 | dedd1 |
| akap13 | 7.2E-16 | 0.5900128 | 0.344 | 0.319 | 2.33E-11 | DF 1 | akap13 |
| si:ch211-194b1.1 | 7.5E-16 | 0.6345315 | 0.249 | 0.213 | 2.43E-11 | DF 1 | si:ch211-194b1.1 |
| rock2b.1 | 1E-15 | 0.7302013 | 0.206 | 0.169 | 3.39E-11 | DF 1 | rock2b.1 |
| cebpa | 1.7E-15 | 0.4570689 | 0.216 | 0.172 | 5.63E-11 | DF 1 | cebpa |
| appa | 1.7E-15 | 0.5490861 | 0.312 | 0.28 | 5.64E-11 | DF 1 | appa |
| h1-10 | 1.7E-15 | 0.7518421 | 0.243 | 0.201 | 5.69E-11 | DF 1 | h1-10 |
| cx35.4 | 1.8E-15 | 0.9569054 | 0.105 | 0.074 | 5.91E-11 | DF 1 | cx35.4 |
| plscr3b | 2.6E-15 | 0.6027005 | 0.273 | 0.238 | 8.33E-11 | DF 1 | plscr3b |
| tpm21 | 2.9E-15 | 0.7874776 | 0.149 | 0.113 | 9.37E-11 | DF 1 | tpm2 |
| ece2b | 3.3E-15 | 0.8281011 | 0.132 | 0.098 | 1.08E-10 | DF 1 | ece2b |
| itm2bb | 4.1E-15 | 0.7735544 | 0.121 | 0.087 | 1.33E-10 | DF 1 | itm2bb |
| caska | 4.3E-15 | 0.9584082 | 0.116 | 0.084 | 1.39E-10 | DF 1 | caska |
| tmtc1 | 4.9E-15 | 0.772618 | 0.15 | 0.114 | 1.6E-10 | DF 1 | tmtc1 |
| lmo4b | 5.1E-15 | 0.3707868 | 0.353 | 0.316 | 1.65E-10 | DF 1 | lmo4b |
| meis1b | 5.3E-15 | 0.4907524 | 0.352 | 0.322 | 1.73E-10 | DF 1 | meis1b |
| abcc5 | 5.8E-15 | 0.7472912 | 0.177 | 0.141 | 1.9E-10 | DF 1 | abcc5 |
| smc3 | 7.1E-15 | 0.6961652 | 0.18 | 0.143 | 2.3E-10 | DF 1 | smc3 |
| setbp11 | 7.6E-15 | 0.6258295 | 0.279 | 0.246 | 2.46E-10 | DF 1 | setbp1 |
| atl3 | 7.8E-15 | 0.5516145 | 0.344 | 0.321 | 2.55E-10 | DF 1 | atl3 |
| acin1a | 8.2E-15 | 0.3221458 | 0.588 | 0.594 | 2.66E-10 | DF 1 | acin1a |
| nuak1b1 | 1.1E-14 | 0.6181494 | 0.309 | 0.28 | 3.73E-10 | DF 1 | nuak1b |
| znf609b | 1.6E-14 | 0.5541361 | 0.302 | 0.271 | 5.34E-10 | DF 1 | znf609b |
| DDX17 | 1.7E-14 | 0.478558 | 0.329 | 0.298 | 5.61E-10 | DF 1 | DDX17 |
| CU984600.11 | 1.9E-14 | 0.6565868 | 0.203 | 0.167 | 6.28E-10 | DF 1 | CU984600.1 |
| fbxl7 | 2.7E-14 | 0.9451973 | 0.104 | 0.074 | 8.62E-10 | DF 1 | fbxl7 |
| rbm25a | 2.8E-14 | 0.8742773 | 0.109 | 0.078 | 8.97E-10 | DF 1 | rbm25a |
| si:ch211-212k18.9 | 3.7E-14 | 0.5983595 | 0.211 | 0.174 | 1.2E-09 | DF 1 | si:ch211-212k18.9 |
| fstl1a | 3.9E-14 | 0.6194422 | 0.139 | 0.104 | 1.26E-09 | DF 1 | fstl1a |

| | | | | | | | |
|---|---|---|---|---|---|---|---|
| jag1b | 4.2E-14 | 0.595085 | 0.249 | 0.213 | 1.36E-09 | DF 1 | jag1b |
| si:ch73-386h18.1 | 4.4E-14 | 0.7096595 | 0.18 | 0.145 | 1.45E-09 | DF 1 | si:ch73-386h18.1 |
| srrm1 | 4.4E-14 | 0.4247323 | 0.423 | 0.407 | 1.45E-09 | DF 1 | srrm1 |
| zfhx3 | 6.4E-14 | 0.2850836 | 0.503 | 0.486 | 2.08E-09 | DF 1 | zfhx3 |
| raraa | 7.4E-14 | 0.4734791 | 0.287 | 0.251 | 2.42E-09 | DF 1 | raraa |
| u2surp | 8E-14 | 0.5075237 | 0.338 | 0.312 | 2.61E-09 | DF 1 | u2surp |
| aldh6a1 | 1.1E-13 | 0.5936436 | 0.298 | 0.272 | 3.62E-09 | DF 1 | aldh6a1 |
| chd3 | 1.5E-13 | 0.6568157 | 0.206 | 0.172 | 4.73E-09 | DF 1 | chd3 |
| pcdh2g28 | 1.6E-13 | 0.5391357 | 0.291 | 0.26 | 5.11E-09 | DF 1 | pcdh2g28 |
| sorbs1 | 1.7E-13 | 0.7385438 | 0.165 | 0.132 | 5.55E-09 | DF 1 | sorbs1 |
| diaph2 | 2E-13 | 0.7591333 | 0.146 | 0.114 | 6.61E-09 | DF 1 | diaph2 |
| cmtm3 | 3.4E-13 | 0.5974514 | 0.225 | 0.192 | 1.1E-08 | DF 1 | cmtm3 |
| mrtfab | 3.6E-13 | 0.8196503 | 0.114 | 0.084 | 1.18E-08 | DF 1 | mrtfab |
| antxr2a | 5.6E-13 | 0.47166 | 0.356 | 0.334 | 1.83E-08 | DF 1 | antxr2a |
| kera | 5.9E-13 | 0.471669 | 0.208 | 0.169 | 1.93E-08 | DF 1 | kera |
| slka | 6.7E-13 | 0.5385565 | 0.325 | 0.302 | 2.19E-08 | DF 1 | slka |
| calm1a | 8.5E-13 | 0.2221494 | 0.624 | 0.619 | 2.77E-08 | DF 1 | calm1a |
| flot2a | 9E-13 | 0.7338434 | 0.146 | 0.115 | 2.94E-08 | DF 1 | flot2a |
| csnk1da | 1E-12 | 0.568775 | 0.243 | 0.21 | 3.4E-08 | DF 1 | csnk1da |
| thrb | 1.1E-12 | 0.7073271 | 0.196 | 0.163 | 3.46E-08 | DF 1 | thrb |
| fbxo25 | 1.2E-12 | 0.8918431 | 0.105 | 0.077 | 3.89E-08 | DF 1 | fbxo25 |
| crim1 | 1.4E-12 | 0.6895628 | 0.174 | 0.141 | 4.61E-08 | DF 1 | crim1 |
| tmem176 | 1.5E-12 | 0.336438 | 0.42 | 0.392 | 4.92E-08 | DF 1 | tmem176 |
| wls | 1.6E-12 | 0.4055468 | 0.41 | 0.394 | 5.29E-08 | DF 1 | wls |
| suco | 2.2E-12 | 0.6606892 | 0.203 | 0.17 | 7.15E-08 | DF 1 | suco |
| ghrb | 2.5E-12 | 0.7644721 | 0.137 | 0.106 | 8.12E-08 | DF 1 | ghrb |
| kifap3a | 2.7E-12 | 0.8367941 | 0.119 | 0.09 | 8.92E-08 | DF 1 | kifap3a |
| hlfa | 2.8E-12 | 0.6825479 | 0.161 | 0.129 | 9E-08 | DF 1 | hlfa |
| trps1 | 3.3E-12 | 0.5974781 | 0.204 | 0.171 | 1.06E-07 | DF 1 | trps1 |
| utrn | 3.3E-12 | 0.6605784 | 0.217 | 0.186 | 1.06E-07 | DF 1 | utrn |
| nedd4a | 3.3E-12 | 0.795227 | 0.118 | 0.089 | 1.07E-07 | DF 1 | nedd4a |
| mid1ip1b | 4.7E-12 | 0.9672983 | 0.147 | 0.117 | 1.53E-07 | DF 1 | mid1ip1b |
| klf7b | 4.8E-12 | 0.4959283 | 0.258 | 0.225 | 1.58E-07 | DF 1 | klf7b |
| si:ch211-166a6.5 | 5E-12 | 0.6742116 | 0.187 | 0.155 | 1.62E-07 | DF 1 | si:ch211-166a6.5 |
| agmo | 5E-12 | 0.8848621 | 0.108 | 0.08 | 1.64E-07 | DF 1 | agmo |
| ubap2a | 6.2E-12 | 0.5893988 | 0.223 | 0.192 | 2.03E-07 | DF 1 | ubap2a |
| ddr1 | 6.8E-12 | 0.6765786 | 0.113 | 0.084 | 2.22E-07 | DF 1 | ddr1 |
| laptm4a | 7.9E-12 | 0.4834789 | 0.326 | 0.305 | 2.59E-07 | DF 1 | laptm4a |
| ntd5 | 9.9E-12 | 0.4730657 | 0.174 | 0.139 | 3.22E-07 | DF 1 | ntd5 |
| arhgap10 | 1.1E-11 | 0.7251036 | 0.175 | 0.146 | 3.61E-07 | DF 1 | arhgap10 |
| tgm1l1 | 1.4E-11 | 0.5913786 | 0.157 | 0.125 | 4.5E-07 | DF 1 | tgm1l1 |
| cpxm1b | 1.6E-11 | 0.7558716 | 0.16 | 0.13 | 5.13E-07 | DF 1 | cpxm1b |
| zgc:174904 | 2.6E-11 | 0.2377707 | 0.156 | 0.122 | 8.59E-07 | DF 1 | zgc:174904 |
| zgc:1135311 | 3E-11 | 0.3876284 | 0.306 | 0.276 | 9.91E-07 | DF 1 | zgc:113531 |
| cebpg | 3.2E-11 | 0.6850512 | 0.165 | 0.136 | 1.05E-06 | DF 1 | cebpg |
| adgrl2a | 4E-11 | 0.4609711 | 0.221 | 0.188 | 1.29E-06 | DF 1 | adgrl2a |
| dip2ca | 4.4E-11 | 0.7807453 | 0.112 | 0.084 | 1.44E-06 | DF 1 | dip2ca |
| arhgap21a | 4.6E-11 | 0.6500808 | 0.133 | 0.104 | 1.5E-06 | DF 1 | arhgap21a |

| | | | | | | | |
|---|---|---|---|---|---|---|---|
| slc44a2 | 5.8E-11 | 0.7322671 | 0.141 | 0.113 | 1.9E-06 | DF 1 | slc44a2 |
| znf609a | 6.5E-11 | 0.674021 | 0.201 | 0.173 | 2.12E-06 | DF 1 | znf609a |
| mob1ba | 7.1E-11 | 0.792274 | 0.126 | 0.099 | 2.3E-06 | DF 1 | mob1ba |
| piezo1 | 7.9E-11 | 0.7626365 | 0.15 | 0.122 | 2.57E-06 | DF 1 | piezo1 |
| gstk1 | 8.7E-11 | 0.668122 | 0.187 | 0.159 | 2.84E-06 | DF 1 | gstk1 |
| tpst1 | 8.8E-11 | 0.565976 | 0.204 | 0.174 | 2.86E-06 | DF 1 | tpst1 |
| stat1a | 1.3E-10 | 0.4612613 | 0.354 | 0.339 | 4.2E-06 | DF 1 | stat1a |
| kirrel1a | 1.8E-10 | 0.3232306 | 0.351 | 0.324 | 5.71E-06 | DF 1 | kirrel1a |
| HACD4 | 2.2E-10 | 0.6414456 | 0.161 | 0.132 | 7.19E-06 | DF 1 | HACD4 |
| krt94 | 2.7E-10 | 0.4291133 | 0.328 | 0.294 | 8.83E-06 | DF 1 | krt94 |
| pnn | 3.8E-10 | 0.5237618 | 0.259 | 0.234 | 1.25E-05 | DF 1 | pnn |
| sgtb | 5E-10 | 0.6261105 | 0.151 | 0.122 | 1.62E-05 | DF 1 | sgtb |
| ognb | 6.3E-10 | 0.2619672 | 0.162 | 0.13 | 2.05E-05 | DF 1 | ognb |
| nkx3-2 | 6.5E-10 | 0.5059835 | 0.223 | 0.194 | 2.11E-05 | DF 1 | nkx3-2 |
| tlcd4b | 7.1E-10 | 0.7690848 | 0.106 | 0.081 | 2.3E-05 | DF 1 | tlcd4b |
| rprd2b | 7.2E-10 | 0.6471404 | 0.148 | 0.12 | 2.34E-05 | DF 1 | rprd2b |
| ada | 1.2E-09 | 0.4590171 | 0.248 | 0.219 | 3.82E-05 | DF 1 | ada |
| calm3a | 1.3E-09 | 0.2306871 | 0.501 | 0.503 | 4.2E-05 | DF 1 | calm3a |
| lamc1 | 1.5E-09 | 0.5729559 | 0.154 | 0.127 | 4.94E-05 | DF 1 | lamc1 |
| zmp:0000000711 | 2.3E-09 | 0.7317989 | 0.105 | 0.081 | 7.57E-05 | DF 1 | zmp:0000000711 |
| fblim1 | 2.4E-09 | 0.695269 | 0.135 | 0.109 | 7.81E-05 | DF 1 | fblim1 |
| tmem198b | 2.5E-09 | 0.597379 | 0.214 | 0.188 | 8.1E-05 | DF 1 | tmem198b |
| nktr | 2.6E-09 | 0.6149117 | 0.183 | 0.157 | 8.6E-05 | DF 1 | nktr |
| foxj3 | 3.3E-09 | 0.6391222 | 0.164 | 0.138 | 0.000107 | DF 1 | foxj3 |
| sh3gl1b | 3.4E-09 | 0.5390783 | 0.263 | 0.243 | 0.00011 | DF 1 | sh3gl1b |
| il11ra1 | 4.2E-09 | 0.48782 | 0.263 | 0.238 | 0.000137 | DF 1 | il11ra |
| mustn1a | 4.6E-09 | 0.5857332 | 0.105 | 0.08 | 0.000149 | DF 1 | mustn1a |
| sod3b | 5.5E-09 | 0.5062367 | 0.149 | 0.121 | 0.000178 | DF 1 | sod3b |
| mhc1uba | 5.5E-09 | 0.4783828 | 0.239 | 0.213 | 0.00018 | DF 1 | mhc1uba |
| phldb1b | 7.9E-09 | 0.8880284 | 0.109 | 0.087 | 0.000257 | DF 1 | phldb1b |
| creg2 | 9.1E-09 | 0.4041233 | 0.338 | 0.325 | 0.000296 | DF 1 | creg2 |
| slc44a1b | 9.5E-09 | 0.6189765 | 0.143 | 0.118 | 0.000308 | DF 1 | slc44a1b |
| fabp3 | 1.1E-08 | 0.2308344 | 0.528 | 0.532 | 0.000358 | DF 1 | fabp3 |
| lrp1ab | 1.3E-08 | 0.3752054 | 0.356 | 0.34 | 0.000438 | DF 1 | lrp1ab |
| rnd3a | 1.4E-08 | 0.3346603 | 0.309 | 0.284 | 0.00045 | DF 1 | rnd3a |
| dlx3b | 1.5E-08 | 0.3153277 | 0.378 | 0.359 | 0.000474 | DF 1 | dlx3b |
| smarca1 | 1.5E-08 | 0.7670946 | 0.102 | 0.08 | 0.000496 | DF 1 | smarca1 |
| si:ch73-138n13.1 | 1.6E-08 | 0.3422628 | 0.389 | 0.377 | 0.000519 | DF 1 | si:ch73-138n13.1 |
| tjp1a | 1.8E-08 | 0.6150021 | 0.177 | 0.152 | 0.000596 | DF 1 | tjp1a |
| zgc:152791 | 2.3E-08 | 0.3510859 | 0.13 | 0.104 | 0.000749 | DF 1 | zgc:152791 |
| sypl2a | 2.9E-08 | 0.5874081 | 0.187 | 0.164 | 0.000936 | DF 1 | sypl2a |
| entpd1 | 3.2E-08 | 0.6355406 | 0.103 | 0.08 | 0.001035 | DF 1 | entpd1 |
| axin1 | 6.3E-08 | 0.68036 | 0.134 | 0.112 | 0.002042 | DF 1 | axin1 |
| pbxip1a | 6.4E-08 | 0.6067362 | 0.207 | 0.186 | 0.002087 | DF 1 | pbxip1a |
| taok1b | 6.7E-08 | 0.6102456 | 0.169 | 0.146 | 0.002167 | DF 1 | taok1b |
| prickle1b | 6.9E-08 | 0.5563283 | 0.2 | 0.177 | 0.002256 | DF 1 | prickle1b |
| rock2a | 7.4E-08 | 0.5018645 | 0.218 | 0.195 | 0.002397 | DF 1 | rock2a |
| slc25a36a | 9.2E-08 | 0.6842816 | 0.127 | 0.105 | 0.002998 | DF 1 | slc25a36a |

| | | | | | | | |
|---|---|---|---|---|---|---|---|
| kif26ba | 9.9E-08 | 0.5070694 | 0.13 | 0.106 | 0.003215 | DF 1 | kif26ba |
| antxr1c | 1E-07 | 0.2183209 | 0.535 | 0.538 | 0.003304 | DF 1 | antxr1c |
| dlx4a1 | 1.1E-07 | 0.4559267 | 0.215 | 0.192 | 0.003739 | DF 1 | dlx4a |
| RAPH1 | 1.3E-07 | 0.673867 | 0.11 | 0.089 | 0.004282 | DF 1 | RAPH1 |
| lpp1 | 1.6E-07 | 0.4084714 | 0.271 | 0.252 | 0.005135 | DF 1 | lpp |
| slc7a8a | 1.8E-07 | 0.7510433 | 0.104 | 0.084 | 0.005871 | DF 1 | slc7a8a |
| plekhj1 | 1.9E-07 | 0.6534892 | 0.142 | 0.12 | 0.006194 | DF 1 | plekhj1 |
| msx1b | 1.9E-07 | 0.3415432 | 0.313 | 0.293 | 0.00623 | DF 1 | msx1b |
| arglu1a | 2.1E-07 | 0.5651205 | 0.193 | 0.171 | 0.006919 | DF 1 | arglu1a |
| ndfip1l | 2.3E-07 | 0.4225414 | 0.342 | 0.338 | 0.007628 | DF 1 | ndfip1l |
| sfswap | 2.9E-07 | 0.6764671 | 0.129 | 0.108 | 0.009283 | DF 1 | sfswap |
| napsa | 3.6E-07 | 0.5022917 | 0.191 | 0.168 | 0.011629 | DF 1 | napsa |
| sc:d217 | 5.9E-07 | 0.6195797 | 0.155 | 0.134 | 0.019348 | DF 1 | sc:d217 |
| camk2g2 | 6.3E-07 | 0.646882 | 0.146 | 0.125 | 0.020495 | DF 1 | camk2g2 |
| cmbl | 6.6E-07 | 0.5461074 | 0.194 | 0.174 | 0.021347 | DF 1 | cmbl |
| sik1 | 7.9E-07 | 0.2330406 | 0.278 | 0.252 | 0.025839 | DF 1 | sik1 |
| oga | 8E-07 | 0.6939162 | 0.116 | 0.096 | 0.026172 | DF 1 | oga |
| adh8b | 8.6E-07 | 0.4780849 | 0.25 | 0.234 | 0.027944 | DF 1 | adh8b |
| si:dkey-188i13.10 | 9.4E-07 | 0.4301985 | 0.264 | 0.247 | 0.030631 | DF 1 | si:dkey-188i13.10 |
| boc | 9.7E-07 | 0.3707548 | 0.377 | 0.374 | 0.031684 | DF 1 | boc |
| pnisr | 1.1E-06 | 0.5401622 | 0.204 | 0.185 | 0.035871 | DF 1 | pnisr |
| bptf | 1.4E-06 | 0.5503312 | 0.198 | 0.179 | 0.046839 | DF 1 | bptf |
| si:cabz01007802.1 | 1.6E-06 | 0.6595784 | 0.108 | 0.089 | 0.053631 | DF 1 | si:cabz01007802.1 |
| supt6h | 1.8E-06 | 0.5551232 | 0.213 | 0.196 | 0.058279 | DF 1 | supt6h |
| ptgr1 | 1.8E-06 | 0.7056395 | 0.157 | 0.139 | 0.059539 | DF 1 | ptgr1 |
| CR847566.1 | 1.9E-06 | 0.5985607 | 0.124 | 0.103 | 0.061225 | DF 1 | CR847566.1 |
| iscub | 1.9E-06 | 0.3978364 | 0.212 | 0.192 | 0.061283 | DF 1 | iscub |
| kmt2cb | 2E-06 | 0.4379372 | 0.287 | 0.274 | 0.064418 | DF 1 | kmt2cb |
| add1 | 2.3E-06 | 0.5569725 | 0.156 | 0.137 | 0.073604 | DF 1 | add1 |
| ehbp1 | 2.4E-06 | 0.7162167 | 0.128 | 0.109 | 0.077137 | DF 1 | ehbp1 |
| sorbs3 | 2.5E-06 | 0.4924679 | 0.208 | 0.189 | 0.081609 | DF 1 | sorbs3 |
| ywhag1 | 2.5E-06 | 0.6310758 | 0.129 | 0.109 | 0.082431 | DF 1 | ywhag1 |
| aplp2 | 2.8E-06 | 0.4272735 | 0.328 | 0.324 | 0.091651 | DF 1 | aplp2 |
| efnb3b | 3.2E-06 | 0.6492038 | 0.156 | 0.138 | 0.102747 | DF 1 | efnb3b |
| ctnna1 | 3.6E-06 | 0.5581857 | 0.194 | 0.177 | 0.116145 | DF 1 | ctnna1 |
| megf8 | 3.7E-06 | 0.6198835 | 0.146 | 0.127 | 0.11907 | DF 1 | megf8 |
| cica | 3.7E-06 | 0.5157093 | 0.203 | 0.185 | 0.120269 | DF 1 | cica |
| sult6b1 | 4.1E-06 | 0.4560187 | 0.196 | 0.176 | 0.132518 | DF 1 | sult6b1 |
| slc16a10 | 4.3E-06 | 0.619564 | 0.151 | 0.132 | 0.139093 | DF 1 | slc16a10 |
| dnmbp | 4.8E-06 | 0.6626892 | 0.119 | 0.101 | 0.154956 | DF 1 | dnmbp |
| cnp | 6E-06 | 0.3898752 | 0.301 | 0.292 | 0.194267 | DF 1 | cnp |
| slc48a1a | 6E-06 | 0.7657896 | 0.105 | 0.088 | 0.196456 | DF 1 | slc48a1a |
| nectin3b | 7.8E-06 | 0.5534895 | 0.172 | 0.154 | 0.252308 | DF 1 | nectin3b |
| sesn3 | 8.8E-06 | 0.6715887 | 0.126 | 0.108 | 0.287125 | DF 1 | sesn3 |
| ENSDARG000000713 | 9.9E-06 | 0.7384978 | 0.103 | 0.086 | 0.321074 | DF 1 | ENSDARG00000071 |
| cyba | 9.9E-06 | 0.4202827 | 0.181 | 0.162 | 0.321509 | DF 1 | cyba |
| ppp1r12a | 1E-05 | 0.5581342 | 0.144 | 0.126 | 0.324905 | DF 1 | ppp1r12a |
| stxbp3 | 1.2E-05 | 0.5455686 | 0.176 | 0.159 | 0.401514 | DF 1 | stxbp3 |

| | | | | | | | |
|---|---|---|---|---|---|---|---|
| si:ch211-286o17.1 | 1.7E-05 | 0.3875156 | 0.233 | 0.216 | 0.549778 | DF 1 | si:ch211-286o17.1 |
| zbtb20 | 2.2E-05 | 0.30374 | 0.378 | 0.375 | 0.702167 | DF 1 | zbtb20 |
| npepps | 2.2E-05 | 0.6115833 | 0.159 | 0.143 | 0.707817 | DF 1 | npepps |
| zranb2 | 2.3E-05 | 0.6311957 | 0.146 | 0.129 | 0.747965 | DF 1 | zranb2 |
| gnsa | 2.3E-05 | 0.5456827 | 0.176 | 0.161 | 0.755565 | DF 1 | gnsa |
| pbxip1b | 2.4E-05 | 0.430527 | 0.253 | 0.241 | 0.769621 | DF 1 | pbxip1b |
| slc1a3a | 3.7E-05 | 0.4878497 | 0.165 | 0.149 | 1 | DF 1 | slc1a3a |
| lygl1 | 4.1E-05 | 0.4660292 | 0.181 | 0.165 | 1 | DF 1 | lygl1 |
| fat1a | 4.1E-05 | 0.4261574 | 0.281 | 0.273 | 1 | DF 1 | fat1a |
| dhrs9 | 4.2E-05 | 0.5182023 | 0.156 | 0.14 | 1 | DF 1 | dhrs9 |
| cspg4 | 4.3E-05 | 0.6182823 | 0.129 | 0.113 | 1 | DF 1 | cspg4 |
| gsto21 | 4.4E-05 | 0.6758451 | 0.119 | 0.103 | 1 | DF 1 | gsto2 |
| trim3b | 4.4E-05 | 0.5074601 | 0.11 | 0.094 | 1 | DF 1 | trim3b |
| dock9b | 4.8E-05 | 0.3424145 | 0.116 | 0.099 | 1 | DF 1 | dock9b |
| si:rp71-36a1.5 | 5.4E-05 | 0.5434856 | 0.102 | 0.086 | 1 | DF 1 | si:rp71-36a1.5 |
| phactr2 | 5.5E-05 | 0.5158998 | 0.156 | 0.14 | 1 | DF 1 | phactr2 |
| si:dkey-226m8.10 | 6E-05 | 0.5164777 | 0.115 | 0.099 | 1 | DF 1 | si:dkey-226m8.10 |
| sltm | 6.2E-05 | 0.455115 | 0.285 | 0.282 | 1 | DF 1 | sltm |
| MDFI | 6.3E-05 | 0.3686506 | 0.252 | 0.238 | 1 | DF 1 | MDFI |
| wu:fj16a031 | 6.5E-05 | 0.4970225 | 0.212 | 0.191 | 1 | DF 1 | wu:fj16a03 |
| ptk7a | 6.5E-05 | 0.374919 | 0.319 | 0.315 | 1 | DF 1 | ptk7a |
| gas1b1 | 6.8E-05 | 0.4615015 | 0.209 | 0.195 | 1 | DF 1 | gas1b |
| tpm3 | 6.9E-05 | 0.2179437 | 0.501 | 0.53 | 1 | DF 1 | tpm3 |
| cep290 | 7E-05 | 0.5929232 | 0.127 | 0.111 | 1 | DF 1 | cep290 |
| cwc27 | 8E-05 | 0.6054287 | 0.137 | 0.122 | 1 | DF 1 | cwc27 |
| ctdspl2b | 8E-05 | 0.5275035 | 0.129 | 0.112 | 1 | DF 1 | ctdspl2b |
| CR318588.1 | 8E-05 | 0.2744984 | 0.307 | 0.295 | 1 | DF 1 | CR318588.1 |
| clk4a | 8.9E-05 | 0.438413 | 0.24 | 0.23 | 1 | DF 1 | clk4a |
| etnk1 | 9.2E-05 | 0.4798727 | 0.202 | 0.188 | 1 | DF 1 | etnk1 |
| jmjd1cb | 9.4E-05 | 0.3742663 | 0.321 | 0.319 | 1 | DF 1 | jmjd1cb |
| znf318 | 0.0001 | 0.5463728 | 0.17 | 0.156 | 1 | DF 1 | znf318 |
| iqgap1 | 0.00011 | 0.465728 | 0.235 | 0.226 | 1 | DF 1 | iqgap1 |
| kif1b | 0.00012 | 0.6519443 | 0.114 | 0.1 | 1 | DF 1 | kif1b |
| golgb1 | 0.00012 | 0.3456423 | 0.371 | 0.376 | 1 | DF 1 | golgb1 |
| plekho2 | 0.00015 | 0.382849 | 0.197 | 0.182 | 1 | DF 1 | plekho2 |
| ches1 | 0.00015 | 0.4869764 | 0.195 | 0.182 | 1 | DF 1 | ches1 |
| arrdc2 | 0.00016 | 0.4328219 | 0.255 | 0.242 | 1 | DF 1 | arrdc2 |
| ythdc1 | 0.00016 | 0.4945932 | 0.144 | 0.129 | 1 | DF 1 | ythdc1 |
| plcd1a | 0.00018 | 0.5863608 | 0.11 | 0.095 | 1 | DF 1 | plcd1a |
| ttyh3a | 0.00018 | 0.4932828 | 0.111 | 0.096 | 1 | DF 1 | ttyh3a |
| h3f3c | 0.00019 | 0.2673796 | 0.449 | 0.468 | 1 | DF 1 | h3f3c |
| stat2 | 0.00027 | 0.4944612 | 0.137 | 0.121 | 1 | DF 1 | stat2 |
| lama4 | 0.00029 | 0.2256455 | 0.282 | 0.27 | 1 | DF 1 | lama4 |
| asap1a | 0.0003 | 0.6022148 | 0.105 | 0.092 | 1 | DF 1 | asap1a |
| atf5b | 0.0003 | 0.3675539 | 0.26 | 0.253 | 1 | DF 1 | atf5b |
| asap1b | 0.0003 | 0.4313775 | 0.165 | 0.151 | 1 | DF 1 | asap1b |
| porb | 0.00032 | 0.5249884 | 0.122 | 0.107 | 1 | DF 1 | porb |
| ms4a17a.9 | 0.00036 | 0.3677015 | 0.227 | 0.216 | 1 | DF 1 | ms4a17a.9 |

| | | | | | | | |
|---|---|---|---|---|---|---|---|
| gripap1 | 0.00036 | 0.5927645 | 0.132 | 0.119 | 1 | DF 1 | gripap1 |
| top2b | 0.00037 | 0.4614808 | 0.268 | 0.266 | 1 | DF 1 | top2b |
| cicb | 0.00039 | 0.5822537 | 0.14 | 0.127 | 1 | DF 1 | cicb |
| sgce | 0.00045 | 0.4282189 | 0.282 | 0.283 | 1 | DF 1 | sgce |
| ccdc85b | 0.00051 | 0.384125 | 0.201 | 0.189 | 1 | DF 1 | ccdc85b |
| pdlim2 | 0.00055 | 0.4554795 | 0.169 | 0.156 | 1 | DF 1 | pdlim2 |
| rnf40 | 0.00059 | 0.5956157 | 0.127 | 0.115 | 1 | DF 1 | rnf40 |
| rhbdf1a | 0.0007 | 0.6932451 | 0.106 | 0.094 | 1 | DF 1 | rhbdf1a |
| sptan1 | 0.00077 | 0.2975632 | 0.254 | 0.245 | 1 | DF 1 | sptan1 |
| nckap1 | 0.00079 | 0.4129747 | 0.244 | 0.238 | 1 | DF 1 | nckap1 |
| nhsl2 | 0.00083 | 0.4496263 | 0.115 | 0.101 | 1 | DF 1 | nhsl2 |
| gstt1b | 0.00086 | 0.528017 | 0.134 | 0.121 | 1 | DF 1 | gstt1b |
| tspan14 | 0.00089 | 0.5442088 | 0.133 | 0.121 | 1 | DF 1 | tspan14 |
| eml3 | 0.00094 | 0.5033454 | 0.133 | 0.12 | 1 | DF 1 | eml3 |
| trioa | 0.00106 | 0.6249085 | 0.104 | 0.092 | 1 | DF 1 | trioa |
| ap2a1 | 0.0011 | 0.4845178 | 0.193 | 0.184 | 1 | DF 1 | ap2a1 |
| eif4a2 | 0.00118 | 0.4521124 | 0.201 | 0.193 | 1 | DF 1 | eif4a2 |
| washc2c | 0.00119 | 0.5021678 | 0.135 | 0.123 | 1 | DF 1 | washc2c |
| mab21l1 | 0.00126 | 0.4383701 | 0.151 | 0.139 | 1 | DF 1 | mab21l1 |
| selenot1b | 0.00126 | 0.2022506 | 0.096 | 0.117 | 1 | DF 1 | selenot1b |
| kdm1a | 0.00134 | 0.2027274 | 0.112 | 0.135 | 1 | DF 1 | kdm1a |
| si:ch211-222k6.3 | 0.00138 | 0.5490537 | 0.122 | 0.11 | 1 | DF 1 | si:ch211-222k6.3 |
| gnb2 | 0.00145 | 0.5206469 | 0.117 | 0.105 | 1 | DF 1 | gnb2 |
| st13 | 0.0015 | 0.4421903 | 0.218 | 0.213 | 1 | DF 1 | st13 |
| zfyve9a | 0.00155 | 0.509025 | 0.165 | 0.156 | 1 | DF 1 | zfyve9a |
| grk4 | 0.00166 | 0.3908762 | 0.19 | 0.18 | 1 | DF 1 | grk4 |
| c1qtnf121 | 0.00174 | 0.4698242 | 0.112 | 0.1 | 1 | DF 1 | c1qtnf12 |
| dhrs13a.1 | 0.00178 | 0.6305764 | 0.113 | 0.102 | 1 | DF 1 | dhrs13a.1 |
| slc12a4 | 0.00191 | 0.5737404 | 0.135 | 0.125 | 1 | DF 1 | slc12a4 |
| clk2a | 0.00194 | 0.5041476 | 0.114 | 0.102 | 1 | DF 1 | clk2a |
| tmem30ab | 0.00204 | 0.5406297 | 0.151 | 0.141 | 1 | DF 1 | tmem30ab |
| hspg2 | 0.00216 | 0.4793236 | 0.156 | 0.146 | 1 | DF 1 | hspg2 |
| pgm1 | 0.00219 | 0.5773769 | 0.11 | 0.1 | 1 | DF 1 | pgm1 |
| si:ch211-71n6.4 | 0.00228 | 0.5766604 | 0.124 | 0.113 | 1 | DF 1 | si:ch211-71n6.4 |
| ptprsa.1 | 0.00228 | 0.5444652 | 0.135 | 0.124 | 1 | DF 1 | ptprsa.1 |
| tns1b | 0.00236 | 0.2440372 | 0.361 | 0.369 | 1 | DF 1 | tns1b |
| arhgap35a | 0.00236 | 0.389001 | 0.189 | 0.18 | 1 | DF 1 | arhgap35a |
| CU459056.1 | 0.00241 | 0.4392299 | 0.179 | 0.169 | 1 | DF 1 | CU459056.1 |
| CU638870.1 | 0.00244 | 0.3547032 | 0.324 | 0.33 | 1 | DF 1 | CU638870.1 |
| DAB2 | 0.00245 | 0.2788999 | 0.333 | 0.384 | 1 | DF 1 | DAB2 |
| ctdsp1 | 0.00273 | 0.4872748 | 0.124 | 0.112 | 1 | DF 1 | ctdsp1 |
| gas1a | 0.00274 | 0.3388894 | 0.171 | 0.16 | 1 | DF 1 | gas1a |
| ramp2 | 0.00335 | 0.3020293 | 0.154 | 0.141 | 1 | DF 1 | ramp2 |
| serbp1b | 0.0034 | 0.4402372 | 0.176 | 0.167 | 1 | DF 1 | serbp1b |
| ano6 | 0.00343 | 0.4871934 | 0.135 | 0.125 | 1 | DF 1 | ano6 |
| shroom4 | 0.00377 | 0.5101028 | 0.143 | 0.133 | 1 | DF 1 | shroom4 |
| gdi2 | 0.00474 | 0.2390708 | 0.446 | 0.478 | 1 | DF 1 | gdi2 |
| tax1bp1b | 0.00522 | 0.3666588 | 0.279 | 0.283 | 1 | DF 1 | tax1bp1b |

| | | | | | | | |
|---|---|---|---|---|---|---|---|
| si:ch211-132g1.3 | 0.00579 | 0.5696616 | 0.113 | 0.103 | 1 | DF 1 | si:ch211-132g1.3 |
| slain21 | 0.00601 | 0.2232376 | 0.094 | 0.112 | 1 | DF 1 | slain2 |
| CABZ01113192.1 | 0.00603 | 0.4517776 | 0.143 | 0.134 | 1 | DF 1 | CABZ01113192.1 |
| tmem131l | 0.00612 | 0.4745218 | 0.141 | 0.132 | 1 | DF 1 | tmem131l |
| phldb2b | 0.00623 | 0.3553796 | 0.154 | 0.143 | 1 | DF 1 | phldb2b |
| pbx3b | 0.00656 | 0.3153596 | 0.154 | 0.143 | 1 | DF 1 | pbx3b |
| ankrd11 | 0.00713 | 0.2537844 | 0.424 | 0.446 | 1 | DF 1 | ankrd11 |
| tasor2 | 0.00714 | 0.4794722 | 0.156 | 0.148 | 1 | DF 1 | tasor2 |
| ptbp1b | 0.00841 | 0.3995713 | 0.195 | 0.19 | 1 | DF 1 | ptbp1b |
| irs1 | 0.00878 | 0.430819 | 0.142 | 0.133 | 1 | DF 1 | irs1 |
| nap1l4b | 0.0089 | 0.2621346 | 0.1 | 0.118 | 1 | DF 1 | nap1l4b |
| p3h31 | 0.00909 | 0.2183692 | 0.117 | 0.138 | 1 | DF 1 | p3h3 |
| ciarta | 0.00938 | 0.3688093 | 0.281 | 0.285 | 1 | DF 1 | ciarta |
| ufd1l | 0.00956 | 0.2072063 | 0.092 | 0.109 | 1 | DF 1 | ufd1l |
| fermt2 | 0.00983 | 0.5168376 | 0.106 | 0.097 | 1 | DF 1 | fermt2 |
| cbr1l | 0.00995 | 0.4137513 | 0.217 | 0.215 | 1 | DF 1 | cbr1l |
| si:ch211-251b21.11 | 0 | 1.2392301 | 0.78 | 0.423 | 0 | DF 2 | si:ch211-251b21.1 |
| fgfbp2a2 | 0 | 1.0532666 | 0.83 | 0.51 | 0 | DF 2 | fgfbp2a |
| igsf101 | 0 | 1.705984 | 0.545 | 0.255 | 0 | DF 2 | igsf10 |
| ptx3a1 | 0 | 1.3021127 | 0.734 | 0.466 | 0 | DF 2 | ptx3a |
| tnc2 | 3E-244 | 1.0966772 | 0.697 | 0.451 | 8.2E-240 | DF 2 | tnc |
| enpp21 | 3E-226 | 1.352949 | 0.619 | 0.43 | 8.5E-222 | DF 2 | enpp2 |
| lxn1 | 1E-201 | 0.7866195 | 0.829 | 0.691 | 4.5E-197 | DF 2 | lxn |
| krt911 | 6E-182 | 0.6369196 | 0.957 | 0.897 | 1.8E-177 | DF 2 | krt91 |
| timp4.3 | 1E-178 | 0.703544 | 0.637 | 0.424 | 4.6E-174 | DF 2 | timp4.3 |
| cfp1 | 3E-175 | 1.1783481 | 0.519 | 0.325 | 9.3E-171 | DF 2 | cfp |
| NC-002333.41 | 2E-171 | 0.4858006 | 1 | 1 | 5.4E-167 | DF 2 | NC-002333.4 |
| CABZ01092746.11 | 1E-166 | 0.6837918 | 0.836 | 0.699 | 4.7E-162 | DF 2 | CABZ01092746.1 |
| si:ch211-106h4.12 | 7E-164 | 0.6519244 | 0.662 | 0.441 | 2.3E-159 | DF 2 | si:ch211-106h4.12 |
| vmo1a1 | 3E-146 | 0.685527 | 0.823 | 0.69 | 1E-141 | DF 2 | vmo1a |
| krt81 | 2E-134 | 0.3867376 | 0.99 | 0.984 | 8E-130 | DF 2 | krt8 |
| ptgdsb.21 | 9E-134 | 0.5847348 | 0.822 | 0.655 | 3.1E-129 | DF 2 | ptgdsb.2 |
| spock31 | 3E-132 | 0.7263981 | 0.595 | 0.397 | 1E-127 | DF 2 | spock3 |
| prrx1a1 | 2E-127 | 0.8158674 | 0.684 | 0.578 | 6E-123 | DF 2 | prrx1a |
| glulb | 2E-117 | 0.7059526 | 0.687 | 0.56 | 5.2E-113 | DF 2 | glulb |
| glula1 | 1E-115 | 0.6861458 | 0.733 | 0.609 | 4.3E-111 | DF 2 | glula |
| ca61 | 6E-114 | 0.5200608 | 0.522 | 0.333 | 1.8E-109 | DF 2 | ca6 |
| crybg1b1 | 1E-101 | 0.8360379 | 0.588 | 0.471 | 4.86E-97 | DF 2 | crybg1b |
| si:dkey-85k7.71 | 1.4E-95 | 0.9998712 | 0.412 | 0.27 | 4.57E-91 | DF 2 | si:dkey-85k7.7 |
| egfl61 | 3.3E-93 | 1.7697962 | 0.13 | 0.049 | 1.06E-88 | DF 2 | egfl6 |
| si:ch73-237c6.11 | 1.9E-84 | 0.886024 | 0.481 | 0.356 | 6.06E-80 | DF 2 | si:ch73-237c6.1 |
| NC-002333.171 | 2.9E-84 | 0.3881927 | 0.999 | 0.999 | 9.34E-80 | DF 2 | NC-002333.17 |
| fmoda | 1.6E-79 | 0.9296085 | 0.296 | 0.174 | 5.32E-75 | DF 2 | fmoda |
| prrx1b1 | 2E-79 | 0.8417021 | 0.486 | 0.371 | 6.59E-75 | DF 2 | prrx1b |
| bhmt1 | 1.7E-77 | 0.3250266 | 0.816 | 0.699 | 5.38E-73 | DF 2 | bhmt |
| fstb1 | 1.2E-75 | 0.6358556 | 0.491 | 0.351 | 3.86E-71 | DF 2 | fstb |
| emilin1a1 | 6.6E-72 | 1.5314555 | 0.175 | 0.087 | 2.13E-67 | DF 2 | emilin1a |
| ecm1b1 | 1.4E-71 | 0.6944509 | 0.567 | 0.464 | 4.58E-67 | DF 2 | ecm1b |

| | | | | | | | |
|---|---|---|---|---|---|---|---|
| kazald32 | 1.4E-71 | 1.0222923 | 0.367 | 0.252 | 4.62E-67 | DF 2 | kazald3 |
| zgc:1943981 | 2.7E-71 | 0.6994333 | 0.496 | 0.374 | 8.69E-67 | DF 2 | zgc:194398 |
| CR383676.21 | 3E-70 | 0.2309858 | 1 | 1 | 9.89E-66 | DF 2 | CR383676.2 |
| fam169aa1 | 3.6E-70 | 0.6600305 | 0.517 | 0.395 | 1.16E-65 | DF 2 | fam169aa |
| ptgdsb.11 | 2.3E-67 | 0.5741195 | 0.587 | 0.453 | 7.56E-63 | DF 2 | ptgdsb.1 |
| cavin1b1 | 8.8E-67 | 0.5474565 | 0.715 | 0.641 | 2.87E-62 | DF 2 | cavin1b |
| ifitm11 | 1.1E-66 | 0.3013814 | 0.919 | 0.779 | 3.66E-62 | DF 2 | ifitm1 |
| sfrp2 | 4.6E-66 | 0.4681727 | 0.521 | 0.388 | 1.5E-61 | DF 2 | sfrp2 |
| ahnak1 | 9.4E-63 | 0.2510601 | 0.971 | 0.961 | 3.07E-58 | DF 2 | ahnak |
| CABZ01068367.11 | 1.6E-62 | 0.8633795 | 0.478 | 0.387 | 5.29E-58 | DF 2 | CABZ01068367.1 |
| lmo4b1 | 2.1E-61 | 0.9441615 | 0.409 | 0.311 | 6.95E-57 | DF 2 | lmo4b |
| slc7a21 | 1.2E-60 | 1.1582654 | 0.247 | 0.152 | 3.87E-56 | DF 2 | slc7a2 |
| si:ch211-160j14.32 | 1.8E-58 | 0.6868529 | 0.419 | 0.306 | 5.96E-54 | DF 2 | si:ch211-160j14.3 |
| nfil3-61 | 9.4E-58 | 0.8178376 | 0.402 | 0.298 | 3.04E-53 | DF 2 | nfil3-6 |
| hhip | 4.1E-56 | 1.5192963 | 0.158 | 0.083 | 1.33E-51 | DF 2 | hhip |
| hpdb1 | 4.6E-56 | 0.2343576 | 0.652 | 0.497 | 1.49E-51 | DF 2 | hpdb |
| ndrg4 | 3.7E-52 | 1.0715136 | 0.236 | 0.148 | 1.19E-47 | DF 2 | ndrg4 |
| postna1 | 9.6E-52 | 0.3183461 | 0.721 | 0.58 | 3.13E-47 | DF 2 | postna |
| BX663503.31 | 4.7E-50 | 0.5987303 | 0.492 | 0.396 | 1.53E-45 | DF 2 | BX663503.3 |
| vcanb1 | 1.1E-49 | 0.3234027 | 0.583 | 0.463 | 3.6E-45 | DF 2 | vcanb |
| cdh111 | 7E-49 | 0.4759423 | 0.657 | 0.585 | 2.27E-44 | DF 2 | cdh11 |
| kif26ba1 | 1.1E-48 | 1.2699485 | 0.178 | 0.102 | 3.53E-44 | DF 2 | kif26ba |
| notch31 | 2.7E-48 | 0.6615936 | 0.429 | 0.34 | 8.93E-44 | DF 2 | notch3 |
| arrdc21 | 1.1E-46 | 0.9322765 | 0.324 | 0.236 | 3.61E-42 | DF 2 | arrdc2 |
| pcolcea1 | 4.1E-42 | 0.3952073 | 0.636 | 0.563 | 1.34E-37 | DF 2 | pcolcea |
| gyg1b1 | 3E-41 | 0.492429 | 0.362 | 0.259 | 9.76E-37 | DF 2 | gyg1b |
| qsox1 | 5.8E-41 | 1.434061 | 0.122 | 0.065 | 1.88E-36 | DF 2 | qsox1 |
| ttc361 | 7.6E-41 | 0.5000726 | 0.446 | 0.347 | 2.47E-36 | DF 2 | ttc36 |
| tagln2 | 9.2E-41 | 0.376596 | 0.616 | 0.533 | 2.99E-36 | DF 2 | tagln |
| mt-co1 | 1.7E-40 | 0.2319297 | 0.992 | 0.995 | 5.47E-36 | DF 2 | mt-co1 |
| slco2b11 | 6.6E-40 | 0.7940897 | 0.299 | 0.215 | 2.15E-35 | DF 2 | slco2b1 |
| pcm11 | 7.5E-40 | 0.9989091 | 0.224 | 0.147 | 2.42E-35 | DF 2 | pcm1 |
| enpp6 | 7.3E-39 | 1.4807345 | 0.105 | 0.053 | 2.37E-34 | DF 2 | enpp6 |
| zgc:158463 | 9.8E-36 | 0.258383 | 0.996 | 0.997 | 3.18E-31 | DF 2 | zgc:158463 |
| si:ch211-170d8.81 | 1.3E-33 | 0.5843897 | 0.526 | 0.481 | 4.32E-29 | DF 2 | si:ch211-170d8.8 |
| col7a1 | 3.1E-32 | 0.2491661 | 0.226 | 0.15 | 1E-27 | DF 2 | col7a1 |
| apof | 7.1E-31 | 1.2525401 | 0.112 | 0.063 | 2.3E-26 | DF 2 | apof |
| raraa1 | 5.8E-30 | 0.8190582 | 0.312 | 0.249 | 1.89E-25 | DF 2 | raraa |
| capn2b1 | 8.7E-30 | 0.6612086 | 0.379 | 0.318 | 2.82E-25 | DF 2 | capn2b |
| hyal2b1 | 1.6E-28 | 0.9432978 | 0.253 | 0.193 | 5.28E-24 | DF 2 | hyal2b |
| en1b | 2.4E-28 | 1.0344194 | 0.162 | 0.106 | 7.8E-24 | DF 2 | en1b |
| tfap2c | 2.9E-27 | 0.8202216 | 0.253 | 0.192 | 9.51E-23 | DF 2 | tfap2c |
| zgc:565851 | 2.7E-25 | 0.5056008 | 0.389 | 0.324 | 8.87E-21 | DF 2 | zgc:56585 |
| adamts21 | 3.1E-25 | 0.5019543 | 0.426 | 0.37 | 9.94E-21 | DF 2 | adamts2 |
| NPAS3 | 4.2E-25 | 1.1329545 | 0.13 | 0.083 | 1.35E-20 | DF 2 | NPAS3 |
| cebpd1 | 2.3E-24 | 0.3966306 | 0.608 | 0.575 | 7.43E-20 | DF 2 | cebpd |
| cav11 | 5E-24 | 0.3118208 | 0.663 | 0.631 | 1.61E-19 | DF 2 | cav1 |
| camk2n1a | 5.2E-23 | 1.1330958 | 0.125 | 0.08 | 1.7E-18 | DF 2 | camk2n1a |

| | | | | | | | |
|---|---|---|---|---|---|---|---|
| timp4.1 | 7.6E-23 | 0.5243222 | 0.142 | 0.092 | 2.48E-18 | DF 2 | timp4.1 |
| cald1b1 | 3.3E-22 | 0.6053456 | 0.223 | 0.164 | 1.07E-17 | DF 2 | cald1b |
| zgc:921611 | 1.4E-21 | 0.7593747 | 0.246 | 0.19 | 4.55E-17 | DF 2 | zgc:92161 |
| tbx3a | 3.9E-21 | 0.8789493 | 0.17 | 0.121 | 1.27E-16 | DF 2 | tbx3a |
| twist3 | 5.8E-21 | 0.6137927 | 0.305 | 0.25 | 1.88E-16 | DF 2 | twist3 |
| ucp21 | 1.2E-20 | 0.3393975 | 0.465 | 0.412 | 3.89E-16 | DF 2 | ucp2 |
| slc22a21 | 3.7E-20 | 0.6922856 | 0.244 | 0.191 | 1.2E-15 | DF 2 | slc22a2 |
| adgrl2a1 | 5.8E-20 | 0.8435242 | 0.236 | 0.187 | 1.87E-15 | DF 2 | adgrl2a |
| adamts15b.1 | 7.9E-20 | 0.9586226 | 0.134 | 0.09 | 2.58E-15 | DF 2 | adamts15b.1 |
| zgc:162612 | 1.5E-19 | 0.8509791 | 0.16 | 0.113 | 5.03E-15 | DF 2 | zgc:162612 |
| c4 | 1.6E-19 | 0.4235318 | 0.262 | 0.204 | 5.26E-15 | DF 2 | c4 |
| ptk7a1 | 2.6E-19 | 0.6490678 | 0.348 | 0.313 | 8.5E-15 | DF 2 | ptk7a |
| myh9b1 | 2.8E-19 | 0.7751762 | 0.34 | 0.3 | 9.12E-15 | DF 2 | myh9b |
| fgl1 | 4.5E-19 | 0.4597705 | 0.265 | 0.206 | 1.46E-14 | DF 2 | fgl1 |
| zeb2a1 | 2.6E-18 | 0.5412129 | 0.379 | 0.339 | 8.59E-14 | DF 2 | zeb2a |
| si:ch211-286o17.11 | 3.8E-18 | 0.6720851 | 0.262 | 0.213 | 1.22E-13 | DF 2 | si:ch211-286o17.1 |
| btf31 | 9.5E-18 | 0.209246 | 0.855 | 0.878 | 3.08E-13 | DF 2 | btf3 |
| cast1 | 1.3E-17 | 0.3340534 | 0.59 | 0.592 | 4.3E-13 | DF 2 | cast |
| cd82a2 | 2.7E-17 | 0.4644526 | 0.394 | 0.353 | 8.62E-13 | DF 2 | cd82a |
| capn121 | 8.1E-17 | 0.3996544 | 0.278 | 0.222 | 2.63E-12 | DF 2 | capn12 |
| cpe | 2E-16 | 0.6639252 | 0.245 | 0.201 | 6.58E-12 | DF 2 | cpe |
| fhl2a1 | 2.2E-16 | 0.7146753 | 0.177 | 0.132 | 7.19E-12 | DF 2 | fhl2a |
| anxa41 | 6.4E-15 | 0.221028 | 0.687 | 0.707 | 2.07E-10 | DF 2 | anxa4 |
| si:ch73-1a9.31 | 7.1E-15 | 0.2401163 | 0.705 | 0.721 | 2.32E-10 | DF 2 | si:ch73-1a9.3 |
| thrap3a1 | 1.8E-14 | 0.5383402 | 0.404 | 0.383 | 6.01E-10 | DF 2 | thrap3a |
| bco2a | 2.2E-14 | 0.8270721 | 0.186 | 0.147 | 7.06E-10 | DF 2 | bco2a |
| CABZ01068358.1 | 4.4E-14 | 0.7659793 | 0.243 | 0.206 | 1.44E-09 | DF 2 | CABZ01068358.1 |
| zfhx31 | 7.1E-14 | 0.4263972 | 0.498 | 0.487 | 2.3E-09 | DF 2 | zfhx3 |
| zgc:174888 | 8.5E-14 | 0.623787 | 0.253 | 0.215 | 2.76E-09 | DF 2 | zgc:174888 |
| ms4a17a.91 | 1E-13 | 0.6745711 | 0.251 | 0.214 | 3.35E-09 | DF 2 | ms4a17a.9 |
| krt5 | 1.7E-13 | 0.3283848 | 0.5 | 0.49 | 5.37E-09 | DF 2 | krt5 |
| tspan35 | 1.8E-13 | 0.7497877 | 0.159 | 0.121 | 5.81E-09 | DF 2 | tspan35 |
| CR759885.11 | 1.9E-13 | 0.8183619 | 0.134 | 0.099 | 6.16E-09 | DF 2 | CR759885.1 |
| mab21l2 | 3.5E-13 | 0.5242397 | 0.356 | 0.33 | 1.12E-08 | DF 2 | mab21l2 |
| pdgfra | 3.5E-13 | 0.3496776 | 0.48 | 0.46 | 1.15E-08 | DF 2 | pdgfra |
| gstr | 5.7E-13 | 0.3847744 | 0.437 | 0.415 | 1.85E-08 | DF 2 | gstr |
| MDFI1 | 7.7E-13 | 0.640009 | 0.271 | 0.237 | 2.52E-08 | DF 2 | MDFI |
| gbe1a1 | 9.7E-13 | 0.7395711 | 0.146 | 0.11 | 3.15E-08 | DF 2 | gbe1a |
| cadm4 | 1.9E-12 | 0.9781143 | 0.104 | 0.074 | 6.18E-08 | DF 2 | cadm4 |
| ITLN11 | 3E-12 | 0.5757856 | 0.247 | 0.208 | 9.67E-08 | DF 2 | ITLN1 |
| sfrp1a1 | 3.1E-12 | 0.2195681 | 0.475 | 0.436 | 1.01E-07 | DF 2 | sfrp1a |
| pcbd11 | 3.8E-12 | 0.325256 | 0.356 | 0.314 | 1.24E-07 | DF 2 | pcbd1 |
| rn7sk1 | 4E-12 | 0.3813559 | 0.474 | 0.455 | 1.29E-07 | DF 2 | rn7sk |
| gstm.11 | 4.6E-12 | 0.2744265 | 0.598 | 0.598 | 1.5E-07 | DF 2 | gstm.1 |
| rsl24d11 | 4.9E-11 | 0.2034625 | 0.73 | 0.751 | 1.59E-06 | DF 2 | rsl24d1 |
| FQ323119.1 | 9.6E-11 | 0.7524578 | 0.133 | 0.102 | 3.13E-06 | DF 2 | FQ323119.1 |
| pcdh11 | 2.3E-10 | 0.8578789 | 0.101 | 0.074 | 7.44E-06 | DF 2 | pcdh11 |
| si:ch73-386h18.11 | 2.8E-10 | 0.8193943 | 0.176 | 0.146 | 9.26E-06 | DF 2 | si:ch73-386h18.1 |

| | | | | | | | |
|---|---|---|---|---|---|---|---|
| suco1 | 4.6E-10 | 0.7261348 | 0.201 | 0.171 | 1.51E-05 | DF 2 | suco |
| FQ323156.11 | 5.8E-10 | 0.5807329 | 0.227 | 0.193 | 1.89E-05 | DF 2 | FQ323156.1 |
| slc44a1b1 | 2.4E-09 | 0.8698002 | 0.146 | 0.118 | 7.93E-05 | DF 2 | slc44a1b |
| fxyd6l | 8.4E-09 | 0.468735 | 0.277 | 0.253 | 0.000275 | DF 2 | fxyd6l |
| ednraa | 8.8E-09 | 0.4706399 | 0.242 | 0.212 | 0.000285 | DF 2 | ednraa |
| slc1a3a1 | 2.2E-08 | 0.7498843 | 0.175 | 0.149 | 0.000711 | DF 2 | slc1a3a |
| ppl1 | 2.9E-08 | 0.5272564 | 0.182 | 0.153 | 0.000931 | DF 2 | ppl |
| si:ch211-183d21.11 | 3.4E-08 | 0.4217552 | 0.393 | 0.382 | 0.001104 | DF 2 | si:ch211-183d21.1 |
| lrp1ab1 | 4.5E-08 | 0.4744315 | 0.353 | 0.341 | 0.001456 | DF 2 | lrp1ab |
| myh101 | 5.3E-08 | 0.4457115 | 0.348 | 0.335 | 0.001733 | DF 2 | myh10 |
| ccdc80l11 | 6E-08 | 0.732354 | 0.101 | 0.077 | 0.001939 | DF 2 | ccdc80l1 |
| fbln7 | 8.5E-08 | 0.4794869 | 0.125 | 0.099 | 0.00275 | DF 2 | fbln7 |
| si:dkey-226m8.101 | 1.2E-07 | 0.73682 | 0.123 | 0.098 | 0.004053 | DF 2 | si:dkey-226m8.10 |
| id1 | 1.8E-07 | 0.2924505 | 0.628 | 0.644 | 0.005713 | DF 2 | id1 |
| LO018188.11 | 2E-07 | 0.2974412 | 0.48 | 0.479 | 0.00658 | DF 2 | LO018188.1 |
| adamts15a1 | 4.6E-07 | 0.6445053 | 0.167 | 0.142 | 0.014846 | DF 2 | adamts15a |
| scinlb1 | 5.7E-07 | 0.45897 | 0.319 | 0.308 | 0.018527 | DF 2 | scinlb |
| cyba1 | 8.6E-07 | 0.5401144 | 0.186 | 0.162 | 0.027952 | DF 2 | cyba |
| mocs2 | 9.4E-07 | 0.8461862 | 0.107 | 0.087 | 0.030682 | DF 2 | mocs2 |
| slc51a1 | 1.2E-06 | 0.4407699 | 0.184 | 0.157 | 0.038171 | DF 2 | slc51a |
| ephb2b | 1.3E-06 | 0.6112183 | 0.218 | 0.199 | 0.042998 | DF 2 | ephb2b |
| flna1 | 1.8E-06 | 0.3686619 | 0.525 | 0.546 | 0.059137 | DF 2 | flna |
| col21a11 | 2.2E-06 | 0.7059306 | 0.1 | 0.079 | 0.070788 | DF 2 | col21a1 |
| neu3.21 | 5E-06 | 0.4390721 | 0.243 | 0.221 | 0.161813 | DF 2 | neu3.2 |
| lhfpl61 | 5.2E-06 | 0.3777874 | 0.414 | 0.419 | 0.170662 | DF 2 | lhfpl6 |
| si:ch73-366l1.51 | 6.1E-06 | 0.7135593 | 0.101 | 0.082 | 0.198438 | DF 2 | si:ch73-366l1.5 |
| c1qtnf122 | 7.2E-06 | 0.5379455 | 0.121 | 0.1 | 0.233021 | DF 2 | c1qtnf12 |
| abhd15a | 1.1E-05 | 0.5653854 | 0.153 | 0.132 | 0.358312 | DF 2 | abhd15a |
| tnfsf10l1 | 1.5E-05 | 0.4842138 | 0.24 | 0.221 | 0.479898 | DF 2 | tnfsf10l |
| fbp1a1 | 2.6E-05 | 0.4539085 | 0.287 | 0.277 | 0.861655 | DF 2 | fbp1a |
| histh1l | 2.8E-05 | 0.2294274 | 0.173 | 0.215 | 0.922348 | DF 2 | histh1l |
| kank41 | 3.3E-05 | 0.6207632 | 0.121 | 0.102 | 1 | DF 2 | kank4 |
| CR847566.11 | 4.1E-05 | 0.6755881 | 0.122 | 0.104 | 1 | DF 2 | CR847566.1 |
| sdc2 | 4.3E-05 | 0.2520345 | 0.552 | 0.592 | 1 | DF 2 | sdc2 |
| sfxn5b1 | 5E-05 | 0.588759 | 0.155 | 0.137 | 1 | DF 2 | sfxn5b |
| actn12 | 6.5E-05 | 0.5448455 | 0.319 | 0.318 | 1 | DF 2 | actn1 |
| eif4ebp3 | 6.5E-05 | 0.5922727 | 0.189 | 0.173 | 1 | DF 2 | eif4ebp3 |
| smarcd2 | 6.6E-05 | 0.569169 | 0.181 | 0.164 | 1 | DF 2 | smarcd2 |
| dazap2 | 7.5E-05 | 0.2334624 | 0.117 | 0.15 | 1 | DF 2 | dazap2 |
| twist21 | 0.00011 | 0.2474272 | 0.354 | 0.341 | 1 | DF 2 | twist2 |
| irs2b | 0.00013 | 0.5788172 | 0.24 | 0.23 | 1 | DF 2 | irs2b |
| stat21 | 0.00017 | 0.7124179 | 0.138 | 0.122 | 1 | DF 2 | stat2 |
| ms4a17a.7 | 0.00017 | 0.409446 | 0.302 | 0.297 | 1 | DF 2 | ms4a17a.7 |
| si:dkey-164f24.22 | 0.00018 | 0.3043356 | 0.19 | 0.169 | 1 | DF 2 | si:dkey-164f24.2 |
| zgc:1951731 | 0.00024 | 0.2486516 | 0.397 | 0.398 | 1 | DF 2 | zgc:195173 |
| rflnb1 | 0.00025 | 0.7231767 | 0.126 | 0.111 | 1 | DF 2 | rflnb |
| caprin1b1 | 0.00029 | 0.2001812 | 0.246 | 0.302 | 1 | DF 2 | caprin1b |
| gnl3 | 0.0003 | 0.2040457 | 0.115 | 0.144 | 1 | DF 2 | gnl3 |

| | | | | | | | |
|---|---|---|---|---|---|---|---|
| cpn11 | 0.00031 | 0.4458621 | 0.224 | 0.212 | 1 | DF 2 | cpn1 |
| si:dkeyp-69e1.8 | 0.00034 | 0.2064978 | 0.087 | 0.111 | 1 | DF 2 | si:dkeyp-69e1.8 |
| slc1a51 | 0.0004 | 0.4911037 | 0.264 | 0.258 | 1 | DF 2 | slc1a5 |
| si:dkeyp-44a8.41 | 0.00044 | 0.3786424 | 0.182 | 0.165 | 1 | DF 2 | si:dkeyp-44a8.4 |
| ehbp1l1a | 0.00049 | 0.6878907 | 0.108 | 0.094 | 1 | DF 2 | ehbp1l1a |
| tnrc181 | 0.00051 | 0.3672657 | 0.394 | 0.41 | 1 | DF 2 | tnrc18 |
| auts2a | 0.00057 | 0.4784043 | 0.232 | 0.223 | 1 | DF 2 | auts2a |
| si:dkey-276j7.1 | 0.0006 | 0.2268261 | 0.139 | 0.171 | 1 | DF 2 | si:dkey-276j7.1 |
| dpyda.11 | 0.00073 | 0.5982736 | 0.125 | 0.111 | 1 | DF 2 | dpyda.1 |
| usp8 | 0.0008 | 0.2156331 | 0.079 | 0.1 | 1 | DF 2 | usp8 |
| chd6 | 0.00082 | 0.2344325 | 0.149 | 0.182 | 1 | DF 2 | chd6 |
| si:dkey-51e6.1 | 0.00094 | 0.4751442 | 0.263 | 0.259 | 1 | DF 2 | si:dkey-51e6.1 |
| si:cabz01007802.11 | 0.00096 | 0.6433912 | 0.103 | 0.089 | 1 | DF 2 | si:cabz01007802.1 |
| nhp2 | 0.00101 | 0.2082737 | 0.103 | 0.128 | 1 | DF 2 | nhp2 |
| fgfr22 | 0.00103 | 0.2791251 | 0.348 | 0.35 | 1 | DF 2 | fgfr2 |
| angptl5 | 0.00107 | 0.3367595 | 0.188 | 0.172 | 1 | DF 2 | angptl5 |
| snai22 | 0.00114 | 0.6273638 | 0.118 | 0.105 | 1 | DF 2 | snai2 |
| mphosph8 | 0.00115 | 0.2181178 | 0.233 | 0.281 | 1 | DF 2 | mphosph8 |
| enah1 | 0.00119 | 0.6506128 | 0.128 | 0.115 | 1 | DF 2 | enah |
| ppp2r2d | 0.0012 | 0.2178993 | 0.119 | 0.145 | 1 | DF 2 | ppp2r2d |
| zgc:123010 | 0.0012 | 0.2099622 | 0.095 | 0.118 | 1 | DF 2 | zgc:123010 |
| fgfr1b | 0.00123 | 0.2033564 | 0.083 | 0.104 | 1 | DF 2 | fgfr1b |
| klf7b1 | 0.00131 | 0.4432361 | 0.236 | 0.228 | 1 | DF 2 | klf7b |
| slc25a36a1 | 0.00135 | 0.2240639 | 0.087 | 0.109 | 1 | DF 2 | slc25a36a |
| rgmb | 0.00137 | 0.2174954 | 0.099 | 0.122 | 1 | DF 2 | rgmb |
| mgat1b | 0.0014 | 0.2096945 | 0.08 | 0.1 | 1 | DF 2 | mgat1b |
| setd2 | 0.0014 | 0.2175234 | 0.102 | 0.126 | 1 | DF 2 | setd2 |
| CR383676.31 | 0.00147 | 0.3031007 | 0.446 | 0.469 | 1 | DF 2 | CR383676.3 |
| kank1a1 | 0.00153 | 0.2048086 | 0.093 | 0.115 | 1 | DF 2 | kank1a |
| tmem176l.3a | 0.00187 | 0.2601132 | 0.125 | 0.153 | 1 | DF 2 | tmem176l.3a |
| ghrb1 | 0.00203 | 0.6178941 | 0.12 | 0.108 | 1 | DF 2 | ghrb |
| hyal41 | 0.00205 | 0.3715981 | 0.12 | 0.105 | 1 | DF 2 | hyal4 |
| cd248a1 | 0.00218 | 0.4308435 | 0.13 | 0.116 | 1 | DF 2 | cd248a |
| myofl1 | 0.00218 | 0.5562692 | 0.17 | 0.16 | 1 | DF 2 | myofl |
| psip1b | 0.00221 | 0.2021648 | 0.162 | 0.193 | 1 | DF 2 | psip1b |
| slc25a24 | 0.00247 | 0.2044074 | 0.186 | 0.223 | 1 | DF 2 | slc25a24 |
| chd2 | 0.00248 | 0.2131064 | 0.152 | 0.182 | 1 | DF 2 | chd2 |
| cnot3a | 0.00296 | 0.2352176 | 0.098 | 0.119 | 1 | DF 2 | cnot3a |
| pawr1 | 0.00303 | 0.3665474 | 0.199 | 0.187 | 1 | DF 2 | pawr |
| ube2ia | 0.00337 | 0.2826377 | 0.125 | 0.151 | 1 | DF 2 | ube2ia |
| znf609b1 | 0.00361 | 0.4347038 | 0.273 | 0.275 | 1 | DF 2 | znf609b |
| kmt2a | 0.00376 | 0.2653341 | 0.137 | 0.165 | 1 | DF 2 | kmt2a |
| gpatch81 | 0.00394 | 0.4222989 | 0.322 | 0.331 | 1 | DF 2 | gpatch8 |
| tnrc6b | 0.0044 | 0.2246083 | 0.118 | 0.142 | 1 | DF 2 | tnrc6b |
| ep300a | 0.00442 | 0.2077867 | 0.102 | 0.123 | 1 | DF 2 | ep300a |
| si:dkey-250d21.1 | 0.00446 | 0.2183881 | 0.123 | 0.148 | 1 | DF 2 | si:dkey-250d21.1 |
| znf609a1 | 0.00451 | 0.52419 | 0.184 | 0.175 | 1 | DF 2 | znf609a |
| rtf2 | 0.00452 | 0.2826294 | 0.1 | 0.121 | 1 | DF 2 | rtf2 |

| | | | | | | | |
|---|---|---|---|---|---|---|---|
| rpl261 | 0.00461 | 0.213868 | 0.133 | 0.159 | 1 | DF 2 | rpl26 |
| arid1aa | 0.00485 | 0.2456872 | 0.099 | 0.12 | 1 | DF 2 | arid1aa |
| glyr1 | 0.00486 | 0.2102605 | 0.085 | 0.104 | 1 | DF 2 | glyr1 |
| akt1s1 | 0.00507 | 0.2320569 | 0.133 | 0.159 | 1 | DF 2 | akt1s1 |
| sik11 | 0.00515 | 0.3154315 | 0.261 | 0.254 | 1 | DF 2 | sik1 |
| ddx18 | 0.00526 | 0.2264525 | 0.083 | 0.101 | 1 | DF 2 | ddx18 |
| barx1 | 0.00555 | 0.5828371 | 0.139 | 0.128 | 1 | DF 2 | barx1 |
| cldnb | 0.00556 | 0.3580587 | 0.159 | 0.148 | 1 | DF 2 | cldnb |
| mdc1 | 0.00556 | 0.5684109 | 0.116 | 0.105 | 1 | DF 2 | mdc1 |
| fnbp4 | 0.00572 | 0.2348985 | 0.132 | 0.158 | 1 | DF 2 | fnbp4 |
| lrrfip2 | 0.00594 | 0.2008312 | 0.088 | 0.107 | 1 | DF 2 | lrrfip2 |
| ctnnb2 | 0.00622 | 0.2100289 | 0.268 | 0.317 | 1 | DF 2 | ctnnb2 |
| trps11 | 0.00624 | 0.4832379 | 0.182 | 0.174 | 1 | DF 2 | trps1 |
| chtopa | 0.00636 | 0.2081296 | 0.117 | 0.139 | 1 | DF 2 | chtopa |
| rp91 | 0.00646 | 0.2061088 | 0.092 | 0.11 | 1 | DF 2 | rp9 |
| ngdn1 | 0.00658 | 0.2381397 | 0.135 | 0.16 | 1 | DF 2 | ngdn |
| gse11 | 0.00673 | 0.6089537 | 0.135 | 0.125 | 1 | DF 2 | gse1 |
| supt16h | 0.00682 | 0.2120521 | 0.123 | 0.146 | 1 | DF 2 | supt16h |
| acvrl11 | 0.00722 | 0.4862974 | 0.246 | 0.247 | 1 | DF 2 | acvrl1 |
| cd99l2 | 0.00775 | 0.4467224 | 0.213 | 0.208 | 1 | DF 2 | cd99l2 |
| srsf5b | 0.00851 | 0.2170632 | 0.099 | 0.118 | 1 | DF 2 | srsf5b |
| slc16a1b1 | 0.00854 | 0.2308592 | 0.112 | 0.133 | 1 | DF 2 | slc16a1b |
| zyx1 | 0.00917 | 0.3733039 | 0.313 | 0.319 | 1 | DF 2 | zyx |
| ms4a17a.8 | 0.00947 | 0.2445408 | 0.097 | 0.116 | 1 | DF 2 | ms4a17a.8 |
| inka1a | 0.0097 | 0.4787776 | 0.182 | 0.175 | 1 | DF 2 | inka1a |
| lpp2 | 0.00994 | 0.5131624 | 0.254 | 0.254 | 1 | DF 2 | lpp |
| hsp70.2 | 0 | 1.9064138 | 0.832 | 0.465 | 0 | DF 3 | hsp70.2 |
| ptx3a2 | 0 | 0.8265643 | 0.808 | 0.46 | 0 | DF 3 | ptx3a |
| gyg1b2 | 0 | 1.5888501 | 0.587 | 0.24 | 0 | DF 3 | gyg1b |
| dnajb1b | 0 | 1.622722 | 0.761 | 0.439 | 0 | DF 3 | dnajb1b |
| hsp70.1 | 0 | 1.8670125 | 0.629 | 0.317 | 0 | DF 3 | hsp70.1 |
| hsp70.3 | 0 | 1.6728234 | 0.833 | 0.531 | 0 | DF 3 | hsp70.3 |
| jdp2b | 0 | 1.3303556 | 0.844 | 0.548 | 0 | DF 3 | jdp2b |
| her6 | 0 | 1.5162039 | 0.693 | 0.411 | 0 | DF 3 | her6 |
| rsrp1 | 0 | 1.0719605 | 0.787 | 0.522 | 0 | DF 3 | rsrp1 |
| hsp90aa1.2 | 0 | 1.7309175 | 0.843 | 0.579 | 0 | DF 3 | hsp90aa1.2 |
| ubb | 0 | 1.3941151 | 0.916 | 0.661 | 0 | DF 3 | ubb |
| mycb | 0 | 1.1455686 | 0.873 | 0.645 | 0 | DF 3 | mycb |
| mcl1a | 0 | 0.910876 | 0.924 | 0.758 | 0 | DF 3 | mcl1a |
| aldh1a31 | 7E-305 | 1.6318551 | 0.307 | 0.094 | 2.3E-300 | DF 3 | aldh1a3 |
| cebpa1 | 2E-302 | 1.5023137 | 0.407 | 0.156 | 5.1E-298 | DF 3 | cebpa |
| vcanb2 | 2E-302 | 0.9658011 | 0.755 | 0.448 | 7.5E-298 | DF 3 | vcanb |
| hsp70l | 3E-302 | 1.7060047 | 0.558 | 0.281 | 1E-297 | DF 3 | hsp70l |
| ptgdsb.22 | 6E-301 | 0.7907754 | 0.899 | 0.648 | 1.9E-296 | DF 3 | ptgdsb.2 |
| atf3 | 1E-300 | 1.4214059 | 0.605 | 0.324 | 4.3E-296 | DF 3 | atf3 |
| ddit3 | 4E-293 | 1.0403209 | 0.8 | 0.564 | 1.2E-288 | DF 3 | ddit3 |
| cxcl8a1 | 2E-288 | 1.1183552 | 0.528 | 0.244 | 8.1E-284 | DF 3 | cxcl8a |
| tob1b1 | 9E-286 | 0.8926477 | 0.871 | 0.688 | 2.8E-281 | DF 3 | tob1b |

| | | | | | | | |
|---|---|---|---|---|---|---|---|
| si:ch73-335l21.4 | 8E-280 | 1.1955915 | 0.787 | 0.552 | 2.6E-275 | DF 3 | si:ch73-335l21.4 |
| cxcl18b | 1E-277 | 0.7226088 | 0.91 | 0.684 | 4E-273 | DF 3 | cxcl18b |
| fosab | 2E-272 | 0.8594497 | 0.983 | 0.888 | 6.5E-268 | DF 3 | fosab |
| jun | 1E-265 | 0.8122771 | 0.973 | 0.844 | 3.9E-261 | DF 3 | jun |
| ptgdsb.12 | 9E-263 | 0.8586934 | 0.736 | 0.44 | 2.9E-258 | DF 3 | ptgdsb.1 |
| hapln1a1 | 1E-261 | 0.8111283 | 0.754 | 0.455 | 4.5E-257 | DF 3 | hapln1a |
| UBB | 1E-255 | 1.1082747 | 0.765 | 0.545 | 4.7E-251 | DF 3 | UBB |
| thy11 | 3E-254 | 1.3424286 | 0.462 | 0.212 | 1.1E-249 | DF 3 | thy1 |
| ZFP36L1.1 | 4E-253 | 0.7584863 | 0.966 | 0.866 | 1.3E-248 | DF 3 | ZFP36L1.1 |
| ier2b | 3E-250 | 0.9301578 | 0.844 | 0.633 | 1.1E-245 | DF 3 | ier2b |
| icn1 | 2E-247 | 0.5647986 | 0.998 | 0.996 | 7.5E-243 | DF 3 | icn |
| fosb1 | 4E-247 | 1.0191792 | 0.869 | 0.678 | 1.4E-242 | DF 3 | fosb |
| junba | 7E-247 | 0.6891136 | 0.971 | 0.835 | 2.1E-242 | DF 3 | junba |
| junbb | 5E-246 | 0.9636258 | 0.809 | 0.584 | 1.6E-241 | DF 3 | junbb |
| tagln3 | 7E-239 | 0.6366024 | 0.781 | 0.518 | 2.1E-234 | DF 3 | tagln |
| fgfbp2a3 | 8E-239 | 0.4056837 | 0.838 | 0.51 | 2.6E-234 | DF 3 | fgfbp2a |
| ier5 | 2E-237 | 1.0214376 | 0.699 | 0.451 | 7.7E-233 | DF 3 | ier5 |
| ifitm12 | 9E-235 | 0.6054556 | 0.959 | 0.776 | 3E-230 | DF 3 | ifitm1 |
| serpine11 | 2E-231 | 0.7650014 | 0.913 | 0.749 | 5E-227 | DF 3 | serpine1 |
| msx1b1 | 6E-227 | 1.2081994 | 0.518 | 0.275 | 2.1E-222 | DF 3 | msx1b |
| id3 | 1E-223 | 0.9488694 | 0.728 | 0.494 | 3.9E-219 | DF 3 | id3 |
| ppp1r10 | 1E-216 | 1.0116454 | 0.694 | 0.467 | 3.9E-212 | DF 3 | ppp1r10 |
| ccl25b1 | 2E-213 | 0.3383681 | 0.91 | 0.643 | 7E-209 | DF 3 | ccl25b |
| ppdpfb1 | 4E-205 | 0.7151165 | 0.877 | 0.77 | 1.3E-200 | DF 3 | ppdpfb |
| krt912 | 1E-196 | 0.5885335 | 0.96 | 0.897 | 3.5E-192 | DF 3 | krt91 |
| lpl1 | 1E-196 | 1.4328211 | 0.271 | 0.1 | 3.7E-192 | DF 3 | lpl |
| nfkbiab | 5E-195 | 0.9108319 | 0.764 | 0.568 | 1.5E-190 | DF 3 | nfkbiab |
| btg2 | 3E-192 | 0.7981801 | 0.861 | 0.686 | 8.4E-188 | DF 3 | btg2 |
| tmsb4x1 | 2E-190 | 0.3836294 | 0.998 | 0.991 | 8E-186 | DF 3 | tmsb4x |
| si:dkey-85k7.72 | 1E-180 | 0.9612857 | 0.495 | 0.263 | 3.3E-176 | DF 3 | si:dkey-85k7.7 |
| ccn1 | 2E-179 | 0.9488452 | 0.805 | 0.629 | 5.1E-175 | DF 3 | ccn1 |
| eif1 | 3E-179 | 0.6832881 | 0.895 | 0.824 | 1E-174 | DF 3 | eif1 |
| egr1 | 3E-178 | 0.8006144 | 0.808 | 0.613 | 1.1E-173 | DF 3 | egr1 |
| bzw1b | 7E-178 | 1.0626953 | 0.67 | 0.477 | 2.1E-173 | DF 3 | bzw1b |
| brd2a | 3E-175 | 1.0721251 | 0.631 | 0.432 | 9.4E-171 | DF 3 | brd2a |
| has21 | 2E-173 | 1.3950377 | 0.262 | 0.101 | 6.7E-169 | DF 3 | has2 |
| hpdb2 | 8E-173 | 0.5722216 | 0.748 | 0.489 | 2.5E-168 | DF 3 | hpdb |
| trib3 | 3E-169 | 1.4088936 | 0.343 | 0.162 | 1.1E-164 | DF 3 | trib3 |
| cxcl12a1 | 3E-167 | 0.3582544 | 0.902 | 0.664 | 1E-162 | DF 3 | cxcl12a |
| plpp1a1 | 9E-166 | 0.8132782 | 0.589 | 0.358 | 2.9E-161 | DF 3 | plpp1a |
| fosl1a | 1E-164 | 0.9283723 | 0.632 | 0.422 | 3.2E-160 | DF 3 | fosl1a |
| lxn2 | 2E-163 | 0.6005353 | 0.855 | 0.689 | 8.1E-159 | DF 3 | lxn |
| C18H15orf481 | 7E-161 | 0.7852893 | 0.519 | 0.293 | 2.2E-156 | DF 3 | C18H15orf48 |
| tnfaip61 | 2E-160 | 0.671317 | 0.64 | 0.414 | 6.3E-156 | DF 3 | tnfaip6 |
| fstl3 | 6E-157 | 1.0767708 | 0.513 | 0.306 | 1.9E-152 | DF 3 | fstl3 |
| tob1a | 2E-156 | 0.7894596 | 0.764 | 0.597 | 5.5E-152 | DF 3 | tob1a |
| sdc4 | 9E-152 | 0.67761 | 0.834 | 0.693 | 3E-147 | DF 3 | sdc4 |
| prelid3b | 2E-148 | 1.0969685 | 0.528 | 0.344 | 5E-144 | DF 3 | prelid3b |

| | | | | | | | |
|---|---|---|---|---|---|---|---|
| CABZ01092746.12 | 2E-148 | 0.5883454 | 0.855 | 0.697 | 5.6E-144 | DF 3 | CABZ01092746.1 |
| hgd1 | 3E-148 | 0.7165904 | 0.635 | 0.412 | 8.6E-144 | DF 3 | hgd |
| CR383676.22 | 2E-146 | 0.3066442 | 1 | 1 | 6.3E-142 | DF 3 | CR383676.2 |
| btg1 | 6E-143 | 0.5763161 | 0.899 | 0.808 | 1.9E-138 | DF 3 | btg1 |
| syt11a1 | 2E-141 | 0.9506305 | 0.326 | 0.152 | 6.4E-137 | DF 3 | syt11a |
| pnrc2 | 1E-139 | 0.5068097 | 0.936 | 0.877 | 3.2E-135 | DF 3 | pnrc2 |
| hmox1a | 2E-139 | 1.0012869 | 0.331 | 0.161 | 6E-135 | DF 3 | hmox1a |
| bhmt2 | 1E-137 | 0.4685779 | 0.866 | 0.695 | 4.3E-133 | DF 3 | bhmt |
| glulb1 | 6E-137 | 0.6948151 | 0.736 | 0.556 | 2.1E-132 | DF 3 | glulb |
| nfkbiaa | 4E-136 | 0.9262065 | 0.601 | 0.409 | 1.4E-131 | DF 3 | nfkbiaa |
| tagapa | 9E-134 | 1.6543652 | 0.183 | 0.067 | 2.9E-129 | DF 3 | tagapa |
| dusp1 | 1E-130 | 0.9898045 | 0.552 | 0.366 | 3.6E-126 | DF 3 | dusp1 |
| spock32 | 4E-130 | 0.3539311 | 0.653 | 0.392 | 1.2E-125 | DF 3 | spock3 |
| si:ch73-173h19.31 | 2E-129 | 0.7692431 | 0.417 | 0.225 | 6.5E-125 | DF 3 | si:ch73-173h19.3 |
| pcolcea2 | 3E-129 | 0.5511819 | 0.746 | 0.554 | 8.8E-125 | DF 3 | pcolcea |
| sik12 | 3E-129 | 0.9254825 | 0.425 | 0.239 | 1E-124 | DF 3 | sik1 |
| krt41 | 9E-128 | 0.4288903 | 0.897 | 0.747 | 2.8E-123 | DF 3 | krt4 |
| slc7a22 | 1E-123 | 0.9433147 | 0.307 | 0.147 | 3.9E-119 | DF 3 | slc7a2 |
| efna1b | 2E-123 | 1.0902182 | 0.281 | 0.132 | 5.6E-119 | DF 3 | efna1b |
| mknk2b | 5E-122 | 0.6579962 | 0.761 | 0.595 | 1.8E-117 | DF 3 | mknk2b |
| rnd3a1 | 7E-122 | 0.8423628 | 0.455 | 0.272 | 2.2E-117 | DF 3 | rnd3a |
| slc25a25a | 2E-120 | 1.0581622 | 0.37 | 0.207 | 7.9E-116 | DF 3 | slc25a25a |
| midn | 7E-120 | 0.9647212 | 0.587 | 0.422 | 2.3E-115 | DF 3 | midn |
| vim1 | 6E-118 | 0.5936672 | 0.558 | 0.355 | 1.9E-113 | DF 3 | vim |
| apoeb1 | 7E-117 | 0.9455876 | 0.409 | 0.235 | 2.2E-112 | DF 3 | apoeb |
| kcne41 | 2E-116 | 1.4041402 | 0.153 | 0.054 | 5.4E-112 | DF 3 | kcne4 |
| capn122 | 2E-116 | 0.620603 | 0.398 | 0.212 | 7.4E-112 | DF 3 | capn12 |
| dlx3b1 | 2E-116 | 0.8209533 | 0.526 | 0.347 | 7.5E-112 | DF 3 | dlx3b |
| cbsb1 | 8E-116 | 1.113869 | 0.271 | 0.128 | 2.5E-111 | DF 3 | cbsb |
| s100a10b1 | 1E-115 | 0.2697416 | 0.997 | 0.994 | 3.3E-111 | DF 3 | s100a10b |
| csnk1da1 | 5E-115 | 1.0819636 | 0.359 | 0.201 | 1.6E-110 | DF 3 | csnk1da |
| postna2 | 3E-113 | 0.2682501 | 0.834 | 0.571 | 9.1E-109 | DF 3 | postna |
| jund1 | 4E-113 | 0.7010886 | 0.718 | 0.558 | 1.3E-108 | DF 3 | jund |
| metrn | 8E-112 | 1.2128768 | 0.237 | 0.107 | 2.6E-107 | DF 3 | metrn |
| angptl4 | 1E-110 | 0.8112943 | 0.615 | 0.443 | 3.5E-106 | DF 3 | angptl4 |
| dusp5 | 4E-110 | 1.3409883 | 0.335 | 0.187 | 1.2E-105 | DF 3 | dusp5 |
| fthl271 | 4E-110 | 0.4569836 | 0.908 | 0.834 | 1.3E-105 | DF 3 | fthl27 |
| sox11a | 5E-110 | 0.8802068 | 0.503 | 0.33 | 1.5E-105 | DF 3 | sox11a |
| cited4b | 4E-109 | 1.0841774 | 0.28 | 0.139 | 1.2E-104 | DF 3 | cited4b |
| tnc3 | 2E-107 | 0.4654597 | 0.668 | 0.454 | 5.2E-103 | DF 3 | tnc |
| atf4b | 3E-106 | 0.5448602 | 0.764 | 0.623 | 1E-101 | DF 3 | atf4b |
| ntd51 | 3E-105 | 0.9818027 | 0.271 | 0.132 | 9.8E-101 | DF 3 | ntd5 |
| tuft1a1 | 7E-104 | 0.628263 | 0.73 | 0.58 | 2.4E-99 | DF 3 | tuft1a |
| pawr2 | 6E-99 | 0.7996295 | 0.327 | 0.176 | 1.81E-94 | DF 3 | pawr |
| gsta.11 | 4.2E-98 | 0.7116998 | 0.515 | 0.344 | 1.37E-93 | DF 3 | gsta.1 |
| aoc21 | 5.1E-98 | 0.7519574 | 0.393 | 0.229 | 1.65E-93 | DF 3 | aoc2 |
| twist1a | 5.7E-98 | 0.6363831 | 0.601 | 0.425 | 1.85E-93 | DF 3 | twist1a |
| si:ch211-105c13.31 | 5.4E-97 | 0.3323919 | 0.744 | 0.555 | 1.75E-92 | DF 3 | si:ch211-105c13.3 |

| | | | | | | | |
|---|---|---|---|---|---|---|---|
| sned11 | 9.3E-97 | 1.0651826 | 0.149 | 0.056 | 3.02E-92 | DF 3 | sned1 |
| lhfpl62 | 1.1E-96 | 0.6549313 | 0.574 | 0.405 | 3.68E-92 | DF 3 | lhfpl6 |
| gbe1a2 | 1E-95 | 0.9092521 | 0.225 | 0.103 | 3.35E-91 | DF 3 | gbe1a |
| klf6a | 6.2E-95 | 0.5928725 | 0.838 | 0.739 | 2.01E-90 | DF 3 | klf6a |
| rbp41 | 7.5E-93 | 0.3000717 | 0.968 | 0.905 | 2.45E-88 | DF 3 | rbp4 |
| gch11 | 1.6E-91 | 0.9203939 | 0.252 | 0.126 | 5.18E-87 | DF 3 | gch1 |
| si:ch211-262i1.41 | 4.6E-91 | 0.9613245 | 0.175 | 0.073 | 1.5E-86 | DF 3 | si:ch211-262i1.4 |
| FQ323156.12 | 4.9E-91 | 0.8140478 | 0.33 | 0.184 | 1.6E-86 | DF 3 | FQ323156.1 |
| brdt | 5.9E-91 | 0.9720738 | 0.446 | 0.3 | 1.91E-86 | DF 3 | brdt |
| actb1 | 3.7E-90 | 0.33172 | 0.944 | 0.86 | 1.21E-85 | DF 3 | actb1 |
| krt151 | 4.7E-90 | 0.3542452 | 0.835 | 0.69 | 1.52E-85 | DF 3 | krt15 |
| adgre10 | 9.4E-90 | 1.3448707 | 0.164 | 0.069 | 3.05E-85 | DF 3 | adgre10 |
| mfap51 | 1.4E-88 | 0.3796826 | 0.726 | 0.531 | 4.57E-84 | DF 3 | mfap5 |
| mlip1 | 8.9E-88 | 1.1372859 | 0.161 | 0.066 | 2.88E-83 | DF 3 | mlip |
| xbp1 | 6.3E-87 | 0.5155205 | 0.842 | 0.755 | 2.04E-82 | DF 3 | xbp1 |
| sgk1 | 6.5E-87 | 0.722697 | 0.513 | 0.355 | 2.13E-82 | DF 3 | sgk1 |
| si:ch211-195b13.1 | 6.9E-87 | 0.8228207 | 0.379 | 0.232 | 2.24E-82 | DF 3 | si:ch211-195b13.1 |
| si:dkey-164f24.23 | 2.6E-86 | 0.8086161 | 0.295 | 0.16 | 8.39E-82 | DF 3 | si:dkey-164f24.2 |
| ucp22 | 3.1E-86 | 0.4877157 | 0.573 | 0.402 | 1.01E-81 | DF 3 | ucp2 |
| pah1 | 4.5E-86 | 0.3101699 | 0.53 | 0.336 | 1.46E-81 | DF 3 | pah |
| qdpra1 | 8.5E-86 | 0.3520756 | 0.613 | 0.411 | 2.76E-81 | DF 3 | qdpra |
| C25H11orf96 | 2.2E-84 | 0.5190589 | 0.547 | 0.37 | 7.03E-80 | DF 3 | C25H11orf96 |
| ccng2 | 1.1E-83 | 1.0180226 | 0.348 | 0.213 | 3.48E-79 | DF 3 | ccng2 |
| si:ch73-86n18.11 | 2.1E-83 | 0.7908496 | 0.27 | 0.145 | 6.88E-79 | DF 3 | si:ch73-86n18.1 |
| nfil3-5 | 2.4E-82 | 1.2079003 | 0.245 | 0.13 | 7.94E-78 | DF 3 | nfil3-5 |
| ebf1b | 6.9E-82 | 1.3925648 | 0.113 | 0.04 | 2.25E-77 | DF 3 | ebf1b |
| fam169aa2 | 7.8E-82 | 0.4575651 | 0.568 | 0.391 | 2.55E-77 | DF 3 | fam169aa |
| pim21 | 9.9E-82 | 0.8436124 | 0.356 | 0.216 | 3.21E-77 | DF 3 | pim2 |
| lmo4b2 | 2.5E-81 | 0.6242537 | 0.464 | 0.307 | 8.11E-77 | DF 3 | lmo4b |
| zfand5b | 7E-81 | 0.8530242 | 0.403 | 0.262 | 2.26E-76 | DF 3 | zfand5b |
| slc51a2 | 1.5E-80 | 0.6872262 | 0.279 | 0.149 | 4.81E-76 | DF 3 | slc51a |
| ppp1r3ca1 | 1.5E-79 | 0.839189 | 0.335 | 0.198 | 4.85E-75 | DF 3 | ppp1r3ca |
| sat1a.2 | 2.1E-78 | 0.6424742 | 0.588 | 0.447 | 6.92E-74 | DF 3 | sat1a.2 |
| id11 | 1.1E-77 | 0.5130125 | 0.774 | 0.631 | 3.43E-73 | DF 3 | id1 |
| hist2h2l | 1.8E-77 | 0.7086777 | 0.508 | 0.364 | 5.88E-73 | DF 3 | hist2h2l |
| ecm1b2 | 8.8E-77 | 0.4530036 | 0.625 | 0.459 | 2.86E-72 | DF 3 | ecm1b |
| gadd45ba | 1.1E-76 | 0.5196534 | 0.882 | 0.797 | 3.6E-72 | DF 3 | gadd45ba |
| agtr21 | 5.3E-76 | 0.8858264 | 0.217 | 0.108 | 1.71E-71 | DF 3 | agtr2 |
| slc40a11 | 3E-75 | 0.6998454 | 0.326 | 0.192 | 9.6E-71 | DF 3 | slc40a1 |
| gadd45ab | 8E-75 | 1.0604532 | 0.286 | 0.167 | 2.61E-70 | DF 3 | gadd45ab |
| BX663503.32 | 5.1E-74 | 0.4994483 | 0.558 | 0.39 | 1.65E-69 | DF 3 | BX663503.3 |
| cebpd2 | 6.4E-74 | 0.5053075 | 0.714 | 0.566 | 2.09E-69 | DF 3 | cebpd |
| ppap2d1 | 1.6E-73 | 0.6492617 | 0.361 | 0.219 | 5.26E-69 | DF 3 | ppap2d |
| aqp3a1 | 1.8E-73 | 0.2318342 | 0.478 | 0.307 | 5.83E-69 | DF 3 | aqp3a |
| klf11b | 7.9E-73 | 0.8316308 | 0.358 | 0.227 | 2.56E-68 | DF 3 | klf11b |
| tubb4b | 1.3E-72 | 0.3512787 | 0.876 | 0.804 | 4.2E-68 | DF 3 | tubb4b |
| si:dkey-79d12.5 | 1.6E-72 | 1.1786499 | 0.227 | 0.122 | 5.09E-68 | DF 3 | si:dkey-79d12.5 |
| si:dkey-195m11.8 | 6.1E-72 | 1.0918709 | 0.225 | 0.12 | 1.97E-67 | DF 3 | si:dkey-195m11.8 |

| | | | | | | | |
|---|---|---|---|---|---|---|---|
| ttc362 | 3.1E-71 | 0.4048432 | 0.516 | 0.341 | 9.98E-67 | DF 3 | ttc36 |
| ip6k2b | 6.1E-71 | 0.7122428 | 0.47 | 0.333 | 2E-66 | DF 3 | ip6k2b |
| pmaip1 | 6.6E-71 | 1.2749347 | 0.18 | 0.088 | 2.13E-66 | DF 3 | pmaip1 |
| sfrp1b1 | 7.5E-71 | 0.3061818 | 0.525 | 0.349 | 2.44E-66 | DF 3 | sfrp1b |
| cfp2 | 1.6E-70 | 0.4701924 | 0.491 | 0.328 | 5.2E-66 | DF 3 | cfp |
| zgc:1951732 | 3.9E-70 | 0.5180611 | 0.537 | 0.386 | 1.28E-65 | DF 3 | zgc:195173 |
| slc22a22 | 4.5E-70 | 0.6247612 | 0.317 | 0.185 | 1.47E-65 | DF 3 | slc22a2 |
| slco2b12 | 3.7E-68 | 0.5376061 | 0.349 | 0.211 | 1.2E-63 | DF 3 | slco2b1 |
| cdon1 | 4.8E-68 | 0.8018433 | 0.3 | 0.179 | 1.58E-63 | DF 3 | cdon |
| nfil3-62 | 1.6E-67 | 0.5761795 | 0.442 | 0.295 | 5.2E-63 | DF 3 | nfil3-6 |
| zgc:1943982 | 2.1E-66 | 0.3877443 | 0.532 | 0.371 | 6.68E-62 | DF 3 | zgc:194398 |
| plpp31 | 6.9E-66 | 0.3435545 | 0.613 | 0.442 | 2.26E-61 | DF 3 | plpp3 |
| zgc:122979 | 4.3E-65 | 1.1955049 | 0.157 | 0.074 | 1.4E-60 | DF 3 | zgc:122979 |
| hbegfa | 2E-64 | 0.615247 | 0.524 | 0.392 | 6.55E-60 | DF 3 | hbegfa |
| fstb2 | 3.8E-64 | 0.3864623 | 0.511 | 0.35 | 1.22E-59 | DF 3 | fstb |
| prx1 | 1.3E-62 | 0.7251223 | 0.136 | 0.06 | 4.15E-58 | DF 3 | prx |
| ubc | 2E-62 | 0.4890092 | 0.756 | 0.655 | 6.6E-58 | DF 3 | ubc |
| arl5c1 | 7.8E-62 | 1.0648976 | 0.21 | 0.115 | 2.53E-57 | DF 3 | arl5c |
| olfml2ba1 | 1.3E-61 | 0.516951 | 0.222 | 0.118 | 4.32E-57 | DF 3 | olfml2ba |
| selenow11 | 2.6E-61 | 0.299325 | 0.926 | 0.895 | 8.59E-57 | DF 3 | selenow1 |
| mfap21 | 4.8E-61 | 0.4974807 | 0.519 | 0.373 | 1.57E-56 | DF 3 | mfap2 |
| anxa42 | 6.4E-61 | 0.3625206 | 0.786 | 0.698 | 2.07E-56 | DF 3 | anxa4 |
| capn2b2 | 9.9E-61 | 0.5442247 | 0.45 | 0.312 | 3.22E-56 | DF 3 | capn2b |
| chac1 | 2.7E-60 | 0.618348 | 0.472 | 0.345 | 8.68E-56 | DF 3 | chac1 |
| igfbp5b1 | 9E-60 | 0.4224654 | 0.689 | 0.562 | 2.94E-55 | DF 3 | igfbp5b |
| gstp12 | 1.3E-58 | 0.3981251 | 0.751 | 0.635 | 4.11E-54 | DF 3 | gstp1 |
| fosl2 | 1.8E-58 | 0.5908255 | 0.542 | 0.42 | 5.74E-54 | DF 3 | fosl2 |
| cebpb | 9E-57 | 0.4363685 | 0.569 | 0.439 | 2.92E-52 | DF 3 | cebpb |
| hspa4a | 1.3E-56 | 1.1145994 | 0.239 | 0.142 | 4.18E-52 | DF 3 | hspa4a |
| si:ch73-380l3.21 | 1.4E-56 | 0.4526196 | 0.545 | 0.406 | 4.71E-52 | DF 3 | si:ch73-380l3.2 |
| meis1b1 | 1.7E-56 | 0.565466 | 0.442 | 0.315 | 5.44E-52 | DF 3 | meis1b |
| crhr11 | 1.9E-56 | 0.8919586 | 0.151 | 0.073 | 6.28E-52 | DF 3 | crhr1 |
| ddx5 | 9E-56 | 0.5569273 | 0.687 | 0.59 | 2.94E-51 | DF 3 | ddx5 |
| ctbp2a1 | 1.5E-55 | 0.6433946 | 0.34 | 0.222 | 4.91E-51 | DF 3 | ctbp2a |
| actb22 | 1.6E-55 | 0.2159409 | 0.993 | 0.991 | 5.31E-51 | DF 3 | actb2 |
| il6st | 9E-55 | 0.6481991 | 0.525 | 0.416 | 2.94E-50 | DF 3 | il6st |
| ppl2 | 6E-54 | 0.6319021 | 0.251 | 0.147 | 1.96E-49 | DF 3 | ppl |
| socs3b | 3.5E-53 | 0.4052524 | 0.595 | 0.461 | 1.15E-48 | DF 3 | socs3b |
| gstz11 | 4E-53 | 0.5178822 | 0.461 | 0.335 | 1.31E-48 | DF 3 | gstz1 |
| igsf21a1 | 4.2E-53 | 1.0696708 | 0.12 | 0.054 | 1.36E-48 | DF 3 | igsf21a |
| clec3ba1 | 7.1E-53 | 0.7935121 | 0.212 | 0.12 | 2.31E-48 | DF 3 | clec3ba |
| pptc7a | 1.5E-52 | 0.7668773 | 0.325 | 0.216 | 4.97E-48 | DF 3 | pptc7a |
| pcbd12 | 3.4E-52 | 0.3482767 | 0.446 | 0.306 | 1.11E-47 | DF 3 | pcbd1 |
| si:ch73-237c6.12 | 4E-52 | 0.4338062 | 0.492 | 0.355 | 1.31E-47 | DF 3 | si:ch73-237c6.1 |
| pthlha1 | 1.1E-51 | 0.9199518 | 0.138 | 0.067 | 3.5E-47 | DF 3 | pthlha |
| rem1 | 1.1E-51 | 1.0587831 | 0.126 | 0.059 | 3.72E-47 | DF 3 | rem1 |
| snai1b1 | 2.6E-51 | 0.8425869 | 0.191 | 0.106 | 8.46E-47 | DF 3 | snai1b |
| crybg1b2 | 9.5E-51 | 0.3887742 | 0.6 | 0.47 | 3.08E-46 | DF 3 | crybg1b |

| | | | | | | | |
|---|---|---|---|---|---|---|---|
| tat1 | 2.1E-50 | 0.3944668 | 0.359 | 0.236 | 6.9E-46 | DF 3 | tat |
| si:ch211-195b15.8 | 4.8E-50 | 1.0098791 | 0.161 | 0.085 | 1.57E-45 | DF 3 | si:ch211-195b15.8 |
| anxa1a1 | 6.6E-50 | 0.2857109 | 0.937 | 0.902 | 2.14E-45 | DF 3 | anxa1a |
| si:dkey-16p21.81 | 9.4E-50 | 0.5986073 | 0.313 | 0.201 | 3.07E-45 | DF 3 | si:dkey-16p21.8 |
| tubb2b | 1E-48 | 0.5079678 | 0.45 | 0.333 | 3.32E-44 | DF 3 | tubb2b |
| vmp1 | 2.5E-48 | 0.4892718 | 0.431 | 0.31 | 8.1E-44 | DF 3 | vmp1 |
| ryr31 | 3.5E-48 | 0.7682757 | 0.16 | 0.084 | 1.13E-43 | DF 3 | ryr3 |
| il15l1 | 5.1E-48 | 0.9768951 | 0.197 | 0.112 | 1.65E-43 | DF 3 | il15l |
| mid1ip1l | 5.7E-48 | 0.7716638 | 0.282 | 0.183 | 1.84E-43 | DF 3 | mid1ip1l |
| celf21 | 6.7E-48 | 0.491254 | 0.301 | 0.19 | 2.19E-43 | DF 3 | celf2 |
| cyp2aa7 | 1.9E-47 | 1.1588007 | 0.112 | 0.052 | 6.12E-43 | DF 3 | cyp2aa7 |
| vegfc1 | 2.5E-47 | 0.87352 | 0.158 | 0.083 | 8.29E-43 | DF 3 | vegfc |
| ier2a | 3.6E-47 | 0.5107697 | 0.591 | 0.477 | 1.17E-42 | DF 3 | ier2a |
| peli31 | 5.3E-47 | 0.9606497 | 0.127 | 0.062 | 1.73E-42 | DF 3 | peli3 |
| ppp1r15a | 1.9E-46 | 0.760826 | 0.302 | 0.203 | 6.29E-42 | DF 3 | ppp1r15a |
| si:dkey-23i12.7 | 2.7E-46 | 0.9703495 | 0.22 | 0.134 | 8.87E-42 | DF 3 | si:dkey-23i12.7 |
| gpc6a1 | 1.3E-45 | 0.5465867 | 0.3 | 0.195 | 4.21E-41 | DF 3 | gpc6a |
| nuak1b2 | 1.7E-45 | 0.5427374 | 0.386 | 0.274 | 5.39E-41 | DF 3 | nuak1b |
| kazald33 | 1.8E-45 | 0.5634964 | 0.364 | 0.253 | 5.88E-41 | DF 3 | kazald3 |
| si:ch211-63p21.11 | 7.9E-45 | 1.0164006 | 0.102 | 0.046 | 2.57E-40 | DF 3 | si:ch211-63p21.1 |
| dchs1b1 | 2.2E-44 | 0.8969865 | 0.158 | 0.086 | 7.21E-40 | DF 3 | dchs1b |
| shox | 2.2E-44 | 1.063738 | 0.109 | 0.052 | 7.26E-40 | DF 3 | shox |
| sult6b11 | 3.2E-44 | 0.695703 | 0.265 | 0.17 | 1.05E-39 | DF 3 | sult6b1 |
| ier5l | 4.2E-44 | 0.6041972 | 0.421 | 0.312 | 1.36E-39 | DF 3 | ier5l |
| slc12a7b1 | 7.2E-44 | 0.6093775 | 0.237 | 0.145 | 2.36E-39 | DF 3 | slc12a7b |
| rpz51 | 4.4E-42 | 0.5962933 | 0.404 | 0.3 | 1.42E-37 | DF 3 | rpz5 |
| dio11 | 2.3E-41 | 0.6434746 | 0.197 | 0.117 | 7.51E-37 | DF 3 | dio1 |
| smoc11 | 1.1E-40 | 0.3709081 | 0.305 | 0.2 | 3.43E-36 | DF 3 | smoc1 |
| si:dkey-11f4.201 | 1.3E-40 | 0.3315104 | 0.596 | 0.48 | 4.34E-36 | DF 3 | si:dkey-11f4.20 |
| tuba1b | 1.4E-40 | 0.5452211 | 0.405 | 0.301 | 4.39E-36 | DF 3 | tuba1b |
| mab21l11 | 1.7E-40 | 0.7871459 | 0.215 | 0.134 | 5.38E-36 | DF 3 | mab21l1 |
| madd1 | 3.4E-40 | 0.9600588 | 0.118 | 0.059 | 1.11E-35 | DF 3 | madd |
| aff21 | 4.4E-40 | 0.6509062 | 0.263 | 0.172 | 1.42E-35 | DF 3 | aff2 |
| zcchc241 | 5.4E-40 | 0.4349112 | 0.356 | 0.249 | 1.77E-35 | DF 3 | zcchc24 |
| lmna1 | 6.3E-40 | 0.2098076 | 0.86 | 0.774 | 2.05E-35 | DF 3 | lmna |
| metrnla1 | 7.3E-40 | 0.5105147 | 0.411 | 0.308 | 2.37E-35 | DF 3 | metrnla |
| id2a1 | 8E-40 | 0.3473423 | 0.611 | 0.495 | 2.6E-35 | DF 3 | id2a |
| pmp22b1 | 1.1E-39 | 0.5093634 | 0.463 | 0.354 | 3.46E-35 | DF 3 | pmp22b |
| ly751 | 1.2E-39 | 0.7910805 | 0.127 | 0.066 | 3.82E-35 | DF 3 | ly75 |
| zfand5a | 1.4E-39 | 0.7009292 | 0.271 | 0.182 | 4.58E-35 | DF 3 | zfand5a |
| tuba8l4 | 1.5E-39 | 0.2440535 | 0.813 | 0.727 | 4.83E-35 | DF 3 | tuba8l4 |
| slc48a1b1 | 2E-39 | 0.7625869 | 0.216 | 0.135 | 6.53E-35 | DF 3 | slc48a1b |
| hspa8b | 2.3E-39 | 0.8448998 | 0.158 | 0.088 | 7.33E-35 | DF 3 | hspa8b |
| nr1d2a | 2.4E-39 | 0.5551904 | 0.344 | 0.244 | 7.81E-35 | DF 3 | nr1d2a |
| pdcd4b | 2.9E-39 | 0.9475309 | 0.174 | 0.103 | 9.34E-35 | DF 3 | pdcd4b |
| dlx4a2 | 5.6E-39 | 0.5833573 | 0.279 | 0.187 | 1.81E-34 | DF 3 | dlx4a |
| tuba8l2 | 7.2E-39 | 0.6455752 | 0.304 | 0.211 | 2.35E-34 | DF 3 | tuba8l2 |
| zgc:565852 | 1E-38 | 0.2735796 | 0.45 | 0.318 | 3.3E-34 | DF 3 | zgc:56585 |

| | | | | | | | |
|---|---|---|---|---|---|---|---|
| tfap2c1 | 1.6E-38 | 0.4387757 | 0.286 | 0.189 | 5.17E-34 | DF 3 | tfap2c |
| kera1 | 2.1E-38 | 0.436454 | 0.258 | 0.165 | 6.9E-34 | DF 3 | kera |
| nabp1a | 3E-38 | 0.6857901 | 0.26 | 0.174 | 9.66E-34 | DF 3 | nabp1a |
| klf7b2 | 3.8E-38 | 0.5159104 | 0.318 | 0.221 | 1.24E-33 | DF 3 | klf7b |
| mustn1a1 | 6.8E-38 | 0.8484291 | 0.142 | 0.078 | 2.2E-33 | DF 3 | mustn1a |
| srsf5a | 8.6E-38 | 0.3408126 | 0.664 | 0.573 | 2.81E-33 | DF 3 | srsf5a |
| klf151 | 1E-37 | 0.504216 | 0.297 | 0.202 | 3.33E-33 | DF 3 | klf15 |
| chka1 | 1.9E-37 | 0.7944331 | 0.152 | 0.086 | 6.13E-33 | DF 3 | chka |
| fabp31 | 2.2E-37 | 0.3378375 | 0.627 | 0.523 | 7.19E-33 | DF 3 | fabp3 |
| inka1b | 3.1E-37 | 0.838352 | 0.153 | 0.087 | 1.01E-32 | DF 3 | inka1b |
| hspb1 | 4.3E-37 | 1.2329543 | 0.26 | 0.174 | 1.38E-32 | DF 3 | hspb1 |
| gpr1431 | 6.6E-37 | 0.7392519 | 0.126 | 0.066 | 2.13E-32 | DF 3 | gpr143 |
| RALGDS | 4.3E-36 | 0.8235919 | 0.202 | 0.128 | 1.41E-31 | DF 3 | RALGDS |
| h2ax1 | 8.3E-36 | 0.347285 | 0.708 | 0.622 | 2.71E-31 | DF 3 | h2ax1 |
| si:dkey-192g7.3 | 1.5E-35 | 0.871482 | 0.105 | 0.053 | 4.74E-31 | DF 3 | si:dkey-192g7.3 |
| fam43a | 1.7E-35 | 0.8729793 | 0.121 | 0.064 | 5.41E-31 | DF 3 | fam43a |
| polr2a | 3.6E-35 | 0.604461 | 0.448 | 0.357 | 1.18E-30 | DF 3 | polr2a |
| hspa5 | 3.7E-35 | 0.3953273 | 0.585 | 0.485 | 1.2E-30 | DF 3 | hspa5 |
| herc11 | 5E-35 | 0.6687731 | 0.161 | 0.094 | 1.63E-30 | DF 3 | herc1 |
| tnfsf101 | 6.4E-35 | 0.6338929 | 0.161 | 0.094 | 2.08E-30 | DF 3 | tnfsf10 |
| h3f3b.1 | 8.6E-35 | 0.3811205 | 0.579 | 0.489 | 2.79E-30 | DF 3 | h3f3b.1 |
| ephb3a1 | 9.6E-35 | 0.5967537 | 0.172 | 0.101 | 3.12E-30 | DF 3 | ephb3a |
| ifrd1 | 2.1E-34 | 0.5906484 | 0.365 | 0.276 | 6.76E-30 | DF 3 | ifrd1 |
| prdm1a | 2.1E-34 | 0.8892422 | 0.132 | 0.073 | 6.82E-30 | DF 3 | prdm1a |
| ets2 | 2.8E-34 | 0.6982436 | 0.231 | 0.154 | 9.09E-30 | DF 3 | ets2 |
| si:dkey-177p2.6 | 3.2E-34 | 0.5584477 | 0.347 | 0.257 | 1.05E-29 | DF 3 | si:dkey-177p2.6 |
| skia1 | 4.8E-34 | 0.3237192 | 0.545 | 0.442 | 1.56E-29 | DF 3 | skia |
| sh3bp5la | 1.6E-33 | 0.8902209 | 0.166 | 0.101 | 5.21E-29 | DF 3 | sh3bp5la |
| zgc:1530181 | 1.9E-33 | 0.7093968 | 0.311 | 0.226 | 6.34E-29 | DF 3 | zgc:153018 |
| ebf3a1 | 2.2E-33 | 0.4373453 | 0.302 | 0.209 | 7.23E-29 | DF 3 | ebf3a |
| fth1a | 2.8E-33 | 0.3121454 | 0.769 | 0.715 | 8.98E-29 | DF 3 | fth1a |
| pcdh18a1 | 3.2E-33 | 0.6286181 | 0.139 | 0.078 | 1.03E-28 | DF 3 | pcdh18a |
| rgs5b1 | 5.1E-33 | 0.2348843 | 0.628 | 0.515 | 1.67E-28 | DF 3 | rgs5b |
| sema3e1 | 6.5E-33 | 0.5540721 | 0.155 | 0.09 | 2.11E-28 | DF 3 | sema3e |
| myclb | 9.6E-33 | 0.6182931 | 0.209 | 0.135 | 3.11E-28 | DF 3 | myclb |
| cav12 | 2.9E-32 | 0.2654005 | 0.723 | 0.626 | 9.31E-28 | DF 3 | cav1 |
| tcima | 3.7E-32 | 0.581516 | 0.305 | 0.221 | 1.2E-27 | DF 3 | tcima |
| sqstm1 | 1.2E-31 | 0.6461336 | 0.389 | 0.299 | 4.01E-27 | DF 3 | sqstm1 |
| suco2 | 1.9E-31 | 0.5467772 | 0.247 | 0.167 | 6.2E-27 | DF 3 | suco |
| si:ch211-194e15.51 | 4E-31 | 0.5693063 | 0.164 | 0.099 | 1.3E-26 | DF 3 | si:ch211-194e15.5 |
| dop1a | 6E-31 | 0.9029989 | 0.128 | 0.073 | 1.95E-26 | DF 3 | dop1a |
| twist22 | 1.1E-30 | 0.3280184 | 0.441 | 0.333 | 3.46E-26 | DF 3 | twist2 |
| nr4a1 | 1.4E-30 | 0.9544847 | 0.158 | 0.098 | 4.61E-26 | DF 3 | nr4a1 |
| hspa12b1 | 1.6E-30 | 0.5161003 | 0.188 | 0.118 | 5.23E-26 | DF 3 | hspa12b |
| cited4a | 2.3E-30 | 0.776582 | 0.175 | 0.111 | 7.49E-26 | DF 3 | cited4a |
| slc20a1b | 4.4E-30 | 0.5537999 | 0.374 | 0.288 | 1.42E-25 | DF 3 | slc20a1b |
| slc1a52 | 7.4E-30 | 0.4153772 | 0.339 | 0.252 | 2.4E-25 | DF 3 | slc1a5 |
| DAB21 | 8E-30 | 0.4311444 | 0.455 | 0.372 | 2.61E-25 | DF 3 | DAB2 |

| | | | | | | | |
|---|---|---|---|---|---|---|---|
| tmem1761 | 8.4E-30 | 0.3124953 | 0.488 | 0.387 | 2.72E-25 | DF 3 | tmem176 |
| nfil3 | 9E-30 | 0.8102047 | 0.133 | 0.077 | 2.94E-25 | DF 3 | nfil3 |
| zgc:158343 | 1.8E-29 | 0.2101307 | 0.47 | 0.371 | 5.97E-25 | DF 3 | zgc:158343 |
| ciarta1 | 2.2E-29 | 0.5123924 | 0.361 | 0.278 | 7.09E-25 | DF 3 | ciarta |
| cd248a2 | 2.2E-29 | 0.4623249 | 0.18 | 0.112 | 7.31E-25 | DF 3 | cd248a |
| klf2a | 2.6E-29 | 0.5512838 | 0.365 | 0.28 | 8.54E-25 | DF 3 | klf2a |
| caly | 2.7E-29 | 0.7740433 | 0.122 | 0.069 | 8.82E-25 | DF 3 | caly |
| PAMR11 | 2.7E-29 | 0.6234655 | 0.181 | 0.115 | 8.84E-25 | DF 3 | PAMR1 |
| zgc:1530311 | 3.6E-29 | 0.7569025 | 0.108 | 0.059 | 1.18E-24 | DF 3 | zgc:153031 |
| b2m1 | 3.9E-29 | 0.4425029 | 0.737 | 0.682 | 1.27E-24 | DF 3 | b2m |
| cdkn1bb | 4.4E-29 | 0.4529316 | 0.378 | 0.292 | 1.44E-24 | DF 3 | cdkn1bb |
| tmeff1b1 | 4.8E-29 | 0.6182039 | 0.164 | 0.102 | 1.55E-24 | DF 3 | tmeff1b |
| kng11 | 5.5E-29 | 0.6393801 | 0.132 | 0.077 | 1.79E-24 | DF 3 | kng1 |
| cdr2l | 7.9E-29 | 0.7539226 | 0.123 | 0.07 | 2.56E-24 | DF 3 | cdr2l |
| ak41 | 1.1E-28 | 0.501004 | 0.257 | 0.18 | 3.69E-24 | DF 3 | ak4 |
| fblim11 | 1.6E-28 | 0.632007 | 0.169 | 0.106 | 5.29E-24 | DF 3 | fblim1 |
| zfand2a | 3.3E-28 | 1.0168238 | 0.113 | 0.064 | 1.08E-23 | DF 3 | zfand2a |
| csrnp1b1 | 4.2E-28 | 0.5680666 | 0.256 | 0.182 | 1.37E-23 | DF 3 | csrnp1b |
| ms4a17a.92 | 5.1E-28 | 0.4528499 | 0.292 | 0.211 | 1.65E-23 | DF 3 | ms4a17a.9 |
| adipor2 | 6.1E-28 | 0.5626098 | 0.245 | 0.172 | 1.98E-23 | DF 3 | adipor2 |
| tril1 | 7.6E-28 | 0.4558041 | 0.166 | 0.102 | 2.47E-23 | DF 3 | tril |
| adma | 7.8E-28 | 0.2697664 | 0.317 | 0.228 | 2.54E-23 | DF 3 | adma |
| lrrc58b | 8.1E-28 | 0.6622664 | 0.182 | 0.118 | 2.63E-23 | DF 3 | lrrc58b |
| pim1 | 8.7E-28 | 0.3613013 | 0.622 | 0.539 | 2.82E-23 | DF 3 | pim1 |
| h3f3d | 1.5E-27 | 0.2938578 | 0.777 | 0.725 | 4.95E-23 | DF 3 | h3f3d |
| hist1h4l.3 | 1.6E-27 | 0.5656126 | 0.206 | 0.139 | 5.36E-23 | DF 3 | hist1h4l.3 |
| creb5a | 2.2E-27 | 0.9096548 | 0.11 | 0.062 | 7.06E-23 | DF 3 | creb5a |
| prickle1b1 | 2.5E-27 | 0.478874 | 0.249 | 0.173 | 8.22E-23 | DF 3 | prickle1b |
| tmc6b | 2.7E-27 | 0.7344418 | 0.12 | 0.069 | 8.65E-23 | DF 3 | tmc6b |
| zyx2 | 4E-27 | 0.3314537 | 0.404 | 0.311 | 1.3E-22 | DF 3 | zyx |
| si:dkeyp-23e4.31 | 5.5E-27 | 0.5594946 | 0.138 | 0.082 | 1.79E-22 | DF 3 | si:dkeyp-23e4.3 |
| ENSDARG000000713 | 6.7E-27 | 0.7565827 | 0.137 | 0.083 | 2.17E-22 | DF 3 | ENSDARG00000071 |
| raraa2 | 6.8E-27 | 0.3565877 | 0.335 | 0.247 | 2.21E-22 | DF 3 | raraa |
| trim3b1 | 1E-26 | 0.5965406 | 0.148 | 0.091 | 3.39E-22 | DF 3 | trim3b |
| ube2al | 1.5E-26 | 0.6020487 | 0.266 | 0.194 | 4.95E-22 | DF 3 | ube2al |
| aldob2 | 1.9E-26 | 0.3664529 | 0.396 | 0.305 | 6.34E-22 | DF 3 | aldob |
| zfp36l1b | 2E-26 | 0.3738594 | 0.626 | 0.54 | 6.66E-22 | DF 3 | zfp36l1b |
| elmsan1b | 2.3E-26 | 0.3749059 | 0.438 | 0.351 | 7.36E-22 | DF 3 | elmsan1b |
| lsp1b1 | 2.5E-26 | 0.2686723 | 0.344 | 0.252 | 8.16E-22 | DF 3 | lsp1b |
| mknk2a1 | 3.1E-26 | 0.5062642 | 0.3 | 0.224 | 1.02E-21 | DF 3 | mknk2a |
| tmtc11 | 6.4E-26 | 0.5499452 | 0.174 | 0.112 | 2.09E-21 | DF 3 | tmtc1 |
| hexim1 | 6.4E-26 | 0.5847151 | 0.302 | 0.229 | 2.09E-21 | DF 3 | hexim1 |
| scara51 | 6.6E-26 | 0.40711 | 0.218 | 0.147 | 2.16E-21 | DF 3 | scara5 |
| serpinb141 | 1E-25 | 0.4593208 | 0.218 | 0.148 | 3.37E-21 | DF 3 | serpinb14 |
| nkx3-21 | 1.2E-25 | 0.4083157 | 0.266 | 0.19 | 3.98E-21 | DF 3 | nkx3-2 |
| si:dkey-240h12.41 | 1.8E-25 | 0.4523194 | 0.176 | 0.114 | 5.75E-21 | DF 3 | si:dkey-240h12.4 |
| nfkb2 | 2.2E-25 | 0.6891236 | 0.153 | 0.097 | 7.3E-21 | DF 3 | nfkb2 |
| si:ch73-386h18.12 | 2.5E-25 | 0.534632 | 0.21 | 0.143 | 8.26E-21 | DF 3 | si:ch73-386h18.1 |

| | | | | | | | |
|---|---|---|---|---|---|---|---|
| grk41 | 3E-25 | 0.4330752 | 0.248 | 0.175 | 9.67E-21 | DF 3 | grk4 |
| letm2 | 4.2E-25 | 0.6104163 | 0.253 | 0.184 | 1.37E-20 | DF 3 | letm2 |
| si:dkey-40c11.21 | 2E-24 | 0.2942238 | 0.401 | 0.31 | 6.38E-20 | DF 3 | si:dkey-40c11.2 |
| ptmab | 2.3E-24 | 0.3340729 | 0.764 | 0.712 | 7.58E-20 | DF 3 | ptmab |
| sfxn5b2 | 2.5E-24 | 0.4277539 | 0.198 | 0.133 | 8.28E-20 | DF 3 | sfxn5b |
| hnrnpa0l.1 | 2.7E-24 | 0.2917752 | 0.729 | 0.663 | 8.79E-20 | DF 3 | hnrnpa0l.1 |
| eppk11 | 3.6E-24 | 0.2580481 | 0.285 | 0.206 | 1.16E-19 | DF 3 | eppk1 |
| adgrl2a2 | 3.7E-24 | 0.3695185 | 0.26 | 0.185 | 1.22E-19 | DF 3 | adgrl2a |
| tmem200b1 | 4E-24 | 0.5385623 | 0.145 | 0.091 | 1.29E-19 | DF 3 | tmem200b |
| plk3 | 4.8E-24 | 0.5539807 | 0.268 | 0.198 | 1.57E-19 | DF 3 | plk3 |
| ece2b1 | 5.1E-24 | 0.5487967 | 0.153 | 0.097 | 1.65E-19 | DF 3 | ece2b |
| ddit4 | 8.3E-24 | 0.7431802 | 0.128 | 0.078 | 2.69E-19 | DF 3 | ddit4 |
| arrdc22 | 1E-23 | 0.4226997 | 0.317 | 0.237 | 3.36E-19 | DF 3 | arrdc2 |
| wsb11 | 1.5E-23 | 0.4793533 | 0.252 | 0.183 | 4.99E-19 | DF 3 | wsb1 |
| nfe2l2a1 | 2.6E-23 | 0.315226 | 0.425 | 0.338 | 8.37E-19 | DF 3 | nfe2l2a |
| pim3 | 3.1E-23 | 0.7284808 | 0.157 | 0.103 | 1E-18 | DF 3 | pim3 |
| pcdh111 | 3.7E-23 | 0.5581419 | 0.12 | 0.072 | 1.21E-18 | DF 3 | pcdh11 |
| tuba8l | 3.7E-23 | 0.3240399 | 0.483 | 0.405 | 1.22E-18 | DF 3 | tuba8l |
| si:ch73-366l1.52 | 5.8E-23 | 0.7144992 | 0.128 | 0.079 | 1.88E-18 | DF 3 | si:ch73-366l1.5 |
| mcl1b | 8E-23 | 0.3983653 | 0.547 | 0.478 | 2.6E-18 | DF 3 | mcl1b |
| sesn31 | 1E-22 | 0.6123792 | 0.16 | 0.106 | 3.32E-18 | DF 3 | sesn3 |
| phc2b | 1.1E-22 | 0.8504834 | 0.119 | 0.072 | 3.42E-18 | DF 3 | phc2b |
| ywhag11 | 1.7E-22 | 0.4486797 | 0.164 | 0.107 | 5.66E-18 | DF 3 | ywhag1 |
| wnk1b1 | 2.3E-22 | 0.6131588 | 0.181 | 0.123 | 7.64E-18 | DF 3 | wnk1b |
| zfp36l1a1 | 2.6E-22 | 0.3996139 | 0.274 | 0.202 | 8.53E-18 | DF 3 | zfp36l1a |
| f3b1 | 2.7E-22 | 0.4483971 | 0.451 | 0.375 | 8.85E-18 | DF 3 | f3b |
| rnd1b1 | 2.9E-22 | 0.6683135 | 0.152 | 0.099 | 9.4E-18 | DF 3 | rnd1b |
| higd1a | 7.3E-22 | 0.3565387 | 0.524 | 0.453 | 2.37E-17 | DF 3 | higd1a |
| zgc:194989 | 1.1E-21 | 0.7116086 | 0.136 | 0.087 | 3.7E-17 | DF 3 | zgc:194989 |
| AL929435.1 | 1.2E-21 | 0.7556643 | 0.154 | 0.103 | 4.06E-17 | DF 3 | AL929435.1 |
| cgnl11 | 1.5E-21 | 0.5698227 | 0.114 | 0.069 | 5.03E-17 | DF 3 | cgnl1 |
| hyal2b2 | 2.1E-21 | 0.2902075 | 0.263 | 0.192 | 6.68E-17 | DF 3 | hyal2b |
| si:ch211-225b11.1 | 6E-21 | 0.7194032 | 0.116 | 0.072 | 1.96E-16 | DF 3 | si:ch211-225b11.1 |
| hist1h4l.18 | 6.4E-21 | 0.4044002 | 0.173 | 0.117 | 2.09E-16 | DF 3 | hist1h4l.18 |
| stmn1b1 | 6.7E-21 | 0.2525393 | 0.178 | 0.12 | 2.19E-16 | DF 3 | stmn1b |
| mepce | 6.9E-21 | 0.7205471 | 0.15 | 0.1 | 2.26E-16 | DF 3 | mepce |
| acsl4a | 1E-20 | 0.7411287 | 0.12 | 0.075 | 3.36E-16 | DF 3 | acsl4a |
| ghrb2 | 1.3E-20 | 0.5287174 | 0.158 | 0.105 | 4.31E-16 | DF 3 | ghrb |
| pdk2b1 | 3E-20 | 0.6058094 | 0.257 | 0.195 | 9.75E-16 | DF 3 | pdk2b |
| h1-101 | 4.8E-20 | 0.4392124 | 0.265 | 0.201 | 1.56E-15 | DF 3 | h1-10 |
| kdm6ba | 4.8E-20 | 0.5205161 | 0.281 | 0.217 | 1.57E-15 | DF 3 | kdm6ba |
| kdm6bb | 5.5E-20 | 0.5027598 | 0.194 | 0.137 | 1.8E-15 | DF 3 | kdm6bb |
| il13ra1 | 8.7E-20 | 0.6694768 | 0.127 | 0.082 | 2.83E-15 | DF 3 | il13ra1 |
| tsc22d3 | 1.2E-19 | 0.2835534 | 0.475 | 0.397 | 4.01E-15 | DF 3 | tsc22d3 |
| irs2b1 | 1.3E-19 | 0.4168249 | 0.293 | 0.226 | 4.15E-15 | DF 3 | irs2b |
| zeb2a2 | 1.3E-19 | 0.2267705 | 0.418 | 0.336 | 4.28E-15 | DF 3 | zeb2a |
| sik2b | 1.6E-19 | 0.3449106 | 0.27 | 0.203 | 5.35E-15 | DF 3 | sik2b |
| ccnl1a | 1.7E-19 | 0.3832858 | 0.326 | 0.258 | 5.5E-15 | DF 3 | ccnl1a |

| | | | | | | | |
|---|---|---|---|---|---|---|---|
| si:ch211-212k18.91 | 3.6E-19 | 0.3811137 | 0.237 | 0.173 | 1.17E-14 | DF 3 | si:ch211-212k18.9 |
| nocta | 4E-19 | 0.8719167 | 0.159 | 0.11 | 1.29E-14 | DF 3 | nocta |
| rbbp6 | 4.8E-19 | 0.4253298 | 0.378 | 0.312 | 1.55E-14 | DF 3 | rbbp6 |
| si:ch211-110e21.31 | 8.7E-19 | 0.5476889 | 0.117 | 0.074 | 2.83E-14 | DF 3 | si:ch211-110e21.3 |
| mao1 | 1.1E-18 | 0.5354171 | 0.112 | 0.07 | 3.51E-14 | DF 3 | mao |
| itga6b1 | 1.7E-18 | 0.2391096 | 0.371 | 0.291 | 5.61E-14 | DF 3 | itga6b |
| gstt1a1 | 2.2E-18 | 0.6274399 | 0.114 | 0.073 | 7.29E-14 | DF 3 | gstt1a |
| heyl1 | 2.5E-18 | 0.4999152 | 0.111 | 0.069 | 8.08E-14 | DF 3 | heyl |
| GNAZ1 | 3.3E-18 | 0.3412102 | 0.188 | 0.132 | 1.07E-13 | DF 3 | GNAZ |
| slc7a8a1 | 3.3E-18 | 0.6393151 | 0.126 | 0.082 | 1.07E-13 | DF 3 | slc7a8a |
| CU984600.12 | 4.2E-18 | 0.3659034 | 0.227 | 0.166 | 1.37E-13 | DF 3 | CU984600.1 |
| lgals3b1 | 5.3E-18 | 0.242641 | 0.475 | 0.403 | 1.73E-13 | DF 3 | lgals3b |
| dpyda.12 | 5.8E-18 | 0.5092523 | 0.157 | 0.108 | 1.88E-13 | DF 3 | dpyda.1 |
| pdk2a | 1.3E-17 | 0.6892449 | 0.122 | 0.08 | 4.22E-13 | DF 3 | pdk2a |
| adamts15a2 | 1.3E-17 | 0.339196 | 0.195 | 0.14 | 4.3E-13 | DF 3 | adamts15a |
| camk2n1a1 | 1.6E-17 | 0.63318 | 0.123 | 0.081 | 5.3E-13 | DF 3 | camk2n1a |
| tspan311 | 1.6E-17 | 0.3725113 | 0.287 | 0.222 | 5.32E-13 | DF 3 | tspan31 |
| mrtfab1 | 3.4E-17 | 0.5618182 | 0.126 | 0.083 | 1.12E-12 | DF 3 | mrtfab |
| unc119.1 | 4.3E-17 | 0.6578028 | 0.116 | 0.075 | 1.4E-12 | DF 3 | unc119.1 |
| cx35.41 | 4.4E-17 | 0.5579071 | 0.115 | 0.074 | 1.42E-12 | DF 3 | cx35.4 |
| rab40b | 4.7E-17 | 0.6017106 | 0.117 | 0.076 | 1.52E-12 | DF 3 | rab40b |
| slc25a33 | 5E-17 | 0.6382704 | 0.161 | 0.114 | 1.62E-12 | DF 3 | slc25a33 |
| aldh9a1a.12 | 5.8E-17 | 0.2162638 | 0.55 | 0.476 | 1.87E-12 | DF 3 | aldh9a1a.1 |
| oser1 | 6.5E-17 | 0.7132339 | 0.125 | 0.084 | 2.11E-12 | DF 3 | oser1 |
| ramp21 | 8E-17 | 0.3124467 | 0.192 | 0.138 | 2.6E-12 | DF 3 | ramp2 |
| si:ch211-212k18.81 | 1E-16 | 0.4324205 | 0.191 | 0.138 | 3.32E-12 | DF 3 | si:ch211-212k18.8 |
| efna1a1 | 1.1E-16 | 0.3985477 | 0.229 | 0.173 | 3.59E-12 | DF 3 | efna1a |
| map2k6 | 2.4E-16 | 0.4051531 | 0.243 | 0.186 | 7.89E-12 | DF 3 | map2k6 |
| map3k14a1 | 2.5E-16 | 0.7112722 | 0.112 | 0.074 | 7.98E-12 | DF 3 | map3k14a |
| rgs4 | 5.4E-16 | 0.4358069 | 0.106 | 0.068 | 1.74E-11 | DF 3 | rgs4 |
| pik3r3a | 6.2E-16 | 0.4821145 | 0.181 | 0.132 | 2E-11 | DF 3 | pik3r3a |
| atp1a1a.1 | 8.4E-16 | 0.3762555 | 0.379 | 0.319 | 2.72E-11 | DF 3 | atp1a1a.1 |
| glud1a1 | 8.8E-16 | 0.2166365 | 0.507 | 0.436 | 2.88E-11 | DF 3 | glud1a |
| slc44a1b2 | 1E-15 | 0.391116 | 0.164 | 0.116 | 3.25E-11 | DF 3 | slc44a1b |
| zgc:112234 | 1E-15 | 0.6271254 | 0.109 | 0.072 | 3.35E-11 | DF 3 | zgc:112234 |
| ip6k2a | 1.1E-15 | 0.392408 | 0.249 | 0.193 | 3.54E-11 | DF 3 | ip6k2a |
| gadd45bb | 1.2E-15 | 0.4253527 | 0.272 | 0.217 | 3.82E-11 | DF 3 | gadd45bb |
| enah2 | 1.7E-15 | 0.3562482 | 0.16 | 0.112 | 5.55E-11 | DF 3 | enah |
| tsc22d1 | 1.8E-15 | 0.5360069 | 0.157 | 0.112 | 5.95E-11 | DF 3 | tsc22d1 |
| zgc:92140 | 2.1E-15 | 0.4420728 | 0.156 | 0.111 | 6.78E-11 | DF 3 | zgc:92140 |
| anxa11a1 | 3.9E-15 | 0.219518 | 0.473 | 0.406 | 1.25E-10 | DF 3 | anxa11a |
| NPAS31 | 3.9E-15 | 0.4355295 | 0.124 | 0.083 | 1.26E-10 | DF 3 | NPAS3 |
| tiparp | 4.4E-15 | 0.464626 | 0.169 | 0.122 | 1.45E-10 | DF 3 | tiparp |
| dcbld21 | 6.7E-15 | 0.351606 | 0.169 | 0.122 | 2.19E-10 | DF 3 | dcbld2 |
| slc35c21 | 6.8E-15 | 0.3759288 | 0.198 | 0.148 | 2.21E-10 | DF 3 | slc35c2 |
| alas1 | 1E-14 | 0.6774962 | 0.145 | 0.104 | 3.31E-10 | DF 3 | alas1 |
| kctd11 | 1.1E-14 | 0.4794737 | 0.115 | 0.077 | 3.64E-10 | DF 3 | kctd1 |
| mvp | 1.2E-14 | 0.3928074 | 0.368 | 0.313 | 3.74E-10 | DF 3 | mvp |

| | | | | | | | |
|---|---|---|---|---|---|---|---|
| abca1a | 1.2E-14 | 0.3023063 | 0.25 | 0.193 | 3.76E-10 | DF 3 | abca1a |
| ehbp11 | 1.3E-14 | 0.4409382 | 0.151 | 0.108 | 4.14E-10 | DF 3 | ehbp1 |
| zgc:110339 | 1.5E-14 | 0.347351 | 0.267 | 0.212 | 4.76E-10 | DF 3 | zgc:110339 |
| emx2 | 2.1E-14 | 0.5430377 | 0.129 | 0.09 | 6.75E-10 | DF 3 | emx2 |
| lef1 | 2.1E-14 | 0.4109268 | 0.149 | 0.105 | 6.79E-10 | DF 3 | lef1 |
| gadd45ga | 2.2E-14 | 0.3886588 | 0.331 | 0.273 | 7.08E-10 | DF 3 | gadd45ga |
| CPNE81 | 2.5E-14 | 0.263413 | 0.346 | 0.282 | 8.17E-10 | DF 3 | CPNE8 |
| kif26ba2 | 2.8E-14 | 0.3449956 | 0.149 | 0.105 | 9.12E-10 | DF 3 | kif26ba |
| abcf2a | 2.9E-14 | 0.51391 | 0.224 | 0.175 | 9.38E-10 | DF 3 | abcf2a |
| ptpn3 | 2.9E-14 | 0.5319118 | 0.102 | 0.067 | 9.56E-10 | DF 3 | ptpn3 |
| mta1 | 4.8E-14 | 0.4792866 | 0.126 | 0.088 | 1.57E-09 | DF 3 | mta1 |
| arid5b1 | 5E-14 | 0.3475846 | 0.18 | 0.132 | 1.61E-09 | DF 3 | arid5b |
| pdzrn3b | 7.8E-14 | 0.4532603 | 0.114 | 0.077 | 2.55E-09 | DF 3 | pdzrn3b |
| ehbp1l1a1 | 1E-13 | 0.4211509 | 0.131 | 0.092 | 3.36E-09 | DF 3 | ehbp1l1a |
| maff | 1.3E-13 | 0.4154969 | 0.271 | 0.22 | 4.29E-09 | DF 3 | maff |
| myo1ca1 | 1.4E-13 | 0.2004476 | 0.257 | 0.199 | 4.52E-09 | DF 3 | myo1ca |
| birc2 | 1.4E-13 | 0.4807119 | 0.177 | 0.133 | 4.7E-09 | DF 3 | birc2 |
| slc12a21 | 1.8E-13 | 0.2913614 | 0.189 | 0.141 | 5.75E-09 | DF 3 | slc12a2 |
| igf2a1 | 2E-13 | 0.3301944 | 0.129 | 0.09 | 6.53E-09 | DF 3 | igf2a |
| pik3ip1 | 2E-13 | 0.4952732 | 0.108 | 0.073 | 6.64E-09 | DF 3 | pik3ip1 |
| cish | 2.4E-13 | 0.487923 | 0.196 | 0.151 | 7.75E-09 | DF 3 | cish |
| rac3b1 | 2.7E-13 | 0.4217002 | 0.117 | 0.08 | 8.62E-09 | DF 3 | rac3b |
| uchl1 | 3.2E-13 | 0.4486832 | 0.168 | 0.125 | 1.04E-08 | DF 3 | uchl1 |
| si:ch211-286o17.12 | 3.5E-13 | 0.235656 | 0.269 | 0.213 | 1.14E-08 | DF 3 | si:ch211-286o17.1 |
| scarb2a | 3.9E-13 | 0.3322282 | 0.236 | 0.185 | 1.27E-08 | DF 3 | scarb2a |
| tead1b1 | 4.8E-13 | 0.2840092 | 0.347 | 0.289 | 1.56E-08 | DF 3 | tead1b |
| slc16a101 | 1.1E-12 | 0.3903067 | 0.174 | 0.13 | 3.42E-08 | DF 3 | slc16a10 |
| jmjd1cb1 | 1.2E-12 | 0.2683446 | 0.371 | 0.315 | 4.01E-08 | DF 3 | jmjd1cb |
| sema3aa1 | 1.3E-12 | 0.2114608 | 0.148 | 0.106 | 4.28E-08 | DF 3 | sema3aa |
| ptbp31 | 1.3E-12 | 0.3660774 | 0.176 | 0.132 | 4.36E-08 | DF 3 | ptbp3 |
| tmem59l1 | 1.3E-12 | 0.372556 | 0.144 | 0.104 | 4.36E-08 | DF 3 | tmem59l |
| ddx61 | 1.6E-12 | 0.2381454 | 0.404 | 0.344 | 5.14E-08 | DF 3 | ddx6 |
| rif1 | 1.7E-12 | 0.6039274 | 0.119 | 0.084 | 5.44E-08 | DF 3 | rif1 |
| zswim5 | 1.8E-12 | 0.3454473 | 0.225 | 0.177 | 5.81E-08 | DF 3 | zswim5 |
| fbxl71 | 2.1E-12 | 0.3931647 | 0.109 | 0.074 | 6.84E-08 | DF 3 | fbxl7 |
| c1qtnf123 | 2.3E-12 | 0.2508218 | 0.138 | 0.098 | 7.44E-08 | DF 3 | c1qtnf12 |
| mych1 | 2.3E-12 | 0.2461787 | 0.435 | 0.376 | 7.63E-08 | DF 3 | mych |
| ponzr1 | 2.6E-12 | 0.2441343 | 0.286 | 0.23 | 8.32E-08 | DF 3 | ponzr1 |
| mylipb | 2.7E-12 | 0.5831671 | 0.112 | 0.078 | 8.76E-08 | DF 3 | mylipb |
| nedd4a1 | 2.9E-12 | 0.4698024 | 0.125 | 0.089 | 9.55E-08 | DF 3 | nedd4a |
| dusp21 | 3.7E-12 | 0.4329845 | 0.266 | 0.22 | 1.21E-07 | DF 3 | dusp2 |
| nrp2a1 | 3.9E-12 | 0.4132023 | 0.167 | 0.126 | 1.27E-07 | DF 3 | nrp2a |
| lpar11 | 4E-12 | 0.2266452 | 0.23 | 0.179 | 1.3E-07 | DF 3 | lpar1 |
| lmo4a | 6E-12 | 0.3714142 | 0.164 | 0.123 | 1.95E-07 | DF 3 | lmo4a |
| alyref | 6.5E-12 | 0.2370765 | 0.399 | 0.344 | 2.12E-07 | DF 3 | alyref |
| tspan18b2 | 6.6E-12 | 0.3150755 | 0.183 | 0.139 | 2.13E-07 | DF 3 | tspan18b |
| itgb21 | 7E-12 | 0.2945275 | 0.143 | 0.103 | 2.29E-07 | DF 3 | itgb2 |
| lrrc151 | 8.2E-12 | 0.2581221 | 0.176 | 0.132 | 2.68E-07 | DF 3 | lrrc15 |

| | | | | | | | |
|---|---|---|---|---|---|---|---|
| fryl | 9E-12 | 0.4517083 | 0.142 | 0.104 | 2.91E-07 | DF 3 | fryl |
| arg2 | 9.1E-12 | 0.6021021 | 0.203 | 0.161 | 2.97E-07 | DF 3 | arg2 |
| myca | 9.7E-12 | 0.4734218 | 0.197 | 0.155 | 3.16E-07 | DF 3 | myca |
| dhrs91 | 1.5E-11 | 0.2223135 | 0.183 | 0.138 | 4.74E-07 | DF 3 | dhrs9 |
| adh8b1 | 1.5E-11 | 0.2625168 | 0.282 | 0.231 | 4.79E-07 | DF 3 | adh8b |
| irs11 | 1.5E-11 | 0.4297809 | 0.171 | 0.13 | 4.79E-07 | DF 3 | irs1 |
| ms4a17a.81 | 1.7E-11 | 0.4110024 | 0.149 | 0.111 | 5.68E-07 | DF 3 | ms4a17a.8 |
| mhc1zaa1 | 1.9E-11 | 0.2514846 | 0.128 | 0.091 | 6.08E-07 | DF 3 | mhc1zaa |
| relb | 2E-11 | 0.4435576 | 0.114 | 0.08 | 6.55E-07 | DF 3 | relb |
| nr1d1 | 2.1E-11 | 0.4318307 | 0.173 | 0.133 | 6.95E-07 | DF 3 | nr1d1 |
| nck1a | 2.6E-11 | 0.413319 | 0.11 | 0.077 | 8.49E-07 | DF 3 | nck1a |
| zgc:114046 | 2.7E-11 | 0.4639606 | 0.134 | 0.098 | 8.81E-07 | DF 3 | zgc:114046 |
| ivns1abpa1 | 2.8E-11 | 0.3925409 | 0.33 | 0.284 | 8.98E-07 | DF 3 | ivns1abpa |
| tgm1l11 | 3.1E-11 | 0.2057847 | 0.167 | 0.125 | 1E-06 | DF 3 | tgm1l1 |
| stat1a1 | 4.6E-11 | 0.256223 | 0.389 | 0.336 | 1.51E-06 | DF 3 | stat1a |
| rnd3b1 | 5E-11 | 0.221802 | 0.382 | 0.328 | 1.61E-06 | DF 3 | rnd3b |
| BX548028.1 | 5E-11 | 0.5814025 | 0.103 | 0.072 | 1.62E-06 | DF 3 | BX548028.1 |
| ccdc1251 | 5.3E-11 | 0.3937914 | 0.105 | 0.073 | 1.72E-06 | DF 3 | ccdc125 |
| eif4a1a | 5.9E-11 | 0.233209 | 0.418 | 0.362 | 1.93E-06 | DF 3 | eif4a1a |
| ITLN12 | 9.2E-11 | 0.2393443 | 0.258 | 0.208 | 2.99E-06 | DF 3 | ITLN1 |
| mrtfbb | 9.7E-11 | 0.4469372 | 0.101 | 0.07 | 3.15E-06 | DF 3 | mrtfbb |
| entpd11 | 9.9E-11 | 0.3106761 | 0.113 | 0.08 | 3.23E-06 | DF 3 | entpd1 |
| foxj31 | 1.3E-10 | 0.3618345 | 0.177 | 0.138 | 4.28E-06 | DF 3 | foxj3 |
| tpst11 | 1.6E-10 | 0.2523728 | 0.219 | 0.173 | 5.34E-06 | DF 3 | tpst1 |
| rgmd | 1.8E-10 | 0.3662785 | 0.116 | 0.083 | 5.98E-06 | DF 3 | rgmd |
| snai1a | 1.9E-10 | 0.2741467 | 0.319 | 0.269 | 6.22E-06 | DF 3 | snai1a |
| sox11b1 | 1.9E-10 | 0.3249927 | 0.253 | 0.207 | 6.29E-06 | DF 3 | sox11b |
| kras | 2E-10 | 0.436518 | 0.216 | 0.176 | 6.41E-06 | DF 3 | kras |
| si:dkey-81l17.6 | 2.1E-10 | 0.4434824 | 0.107 | 0.076 | 6.72E-06 | DF 3 | si:dkey-81l17.6 |
| plcd4b1 | 2.1E-10 | 0.3788959 | 0.111 | 0.079 | 6.9E-06 | DF 3 | plcd4b |
| pfkfb3 | 2.7E-10 | 0.4955763 | 0.105 | 0.074 | 8.62E-06 | DF 3 | pfkfb3 |
| cpeb4b | 2.7E-10 | 0.4391875 | 0.205 | 0.165 | 8.63E-06 | DF 3 | cpeb4b |
| si:dkey-23i12.5 | 2.7E-10 | 0.3533948 | 0.26 | 0.216 | 8.69E-06 | DF 3 | si:dkey-23i12.5 |
| dusp6 | 2.7E-10 | 0.4159222 | 0.183 | 0.144 | 8.79E-06 | DF 3 | dusp6 |
| bach1b | 3.4E-10 | 0.541778 | 0.123 | 0.091 | 1.11E-05 | DF 3 | bach1b |
| ddr11 | 4E-10 | 0.2601024 | 0.117 | 0.084 | 1.3E-05 | DF 3 | ddr1 |
| crsp7 | 5.2E-10 | 0.5328594 | 0.1 | 0.071 | 1.68E-05 | DF 3 | crsp7 |
| rhoq | 5.9E-10 | 0.2477368 | 0.128 | 0.094 | 1.93E-05 | DF 3 | rhoq |
| si:ch211-15b10.6 | 6.9E-10 | 0.4921 | 0.131 | 0.098 | 2.25E-05 | DF 3 | si:ch211-15b10.6 |
| mapk6 | 7.1E-10 | 0.5471476 | 0.116 | 0.086 | 2.31E-05 | DF 3 | mapk6 |
| gstt1b1 | 9.1E-10 | 0.2946126 | 0.157 | 0.12 | 2.96E-05 | DF 3 | gstt1b |
| slc43a3b1 | 9.7E-10 | 0.3443071 | 0.107 | 0.077 | 3.16E-05 | DF 3 | slc43a3b |
| xpo1b | 1E-09 | 0.3736169 | 0.133 | 0.1 | 3.32E-05 | DF 3 | xpo1b |
| diaph21 | 1.3E-09 | 0.3077755 | 0.15 | 0.114 | 4.15E-05 | DF 3 | diaph2 |
| rdh10b | 1.4E-09 | 0.5376087 | 0.101 | 0.073 | 4.49E-05 | DF 3 | rdh10b |
| amd1 | 1.4E-09 | 0.5627728 | 0.125 | 0.094 | 4.71E-05 | DF 3 | amd1 |
| cdkn1a | 1.6E-09 | 0.5159447 | 0.154 | 0.12 | 5.19E-05 | DF 3 | cdkn1a |
| psd21 | 1.9E-09 | 0.2823592 | 0.128 | 0.095 | 6.02E-05 | DF 3 | psd2 |

| | | | | | | | |
|---|---|---|---|---|---|---|---|
| tnfrsf19 | 1.9E-09 | 0.2479726 | 0.142 | 0.106 | 6.22E-05 | DF 3 | tnfrsf19 |
| zgc:77486.1 | 2E-09 | 0.3429606 | 0.207 | 0.167 | 6.42E-05 | DF 3 | zgc:77486.1 |
| rnf111 | 2.6E-09 | 0.4185811 | 0.123 | 0.091 | 8.3E-05 | DF 3 | rnf111 |
| tet3 | 2.6E-09 | 0.3865417 | 0.129 | 0.097 | 8.61E-05 | DF 3 | tet3 |
| myd88 | 2.9E-09 | 0.4903616 | 0.105 | 0.076 | 9.29E-05 | DF 3 | myd88 |
| ctdsp2 | 3.3E-09 | 0.2196436 | 0.232 | 0.188 | 0.000106 | DF 3 | ctdsp2 |
| mapk14a | 4.3E-09 | 0.4598147 | 0.111 | 0.082 | 0.000141 | DF 3 | mapk14a |
| im:7152348 | 4.5E-09 | 0.2771582 | 0.276 | 0.234 | 0.000146 | DF 3 | im:7152348 |
| hlfa1 | 5E-09 | 0.2773964 | 0.166 | 0.13 | 0.000162 | DF 3 | hlfa |
| tlcd4b1 | 5.1E-09 | 0.3743224 | 0.111 | 0.081 | 0.000165 | DF 3 | tlcd4b |
| cadm41 | 5.2E-09 | 0.2333311 | 0.103 | 0.074 | 0.00017 | DF 3 | cadm4 |
| trim8a | 5.8E-09 | 0.4604012 | 0.11 | 0.081 | 0.00019 | DF 3 | trim8a |
| ubap2a1 | 6.2E-09 | 0.2500007 | 0.233 | 0.192 | 0.000203 | DF 3 | ubap2a |
| si:ch211-57n23.11 | 7.2E-09 | 0.2212717 | 0.26 | 0.215 | 0.000236 | DF 3 | si:ch211-57n23.1 |
| si:ch211-178n15.1 | 7.6E-09 | 0.3537041 | 0.205 | 0.167 | 0.000248 | DF 3 | si:ch211-178n15.1 |
| adam8b | 8.5E-09 | 0.6301526 | 0.105 | 0.078 | 0.000277 | DF 3 | adam8b |
| mpp6a | 8.7E-09 | 0.4220218 | 0.108 | 0.08 | 0.000284 | DF 3 | mpp6a |
| her9 | 8.8E-09 | 0.445965 | 0.263 | 0.226 | 0.000285 | DF 3 | her9 |
| agmo1 | 1.1E-08 | 0.3221483 | 0.109 | 0.081 | 0.000373 | DF 3 | agmo |
| socs1a1 | 1.9E-08 | 0.2896557 | 0.156 | 0.122 | 0.000614 | DF 3 | socs1a |
| si:dkey-51e6.11 | 2.2E-08 | 0.2005063 | 0.302 | 0.255 | 0.000704 | DF 3 | si:dkey-51e6.1 |
| cmtm31 | 2.3E-08 | 0.274891 | 0.231 | 0.192 | 0.000737 | DF 3 | cmtm3 |
| clk4a1 | 2.5E-08 | 0.2504143 | 0.269 | 0.227 | 0.000809 | DF 3 | clk4a |
| lima1a | 3.1E-08 | 0.3398936 | 0.117 | 0.088 | 0.001014 | DF 3 | lima1a |
| ndfip21 | 3.1E-08 | 0.2400778 | 0.326 | 0.284 | 0.001019 | DF 3 | ndfip2 |
| tent4b | 3.5E-08 | 0.2685043 | 0.24 | 0.202 | 0.001126 | DF 3 | tent4b |
| BX005309.1 | 4.4E-08 | 0.3633569 | 0.108 | 0.081 | 0.001416 | DF 3 | BX005309.1 |
| dip2ca1 | 4.4E-08 | 0.3142537 | 0.113 | 0.085 | 0.001426 | DF 3 | dip2ca |
| bahcc1a | 4.4E-08 | 0.4466641 | 0.113 | 0.085 | 0.001439 | DF 3 | bahcc1a |
| rhpn21 | 7.1E-08 | 0.3566785 | 0.174 | 0.141 | 0.002306 | DF 3 | rhpn2 |
| mhc1uba1 | 7.6E-08 | 0.2685831 | 0.252 | 0.212 | 0.002469 | DF 3 | mhc1uba |
| zmp:00000007111 | 8.3E-08 | 0.3813323 | 0.107 | 0.081 | 0.002683 | DF 3 | zmp:0000000711 |
| grinab | 8.9E-08 | 0.2367543 | 0.248 | 0.21 | 0.002882 | DF 3 | grinab |
| efnb3b1 | 9.5E-08 | 0.237022 | 0.171 | 0.137 | 0.0031 | DF 3 | efnb3b |
| ptenb | 9.9E-08 | 0.2137561 | 0.21 | 0.173 | 0.003205 | DF 3 | ptenb |
| si:dkey-112a7.4 | 1E-07 | 0.379595 | 0.114 | 0.087 | 0.003369 | DF 3 | si:dkey-112a7.4 |
| top11 | 1.1E-07 | 0.2433557 | 0.108 | 0.08 | 0.003558 | DF 3 | top1 |
| stard13b | 1.2E-07 | 0.3327229 | 0.109 | 0.082 | 0.003749 | DF 3 | stard13b |
| mob1ba1 | 1.4E-07 | 0.3519639 | 0.128 | 0.099 | 0.004617 | DF 3 | mob1ba |
| tspan351 | 1.5E-07 | 0.2435901 | 0.154 | 0.122 | 0.004806 | DF 3 | tspan35 |
| itga91 | 1.5E-07 | 0.2393007 | 0.158 | 0.125 | 0.004833 | DF 3 | itga9 |
| gpt2l1 | 2E-07 | 0.2301105 | 0.16 | 0.128 | 0.006505 | DF 3 | gpt2l |
| creb1a | 2.1E-07 | 0.3216313 | 0.193 | 0.161 | 0.006802 | DF 3 | creb1a |
| rasl11b | 2.1E-07 | 0.2987555 | 0.251 | 0.215 | 0.006921 | DF 3 | rasl11b |
| slc44a21 | 2.7E-07 | 0.2275087 | 0.143 | 0.113 | 0.008843 | DF 3 | slc44a2 |
| ttyh3a1 | 3.7E-07 | 0.2716163 | 0.123 | 0.095 | 0.012171 | DF 3 | ttyh3a |
| slc25a481 | 3.8E-07 | 0.3158122 | 0.168 | 0.137 | 0.012259 | DF 3 | slc25a48 |
| tnfaip2a | 3.9E-07 | 0.4064058 | 0.113 | 0.087 | 0.012594 | DF 3 | tnfaip2a |

| | | | | | | | |
|---|---|---|---|---|---|---|---|
| peli1b | 3.9E-07 | 0.3315794 | 0.115 | 0.089 | 0.012738 | DF 3 | peli1b |
| zgc:1008291 | 5.1E-07 | 0.3248554 | 0.146 | 0.118 | 0.01664 | DF 3 | zgc:100829 |
| gcdha | 6.3E-07 | 0.3155787 | 0.135 | 0.107 | 0.020344 | DF 3 | gcdha |
| zgc:1527911 | 7.6E-07 | 0.2372189 | 0.132 | 0.104 | 0.024776 | DF 3 | zgc:152791 |
| oaz1b | 8E-07 | 0.2709005 | 0.34 | 0.306 | 0.026082 | DF 3 | oaz1b |
| phip | 8.2E-07 | 0.3020032 | 0.187 | 0.156 | 0.026718 | DF 3 | phip |
| slc3a2b | 8.3E-07 | 0.3394099 | 0.166 | 0.137 | 0.026908 | DF 3 | slc3a2b |
| ulk2 | 8.9E-07 | 0.374339 | 0.101 | 0.077 | 0.028948 | DF 3 | ulk2 |
| piezo11 | 9.6E-07 | 0.2683362 | 0.152 | 0.122 | 0.031137 | DF 3 | piezo1 |
| crybg1a1 | 1E-06 | 0.2545588 | 0.252 | 0.218 | 0.032855 | DF 3 | crybg1a |
| gtf2b | 1E-06 | 0.562817 | 0.1 | 0.078 | 0.032944 | DF 3 | gtf2b |
| ube2d2l | 1.1E-06 | 0.2801699 | 0.178 | 0.147 | 0.034395 | DF 3 | ube2d2l |
| rassf1 | 1.4E-06 | 0.4471981 | 0.146 | 0.119 | 0.043907 | DF 3 | rassf1 |
| zgc:92242 | 1.4E-06 | 0.2418847 | 0.137 | 0.109 | 0.043984 | DF 3 | zgc:92242 |
| apmap | 1.4E-06 | 0.3899765 | 0.102 | 0.079 | 0.046016 | DF 3 | apmap |
| vamp5 | 1.5E-06 | 0.2617421 | 0.103 | 0.079 | 0.049969 | DF 3 | vamp5 |
| enc11 | 1.6E-06 | 0.3583861 | 0.151 | 0.123 | 0.053135 | DF 3 | enc1 |
| spen | 1.7E-06 | 0.3164412 | 0.229 | 0.196 | 0.055243 | DF 3 | spen |
| aff4 | 1.7E-06 | 0.3173086 | 0.109 | 0.084 | 0.05576 | DF 3 | aff4 |
| rprd2b1 | 2.1E-06 | 0.2994177 | 0.149 | 0.121 | 0.068624 | DF 3 | rprd2b |
| nr1d2b | 2.5E-06 | 0.2949893 | 0.18 | 0.151 | 0.081289 | DF 3 | nr1d2b |
| si:ch1073-185p12.2 | 2.5E-06 | 0.3625512 | 0.105 | 0.082 | 0.081636 | DF 3 | si:ch1073-185p12.2 |
| zgc:1107961 | 2.5E-06 | 0.3890011 | 0.102 | 0.08 | 0.082757 | DF 3 | zgc:110796 |
| rmdn1 | 2.6E-06 | 0.3008861 | 0.117 | 0.093 | 0.083296 | DF 3 | rmdn1 |
| ep300b | 2.6E-06 | 0.2384107 | 0.159 | 0.13 | 0.084761 | DF 3 | ep300b |
| sorbs11 | 2.7E-06 | 0.2502115 | 0.162 | 0.133 | 0.087419 | DF 3 | sorbs1 |
| lonrf1l | 2.8E-06 | 0.3160034 | 0.18 | 0.15 | 0.090588 | DF 3 | lonrf1l |
| vegfaa | 2.9E-06 | 0.2517898 | 0.142 | 0.115 | 0.094286 | DF 3 | vegfaa |
| dennd4a | 3.6E-06 | 0.3301045 | 0.109 | 0.085 | 0.115816 | DF 3 | dennd4a |
| atf4a | 3.6E-06 | 0.2519572 | 0.328 | 0.296 | 0.116818 | DF 3 | atf4a |
| arl4ab | 5.4E-06 | 0.3904548 | 0.134 | 0.109 | 0.175497 | DF 3 | arl4ab |
| galnt7 | 5.7E-06 | 0.3490839 | 0.142 | 0.116 | 0.185176 | DF 3 | galnt7 |
| znf706 | 5.8E-06 | 0.29437 | 0.119 | 0.095 | 0.190042 | DF 3 | znf706 |
| tshz11 | 7E-06 | 0.2182315 | 0.163 | 0.135 | 0.228834 | DF 3 | tshz1 |
| tgif1 | 7.1E-06 | 0.3602535 | 0.232 | 0.204 | 0.230205 | DF 3 | tgif1 |
| fam171a2a | 1.1E-05 | 0.2313732 | 0.104 | 0.082 | 0.353953 | DF 3 | fam171a2a |
| rin3 | 1.3E-05 | 0.2377946 | 0.181 | 0.153 | 0.416954 | DF 3 | rin3 |
| wtap | 1.4E-05 | 0.2427744 | 0.151 | 0.125 | 0.440492 | DF 3 | wtap |
| per2 | 1.5E-05 | 0.3623426 | 0.127 | 0.104 | 0.478778 | DF 3 | per2 |
| mllt10 | 1.6E-05 | 0.3551757 | 0.102 | 0.081 | 0.514984 | DF 3 | mllt10 |
| coq10b | 1.6E-05 | 0.3015944 | 0.195 | 0.167 | 0.522483 | DF 3 | coq10b |
| mnta | 2E-05 | 0.3569465 | 0.123 | 0.101 | 0.643546 | DF 3 | mnta |
| ctnnb21 | 2.3E-05 | 0.2244235 | 0.34 | 0.311 | 0.740679 | DF 3 | ctnnb2 |
| pcf11 | 2.3E-05 | 0.2020252 | 0.283 | 0.251 | 0.745199 | DF 3 | pcf11 |
| plekhj11 | 2.3E-05 | 0.2514811 | 0.145 | 0.121 | 0.750913 | DF 3 | plekhj1 |
| rpz2 | 2.4E-05 | 0.3191754 | 0.204 | 0.178 | 0.769517 | DF 3 | rpz2 |
| dusp8a | 2.5E-05 | 0.2771389 | 0.111 | 0.089 | 0.820083 | DF 3 | dusp8a |
| pitpnc1b | 3E-05 | 0.2548362 | 0.102 | 0.081 | 0.981713 | DF 3 | pitpnc1b |

| | | | | | | | |
|---|---|---|---|---|---|---|---|
| taok1b1 | 3.3E-05 | 0.2076968 | 0.172 | 0.146 | 1 | DF 3 | taok1b |
| acap11 | 3.3E-05 | 0.2557572 | 0.102 | 0.081 | 1 | DF 3 | acap1 |
| ENSDARG0000005970 | 3.6E-05 | 0.2181318 | 0.105 | 0.084 | 1 | DF 3 | ENSDARG00000059 |
| oaz1a | 4E-05 | 0.2005762 | 0.474 | 0.451 | 1 | DF 3 | oaz1a |
| arl6ip1 | 4.9E-05 | 0.2101209 | 0.186 | 0.16 | 1 | DF 3 | arl6ip1 |
| dyrk1b | 5.1E-05 | 0.3382574 | 0.113 | 0.092 | 1 | DF 3 | dyrk1b |
| pgm11 | 5.3E-05 | 0.3166846 | 0.12 | 0.099 | 1 | DF 3 | pgm1 |
| klf13 | 6.3E-05 | 0.2347575 | 0.15 | 0.126 | 1 | DF 3 | klf13 |
| zc3h4 | 6.5E-05 | 0.2626534 | 0.161 | 0.138 | 1 | DF 3 | zc3h4 |
| cep131 | 6.5E-05 | 0.2676729 | 0.117 | 0.096 | 1 | DF 3 | cep131 |
| atp1b2a | 7.3E-05 | 0.2935902 | 0.1 | 0.08 | 1 | DF 3 | atp1b2a |
| cicb1 | 7.6E-05 | 0.314952 | 0.149 | 0.126 | 1 | DF 3 | cicb |
| ccnl1b | 8.3E-05 | 0.3956005 | 0.102 | 0.083 | 1 | DF 3 | ccnl1b |
| crlf3 | 0.00011 | 0.205806 | 0.107 | 0.087 | 1 | DF 3 | crlf3 |
| rnf169 | 0.00011 | 0.3008107 | 0.151 | 0.128 | 1 | DF 3 | rnf169 |
| rad54l2 | 0.00011 | 0.2887601 | 0.101 | 0.082 | 1 | DF 3 | rad54l2 |
| foxo3b | 0.00012 | 0.2852453 | 0.1 | 0.081 | 1 | DF 3 | foxo3b |
| kifap3a1 | 0.00013 | 0.2315786 | 0.111 | 0.091 | 1 | DF 3 | kifap3a |
| kdm2ba | 0.00013 | 0.3104387 | 0.16 | 0.137 | 1 | DF 3 | kdm2ba |
| id2b1 | 0.00013 | 0.2612032 | 0.214 | 0.189 | 1 | DF 3 | id2b |
| nt5c3a | 0.00019 | 0.2020351 | 0.193 | 0.169 | 1 | DF 3 | nt5c3a |
| ano5b | 0.0002 | 0.3340346 | 0.105 | 0.086 | 1 | DF 3 | ano5b |
| irf2 | 0.00022 | 0.255438 | 0.171 | 0.148 | 1 | DF 3 | irf2 |
| park7 | 0.00024 | 0.2547127 | 0.172 | 0.15 | 1 | DF 3 | park7 |
| slc25a36a2 | 0.00028 | 0.2136868 | 0.125 | 0.105 | 1 | DF 3 | slc25a36a |
| klhl24b | 0.00033 | 0.261328 | 0.107 | 0.089 | 1 | DF 3 | klhl24b |
| oga1 | 0.0004 | 0.2336936 | 0.115 | 0.097 | 1 | DF 3 | oga |
| ppp1r13l | 0.00054 | 0.2119905 | 0.104 | 0.087 | 1 | DF 3 | ppp1r13l |
| glsb | 0.00059 | 0.242867 | 0.135 | 0.116 | 1 | DF 3 | glsb |
| cry2 | 0.00064 | 0.2308129 | 0.187 | 0.166 | 1 | DF 3 | cry2 |
| caska1 | 0.00071 | 0.2601235 | 0.103 | 0.086 | 1 | DF 3 | caska |
| kifbp | 0.00072 | 0.2851616 | 0.106 | 0.088 | 1 | DF 3 | kifbp |
| si:ch211-222k6.31 | 0.00091 | 0.2033531 | 0.129 | 0.11 | 1 | DF 3 | si:ch211-222k6.3 |
| ak11 | 0.00096 | 0.2355421 | 0.145 | 0.126 | 1 | DF 3 | ak1 |
| rgs3a1 | 0.00121 | 0.2799251 | 0.109 | 0.093 | 1 | DF 3 | rgs3a |
| clk2a1 | 0.00123 | 0.2631838 | 0.12 | 0.102 | 1 | DF 3 | clk2a |
| fam193a | 0.00128 | 0.2138284 | 0.12 | 0.103 | 1 | DF 3 | fam193a |
| rtca | 0.00162 | 0.2522077 | 0.207 | 0.187 | 1 | DF 3 | rtca |
| crema | 0.00201 | 0.2458266 | 0.125 | 0.108 | 1 | DF 3 | crema |
| brd1b | 0.00201 | 0.292123 | 0.114 | 0.098 | 1 | DF 3 | brd1b |
| ell | 0.0025 | 0.2006361 | 0.108 | 0.092 | 1 | DF 3 | ell |
| frmd6 | 0.00252 | 0.203388 | 0.208 | 0.189 | 1 | DF 3 | frmd6 |
| sptlc2b | 0.00396 | 0.2159566 | 0.134 | 0.119 | 1 | DF 3 | sptlc2b |
| yaf2 | 0.00424 | 0.2340302 | 0.124 | 0.109 | 1 | DF 3 | yaf2 |
| CU104709.1 | 0.0044 | 0.2275388 | 0.132 | 0.116 | 1 | DF 3 | CU104709.1 |
| rcan1a | 0.00558 | 0.2052508 | 0.247 | 0.23 | 1 | DF 3 | rcan1a |
| lysmd3 | 0.00611 | 0.2130138 | 0.119 | 0.105 | 1 | DF 3 | lysmd3 |
| slc25a55a | 0.00694 | 0.2885371 | 0.212 | 0.197 | 1 | DF 3 | slc25a55a |

| | | | | | | | |
|---|---|---|---|---|---|---|---|
| nfe2l1b | 0.00761 | 0.2066379 | 0.1 | 0.087 | 1 | DF 3 | nfe2l1b |
| rexo1 | 0.0078 | 0.309992 | 0.132 | 0.118 | 1 | DF 3 | rexo1 |
| egr2a1 | 0.00841 | 0.3238496 | 0.107 | 0.095 | 1 | DF 3 | egr2a |
| si:dkey-102m7.3 | 0.00947 | 0.2328992 | 0.134 | 0.12 | 1 | DF 3 | si:dkey-102m7.3 |
| col10a1a | 0 | 7.9462939 | 0.892 | 0.054 | 0 | Osteoblasts | col10a1a |
| bgna | 0 | 5.4389463 | 0.833 | 0.06 | 0 | Osteoblasts | bgna |
| ifitm5 | 0 | 8.257794 | 0.768 | 0.009 | 0 | Osteoblasts | ifitm5 |
| col5a2b | 0 | 7.3580213 | 0.762 | 0.013 | 0 | Osteoblasts | col5a2b |
| col11a2 | 0 | 3.0906592 | 0.76 | 0.058 | 0 | Osteoblasts | col11a2 |
| col10a1b | 0 | 9.3483682 | 0.714 | 0.03 | 0 | Osteoblasts | col10a1b |
| cd81b | 0 | 8.0214234 | 0.659 | 0.006 | 0 | Osteoblasts | cd81b |
| spp1 | 0 | 5.408385 | 0.637 | 0.026 | 0 | Osteoblasts | spp1 |
| loxl4 | 0 | 5.9820737 | 0.623 | 0.013 | 0 | Osteoblasts | loxl4 |
| itga10 | 0 | 2.6083677 | 0.816 | 0.208 | 0 | Osteoblasts | itga10 |
| bmp8a | 0 | 6.7926353 | 0.611 | 0.009 | 0 | Osteoblasts | bmp8a |
| fgfr31 | 0 | 2.7310444 | 0.819 | 0.225 | 0 | Osteoblasts | fgfr3 |
| bglap | 0 | 8.03703 | 0.608 | 0.019 | 0 | Osteoblasts | bglap |
| col22a1 | 0 | 5.3149475 | 0.605 | 0.017 | 0 | Osteoblasts | col22a1 |
| pcolceb1 | 0 | 3.3289283 | 0.703 | 0.13 | 0 | Osteoblasts | pcolceb |
| emid11 | 0 | 3.4934788 | 0.637 | 0.099 | 0 | Osteoblasts | emid1 |
| LOXL4 | 0 | 5.1328976 | 0.559 | 0.024 | 0 | Osteoblasts | LOXL4 |
| si:ch211-76l23.41 | 0 | 1.774271 | 0.666 | 0.148 | 0 | Osteoblasts | si:ch211-76l23.4 |
| cadm1b | 0 | 7.6295192 | 0.515 | 0.003 | 0 | Osteoblasts | cadm1b |
| tnfrsf11a1 | 0 | 2.449237 | 0.653 | 0.146 | 0 | Osteoblasts | tnfrsf11a |
| tpm11 | 0 | 1.6807474 | 0.798 | 0.299 | 0 | Osteoblasts | tpm1 |
| fpr1 | 0 | 2.2638959 | 0.669 | 0.181 | 0 | Osteoblasts | fpr1 |
| panx3 | 0 | 7.976516 | 0.487 | 0.003 | 0 | Osteoblasts | panx3 |
| kank1b1 | 0 | 3.6613855 | 0.527 | 0.053 | 0 | Osteoblasts | kank1b |
| mef2cb | 0 | 3.9419247 | 0.508 | 0.035 | 0 | Osteoblasts | mef2cb |
| aqp3b1 | 0 | 3.9144463 | 0.518 | 0.046 | 0 | Osteoblasts | aqp3b |
| mef2aa | 0 | 2.3901977 | 0.653 | 0.187 | 0 | Osteoblasts | mef2aa |
| CABZ01080702.11 | 0 | 2.0114116 | 0.723 | 0.267 | 0 | Osteoblasts | CABZ01080702.1 |
| marcksl1b1 | 0 | 2.3217804 | 0.669 | 0.221 | 0 | Osteoblasts | marcksl1b |
| bgnb | 0 | 2.4122251 | 0.539 | 0.106 | 0 | Osteoblasts | bgnb |
| phex | 0 | 3.3404531 | 0.471 | 0.046 | 0 | Osteoblasts | phex |
| phospho1 | 0 | 5.4370791 | 0.429 | 0.009 | 0 | Osteoblasts | phospho1 |
| rpz52 | 0 | 0.9626993 | 0.684 | 0.277 | 0 | Osteoblasts | rpz5 |
| efhd1 | 0 | 3.6916573 | 0.442 | 0.035 | 0 | Osteoblasts | efhd1 |
| bambib1 | 0 | 3.7380681 | 0.438 | 0.037 | 0 | Osteoblasts | bambib |
| col11a1a | 0 | 0.6739355 | 0.688 | 0.292 | 0 | Osteoblasts | col11a1a |
| tent5ba | 0 | 1.9797172 | 0.507 | 0.114 | 0 | Osteoblasts | tent5ba |
| rgcc | 0 | 1.6607482 | 0.559 | 0.17 | 0 | Osteoblasts | rgcc |
| bglapl | 0 | 8.6531569 | 0.421 | 0.032 | 0 | Osteoblasts | bglapl |
| sod3b1 | 0 | 2.3203283 | 0.476 | 0.095 | 0 | Osteoblasts | sod3b |
| tspan4a | 0 | 2.29912 | 0.477 | 0.11 | 0 | Osteoblasts | tspan4a |
| sgms2a | 0 | 6.6017471 | 0.355 | 0.004 | 0 | Osteoblasts | sgms2a |
| mibp2 | 0 | 2.642133 | 0.936 | 0.586 | 0 | Osteoblasts | mibp2 |
| mfsd2b | 0 | 7.7543712 | 0.343 | 0.002 | 0 | Osteoblasts | mfsd2b |

| | | | | | | | |
|---|---|---|---|---|---|---|---|
| p4ha1b1 | 0 | 1.8153742 | 0.456 | 0.115 | 0 | Osteoblasts | p4ha1b |
| ccdc187 | 0 | 2.0711711 | 0.464 | 0.123 | 0 | Osteoblasts | ccdc187 |
| ptk2ab | 0 | 3.6572193 | 0.366 | 0.032 | 0 | Osteoblasts | ptk2ab |
| sp7 | 0 | 4.2903737 | 0.343 | 0.018 | 0 | Osteoblasts | sp7 |
| znf385a | 0 | 3.2178792 | 0.351 | 0.027 | 0 | Osteoblasts | znf385a |
| spns2 | 0 | 4.6121935 | 0.336 | 0.013 | 0 | Osteoblasts | spns2 |
| si:dkey-12l12.11 | 0 | 1.6752101 | 0.879 | 0.556 | 0 | Osteoblasts | si:dkey-12l12.1 |
| nupr1b | 0 | 1.7728393 | 0.452 | 0.14 | 0 | Osteoblasts | nupr1b |
| slc35c22 | 0 | 1.5443606 | 0.44 | 0.128 | 0 | Osteoblasts | slc35c2 |
| phlda31 | 0 | 1.5847634 | 0.466 | 0.157 | 0 | Osteoblasts | phlda3 |
| tox | 0 | 2.9828968 | 0.344 | 0.036 | 0 | Osteoblasts | tox |
| tnfsf121 | 0 | 1.7631232 | 0.43 | 0.123 | 0 | Osteoblasts | tnfsf12 |
| ghra1 | 0 | 1.2547055 | 0.509 | 0.205 | 0 | Osteoblasts | ghra |
| cthrc1a | 0 | 1.8576893 | 0.434 | 0.131 | 0 | Osteoblasts | cthrc1a |
| dap1b.11 | 0 | 1.4437889 | 0.787 | 0.485 | 0 | Osteoblasts | dap1b.1 |
| rasgef1ba1 | 0 | 1.5713922 | 0.438 | 0.138 | 0 | Osteoblasts | rasgef1ba |
| bambia1 | 0 | 1.468452 | 0.425 | 0.126 | 0 | Osteoblasts | bambia |
| thsd7aa | 0 | 5.0078454 | 0.305 | 0.008 | 0 | Osteoblasts | thsd7aa |
| cx431 | 0 | 1.3415486 | 0.444 | 0.148 | 0 | Osteoblasts | cx43 |
| sept15 | 0 | 1.1956435 | 0.478 | 0.182 | 0 | Osteoblasts | sept15 |
| tgfb51 | 0 | 2.0188083 | 0.383 | 0.092 | 0 | Osteoblasts | tgfb5 |
| entpd5a1 | 0 | 3.5329709 | 0.32 | 0.031 | 0 | Osteoblasts | entpd5a |
| zgc:92162 | 0 | 5.1247475 | 0.297 | 0.01 | 0 | Osteoblasts | zgc:92162 |
| jcada1 | 0 | 1.7140653 | 0.385 | 0.1 | 0 | Osteoblasts | jcada |
| marcksb | 0 | 1.916428 | 0.358 | 0.08 | 0 | Osteoblasts | marcksb |
| fgfr41 | 0 | 2.3248108 | 0.333 | 0.056 | 0 | Osteoblasts | fgfr4 |
| si:dkey-32e6.6 | 0 | 4.5685129 | 0.289 | 0.013 | 0 | Osteoblasts | si:dkey-32e6.6 |
| si:ch73-217n20.1 | 0 | 2.9520053 | 0.306 | 0.032 | 0 | Osteoblasts | si:ch73-217n20.1 |
| scpp1 | 0 | 6.1453602 | 0.278 | 0.005 | 0 | Osteoblasts | scpp1 |
| dlx2b | 0 | 6.0548557 | 0.275 | 0.003 | 0 | Osteoblasts | dlx2b |
| ccn4b | 0 | 3.0936585 | 0.303 | 0.031 | 0 | Osteoblasts | ccn4b |
| cntfr | 0 | 3.0667092 | 0.306 | 0.035 | 0 | Osteoblasts | cntfr |
| furina | 0 | 1.3976473 | 0.404 | 0.136 | 0 | Osteoblasts | furina |
| pkig | 0 | 1.5327903 | 0.363 | 0.098 | 0 | Osteoblasts | pkig |
| cgref1 | 0 | 1.6197151 | 0.417 | 0.153 | 0 | Osteoblasts | cgref1 |
| eva1ba | 0 | 1.9036028 | 0.351 | 0.088 | 0 | Osteoblasts | eva1ba |
| fkbp11 | 0 | 1.488534 | 0.402 | 0.14 | 0 | Osteoblasts | fkbp11 |
| prdx5 | 0 | 1.0690886 | 0.833 | 0.571 | 0 | Osteoblasts | prdx5 |
| limch1a1 | 0 | 2.1726653 | 0.319 | 0.059 | 0 | Osteoblasts | limch1a |
| kalrnb | 0 | 1.9514432 | 0.33 | 0.073 | 0 | Osteoblasts | kalrnb |
| abcc6a | 0 | 2.8323893 | 0.313 | 0.057 | 0 | Osteoblasts | abcc6a |
| farp21 | 0 | 1.6351455 | 0.351 | 0.096 | 0 | Osteoblasts | farp2 |
| bmp4 | 0 | 2.1264329 | 0.297 | 0.047 | 0 | Osteoblasts | bmp4 |
| cfl2 | 0 | 2.8606421 | 0.287 | 0.038 | 0 | Osteoblasts | cfl2 |
| eva1bb | 0 | 3.5247623 | 0.266 | 0.022 | 0 | Osteoblasts | eva1bb |
| bmp2b1 | 0 | 1.7513053 | 0.328 | 0.084 | 0 | Osteoblasts | bmp2b |
| bmp2a | 0 | 3.5305724 | 0.249 | 0.018 | 0 | Osteoblasts | bmp2a |
| slc8a4b | 0 | 6.4271831 | 0.23 | 0.003 | 0 | Osteoblasts | slc8a4b |

| | | | | | | | |
|---|---|---|---|---|---|---|---|
| fhl3b1 | 0 | 1.4123535 | 0.322 | 0.095 | 0 | Osteoblasts | fhl3b |
| mob2b | 0 | 1.8069997 | 0.287 | 0.064 | 0 | Osteoblasts | mob2b |
| spon1a | 0 | 3.3271481 | 0.237 | 0.017 | 0 | Osteoblasts | spon1a |
| bmp3 | 0 | 5.730953 | 0.217 | 0.004 | 0 | Osteoblasts | bmp3 |
| fat3a1 | 0 | 2.6545027 | 0.246 | 0.034 | 0 | Osteoblasts | fat3a |
| bmper | 0 | 4.7912998 | 0.217 | 0.006 | 0 | Osteoblasts | bmper |
| swap70b | 0 | 2.205049 | 0.25 | 0.043 | 0 | Osteoblasts | swap70b |
| rpl22l11 | 0 | 0.9336658 | 0.936 | 0.733 | 0 | Osteoblasts | rpl22l1 |
| impdh1b1 | 0 | 1.8396103 | 0.27 | 0.068 | 0 | Osteoblasts | impdh1b |
| adcy2b | 0 | 4.4188915 | 0.204 | 0.007 | 0 | Osteoblasts | adcy2b |
| rtn4rl2a | 0 | 3.6303122 | 0.207 | 0.014 | 0 | Osteoblasts | rtn4rl2a |
| satb2 | 0 | 4.0877493 | 0.201 | 0.009 | 0 | Osteoblasts | satb2 |
| klf2b | 0 | 1.7282439 | 0.232 | 0.047 | 0 | Osteoblasts | klf2b |
| rnf145a | 0 | 2.537411 | 0.214 | 0.029 | 0 | Osteoblasts | rnf145a |
| glsl | 0 | 3.8717105 | 0.19 | 0.011 | 0 | Osteoblasts | glsl |
| csk | 0 | 1.6145081 | 0.225 | 0.051 | 0 | Osteoblasts | csk |
| tnika | 0 | 3.099854 | 0.189 | 0.018 | 0 | Osteoblasts | tnika |
| map4l | 0 | 3.3016612 | 0.186 | 0.016 | 0 | Osteoblasts | map4l |
| mafa1 | 0 | 1.9339953 | 0.194 | 0.037 | 0 | Osteoblasts | mafa |
| mef2ca | 0 | 2.7667656 | 0.18 | 0.024 | 0 | Osteoblasts | mef2ca |
| krt18a.11 | 0 | 0.8558164 | 0.985 | 0.83 | 0 | Osteoblasts | krt18a.1 |
| b4galnt3b | 0 | 4.7183686 | 0.157 | 0.004 | 0 | Osteoblasts | b4galnt3b |
| fzd8a | 0 | 2.7580045 | 0.174 | 0.021 | 0 | Osteoblasts | fzd8a |
| adamts18 | 0 | 6.9887191 | 0.152 | 0.001 | 0 | Osteoblasts | adamts18 |
| VIT | 0 | 2.9861602 | 0.164 | 0.016 | 0 | Osteoblasts | VIT |
| si:ch73-288o11.5 | 0 | 3.0761561 | 0.157 | 0.014 | 0 | Osteoblasts | si:ch73-288o11.5 |
| zbtb7c | 0 | 4.1358368 | 0.148 | 0.006 | 0 | Osteoblasts | zbtb7c |
| chst3b | 0 | 7.9810353 | 0.14 | 0 | 0 | Osteoblasts | chst3b |
| adgrv1 | 0 | 2.4281321 | 0.16 | 0.022 | 0 | Osteoblasts | adgrv1 |
| arhgap27l | 0 | 3.0362068 | 0.148 | 0.013 | 0 | Osteoblasts | arhgap27l |
| cpz | 0 | 2.5119306 | 0.154 | 0.021 | 0 | Osteoblasts | cpz |
| fyna | 0 | 3.193438 | 0.145 | 0.012 | 0 | Osteoblasts | fyna |
| frmd4bb | 0 | 3.0428506 | 0.145 | 0.013 | 0 | Osteoblasts | frmd4bb |
| tmem119a | 0 | 5.9905488 | 0.13 | 0.001 | 0 | Osteoblasts | tmem119a |
| pltp | 0 | 4.1277949 | 0.132 | 0.007 | 0 | Osteoblasts | pltp |
| ninj2 | 0 | 2.2461756 | 0.137 | 0.015 | 0 | Osteoblasts | ninj2 |
| sh3d21 | 0 | 1.9418254 | 0.138 | 0.018 | 0 | Osteoblasts | sh3d21 |
| rgs2 | 0 | 4.8873505 | 0.121 | 0.003 | 0 | Osteoblasts | rgs2 |
| sdk2b | 0 | 3.5472155 | 0.121 | 0.008 | 0 | Osteoblasts | sdk2b |
| zgc:110699 | 0 | 4.5110441 | 0.106 | 0.003 | 0 | Osteoblasts | zgc:110699 |
| sc:d189 | 0 | 3.3061614 | 0.109 | 0.008 | 0 | Osteoblasts | sc:d189 |
| si:dkey-35i13.1 | 0 | 4.0420101 | 0.102 | 0.004 | 0 | Osteoblasts | si:dkey-35i13.1 |
| fam167ab | 0 | 3.1084 | 0.101 | 0.009 | 0 | Osteoblasts | fam167ab |
| col1a1a1 | 0 | 1.6648881 | 0.997 | 0.979 | 0 | Osteoblasts | col1a1a |
| rpsa1 | 0 | 0.6188204 | 0.999 | 0.984 | 0 | Osteoblasts | rpsa |
| sparc1 | 0 | 1.885187 | 0.994 | 0.979 | 0 | Osteoblasts | sparc |
| rps121 | 0 | 0.5983304 | 0.992 | 0.977 | 0 | Osteoblasts | rps12 |
| rps201 | 0 | 0.5686836 | 0.995 | 0.985 | 0 | Osteoblasts | rps20 |

| | | | | | | | |
|---|---|---|---|---|---|---|---|
| col1a1b1 | 0 | 1.3610392 | 0.996 | 0.986 | 0 | Osteoblasts | col1a1b |
| col1a21 | 0 | 1.4368129 | 0.998 | 0.991 | 0 | Osteoblasts | col1a2 |
| eef1a1l11 | 0 | 0.4595889 | 0.999 | 0.996 | 0 | Osteoblasts | eef1a1l1 |
| rplp11 | 0 | 0.425079 | 1 | 0.999 | 0 | Osteoblasts | rplp1 |
| pabpc4 | 2E-307 | 2.5037982 | 0.124 | 0.016 | 5E-303 | Osteoblasts | pabpc4 |
| ptprfb | 7E-307 | 1.2005217 | 0.545 | 0.231 | 2.2E-302 | Osteoblasts | ptprfb |
| eef1g1 | 8E-307 | 0.609732 | 0.99 | 0.949 | 2.7E-302 | Osteoblasts | eef1g |
| cdh112 | 6E-305 | 0.7969944 | 0.873 | 0.567 | 2.1E-300 | Osteoblasts | cdh11 |
| lmo7a1 | 2E-302 | 1.2401578 | 0.393 | 0.134 | 7.8E-298 | Osteoblasts | lmo7a |
| pth1r | 3E-301 | 2.6994226 | 0.119 | 0.016 | 1.1E-296 | Osteoblasts | pth1r |
| fkbp1b | 3E-298 | 1.6189721 | 0.243 | 0.061 | 9.1E-294 | Osteoblasts | fkbp1b |
| si:ch211-253b8.5 | 6E-296 | 2.7327898 | 0.102 | 0.011 | 1.9E-291 | Osteoblasts | si:ch211-253b8.5 |
| rplp2l1 | 2E-295 | 0.4687346 | 0.998 | 0.986 | 5.3E-291 | Osteoblasts | rplp2l |
| ppfibp2a | 2E-290 | 2.1758958 | 0.159 | 0.029 | 5.8E-286 | Osteoblasts | ppfibp2a |
| rps26l1 | 6E-290 | 0.4869418 | 0.995 | 0.973 | 1.8E-285 | Osteoblasts | rps26l |
| creb3l11 | 6E-289 | 1.4575122 | 0.315 | 0.097 | 1.8E-284 | Osteoblasts | creb3l1 |
| cox6b1 | 7E-288 | 1.7348586 | 0.231 | 0.057 | 2.1E-283 | Osteoblasts | cox6b1 |
| tpm4b1 | 2E-287 | 1.0648244 | 0.261 | 0.07 | 5.2E-283 | Osteoblasts | tpm4b |
| xirp2a | 2E-284 | 1.0325211 | 0.541 | 0.235 | 6.7E-280 | Osteoblasts | xirp2a |
| mmp13b | 6E-284 | 3.7766742 | 0.14 | 0.024 | 2E-279 | Osteoblasts | mmp13b |
| lims2 | 4E-281 | 2.3913065 | 0.106 | 0.013 | 1.2E-276 | Osteoblasts | lims2 |
| eif4ebp3l1 | 8E-278 | 1.2138878 | 0.383 | 0.136 | 2.8E-273 | Osteoblasts | eif4ebp3l |
| dhrs3b1 | 2E-277 | 1.1562665 | 0.32 | 0.1 | 5.8E-273 | Osteoblasts | dhrs3b |
| rps4x1 | 2E-277 | 0.5069589 | 0.994 | 0.965 | 6.8E-273 | Osteoblasts | rps4x |
| shroom2a | 3E-276 | 1.5431067 | 0.287 | 0.086 | 1.1E-271 | Osteoblasts | shroom2a |
| si:ch73-22o12.1 | 9E-276 | 1.898382 | 0.147 | 0.025 | 2.9E-271 | Osteoblasts | si:ch73-22o12.1 |
| loxa1 | 3E-275 | 1.309716 | 0.41 | 0.155 | 9.7E-271 | Osteoblasts | loxa |
| rpl151 | 6E-274 | 0.5162702 | 0.997 | 0.969 | 2.1E-269 | Osteoblasts | rpl15 |
| mknk2a2 | 9E-269 | 1.2222613 | 0.491 | 0.208 | 2.8E-264 | Osteoblasts | mknk2a |
| serpinh1a1 | 4E-268 | 1.2133496 | 0.496 | 0.218 | 1.4E-263 | Osteoblasts | serpinh1a |
| rps251 | 3E-267 | 0.3908269 | 0.999 | 0.994 | 1.1E-262 | Osteoblasts | rps25 |
| kremen1 | 4E-261 | 1.5864198 | 0.244 | 0.067 | 1.5E-256 | Osteoblasts | kremen1 |
| rpl101 | 3E-257 | 0.5047656 | 0.99 | 0.967 | 9.9E-253 | Osteoblasts | rpl10 |
| rps241 | 1E-256 | 0.3913006 | 0.999 | 0.996 | 3.2E-252 | Osteoblasts | rps24 |
| rpl391 | 2E-256 | 0.4531568 | 0.996 | 0.983 | 6.9E-252 | Osteoblasts | rpl39 |
| nuak1b3 | 3E-256 | 1.1659614 | 0.55 | 0.261 | 9.1E-252 | Osteoblasts | nuak1b |
| rps71 | 9E-255 | 0.4203406 | 0.998 | 0.991 | 2.9E-250 | Osteoblasts | rps7 |
| sema5a | 2E-250 | 1.1554701 | 0.247 | 0.068 | 7.9E-246 | Osteoblasts | sema5a |
| endouc | 4E-249 | 1.2046263 | 0.482 | 0.217 | 1.1E-244 | Osteoblasts | endouc |
| vasnb1 | 1E-245 | 1.22934 | 0.353 | 0.127 | 4.6E-241 | Osteoblasts | vasnb |
| rplp01 | 2E-245 | 0.4745204 | 0.996 | 0.981 | 6.4E-241 | Osteoblasts | rplp0 |
| si:dkey-151g10.61 | 1E-243 | 0.3866924 | 1 | 0.999 | 3.2E-239 | Osteoblasts | si:dkey-151g10.6 |
| nme2b.11 | 2E-243 | 0.5131083 | 0.983 | 0.913 | 7.8E-239 | Osteoblasts | nme2b.1 |
| CR855311.11 | 7E-242 | 2.0043706 | 0.354 | 0.138 | 2.4E-237 | Osteoblasts | CR855311.1 |
| anks1b | 1E-241 | 1.6816653 | 0.197 | 0.049 | 4.2E-237 | Osteoblasts | anks1b |
| bcl2l101 | 2E-238 | 1.1928462 | 0.403 | 0.162 | 7.7E-234 | Osteoblasts | bcl2l10 |
| tcea3 | 5E-233 | 2.0577251 | 0.118 | 0.02 | 1.6E-228 | Osteoblasts | tcea3 |
| sost | 9E-232 | 1.2281129 | 0.231 | 0.066 | 2.8E-227 | Osteoblasts | sost |

| | | | | | | | |
|---|---|---|---|---|---|---|---|
| rpl181 | 3E-231 | 0.4481187 | 0.996 | 0.981 | 9.6E-227 | Osteoblasts | rpl18 |
| rps15a1 | 3E-230 | 0.3873281 | 0.998 | 0.989 | 1.1E-225 | Osteoblasts | rps15a |
| rpl111 | 2E-228 | 0.3958436 | 0.997 | 0.988 | 6.7E-224 | Osteoblasts | rpl11 |
| rps27.11 | 1E-225 | 0.415394 | 0.995 | 0.976 | 3.9E-221 | Osteoblasts | rps27.1 |
| ssr31 | 2E-225 | 0.7531949 | 0.819 | 0.559 | 7E-221 | Osteoblasts | ssr3 |
| rps3a1 | 6E-224 | 0.4519377 | 0.996 | 0.98 | 2E-219 | Osteoblasts | rps3a |
| rps27a1 | 2E-223 | 0.3874627 | 0.998 | 0.992 | 5.9E-219 | Osteoblasts | rps27a |
| ssr21 | 4E-223 | 0.669572 | 0.883 | 0.676 | 1.2E-218 | Osteoblasts | ssr2 |
| cox17 | 1E-221 | 1.0980463 | 0.355 | 0.135 | 3.9E-217 | Osteoblasts | cox17 |
| rftn2 | 3E-221 | 2.1525669 | 0.114 | 0.019 | 8.6E-217 | Osteoblasts | rftn2 |
| fam20a | 1E-220 | 2.472605 | 0.105 | 0.016 | 3.6E-216 | Osteoblasts | fam20a |
| dap | 3E-220 | 0.9899041 | 0.425 | 0.177 | 9.7E-216 | Osteoblasts | dap |
| rps231 | 4E-219 | 0.4019699 | 0.994 | 0.981 | 1.5E-214 | Osteoblasts | rps23 |
| crip3 | 1E-216 | 2.0291187 | 0.117 | 0.021 | 3.1E-212 | Osteoblasts | crip3 |
| rpl5a1 | 2E-216 | 0.4855435 | 0.98 | 0.914 | 5.6E-212 | Osteoblasts | rpl5a |
| sec61a11 | 2E-216 | 0.8836462 | 0.678 | 0.4 | 6.3E-212 | Osteoblasts | sec61a1 |
| fhl1b | 7E-216 | 1.2349569 | 0.257 | 0.082 | 2.2E-211 | Osteoblasts | fhl1b |
| intu | 7E-216 | 1.9941033 | 0.133 | 0.026 | 2.3E-211 | Osteoblasts | intu |
| rps22 | 7E-216 | 0.4081346 | 0.999 | 0.99 | 2.3E-211 | Osteoblasts | rps2 |
| rps8a1 | 2E-215 | 0.3881366 | 0.999 | 0.991 | 4.9E-211 | Osteoblasts | rps8a |
| sept6 | 2E-215 | 1.0793369 | 0.347 | 0.131 | 7.8E-211 | Osteoblasts | sept6 |
| chst11 | 3E-215 | 1.7726464 | 0.154 | 0.035 | 8.2E-211 | Osteoblasts | chst11 |
| serpine2 | 3E-215 | 1.1511482 | 0.58 | 0.295 | 1.1E-210 | Osteoblasts | serpine2 |
| rpl321 | 4E-215 | 0.3997175 | 0.997 | 0.988 | 1.4E-210 | Osteoblasts | rpl32 |
| eef2b1 | 7E-215 | 0.484517 | 0.98 | 0.904 | 2.3E-210 | Osteoblasts | eef2b |
| baiap2a | 1E-214 | 1.6918868 | 0.148 | 0.032 | 3.9E-210 | Osteoblasts | baiap2a |
| rps51 | 9E-214 | 0.4319597 | 0.994 | 0.971 | 2.8E-209 | Osteoblasts | rps5 |
| rps181 | 3E-213 | 0.4642631 | 0.994 | 0.961 | 1E-208 | Osteoblasts | rps18 |
| rps141 | 2E-212 | 0.3781658 | 1 | 0.993 | 5.1E-208 | Osteoblasts | rps14 |
| faua1 | 9E-212 | 0.4198559 | 0.993 | 0.972 | 2.8E-207 | Osteoblasts | faua |
| ncam1a1 | 3E-211 | 1.3243033 | 0.212 | 0.061 | 9.4E-207 | Osteoblasts | ncam1a |
| rpl35a1 | 1E-209 | 0.37238 | 0.998 | 0.988 | 4.1E-205 | Osteoblasts | rpl35a |
| rpl71 | 6E-209 | 0.4563123 | 0.993 | 0.971 | 1.8E-204 | Osteoblasts | rpl7 |
| tcf7l2 | 5E-208 | 1.0408773 | 0.37 | 0.147 | 1.5E-203 | Osteoblasts | tcf7l2 |
| rpl41 | 2E-206 | 0.5076294 | 0.972 | 0.89 | 5.9E-202 | Osteoblasts | rpl4 |
| rcn1 | 3E-206 | 1.4990162 | 0.307 | 0.116 | 8.4E-202 | Osteoblasts | rcn1 |
| si:ch211-136a13.1 | 1E-204 | 1.2103299 | 0.238 | 0.074 | 4.4E-200 | Osteoblasts | si:ch211-136a13.1 |
| rpl81 | 5E-204 | 0.4121748 | 0.996 | 0.977 | 1.6E-199 | Osteoblasts | rpl8 |
| niban2a1 | 1E-202 | 1.5231492 | 0.169 | 0.042 | 3.7E-198 | Osteoblasts | niban2a |
| rps61 | 2E-202 | 0.4327354 | 0.992 | 0.958 | 6.8E-198 | Osteoblasts | rps6 |
| csrp1a1 | 5E-202 | 1.0064932 | 0.494 | 0.237 | 1.5E-197 | Osteoblasts | csrp1a |
| nupr1a1 | 1E-198 | 0.9358232 | 0.495 | 0.238 | 3.8E-194 | Osteoblasts | nupr1a |
| gmps1 | 2E-197 | 1.0823279 | 0.322 | 0.122 | 5E-193 | Osteoblasts | gmps |
| ddx211 | 3E-197 | 0.907379 | 0.598 | 0.323 | 8.3E-193 | Osteoblasts | ddx21 |
| rpl211 | 7E-197 | 0.3734786 | 0.997 | 0.984 | 2.3E-192 | Osteoblasts | rpl21 |
| rpl171 | 2E-196 | 0.3805291 | 0.999 | 0.988 | 5.7E-192 | Osteoblasts | rpl17 |
| fat1a1 | 2E-196 | 0.8906349 | 0.518 | 0.253 | 7.1E-192 | Osteoblasts | fat1a |
| slc20a2 | 7E-196 | 0.9232889 | 0.517 | 0.253 | 2.4E-191 | Osteoblasts | slc20a2 |

| | | | | | | | |
|---|---|---|---|---|---|---|---|
| timp2b | 1E-194 | 2.8562156 | 0.452 | 0.236 | 3.5E-190 | Osteoblasts | timp2b |
| slc23a2 | 2E-194 | 1.5824344 | 0.185 | 0.051 | 6.3E-190 | Osteoblasts | slc23a2 |
| rpl18a1 | 3E-194 | 0.3735377 | 0.998 | 0.983 | 1.1E-189 | Osteoblasts | rpl18a |
| rps101 | 8E-193 | 0.3435789 | 0.999 | 0.994 | 2.7E-188 | Osteoblasts | rps10 |
| smad6a1 | 2E-192 | 1.2595344 | 0.185 | 0.051 | 7.4E-188 | Osteoblasts | smad6a |
| rack11 | 1E-190 | 0.4278325 | 0.987 | 0.945 | 3.7E-186 | Osteoblasts | rack1 |
| dipk1c1 | 3E-190 | 0.9771052 | 0.284 | 0.102 | 1E-185 | Osteoblasts | dipk1c |
| enpp11 | 3E-189 | 1.2387414 | 0.386 | 0.171 | 1E-184 | Osteoblasts | enpp1 |
| rps91 | 8E-189 | 0.3715565 | 0.998 | 0.986 | 2.6E-184 | Osteoblasts | rps9 |
| wnt9a | 2E-187 | 1.7689919 | 0.115 | 0.023 | 6E-183 | Osteoblasts | wnt9a |
| idh21 | 1E-186 | 0.8447573 | 0.588 | 0.325 | 4.5E-182 | Osteoblasts | idh2 |
| cdc42ep1a | 1E-186 | 1.6028674 | 0.135 | 0.03 | 4.8E-182 | Osteoblasts | cdc42ep1a |
| cd276 | 9E-186 | 1.0480233 | 0.31 | 0.119 | 3E-181 | Osteoblasts | cd276 |
| afap1l2 | 1E-185 | 1.6317734 | 0.133 | 0.03 | 4E-181 | Osteoblasts | afap1l2 |
| fgfr1a | 2E-184 | 1.0221616 | 0.347 | 0.142 | 6.8E-180 | Osteoblasts | fgfr1a |
| rpl13a1 | 2E-183 | 0.3744434 | 0.998 | 0.984 | 7.7E-179 | Osteoblasts | rpl13a |
| usp6nl | 4E-183 | 1.6080485 | 0.136 | 0.031 | 1.3E-178 | Osteoblasts | usp6nl |
| tpt11 | 2E-182 | 0.4575371 | 0.977 | 0.923 | 5.7E-178 | Osteoblasts | tpt1 |
| rpl231 | 3E-181 | 0.348991 | 0.999 | 0.992 | 8.3E-177 | Osteoblasts | rpl23 |
| cited4a1 | 3E-180 | 1.0929851 | 0.279 | 0.102 | 1E-175 | Osteoblasts | cited4a |
| eef1b21 | 1E-179 | 0.4452491 | 0.98 | 0.925 | 4.6E-175 | Osteoblasts | eef1b2 |
| rpl10a1 | 5E-179 | 0.4126478 | 0.995 | 0.977 | 1.5E-174 | Osteoblasts | rpl10a |
| zgc:153675 | 1E-178 | 0.970833 | 0.483 | 0.245 | 3.3E-174 | Osteoblasts | zgc:153675 |
| lmf2a | 2E-177 | 1.5106333 | 0.138 | 0.033 | 5.7E-173 | Osteoblasts | lmf2a |
| frmd61 | 7E-177 | 0.9535126 | 0.393 | 0.173 | 2.3E-172 | Osteoblasts | frmd6 |
| mmp9 | 2E-176 | 2.0608445 | 0.2 | 0.064 | 6.8E-172 | Osteoblasts | mmp9 |
| rpl131 | 4E-176 | 0.342415 | 1 | 0.993 | 1.3E-171 | Osteoblasts | rpl13 |
| arhgap29b | 2E-175 | 1.3540566 | 0.172 | 0.048 | 5.6E-171 | Osteoblasts | arhgap29b |
| rpl191 | 3E-174 | 0.3933168 | 0.996 | 0.982 | 9.1E-170 | Osteoblasts | rpl19 |
| dixdc1a | 7E-174 | 1.3901446 | 0.166 | 0.046 | 2.2E-169 | Osteoblasts | dixdc1a |
| MYO1D | 1E-173 | 1.0339349 | 0.256 | 0.091 | 4.6E-169 | Osteoblasts | MYO1D |
| olfml2a | 7E-173 | 1.6008542 | 0.134 | 0.032 | 2.2E-168 | Osteoblasts | olfml2a |
| cndp2 | 8E-173 | 1.3271061 | 0.266 | 0.099 | 2.4E-168 | Osteoblasts | cndp2 |
| pno11 | 6E-172 | 0.7923014 | 0.655 | 0.388 | 1.9E-167 | Osteoblasts | pno1 |
| arvcfb1 | 7E-171 | 0.9828911 | 0.255 | 0.091 | 2.4E-166 | Osteoblasts | arvcfb |
| nedd91 | 5E-170 | 1.2704746 | 0.2 | 0.063 | 1.8E-165 | Osteoblasts | nedd9 |
| acvrl12 | 1E-169 | 0.8611152 | 0.469 | 0.229 | 3.2E-165 | Osteoblasts | acvrl1 |
| rpl23a1 | 1E-169 | 0.3869713 | 0.995 | 0.969 | 3.6E-165 | Osteoblasts | rpl23a |
| mfge8a1 | 2E-169 | 1.2824176 | 0.278 | 0.107 | 5.4E-165 | Osteoblasts | mfge8a |
| rpl7a1 | 2E-169 | 0.3549306 | 0.999 | 0.99 | 5.6E-165 | Osteoblasts | rpl7a |
| tbx2b1 | 4E-167 | 1.0057649 | 0.291 | 0.114 | 1.2E-162 | Osteoblasts | tbx2b |
| rpl141 | 5E-167 | 0.2957315 | 1 | 0.995 | 1.6E-162 | Osteoblasts | rpl14 |
| cdc42ep3 | 3E-166 | 1.028225 | 0.179 | 0.053 | 1.1E-161 | Osteoblasts | cdc42ep3 |
| fscn1a1 | 4E-166 | 0.9820697 | 0.369 | 0.163 | 1.4E-161 | Osteoblasts | fscn1a |
| rps31 | 1E-165 | 0.4045367 | 0.984 | 0.944 | 3.8E-161 | Osteoblasts | rps3 |
| rpl5b1 | 9E-164 | 0.4986957 | 0.951 | 0.842 | 2.9E-159 | Osteoblasts | rpl5b |
| rpl241 | 3E-163 | 0.3263894 | 1 | 0.993 | 9.4E-159 | Osteoblasts | rpl24 |
| chp1 | 2E-162 | 1.4604586 | 0.162 | 0.046 | 5.8E-158 | Osteoblasts | chp1 |

| | | | | | | | |
|---|---|---|---|---|---|---|---|
| copz2 | 3E-161 | 1.007072 | 0.376 | 0.173 | 9E-157 | Osteoblasts | copz2 |
| calm2a | 8E-161 | 0.7005012 | 0.562 | 0.303 | 2.7E-156 | Osteoblasts | calm2a |
| hdac9b | 2E-160 | 1.0731218 | 0.192 | 0.06 | 7.5E-156 | Osteoblasts | hdac9b |
| zgc:77158 | 7E-160 | 1.3842551 | 0.175 | 0.053 | 2.4E-155 | Osteoblasts | zgc:77158 |
| ahcy1 | 1E-159 | 0.8178907 | 0.377 | 0.17 | 4.5E-155 | Osteoblasts | ahcy |
| ssr41 | 4E-159 | 0.6470535 | 0.812 | 0.592 | 1.4E-154 | Osteoblasts | ssr4 |
| usta | 4E-158 | 1.6664232 | 0.115 | 0.026 | 1.3E-153 | Osteoblasts | usta |
| runx31 | 8E-158 | 0.7715993 | 0.496 | 0.254 | 2.6E-153 | Osteoblasts | runx3 |
| wipi1 | 7E-157 | 0.9189277 | 0.28 | 0.109 | 2.2E-152 | Osteoblasts | wipi1 |
| atp5mc3b | 2E-156 | 0.6147278 | 0.828 | 0.606 | 6.2E-152 | Osteoblasts | atp5mc3b |
| specc1 | 8E-155 | 1.3904351 | 0.127 | 0.031 | 2.6E-150 | Osteoblasts | specc1 |
| myca1 | 5E-154 | 0.9064751 | 0.332 | 0.144 | 1.7E-149 | Osteoblasts | myca |
| slc38a5b1 | 1E-151 | 0.8374473 | 0.579 | 0.335 | 3.4E-147 | Osteoblasts | slc38a5b |
| rps151 | 6E-148 | 0.3128712 | 0.998 | 0.988 | 1.9E-143 | Osteoblasts | rps15 |
| elmo11 | 2E-147 | 1.13957 | 0.229 | 0.084 | 5.4E-143 | Osteoblasts | elmo1 |
| rps131 | 2E-147 | 0.3079678 | 0.998 | 0.99 | 8.1E-143 | Osteoblasts | rps13 |
| tomm20b1 | 4E-147 | 0.772486 | 0.569 | 0.326 | 1.3E-142 | Osteoblasts | tomm20b |
| rpl301 | 3E-146 | 0.3284022 | 0.995 | 0.982 | 9.4E-142 | Osteoblasts | rpl30 |
| igfbp3 | 5E-144 | 1.1434381 | 0.25 | 0.099 | 1.5E-139 | Osteoblasts | igfbp3 |
| CR381618.11 | 6E-144 | 0.6471786 | 0.431 | 0.211 | 1.9E-139 | Osteoblasts | CR381618.1 |
| rpl121 | 2E-143 | 0.3100334 | 0.995 | 0.981 | 7.8E-139 | Osteoblasts | rpl12 |
| aif1l | 1E-142 | 1.1844579 | 0.211 | 0.076 | 3.9E-138 | Osteoblasts | aif1l |
| kctd12.21 | 2E-142 | 0.73237 | 0.422 | 0.211 | 4.9E-138 | Osteoblasts | kctd12.2 |
| crip2 | 1E-141 | 0.500995 | 0.198 | 0.068 | 4.8E-137 | Osteoblasts | crip2 |
| c1qtnf6b1 | 3E-141 | 1.1609997 | 0.195 | 0.067 | 9.2E-137 | Osteoblasts | c1qtnf6b |
| glg1b | 7E-141 | 1.430019 | 0.106 | 0.024 | 2.4E-136 | Osteoblasts | glg1b |
| sgip1a | 7E-139 | 1.1800446 | 0.15 | 0.045 | 2.3E-134 | Osteoblasts | sgip1a |
| evla | 2E-138 | 1.0626578 | 0.213 | 0.078 | 6.5E-134 | Osteoblasts | evla |
| fhod3b | 1E-137 | 1.2433379 | 0.124 | 0.033 | 4.6E-133 | Osteoblasts | fhod3b |
| rpl271 | 7E-137 | 0.3237641 | 0.996 | 0.977 | 2.2E-132 | Osteoblasts | rpl27 |
| fam162a | 9E-136 | 0.7894849 | 0.436 | 0.223 | 2.9E-131 | Osteoblasts | fam162a |
| col11a1b1 | 3E-135 | 0.6117627 | 0.755 | 0.527 | 8.5E-131 | Osteoblasts | col11a1b |
| plpp2b | 3E-135 | 0.9137913 | 0.276 | 0.115 | 9.6E-131 | Osteoblasts | plpp2b |
| gpc6a2 | 2E-134 | 0.8041882 | 0.387 | 0.188 | 6E-130 | Osteoblasts | gpc6a |
| ldlrap1a | 2E-133 | 1.3303372 | 0.177 | 0.06 | 7.8E-129 | Osteoblasts | ldlrap1a |
| adka | 5E-133 | 0.6297672 | 0.294 | 0.125 | 1.7E-128 | Osteoblasts | adka |
| aopep1 | 5E-133 | 1.2225066 | 0.189 | 0.067 | 1.7E-128 | Osteoblasts | aopep |
| ybx1 | 3E-131 | 0.2948589 | 0.991 | 0.968 | 8.4E-127 | Osteoblasts | ybx1 |
| serpinf11 | 8E-130 | 0.8549068 | 0.712 | 0.542 | 2.6E-125 | Osteoblasts | serpinf1 |
| pard6gb | 2E-129 | 1.3667236 | 0.143 | 0.043 | 6.7E-125 | Osteoblasts | pard6gb |
| sh3pxd2aa | 9E-129 | 0.8526269 | 0.297 | 0.131 | 3E-124 | Osteoblasts | sh3pxd2aa |
| rps161 | 2E-128 | 0.3008042 | 0.997 | 0.988 | 5.4E-124 | Osteoblasts | rps16 |
| rpl33 | 2E-128 | 0.3195075 | 0.995 | 0.976 | 5.8E-124 | Osteoblasts | rpl3 |
| tomm7 | 3E-128 | 0.7154124 | 0.375 | 0.181 | 1E-123 | Osteoblasts | tomm7 |
| rpl341 | 3E-126 | 0.3041049 | 0.997 | 0.984 | 8.8E-122 | Osteoblasts | rpl34 |
| srgap1a | 5E-126 | 1.3863304 | 0.112 | 0.029 | 1.5E-121 | Osteoblasts | srgap1a |
| cox5aa | 5E-126 | 0.6162485 | 0.498 | 0.269 | 1.6E-121 | Osteoblasts | cox5aa |
| slc39a131 | 5E-125 | 0.8431076 | 0.293 | 0.13 | 1.8E-120 | Osteoblasts | slc39a13 |

| | | | | | | | |
|---|---|---|---|---|---|---|---|
| rasl11b1 | 6E-125 | 0.6315483 | 0.402 | 0.202 | 2.1E-120 | Osteoblasts | rasl11b |
| rps191 | 9E-125 | 0.2646219 | 0.999 | 0.996 | 2.9E-120 | Osteoblasts | rps19 |
| sec61b | 1E-124 | 0.5011516 | 0.79 | 0.567 | 3.4E-120 | Osteoblasts | sec61b |
| CU929070.51 | 8E-124 | 0.7146067 | 0.433 | 0.225 | 2.5E-119 | Osteoblasts | CU929070.5 |
| ak12 | 9E-124 | 0.7246966 | 0.271 | 0.115 | 2.9E-119 | Osteoblasts | ak1 |
| rpl91 | 1E-123 | 0.288897 | 0.997 | 0.987 | 4.3E-119 | Osteoblasts | rpl9 |
| bmp6 | 2E-123 | 0.7485894 | 0.16 | 0.052 | 5.4E-119 | Osteoblasts | bmp6 |
| AL954322.2 | 2E-123 | 1.4336038 | 0.129 | 0.038 | 5.7E-119 | Osteoblasts | AL954322.2 |
| csrp21 | 1E-120 | 0.8516837 | 0.256 | 0.109 | 4.4E-116 | Osteoblasts | csrp2 |
| phldb1a | 3E-119 | 1.1916738 | 0.124 | 0.036 | 9.3E-115 | Osteoblasts | phldb1a |
| vat11 | 5E-119 | 0.5933279 | 0.525 | 0.299 | 1.6E-114 | Osteoblasts | vat1 |
| shc1 | 4E-118 | 1.0932058 | 0.154 | 0.051 | 1.4E-113 | Osteoblasts | shc1 |
| srprb | 3E-116 | 0.678606 | 0.348 | 0.168 | 9.1E-112 | Osteoblasts | srprb |
| fam120a1 | 1E-115 | 0.8647279 | 0.34 | 0.169 | 4.1E-111 | Osteoblasts | fam120a |
| kdelr2b | 3E-115 | 0.6322066 | 0.525 | 0.304 | 1.1E-110 | Osteoblasts | kdelr2b |
| si:dkey-56m19.51 | 5E-115 | 0.9314139 | 0.199 | 0.077 | 1.5E-110 | Osteoblasts | si:dkey-56m19.5 |
| git1 | 1E-114 | 0.9821379 | 0.167 | 0.058 | 3.1E-110 | Osteoblasts | git1 |
| pof1b | 3E-114 | 1.4190854 | 0.116 | 0.033 | 1E-109 | Osteoblasts | pof1b |
| gphna | 1E-112 | 0.9339106 | 0.146 | 0.048 | 4E-108 | Osteoblasts | gphna |
| p4hb | 2E-112 | 0.5991905 | 0.777 | 0.558 | 6.3E-108 | Osteoblasts | p4hb |
| pkp3b | 8E-112 | 1.4123044 | 0.113 | 0.033 | 2.7E-107 | Osteoblasts | pkp3b |
| runx2b | 7E-111 | 1.4193073 | 0.104 | 0.028 | 2.1E-106 | Osteoblasts | runx2b |
| phf21aa | 2E-110 | 1.4091885 | 0.103 | 0.028 | 5.1E-106 | Osteoblasts | phf21aa |
| tfpia1 | 9E-110 | 0.4373203 | 0.345 | 0.166 | 2.8E-105 | Osteoblasts | tfpia |
| rpl36a1 | 2E-109 | 0.2668051 | 0.998 | 0.994 | 7.9E-105 | Osteoblasts | rpl36a |
| ggcx1 | 3E-109 | 0.6492302 | 0.321 | 0.155 | 1.1E-104 | Osteoblasts | ggcx |
| musk | 9E-109 | 1.2807599 | 0.113 | 0.033 | 3E-104 | Osteoblasts | musk |
| pik3r3b1 | 1E-108 | 0.9219529 | 0.451 | 0.26 | 4.5E-104 | Osteoblasts | pik3r3b |
| chsy1 | 2E-108 | 0.9967318 | 0.163 | 0.058 | 6E-104 | Osteoblasts | chsy1 |
| atp5pd | 1E-107 | 0.5841847 | 0.581 | 0.352 | 3.1E-103 | Osteoblasts | atp5pd |
| selenom1 | 4E-107 | 0.7131197 | 0.326 | 0.16 | 1.4E-102 | Osteoblasts | selenom |
| si:ch211-217k17.7 | 3E-106 | 0.6261878 | 0.473 | 0.267 | 9.5E-102 | Osteoblasts | si:ch211-217k17.7 |
| p3h1 | 7E-106 | 0.6137887 | 0.384 | 0.2 | 2.2E-101 | Osteoblasts | p3h1 |
| her91 | 3E-105 | 0.2899878 | 0.404 | 0.215 | 1.1E-100 | Osteoblasts | her9 |
| serf21 | 1E-104 | 0.4087078 | 0.898 | 0.762 | 3.8E-100 | Osteoblasts | serf2 |
| hint1 | 4E-104 | 0.5358787 | 0.594 | 0.37 | 1.3E-99 | Osteoblasts | hint1 |
| lrrc171 | 7E-104 | 0.4738212 | 0.609 | 0.38 | 2.1E-99 | Osteoblasts | lrrc17 |
| serp11 | 2E-103 | 0.680197 | 0.504 | 0.301 | 5.9E-99 | Osteoblasts | serp1 |
| kdelr3 | 2E-103 | 0.8549742 | 0.332 | 0.171 | 6E-99 | Osteoblasts | kdelr3 |
| RPS171 | 4E-103 | 0.2662669 | 0.998 | 0.986 | 1.37E-98 | Osteoblasts | RPS17 |
| rpl281 | 4E-102 | 0.2435589 | 0.999 | 0.991 | 1.37E-97 | Osteoblasts | rpl28 |
| cbfb1 | 7E-102 | 0.8195917 | 0.252 | 0.114 | 2.24E-97 | Osteoblasts | cbfb |
| rpl351 | 5E-101 | 0.2597671 | 0.999 | 0.991 | 1.52E-96 | Osteoblasts | rpl35 |
| fut8a1 | 7E-101 | 0.7521462 | 0.172 | 0.065 | 2.12E-96 | Osteoblasts | fut8a |
| cct2.11 | 2E-100 | 0.5133612 | 0.639 | 0.407 | 5.38E-96 | Osteoblasts | cct2.1 |
| ppib1 | 2E-100 | 0.3114377 | 0.971 | 0.913 | 5.42E-96 | Osteoblasts | ppib |
| zgc:1949811 | 4E-100 | 0.8089666 | 0.186 | 0.073 | 1.38E-95 | Osteoblasts | zgc:194981 |
| tent5bb1 | 5E-100 | 0.5098824 | 0.282 | 0.132 | 1.7E-95 | Osteoblasts | tent5bb |

| | | | | | | | |
|---|---|---|---|---|---|---|---|
| gng10 | 8E-100 | 0.9440404 | 0.156 | 0.057 | 2.59E-95 | Osteoblasts | gng10 |
| rpl311 | 8E-100 | 0.2428734 | 0.998 | 0.993 | 2.76E-95 | Osteoblasts | rpl31 |
| tgfbr2b | 6E-99 | 0.4726183 | 0.283 | 0.133 | 2.11E-94 | Osteoblasts | tgfbr2b |
| slc30a7 | 8E-99 | 0.785672 | 0.233 | 0.103 | 2.6E-94 | Osteoblasts | slc30a7 |
| epha2b1 | 1.8E-98 | 1.071806 | 0.139 | 0.048 | 5.95E-94 | Osteoblasts | epha2b |
| ENSDARG000001152 | 8.8E-98 | 0.2505097 | 0.999 | 0.993 | 2.86E-93 | Osteoblasts | ENSDARG00000115 |
| dusp22 | 2E-96 | 0.7086422 | 0.38 | 0.21 | 6.46E-92 | Osteoblasts | dusp2 |
| dab2ipb | 8.3E-96 | 0.5889734 | 0.235 | 0.104 | 2.69E-91 | Osteoblasts | dab2ipb |
| zfp36l2 | 4.7E-95 | 0.6191716 | 0.446 | 0.254 | 1.53E-90 | Osteoblasts | zfp36l2 |
| uba521 | 5.1E-95 | 0.2287645 | 0.998 | 0.991 | 1.65E-90 | Osteoblasts | uba52 |
| nop10 | 6.1E-95 | 0.5617122 | 0.338 | 0.174 | 1.99E-90 | Osteoblasts | nop10 |
| tnfrsf9a | 5.6E-94 | 1.7830942 | 0.126 | 0.044 | 1.83E-89 | Osteoblasts | tnfrsf9a |
| snu13b | 8.3E-93 | 0.4650773 | 0.333 | 0.17 | 2.7E-88 | Osteoblasts | snu13b |
| cdo11 | 1.1E-92 | 0.5887911 | 0.506 | 0.315 | 3.74E-88 | Osteoblasts | cdo1 |
| sept7a1 | 1.2E-92 | 0.4783508 | 0.558 | 0.346 | 3.98E-88 | Osteoblasts | sept7a |
| gtpbp41 | 3.9E-92 | 0.5852297 | 0.6 | 0.396 | 1.26E-87 | Osteoblasts | gtpbp4 |
| phc2a | 6.7E-92 | 0.9750983 | 0.161 | 0.063 | 2.18E-87 | Osteoblasts | phc2a |
| prdx1 | 1.6E-91 | 1.0961842 | 0.156 | 0.061 | 5.09E-87 | Osteoblasts | prdx1 |
| npm1a | 1E-90 | 0.3158801 | 0.561 | 0.344 | 3.3E-86 | Osteoblasts | npm1a |
| btbd6a | 1.3E-90 | 0.8712976 | 0.177 | 0.072 | 4.16E-86 | Osteoblasts | btbd6a |
| ctnnb1 | 2.7E-90 | 0.5593081 | 0.701 | 0.49 | 8.91E-86 | Osteoblasts | ctnnb1 |
| inppl1a | 9.9E-90 | 0.790302 | 0.19 | 0.08 | 3.22E-85 | Osteoblasts | inppl1a |
| lman1 | 1.9E-89 | 0.7374939 | 0.245 | 0.116 | 6.2E-85 | Osteoblasts | lman1 |
| eif3ea1 | 2.4E-89 | 0.4745647 | 0.743 | 0.546 | 7.69E-85 | Osteoblasts | eif3ea |
| ankha | 5E-89 | 1.4870838 | 0.102 | 0.032 | 1.62E-84 | Osteoblasts | ankha |
| bccip | 1.2E-88 | 0.5344472 | 0.347 | 0.183 | 3.82E-84 | Osteoblasts | bccip |
| pdcd4a1 | 2.2E-88 | 0.6915445 | 0.191 | 0.08 | 7.09E-84 | Osteoblasts | pdcd4a |
| gars1 | 3.1E-88 | 0.6307493 | 0.281 | 0.139 | 1.01E-83 | Osteoblasts | gars1 |
| nop531 | 3.7E-88 | 0.6011556 | 0.639 | 0.444 | 1.21E-83 | Osteoblasts | nop53 |
| pcbp2 | 1.2E-87 | 0.4792469 | 0.39 | 0.212 | 3.81E-83 | Osteoblasts | pcbp2 |
| bicc1a | 4.7E-87 | 0.8597822 | 0.134 | 0.048 | 1.54E-82 | Osteoblasts | bicc1a |
| mpc1 | 7.3E-87 | 0.5883788 | 0.304 | 0.155 | 2.37E-82 | Osteoblasts | mpc1 |
| CU929259.1 | 1.2E-86 | 0.4546654 | 0.512 | 0.303 | 3.87E-82 | Osteoblasts | CU929259.1 |
| pfn2 | 2.6E-86 | 0.4438376 | 0.742 | 0.523 | 8.33E-82 | Osteoblasts | pfn2 |
| rsl24d12 | 1.3E-85 | 0.5309233 | 0.865 | 0.739 | 4.1E-81 | Osteoblasts | rsl24d1 |
| rsl1d1 | 1.4E-85 | 0.5357897 | 0.566 | 0.36 | 4.44E-81 | Osteoblasts | rsl1d1 |
| rcn31 | 3.5E-85 | 0.6803653 | 0.42 | 0.251 | 1.15E-80 | Osteoblasts | rcn3 |
| rock2b.11 | 5.4E-85 | 0.3788879 | 0.317 | 0.161 | 1.75E-80 | Osteoblasts | rock2b.1 |
| atp5f1c | 6.3E-85 | 0.5399966 | 0.425 | 0.243 | 2.06E-80 | Osteoblasts | atp5f1c |
| serbp1a1 | 2.3E-84 | 0.3633001 | 0.957 | 0.905 | 7.63E-80 | Osteoblasts | serbp1a |
| rps111 | 9.9E-84 | 0.2218066 | 0.998 | 0.988 | 3.23E-79 | Osteoblasts | rps11 |
| hif1al | 2.6E-82 | 0.7876108 | 0.221 | 0.104 | 8.32E-78 | Osteoblasts | hif1al |
| selenon1 | 6.1E-82 | 0.6630155 | 0.286 | 0.147 | 1.98E-77 | Osteoblasts | selenon |
| vdac21 | 8.1E-82 | 0.4523952 | 0.714 | 0.504 | 2.65E-77 | Osteoblasts | vdac2 |
| sdf4 | 1.1E-81 | 0.4491869 | 0.483 | 0.29 | 3.73E-77 | Osteoblasts | sdf4 |
| gpx8 | 2.4E-81 | 0.552574 | 0.312 | 0.164 | 7.84E-77 | Osteoblasts | gpx8 |
| nanos11 | 3.2E-81 | 0.3570196 | 0.358 | 0.193 | 1.04E-76 | Osteoblasts | nanos1 |
| nutf2l | 1.3E-80 | 0.7129783 | 0.205 | 0.094 | 4.17E-76 | Osteoblasts | nutf2l |

| | | | | | | | |
|---|---|---|---|---|---|---|---|
| rpl61 | 1.7E-80 | 0.308014 | 0.989 | 0.956 | 5.63E-76 | Osteoblasts | rpl6 |
| sec22bb | 2.2E-80 | 0.8901922 | 0.125 | 0.046 | 7.25E-76 | Osteoblasts | sec22bb |
| elof1 | 3.8E-80 | 0.5322809 | 0.27 | 0.135 | 1.25E-75 | Osteoblasts | elof1 |
| inppl1b | 1E-79 | 1.1691567 | 0.103 | 0.034 | 3.38E-75 | Osteoblasts | inppl1b |
| schip1 | 7E-79 | 0.5119365 | 0.271 | 0.136 | 2.29E-74 | Osteoblasts | schip1 |
| rpl361 | 7.3E-79 | 0.2421064 | 0.996 | 0.985 | 2.38E-74 | Osteoblasts | rpl36 |
| luzp11 | 2.4E-78 | 0.4627586 | 0.312 | 0.164 | 7.67E-74 | Osteoblasts | luzp1 |
| s1pr11 | 3.1E-78 | 0.9467713 | 0.188 | 0.085 | 1.01E-73 | Osteoblasts | s1pr1 |
| cox7b | 4.1E-78 | 0.449228 | 0.593 | 0.381 | 1.33E-73 | Osteoblasts | cox7b |
| slc25a51 | 5.1E-78 | 0.2476298 | 0.988 | 0.961 | 1.66E-73 | Osteoblasts | slc25a5 |
| mfge8b | 1.4E-77 | 0.8622273 | 0.133 | 0.051 | 4.53E-73 | Osteoblasts | mfge8b |
| tmem45a | 4.3E-77 | 0.4805021 | 0.288 | 0.148 | 1.38E-72 | Osteoblasts | tmem45a |
| tfe3a1 | 4.9E-77 | 0.7246213 | 0.193 | 0.087 | 1.59E-72 | Osteoblasts | tfe3a |
| snd1 | 1.1E-76 | 0.452912 | 0.4 | 0.228 | 3.45E-72 | Osteoblasts | snd1 |
| lrrc59 | 2.4E-76 | 0.6080173 | 0.26 | 0.131 | 7.69E-72 | Osteoblasts | lrrc59 |
| cox7a2a | 3.6E-76 | 0.4337928 | 0.437 | 0.257 | 1.18E-71 | Osteoblasts | cox7a2a |
| serinc51 | 7.7E-76 | 0.4187046 | 0.398 | 0.228 | 2.52E-71 | Osteoblasts | serinc5 |
| slc25a3b | 8.2E-76 | 0.4193316 | 0.74 | 0.536 | 2.66E-71 | Osteoblasts | slc25a3b |
| pld71 | 8.7E-76 | 0.5591761 | 0.276 | 0.143 | 2.84E-71 | Osteoblasts | pld7 |
| ajuba | 2.1E-75 | 0.7788603 | 0.166 | 0.071 | 6.69E-71 | Osteoblasts | ajuba |
| hdlbpa | 2.6E-75 | 0.3754395 | 0.835 | 0.663 | 8.39E-71 | Osteoblasts | hdlbpa |
| eif1ad | 4.6E-75 | 0.5096686 | 0.259 | 0.131 | 1.49E-70 | Osteoblasts | eif1ad |
| s1pr5a1 | 8.4E-75 | 0.8891557 | 0.139 | 0.056 | 2.73E-70 | Osteoblasts | s1pr5a |
| nfkbib | 1.6E-74 | 0.7014787 | 0.218 | 0.105 | 5.2E-70 | Osteoblasts | nfkbib |
| rps281 | 5.8E-74 | 0.2665339 | 0.994 | 0.977 | 1.87E-69 | Osteoblasts | rps28 |
| eif2s31 | 5.8E-74 | 0.4375521 | 0.678 | 0.481 | 1.89E-69 | Osteoblasts | eif2s3 |
| fam114a1 | 1.5E-73 | 0.7249502 | 0.213 | 0.102 | 5.02E-69 | Osteoblasts | fam114a1 |
| dnajc1 | 1.9E-73 | 0.3810529 | 0.454 | 0.27 | 6.15E-69 | Osteoblasts | dnajc1 |
| nsa21 | 8.1E-73 | 0.5127368 | 0.784 | 0.62 | 2.63E-68 | Osteoblasts | nsa2 |
| fam49a | 9.5E-73 | 0.6835911 | 0.161 | 0.069 | 3.1E-68 | Osteoblasts | fam49a |
| mgst3a | 9.8E-73 | 0.4251612 | 0.416 | 0.243 | 3.18E-68 | Osteoblasts | mgst3a |
| si:ch1073-325m22.2 | 1.3E-71 | 0.4158666 | 0.476 | 0.289 | 4.11E-67 | Osteoblasts | si:ch1073-325m22. |
| plekhg5a1 | 1.6E-71 | 0.6036564 | 0.156 | 0.066 | 5.06E-67 | Osteoblasts | plekhg5a |
| gtdc1 | 3.7E-71 | 0.9579617 | 0.107 | 0.038 | 1.22E-66 | Osteoblasts | gtdc1 |
| ksr1b | 1.7E-70 | 1.0863987 | 0.116 | 0.044 | 5.56E-66 | Osteoblasts | ksr1b |
| cd9b1 | 4.9E-70 | 0.4199011 | 0.196 | 0.093 | 1.6E-65 | Osteoblasts | cd9b |
| plekha6 | 5.7E-70 | 0.8531631 | 0.132 | 0.053 | 1.86E-65 | Osteoblasts | plekha6 |
| bysl | 5.7E-70 | 0.5563937 | 0.218 | 0.106 | 1.86E-65 | Osteoblasts | bysl |
| eif3d | 1.6E-69 | 0.4422085 | 0.605 | 0.407 | 5.25E-65 | Osteoblasts | eif3d |
| pkp4 | 2.4E-69 | 0.7787634 | 0.12 | 0.046 | 7.7E-65 | Osteoblasts | pkp4 |
| smad6b1 | 6E-69 | 0.5711764 | 0.201 | 0.096 | 1.97E-64 | Osteoblasts | smad6b |
| cct4 | 7.4E-69 | 0.4093236 | 0.482 | 0.3 | 2.41E-64 | Osteoblasts | cct4 |
| mgat1b1 | 8.4E-69 | 0.5415593 | 0.192 | 0.09 | 2.75E-64 | Osteoblasts | mgat1b |
| brf1a | 1.5E-68 | 0.8655434 | 0.118 | 0.046 | 4.78E-64 | Osteoblasts | brf1a |
| tanc2a1 | 2.2E-68 | 0.6755962 | 0.163 | 0.072 | 7.31E-64 | Osteoblasts | tanc2a |
| smim15 | 6.5E-68 | 0.5224857 | 0.295 | 0.16 | 2.1E-63 | Osteoblasts | smim15 |
| egr2a2 | 8E-68 | 0.6248203 | 0.187 | 0.088 | 2.59E-63 | Osteoblasts | egr2a |
| zgc:66433 | 8.1E-68 | 0.6748696 | 0.214 | 0.105 | 2.65E-63 | Osteoblasts | zgc:66433 |

| | | | | | | | |
|---|---|---|---|---|---|---|---|
| ppm1ba | 9.7E-68 | 0.6630273 | 0.16 | 0.07 | 3.15E-63 | Osteoblasts | ppm1ba |
| tmed31 | 1.1E-67 | 0.6741542 | 0.19 | 0.089 | 3.44E-63 | Osteoblasts | tmed3 |
| hbegfa1 | 2.3E-67 | 0.3920668 | 0.578 | 0.389 | 7.62E-63 | Osteoblasts | hbegfa |
| foxn3 | 3.7E-67 | 0.5589966 | 0.16 | 0.07 | 1.2E-62 | Osteoblasts | foxn3 |
| pycr1b | 4.2E-67 | 0.7806815 | 0.123 | 0.049 | 1.37E-62 | Osteoblasts | pycr1b |
| eif3s6ip1 | 6.3E-67 | 0.4257764 | 0.488 | 0.306 | 2.04E-62 | Osteoblasts | eif3s6ip |
| srp9 | 8.2E-67 | 0.5080475 | 0.207 | 0.101 | 2.66E-62 | Osteoblasts | srp9 |
| tram1 | 1.1E-66 | 0.3996482 | 0.448 | 0.275 | 3.44E-62 | Osteoblasts | tram1 |
| slc31a1 | 1.6E-66 | 0.4300775 | 0.333 | 0.187 | 5.07E-62 | Osteoblasts | slc31a1 |
| pik3r1 | 2.6E-66 | 0.440097 | 0.324 | 0.181 | 8.54E-62 | Osteoblasts | pik3r1 |
| aars1 | 5.1E-66 | 0.5234237 | 0.249 | 0.13 | 1.65E-61 | Osteoblasts | aars1 |
| pa2g4b | 6.6E-66 | 0.3831738 | 0.403 | 0.239 | 2.13E-61 | Osteoblasts | pa2g4b |
| kctd5a | 1.2E-65 | 0.3383611 | 0.493 | 0.306 | 3.86E-61 | Osteoblasts | kctd5a |
| c1qtnf6a1 | 2.3E-65 | 0.7586905 | 0.134 | 0.056 | 7.49E-61 | Osteoblasts | c1qtnf6a |
| CABZ01080375.1 | 4.4E-65 | 0.6687291 | 0.143 | 0.061 | 1.42E-60 | Osteoblasts | CABZ01080375.1 |
| furinb | 5.3E-65 | 0.5286134 | 0.125 | 0.05 | 1.74E-60 | Osteoblasts | furinb |
| tmem39a | 5.8E-65 | 0.5180548 | 0.265 | 0.141 | 1.88E-60 | Osteoblasts | tmem39a |
| hyal1 | 7.8E-65 | 0.7755212 | 0.125 | 0.051 | 2.53E-60 | Osteoblasts | hyal1 |
| mrpl36 | 9.3E-65 | 0.47412 | 0.249 | 0.13 | 3.03E-60 | Osteoblasts | mrpl36 |
| uqcrh | 1.9E-64 | 0.3309898 | 0.566 | 0.37 | 6.16E-60 | Osteoblasts | uqcrh |
| cox6a1 | 9.4E-64 | 0.3935543 | 0.678 | 0.477 | 3.05E-59 | Osteoblasts | cox6a1 |
| scxa1 | 1E-63 | 0.6798009 | 0.166 | 0.077 | 3.26E-59 | Osteoblasts | scxa |
| cnpy2 | 2E-63 | 0.9814319 | 0.106 | 0.04 | 6.36E-59 | Osteoblasts | cnpy2 |
| vstm4a | 2.2E-63 | 0.2561929 | 0.26 | 0.136 | 7.19E-59 | Osteoblasts | vstm4a |
| COX5B.1 | 2.7E-63 | 0.3200188 | 0.739 | 0.526 | 8.82E-59 | Osteoblasts | COX5B.1 |
| rassf2a | 3.5E-63 | 0.6484137 | 0.228 | 0.118 | 1.14E-58 | Osteoblasts | rassf2a |
| usp36 | 4.1E-63 | 0.4559005 | 0.332 | 0.191 | 1.33E-58 | Osteoblasts | usp36 |
| kank1a2 | 9.9E-63 | 0.5938893 | 0.209 | 0.105 | 3.22E-58 | Osteoblasts | kank1a |
| ola11 | 1.4E-62 | 0.3891679 | 0.688 | 0.492 | 4.66E-58 | Osteoblasts | ola1 |
| rpl221 | 3.8E-62 | 0.2219597 | 0.992 | 0.972 | 1.22E-57 | Osteoblasts | rpl22 |
| eif2s1b1 | 4.3E-62 | 0.3660369 | 0.568 | 0.373 | 1.39E-57 | Osteoblasts | eif2s1b |
| tdh1 | 7E-62 | 0.4446745 | 0.38 | 0.23 | 2.27E-57 | Osteoblasts | tdh |
| eif3f1 | 8.3E-62 | 0.3205722 | 0.854 | 0.719 | 2.7E-57 | Osteoblasts | eif3f |
| si:dkey-54n8.2 | 3.5E-61 | 0.6860932 | 0.122 | 0.05 | 1.13E-56 | Osteoblasts | si:dkey-54n8.2 |
| slc39a7 | 7E-61 | 0.5428137 | 0.215 | 0.11 | 2.27E-56 | Osteoblasts | slc39a7 |
| cox5ab | 8.1E-61 | 0.4564141 | 0.353 | 0.21 | 2.63E-56 | Osteoblasts | cox5ab |
| naca1 | 6.3E-60 | 0.2522838 | 0.984 | 0.959 | 2.05E-55 | Osteoblasts | naca |
| hmgb3b | 6.9E-60 | 0.5055852 | 0.214 | 0.11 | 2.24E-55 | Osteoblasts | hmgb3b |
| trmt112 | 7.9E-60 | 0.5047728 | 0.221 | 0.114 | 2.57E-55 | Osteoblasts | trmt112 |
| ikbip | 2.3E-59 | 0.4212291 | 0.218 | 0.112 | 7.53E-55 | Osteoblasts | ikbip |
| ndufab1b | 3.5E-59 | 0.3240224 | 0.336 | 0.193 | 1.13E-54 | Osteoblasts | ndufab1b |
| arhgap23a1 | 3.5E-59 | 0.6285151 | 0.17 | 0.081 | 1.15E-54 | Osteoblasts | arhgap23a |
| znf593 | 6.3E-59 | 0.4941458 | 0.199 | 0.1 | 2.04E-54 | Osteoblasts | znf593 |
| fkbp1aa | 8.5E-59 | 0.3393208 | 0.695 | 0.491 | 2.77E-54 | Osteoblasts | fkbp1aa |
| svep11 | 1.1E-58 | 0.4285148 | 0.253 | 0.136 | 3.63E-54 | Osteoblasts | svep1 |
| slc29a1a1 | 1.3E-58 | 0.4269957 | 0.324 | 0.188 | 4.37E-54 | Osteoblasts | slc29a1a |
| etv5a | 1.7E-58 | 0.6719419 | 0.193 | 0.098 | 5.51E-54 | Osteoblasts | etv5a |
| rps291 | 2.2E-58 | 0.2397782 | 0.994 | 0.968 | 7.15E-54 | Osteoblasts | rps29 |

| | | | | | | | |
|---|---|---|---|---|---|---|---|
| dab2ipa | 4E-58 | 0.4801897 | 0.207 | 0.106 | 1.31E-53 | Osteoblasts | dab2ipa |
| rfxap | 4.5E-58 | 0.4355185 | 0.226 | 0.118 | 1.48E-53 | Osteoblasts | rfxap |
| sav1 | 6.1E-58 | 0.361264 | 0.226 | 0.117 | 1.98E-53 | Osteoblasts | sav1 |
| atox11 | 1E-57 | 0.3293894 | 0.405 | 0.247 | 3.37E-53 | Osteoblasts | atox1 |
| fam13a | 1.5E-57 | 0.7949805 | 0.131 | 0.057 | 4.75E-53 | Osteoblasts | fam13a |
| eif5a1 | 2.6E-57 | 0.2971822 | 0.863 | 0.709 | 8.46E-53 | Osteoblasts | eif5a |
| btg3 | 3.3E-57 | 0.4032519 | 0.227 | 0.119 | 1.06E-52 | Osteoblasts | btg3 |
| ptprsa.11 | 4.4E-57 | 0.4086761 | 0.224 | 0.117 | 1.42E-52 | Osteoblasts | ptprsa.1 |
| cct7 | 4.5E-57 | 0.3737744 | 0.54 | 0.359 | 1.47E-52 | Osteoblasts | cct7 |
| si:ch211-160o17.2 | 9.1E-57 | 0.776437 | 0.121 | 0.052 | 2.96E-52 | Osteoblasts | si:ch211-160o17.2 |
| atf4a1 | 9.5E-57 | 0.3268297 | 0.453 | 0.286 | 3.09E-52 | Osteoblasts | atf4a |
| rpl381 | 1E-56 | 0.2799779 | 0.959 | 0.907 | 3.39E-52 | Osteoblasts | rpl38 |
| ndufs8a | 1.5E-56 | 0.6547418 | 0.175 | 0.086 | 4.88E-52 | Osteoblasts | ndufs8a |
| cd151 | 2.3E-56 | 0.7135283 | 0.184 | 0.093 | 7.49E-52 | Osteoblasts | cd151 |
| BX511129.2 | 3.2E-56 | 0.4029336 | 0.172 | 0.083 | 1.05E-51 | Osteoblasts | BX511129.2 |
| emc8 | 3.8E-56 | 0.6913379 | 0.115 | 0.048 | 1.23E-51 | Osteoblasts | emc8 |
| ndufa4l | 4.1E-56 | 0.3018973 | 0.885 | 0.742 | 1.32E-51 | Osteoblasts | ndufa4l |
| rbbp9 | 5.7E-56 | 0.4679781 | 0.223 | 0.118 | 1.84E-51 | Osteoblasts | rbbp9 |
| tns2a | 7.4E-56 | 0.7812794 | 0.126 | 0.055 | 2.42E-51 | Osteoblasts | tns2a |
| tiam2a | 9.2E-56 | 0.6948098 | 0.129 | 0.057 | 3E-51 | Osteoblasts | tiam2a |
| nav1b | 9.3E-56 | 0.4427799 | 0.145 | 0.066 | 3.03E-51 | Osteoblasts | nav1b |
| prss351 | 1.1E-55 | 1.0254763 | 0.172 | 0.087 | 3.6E-51 | Osteoblasts | prss35 |
| tmem184ba | 1.6E-55 | 0.5031455 | 0.141 | 0.064 | 5.06E-51 | Osteoblasts | tmem184ba |
| ism1 | 2.3E-55 | 0.5079076 | 0.221 | 0.117 | 7.45E-51 | Osteoblasts | ism1 |
| cdkn1ba1 | 7.2E-55 | 0.3576927 | 0.212 | 0.11 | 2.33E-50 | Osteoblasts | cdkn1ba |
| si:ch73-238c9.1 | 1.9E-54 | 0.5056115 | 0.191 | 0.097 | 6.28E-50 | Osteoblasts | si:ch73-238c9.1 |
| cx43.41 | 4.1E-54 | 0.3712426 | 0.35 | 0.212 | 1.34E-49 | Osteoblasts | cx43.4 |
| si:dkey-184p18.2 | 1E-53 | 0.843522 | 0.121 | 0.053 | 3.39E-49 | Osteoblasts | si:dkey-184p18.2 |
| myl9a1 | 1.2E-53 | 0.244461 | 0.548 | 0.369 | 4.06E-49 | Osteoblasts | myl9a |
| tacc11 | 1.9E-53 | 0.3659961 | 0.344 | 0.207 | 6.32E-49 | Osteoblasts | tacc1 |
| sort1a1 | 2E-53 | 0.2920581 | 0.206 | 0.107 | 6.41E-49 | Osteoblasts | sort1a |
| tomm20a.1 | 2.6E-53 | 0.3742509 | 0.269 | 0.152 | 8.54E-49 | Osteoblasts | tomm20a.1 |
| zgc:194659 | 7.4E-53 | 0.7382036 | 0.124 | 0.055 | 2.41E-48 | Osteoblasts | zgc:194659 |
| lrrfip1a | 1.1E-52 | 0.5618323 | 0.146 | 0.069 | 3.51E-48 | Osteoblasts | lrrfip1a |
| ppid | 1.2E-52 | 0.6181365 | 0.156 | 0.076 | 3.87E-48 | Osteoblasts | ppid |
| aplp21 | 1.7E-52 | 0.2508939 | 0.486 | 0.311 | 5.37E-48 | Osteoblasts | aplp2 |
| krtcap2 | 1.7E-52 | 0.3643781 | 0.411 | 0.259 | 5.55E-48 | Osteoblasts | krtcap2 |
| slc43a1b | 1.7E-52 | 0.6287404 | 0.109 | 0.045 | 5.56E-48 | Osteoblasts | slc43a1b |
| myl12.2 | 2.2E-52 | 0.3258584 | 0.281 | 0.16 | 7.23E-48 | Osteoblasts | myl12.2 |
| eif4a1b1 | 2.4E-52 | 0.3169148 | 0.764 | 0.599 | 7.9E-48 | Osteoblasts | eif4a1b |
| triqk | 5.6E-52 | 0.3273797 | 0.271 | 0.154 | 1.81E-47 | Osteoblasts | triqk |
| myo1b | 1.5E-51 | 0.3600626 | 0.208 | 0.11 | 4.9E-47 | Osteoblasts | myo1b |
| rps171 | 2E-51 | 0.2028495 | 0.983 | 0.942 | 6.51E-47 | Osteoblasts | rps17 |
| traf4a | 2.3E-51 | 0.4923173 | 0.147 | 0.07 | 7.57E-47 | Osteoblasts | traf4a |
| ppfibp1b | 2.4E-51 | 0.4509001 | 0.242 | 0.136 | 7.84E-47 | Osteoblasts | ppfibp1b |
| ctdspla | 3.3E-51 | 0.5334231 | 0.204 | 0.108 | 1.06E-46 | Osteoblasts | ctdspla |
| pdap1a | 3.5E-51 | 0.3762483 | 0.345 | 0.21 | 1.14E-46 | Osteoblasts | pdap1a |
| ufm1 | 5.3E-51 | 0.4023841 | 0.304 | 0.18 | 1.71E-46 | Osteoblasts | ufm1 |

| | | | | | | | |
|---|---|---|---|---|---|---|---|
| pbdc1 | 7.1E-51 | 0.2715482 | 0.418 | 0.261 | 2.32E-46 | Osteoblasts | pbdc1 |
| dad1 | 8.4E-51 | 0.3195081 | 0.565 | 0.386 | 2.73E-46 | Osteoblasts | dad1 |
| arhgef10 | 8.4E-51 | 0.812351 | 0.103 | 0.043 | 2.74E-46 | Osteoblasts | arhgef10 |
| ak21 | 1.1E-50 | 0.3062833 | 0.547 | 0.367 | 3.47E-46 | Osteoblasts | ak2 |
| sec61g1 | 1.6E-50 | 0.2496098 | 0.962 | 0.921 | 5.24E-46 | Osteoblasts | sec61g |
| gulp1a | 2.2E-50 | 0.4024586 | 0.302 | 0.179 | 7.09E-46 | Osteoblasts | gulp1a |
| tcp1 | 3.3E-50 | 0.3019643 | 0.476 | 0.311 | 1.07E-45 | Osteoblasts | tcp1 |
| vgll4b | 5.1E-50 | 0.3973345 | 0.27 | 0.155 | 1.66E-45 | Osteoblasts | vgll4b |
| smox | 1.6E-49 | 0.5363495 | 0.154 | 0.076 | 5.34E-45 | Osteoblasts | smox |
| higd2a | 1.8E-49 | 0.3442897 | 0.233 | 0.128 | 5.73E-45 | Osteoblasts | higd2a |
| ndufa8 | 2.5E-49 | 0.3188188 | 0.364 | 0.223 | 8.13E-45 | Osteoblasts | ndufa8 |
| ppa1b | 3.7E-49 | 0.4465362 | 0.234 | 0.131 | 1.21E-44 | Osteoblasts | ppa1b |
| zdhhc8b | 5.4E-49 | 0.3846758 | 0.153 | 0.075 | 1.77E-44 | Osteoblasts | zdhhc8b |
| nop16 | 1E-48 | 0.5172824 | 0.128 | 0.059 | 3.27E-44 | Osteoblasts | nop16 |
| nfkbiaa1 | 1.5E-48 | 0.2685588 | 0.572 | 0.412 | 4.83E-44 | Osteoblasts | nfkbiaa |
| grb10a | 3E-48 | 0.3872222 | 0.194 | 0.103 | 9.63E-44 | Osteoblasts | grb10a |
| sec22ba | 3.3E-48 | 0.4142981 | 0.205 | 0.111 | 1.06E-43 | Osteoblasts | sec22ba |
| irx5a | 3.5E-48 | 0.3902686 | 0.167 | 0.085 | 1.13E-43 | Osteoblasts | irx5a |
| steap41 | 4.2E-48 | 1.5337029 | 0.277 | 0.177 | 1.35E-43 | Osteoblasts | steap4 |
| fkbp4 | 7.8E-48 | 0.2485949 | 0.263 | 0.15 | 2.54E-43 | Osteoblasts | fkbp4 |
| rpe65b1 | 1.3E-47 | 0.9031006 | 0.134 | 0.065 | 4.15E-43 | Osteoblasts | rpe65b |
| ppp1cb | 1.6E-47 | 0.3143921 | 0.312 | 0.187 | 5.24E-43 | Osteoblasts | ppp1cb |
| rpl291 | 3.7E-47 | 0.2435828 | 0.972 | 0.928 | 1.2E-42 | Osteoblasts | rpl29 |
| eif3ba | 5.3E-47 | 0.3279768 | 0.587 | 0.416 | 1.72E-42 | Osteoblasts | eif3ba |
| ndufaf3 | 6.1E-47 | 0.4668579 | 0.193 | 0.103 | 1.97E-42 | Osteoblasts | ndufaf3 |
| rin21 | 6.3E-47 | 0.4889584 | 0.196 | 0.106 | 2.05E-42 | Osteoblasts | rin2 |
| tenm31 | 7.5E-47 | 0.3187724 | 0.324 | 0.198 | 2.45E-42 | Osteoblasts | tenm3 |
| p4ha1a | 7.8E-47 | 0.4532099 | 0.175 | 0.091 | 2.54E-42 | Osteoblasts | p4ha1a |
| mrc21 | 1.2E-46 | 0.4779691 | 0.163 | 0.083 | 4.03E-42 | Osteoblasts | mrc2 |
| stk17al | 1.4E-46 | 0.4981359 | 0.145 | 0.071 | 4.43E-42 | Osteoblasts | stk17al |
| ranbp1 | 1.8E-46 | 0.2522755 | 0.506 | 0.337 | 5.94E-42 | Osteoblasts | ranbp1 |
| STMP1 | 2.8E-46 | 0.2796486 | 0.391 | 0.247 | 9.13E-42 | Osteoblasts | STMP1 |
| golim4b | 3.3E-46 | 0.3954596 | 0.191 | 0.103 | 1.08E-41 | Osteoblasts | golim4b |
| egr11 | 5.6E-46 | 0.2033383 | 0.778 | 0.616 | 1.82E-41 | Osteoblasts | egr1 |
| uacab1 | 9.7E-46 | 0.6219295 | 0.139 | 0.068 | 3.16E-41 | Osteoblasts | uacab |
| plgrkt | 4.7E-45 | 0.6643805 | 0.113 | 0.051 | 1.53E-40 | Osteoblasts | plgrkt |
| FOXK2 | 4.7E-45 | 0.3496048 | 0.229 | 0.13 | 1.53E-40 | Osteoblasts | FOXK2 |
| ier3ip1 | 5.1E-45 | 0.2681872 | 0.362 | 0.227 | 1.67E-40 | Osteoblasts | ier3ip1 |
| p4ha2 | 6.9E-45 | 0.3405616 | 0.237 | 0.136 | 2.26E-40 | Osteoblasts | p4ha2 |
| atp5f1e | 1.4E-44 | 0.2779789 | 0.576 | 0.4 | 4.54E-40 | Osteoblasts | atp5f1e |
| ndufs6 | 1.6E-44 | 0.3385029 | 0.235 | 0.135 | 5.19E-40 | Osteoblasts | ndufs6 |
| cux1a | 1.7E-44 | 0.6050484 | 0.132 | 0.064 | 5.61E-40 | Osteoblasts | cux1a |
| atp6v1e1b | 2.2E-44 | 0.4902263 | 0.176 | 0.093 | 7.08E-40 | Osteoblasts | atp6v1e1b |
| atp5po1 | 2.3E-44 | 0.2696533 | 0.595 | 0.412 | 7.59E-40 | Osteoblasts | atp5po |
| pdcd5 | 2.7E-44 | 0.3100174 | 0.26 | 0.152 | 8.73E-40 | Osteoblasts | pdcd5 |
| atp5f1b | 2.8E-44 | 0.3107017 | 0.726 | 0.545 | 9.05E-40 | Osteoblasts | atp5f1b |
| plxna3 | 3E-44 | 0.4890886 | 0.162 | 0.084 | 9.91E-40 | Osteoblasts | plxna3 |
| smad3a | 3.5E-44 | 0.3002117 | 0.446 | 0.295 | 1.15E-39 | Osteoblasts | smad3a |

| | | | | | | | |
|---|---|---|---|---|---|---|---|
| CR936321.5 | 6.4E-44 | 0.4697253 | 0.169 | 0.089 | 2.07E-39 | Osteoblasts | CR936321.5 |
| scinlb2 | 1.4E-43 | 0.2179359 | 0.451 | 0.297 | 4.67E-39 | Osteoblasts | scinlb |
| fndc3ba | 2.1E-43 | 0.3592077 | 0.23 | 0.132 | 6.79E-39 | Osteoblasts | fndc3ba |
| alx4b | 4.2E-43 | 0.5314723 | 0.116 | 0.054 | 1.37E-38 | Osteoblasts | alx4b |
| lbr | 5.1E-43 | 0.6681304 | 0.134 | 0.066 | 1.67E-38 | Osteoblasts | lbr |
| aimp2 | 5.8E-43 | 0.3047239 | 0.275 | 0.164 | 1.9E-38 | Osteoblasts | aimp2 |
| atp5pf | 7.8E-43 | 0.2446282 | 0.566 | 0.392 | 2.53E-38 | Osteoblasts | atp5pf |
| uck2b | 9.6E-43 | 0.6587324 | 0.109 | 0.05 | 3.12E-38 | Osteoblasts | uck2b |
| FBXO48 | 1.3E-42 | 0.5902568 | 0.119 | 0.056 | 4.26E-38 | Osteoblasts | FBXO48 |
| vamp3 | 1.4E-42 | 0.2251896 | 0.379 | 0.24 | 4.39E-38 | Osteoblasts | vamp3 |
| eif4ebp1 | 1.4E-42 | 0.3007698 | 0.25 | 0.146 | 4.55E-38 | Osteoblasts | eif4ebp1 |
| rnaseka | 1.5E-42 | 0.4261285 | 0.184 | 0.1 | 5.02E-38 | Osteoblasts | rnaseka |
| cyp1c11 | 1.7E-42 | 0.2877437 | 0.18 | 0.097 | 5.5E-38 | Osteoblasts | cyp1c1 |
| timp4.11 | 2.6E-42 | 2.2993172 | 0.161 | 0.091 | 8.55E-38 | Osteoblasts | timp4.1 |
| tram2 | 2.6E-42 | 0.4067983 | 0.19 | 0.104 | 8.58E-38 | Osteoblasts | tram2 |
| eprs1 | 2.9E-42 | 0.3839461 | 0.167 | 0.088 | 9.53E-38 | Osteoblasts | eprs1 |
| esd | 3E-42 | 0.2273253 | 0.228 | 0.129 | 9.77E-38 | Osteoblasts | esd |
| slc66a3 | 3.5E-42 | 0.2594495 | 0.204 | 0.113 | 1.13E-37 | Osteoblasts | slc66a3 |
| tcta | 4.9E-42 | 0.4321119 | 0.161 | 0.085 | 1.59E-37 | Osteoblasts | tcta |
| ndrg3b | 6.1E-42 | 0.4323459 | 0.113 | 0.052 | 1.99E-37 | Osteoblasts | ndrg3b |
| tmem181 | 1.1E-41 | 0.4049442 | 0.158 | 0.083 | 3.58E-37 | Osteoblasts | tmem18 |
| zgc:162944 | 1.2E-41 | 0.223242 | 0.433 | 0.283 | 3.81E-37 | Osteoblasts | zgc:162944 |
| nme31 | 1.3E-41 | 0.2634358 | 0.611 | 0.437 | 4.25E-37 | Osteoblasts | nme3 |
| fam114a2 | 1.8E-41 | 0.3704995 | 0.196 | 0.109 | 5.7E-37 | Osteoblasts | fam114a2 |
| vdac11 | 3.1E-41 | 0.2379458 | 0.469 | 0.312 | 1.02E-36 | Osteoblasts | vdac1 |
| si:dkey-239h2.3 | 4E-41 | 0.3054131 | 0.251 | 0.148 | 1.3E-36 | Osteoblasts | si:dkey-239h2.3 |
| cdc42 | 7.2E-41 | 0.2526266 | 0.529 | 0.364 | 2.33E-36 | Osteoblasts | cdc42 |
| c1qbp | 7.6E-41 | 0.2175341 | 0.337 | 0.211 | 2.48E-36 | Osteoblasts | c1qbp |
| lygl11 | 8.7E-41 | 0.2780071 | 0.264 | 0.158 | 2.82E-36 | Osteoblasts | lygl1 |
| mrpl34 | 9.6E-41 | 0.3268639 | 0.215 | 0.123 | 3.12E-36 | Osteoblasts | mrpl34 |
| fam32a | 9.8E-41 | 0.2325319 | 0.32 | 0.198 | 3.18E-36 | Osteoblasts | fam32a |
| cct5 | 1E-40 | 0.2575389 | 0.533 | 0.37 | 3.36E-36 | Osteoblasts | cct5 |
| chad | 1.1E-40 | 0.4064683 | 0.105 | 0.049 | 3.48E-36 | Osteoblasts | chad |
| prdx41 | 2.2E-40 | 0.2776507 | 0.507 | 0.354 | 7.23E-36 | Osteoblasts | prdx4 |
| dspa | 2.9E-40 | 0.4971157 | 0.207 | 0.12 | 9.4E-36 | Osteoblasts | dspa |
| caska2 | 3.9E-40 | 0.3899931 | 0.156 | 0.082 | 1.26E-35 | Osteoblasts | caska |
| atp5f1d | 6.2E-40 | 0.2358241 | 0.48 | 0.323 | 2.02E-35 | Osteoblasts | atp5f1d |
| timp2a1 | 7.8E-40 | 0.2468494 | 0.881 | 0.745 | 2.54E-35 | Osteoblasts | timp2a |
| eif3i1 | 8.2E-40 | 0.2871971 | 0.486 | 0.334 | 2.67E-35 | Osteoblasts | eif3i |
| si:ch211-110e21.32 | 8.5E-40 | 0.2316433 | 0.142 | 0.072 | 2.77E-35 | Osteoblasts | si:ch211-110e21.3 |
| fam207a | 9.2E-40 | 0.4082219 | 0.143 | 0.074 | 2.98E-35 | Osteoblasts | fam207a |
| arl4d1 | 3E-39 | 0.5377051 | 0.13 | 0.066 | 9.89E-35 | Osteoblasts | arl4d |
| dpt | 3.6E-39 | 0.9566106 | 0.234 | 0.149 | 1.19E-34 | Osteoblasts | dpt |
| foxc1b | 3.9E-39 | 0.5675454 | 0.12 | 0.059 | 1.26E-34 | Osteoblasts | foxc1b |
| irf1b1 | 3.9E-39 | 0.3480768 | 0.195 | 0.112 | 1.26E-34 | Osteoblasts | irf1b |
| cmc1 | 4E-39 | 0.3088631 | 0.169 | 0.092 | 1.29E-34 | Osteoblasts | cmc1 |
| trib21 | 5.7E-39 | 0.3498886 | 0.157 | 0.083 | 1.84E-34 | Osteoblasts | trib2 |
| snx4 | 1.9E-38 | 0.3672124 | 0.162 | 0.087 | 6.03E-34 | Osteoblasts | snx4 |

| | | | | | | | |
|---|---|---|---|---|---|---|---|
| fbl | 1.9E-38 | 0.2155943 | 0.332 | 0.212 | 6.23E-34 | Osteoblasts | fbl |
| sars1 | 2.1E-38 | 0.2964885 | 0.235 | 0.14 | 6.8E-34 | Osteoblasts | sars1 |
| ift20 | 2.2E-38 | 0.5006274 | 0.115 | 0.056 | 7.15E-34 | Osteoblasts | ift20 |
| aamp1 | 2.5E-38 | 0.2048931 | 0.248 | 0.147 | 8.12E-34 | Osteoblasts | aamp |
| srp19 | 2.7E-38 | 0.2788395 | 0.228 | 0.134 | 8.68E-34 | Osteoblasts | srp19 |
| rab5if | 3E-38 | 0.3370913 | 0.2 | 0.115 | 9.79E-34 | Osteoblasts | rab5if |
| ptbp1a | 5.8E-38 | 0.2093209 | 0.345 | 0.22 | 1.88E-33 | Osteoblasts | ptbp1a |
| palm2akap2.11 | 6.4E-38 | 0.2981428 | 0.152 | 0.08 | 2.09E-33 | Osteoblasts | palm2akap2.1 |
| vdac31 | 6.8E-38 | 0.2398764 | 0.682 | 0.515 | 2.21E-33 | Osteoblasts | vdac3 |
| TMEM19 | 7.7E-38 | 0.4422147 | 0.129 | 0.066 | 2.5E-33 | Osteoblasts | TMEM19 |
| aldoaa | 1.4E-37 | 0.2123131 | 0.512 | 0.358 | 4.49E-33 | Osteoblasts | aldoaa |
| klf131 | 1.5E-37 | 0.2156843 | 0.21 | 0.121 | 4.78E-33 | Osteoblasts | klf13 |
| uqcrc2a | 1.5E-37 | 0.4389725 | 0.175 | 0.098 | 4.82E-33 | Osteoblasts | uqcrc2a |
| cox6c | 2.2E-37 | 0.2321975 | 0.578 | 0.402 | 7.3E-33 | Osteoblasts | cox6c |
| kars1 | 2.7E-37 | 0.224986 | 0.261 | 0.159 | 8.66E-33 | Osteoblasts | kars1 |
| gemin7 | 3.1E-37 | 0.4736747 | 0.128 | 0.065 | 1.02E-32 | Osteoblasts | gemin7 |
| pinx1 | 4.1E-37 | 0.2538922 | 0.346 | 0.225 | 1.33E-32 | Osteoblasts | pinx1 |
| il34 | 5.4E-37 | 1.7920695 | 0.101 | 0.049 | 1.74E-32 | Osteoblasts | il34 |
| dkc1 | 5.5E-37 | 0.2298303 | 0.291 | 0.183 | 1.78E-32 | Osteoblasts | dkc1 |
| slc38a5a | 5.9E-37 | 0.367306 | 0.13 | 0.066 | 1.93E-32 | Osteoblasts | slc38a5a |
| dynll1 | 7.2E-37 | 0.2591397 | 0.565 | 0.416 | 2.35E-32 | Osteoblasts | dynll1 |
| vps29 | 8E-37 | 0.2167728 | 0.307 | 0.193 | 2.62E-32 | Osteoblasts | vps29 |
| ppp1r14bb1 | 9.2E-37 | 0.3297135 | 0.236 | 0.143 | 2.99E-32 | Osteoblasts | ppp1r14bb |
| nudcd3 | 9.6E-37 | 0.3262555 | 0.149 | 0.079 | 3.11E-32 | Osteoblasts | nudcd3 |
| atp5if1b | 1.6E-36 | 0.2040152 | 0.473 | 0.322 | 5.09E-32 | Osteoblasts | atp5if1b |
| si:dkey-112e7.2 | 2.7E-36 | 0.246487 | 0.337 | 0.218 | 8.9E-32 | Osteoblasts | si:dkey-112e7.2 |
| ccng1 | 3.2E-36 | 0.2230196 | 0.652 | 0.494 | 1.05E-31 | Osteoblasts | ccng1 |
| adi1 | 4E-36 | 0.3363384 | 0.157 | 0.086 | 1.31E-31 | Osteoblasts | adi1 |
| tmem208 | 8.1E-36 | 0.4426125 | 0.122 | 0.062 | 2.64E-31 | Osteoblasts | tmem208 |
| pon3.2.1 | 1.1E-35 | 0.3378281 | 0.1 | 0.047 | 3.59E-31 | Osteoblasts | pon3.2.1 |
| prdx3 | 1.9E-35 | 0.2962428 | 0.225 | 0.135 | 6.07E-31 | Osteoblasts | prdx3 |
| dchs1a1 | 3.1E-35 | 0.2651299 | 0.143 | 0.076 | 1.01E-30 | Osteoblasts | dchs1a |
| si:dkeyp-120h9.1 | 4.8E-35 | 0.3360725 | 0.147 | 0.08 | 1.56E-30 | Osteoblasts | si:dkeyp-120h9.1 |
| naa10 | 4.9E-35 | 0.2559593 | 0.4 | 0.269 | 1.61E-30 | Osteoblasts | naa10 |
| lamtor2 | 6.1E-35 | 0.3238435 | 0.197 | 0.115 | 1.99E-30 | Osteoblasts | lamtor2 |
| dchs1b2 | 6.4E-35 | 0.2244712 | 0.158 | 0.086 | 2.07E-30 | Osteoblasts | dchs1b |
| rbpjb1 | 6.7E-35 | 0.3906866 | 0.165 | 0.092 | 2.16E-30 | Osteoblasts | rbpjb |
| MYG1 | 8.4E-35 | 0.3354697 | 0.174 | 0.099 | 2.72E-30 | Osteoblasts | MYG1 |
| gskip | 9.1E-35 | 0.2593561 | 0.251 | 0.155 | 2.95E-30 | Osteoblasts | gskip |
| cope | 1.1E-34 | 0.2219439 | 0.374 | 0.248 | 3.49E-30 | Osteoblasts | cope |
| nrp2b1 | 1.2E-34 | 0.2870206 | 0.203 | 0.12 | 3.78E-30 | Osteoblasts | nrp2b |
| plod2 | 1.3E-34 | 0.5372925 | 0.129 | 0.068 | 4.19E-30 | Osteoblasts | plod2 |
| crfb4 | 1.8E-34 | 0.3373569 | 0.176 | 0.1 | 5.73E-30 | Osteoblasts | crfb4 |
| eef1db1 | 3.4E-34 | 0.2561374 | 0.735 | 0.569 | 1.11E-29 | Osteoblasts | eef1db |
| polr1d1 | 4.9E-34 | 0.3372465 | 0.148 | 0.081 | 1.6E-29 | Osteoblasts | polr1d |
| tsr2 | 6.6E-34 | 0.4138358 | 0.127 | 0.066 | 2.15E-29 | Osteoblasts | tsr2 |
| got2b | 7E-34 | 0.3580489 | 0.151 | 0.083 | 2.27E-29 | Osteoblasts | got2b |
| sox61 | 1.8E-33 | 0.395646 | 0.17 | 0.098 | 5.75E-29 | Osteoblasts | sox6 |

| | | | | | | | |
|---|---|---|---|---|---|---|---|
| ngdn2 | 1.9E-33 | 0.2586801 | 0.244 | 0.151 | 6.03E-29 | Osteoblasts | ngdn |
| zgc:56525 | 2.3E-33 | 0.2891983 | 0.24 | 0.148 | 7.34E-29 | Osteoblasts | zgc:56525 |
| si:dkey-51e6.12 | 2.4E-33 | 0.2086086 | 0.373 | 0.25 | 7.91E-29 | Osteoblasts | si:dkey-51e6.1 |
| jupa | 2.5E-33 | 0.2549628 | 0.207 | 0.123 | 8.28E-29 | Osteoblasts | jupa |
| golt1bb | 2.6E-33 | 0.3192722 | 0.169 | 0.096 | 8.6E-29 | Osteoblasts | golt1bb |
| hars | 2.7E-33 | 0.3736027 | 0.152 | 0.084 | 8.68E-29 | Osteoblasts | hars |
| snrnp27 | 3.1E-33 | 0.2266355 | 0.219 | 0.132 | 9.97E-29 | Osteoblasts | snrnp27 |
| pin4 | 3.3E-33 | 0.2076488 | 0.233 | 0.142 | 1.06E-28 | Osteoblasts | pin4 |
| arhgef4 | 3.6E-33 | 0.3135686 | 0.134 | 0.072 | 1.18E-28 | Osteoblasts | arhgef4 |
| smim7 | 4E-33 | 0.2249768 | 0.214 | 0.129 | 1.29E-28 | Osteoblasts | smim7 |
| phlda2 | 9.2E-33 | 0.2510283 | 0.308 | 0.213 | 3E-28 | Osteoblasts | phlda2 |
| rnf170 | 9.3E-33 | 0.3350284 | 0.142 | 0.077 | 3.04E-28 | Osteoblasts | rnf170 |
| cab39l | 9.8E-33 | 0.3340929 | 0.196 | 0.116 | 3.19E-28 | Osteoblasts | cab39l |
| banf1 | 1E-32 | 0.2732074 | 0.204 | 0.122 | 3.25E-28 | Osteoblasts | banf1 |
| eif3ha1 | 1E-32 | 0.2092691 | 0.571 | 0.41 | 3.29E-28 | Osteoblasts | eif3ha |
| timm9 | 1.4E-32 | 0.3035708 | 0.163 | 0.093 | 4.46E-28 | Osteoblasts | timm9 |
| igfbp1a | 1.5E-32 | 0.8022367 | 0.125 | 0.068 | 5.02E-28 | Osteoblasts | igfbp1a |
| itgb5 | 1.7E-32 | 0.2767091 | 0.146 | 0.08 | 5.41E-28 | Osteoblasts | itgb5 |
| map2k1 | 1.8E-32 | 0.5185638 | 0.11 | 0.056 | 5.69E-28 | Osteoblasts | map2k1 |
| eif3g1 | 2.2E-32 | 0.2490611 | 0.676 | 0.515 | 7.02E-28 | Osteoblasts | eif3g |
| ufc1 | 2.3E-32 | 0.2147887 | 0.211 | 0.127 | 7.62E-28 | Osteoblasts | ufc1 |
| gnai2b1 | 3.8E-32 | 0.4609309 | 0.115 | 0.059 | 1.24E-27 | Osteoblasts | gnai2b |
| llph | 5E-32 | 0.4146203 | 0.144 | 0.08 | 1.62E-27 | Osteoblasts | llph |
| pdap1b | 6.5E-32 | 0.2477691 | 0.16 | 0.091 | 2.11E-27 | Osteoblasts | pdap1b |
| srsf4 | 8.3E-32 | 0.2156259 | 0.249 | 0.156 | 2.69E-27 | Osteoblasts | srsf4 |
| histh1l1 | 1.1E-31 | 0.3350316 | 0.303 | 0.204 | 3.56E-27 | Osteoblasts | histh1l |
| magt1 | 1.6E-31 | 0.2157469 | 0.375 | 0.253 | 5.13E-27 | Osteoblasts | magt1 |
| atp5mc1 | 1.7E-31 | 0.2884996 | 0.428 | 0.302 | 5.63E-27 | Osteoblasts | atp5mc1 |
| prrc1 | 1.8E-31 | 0.3882545 | 0.136 | 0.074 | 5.73E-27 | Osteoblasts | prrc1 |
| btf3l4 | 2.6E-31 | 0.2000604 | 0.308 | 0.201 | 8.41E-27 | Osteoblasts | btf3l4 |
| ndufa11 | 2.7E-31 | 0.2928888 | 0.211 | 0.129 | 8.93E-27 | Osteoblasts | ndufa11 |
| tmem258 | 3.4E-31 | 0.2132745 | 0.744 | 0.576 | 1.11E-26 | Osteoblasts | tmem258 |
| mbpb | 4.6E-31 | 0.2252305 | 0.108 | 0.055 | 1.5E-26 | Osteoblasts | mbpb |
| mrps34 | 4.9E-31 | 0.4119073 | 0.107 | 0.055 | 1.59E-26 | Osteoblasts | mrps34 |
| mxd4 | 5E-31 | 0.2842145 | 0.153 | 0.087 | 1.64E-26 | Osteoblasts | mxd4 |
| sat1b1 | 7.5E-31 | 0.2619268 | 0.179 | 0.105 | 2.45E-26 | Osteoblasts | sat1b |
| selenof | 8.9E-31 | 0.2519717 | 0.611 | 0.456 | 2.9E-26 | Osteoblasts | selenof |
| bcl7bb | 9.9E-31 | 0.21927 | 0.153 | 0.086 | 3.23E-26 | Osteoblasts | bcl7bb |
| timm8a | 1.4E-30 | 0.3152918 | 0.152 | 0.086 | 4.68E-26 | Osteoblasts | timm8a |
| tmem126a | 1.6E-30 | 0.3368916 | 0.117 | 0.061 | 5.1E-26 | Osteoblasts | tmem126a |
| cd2ap | 1.9E-30 | 0.2902846 | 0.144 | 0.081 | 6.19E-26 | Osteoblasts | cd2ap |
| arcn1b | 2.3E-30 | 0.2429648 | 0.196 | 0.118 | 7.33E-26 | Osteoblasts | arcn1b |
| emc10 | 3.4E-30 | 0.2200924 | 0.224 | 0.139 | 1.1E-25 | Osteoblasts | emc10 |
| cox8a | 3.4E-30 | 0.214269 | 0.72 | 0.551 | 1.11E-25 | Osteoblasts | cox8a |
| map7d1b | 3.8E-30 | 0.2039761 | 0.281 | 0.183 | 1.24E-25 | Osteoblasts | map7d1b |
| ost4 | 3.8E-30 | 0.3104703 | 0.23 | 0.145 | 1.25E-25 | Osteoblasts | ost4 |
| eva1a1 | 5E-30 | 0.5021492 | 0.104 | 0.054 | 1.61E-25 | Osteoblasts | eva1a |
| trappc3 | 5.1E-30 | 0.3364911 | 0.172 | 0.102 | 1.65E-25 | Osteoblasts | trappc3 |

| | | | | | | | |
|---|---|---|---|---|---|---|---|
| unc50 | 5.8E-30 | 0.2841402 | 0.138 | 0.077 | 1.89E-25 | Osteoblasts | unc50 |
| agpat3 | 1.1E-29 | 0.2537694 | 0.137 | 0.076 | 3.51E-25 | Osteoblasts | agpat3 |
| bsdc1 | 1.2E-29 | 0.2514241 | 0.157 | 0.09 | 4.04E-25 | Osteoblasts | bsdc1 |
| exosc3 | 1.5E-29 | 0.3256185 | 0.126 | 0.069 | 4.91E-25 | Osteoblasts | exosc3 |
| copz1 | 1.6E-29 | 0.4708273 | 0.226 | 0.144 | 5.25E-25 | Osteoblasts | copz1 |
| zgc:1533721 | 2.9E-29 | 0.2352216 | 0.146 | 0.083 | 9.58E-25 | Osteoblasts | zgc:153372 |
| tnmd1 | 3.7E-29 | 0.2418309 | 0.226 | 0.146 | 1.21E-24 | Osteoblasts | tnmd |
| kifc3 | 3.8E-29 | 0.3426517 | 0.118 | 0.063 | 1.24E-24 | Osteoblasts | kifc3 |
| rnf185 | 4.4E-29 | 0.2243972 | 0.243 | 0.154 | 1.44E-24 | Osteoblasts | rnf185 |
| ccnd11 | 4.8E-29 | 0.2785025 | 0.132 | 0.073 | 1.57E-24 | Osteoblasts | ccnd1 |
| nip7 | 6.1E-29 | 0.3350042 | 0.113 | 0.06 | 1.98E-24 | Osteoblasts | nip7 |
| timm23a | 6.7E-29 | 0.2369033 | 0.163 | 0.095 | 2.19E-24 | Osteoblasts | timm23a |
| gorasp2 | 8.5E-29 | 0.2418258 | 0.177 | 0.106 | 2.76E-24 | Osteoblasts | gorasp2 |
| fgfr1op2 | 9.1E-29 | 0.3088895 | 0.137 | 0.077 | 2.96E-24 | Osteoblasts | fgfr1op2 |
| scfd1 | 1.1E-28 | 0.2635958 | 0.169 | 0.1 | 3.64E-24 | Osteoblasts | scfd1 |
| chid11 | 1.4E-28 | 0.3661431 | 0.143 | 0.081 | 4.59E-24 | Osteoblasts | chid1 |
| gpx1a1 | 2.9E-28 | 0.2101068 | 0.282 | 0.186 | 9.56E-24 | Osteoblasts | gpx1a |
| mafba1 | 4.4E-28 | 0.2839603 | 0.524 | 0.38 | 1.45E-23 | Osteoblasts | mafba |
| MRPS12 | 4.7E-28 | 0.3300081 | 0.147 | 0.085 | 1.53E-23 | Osteoblasts | MRPS12 |
| mrpl18 | 6.2E-28 | 0.315883 | 0.116 | 0.063 | 2.02E-23 | Osteoblasts | mrpl18 |
| atp6v1g1 | 1.3E-27 | 0.213663 | 0.204 | 0.127 | 4.08E-23 | Osteoblasts | atp6v1g1 |
| CU914776.1 | 1.4E-27 | 1.4594274 | 0.135 | 0.081 | 4.5E-23 | Osteoblasts | CU914776.1 |
| tcea2 | 3.6E-27 | 0.2581267 | 0.132 | 0.074 | 1.18E-22 | Osteoblasts | tcea2 |
| gsc1 | 4.1E-27 | 0.2575656 | 0.182 | 0.112 | 1.35E-22 | Osteoblasts | gsc |
| relch1 | 4.1E-27 | 0.504226 | 0.119 | 0.066 | 1.35E-22 | Osteoblasts | relch |
| pwp2h | 1.5E-26 | 0.3403075 | 0.136 | 0.079 | 4.97E-22 | Osteoblasts | pwp2h |
| snx71 | 1.5E-26 | 0.3404057 | 0.128 | 0.073 | 5.04E-22 | Osteoblasts | snx7 |
| rrs1 | 2.3E-26 | 0.2080645 | 0.174 | 0.106 | 7.34E-22 | Osteoblasts | rrs1 |
| psmg2 | 2.6E-26 | 0.3412232 | 0.167 | 0.102 | 8.29E-22 | Osteoblasts | psmg2 |
| uqcrc2b | 2.7E-26 | 0.2517746 | 0.179 | 0.109 | 8.69E-22 | Osteoblasts | uqcrc2b |
| aldh18a1 | 3.1E-26 | 0.3183994 | 0.109 | 0.059 | 9.95E-22 | Osteoblasts | aldh18a1 |
| tiam1a | 3.4E-26 | 0.3820261 | 0.108 | 0.059 | 1.12E-21 | Osteoblasts | tiam1a |
| eif3m1 | 4.1E-26 | 0.2266798 | 0.751 | 0.617 | 1.34E-21 | Osteoblasts | eif3m |
| si:dkey-248g15.3 | 4.7E-26 | 0.5303933 | 0.278 | 0.194 | 1.52E-21 | Osteoblasts | si:dkey-248g15.3 |
| mrpl30 | 6.6E-26 | 0.2555073 | 0.17 | 0.104 | 2.15E-21 | Osteoblasts | mrpl30 |
| arhgap1 | 1.6E-25 | 0.2465773 | 0.133 | 0.077 | 5.29E-21 | Osteoblasts | arhgap1 |
| nifk | 2.2E-25 | 0.2117999 | 0.145 | 0.085 | 7.12E-21 | Osteoblasts | nifk |
| mkrn11 | 3.7E-25 | 0.2450929 | 0.141 | 0.083 | 1.21E-20 | Osteoblasts | mkrn1 |
| eif2b2 | 3.8E-25 | 0.335351 | 0.119 | 0.067 | 1.25E-20 | Osteoblasts | eif2b2 |
| itgav | 4.8E-25 | 0.3535186 | 0.149 | 0.09 | 1.57E-20 | Osteoblasts | itgav |
| cldni | 6E-25 | 0.7837892 | 0.105 | 0.059 | 1.95E-20 | Osteoblasts | cldni |
| vbp1 | 1E-24 | 0.201509 | 0.194 | 0.123 | 3.4E-20 | Osteoblasts | vbp1 |
| ube2e1 | 1.2E-24 | 0.2357805 | 0.18 | 0.112 | 4E-20 | Osteoblasts | ube2e1 |
| cygb11 | 1.7E-24 | 1.4775868 | 0.459 | 0.42 | 5.59E-20 | Osteoblasts | cygb1 |
| rimkla | 1.7E-24 | 0.2941105 | 0.115 | 0.065 | 5.64E-20 | Osteoblasts | rimkla |
| vaspb1 | 2.3E-24 | 0.3788217 | 0.156 | 0.096 | 7.5E-20 | Osteoblasts | vaspb |
| hsdl2 | 3.7E-24 | 0.338419 | 0.123 | 0.071 | 1.22E-19 | Osteoblasts | hsdl2 |
| bud23 | 4E-24 | 0.2610784 | 0.105 | 0.057 | 1.29E-19 | Osteoblasts | bud23 |

| | | | | | | | |
|---|---|---|---|---|---|---|---|
| ndufa2 | 4.3E-24 | 0.2666582 | 0.148 | 0.09 | 1.4E-19 | Osteoblasts | ndufa2 |
| taf10 | 4.7E-24 | 0.3654149 | 0.124 | 0.072 | 1.53E-19 | Osteoblasts | taf10 |
| urm1 | 6.1E-24 | 0.2812863 | 0.116 | 0.066 | 1.99E-19 | Osteoblasts | urm1 |
| snrpg | 7.8E-24 | 0.2028232 | 0.262 | 0.177 | 2.55E-19 | Osteoblasts | snrpg |
| plpp5 | 9.6E-24 | 0.246426 | 0.111 | 0.062 | 3.12E-19 | Osteoblasts | plpp5 |
| dpm3 | 9.7E-24 | 0.2101249 | 0.149 | 0.09 | 3.14E-19 | Osteoblasts | dpm3 |
| nol7 | 1.2E-23 | 0.2736378 | 0.111 | 0.062 | 3.9E-19 | Osteoblasts | nol7 |
| agap1 | 1.8E-23 | 0.2845344 | 0.125 | 0.073 | 5.92E-19 | Osteoblasts | agap1 |
| mff | 1.9E-23 | 0.3502947 | 0.117 | 0.067 | 6.18E-19 | Osteoblasts | mff |
| ccdc167 | 1.9E-23 | 0.3006518 | 0.11 | 0.062 | 6.21E-19 | Osteoblasts | ccdc167 |
| slc7a10a1 | 2.2E-23 | 0.2486594 | 0.166 | 0.104 | 7.27E-19 | Osteoblasts | slc7a10a |
| utp11 | 2.3E-23 | 0.2615596 | 0.11 | 0.062 | 7.58E-19 | Osteoblasts | utp11 |
| zgc:1008292 | 2.4E-23 | 0.3618791 | 0.18 | 0.115 | 7.71E-19 | Osteoblasts | zgc:100829 |
| tnip1 | 3E-23 | 0.2062248 | 0.262 | 0.177 | 9.73E-19 | Osteoblasts | tnip1 |
| cited4b1 | 3E-23 | 0.289092 | 0.219 | 0.145 | 9.84E-19 | Osteoblasts | cited4b |
| bloc1s61 | 3E-23 | 0.2105829 | 0.257 | 0.174 | 9.86E-19 | Osteoblasts | bloc1s6 |
| farsa | 3.5E-23 | 0.2949303 | 0.132 | 0.078 | 1.15E-18 | Osteoblasts | farsa |
| wdr5 | 3.6E-23 | 0.3048832 | 0.113 | 0.065 | 1.19E-18 | Osteoblasts | wdr5 |
| gar1 | 5E-23 | 0.2909656 | 0.163 | 0.102 | 1.62E-18 | Osteoblasts | gar1 |
| ythdf2 | 6.5E-23 | 0.2917719 | 0.162 | 0.101 | 2.11E-18 | Osteoblasts | ythdf2 |
| FP074902.21 | 6.8E-23 | 0.3742418 | 0.12 | 0.07 | 2.21E-18 | Osteoblasts | FP074902.2 |
| sra1 | 1.9E-22 | 0.2664696 | 0.108 | 0.062 | 6.25E-18 | Osteoblasts | sra1 |
| rrp36 | 2.3E-22 | 0.2206534 | 0.137 | 0.083 | 7.34E-18 | Osteoblasts | rrp36 |
| sf3b5 | 2.9E-22 | 0.31071 | 0.155 | 0.097 | 9.53E-18 | Osteoblasts | sf3b5 |
| mrpl14 | 3.4E-22 | 0.3384195 | 0.118 | 0.069 | 1.1E-17 | Osteoblasts | mrpl14 |
| MYO9B | 3.8E-22 | 0.3079394 | 0.183 | 0.119 | 1.22E-17 | Osteoblasts | MYO9B |
| micall2b | 4E-22 | 0.2506198 | 0.11 | 0.063 | 1.3E-17 | Osteoblasts | micall2b |
| zgc:55943 | 4.2E-22 | 0.234468 | 0.12 | 0.07 | 1.37E-17 | Osteoblasts | zgc:55943 |
| pabpc1b1 | 4.4E-22 | 0.250575 | 0.686 | 0.579 | 1.42E-17 | Osteoblasts | pabpc1b |
| mrps6 | 6.5E-22 | 0.2251818 | 0.131 | 0.078 | 2.1E-17 | Osteoblasts | mrps6 |
| mbnl1 | 7.1E-22 | 0.2426062 | 0.115 | 0.067 | 2.3E-17 | Osteoblasts | mbnl1 |
| psma2 | 8.1E-22 | 0.2241157 | 0.281 | 0.196 | 2.64E-17 | Osteoblasts | psma2 |
| btf32 | 8.9E-22 | 0.2037931 | 0.94 | 0.871 | 2.9E-17 | Osteoblasts | btf3 |
| cd151l | 9.9E-22 | 0.4691881 | 0.119 | 0.071 | 3.22E-17 | Osteoblasts | cd151l |
| dnajb11 | 1E-21 | 0.2697717 | 0.161 | 0.102 | 3.38E-17 | Osteoblasts | dnajb11 |
| mrpl54 | 1.2E-21 | 0.2193737 | 0.145 | 0.089 | 3.9E-17 | Osteoblasts | mrpl54 |
| mrpl13 | 1.7E-21 | 0.3665032 | 0.109 | 0.063 | 5.47E-17 | Osteoblasts | mrpl13 |
| rad23b | 2.3E-21 | 0.233552 | 0.152 | 0.094 | 7.5E-17 | Osteoblasts | rad23b |
| lrch1 | 2.5E-21 | 0.2367313 | 0.101 | 0.057 | 8.14E-17 | Osteoblasts | lrch1 |
| si:ch211-235e9.6 | 2.9E-21 | 0.2117009 | 0.145 | 0.09 | 9.4E-17 | Osteoblasts | si:ch211-235e9.6 |
| rnaset21 | 4.1E-21 | 0.2112154 | 0.192 | 0.126 | 1.32E-16 | Osteoblasts | rnaset2 |
| arf4a.1 | 4.1E-21 | 0.2930237 | 0.12 | 0.072 | 1.35E-16 | Osteoblasts | arf4a.1 |
| bcar1 | 6.4E-21 | 0.2580064 | 0.144 | 0.09 | 2.09E-16 | Osteoblasts | bcar1 |
| acadsb | 8.3E-21 | 0.2893364 | 0.106 | 0.061 | 2.7E-16 | Osteoblasts | acadsb |
| C2H1orf35 | 1E-20 | 0.2385469 | 0.135 | 0.082 | 3.36E-16 | Osteoblasts | C2H1orf35 |
| PYURF | 3E-20 | 0.2223731 | 0.114 | 0.068 | 9.74E-16 | Osteoblasts | PYURF |
| CABZ01058650.1 | 3.6E-20 | 0.2848758 | 0.117 | 0.07 | 1.18E-15 | Osteoblasts | CABZ01058650.1 |
| gnpnat1 | 5.9E-20 | 0.2235037 | 0.139 | 0.086 | 1.91E-15 | Osteoblasts | gnpnat1 |

| | | | | | | | |
|---|---|---|---|---|---|---|---|
| rn7sk2 | 7.3E-20 | 0.2615524 | 0.571 | 0.447 | 2.39E-15 | Osteoblasts | rn7sk |
| tmco1 | 2.8E-19 | 0.200185 | 0.128 | 0.079 | 8.98E-15 | Osteoblasts | tmco1 |
| nt5c2b | 3.5E-19 | 0.2229252 | 0.1 | 0.058 | 1.14E-14 | Osteoblasts | nt5c2b |
| pop7 | 3.6E-19 | 0.4129593 | 0.111 | 0.067 | 1.17E-14 | Osteoblasts | pop7 |
| mrpl16 | 7.9E-19 | 0.2462598 | 0.101 | 0.059 | 2.55E-14 | Osteoblasts | mrpl16 |
| mrpl20 | 1.6E-18 | 0.3561863 | 0.114 | 0.07 | 5.08E-14 | Osteoblasts | mrpl20 |
| slc43a1a | 2.9E-18 | 0.2111077 | 0.127 | 0.079 | 9.33E-14 | Osteoblasts | slc43a1a |
| epcam | 4.5E-18 | 0.50578 | 0.171 | 0.117 | 1.48E-13 | Osteoblasts | epcam |
| ndufs2 | 4.8E-18 | 0.2083863 | 0.166 | 0.109 | 1.56E-13 | Osteoblasts | ndufs2 |
| cfl1l | 4.8E-18 | 0.5116693 | 0.204 | 0.144 | 1.57E-13 | Osteoblasts | cfl1l |
| mrto4 | 1.3E-17 | 0.3129132 | 0.101 | 0.061 | 4.12E-13 | Osteoblasts | mrto4 |
| txn | 2.1E-17 | 0.218423 | 0.469 | 0.366 | 6.88E-13 | Osteoblasts | txn |
| ncln | 3.9E-17 | 0.2106388 | 0.113 | 0.07 | 1.28E-12 | Osteoblasts | ncln |
| dsc2l | 9E-16 | 0.2587894 | 0.106 | 0.066 | 2.93E-11 | Osteoblasts | dsc2l |
| cybrd1 | 6.6E-15 | 0.2398505 | 0.123 | 0.081 | 2.14E-10 | Osteoblasts | cybrd1 |
| nr4a11 | 1.2E-14 | 0.4303545 | 0.144 | 0.099 | 3.96E-10 | Osteoblasts | nr4a1 |
| ndnf1 | 6E-14 | 0.4724371 | 0.146 | 0.102 | 1.95E-09 | Osteoblasts | ndnf |
| jam3b1 | 4.2E-13 | 0.2373129 | 0.283 | 0.217 | 1.35E-08 | Osteoblasts | jam3b |
| mydgf | 4.2E-13 | 0.2149528 | 0.116 | 0.078 | 1.38E-08 | Osteoblasts | mydgf |
| alcama | 6.3E-11 | 0.2685732 | 0.107 | 0.073 | 2.05E-06 | Osteoblasts | alcama |
| txnl4a | 1.1E-09 | 0.2017324 | 0.111 | 0.079 | 3.55E-05 | Osteoblasts | txnl4a |
| efemp2a | 2.1E-09 | 0.2508107 | 0.309 | 0.367 | 6.68E-05 | Osteoblasts | efemp2a |
| si:dkey-193p11.2 | 9.3E-08 | 0.4694661 | 0.314 | 0.254 | 0.003009 | Osteoblasts | si:dkey-193p11.2 |
| cldnb1 | 2.5E-07 | 0.2937001 | 0.183 | 0.146 | 0.008244 | Osteoblasts | cldnb |
| odc1 | 0.00042 | 0.4454144 | 0.146 | 0.123 | 1 | Osteoblasts | odc1 |
| C18H15orf482 | 0.00297 | 0.3557318 | 0.343 | 0.308 | 1 | Osteoblasts | C18H15orf48 |
| abi3bpa1 | 0 | 1.7330245 | 0.615 | 0.28 | 0 | Stroma 1 | abi3bpa |
| sfrp5 | 0 | 2.4254721 | 0.475 | 0.193 | 0 | Stroma 1 | sfrp5 |
| si:dkey-282h22.5 | 0 | 3.1043873 | 0.284 | 0.057 | 0 | Stroma 1 | si:dkey-282h22.5 |
| nova2 | 2E-301 | 1.739461 | 0.489 | 0.197 | 6.8E-297 | Stroma 1 | nova2 |
| scara3 | 3E-299 | 2.0241392 | 0.369 | 0.12 | 9.1E-295 | Stroma 1 | scara3 |
| dcn1 | 2E-293 | 0.856006 | 0.971 | 0.862 | 6.5E-289 | Stroma 1 | dcn |
| angptl7 | 2E-284 | 1.8974615 | 0.465 | 0.187 | 5.5E-280 | Stroma 1 | angptl7 |
| thbs2b1 | 4E-274 | 1.6497262 | 0.524 | 0.234 | 1.4E-269 | Stroma 1 | thbs2b |
| angptl1a | 2E-256 | 1.4602725 | 0.653 | 0.377 | 7.2E-252 | Stroma 1 | angptl1a |
| apcdd1l1 | 3E-243 | 1.909799 | 0.307 | 0.098 | 9.3E-239 | Stroma 1 | apcdd1l |
| abi3bpb | 1E-233 | 1.1520282 | 0.806 | 0.572 | 3.6E-229 | Stroma 1 | abi3bpb |
| si:ch211-243a20.3 | 1E-227 | 1.3394079 | 0.548 | 0.266 | 3.7E-223 | Stroma 1 | si:ch211-243a20.3 |
| col5a11 | 2E-227 | 0.6640574 | 0.995 | 0.963 | 5.5E-223 | Stroma 1 | col5a1 |
| MYADM.11 | 6E-227 | 1.1666913 | 0.483 | 0.214 | 1.9E-222 | Stroma 1 | MYADM.1 |
| cilp | 3E-220 | 1.9641496 | 0.388 | 0.159 | 8.5E-216 | Stroma 1 | cilp |
| si:ch1073-459j12.11 | 6E-220 | 0.7656261 | 0.947 | 0.803 | 1.9E-215 | Stroma 1 | si:ch1073-459j12.1 |
| fbln2 | 4E-201 | 1.7573733 | 0.31 | 0.111 | 1.3E-196 | Stroma 1 | fbln2 |
| si:busm1-57f23.11 | 1E-200 | 0.8533994 | 0.805 | 0.56 | 4.3E-196 | Stroma 1 | si:busm1-57f23.1 |
| tppp3 | 1E-196 | 1.8373214 | 0.195 | 0.051 | 4.5E-192 | Stroma 1 | tppp3 |
| fn1b | 8E-192 | 0.8901955 | 0.67 | 0.385 | 2.5E-187 | Stroma 1 | fn1b |
| adh8a | 1E-191 | 1.8574673 | 0.429 | 0.198 | 4E-187 | Stroma 1 | adh8a |
| col8a1b | 4E-189 | 0.2343462 | 0.338 | 0.129 | 1.4E-184 | Stroma 1 | col8a1b |

| | | | | | | | |
|---|---|---|---|---|---|---|---|
| krt152 | 2E-187 | 0.9249256 | 0.864 | 0.691 | 5.1E-183 | Stroma 1 | krt15 |
| frzb1 | 3E-187 | 2.1348009 | 0.311 | 0.118 | 1.1E-182 | Stroma 1 | frzb |
| foxp41 | 2E-174 | 1.4137495 | 0.407 | 0.191 | 7.8E-170 | Stroma 1 | foxp4 |
| mmp21 | 4E-169 | 0.7161519 | 0.886 | 0.736 | 1.3E-164 | Stroma 1 | mmp2 |
| lgals2a1 | 1E-161 | 0.9226268 | 0.818 | 0.63 | 3.3E-157 | Stroma 1 | lgals2a |
| angptl2a1 | 1E-160 | 1.317422 | 0.533 | 0.306 | 4.7E-156 | Stroma 1 | angptl2a |
| cilp2 | 3E-150 | 1.8364498 | 0.183 | 0.055 | 9.7E-146 | Stroma 1 | cilp2 |
| thbs4a | 9E-150 | 2.2827535 | 0.213 | 0.072 | 2.9E-145 | Stroma 1 | thbs4a |
| colec121 | 3E-147 | 1.2262169 | 0.619 | 0.428 | 1.1E-142 | Stroma 1 | colec12 |
| prelp | 1E-145 | 1.412545 | 0.298 | 0.123 | 3.2E-141 | Stroma 1 | prelp |
| plpp1a2 | 3E-145 | 1.135697 | 0.583 | 0.363 | 8.7E-141 | Stroma 1 | plpp1a |
| ntn1b | 1E-142 | 2.6444127 | 0.112 | 0.025 | 4.3E-138 | Stroma 1 | ntn1b |
| col1a22 | 4E-142 | 0.523809 | 0.998 | 0.991 | 1.3E-137 | Stroma 1 | col1a2 |
| CR383676.23 | 2E-141 | 0.4044888 | 1 | 1 | 5.8E-137 | Stroma 1 | CR383676.2 |
| col5a2a1 | 4E-139 | 0.673324 | 0.894 | 0.786 | 1.2E-134 | Stroma 1 | col5a2a |
| anxa2a | 8E-136 | 0.5311386 | 0.953 | 0.887 | 2.7E-131 | Stroma 1 | anxa2a |
| sgcg1 | 6E-135 | 1.6131245 | 0.227 | 0.083 | 2E-130 | Stroma 1 | sgcg |
| col16a11 | 9E-135 | 0.9886707 | 0.64 | 0.444 | 2.8E-130 | Stroma 1 | col16a1 |
| fndc11 | 3E-132 | 1.3410343 | 0.338 | 0.157 | 9.2E-128 | Stroma 1 | fndc1 |
| sostdc1a | 4E-132 | 2.3275604 | 0.12 | 0.029 | 1.2E-127 | Stroma 1 | sostdc1a |
| thbs3a | 4E-132 | 1.1118858 | 0.503 | 0.297 | 1.4E-127 | Stroma 1 | thbs3a |
| hsd11b21 | 7E-127 | 1.5733501 | 0.227 | 0.086 | 2.3E-122 | Stroma 1 | hsd11b2 |
| rgs5a | 2E-124 | 0.9548181 | 0.315 | 0.142 | 5.4E-120 | Stroma 1 | rgs5a |
| pmp22a | 5E-124 | 0.6536799 | 0.817 | 0.682 | 1.7E-119 | Stroma 1 | pmp22a |
| synm1 | 3E-123 | 1.185578 | 0.411 | 0.22 | 9.6E-119 | Stroma 1 | synm |
| col1a1b2 | 3E-122 | 0.4543552 | 0.998 | 0.986 | 9.6E-118 | Stroma 1 | col1a1b |
| vim2 | 2E-121 | 0.9747244 | 0.562 | 0.359 | 7.8E-117 | Stroma 1 | vim |
| fgl2b | 6E-121 | 0.9292073 | 0.272 | 0.114 | 1.9E-116 | Stroma 1 | fgl2b |
| KCTD151 | 1E-120 | 1.0071378 | 0.527 | 0.331 | 3.5E-116 | Stroma 1 | KCTD15 |
| si:ch211-105c13.32 | 2E-118 | 0.750953 | 0.735 | 0.56 | 6.8E-114 | Stroma 1 | si:ch211-105c13.3 |
| col1a1a2 | 2E-117 | 0.4325153 | 0.995 | 0.979 | 5.8E-113 | Stroma 1 | col1a1a |
| CABZ01092746.13 | 1E-116 | 0.6784515 | 0.838 | 0.702 | 4.7E-112 | Stroma 1 | CABZ01092746.1 |
| fgf18a | 6E-114 | 1.9534669 | 0.134 | 0.039 | 1.9E-109 | Stroma 1 | fgf18a |
| si:ch73-138n13.11 | 1E-113 | 1.0039322 | 0.552 | 0.367 | 3.3E-109 | Stroma 1 | si:ch73-138n13.1 |
| ckba | 8E-113 | 0.6340139 | 0.774 | 0.633 | 2.6E-108 | Stroma 1 | ckba |
| c1qtnf2 | 1E-106 | 1.4746907 | 0.213 | 0.085 | 4.5E-102 | Stroma 1 | c1qtnf2 |
| fstl1b | 9E-106 | 0.6634959 | 0.846 | 0.751 | 3E-101 | Stroma 1 | fstl1b |
| FP102018.11 | 2E-103 | 1.0076258 | 0.548 | 0.382 | 6.7E-99 | Stroma 1 | FP102018.1 |
| serpinf12 | 4E-102 | 0.7787527 | 0.703 | 0.545 | 1.19E-97 | Stroma 1 | serpinf1 |
| aspn1 | 1E-99 | 0.5529704 | 0.595 | 0.392 | 3.76E-95 | Stroma 1 | aspn |
| thbs4b | 1.3E-98 | 1.1321686 | 0.424 | 0.25 | 4.23E-94 | Stroma 1 | thbs4b |
| ognb1 | 9E-97 | 1.3340124 | 0.265 | 0.124 | 2.92E-92 | Stroma 1 | ognb |
| six1a1 | 4.5E-96 | 1.4579148 | 0.259 | 0.123 | 1.46E-91 | Stroma 1 | six1a |
| nog31 | 8.8E-89 | 1.6914028 | 0.185 | 0.076 | 2.88E-84 | Stroma 1 | nog3 |
| lum1 | 2.9E-88 | 0.8922399 | 0.821 | 0.755 | 9.52E-84 | Stroma 1 | lum |
| pbx3b1 | 4.7E-87 | 1.4948074 | 0.268 | 0.136 | 1.52E-82 | Stroma 1 | pbx3b |
| scara52 | 4.5E-85 | 1.2966681 | 0.28 | 0.144 | 1.47E-80 | Stroma 1 | scara5 |
| tsku1 | 1.6E-84 | 1.0534787 | 0.465 | 0.31 | 5.14E-80 | Stroma 1 | tsku |

| | | | | | | | |
|---|---|---|---|---|---|---|---|
| ddr2a | 3.8E-83 | 1.1898467 | 0.345 | 0.2 | 1.23E-78 | Stroma 1 | ddr2a |
| col6a1 | 4.6E-83 | 0.3814821 | 0.94 | 0.871 | 1.5E-78 | Stroma 1 | col6a1 |
| loxl11 | 1.1E-81 | 1.2028451 | 0.339 | 0.193 | 3.55E-77 | Stroma 1 | loxl1 |
| cd991 | 1.7E-81 | 0.5304479 | 0.811 | 0.721 | 5.57E-77 | Stroma 1 | cd99 |
| myo7ab | 2.9E-77 | 1.8480525 | 0.12 | 0.041 | 9.39E-73 | Stroma 1 | myo7ab |
| dpysl3 | 1.3E-76 | 1.1023231 | 0.287 | 0.154 | 4.37E-72 | Stroma 1 | dpysl3 |
| cd9b2 | 1.8E-76 | 1.1972985 | 0.206 | 0.094 | 5.81E-72 | Stroma 1 | cd9b |
| epas1b | 3.3E-76 | 1.8477626 | 0.106 | 0.034 | 1.06E-71 | Stroma 1 | epas1b |
| fstl1a1 | 1.2E-74 | 1.4227739 | 0.212 | 0.101 | 3.74E-70 | Stroma 1 | fstl1a |
| lrp1aa | 3.5E-74 | 0.5782354 | 0.775 | 0.668 | 1.13E-69 | Stroma 1 | lrp1aa |
| fgfrl1a | 2E-73 | 2.1453168 | 0.107 | 0.035 | 6.57E-69 | Stroma 1 | fgfrl1a |
| htra1b | 4.7E-73 | 0.8044625 | 0.551 | 0.404 | 1.53E-68 | Stroma 1 | htra1b |
| tnmd2 | 2.5E-72 | 1.1510593 | 0.271 | 0.145 | 8.25E-68 | Stroma 1 | tnmd |
| cd81a1 | 2.7E-72 | 0.3998611 | 0.932 | 0.901 | 8.84E-68 | Stroma 1 | cd81a |
| bcan1 | 2.4E-70 | 1.0719012 | 0.204 | 0.096 | 7.78E-66 | Stroma 1 | bcan |
| nfixb1 | 6.2E-70 | 0.9069179 | 0.428 | 0.287 | 2E-65 | Stroma 1 | nfixb |
| ogna1 | 1.3E-68 | 1.1571421 | 0.316 | 0.182 | 4.14E-64 | Stroma 1 | ogna |
| rspo12 | 7.7E-68 | 1.8555932 | 0.143 | 0.059 | 2.5E-63 | Stroma 1 | rspo1 |
| zeb1b | 7.9E-68 | 1.4131437 | 0.181 | 0.083 | 2.57E-63 | Stroma 1 | zeb1b |
| col27a1b1 | 1.2E-66 | 0.8169733 | 0.474 | 0.329 | 3.97E-62 | Stroma 1 | col27a1b |
| tgfb31 | 3.3E-66 | 1.377109 | 0.169 | 0.075 | 1.07E-61 | Stroma 1 | tgfb3 |
| clec19a | 7.9E-66 | 1.0514417 | 0.145 | 0.059 | 2.58E-61 | Stroma 1 | clec19a |
| celf22 | 1.3E-65 | 0.9953082 | 0.323 | 0.191 | 4.38E-61 | Stroma 1 | celf2 |
| col6a21 | 2.2E-65 | 0.3375289 | 0.933 | 0.874 | 7.1E-61 | Stroma 1 | col6a2 |
| add3a1 | 6.2E-65 | 0.7186292 | 0.573 | 0.441 | 2.03E-60 | Stroma 1 | add3a |
| CR318588.4 | 1.3E-64 | 0.2817548 | 0.99 | 0.986 | 4.12E-60 | Stroma 1 | CR318588.4 |
| adamts15b | 3.9E-63 | 1.7092472 | 0.125 | 0.049 | 1.28E-58 | Stroma 1 | adamts15b |
| si:dkey-261h17.11 | 1.3E-62 | 0.630539 | 0.427 | 0.283 | 4.17E-58 | Stroma 1 | si:dkey-261h17.1 |
| crabp2b | 8.1E-60 | 0.6941359 | 0.548 | 0.412 | 2.64E-55 | Stroma 1 | crabp2b |
| crabp2a | 8.6E-60 | 0.7708803 | 0.278 | 0.155 | 2.81E-55 | Stroma 1 | crabp2a |
| loxl2b1 | 9.3E-60 | 0.8745111 | 0.435 | 0.31 | 3.03E-55 | Stroma 1 | loxl2b |
| col12a1b | 2.9E-58 | 0.5767742 | 0.737 | 0.617 | 9.54E-54 | Stroma 1 | col12a1b |
| cobll1b1 | 7.4E-58 | 1.3342572 | 0.138 | 0.059 | 2.4E-53 | Stroma 1 | cobll1b |
| thbs2a1 | 1.9E-57 | 1.0206814 | 0.316 | 0.197 | 6.24E-53 | Stroma 1 | thbs2a |
| gas1a1 | 3.1E-56 | 1.0723215 | 0.266 | 0.154 | 1.02E-51 | Stroma 1 | gas1a |
| rgs5b2 | 1.6E-55 | 0.4501049 | 0.659 | 0.515 | 5.14E-51 | Stroma 1 | rgs5b |
| CR925728.1 | 1.7E-55 | 1.224209 | 0.18 | 0.089 | 5.44E-51 | Stroma 1 | CR925728.1 |
| cacna1g1 | 2.1E-54 | 1.2546845 | 0.183 | 0.092 | 6.9E-50 | Stroma 1 | cacna1g |
| tgfbr2b1 | 2.3E-54 | 1.0543154 | 0.244 | 0.138 | 7.49E-50 | Stroma 1 | tgfbr2b |
| adamtsl3 | 5.9E-54 | 1.3025896 | 0.145 | 0.065 | 1.92E-49 | Stroma 1 | adamtsl3 |
| loxa2 | 6.8E-54 | 0.9446288 | 0.282 | 0.168 | 2.21E-49 | Stroma 1 | loxa |
| fxyd11 | 5.7E-53 | 0.4045348 | 0.689 | 0.566 | 1.84E-48 | Stroma 1 | fxyd1 |
| peli1a1 | 8.2E-53 | 1.0155141 | 0.225 | 0.123 | 2.68E-48 | Stroma 1 | peli1a |
| c6.1 | 3.3E-52 | 0.8013739 | 0.318 | 0.199 | 1.08E-47 | Stroma 1 | c6.1 |
| vasna | 4.4E-52 | 1.2095163 | 0.205 | 0.11 | 1.42E-47 | Stroma 1 | vasna |
| lhx6 | 5.2E-52 | 1.6050511 | 0.109 | 0.044 | 1.68E-47 | Stroma 1 | lhx6 |
| pdgfrl | 3.9E-50 | 0.5710867 | 0.665 | 0.581 | 1.28E-45 | Stroma 1 | pdgfrl |
| timp2a2 | 6.1E-50 | 0.3055142 | 0.833 | 0.751 | 1.99E-45 | Stroma 1 | timp2a |

| | | | | | | | |
|---|---|---|---|---|---|---|---|
| hivep3a | 7.6E-50 | 1.1444761 | 0.196 | 0.105 | 2.47E-45 | Stroma 1 | hivep3a |
| antxr1b | 1.3E-47 | 1.3114755 | 0.113 | 0.048 | 4.14E-43 | Stroma 1 | antxr1b |
| col4a1 | 2.2E-47 | 0.2888906 | 0.343 | 0.22 | 7.03E-43 | Stroma 1 | col4a1 |
| zyx3 | 5E-46 | 0.8092585 | 0.42 | 0.312 | 1.61E-41 | Stroma 1 | zyx |
| pear11 | 9.8E-46 | 0.9899591 | 0.22 | 0.126 | 3.2E-41 | Stroma 1 | pear1 |
| CU914487.1 | 2.5E-45 | 1.1393874 | 0.123 | 0.055 | 8.16E-41 | Stroma 1 | CU914487.1 |
| ccn2b | 9.9E-45 | 0.8575296 | 0.309 | 0.2 | 3.21E-40 | Stroma 1 | ccn2b |
| ddr12 | 1.5E-44 | 1.2119794 | 0.161 | 0.082 | 4.96E-40 | Stroma 1 | ddr1 |
| dipk1c2 | 1.8E-43 | 1.4615644 | 0.193 | 0.111 | 5.88E-39 | Stroma 1 | dipk1c |
| ackr3b1 | 9.6E-43 | 1.0234968 | 0.202 | 0.114 | 3.11E-38 | Stroma 1 | ackr3b |
| mast2 | 2.7E-42 | 0.9976568 | 0.255 | 0.159 | 8.8E-38 | Stroma 1 | mast2 |
| meox1 | 2.8E-42 | 0.9535398 | 0.128 | 0.06 | 9E-38 | Stroma 1 | meox1 |
| c6 | 5.1E-41 | 1.1978555 | 0.139 | 0.069 | 1.65E-36 | Stroma 1 | c6 |
| col4a2 | 4.2E-40 | 0.3947116 | 0.322 | 0.213 | 1.36E-35 | Stroma 1 | col4a2 |
| olfml3a1 | 4.9E-40 | 0.9522553 | 0.353 | 0.253 | 1.59E-35 | Stroma 1 | olfml3a |
| crip1 | 2.5E-39 | 0.567144 | 0.565 | 0.456 | 8.05E-35 | Stroma 1 | crip1 |
| mfap22 | 2.2E-38 | 0.6773449 | 0.478 | 0.379 | 7.2E-34 | Stroma 1 | mfap2 |
| rspo31 | 3.5E-38 | 1.593719 | 0.124 | 0.061 | 1.14E-33 | Stroma 1 | rspo3 |
| si:dkey-238i5.2 | 4.7E-38 | 1.3510651 | 0.139 | 0.072 | 1.54E-33 | Stroma 1 | si:dkey-238i5.2 |
| si:dkey-40c11.22 | 8.5E-38 | 0.7522325 | 0.411 | 0.312 | 2.75E-33 | Stroma 1 | si:dkey-40c11.2 |
| tgfb52 | 2.4E-37 | 1.0207664 | 0.189 | 0.11 | 7.78E-33 | Stroma 1 | tgfb5 |
| socs3b1 | 3.2E-36 | 0.7046051 | 0.543 | 0.468 | 1.06E-31 | Stroma 1 | socs3b |
| tns1b1 | 3.7E-35 | 0.6151676 | 0.457 | 0.363 | 1.21E-30 | Stroma 1 | tns1b |
| skia2 | 7.1E-35 | 0.6104789 | 0.526 | 0.445 | 2.3E-30 | Stroma 1 | skia |
| lrp51 | 1.1E-34 | 1.0479558 | 0.184 | 0.108 | 3.46E-30 | Stroma 1 | lrp5 |
| mfap52 | 1.8E-34 | 0.3932187 | 0.644 | 0.54 | 6.01E-30 | Stroma 1 | mfap5 |
| twist1b | 4.1E-33 | 0.897132 | 0.238 | 0.155 | 1.35E-28 | Stroma 1 | twist1b |
| CU638870.11 | 5.3E-33 | 0.7220709 | 0.414 | 0.324 | 1.73E-28 | Stroma 1 | CU638870.1 |
| mt-co31 | 1.2E-32 | 0.2714404 | 0.997 | 0.997 | 3.95E-28 | Stroma 1 | mt-co3 |
| tenm32 | 9.5E-32 | 0.8031319 | 0.29 | 0.203 | 3.09E-27 | Stroma 1 | tenm3 |
| mgst1.2 | 3.3E-31 | 0.6429602 | 0.387 | 0.296 | 1.08E-26 | Stroma 1 | mgst1.2 |
| postna3 | 4E-31 | 0.4059175 | 0.697 | 0.585 | 1.3E-26 | Stroma 1 | postna |
| akap131 | 4.4E-31 | 0.6188517 | 0.409 | 0.316 | 1.42E-26 | Stroma 1 | akap13 |
| rbfox11 | 1.1E-30 | 0.6011663 | 0.482 | 0.402 | 3.6E-26 | Stroma 1 | rbfox1 |
| nfic | 2.1E-30 | 1.0144205 | 0.137 | 0.075 | 6.68E-26 | Stroma 1 | nfic |
| hmcn1 | 6.8E-30 | 1.1288988 | 0.128 | 0.069 | 2.21E-25 | Stroma 1 | hmcn1 |
| clu1 | 7.4E-30 | 0.8324676 | 0.471 | 0.379 | 2.39E-25 | Stroma 1 | clu |
| si:ch211-132b12.7 | 8.7E-30 | 0.8355387 | 0.231 | 0.152 | 2.84E-25 | Stroma 1 | si:ch211-132b12.7 |
| epha4a1 | 1.2E-29 | 1.1614315 | 0.121 | 0.065 | 3.82E-25 | Stroma 1 | epha4a |
| sost1 | 1.4E-29 | 1.3827181 | 0.136 | 0.075 | 4.4E-25 | Stroma 1 | sost |
| tmsb2 | 2E-29 | 0.8095611 | 0.161 | 0.094 | 6.45E-25 | Stroma 1 | tmsb2 |
| sema3fa1 | 3.6E-29 | 1.2139693 | 0.116 | 0.061 | 1.17E-24 | Stroma 1 | sema3fa |
| soul51 | 4.9E-29 | 1.0969317 | 0.354 | 0.273 | 1.59E-24 | Stroma 1 | soul5 |
| fbn2b | 5.6E-29 | 1.1171968 | 0.124 | 0.067 | 1.83E-24 | Stroma 1 | fbn2b |
| igfbp5b2 | 1.7E-28 | 0.6673605 | 0.646 | 0.567 | 5.56E-24 | Stroma 1 | igfbp5b |
| itga11a1 | 1.7E-28 | 0.8807118 | 0.175 | 0.107 | 5.58E-24 | Stroma 1 | itga11a |
| mmp14b1 | 3.4E-28 | 0.4396533 | 0.621 | 0.546 | 1.1E-23 | Stroma 1 | mmp14b |
| flrt21 | 6.4E-28 | 1.099554 | 0.13 | 0.073 | 2.07E-23 | Stroma 1 | flrt2 |

| | | | | | | | |
|---|---|---|---|---|---|---|---|
| pcolce2b | 1E-27 | 0.5200886 | 0.383 | 0.292 | 3.28E-23 | Stroma 1 | pcolce2b |
| prss231 | 1.7E-27 | 0.7270353 | 0.278 | 0.196 | 5.49E-23 | Stroma 1 | prss23 |
| rfx21 | 3.3E-27 | 0.9752689 | 0.124 | 0.068 | 1.06E-22 | Stroma 1 | rfx2 |
| ifitm13 | 1.4E-26 | 0.2517933 | 0.877 | 0.785 | 4.41E-22 | Stroma 1 | ifitm1 |
| myoc | 7.9E-26 | 1.0672828 | 0.119 | 0.066 | 2.57E-21 | Stroma 1 | myoc |
| egfra | 1.3E-25 | 0.6641844 | 0.335 | 0.255 | 4.32E-21 | Stroma 1 | egfra |
| vasnb2 | 2.8E-25 | 0.8343794 | 0.21 | 0.141 | 9.04E-21 | Stroma 1 | vasnb |
| mcf2l2 | 8.9E-25 | 0.9044077 | 0.166 | 0.104 | 2.88E-20 | Stroma 1 | mcf2l2 |
| mt-co2 | 3.9E-24 | 0.225965 | 0.998 | 0.998 | 1.26E-19 | Stroma 1 | mt-co2 |
| fn1a | 6.1E-24 | 0.3599823 | 0.406 | 0.313 | 2E-19 | Stroma 1 | fn1a |
| dock9b1 | 7.8E-24 | 0.7281768 | 0.158 | 0.097 | 2.54E-19 | Stroma 1 | dock9b |
| col11a1b2 | 1.3E-23 | 0.5614298 | 0.589 | 0.542 | 4.16E-19 | Stroma 1 | col11a1b |
| nfia | 1.7E-23 | 0.7114039 | 0.239 | 0.166 | 5.61E-19 | Stroma 1 | nfia |
| meis1b2 | 2.1E-23 | 0.819527 | 0.385 | 0.321 | 6.67E-19 | Stroma 1 | meis1b |
| b2m2 | 4.5E-23 | 0.2909888 | 0.732 | 0.684 | 1.46E-18 | Stroma 1 | b2m |
| rabac1 | 4.8E-23 | 0.4201663 | 0.47 | 0.393 | 1.58E-18 | Stroma 1 | rabac1 |
| lsp1a | 8.2E-23 | 0.852798 | 0.275 | 0.206 | 2.68E-18 | Stroma 1 | lsp1a |
| kirrel1a1 | 9.8E-23 | 0.4813408 | 0.401 | 0.322 | 3.2E-18 | Stroma 1 | kirrel1a |
| C25H11orf961 | 1.2E-22 | 0.6170566 | 0.455 | 0.38 | 3.94E-18 | Stroma 1 | C25H11orf96 |
| ms4a17a.11 | 1.8E-22 | 0.7282458 | 0.319 | 0.249 | 5.92E-18 | Stroma 1 | ms4a17a.11 |
| cavin2b1 | 2.7E-22 | 0.4407988 | 0.588 | 0.528 | 8.85E-18 | Stroma 1 | cavin2b |
| c7a1 | 4.2E-22 | 0.6408029 | 0.242 | 0.171 | 1.37E-17 | Stroma 1 | c7a |
| stmn1b2 | 4.5E-22 | 0.7493683 | 0.184 | 0.12 | 1.45E-17 | Stroma 1 | stmn1b |
| mab21l21 | 5.6E-22 | 0.6156448 | 0.398 | 0.328 | 1.81E-17 | Stroma 1 | mab21l2 |
| fam107b | 5.9E-22 | 0.6922288 | 0.278 | 0.206 | 1.91E-17 | Stroma 1 | fam107b |
| olfml2bb | 1.4E-21 | 0.686016 | 0.144 | 0.088 | 4.57E-17 | Stroma 1 | olfml2bb |
| lfng1 | 2.3E-21 | 0.6212301 | 0.322 | 0.251 | 7.44E-17 | Stroma 1 | lfng |
| pcdh18b | 4.2E-21 | 0.661634 | 0.286 | 0.216 | 1.36E-16 | Stroma 1 | pcdh18b |
| chd5 | 4.6E-21 | 0.8107437 | 0.184 | 0.124 | 1.49E-16 | Stroma 1 | chd5 |
| cald1b2 | 5.7E-21 | 0.6577387 | 0.233 | 0.164 | 1.84E-16 | Stroma 1 | cald1b |
| rrbp1a1 | 9.2E-21 | 0.4430241 | 0.492 | 0.428 | 3E-16 | Stroma 1 | rrbp1a |
| LO018188.12 | 1.4E-20 | 0.6700181 | 0.526 | 0.476 | 4.55E-16 | Stroma 1 | LO018188.1 |
| alcama1 | 6.4E-20 | 1.0081792 | 0.121 | 0.073 | 2.08E-15 | Stroma 1 | alcama |
| emilin2a1 | 1.1E-19 | 0.7007611 | 0.175 | 0.117 | 3.43E-15 | Stroma 1 | emilin2a |
| ano61 | 1.2E-19 | 0.7581219 | 0.181 | 0.122 | 3.85E-15 | Stroma 1 | ano6 |
| lmna2 | 1.2E-19 | 0.2052337 | 0.821 | 0.779 | 4.02E-15 | Stroma 1 | lmna |
| tfpia2 | 1.4E-19 | 0.5285229 | 0.243 | 0.176 | 4.57E-15 | Stroma 1 | tfpia |
| serpinh1b | 1.9E-19 | 0.3516486 | 0.64 | 0.579 | 6.09E-15 | Stroma 1 | serpinh1b |
| PAMR12 | 2.4E-19 | 0.6353115 | 0.176 | 0.117 | 7.79E-15 | Stroma 1 | PAMR1 |
| rn7sk3 | 1.3E-18 | 0.3704439 | 0.514 | 0.453 | 4.26E-14 | Stroma 1 | rn7sk |
| srgn | 1.8E-18 | 0.348989 | 0.514 | 0.449 | 5.71E-14 | Stroma 1 | srgn |
| angptl51 | 3E-18 | 0.6934035 | 0.233 | 0.17 | 9.82E-14 | Stroma 1 | angptl5 |
| si:ch73-46j18.5 | 6E-18 | 0.3042734 | 0.644 | 0.601 | 1.94E-13 | Stroma 1 | si:ch73-46j18.5 |
| prss352 | 1.3E-17 | 0.395636 | 0.142 | 0.091 | 4.25E-13 | Stroma 1 | prss35 |
| pdia71 | 1.5E-17 | 0.7169331 | 0.199 | 0.142 | 4.79E-13 | Stroma 1 | pdia7 |
| arhgef40 | 3E-17 | 0.6143149 | 0.251 | 0.188 | 9.6E-13 | Stroma 1 | arhgef40 |
| htra1a1 | 4.4E-17 | 0.8109785 | 0.283 | 0.225 | 1.43E-12 | Stroma 1 | htra1a |
| utrn1 | 7.8E-17 | 0.6400816 | 0.244 | 0.185 | 2.53E-12 | Stroma 1 | utrn |

| | | | | | | | |
|---|---|---|---|---|---|---|---|
| myh102 | 1.2E-16 | 0.6246481 | 0.389 | 0.332 | 3.78E-12 | Stroma 1 | myh10 |
| si:dkeyp-23e4.32 | 2.1E-16 | 0.7877465 | 0.129 | 0.084 | 6.72E-12 | Stroma 1 | si:dkeyp-23e4.3 |
| socs3a | 3.8E-16 | 0.2700325 | 0.812 | 0.794 | 1.24E-11 | Stroma 1 | socs3a |
| emilin1b1 | 4.9E-16 | 0.7201079 | 0.13 | 0.084 | 1.59E-11 | Stroma 1 | emilin1b |
| pcdh2g281 | 6.2E-16 | 0.5060068 | 0.322 | 0.26 | 2.01E-11 | Stroma 1 | pcdh2g28 |
| si:ch211-156j16.1 | 6.4E-16 | 0.2053193 | 0.211 | 0.15 | 2.09E-11 | Stroma 1 | si:ch211-156j16.1 |
| CR759885.12 | 9.9E-16 | 0.8007421 | 0.146 | 0.099 | 3.22E-11 | Stroma 1 | CR759885.1 |
| fgl2a1 | 1.2E-15 | 0.6294287 | 0.212 | 0.155 | 4.06E-11 | Stroma 1 | fgl2a |
| calm3a1 | 1.3E-15 | 0.2690549 | 0.559 | 0.499 | 4.35E-11 | Stroma 1 | calm3a |
| ch25hl3 | 2.7E-15 | 0.8210556 | 0.152 | 0.105 | 8.72E-11 | Stroma 1 | ch25hl3 |
| cxcl12a2 | 3E-15 | 0.2062496 | 0.775 | 0.677 | 9.6E-11 | Stroma 1 | cxcl12a |
| zfp36l1b1 | 4.7E-15 | 0.3400761 | 0.59 | 0.544 | 1.54E-10 | Stroma 1 | zfp36l1b |
| mmp14a1 | 7.9E-15 | 0.240366 | 0.677 | 0.646 | 2.56E-10 | Stroma 1 | mmp14a |
| plekho21 | 9.2E-15 | 0.5568613 | 0.236 | 0.18 | 2.99E-10 | Stroma 1 | plekho2 |
| CABZ01068367.12 | 9.6E-15 | 0.3637017 | 0.455 | 0.39 | 3.11E-10 | Stroma 1 | CABZ01068367.1 |
| dedd11 | 9.7E-15 | 0.3251484 | 0.593 | 0.55 | 3.16E-10 | Stroma 1 | dedd1 |
| CR383676.32 | 1.3E-14 | 0.4057904 | 0.51 | 0.464 | 4.31E-10 | Stroma 1 | CR383676.3 |
| hivep1 | 2.2E-14 | 0.674597 | 0.203 | 0.151 | 7.17E-10 | Stroma 1 | hivep1 |
| fbxo341 | 3.5E-14 | 0.8283915 | 0.131 | 0.088 | 1.13E-09 | Stroma 1 | fbxo34 |
| svep12 | 3.7E-14 | 0.724027 | 0.191 | 0.143 | 1.21E-09 | Stroma 1 | svep1 |
| serinc52 | 5E-14 | 0.6065418 | 0.291 | 0.237 | 1.62E-09 | Stroma 1 | serinc5 |
| tspan4b1 | 1.2E-13 | 0.4000823 | 0.391 | 0.334 | 3.93E-09 | Stroma 1 | tspan4b |
| si:dkey-11f4.202 | 2.5E-13 | 0.3919241 | 0.527 | 0.487 | 8.25E-09 | Stroma 1 | si:dkey-11f4.20 |
| efna5a1 | 2.7E-13 | 0.848184 | 0.115 | 0.076 | 8.63E-09 | Stroma 1 | efna5a |
| si:dkey-27i16.2 | 3.9E-13 | 0.4893834 | 0.163 | 0.115 | 1.28E-08 | Stroma 1 | si:dkey-27i16.2 |
| ppap2d2 | 4.1E-13 | 0.4250403 | 0.287 | 0.227 | 1.35E-08 | Stroma 1 | ppap2d |
| dusp8a1 | 4.9E-13 | 0.8959879 | 0.128 | 0.088 | 1.58E-08 | Stroma 1 | dusp8a |
| olfml3b1 | 5.8E-13 | 0.3741624 | 0.544 | 0.513 | 1.88E-08 | Stroma 1 | olfml3b |
| cdc42ep4b1 | 7.3E-13 | 0.7145626 | 0.124 | 0.084 | 2.37E-08 | Stroma 1 | cdc42ep4b |
| ctsk1 | 1.1E-12 | 0.4026088 | 0.622 | 0.592 | 3.66E-08 | Stroma 1 | ctsk |
| il15l2 | 1.2E-12 | 0.6916032 | 0.161 | 0.116 | 3.82E-08 | Stroma 1 | il15l |
| phldb1b1 | 1.9E-12 | 0.7502921 | 0.126 | 0.087 | 6.21E-08 | Stroma 1 | phldb1b |
| krt941 | 2.5E-12 | 0.2154047 | 0.351 | 0.294 | 7.99E-08 | Stroma 1 | krt94 |
| rasgef1bb | 2.6E-12 | 0.510782 | 0.201 | 0.153 | 8.44E-08 | Stroma 1 | rasgef1bb |
| pld1a | 2.8E-12 | 0.6826376 | 0.131 | 0.091 | 9.12E-08 | Stroma 1 | pld1a |
| pard3bb | 3E-12 | 0.5579339 | 0.284 | 0.235 | 9.89E-08 | Stroma 1 | pard3bb |
| s100w1 | 3.6E-12 | 0.2857212 | 0.485 | 0.431 | 1.18E-07 | Stroma 1 | s100w |
| si:dkey-7j14.6 | 4.1E-12 | 0.2032488 | 0.789 | 0.777 | 1.34E-07 | Stroma 1 | si:dkey-7j14.6 |
| ugt2a7 | 4.2E-12 | 0.8202667 | 0.131 | 0.092 | 1.36E-07 | Stroma 1 | ugt2a7 |
| spon2b1 | 4.5E-12 | 0.7293195 | 0.171 | 0.126 | 1.46E-07 | Stroma 1 | spon2b |
| sptbn1 | 4.5E-12 | 0.3967439 | 0.3 | 0.247 | 1.47E-07 | Stroma 1 | sptbn1 |
| ahi1 | 4.9E-12 | 0.6927811 | 0.153 | 0.111 | 1.58E-07 | Stroma 1 | ahi1 |
| dlx4a3 | 7.7E-12 | 0.5987035 | 0.239 | 0.191 | 2.5E-07 | Stroma 1 | dlx4a |
| camsap2b | 1.2E-11 | 0.7270499 | 0.146 | 0.106 | 3.76E-07 | Stroma 1 | camsap2b |
| col14a1a1 | 1.4E-11 | 0.7950098 | 0.108 | 0.072 | 4.45E-07 | Stroma 1 | col14a1a |
| CR848833.1 | 1.4E-11 | 0.9098176 | 0.105 | 0.071 | 4.46E-07 | Stroma 1 | CR848833.1 |
| EPB41L2 | 1.7E-11 | 0.6465742 | 0.166 | 0.125 | 5.52E-07 | Stroma 1 | EPB41L2 |
| slit2 | 2.6E-11 | 0.7953375 | 0.102 | 0.068 | 8.49E-07 | Stroma 1 | slit2 |

| | | | | | | | |
|---|---|---|---|---|---|---|---|
| dhrs3a | 2.7E-11 | 0.7574324 | 0.101 | 0.067 | 8.84E-07 | Stroma 1 | dhrs3a |
| lmod1b1 | 3.6E-11 | 0.3902449 | 0.135 | 0.095 | 1.18E-06 | Stroma 1 | lmod1b |
| ches11 | 4E-11 | 0.4979895 | 0.228 | 0.18 | 1.29E-06 | Stroma 1 | ches1 |
| ywhag12 | 4.9E-11 | 0.6428579 | 0.149 | 0.109 | 1.59E-06 | Stroma 1 | ywhag1 |
| zfp36l1a2 | 5.1E-11 | 0.5207157 | 0.252 | 0.205 | 1.67E-06 | Stroma 1 | zfp36l1a |
| plod21 | 5.3E-11 | 1.0126493 | 0.103 | 0.071 | 1.73E-06 | Stroma 1 | plod2 |
| si:dkey-172h23.2 | 6E-11 | 0.5565878 | 0.189 | 0.145 | 1.97E-06 | Stroma 1 | si:dkey-172h23.2 |
| arl6ip5b1 | 7.7E-11 | 0.2850429 | 0.541 | 0.508 | 2.52E-06 | Stroma 1 | arl6ip5b |
| tnfrsf1a | 7.9E-11 | 0.4770971 | 0.297 | 0.25 | 2.58E-06 | Stroma 1 | tnfrsf1a |
| nucb2a | 8.7E-11 | 0.3149851 | 0.556 | 0.523 | 2.84E-06 | Stroma 1 | nucb2a |
| emx21 | 1.4E-10 | 0.8000571 | 0.127 | 0.091 | 4.6E-06 | Stroma 1 | emx2 |
| RAPH11 | 1.7E-10 | 0.6223713 | 0.125 | 0.089 | 5.45E-06 | Stroma 1 | RAPH1 |
| ndnf2 | 2.1E-10 | 0.6427494 | 0.141 | 0.103 | 6.83E-06 | Stroma 1 | ndnf |
| srpx1 | 2.2E-10 | 0.4575791 | 0.21 | 0.165 | 7.26E-06 | Stroma 1 | srpx |
| ablim1b1 | 2.3E-10 | 0.3070447 | 0.308 | 0.254 | 7.52E-06 | Stroma 1 | ablim1b |
| mxra8a1 | 4.3E-10 | 0.4628987 | 0.297 | 0.252 | 1.41E-05 | Stroma 1 | mxra8a |
| igfbp2b | 4.6E-10 | 0.6194319 | 0.101 | 0.069 | 1.5E-05 | Stroma 1 | igfbp2b |
| mdka | 4.9E-10 | 0.8044689 | 0.541 | 0.522 | 1.58E-05 | Stroma 1 | mdka |
| myo9aa | 5.6E-10 | 0.7958394 | 0.103 | 0.072 | 1.81E-05 | Stroma 1 | myo9aa |
| zcchc242 | 6.2E-10 | 0.5760354 | 0.297 | 0.255 | 2.01E-05 | Stroma 1 | zcchc24 |
| c1qtnf11 | 6.9E-10 | 0.7460126 | 0.106 | 0.074 | 2.26E-05 | Stroma 1 | c1qtnf1 |
| si:ch211-57n23.12 | 7.1E-10 | 0.4036111 | 0.263 | 0.216 | 2.32E-05 | Stroma 1 | si:ch211-57n23.1 |
| uap1 | 7.2E-10 | 0.568038 | 0.17 | 0.13 | 2.33E-05 | Stroma 1 | uap1 |
| eya11 | 1.3E-09 | 0.6228586 | 0.173 | 0.134 | 4.13E-05 | Stroma 1 | eya1 |
| gsc2 | 1.9E-09 | 0.5699841 | 0.153 | 0.116 | 6.19E-05 | Stroma 1 | gsc |
| boc1 | 2.2E-09 | 0.2780336 | 0.415 | 0.371 | 7.19E-05 | Stroma 1 | boc |
| itga51 | 2.7E-09 | 0.4329278 | 0.297 | 0.255 | 8.88E-05 | Stroma 1 | itga5 |
| six2a1 | 3.3E-09 | 0.5582047 | 0.135 | 0.1 | 0.000108 | Stroma 1 | six2a |
| svila1 | 3.4E-09 | 0.5099657 | 0.187 | 0.147 | 0.000111 | Stroma 1 | svila |
| pbx1a1 | 3.6E-09 | 0.4345876 | 0.273 | 0.231 | 0.000116 | Stroma 1 | pbx1a |
| f101 | 3.9E-09 | 0.3106549 | 0.436 | 0.398 | 0.000126 | Stroma 1 | f10 |
| fibina1 | 4.1E-09 | 0.7373964 | 0.251 | 0.211 | 0.000134 | Stroma 1 | fibina |
| asap1b1 | 4.4E-09 | 0.5383352 | 0.189 | 0.15 | 0.000144 | Stroma 1 | asap1b |
| atp10a1 | 6.5E-09 | 0.7914225 | 0.128 | 0.096 | 0.000212 | Stroma 1 | atp10a |
| net11 | 6.6E-09 | 0.3862355 | 0.408 | 0.372 | 0.000214 | Stroma 1 | net1 |
| ctdspl2b1 | 8.5E-09 | 0.5459867 | 0.147 | 0.112 | 0.000277 | Stroma 1 | ctdspl2b |
| mgst1.11 | 1.1E-08 | 0.3999335 | 0.29 | 0.249 | 0.00035 | Stroma 1 | mgst1.1 |
| slc12a22 | 1.2E-08 | 0.6055524 | 0.18 | 0.142 | 0.000388 | Stroma 1 | slc12a2 |
| arrdc3a | 1.5E-08 | 0.5632916 | 0.202 | 0.163 | 0.000484 | Stroma 1 | arrdc3a |
| tlr5b | 1.6E-08 | 0.7377291 | 0.102 | 0.073 | 0.000506 | Stroma 1 | tlr5b |
| asap3 | 1.8E-08 | 0.5809439 | 0.16 | 0.125 | 0.000599 | Stroma 1 | asap3 |
| icam3 | 1.9E-08 | 0.5748724 | 0.112 | 0.082 | 0.000611 | Stroma 1 | icam3 |
| b3gnt5a1 | 2.2E-08 | 0.64651 | 0.144 | 0.111 | 0.000704 | Stroma 1 | b3gnt5a |
| si:ch211-108d22.2 | 2.8E-08 | 0.407023 | 0.25 | 0.211 | 0.000901 | Stroma 1 | si:ch211-108d22.2 |
| SBNO2 | 2.8E-08 | 0.4652835 | 0.265 | 0.226 | 0.000907 | Stroma 1 | SBNO2 |
| akap12b | 3E-08 | 0.5056367 | 0.177 | 0.142 | 0.000985 | Stroma 1 | akap12b |
| hdac4 | 3.7E-08 | 0.5675203 | 0.153 | 0.119 | 0.00121 | Stroma 1 | hdac4 |
| setbp12 | 4.7E-08 | 0.3669077 | 0.287 | 0.247 | 0.001531 | Stroma 1 | setbp1 |

| | | | | | | | |
|---|---|---|---|---|---|---|---|
| calr3a | 5.4E-08 | 0.3752748 | 0.393 | 0.361 | 0.001752 | Stroma 1 | calr3a |
| si:dkey-17m8.1 | 5.9E-08 | 0.5036296 | 0.156 | 0.121 | 0.001916 | Stroma 1 | si:dkey-17m8.1 |
| il11ra2 | 6E-08 | 0.3948136 | 0.277 | 0.238 | 0.001937 | Stroma 1 | il11ra |
| pamr11 | 8.2E-08 | 0.5335319 | 0.141 | 0.108 | 0.002676 | Stroma 1 | pamr1 |
| lmcd1 | 9.1E-08 | 0.7348483 | 0.116 | 0.087 | 0.002957 | Stroma 1 | lmcd1 |
| plxna31 | 9.5E-08 | 0.6618089 | 0.117 | 0.088 | 0.003073 | Stroma 1 | plxna3 |
| dhrs92 | 1.8E-07 | 0.4825146 | 0.173 | 0.139 | 0.0058 | Stroma 1 | dhrs9 |
| qkia | 2.4E-07 | 0.3140547 | 0.483 | 0.468 | 0.007934 | Stroma 1 | qkia |
| calua | 2.9E-07 | 0.3545572 | 0.36 | 0.325 | 0.00944 | Stroma 1 | calua |
| zgc:1102391 | 3E-07 | 0.4500943 | 0.263 | 0.231 | 0.009779 | Stroma 1 | zgc:110239 |
| olfml2ba2 | 4.4E-07 | 0.4919247 | 0.157 | 0.124 | 0.014401 | Stroma 1 | olfml2ba |
| kdelr31 | 6.1E-07 | 0.4261089 | 0.215 | 0.181 | 0.019707 | Stroma 1 | kdelr3 |
| gas1b2 | 7E-07 | 0.455687 | 0.229 | 0.195 | 0.022866 | Stroma 1 | gas1b |
| si:ch1073-392o20.2 | 9.3E-07 | 0.3534079 | 0.277 | 0.241 | 0.030276 | Stroma 1 | si:ch1073-392o20.2 |
| fmnl2a | 9.9E-07 | 0.5380556 | 0.138 | 0.109 | 0.032187 | Stroma 1 | fmnl2a |
| zgc:1135312 | 1.1E-06 | 0.3043521 | 0.316 | 0.277 | 0.037267 | Stroma 1 | zgc:113531 |
| niban1b | 1.2E-06 | 0.3924712 | 0.123 | 0.094 | 0.03801 | Stroma 1 | niban1b |
| si:dkey-16l2.16 | 1.2E-06 | 0.4882528 | 0.146 | 0.117 | 0.039711 | Stroma 1 | si:dkey-16l2.16 |
| sertad4 | 1.4E-06 | 0.4825178 | 0.226 | 0.194 | 0.043996 | Stroma 1 | sertad4 |
| bcl2a | 2E-06 | 0.5058844 | 0.184 | 0.154 | 0.066333 | Stroma 1 | bcl2a |
| bhlhe41 | 2.2E-06 | 0.570152 | 0.157 | 0.128 | 0.072144 | Stroma 1 | bhlhe41 |
| FO704772.3 | 2.2E-06 | 0.3971306 | 0.358 | 0.329 | 0.072701 | Stroma 1 | FO704772.3 |
| podn1 | 2.9E-06 | 0.3704785 | 0.204 | 0.171 | 0.092828 | Stroma 1 | podn |
| coro1ca1 | 3E-06 | 0.3964064 | 0.268 | 0.237 | 0.097717 | Stroma 1 | coro1ca |
| cpne3 | 3.1E-06 | 0.5491998 | 0.134 | 0.107 | 0.099294 | Stroma 1 | cpne3 |
| tnrc6c1 | 3.2E-06 | 0.2597136 | 0.502 | 0.486 | 0.102515 | Stroma 1 | tnrc6c1 |
| bnc2 | 3.4E-06 | 0.4835645 | 0.107 | 0.081 | 0.110928 | Stroma 1 | bnc2 |
| bmpr2a | 4.3E-06 | 0.4071301 | 0.208 | 0.177 | 0.1398 | Stroma 1 | bmpr2a |
| SCARF21 | 4.9E-06 | 0.3829887 | 0.297 | 0.267 | 0.16023 | Stroma 1 | SCARF2 |
| mertka | 5.8E-06 | 0.5059101 | 0.148 | 0.12 | 0.189434 | Stroma 1 | mertka |
| arl4ab1 | 6E-06 | 0.4782296 | 0.137 | 0.109 | 0.196348 | Stroma 1 | arl4ab |
| ntn1a | 6.3E-06 | 0.4834399 | 0.15 | 0.12 | 0.205935 | Stroma 1 | ntn1a |
| zgc:175088 | 6.4E-06 | 0.3909203 | 0.301 | 0.273 | 0.207998 | Stroma 1 | zgc:175088 |
| pde7a | 7.4E-06 | 0.6553278 | 0.116 | 0.091 | 0.239935 | Stroma 1 | pde7a |
| kif1b1 | 7.6E-06 | 0.4625919 | 0.126 | 0.1 | 0.245647 | Stroma 1 | kif1b |
| sfrp21 | 8.2E-06 | 0.6675473 | 0.414 | 0.398 | 0.265529 | Stroma 1 | sfrp2 |
| mmp15a1 | 1.1E-05 | 0.4331839 | 0.107 | 0.083 | 0.345847 | Stroma 1 | mmp15a |
| fhod11 | 1.2E-05 | 0.6428778 | 0.104 | 0.081 | 0.39773 | Stroma 1 | fhod1 |
| sik2b1 | 1.3E-05 | 0.5232916 | 0.232 | 0.206 | 0.418259 | Stroma 1 | sik2b |
| zfhx41 | 1.4E-05 | 0.2914598 | 0.463 | 0.45 | 0.46344 | Stroma 1 | zfhx4 |
| reep5 | 1.4E-05 | 0.2404082 | 0.546 | 0.541 | 0.46734 | Stroma 1 | reep5 |
| lgmn | 1.5E-05 | 0.3811998 | 0.2 | 0.171 | 0.485418 | Stroma 1 | lgmn |
| hivep2a | 1.5E-05 | 0.4760724 | 0.242 | 0.215 | 0.488919 | Stroma 1 | hivep2a |
| si:dkey-235d18.5 | 1.6E-05 | 0.4855602 | 0.122 | 0.097 | 0.522012 | Stroma 1 | si:dkey-235d18.5 |
| jam3b2 | 1.6E-05 | 0.4632222 | 0.247 | 0.22 | 0.529132 | Stroma 1 | jam3b |
| si:ch73-380l3.22 | 1.7E-05 | 0.2829171 | 0.442 | 0.416 | 0.555606 | Stroma 1 | si:ch73-380l3.2 |
| si:ch211-133n4.41 | 1.9E-05 | 0.2091607 | 0.47 | 0.447 | 0.611832 | Stroma 1 | si:ch211-133n4.4 |
| lhfpl2a | 2E-05 | 0.6262815 | 0.13 | 0.107 | 0.645431 | Stroma 1 | lhfpl2a |

| | | | | | | | |
|---|---|---|---|---|---|---|---|
| pdia5 | 2E-05 | 0.4804048 | 0.192 | 0.165 | 0.664796 | Stroma 1 | pdia5 |
| ehbp12 | 2.1E-05 | 0.4134948 | 0.136 | 0.11 | 0.678188 | Stroma 1 | ehbp1 |
| sesn11 | 2.2E-05 | 0.4665523 | 0.246 | 0.217 | 0.71182 | Stroma 1 | sesn1 |
| ccdc85b1 | 2.2E-05 | 0.404139 | 0.217 | 0.189 | 0.713424 | Stroma 1 | ccdc85b |
| smarcc2 | 3E-05 | 0.4845914 | 0.198 | 0.172 | 0.964563 | Stroma 1 | smarcc2 |
| lix1l1 | 3.5E-05 | 0.4978764 | 0.199 | 0.174 | 1 | Stroma 1 | lix1l |
| nhsl21 | 3.8E-05 | 0.4983513 | 0.125 | 0.101 | 1 | Stroma 1 | nhsl2 |
| si:ch73-44m9.3.1 | 3.9E-05 | 0.4773103 | 0.122 | 0.098 | 1 | Stroma 1 | si:ch73-44m9.3.1 |
| rnd3a2 | 4.8E-05 | 0.4123036 | 0.308 | 0.285 | 1 | Stroma 1 | rnd3a |
| adam10a1 | 5.5E-05 | 0.2066518 | 0.403 | 0.381 | 1 | Stroma 1 | adam10a |
| osr1 | 5.8E-05 | 0.5097435 | 0.197 | 0.172 | 1 | Stroma 1 | osr1 |
| zgc:1584631 | 6.4E-05 | 0.2197094 | 0.997 | 0.997 | 1 | Stroma 1 | zgc:158463 |
| ctsba | 6.6E-05 | 0.3035868 | 0.402 | 0.384 | 1 | Stroma 1 | ctsba |
| itgb1a1 | 6.8E-05 | 0.3062139 | 0.289 | 0.264 | 1 | Stroma 1 | itgb1a |
| reep3a | 7.1E-05 | 0.4480543 | 0.189 | 0.165 | 1 | Stroma 1 | reep3a |
| cops9 | 7.4E-05 | 0.4781126 | 0.177 | 0.153 | 1 | Stroma 1 | cops9 |
| smo | 7.8E-05 | 0.2702755 | 0.284 | 0.257 | 1 | Stroma 1 | smo |
| chka2 | 7.9E-05 | 0.562589 | 0.111 | 0.09 | 1 | Stroma 1 | chka |
| antxr2a1 | 0.0001 | 0.2971381 | 0.357 | 0.335 | 1 | Stroma 1 | antxr2a |
| DAB22 | 0.00011 | 0.2922655 | 0.403 | 0.378 | 1 | Stroma 1 | DAB2 |
| per3 | 0.00011 | 0.2872647 | 0.367 | 0.347 | 1 | Stroma 1 | per3 |
| ptprfb1 | 0.00011 | 0.3365888 | 0.278 | 0.254 | 1 | Stroma 1 | ptprfb |
| dnajc4 | 0.00012 | 0.5224739 | 0.116 | 0.095 | 1 | Stroma 1 | dnajc4 |
| ebf3a2 | 0.00013 | 0.259423 | 0.244 | 0.215 | 1 | Stroma 1 | ebf3a |
| lrrc172 | 0.00013 | 0.3364465 | 0.415 | 0.396 | 1 | Stroma 1 | lrrc17 |
| nkx3-22 | 0.00013 | 0.4953195 | 0.219 | 0.195 | 1 | Stroma 1 | nkx3-2 |
| cd164 | 0.00015 | 0.2996409 | 0.357 | 0.337 | 1 | Stroma 1 | cd164 |
| lsp1b2 | 0.00015 | 0.469195 | 0.281 | 0.258 | 1 | Stroma 1 | lsp1b |
| prnprs3 | 0.00015 | 0.4613576 | 0.117 | 0.096 | 1 | Stroma 1 | prnprs3 |
| nfatc3a | 0.00017 | 0.4726915 | 0.114 | 0.093 | 1 | Stroma 1 | nfatc3a |
| rbm4.1 | 0.00019 | 0.5413981 | 0.121 | 0.1 | 1 | Stroma 1 | rbm4.1 |
| ahr21 | 0.00019 | 0.2945167 | 0.398 | 0.384 | 1 | Stroma 1 | ahr2 |
| ccdc3b1 | 0.00019 | 0.4512359 | 0.146 | 0.124 | 1 | Stroma 1 | ccdc3b |
| skib | 0.00021 | 0.3835641 | 0.225 | 0.202 | 1 | Stroma 1 | skib |
| tjp1b | 0.00023 | 0.4389701 | 0.165 | 0.142 | 1 | Stroma 1 | tjp1b |
| mt-nd3 | 0.00024 | 0.3064228 | 0.374 | 0.361 | 1 | Stroma 1 | mt-nd3 |
| rarga | 0.00029 | 0.2442149 | 0.3 | 0.277 | 1 | Stroma 1 | rarga |
| clk4a2 | 0.0003 | 0.3478338 | 0.251 | 0.229 | 1 | Stroma 1 | clk4a |
| paxbp1 | 0.00039 | 0.3479465 | 0.139 | 0.117 | 1 | Stroma 1 | paxbp1 |
| ITLN13 | 0.0004 | 0.2396678 | 0.237 | 0.21 | 1 | Stroma 1 | ITLN1 |
| ppt1 | 0.00041 | 0.2671437 | 0.362 | 0.344 | 1 | Stroma 1 | ppt1 |
| arglu1a1 | 0.00043 | 0.392226 | 0.195 | 0.172 | 1 | Stroma 1 | arglu1a |
| stat3 | 0.00044 | 0.4683167 | 0.17 | 0.15 | 1 | Stroma 1 | stat3 |
| ptgs2a1 | 0.00048 | 0.4618057 | 0.152 | 0.13 | 1 | Stroma 1 | ptgs2a |
| trib22 | 0.00049 | 0.4512566 | 0.107 | 0.088 | 1 | Stroma 1 | trib2 |
| si:ch73-173h19.32 | 0.00051 | 0.2700677 | 0.267 | 0.238 | 1 | Stroma 1 | si:ch73-173h19.3 |
| rnf24 | 0.00051 | 0.4864166 | 0.111 | 0.092 | 1 | Stroma 1 | rnf24 |
| nos1apa1 | 0.00053 | 0.4744642 | 0.128 | 0.108 | 1 | Stroma 1 | nos1apa |

| | | | | | | | |
|---|---|---|---|---|---|---|---|
| calcoco1a | 0.00057 | 0.3920031 | 0.121 | 0.101 | 1 | Stroma 1 | calcoco1a |
| knop1 | 0.0006 | 0.4409091 | 0.184 | 0.162 | 1 | Stroma 1 | knop1 |
| calr3b | 0.0007 | 0.2484144 | 0.295 | 0.27 | 1 | Stroma 1 | calr3b |
| tmem9b | 0.00075 | 0.5109023 | 0.121 | 0.102 | 1 | Stroma 1 | tmem9b |
| serinc2 | 0.00079 | 0.2626149 | 0.334 | 0.317 | 1 | Stroma 1 | serinc2 |
| rai141 | 0.00082 | 0.5716614 | 0.103 | 0.086 | 1 | Stroma 1 | rai14 |
| atrnl1b | 0.00082 | 0.4601555 | 0.128 | 0.109 | 1 | Stroma 1 | atrnl1b |
| si:ch211-196l7.41 | 0.00088 | 0.2058345 | 0.481 | 0.476 | 1 | Stroma 1 | si:ch211-196l7.4 |
| klf2a1 | 0.00096 | 0.3712857 | 0.303 | 0.285 | 1 | Stroma 1 | klf2a |
| tmem119b | 0.001 | 0.2715886 | 0.23 | 0.206 | 1 | Stroma 1 | tmem119b |
| zgc:113208 | 0.00122 | 0.3922791 | 0.21 | 0.191 | 1 | Stroma 1 | zgc:113208 |
| ecm21 | 0.00128 | 0.2343735 | 0.32 | 0.298 | 1 | Stroma 1 | ecm2 |
| exoc6 | 0.00143 | 0.4020034 | 0.117 | 0.099 | 1 | Stroma 1 | exoc6 |
| nrp2b2 | 0.00151 | 0.4832959 | 0.143 | 0.125 | 1 | Stroma 1 | nrp2b |
| mex3b | 0.00155 | 0.3228928 | 0.192 | 0.171 | 1 | Stroma 1 | mex3b |
| atl31 | 0.00161 | 0.2410997 | 0.34 | 0.322 | 1 | Stroma 1 | atl3 |
| zmiz1a | 0.00163 | 0.552396 | 0.137 | 0.12 | 1 | Stroma 1 | zmiz1a |
| CABZ01074397.11 | 0.00168 | 0.324889 | 0.168 | 0.149 | 1 | Stroma 1 | CABZ01074397.1 |
| dpysl2b1 | 0.00174 | 0.2997311 | 0.182 | 0.161 | 1 | Stroma 1 | dpysl2b |
| bmp5 | 0.00181 | 0.5386682 | 0.105 | 0.088 | 1 | Stroma 1 | bmp5 |
| si:dkey-199f5.81 | 0.00193 | 0.3381761 | 0.184 | 0.164 | 1 | Stroma 1 | si:dkey-199f5.8 |
| rnd1b2 | 0.00203 | 0.4082568 | 0.12 | 0.103 | 1 | Stroma 1 | rnd1b |
| bco2l | 0.00215 | 0.2909289 | 0.143 | 0.124 | 1 | Stroma 1 | bco2l |
| waslb | 0.00224 | 0.3594972 | 0.194 | 0.175 | 1 | Stroma 1 | waslb |
| MDFIC | 0.00226 | 0.3657211 | 0.204 | 0.186 | 1 | Stroma 1 | MDFIC |
| trim3b2 | 0.00229 | 0.489857 | 0.111 | 0.094 | 1 | Stroma 1 | trim3b |
| kdm6bb1 | 0.00244 | 0.3055571 | 0.16 | 0.14 | 1 | Stroma 1 | kdm6bb |
| glis2a | 0.00266 | 0.3644565 | 0.221 | 0.204 | 1 | Stroma 1 | glis2a |
| chd31 | 0.00278 | 0.2588826 | 0.195 | 0.174 | 1 | Stroma 1 | chd3 |
| fkbp10a1 | 0.00278 | 0.3536971 | 0.153 | 0.134 | 1 | Stroma 1 | fkbp10a |
| myo1ca2 | 0.00278 | 0.2409951 | 0.224 | 0.202 | 1 | Stroma 1 | myo1ca |
| cav21 | 0.00282 | 0.3689479 | 0.19 | 0.172 | 1 | Stroma 1 | cav2 |
| pkd2 | 0.00286 | 0.386602 | 0.125 | 0.107 | 1 | Stroma 1 | pkd2 |
| dbpb | 0.003 | 0.322888 | 0.131 | 0.113 | 1 | Stroma 1 | dbpb |
| jak1 | 0.00312 | 0.2400306 | 0.441 | 0.439 | 1 | Stroma 1 | jak1 |
| u2surp1 | 0.00337 | 0.2531899 | 0.329 | 0.313 | 1 | Stroma 1 | u2surp |
| golga4 | 0.00339 | 0.3478503 | 0.179 | 0.162 | 1 | Stroma 1 | golga4 |
| ezh1 | 0.00354 | 0.411194 | 0.158 | 0.141 | 1 | Stroma 1 | ezh1 |
| scinla | 0.00373 | 0.2388223 | 0.426 | 0.409 | 1 | Stroma 1 | scinla |
| gng2 | 0.00375 | 0.3658551 | 0.126 | 0.109 | 1 | Stroma 1 | gng2 |
| creb3l3l | 0.00432 | 0.2558119 | 0.266 | 0.249 | 1 | Stroma 1 | creb3l3l |
| mia3 | 0.00432 | 0.2937097 | 0.213 | 0.196 | 1 | Stroma 1 | mia3 |
| dvl2 | 0.00504 | 0.4112883 | 0.145 | 0.129 | 1 | Stroma 1 | dvl2 |
| mrc22 | 0.00544 | 0.4017617 | 0.103 | 0.088 | 1 | Stroma 1 | mrc2 |
| tent5bb2 | 0.00612 | 0.4187482 | 0.158 | 0.142 | 1 | Stroma 1 | tent5bb |
| sorbs31 | 0.00655 | 0.2381494 | 0.208 | 0.19 | 1 | Stroma 1 | sorbs3 |
| nucb2b | 0.00678 | 0.3768711 | 0.134 | 0.118 | 1 | Stroma 1 | nucb2b |
| fto | 0.00692 | 0.3880947 | 0.108 | 0.094 | 1 | Stroma 1 | fto |

| | | | | | | | |
|---|---|---|---|---|---|---|---|
| RALGDS1 | 0.00727 | 0.3796531 | 0.149 | 0.133 | 1 | Stroma 1 | RALGDS |
| sntb2 | 0.00755 | 0.2887135 | 0.251 | 0.237 | 1 | Stroma 1 | sntb2 |
| fbxl6 | 0.00802 | 0.4567742 | 0.108 | 0.094 | 1 | Stroma 1 | fbxl6 |
| fzd7b | 0.00814 | 0.433994 | 0.124 | 0.109 | 1 | Stroma 1 | fzd7b |
| snx17 | 0.00842 | 0.2832211 | 0.257 | 0.243 | 1 | Stroma 1 | snx17 |
| cnpy3 | 0.00898 | 0.4817487 | 0.102 | 0.088 | 1 | Stroma 1 | cnpy3 |
| hmgn31 | 0.00899 | 0.2126721 | 0.179 | 0.162 | 1 | Stroma 1 | hmgn3 |
| mapkapk2a | 0.00937 | 0.2590885 | 0.238 | 0.222 | 1 | Stroma 1 | mapkapk2a |
| stom | 0.00946 | 0.2511146 | 0.282 | 0.27 | 1 | Stroma 1 | stom |
| porb1 | 0.00965 | 0.3035783 | 0.123 | 0.108 | 1 | Stroma 1 | porb |
| notch2 | 0.00981 | 0.2236017 | 0.387 | 0.379 | 1 | Stroma 1 | notch2 |
| txn1 | 0 | 2.3507416 | 0.868 | 0.342 | 0 | Regenerative Mesenchyme | txn |
| vmp11 | 0 | 2.8602371 | 0.753 | 0.292 | 0 | Regenerative Mesenchyme | vmp1 |
| hmga1a | 0 | 2.9777499 | 0.585 | 0.163 | 0 | Regenerative Mesenchyme | hmga1a |
| snrpg1 | 0 | 2.6267132 | 0.579 | 0.159 | 0 | Regenerative Mesenchyme | snrpg |
| snrpd1 | 0 | 2.5459821 | 0.579 | 0.173 | 0 | Regenerative Mesenchyme | snrpd1 |
| snrpf | 0 | 2.2559541 | 0.692 | 0.291 | 0 | Regenerative Mesenchyme | snrpf |
| tuba1b1 | 0 | 1.8819898 | 0.684 | 0.286 | 0 | Regenerative Mesenchyme | tuba1b |
| atp5mc11 | 0 | 1.9950596 | 0.683 | 0.288 | 0 | Regenerative Mesenchyme | atp5mc1 |
| npm1a1 | 0 | 2.5262311 | 0.732 | 0.337 | 0 | Regenerative Mesenchyme | npm1a |
| snrpe | 0 | 2.2408403 | 0.612 | 0.225 | 0 | Regenerative Mesenchyme | snrpe |
| cycsb | 0 | 2.3869331 | 0.738 | 0.352 | 0 | Regenerative Mesenchyme | cycsb |
| adamtsl7 | 0 | 2.7635043 | 0.506 | 0.124 | 0 | Regenerative Mesenchyme | adamtsl7 |
| cox7a2a1 | 0 | 2.2439139 | 0.63 | 0.248 | 0 | Regenerative Mesenchyme | cox7a2a |
| cygb12 | 0 | 1.3628904 | 0.772 | 0.401 | 0 | Regenerative Mesenchyme | cygb1 |
| h2az2b | 0 | 1.9748431 | 0.746 | 0.376 | 0 | Regenerative Mesenchyme | h2az2b |
| tma7 | 0 | 1.9040822 | 0.712 | 0.354 | 0 | Regenerative Mesenchyme | tma7 |
| pdap1a1 | 0 | 2.1850809 | 0.552 | 0.199 | 0 | Regenerative Mesenchyme | pdap1a |
| hspe1 | 0 | 2.0464622 | 0.627 | 0.274 | 0 | Regenerative Mesenchyme | hspe1 |
| psmb1 | 0 | 1.9887413 | 0.595 | 0.243 | 0 | Regenerative Mesenchyme | psmb1 |
| snrpb | 0 | 1.8879508 | 0.629 | 0.28 | 0 | Regenerative Mesenchyme | snrpb |
| calr3b1 | 0 | 1.6429866 | 0.598 | 0.25 | 0 | Regenerative Mesenchyme | calr3b |
| tubb5 | 0 | 2.4722877 | 0.475 | 0.13 | 0 | Regenerative Mesenchyme | tubb5 |
| si:ch211-222l21.1 | 0 | 1.0983411 | 0.879 | 0.535 | 0 | Regenerative Mesenchyme | si:ch211-222l21.1 |
| tubb2b1 | 0 | 1.7393947 | 0.664 | 0.322 | 0 | Regenerative Mesenchyme | tubb2b |
| abracl | 0 | 2.6866467 | 0.431 | 0.09 | 0 | Regenerative Mesenchyme | abracl |
| snu13b1 | 0 | 2.4411953 | 0.503 | 0.162 | 0 | Regenerative Mesenchyme | snu13b |
| psmb7 | 0 | 1.9603546 | 0.576 | 0.235 | 0 | Regenerative Mesenchyme | psmb7 |
| fn1a1 | 0 | 1.8100379 | 0.638 | 0.298 | 0 | Regenerative Mesenchyme | fn1a |
| manf | 0 | 2.1060895 | 0.505 | 0.166 | 0 | Regenerative Mesenchyme | manf |
| snrpd2 | 0 | 2.0532545 | 0.52 | 0.182 | 0 | Regenerative Mesenchyme | snrpd2 |
| cox5ab1 | 0 | 1.9368041 | 0.539 | 0.201 | 0 | Regenerative Mesenchyme | cox5ab |
| abcf1 | 0 | 1.7137375 | 0.628 | 0.29 | 0 | Regenerative Mesenchyme | abcf1 |
| tnfaip62 | 0 | 1.7445983 | 0.748 | 0.412 | 0 | Regenerative Mesenchyme | tnfaip6 |
| atp5if1a | 0 | 2.0287449 | 0.642 | 0.307 | 0 | Regenerative Mesenchyme | atp5if1a |
| mdka1 | 0 | 1.5577173 | 0.835 | 0.503 | 0 | Regenerative Mesenchyme | mdka |
| chaf1a | 0 | 3.9757661 | 0.366 | 0.034 | 0 | Regenerative Mesenchyme | chaf1a |
| fbl1 | 0 | 2.0784862 | 0.531 | 0.201 | 0 | Regenerative Mesenchyme | fbl |

| | | | | | | | |
|---|---|---|---|---|---|---|---|
| nutf2l1 | 0 | 2.9139162 | 0.411 | 0.082 | 0 | Regenerative Mesenchyme | nutf2l |
| eif5a2 | 0 | 1.6674688 | 0.777 | 0.449 | 0 | Regenerative Mesenchyme | eif5a2 |
| hmgb2b | 0 | 1.9461896 | 0.671 | 0.345 | 0 | Regenerative Mesenchyme | hmgb2b |
| pomp | 0 | 1.6972915 | 0.647 | 0.323 | 0 | Regenerative Mesenchyme | pomp |
| dad11 | 0 | 1.5448061 | 0.705 | 0.381 | 0 | Regenerative Mesenchyme | dad1 |
| ptges3b | 0 | 2.30272 | 0.447 | 0.124 | 0 | Regenerative Mesenchyme | ptges3b |
| hnrnpa0b | 0 | 1.5214031 | 0.707 | 0.384 | 0 | Regenerative Mesenchyme | hnrnpa0b |
| hsp90b1 | 0 | 1.6249323 | 0.649 | 0.327 | 0 | Regenerative Mesenchyme | hsp90b1 |
| denr | 0 | 1.8924013 | 0.525 | 0.205 | 0 | Regenerative Mesenchyme | denr |
| tomm6 | 0 | 2.2158279 | 0.461 | 0.143 | 0 | Regenerative Mesenchyme | tomm6 |
| hmgb2a | 0 | 2.0078591 | 0.744 | 0.427 | 0 | Regenerative Mesenchyme | hmgb2a |
| psma3 | 0 | 1.7784993 | 0.545 | 0.231 | 0 | Regenerative Mesenchyme | psma3 |
| psmb3 | 0 | 1.7897069 | 0.547 | 0.238 | 0 | Regenerative Mesenchyme | psmb3 |
| pfdn4 | 0 | 1.80507 | 0.548 | 0.24 | 0 | Regenerative Mesenchyme | pfdn4 |
| cox7b1 | 0 | 1.5052415 | 0.684 | 0.379 | 0 | Regenerative Mesenchyme | cox7b |
| dbi | 0 | 1.369646 | 0.729 | 0.425 | 0 | Regenerative Mesenchyme | dbi |
| uqcr10 | 0 | 1.8160105 | 0.524 | 0.221 | 0 | Regenerative Mesenchyme | uqcr10 |
| hint11 | 0 | 1.4397283 | 0.671 | 0.369 | 0 | Regenerative Mesenchyme | hint1 |
| atp5pf1 | 0 | 1.6731021 | 0.687 | 0.388 | 0 | Regenerative Mesenchyme | atp5pf |
| atp5mf | 0 | 1.5506897 | 0.65 | 0.355 | 0 | Regenerative Mesenchyme | atp5mf |
| pcna | 0 | 2.965034 | 0.389 | 0.094 | 0 | Regenerative Mesenchyme | pcna |
| hyou1 | 0 | 2.0102277 | 0.427 | 0.136 | 0 | Regenerative Mesenchyme | hyou1 |
| lsm6 | 0 | 1.9028586 | 0.445 | 0.156 | 0 | Regenerative Mesenchyme | lsm6 |
| cox6a11 | 0 | 1.434908 | 0.764 | 0.475 | 0 | Regenerative Mesenchyme | cox6a1 |
| cmpk | 0 | 1.876951 | 0.431 | 0.148 | 0 | Regenerative Mesenchyme | cmpk |
| odc11 | 0 | 2.0945709 | 0.388 | 0.108 | 0 | Regenerative Mesenchyme | odc1 |
| marcksb1 | 0 | 2.4073253 | 0.362 | 0.085 | 0 | Regenerative Mesenchyme | marcksb |
| fosl1b | 0 | 4.4695194 | 0.292 | 0.016 | 0 | Regenerative Mesenchyme | fosl1b |
| tomm20a.11 | 0 | 1.9631025 | 0.416 | 0.145 | 0 | Regenerative Mesenchyme | tomm20a.1 |
| atp5fa1 | 0 | 1.4737881 | 0.714 | 0.446 | 0 | Regenerative Mesenchyme | atp5fa1 |
| banf11 | 0 | 2.1312852 | 0.371 | 0.113 | 0 | Regenerative Mesenchyme | banf1 |
| magoh | 0 | 2.0084511 | 0.374 | 0.117 | 0 | Regenerative Mesenchyme | magoh |
| rpa3 | 0 | 3.4474767 | 0.295 | 0.038 | 0 | Regenerative Mesenchyme | rpa3 |
| COX5B.11 | 0 | 1.506461 | 0.784 | 0.527 | 0 | Regenerative Mesenchyme | COX5B.1 |
| s100u1 | 0 | 1.5090579 | 0.759 | 0.504 | 0 | Regenerative Mesenchyme | s100u |
| phf5a | 0 | 2.0340133 | 0.35 | 0.096 | 0 | Regenerative Mesenchyme | phf5a |
| fen1 | 0 | 3.641525 | 0.281 | 0.031 | 0 | Regenerative Mesenchyme | fen1 |
| mrpl12 | 0 | 2.0546143 | 0.358 | 0.109 | 0 | Regenerative Mesenchyme | mrpl12 |
| rbbp4 | 0 | 2.33327 | 0.324 | 0.079 | 0 | Regenerative Mesenchyme | rbbp4 |
| rpa2 | 0 | 3.320643 | 0.269 | 0.034 | 0 | Regenerative Mesenchyme | rpa2 |
| cirbpa | 0 | 1.473428 | 0.881 | 0.648 | 0 | Regenerative Mesenchyme | cirbpa |
| atp5f1b1 | 0 | 1.3728625 | 0.772 | 0.545 | 0 | Regenerative Mesenchyme | atp5f1b |
| rpl22l12 | 0 | 1.356868 | 0.96 | 0.735 | 0 | Regenerative Mesenchyme | rpl22l1 |
| si:ch211-156b7.4 | 0 | 3.1019116 | 0.264 | 0.041 | 0 | Regenerative Mesenchyme | si:ch211-156b7.4 |
| atp5mc3b1 | 0 | 1.3244611 | 0.832 | 0.61 | 0 | Regenerative Mesenchyme | atp5mc3b |
| cox8a1 | 0 | 1.3061105 | 0.771 | 0.551 | 0 | Regenerative Mesenchyme | cox8a |
| dut | 0 | 3.5000079 | 0.256 | 0.04 | 0 | Regenerative Mesenchyme | dut |
| mcm2 | 0 | 4.0627379 | 0.233 | 0.018 | 0 | Regenerative Mesenchyme | mcm2 |

| | | | | | | | |
|---|---|---|---|---|---|---|---|
| atp5l | 0 | 1.2496331 | 0.795 | 0.582 | 0 | Regenerative Mesenchyme | atp5l |
| mcm6 | 0 | 3.9006062 | 0.228 | 0.02 | 0 | Regenerative Mesenchyme | mcm6 |
| nasp | 0 | 2.8464611 | 0.25 | 0.043 | 0 | Regenerative Mesenchyme | nasp |
| tubb6 | 0 | 3.5554785 | 0.229 | 0.022 | 0 | Regenerative Mesenchyme | tubb6 |
| mcm3 | 0 | 4.2385257 | 0.219 | 0.014 | 0 | Regenerative Mesenchyme | mcm3 |
| ssrp1a | 0 | 2.6381049 | 0.249 | 0.046 | 0 | Regenerative Mesenchyme | ssrp1a |
| mcm5 | 0 | 4.0365426 | 0.218 | 0.018 | 0 | Regenerative Mesenchyme | mcm5 |
| mki67 | 0 | 5.1700378 | 0.207 | 0.01 | 0 | Regenerative Mesenchyme | mki67 |
| tnfrsf9a1 | 0 | 3.1480685 | 0.235 | 0.038 | 0 | Regenerative Mesenchyme | tnfrsf9a |
| khdrbs1a | 0 | 1.1832941 | 0.837 | 0.642 | 0 | Regenerative Mesenchyme | khdrbs1a |
| lig1 | 0 | 3.2526172 | 0.221 | 0.027 | 0 | Regenerative Mesenchyme | lig1 |
| ppil1 | 0 | 2.8012752 | 0.224 | 0.034 | 0 | Regenerative Mesenchyme | ppil1 |
| yars1 | 0 | 2.4210174 | 0.244 | 0.054 | 0 | Regenerative Mesenchyme | yars1 |
| mcm4 | 0 | 3.9565106 | 0.2 | 0.015 | 0 | Regenerative Mesenchyme | mcm4 |
| jpt1b | 0 | 1.2512788 | 0.865 | 0.681 | 0 | Regenerative Mesenchyme | jpt1b |
| pclaf | 0 | 4.3022955 | 0.197 | 0.016 | 0 | Regenerative Mesenchyme | pclaf |
| hells | 0 | 4.4143582 | 0.191 | 0.011 | 0 | Regenerative Mesenchyme | hells |
| cenpx | 0 | 2.5243671 | 0.222 | 0.045 | 0 | Regenerative Mesenchyme | cenpx |
| TXN | 0 | 1.2285496 | 0.903 | 0.738 | 0 | Regenerative Mesenchyme | TXN |
| CABZ01005379.1 | 0 | 4.1360418 | 0.175 | 0.012 | 0 | Regenerative Mesenchyme | CABZ01005379.1 |
| tuba8l41 | 0 | 1.0900846 | 0.885 | 0.724 | 0 | Regenerative Mesenchyme | tuba8l4 |
| tyms | 0 | 5.0640619 | 0.159 | 0.006 | 0 | Regenerative Mesenchyme | tyms |
| uhrf1 | 0 | 4.0624129 | 0.161 | 0.011 | 0 | Regenerative Mesenchyme | uhrf1 |
| tnfrsf11b | 0 | 3.1876669 | 0.172 | 0.023 | 0 | Regenerative Mesenchyme | tnfrsf11b |
| CKS2 | 0 | 2.7855012 | 0.171 | 0.026 | 0 | Regenerative Mesenchyme | CKS2 |
| cfl1 | 0 | 1.3391267 | 0.971 | 0.836 | 0 | Regenerative Mesenchyme | cfl1 |
| ing5b | 0 | 3.3742986 | 0.147 | 0.015 | 0 | Regenerative Mesenchyme | ing5b |
| cks1b | 0 | 4.5088563 | 0.138 | 0.008 | 0 | Regenerative Mesenchyme | cks1b |
| CABZ01058261.1 | 0 | 5.6623512 | 0.133 | 0.004 | 0 | Regenerative Mesenchyme | CABZ01058261.1 |
| smc2 | 0 | 4.992177 | 0.133 | 0.005 | 0 | Regenerative Mesenchyme | smc2 |
| top2a | 0 | 6.136106 | 0.129 | 0.003 | 0 | Regenerative Mesenchyme | top2a |
| rrm1 | 0 | 3.1411702 | 0.143 | 0.018 | 0 | Regenerative Mesenchyme | rrm1 |
| smc4 | 0 | 4.1645331 | 0.131 | 0.01 | 0 | Regenerative Mesenchyme | smc4 |
| ube2c.1 | 0 | 5.5941855 | 0.125 | 0.004 | 0 | Regenerative Mesenchyme | ube2c.1 |
| atad2 | 0 | 3.382883 | 0.13 | 0.013 | 0 | Regenerative Mesenchyme | atad2 |
| kpna2 | 0 | 5.2251365 | 0.123 | 0.006 | 0 | Regenerative Mesenchyme | kpna2 |
| cdk1 | 0 | 6.4851162 | 0.118 | 0.002 | 0 | Regenerative Mesenchyme | cdk1 |
| rrm2.1 | 0 | 4.4968169 | 0.123 | 0.008 | 0 | Regenerative Mesenchyme | rrm2.1 |
| si:dkey-30c15.10 | 0 | 3.9561061 | 0.122 | 0.01 | 0 | Regenerative Mesenchyme | si:dkey-30c15.10 |
| actb11 | 0 | 0.9609934 | 0.97 | 0.86 | 0 | Regenerative Mesenchyme | actb1 |
| birc5a | 0 | 4.054028 | 0.117 | 0.009 | 0 | Regenerative Mesenchyme | birc5a |
| rrm2 | 0 | 4.5646256 | 0.114 | 0.006 | 0 | Regenerative Mesenchyme | rrm2 |
| f8 | 0 | 3.8413827 | 0.112 | 0.009 | 0 | Regenerative Mesenchyme | f8 |
| dnph1 | 0 | 3.7698211 | 0.112 | 0.011 | 0 | Regenerative Mesenchyme | dnph1 |
| orc4 | 0 | 3.6446971 | 0.109 | 0.011 | 0 | Regenerative Mesenchyme | orc4 |
| tpx2 | 0 | 5.0173324 | 0.101 | 0.005 | 0 | Regenerative Mesenchyme | tpx2 |
| nme2b.12 | 0 | 0.9167423 | 0.978 | 0.915 | 0 | Regenerative Mesenchyme | nme2b.1 |
| mt-atp6 | 0 | 1.0508774 | 0.994 | 0.964 | 0 | Regenerative Mesenchyme | mt-atp6 |

| | | | | | | | |
|---|---|---|---|---|---|---|---|
| rps202 | 0 | 0.6255394 | 0.998 | 0.985 | 0 | Regenerative Mesenchyme | rps20 |
| rplp2l2 | 0 | 0.8704954 | 0.998 | 0.986 | 0 | Regenerative Mesenchyme | rplp2l |
| tuba8l1 | 2E-307 | 1.5424422 | 0.684 | 0.393 | 7.8E-303 | Regenerative Mesenchyme | tuba8l |
| ran | 3E-305 | 1.0903565 | 0.847 | 0.69 | 9E-301 | Regenerative Mesenchyme | ran |
| ppa1b1 | 4E-304 | 1.9848763 | 0.381 | 0.124 | 1.1E-299 | Regenerative Mesenchyme | ppa1b |
| bxdc2 | 1E-303 | 2.2303607 | 0.295 | 0.078 | 3.5E-299 | Regenerative Mesenchyme | bxdc2 |
| uqcrh1 | 3E-303 | 1.6171291 | 0.643 | 0.369 | 1E-298 | Regenerative Mesenchyme | uqcrh |
| chchd10 | 5E-303 | 2.8911143 | 0.155 | 0.023 | 1.5E-298 | Regenerative Mesenchyme | chchd10 |
| eif4e1c | 6E-303 | 1.9371725 | 0.391 | 0.129 | 2E-298 | Regenerative Mesenchyme | eif4e1c |
| dkc11 | 1E-301 | 1.8979744 | 0.454 | 0.174 | 3.8E-297 | Regenerative Mesenchyme | dkc1 |
| hspa51 | 2E-301 | 1.3820532 | 0.759 | 0.476 | 5.4E-297 | Regenerative Mesenchyme | hspa5 |
| dnajc9 | 1E-299 | 2.5489351 | 0.22 | 0.046 | 4E-295 | Regenerative Mesenchyme | dnajc9 |
| dynll11 | 3E-299 | 1.396574 | 0.707 | 0.409 | 9.5E-295 | Regenerative Mesenchyme | dynll1 |
| ilf2 | 8E-299 | 1.9232234 | 0.413 | 0.145 | 2.6E-294 | Regenerative Mesenchyme | ilf2 |
| ndufab1a | 6E-298 | 1.9583162 | 0.362 | 0.114 | 2.1E-293 | Regenerative Mesenchyme | ndufab1a |
| ssr32 | 2E-293 | 1.2666911 | 0.796 | 0.565 | 8E-289 | Regenerative Mesenchyme | ssr3 |
| ddx39ab | 2E-293 | 1.8916359 | 0.425 | 0.153 | 8E-289 | Regenerative Mesenchyme | ddx39ab |
| dnajb111 | 4E-293 | 2.0192006 | 0.322 | 0.092 | 1.1E-288 | Regenerative Mesenchyme | dnajb11 |
| eif1b | 6E-292 | 1.7847104 | 0.477 | 0.193 | 2E-287 | Regenerative Mesenchyme | eif1b |
| timm8b | 1E-291 | 2.0854031 | 0.318 | 0.091 | 4E-287 | Regenerative Mesenchyme | timm8b |
| got2b1 | 4E-291 | 2.338363 | 0.284 | 0.076 | 1.4E-286 | Regenerative Mesenchyme | got2b |
| atp5mc3a | 1E-290 | 1.6436836 | 0.551 | 0.254 | 3.2E-286 | Regenerative Mesenchyme | atp5mc3a |
| glipr2l | 5E-290 | 2.6949511 | 0.179 | 0.031 | 1.5E-285 | Regenerative Mesenchyme | glipr2l |
| cox5aa1 | 1E-289 | 1.7195984 | 0.559 | 0.27 | 3.5E-285 | Regenerative Mesenchyme | cox5aa |
| psma8 | 3E-289 | 1.6701455 | 0.531 | 0.24 | 1.1E-284 | Regenerative Mesenchyme | psma8 |
| rbx1 | 8E-289 | 1.3574521 | 0.698 | 0.439 | 2.8E-284 | Regenerative Mesenchyme | rbx1 |
| cgref11 | 8E-285 | 1.7814717 | 0.427 | 0.157 | 2.7E-280 | Regenerative Mesenchyme | cgref1 |
| stmn1a | 9E-284 | 3.0979593 | 0.168 | 0.029 | 2.8E-279 | Regenerative Mesenchyme | stmn1a |
| golt1bb1 | 3E-281 | 2.0463639 | 0.31 | 0.089 | 8.2E-277 | Regenerative Mesenchyme | golt1bb |
| hypk | 1E-280 | 1.6334326 | 0.559 | 0.268 | 4.1E-276 | Regenerative Mesenchyme | hypk |
| msh2 | 3E-279 | 3.1244733 | 0.117 | 0.014 | 1.1E-274 | Regenerative Mesenchyme | msh2 |
| ppih | 4E-279 | 2.3087928 | 0.221 | 0.048 | 1.3E-274 | Regenerative Mesenchyme | ppih |
| psmd1 | 8E-279 | 1.7433577 | 0.437 | 0.165 | 2.6E-274 | Regenerative Mesenchyme | psmd1 |
| ddost | 1E-276 | 1.6707541 | 0.494 | 0.208 | 3.6E-272 | Regenerative Mesenchyme | ddost |
| mt-nd5 | 1E-276 | 1.3009351 | 0.755 | 0.474 | 4E-272 | Regenerative Mesenchyme | mt-nd5 |
| eif2s2 | 1E-276 | 1.3145477 | 0.71 | 0.449 | 4.7E-272 | Regenerative Mesenchyme | eif2s2 |
| slirp | 4E-276 | 1.8607323 | 0.378 | 0.129 | 1.2E-271 | Regenerative Mesenchyme | slirp |
| si:ch1073-325m22.2 | 3E-275 | 1.5591497 | 0.573 | 0.286 | 1E-270 | Regenerative Mesenchyme | si:ch1073-325m22. |
| hars1 | 6E-275 | 2.1275148 | 0.283 | 0.077 | 2.1E-270 | Regenerative Mesenchyme | hars |
| prdx2 | 2E-274 | 1.5268063 | 0.607 | 0.313 | 7E-270 | Regenerative Mesenchyme | prdx2 |
| hmgn2 | 1E-273 | 2.1942952 | 0.761 | 0.554 | 3.7E-269 | Regenerative Mesenchyme | hmgn2 |
| gapdhs | 2E-273 | 1.1255675 | 0.814 | 0.619 | 6.1E-269 | Regenerative Mesenchyme | gapdhs |
| sae1 | 3E-272 | 2.8266037 | 0.148 | 0.023 | 8.7E-268 | Regenerative Mesenchyme | sae1 |
| selenom2 | 5E-272 | 1.7861676 | 0.419 | 0.157 | 1.5E-267 | Regenerative Mesenchyme | selenom |
| tuba8l3 | 6E-271 | 1.4238887 | 0.677 | 0.41 | 2E-266 | Regenerative Mesenchyme | tuba8l3 |
| nop58 | 5E-270 | 1.7058875 | 0.55 | 0.272 | 1.8E-265 | Regenerative Mesenchyme | nop58 |
| eny2 | 1E-269 | 1.9086774 | 0.318 | 0.095 | 3.2E-265 | Regenerative Mesenchyme | eny2 |
| pola1 | 8E-268 | 3.2421506 | 0.107 | 0.012 | 2.5E-263 | Regenerative Mesenchyme | pola1 |

| | | | | | | | |
|---|---|---|---|---|---|---|---|
| ufm11 | 7E-267 | 1.6667479 | 0.443 | 0.174 | 2.2E-262 | Regenerative Mesenchyme | ufm1 |
| cthrc1a1 | 2E-264 | 1.6488335 | 0.394 | 0.139 | 5E-260 | Regenerative Mesenchyme | cthrc1a |
| rpl112 | 3E-264 | 0.5260116 | 0.996 | 0.988 | 1E-259 | Regenerative Mesenchyme | rpl11 |
| arpc5a | 4E-264 | 1.7445423 | 0.394 | 0.14 | 1.2E-259 | Regenerative Mesenchyme | arpc5a |
| zgc:110425 | 2E-262 | 4.1086034 | 0.105 | 0.012 | 5.1E-258 | Regenerative Mesenchyme | zgc:110425 |
| si:ch211-156j16.11 | 3E-262 | 1.1680956 | 0.4 | 0.138 | 8.4E-258 | Regenerative Mesenchyme | si:ch211-156j16.1 |
| asph | 9E-262 | 1.5038591 | 0.643 | 0.362 | 2.8E-257 | Regenerative Mesenchyme | asph |
| mrto41 | 5E-259 | 2.2301268 | 0.226 | 0.053 | 1.7E-254 | Regenerative Mesenchyme | mrto4 |
| ppiaa1 | 1E-258 | 0.5809424 | 0.99 | 0.958 | 4.6E-254 | Regenerative Mesenchyme | ppiaa |
| si:ch211-11k18.4 | 2E-258 | 2.3096994 | 0.216 | 0.049 | 8.1E-254 | Regenerative Mesenchyme | si:ch211-11k18.4 |
| cbx3a | 2E-257 | 1.4773472 | 0.582 | 0.299 | 6.2E-253 | Regenerative Mesenchyme | cbx3a |
| rcn11 | 6E-257 | 1.9168189 | 0.348 | 0.116 | 2E-252 | Regenerative Mesenchyme | rcn1 |
| fkbp2 | 2E-256 | 1.635821 | 0.457 | 0.188 | 7E-252 | Regenerative Mesenchyme | fkbp2 |
| hnrnpaba | 1E-255 | 1.0465118 | 0.796 | 0.593 | 3.4E-251 | Regenerative Mesenchyme | hnrnpaba |
| psmb6 | 4E-255 | 1.5273676 | 0.572 | 0.299 | 1.2E-250 | Regenerative Mesenchyme | psmb6 |
| kpnb3 | 1E-254 | 1.8715203 | 0.373 | 0.132 | 4.6E-250 | Regenerative Mesenchyme | kpnb3 |
| aldoaa1 | 3E-254 | 1.4848889 | 0.624 | 0.353 | 1E-249 | Regenerative Mesenchyme | aldoaa |
| mcts1 | 4E-254 | 1.6858618 | 0.41 | 0.157 | 1.2E-249 | Regenerative Mesenchyme | mcts1 |
| cdca7b | 4E-254 | 3.0483524 | 0.121 | 0.016 | 1.3E-249 | Regenerative Mesenchyme | cdca7b |
| tmem2581 | 1E-253 | 1.1381821 | 0.778 | 0.577 | 4E-249 | Regenerative Mesenchyme | tmem258 |
| dhfr | 1E-251 | 2.8069989 | 0.129 | 0.019 | 4.9E-247 | Regenerative Mesenchyme | dhfr |
| anp32b | 3E-251 | 1.6326218 | 0.459 | 0.191 | 8.7E-247 | Regenerative Mesenchyme | anp32b |
| snrpd3l | 4E-250 | 1.7479304 | 0.337 | 0.11 | 1.3E-245 | Regenerative Mesenchyme | snrpd3l |
| tubb4b1 | 1E-249 | 0.9038385 | 0.899 | 0.804 | 3.7E-245 | Regenerative Mesenchyme | tubb4b |
| kdelr2b1 | 1E-248 | 1.6154937 | 0.57 | 0.305 | 4.9E-244 | Regenerative Mesenchyme | kdelr2b |
| atp5f1d1 | 2E-247 | 1.5039355 | 0.585 | 0.319 | 5.3E-243 | Regenerative Mesenchyme | atp5f1d |
| thop1 | 2E-247 | 2.1906903 | 0.205 | 0.046 | 5.6E-243 | Regenerative Mesenchyme | thop1 |
| nap1l1 | 4E-247 | 1.2663169 | 0.731 | 0.514 | 1.2E-242 | Regenerative Mesenchyme | nap1l1 |
| lsm8 | 7E-247 | 1.9848619 | 0.263 | 0.072 | 2.2E-242 | Regenerative Mesenchyme | lsm8 |
| rtn4a | 2E-245 | 1.2672239 | 0.689 | 0.436 | 7.2E-241 | Regenerative Mesenchyme | rtn4a |
| bcam | 1E-243 | 3.3221559 | 0.111 | 0.015 | 3.5E-239 | Regenerative Mesenchyme | bcam |
| hnrnpabb | 2E-243 | 1.2889375 | 0.698 | 0.466 | 6.8E-239 | Regenerative Mesenchyme | hnrnpabb |
| micos10 | 3E-243 | 1.7308857 | 0.405 | 0.157 | 9.2E-239 | Regenerative Mesenchyme | micos10 |
| psmb4 | 3E-242 | 1.5219308 | 0.495 | 0.223 | 9.2E-238 | Regenerative Mesenchyme | psmb4 |
| ssb | 3E-242 | 1.8780641 | 0.348 | 0.121 | 9.5E-238 | Regenerative Mesenchyme | ssb |
| ywhaqa | 2E-241 | 1.4608075 | 0.537 | 0.257 | 5E-237 | Regenerative Mesenchyme | ywhaqa |
| atp5pd1 | 2E-239 | 1.3382111 | 0.618 | 0.353 | 5.5E-235 | Regenerative Mesenchyme | atp5pd |
| phb | 1E-238 | 1.8404624 | 0.335 | 0.114 | 3.7E-234 | Regenerative Mesenchyme | phb |
| c1qbp1 | 3E-238 | 1.6807919 | 0.461 | 0.206 | 1.1E-233 | Regenerative Mesenchyme | c1qbp |
| mt-nd1 | 6E-238 | 1.0065937 | 0.885 | 0.708 | 2E-233 | Regenerative Mesenchyme | mt-nd1 |
| eif4a1a1 | 9E-238 | 1.3905051 | 0.623 | 0.35 | 2.8E-233 | Regenerative Mesenchyme | eif4a1a |
| aimp1a | 1E-237 | 1.7872494 | 0.36 | 0.129 | 4.3E-233 | Regenerative Mesenchyme | aimp1a |
| ahcy2 | 3E-236 | 1.6399015 | 0.422 | 0.17 | 1.1E-231 | Regenerative Mesenchyme | ahcy |
| psmd7 | 1E-235 | 1.4964125 | 0.501 | 0.231 | 3.4E-231 | Regenerative Mesenchyme | psmd7 |
| polr2h | 1E-235 | 2.5603433 | 0.137 | 0.022 | 4.7E-231 | Regenerative Mesenchyme | polr2h |
| gadd45gip1 | 9E-235 | 2.0086476 | 0.214 | 0.051 | 2.8E-230 | Regenerative Mesenchyme | gadd45gip1 |
| dnmt1 | 1E-234 | 2.774812 | 0.127 | 0.02 | 3.1E-230 | Regenerative Mesenchyme | dnmt1 |
| rnaseh2a | 2E-234 | 2.6228773 | 0.159 | 0.031 | 6.8E-230 | Regenerative Mesenchyme | rnaseh2a |

| | | | | | | | |
|---|---|---|---|---|---|---|---|
| ywhaqb | 9E-233 | 1.0024492 | 0.759 | 0.524 | 3E-228 | Regenerative Mesenchyme | ywhaqb |
| rps172 | 1E-232 | 0.5590792 | 0.979 | 0.943 | 4.1E-228 | Regenerative Mesenchyme | rps17 |
| pmm2 | 5E-231 | 2.5010571 | 0.158 | 0.03 | 1.8E-226 | Regenerative Mesenchyme | pmm2 |
| timm10 | 8E-231 | 2.039003 | 0.232 | 0.06 | 2.5E-226 | Regenerative Mesenchyme | timm10 |
| CT027638.1 | 1E-230 | 0.7927653 | 1 | 0.995 | 3.7E-226 | Regenerative Mesenchyme | CT027638.1 |
| rfc4 | 1E-230 | 2.8823492 | 0.119 | 0.018 | 4.1E-226 | Regenerative Mesenchyme | rfc4 |
| CABZ01045617.1 | 1E-230 | 3.2248994 | 0.104 | 0.013 | 4.6E-226 | Regenerative Mesenchyme | CABZ01045617.1 |
| fkbp91 | 2E-229 | 1.3817791 | 0.63 | 0.369 | 6E-225 | Regenerative Mesenchyme | fkbp9 |
| cct41 | 7E-229 | 1.5104318 | 0.558 | 0.298 | 2.2E-224 | Regenerative Mesenchyme | cct4 |
| mrpl27 | 6E-228 | 2.0946851 | 0.208 | 0.05 | 2.1E-223 | Regenerative Mesenchyme | mrpl27 |
| slc39a71 | 7E-228 | 1.8580793 | 0.316 | 0.105 | 2.3E-223 | Regenerative Mesenchyme | slc39a7 |
| msna | 6E-227 | 0.7794083 | 0.884 | 0.743 | 2.1E-222 | Regenerative Mesenchyme | msna |
| uchl3 | 5E-226 | 1.8574471 | 0.302 | 0.097 | 1.5E-221 | Regenerative Mesenchyme | uchl3 |
| quo | 6E-226 | 1.8854329 | 0.226 | 0.058 | 2E-221 | Regenerative Mesenchyme | quo |
| ebna1bp2 | 2E-225 | 1.6863526 | 0.355 | 0.129 | 6.2E-221 | Regenerative Mesenchyme | ebna1bp2 |
| lsm3 | 7E-225 | 1.5574274 | 0.447 | 0.192 | 2.4E-220 | Regenerative Mesenchyme | lsm3 |
| abce1 | 2E-224 | 1.7483447 | 0.325 | 0.111 | 5.2E-220 | Regenerative Mesenchyme | abce1 |
| psma5 | 3E-224 | 1.6604821 | 0.425 | 0.18 | 9.1E-220 | Regenerative Mesenchyme | psma5 |
| psma6a | 3E-224 | 1.7180315 | 0.338 | 0.118 | 1.1E-219 | Regenerative Mesenchyme | psma6a |
| sec61b1 | 6E-224 | 1.2376318 | 0.752 | 0.573 | 2.1E-219 | Regenerative Mesenchyme | sec61b |
| ranbp11 | 1E-223 | 1.419946 | 0.588 | 0.335 | 3.1E-219 | Regenerative Mesenchyme | ranbp1 |
| cnbpa | 4E-223 | 1.4100537 | 0.59 | 0.334 | 1.2E-218 | Regenerative Mesenchyme | cnbpa |
| polr2l | 4E-223 | 1.61616 | 0.428 | 0.18 | 1.2E-218 | Regenerative Mesenchyme | polr2l |
| cox7a2l | 8E-223 | 1.7572947 | 0.374 | 0.145 | 2.4E-218 | Regenerative Mesenchyme | cox7a2l |
| ostc | 9E-222 | 1.4043273 | 0.541 | 0.276 | 3E-217 | Regenerative Mesenchyme | ostc |
| dkk3b1 | 1E-221 | 1.1373553 | 0.509 | 0.232 | 3.8E-217 | Regenerative Mesenchyme | dkk3b |
| taf15 | 7E-221 | 1.5422839 | 0.455 | 0.2 | 2.2E-216 | Regenerative Mesenchyme | taf15 |
| dbn1 | 8E-221 | 1.5476731 | 0.439 | 0.186 | 2.7E-216 | Regenerative Mesenchyme | dbn1 |
| cdv3 | 1E-220 | 1.4264587 | 0.486 | 0.223 | 3.4E-216 | Regenerative Mesenchyme | cdv3 |
| nhp21 | 2E-219 | 1.7883338 | 0.324 | 0.113 | 6.8E-215 | Regenerative Mesenchyme | nhp2 |
| tars1 | 2E-219 | 2.081541 | 0.234 | 0.064 | 7.5E-215 | Regenerative Mesenchyme | tars1 |
| ndufs5 | 4E-219 | 1.5349192 | 0.407 | 0.164 | 1.2E-214 | Regenerative Mesenchyme | ndufs5 |
| cox7c | 3E-218 | 1.4136573 | 0.545 | 0.285 | 9.4E-214 | Regenerative Mesenchyme | cox7c |
| hspd1 | 6E-218 | 1.6505595 | 0.448 | 0.2 | 2E-213 | Regenerative Mesenchyme | hspd1 |
| pfdn6 | 7E-218 | 1.7060661 | 0.347 | 0.127 | 2.2E-213 | Regenerative Mesenchyme | pfdn6 |
| ndufb3 | 1E-217 | 1.408143 | 0.522 | 0.264 | 4.5E-213 | Regenerative Mesenchyme | ndufb3 |
| tomm40l | 1E-217 | 2.4882229 | 0.16 | 0.033 | 4.7E-213 | Regenerative Mesenchyme | tomm40l |
| rwdd1 | 1E-216 | 1.3985234 | 0.51 | 0.248 | 3.2E-212 | Regenerative Mesenchyme | rwdd1 |
| psmd3 | 2E-216 | 1.6783113 | 0.367 | 0.14 | 6.8E-212 | Regenerative Mesenchyme | psmd3 |
| tnfrsf21 | 3E-216 | 2.2725352 | 0.154 | 0.03 | 1.1E-211 | Regenerative Mesenchyme | tnfrsf21 |
| psmd8 | 4E-216 | 1.6816452 | 0.366 | 0.139 | 1.2E-211 | Regenerative Mesenchyme | psmd8 |
| selenof1 | 8E-216 | 1.1579886 | 0.692 | 0.454 | 2.5E-211 | Regenerative Mesenchyme | selenof |
| phb2b | 2E-215 | 2.0124135 | 0.236 | 0.066 | 8.1E-211 | Regenerative Mesenchyme | phb2b |
| timp2b1 | 5E-215 | 1.3073811 | 0.498 | 0.237 | 1.6E-210 | Regenerative Mesenchyme | timp2b |
| uqcrfs1 | 6E-214 | 1.5884835 | 0.453 | 0.207 | 2.1E-209 | Regenerative Mesenchyme | uqcrfs1 |
| sub1a | 2E-213 | 1.3406534 | 0.575 | 0.319 | 6.5E-209 | Regenerative Mesenchyme | sub1a |
| lifrb | 7E-213 | 1.4223289 | 0.436 | 0.185 | 2.3E-208 | Regenerative Mesenchyme | lifrb |
| psme3 | 2E-212 | 1.9232179 | 0.265 | 0.081 | 5.7E-208 | Regenerative Mesenchyme | psme3 |

| | | | | | | | | |
|---|---|---|---|---|---|---|---|---|
| pfn2l | 1E-211 | 1.075929 | 0.703 | 0.464 | 3.5E-207 | Regenerative | Mesenchyme | pfn2l |
| eif4h | 4E-211 | 1.6020807 | 0.428 | 0.187 | 1.2E-206 | Regenerative | Mesenchyme | eif4h |
| mrpl52 | 2E-210 | 2.2812422 | 0.162 | 0.034 | 5.2E-206 | Regenerative | Mesenchyme | mrpl52 |
| fkbp111 | 2E-210 | 1.7998657 | 0.368 | 0.147 | 5.8E-206 | Regenerative | Mesenchyme | fkbp11 |
| bzw1a | 5E-210 | 1.4334265 | 0.565 | 0.302 | 1.6E-205 | Regenerative | Mesenchyme | bzw1a |
| gar11 | 6E-210 | 1.7686282 | 0.289 | 0.095 | 1.8E-205 | Regenerative | Mesenchyme | gar1 |
| ndufa12 | 4E-209 | 1.4202228 | 0.458 | 0.205 | 1.2E-204 | Regenerative | Mesenchyme | ndufa12 |
| zgc:1584632 | 2E-207 | 0.726432 | 0.999 | 0.996 | 8E-203 | Regenerative | Mesenchyme | zgc:158463 |
| tuba1a | 4E-207 | 2.1920466 | 0.255 | 0.079 | 1.3E-202 | Regenerative | Mesenchyme | tuba1a |
| mt-nd2 | 5E-207 | 1.0162659 | 0.829 | 0.619 | 1.6E-202 | Regenerative | Mesenchyme | mt-nd2 |
| psmd13 | 6E-207 | 1.6508297 | 0.347 | 0.13 | 2E-202 | Regenerative | Mesenchyme | psmd13 |
| mrpl11 | 7E-207 | 1.714406 | 0.312 | 0.108 | 2.3E-202 | Regenerative | Mesenchyme | mrpl11 |
| mt-cyb | 1E-206 | 0.6698925 | 0.995 | 0.974 | 4.8E-202 | Regenerative | Mesenchyme | mt-cyb |
| adam8a | 1E-205 | 1.835671 | 0.334 | 0.123 | 3.3E-201 | Regenerative | Mesenchyme | adam8a |
| mrpl141 | 1E-205 | 1.8945272 | 0.227 | 0.063 | 3.9E-201 | Regenerative | Mesenchyme | mrpl14 |
| imp3 | 1E-205 | 2.0276335 | 0.205 | 0.053 | 4.9E-201 | Regenerative | Mesenchyme | imp3 |
| ndufa6 | 4E-205 | 1.4769061 | 0.463 | 0.217 | 1.4E-200 | Regenerative | Mesenchyme | ndufa6 |
| psmc5 | 1E-204 | 1.5186305 | 0.394 | 0.16 | 3.9E-200 | Regenerative | Mesenchyme | psmc5 |
| mrpl33 | 4E-203 | 1.5810325 | 0.364 | 0.141 | 1.4E-198 | Regenerative | Mesenchyme | mrpl33 |
| msh6 | 4E-203 | 2.1469248 | 0.177 | 0.041 | 1.4E-198 | Regenerative | Mesenchyme | msh6 |
| gale | 9E-203 | 2.1477545 | 0.187 | 0.045 | 2.8E-198 | Regenerative | Mesenchyme | gale |
| cd2bp2 | 1E-202 | 1.5609241 | 0.312 | 0.107 | 3.4E-198 | Regenerative | Mesenchyme | cd2bp2 |
| snrpc | 2E-202 | 1.6644572 | 0.282 | 0.092 | 5.5E-198 | Regenerative | Mesenchyme | snrpc |
| prmt1 | 9E-202 | 1.6469076 | 0.409 | 0.176 | 3E-197 | Regenerative | Mesenchyme | prmt1 |
| nop101 | 4E-201 | 1.5759563 | 0.403 | 0.173 | 1.4E-196 | Regenerative | Mesenchyme | nop10 |
| iars1 | 5E-201 | 2.279914 | 0.192 | 0.048 | 1.6E-196 | Regenerative | Mesenchyme | iars1 |
| nutf2 | 1E-200 | 1.6498997 | 0.331 | 0.122 | 3.8E-196 | Regenerative | Mesenchyme | nutf2 |
| fkbp1aa1 | 2E-200 | 1.041139 | 0.722 | 0.493 | 6.7E-196 | Regenerative | Mesenchyme | fkbp1aa |
| sars11 | 3E-200 | 1.713652 | 0.347 | 0.134 | 1.1E-195 | Regenerative | Mesenchyme | sars1 |
| hadhab | 5E-200 | 1.8995893 | 0.215 | 0.058 | 1.7E-195 | Regenerative | Mesenchyme | hadhab |
| h3f3b.11 | 8E-200 | 1.2388858 | 0.715 | 0.482 | 2.6E-195 | Regenerative | Mesenchyme | h3f3b.1 |
| cfl1l1 | 1E-199 | 1.9862386 | 0.346 | 0.136 | 3.2E-195 | Regenerative | Mesenchyme | cfl1l |
| pdia6 | 5E-199 | 1.5181487 | 0.424 | 0.186 | 1.6E-194 | Regenerative | Mesenchyme | pdia6 |
| ndufab1b1 | 6E-199 | 1.5251676 | 0.424 | 0.19 | 1.9E-194 | Regenerative | Mesenchyme | ndufab1b |
| rpl232 | 2E-197 | 0.4279579 | 0.998 | 0.992 | 6.3E-193 | Regenerative | Mesenchyme | rpl23 |
| ppm1g | 3E-197 | 1.6035343 | 0.364 | 0.143 | 8.4E-193 | Regenerative | Mesenchyme | ppm1g |
| has1 | 4E-197 | 2.3292578 | 0.141 | 0.028 | 1.2E-192 | Regenerative | Mesenchyme | has1 |
| canx | 8E-197 | 1.4168291 | 0.487 | 0.237 | 2.5E-192 | Regenerative | Mesenchyme | canx |
| ptrhd1 | 1E-195 | 2.2823416 | 0.166 | 0.038 | 3.3E-191 | Regenerative | Mesenchyme | ptrhd1 |
| znf5931 | 2E-195 | 1.7490928 | 0.285 | 0.096 | 5.8E-191 | Regenerative | Mesenchyme | znf593 |
| pgam1a | 1E-194 | 1.4351258 | 0.407 | 0.174 | 4.3E-190 | Regenerative | Mesenchyme | pgam1a |
| psma4 | 1E-194 | 1.3664559 | 0.519 | 0.276 | 4.3E-190 | Regenerative | Mesenchyme | psma4 |
| cox4i1 | 6E-194 | 0.7394128 | 0.863 | 0.773 | 2.1E-189 | Regenerative | Mesenchyme | cox4i1 |
| sf3b51 | 1E-193 | 1.717337 | 0.274 | 0.09 | 4.1E-189 | Regenerative | Mesenchyme | sf3b5 |
| pa2g4a | 3E-193 | 1.5928779 | 0.398 | 0.173 | 1.1E-188 | Regenerative | Mesenchyme | pa2g4a |
| dpy30 | 7E-193 | 1.5685239 | 0.372 | 0.152 | 2.4E-188 | Regenerative | Mesenchyme | dpy30 |
| bub3 | 6E-192 | 2.2600786 | 0.175 | 0.042 | 2.1E-187 | Regenerative | Mesenchyme | bub3 |
| mrpl57 | 1E-191 | 1.6811622 | 0.33 | 0.125 | 3.2E-187 | Regenerative | Mesenchyme | mrpl57 |

| | | | | | | | |
|---|---|---|---|---|---|---|---|
| txnl4a1 | 1E-191 | 1.7984742 | 0.238 | 0.071 | 3.6E-187 | Regenerative Mesenchyme | txnl4a |
| ssr22 | 2E-191 | 0.9249404 | 0.83 | 0.683 | 4.9E-187 | Regenerative Mesenchyme | ssr2 |
| idh22 | 2E-191 | 1.2481682 | 0.581 | 0.33 | 5.1E-187 | Regenerative Mesenchyme | idh2 |
| rbm8a | 7E-191 | 1.2635028 | 0.58 | 0.34 | 2.1E-186 | Regenerative Mesenchyme | rbm8a |
| jac4 | 1E-190 | 2.6162562 | 0.126 | 0.024 | 3.3E-186 | Regenerative Mesenchyme | jac4 |
| MYG11 | 1E-190 | 1.7590494 | 0.278 | 0.094 | 3.8E-186 | Regenerative Mesenchyme | MYG1 |
| sec11a | 5E-189 | 1.2667213 | 0.573 | 0.329 | 1.6E-184 | Regenerative Mesenchyme | sec11a |
| eif5b | 7E-189 | 1.1625668 | 0.588 | 0.334 | 2.2E-184 | Regenerative Mesenchyme | eif5b |
| rbm4.3 | 7E-189 | 1.4260717 | 0.443 | 0.204 | 2.2E-184 | Regenerative Mesenchyme | rbm4.3 |
| snrpb2 | 2E-188 | 1.7946919 | 0.22 | 0.063 | 5.5E-184 | Regenerative Mesenchyme | snrpb2 |
| polr2eb | 2E-187 | 1.5647668 | 0.351 | 0.139 | 6.3E-183 | Regenerative Mesenchyme | polr2eb |
| trmt1121 | 4E-187 | 1.641393 | 0.306 | 0.111 | 1.4E-182 | Regenerative Mesenchyme | trmt112 |
| CABZ01115113.1 | 9E-187 | 1.670157 | 0.291 | 0.102 | 2.9E-182 | Regenerative Mesenchyme | CABZ01115113.1 |
| cox6c1 | 2E-186 | 1.2055162 | 0.63 | 0.402 | 6.8E-182 | Regenerative Mesenchyme | cox6c |
| psmd14 | 3E-186 | 1.6599313 | 0.324 | 0.123 | 1.1E-181 | Regenerative Mesenchyme | psmd14 |
| prdx11 | 3E-186 | 1.5611457 | 0.211 | 0.059 | 1.1E-181 | Regenerative Mesenchyme | prdx1 |
| lyar | 3E-185 | 1.8794177 | 0.213 | 0.061 | 9.9E-181 | Regenerative Mesenchyme | lyar |
| abt1 | 9E-185 | 1.7390306 | 0.23 | 0.069 | 2.8E-180 | Regenerative Mesenchyme | abt1 |
| mrpl17 | 1E-184 | 1.7975898 | 0.212 | 0.06 | 3.8E-180 | Regenerative Mesenchyme | mrpl17 |
| pum3 | 1E-183 | 1.704705 | 0.236 | 0.072 | 4.5E-179 | Regenerative Mesenchyme | pum3 |
| mrpl201 | 2E-183 | 1.8793475 | 0.219 | 0.064 | 6.9E-179 | Regenerative Mesenchyme | mrpl20 |
| krt942 | 4E-183 | 0.477997 | 0.576 | 0.279 | 1.3E-178 | Regenerative Mesenchyme | krt94 |
| cyc1 | 5E-183 | 1.5453504 | 0.374 | 0.158 | 1.6E-178 | Regenerative Mesenchyme | cyc1 |
| pole4 | 8E-183 | 2.0056759 | 0.192 | 0.051 | 2.5E-178 | Regenerative Mesenchyme | pole4 |
| lsm7 | 3E-182 | 1.3682191 | 0.488 | 0.247 | 1.1E-177 | Regenerative Mesenchyme | lsm7 |
| selenoh | 4E-182 | 1.7665434 | 0.303 | 0.113 | 1.2E-177 | Regenerative Mesenchyme | selenoh |
| pfn21 | 5E-182 | 0.9414178 | 0.745 | 0.527 | 1.6E-177 | Regenerative Mesenchyme | pfn2 |
| rps102 | 3E-181 | 0.3941655 | 0.998 | 0.994 | 8.3E-177 | Regenerative Mesenchyme | rps10 |
| nars1 | 5E-181 | 1.5659691 | 0.343 | 0.137 | 1.6E-176 | Regenerative Mesenchyme | nars1 |
| nop161 | 1E-180 | 1.8480289 | 0.201 | 0.056 | 4.1E-176 | Regenerative Mesenchyme | nop16 |
| nop2 | 1E-180 | 1.4670854 | 0.415 | 0.188 | 4.3E-176 | Regenerative Mesenchyme | nop2 |
| sap18 | 2E-180 | 1.3748017 | 0.419 | 0.189 | 6.1E-176 | Regenerative Mesenchyme | sap18 |
| krtcap21 | 3E-180 | 1.4003186 | 0.49 | 0.257 | 9.6E-176 | Regenerative Mesenchyme | krtcap2 |
| mrpl131 | 8E-180 | 1.8631438 | 0.205 | 0.058 | 2.6E-175 | Regenerative Mesenchyme | mrpl13 |
| cct51 | 1E-179 | 1.2912011 | 0.592 | 0.369 | 4.9E-175 | Regenerative Mesenchyme | cct5 |
| acp1 | 2E-179 | 1.8399441 | 0.225 | 0.068 | 5.2E-175 | Regenerative Mesenchyme | acp1 |
| vcp | 4E-179 | 1.248782 | 0.544 | 0.304 | 1.4E-174 | Regenerative Mesenchyme | vcp |
| czib | 5E-178 | 1.6554325 | 0.259 | 0.086 | 1.8E-173 | Regenerative Mesenchyme | czib |
| aurkaip1 | 6E-178 | 1.8950338 | 0.209 | 0.06 | 1.9E-173 | Regenerative Mesenchyme | aurkaip1 |
| sec13 | 9E-177 | 1.6046875 | 0.308 | 0.115 | 2.9E-172 | Regenerative Mesenchyme | sec13 |
| pa2g4b1 | 9E-177 | 1.4032084 | 0.466 | 0.238 | 2.9E-172 | Regenerative Mesenchyme | pa2g4b |
| hspa8 | 1E-176 | 0.5064054 | 0.984 | 0.934 | 4.4E-172 | Regenerative Mesenchyme | hspa8 |
| chchd1 | 1E-176 | 1.9756442 | 0.17 | 0.042 | 4.6E-172 | Regenerative Mesenchyme | chchd1 |
| cct3 | 5E-176 | 1.2866268 | 0.548 | 0.318 | 1.5E-171 | Regenerative Mesenchyme | cct3 |
| myl12.1 | 6E-176 | 0.9629741 | 0.722 | 0.524 | 1.8E-171 | Regenerative Mesenchyme | myl12.1 |
| rpa1 | 9E-176 | 2.1116277 | 0.167 | 0.041 | 2.8E-171 | Regenerative Mesenchyme | rpa1 |
| ccdc3a | 3E-175 | 1.5670938 | 0.209 | 0.06 | 1.1E-170 | Regenerative Mesenchyme | ccdc3a |
| anxa2a1 | 6E-175 | 0.6344386 | 0.958 | 0.887 | 1.9E-170 | Regenerative Mesenchyme | anxa2a |

| | | | | | | | |
|---|---|---|---|---|---|---|---|
| psmc1a | 1E-174 | 1.5930484 | 0.288 | 0.104 | 4.3E-170 | Regenerative Mesenchyme | psmc1a |
| C25H12orf73 | 3E-174 | 2.0921942 | 0.154 | 0.036 | 8.8E-170 | Regenerative Mesenchyme | C25H12orf73 |
| syncrip | 4E-174 | 1.0281358 | 0.694 | 0.484 | 1.2E-169 | Regenerative Mesenchyme | syncrip |
| psma6b | 4E-174 | 1.5605353 | 0.344 | 0.141 | 1.2E-169 | Regenerative Mesenchyme | psma6b |
| pin41 | 4E-174 | 1.5243987 | 0.34 | 0.137 | 1.5E-169 | Regenerative Mesenchyme | pin4 |
| eif6 | 5E-174 | 1.453978 | 0.378 | 0.164 | 1.7E-169 | Regenerative Mesenchyme | eif6 |
| eef1a1l12 | 7E-174 | 0.4059641 | 0.999 | 0.996 | 2.2E-169 | Regenerative Mesenchyme | eef1a1l1 |
| chd4a | 3E-173 | 1.2798297 | 0.451 | 0.215 | 8.3E-169 | Regenerative Mesenchyme | chd4a |
| cldni1 | 1E-172 | 2.3960836 | 0.192 | 0.054 | 4.2E-168 | Regenerative Mesenchyme | cldni |
| tram11 | 2E-172 | 1.3501249 | 0.507 | 0.274 | 6.9E-168 | Regenerative Mesenchyme | tram1 |
| eno1a | 3E-172 | 1.3487998 | 0.556 | 0.327 | 9.6E-168 | Regenerative Mesenchyme | eno1a |
| gars11 | 2E-171 | 1.6340885 | 0.335 | 0.138 | 6.1E-167 | Regenerative Mesenchyme | gars1 |
| lman11 | 4E-171 | 1.6627473 | 0.302 | 0.114 | 1.3E-166 | Regenerative Mesenchyme | lman1 |
| lipg | 9E-171 | 2.7318353 | 0.108 | 0.019 | 3E-166 | Regenerative Mesenchyme | lipg |
| srp191 | 5E-170 | 1.5064813 | 0.327 | 0.13 | 1.7E-165 | Regenerative Mesenchyme | srp19 |
| rars1 | 6E-170 | 1.8268297 | 0.198 | 0.056 | 2E-165 | Regenerative Mesenchyme | rars1 |
| nsfl1c | 1E-169 | 1.5441932 | 0.305 | 0.116 | 3.3E-165 | Regenerative Mesenchyme | nsfl1c |
| usp14 | 1E-169 | 1.9764046 | 0.185 | 0.05 | 4.3E-165 | Regenerative Mesenchyme | usp14 |
| mrps33 | 2E-169 | 1.5019814 | 0.297 | 0.111 | 6E-165 | Regenerative Mesenchyme | mrps33 |
| atad5a | 2E-169 | 2.4074641 | 0.111 | 0.02 | 6.1E-165 | Regenerative Mesenchyme | atad5a |
| hspa4b | 8E-169 | 1.5013513 | 0.33 | 0.132 | 2.7E-164 | Regenerative Mesenchyme | hspa4b |
| kars11 | 8E-169 | 1.5393577 | 0.36 | 0.154 | 2.7E-164 | Regenerative Mesenchyme | kars1 |
| tcp11 | 1E-168 | 1.3082173 | 0.532 | 0.31 | 4.4E-164 | Regenerative Mesenchyme | tcp1 |
| mrps24 | 2E-168 | 1.8387341 | 0.212 | 0.064 | 7.1E-164 | Regenerative Mesenchyme | mrps24 |
| ptmab1 | 7E-168 | 0.5982723 | 0.899 | 0.704 | 2.4E-163 | Regenerative Mesenchyme | ptmab |
| ENSDARG00000356 | 2E-167 | 1.3386651 | 0.427 | 0.205 | 6.8E-163 | Regenerative Mesenchyme | ENSDARG00000035 |
| glrx3 | 2E-167 | 1.5481101 | 0.309 | 0.119 | 8.1E-163 | Regenerative Mesenchyme | glrx3 |
| ncl | 3E-167 | 1.1609184 | 0.739 | 0.597 | 9.1E-163 | Regenerative Mesenchyme | ncl |
| sumo3b | 7E-167 | 1.7002555 | 0.297 | 0.114 | 2.2E-162 | Regenerative Mesenchyme | sumo3b |
| llph1 | 2E-166 | 1.7234076 | 0.233 | 0.075 | 5.4E-162 | Regenerative Mesenchyme | llph |
| lmnb2 | 2E-166 | 1.733015 | 0.259 | 0.09 | 6.5E-162 | Regenerative Mesenchyme | lmnb2 |
| timm91 | 5E-166 | 1.6288542 | 0.256 | 0.088 | 1.8E-161 | Regenerative Mesenchyme | timm9 |
| ndufc2 | 6E-166 | 1.2037421 | 0.5 | 0.264 | 2E-161 | Regenerative Mesenchyme | ndufc2 |
| ost41 | 8E-166 | 1.5449478 | 0.338 | 0.14 | 2.5E-161 | Regenerative Mesenchyme | ost4 |
| mrpl21 | 1E-165 | 1.9688199 | 0.159 | 0.039 | 4.1E-161 | Regenerative Mesenchyme | mrpl21 |
| vars1 | 1E-165 | 1.7748744 | 0.204 | 0.06 | 4.8E-161 | Regenerative Mesenchyme | vars1 |
| smarcad1a | 3E-165 | 2.2048681 | 0.125 | 0.026 | 9.8E-161 | Regenerative Mesenchyme | smarcad1a |
| ckba1 | 5E-165 | 0.6996508 | 0.825 | 0.63 | 1.5E-160 | Regenerative Mesenchyme | ckba |
| CT573476.1 | 6E-165 | 2.0531889 | 0.151 | 0.036 | 1.9E-160 | Regenerative Mesenchyme | CT573476.1 |
| nansa | 1E-164 | 1.7757901 | 0.213 | 0.065 | 3.9E-160 | Regenerative Mesenchyme | nansa |
| calm3a2 | 2E-164 | 0.8404513 | 0.721 | 0.489 | 5.8E-160 | Regenerative Mesenchyme | calm3a |
| npm3 | 4E-164 | 1.5266401 | 0.298 | 0.113 | 1.4E-159 | Regenerative Mesenchyme | npm3 |
| calua1 | 2E-163 | 1.149092 | 0.549 | 0.313 | 5.7E-159 | Regenerative Mesenchyme | calua |
| pwp1 | 3E-163 | 1.5766745 | 0.217 | 0.067 | 1E-158 | Regenerative Mesenchyme | pwp1 |
| NC-002333.172 | 6E-163 | 0.6627505 | 1 | 0.998 | 1.9E-158 | Regenerative Mesenchyme | NC-002333.17 |
| psmc3 | 9E-163 | 1.6334546 | 0.254 | 0.088 | 2.8E-158 | Regenerative Mesenchyme | psmc3 |
| psmb2 | 2E-162 | 1.3614274 | 0.473 | 0.254 | 6.2E-158 | Regenerative Mesenchyme | psmb2 |
| ndufs61 | 2E-162 | 1.4583296 | 0.325 | 0.131 | 7.9E-158 | Regenerative Mesenchyme | ndufs6 |

| | | | | | | | |
|---|---|---|---|---|---|---|---|
| cad | 4E-162 | 1.7444915 | 0.225 | 0.072 | 1.4E-157 | Regenerative Mesenchyme | cad |
| si:ch211-235e9.61 | 2E-161 | 1.699637 | 0.247 | 0.084 | 7.6E-157 | Regenerative Mesenchyme | si:ch211-235e9.6 |
| psma1 | 3E-161 | 1.1665329 | 0.53 | 0.298 | 8.7E-157 | Regenerative Mesenchyme | psma1 |
| flna2 | 6E-161 | 0.9294585 | 0.74 | 0.532 | 1.8E-156 | Regenerative Mesenchyme | flna |
| rcn32 | 6E-161 | 1.3247271 | 0.476 | 0.25 | 1.9E-156 | Regenerative Mesenchyme | rcn3 |
| mrpl181 | 7E-161 | 1.8796301 | 0.198 | 0.059 | 2.3E-156 | Regenerative Mesenchyme | mrpl18 |
| rab5if1 | 2E-160 | 1.6278967 | 0.288 | 0.111 | 6E-156 | Regenerative Mesenchyme | rab5if |
| timm8a1 | 3E-160 | 1.6853963 | 0.241 | 0.082 | 8.7E-156 | Regenerative Mesenchyme | timm8a |
| ppid1 | 3E-160 | 1.8063933 | 0.225 | 0.073 | 1.1E-155 | Regenerative Mesenchyme | ppid |
| ssr42 | 4E-160 | 0.8591551 | 0.772 | 0.598 | 1.3E-155 | Regenerative Mesenchyme | ssr4 |
| krt18a.12 | 8E-160 | 0.4812464 | 0.966 | 0.834 | 2.5E-155 | Regenerative Mesenchyme | krt18a.1 |
| slc25a3b1 | 1E-159 | 0.9412149 | 0.726 | 0.541 | 3.8E-155 | Regenerative Mesenchyme | slc25a3b |
| sgk2b | 2E-159 | 2.3749835 | 0.139 | 0.033 | 6.8E-155 | Regenerative Mesenchyme | sgk2b |
| rpn1 | 2E-159 | 1.375725 | 0.417 | 0.201 | 7.7E-155 | Regenerative Mesenchyme | rpn1 |
| uqcrc1 | 3E-159 | 1.4319676 | 0.376 | 0.17 | 9.9E-155 | Regenerative Mesenchyme | uqcrc1 |
| gnl31 | 3E-158 | 1.559395 | 0.317 | 0.13 | 1.1E-153 | Regenerative Mesenchyme | gnl3 |
| glrx | 1E-157 | 1.0599937 | 0.582 | 0.344 | 3.6E-153 | Regenerative Mesenchyme | glrx |
| psmc2 | 1E-157 | 1.5509259 | 0.297 | 0.116 | 3.8E-153 | Regenerative Mesenchyme | psmc2 |
| uqcrq | 1E-157 | 1.2498564 | 0.479 | 0.257 | 4.6E-153 | Regenerative Mesenchyme | uqcrq |
| g3bp1 | 2E-157 | 1.4057573 | 0.36 | 0.158 | 5.4E-153 | Regenerative Mesenchyme | g3bp1 |
| cers2b | 6E-157 | 1.8332361 | 0.199 | 0.06 | 1.9E-152 | Regenerative Mesenchyme | cers2b |
| rpl152 | 7E-157 | 0.4318261 | 0.991 | 0.969 | 2.2E-152 | Regenerative Mesenchyme | rpl15 |
| tbca | 1E-156 | 1.1421456 | 0.533 | 0.303 | 3.6E-152 | Regenerative Mesenchyme | tbca |
| mrpl47 | 1E-156 | 1.763862 | 0.194 | 0.057 | 3.8E-152 | Regenerative Mesenchyme | mrpl47 |
| si:dkey-34d22.1 | 2E-156 | 2.1876539 | 0.137 | 0.032 | 5.3E-152 | Regenerative Mesenchyme | si:dkey-34d22.1 |
| mrpl28 | 3E-156 | 1.818708 | 0.204 | 0.063 | 9.6E-152 | Regenerative Mesenchyme | mrpl28 |
| emc6 | 5E-156 | 1.5827706 | 0.26 | 0.094 | 1.7E-151 | Regenerative Mesenchyme | emc6 |
| psmb5 | 9E-156 | 1.5311782 | 0.314 | 0.127 | 2.8E-151 | Regenerative Mesenchyme | psmb5 |
| atp5f1c1 | 9E-156 | 1.3916404 | 0.46 | 0.244 | 3E-151 | Regenerative Mesenchyme | atp5f1c |
| h2az2a | 1E-155 | 1.4723311 | 0.341 | 0.145 | 4.1E-151 | Regenerative Mesenchyme | h2az2a |
| parp1 | 1E-155 | 1.9328235 | 0.169 | 0.046 | 4.2E-151 | Regenerative Mesenchyme | parp1 |
| ndufb8 | 2E-155 | 1.1325039 | 0.549 | 0.324 | 6.8E-151 | Regenerative Mesenchyme | ndufb8 |
| suclg1 | 6E-155 | 1.3891345 | 0.366 | 0.164 | 1.8E-150 | Regenerative Mesenchyme | suclg1 |
| atp5pb | 2E-154 | 1.0467016 | 0.642 | 0.454 | 5.5E-150 | Regenerative Mesenchyme | atp5pb |
| mrps26 | 4E-154 | 1.776157 | 0.177 | 0.05 | 1.3E-149 | Regenerative Mesenchyme | mrps26 |
| sdhdb | 4E-154 | 1.5524427 | 0.298 | 0.118 | 1.4E-149 | Regenerative Mesenchyme | sdhdb |
| eif4g1a | 2E-153 | 1.1445251 | 0.569 | 0.343 | 7.7E-149 | Regenerative Mesenchyme | eif4g1a |
| psma21 | 2E-153 | 1.3835898 | 0.399 | 0.19 | 8E-149 | Regenerative Mesenchyme | psma2 |
| cox6b21 | 4E-153 | 0.624222 | 0.883 | 0.811 | 1.2E-148 | Regenerative Mesenchyme | cox6b2 |
| tmsb21 | 6E-153 | 1.8460958 | 0.245 | 0.088 | 1.9E-148 | Regenerative Mesenchyme | tmsb2 |
| cd2761 | 1E-152 | 1.5792506 | 0.304 | 0.123 | 3.5E-148 | Regenerative Mesenchyme | cd276 |
| adtrp1 | 1E-152 | 1.6555272 | 0.223 | 0.073 | 4.3E-148 | Regenerative Mesenchyme | adtrp1 |
| zgc:1629441 | 2E-152 | 1.1785175 | 0.505 | 0.281 | 5.6E-148 | Regenerative Mesenchyme | zgc:162944 |
| psmd4a | 2E-152 | 1.5376067 | 0.255 | 0.091 | 6.8E-148 | Regenerative Mesenchyme | psmd4a |
| BOLA2 | 3E-152 | 1.5707234 | 0.248 | 0.087 | 1E-147 | Regenerative Mesenchyme | BOLA2 |
| pcdh19 | 6E-152 | 1.7317007 | 0.169 | 0.046 | 2E-147 | Regenerative Mesenchyme | pcdh19 |
| nip71 | 9E-152 | 1.8793047 | 0.188 | 0.056 | 2.8E-147 | Regenerative Mesenchyme | nip7 |
| nedd8 | 1E-151 | 1.5470999 | 0.267 | 0.098 | 4.6E-147 | Regenerative Mesenchyme | nedd8 |

| | | | | | | | |
|---|---|---|---|---|---|---|---|
| sem1 | 4E-151 | 1.0364769 | 0.614 | 0.406 | 1.3E-146 | Regenerative Mesenchyme | sem1 |
| ndufs7 | 8E-151 | 1.3582484 | 0.369 | 0.167 | 2.5E-146 | Regenerative Mesenchyme | ndufs7 |
| ilf3b | 2E-150 | 1.2694529 | 0.384 | 0.176 | 5.7E-146 | Regenerative Mesenchyme | ilf3b |
| cdkn2aipnl | 2E-150 | 1.7980816 | 0.187 | 0.055 | 7.9E-146 | Regenerative Mesenchyme | cdkn2aipnl |
| ptrh2 | 1E-149 | 1.9907265 | 0.148 | 0.037 | 3.6E-145 | Regenerative Mesenchyme | ptrh2 |
| psmc6 | 1E-149 | 1.4840844 | 0.306 | 0.125 | 4E-145 | Regenerative Mesenchyme | psmc6 |
| creld2 | 2E-149 | 2.3911003 | 0.11 | 0.022 | 6E-145 | Regenerative Mesenchyme | creld2 |
| noc2l | 1E-148 | 1.4476175 | 0.287 | 0.112 | 3.4E-144 | Regenerative Mesenchyme | noc2l |
| enc12 | 1E-148 | 1.3168032 | 0.292 | 0.114 | 4.1E-144 | Regenerative Mesenchyme | enc1 |
| ugdh | 2E-148 | 1.267938 | 0.368 | 0.167 | 5.9E-144 | Regenerative Mesenchyme | ugdh |
| ndufs4 | 3E-148 | 1.5224785 | 0.302 | 0.123 | 1E-143 | Regenerative Mesenchyme | ndufs4 |
| bms1 | 1E-147 | 1.5135112 | 0.258 | 0.095 | 3.9E-143 | Regenerative Mesenchyme | bms1 |
| samm50 | 7E-147 | 2.0012754 | 0.137 | 0.033 | 2.2E-142 | Regenerative Mesenchyme | samm50 |
| tmed10 | 1E-146 | 1.1284575 | 0.545 | 0.325 | 3.7E-142 | Regenerative Mesenchyme | tmed10 |
| eprs11 | 2E-146 | 1.6206516 | 0.24 | 0.085 | 7E-142 | Regenerative Mesenchyme | eprs1 |
| rpl272 | 2E-145 | 0.3973857 | 0.992 | 0.977 | 5.6E-141 | Regenerative Mesenchyme | rpl27 |
| rplp12 | 2E-145 | 0.3054845 | 1 | 0.999 | 6.7E-141 | Regenerative Mesenchyme | rplp1 |
| si:dkeyp-32g11.8 | 3E-145 | 2.5642936 | 0.102 | 0.02 | 8.6E-141 | Regenerative Mesenchyme | si:dkeyp-32g11.8 |
| mrps21 | 4E-145 | 1.5687496 | 0.211 | 0.069 | 1.2E-140 | Regenerative Mesenchyme | mrps21 |
| si:ch211-51e12.7 | 6E-145 | 1.396519 | 0.312 | 0.129 | 2.1E-140 | Regenerative Mesenchyme | si:ch211-51e12.7 |
| ndufb7 | 6E-145 | 1.1056711 | 0.522 | 0.302 | 2.1E-140 | Regenerative Mesenchyme | ndufb7 |
| gpx4b | 2E-144 | 1.1279055 | 0.504 | 0.283 | 6.9E-140 | Regenerative Mesenchyme | gpx4b |
| hat1 | 4E-144 | 2.1066328 | 0.137 | 0.034 | 1.2E-139 | Regenerative Mesenchyme | hat1 |
| calm3b | 4E-144 | 1.0046981 | 0.616 | 0.405 | 1.2E-139 | Regenerative Mesenchyme | calm3b |
| mrps17 | 6E-144 | 2.1393814 | 0.113 | 0.024 | 1.8E-139 | Regenerative Mesenchyme | mrps17 |
| mars1 | 7E-144 | 1.8047462 | 0.184 | 0.056 | 2.2E-139 | Regenerative Mesenchyme | mars1 |
| inhbaa | 2E-143 | 1.1490714 | 0.485 | 0.26 | 8.1E-139 | Regenerative Mesenchyme | inhbaa |
| tomm22 | 3E-143 | 1.6053917 | 0.247 | 0.09 | 8.3E-139 | Regenerative Mesenchyme | tomm22 |
| gpatch4 | 3E-143 | 1.6403496 | 0.196 | 0.061 | 8.7E-139 | Regenerative Mesenchyme | gpatch4 |
| ywhag2 | 3E-143 | 1.3865158 | 0.35 | 0.157 | 1E-138 | Regenerative Mesenchyme | ywhag2 |
| rps8a2 | 2E-142 | 0.3663728 | 0.998 | 0.991 | 5.8E-138 | Regenerative Mesenchyme | rps8a |
| ssbp1 | 2E-142 | 1.7407669 | 0.173 | 0.05 | 7E-138 | Regenerative Mesenchyme | ssbp1 |
| hnrnph1l | 5E-142 | 1.4082528 | 0.302 | 0.124 | 1.6E-137 | Regenerative Mesenchyme | hnrnph1l |
| cct71 | 8E-142 | 1.0690603 | 0.569 | 0.361 | 2.7E-137 | Regenerative Mesenchyme | cct7 |
| rps252 | 1E-141 | 0.3387566 | 0.997 | 0.994 | 4.3E-137 | Regenerative Mesenchyme | rps25 |
| si:dkey-248g15.31 | 1E-141 | 1.7413451 | 0.383 | 0.189 | 4.8E-137 | Regenerative Mesenchyme | si:dkey-248g15.3 |
| arf5 | 6E-141 | 1.1844376 | 0.509 | 0.298 | 1.9E-136 | Regenerative Mesenchyme | arf5 |
| rcc1 | 2E-140 | 2.2544646 | 0.105 | 0.021 | 7.1E-136 | Regenerative Mesenchyme | rcc1 |
| cct2.12 | 2E-140 | 1.050787 | 0.606 | 0.413 | 7.5E-136 | Regenerative Mesenchyme | cct2.1 |
| qars1 | 3E-140 | 1.5173026 | 0.219 | 0.074 | 1E-135 | Regenerative Mesenchyme | qars1 |
| alyref1 | 4E-140 | 1.0495852 | 0.55 | 0.336 | 1.2E-135 | Regenerative Mesenchyme | alyref |
| rps242 | 2E-139 | 0.3080586 | 0.999 | 0.996 | 5.8E-135 | Regenerative Mesenchyme | rps24 |
| lrpprc | 2E-139 | 1.8731814 | 0.151 | 0.04 | 6.3E-135 | Regenerative Mesenchyme | lrpprc |
| ppib2 | 2E-139 | 0.5212166 | 0.952 | 0.915 | 7.4E-135 | Regenerative Mesenchyme | ppib |
| psph | 3E-139 | 2.1601096 | 0.123 | 0.029 | 9.2E-135 | Regenerative Mesenchyme | psph |
| si:ch211-114n24.6 | 5E-139 | 1.9789372 | 0.199 | 0.066 | 1.8E-134 | Regenerative Mesenchyme | si:ch211-114n24.6 |
| lrrc591 | 7E-139 | 1.4221253 | 0.309 | 0.131 | 2.3E-134 | Regenerative Mesenchyme | lrrc59 |
| bop1 | 1E-138 | 1.8313734 | 0.159 | 0.044 | 4.5E-134 | Regenerative Mesenchyme | bop1 |

| | | | | | | | | |
|---|---|---|---|---|---|---|---|---|
| rpl7l1 | 1E-138 | 1.4567682 | 0.3 | 0.125 | 4.7E-134 | Regenerative | Mesenchyme | rpl7l1 |
| arpp19b | 2E-138 | 1.6861807 | 0.261 | 0.102 | 5.8E-134 | Regenerative | Mesenchyme | arpp19b |
| lmnb1 | 4E-138 | 2.0270933 | 0.157 | 0.044 | 1.2E-133 | Regenerative | Mesenchyme | lmnb1 |
| cst14a.2 | 1E-137 | 0.9497675 | 0.573 | 0.351 | 3.9E-133 | Regenerative | Mesenchyme | cst14a.2 |
| smarca5 | 1E-137 | 1.8964576 | 0.152 | 0.042 | 4.2E-133 | Regenerative | Mesenchyme | smarca5 |
| psmc4 | 2E-137 | 1.3500781 | 0.342 | 0.155 | 5.6E-133 | Regenerative | Mesenchyme | psmc4 |
| ppig | 3E-137 | 1.2193976 | 0.299 | 0.121 | 8.8E-133 | Regenerative | Mesenchyme | ppig |
| bysl1 | 3E-137 | 1.6043702 | 0.265 | 0.105 | 9.5E-133 | Regenerative | Mesenchyme | bysl |
| rpl92 | 8E-137 | 0.3451387 | 0.996 | 0.987 | 2.6E-132 | Regenerative | Mesenchyme | rpl9 |
| prmt5 | 9E-137 | 2.044205 | 0.126 | 0.03 | 2.9E-132 | Regenerative | Mesenchyme | prmt5 |
| mapre1b | 2E-136 | 1.4870112 | 0.281 | 0.113 | 6.1E-132 | Regenerative | Mesenchyme | mapre1b |
| ppiab | 3E-136 | 0.4762555 | 0.953 | 0.893 | 1E-131 | Regenerative | Mesenchyme | ppiab |
| pdia4 | 6E-136 | 1.4303536 | 0.318 | 0.139 | 2E-131 | Regenerative | Mesenchyme | pdia4 |
| cope1 | 8E-136 | 1.2487176 | 0.451 | 0.245 | 2.5E-131 | Regenerative | Mesenchyme | cope |
| rps192 | 1E-135 | 0.3163993 | 0.998 | 0.996 | 3.8E-131 | Regenerative | Mesenchyme | rps19 |
| ywhaba | 1E-135 | 0.8671143 | 0.687 | 0.496 | 4.2E-131 | Regenerative | Mesenchyme | ywhaba |
| hnrnpa0l.11 | 2E-135 | 0.7149601 | 0.792 | 0.66 | 5.2E-131 | Regenerative | Mesenchyme | hnrnpa0l.1 |
| exosc9 | 2E-135 | 2.1638832 | 0.105 | 0.022 | 5.7E-131 | Regenerative | Mesenchyme | exosc9 |
| ddx27 | 2E-135 | 1.3310421 | 0.368 | 0.176 | 7.6E-131 | Regenerative | Mesenchyme | ddx27 |
| hnrnpa1b | 3E-135 | 1.0233098 | 0.603 | 0.404 | 1E-130 | Regenerative | Mesenchyme | hnrnpa1b |
| kdelr2a | 3E-135 | 1.0429561 | 0.59 | 0.379 | 1E-130 | Regenerative | Mesenchyme | kdelr2a |
| rps132 | 3E-135 | 0.3423214 | 0.996 | 0.99 | 1.1E-130 | Regenerative | Mesenchyme | rps13 |
| hm13 | 5E-135 | 1.409271 | 0.307 | 0.132 | 1.7E-130 | Regenerative | Mesenchyme | hm13 |
| ruvbl1 | 9E-135 | 1.7132209 | 0.169 | 0.05 | 2.9E-130 | Regenerative | Mesenchyme | ruvbl1 |
| vcana | 2E-134 | 1.6128908 | 0.115 | 0.026 | 5.5E-130 | Regenerative | Mesenchyme | vcana |
| fhl3a | 3E-134 | 1.8257866 | 0.189 | 0.061 | 9.3E-130 | Regenerative | Mesenchyme | fhl3a |
| pdlim4 | 4E-134 | 1.9450362 | 0.136 | 0.035 | 1.3E-129 | Regenerative | Mesenchyme | pdlim4 |
| dnajc2 | 4E-134 | 1.7373895 | 0.153 | 0.042 | 1.3E-129 | Regenerative | Mesenchyme | dnajc2 |
| tomm71 | 4E-134 | 1.3534578 | 0.375 | 0.184 | 1.3E-129 | Regenerative | Mesenchyme | tomm7 |
| adssl | 3E-133 | 1.6026243 | 0.214 | 0.074 | 1E-128 | Regenerative | Mesenchyme | adssl |
| col17a1b | 3E-133 | 2.9350853 | 0.105 | 0.023 | 1E-128 | Regenerative | Mesenchyme | col17a1b |
| h3f3d1 | 4E-133 | 0.6161975 | 0.832 | 0.722 | 1.2E-128 | Regenerative | Mesenchyme | h3f3d |
| tsr21 | 5E-133 | 1.5871181 | 0.194 | 0.063 | 1.6E-128 | Regenerative | Mesenchyme | tsr2 |
| rps232 | 7E-133 | 0.3783786 | 0.995 | 0.981 | 2.2E-128 | Regenerative | Mesenchyme | rps23 |
| ndufa7 | 7E-133 | 1.7568258 | 0.155 | 0.044 | 2.4E-128 | Regenerative | Mesenchyme | ndufa7 |
| rrp1 | 1E-132 | 1.3468079 | 0.318 | 0.139 | 3.2E-128 | Regenerative | Mesenchyme | rrp1 |
| coa6 | 1E-132 | 1.4056662 | 0.247 | 0.093 | 4.7E-128 | Regenerative | Mesenchyme | coa6 |
| actl6a | 2E-132 | 1.8130659 | 0.173 | 0.053 | 7.5E-128 | Regenerative | Mesenchyme | actl6a |
| aco2 | 3E-132 | 1.6228637 | 0.213 | 0.074 | 9.5E-128 | Regenerative | Mesenchyme | aco2 |
| mybbp1a | 4E-132 | 1.4035064 | 0.283 | 0.116 | 1.4E-127 | Regenerative | Mesenchyme | mybbp1a |
| snrpa1 | 5E-132 | 1.4485725 | 0.242 | 0.09 | 1.6E-127 | Regenerative | Mesenchyme | snrpa1 |
| arpc3 | 1E-131 | 0.9997784 | 0.584 | 0.373 | 3.8E-127 | Regenerative | Mesenchyme | arpc3 |
| nav3 | 3E-131 | 1.5694929 | 0.214 | 0.075 | 9.3E-127 | Regenerative | Mesenchyme | nav3 |
| map7d1b1 | 2E-130 | 1.2314509 | 0.372 | 0.179 | 5.4E-126 | Regenerative | Mesenchyme | map7d1b |
| ftsj3 | 2E-130 | 1.5862569 | 0.186 | 0.06 | 6.7E-126 | Regenerative | Mesenchyme | ftsj3 |
| rps72 | 3E-130 | 0.3578986 | 0.998 | 0.991 | 9.4E-126 | Regenerative | Mesenchyme | rps7 |
| gcsha | 7E-130 | 1.4986067 | 0.245 | 0.093 | 2.1E-125 | Regenerative | Mesenchyme | gcsha |
| hibadhb | 1E-129 | 1.5109317 | 0.238 | 0.089 | 3.3E-125 | Regenerative | Mesenchyme | hibadhb |

| | | | | | | | |
|---|---|---|---|---|---|---|---|
| hnrnpa1a | 1E-129 | 1.4118772 | 0.297 | 0.127 | 3.5E-125 | Regenerative Mesenchyme | hnrnpa1a |
| ube2na | 3E-129 | 1.0539847 | 0.485 | 0.271 | 9.3E-125 | Regenerative Mesenchyme | ube2na |
| rrs11 | 4E-129 | 1.5069144 | 0.258 | 0.102 | 1.2E-124 | Regenerative Mesenchyme | rrs1 |
| timm13 | 5E-129 | 1.6647657 | 0.181 | 0.058 | 1.7E-124 | Regenerative Mesenchyme | timm13 |
| nolc1 | 6E-129 | 1.4599956 | 0.264 | 0.106 | 1.9E-124 | Regenerative Mesenchyme | nolc1 |
| ddx24 | 8E-129 | 1.3853354 | 0.255 | 0.099 | 2.6E-124 | Regenerative Mesenchyme | ddx24 |
| si:dkeyp-27c8.2 | 2E-128 | 1.5797712 | 0.152 | 0.043 | 5.9E-124 | Regenerative Mesenchyme | si:dkeyp-27c8.2 |
| smarce1 | 1E-127 | 1.364048 | 0.316 | 0.141 | 3.8E-123 | Regenerative Mesenchyme | smarce1 |
| rab1ab | 4E-127 | 0.9240168 | 0.611 | 0.409 | 1.4E-122 | Regenerative Mesenchyme | rab1ab |
| pwp2h1 | 5E-127 | 1.5562244 | 0.211 | 0.075 | 1.7E-122 | Regenerative Mesenchyme | pwp2h |
| sf3a2 | 9E-127 | 1.3036692 | 0.329 | 0.15 | 2.8E-122 | Regenerative Mesenchyme | sf3a2 |
| hnrnpl | 2E-126 | 1.4258402 | 0.243 | 0.093 | 5.6E-122 | Regenerative Mesenchyme | hnrnpl |
| bud31 | 3E-126 | 1.2245044 | 0.39 | 0.197 | 9E-122 | Regenerative Mesenchyme | bud31 |
| anapc11 | 4E-126 | 1.3393685 | 0.285 | 0.12 | 1.2E-121 | Regenerative Mesenchyme | anapc11 |
| ipo4 | 5E-126 | 1.9429752 | 0.111 | 0.025 | 1.5E-121 | Regenerative Mesenchyme | ipo4 |
| uba1 | 4E-125 | 1.202614 | 0.395 | 0.201 | 1.3E-120 | Regenerative Mesenchyme | uba1 |
| mrpl161 | 1E-124 | 1.6456148 | 0.174 | 0.055 | 4.9E-120 | Regenerative Mesenchyme | mrpl16 |
| marcksl1a | 5E-124 | 0.6208119 | 0.853 | 0.699 | 1.6E-119 | Regenerative Mesenchyme | marcksl1a |
| psmd11b | 6E-124 | 1.8384843 | 0.145 | 0.041 | 2E-119 | Regenerative Mesenchyme | psmd11b |
| mrpl341 | 2E-123 | 1.382328 | 0.282 | 0.12 | 6.8E-119 | Regenerative Mesenchyme | mrpl34 |
| rps15a2 | 5E-123 | 0.3308195 | 0.994 | 0.989 | 1.7E-118 | Regenerative Mesenchyme | rps15a |
| ncbp2 | 7E-123 | 1.3503733 | 0.274 | 0.114 | 2.4E-118 | Regenerative Mesenchyme | ncbp2 |
| nfkbiaa2 | 8E-123 | 0.8189705 | 0.634 | 0.411 | 2.6E-118 | Regenerative Mesenchyme | nfkbiaa |
| lin52 | 2E-122 | 1.5777836 | 0.179 | 0.058 | 5.6E-118 | Regenerative Mesenchyme | lin52 |
| blmh | 3E-122 | 1.6907324 | 0.167 | 0.052 | 8.5E-118 | Regenerative Mesenchyme | blmh |
| ruvbl2 | 6E-122 | 1.9865251 | 0.106 | 0.024 | 1.9E-117 | Regenerative Mesenchyme | ruvbl2 |
| rbb4l | 6E-122 | 1.4516608 | 0.223 | 0.082 | 2E-117 | Regenerative Mesenchyme | rbb4l |
| etf1b | 8E-122 | 1.0010995 | 0.573 | 0.383 | 2.6E-117 | Regenerative Mesenchyme | etf1b |
| ndufa21 | 9E-122 | 1.4444315 | 0.227 | 0.086 | 2.9E-117 | Regenerative Mesenchyme | ndufa2 |
| elocb | 9E-122 | 1.061126 | 0.488 | 0.285 | 2.9E-117 | Regenerative Mesenchyme | elocb |
| si:dkey-102m7.31 | 1E-121 | 1.4171251 | 0.268 | 0.112 | 3.6E-117 | Regenerative Mesenchyme | si:dkey-102m7.3 |
| tln2a | 2E-121 | 1.6278441 | 0.2 | 0.07 | 5.8E-117 | Regenerative Mesenchyme | tln2a |
| mrpl43 | 2E-121 | 1.7319925 | 0.141 | 0.04 | 7.2E-117 | Regenerative Mesenchyme | mrpl43 |
| nop56 | 2E-121 | 1.0655283 | 0.601 | 0.426 | 8E-117 | Regenerative Mesenchyme | nop56 |
| eif1axb | 4E-121 | 1.0880399 | 0.49 | 0.293 | 1.3E-116 | Regenerative Mesenchyme | eif1axb |
| tsr1 | 6E-121 | 1.7479186 | 0.162 | 0.05 | 2E-116 | Regenerative Mesenchyme | tsr1 |
| pprc11 | 9E-121 | 1.4939412 | 0.262 | 0.108 | 3.1E-116 | Regenerative Mesenchyme | pprc1 |
| si:ch211-288g17.3 | 1E-120 | 0.9137901 | 0.598 | 0.392 | 3.4E-116 | Regenerative Mesenchyme | si:ch211-288g17.3 |
| fkbp71 | 1E-120 | 0.8292595 | 0.746 | 0.605 | 3.6E-116 | Regenerative Mesenchyme | fkbp7 |
| polr2d | 1E-120 | 1.2973926 | 0.309 | 0.139 | 4E-116 | Regenerative Mesenchyme | polr2d |
| ciapin1 | 3E-120 | 1.6686829 | 0.172 | 0.056 | 1E-115 | Regenerative Mesenchyme | ciapin1 |
| snrnp70 | 4E-120 | 1.1739427 | 0.336 | 0.156 | 1.4E-115 | Regenerative Mesenchyme | snrnp70 |
| CR936321.51 | 1E-119 | 1.4493914 | 0.228 | 0.087 | 4.3E-115 | Regenerative Mesenchyme | CR936321.5 |
| adi11 | 1E-119 | 1.544008 | 0.22 | 0.083 | 4.5E-115 | Regenerative Mesenchyme | adi1 |
| c1d | 3E-119 | 1.4444572 | 0.222 | 0.083 | 9.3E-115 | Regenerative Mesenchyme | c1d |
| kpnb1 | 6E-119 | 1.306138 | 0.301 | 0.135 | 2.1E-114 | Regenerative Mesenchyme | kpnb1 |
| polr1d2 | 7E-119 | 1.54587 | 0.212 | 0.078 | 2.1E-114 | Regenerative Mesenchyme | polr1d |
| UTP14C | 1E-118 | 1.344976 | 0.242 | 0.095 | 4E-114 | Regenerative Mesenchyme | UTP14C |

| | | | | | | | |
|---|---|---|---|---|---|---|---|
| spcs1 | 9E-118 | 0.9191267 | 0.607 | 0.426 | 3E-113 | Regenerative Mesenchyme | spcs1 |
| lsm4 | 1E-117 | 1.1232932 | 0.4 | 0.207 | 4.3E-113 | Regenerative Mesenchyme | lsm4 |
| ndufa4l1 | 2E-117 | 0.5675473 | 0.845 | 0.748 | 5.5E-113 | Regenerative Mesenchyme | ndufa4l |
| ndufb10 | 3E-117 | 1.11169 | 0.417 | 0.225 | 9.6E-113 | Regenerative Mesenchyme | ndufb10 |
| pbdc11 | 3E-117 | 1.0913399 | 0.457 | 0.261 | 1E-112 | Regenerative Mesenchyme | pbdc1 |
| rps122 | 1E-116 | 0.3872506 | 0.991 | 0.977 | 4.3E-112 | Regenerative Mesenchyme | rps12 |
| atp5md | 2E-116 | 1.372931 | 0.298 | 0.135 | 5.6E-112 | Regenerative Mesenchyme | atp5md |
| chst111 | 2E-116 | 2.1548376 | 0.134 | 0.038 | 5.8E-112 | Regenerative Mesenchyme | chst11 |
| pqbp1 | 2E-116 | 1.8874406 | 0.123 | 0.033 | 7.6E-112 | Regenerative Mesenchyme | pqbp1 |
| ubxn1 | 3E-116 | 1.3626734 | 0.231 | 0.089 | 8.2E-112 | Regenerative Mesenchyme | ubxn1 |
| hk2 | 3E-116 | 1.7496565 | 0.153 | 0.047 | 8.7E-112 | Regenerative Mesenchyme | hk2 |
| nup93 | 3E-116 | 1.9688692 | 0.113 | 0.028 | 8.7E-112 | Regenerative Mesenchyme | nup93 |
| coa5 | 4E-116 | 1.5964037 | 0.179 | 0.061 | 1.4E-111 | Regenerative Mesenchyme | coa5 |
| nudcd2 | 8E-116 | 1.5573861 | 0.195 | 0.069 | 2.6E-111 | Regenerative Mesenchyme | nudcd2 |
| ap2m1a | 3E-115 | 1.004038 | 0.493 | 0.294 | 9.7E-111 | Regenerative Mesenchyme | ap2m1a |
| lsm5 | 4E-115 | 1.3806579 | 0.261 | 0.11 | 1.2E-110 | Regenerative Mesenchyme | lsm5 |
| adrm1 | 7E-115 | 1.47349 | 0.218 | 0.083 | 2.4E-110 | Regenerative Mesenchyme | adrm1 |
| ewsr1a | 9E-115 | 1.1695298 | 0.311 | 0.141 | 3E-110 | Regenerative Mesenchyme | ewsr1a |
| glod5 | 2E-114 | 1.7535918 | 0.162 | 0.052 | 4.9E-110 | Regenerative Mesenchyme | glod5 |
| nup43 | 5E-114 | 1.8708195 | 0.12 | 0.032 | 1.5E-109 | Regenerative Mesenchyme | nup43 |
| ccdc58 | 6E-114 | 1.8796298 | 0.119 | 0.031 | 1.8E-109 | Regenerative Mesenchyme | ccdc58 |
| uggt2 | 6E-114 | 1.8397994 | 0.155 | 0.049 | 2.1E-109 | Regenerative Mesenchyme | uggt2 |
| sec31a | 9E-114 | 1.4060106 | 0.246 | 0.101 | 3E-109 | Regenerative Mesenchyme | sec31a |
| ndufb6 | 2E-113 | 1.1621195 | 0.35 | 0.172 | 5.1E-109 | Regenerative Mesenchyme | ndufb6 |
| ddx3xb | 2E-113 | 1.196748 | 0.318 | 0.148 | 5.9E-109 | Regenerative Mesenchyme | ddx3xb |
| lepa | 2E-113 | 0.8362396 | 0.168 | 0.054 | 6.5E-109 | Regenerative Mesenchyme | lepa |
| ldha | 3E-113 | 1.149995 | 0.406 | 0.22 | 9.2E-109 | Regenerative Mesenchyme | ldha |
| eif4bb | 4E-113 | 1.1242722 | 0.465 | 0.274 | 1.2E-108 | Regenerative Mesenchyme | eif4bb |
| cd63 | 4E-113 | 0.5847083 | 0.873 | 0.825 | 1.3E-108 | Regenerative Mesenchyme | cd63 |
| tufm | 6E-113 | 1.3116192 | 0.277 | 0.121 | 2E-108 | Regenerative Mesenchyme | tufm |
| BX284638.2 | 9E-113 | 1.6161433 | 0.126 | 0.034 | 2.8E-108 | Regenerative Mesenchyme | BX284638.2 |
| ndufb5 | 3E-112 | 1.1738832 | 0.356 | 0.178 | 8.2E-108 | Regenerative Mesenchyme | ndufb5 |
| dpp3 | 6E-112 | 1.982119 | 0.125 | 0.034 | 1.9E-107 | Regenerative Mesenchyme | dpp3 |
| rpl242 | 6E-112 | 0.332494 | 0.998 | 0.993 | 2E-107 | Regenerative Mesenchyme | rpl24 |
| atp5if1b1 | 7E-112 | 0.9879639 | 0.518 | 0.322 | 2.2E-107 | Regenerative Mesenchyme | atp5if1b |
| polr2k | 8E-112 | 1.6103727 | 0.16 | 0.051 | 2.7E-107 | Regenerative Mesenchyme | polr2k |
| psmd12 | 1E-111 | 1.3709059 | 0.255 | 0.108 | 4.3E-107 | Regenerative Mesenchyme | psmd12 |
| coa3a | 2E-111 | 1.2641992 | 0.275 | 0.12 | 7.2E-107 | Regenerative Mesenchyme | coa3a |
| cirbpb1 | 1E-110 | 0.5182723 | 0.895 | 0.83 | 4.3E-106 | Regenerative Mesenchyme | cirbpb |
| tmem167a | 2E-110 | 1.0815952 | 0.431 | 0.243 | 5.4E-106 | Regenerative Mesenchyme | tmem167a |
| prkcsh | 2E-110 | 1.4069059 | 0.282 | 0.127 | 7.2E-106 | Regenerative Mesenchyme | prkcsh |
| rrp15 | 2E-110 | 1.4686465 | 0.175 | 0.06 | 8.1E-106 | Regenerative Mesenchyme | rrp15 |
| srsf1a | 3E-110 | 1.2054414 | 0.331 | 0.16 | 8.3E-106 | Regenerative Mesenchyme | srsf1a |
| nup50 | 4E-110 | 1.5433603 | 0.177 | 0.061 | 1.2E-105 | Regenerative Mesenchyme | nup50 |
| copb1 | 5E-110 | 1.2639508 | 0.299 | 0.138 | 1.5E-105 | Regenerative Mesenchyme | copb1 |
| rps26l2 | 1E-109 | 0.3360131 | 0.989 | 0.974 | 4.5E-105 | Regenerative Mesenchyme | rps26l |
| igfbp1a1 | 2E-109 | 1.1351529 | 0.185 | 0.065 | 6E-105 | Regenerative Mesenchyme | igfbp1a |
| sf3b4 | 2E-109 | 1.9345231 | 0.119 | 0.032 | 6.3E-105 | Regenerative Mesenchyme | sf3b4 |

| | | | | | | | |
|---|---|---|---|---|---|---|---|
| rpl132 | 3E-109 | 0.3099816 | 0.999 | 0.993 | 1.1E-104 | Regenerative Mesenchyme | rpl13 |
| rpl142 | 4E-109 | 0.2791274 | 0.996 | 0.996 | 1.3E-104 | Regenerative Mesenchyme | rpl14 |
| slc31a11 | 4E-109 | 1.1481324 | 0.366 | 0.188 | 1.3E-104 | Regenerative Mesenchyme | slc31a1 |
| eef2b2 | 7E-109 | 0.4104757 | 0.962 | 0.906 | 2.4E-104 | Regenerative Mesenchyme | eef2b |
| cse1l | 1E-108 | 1.5541781 | 0.167 | 0.056 | 3.2E-104 | Regenerative Mesenchyme | cse1l |
| mt-nd41 | 2E-108 | 0.5393491 | 0.929 | 0.852 | 5.2E-104 | Regenerative Mesenchyme | mt-nd4 |
| ier3ip11 | 2E-108 | 1.1431405 | 0.409 | 0.226 | 7.5E-104 | Regenerative Mesenchyme | ier3ip1 |
| ctps1a | 3E-108 | 1.6276754 | 0.151 | 0.048 | 9.4E-104 | Regenerative Mesenchyme | ctps1a |
| mrpl22 | 4E-108 | 1.9523431 | 0.106 | 0.027 | 1.4E-103 | Regenerative Mesenchyme | mrpl22 |
| CU467861.6 | 5E-108 | 1.420687 | 0.243 | 0.101 | 1.8E-103 | Regenerative Mesenchyme | CU467861.6 |
| ube2v2 | 2E-107 | 0.7171252 | 0.693 | 0.521 | 5.8E-103 | Regenerative Mesenchyme | ube2v2 |
| fam210aa | 2E-107 | 1.3968838 | 0.2 | 0.074 | 6.3E-103 | Regenerative Mesenchyme | fam210aa |
| abi1b | 2E-107 | 1.9020389 | 0.11 | 0.028 | 7.6E-103 | Regenerative Mesenchyme | abi1b |
| eif2s1a | 4E-107 | 1.2231826 | 0.294 | 0.135 | 1.3E-102 | Regenerative Mesenchyme | eif2s1a |
| ppp5c | 6E-107 | 1.749597 | 0.129 | 0.037 | 2.1E-102 | Regenerative Mesenchyme | ppp5c |
| hmgb3b1 | 1E-106 | 1.3293596 | 0.255 | 0.109 | 3.7E-102 | Regenerative Mesenchyme | hmgb3b |
| nom1 | 3E-106 | 1.7340539 | 0.129 | 0.037 | 9.9E-102 | Regenerative Mesenchyme | nom1 |
| h2ax | 1E-105 | 1.4912241 | 0.18 | 0.064 | 4E-101 | Regenerative Mesenchyme | h2ax |
| mphosph10 | 3E-105 | 1.2707523 | 0.23 | 0.093 | 8.7E-101 | Regenerative Mesenchyme | mphosph10 |
| ndufb4 | 4E-105 | 0.9521521 | 0.504 | 0.315 | 1.4E-100 | Regenerative Mesenchyme | ndufb4 |
| srsf3a.1 | 5E-105 | 1.417121 | 0.202 | 0.076 | 1.7E-100 | Regenerative Mesenchyme | srsf3a.1 |
| polr3k | 6E-105 | 1.6707735 | 0.149 | 0.048 | 2E-100 | Regenerative Mesenchyme | polr3k |
| ndufb9 | 2E-104 | 0.9441879 | 0.489 | 0.299 | 5.7E-100 | Regenerative Mesenchyme | ndufb9 |
| tardbp | 3E-104 | 1.1106279 | 0.316 | 0.151 | 8.9E-100 | Regenerative Mesenchyme | tardbp |
| mesd | 3E-104 | 1.1615086 | 0.302 | 0.141 | 1E-99 | Regenerative Mesenchyme | mesd |
| clta | 3E-104 | 1.1812038 | 0.296 | 0.138 | 1.1E-99 | Regenerative Mesenchyme | clta |
| txnl1 | 3E-104 | 1.4353062 | 0.198 | 0.075 | 1.1E-99 | Regenerative Mesenchyme | txnl1 |
| rer1 | 4E-104 | 1.066041 | 0.375 | 0.196 | 1.3E-99 | Regenerative Mesenchyme | rer1 |
| cars1 | 2E-103 | 1.5468835 | 0.196 | 0.074 | 6.8E-99 | Regenerative Mesenchyme | cars1 |
| mrps18a | 4E-103 | 1.5706481 | 0.159 | 0.053 | 1.23E-98 | Regenerative Mesenchyme | mrps18a |
| cyb5b | 6E-103 | 1.8614717 | 0.113 | 0.03 | 2.1E-98 | Regenerative Mesenchyme | cyb5b |
| copz11 | 8E-103 | 1.2064458 | 0.3 | 0.141 | 2.54E-98 | Regenerative Mesenchyme | copz1 |
| nomo | 1E-102 | 1.8536827 | 0.134 | 0.041 | 3.14E-98 | Regenerative Mesenchyme | nomo |
| rpl42 | 1E-102 | 0.4121275 | 0.935 | 0.894 | 4.21E-98 | Regenerative Mesenchyme | rpl4 |
| NC-002333.42 | 1E-102 | 0.4146509 | 1 | 1 | 4.43E-98 | Regenerative Mesenchyme | NC-002333.4 |
| ppan | 1E-102 | 1.2885543 | 0.245 | 0.104 | 4.48E-98 | Regenerative Mesenchyme | ppan |
| surf4l | 1E-102 | 1.3213179 | 0.265 | 0.118 | 4.57E-98 | Regenerative Mesenchyme | surf4l |
| ddx181 | 2E-102 | 1.2893922 | 0.225 | 0.092 | 5.79E-98 | Regenerative Mesenchyme | ddx18 |
| wdr3 | 2E-102 | 1.4774481 | 0.187 | 0.069 | 7.8E-98 | Regenerative Mesenchyme | wdr3 |
| usp10 | 2E-102 | 1.0259359 | 0.423 | 0.239 | 8.03E-98 | Regenerative Mesenchyme | usp10 |
| commd9 | 1E-101 | 1.4912935 | 0.168 | 0.058 | 3.14E-97 | Regenerative Mesenchyme | commd9 |
| zgc:153867 | 1E-101 | 0.5455511 | 0.853 | 0.747 | 3.26E-97 | Regenerative Mesenchyme | zgc:153867 |
| copb2 | 1E-101 | 1.0598633 | 0.441 | 0.26 | 4.31E-97 | Regenerative Mesenchyme | copb2 |
| eef1b22 | 3E-101 | 0.3875187 | 0.964 | 0.927 | 9.31E-97 | Regenerative Mesenchyme | eef1b2 |
| ndufa81 | 3E-101 | 1.0639844 | 0.4 | 0.223 | 9.58E-97 | Regenerative Mesenchyme | ndufa8 |
| srsf3b | 3E-101 | 0.9148539 | 0.457 | 0.263 | 1.09E-96 | Regenerative Mesenchyme | srsf3b |
| ywhae1 | 3E-101 | 0.9111402 | 0.514 | 0.319 | 1.1E-96 | Regenerative Mesenchyme | ywhae1 |
| calr3a1 | 4E-101 | 0.8726208 | 0.547 | 0.352 | 1.16E-96 | Regenerative Mesenchyme | calr3a |

| | | | | | | | |
|---|---|---|---|---|---|---|---|
| mrpl53 | 4E-101 | 1.5091286 | 0.133 | 0.04 | 1.3E-96 | Regenerative Mesenchyme | mrpl53 |
| ndufa5 | 5E-101 | 0.9919378 | 0.378 | 0.2 | 1.57E-96 | Regenerative Mesenchyme | ndufa5 |
| fcf1 | 5E-101 | 1.4659432 | 0.159 | 0.054 | 1.62E-96 | Regenerative Mesenchyme | fcf1 |
| emg1 | 7E-101 | 1.7040691 | 0.13 | 0.039 | 2.24E-96 | Regenerative Mesenchyme | emg1 |
| psmd6 | 8E-101 | 1.2765202 | 0.255 | 0.112 | 2.53E-96 | Regenerative Mesenchyme | psmd6 |
| btf3l41 | 1E-100 | 1.065303 | 0.373 | 0.199 | 3.92E-96 | Regenerative Mesenchyme | btf3l4 |
| esf1 | 2E-100 | 1.0389824 | 0.314 | 0.15 | 5.01E-96 | Regenerative Mesenchyme | esf1 |
| psme1 | 2E-100 | 1.1831868 | 0.247 | 0.105 | 5.76E-96 | Regenerative Mesenchyme | psme1 |
| ywhah | 2E-100 | 1.1342346 | 0.313 | 0.152 | 6.12E-96 | Regenerative Mesenchyme | ywhah |
| mrps18c | 2E-100 | 1.3776754 | 0.193 | 0.073 | 6.65E-96 | Regenerative Mesenchyme | mrps18c |
| rrp7a | 6E-100 | 1.7926198 | 0.11 | 0.03 | 1.79E-95 | Regenerative Mesenchyme | rrp7a |
| spcs3 | 7E-100 | 1.0125223 | 0.443 | 0.26 | 2.4E-95 | Regenerative Mesenchyme | spcs3 |
| yeats4 | 8E-100 | 1.4406466 | 0.177 | 0.064 | 2.45E-95 | Regenerative Mesenchyme | yeats4 |
| sub1b | 9E-100 | 1.0091348 | 0.451 | 0.268 | 2.81E-95 | Regenerative Mesenchyme | sub1b |
| rpl282 | 1E-99 | 0.2992832 | 0.998 | 0.991 | 4.04E-95 | Regenerative Mesenchyme | rpl28 |
| fam162a1 | 2E-99 | 1.0830129 | 0.404 | 0.229 | 5.9E-95 | Regenerative Mesenchyme | fam162a |
| smarca4a | 4E-99 | 1.3332447 | 0.218 | 0.088 | 1.21E-94 | Regenerative Mesenchyme | smarca4a |
| copg2 | 1E-98 | 1.2507108 | 0.294 | 0.141 | 3.19E-94 | Regenerative Mesenchyme | copg2 |
| mrps10 | 1E-98 | 1.3056323 | 0.239 | 0.103 | 3.35E-94 | Regenerative Mesenchyme | mrps10 |
| clpp | 1.1E-98 | 1.445556 | 0.19 | 0.072 | 3.54E-94 | Regenerative Mesenchyme | clpp |
| si:ch73-308l14.2 | 1.7E-98 | 1.0538753 | 0.241 | 0.102 | 5.38E-94 | Regenerative Mesenchyme | si:ch73-308l14.2 |
| cd3eap | 3.6E-98 | 1.1589297 | 0.254 | 0.112 | 1.18E-93 | Regenerative Mesenchyme | cd3eap |
| mrpl9 | 3.7E-98 | 1.5947053 | 0.128 | 0.039 | 1.22E-93 | Regenerative Mesenchyme | mrpl9 |
| mrpl39 | 3.9E-98 | 1.6629003 | 0.121 | 0.035 | 1.26E-93 | Regenerative Mesenchyme | mrpl39 |
| nudc | 3.9E-98 | 0.9787315 | 0.344 | 0.175 | 1.27E-93 | Regenerative Mesenchyme | nudc |
| si:ch211-217k17.71 | 4.1E-98 | 0.9946342 | 0.453 | 0.272 | 1.33E-93 | Regenerative Mesenchyme | si:ch211-217k17.7 |
| txndc17 | 5.9E-98 | 1.2411422 | 0.252 | 0.111 | 1.91E-93 | Regenerative Mesenchyme | txndc17 |
| zgc:193541 | 6.7E-98 | 0.8752834 | 0.566 | 0.384 | 2.18E-93 | Regenerative Mesenchyme | zgc:193541 |
| psmd11a | 7E-98 | 1.2851318 | 0.215 | 0.087 | 2.26E-93 | Regenerative Mesenchyme | psmd11a |
| setb | 1.1E-97 | 0.9474256 | 0.541 | 0.366 | 3.49E-93 | Regenerative Mesenchyme | setb |
| mrpl46 | 1.1E-97 | 1.5277965 | 0.137 | 0.043 | 3.5E-93 | Regenerative Mesenchyme | mrpl46 |
| BX908782.2 | 1.2E-97 | 0.342199 | 0.153 | 0.051 | 3.97E-93 | Regenerative Mesenchyme | BX908782.2 |
| pdcd11 | 4.8E-97 | 1.2524 | 0.242 | 0.105 | 1.55E-92 | Regenerative Mesenchyme | pdcd11 |
| cmss1 | 6.7E-97 | 1.5503525 | 0.149 | 0.05 | 2.18E-92 | Regenerative Mesenchyme | cmss1 |
| rrp12 | 7.6E-97 | 1.6179178 | 0.167 | 0.06 | 2.49E-92 | Regenerative Mesenchyme | rrp12 |
| ubr7 | 7.7E-97 | 1.3232079 | 0.214 | 0.087 | 2.51E-92 | Regenerative Mesenchyme | ubr7 |
| meox11 | 9E-97 | 1.3947357 | 0.163 | 0.057 | 2.92E-92 | Regenerative Mesenchyme | meox1 |
| prdx6 | 9.1E-97 | 0.8832898 | 0.514 | 0.334 | 2.97E-92 | Regenerative Mesenchyme | prdx6 |
| exosc2 | 1E-96 | 1.7019614 | 0.114 | 0.032 | 3.34E-92 | Regenerative Mesenchyme | exosc2 |
| hdac1 | 1.4E-96 | 1.0086741 | 0.445 | 0.262 | 4.6E-92 | Regenerative Mesenchyme | hdac1 |
| apex1 | 1.4E-96 | 1.8737514 | 0.113 | 0.032 | 4.67E-92 | Regenerative Mesenchyme | apex1 |
| mrpl301 | 2E-96 | 1.2605223 | 0.236 | 0.101 | 6.5E-92 | Regenerative Mesenchyme | mrpl30 |
| mrps16 | 2.2E-96 | 1.4432482 | 0.188 | 0.072 | 7.23E-92 | Regenerative Mesenchyme | mrps16 |
| C7H16orf871 | 4.3E-96 | 1.2047544 | 0.262 | 0.119 | 1.41E-91 | Regenerative Mesenchyme | C7H16orf87 |
| lypla2 | 6E-96 | 1.69449 | 0.149 | 0.05 | 1.96E-91 | Regenerative Mesenchyme | lypla2 |
| mtdha | 6.1E-96 | 0.7619106 | 0.692 | 0.556 | 1.97E-91 | Regenerative Mesenchyme | mtdha |
| eif2a | 6.2E-96 | 1.4096919 | 0.195 | 0.077 | 2.03E-91 | Regenerative Mesenchyme | eif2a |
| si:ch211-66k16.27 | 1.2E-95 | 1.417891 | 0.198 | 0.078 | 3.85E-91 | Regenerative Mesenchyme | si:ch211-66k16.27 |

| | | | | | | | |
|---|---|---|---|---|---|---|---|
| mak16 | 1.3E-95 | 1.3251968 | 0.223 | 0.094 | 4.12E-91 | Regenerative Mesenchyme | mak16 |
| STMP11 | 3.1E-95 | 1.0014425 | 0.423 | 0.248 | 1.01E-90 | Regenerative Mesenchyme | STMP1 |
| sf3b3 | 4.7E-95 | 1.3888288 | 0.198 | 0.078 | 1.52E-90 | Regenerative Mesenchyme | sf3b3 |
| si:dkey-151g10.62 | 4.8E-95 | 0.258244 | 0.999 | 0.999 | 1.57E-90 | Regenerative Mesenchyme | si:dkey-151g10.6 |
| cdh1 | 5.1E-95 | 2.3190542 | 0.119 | 0.036 | 1.65E-90 | Regenerative Mesenchyme | cdh1 |
| srsf10b1 | 5.2E-95 | 1.2896494 | 0.201 | 0.08 | 1.69E-90 | Regenerative Mesenchyme | srsf10b |
| mydgf1 | 7.2E-95 | 1.4766194 | 0.189 | 0.074 | 2.34E-90 | Regenerative Mesenchyme | mydgf |
| lsp1a1 | 8.1E-95 | 1.0712337 | 0.37 | 0.2 | 2.62E-90 | Regenerative Mesenchyme | lsp1a |
| rps92 | 2E-94 | 0.3057111 | 0.995 | 0.987 | 6.42E-90 | Regenerative Mesenchyme | rps9 |
| uqcrc2a1 | 2.3E-94 | 1.3673139 | 0.225 | 0.096 | 7.37E-90 | Regenerative Mesenchyme | uqcrc2a |
| polr2f | 2.3E-94 | 1.005311 | 0.34 | 0.175 | 7.62E-90 | Regenerative Mesenchyme | polr2f |
| vdac22 | 4.5E-94 | 0.802024 | 0.653 | 0.512 | 1.48E-89 | Regenerative Mesenchyme | vdac2 |
| ipo7 | 7E-94 | 1.0331162 | 0.413 | 0.238 | 2.28E-89 | Regenerative Mesenchyme | ipo7 |
| pdia31 | 7.5E-94 | 0.6093981 | 0.789 | 0.68 | 2.44E-89 | Regenerative Mesenchyme | pdia3 |
| coil | 1.1E-93 | 1.3412225 | 0.143 | 0.047 | 3.49E-89 | Regenerative Mesenchyme | coil |
| sar1b | 1.2E-93 | 0.9348751 | 0.47 | 0.29 | 4E-89 | Regenerative Mesenchyme | sar1b |
| dnajc21 | 1.7E-93 | 1.7043887 | 0.104 | 0.028 | 5.56E-89 | Regenerative Mesenchyme | dnajc21 |
| pcolce2b1 | 2.3E-93 | 0.7013783 | 0.483 | 0.286 | 7.36E-89 | Regenerative Mesenchyme | pcolce2b |
| mtdhb | 2.5E-93 | 1.0588202 | 0.299 | 0.145 | 8.14E-89 | Regenerative Mesenchyme | mtdhb |
| rpsa2 | 3.8E-93 | 0.3128196 | 0.994 | 0.984 | 1.23E-88 | Regenerative Mesenchyme | rpsa |
| si:ch211-198c19.3 | 3.9E-93 | 0.5113311 | 0.384 | 0.201 | 1.27E-88 | Regenerative Mesenchyme | si:ch211-198c19.3 |
| mrpl24 | 3.9E-93 | 1.7054143 | 0.148 | 0.051 | 1.28E-88 | Regenerative Mesenchyme | mrpl24 |
| nat10 | 5.3E-93 | 1.3452146 | 0.161 | 0.057 | 1.71E-88 | Regenerative Mesenchyme | nat10 |
| ufc11 | 5.5E-93 | 1.1773536 | 0.269 | 0.125 | 1.79E-88 | Regenerative Mesenchyme | ufc1 |
| zc3h15 | 5.9E-93 | 1.1239117 | 0.281 | 0.133 | 1.92E-88 | Regenerative Mesenchyme | zc3h15 |
| psmd10 | 5.9E-93 | 1.3093365 | 0.164 | 0.059 | 1.93E-88 | Regenerative Mesenchyme | psmd10 |
| prdx31 | 6.5E-93 | 1.139355 | 0.28 | 0.133 | 2.11E-88 | Regenerative Mesenchyme | prdx3 |
| ddx611 | 1E-92 | 0.8187997 | 0.555 | 0.371 | 3.38E-88 | Regenerative Mesenchyme | ddx61 |
| arf2a | 1.9E-92 | 1.0267811 | 0.408 | 0.234 | 6.06E-88 | Regenerative Mesenchyme | arf2a |
| snrpa | 2.4E-92 | 1.1990071 | 0.23 | 0.099 | 7.73E-88 | Regenerative Mesenchyme | snrpa |
| nifk1 | 4.3E-92 | 1.3211828 | 0.203 | 0.082 | 1.38E-87 | Regenerative Mesenchyme | nifk |
| lgals2a2 | 7.6E-92 | 0.4716477 | 0.797 | 0.632 | 2.48E-87 | Regenerative Mesenchyme | lgals2a |
| sec23a | 9.7E-92 | 1.2147003 | 0.287 | 0.139 | 3.15E-87 | Regenerative Mesenchyme | sec23a |
| tes | 1.1E-91 | 1.4830206 | 0.179 | 0.069 | 3.55E-87 | Regenerative Mesenchyme | tes |
| pkma | 1.2E-91 | 1.1299543 | 0.368 | 0.204 | 3.85E-87 | Regenerative Mesenchyme | pkma |
| zgc:171480 | 1.5E-91 | 1.1741916 | 0.221 | 0.093 | 4.88E-87 | Regenerative Mesenchyme | zgc:171480 |
| itpa | 2.4E-91 | 1.8453531 | 0.109 | 0.031 | 7.95E-87 | Regenerative Mesenchyme | itpa |
| dnajc11 | 2.5E-91 | 0.923703 | 0.454 | 0.273 | 8.09E-87 | Regenerative Mesenchyme | dnajc1 |
| gnpnat11 | 8.5E-91 | 1.5144984 | 0.202 | 0.083 | 2.76E-86 | Regenerative Mesenchyme | gnpnat1 |
| ndufa10 | 9.8E-91 | 1.3732515 | 0.245 | 0.112 | 3.19E-86 | Regenerative Mesenchyme | ndufa10 |
| triap1 | 1.3E-90 | 1.5100195 | 0.115 | 0.034 | 4.23E-86 | Regenerative Mesenchyme | triap1 |
| hadhb | 1.5E-90 | 1.1997084 | 0.264 | 0.124 | 4.81E-86 | Regenerative Mesenchyme | hadhb |
| mrps11 | 1.9E-90 | 1.3682078 | 0.21 | 0.089 | 6.16E-86 | Regenerative Mesenchyme | mrps11 |
| p4hb1 | 2.4E-90 | 0.8352228 | 0.717 | 0.566 | 7.65E-86 | Regenerative Mesenchyme | p4hb |
| smfn | 2.5E-90 | 1.4485452 | 0.154 | 0.055 | 8.16E-86 | Regenerative Mesenchyme | smfn |
| calr | 3.5E-90 | 1.0768813 | 0.352 | 0.189 | 1.12E-85 | Regenerative Mesenchyme | calr |
| htra1a2 | 4.8E-90 | 1.0685864 | 0.383 | 0.219 | 1.57E-85 | Regenerative Mesenchyme | htra1a |
| chchd2 | 5.7E-90 | 1.298723 | 0.21 | 0.087 | 1.84E-85 | Regenerative Mesenchyme | chchd2 |

| | | | | | | | |
|---|---|---|---|---|---|---|---|
| tspan361 | 7.2E-90 | 1.0328294 | 0.306 | 0.154 | 2.33E-85 | Regenerative Mesenchyme | tspan36 |
| mrps341 | 1.1E-89 | 1.491424 | 0.151 | 0.053 | 3.52E-85 | Regenerative Mesenchyme | mrps34 |
| her92 | 1.2E-89 | 1.0255309 | 0.386 | 0.219 | 3.81E-85 | Regenerative Mesenchyme | her9 |
| psmc1b | 1.5E-89 | 1.3610479 | 0.199 | 0.081 | 4.95E-85 | Regenerative Mesenchyme | psmc1b |
| crabp2a1 | 2.5E-89 | 0.7127687 | 0.312 | 0.153 | 8.06E-85 | Regenerative Mesenchyme | crabp2a |
| hirip3 | 3.5E-89 | 1.1317973 | 0.257 | 0.119 | 1.13E-84 | Regenerative Mesenchyme | hirip3 |
| atp5f1e1 | 4.8E-89 | 0.8423417 | 0.573 | 0.404 | 1.55E-84 | Regenerative Mesenchyme | atp5f1e |
| gfpt1 | 8.6E-89 | 1.4888174 | 0.15 | 0.053 | 2.81E-84 | Regenerative Mesenchyme | gfpt1 |
| tmed9 | 9.4E-89 | 0.9974661 | 0.388 | 0.218 | 3.05E-84 | Regenerative Mesenchyme | tmed9 |
| mrpl541 | 1.2E-88 | 1.2880306 | 0.206 | 0.086 | 3.88E-84 | Regenerative Mesenchyme | mrpl54 |
| exosc8 | 2.1E-88 | 1.58081 | 0.111 | 0.033 | 6.81E-84 | Regenerative Mesenchyme | exosc8 |
| dnajc19 | 2.6E-88 | 1.4043564 | 0.159 | 0.058 | 8.46E-84 | Regenerative Mesenchyme | dnajc19 |
| ubqln4 | 3.1E-88 | 1.029098 | 0.278 | 0.132 | 9.93E-84 | Regenerative Mesenchyme | ubqln4 |
| psmd4b | 3.5E-88 | 1.0957664 | 0.286 | 0.139 | 1.13E-83 | Regenerative Mesenchyme | psmd4b |
| wdr18 | 3.9E-88 | 1.5846746 | 0.126 | 0.04 | 1.27E-83 | Regenerative Mesenchyme | wdr18 |
| dpcd | 5.8E-88 | 1.6485971 | 0.105 | 0.03 | 1.89E-83 | Regenerative Mesenchyme | dpcd |
| rtcb | 8E-88 | 1.6241282 | 0.133 | 0.044 | 2.59E-83 | Regenerative Mesenchyme | rtcb |
| pfdn2 | 8E-88 | 1.373901 | 0.171 | 0.065 | 2.59E-83 | Regenerative Mesenchyme | pfdn2 |
| copa | 1.3E-87 | 1.0911581 | 0.338 | 0.181 | 4.07E-83 | Regenerative Mesenchyme | copa |
| mphosph6 | 1.8E-87 | 1.3881733 | 0.169 | 0.064 | 5.9E-83 | Regenerative Mesenchyme | mphosph6 |
| ccdc43 | 2.2E-87 | 1.5678235 | 0.128 | 0.041 | 7.11E-83 | Regenerative Mesenchyme | ccdc43 |
| si:ch73-281n10.2 | 2.2E-87 | 1.1139912 | 0.577 | 0.434 | 7.23E-83 | Regenerative Mesenchyme | si:ch73-281n10.2 |
| uchl5 | 3.9E-87 | 1.3433914 | 0.18 | 0.071 | 1.28E-82 | Regenerative Mesenchyme | uchl5 |
| cbx5 | 5.4E-87 | 0.9904241 | 0.384 | 0.217 | 1.74E-82 | Regenerative Mesenchyme | cbx5 |
| cfl21 | 1.3E-86 | 1.6252169 | 0.146 | 0.052 | 4.32E-82 | Regenerative Mesenchyme | cfl2 |
| mrpl45 | 1.3E-86 | 1.5877045 | 0.131 | 0.043 | 4.33E-82 | Regenerative Mesenchyme | mrpl45 |
| ccdc59 | 1.7E-86 | 1.3817447 | 0.151 | 0.054 | 5.69E-82 | Regenerative Mesenchyme | ccdc59 |
| hspa14 | 2.3E-86 | 1.4514265 | 0.145 | 0.05 | 7.56E-82 | Regenerative Mesenchyme | hspa14 |
| rpl62 | 2.5E-86 | 0.3065447 | 0.976 | 0.958 | 8.02E-82 | Regenerative Mesenchyme | rpl6 |
| serp12 | 2.7E-86 | 0.9685629 | 0.473 | 0.306 | 8.84E-82 | Regenerative Mesenchyme | serp1 |
| uba5 | 2.8E-86 | 1.2101399 | 0.227 | 0.1 | 9.25E-82 | Regenerative Mesenchyme | uba5 |
| ndufs21 | 3.5E-86 | 1.3627964 | 0.234 | 0.106 | 1.13E-81 | Regenerative Mesenchyme | ndufs2 |
| myl6 | 6.4E-86 | 1.1366712 | 0.365 | 0.207 | 2.07E-81 | Regenerative Mesenchyme | myl6 |
| elavl1a | 6.9E-86 | 1.12024 | 0.244 | 0.111 | 2.24E-81 | Regenerative Mesenchyme | elavl1a |
| inip | 8.1E-86 | 1.3017004 | 0.175 | 0.067 | 2.63E-81 | Regenerative Mesenchyme | inip |
| tfam | 9.2E-86 | 1.3247192 | 0.184 | 0.073 | 3E-81 | Regenerative Mesenchyme | tfam |
| rrp9 | 9.5E-86 | 1.5742849 | 0.112 | 0.034 | 3.09E-81 | Regenerative Mesenchyme | rrp9 |
| oxsr1a | 1.1E-85 | 0.9429442 | 0.366 | 0.201 | 3.44E-81 | Regenerative Mesenchyme | oxsr1a |
| eftud2 | 1.4E-85 | 1.5763467 | 0.113 | 0.034 | 4.62E-81 | Regenerative Mesenchyme | eftud2 |
| hdlbpa1 | 1.5E-85 | 0.6437902 | 0.772 | 0.67 | 5.02E-81 | Regenerative Mesenchyme | hdlbpa |
| eif2b1 | 2.4E-85 | 1.8155922 | 0.112 | 0.034 | 7.93E-81 | Regenerative Mesenchyme | eif2b1 |
| rab41 | 2.5E-85 | 1.3330419 | 0.188 | 0.076 | 8.16E-81 | Regenerative Mesenchyme | rab41 |
| slc43a1a1 | 2.7E-85 | 1.3667616 | 0.188 | 0.076 | 8.91E-81 | Regenerative Mesenchyme | slc43a1a |
| phgdh | 4E-85 | 1.6251968 | 0.118 | 0.037 | 1.3E-80 | Regenerative Mesenchyme | phgdh |
| pfdn1 | 4.3E-85 | 1.0082829 | 0.344 | 0.186 | 1.39E-80 | Regenerative Mesenchyme | pfdn1 |
| tbl2 | 5E-85 | 1.5303681 | 0.136 | 0.046 | 1.63E-80 | Regenerative Mesenchyme | tbl2 |
| ndufa3 | 5.2E-85 | 1.0536584 | 0.276 | 0.133 | 1.69E-80 | Regenerative Mesenchyme | ndufa3 |
| dynlrb1.1 | 5.3E-85 | 0.8730579 | 0.481 | 0.308 | 1.73E-80 | Regenerative Mesenchyme | dynlrb1.1 |

| | | | | | | | |
|---|---|---|---|---|---|---|---|
| lars1b | 6.3E-85 | 1.5777679 | 0.125 | 0.041 | 2.05E-80 | Regenerative Mesenchyme | lars1b |
| bcas2 | 6.7E-85 | 1.0321391 | 0.315 | 0.163 | 2.17E-80 | Regenerative Mesenchyme | bcas2 |
| nol71 | 1.6E-84 | 1.3334003 | 0.161 | 0.06 | 5.06E-80 | Regenerative Mesenchyme | nol7 |
| chd7 | 1.7E-84 | 1.8302756 | 0.116 | 0.037 | 5.4E-80 | Regenerative Mesenchyme | chd7 |
| baz1b | 2E-84 | 1.049783 | 0.32 | 0.166 | 6.54E-80 | Regenerative Mesenchyme | baz1b |
| macf1a | 2.7E-84 | 1.021674 | 0.32 | 0.168 | 8.74E-80 | Regenerative Mesenchyme | macf1a |
| lbr1 | 3.7E-84 | 1.5869529 | 0.167 | 0.065 | 1.21E-79 | Regenerative Mesenchyme | lbr |
| utp111 | 5.7E-84 | 1.3210602 | 0.161 | 0.06 | 1.86E-79 | Regenerative Mesenchyme | utp11 |
| ndufa111 | 1.2E-83 | 1.0889489 | 0.265 | 0.127 | 3.9E-79 | Regenerative Mesenchyme | ndufa11 |
| rack12 | 1.6E-83 | 0.3324572 | 0.974 | 0.947 | 5.13E-79 | Regenerative Mesenchyme | rack1 |
| phb2a | 2.5E-83 | 1.1608722 | 0.231 | 0.105 | 8.12E-79 | Regenerative Mesenchyme | phb2a |
| gadd45bb1 | 2.6E-83 | 0.7998635 | 0.377 | 0.211 | 8.39E-79 | Regenerative Mesenchyme | gadd45bb |
| rfc3 | 3.2E-83 | 2.0526344 | 0.1 | 0.029 | 1.03E-78 | Regenerative Mesenchyme | rfc3 |
| mrpl361 | 5.1E-83 | 1.1705701 | 0.267 | 0.131 | 1.66E-78 | Regenerative Mesenchyme | mrpl36 |
| ptges3a | 9.8E-83 | 1.1348792 | 0.249 | 0.116 | 3.17E-78 | Regenerative Mesenchyme | ptges3a |
| nle1 | 1E-82 | 1.6205993 | 0.125 | 0.041 | 3.24E-78 | Regenerative Mesenchyme | nle1 |
| wdr83os | 1.3E-82 | 1.0094108 | 0.329 | 0.175 | 4.38E-78 | Regenerative Mesenchyme | wdr83os |
| tmco11 | 1.4E-82 | 1.3615428 | 0.185 | 0.076 | 4.55E-78 | Regenerative Mesenchyme | tmco1 |
| cnpy21 | 1.4E-82 | 1.7903872 | 0.123 | 0.041 | 4.56E-78 | Regenerative Mesenchyme | cnpy2 |
| tmed7 | 1.5E-82 | 0.8693989 | 0.479 | 0.307 | 5.02E-78 | Regenerative Mesenchyme | tmed7 |
| snd11 | 1.6E-82 | 1.0449613 | 0.392 | 0.232 | 5.28E-78 | Regenerative Mesenchyme | snd1 |
| tpm31 | 1.7E-82 | 0.6814946 | 0.668 | 0.518 | 5.57E-78 | Regenerative Mesenchyme | tpm3 |
| abcf2a1 | 2.4E-82 | 1.0120357 | 0.322 | 0.17 | 7.88E-78 | Regenerative Mesenchyme | abcf2a |
| ndufb2 | 3.3E-82 | 0.8987863 | 0.412 | 0.246 | 1.08E-77 | Regenerative Mesenchyme | ndufb2 |
| tmem263 | 4E-82 | 1.1872502 | 0.288 | 0.147 | 1.29E-77 | Regenerative Mesenchyme | tmem263 |
| rida | 4.8E-82 | 1.4722745 | 0.184 | 0.076 | 1.57E-77 | Regenerative Mesenchyme | rida |
| pam16 | 4.8E-82 | 1.1413847 | 0.228 | 0.103 | 1.58E-77 | Regenerative Mesenchyme | pam16 |
| gng5 | 6.4E-82 | 0.850182 | 0.425 | 0.254 | 2.07E-77 | Regenerative Mesenchyme | gng5 |
| srprb1 | 8.3E-82 | 1.1345732 | 0.321 | 0.173 | 2.71E-77 | Regenerative Mesenchyme | srprb |
| surf6 | 1.1E-81 | 1.5956883 | 0.106 | 0.032 | 3.7E-77 | Regenerative Mesenchyme | surf6 |
| ndufs3 | 1.3E-81 | 1.1612113 | 0.256 | 0.123 | 4.29E-77 | Regenerative Mesenchyme | ndufs3 |
| mapre1a | 1.8E-81 | 1.5642526 | 0.134 | 0.046 | 5.76E-77 | Regenerative Mesenchyme | mapre1a |
| dnajc8 | 2.5E-81 | 1.207986 | 0.196 | 0.082 | 8.21E-77 | Regenerative Mesenchyme | dnajc8 |
| ywhae2 | 2.8E-81 | 0.975453 | 0.276 | 0.135 | 8.97E-77 | Regenerative Mesenchyme | ywhae2 |
| u2af1 | 5.4E-81 | 1.0575847 | 0.315 | 0.166 | 1.75E-76 | Regenerative Mesenchyme | u2af1 |
| rpf2 | 6.8E-81 | 1.2913162 | 0.193 | 0.081 | 2.2E-76 | Regenerative Mesenchyme | rpf2 |
| myef2 | 8.3E-81 | 1.1519494 | 0.213 | 0.093 | 2.71E-76 | Regenerative Mesenchyme | myef2 |
| foxp42 | 1.1E-80 | 0.9153831 | 0.351 | 0.195 | 3.63E-76 | Regenerative Mesenchyme | foxp4 |
| exosc6 | 1.3E-80 | 1.6124229 | 0.109 | 0.034 | 4.33E-76 | Regenerative Mesenchyme | exosc6 |
| rtraf | 1.4E-80 | 1.1103254 | 0.221 | 0.099 | 4.6E-76 | Regenerative Mesenchyme | rtraf |
| shmt1 | 1.5E-80 | 1.7761205 | 0.119 | 0.039 | 4.92E-76 | Regenerative Mesenchyme | shmt1 |
| prelid1a | 2.2E-80 | 1.1657192 | 0.247 | 0.118 | 7.25E-76 | Regenerative Mesenchyme | prelid1a |
| ube2d2l1 | 2.4E-80 | 1.0734806 | 0.281 | 0.141 | 7.7E-76 | Regenerative Mesenchyme | ube2d2l |
| eif3ha2 | 3E-80 | 0.8084938 | 0.563 | 0.413 | 9.76E-76 | Regenerative Mesenchyme | eif3ha |
| mpdu1b | 3.3E-80 | 1.4727696 | 0.128 | 0.043 | 1.06E-75 | Regenerative Mesenchyme | mpdu1b |
| lsm12a | 3.5E-80 | 1.3088136 | 0.172 | 0.069 | 1.15E-75 | Regenerative Mesenchyme | lsm12a |
| sulf2b | 3.7E-80 | 1.0447169 | 0.301 | 0.157 | 1.22E-75 | Regenerative Mesenchyme | sulf2b |
| mrpl35 | 3.8E-80 | 1.4474167 | 0.143 | 0.051 | 1.24E-75 | Regenerative Mesenchyme | mrpl35 |

| | | | | | | | |
|---|---|---|---|---|---|---|---|
| mrps15 | 4.5E-80 | 1.6182654 | 0.121 | 0.04 | 1.46E-75 | Regenerative Mesenchyme | mrps15 |
| hnrnpub | 8.8E-80 | 0.8489055 | 0.486 | 0.312 | 2.87E-75 | Regenerative Mesenchyme | hnrnpub |
| pelo | 1.3E-79 | 1.7910585 | 0.101 | 0.03 | 4.34E-75 | Regenerative Mesenchyme | pelo |
| aars11 | 1.5E-79 | 1.1863284 | 0.263 | 0.131 | 4.93E-75 | Regenerative Mesenchyme | aars1 |
| si:dkey-88p24.11 | 1.9E-79 | 1.1535828 | 0.184 | 0.076 | 6.33E-75 | Regenerative Mesenchyme | si:dkey-88p24.11 |
| dpm1 | 2.1E-79 | 1.1970232 | 0.192 | 0.081 | 6.92E-75 | Regenerative Mesenchyme | dpm1 |
| psmg1 | 3E-79 | 1.1211743 | 0.226 | 0.103 | 9.61E-75 | Regenerative Mesenchyme | psmg1 |
| exosc5 | 4.8E-79 | 1.4794673 | 0.108 | 0.033 | 1.57E-74 | Regenerative Mesenchyme | exosc5 |
| mrps25 | 5.3E-79 | 1.227238 | 0.206 | 0.09 | 1.74E-74 | Regenerative Mesenchyme | mrps25 |
| psmd9 | 5.9E-79 | 1.2761432 | 0.179 | 0.073 | 1.91E-74 | Regenerative Mesenchyme | psmd9 |
| drg1 | 7.3E-79 | 1.4033787 | 0.163 | 0.064 | 2.37E-74 | Regenerative Mesenchyme | drg1 |
| nono | 8.6E-79 | 1.0824747 | 0.34 | 0.189 | 2.8E-74 | Regenerative Mesenchyme | nono |
| cdc42l | 1.1E-78 | 0.7448697 | 0.565 | 0.395 | 3.55E-74 | Regenerative Mesenchyme | cdc42l |
| aprt | 1.1E-78 | 1.7874457 | 0.113 | 0.036 | 3.7E-74 | Regenerative Mesenchyme | aprt |
| cltcb | 2E-78 | 1.1996074 | 0.223 | 0.102 | 6.62E-74 | Regenerative Mesenchyme | cltcb |
| rps62 | 3.1E-78 | 0.2965376 | 0.983 | 0.959 | 9.98E-74 | Regenerative Mesenchyme | rps6 |
| ndufa1 | 3.9E-78 | 0.953854 | 0.351 | 0.196 | 1.26E-73 | Regenerative Mesenchyme | ndufa1 |
| uap11 | 4.8E-78 | 0.9851169 | 0.258 | 0.125 | 1.56E-73 | Regenerative Mesenchyme | uap1 |
| pes | 5.8E-78 | 1.1594695 | 0.226 | 0.104 | 1.88E-73 | Regenerative Mesenchyme | pes |
| ctnnbip1 | 8E-78 | 1.6546809 | 0.117 | 0.039 | 2.61E-73 | Regenerative Mesenchyme | ctnnbip1 |
| ppp2cb | 1.3E-77 | 0.8396188 | 0.452 | 0.283 | 4.17E-73 | Regenerative Mesenchyme | ppp2cb |
| AK6 | 1.5E-77 | 1.0963006 | 0.229 | 0.106 | 4.87E-73 | Regenerative Mesenchyme | AK6 |
| tfg | 1.9E-77 | 1.0267756 | 0.33 | 0.182 | 6.05E-73 | Regenerative Mesenchyme | tfg |
| rpl36a2 | 2.4E-77 | 0.2266858 | 0.997 | 0.994 | 7.81E-73 | Regenerative Mesenchyme | rpl36a |
| dab2ipb1 | 2.5E-77 | 1.1530508 | 0.229 | 0.107 | 8.04E-73 | Regenerative Mesenchyme | dab2ipb |
| cpsf2 | 2.8E-77 | 1.4272131 | 0.133 | 0.047 | 9.11E-73 | Regenerative Mesenchyme | cpsf2 |
| sgta | 2.8E-77 | 1.0352676 | 0.264 | 0.13 | 9.13E-73 | Regenerative Mesenchyme | sgta |
| trappc2l | 2.9E-77 | 1.2225744 | 0.163 | 0.064 | 9.3E-73 | Regenerative Mesenchyme | trappc2l |
| imp4 | 3E-77 | 1.5821351 | 0.1 | 0.03 | 9.62E-73 | Regenerative Mesenchyme | imp4 |
| hk1 | 3.4E-77 | 1.1530808 | 0.236 | 0.112 | 1.1E-72 | Regenerative Mesenchyme | hk1 |
| btg31 | 3.5E-77 | 1.201771 | 0.246 | 0.119 | 1.15E-72 | Regenerative Mesenchyme | btg3 |
| ddx56 | 3.7E-77 | 1.534173 | 0.131 | 0.046 | 1.21E-72 | Regenerative Mesenchyme | ddx56 |
| mcmbp | 3.9E-77 | 1.6230953 | 0.101 | 0.031 | 1.28E-72 | Regenerative Mesenchyme | mcmbp |
| ddx52 | 5.4E-77 | 1.5452648 | 0.115 | 0.037 | 1.75E-72 | Regenerative Mesenchyme | ddx52 |
| pak1ip1 | 6.1E-77 | 1.3176923 | 0.144 | 0.053 | 1.97E-72 | Regenerative Mesenchyme | pak1ip1 |
| prpf19 | 1E-76 | 1.3250089 | 0.169 | 0.068 | 3.39E-72 | Regenerative Mesenchyme | prpf19 |
| rrp361 | 1.6E-76 | 1.1362142 | 0.19 | 0.081 | 5.06E-72 | Regenerative Mesenchyme | rrp36 |
| tomm40 | 1.7E-76 | 1.1937013 | 0.207 | 0.092 | 5.61E-72 | Regenerative Mesenchyme | tomm40 |
| tmem147 | 1.8E-76 | 1.0688229 | 0.282 | 0.145 | 5.78E-72 | Regenerative Mesenchyme | tmem147 |
| tmed2 | 1.8E-76 | 0.7794484 | 0.546 | 0.382 | 5.86E-72 | Regenerative Mesenchyme | tmed2 |
| rpl102 | 2E-76 | 0.3400844 | 0.986 | 0.967 | 6.45E-72 | Regenerative Mesenchyme | rpl10 |
| khsrp | 3E-76 | 1.0997844 | 0.246 | 0.118 | 9.87E-72 | Regenerative Mesenchyme | khsrp |
| uqcc2 | 3.3E-76 | 1.0299758 | 0.241 | 0.114 | 1.08E-71 | Regenerative Mesenchyme | uqcc2 |
| actr2a | 4.6E-76 | 0.9461874 | 0.35 | 0.197 | 1.49E-71 | Regenerative Mesenchyme | actr2a |
| smim151 | 6.1E-76 | 1.0157595 | 0.305 | 0.162 | 1.99E-71 | Regenerative Mesenchyme | smim15 |
| pdcd51 | 6.7E-76 | 1.0032704 | 0.292 | 0.152 | 2.19E-71 | Regenerative Mesenchyme | pdcd5 |
| tmem14ca | 7.9E-76 | 1.0361167 | 0.242 | 0.115 | 2.58E-71 | Regenerative Mesenchyme | tmem14ca |
| nnt | 1.3E-75 | 1.4116994 | 0.146 | 0.055 | 4.1E-71 | Regenerative Mesenchyme | nnt |

| | | | | | | | |
|---|---|---|---|---|---|---|---|
| clip2 | 1.4E-75 | 1.1612052 | 0.213 | 0.097 | 4.41E-71 | Regenerative Mesenchyme | clip2 |
| nmt1a | 1.8E-75 | 1.4008786 | 0.141 | 0.052 | 5.92E-71 | Regenerative Mesenchyme | nmt1a |
| atp6v0ca | 2.7E-75 | 0.7787894 | 0.501 | 0.33 | 8.71E-71 | Regenerative Mesenchyme | atp6v0ca |
| chmp5b | 4.3E-75 | 0.9134892 | 0.378 | 0.221 | 1.39E-70 | Regenerative Mesenchyme | chmp5b |
| ywhaz | 5.1E-75 | 0.6744975 | 0.674 | 0.522 | 1.64E-70 | Regenerative Mesenchyme | ywhaz |
| utp3 | 5.3E-75 | 0.9782849 | 0.26 | 0.128 | 1.72E-70 | Regenerative Mesenchyme | utp3 |
| lsm1 | 7.8E-75 | 1.4309357 | 0.117 | 0.039 | 2.55E-70 | Regenerative Mesenchyme | lsm1 |
| tmem214 | 9.9E-75 | 1.1258823 | 0.245 | 0.12 | 3.21E-70 | Regenerative Mesenchyme | tmem214 |
| timm44 | 1.9E-74 | 1.4642932 | 0.124 | 0.043 | 6.07E-70 | Regenerative Mesenchyme | timm44 |
| srsf2a | 3.2E-74 | 0.9111514 | 0.372 | 0.217 | 1.04E-69 | Regenerative Mesenchyme | srsf2a |
| cotl1 | 3.3E-74 | 1.3352665 | 0.209 | 0.096 | 1.07E-69 | Regenerative Mesenchyme | cotl1 |
| ddx39aa | 4.2E-74 | 1.0087679 | 0.318 | 0.174 | 1.37E-69 | Regenerative Mesenchyme | ddx39aa |
| slbp | 4.2E-74 | 1.8703896 | 0.106 | 0.034 | 1.37E-69 | Regenerative Mesenchyme | slbp |
| dcaf13 | 4.5E-74 | 1.3026464 | 0.141 | 0.053 | 1.47E-69 | Regenerative Mesenchyme | dcaf13 |
| naa101 | 4.8E-74 | 0.892124 | 0.427 | 0.269 | 1.55E-69 | Regenerative Mesenchyme | naa10 |
| srrt | 5E-74 | 0.9745516 | 0.249 | 0.12 | 1.62E-69 | Regenerative Mesenchyme | srrt |
| seta | 5.1E-74 | 0.8555905 | 0.471 | 0.309 | 1.65E-69 | Regenerative Mesenchyme | seta |
| eif3ja | 5.4E-74 | 1.0675332 | 0.227 | 0.106 | 1.77E-69 | Regenerative Mesenchyme | eif3ja |
| tmem242 | 6.6E-74 | 1.4858348 | 0.109 | 0.035 | 2.13E-69 | Regenerative Mesenchyme | tmem242 |
| micos13 | 7.3E-74 | 1.1689224 | 0.239 | 0.116 | 2.37E-69 | Regenerative Mesenchyme | micos13 |
| farsb | 8.1E-74 | 1.2860326 | 0.189 | 0.082 | 2.63E-69 | Regenerative Mesenchyme | farsb |
| gmppb | 1.4E-73 | 1.2136968 | 0.236 | 0.115 | 4.53E-69 | Regenerative Mesenchyme | gmppb |
| angptl41 | 2E-73 | 0.5307808 | 0.619 | 0.447 | 6.67E-69 | Regenerative Mesenchyme | angptl4 |
| ddx19 | 4.7E-73 | 1.3754973 | 0.141 | 0.053 | 1.51E-68 | Regenerative Mesenchyme | ddx19 |
| c7a2 | 5.5E-73 | 0.7312592 | 0.312 | 0.167 | 1.78E-68 | Regenerative Mesenchyme | c7a |
| nedd8l | 6E-73 | 1.2820852 | 0.159 | 0.063 | 1.95E-68 | Regenerative Mesenchyme | nedd8l |
| rpl18a2 | 7.6E-73 | 0.2691409 | 0.99 | 0.984 | 2.47E-68 | Regenerative Mesenchyme | rpl18a |
| oaz1a1 | 9.5E-73 | 0.6173931 | 0.607 | 0.443 | 3.09E-68 | Regenerative Mesenchyme | oaz1a |
| mpc11 | 1E-72 | 1.0704658 | 0.295 | 0.158 | 3.36E-68 | Regenerative Mesenchyme | mpc1 |
| rbmx2 | 1.1E-72 | 1.1174228 | 0.166 | 0.067 | 3.42E-68 | Regenerative Mesenchyme | rbmx2 |
| clns1a | 1.2E-72 | 1.3169816 | 0.172 | 0.072 | 3.92E-68 | Regenerative Mesenchyme | clns1a |
| arpc1a | 1.7E-72 | 0.831489 | 0.471 | 0.309 | 5.67E-68 | Regenerative Mesenchyme | arpc1a |
| si:dkey-230p4.1 | 1.8E-72 | 1.3840957 | 0.153 | 0.061 | 5.93E-68 | Regenerative Mesenchyme | si:dkey-230p4.1 |
| dpm31 | 1.8E-72 | 1.043513 | 0.199 | 0.088 | 6E-68 | Regenerative Mesenchyme | dpm3 |
| mrps23 | 2.4E-72 | 1.4777353 | 0.112 | 0.037 | 7.73E-68 | Regenerative Mesenchyme | mrps23 |
| top1l | 3.5E-72 | 0.7155105 | 0.553 | 0.384 | 1.15E-67 | Regenerative Mesenchyme | top1l |
| rpl23a2 | 4.1E-72 | 0.2614652 | 0.986 | 0.97 | 1.35E-67 | Regenerative Mesenchyme | rpl23a |
| tomm70a | 4.8E-72 | 0.9273139 | 0.309 | 0.167 | 1.56E-67 | Regenerative Mesenchyme | tomm70a |
| stt3b | 6.1E-72 | 1.2254698 | 0.217 | 0.102 | 1.99E-67 | Regenerative Mesenchyme | stt3b |
| metap2b | 4E-71 | 0.9657005 | 0.322 | 0.179 | 1.31E-66 | Regenerative Mesenchyme | metap2b |
| pycr1b1 | 4.1E-71 | 1.5450465 | 0.133 | 0.05 | 1.33E-66 | Regenerative Mesenchyme | pycr1b |
| rrbp1a2 | 6E-71 | 0.6712209 | 0.584 | 0.422 | 1.95E-66 | Regenerative Mesenchyme | rrbp1a |
| aldh18a11 | 8.9E-71 | 1.3852209 | 0.147 | 0.057 | 2.91E-66 | Regenerative Mesenchyme | aldh18a1 |
| tia1l | 1.4E-70 | 1.1265749 | 0.258 | 0.132 | 4.63E-66 | Regenerative Mesenchyme | tia1l |
| mad1l1 | 1.8E-70 | 1.5533049 | 0.11 | 0.037 | 5.88E-66 | Regenerative Mesenchyme | mad1l1 |
| arfgap2 | 1.8E-70 | 1.2931445 | 0.211 | 0.099 | 5.96E-66 | Regenerative Mesenchyme | arfgap2 |
| uqcrc2b1 | 1.9E-70 | 1.1186787 | 0.225 | 0.108 | 6.08E-66 | Regenerative Mesenchyme | uqcrc2b |
| cnp1 | 2E-70 | 0.7729174 | 0.447 | 0.283 | 6.37E-66 | Regenerative Mesenchyme | cnp |

| | | | | | | | |
|---|---|---|---|---|---|---|---|
| eef1db2 | 2.3E-70 | 0.5703345 | 0.695 | 0.574 | 7.34E-66 | Regenerative Mesenchyme | eef1db |
| lrpap1 | 3.6E-70 | 0.876094 | 0.316 | 0.173 | 1.17E-65 | Regenerative Mesenchyme | lrpap1 |
| arcn1a | 3.7E-70 | 1.2239186 | 0.202 | 0.092 | 1.21E-65 | Regenerative Mesenchyme | arcn1a |
| cox6b11 | 4E-70 | 1.7221591 | 0.157 | 0.065 | 1.32E-65 | Regenerative Mesenchyme | cox6b1 |
| ctnnbl1 | 5.7E-70 | 1.5114535 | 0.107 | 0.036 | 1.86E-65 | Regenerative Mesenchyme | ctnnbl1 |
| pfn1 | 6.1E-70 | 1.2627735 | 0.452 | 0.3 | 1.98E-65 | Regenerative Mesenchyme | pfn1 |
| hmgb1b | 6.3E-70 | 0.9014764 | 0.398 | 0.244 | 2.04E-65 | Regenerative Mesenchyme | hmgb1b |
| carm1 | 6.4E-70 | 1.1764659 | 0.179 | 0.077 | 2.1E-65 | Regenerative Mesenchyme | carm1 |
| lima1a1 | 8.8E-70 | 1.1664802 | 0.191 | 0.084 | 2.85E-65 | Regenerative Mesenchyme | lima1a |
| cyp20a1 | 1E-69 | 1.0983338 | 0.222 | 0.106 | 3.37E-65 | Regenerative Mesenchyme | cyp20a1 |
| loxl2a | 1.1E-69 | 0.6985288 | 0.369 | 0.216 | 3.47E-65 | Regenerative Mesenchyme | loxl2a |
| grwd11 | 1.2E-69 | 0.9507976 | 0.263 | 0.135 | 3.87E-65 | Regenerative Mesenchyme | grwd1 |
| mtap | 1.5E-69 | 0.905927 | 0.341 | 0.195 | 4.99E-65 | Regenerative Mesenchyme | mtap |
| fam114a11 | 1.7E-69 | 1.1015311 | 0.219 | 0.104 | 5.42E-65 | Regenerative Mesenchyme | fam114a1 |
| snrnp40 | 1.9E-69 | 1.6107568 | 0.1 | 0.032 | 6.26E-65 | Regenerative Mesenchyme | snrnp40 |
| eif3k | 2.4E-69 | 0.7282399 | 0.592 | 0.467 | 7.73E-65 | Regenerative Mesenchyme | eif3k |
| rbm34 | 3E-69 | 1.1538419 | 0.161 | 0.066 | 9.67E-65 | Regenerative Mesenchyme | rbm34 |
| atp5meb | 3E-69 | 0.6505564 | 0.65 | 0.531 | 9.77E-65 | Regenerative Mesenchyme | atp5meb |
| ywhabl | 5.6E-69 | 0.7484776 | 0.561 | 0.41 | 1.83E-64 | Regenerative Mesenchyme | ywhabl |
| ube2l3b | 8E-69 | 0.8783723 | 0.348 | 0.202 | 2.6E-64 | Regenerative Mesenchyme | ube2l3b |
| dlst | 8.4E-69 | 0.9576874 | 0.324 | 0.183 | 2.75E-64 | Regenerative Mesenchyme | dlst |
| igfbp6b | 1.3E-68 | 1.4452439 | 0.142 | 0.056 | 4.21E-64 | Regenerative Mesenchyme | igfbp6b |
| rps152 | 1.5E-68 | 0.2171008 | 0.992 | 0.989 | 4.73E-64 | Regenerative Mesenchyme | rps15 |
| rps52 | 1.8E-68 | 0.3207327 | 0.983 | 0.972 | 5.91E-64 | Regenerative Mesenchyme | rps5 |
| rwdd | 2.3E-68 | 1.3527499 | 0.135 | 0.051 | 7.35E-64 | Regenerative Mesenchyme | rwdd |
| hnrpkl | 3.1E-68 | 1.0545536 | 0.263 | 0.136 | 1.01E-63 | Regenerative Mesenchyme | hnrpkl |
| calm2a1 | 1.9E-67 | 0.7285633 | 0.473 | 0.313 | 6.2E-63 | Regenerative Mesenchyme | calm2a |
| dcun1d5 | 2.3E-67 | 1.2566056 | 0.164 | 0.069 | 7.5E-63 | Regenerative Mesenchyme | dcun1d5 |
| uck2a | 2.6E-67 | 1.5709873 | 0.117 | 0.042 | 8.61E-63 | Regenerative Mesenchyme | uck2a |
| epcam1 | 2.6E-67 | 1.5244054 | 0.227 | 0.114 | 8.61E-63 | Regenerative Mesenchyme | epcam |
| smim4 | 3E-67 | 1.2657761 | 0.139 | 0.054 | 9.6E-63 | Regenerative Mesenchyme | smim4 |
| ccdc137 | 3E-67 | 1.3934691 | 0.118 | 0.042 | 9.64E-63 | Regenerative Mesenchyme | ccdc137 |
| mrps35 | 3.7E-67 | 1.3102117 | 0.133 | 0.05 | 1.21E-62 | Regenerative Mesenchyme | mrps35 |
| ddx54 | 5E-67 | 1.2078897 | 0.14 | 0.054 | 1.64E-62 | Regenerative Mesenchyme | ddx54 |
| rfc5 | 9.4E-67 | 1.4699846 | 0.109 | 0.037 | 3.07E-62 | Regenerative Mesenchyme | rfc5 |
| sat1a.21 | 1E-66 | 0.4474987 | 0.611 | 0.448 | 3.28E-62 | Regenerative Mesenchyme | sat1a.2 |
| drg2 | 1.2E-66 | 1.1244831 | 0.228 | 0.112 | 3.74E-62 | Regenerative Mesenchyme | drg2 |
| dap3 | 2.1E-66 | 1.4695516 | 0.108 | 0.037 | 6.86E-62 | Regenerative Mesenchyme | dap3 |
| mcfd2 | 3.6E-66 | 1.047246 | 0.24 | 0.121 | 1.18E-61 | Regenerative Mesenchyme | mcfd2 |
| ndufa9a | 3.8E-66 | 1.1763081 | 0.207 | 0.099 | 1.24E-61 | Regenerative Mesenchyme | ndufa9a |
| rabif | 4.2E-66 | 1.3761443 | 0.147 | 0.059 | 1.36E-61 | Regenerative Mesenchyme | rabif |
| arf1 | 4.2E-66 | 0.9123789 | 0.272 | 0.143 | 1.38E-61 | Regenerative Mesenchyme | arf1 |
| rps292 | 5E-66 | 0.2847519 | 0.982 | 0.969 | 1.63E-61 | Regenerative Mesenchyme | rps29 |
| eif4a3 | 5.4E-66 | 0.9293758 | 0.314 | 0.177 | 1.74E-61 | Regenerative Mesenchyme | eif4a3 |
| mif | 6.6E-66 | 0.7407569 | 0.542 | 0.392 | 2.14E-61 | Regenerative Mesenchyme | mif |
| CU694197.1 | 1.4E-65 | 0.8999928 | 0.339 | 0.199 | 4.51E-61 | Regenerative Mesenchyme | CU694197.1 |
| ola12 | 1.4E-65 | 0.6283758 | 0.628 | 0.5 | 4.57E-61 | Regenerative Mesenchyme | ola1 |
| twistnb | 1.4E-65 | 1.0218824 | 0.211 | 0.101 | 4.64E-61 | Regenerative Mesenchyme | twistnb |

| | | | | | | | |
|---|---|---|---|---|---|---|---|
| emc9 | 3E-65 | 1.3358234 | 0.126 | 0.047 | 9.83E-61 | Regenerative Mesenchyme | emc9 |
| phpt1 | 3.6E-65 | 1.2441529 | 0.143 | 0.057 | 1.18E-60 | Regenerative Mesenchyme | phpt1 |
| prpf8 | 5.5E-65 | 1.0581657 | 0.215 | 0.103 | 1.8E-60 | Regenerative Mesenchyme | prpf8 |
| pxna | 6.3E-65 | 1.1508862 | 0.213 | 0.103 | 2.06E-60 | Regenerative Mesenchyme | pxna |
| plgrkt1 | 7.6E-65 | 1.3716066 | 0.132 | 0.051 | 2.47E-60 | Regenerative Mesenchyme | plgrkt |
| tpra | 1.1E-64 | 1.2211889 | 0.159 | 0.067 | 3.56E-60 | Regenerative Mesenchyme | tpra |
| si:ch211-282b22.1 | 1.4E-64 | 1.0670036 | 0.187 | 0.085 | 4.64E-60 | Regenerative Mesenchyme | si:ch211-282b22.1 |
| zgc:91910 | 1.4E-64 | 0.8001512 | 0.395 | 0.244 | 4.69E-60 | Regenerative Mesenchyme | zgc:91910 |
| mrps9 | 1.6E-64 | 1.2744129 | 0.121 | 0.045 | 5.34E-60 | Regenerative Mesenchyme | mrps9 |
| ccdc25 | 1.9E-64 | 0.9155028 | 0.305 | 0.171 | 6.31E-60 | Regenerative Mesenchyme | ccdc25 |
| atp13a1 | 2.7E-64 | 1.2484522 | 0.129 | 0.049 | 8.83E-60 | Regenerative Mesenchyme | atp13a1 |
| cep170aa | 2.8E-64 | 1.3602603 | 0.14 | 0.056 | 9.21E-60 | Regenerative Mesenchyme | cep170aa |
| ncln1 | 3.2E-64 | 1.1871045 | 0.16 | 0.068 | 1.03E-59 | Regenerative Mesenchyme | ncln |
| sec22ba1 | 4.5E-64 | 1.0654144 | 0.225 | 0.112 | 1.45E-59 | Regenerative Mesenchyme | sec22ba |
| mrps61 | 8.2E-64 | 1.1775336 | 0.173 | 0.077 | 2.66E-59 | Regenerative Mesenchyme | mrps6 |
| rai142 | 9.1E-64 | 1.1607788 | 0.18 | 0.081 | 2.97E-59 | Regenerative Mesenchyme | rai14 |
| si:dkey-286j15.1 | 1.3E-63 | 0.9587904 | 0.277 | 0.15 | 4.18E-59 | Regenerative Mesenchyme | si:dkey-286j15.1 |
| ube2v1 | 3.2E-63 | 0.8168108 | 0.408 | 0.258 | 1.05E-58 | Regenerative Mesenchyme | ube2v1 |
| rrbp1b1 | 3.4E-63 | 0.8842018 | 0.374 | 0.231 | 1.12E-58 | Regenerative Mesenchyme | rrbp1b |
| dusp51 | 3.6E-63 | 0.7872513 | 0.327 | 0.191 | 1.18E-58 | Regenerative Mesenchyme | dusp5 |
| ndc1 | 5.9E-63 | 1.25259 | 0.137 | 0.055 | 1.93E-58 | Regenerative Mesenchyme | ndc1 |
| thoc7 | 7.6E-63 | 1.1762876 | 0.146 | 0.06 | 2.48E-58 | Regenerative Mesenchyme | thoc7 |
| RPS172 | 9.8E-63 | 0.2207202 | 0.997 | 0.987 | 3.18E-58 | Regenerative Mesenchyme | RPS17 |
| hbegfa2 | 1.3E-62 | 0.5731922 | 0.559 | 0.393 | 4.28E-58 | Regenerative Mesenchyme | hbegfa |
| mrps36 | 1.4E-62 | 0.9754775 | 0.274 | 0.15 | 4.57E-58 | Regenerative Mesenchyme | mrps36 |
| cbx1a | 2.2E-62 | 0.8862928 | 0.333 | 0.197 | 7.04E-58 | Regenerative Mesenchyme | cbx1a |
| igf2b | 2.7E-62 | 0.6310974 | 0.577 | 0.428 | 8.65E-58 | Regenerative Mesenchyme | igf2b |
| srsf6a | 2.8E-62 | 0.9686703 | 0.236 | 0.119 | 9.11E-58 | Regenerative Mesenchyme | srsf6a |
| mdn1 | 5.6E-62 | 1.0807659 | 0.214 | 0.105 | 1.82E-57 | Regenerative Mesenchyme | mdn1 |
| rad50 | 6E-62 | 1.2165127 | 0.144 | 0.059 | 1.94E-57 | Regenerative Mesenchyme | rad50 |
| naa50 | 7E-62 | 0.9649718 | 0.25 | 0.13 | 2.27E-57 | Regenerative Mesenchyme | naa50 |
| mrpl321 | 7.4E-62 | 0.8775369 | 0.291 | 0.162 | 2.41E-57 | Regenerative Mesenchyme | mrpl32 |
| srp91 | 7.9E-62 | 1.0841786 | 0.209 | 0.102 | 2.58E-57 | Regenerative Mesenchyme | srp9 |
| mrpl37 | 1E-61 | 1.3498285 | 0.112 | 0.04 | 3.31E-57 | Regenerative Mesenchyme | mrpl37 |
| rpl172 | 1.3E-61 | 0.2390735 | 0.995 | 0.989 | 4.34E-57 | Regenerative Mesenchyme | rpl17 |
| naa20 | 1.4E-61 | 0.906793 | 0.267 | 0.143 | 4.71E-57 | Regenerative Mesenchyme | naa20 |
| hnrnpa0a | 1.6E-61 | 0.8073295 | 0.369 | 0.224 | 5.16E-57 | Regenerative Mesenchyme | hnrnpa0a |
| caprin1a | 1.7E-61 | 1.0366843 | 0.202 | 0.097 | 5.52E-57 | Regenerative Mesenchyme | caprin1a |
| rpl5a2 | 3.7E-61 | 0.2981934 | 0.957 | 0.917 | 1.21E-56 | Regenerative Mesenchyme | rpl5a |
| ufsp2 | 3.8E-61 | 1.1693474 | 0.155 | 0.066 | 1.22E-56 | Regenerative Mesenchyme | ufsp2 |
| rps4x2 | 4.6E-61 | 0.2681659 | 0.986 | 0.966 | 1.51E-56 | Regenerative Mesenchyme | rps4x |
| pinx11 | 6.1E-61 | 0.791891 | 0.367 | 0.226 | 1.99E-56 | Regenerative Mesenchyme | pinx1 |
| rpl35a2 | 6.4E-61 | 0.2037349 | 0.996 | 0.989 | 2.08E-56 | Regenerative Mesenchyme | rpl35a |
| eif4eb | 8E-61 | 1.0170641 | 0.205 | 0.099 | 2.62E-56 | Regenerative Mesenchyme | eif4eb |
| lmf2b | 8.4E-61 | 0.9019528 | 0.25 | 0.131 | 2.75E-56 | Regenerative Mesenchyme | lmf2b |
| ptcd3 | 9.1E-61 | 1.4651334 | 0.1 | 0.035 | 2.98E-56 | Regenerative Mesenchyme | ptcd3 |
| gspt1l | 1E-60 | 0.8598754 | 0.356 | 0.218 | 3.33E-56 | Regenerative Mesenchyme | gspt1l |
| mrps7 | 1E-60 | 1.101513 | 0.166 | 0.073 | 3.35E-56 | Regenerative Mesenchyme | mrps7 |

| | | | | | | | |
|---|---|---|---|---|---|---|---|
| rrad | 1.2E-60 | 1.0381547 | 0.153 | 0.066 | 4.03E-56 | Regenerative Mesenchyme | rrad |
| plekha1a | 1.6E-60 | 1.3527835 | 0.134 | 0.054 | 5.19E-56 | Regenerative Mesenchyme | plekha1a |
| pop71 | 1.7E-60 | 1.09043 | 0.153 | 0.065 | 5.4E-56 | Regenerative Mesenchyme | pop7 |
| ywhabb | 1.9E-60 | 0.6589191 | 0.539 | 0.389 | 6.3E-56 | Regenerative Mesenchyme | ywhabb |
| krr1 | 2E-60 | 1.2132925 | 0.129 | 0.051 | 6.59E-56 | Regenerative Mesenchyme | krr1 |
| eed | 2.2E-60 | 1.3079911 | 0.118 | 0.045 | 7.11E-56 | Regenerative Mesenchyme | eed |
| fkbp1ab | 4.1E-60 | 1.3798623 | 0.101 | 0.035 | 1.32E-55 | Regenerative Mesenchyme | fkbp1ab |
| pole3 | 6.7E-60 | 1.0436461 | 0.215 | 0.107 | 2.18E-55 | Regenerative Mesenchyme | pole3 |
| tmem39a1 | 1.2E-59 | 1.0009364 | 0.263 | 0.144 | 4.01E-55 | Regenerative Mesenchyme | tmem39a |
| mrpl40 | 1.8E-59 | 1.2911741 | 0.111 | 0.041 | 5.72E-55 | Regenerative Mesenchyme | mrpl40 |
| nucb2b1 | 2.1E-59 | 1.0890086 | 0.221 | 0.112 | 6.9E-55 | Regenerative Mesenchyme | nucb2b |
| wdr12 | 2.2E-59 | 1.2351946 | 0.129 | 0.051 | 7.2E-55 | Regenerative Mesenchyme | wdr12 |
| sap30l | 2.5E-59 | 0.8195178 | 0.348 | 0.209 | 8.18E-55 | Regenerative Mesenchyme | sap30l |
| psme2 | 3.4E-59 | 1.1168178 | 0.162 | 0.072 | 1.1E-54 | Regenerative Mesenchyme | psme2 |
| ndufv1 | 3.8E-59 | 1.0096175 | 0.239 | 0.126 | 1.24E-54 | Regenerative Mesenchyme | ndufv1 |
| trmt6 | 4.1E-59 | 1.1583741 | 0.138 | 0.057 | 1.34E-54 | Regenerative Mesenchyme | trmt6 |
| cacybp | 5.1E-59 | 0.8317994 | 0.27 | 0.146 | 1.67E-54 | Regenerative Mesenchyme | cacybp |
| rplp02 | 5.3E-59 | 0.2769843 | 0.994 | 0.982 | 1.71E-54 | Regenerative Mesenchyme | rplp0 |
| tcnbb | 9.9E-59 | 0.9790259 | 0.115 | 0.044 | 3.23E-54 | Regenerative Mesenchyme | tcnbb |
| farsa1 | 1.2E-58 | 1.2137096 | 0.168 | 0.077 | 3.98E-54 | Regenerative Mesenchyme | farsa |
| dph5 | 1.2E-58 | 1.4031637 | 0.11 | 0.041 | 4.03E-54 | Regenerative Mesenchyme | dph5 |
| arpc4 | 1.3E-58 | 0.9038419 | 0.3 | 0.171 | 4.21E-54 | Regenerative Mesenchyme | arpc4 |
| atf4a2 | 1.4E-58 | 0.7023495 | 0.441 | 0.29 | 4.7E-54 | Regenerative Mesenchyme | atf4a |
| slc30a71 | 1.7E-58 | 1.0692811 | 0.212 | 0.107 | 5.58E-54 | Regenerative Mesenchyme | slc30a7 |
| tpm4a1 | 1.8E-58 | 0.4617933 | 0.665 | 0.547 | 5.7E-54 | Regenerative Mesenchyme | tpm4a |
| supt4h1 | 1.8E-58 | 0.9494834 | 0.245 | 0.129 | 5.99E-54 | Regenerative Mesenchyme | supt4h1 |
| hsp90ab1 | 2.8E-58 | 0.2120856 | 0.996 | 0.992 | 8.98E-54 | Regenerative Mesenchyme | hsp90ab1 |
| CABZ01102240.1 | 3.9E-58 | 1.4519314 | 0.105 | 0.038 | 1.28E-53 | Regenerative Mesenchyme | CABZ01102240.1 |
| syne1a | 4.3E-58 | 1.2445782 | 0.136 | 0.056 | 1.39E-53 | Regenerative Mesenchyme | syne1a |
| nudt5 | 4.7E-58 | 1.3849622 | 0.116 | 0.045 | 1.53E-53 | Regenerative Mesenchyme | nudt5 |
| ewsr1b | 5.4E-58 | 0.8037172 | 0.364 | 0.224 | 1.77E-53 | Regenerative Mesenchyme | ewsr1b |
| gorasp21 | 8.8E-58 | 0.9775668 | 0.211 | 0.105 | 2.87E-53 | Regenerative Mesenchyme | gorasp2 |
| eif3s10 | 9.2E-58 | 0.6252814 | 0.6 | 0.466 | 3.01E-53 | Regenerative Mesenchyme | eif3s10 |
| eif3jb | 1.1E-57 | 0.7693986 | 0.356 | 0.218 | 3.57E-53 | Regenerative Mesenchyme | eif3jb |
| prpf40a | 1.1E-57 | 0.8879171 | 0.229 | 0.118 | 3.61E-53 | Regenerative Mesenchyme | prpf40a |
| vaspb2 | 1.2E-57 | 1.2357926 | 0.193 | 0.095 | 3.82E-53 | Regenerative Mesenchyme | vaspb |
| ilf3a | 1.3E-57 | 1.003528 | 0.148 | 0.063 | 4.25E-53 | Regenerative Mesenchyme | ilf3a |
| cnpy1 | 1.7E-57 | 1.1627105 | 0.166 | 0.076 | 5.53E-53 | Regenerative Mesenchyme | cnpy1 |
| pdia51 | 2E-57 | 0.825206 | 0.284 | 0.159 | 6.5E-53 | Regenerative Mesenchyme | pdia5 |
| si:dkey-250d21.11 | 2.1E-57 | 0.8562457 | 0.258 | 0.138 | 6.84E-53 | Regenerative Mesenchyme | si:dkey-250d21.1 |
| sumo3a | 2.4E-57 | 0.7785229 | 0.42 | 0.279 | 7.95E-53 | Regenerative Mesenchyme | sumo3a |
| wdr46 | 2.9E-57 | 1.2180425 | 0.124 | 0.049 | 9.41E-53 | Regenerative Mesenchyme | wdr46 |
| ssr1 | 5E-57 | 1.3540776 | 0.128 | 0.052 | 1.61E-52 | Regenerative Mesenchyme | ssr1 |
| syncripl | 5.3E-57 | 0.9155267 | 0.224 | 0.115 | 1.73E-52 | Regenerative Mesenchyme | syncripl |
| lrrc47 | 5.9E-57 | 1.0246394 | 0.154 | 0.068 | 1.92E-52 | Regenerative Mesenchyme | lrrc47 |
| riok3 | 6E-57 | 1.2925274 | 0.118 | 0.046 | 1.94E-52 | Regenerative Mesenchyme | riok3 |
| lyrm4 | 6.4E-57 | 1.0969538 | 0.125 | 0.05 | 2.09E-52 | Regenerative Mesenchyme | lyrm4 |
| rbm28 | 6.5E-57 | 0.711085 | 0.379 | 0.233 | 2.11E-52 | Regenerative Mesenchyme | rbm28 |

| | | | | | | | |
|---|---|---|---|---|---|---|---|
| rangap1a | 7.2E-57 | 1.156035 | 0.137 | 0.057 | 2.35E-52 | Regenerative Mesenchyme | rangap1a |
| ap1m1 | 8.3E-57 | 0.9781073 | 0.228 | 0.119 | 2.69E-52 | Regenerative Mesenchyme | ap1m1 |
| eif5 | 1E-56 | 0.5913698 | 0.602 | 0.473 | 3.29E-52 | Regenerative Mesenchyme | eif5 |
| u2af2b | 1.5E-56 | 0.7003304 | 0.444 | 0.298 | 4.91E-52 | Regenerative Mesenchyme | u2af2b |
| ntmt1 | 1.7E-56 | 1.5527397 | 0.108 | 0.041 | 5.59E-52 | Regenerative Mesenchyme | ntmt1 |
| atp2a2b | 2.2E-56 | 1.1232656 | 0.196 | 0.097 | 7.07E-52 | Regenerative Mesenchyme | atp2a2b |
| fibina2 | 2.3E-56 | 0.6756753 | 0.34 | 0.206 | 7.52E-52 | Regenerative Mesenchyme | fibina |
| rps19bp1 | 2.4E-56 | 1.1203747 | 0.131 | 0.054 | 7.65E-52 | Regenerative Mesenchyme | rps19bp1 |
| nucb1 | 3.6E-56 | 0.7974534 | 0.334 | 0.2 | 1.17E-51 | Regenerative Mesenchyme | nucb1 |
| gmds | 3.8E-56 | 1.2797162 | 0.121 | 0.048 | 1.23E-51 | Regenerative Mesenchyme | gmds |
| dnaja2a | 4.7E-56 | 0.6644575 | 0.494 | 0.346 | 1.53E-51 | Regenerative Mesenchyme | dnaja2a |
| ezrb | 8.7E-56 | 0.6085893 | 0.316 | 0.184 | 2.84E-51 | Regenerative Mesenchyme | ezrb |
| ap2s1 | 9.4E-56 | 0.8620643 | 0.29 | 0.166 | 3.07E-51 | Regenerative Mesenchyme | ap2s1 |
| mrpl44 | 1.3E-55 | 1.3289813 | 0.121 | 0.049 | 4.19E-51 | Regenerative Mesenchyme | mrpl44 |
| atp5po2 | 1.3E-55 | 0.6745116 | 0.551 | 0.418 | 4.29E-51 | Regenerative Mesenchyme | atp5po |
| erh | 3.3E-55 | 0.6722022 | 0.537 | 0.402 | 1.07E-50 | Regenerative Mesenchyme | erh |
| ubfd1 | 3.8E-55 | 0.9633193 | 0.203 | 0.102 | 1.23E-50 | Regenerative Mesenchyme | ubfd1 |
| ndnl2 | 4.2E-55 | 1.1952446 | 0.129 | 0.053 | 1.37E-50 | Regenerative Mesenchyme | ndnl2 |
| vrk1 | 4.4E-55 | 1.3037854 | 0.117 | 0.046 | 1.41E-50 | Regenerative Mesenchyme | vrk1 |
| ralaa | 6.6E-55 | 1.2522697 | 0.107 | 0.04 | 2.14E-50 | Regenerative Mesenchyme | ralaa |
| arg21 | 6.7E-55 | 0.7426189 | 0.278 | 0.157 | 2.17E-50 | Regenerative Mesenchyme | arg2 |
| rpl182 | 7.6E-55 | 0.2180809 | 0.99 | 0.981 | 2.47E-50 | Regenerative Mesenchyme | rpl18 |
| mrps31 | 1E-54 | 0.9876485 | 0.155 | 0.069 | 3.24E-50 | Regenerative Mesenchyme | mrps31 |
| sf3b2 | 1.1E-54 | 0.7445998 | 0.33 | 0.196 | 3.44E-50 | Regenerative Mesenchyme | sf3b2 |
| rpl82 | 1.4E-54 | 0.2598899 | 0.99 | 0.977 | 4.69E-50 | Regenerative Mesenchyme | rpl8 |
| polr2b | 1.5E-54 | 1.2737584 | 0.125 | 0.051 | 4.74E-50 | Regenerative Mesenchyme | polr2b |
| ube2ib | 1.5E-54 | 0.8279045 | 0.291 | 0.168 | 5E-50 | Regenerative Mesenchyme | ube2ib |
| mid1ip1b1 | 2E-54 | 0.7872761 | 0.222 | 0.114 | 6.45E-50 | Regenerative Mesenchyme | mid1ip1b |
| cpsf1 | 3.1E-54 | 1.3052423 | 0.1 | 0.037 | 1.02E-49 | Regenerative Mesenchyme | cpsf1 |
| faub | 3.7E-54 | 1.1777131 | 0.171 | 0.081 | 1.21E-49 | Regenerative Mesenchyme | faub |
| creb3l12 | 4.3E-54 | 1.0287878 | 0.209 | 0.108 | 1.4E-49 | Regenerative Mesenchyme | creb3l1 |
| baz2ba | 6.3E-54 | 0.7676796 | 0.287 | 0.163 | 2.04E-49 | Regenerative Mesenchyme | baz2ba |
| si:dkey-67c22.2 | 7E-54 | 0.7016028 | 0.359 | 0.221 | 2.27E-49 | Regenerative Mesenchyme | si:dkey-67c22.2 |
| urm11 | 7.8E-54 | 1.1983827 | 0.147 | 0.065 | 2.53E-49 | Regenerative Mesenchyme | urm1 |
| nrd1a | 8.8E-54 | 0.9727174 | 0.177 | 0.085 | 2.88E-49 | Regenerative Mesenchyme | nrd1a |
| emc101 | 1.2E-53 | 0.9073216 | 0.252 | 0.139 | 4.01E-49 | Regenerative Mesenchyme | emc10 |
| mrps30 | 1.3E-53 | 1.1996097 | 0.113 | 0.044 | 4.21E-49 | Regenerative Mesenchyme | mrps30 |
| ccnd12 | 1.5E-53 | 1.360075 | 0.157 | 0.073 | 4.77E-49 | Regenerative Mesenchyme | ccnd1 |
| hdgfl2 | 1.6E-53 | 0.6813528 | 0.299 | 0.171 | 5.22E-49 | Regenerative Mesenchyme | hdgfl2 |
| dnttip2 | 2.4E-53 | 1.1437871 | 0.154 | 0.07 | 7.88E-49 | Regenerative Mesenchyme | dnttip2 |
| pgk1 | 3E-53 | 0.8359077 | 0.328 | 0.201 | 9.6E-49 | Regenerative Mesenchyme | pgk1 |
| prdm1a1 | 3E-53 | 0.8483715 | 0.16 | 0.073 | 9.63E-49 | Regenerative Mesenchyme | prdm1a |
| ganab | 3.7E-53 | 1.2605703 | 0.133 | 0.056 | 1.21E-48 | Regenerative Mesenchyme | ganab |
| parp2 | 3.9E-53 | 1.0656135 | 0.139 | 0.06 | 1.27E-48 | Regenerative Mesenchyme | parp2 |
| csnk2b | 4.9E-53 | 0.761861 | 0.382 | 0.246 | 1.6E-48 | Regenerative Mesenchyme | csnk2b |
| arhgdia1 | 5E-53 | 0.5688082 | 0.547 | 0.404 | 1.62E-48 | Regenerative Mesenchyme | arhgdia |
| tprb | 5.2E-53 | 0.8322587 | 0.197 | 0.098 | 1.68E-48 | Regenerative Mesenchyme | tprb |
| mrpl51 | 5.6E-53 | 1.2392184 | 0.101 | 0.037 | 1.82E-48 | Regenerative Mesenchyme | mrpl51 |

| | | | | | | | |
|---|---|---|---|---|---|---|---|
| faf2 | 6.1E-53 | 0.8594118 | 0.247 | 0.134 | 1.97E-48 | Regenerative Mesenchyme | faf2 |
| mrpl42 | 6.1E-53 | 1.263765 | 0.113 | 0.044 | 1.98E-48 | Regenerative Mesenchyme | mrpl42 |
| pdha1a | 7.4E-53 | 0.9228301 | 0.241 | 0.13 | 2.4E-48 | Regenerative Mesenchyme | pdha1a |
| aldh1a2 | 1E-52 | 0.8206654 | 0.124 | 0.051 | 3.32E-48 | Regenerative Mesenchyme | aldh1a2 |
| pfdn5 | 1.2E-52 | 0.8861848 | 0.29 | 0.17 | 3.74E-48 | Regenerative Mesenchyme | pfdn5 |
| utp25 | 1.3E-52 | 1.0883147 | 0.126 | 0.052 | 4.11E-48 | Regenerative Mesenchyme | utp25 |
| txndc5 | 1.3E-52 | 0.8480722 | 0.326 | 0.2 | 4.34E-48 | Regenerative Mesenchyme | txndc5 |
| dlat | 1.5E-52 | 1.1587056 | 0.157 | 0.073 | 4.74E-48 | Regenerative Mesenchyme | dlat |
| sigmar1 | 1.9E-52 | 1.0440233 | 0.185 | 0.091 | 6.21E-48 | Regenerative Mesenchyme | sigmar1 |
| mtch2 | 2.4E-52 | 0.9806275 | 0.193 | 0.096 | 7.66E-48 | Regenerative Mesenchyme | mtch2 |
| stip1 | 3.3E-52 | 0.7975632 | 0.272 | 0.154 | 1.06E-47 | Regenerative Mesenchyme | stip1 |
| nudt21 | 4.9E-52 | 0.9935078 | 0.195 | 0.098 | 1.6E-47 | Regenerative Mesenchyme | nudt21 |
| tnpo2 | 5.9E-52 | 1.0434567 | 0.171 | 0.082 | 1.9E-47 | Regenerative Mesenchyme | tnpo2 |
| cox14 | 2.8E-51 | 1.0586053 | 0.167 | 0.08 | 9.04E-47 | Regenerative Mesenchyme | cox14 |
| atxn2l | 2.9E-51 | 0.8937001 | 0.18 | 0.088 | 9.27E-47 | Regenerative Mesenchyme | atxn2l |
| tcima1 | 3.3E-51 | 0.4701701 | 0.353 | 0.219 | 1.07E-46 | Regenerative Mesenchyme | tcima |
| ndufv3 | 4E-51 | 0.866717 | 0.23 | 0.124 | 1.3E-46 | Regenerative Mesenchyme | ndufv3 |
| bola3 | 6.2E-51 | 0.9757674 | 0.134 | 0.058 | 2.02E-46 | Regenerative Mesenchyme | bola3 |
| txnrd3 | 9.1E-51 | 1.0214749 | 0.185 | 0.093 | 2.95E-46 | Regenerative Mesenchyme | txnrd3 |
| ikbip1 | 1.2E-50 | 1.0385895 | 0.214 | 0.114 | 3.78E-46 | Regenerative Mesenchyme | ikbip |
| fip1l1b | 1.2E-50 | 1.0434496 | 0.162 | 0.077 | 3.98E-46 | Regenerative Mesenchyme | fip1l1b |
| klhdc4 | 1.3E-50 | 1.1335142 | 0.124 | 0.052 | 4.12E-46 | Regenerative Mesenchyme | klhdc4 |
| si:dkeyp-55f12.3 | 1.3E-50 | 1.0815804 | 0.142 | 0.064 | 4.27E-46 | Regenerative Mesenchyme | si:dkeyp-55f12.3 |
| sdhb | 1.4E-50 | 0.8943001 | 0.286 | 0.17 | 4.52E-46 | Regenerative Mesenchyme | sdhb |
| hspa4a1 | 1.6E-50 | 0.7284977 | 0.257 | 0.143 | 5.19E-46 | Regenerative Mesenchyme | hspa4a |
| ndufv2 | 1.6E-50 | 0.9958003 | 0.196 | 0.101 | 5.34E-46 | Regenerative Mesenchyme | ndufv2 |
| znrd2 | 3.1E-50 | 1.1453824 | 0.149 | 0.069 | 1.02E-45 | Regenerative Mesenchyme | znrd2 |
| rbmx | 3.3E-50 | 0.8270117 | 0.228 | 0.122 | 1.08E-45 | Regenerative Mesenchyme | rbmx |
| fibinb1 | 3.6E-50 | 0.4244773 | 0.412 | 0.267 | 1.17E-45 | Regenerative Mesenchyme | fibinb |
| hadhaa | 3.7E-50 | 1.2783158 | 0.107 | 0.042 | 1.19E-45 | Regenerative Mesenchyme | hadhaa |
| sarnp | 4E-50 | 0.7290563 | 0.401 | 0.266 | 1.31E-45 | Regenerative Mesenchyme | sarnp |
| polr2c | 5.3E-50 | 1.0967816 | 0.116 | 0.047 | 1.72E-45 | Regenerative Mesenchyme | polr2c |
| dnajc11a | 5.3E-50 | 1.3412987 | 0.102 | 0.039 | 1.73E-45 | Regenerative Mesenchyme | dnajc11a |
| suclg2 | 7.7E-50 | 1.3082342 | 0.106 | 0.042 | 2.52E-45 | Regenerative Mesenchyme | suclg2 |
| C20H12orf57 | 7.9E-50 | 0.9938063 | 0.177 | 0.087 | 2.56E-45 | Regenerative Mesenchyme | C20H12orf57 |
| gnl2 | 1E-49 | 1.030229 | 0.147 | 0.067 | 3.25E-45 | Regenerative Mesenchyme | gnl2 |
| rnf121 | 1.4E-49 | 1.2722785 | 0.107 | 0.043 | 4.53E-45 | Regenerative Mesenchyme | rnf121 |
| timm23a1 | 1.5E-49 | 0.9497107 | 0.188 | 0.095 | 4.81E-45 | Regenerative Mesenchyme | timm23a |
| commd4 | 1.5E-49 | 1.0001026 | 0.146 | 0.066 | 4.82E-45 | Regenerative Mesenchyme | commd4 |
| phlda21 | 1.6E-49 | 0.3220209 | 0.346 | 0.212 | 5.24E-45 | Regenerative Mesenchyme | phlda2 |
| pim11 | 2.6E-49 | 0.2802688 | 0.693 | 0.537 | 8.57E-45 | Regenerative Mesenchyme | pim1 |
| cdh21 | 2.6E-49 | 0.6548894 | 0.426 | 0.285 | 8.6E-45 | Regenerative Mesenchyme | cdh2 |
| ddx47 | 3E-49 | 1.1797776 | 0.129 | 0.056 | 9.71E-45 | Regenerative Mesenchyme | ddx47 |
| smdt1b | 3.4E-49 | 0.7270468 | 0.375 | 0.245 | 1.1E-44 | Regenerative Mesenchyme | smdt1b |
| uck2b1 | 4.5E-49 | 1.3987984 | 0.119 | 0.05 | 1.48E-44 | Regenerative Mesenchyme | uck2b |
| psmb10 | 5.2E-49 | 1.1852339 | 0.137 | 0.061 | 1.68E-44 | Regenerative Mesenchyme | psmb10 |
| sdhc | 5.7E-49 | 0.7347436 | 0.312 | 0.189 | 1.87E-44 | Regenerative Mesenchyme | sdhc |
| angptl2b1 | 6E-49 | 0.5499024 | 0.244 | 0.135 | 1.96E-44 | Regenerative Mesenchyme | angptl2b |

| | | | | | | | |
|---|---|---|---|---|---|---|---|
| acad9 | 8.7E-49 | 1.2926042 | 0.12 | 0.051 | 2.81E-44 | Regenerative Mesenchyme | acad9 |
| loxa3 | 9.1E-49 | 0.6028136 | 0.286 | 0.168 | 2.97E-44 | Regenerative Mesenchyme | loxa |
| rrp8 | 1.3E-48 | 0.8740603 | 0.164 | 0.078 | 4.38E-44 | Regenerative Mesenchyme | rrp8 |
| nup62l | 1.6E-48 | 1.0757556 | 0.132 | 0.058 | 5.04E-44 | Regenerative Mesenchyme | nup62l |
| apmap1 | 1.8E-48 | 0.9539095 | 0.159 | 0.076 | 5.94E-44 | Regenerative Mesenchyme | apmap |
| cap1 | 3.4E-48 | 1.3774169 | 0.112 | 0.046 | 1.1E-43 | Regenerative Mesenchyme | cap1 |
| dtd1 | 3.4E-48 | 1.1701565 | 0.111 | 0.045 | 1.11E-43 | Regenerative Mesenchyme | dtd1 |
| serpinh1a2 | 3.5E-48 | 0.8109486 | 0.357 | 0.232 | 1.13E-43 | Regenerative Mesenchyme | serpinh1a |
| rps27.12 | 3.7E-48 | 0.2031143 | 0.987 | 0.977 | 1.22E-43 | Regenerative Mesenchyme | rps27.1 |
| phf14 | 4.5E-48 | 0.8174539 | 0.211 | 0.112 | 1.45E-43 | Regenerative Mesenchyme | phf14 |
| chordc1b | 4.8E-48 | 0.9050632 | 0.19 | 0.097 | 1.58E-43 | Regenerative Mesenchyme | chordc1b |
| knop11 | 9.3E-48 | 0.6612783 | 0.273 | 0.157 | 3.02E-43 | Regenerative Mesenchyme | knop1 |
| nucks1a | 1.2E-47 | 0.638558 | 0.494 | 0.355 | 3.8E-43 | Regenerative Mesenchyme | nucks1a |
| tktb | 1.2E-47 | 0.8461182 | 0.239 | 0.134 | 3.85E-43 | Regenerative Mesenchyme | tktb |
| kri1 | 1.3E-47 | 0.8196544 | 0.19 | 0.097 | 4.18E-43 | Regenerative Mesenchyme | kri1 |
| ccdc12 | 1.4E-47 | 0.9570399 | 0.161 | 0.077 | 4.55E-43 | Regenerative Mesenchyme | ccdc12 |
| mrpl38 | 1.5E-47 | 1.0356095 | 0.131 | 0.058 | 4.75E-43 | Regenerative Mesenchyme | mrpl38 |
| idh3a | 1.8E-47 | 1.0578081 | 0.157 | 0.075 | 5.75E-43 | Regenerative Mesenchyme | idh3a |
| api5 | 2.3E-47 | 0.8667462 | 0.239 | 0.133 | 7.39E-43 | Regenerative Mesenchyme | api5 |
| ifi45 | 2.8E-47 | 0.473426 | 0.352 | 0.222 | 9.13E-43 | Regenerative Mesenchyme | ifi45 |
| arl14ep | 3.5E-47 | 1.1478271 | 0.12 | 0.052 | 1.13E-42 | Regenerative Mesenchyme | arl14ep |
| dap1 | 4.7E-47 | 0.7664614 | 0.308 | 0.189 | 1.52E-42 | Regenerative Mesenchyme | dap |
| ccdc9 | 5.2E-47 | 0.9033339 | 0.157 | 0.075 | 1.68E-42 | Regenerative Mesenchyme | ccdc9 |
| zcchc17 | 5.6E-47 | 0.8886704 | 0.119 | 0.05 | 1.84E-42 | Regenerative Mesenchyme | zcchc17 |
| lrrc173 | 6.1E-47 | 0.6553673 | 0.518 | 0.39 | 1.99E-42 | Regenerative Mesenchyme | lrrc17 |
| dip2ba | 7.9E-47 | 1.1921545 | 0.149 | 0.071 | 2.56E-42 | Regenerative Mesenchyme | dip2ba |
| cdk11b | 8.3E-47 | 1.0083523 | 0.137 | 0.062 | 2.7E-42 | Regenerative Mesenchyme | cdk11b |
| emc1 | 8.7E-47 | 1.2708913 | 0.109 | 0.045 | 2.83E-42 | Regenerative Mesenchyme | emc1 |
| atpaf1 | 1.1E-46 | 1.1071057 | 0.122 | 0.053 | 3.61E-42 | Regenerative Mesenchyme | atpaf1 |
| chd61 | 1.4E-46 | 0.6750957 | 0.289 | 0.173 | 4.59E-42 | Regenerative Mesenchyme | chd6 |
| ttf1 | 1.4E-46 | 0.996975 | 0.109 | 0.045 | 4.67E-42 | Regenerative Mesenchyme | ttf1 |
| ddx23 | 1.5E-46 | 0.722667 | 0.27 | 0.157 | 4.94E-42 | Regenerative Mesenchyme | ddx23 |
| hsbp1b | 2E-46 | 0.6904654 | 0.395 | 0.264 | 6.51E-42 | Regenerative Mesenchyme | hsbp1b |
| myof1 | 3.2E-46 | 0.6990221 | 0.239 | 0.133 | 1.05E-41 | Regenerative Mesenchyme | myof |
| polr2i | 3.6E-46 | 1.1637487 | 0.13 | 0.058 | 1.16E-41 | Regenerative Mesenchyme | polr2i |
| rpap3 | 3.8E-46 | 1.136725 | 0.112 | 0.047 | 1.25E-41 | Regenerative Mesenchyme | rpap3 |
| prrc2c | 4.3E-46 | 0.8015963 | 0.21 | 0.112 | 1.41E-41 | Regenerative Mesenchyme | prrc2c |
| mtpn | 4.5E-46 | 0.5917927 | 0.442 | 0.305 | 1.46E-41 | Regenerative Mesenchyme | mtpn |
| immt | 5.1E-46 | 0.9437564 | 0.17 | 0.085 | 1.65E-41 | Regenerative Mesenchyme | immt |
| tomm34 | 5.2E-46 | 1.3917459 | 0.103 | 0.042 | 1.71E-41 | Regenerative Mesenchyme | tomm34 |
| marveld1 | 7E-46 | 0.685943 | 0.325 | 0.203 | 2.29E-41 | Regenerative Mesenchyme | marveld1 |
| polr3h | 8.4E-46 | 1.1004597 | 0.138 | 0.064 | 2.73E-41 | Regenerative Mesenchyme | polr3h |
| vkorc1l1 | 9.6E-46 | 0.9419657 | 0.172 | 0.086 | 3.12E-41 | Regenerative Mesenchyme | vkorc1l1 |
| srp72 | 1.4E-45 | 0.8136838 | 0.251 | 0.145 | 4.65E-41 | Regenerative Mesenchyme | srp72 |
| eif4g2a | 1.8E-45 | 0.7686083 | 0.293 | 0.178 | 5.71E-41 | Regenerative Mesenchyme | eif4g2a |
| cox171 | 1.8E-45 | 0.8966634 | 0.25 | 0.145 | 5.9E-41 | Regenerative Mesenchyme | cox17 |
| gnl3l | 1.9E-45 | 1.0836215 | 0.116 | 0.05 | 6.06E-41 | Regenerative Mesenchyme | gnl3l |
| rcl1 | 2E-45 | 1.1291694 | 0.113 | 0.048 | 6.58E-41 | Regenerative Mesenchyme | rcl1 |

| | | | | | | | |
|---|---|---|---|---|---|---|---|
| mycbp | 2.2E-45 | 0.6599322 | 0.369 | 0.242 | 7.19E-41 | Regenerative Mesenchyme | mycbp |
| capza1b | 2.6E-45 | 0.6448854 | 0.49 | 0.36 | 8.32E-41 | Regenerative Mesenchyme | capza1b |
| zgc:162025 | 2.8E-45 | 0.9091753 | 0.163 | 0.08 | 9.04E-41 | Regenerative Mesenchyme | zgc:162025 |
| ak22 | 4.5E-45 | 0.5841309 | 0.504 | 0.373 | 1.48E-40 | Regenerative Mesenchyme | ak2 |
| hnrnpl2 | 5.9E-45 | 0.7943448 | 0.249 | 0.143 | 1.92E-40 | Regenerative Mesenchyme | hnrnpl2 |
| mrpl48 | 7.2E-45 | 1.0855079 | 0.122 | 0.054 | 2.34E-40 | Regenerative Mesenchyme | mrpl48 |
| tia1 | 9E-45 | 0.8393474 | 0.175 | 0.088 | 2.93E-40 | Regenerative Mesenchyme | tia1 |
| rab34a1 | 1.1E-44 | 0.9501792 | 0.208 | 0.114 | 3.63E-40 | Regenerative Mesenchyme | rab34a |
| prps1a | 1.3E-44 | 0.9206826 | 0.205 | 0.111 | 4.28E-40 | Regenerative Mesenchyme | prps1a |
| srp54 | 1.8E-44 | 0.6939391 | 0.284 | 0.169 | 5.77E-40 | Regenerative Mesenchyme | srp54 |
| COA4 | 2E-44 | 1.163869 | 0.11 | 0.047 | 6.49E-40 | Regenerative Mesenchyme | COA4 |
| usp5 | 2E-44 | 0.9355127 | 0.192 | 0.102 | 6.63E-40 | Regenerative Mesenchyme | usp5 |
| cldnb2 | 3.1E-44 | 1.0354237 | 0.241 | 0.143 | 9.94E-40 | Regenerative Mesenchyme | cldnb |
| m6pr | 3.7E-44 | 0.813737 | 0.216 | 0.119 | 1.19E-39 | Regenerative Mesenchyme | m6pr |
| dhx9 | 3.9E-44 | 0.9769847 | 0.147 | 0.07 | 1.26E-39 | Regenerative Mesenchyme | dhx9 |
| htatsf1 | 5.2E-44 | 0.6876379 | 0.207 | 0.111 | 1.68E-39 | Regenerative Mesenchyme | htatsf1 |
| ankzf1 | 5.4E-44 | 1.1361434 | 0.104 | 0.043 | 1.74E-39 | Regenerative Mesenchyme | ankzf1 |
| etfb | 6.2E-44 | 0.7885498 | 0.274 | 0.163 | 2.01E-39 | Regenerative Mesenchyme | etfb |
| rap1aa | 7.7E-44 | 0.783583 | 0.228 | 0.128 | 2.52E-39 | Regenerative Mesenchyme | rap1aa |
| mrps28 | 8.9E-44 | 1.0563395 | 0.119 | 0.052 | 2.9E-39 | Regenerative Mesenchyme | mrps28 |
| leo1 | 1E-43 | 0.6966906 | 0.256 | 0.148 | 3.34E-39 | Regenerative Mesenchyme | leo1 |
| ube2e11 | 1.1E-43 | 0.8835627 | 0.206 | 0.112 | 3.69E-39 | Regenerative Mesenchyme | ube2e1 |
| aimp21 | 1.2E-43 | 0.7812095 | 0.275 | 0.166 | 4.05E-39 | Regenerative Mesenchyme | aimp2 |
| ppp2r1bb | 1.3E-43 | 0.9557063 | 0.175 | 0.09 | 4.18E-39 | Regenerative Mesenchyme | ppp2r1bb |
| mrpl4 | 1.4E-43 | 1.0184348 | 0.131 | 0.06 | 4.43E-39 | Regenerative Mesenchyme | mrpl4 |
| timm17a | 1.4E-43 | 0.8853019 | 0.191 | 0.101 | 4.46E-39 | Regenerative Mesenchyme | timm17a |
| sept2 | 1.4E-43 | 0.6879497 | 0.379 | 0.252 | 4.48E-39 | Regenerative Mesenchyme | sept2 |
| nol11 | 1.5E-43 | 1.2502359 | 0.1 | 0.041 | 4.87E-39 | Regenerative Mesenchyme | nol11 |
| srek1ip1 | 1.5E-43 | 1.0795826 | 0.104 | 0.043 | 4.89E-39 | Regenerative Mesenchyme | srek1ip1 |
| lamtor21 | 1.5E-43 | 0.9230974 | 0.21 | 0.116 | 4.97E-39 | Regenerative Mesenchyme | lamtor2 |
| zgc:101858 | 1.6E-43 | 1.1180448 | 0.115 | 0.05 | 5.04E-39 | Regenerative Mesenchyme | zgc:101858 |
| atp6v1g11 | 1.7E-43 | 0.8413513 | 0.225 | 0.127 | 5.53E-39 | Regenerative Mesenchyme | atp6v1g1 |
| efna5a2 | 1.7E-43 | 1.0332144 | 0.151 | 0.074 | 5.68E-39 | Regenerative Mesenchyme | efna5a |
| cebpz | 2.2E-43 | 0.9328586 | 0.137 | 0.064 | 7.21E-39 | Regenerative Mesenchyme | cebpz |
| ap2a11 | 2.3E-43 | 0.6833458 | 0.293 | 0.178 | 7.56E-39 | Regenerative Mesenchyme | ap2a1 |
| eif3c | 2.9E-43 | 0.5318663 | 0.613 | 0.502 | 9.46E-39 | Regenerative Mesenchyme | eif3c |
| FO704772.2 | 3.4E-43 | 0.8064823 | 0.356 | 0.237 | 1.1E-38 | Regenerative Mesenchyme | FO704772.2 |
| arf6a | 3.5E-43 | 0.9204278 | 0.195 | 0.105 | 1.15E-38 | Regenerative Mesenchyme | arf6a |
| cnih1 | 4.8E-43 | 0.941022 | 0.177 | 0.092 | 1.56E-38 | Regenerative Mesenchyme | cnih1 |
| marcksl1b2 | 5.9E-43 | 0.3302927 | 0.38 | 0.247 | 1.92E-38 | Regenerative Mesenchyme | marcksl1b |
| b3gnt5a2 | 6.4E-43 | 0.8795282 | 0.198 | 0.107 | 2.09E-38 | Regenerative Mesenchyme | b3gnt5a |
| snrnp200 | 7.4E-43 | 0.862198 | 0.2 | 0.108 | 2.4E-38 | Regenerative Mesenchyme | snrnp200 |
| xpot | 7.8E-43 | 1.1470884 | 0.123 | 0.055 | 2.52E-38 | Regenerative Mesenchyme | xpot |
| nsmce2 | 8.9E-43 | 1.1715988 | 0.111 | 0.048 | 2.91E-38 | Regenerative Mesenchyme | nsmce2 |
| si:dkey-238c7.161 | 1.1E-42 | 0.6414453 | 0.27 | 0.159 | 3.45E-38 | Regenerative Mesenchyme | si:dkey-238c7.16 |
| dhx37 | 1.1E-42 | 0.9317212 | 0.127 | 0.058 | 3.66E-38 | Regenerative Mesenchyme | dhx37 |
| ergic3 | 1.3E-42 | 1.2276417 | 0.125 | 0.058 | 4.08E-38 | Regenerative Mesenchyme | ergic3 |
| npepl1 | 1.3E-42 | 0.8064274 | 0.199 | 0.108 | 4.33E-38 | Regenerative Mesenchyme | npepl1 |

| | | | | | | | |
|---|---|---|---|---|---|---|---|
| arf4a.11 | 1.5E-42 | 1.0620355 | 0.146 | 0.071 | 5E-38 | Regenerative Mesenchyme | arf4a.1 |
| MZT1 | 1.7E-42 | 1.0812721 | 0.148 | 0.072 | 5.39E-38 | Regenerative Mesenchyme | MZT1 |
| klf12a1 | 2.2E-42 | 0.9433474 | 0.231 | 0.133 | 7.13E-38 | Regenerative Mesenchyme | klf12a |
| csde1 | 2.4E-42 | 0.6342279 | 0.515 | 0.385 | 7.67E-38 | Regenerative Mesenchyme | csde1 |
| cnot9 | 2.7E-42 | 1.0217054 | 0.151 | 0.074 | 8.9E-38 | Regenerative Mesenchyme | cnot9 |
| trappc31 | 2.7E-42 | 0.8815351 | 0.19 | 0.102 | 8.93E-38 | Regenerative Mesenchyme | trappc3 |
| hsd17b10 | 3.5E-42 | 0.9307777 | 0.138 | 0.065 | 1.13E-37 | Regenerative Mesenchyme | hsd17b10 |
| si:ch211-22i13.2 | 3.8E-42 | 0.9448324 | 0.145 | 0.071 | 1.25E-37 | Regenerative Mesenchyme | si:ch211-22i13.2 |
| noc3l | 4.5E-42 | 1.0886842 | 0.102 | 0.042 | 1.46E-37 | Regenerative Mesenchyme | noc3l |
| derl2 | 5.2E-42 | 0.9316078 | 0.171 | 0.088 | 1.68E-37 | Regenerative Mesenchyme | derl2 |
| stx18 | 5.9E-42 | 1.0859105 | 0.115 | 0.051 | 1.91E-37 | Regenerative Mesenchyme | stx18 |
| bet1 | 7.3E-42 | 0.8273971 | 0.18 | 0.094 | 2.37E-37 | Regenerative Mesenchyme | bet1 |
| rbms1a | 7.6E-42 | 0.9732779 | 0.147 | 0.072 | 2.47E-37 | Regenerative Mesenchyme | rbms1a |
| rbm19 | 8E-42 | 0.7404568 | 0.214 | 0.119 | 2.59E-37 | Regenerative Mesenchyme | rbm19 |
| plod22 | 8.1E-42 | 1.1783175 | 0.14 | 0.068 | 2.62E-37 | Regenerative Mesenchyme | plod2 |
| mrps14 | 1.1E-41 | 0.8559023 | 0.222 | 0.127 | 3.61E-37 | Regenerative Mesenchyme | mrps14 |
| wnk1a | 1.1E-41 | 1.0139186 | 0.125 | 0.057 | 3.7E-37 | Regenerative Mesenchyme | wnk1a |
| rps32 | 1.2E-41 | 0.2304032 | 0.973 | 0.946 | 3.8E-37 | Regenerative Mesenchyme | rps3 |
| bcap31 | 1.2E-41 | 0.7945837 | 0.184 | 0.097 | 3.93E-37 | Regenerative Mesenchyme | bcap31 |
| ppp1r14bb2 | 1.5E-41 | 0.7834828 | 0.245 | 0.144 | 4.98E-37 | Regenerative Mesenchyme | ppp1r14bb |
| actn4 | 1.9E-41 | 0.8956175 | 0.193 | 0.105 | 6.28E-37 | Regenerative Mesenchyme | actn4 |
| si:dkey-12j5.1 | 2.3E-41 | 1.0971335 | 0.114 | 0.051 | 7.59E-37 | Regenerative Mesenchyme | si:dkey-12j5.1 |
| rpl382 | 2.6E-41 | 0.3059258 | 0.932 | 0.91 | 8.41E-37 | Regenerative Mesenchyme | rpl38 |
| p4ha1b2 | 2.9E-41 | 0.9315602 | 0.231 | 0.135 | 9.5E-37 | Regenerative Mesenchyme | p4ha1b |
| guk1a | 3.2E-41 | 0.8395181 | 0.228 | 0.132 | 1.03E-36 | Regenerative Mesenchyme | guk1a |
| arcn1b1 | 4.8E-41 | 0.8776413 | 0.211 | 0.118 | 1.58E-36 | Regenerative Mesenchyme | arcn1b |
| cdk8 | 9.8E-41 | 1.022654 | 0.114 | 0.051 | 3.19E-36 | Regenerative Mesenchyme | cdk8 |
| prrc11 | 1.1E-40 | 1.053523 | 0.15 | 0.075 | 3.7E-36 | Regenerative Mesenchyme | prrc1 |
| p3h11 | 1.2E-40 | 0.7022048 | 0.321 | 0.207 | 4.02E-36 | Regenerative Mesenchyme | p3h1 |
| pitpnb | 1.5E-40 | 0.9792256 | 0.105 | 0.045 | 5.01E-36 | Regenerative Mesenchyme | pitpnb |
| vbp11 | 1.6E-40 | 0.8315123 | 0.215 | 0.123 | 5.14E-36 | Regenerative Mesenchyme | vbp1 |
| pagr1 | 1.6E-40 | 1.0894471 | 0.127 | 0.059 | 5.32E-36 | Regenerative Mesenchyme | pagr1 |
| ddx51 | 1.8E-40 | 0.8473053 | 0.117 | 0.052 | 5.88E-36 | Regenerative Mesenchyme | ddx51 |
| mt-co11 | 1.8E-40 | 0.271388 | 0.996 | 0.995 | 5.98E-36 | Regenerative Mesenchyme | mt-co1 |
| taf13 | 1.9E-40 | 0.659062 | 0.328 | 0.212 | 6.04E-36 | Regenerative Mesenchyme | taf13 |
| dohh1 | 1.9E-40 | 0.8304342 | 0.242 | 0.144 | 6.19E-36 | Regenerative Mesenchyme | dohh |
| atp6v1e1b1 | 2E-40 | 0.9278076 | 0.178 | 0.095 | 6.53E-36 | Regenerative Mesenchyme | atp6v1e1b |
| cops5 | 2.1E-40 | 0.9149631 | 0.129 | 0.06 | 6.67E-36 | Regenerative Mesenchyme | cops5 |
| errfi1a | 2.1E-40 | 1.1956958 | 0.109 | 0.048 | 6.98E-36 | Regenerative Mesenchyme | errfi1a |
| isg20 | 2.2E-40 | 1.1935976 | 0.107 | 0.047 | 7.24E-36 | Regenerative Mesenchyme | isg20 |
| rpf1 | 2.6E-40 | 0.8559478 | 0.161 | 0.082 | 8.59E-36 | Regenerative Mesenchyme | rpf1 |
| ppp1r2 | 4.5E-40 | 0.9559259 | 0.12 | 0.055 | 1.46E-35 | Regenerative Mesenchyme | ppp1r2 |
| cdipt | 6.6E-40 | 0.7778007 | 0.199 | 0.11 | 2.16E-35 | Regenerative Mesenchyme | cdipt |
| eif4e2rs1 | 7.1E-40 | 0.9517982 | 0.127 | 0.059 | 2.3E-35 | Regenerative Mesenchyme | eif4e2rs1 |
| atic | 7.5E-40 | 1.2024307 | 0.118 | 0.054 | 2.45E-35 | Regenerative Mesenchyme | atic |
| mt-nd31 | 9.4E-40 | 0.6058446 | 0.48 | 0.354 | 3.05E-35 | Regenerative Mesenchyme | mt-nd3 |
| edf1 | 1.1E-39 | 0.4743393 | 0.614 | 0.507 | 3.58E-35 | Regenerative Mesenchyme | edf1 |
| itgb51 | 1.1E-39 | 0.8390719 | 0.158 | 0.081 | 3.69E-35 | Regenerative Mesenchyme | itgb5 |

| | | | | | | | |
|---|---|---|---|---|---|---|---|
| cndp21 | 1.2E-39 | 0.8982894 | 0.193 | 0.107 | 3.99E-35 | Regenerative Mesenchyme | cndp2 |
| hnrnpc | 1.6E-39 | 0.8543917 | 0.177 | 0.095 | 5.16E-35 | Regenerative Mesenchyme | hnrnpc |
| sdad1 | 1.9E-39 | 0.7128556 | 0.193 | 0.106 | 6.02E-35 | Regenerative Mesenchyme | sdad1 |
| znf330 | 2.2E-39 | 1.1740443 | 0.113 | 0.051 | 7.26E-35 | Regenerative Mesenchyme | znf330 |
| skp1 | 2.3E-39 | 0.4858713 | 0.606 | 0.493 | 7.32E-35 | Regenerative Mesenchyme | skp1 |
| cd151l1 | 2.3E-39 | 1.249922 | 0.141 | 0.07 | 7.49E-35 | Regenerative Mesenchyme | cd151l |
| ppp1caa | 2.7E-39 | 0.9329006 | 0.223 | 0.131 | 8.88E-35 | Regenerative Mesenchyme | ppp1caa |
| nap1l4a | 2.8E-39 | 0.837643 | 0.17 | 0.089 | 9.15E-35 | Regenerative Mesenchyme | nap1l4a |
| spaca4l | 3.5E-39 | 0.9114201 | 0.153 | 0.079 | 1.12E-34 | Regenerative Mesenchyme | spaca4l |
| hspa9 | 3.5E-39 | 0.7486934 | 0.29 | 0.184 | 1.13E-34 | Regenerative Mesenchyme | hspa9 |
| parp3 | 5.3E-39 | 1.0059232 | 0.16 | 0.083 | 1.72E-34 | Regenerative Mesenchyme | parp3 |
| rac1a | 5.7E-39 | 0.7789226 | 0.288 | 0.182 | 1.85E-34 | Regenerative Mesenchyme | rac1a |
| pabpn1 | 6.2E-39 | 0.518299 | 0.528 | 0.404 | 2.01E-34 | Regenerative Mesenchyme | pabpn1 |
| ybx11 | 6.7E-39 | 0.2055446 | 0.984 | 0.969 | 2.19E-34 | Regenerative Mesenchyme | ybx1 |
| xirp2a1 | 8E-39 | 0.5255655 | 0.374 | 0.251 | 2.6E-34 | Regenerative Mesenchyme | xirp2a |
| gtf2f1 | 1E-38 | 0.6153494 | 0.274 | 0.168 | 3.24E-34 | Regenerative Mesenchyme | gtf2f1 |
| ntn1a1 | 1E-38 | 0.4339201 | 0.209 | 0.116 | 3.29E-34 | Regenerative Mesenchyme | ntn1a |
| sf3a3 | 1E-38 | 0.8746099 | 0.139 | 0.068 | 3.41E-34 | Regenerative Mesenchyme | sf3a3 |
| gpia | 1.1E-38 | 0.9035734 | 0.214 | 0.123 | 3.62E-34 | Regenerative Mesenchyme | gpia |
| aifm1 | 1.1E-38 | 1.2064787 | 0.111 | 0.05 | 3.66E-34 | Regenerative Mesenchyme | aifm1 |
| hprt1 | 1.2E-38 | 0.9989407 | 0.12 | 0.056 | 3.78E-34 | Regenerative Mesenchyme | hprt1 |
| GCA | 1.2E-38 | 0.6347063 | 0.407 | 0.29 | 3.94E-34 | Regenerative Mesenchyme | GCA |
| cct6a | 1.2E-38 | 0.5237342 | 0.576 | 0.474 | 3.96E-34 | Regenerative Mesenchyme | cct6a |
| dr1 | 1.3E-38 | 1.0394543 | 0.137 | 0.068 | 4.2E-34 | Regenerative Mesenchyme | dr1 |
| ngfb | 1.3E-38 | 0.9510549 | 0.153 | 0.078 | 4.22E-34 | Regenerative Mesenchyme | ngfb |
| hnrnpm | 1.8E-38 | 0.6544613 | 0.286 | 0.178 | 5.77E-34 | Regenerative Mesenchyme | hnrnpm |
| smu1a | 1.8E-38 | 1.0793714 | 0.117 | 0.054 | 5.89E-34 | Regenerative Mesenchyme | smu1a |
| mrpl15 | 1.9E-38 | 0.9242849 | 0.132 | 0.064 | 6.27E-34 | Regenerative Mesenchyme | mrpl15 |
| lsm12b | 1.9E-38 | 0.5276997 | 0.455 | 0.328 | 6.32E-34 | Regenerative Mesenchyme | lsm12b |
| srsf11 | 2.1E-38 | 0.6161604 | 0.357 | 0.237 | 6.76E-34 | Regenerative Mesenchyme | srsf11 |
| ccdc801 | 2.5E-38 | 0.2375587 | 0.524 | 0.379 | 8.05E-34 | Regenerative Mesenchyme | ccdc80 |
| smarcc1a | 2.7E-38 | 0.9469671 | 0.14 | 0.069 | 8.91E-34 | Regenerative Mesenchyme | smarcc1a |
| wdr43 | 2.8E-38 | 0.8745158 | 0.243 | 0.148 | 9.15E-34 | Regenerative Mesenchyme | wdr43 |
| fnbp1l | 3.3E-38 | 0.876468 | 0.133 | 0.064 | 1.07E-33 | Regenerative Mesenchyme | fnbp1l |
| CABZ01044053.1 | 4.7E-38 | 0.8450677 | 0.152 | 0.077 | 1.54E-33 | Regenerative Mesenchyme | CABZ01044053.1 |
| npm1b | 5E-38 | 0.8716206 | 0.181 | 0.098 | 1.63E-33 | Regenerative Mesenchyme | npm1b |
| surf2 | 5.9E-38 | 1.0421491 | 0.111 | 0.05 | 1.92E-33 | Regenerative Mesenchyme | surf2 |
| rbbp91 | 6.2E-38 | 0.7766878 | 0.21 | 0.12 | 2.01E-33 | Regenerative Mesenchyme | rbbp9 |
| COX7A2 | 7.5E-38 | 0.7911925 | 0.218 | 0.126 | 2.44E-33 | Regenerative Mesenchyme | COX7A2 |
| wdr36 | 1E-37 | 0.8350911 | 0.1 | 0.043 | 3.26E-33 | Regenerative Mesenchyme | wdr36 |
| gtf2f2a | 1.4E-37 | 0.8879714 | 0.219 | 0.129 | 4.41E-33 | Regenerative Mesenchyme | gtf2f2a |
| smim20 | 1.5E-37 | 0.8837764 | 0.151 | 0.077 | 4.96E-33 | Regenerative Mesenchyme | smim20 |
| adam9 | 1.8E-37 | 0.9652325 | 0.131 | 0.064 | 5.84E-33 | Regenerative Mesenchyme | adam9 |
| pld72 | 2.1E-37 | 0.6990655 | 0.245 | 0.148 | 6.8E-33 | Regenerative Mesenchyme | pld7 |
| uba2 | 2.5E-37 | 1.0531687 | 0.107 | 0.048 | 8.05E-33 | Regenerative Mesenchyme | uba2 |
| hcfc1a | 2.7E-37 | 0.909415 | 0.169 | 0.091 | 8.85E-33 | Regenerative Mesenchyme | hcfc1a |
| gtf2a2 | 3.2E-37 | 0.7497059 | 0.191 | 0.106 | 1.03E-32 | Regenerative Mesenchyme | gtf2a2 |
| sap30bp | 3.2E-37 | 0.7732378 | 0.159 | 0.083 | 1.04E-32 | Regenerative Mesenchyme | sap30bp |

| | | | | | | | |
|---|---|---|---|---|---|---|---|
| arpc4l | 3.3E-37 | 0.5481031 | 0.479 | 0.357 | 1.07E-32 | Regenerative Mesenchyme | arpc4l |
| eif3ba1 | 3.3E-37 | 0.5684869 | 0.533 | 0.423 | 1.07E-32 | Regenerative Mesenchyme | eif3ba |
| si:dkeyp-120h9.11 | 3.4E-37 | 1.0071782 | 0.154 | 0.08 | 1.12E-32 | Regenerative Mesenchyme | si:dkeyp-120h9.1 |
| acadsb1 | 5.1E-37 | 0.9690261 | 0.126 | 0.061 | 1.67E-32 | Regenerative Mesenchyme | acadsb |
| dio3b | 5.1E-37 | 1.0203948 | 0.113 | 0.052 | 1.67E-32 | Regenerative Mesenchyme | dio3b |
| mt-nd4l | 5.4E-37 | 0.9130087 | 0.143 | 0.073 | 1.76E-32 | Regenerative Mesenchyme | mt-nd4l |
| rsl1d11 | 9.4E-37 | 0.5080586 | 0.487 | 0.368 | 3.05E-32 | Regenerative Mesenchyme | rsl1d1 |
| ndufs8a1 | 1.1E-36 | 1.0272928 | 0.164 | 0.088 | 3.49E-32 | Regenerative Mesenchyme | ndufs8a |
| arl1 | 1.3E-36 | 0.7001143 | 0.26 | 0.16 | 4.13E-32 | Regenerative Mesenchyme | arl1 |
| sdf2 | 1.5E-36 | 1.0005774 | 0.117 | 0.055 | 4.72E-32 | Regenerative Mesenchyme | sdf2 |
| krt51 | 1.8E-36 | 1.0643538 | 0.579 | 0.486 | 5.79E-32 | Regenerative Mesenchyme | krt5 |
| riok2 | 1.8E-36 | 0.9933999 | 0.11 | 0.05 | 6E-32 | Regenerative Mesenchyme | riok2 |
| rbm25b | 2E-36 | 0.6148908 | 0.314 | 0.204 | 6.41E-32 | Regenerative Mesenchyme | rbm25b |
| rpe | 2.4E-36 | 0.7966327 | 0.167 | 0.09 | 7.94E-32 | Regenerative Mesenchyme | rpe |
| sri | 2.9E-36 | 0.5918405 | 0.382 | 0.265 | 9.53E-32 | Regenerative Mesenchyme | sri |
| ctnnb11 | 3.4E-36 | 0.4558579 | 0.61 | 0.499 | 1.11E-31 | Regenerative Mesenchyme | ctnnb1 |
| arhgap42a | 4.6E-36 | 0.9513418 | 0.137 | 0.069 | 1.51E-31 | Regenerative Mesenchyme | arhgap42a |
| gemin71 | 5.3E-36 | 0.9610071 | 0.133 | 0.066 | 1.71E-31 | Regenerative Mesenchyme | gemin7 |
| elovl1b | 5.8E-36 | 0.4237472 | 0.474 | 0.345 | 1.88E-31 | Regenerative Mesenchyme | elovl1b |
| gnb1b | 7.4E-36 | 0.4237079 | 0.654 | 0.563 | 2.42E-31 | Regenerative Mesenchyme | gnb1b |
| mt-co21 | 7.6E-36 | 0.221685 | 1 | 0.998 | 2.47E-31 | Regenerative Mesenchyme | mt-co2 |
| nkap | 1.3E-35 | 0.5761159 | 0.288 | 0.182 | 4.1E-31 | Regenerative Mesenchyme | nkap |
| lepr | 1.6E-35 | 0.9535799 | 0.124 | 0.06 | 5.21E-31 | Regenerative Mesenchyme | lepr |
| wdr82 | 1.6E-35 | 1.1173527 | 0.118 | 0.057 | 5.33E-31 | Regenerative Mesenchyme | wdr82 |
| mrpl2 | 2.6E-35 | 0.9541582 | 0.158 | 0.085 | 8.36E-31 | Regenerative Mesenchyme | mrpl2 |
| zgc:1536751 | 2.8E-35 | 0.6459947 | 0.366 | 0.257 | 9.06E-31 | Regenerative Mesenchyme | zgc:153675 |
| atf6 | 2.9E-35 | 0.6910323 | 0.199 | 0.113 | 9.3E-31 | Regenerative Mesenchyme | atf6 |
| ptp4a2b | 2.9E-35 | 0.5369037 | 0.416 | 0.298 | 9.32E-31 | Regenerative Mesenchyme | ptp4a2b |
| cdk5rap3 | 2.9E-35 | 0.9512819 | 0.11 | 0.051 | 9.32E-31 | Regenerative Mesenchyme | cdk5rap3 |
| cmc11 | 3.4E-35 | 0.9527623 | 0.169 | 0.093 | 1.09E-30 | Regenerative Mesenchyme | cmc1 |
| eif1axa | 3.7E-35 | 0.7752488 | 0.172 | 0.094 | 1.22E-30 | Regenerative Mesenchyme | eif1axa |
| EMC4 | 4.7E-35 | 1.0236135 | 0.117 | 0.056 | 1.54E-30 | Regenerative Mesenchyme | EMC4 |
| cyth1a.1 | 7.3E-35 | 0.7658919 | 0.221 | 0.131 | 2.38E-30 | Regenerative Mesenchyme | cyth1a.1 |
| sil1 | 9.2E-35 | 0.9271844 | 0.127 | 0.063 | 2.99E-30 | Regenerative Mesenchyme | sil1 |
| skib1 | 1.1E-34 | 0.6099114 | 0.301 | 0.197 | 3.54E-30 | Regenerative Mesenchyme | skib |
| nck1b | 1.1E-34 | 0.8597482 | 0.17 | 0.093 | 3.74E-30 | Regenerative Mesenchyme | nck1b |
| gfpt2 | 1.3E-34 | 0.9330205 | 0.13 | 0.065 | 4.15E-30 | Regenerative Mesenchyme | gfpt2 |
| rps27l | 1.4E-34 | 0.7580749 | 0.392 | 0.283 | 4.5E-30 | Regenerative Mesenchyme | rps27l |
| jagn1b | 1.4E-34 | 0.6327862 | 0.253 | 0.157 | 4.65E-30 | Regenerative Mesenchyme | jagn1b |
| fabp11a | 1.9E-34 | 0.4847691 | 0.137 | 0.07 | 6.27E-30 | Regenerative Mesenchyme | fabp11a |
| erlec1 | 2.3E-34 | 0.8100236 | 0.198 | 0.115 | 7.34E-30 | Regenerative Mesenchyme | erlec1 |
| pmpcb | 2.3E-34 | 0.9629344 | 0.12 | 0.058 | 7.44E-30 | Regenerative Mesenchyme | pmpcb |
| fam207a1 | 2.4E-34 | 0.7960271 | 0.144 | 0.075 | 7.91E-30 | Regenerative Mesenchyme | fam207a |
| ubap2b | 2.5E-34 | 0.6995492 | 0.252 | 0.157 | 8E-30 | Regenerative Mesenchyme | ubap2b |
| hsdl21 | 3E-34 | 1.0469849 | 0.138 | 0.071 | 9.69E-30 | Regenerative Mesenchyme | hsdl2 |
| si:ch211-218c6.8 | 3.4E-34 | 0.8011193 | 0.184 | 0.104 | 1.11E-29 | Regenerative Mesenchyme | si:ch211-218c6.8 |
| wdr75 | 4.2E-34 | 1.0663684 | 0.111 | 0.053 | 1.38E-29 | Regenerative Mesenchyme | wdr75 |
| xrcc4 | 4.7E-34 | 0.7992601 | 0.11 | 0.052 | 1.52E-29 | Regenerative Mesenchyme | xrcc4 |

| | | | | | | | |
|---|---|---|---|---|---|---|---|
| dhx15 | 6.7E-34 | 0.7842944 | 0.187 | 0.107 | 2.19E-29 | Regenerative Mesenchyme | dhx15 |
| hnrnpul1 | 7.7E-34 | 1.0444783 | 0.124 | 0.062 | 2.51E-29 | Regenerative Mesenchyme | hnrnpul1 |
| si:ch211-162e15.3 | 8.8E-34 | 0.7674799 | 0.153 | 0.081 | 2.86E-29 | Regenerative Mesenchyme | si:ch211-162e15.3 |
| ube2nb | 9.1E-34 | 0.6461517 | 0.225 | 0.134 | 2.98E-29 | Regenerative Mesenchyme | ube2nb |
| hikeshi | 1.1E-33 | 0.853438 | 0.177 | 0.1 | 3.42E-29 | Regenerative Mesenchyme | hikeshi |
| pgd | 1.1E-33 | 0.9391736 | 0.144 | 0.075 | 3.56E-29 | Regenerative Mesenchyme | pgd |
| nbas | 1.4E-33 | 0.7926947 | 0.171 | 0.095 | 4.4E-29 | Regenerative Mesenchyme | nbas |
| smndc1 | 2.4E-33 | 0.5187535 | 0.309 | 0.202 | 7.96E-29 | Regenerative Mesenchyme | smndc1 |
| acbd3 | 3.8E-33 | 0.8319576 | 0.217 | 0.132 | 1.24E-28 | Regenerative Mesenchyme | acbd3 |
| cstf2 | 4.5E-33 | 0.8217072 | 0.171 | 0.095 | 1.45E-28 | Regenerative Mesenchyme | cstf2 |
| kctd5a1 | 7.4E-33 | 0.5017981 | 0.429 | 0.313 | 2.42E-28 | Regenerative Mesenchyme | kctd5a |
| si:ch1073-314i13.4 | 7.4E-33 | 0.8170489 | 0.141 | 0.074 | 2.42E-28 | Regenerative Mesenchyme | si:ch1073-314i13.4 |
| arpc2 | 8.2E-33 | 0.4992464 | 0.546 | 0.438 | 2.66E-28 | Regenerative Mesenchyme | arpc2 |
| med19a | 9.9E-33 | 0.7338701 | 0.213 | 0.128 | 3.22E-28 | Regenerative Mesenchyme | med19a |
| srfbp1 | 1.1E-32 | 0.6794188 | 0.166 | 0.092 | 3.47E-28 | Regenerative Mesenchyme | srfbp1 |
| rhocb | 1.1E-32 | 0.6918619 | 0.232 | 0.143 | 3.54E-28 | Regenerative Mesenchyme | rhocb |
| mtrex | 1.2E-32 | 0.88836 | 0.116 | 0.057 | 3.87E-28 | Regenerative Mesenchyme | mtrex |
| epha7 | 1.3E-32 | 1.1323876 | 0.104 | 0.049 | 4.26E-28 | Regenerative Mesenchyme | epha7 |
| glrx5 | 1.3E-32 | 1.1569046 | 0.104 | 0.049 | 4.35E-28 | Regenerative Mesenchyme | glrx5 |
| krt18b1 | 1.3E-32 | 0.2758942 | 0.922 | 0.89 | 4.35E-28 | Regenerative Mesenchyme | krt18b |
| praf2 | 1.4E-32 | 0.6528949 | 0.222 | 0.134 | 4.6E-28 | Regenerative Mesenchyme | praf2 |
| pitpnaa | 1.5E-32 | 0.9296503 | 0.108 | 0.051 | 4.83E-28 | Regenerative Mesenchyme | pitpnaa |
| srpk1b | 1.5E-32 | 0.8452113 | 0.119 | 0.058 | 4.84E-28 | Regenerative Mesenchyme | srpk1b |
| tmpoa | 1.6E-32 | 1.0114555 | 0.103 | 0.049 | 5.29E-28 | Regenerative Mesenchyme | tmpoa |
| fam3c1 | 1.9E-32 | 0.8508853 | 0.173 | 0.098 | 6.1E-28 | Regenerative Mesenchyme | fam3c |
| tcea1 | 3.2E-32 | 0.6999497 | 0.18 | 0.102 | 1.04E-27 | Regenerative Mesenchyme | tcea1 |
| slc3a2b1 | 3.8E-32 | 0.5941738 | 0.221 | 0.134 | 1.24E-27 | Regenerative Mesenchyme | slc3a2b |
| mvp1 | 4E-32 | 0.3039309 | 0.433 | 0.31 | 1.3E-27 | Regenerative Mesenchyme | mvp |
| puf60b | 4.2E-32 | 0.7567557 | 0.168 | 0.094 | 1.37E-27 | Regenerative Mesenchyme | puf60b |
| cltca | 5.1E-32 | 0.6259368 | 0.29 | 0.192 | 1.66E-27 | Regenerative Mesenchyme | cltca |
| papss2b | 5.6E-32 | 0.7808273 | 0.168 | 0.094 | 1.81E-27 | Regenerative Mesenchyme | papss2b |
| eif2s1b2 | 5.6E-32 | 0.5224906 | 0.489 | 0.382 | 1.81E-27 | Regenerative Mesenchyme | eif2s1b |
| txn2 | 6.5E-32 | 0.7694561 | 0.186 | 0.108 | 2.1E-27 | Regenerative Mesenchyme | txn2 |
| zc3h13 | 9.7E-32 | 0.4093637 | 0.341 | 0.228 | 3.14E-27 | Regenerative Mesenchyme | zc3h13 |
| nrp1a | 1.1E-31 | 0.6165719 | 0.211 | 0.127 | 3.66E-27 | Regenerative Mesenchyme | nrp1a |
| chchd6b | 1.1E-31 | 0.7094525 | 0.161 | 0.088 | 3.72E-27 | Regenerative Mesenchyme | chchd6b |
| atox12 | 1.5E-31 | 0.6386552 | 0.356 | 0.253 | 4.8E-27 | Regenerative Mesenchyme | atox1 |
| zgc:112496 | 1.5E-31 | 0.9081372 | 0.106 | 0.051 | 4.92E-27 | Regenerative Mesenchyme | zgc:112496 |
| uggt1 | 1.7E-31 | 1.0825693 | 0.105 | 0.05 | 5.67E-27 | Regenerative Mesenchyme | uggt1 |
| dek | 1.8E-31 | 0.8715122 | 0.202 | 0.122 | 5.8E-27 | Regenerative Mesenchyme | dek |
| scfd11 | 3E-31 | 0.8228435 | 0.176 | 0.101 | 9.89E-27 | Regenerative Mesenchyme | scfd1 |
| slc7a10a2 | 3.1E-31 | 0.8410099 | 0.179 | 0.104 | 9.93E-27 | Regenerative Mesenchyme | slc7a10a |
| eif4g2b | 3.7E-31 | 0.6930098 | 0.283 | 0.186 | 1.21E-26 | Regenerative Mesenchyme | eif4g2b |
| cnn21 | 3.9E-31 | 0.270836 | 0.563 | 0.439 | 1.28E-26 | Regenerative Mesenchyme | cnn2 |
| srrm2 | 3.9E-31 | 0.4376118 | 0.423 | 0.306 | 1.28E-26 | Regenerative Mesenchyme | srrm2 |
| vps25 | 4.8E-31 | 0.7558219 | 0.164 | 0.092 | 1.57E-26 | Regenerative Mesenchyme | vps25 |
| eif4ba | 4.9E-31 | 0.6340027 | 0.199 | 0.118 | 1.58E-26 | Regenerative Mesenchyme | eif4ba |
| zmat2 | 5.2E-31 | 0.6038555 | 0.311 | 0.21 | 1.7E-26 | Regenerative Mesenchyme | zmat2 |

| | | | | | | | |
|---|---|---|---|---|---|---|---|
| tex261 | 6.1E-31 | 0.7310276 | 0.192 | 0.113 | 1.99E-26 | Regenerative Mesenchyme | tex261 |
| gmfb | 6.9E-31 | 0.7534933 | 0.226 | 0.14 | 2.24E-26 | Regenerative Mesenchyme | gmfb |
| akirin1 | 7.6E-31 | 0.8221143 | 0.166 | 0.094 | 2.48E-26 | Regenerative Mesenchyme | akirin1 |
| tgif11 | 1.5E-30 | 0.5202419 | 0.3 | 0.2 | 4.79E-26 | Regenerative Mesenchyme | tgif1 |
| myo1ea1 | 1.5E-30 | 0.8699792 | 0.145 | 0.079 | 5.02E-26 | Regenerative Mesenchyme | myo1ea |
| BX548028.11 | 2E-30 | 1.4394675 | 0.132 | 0.071 | 6.4E-26 | Regenerative Mesenchyme | BX548028.1 |
| ppp4r3b | 2.4E-30 | 0.8746522 | 0.114 | 0.057 | 7.91E-26 | Regenerative Mesenchyme | ppp4r3b |
| sec61a12 | 2.7E-30 | 0.5550911 | 0.507 | 0.416 | 8.66E-26 | Regenerative Mesenchyme | sec61a1 |
| eif3s6ip2 | 2.9E-30 | 0.6003512 | 0.415 | 0.314 | 9.45E-26 | Regenerative Mesenchyme | eif3s6ip |
| MRPS121 | 3.3E-30 | 0.7725029 | 0.155 | 0.086 | 1.07E-25 | Regenerative Mesenchyme | MRPS12 |
| csnk2a1 | 3.6E-30 | 0.5521982 | 0.358 | 0.251 | 1.16E-25 | Regenerative Mesenchyme | csnk2a1 |
| cct8 | 4.3E-30 | 0.5962065 | 0.37 | 0.266 | 1.41E-25 | Regenerative Mesenchyme | cct8 |
| vegfaa1 | 4.4E-30 | 1.0446402 | 0.186 | 0.113 | 1.42E-25 | Regenerative Mesenchyme | vegfaa |
| psat1 | 5.8E-30 | 1.0986132 | 0.125 | 0.066 | 1.88E-25 | Regenerative Mesenchyme | psat1 |
| metap1 | 6.3E-30 | 0.7907232 | 0.129 | 0.067 | 2.06E-25 | Regenerative Mesenchyme | metap1 |
| srsf41 | 7.7E-30 | 0.6120913 | 0.247 | 0.158 | 2.5E-25 | Regenerative Mesenchyme | srsf4 |
| cnn3a1 | 7.8E-30 | 0.3764996 | 0.581 | 0.471 | 2.53E-25 | Regenerative Mesenchyme | cnn3a |
| dolpp1 | 7.9E-30 | 0.7241943 | 0.167 | 0.095 | 2.58E-25 | Regenerative Mesenchyme | dolpp1 |
| mrgbp | 9.3E-30 | 0.7281518 | 0.149 | 0.082 | 3.03E-25 | Regenerative Mesenchyme | mrgbp |
| PYURF1 | 9.5E-30 | 0.8343073 | 0.129 | 0.068 | 3.08E-25 | Regenerative Mesenchyme | PYURF |
| ehd1a | 1.2E-29 | 0.7578531 | 0.121 | 0.062 | 3.77E-25 | Regenerative Mesenchyme | ehd1a |
| MYADM | 1.3E-29 | 0.728325 | 0.165 | 0.094 | 4.17E-25 | Regenerative Mesenchyme | MYADM |
| plod3 | 1.4E-29 | 0.6989311 | 0.231 | 0.147 | 4.4E-25 | Regenerative Mesenchyme | plod3 |
| sod2 | 1.8E-29 | 0.7424997 | 0.204 | 0.125 | 5.82E-25 | Regenerative Mesenchyme | sod2 |
| sdhaf2 | 2.2E-29 | 0.7137008 | 0.152 | 0.084 | 7.25E-25 | Regenerative Mesenchyme | sdhaf2 |
| znf346 | 2.5E-29 | 0.7173431 | 0.157 | 0.088 | 8.07E-25 | Regenerative Mesenchyme | znf346 |
| zgc:86598 | 2.8E-29 | 0.6565671 | 0.245 | 0.157 | 9.17E-25 | Regenerative Mesenchyme | zgc:86598 |
| abhd16a | 2.9E-29 | 0.845462 | 0.127 | 0.066 | 9.28E-25 | Regenerative Mesenchyme | abhd16a |
| smap1 | 3.4E-29 | 0.5695349 | 0.239 | 0.15 | 1.12E-24 | Regenerative Mesenchyme | smap1 |
| serpinb1l3 | 3.6E-29 | 0.6774326 | 0.153 | 0.085 | 1.17E-24 | Regenerative Mesenchyme | serpinb1l3 |
| ckap41 | 4.1E-29 | 0.4820485 | 0.592 | 0.515 | 1.35E-24 | Regenerative Mesenchyme | ckap4 |
| C2H1orf351 | 5.6E-29 | 0.7297474 | 0.149 | 0.083 | 1.81E-24 | Regenerative Mesenchyme | C2H1orf35 |
| smc1al | 6.7E-29 | 0.5713129 | 0.372 | 0.269 | 2.19E-24 | Regenerative Mesenchyme | smc1al |
| uqcrb | 8.7E-29 | 0.9912435 | 0.118 | 0.061 | 2.84E-24 | Regenerative Mesenchyme | uqcrb |
| txndc15 | 9E-29 | 0.7145684 | 0.178 | 0.105 | 2.94E-24 | Regenerative Mesenchyme | txndc15 |
| mt-nd6 | 1E-28 | 1.0372884 | 0.103 | 0.051 | 3.38E-24 | Regenerative Mesenchyme | mt-nd6 |
| tram21 | 1.1E-28 | 0.7903336 | 0.179 | 0.106 | 3.58E-24 | Regenerative Mesenchyme | tram2 |
| wdr51 | 1.3E-28 | 0.9211278 | 0.123 | 0.065 | 4.27E-24 | Regenerative Mesenchyme | wdr5 |
| cops8 | 1.4E-28 | 0.7230474 | 0.152 | 0.085 | 4.67E-24 | Regenerative Mesenchyme | cops8 |
| chtopa1 | 1.4E-28 | 0.5667871 | 0.215 | 0.132 | 4.69E-24 | Regenerative Mesenchyme | chtopa |
| STEEP1 | 1.8E-28 | 0.7105072 | 0.129 | 0.068 | 5.71E-24 | Regenerative Mesenchyme | STEEP1 |
| zgc:77056 | 2.1E-28 | 0.7307992 | 0.125 | 0.066 | 6.94E-24 | Regenerative Mesenchyme | zgc:77056 |
| brd7 | 2.2E-28 | 0.7272497 | 0.113 | 0.057 | 7.19E-24 | Regenerative Mesenchyme | brd7 |
| BX927308.1 | 2.4E-28 | 0.9961007 | 0.117 | 0.061 | 7.77E-24 | Regenerative Mesenchyme | BX927308.1 |
| smarcb1a | 2.5E-28 | 0.6439564 | 0.182 | 0.107 | 8.13E-24 | Regenerative Mesenchyme | smarcb1a |
| dldh | 2.5E-28 | 0.6588378 | 0.287 | 0.195 | 8.2E-24 | Regenerative Mesenchyme | dldh |
| CABZ01058650.11 | 3.5E-28 | 0.8370089 | 0.13 | 0.07 | 1.15E-23 | Regenerative Mesenchyme | CABZ01058650.1 |
| pak2a | 3.8E-28 | 0.6361796 | 0.233 | 0.148 | 1.22E-23 | Regenerative Mesenchyme | pak2a |

| | | | | | | | |
|---|---|---|---|---|---|---|---|
| slc35e1 | 4.2E-28 | 0.8735555 | 0.108 | 0.054 | 1.38E-23 | Regenerative Mesenchyme | slc35e1 |
| riok1 | 4.3E-28 | 0.8107488 | 0.123 | 0.064 | 1.4E-23 | Regenerative Mesenchyme | riok1 |
| cux1a1 | 4.3E-28 | 0.9194246 | 0.123 | 0.066 | 1.41E-23 | Regenerative Mesenchyme | cux1a |
| spcs2 | 4.7E-28 | 0.4772016 | 0.386 | 0.281 | 1.52E-23 | Regenerative Mesenchyme | spcs2 |
| baxa.1 | 5.3E-28 | 0.7745696 | 0.168 | 0.098 | 1.71E-23 | Regenerative Mesenchyme | baxa.1 |
| zgc:153115 | 5.8E-28 | 0.8222052 | 0.143 | 0.079 | 1.9E-23 | Regenerative Mesenchyme | zgc:153115 |
| tp53 | 6.1E-28 | 0.7352404 | 0.239 | 0.156 | 1.98E-23 | Regenerative Mesenchyme | tp53 |
| si:dkey-23i12.51 | 6.4E-28 | 0.6361464 | 0.307 | 0.214 | 2.08E-23 | Regenerative Mesenchyme | si:dkey-23i12.5 |
| rad23b1 | 7.4E-28 | 0.7953776 | 0.163 | 0.095 | 2.4E-23 | Regenerative Mesenchyme | rad23b |
| tfpia3 | 7.5E-28 | 0.7921467 | 0.256 | 0.175 | 2.43E-23 | Regenerative Mesenchyme | tfpia |
| psmg21 | 8.4E-28 | 0.7715325 | 0.173 | 0.102 | 2.74E-23 | Regenerative Mesenchyme | psmg2 |
| ccdc47 | 9.1E-28 | 0.6850908 | 0.215 | 0.135 | 2.97E-23 | Regenerative Mesenchyme | ccdc47 |
| fus | 1.1E-27 | 0.7401907 | 0.108 | 0.054 | 3.43E-23 | Regenerative Mesenchyme | fus |
| ctsk2 | 1.1E-27 | 0.3220813 | 0.677 | 0.588 | 3.45E-23 | Regenerative Mesenchyme | ctsk |
| aif1l1 | 1.1E-27 | 0.6911626 | 0.148 | 0.083 | 3.74E-23 | Regenerative Mesenchyme | aif1l |
| rab12 | 1.2E-27 | 0.7369644 | 0.124 | 0.066 | 3.84E-23 | Regenerative Mesenchyme | rab12 |
| emc2 | 1.4E-27 | 0.5807785 | 0.291 | 0.198 | 4.54E-23 | Regenerative Mesenchyme | emc2 |
| gcat | 1.6E-27 | 0.6693595 | 0.135 | 0.074 | 5.22E-23 | Regenerative Mesenchyme | gcat |
| timeless | 1.9E-27 | 0.7885664 | 0.108 | 0.055 | 6.06E-23 | Regenerative Mesenchyme | timeless |
| lgals3a | 2.3E-27 | 0.4903142 | 0.375 | 0.272 | 7.64E-23 | Regenerative Mesenchyme | lgals3a |
| ddx46 | 2.5E-27 | 0.7982383 | 0.146 | 0.082 | 8.29E-23 | Regenerative Mesenchyme | ddx46 |
| idh3g | 2.6E-27 | 0.960128 | 0.112 | 0.058 | 8.38E-23 | Regenerative Mesenchyme | idh3g |
| mfap1 | 3.1E-27 | 0.5591003 | 0.215 | 0.134 | 1.02E-22 | Regenerative Mesenchyme | mfap1 |
| sptssa | 3.3E-27 | 0.6779554 | 0.185 | 0.111 | 1.07E-22 | Regenerative Mesenchyme | sptssa |
| rcc2 | 3.6E-27 | 0.9262977 | 0.131 | 0.072 | 1.17E-22 | Regenerative Mesenchyme | rcc2 |
| zgc:77486.11 | 4E-27 | 0.6344797 | 0.25 | 0.165 | 1.3E-22 | Regenerative Mesenchyme | zgc:77486.1 |
| rif11 | 4.1E-27 | 0.7434244 | 0.146 | 0.083 | 1.32E-22 | Regenerative Mesenchyme | rif1 |
| med28 | 5.3E-27 | 0.6954817 | 0.137 | 0.076 | 1.72E-22 | Regenerative Mesenchyme | med28 |
| lhfpl2a1 | 6E-27 | 0.55536 | 0.175 | 0.104 | 1.94E-22 | Regenerative Mesenchyme | lhfpl2a |
| polr2j | 6.5E-27 | 0.7181313 | 0.16 | 0.093 | 2.11E-22 | Regenerative Mesenchyme | polr2j |
| fam76b | 7.3E-27 | 0.8174221 | 0.145 | 0.082 | 2.36E-22 | Regenerative Mesenchyme | fam76b |
| sec63 | 8.4E-27 | 0.5582916 | 0.307 | 0.213 | 2.72E-22 | Regenerative Mesenchyme | sec63 |
| rhoac | 9.6E-27 | 0.4934296 | 0.394 | 0.29 | 3.11E-22 | Regenerative Mesenchyme | rhoac |
| kpna4 | 9.7E-27 | 0.5496589 | 0.221 | 0.14 | 3.14E-22 | Regenerative Mesenchyme | kpna4 |
| C11H1orf50 | 1E-26 | 0.8624243 | 0.101 | 0.051 | 3.27E-22 | Regenerative Mesenchyme | C11H1orf50 |
| naa15a | 1E-26 | 0.6567375 | 0.177 | 0.105 | 3.33E-22 | Regenerative Mesenchyme | naa15a |
| selenon2 | 1E-26 | 0.5723062 | 0.235 | 0.153 | 3.41E-22 | Regenerative Mesenchyme | selenon |
| cand1 | 1.1E-26 | 0.5539994 | 0.185 | 0.111 | 3.52E-22 | Regenerative Mesenchyme | cand1 |
| hnrnpr | 1.1E-26 | 0.6629508 | 0.191 | 0.117 | 3.53E-22 | Regenerative Mesenchyme | hnrnpr |
| ilk | 1.1E-26 | 0.6456742 | 0.263 | 0.176 | 3.59E-22 | Regenerative Mesenchyme | ilk |
| sart1 | 1.2E-26 | 0.5455176 | 0.249 | 0.163 | 3.98E-22 | Regenerative Mesenchyme | sart1 |
| rad21a | 1.5E-26 | 0.5945073 | 0.287 | 0.197 | 5.01E-22 | Regenerative Mesenchyme | rad21a |
| cnot6a | 1.7E-26 | 0.7450846 | 0.155 | 0.089 | 5.68E-22 | Regenerative Mesenchyme | cnot6a |
| strap | 1.8E-26 | 0.8696141 | 0.151 | 0.087 | 5.83E-22 | Regenerative Mesenchyme | strap |
| prpf39 | 2.1E-26 | 0.5513024 | 0.204 | 0.126 | 6.7E-22 | Regenerative Mesenchyme | prpf39 |
| ralba | 2.1E-26 | 0.7028315 | 0.152 | 0.087 | 6.82E-22 | Regenerative Mesenchyme | ralba |
| mdh2 | 2.2E-26 | 0.4779001 | 0.528 | 0.435 | 7.09E-22 | Regenerative Mesenchyme | mdh2 |
| cyb5r1 | 3.9E-26 | 0.5443612 | 0.287 | 0.197 | 1.26E-21 | Regenerative Mesenchyme | cyb5r1 |

| | | | | | | | |
|---|---|---|---|---|---|---|---|
| dnajc3b | 4.2E-26 | 0.8490141 | 0.112 | 0.059 | 1.35E-21 | Regenerative Mesenchyme | dnajc3b |
| slc38a2 | 5.7E-26 | 0.8520033 | 0.136 | 0.077 | 1.85E-21 | Regenerative Mesenchyme | slc38a2 |
| calub | 6.2E-26 | 0.9819517 | 0.118 | 0.064 | 2.01E-21 | Regenerative Mesenchyme | calub |
| rpn2 | 6.2E-26 | 0.7543164 | 0.116 | 0.062 | 2.01E-21 | Regenerative Mesenchyme | rpn2 |
| eif4ebp11 | 6.4E-26 | 0.6164772 | 0.229 | 0.149 | 2.08E-21 | Regenerative Mesenchyme | eif4ebp1 |
| myl12.21 | 7E-26 | 0.6626833 | 0.246 | 0.164 | 2.29E-21 | Regenerative Mesenchyme | myl12.2 |
| mrpl411 | 7.5E-26 | 0.7144819 | 0.143 | 0.082 | 2.43E-21 | Regenerative Mesenchyme | mrpl41 |
| smarcd1 | 7.6E-26 | 0.9446034 | 0.121 | 0.066 | 2.46E-21 | Regenerative Mesenchyme | smarcd1 |
| cops2 | 7.7E-26 | 0.7106898 | 0.191 | 0.118 | 2.49E-21 | Regenerative Mesenchyme | cops2 |
| slc38a5a1 | 1.1E-25 | 0.7197521 | 0.124 | 0.068 | 3.5E-21 | Regenerative Mesenchyme | slc38a5a |
| snrnp271 | 1.1E-25 | 0.66343 | 0.211 | 0.134 | 3.52E-21 | Regenerative Mesenchyme | snrnp27 |
| prpf6 | 1.1E-25 | 0.6618442 | 0.156 | 0.091 | 3.55E-21 | Regenerative Mesenchyme | prpf6 |
| smg1 | 1.1E-25 | 0.7552635 | 0.12 | 0.064 | 3.64E-21 | Regenerative Mesenchyme | smg1 |
| anp32a | 1.6E-25 | 0.5056216 | 0.449 | 0.351 | 5.32E-21 | Regenerative Mesenchyme | anp32a |
| hist1h4l.181 | 1.6E-25 | 1.5684347 | 0.182 | 0.118 | 5.34E-21 | Regenerative Mesenchyme | hist1h4l.18 |
| p3h2 | 2E-25 | 0.8346424 | 0.113 | 0.06 | 6.36E-21 | Regenerative Mesenchyme | p3h2 |
| med21 | 2E-25 | 0.7560722 | 0.125 | 0.068 | 6.46E-21 | Regenerative Mesenchyme | med21 |
| pet100 | 2.6E-25 | 0.6714144 | 0.142 | 0.081 | 8.52E-21 | Regenerative Mesenchyme | pet100 |
| max | 2.8E-25 | 0.4924872 | 0.386 | 0.285 | 9.21E-21 | Regenerative Mesenchyme | max |
| atp6v0cb | 3.1E-25 | 0.7968421 | 0.141 | 0.08 | 9.99E-21 | Regenerative Mesenchyme | atp6v0cb |
| selenot1a | 3.8E-25 | 0.591643 | 0.248 | 0.165 | 1.23E-20 | Regenerative Mesenchyme | selenot1a |
| kif5ba | 3.8E-25 | 0.7373258 | 0.159 | 0.094 | 1.23E-20 | Regenerative Mesenchyme | kif5ba |
| tmem189 | 3.8E-25 | 0.5530678 | 0.203 | 0.128 | 1.24E-20 | Regenerative Mesenchyme | tmem189 |
| cfap20 | 4.3E-25 | 0.8093496 | 0.125 | 0.069 | 1.41E-20 | Regenerative Mesenchyme | cfap20 |
| bnc21 | 7.7E-25 | 0.6293583 | 0.139 | 0.079 | 2.52E-20 | Regenerative Mesenchyme | bnc2 |
| zmp:0000000735 | 7.9E-25 | 0.5501016 | 0.109 | 0.057 | 2.57E-20 | Regenerative Mesenchyme | zmp:0000000735 |
| rcan1a1 | 8.9E-25 | 0.5428821 | 0.314 | 0.226 | 2.9E-20 | Regenerative Mesenchyme | rcan1a |
| naa15b | 9.1E-25 | 0.6190264 | 0.16 | 0.095 | 2.97E-20 | Regenerative Mesenchyme | naa15b |
| irf2a | 1.1E-24 | 0.8044901 | 0.141 | 0.082 | 3.6E-20 | Regenerative Mesenchyme | irf2a |
| tm9sf4 | 1.1E-24 | 0.78241 | 0.129 | 0.072 | 3.63E-20 | Regenerative Mesenchyme | tm9sf4 |
| dhx16 | 1.3E-24 | 0.4466549 | 0.175 | 0.105 | 4.18E-20 | Regenerative Mesenchyme | dhx16 |
| lamtor3 | 1.4E-24 | 0.6782318 | 0.137 | 0.078 | 4.54E-20 | Regenerative Mesenchyme | lamtor3 |
| srpra | 1.7E-24 | 0.5357716 | 0.305 | 0.215 | 5.48E-20 | Regenerative Mesenchyme | srpra |
| atp1a1a.11 | 2.3E-24 | 0.3974559 | 0.418 | 0.318 | 7.43E-20 | Regenerative Mesenchyme | atp1a1a.1 |
| fam3a | 2.3E-24 | 0.6066104 | 0.213 | 0.137 | 7.58E-20 | Regenerative Mesenchyme | fam3a |
| yif1a | 2.4E-24 | 0.7307196 | 0.192 | 0.121 | 7.91E-20 | Regenerative Mesenchyme | yif1a |
| idh3b | 2.9E-24 | 0.805791 | 0.127 | 0.072 | 9.43E-20 | Regenerative Mesenchyme | idh3b |
| zranb21 | 3.1E-24 | 0.461366 | 0.201 | 0.126 | 9.92E-20 | Regenerative Mesenchyme | zranb2 |
| ptgesl | 3.6E-24 | 0.8142876 | 0.113 | 0.061 | 1.19E-19 | Regenerative Mesenchyme | ptgesl |
| kmt5aa | 4.3E-24 | 0.6993457 | 0.179 | 0.111 | 1.41E-19 | Regenerative Mesenchyme | kmt5aa |
| map1ab | 5.2E-24 | 0.6000805 | 0.272 | 0.193 | 1.69E-19 | Regenerative Mesenchyme | map1ab |
| pitpnbl | 6E-24 | 0.7365155 | 0.18 | 0.112 | 1.95E-19 | Regenerative Mesenchyme | pitpnbl |
| trir | 6.4E-24 | 0.7296574 | 0.126 | 0.07 | 2.09E-19 | Regenerative Mesenchyme | trir |
| ap1b1 | 6.6E-24 | 0.7262894 | 0.101 | 0.052 | 2.14E-19 | Regenerative Mesenchyme | ap1b1 |
| ssu72 | 8E-24 | 0.6360271 | 0.154 | 0.092 | 2.62E-19 | Regenerative Mesenchyme | ssu72 |
| impdh1b2 | 1.2E-23 | 0.8078868 | 0.138 | 0.08 | 3.77E-19 | Regenerative Mesenchyme | impdh1b |
| tmsb11 | 1.2E-23 | 0.7716333 | 0.538 | 0.445 | 3.87E-19 | Regenerative Mesenchyme | tmsb1 |
| dync1li2 | 1.4E-23 | 0.5806055 | 0.24 | 0.161 | 4.61E-19 | Regenerative Mesenchyme | dync1li2 |

| | | | | | | | |
|---|---|---|---|---|---|---|---|
| pnn1 | 1.4E-23 | 0.3830523 | 0.328 | 0.231 | 4.66E-19 | Regenerative Mesenchyme | pnn |
| actr1 | 1.4E-23 | 0.6301983 | 0.216 | 0.141 | 4.68E-19 | Regenerative Mesenchyme | actr1 |
| emd | 1.7E-23 | 0.4532862 | 0.453 | 0.357 | 5.42E-19 | Regenerative Mesenchyme | emd |
| larp7 | 1.9E-23 | 0.6795792 | 0.137 | 0.079 | 6.04E-19 | Regenerative Mesenchyme | larp7 |
| exosc31 | 1.9E-23 | 0.7259954 | 0.124 | 0.07 | 6.2E-19 | Regenerative Mesenchyme | exosc3 |
| itga52 | 2E-23 | 0.3643674 | 0.35 | 0.252 | 6.53E-19 | Regenerative Mesenchyme | itga5 |
| tmem126a1 | 2.9E-23 | 0.879876 | 0.114 | 0.063 | 9.53E-19 | Regenerative Mesenchyme | tmem126a |
| si:dkey-19e4.5 | 3E-23 | 0.6100408 | 0.179 | 0.112 | 9.64E-19 | Regenerative Mesenchyme | si:dkey-19e4.5 |
| szrd1 | 3E-23 | 0.5250197 | 0.235 | 0.155 | 9.84E-19 | Regenerative Mesenchyme | szrd1 |
| zc3h18 | 3.8E-23 | 0.4624446 | 0.211 | 0.135 | 1.23E-18 | Regenerative Mesenchyme | zc3h18 |
| brk1 | 4.5E-23 | 0.5546907 | 0.283 | 0.2 | 1.45E-18 | Regenerative Mesenchyme | brk1 |
| gatd3a | 4.6E-23 | 0.7074387 | 0.108 | 0.058 | 1.51E-18 | Regenerative Mesenchyme | gatd3a |
| yipf5 | 5E-23 | 0.5967776 | 0.189 | 0.12 | 1.63E-18 | Regenerative Mesenchyme | yipf5 |
| tmem11 | 5.4E-23 | 0.6128755 | 0.164 | 0.1 | 1.75E-18 | Regenerative Mesenchyme | tmem11 |
| fkbp141 | 5.4E-23 | 0.5040131 | 0.42 | 0.327 | 1.77E-18 | Regenerative Mesenchyme | fkbp14 |
| ltv1 | 5.6E-23 | 0.7146355 | 0.124 | 0.07 | 1.82E-18 | Regenerative Mesenchyme | ltv1 |
| ciao2b | 6.9E-23 | 0.6717631 | 0.2 | 0.129 | 2.24E-18 | Regenerative Mesenchyme | ciao2b |
| sdf41 | 7.2E-23 | 0.3751439 | 0.398 | 0.299 | 2.34E-18 | Regenerative Mesenchyme | sdf4 |
| stx5a | 8.1E-23 | 0.7026921 | 0.157 | 0.096 | 2.63E-18 | Regenerative Mesenchyme | stx5a |
| naa38 | 8.2E-23 | 0.8073767 | 0.112 | 0.062 | 2.67E-18 | Regenerative Mesenchyme | naa38 |
| cpne2 | 8.2E-23 | 0.7125473 | 0.116 | 0.064 | 2.68E-18 | Regenerative Mesenchyme | cpne2 |
| CABZ01064941.1 | 8.8E-23 | 0.8102245 | 0.139 | 0.083 | 2.85E-18 | Regenerative Mesenchyme | CABZ01064941.1 |
| dync1i2a | 9.5E-23 | 0.6721853 | 0.176 | 0.11 | 3.09E-18 | Regenerative Mesenchyme | dync1i2a |
| wu:fj16a032 | 9.9E-23 | 0.2829408 | 0.272 | 0.188 | 3.23E-18 | Regenerative Mesenchyme | wu:fj16a03 |
| scamp4 | 1E-22 | 0.7651164 | 0.13 | 0.075 | 3.39E-18 | Regenerative Mesenchyme | scamp4 |
| pin1 | 1.1E-22 | 0.5048638 | 0.348 | 0.258 | 3.61E-18 | Regenerative Mesenchyme | pin1 |
| commd8 | 1.2E-22 | 0.7608847 | 0.124 | 0.07 | 3.79E-18 | Regenerative Mesenchyme | commd8 |
| pcyt2 | 1.2E-22 | 0.7435963 | 0.154 | 0.093 | 3.89E-18 | Regenerative Mesenchyme | pcyt2 |
| pdap1b1 | 1.2E-22 | 0.650099 | 0.153 | 0.092 | 3.93E-18 | Regenerative Mesenchyme | pdap1b |
| MYO9B1 | 1.2E-22 | 0.5779122 | 0.189 | 0.12 | 3.97E-18 | Regenerative Mesenchyme | MYO9B |
| ms4a17a.82 | 1.3E-22 | 0.5655993 | 0.177 | 0.11 | 4.08E-18 | Regenerative Mesenchyme | ms4a17a.8 |
| fam133b | 1.5E-22 | 0.4233573 | 0.338 | 0.247 | 4.78E-18 | Regenerative Mesenchyme | fam133b |
| pak1 | 1.6E-22 | 0.6534845 | 0.118 | 0.066 | 5.05E-18 | Regenerative Mesenchyme | pak1 |
| arl2 | 1.7E-22 | 0.5915158 | 0.128 | 0.073 | 5.51E-18 | Regenerative Mesenchyme | arl2 |
| aip | 1.9E-22 | 0.5193928 | 0.28 | 0.197 | 6.2E-18 | Regenerative Mesenchyme | aip |
| cog4 | 2.2E-22 | 0.8758169 | 0.101 | 0.054 | 7.25E-18 | Regenerative Mesenchyme | cog4 |
| si:dkey-33c12.4 | 2.6E-22 | 0.3824487 | 0.437 | 0.337 | 8.36E-18 | Regenerative Mesenchyme | si:dkey-33c12.4 |
| pelp1 | 3E-22 | 0.7477329 | 0.127 | 0.073 | 9.85E-18 | Regenerative Mesenchyme | pelp1 |
| rab11a | 3.2E-22 | 0.5364677 | 0.28 | 0.196 | 1.04E-17 | Regenerative Mesenchyme | rab11a |
| med10 | 3.3E-22 | 0.6425253 | 0.141 | 0.083 | 1.08E-17 | Regenerative Mesenchyme | med10 |
| c1galt1c1 | 3.9E-22 | 0.7411106 | 0.119 | 0.067 | 1.27E-17 | Regenerative Mesenchyme | c1galt1c1 |
| fxr2 | 4.4E-22 | 0.6521373 | 0.148 | 0.089 | 1.44E-17 | Regenerative Mesenchyme | fxr2 |
| akirin2 | 5.2E-22 | 1.015223 | 0.118 | 0.067 | 1.68E-17 | Regenerative Mesenchyme | akirin2 |
| selenos | 6.5E-22 | 0.494225 | 0.234 | 0.157 | 2.12E-17 | Regenerative Mesenchyme | selenos |
| ube2g1b | 8.1E-22 | 0.6305403 | 0.153 | 0.093 | 2.64E-17 | Regenerative Mesenchyme | ube2g1b |
| copz21 | 8.1E-22 | 0.5493993 | 0.263 | 0.184 | 2.64E-17 | Regenerative Mesenchyme | copz2 |
| hivep3a1 | 8.7E-22 | 0.6544033 | 0.169 | 0.106 | 2.82E-17 | Regenerative Mesenchyme | hivep3a |
| eif4ebp2 | 9.8E-22 | 0.560034 | 0.296 | 0.212 | 3.17E-17 | Regenerative Mesenchyme | eif4ebp2 |

| | | | | | | | |
|---|---|---|---|---|---|---|---|
| pcnt | 9.8E-22 | 0.9250986 | 0.111 | 0.062 | 3.19E-17 | Regenerative Mesenchyme | pcnt |
| rab2a | 1.1E-21 | 0.346918 | 0.606 | 0.517 | 3.63E-17 | Regenerative Mesenchyme | rab2a |
| akap8l | 1.1E-21 | 0.4649366 | 0.276 | 0.192 | 3.64E-17 | Regenerative Mesenchyme | akap8l |
| sra11 | 1.1E-21 | 0.765132 | 0.111 | 0.062 | 3.7E-17 | Regenerative Mesenchyme | sra1 |
| mapk61 | 1.1E-21 | 0.7212651 | 0.142 | 0.085 | 3.73E-17 | Regenerative Mesenchyme | mapk6 |
| fnta | 1.3E-21 | 0.7043834 | 0.117 | 0.066 | 4.26E-17 | Regenerative Mesenchyme | fnta |
| rab10 | 1.5E-21 | 0.3484695 | 0.542 | 0.453 | 4.77E-17 | Regenerative Mesenchyme | rab10 |
| sh3kbp1 | 1.6E-21 | 0.509172 | 0.199 | 0.129 | 5.2E-17 | Regenerative Mesenchyme | sh3kbp1 |
| bud231 | 1.6E-21 | 0.8065962 | 0.106 | 0.058 | 5.28E-17 | Regenerative Mesenchyme | bud23 |
| csrp1a2 | 1.8E-21 | 0.4311067 | 0.341 | 0.252 | 5.78E-17 | Regenerative Mesenchyme | csrp1a |
| G3BP2 | 2E-21 | 0.6514361 | 0.171 | 0.108 | 6.61E-17 | Regenerative Mesenchyme | G3BP2 |
| atp6v1ba | 2.1E-21 | 0.9279624 | 0.101 | 0.055 | 6.84E-17 | Regenerative Mesenchyme | atp6v1ba |
| rcor1 | 2.4E-21 | 0.5873358 | 0.2 | 0.13 | 7.66E-17 | Regenerative Mesenchyme | rcor1 |
| srp68 | 2.7E-21 | 0.6068969 | 0.148 | 0.09 | 8.71E-17 | Regenerative Mesenchyme | srp68 |
| rhebl1 | 3.2E-21 | 0.8517391 | 0.11 | 0.062 | 1.04E-16 | Regenerative Mesenchyme | rhebl1 |
| sart3 | 3.7E-21 | 0.6911293 | 0.142 | 0.086 | 1.21E-16 | Regenerative Mesenchyme | sart3 |
| tmtc3 | 3.8E-21 | 0.6636651 | 0.125 | 0.072 | 1.23E-16 | Regenerative Mesenchyme | tmtc3 |
| ctsd | 3.9E-21 | 0.6476199 | 0.222 | 0.151 | 1.26E-16 | Regenerative Mesenchyme | ctsd |
| sept7b1 | 4.3E-21 | 0.510462 | 0.275 | 0.194 | 1.41E-16 | Regenerative Mesenchyme | sept7b |
| ube2l3a | 4.4E-21 | 0.5907887 | 0.22 | 0.148 | 1.45E-16 | Regenerative Mesenchyme | ube2l3a |
| zgc:921612 | 5E-21 | 0.577177 | 0.266 | 0.19 | 1.62E-16 | Regenerative Mesenchyme | zgc:92161 |
| sf3a1 | 5.4E-21 | 0.66246 | 0.148 | 0.09 | 1.75E-16 | Regenerative Mesenchyme | sf3a1 |
| dnajc7 | 5.9E-21 | 0.4472414 | 0.276 | 0.194 | 1.91E-16 | Regenerative Mesenchyme | dnajc7 |
| supt16h1 | 6.3E-21 | 0.5445337 | 0.211 | 0.14 | 2.05E-16 | Regenerative Mesenchyme | supt16h |
| commd7 | 8.7E-21 | 0.7579117 | 0.137 | 0.083 | 2.81E-16 | Regenerative Mesenchyme | commd7 |
| pak2b | 9.8E-21 | 0.5234354 | 0.217 | 0.145 | 3.17E-16 | Regenerative Mesenchyme | pak2b |
| ap1g1 | 1.4E-20 | 0.5152832 | 0.145 | 0.088 | 4.4E-16 | Regenerative Mesenchyme | ap1g1 |
| irf21 | 1.4E-20 | 0.4033482 | 0.219 | 0.146 | 4.67E-16 | Regenerative Mesenchyme | irf2 |
| pdcd10a | 1.5E-20 | 0.5899358 | 0.151 | 0.093 | 4.75E-16 | Regenerative Mesenchyme | pdcd10a |
| trappc1 | 1.7E-20 | 0.5527197 | 0.213 | 0.142 | 5.45E-16 | Regenerative Mesenchyme | trappc1 |
| sumo2b | 2E-20 | 0.3895222 | 0.481 | 0.389 | 6.59E-16 | Regenerative Mesenchyme | sumo2b |
| rab5c | 2.1E-20 | 0.4657729 | 0.313 | 0.226 | 6.92E-16 | Regenerative Mesenchyme | rab5c |
| adam8b1 | 2.3E-20 | 0.632359 | 0.13 | 0.077 | 7.41E-16 | Regenerative Mesenchyme | adam8b |
| clic1 | 2.4E-20 | 0.5670682 | 0.163 | 0.102 | 7.69E-16 | Regenerative Mesenchyme | clic1 |
| mmgt1 | 2.4E-20 | 0.5497369 | 0.178 | 0.114 | 7.7E-16 | Regenerative Mesenchyme | mmgt1 |
| rap1ab | 2.4E-20 | 0.5740076 | 0.115 | 0.066 | 7.7E-16 | Regenerative Mesenchyme | rap1ab |
| atp1b2a1 | 2.4E-20 | 0.5823006 | 0.133 | 0.079 | 7.89E-16 | Regenerative Mesenchyme | atp1b2a |
| usp4 | 2.7E-20 | 0.5833284 | 0.151 | 0.093 | 8.81E-16 | Regenerative Mesenchyme | usp4 |
| rpia | 2.8E-20 | 0.5355645 | 0.153 | 0.094 | 9.12E-16 | Regenerative Mesenchyme | rpia |
| fkbp10b1 | 3.2E-20 | 0.5581381 | 0.33 | 0.251 | 1.03E-15 | Regenerative Mesenchyme | fkbp10b |
| rgmb1 | 3.4E-20 | 0.5630972 | 0.18 | 0.116 | 1.1E-15 | Regenerative Mesenchyme | rgmb |
| ppp4cb | 3.8E-20 | 0.6979579 | 0.138 | 0.084 | 1.24E-15 | Regenerative Mesenchyme | ppp4cb |
| drap1 | 4.7E-20 | 0.4876319 | 0.195 | 0.128 | 1.52E-15 | Regenerative Mesenchyme | drap1 |
| tmem2081 | 4.7E-20 | 0.704668 | 0.111 | 0.063 | 1.53E-15 | Regenerative Mesenchyme | tmem208 |
| zgc:565251 | 4.7E-20 | 0.627935 | 0.22 | 0.151 | 1.54E-15 | Regenerative Mesenchyme | zgc:56525 |
| pds5a | 5E-20 | 0.5770086 | 0.147 | 0.09 | 1.62E-15 | Regenerative Mesenchyme | pds5a |
| scyl1 | 5.2E-20 | 0.6235777 | 0.127 | 0.075 | 1.69E-15 | Regenerative Mesenchyme | scyl1 |
| gorab | 6.4E-20 | 0.6967303 | 0.117 | 0.068 | 2.09E-15 | Regenerative Mesenchyme | gorab |

| | | | | | | | |
|---|---|---|---|---|---|---|---|
| cpne1 | 7.8E-20 | 0.3825638 | 0.551 | 0.466 | 2.55E-15 | Regenerative Mesenchyme | cpne1 |
| eci2 | 7.9E-20 | 0.7563351 | 0.109 | 0.062 | 2.57E-15 | Regenerative Mesenchyme | eci2 |
| hcfc1b | 7.9E-20 | 0.5615401 | 0.185 | 0.121 | 2.57E-15 | Regenerative Mesenchyme | hcfc1b |
| opa1 | 8E-20 | 0.5850113 | 0.128 | 0.076 | 2.6E-15 | Regenerative Mesenchyme | opa1 |
| si:ch211-107o10.31 | 8.1E-20 | 0.509341 | 0.149 | 0.091 | 2.62E-15 | Regenerative Mesenchyme | si:ch211-107o10.3 |
| ap3m1 | 9E-20 | 0.6850515 | 0.136 | 0.082 | 2.94E-15 | Regenerative Mesenchyme | ap3m1 |
| MED9 | 9.5E-20 | 0.6326048 | 0.118 | 0.068 | 3.09E-15 | Regenerative Mesenchyme | MED9 |
| eif2b21 | 1E-19 | 0.7230714 | 0.117 | 0.068 | 3.31E-15 | Regenerative Mesenchyme | eif2b2 |
| get3 | 1.2E-19 | 0.7463838 | 0.101 | 0.056 | 3.9E-15 | Regenerative Mesenchyme | get3 |
| rab6a | 1.3E-19 | 0.583605 | 0.254 | 0.181 | 4.07E-15 | Regenerative Mesenchyme | rab6a |
| lsp1b3 | 1.4E-19 | 0.2682956 | 0.347 | 0.253 | 4.58E-15 | Regenerative Mesenchyme | lsp1b |
| cops91 | 1.5E-19 | 0.4714981 | 0.221 | 0.15 | 4.9E-15 | Regenerative Mesenchyme | cops9 |
| ccnt2b | 1.5E-19 | 0.6052533 | 0.157 | 0.099 | 4.91E-15 | Regenerative Mesenchyme | ccnt2b |
| gdi1 | 2.2E-19 | 0.648903 | 0.16 | 0.102 | 7.23E-15 | Regenerative Mesenchyme | gdi1 |
| il4r.1 | 2.6E-19 | 0.4326363 | 0.22 | 0.149 | 8.3E-15 | Regenerative Mesenchyme | il4r.1 |
| actn13 | 2.9E-19 | 0.2416991 | 0.408 | 0.313 | 9.27E-15 | Regenerative Mesenchyme | actn1 |
| zgc:63587 | 2.9E-19 | 0.7058396 | 0.135 | 0.082 | 9.49E-15 | Regenerative Mesenchyme | zgc:63587 |
| tagln3b | 3.1E-19 | 0.6915886 | 0.114 | 0.066 | 9.94E-15 | Regenerative Mesenchyme | tagln3b |
| rnf7 | 3.8E-19 | 0.4370285 | 0.285 | 0.205 | 1.23E-14 | Regenerative Mesenchyme | rnf7 |
| slc25a20 | 3.9E-19 | 0.5864205 | 0.115 | 0.067 | 1.28E-14 | Regenerative Mesenchyme | slc25a20 |
| ctnnb22 | 4E-19 | 0.3457004 | 0.4 | 0.308 | 1.31E-14 | Regenerative Mesenchyme | ctnnb2 |
| txndc12 | 5.2E-19 | 0.7262869 | 0.111 | 0.064 | 1.7E-14 | Regenerative Mesenchyme | txndc12 |
| mbd2 | 5.4E-19 | 0.7925695 | 0.131 | 0.08 | 1.75E-14 | Regenerative Mesenchyme | mbd2 |
| sptlc2b1 | 8E-19 | 0.5616467 | 0.177 | 0.116 | 2.59E-14 | Regenerative Mesenchyme | sptlc2b |
| si:ch211-207i1.2 | 8.5E-19 | 0.9868613 | 0.13 | 0.08 | 2.77E-14 | Regenerative Mesenchyme | si:ch211-207i1.2 |
| yaf21 | 1.3E-18 | 0.6364771 | 0.165 | 0.107 | 4.26E-14 | Regenerative Mesenchyme | yaf2 |
| prkrip1 | 1.3E-18 | 0.6243681 | 0.113 | 0.066 | 4.32E-14 | Regenerative Mesenchyme | prkrip1 |
| pacsin2 | 1.6E-18 | 0.6062571 | 0.107 | 0.061 | 5.17E-14 | Regenerative Mesenchyme | pacsin2 |
| acsl4a1 | 1.6E-18 | 0.6564569 | 0.125 | 0.076 | 5.21E-14 | Regenerative Mesenchyme | acsl4a |
| ptprk | 1.7E-18 | 0.4405998 | 0.115 | 0.067 | 5.5E-14 | Regenerative Mesenchyme | ptprk |
| dpm2 | 1.8E-18 | 0.4508066 | 0.202 | 0.136 | 5.91E-14 | Regenerative Mesenchyme | dpm2 |
| cnep1r1 | 1.9E-18 | 0.5701165 | 0.155 | 0.099 | 6.06E-14 | Regenerative Mesenchyme | cnep1r1 |
| fam114a21 | 1.9E-18 | 0.5545244 | 0.172 | 0.112 | 6.1E-14 | Regenerative Mesenchyme | fam114a2 |
| si:ch211-276i12.11 | 2.1E-18 | 0.6268225 | 0.158 | 0.101 | 6.78E-14 | Regenerative Mesenchyme | si:ch211-276i12.11 |
| ube2f | 3.4E-18 | 0.6032159 | 0.11 | 0.064 | 1.11E-13 | Regenerative Mesenchyme | ube2f |
| tusc2b | 4E-18 | 0.6649008 | 0.125 | 0.076 | 1.3E-13 | Regenerative Mesenchyme | tusc2b |
| seh1l | 4.3E-18 | 0.6635527 | 0.112 | 0.066 | 1.39E-13 | Regenerative Mesenchyme | seh1l |
| ralbp1 | 4.4E-18 | 0.349878 | 0.229 | 0.157 | 1.44E-13 | Regenerative Mesenchyme | ralbp1 |
| srsf7a | 4.9E-18 | 0.4554067 | 0.255 | 0.182 | 1.58E-13 | Regenerative Mesenchyme | srsf7a |
| adam17a | 5.1E-18 | 0.5298878 | 0.151 | 0.096 | 1.64E-13 | Regenerative Mesenchyme | adam17a |
| abi1a | 5.6E-18 | 0.8190391 | 0.111 | 0.066 | 1.81E-13 | Regenerative Mesenchyme | abi1a |
| prpf4bb | 5.7E-18 | 0.5913234 | 0.108 | 0.063 | 1.85E-13 | Regenerative Mesenchyme | prpf4bb |
| gtf2a1 | 5.8E-18 | 0.5567993 | 0.131 | 0.08 | 1.9E-13 | Regenerative Mesenchyme | gtf2a1 |
| sept61 | 6E-18 | 0.605782 | 0.208 | 0.144 | 1.95E-13 | Regenerative Mesenchyme | sept6 |
| cherp | 6.1E-18 | 0.5564976 | 0.106 | 0.061 | 1.98E-13 | Regenerative Mesenchyme | cherp |
| camk2n1a2 | 6.8E-18 | 0.3981575 | 0.132 | 0.081 | 2.2E-13 | Regenerative Mesenchyme | camk2n1a |
| nufip2 | 7E-18 | 0.6297375 | 0.21 | 0.145 | 2.28E-13 | Regenerative Mesenchyme | nufip2 |
| chid12 | 7E-18 | 0.6449385 | 0.134 | 0.083 | 2.29E-13 | Regenerative Mesenchyme | chid1 |

| | | | | | | | |
|---|---|---|---|---|---|---|---|
| ctr9 | 7.2E-18 | 0.6567534 | 0.127 | 0.078 | 2.35E-13 | Regenerative Mesenchyme | ctr9 |
| prpf3 | 7.6E-18 | 0.502667 | 0.121 | 0.072 | 2.46E-13 | Regenerative Mesenchyme | prpf3 |
| mmp91 | 1E-17 | 0.2076453 | 0.12 | 0.072 | 3.34E-13 | Regenerative Mesenchyme | mmp9 |
| afap11 | 1.1E-17 | 0.7843094 | 0.115 | 0.069 | 3.48E-13 | Regenerative Mesenchyme | afap1 |
| gemin8 | 1.1E-17 | 0.5812452 | 0.129 | 0.079 | 3.53E-13 | Regenerative Mesenchyme | gemin8 |
| nfatc11 | 1.1E-17 | 0.6541551 | 0.134 | 0.084 | 3.73E-13 | Regenerative Mesenchyme | nfatc1 |
| map1sa | 1.2E-17 | 0.598176 | 0.151 | 0.097 | 3.81E-13 | Regenerative Mesenchyme | map1sa |
| zbtb8os | 1.2E-17 | 0.6561717 | 0.119 | 0.072 | 3.93E-13 | Regenerative Mesenchyme | zbtb8os |
| med11 | 1.4E-17 | 0.4264185 | 0.207 | 0.14 | 4.53E-13 | Regenerative Mesenchyme | med11 |
| gtf2e2 | 1.6E-17 | 0.6847191 | 0.143 | 0.091 | 5.33E-13 | Regenerative Mesenchyme | gtf2e2 |
| si:dkey-239h2.31 | 1.7E-17 | 0.4719672 | 0.218 | 0.152 | 5.43E-13 | Regenerative Mesenchyme | si:dkey-239h2.3 |
| vps26c | 1.8E-17 | 0.5802073 | 0.121 | 0.074 | 5.9E-13 | Regenerative Mesenchyme | vps26c |
| scaf11 | 2.1E-17 | 0.7635517 | 0.103 | 0.06 | 6.93E-13 | Regenerative Mesenchyme | scaf11 |
| hif1ab | 2.2E-17 | 0.3080067 | 0.461 | 0.37 | 7.11E-13 | Regenerative Mesenchyme | hif1ab |
| rras | 2.4E-17 | 0.5183634 | 0.225 | 0.158 | 7.83E-13 | Regenerative Mesenchyme | rras |
| ap3s2 | 2.5E-17 | 0.806769 | 0.103 | 0.06 | 7.99E-13 | Regenerative Mesenchyme | ap3s2 |
| si:dkey-25e12.3 | 2.5E-17 | 0.6789706 | 0.117 | 0.071 | 8.02E-13 | Regenerative Mesenchyme | si:dkey-25e12.3 |
| lgals9l3 | 3E-17 | 0.412075 | 0.181 | 0.12 | 9.63E-13 | Regenerative Mesenchyme | lgals9l3 |
| ngdn3 | 3.1E-17 | 0.4307708 | 0.222 | 0.154 | 1.02E-12 | Regenerative Mesenchyme | ngdn |
| zfr | 3.2E-17 | 0.5145635 | 0.167 | 0.11 | 1.03E-12 | Regenerative Mesenchyme | zfr |
| wasla | 3.6E-17 | 0.4479397 | 0.245 | 0.174 | 1.18E-12 | Regenerative Mesenchyme | wasla |
| ube2d2 | 4.1E-17 | 0.381894 | 0.469 | 0.385 | 1.33E-12 | Regenerative Mesenchyme | ube2d2 |
| cnot1 | 4.5E-17 | 0.5647997 | 0.161 | 0.105 | 1.47E-12 | Regenerative Mesenchyme | cnot1 |
| cx43.42 | 5.1E-17 | 0.4077699 | 0.292 | 0.218 | 1.65E-12 | Regenerative Mesenchyme | cx43.4 |
| derl1 | 6E-17 | 0.6597974 | 0.116 | 0.07 | 1.95E-12 | Regenerative Mesenchyme | derl1 |
| rnaseka1 | 6.2E-17 | 0.616374 | 0.158 | 0.103 | 2.01E-12 | Regenerative Mesenchyme | rnaseka |
| dbnlb | 6.5E-17 | 0.4746849 | 0.134 | 0.084 | 2.1E-12 | Regenerative Mesenchyme | dbnlb |
| rnf1851 | 6.9E-17 | 0.5768002 | 0.221 | 0.157 | 2.23E-12 | Regenerative Mesenchyme | rnf185 |
| mdc11 | 6.9E-17 | 0.4155271 | 0.158 | 0.103 | 2.24E-12 | Regenerative Mesenchyme | mdc1 |
| tax1bp3 | 6.9E-17 | 0.3944286 | 0.329 | 0.249 | 2.25E-12 | Regenerative Mesenchyme | tax1bp3 |
| lrrc152 | 7.3E-17 | 0.4832285 | 0.193 | 0.132 | 2.36E-12 | Regenerative Mesenchyme | lrrc15 |
| si:ch1073-392o20.21 | 8E-17 | 0.3676218 | 0.316 | 0.239 | 2.61E-12 | Regenerative Mesenchyme | si:ch1073-392o20.2 |
| erp44 | 8.3E-17 | 0.4921348 | 0.252 | 0.183 | 2.71E-12 | Regenerative Mesenchyme | erp44 |
| crtap | 8.6E-17 | 0.4225831 | 0.399 | 0.315 | 2.8E-12 | Regenerative Mesenchyme | crtap |
| snx6 | 9.5E-17 | 0.5532927 | 0.171 | 0.114 | 3.09E-12 | Regenerative Mesenchyme | snx6 |
| gtf2h5 | 1.1E-16 | 0.5426182 | 0.147 | 0.095 | 3.55E-12 | Regenerative Mesenchyme | gtf2h5 |
| vps36 | 1.1E-16 | 0.6539309 | 0.105 | 0.062 | 3.63E-12 | Regenerative Mesenchyme | vps36 |
| eif3i2 | 1.1E-16 | 0.3987149 | 0.422 | 0.341 | 3.71E-12 | Regenerative Mesenchyme | eif3i |
| nat9 | 1.1E-16 | 0.5139541 | 0.135 | 0.085 | 3.74E-12 | Regenerative Mesenchyme | nat9 |
| psip1b1 | 1.3E-16 | 0.2837175 | 0.258 | 0.186 | 4.14E-12 | Regenerative Mesenchyme | psip1b |
| baz1a | 1.3E-16 | 0.4745896 | 0.167 | 0.111 | 4.24E-12 | Regenerative Mesenchyme | baz1a |
| rfc1 | 1.4E-16 | 0.6378339 | 0.107 | 0.064 | 4.41E-12 | Regenerative Mesenchyme | rfc1 |
| smim71 | 1.5E-16 | 0.3830673 | 0.194 | 0.131 | 4.86E-12 | Regenerative Mesenchyme | smim7 |
| znf207b | 1.9E-16 | 0.6553085 | 0.112 | 0.068 | 6.05E-12 | Regenerative Mesenchyme | znf207b |
| etfa | 2.2E-16 | 0.4832929 | 0.221 | 0.157 | 7.07E-12 | Regenerative Mesenchyme | etfa |
| tbcb | 2.2E-16 | 0.3609442 | 0.292 | 0.214 | 7.29E-12 | Regenerative Mesenchyme | tbcb |
| get4 | 2.5E-16 | 0.5913375 | 0.125 | 0.078 | 8.21E-12 | Regenerative Mesenchyme | get4 |
| rabac11 | 2.8E-16 | 0.303387 | 0.476 | 0.393 | 9.07E-12 | Regenerative Mesenchyme | rabac1 |

| | | | | | | | |
|---|---|---|---|---|---|---|---|
| chd51 | 3.8E-16 | 0.3852374 | 0.184 | 0.124 | 1.23E-11 | Regenerative Mesenchyme | chd5 |
| anapc16 | 3.9E-16 | 0.4211554 | 0.207 | 0.144 | 1.27E-11 | Regenerative Mesenchyme | anapc16 |
| syap1 | 4.1E-16 | 0.4200864 | 0.219 | 0.154 | 1.34E-11 | Regenerative Mesenchyme | syap1 |
| prpf38b | 4.2E-16 | 0.49845 | 0.17 | 0.114 | 1.37E-11 | Regenerative Mesenchyme | prpf38b |
| sec62 | 6.4E-16 | 0.3658374 | 0.31 | 0.233 | 2.09E-11 | Regenerative Mesenchyme | sec62 |
| trip11 | 6.5E-16 | 0.523164 | 0.139 | 0.089 | 2.1E-11 | Regenerative Mesenchyme | trip11 |
| si:ch211-165e15.1 | 8.1E-16 | 0.6195836 | 0.107 | 0.065 | 2.64E-11 | Regenerative Mesenchyme | si:ch211-165e15.1 |
| sox4a | 8.7E-16 | 0.5281346 | 0.256 | 0.191 | 2.84E-11 | Regenerative Mesenchyme | sox4a |
| dsc2l1 | 9.1E-16 | 0.9903188 | 0.107 | 0.066 | 2.97E-11 | Regenerative Mesenchyme | dsc2l |
| plp2 | 9.5E-16 | 0.5530643 | 0.171 | 0.115 | 3.09E-11 | Regenerative Mesenchyme | plp2 |
| tnc4 | 1.1E-15 | 0.2106466 | 0.571 | 0.465 | 3.45E-11 | Regenerative Mesenchyme | tnc |
| soga11 | 1.2E-15 | 0.4039867 | 0.182 | 0.124 | 3.78E-11 | Regenerative Mesenchyme | soga1 |
| ammecr1 | 1.5E-15 | 0.4660928 | 0.204 | 0.143 | 4.91E-11 | Regenerative Mesenchyme | ammecr1 |
| ube4b | 1.6E-15 | 0.6035209 | 0.107 | 0.065 | 5.23E-11 | Regenerative Mesenchyme | ube4b |
| sulf1 | 1.7E-15 | 0.3810154 | 0.205 | 0.144 | 5.66E-11 | Regenerative Mesenchyme | sulf1 |
| plpp51 | 1.8E-15 | 0.784769 | 0.105 | 0.064 | 5.74E-11 | Regenerative Mesenchyme | plpp5 |
| arf6b | 1.9E-15 | 0.6638115 | 0.105 | 0.063 | 6.15E-11 | Regenerative Mesenchyme | arf6b |
| golgb11 | 2.2E-15 | 0.3025275 | 0.459 | 0.37 | 7.02E-11 | Regenerative Mesenchyme | golgb1 |
| cdkn1a1 | 2.6E-15 | 0.5051812 | 0.173 | 0.119 | 8.61E-11 | Regenerative Mesenchyme | cdkn1a |
| efemp2a1 | 2.7E-15 | 0.4156117 | 0.437 | 0.358 | 8.83E-11 | Regenerative Mesenchyme | efemp2a |
| fam20b | 3.3E-15 | 0.581998 | 0.104 | 0.063 | 1.06E-10 | Regenerative Mesenchyme | fam20b |
| purba | 3.6E-15 | 0.4552039 | 0.182 | 0.125 | 1.18E-10 | Regenerative Mesenchyme | purba |
| ythdf21 | 3.7E-15 | 0.6260684 | 0.153 | 0.102 | 1.2E-10 | Regenerative Mesenchyme | ythdf2 |
| zswim8 | 3.7E-15 | 0.6294649 | 0.125 | 0.08 | 1.2E-10 | Regenerative Mesenchyme | zswim8 |
| rhoaa | 3.8E-15 | 0.2925232 | 0.501 | 0.417 | 1.25E-10 | Regenerative Mesenchyme | rhoaa |
| tpi1b | 4.2E-15 | 0.2416028 | 0.441 | 0.353 | 1.35E-10 | Regenerative Mesenchyme | tpi1b |
| npc1 | 4.2E-15 | 0.5577864 | 0.116 | 0.072 | 1.37E-10 | Regenerative Mesenchyme | npc1 |
| otud5b | 4.2E-15 | 0.6340931 | 0.129 | 0.083 | 1.38E-10 | Regenerative Mesenchyme | otud5b |
| adamts15a3 | 4.4E-15 | 0.2324677 | 0.203 | 0.14 | 1.44E-10 | Regenerative Mesenchyme | adamts15a |
| niban2b | 6.3E-15 | 0.5367716 | 0.205 | 0.147 | 2.04E-10 | Regenerative Mesenchyme | niban2b |
| btbd10b | 6.4E-15 | 0.5786831 | 0.154 | 0.103 | 2.07E-10 | Regenerative Mesenchyme | btbd10b |
| CABZ01026681.1 | 6.5E-15 | 0.5290155 | 0.127 | 0.081 | 2.13E-10 | Regenerative Mesenchyme | CABZ01026681.1 |
| srcap | 6.8E-15 | 0.3540842 | 0.205 | 0.144 | 2.22E-10 | Regenerative Mesenchyme | srcap |
| ccdc124 | 7.1E-15 | 0.3265155 | 0.233 | 0.167 | 2.31E-10 | Regenerative Mesenchyme | ccdc124 |
| rbm22 | 7.1E-15 | 0.5302906 | 0.152 | 0.101 | 2.32E-10 | Regenerative Mesenchyme | rbm22 |
| ptp4a1 | 8.2E-15 | 0.3603632 | 0.303 | 0.23 | 2.67E-10 | Regenerative Mesenchyme | ptp4a1 |
| atp1b3b | 8.7E-15 | 0.4166946 | 0.122 | 0.077 | 2.84E-10 | Regenerative Mesenchyme | atp1b3b |
| taf7 | 9.1E-15 | 0.3871182 | 0.174 | 0.118 | 2.97E-10 | Regenerative Mesenchyme | taf7 |
| pho | 1.1E-14 | 0.2053709 | 0.295 | 0.22 | 3.47E-10 | Regenerative Mesenchyme | pho |
| morf4l1 | 1.1E-14 | 0.3294841 | 0.241 | 0.175 | 3.64E-10 | Regenerative Mesenchyme | morf4l1 |
| zgc:158803 | 1.3E-14 | 0.5235394 | 0.139 | 0.091 | 4.24E-10 | Regenerative Mesenchyme | zgc:158803 |
| atrx | 1.5E-14 | 0.2781905 | 0.545 | 0.463 | 4.81E-10 | Regenerative Mesenchyme | atrx |
| ccdc1671 | 1.7E-14 | 0.5791112 | 0.104 | 0.064 | 5.51E-10 | Regenerative Mesenchyme | ccdc167 |
| polr1a | 1.7E-14 | 0.5644932 | 0.119 | 0.075 | 5.54E-10 | Regenerative Mesenchyme | polr1a |
| magt11 | 2E-14 | 0.410603 | 0.331 | 0.258 | 6.63E-10 | Regenerative Mesenchyme | magt1 |
| ube2d1a | 2.4E-14 | 0.5508038 | 0.149 | 0.1 | 7.65E-10 | Regenerative Mesenchyme | ube2d1a |
| fdx1 | 2.7E-14 | 0.4453435 | 0.158 | 0.106 | 8.78E-10 | Regenerative Mesenchyme | fdx1 |
| sugt1 | 2.7E-14 | 0.5123914 | 0.159 | 0.108 | 8.78E-10 | Regenerative Mesenchyme | sugt1 |

| | | | | | | | |
|---|---|---|---|---|---|---|---|
| baz2a | 2.8E-14 | 0.4144086 | 0.153 | 0.102 | 9.11E-10 | Regenerative Mesenchyme | baz2a |
| itgav1 | 2.9E-14 | 0.5924069 | 0.139 | 0.092 | 9.29E-10 | Regenerative Mesenchyme | itgav |
| clint1a | 2.9E-14 | 0.3216818 | 0.275 | 0.205 | 9.37E-10 | Regenerative Mesenchyme | clint1a |
| mta3 | 3.7E-14 | 0.5047694 | 0.107 | 0.067 | 1.21E-09 | Regenerative Mesenchyme | mta3 |
| ccdc1871 | 4.1E-14 | 0.3671111 | 0.205 | 0.146 | 1.32E-09 | Regenerative Mesenchyme | ccdc187 |
| agpat4 | 4.2E-14 | 0.4782059 | 0.103 | 0.063 | 1.37E-09 | Regenerative Mesenchyme | agpat4 |
| fkbp1b1 | 4.8E-14 | 0.8004223 | 0.113 | 0.072 | 1.56E-09 | Regenerative Mesenchyme | fkbp1b |
| ss18 | 4.9E-14 | 0.4953167 | 0.132 | 0.086 | 1.58E-09 | Regenerative Mesenchyme | ss18 |
| erlin1 | 4.9E-14 | 0.526351 | 0.138 | 0.091 | 1.6E-09 | Regenerative Mesenchyme | erlin1 |
| safb | 5E-14 | 0.3368385 | 0.296 | 0.224 | 1.63E-09 | Regenerative Mesenchyme | safb |
| gskip1 | 6.5E-14 | 0.4395469 | 0.218 | 0.159 | 2.1E-09 | Regenerative Mesenchyme | gskip |
| map7d3 | 6.5E-14 | 0.608366 | 0.121 | 0.078 | 2.11E-09 | Regenerative Mesenchyme | map7d3 |
| chmp6b | 7E-14 | 0.4572709 | 0.207 | 0.149 | 2.28E-09 | Regenerative Mesenchyme | chmp6b |
| rhoca | 7E-14 | 0.3649012 | 0.328 | 0.254 | 2.29E-09 | Regenerative Mesenchyme | rhoca |
| sec61a1l | 7.1E-14 | 0.4639379 | 0.193 | 0.137 | 2.32E-09 | Regenerative Mesenchyme | sec61a1l |
| taf101 | 7.2E-14 | 0.7410444 | 0.115 | 0.073 | 2.34E-09 | Regenerative Mesenchyme | taf10 |
| birc6 | 8.4E-14 | 0.4299253 | 0.163 | 0.111 | 2.72E-09 | Regenerative Mesenchyme | birc6 |
| kpna3 | 8.8E-14 | 0.328349 | 0.209 | 0.149 | 2.85E-09 | Regenerative Mesenchyme | kpna3 |
| gadd45ab1 | 8.8E-14 | 0.2518821 | 0.237 | 0.173 | 2.87E-09 | Regenerative Mesenchyme | gadd45ab |
| srsf9 | 9.3E-14 | 0.42119 | 0.169 | 0.116 | 3.01E-09 | Regenerative Mesenchyme | srsf9 |
| lamtor5 | 1.1E-13 | 0.5345546 | 0.131 | 0.086 | 3.42E-09 | Regenerative Mesenchyme | lamtor5 |
| anxa13 | 1.4E-13 | 0.2867715 | 0.419 | 0.341 | 4.45E-09 | Regenerative Mesenchyme | anxa13 |
| tomm51 | 1.4E-13 | 0.2818746 | 0.515 | 0.445 | 4.55E-09 | Regenerative Mesenchyme | tomm5 |
| blcap | 1.7E-13 | 0.4059707 | 0.197 | 0.141 | 5.6E-09 | Regenerative Mesenchyme | blcap |
| mrpl231 | 1.8E-13 | 0.4925488 | 0.119 | 0.077 | 5.96E-09 | Regenerative Mesenchyme | mrpl23 |
| paics | 1.9E-13 | 0.5100606 | 0.125 | 0.081 | 6.26E-09 | Regenerative Mesenchyme | paics |
| pls3 | 2E-13 | 0.3568839 | 0.392 | 0.317 | 6.38E-09 | Regenerative Mesenchyme | pls3 |
| puf60a | 2E-13 | 0.4595238 | 0.206 | 0.149 | 6.49E-09 | Regenerative Mesenchyme | puf60a |
| higd1a1 | 2E-13 | 0.2087915 | 0.531 | 0.454 | 6.54E-09 | Regenerative Mesenchyme | higd1a |
| uso1 | 2E-13 | 0.4069815 | 0.168 | 0.116 | 6.66E-09 | Regenerative Mesenchyme | uso1 |
| fam214a | 2.2E-13 | 0.6178748 | 0.137 | 0.092 | 7.17E-09 | Regenerative Mesenchyme | fam214a |
| hsd17b12b | 2.4E-13 | 0.4240337 | 0.133 | 0.088 | 7.64E-09 | Regenerative Mesenchyme | hsd17b12b |
| scxa2 | 2.5E-13 | 0.6378677 | 0.123 | 0.081 | 8E-09 | Regenerative Mesenchyme | scxa |
| pgam5 | 3E-13 | 0.52079 | 0.113 | 0.072 | 9.63E-09 | Regenerative Mesenchyme | pgam5 |
| hnrnpua | 3.1E-13 | 0.3893973 | 0.252 | 0.189 | 1.02E-08 | Regenerative Mesenchyme | hnrnpua |
| zgc:56095 | 3.2E-13 | 0.495292 | 0.136 | 0.091 | 1.03E-08 | Regenerative Mesenchyme | zgc:56095 |
| ebag9 | 3.3E-13 | 0.6444291 | 0.114 | 0.073 | 1.06E-08 | Regenerative Mesenchyme | ebag9 |
| capgb1 | 3.6E-13 | 0.6449015 | 0.147 | 0.101 | 1.17E-08 | Regenerative Mesenchyme | capgb |
| ctbp1 | 4.2E-13 | 0.4555008 | 0.147 | 0.1 | 1.35E-08 | Regenerative Mesenchyme | ctbp1 |
| oaz1b1 | 4.4E-13 | 0.3102052 | 0.382 | 0.304 | 1.43E-08 | Regenerative Mesenchyme | oaz1b |
| chd21 | 5.3E-13 | 0.3018412 | 0.238 | 0.176 | 1.71E-08 | Regenerative Mesenchyme | chd2 |
| BX548011.2 | 5.6E-13 | 0.6229622 | 0.111 | 0.072 | 1.82E-08 | Regenerative Mesenchyme | BX548011.2 |
| eif3ea2 | 6E-13 | 0.2456221 | 0.618 | 0.557 | 1.95E-08 | Regenerative Mesenchyme | eif3ea |
| ube2d1b | 6.2E-13 | 0.4362923 | 0.19 | 0.136 | 2.01E-08 | Regenerative Mesenchyme | ube2d1b |
| rab35b | 6.6E-13 | 0.4241784 | 0.135 | 0.09 | 2.15E-08 | Regenerative Mesenchyme | rab35b |
| ckap5 | 6.8E-13 | 0.5680994 | 0.129 | 0.086 | 2.2E-08 | Regenerative Mesenchyme | ckap5 |
| ap3d1 | 7.5E-13 | 0.4129394 | 0.174 | 0.122 | 2.45E-08 | Regenerative Mesenchyme | ap3d1 |
| ogdha | 8E-13 | 0.3362957 | 0.277 | 0.211 | 2.62E-08 | Regenerative Mesenchyme | ogdha |

| | | | | | | | |
|---|---|---|---|---|---|---|---|
| supt6h1 | 8.8E-13 | 0.2917565 | 0.259 | 0.194 | 2.87E-08 | Regenerative Mesenchyme | supt6h |
| ptges1 | 9.1E-13 | 0.6517567 | 0.153 | 0.107 | 2.96E-08 | Regenerative Mesenchyme | ptges |
| smurf2 | 9.8E-13 | 0.4646013 | 0.103 | 0.065 | 3.18E-08 | Regenerative Mesenchyme | smurf2 |
| plekha8 | 1E-12 | 0.4835746 | 0.108 | 0.069 | 3.33E-08 | Regenerative Mesenchyme | plekha8 |
| si:ch211-194k22.8 | 1.1E-12 | 0.3771839 | 0.138 | 0.093 | 3.47E-08 | Regenerative Mesenchyme | si:ch211-194k22.8 |
| zgc:1750881 | 1.1E-12 | 0.3213054 | 0.34 | 0.271 | 3.54E-08 | Regenerative Mesenchyme | zgc:175088 |
| selenot2 | 1.2E-12 | 0.6059194 | 0.147 | 0.102 | 3.88E-08 | Regenerative Mesenchyme | selenot2 |
| fabp32 | 1.2E-12 | 0.2013194 | 0.61 | 0.527 | 3.98E-08 | Regenerative Mesenchyme | fabp3 |
| thrap3b | 1.4E-12 | 0.2710623 | 0.278 | 0.21 | 4.4E-08 | Regenerative Mesenchyme | thrap3b |
| uchl11 | 1.5E-12 | 0.5083264 | 0.176 | 0.126 | 4.89E-08 | Regenerative Mesenchyme | uchl1 |
| tegt | 1.6E-12 | 0.2658165 | 0.212 | 0.153 | 5.08E-08 | Regenerative Mesenchyme | tegt |
| ppp1cab | 1.8E-12 | 0.6503154 | 0.102 | 0.065 | 5.95E-08 | Regenerative Mesenchyme | ppp1cab |
| rgs12b1 | 1.9E-12 | 0.5435351 | 0.129 | 0.087 | 6.31E-08 | Regenerative Mesenchyme | rgs12b |
| dynlt3 | 2E-12 | 0.3695444 | 0.283 | 0.217 | 6.36E-08 | Regenerative Mesenchyme | dynlt3 |
| mta2 | 3.4E-12 | 0.5028039 | 0.135 | 0.092 | 1.1E-07 | Regenerative Mesenchyme | mta2 |
| ubr5 | 3.7E-12 | 0.4425191 | 0.142 | 0.098 | 1.22E-07 | Regenerative Mesenchyme | ubr5 |
| mcrip1 | 4E-12 | 0.5296372 | 0.112 | 0.073 | 1.29E-07 | Regenerative Mesenchyme | mcrip1 |
| qkia1 | 4.2E-12 | 0.2291631 | 0.539 | 0.465 | 1.37E-07 | Regenerative Mesenchyme | qkia |
| mbd1b | 4.8E-12 | 0.421443 | 0.122 | 0.081 | 1.56E-07 | Regenerative Mesenchyme | mbd1b |
| rnf20 | 5.3E-12 | 0.5203887 | 0.104 | 0.067 | 1.71E-07 | Regenerative Mesenchyme | rnf20 |
| eif3d1 | 5.5E-12 | 0.3018809 | 0.483 | 0.418 | 1.79E-07 | Regenerative Mesenchyme | eif3d |
| ylpm1 | 6.5E-12 | 0.3326714 | 0.167 | 0.118 | 2.1E-07 | Regenerative Mesenchyme | ylpm1 |
| txlna | 6.7E-12 | 0.4587076 | 0.145 | 0.1 | 2.16E-07 | Regenerative Mesenchyme | txlna |
| emc3 | 7E-12 | 0.6070616 | 0.116 | 0.077 | 2.29E-07 | Regenerative Mesenchyme | emc3 |
| ccng11 | 8.4E-12 | 0.2921213 | 0.559 | 0.503 | 2.75E-07 | Regenerative Mesenchyme | ccng1 |
| dhx38 | 8.7E-12 | 0.4374896 | 0.14 | 0.097 | 2.83E-07 | Regenerative Mesenchyme | dhx38 |
| kdelr32 | 9.1E-12 | 0.4212545 | 0.237 | 0.18 | 2.95E-07 | Regenerative Mesenchyme | kdelr3 |
| cs | 1E-11 | 0.4894927 | 0.201 | 0.149 | 3.24E-07 | Regenerative Mesenchyme | cs |
| nucb2a1 | 1.1E-11 | 0.2289644 | 0.59 | 0.521 | 3.65E-07 | Regenerative Mesenchyme | nucb2a |
| si:ch73-37h15.2 | 1.3E-11 | 0.2867022 | 0.153 | 0.106 | 4.3E-07 | Regenerative Mesenchyme | si:ch73-37h15.2 |
| atp5mea | 1.6E-11 | 0.2933215 | 0.317 | 0.251 | 5.18E-07 | Regenerative Mesenchyme | atp5mea |
| CDPF1 | 2E-11 | 0.3477625 | 0.233 | 0.176 | 6.47E-07 | Regenerative Mesenchyme | CDPF1 |
| dlgap4a | 2E-11 | 0.4393297 | 0.133 | 0.091 | 6.61E-07 | Regenerative Mesenchyme | dlgap4a |
| p4ha21 | 2.2E-11 | 0.4896299 | 0.191 | 0.14 | 7.25E-07 | Regenerative Mesenchyme | p4ha2 |
| nckap11 | 2.4E-11 | 0.2669387 | 0.301 | 0.235 | 7.76E-07 | Regenerative Mesenchyme | nckap1 |
| brd4 | 2.4E-11 | 0.2566243 | 0.327 | 0.259 | 7.77E-07 | Regenerative Mesenchyme | brd4 |
| crk | 2.6E-11 | 0.4085495 | 0.134 | 0.092 | 8.48E-07 | Regenerative Mesenchyme | crk |
| faf1 | 2.8E-11 | 0.491007 | 0.105 | 0.069 | 9.26E-07 | Regenerative Mesenchyme | faf1 |
| alas11 | 3.2E-11 | 0.4789305 | 0.149 | 0.105 | 1.05E-06 | Regenerative Mesenchyme | alas1 |
| sephs2 | 3.3E-11 | 0.5521068 | 0.111 | 0.074 | 1.06E-06 | Regenerative Mesenchyme | sephs2 |
| CTBP1 | 3.6E-11 | 0.2646368 | 0.223 | 0.166 | 1.17E-06 | Regenerative Mesenchyme | CTBP1 |
| chd4b | 3.8E-11 | 0.2190016 | 0.326 | 0.256 | 1.25E-06 | Regenerative Mesenchyme | chd4b |
| nipblb | 3.9E-11 | 0.2247263 | 0.249 | 0.188 | 1.26E-06 | Regenerative Mesenchyme | nipblb |
| ube2g1a | 3.9E-11 | 0.4858693 | 0.121 | 0.082 | 1.27E-06 | Regenerative Mesenchyme | ube2g1a |
| scpep11 | 4E-11 | 0.4585973 | 0.177 | 0.13 | 1.32E-06 | Regenerative Mesenchyme | scpep1 |
| si:ch73-238c9.11 | 4.2E-11 | 0.5107083 | 0.145 | 0.102 | 1.35E-06 | Regenerative Mesenchyme | si:ch73-238c9.1 |
| vta1 | 5.2E-11 | 0.5795259 | 0.11 | 0.074 | 1.7E-06 | Regenerative Mesenchyme | vta1 |
| ubl7b | 5.3E-11 | 0.4436146 | 0.111 | 0.074 | 1.74E-06 | Regenerative Mesenchyme | ubl7b |

| | | | | | | | |
|---|---|---|---|---|---|---|---|
| zgc:153911 | 5.8E-11 | 0.7689324 | 0.101 | 0.068 | 1.89E-06 | Regenerative Mesenchyme | zgc:153911 |
| mob3a | 6.3E-11 | 0.4807515 | 0.116 | 0.078 | 2.04E-06 | Regenerative Mesenchyme | mob3a |
| tpm4b2 | 6.6E-11 | 0.5979734 | 0.119 | 0.082 | 2.14E-06 | Regenerative Mesenchyme | tpm4b |
| chd9 | 6.6E-11 | 0.37774 | 0.193 | 0.142 | 2.16E-06 | Regenerative Mesenchyme | chd9 |
| ube2kb | 6.7E-11 | 0.4249827 | 0.119 | 0.081 | 2.17E-06 | Regenerative Mesenchyme | ube2kb |
| snw1 | 6.9E-11 | 0.2306327 | 0.252 | 0.192 | 2.26E-06 | Regenerative Mesenchyme | snw1 |
| plk31 | 7.4E-11 | 0.3192352 | 0.256 | 0.2 | 2.4E-06 | Regenerative Mesenchyme | plk3 |
| dda1 | 7.4E-11 | 0.3025782 | 0.207 | 0.153 | 2.42E-06 | Regenerative Mesenchyme | dda1 |
| fubp3 | 7.9E-11 | 0.4077488 | 0.112 | 0.075 | 2.57E-06 | Regenerative Mesenchyme | fubp3 |
| nipa2 | 8E-11 | 0.5372853 | 0.111 | 0.074 | 2.6E-06 | Regenerative Mesenchyme | nipa2 |
| smco41 | 8.5E-11 | 0.3686686 | 0.189 | 0.139 | 2.76E-06 | Regenerative Mesenchyme | smco4 |
| lamtor4 | 8.7E-11 | 0.3622703 | 0.145 | 0.101 | 2.82E-06 | Regenerative Mesenchyme | lamtor4 |
| ktn11 | 1E-10 | 0.2239357 | 0.338 | 0.269 | 3.29E-06 | Regenerative Mesenchyme | ktn1 |
| si:zfos-1192g2.3 | 1E-10 | 0.3601428 | 0.207 | 0.155 | 3.34E-06 | Regenerative Mesenchyme | si:zfos-1192g2.3 |
| lrch3 | 1.2E-10 | 0.5590433 | 0.113 | 0.077 | 3.86E-06 | Regenerative Mesenchyme | lrch3 |
| ndufaf31 | 1.2E-10 | 0.4044885 | 0.151 | 0.108 | 4.04E-06 | Regenerative Mesenchyme | ndufaf3 |
| dctn3 | 1.3E-10 | 0.4829897 | 0.198 | 0.149 | 4.07E-06 | Regenerative Mesenchyme | dctn3 |
| ergic2 | 1.3E-10 | 0.2818001 | 0.212 | 0.158 | 4.39E-06 | Regenerative Mesenchyme | ergic2 |
| scamp2 | 1.4E-10 | 0.3469253 | 0.166 | 0.12 | 4.57E-06 | Regenerative Mesenchyme | scamp2 |
| cnbpb | 1.6E-10 | 0.2734532 | 0.601 | 0.576 | 5.23E-06 | Regenerative Mesenchyme | cnbpb |
| rexo4 | 1.8E-10 | 0.2461542 | 0.12 | 0.082 | 6.01E-06 | Regenerative Mesenchyme | rexo4 |
| fhl1b1 | 1.9E-10 | 0.592442 | 0.131 | 0.093 | 6.02E-06 | Regenerative Mesenchyme | fhl1b |
| cops7a | 2E-10 | 0.4156492 | 0.154 | 0.111 | 6.52E-06 | Regenerative Mesenchyme | cops7a |
| gdi21 | 2.2E-10 | 0.2434745 | 0.532 | 0.471 | 7.1E-06 | Regenerative Mesenchyme | gdi2 |
| rbm4.2 | 2.3E-10 | 0.422385 | 0.215 | 0.163 | 7.63E-06 | Regenerative Mesenchyme | rbm4.2 |
| cttn | 2.5E-10 | 0.4408418 | 0.103 | 0.069 | 8.28E-06 | Regenerative Mesenchyme | cttn |
| fundc2 | 2.6E-10 | 0.284305 | 0.364 | 0.299 | 8.47E-06 | Regenerative Mesenchyme | fundc2 |
| capza1a | 2.6E-10 | 0.3275328 | 0.331 | 0.269 | 8.5E-06 | Regenerative Mesenchyme | capza1a |
| degs1 | 2.7E-10 | 0.3368133 | 0.241 | 0.186 | 8.63E-06 | Regenerative Mesenchyme | degs1 |
| CR381618.12 | 2.9E-10 | 0.3240425 | 0.281 | 0.225 | 9.43E-06 | Regenerative Mesenchyme | CR381618.1 |
| tnrc6b1 | 3E-10 | 0.3535047 | 0.185 | 0.137 | 9.72E-06 | Regenerative Mesenchyme | tnrc6b |
| vps35 | 3E-10 | 0.2870431 | 0.22 | 0.167 | 9.78E-06 | Regenerative Mesenchyme | vps35 |
| cops4 | 3E-10 | 0.4420881 | 0.152 | 0.11 | 9.85E-06 | Regenerative Mesenchyme | cops4 |
| fscn1a2 | 3.2E-10 | 0.4134393 | 0.227 | 0.176 | 1.05E-05 | Regenerative Mesenchyme | fscn1a |
| selenok | 3.3E-10 | 0.3089386 | 0.232 | 0.178 | 1.06E-05 | Regenerative Mesenchyme | selenok |
| ube2g2 | 3.8E-10 | 0.4303531 | 0.153 | 0.11 | 1.24E-05 | Regenerative Mesenchyme | ube2g2 |
| pdia72 | 4.7E-10 | 0.4306459 | 0.189 | 0.143 | 1.54E-05 | Regenerative Mesenchyme | pdia7 |
| dusp23 | 5.2E-10 | 0.2299101 | 0.274 | 0.22 | 1.69E-05 | Regenerative Mesenchyme | dusp2 |
| sfpq | 5.7E-10 | 0.2847109 | 0.262 | 0.204 | 1.84E-05 | Regenerative Mesenchyme | sfpq |
| hexb | 5.8E-10 | 0.5231244 | 0.127 | 0.09 | 1.88E-05 | Regenerative Mesenchyme | hexb |
| coro1ca2 | 7.3E-10 | 0.3084067 | 0.294 | 0.235 | 2.38E-05 | Regenerative Mesenchyme | coro1ca |
| pcmt | 7.7E-10 | 0.4431134 | 0.13 | 0.091 | 2.52E-05 | Regenerative Mesenchyme | pcmt |
| vps291 | 8E-10 | 0.3457783 | 0.254 | 0.199 | 2.59E-05 | Regenerative Mesenchyme | vps29 |
| yipf6 | 8E-10 | 0.4668194 | 0.111 | 0.076 | 2.59E-05 | Regenerative Mesenchyme | yipf6 |
| aup1 | 8E-10 | 0.431667 | 0.123 | 0.086 | 2.62E-05 | Regenerative Mesenchyme | aup1 |
| arpc5b | 9.4E-10 | 0.4715333 | 0.123 | 0.086 | 3.04E-05 | Regenerative Mesenchyme | arpc5b |
| ppme1 | 9.9E-10 | 0.4978112 | 0.142 | 0.102 | 3.21E-05 | Regenerative Mesenchyme | ppme1 |
| triqk1 | 1.1E-09 | 0.3619385 | 0.208 | 0.16 | 3.53E-05 | Regenerative Mesenchyme | triqk |

| | | | | | | | |
|---|---|---|---|---|---|---|---|
| tm9sf3 | 1.2E-09 | 0.3613828 | 0.22 | 0.17 | 3.94E-05 | Regenerative Mesenchyme | tm9sf3 |
| dcbld22 | 1.5E-09 | 0.4148509 | 0.166 | 0.123 | 4.79E-05 | Regenerative Mesenchyme | dcbld2 |
| strn3 | 1.5E-09 | 0.30506 | 0.231 | 0.179 | 4.93E-05 | Regenerative Mesenchyme | strn3 |
| cnot3a1 | 1.5E-09 | 0.270305 | 0.158 | 0.115 | 4.96E-05 | Regenerative Mesenchyme | cnot3a |
| vdac12 | 1.6E-09 | 0.321217 | 0.378 | 0.321 | 5.2E-05 | Regenerative Mesenchyme | vdac1 |
| ap3b1a | 1.8E-09 | 0.3254532 | 0.119 | 0.083 | 5.74E-05 | Regenerative Mesenchyme | ap3b1a |
| txndc9 | 2E-09 | 0.3420616 | 0.26 | 0.206 | 6.67E-05 | Regenerative Mesenchyme | txndc9 |
| gnai1 | 2.2E-09 | 0.3676176 | 0.205 | 0.157 | 7.07E-05 | Regenerative Mesenchyme | gnai1 |
| si:dkey-6n21.12 | 2.2E-09 | 0.3736178 | 0.172 | 0.128 | 7.23E-05 | Regenerative Mesenchyme | si:dkey-6n21.12 |
| dusp7 | 2.2E-09 | 0.5627088 | 0.105 | 0.073 | 7.25E-05 | Regenerative Mesenchyme | dusp7 |
| xrn2 | 2.3E-09 | 0.2145016 | 0.28 | 0.22 | 7.33E-05 | Regenerative Mesenchyme | xrn2 |
| vma21 | 2.4E-09 | 0.3640634 | 0.154 | 0.112 | 7.84E-05 | Regenerative Mesenchyme | vma21 |
| bloc1s2 | 2.5E-09 | 0.5031394 | 0.105 | 0.072 | 8.02E-05 | Regenerative Mesenchyme | bloc1s2 |
| eif1ad1 | 2.5E-09 | 0.2961124 | 0.184 | 0.138 | 8.18E-05 | Regenerative Mesenchyme | eif1ad |
| snx3 | 2.6E-09 | 0.3157555 | 0.227 | 0.176 | 8.29E-05 | Regenerative Mesenchyme | snx3 |
| ptbp1a1 | 2.6E-09 | 0.3585818 | 0.28 | 0.227 | 8.4E-05 | Regenerative Mesenchyme | ptbp1a |
| gosr2 | 2.8E-09 | 0.4435073 | 0.127 | 0.09 | 9.17E-05 | Regenerative Mesenchyme | gosr2 |
| get1 | 3.6E-09 | 0.4375523 | 0.107 | 0.074 | 0.000118 | Regenerative Mesenchyme | get1 |
| pora | 3.7E-09 | 0.2240672 | 0.15 | 0.109 | 0.000119 | Regenerative Mesenchyme | pora |
| chmp4bb | 4E-09 | 0.3776729 | 0.197 | 0.151 | 0.000131 | Regenerative Mesenchyme | chmp4bb |
| arglu1b | 4.3E-09 | 0.3649689 | 0.15 | 0.109 | 0.000138 | Regenerative Mesenchyme | arglu1b |
| trappc5 | 5.6E-09 | 0.4136343 | 0.152 | 0.112 | 0.000181 | Regenerative Mesenchyme | trappc5 |
| cbfb2 | 6E-09 | 0.5037411 | 0.162 | 0.122 | 0.000194 | Regenerative Mesenchyme | cbfb |
| vapal | 6.7E-09 | 0.3553172 | 0.21 | 0.163 | 0.000219 | Regenerative Mesenchyme | vapal |
| sat2b | 7.2E-09 | 0.2938964 | 0.133 | 0.095 | 0.000235 | Regenerative Mesenchyme | sat2b |
| kank3 | 7.5E-09 | 0.3762164 | 0.112 | 0.079 | 0.000244 | Regenerative Mesenchyme | kank3 |
| uqcc1 | 7.6E-09 | 0.3895368 | 0.119 | 0.085 | 0.000247 | Regenerative Mesenchyme | uqcc1 |
| acsl4b | 7.7E-09 | 0.3399344 | 0.173 | 0.13 | 0.000251 | Regenerative Mesenchyme | acsl4b |
| pafah1b1b | 7.9E-09 | 0.301988 | 0.259 | 0.205 | 0.000257 | Regenerative Mesenchyme | pafah1b1b |
| ap2b1 | 8.4E-09 | 0.3997036 | 0.153 | 0.113 | 0.000272 | Regenerative Mesenchyme | ap2b1 |
| psenen | 9E-09 | 0.3443206 | 0.166 | 0.124 | 0.000292 | Regenerative Mesenchyme | psenen |
| chd1 | 9.3E-09 | 0.4485254 | 0.139 | 0.102 | 0.000303 | Regenerative Mesenchyme | chd1 |
| timmdc1 | 1E-08 | 0.5186784 | 0.122 | 0.088 | 0.000327 | Regenerative Mesenchyme | timmdc1 |
| ccnl1b1 | 1.3E-08 | 0.3729944 | 0.116 | 0.083 | 0.000407 | Regenerative Mesenchyme | ccnl1b |
| usf1l | 1.3E-08 | 0.5282767 | 0.1 | 0.07 | 0.000423 | Regenerative Mesenchyme | usf1l |
| ctsh | 1.3E-08 | 0.3376077 | 0.147 | 0.109 | 0.000426 | Regenerative Mesenchyme | ctsh |
| CABZ01061894.1 | 1.3E-08 | 0.2567634 | 0.134 | 0.097 | 0.000437 | Regenerative Mesenchyme | CABZ01061894.1 |
| AL935174.6 | 1.4E-08 | 0.3548804 | 0.113 | 0.08 | 0.000445 | Regenerative Mesenchyme | AL935174.6 |
| lmcd11 | 1.4E-08 | 0.4866665 | 0.121 | 0.087 | 0.00045 | Regenerative Mesenchyme | lmcd1 |
| si:dkey-28b4.7 | 1.4E-08 | 0.3953254 | 0.245 | 0.197 | 0.000455 | Regenerative Mesenchyme | si:dkey-28b4.7 |
| acadvl | 1.5E-08 | 0.478232 | 0.133 | 0.097 | 0.000476 | Regenerative Mesenchyme | acadvl |
| morc3a | 1.5E-08 | 0.3051492 | 0.103 | 0.071 | 0.000485 | Regenerative Mesenchyme | morc3a |
| anp32e | 1.6E-08 | 0.3817032 | 0.185 | 0.141 | 0.000527 | Regenerative Mesenchyme | anp32e |
| ptgr11 | 1.7E-08 | 0.3375845 | 0.181 | 0.138 | 0.000569 | Regenerative Mesenchyme | ptgr1 |
| slc16a1b2 | 1.8E-08 | 0.2612806 | 0.172 | 0.129 | 0.000585 | Regenerative Mesenchyme | slc16a1b |
| tmx1 | 1.8E-08 | 0.2217198 | 0.159 | 0.117 | 0.000593 | Regenerative Mesenchyme | tmx1 |
| huwe1 | 1.8E-08 | 0.2815679 | 0.233 | 0.183 | 0.000595 | Regenerative Mesenchyme | huwe1 |
| ube2d3 | 2.1E-08 | 0.3559209 | 0.124 | 0.089 | 0.000696 | Regenerative Mesenchyme | ube2d3 |

| | | | | | | | |
|---|---|---|---|---|---|---|---|
| pum1 | 2.3E-08 | 0.4348209 | 0.131 | 0.096 | 0.000763 | Regenerative Mesenchyme | pum1 |
| cwc271 | 2.4E-08 | 0.2080593 | 0.163 | 0.121 | 0.000789 | Regenerative Mesenchyme | cwc27 |
| tor1 | 2.5E-08 | 0.2316544 | 0.145 | 0.106 | 0.000801 | Regenerative Mesenchyme | tor1 |
| chmp7 | 2.5E-08 | 0.4597367 | 0.106 | 0.074 | 0.00081 | Regenerative Mesenchyme | chmp7 |
| snx41 | 2.6E-08 | 0.485331 | 0.125 | 0.091 | 0.000835 | Regenerative Mesenchyme | snx4 |
| ctcf | 2.9E-08 | 0.348567 | 0.16 | 0.12 | 0.00093 | Regenerative Mesenchyme | ctcf |
| kdm5c | 3.2E-08 | 0.3099434 | 0.193 | 0.15 | 0.001056 | Regenerative Mesenchyme | kdm5c |
| fam50a | 3.3E-08 | 0.2541331 | 0.206 | 0.16 | 0.001073 | Regenerative Mesenchyme | fam50a |
| perp | 3.9E-08 | 0.3673484 | 0.284 | 0.235 | 0.00127 | Regenerative Mesenchyme | perp |
| prkag1 | 4.4E-08 | 0.291896 | 0.13 | 0.095 | 0.001421 | Regenerative Mesenchyme | prkag1 |
| ddx212 | 4.8E-08 | 0.223384 | 0.4 | 0.34 | 0.001547 | Regenerative Mesenchyme | ddx21 |
| itsn2b | 4.8E-08 | 0.3840599 | 0.181 | 0.14 | 0.001552 | Regenerative Mesenchyme | itsn2b |
| CU929070.52 | 4.8E-08 | 0.2739117 | 0.289 | 0.238 | 0.00156 | Regenerative Mesenchyme | CU929070.5 |
| rraga | 5.1E-08 | 0.3421968 | 0.104 | 0.073 | 0.001645 | Regenerative Mesenchyme | rraga |
| mpc2 | 5.6E-08 | 0.3485905 | 0.226 | 0.18 | 0.001831 | Regenerative Mesenchyme | mpc2 |
| adipor21 | 6.9E-08 | 0.434242 | 0.219 | 0.175 | 0.002248 | Regenerative Mesenchyme | adipor2 |
| mtrr | 7E-08 | 0.2697037 | 0.125 | 0.091 | 0.002265 | Regenerative Mesenchyme | mtrr |
| cib1 | 7.4E-08 | 0.3187292 | 0.158 | 0.119 | 0.002405 | Regenerative Mesenchyme | cib1 |
| ptp4a2a | 7.6E-08 | 0.4465333 | 0.176 | 0.137 | 0.00246 | Regenerative Mesenchyme | ptp4a2a |
| hnrnpk | 7.7E-08 | 0.4178898 | 0.19 | 0.149 | 0.002506 | Regenerative Mesenchyme | hnrnpk |
| ostf1 | 7.9E-08 | 0.4977994 | 0.111 | 0.08 | 0.002582 | Regenerative Mesenchyme | ostf1 |
| sh3bgrl3 | 8E-08 | 0.3586553 | 0.173 | 0.133 | 0.00259 | Regenerative Mesenchyme | sh3bgrl3 |
| ist1 | 8E-08 | 0.3618345 | 0.106 | 0.075 | 0.002602 | Regenerative Mesenchyme | ist1 |
| si:dkey-188i13.11 | 8.8E-08 | 0.3150366 | 0.248 | 0.201 | 0.002852 | Regenerative Mesenchyme | si:dkey-188i13.11 |
| rbp51 | 9.4E-08 | 0.3020864 | 0.154 | 0.117 | 0.003053 | Regenerative Mesenchyme | rbp5 |
| ankhd1 | 9.5E-08 | 0.2593239 | 0.201 | 0.158 | 0.003089 | Regenerative Mesenchyme | ankhd1 |
| usp1 | 9.9E-08 | 0.2508404 | 0.117 | 0.084 | 0.003218 | Regenerative Mesenchyme | usp1 |
| kctd10 | 1E-07 | 0.4555471 | 0.102 | 0.072 | 0.003297 | Regenerative Mesenchyme | kctd10 |
| u2af2a | 1E-07 | 0.2839129 | 0.142 | 0.105 | 0.003359 | Regenerative Mesenchyme | u2af2a |
| smad5 | 1.1E-07 | 0.472185 | 0.112 | 0.081 | 0.003472 | Regenerative Mesenchyme | smad5 |
| pdcd10b | 1.2E-07 | 0.3448962 | 0.133 | 0.099 | 0.003919 | Regenerative Mesenchyme | pdcd10b |
| fkbp31 | 1.3E-07 | 0.2289135 | 0.413 | 0.355 | 0.004219 | Regenerative Mesenchyme | fkbp3 |
| plcg1 | 1.4E-07 | 0.5265448 | 0.102 | 0.073 | 0.00442 | Regenerative Mesenchyme | plcg1 |
| slc25a55a1 | 1.4E-07 | 0.3246322 | 0.24 | 0.196 | 0.004439 | Regenerative Mesenchyme | slc25a55a |
| supt5h | 1.5E-07 | 0.3231539 | 0.113 | 0.082 | 0.005009 | Regenerative Mesenchyme | supt5h |
| CR848833.11 | 1.6E-07 | 0.2416479 | 0.1 | 0.071 | 0.005149 | Regenerative Mesenchyme | CR848833.1 |
| pde4ba | 1.7E-07 | 0.3845213 | 0.151 | 0.116 | 0.005476 | Regenerative Mesenchyme | pde4ba |
| zgc:1034381 | 1.8E-07 | 0.2689581 | 0.119 | 0.088 | 0.005801 | Regenerative Mesenchyme | zgc:103438 |
| grb2b | 1.9E-07 | 0.3563671 | 0.202 | 0.161 | 0.006154 | Regenerative Mesenchyme | grb2b |
| nhsl1a | 2E-07 | 0.2436902 | 0.109 | 0.078 | 0.006447 | Regenerative Mesenchyme | nhsl1a |
| pwwp2b | 2.2E-07 | 0.2982706 | 0.102 | 0.073 | 0.00728 | Regenerative Mesenchyme | pwwp2b |
| flii1 | 2.4E-07 | 0.3703558 | 0.143 | 0.108 | 0.007706 | Regenerative Mesenchyme | flii |
| C5H9orf78 | 3.1E-07 | 0.3695976 | 0.116 | 0.085 | 0.009971 | Regenerative Mesenchyme | C5H9orf78 |
| rab8b | 3.3E-07 | 0.2744735 | 0.174 | 0.135 | 0.010588 | Regenerative Mesenchyme | rab8b |
| tardbpl | 3.3E-07 | 0.2037808 | 0.197 | 0.154 | 0.01062 | Regenerative Mesenchyme | tardbpl |
| setdb1a | 3.3E-07 | 0.3745164 | 0.103 | 0.074 | 0.01083 | Regenerative Mesenchyme | setdb1a |
| nap1l4b1 | 3.6E-07 | 0.3295061 | 0.149 | 0.114 | 0.01169 | Regenerative Mesenchyme | nap1l4b |
| cpsf6 | 3.7E-07 | 0.207047 | 0.263 | 0.214 | 0.011962 | Regenerative Mesenchyme | cpsf6 |

| | | | | | | | |
|---|---|---|---|---|---|---|---|
| sf1 | 3.8E-07 | 0.2853155 | 0.284 | 0.235 | 0.012326 | Regenerative Mesenchyme | sf1 |
| tssc4 | 3.9E-07 | 0.2721409 | 0.123 | 0.091 | 0.012538 | Regenerative Mesenchyme | tssc4 |
| fubp1 | 3.9E-07 | 0.2691596 | 0.237 | 0.191 | 0.01264 | Regenerative Mesenchyme | fubp1 |
| trip12 | 4.1E-07 | 0.3225754 | 0.136 | 0.102 | 0.013357 | Regenerative Mesenchyme | trip12 |
| uri1 | 4.4E-07 | 0.2344515 | 0.166 | 0.128 | 0.01424 | Regenerative Mesenchyme | uri1 |
| ubap2l | 5.3E-07 | 0.3269652 | 0.19 | 0.15 | 0.017205 | Regenerative Mesenchyme | ubap2l |
| epn2 | 5.8E-07 | 0.2865888 | 0.142 | 0.108 | 0.018856 | Regenerative Mesenchyme | epn2 |
| mafba2 | 5.9E-07 | 0.3215099 | 0.431 | 0.389 | 0.019162 | Regenerative Mesenchyme | mafba |
| parp4 | 6E-07 | 0.271176 | 0.113 | 0.083 | 0.019647 | Regenerative Mesenchyme | parp4 |
| ano62 | 6.1E-07 | 0.2247851 | 0.161 | 0.124 | 0.019824 | Regenerative Mesenchyme | ano6 |
| ARF5 | 6.3E-07 | 0.2202006 | 0.192 | 0.151 | 0.020449 | Regenerative Mesenchyme | ARF5 |
| dchs1b3 | 6.9E-07 | 0.2167621 | 0.121 | 0.09 | 0.022521 | Regenerative Mesenchyme | dchs1b |
| lrrfip21 | 7E-07 | 0.2843493 | 0.137 | 0.103 | 0.022787 | Regenerative Mesenchyme | lrrfip2 |
| tbc1d23 | 7.1E-07 | 0.4036164 | 0.1 | 0.073 | 0.022961 | Regenerative Mesenchyme | tbc1d23 |
| elof11 | 7.1E-07 | 0.3020203 | 0.181 | 0.143 | 0.023151 | Regenerative Mesenchyme | elof1 |
| sf3b6 | 7.5E-07 | 0.2040703 | 0.334 | 0.28 | 0.024348 | Regenerative Mesenchyme | sf3b6 |
| dctn2 | 7.7E-07 | 0.2773675 | 0.163 | 0.126 | 0.025004 | Regenerative Mesenchyme | dctn2 |
| rab1ba | 8.1E-07 | 0.3076983 | 0.177 | 0.138 | 0.026247 | Regenerative Mesenchyme | rab1ba |
| ptgs2a2 | 8.7E-07 | 0.3672237 | 0.162 | 0.129 | 0.028158 | Regenerative Mesenchyme | ptgs2a |
| ctnnd1 | 8.9E-07 | 0.3458336 | 0.205 | 0.165 | 0.029007 | Regenerative Mesenchyme | ctnnd1 |
| rbm7 | 9.4E-07 | 0.3022989 | 0.116 | 0.086 | 0.030482 | Regenerative Mesenchyme | rbm7 |
| usp81 | 1E-06 | 0.2122126 | 0.129 | 0.097 | 0.033963 | Regenerative Mesenchyme | usp8 |
| bmp1a | 1.1E-06 | 0.2973249 | 0.226 | 0.186 | 0.035957 | Regenerative Mesenchyme | bmp1a |
| ppp2r5eb | 1.1E-06 | 0.2823735 | 0.193 | 0.154 | 0.037249 | Regenerative Mesenchyme | ppp2r5eb |
| ccdc6b | 1.2E-06 | 0.2747181 | 0.108 | 0.079 | 0.037618 | Regenerative Mesenchyme | ccdc6b |
| map3k4 | 1.3E-06 | 0.3217617 | 0.112 | 0.083 | 0.041113 | Regenerative Mesenchyme | map3k4 |
| ap1ar | 1.3E-06 | 0.2148134 | 0.111 | 0.082 | 0.043604 | Regenerative Mesenchyme | ap1ar |
| snx5 | 1.9E-06 | 0.2502832 | 0.113 | 0.084 | 0.061118 | Regenerative Mesenchyme | snx5 |
| gusb | 2E-06 | 0.4206804 | 0.162 | 0.128 | 0.066474 | Regenerative Mesenchyme | gusb |
| palm2akap2.12 | 2.2E-06 | 0.3581858 | 0.112 | 0.084 | 0.070293 | Regenerative Mesenchyme | palm2akap2.1 |
| zgc:153409 | 2.3E-06 | 0.3958821 | 0.458 | 0.419 | 0.073708 | Regenerative Mesenchyme | zgc:153409 |
| kdm1a1 | 2.3E-06 | 0.2597749 | 0.165 | 0.13 | 0.075949 | Regenerative Mesenchyme | kdm1a |
| pdhb | 2.4E-06 | 0.3685096 | 0.14 | 0.108 | 0.076445 | Regenerative Mesenchyme | pdhb |
| scocb | 2.4E-06 | 0.47694 | 0.108 | 0.081 | 0.077975 | Regenerative Mesenchyme | scocb |
| nfkbiab1 | 2.7E-06 | 0.2102111 | 0.622 | 0.582 | 0.088953 | Regenerative Mesenchyme | nfkbiab |
| ino80 | 2.9E-06 | 0.2668853 | 0.134 | 0.103 | 0.095599 | Regenerative Mesenchyme | ino80 |
| armh3 | 3.2E-06 | 0.2168306 | 0.113 | 0.085 | 0.104298 | Regenerative Mesenchyme | armh3 |
| lrrc8aa | 3.3E-06 | 0.2772629 | 0.129 | 0.098 | 0.106275 | Regenerative Mesenchyme | lrrc8aa |
| tcta1 | 3.6E-06 | 0.3879968 | 0.117 | 0.089 | 0.118458 | Regenerative Mesenchyme | tcta |
| dspa1 | 3.7E-06 | 0.4896994 | 0.155 | 0.125 | 0.120123 | Regenerative Mesenchyme | dspa |
| naa30 | 3.8E-06 | 0.3074413 | 0.106 | 0.079 | 0.124139 | Regenerative Mesenchyme | naa30 |
| cnpy31 | 3.9E-06 | 0.2134353 | 0.116 | 0.087 | 0.125637 | Regenerative Mesenchyme | cnpy3 |
| cybrd11 | 4.1E-06 | 0.3345575 | 0.109 | 0.082 | 0.132877 | Regenerative Mesenchyme | cybrd1 |
| tmem45a1 | 5.3E-06 | 0.400049 | 0.19 | 0.156 | 0.172737 | Regenerative Mesenchyme | tmem45a |
| kng12 | 5.6E-06 | 0.3407833 | 0.106 | 0.08 | 0.182226 | Regenerative Mesenchyme | kng1 |
| sinhcafl | 6.1E-06 | 0.2589519 | 0.157 | 0.124 | 0.198589 | Regenerative Mesenchyme | sinhcafl |
| kif13ba | 6.5E-06 | 0.3245745 | 0.131 | 0.102 | 0.211662 | Regenerative Mesenchyme | kif13ba |
| ell2 | 6.9E-06 | 0.2371591 | 0.172 | 0.137 | 0.22314 | Regenerative Mesenchyme | ell2 |

| | | | | | | | |
|---|---|---|---|---|---|---|---|
| top2b1 | 8.2E-06 | 0.2124114 | 0.31 | 0.264 | 0.266228 | Regenerative Mesenchyme | top2b |
| srgap2 | 8.8E-06 | 0.3339954 | 0.11 | 0.083 | 0.286593 | Regenerative Mesenchyme | srgap2 |
| tmed32 | 9.4E-06 | 0.2806169 | 0.124 | 0.095 | 0.306415 | Regenerative Mesenchyme | tmed3 |
| unc501 | 1.1E-05 | 0.2623364 | 0.106 | 0.08 | 0.34317 | Regenerative Mesenchyme | unc50 |
| chmp2bb | 1.1E-05 | 0.2795259 | 0.13 | 0.101 | 0.35974 | Regenerative Mesenchyme | chmp2bb |
| bmp51 | 1.1E-05 | 0.3884101 | 0.114 | 0.087 | 0.362458 | Regenerative Mesenchyme | bmp5 |
| mxra8a2 | 1.4E-05 | 0.306346 | 0.29 | 0.252 | 0.464663 | Regenerative Mesenchyme | mxra8a |
| arfip1 | 1.8E-05 | 0.2123129 | 0.125 | 0.097 | 0.572835 | Regenerative Mesenchyme | arfip1 |
| ppfibp1b1 | 2.1E-05 | 0.3435524 | 0.174 | 0.142 | 0.681097 | Regenerative Mesenchyme | ppfibp1b |
| mob4 | 2.1E-05 | 0.2477326 | 0.157 | 0.125 | 0.689019 | Regenerative Mesenchyme | mob4 |
| ubr4 | 2.3E-05 | 0.2606979 | 0.127 | 0.099 | 0.742177 | Regenerative Mesenchyme | ubr4 |
| ctsla1 | 2.3E-05 | 0.2909999 | 0.755 | 0.72 | 0.744755 | Regenerative Mesenchyme | ctsla |
| hist1h4l.31 | 2.4E-05 | 0.9486858 | 0.169 | 0.142 | 0.766504 | Regenerative Mesenchyme | hist1h4l.3 |
| zgc:162698 | 2.5E-05 | 0.2093648 | 0.143 | 0.113 | 0.813122 | Regenerative Mesenchyme | zgc:162698 |
| glyr11 | 3E-05 | 0.2111254 | 0.129 | 0.101 | 0.977627 | Regenerative Mesenchyme | glyr1 |
| wu:fi09b08 | 3.1E-05 | 0.2137908 | 0.113 | 0.087 | 1 | Regenerative Mesenchyme | wu:fi09b08 |
| rassf11 | 3.2E-05 | 0.3458588 | 0.148 | 0.12 | 1 | Regenerative Mesenchyme | rassf1 |
| si:dkey-262k9.2 | 3.3E-05 | 0.6964445 | 0.125 | 0.102 | 1 | Regenerative Mesenchyme | si:dkey-262k9.2 |
| p3h32 | 3.3E-05 | 0.2725456 | 0.165 | 0.134 | 1 | Regenerative Mesenchyme | p3h3 |
| gsk3ba | 3.5E-05 | 0.3054377 | 0.175 | 0.144 | 1 | Regenerative Mesenchyme | gsk3ba |
| lztfl1 | 4.3E-05 | 0.2288471 | 0.106 | 0.081 | 1 | Regenerative Mesenchyme | lztfl1 |
| caap1 | 4.8E-05 | 0.2118023 | 0.102 | 0.078 | 1 | Regenerative Mesenchyme | caap1 |
| smc5 | 5E-05 | 0.2729412 | 0.112 | 0.087 | 1 | Regenerative Mesenchyme | smc5 |
| pef1 | 5.3E-05 | 0.2338758 | 0.189 | 0.156 | 1 | Regenerative Mesenchyme | pef1 |
| aamp2 | 6.9E-05 | 0.2570556 | 0.185 | 0.153 | 1 | Regenerative Mesenchyme | aamp |
| chmp1b | 8E-05 | 0.2463011 | 0.146 | 0.118 | 1 | Regenerative Mesenchyme | chmp1b |
| ccdc6a | 8.2E-05 | 0.2702218 | 0.141 | 0.113 | 1 | Regenerative Mesenchyme | ccdc6a |
| bccip1 | 8.5E-05 | 0.2244821 | 0.229 | 0.193 | 1 | Regenerative Mesenchyme | bccip |
| cdc73 | 9.1E-05 | 0.2257033 | 0.105 | 0.081 | 1 | Regenerative Mesenchyme | cdc73 |
| luc7l3 | 9.1E-05 | 0.2277638 | 0.145 | 0.116 | 1 | Regenerative Mesenchyme | luc7l3 |
| mfap3l | 0.0001 | 0.2251424 | 0.113 | 0.089 | 1 | Regenerative Mesenchyme | mfap3l |
| larp4ab | 0.0001 | 0.32828 | 0.189 | 0.159 | 1 | Regenerative Mesenchyme | larp4ab |
| serpine21 | 0.0001 | 0.5553084 | 0.339 | 0.316 | 1 | Regenerative Mesenchyme | serpine2 |
| mapk14a1 | 0.0001 | 0.2713008 | 0.107 | 0.083 | 1 | Regenerative Mesenchyme | mapk14a |
| echs1 | 0.00013 | 0.2975242 | 0.168 | 0.139 | 1 | Regenerative Mesenchyme | echs1 |
| ch25hl31 | 0.00014 | 0.2093631 | 0.132 | 0.106 | 1 | Regenerative Mesenchyme | ch25hl3 |
| acadl | 0.00014 | 0.2357853 | 0.115 | 0.091 | 1 | Regenerative Mesenchyme | acadl |
| FOXK21 | 0.00015 | 0.2900768 | 0.163 | 0.136 | 1 | Regenerative Mesenchyme | FOXK2 |
| mxra8b1 | 0.00017 | 0.2872634 | 0.438 | 0.407 | 1 | Regenerative Mesenchyme | mxra8b |
| galnt71 | 0.00022 | 0.2013792 | 0.143 | 0.116 | 1 | Regenerative Mesenchyme | galnt7 |
| ubtd2 | 0.00026 | 0.2011748 | 0.128 | 0.103 | 1 | Regenerative Mesenchyme | ubtd2 |
| bcl7bb1 | 0.00036 | 0.3259558 | 0.111 | 0.09 | 1 | Regenerative Mesenchyme | bcl7bb |
| dnaja2b | 0.00039 | 0.2714158 | 0.135 | 0.11 | 1 | Regenerative Mesenchyme | dnaja2b |
| ano5b1 | 0.0004 | 0.2072854 | 0.108 | 0.086 | 1 | Regenerative Mesenchyme | ano5b |
| nfil31 | 0.00045 | 0.3067643 | 0.101 | 0.081 | 1 | Regenerative Mesenchyme | nfil3 |
| clic4 | 0.00046 | 0.2308719 | 0.229 | 0.198 | 1 | Regenerative Mesenchyme | clic4 |
| ufd1l1 | 0.00053 | 0.2053477 | 0.13 | 0.106 | 1 | Regenerative Mesenchyme | ufd1l |
| ppp2r1ba | 0.00055 | 0.2075208 | 0.183 | 0.155 | 1 | Regenerative Mesenchyme | ppp2r1ba |

| | | | | | | | |
|---|---|---|---|---|---|---|---|
| pdcd6ip | 0.00068 | 0.2108544 | 0.157 | 0.131 | 1 | Regenerative Mesenchyme | pdcd6ip |
| tpte1 | 0.00077 | 0.2166168 | 0.108 | 0.087 | 1 | Regenerative Mesenchyme | tpte |
| erbin | 0.00082 | 0.2117537 | 0.1 | 0.08 | 1 | Regenerative Mesenchyme | erbin |
| zgc:1008293 | 0.00104 | 0.2656277 | 0.142 | 0.119 | 1 | Regenerative Mesenchyme | zgc:100829 |
| inppl1a1 | 0.00145 | 0.2751717 | 0.107 | 0.088 | 1 | Regenerative Mesenchyme | inppl1a |
| atp6v1e1a | 0.0028 | 0.2872945 | 0.123 | 0.104 | 1 | Regenerative Mesenchyme | atp6v1e1a |
| p4ha1a1 | 0.00345 | 0.3023879 | 0.114 | 0.096 | 1 | Regenerative Mesenchyme | p4ha1a |
| selenow2a | 0.00416 | 0.2242066 | 0.187 | 0.163 | 1 | Regenerative Mesenchyme | selenow2a |
| nek7 | 0.00448 | 0.215015 | 0.139 | 0.119 | 1 | Regenerative Mesenchyme | nek7 |
| slc66a31 | 0.00483 | 0.3101116 | 0.138 | 0.119 | 1 | Regenerative Mesenchyme | slc66a3 |
| si:dkey-193p11.21 | 0.00514 | 0.5804314 | 0.278 | 0.257 | 1 | Regenerative Mesenchyme | si:dkey-193p11.2 |
| rab11ba | 0.00586 | 0.203437 | 0.207 | 0.183 | 1 | Regenerative Mesenchyme | rab11ba |
| ube2a | 0.00643 | 0.2031335 | 0.1 | 0.083 | 1 | Regenerative Mesenchyme | ube2a |
| atp6ap2 | 0.00865 | 0.30002 | 0.117 | 0.1 | 1 | Regenerative Mesenchyme | atp6ap2 |
| angptl71 | 0 | 2.8785151 | 0.618 | 0.179 | 0 | Stroma 2 | angptl7 |
| dpt1 | 0 | 2.5884797 | 0.561 | 0.131 | 0 | Stroma 2 | dpt |
| mdka2 | 0 | 1.9035692 | 0.899 | 0.5 | 0 | Stroma 2 | mdka |
| abi3bpa2 | 0 | 1.9496922 | 0.667 | 0.278 | 0 | Stroma 2 | abi3bpa |
| sfrp1a2 | 0 | 1.4590285 | 0.771 | 0.418 | 0 | Stroma 2 | sfrp1a |
| ccl25b2 | 0 | 1.7785119 | 0.926 | 0.649 | 0 | Stroma 2 | ccl25b |
| ntn1a2 | 0 | 2.0112609 | 0.372 | 0.106 | 0 | Stroma 2 | ntn1a |
| mdkb | 0 | 3.3804989 | 0.216 | 0.032 | 0 | Stroma 2 | mdkb |
| asip2b | 3E-304 | 2.8029795 | 0.377 | 0.121 | 9.3E-300 | Stroma 2 | asip2b |
| sfrp1b2 | 5E-282 | 1.5904845 | 0.654 | 0.345 | 1.6E-277 | Stroma 2 | sfrp1b |
| crhbp | 4E-231 | 3.1954953 | 0.165 | 0.032 | 1.4E-226 | Stroma 2 | crhbp |
| crip11 | 3E-225 | 0.8843852 | 0.743 | 0.446 | 1.1E-220 | Stroma 2 | crip1 |
| fgl2b1 | 3E-213 | 2.0039188 | 0.319 | 0.112 | 8.2E-209 | Stroma 2 | fgl2b |
| myoc1 | 3E-199 | 2.9130364 | 0.217 | 0.06 | 1.1E-194 | Stroma 2 | myoc |
| c6.11 | 8E-199 | 1.4938228 | 0.437 | 0.192 | 2.7E-194 | Stroma 2 | c6.1 |
| rgs5a1 | 3E-162 | 0.9687484 | 0.344 | 0.141 | 1E-157 | Stroma 2 | rgs5a |
| thbs1b | 5E-162 | 1.2700881 | 0.42 | 0.202 | 1.6E-157 | Stroma 2 | thbs1b |
| hand2 | 4E-149 | 2.1370897 | 0.149 | 0.038 | 1.2E-144 | Stroma 2 | hand2 |
| aqp3a2 | 4E-138 | 1.1223372 | 0.55 | 0.306 | 1.3E-133 | Stroma 2 | aqp3a |
| si:ch211-227m13.1 | 1E-135 | 1.6776421 | 0.165 | 0.047 | 4.6E-131 | Stroma 2 | si:ch211-227m13.1 |
| c4b | 5E-135 | 1.8737988 | 0.167 | 0.049 | 1.7E-130 | Stroma 2 | c4b |
| apcdd1l2 | 2E-134 | 1.4922455 | 0.262 | 0.102 | 5.5E-130 | Stroma 2 | apcdd1l |
| ackr4b | 1E-126 | 2.3291627 | 0.155 | 0.045 | 4.8E-122 | Stroma 2 | ackr4b |
| ponzr11 | 4E-124 | 1.128062 | 0.421 | 0.223 | 1.3E-119 | Stroma 2 | ponzr1 |
| dkk2 | 1E-118 | 1.8093051 | 0.16 | 0.049 | 3.6E-114 | Stroma 2 | dkk2 |
| cxcl12a3 | 6E-118 | 0.8047092 | 0.871 | 0.672 | 1.8E-113 | Stroma 2 | cxcl12a |
| si:dkey-261h17.12 | 2E-116 | 0.9720409 | 0.487 | 0.28 | 7E-112 | Stroma 2 | si:dkey-261h17.1 |
| c41 | 2E-115 | 1.1751258 | 0.389 | 0.197 | 5.9E-111 | Stroma 2 | c4 |
| si:dkey-282h22.51 | 3E-115 | 1.7758646 | 0.186 | 0.064 | 9.3E-111 | Stroma 2 | si:dkey-282h22.5 |
| nova21 | 2E-113 | 1.1550377 | 0.392 | 0.204 | 6.6E-109 | Stroma 2 | nova2 |
| pmp22a1 | 3E-113 | 0.7629012 | 0.818 | 0.682 | 8.6E-109 | Stroma 2 | pmp22a |
| eppk12 | 6E-107 | 1.3586023 | 0.382 | 0.202 | 1.8E-102 | Stroma 2 | eppk1 |
| neu3.5 | 1E-106 | 1.8347626 | 0.179 | 0.063 | 3.7E-102 | Stroma 2 | neu3.5 |
| pcm12 | 4E-105 | 1.5197628 | 0.3 | 0.144 | 1.3E-100 | Stroma 2 | pcm1 |

| | | | | | | | |
|---|---|---|---|---|---|---|---|
| nrn1a | 5E-104 | 1.4111033 | 0.212 | 0.082 | 1.7E-99 | Stroma 2 | nrn1a |
| dlx4b | 2.4E-97 | 1.7924086 | 0.135 | 0.042 | 7.96E-93 | Stroma 2 | dlx4b |
| sfrp22 | 3.1E-96 | 0.9220901 | 0.584 | 0.387 | 1.02E-91 | Stroma 2 | sfrp2 |
| cyp1c12 | 1.3E-94 | 1.2641604 | 0.227 | 0.096 | 4.22E-90 | Stroma 2 | cyp1c1 |
| sulf2b1 | 2E-93 | 1.2083875 | 0.312 | 0.157 | 6.56E-89 | Stroma 2 | sulf2b |
| si:ch211-198c19.31 | 5.4E-90 | 1.0423623 | 0.37 | 0.203 | 1.74E-85 | Stroma 2 | si:ch211-198c19.3 |
| sfrp51 | 2.5E-88 | 1.0275956 | 0.361 | 0.201 | 8.19E-84 | Stroma 2 | sfrp5 |
| ptges2 | 1E-87 | 1.4185255 | 0.229 | 0.102 | 3.29E-83 | Stroma 2 | ptges |
| ognb2 | 2.4E-84 | 1.2065135 | 0.262 | 0.125 | 7.89E-80 | Stroma 2 | ognb |
| nfia1 | 1.3E-82 | 1.1004988 | 0.309 | 0.162 | 4.15E-78 | Stroma 2 | nfia |
| mfap53 | 2.8E-82 | 0.7324449 | 0.718 | 0.536 | 8.96E-78 | Stroma 2 | mfap5 |
| foxd1 | 2.4E-81 | 1.5223539 | 0.163 | 0.062 | 7.89E-77 | Stroma 2 | foxd1 |
| zgc:1527912 | 1.1E-80 | 1.4656916 | 0.221 | 0.099 | 3.43E-76 | Stroma 2 | zgc:152791 |
| dlx3b2 | 2.4E-80 | 0.7983666 | 0.523 | 0.351 | 7.84E-76 | Stroma 2 | dlx3b |
| clec19a1 | 3.1E-78 | 0.8158167 | 0.157 | 0.059 | 1.01E-73 | Stroma 2 | clec19a |
| igf2b1 | 1.9E-73 | 0.6913404 | 0.594 | 0.428 | 6.34E-69 | Stroma 2 | igf2b |
| asip12 | 1.2E-71 | 0.9363793 | 0.312 | 0.17 | 3.97E-67 | Stroma 2 | asip1 |
| krt42 | 3.3E-71 | 0.3552392 | 0.892 | 0.751 | 1.08E-66 | Stroma 2 | krt4 |
| pbx3b2 | 3.9E-71 | 1.1728043 | 0.264 | 0.137 | 1.27E-66 | Stroma 2 | pbx3b |
| fndc12 | 1.3E-70 | 1.4588543 | 0.292 | 0.161 | 4.38E-66 | Stroma 2 | fndc1 |
| rdh10a | 6.5E-69 | 1.5609057 | 0.108 | 0.036 | 2.1E-64 | Stroma 2 | rdh10a |
| si:ch211-105c13.33 | 2.6E-68 | 0.577306 | 0.711 | 0.562 | 8.35E-64 | Stroma 2 | si:ch211-105c13.3 |
| ablim2 | 5.3E-66 | 1.5686685 | 0.121 | 0.044 | 1.72E-61 | Stroma 2 | ablim2 |
| si:ch73-173h19.33 | 8.7E-66 | 0.7262215 | 0.388 | 0.231 | 2.82E-61 | Stroma 2 | si:ch73-173h19.3 |
| ifitm14 | 1.1E-65 | 0.4039441 | 0.928 | 0.782 | 3.67E-61 | Stroma 2 | ifitm1 |
| loxl12 | 1.3E-65 | 0.9327584 | 0.334 | 0.194 | 4.23E-61 | Stroma 2 | loxl1 |
| ttc363 | 6.4E-64 | 0.8165182 | 0.504 | 0.346 | 2.07E-59 | Stroma 2 | ttc36 |
| mast21 | 1.1E-63 | 0.9953251 | 0.285 | 0.158 | 3.68E-59 | Stroma 2 | mast2 |
| zgc:1951733 | 1.4E-63 | 0.6054165 | 0.552 | 0.388 | 4.5E-59 | Stroma 2 | zgc:195173 |
| ablim1b2 | 5.2E-62 | 0.9437735 | 0.391 | 0.249 | 1.68E-57 | Stroma 2 | ablim1b |
| il11ra3 | 1E-60 | 0.8273509 | 0.375 | 0.233 | 3.26E-56 | Stroma 2 | il11ra |
| gfra1a | 2.2E-60 | 1.5991801 | 0.13 | 0.051 | 7.22E-56 | Stroma 2 | gfra1a |
| hpdb3 | 4.4E-60 | 0.6882433 | 0.681 | 0.499 | 1.44E-55 | Stroma 2 | hpdb |
| adh8a1 | 4.9E-60 | 0.8049381 | 0.342 | 0.204 | 1.58E-55 | Stroma 2 | adh8a |
| tshz12 | 8.5E-60 | 1.1024268 | 0.244 | 0.13 | 2.76E-55 | Stroma 2 | tshz1 |
| add3a2 | 1.3E-57 | 0.7533282 | 0.581 | 0.441 | 4.22E-53 | Stroma 2 | add3a |
| zgc:1749041 | 8E-57 | 0.9934776 | 0.229 | 0.119 | 2.61E-52 | Stroma 2 | zgc:174904 |
| marcksl1a1 | 1.9E-56 | 0.4909418 | 0.809 | 0.702 | 6.17E-52 | Stroma 2 | marcksl1a |
| sost2 | 2.1E-56 | 1.0292265 | 0.163 | 0.074 | 6.76E-52 | Stroma 2 | sost |
| wnt111 | 2.9E-56 | 0.6713814 | 0.428 | 0.276 | 9.29E-52 | Stroma 2 | wnt11 |
| cfd | 1.8E-55 | 1.0463809 | 0.192 | 0.094 | 5.76E-51 | Stroma 2 | cfd |
| ednraa1 | 1.8E-55 | 0.7702391 | 0.342 | 0.207 | 5.9E-51 | Stroma 2 | ednraa |
| prelp1 | 3.2E-53 | 0.9069518 | 0.237 | 0.128 | 1.04E-48 | Stroma 2 | prelp |
| c7a3 | 1.5E-52 | 0.9050031 | 0.287 | 0.169 | 4.92E-48 | Stroma 2 | c7a |
| mxra5a1 | 2.4E-52 | 1.4240662 | 0.156 | 0.073 | 7.95E-48 | Stroma 2 | mxra5a |
| dcn2 | 7.9E-52 | 0.2800602 | 0.956 | 0.863 | 2.55E-47 | Stroma 2 | dcn |
| angptl1a1 | 1.8E-51 | 0.4999099 | 0.539 | 0.385 | 5.73E-47 | Stroma 2 | angptl1a |
| rgs41 | 6.2E-51 | 1.3767669 | 0.146 | 0.066 | 2.02E-46 | Stroma 2 | rgs4 |

| | | | | | | | |
|---|---|---|---|---|---|---|---|
| c1qtnf21 | 1.5E-50 | 1.2060158 | 0.177 | 0.088 | 4.81E-46 | Stroma 2 | c1qtnf2 |
| CR925728.11 | 3.3E-50 | 0.6716379 | 0.182 | 0.089 | 1.07E-45 | Stroma 2 | CR925728.1 |
| tat2 | 9.9E-50 | 0.9488232 | 0.365 | 0.239 | 3.22E-45 | Stroma 2 | tat |
| sod3b2 | 1.7E-49 | 1.117077 | 0.218 | 0.118 | 5.62E-45 | Stroma 2 | sod3b |
| gyg1b3 | 8.6E-49 | 0.8397156 | 0.389 | 0.26 | 2.81E-44 | Stroma 2 | gyg1b |
| angptl2a2 | 1.8E-48 | 0.5855916 | 0.454 | 0.311 | 5.93E-44 | Stroma 2 | angptl2a |
| si:ch211-243a20.31 | 8.7E-48 | 0.7471067 | 0.41 | 0.275 | 2.84E-43 | Stroma 2 | si:ch211-243a20.3 |
| si:ch211-196i2.1 | 1.2E-47 | 1.627471 | 0.117 | 0.05 | 3.8E-43 | Stroma 2 | si:ch211-196i2.1 |
| rgs5b3 | 5.3E-47 | 0.3142496 | 0.654 | 0.516 | 1.71E-42 | Stroma 2 | rgs5b |
| klf2a2 | 1.6E-46 | 0.8309613 | 0.404 | 0.279 | 5.25E-42 | Stroma 2 | klf2a |
| nhsl22 | 1E-45 | 1.0587353 | 0.187 | 0.098 | 3.27E-41 | Stroma 2 | nhsl2 |
| aldh9a1a.13 | 1E-45 | 0.5725475 | 0.594 | 0.475 | 3.38E-41 | Stroma 2 | aldh9a1a.1 |
| lrp1aa1 | 4.4E-45 | 0.4722388 | 0.764 | 0.669 | 1.42E-40 | Stroma 2 | lrp1aa |
| col27a1b2 | 6.4E-45 | 0.5298245 | 0.472 | 0.33 | 2.07E-40 | Stroma 2 | col27a1b |
| CU914776.11 | 1.4E-44 | 0.9096467 | 0.162 | 0.08 | 4.49E-40 | Stroma 2 | CU914776.1 |
| nt5e | 2.5E-44 | 1.3060744 | 0.131 | 0.061 | 8.18E-40 | Stroma 2 | nt5e |
| srgn1 | 4.1E-44 | 0.5107108 | 0.576 | 0.445 | 1.34E-39 | Stroma 2 | srgn |
| clec3ba2 | 7E-44 | 0.6369018 | 0.222 | 0.122 | 2.27E-39 | Stroma 2 | clec3ba |
| aldh1a21 | 7.8E-44 | 0.8870837 | 0.119 | 0.051 | 2.55E-39 | Stroma 2 | aldh1a2 |
| itgb1b.22 | 1.5E-43 | 1.1009454 | 0.196 | 0.107 | 4.83E-39 | Stroma 2 | itgb1b.2 |
| rbp42 | 2.2E-43 | 0.2837377 | 0.982 | 0.906 | 7.29E-39 | Stroma 2 | rbp4 |
| pcxa1 | 4E-43 | 1.0411475 | 0.202 | 0.112 | 1.31E-38 | Stroma 2 | pcxa |
| si:ch1073-291c23.21 | 4.6E-43 | 0.4736079 | 0.598 | 0.468 | 1.51E-38 | Stroma 2 | si:ch1073-291c23.2 |
| frzb2 | 4.9E-43 | 0.9712979 | 0.219 | 0.125 | 1.59E-38 | Stroma 2 | frzb |
| hspb11 | 9.3E-43 | 0.7835378 | 0.282 | 0.174 | 3.02E-38 | Stroma 2 | hspb1 |
| dpysl31 | 1.8E-42 | 0.8240081 | 0.26 | 0.156 | 5.7E-38 | Stroma 2 | dpysl3 |
| si:ch211-250c4.4 | 8.3E-42 | 1.2373223 | 0.1 | 0.041 | 2.69E-37 | Stroma 2 | si:ch211-250c4.4 |
| zbtb16b | 1.1E-41 | 1.0703038 | 0.185 | 0.1 | 3.44E-37 | Stroma 2 | zbtb16b |
| htra1b1 | 4.3E-41 | 0.4492762 | 0.528 | 0.406 | 1.39E-36 | Stroma 2 | htra1b |
| adcyap1b | 4.8E-41 | 0.6025732 | 0.102 | 0.043 | 1.57E-36 | Stroma 2 | adcyap1b |
| cald1b3 | 2.4E-40 | 0.4542641 | 0.27 | 0.162 | 7.72E-36 | Stroma 2 | cald1b |
| fbln21 | 4.1E-40 | 0.7825611 | 0.21 | 0.118 | 1.32E-35 | Stroma 2 | fbln2 |
| edil3a1 | 1E-39 | 0.5016029 | 0.559 | 0.429 | 3.26E-35 | Stroma 2 | edil3a |
| hgd2 | 5.5E-39 | 0.4972415 | 0.557 | 0.423 | 1.79E-34 | Stroma 2 | hgd |
| thbs3a1 | 6.3E-39 | 0.6288333 | 0.423 | 0.303 | 2.06E-34 | Stroma 2 | thbs3a |
| zfp36l1b2 | 1E-38 | 0.3887433 | 0.663 | 0.54 | 3.36E-34 | Stroma 2 | zfp36l1b |
| gstz12 | 1.2E-38 | 0.5428642 | 0.462 | 0.338 | 3.93E-34 | Stroma 2 | gstz1 |
| cd631 | 5.7E-38 | 0.4802109 | 0.867 | 0.825 | 1.87E-33 | Stroma 2 | cd63 |
| vasna1 | 1.9E-37 | 0.8716314 | 0.195 | 0.111 | 6.27E-33 | Stroma 2 | vasna |
| endouc1 | 1.8E-36 | 0.7428854 | 0.338 | 0.231 | 5.82E-32 | Stroma 2 | endouc |
| thbs2a2 | 7.2E-36 | 0.5911273 | 0.304 | 0.198 | 2.34E-31 | Stroma 2 | thbs2a |
| adma1 | 8.3E-36 | 0.6233437 | 0.342 | 0.228 | 2.71E-31 | Stroma 2 | adma |
| tmem1762 | 3.4E-35 | 0.4645985 | 0.506 | 0.388 | 1.11E-30 | Stroma 2 | tmem176 |
| sgip1a1 | 6.4E-35 | 1.1565122 | 0.107 | 0.049 | 2.09E-30 | Stroma 2 | sgip1a |
| thbs2b2 | 8.7E-35 | 0.5320816 | 0.359 | 0.246 | 2.83E-30 | Stroma 2 | thbs2b |
| furinb1 | 9.7E-35 | 1.2601207 | 0.111 | 0.052 | 3.16E-30 | Stroma 2 | furinb |
| krt153 | 1E-34 | 0.3176715 | 0.798 | 0.696 | 3.24E-30 | Stroma 2 | krt15 |
| tmsb22 | 1.5E-34 | 0.9275664 | 0.17 | 0.093 | 4.73E-30 | Stroma 2 | tmsb2 |

| | | | | | | | |
|---|---|---|---|---|---|---|---|
| im:71523481 | 2.3E-34 | 0.5328567 | 0.34 | 0.231 | 7.57E-30 | Stroma 2 | im:7152348 |
| zgc:565853 | 3.3E-34 | 0.5436015 | 0.441 | 0.322 | 1.06E-29 | Stroma 2 | zgc:56585 |
| sulf11 | 3.4E-34 | 0.767864 | 0.232 | 0.142 | 1.11E-29 | Stroma 2 | sulf1 |
| bhmt3 | 4.7E-34 | 0.4285084 | 0.833 | 0.701 | 1.51E-29 | Stroma 2 | bhmt |
| plpp1a3 | 5.9E-34 | 0.4463614 | 0.489 | 0.369 | 1.91E-29 | Stroma 2 | plpp1a |
| si:ch73-380l3.23 | 6.8E-34 | 0.4557577 | 0.537 | 0.41 | 2.21E-29 | Stroma 2 | si:ch73-380l3.2 |
| vat12 | 9.8E-34 | 0.4205431 | 0.426 | 0.31 | 3.17E-29 | Stroma 2 | vat1 |
| meis2a.1 | 1.9E-33 | 0.7934057 | 0.214 | 0.129 | 6.05E-29 | Stroma 2 | meis2a.1 |
| gchfr1 | 1.9E-33 | 0.7892384 | 0.25 | 0.158 | 6.3E-29 | Stroma 2 | gchfr |
| barx11 | 2E-33 | 0.7260068 | 0.209 | 0.124 | 6.53E-29 | Stroma 2 | barx1 |
| scara31 | 7.9E-33 | 0.50762 | 0.218 | 0.13 | 2.55E-28 | Stroma 2 | scara3 |
| si:ch211-195b13.11 | 8.5E-33 | 0.5796959 | 0.341 | 0.238 | 2.76E-28 | Stroma 2 | si:ch211-195b13.1 |
| b2m3 | 1.2E-32 | 0.3032933 | 0.779 | 0.681 | 4.02E-28 | Stroma 2 | b2m |
| entpd12 | 1.3E-32 | 1.0779082 | 0.146 | 0.079 | 4.17E-28 | Stroma 2 | entpd1 |
| fgl11 | 3.9E-32 | 0.5550648 | 0.306 | 0.205 | 1.26E-27 | Stroma 2 | fgl1 |
| tnfsf10l2 | 4.4E-32 | 0.6175879 | 0.318 | 0.217 | 1.42E-27 | Stroma 2 | tnfsf10l |
| atp1b3b1 | 7.1E-32 | 1.113304 | 0.141 | 0.076 | 2.3E-27 | Stroma 2 | atp1b3b |
| wu:fj16a033 | 3.4E-31 | 0.4876773 | 0.283 | 0.187 | 1.11E-26 | Stroma 2 | wu:fj16a03 |
| b2ml1 | 1.2E-30 | 0.2286515 | 0.932 | 0.896 | 3.77E-26 | Stroma 2 | b2ml |
| sult1st6 | 1.6E-30 | 0.8149613 | 0.19 | 0.113 | 5.26E-26 | Stroma 2 | sult1st6 |
| adam10a2 | 1.8E-30 | 0.4927132 | 0.478 | 0.377 | 5.93E-26 | Stroma 2 | adam10a |
| ctsk3 | 5.5E-30 | 0.4514094 | 0.669 | 0.589 | 1.8E-25 | Stroma 2 | ctsk |
| col4a11 | 9.3E-30 | 0.3989432 | 0.322 | 0.222 | 3.03E-25 | Stroma 2 | col4a1 |
| nr2f1a | 1.5E-29 | 0.7257811 | 0.232 | 0.148 | 5.02E-25 | Stroma 2 | nr2f1a |
| znf5031 | 2E-29 | 0.8469354 | 0.183 | 0.11 | 6.48E-25 | Stroma 2 | znf503 |
| colec122 | 2.2E-29 | 0.3871291 | 0.545 | 0.433 | 7.18E-25 | Stroma 2 | colec12 |
| c61 | 2.7E-29 | 0.880993 | 0.131 | 0.07 | 8.8E-25 | Stroma 2 | c6 |
| raraa3 | 3.5E-29 | 0.6046686 | 0.348 | 0.249 | 1.14E-24 | Stroma 2 | raraa |
| plpp32 | 7.3E-29 | 0.4038947 | 0.563 | 0.449 | 2.36E-24 | Stroma 2 | plpp3 |
| ccl44 | 7.7E-29 | 0.871803 | 0.203 | 0.124 | 2.49E-24 | Stroma 2 | ccl44 |
| cygb13 | 1.5E-28 | 0.4125575 | 0.524 | 0.417 | 4.86E-24 | Stroma 2 | cygb1 |
| aldob3 | 5.2E-28 | 0.5348758 | 0.406 | 0.306 | 1.71E-23 | Stroma 2 | aldob |
| igfbp1a2 | 1.9E-27 | 1.6299264 | 0.126 | 0.069 | 6.1E-23 | Stroma 2 | igfbp1a |
| meis1b3 | 2.8E-27 | 0.4687119 | 0.421 | 0.319 | 9.11E-23 | Stroma 2 | meis1b |
| hic1 | 5.1E-27 | 0.7955244 | 0.118 | 0.062 | 1.65E-22 | Stroma 2 | hic1 |
| hhla2a.1 | 2.2E-26 | 0.9394773 | 0.139 | 0.079 | 7.05E-22 | Stroma 2 | hhla2a.1 |
| postna4 | 9.1E-26 | 0.2389518 | 0.735 | 0.583 | 2.96E-21 | Stroma 2 | postna |
| dock9b2 | 1.1E-25 | 0.8631658 | 0.162 | 0.097 | 3.57E-21 | Stroma 2 | dock9b |
| qdpra2 | 1.7E-25 | 0.4403037 | 0.54 | 0.42 | 5.49E-21 | Stroma 2 | qdpra |
| anos1b | 2.2E-25 | 0.9273258 | 0.131 | 0.073 | 7.3E-21 | Stroma 2 | anos1b |
| ssh2a | 7.9E-25 | 0.895768 | 0.136 | 0.077 | 2.56E-20 | Stroma 2 | ssh2a |
| tgfbi | 1.2E-24 | 0.343242 | 0.285 | 0.196 | 3.87E-20 | Stroma 2 | tgfbi |
| clic2 | 1.5E-24 | 0.3097074 | 0.336 | 0.243 | 4.73E-20 | Stroma 2 | clic2 |
| si:ch211-106h4.121 | 4.4E-24 | 0.8468435 | 0.561 | 0.453 | 1.44E-19 | Stroma 2 | si:ch211-106h4.12 |
| cxcl18b1 | 5E-24 | 0.4167421 | 0.769 | 0.699 | 1.61E-19 | Stroma 2 | cxcl18b |
| ms4a17a.111 | 6.3E-24 | 0.5639246 | 0.332 | 0.248 | 2.03E-19 | Stroma 2 | ms4a17a.11 |
| gsta.12 | 7.7E-24 | 0.3837923 | 0.449 | 0.353 | 2.51E-19 | Stroma 2 | gsta.1 |
| alx4b1 | 9.6E-24 | 0.9076968 | 0.105 | 0.056 | 3.11E-19 | Stroma 2 | alx4b |

| | | | | | | | |
|---|---|---|---|---|---|---|---|
| fstl31 | 1E-23 | 0.2990799 | 0.423 | 0.317 | 3.27E-19 | Stroma 2 | fstl3 |
| slco2b13 | 1.1E-23 | 0.5643163 | 0.305 | 0.217 | 3.46E-19 | Stroma 2 | slco2b1 |
| pcolcea3 | 1.2E-23 | 0.3310462 | 0.656 | 0.564 | 3.82E-19 | Stroma 2 | pcolcea |
| sptan11 | 1.2E-23 | 0.5343163 | 0.328 | 0.24 | 4.05E-19 | Stroma 2 | sptan1 |
| CR318588.41 | 1.8E-23 | 0.2247325 | 0.994 | 0.986 | 5.7E-19 | Stroma 2 | CR318588.4 |
| thbs1a | 2.2E-23 | 0.7841807 | 0.389 | 0.309 | 7.04E-19 | Stroma 2 | thbs1a |
| pdgfrl1 | 2.9E-23 | 0.3573197 | 0.659 | 0.581 | 9.29E-19 | Stroma 2 | pdgfrl |
| plscr3b1 | 3E-23 | 0.49199 | 0.323 | 0.236 | 9.61E-19 | Stroma 2 | plscr3b |
| si:dkey-240h12.42 | 3.8E-23 | 0.6902733 | 0.182 | 0.115 | 1.24E-18 | Stroma 2 | si:dkey-240h12.4 |
| hmox1a1 | 7E-23 | 0.5436441 | 0.248 | 0.17 | 2.28E-18 | Stroma 2 | hmox1a |
| akap12b1 | 1.1E-22 | 0.6346706 | 0.211 | 0.14 | 3.51E-18 | Stroma 2 | akap12b |
| irx5a1 | 2.3E-22 | 0.8091818 | 0.146 | 0.088 | 7.4E-18 | Stroma 2 | irx5a |
| crabp2b1 | 2.8E-22 | 0.5182615 | 0.495 | 0.416 | 9.06E-18 | Stroma 2 | crabp2b |
| adamts22 | 3.5E-22 | 0.3612902 | 0.467 | 0.369 | 1.15E-17 | Stroma 2 | adamts2 |
| si:dkey-16l2.20 | 5.1E-22 | 0.9826505 | 0.11 | 0.061 | 1.64E-17 | Stroma 2 | si:dkey-16l2.20 |
| cilp21 | 7.4E-22 | 0.5319323 | 0.109 | 0.06 | 2.4E-17 | Stroma 2 | cilp2 |
| ccn11 | 7.6E-22 | 0.4756967 | 0.697 | 0.64 | 2.47E-17 | Stroma 2 | ccn1 |
| col4a21 | 8.4E-22 | 0.205324 | 0.301 | 0.215 | 2.72E-17 | Stroma 2 | col4a2 |
| pdgfra1 | 9.2E-22 | 0.304841 | 0.558 | 0.455 | 3E-17 | Stroma 2 | pdgfra |
| gstm.12 | 9.5E-22 | 0.2823409 | 0.68 | 0.593 | 3.11E-17 | Stroma 2 | gstm.1 |
| ms4a17a.71 | 1.2E-21 | 0.3992577 | 0.383 | 0.292 | 3.78E-17 | Stroma 2 | ms4a17a.7 |
| ptgdsb.13 | 2.4E-21 | 0.2670755 | 0.569 | 0.458 | 7.91E-17 | Stroma 2 | ptgdsb.1 |
| si:dkey-16p21.82 | 3.1E-21 | 0.4605455 | 0.289 | 0.206 | 9.98E-17 | Stroma 2 | si:dkey-16p21.8 |
| sptbn11 | 5.9E-21 | 0.4438891 | 0.328 | 0.246 | 1.93E-16 | Stroma 2 | sptbn1 |
| ecm22 | 9.8E-21 | 0.4369798 | 0.381 | 0.294 | 3.18E-16 | Stroma 2 | ecm2 |
| hsd11b22 | 1.2E-20 | 0.8378331 | 0.146 | 0.091 | 3.77E-16 | Stroma 2 | hsd11b2 |
| ryr32 | 3.4E-20 | 0.7463475 | 0.142 | 0.087 | 1.11E-15 | Stroma 2 | ryr3 |
| twist23 | 3.7E-20 | 0.3125087 | 0.432 | 0.336 | 1.2E-15 | Stroma 2 | twist2 |
| dlx4a4 | 3.8E-20 | 0.550448 | 0.264 | 0.19 | 1.24E-15 | Stroma 2 | dlx4a |
| pde3b | 4.2E-20 | 0.6251999 | 0.18 | 0.118 | 1.38E-15 | Stroma 2 | pde3b |
| CR384075.11 | 6.7E-20 | 0.9741597 | 0.1 | 0.055 | 2.19E-15 | Stroma 2 | CR384075.1 |
| celf23 | 7.9E-20 | 0.523101 | 0.27 | 0.194 | 2.57E-15 | Stroma 2 | celf2 |
| ramp22 | 1E-19 | 0.6238695 | 0.206 | 0.139 | 3.35E-15 | Stroma 2 | ramp2 |
| icam31 | 1.6E-19 | 0.6654225 | 0.133 | 0.08 | 5.25E-15 | Stroma 2 | icam3 |
| pdgfrb1 | 1.8E-19 | 0.3639233 | 0.494 | 0.408 | 5.87E-15 | Stroma 2 | pdgfrb |
| tspan4b2 | 2.1E-19 | 0.3849709 | 0.422 | 0.332 | 6.87E-15 | Stroma 2 | tspan4b |
| lepa1 | 6.9E-19 | 1.4369553 | 0.102 | 0.059 | 2.24E-14 | Stroma 2 | lepa |
| zeb1b1 | 8.3E-19 | 0.9021712 | 0.137 | 0.086 | 2.7E-14 | Stroma 2 | zeb1b |
| f3b2 | 1.2E-18 | 0.5877914 | 0.455 | 0.376 | 4.05E-14 | Stroma 2 | f3b |
| GNAZ2 | 1.3E-18 | 0.6121304 | 0.195 | 0.132 | 4.32E-14 | Stroma 2 | GNAZ |
| bhlhe40 | 1.4E-18 | 0.4994177 | 0.531 | 0.462 | 4.54E-14 | Stroma 2 | bhlhe40 |
| lama41 | 1.7E-18 | 0.3854471 | 0.347 | 0.267 | 5.55E-14 | Stroma 2 | lama4 |
| aff22 | 1.9E-18 | 0.5212812 | 0.246 | 0.176 | 6.06E-14 | Stroma 2 | aff2 |
| chd52 | 2.3E-18 | 0.6533111 | 0.185 | 0.124 | 7.52E-14 | Stroma 2 | chd5 |
| elmsan1b1 | 2.6E-18 | 0.3977678 | 0.431 | 0.353 | 8.36E-14 | Stroma 2 | elmsan1b |
| nfixb2 | 2.7E-18 | 0.3928892 | 0.37 | 0.291 | 8.83E-14 | Stroma 2 | nfixb |
| si:ch73-46j18.51 | 3.1E-18 | 0.4053339 | 0.654 | 0.6 | 1.02E-13 | Stroma 2 | si:ch73-46j18.5 |
| plxna4 | 3.2E-18 | 0.8010549 | 0.101 | 0.057 | 1.05E-13 | Stroma 2 | plxna4 |

| | | | | | | | |
|---|---|---|---|---|---|---|---|
| scara53 | 3.4E-18 | 0.3958867 | 0.217 | 0.149 | 1.11E-13 | Stroma 2 | scara5 |
| si:ch211-212k18.7 | 4.2E-18 | 0.8544909 | 0.106 | 0.062 | 1.36E-13 | Stroma 2 | si:ch211-212k18.7 |
| gpt2l2 | 5.4E-18 | 0.6104401 | 0.187 | 0.127 | 1.75E-13 | Stroma 2 | gpt2l |
| specc1la | 5.4E-18 | 0.6861811 | 0.127 | 0.077 | 1.76E-13 | Stroma 2 | specc1la |
| tnfaip2a1 | 9.3E-18 | 0.6943776 | 0.137 | 0.086 | 3.01E-13 | Stroma 2 | tnfaip2a |
| si:dkey-11f4.203 | 1.2E-17 | 0.3188765 | 0.573 | 0.484 | 3.99E-13 | Stroma 2 | si:dkey-11f4.20 |
| ghrb3 | 1.3E-17 | 0.5506046 | 0.162 | 0.106 | 4.32E-13 | Stroma 2 | ghrb |
| fzd2 | 1.4E-17 | 0.7767342 | 0.149 | 0.096 | 4.44E-13 | Stroma 2 | fzd2 |
| col6a11 | 1.5E-17 | 0.2512328 | 0.93 | 0.872 | 4.87E-13 | Stroma 2 | col6a1 |
| plxdc2 | 2.3E-17 | 0.4473573 | 0.222 | 0.155 | 7.41E-13 | Stroma 2 | plxdc2 |
| si:ch211-183d21.12 | 3.2E-17 | 0.4685256 | 0.443 | 0.379 | 1.05E-12 | Stroma 2 | si:ch211-183d21.1 |
| slc51a3 | 3.3E-17 | 0.341629 | 0.224 | 0.156 | 1.06E-12 | Stroma 2 | slc51a |
| zcchc243 | 5E-17 | 0.4461971 | 0.328 | 0.253 | 1.62E-12 | Stroma 2 | zcchc24 |
| capn2b3 | 5E-17 | 0.3086025 | 0.403 | 0.318 | 1.63E-12 | Stroma 2 | capn2b |
| mhc1zba1 | 5.7E-17 | 0.2547459 | 0.658 | 0.58 | 1.84E-12 | Stroma 2 | mhc1zba |
| jak11 | 9.9E-17 | 0.3145509 | 0.506 | 0.434 | 3.2E-12 | Stroma 2 | jak1 |
| fstb3 | 3E-16 | 0.2161076 | 0.455 | 0.357 | 9.8E-12 | Stroma 2 | fstb |
| ptprk1 | 4.1E-16 | 0.7142475 | 0.111 | 0.068 | 1.34E-11 | Stroma 2 | ptprk |
| runx1t1 | 7E-16 | 0.5643969 | 0.139 | 0.089 | 2.29E-11 | Stroma 2 | runx1t1 |
| mocs21 | 9E-16 | 0.7627173 | 0.133 | 0.086 | 2.91E-11 | Stroma 2 | mocs2 |
| tgm1l12 | 1.3E-15 | 0.5586613 | 0.181 | 0.125 | 4.29E-11 | Stroma 2 | tgm1l1 |
| cyba2 | 1.7E-15 | 0.822894 | 0.219 | 0.161 | 5.37E-11 | Stroma 2 | cyba |
| epn21 | 2.1E-15 | 0.5467118 | 0.159 | 0.107 | 6.97E-11 | Stroma 2 | epn2 |
| BX936337.1 | 2.2E-15 | 0.3708506 | 0.449 | 0.386 | 7.21E-11 | Stroma 2 | BX936337.1 |
| LO018188.13 | 3E-15 | 0.3220781 | 0.545 | 0.475 | 9.62E-11 | Stroma 2 | LO018188.1 |
| cbx7a | 3.4E-15 | 0.4141638 | 0.416 | 0.343 | 1.1E-10 | Stroma 2 | cbx7a |
| CR339041.1 | 3.4E-15 | 0.4965437 | 0.146 | 0.096 | 1.11E-10 | Stroma 2 | CR339041.1 |
| si:cabz01069012.21 | 4.1E-15 | 0.6316295 | 0.151 | 0.101 | 1.33E-10 | Stroma 2 | si:cabz01069012.2 |
| BX663503.33 | 7E-15 | 0.2025391 | 0.49 | 0.398 | 2.26E-10 | Stroma 2 | BX663503.3 |
| KCTD152 | 7.7E-15 | 0.3562942 | 0.411 | 0.339 | 2.51E-10 | Stroma 2 | KCTD15 |
| ifi46 | 1E-14 | 0.6271739 | 0.141 | 0.093 | 3.38E-10 | Stroma 2 | ifi46 |
| rbp52 | 1.1E-14 | 0.6190861 | 0.167 | 0.116 | 3.74E-10 | Stroma 2 | rbp5 |
| zfp36l1a3 | 1.2E-14 | 0.4325179 | 0.268 | 0.204 | 4.05E-10 | Stroma 2 | zfp36l1a |
| nccrp1 | 1.4E-14 | 0.4341762 | 0.338 | 0.273 | 4.63E-10 | Stroma 2 | nccrp1 |
| ITLN14 | 1.4E-14 | 0.3709668 | 0.275 | 0.208 | 4.68E-10 | Stroma 2 | ITLN1 |
| ak42 | 1.7E-14 | 0.5003534 | 0.242 | 0.182 | 5.37E-10 | Stroma 2 | ak4 |
| hdlbpa2 | 2.9E-14 | 0.238514 | 0.727 | 0.673 | 9.55E-10 | Stroma 2 | hdlbpa |
| tsku2 | 3E-14 | 0.3664212 | 0.387 | 0.316 | 9.82E-10 | Stroma 2 | tsku |
| slc25a482 | 3.7E-14 | 0.5584224 | 0.19 | 0.136 | 1.21E-09 | Stroma 2 | slc25a48 |
| tcima2 | 4.7E-14 | 0.8024036 | 0.282 | 0.224 | 1.52E-09 | Stroma 2 | tcima |
| lrp52 | 4.7E-14 | 0.6654624 | 0.158 | 0.11 | 1.53E-09 | Stroma 2 | lrp5 |
| DAB23 | 7.2E-14 | 0.4219684 | 0.434 | 0.376 | 2.36E-09 | Stroma 2 | DAB2 |
| ehbp1l1b | 1.1E-13 | 0.5447233 | 0.176 | 0.125 | 3.7E-09 | Stroma 2 | ehbp1l1b |
| si:dkey-172h23.21 | 1.4E-13 | 0.4805369 | 0.197 | 0.145 | 4.58E-09 | Stroma 2 | si:dkey-172h23.2 |
| cwc272 | 1.5E-13 | 0.6294741 | 0.17 | 0.121 | 5.02E-09 | Stroma 2 | cwc27 |
| CU984600.13 | 2.1E-13 | 0.4177207 | 0.225 | 0.168 | 6.74E-09 | Stroma 2 | CU984600.1 |
| creg21 | 2.1E-13 | 0.3511657 | 0.387 | 0.322 | 6.98E-09 | Stroma 2 | creg2 |
| nr2f2 | 5E-13 | 0.4989883 | 0.162 | 0.113 | 1.64E-08 | Stroma 2 | nr2f2 |

| | | | | | | | |
|---|---|---|---|---|---|---|---|
| twist31 | 5.8E-13 | 0.2591637 | 0.322 | 0.25 | 1.9E-08 | Stroma 2 | twist3 |
| nid1b | 7.6E-13 | 0.5445728 | 0.119 | 0.078 | 2.46E-08 | Stroma 2 | nid1b |
| itgb22 | 1.2E-12 | 0.479052 | 0.151 | 0.104 | 3.9E-08 | Stroma 2 | itgb2 |
| ENSDARG00000090 8 | 1.4E-12 | 0.7111425 | 0.131 | 0.089 | 4.62E-08 | Stroma 2 | ENSDARG00000090 |
| nfe2l2a2 | 1.6E-12 | 0.3673224 | 0.408 | 0.341 | 5.34E-08 | Stroma 2 | nfe2l2a |
| ezh11 | 2.5E-12 | 0.4515362 | 0.19 | 0.139 | 8.23E-08 | Stroma 2 | ezh1 |
| ch25hl32 | 3E-12 | 0.6317122 | 0.149 | 0.105 | 9.6E-08 | Stroma 2 | ch25hl3 |
| si:dkey-40c11.23 | 3.6E-12 | 0.2562074 | 0.385 | 0.314 | 1.17E-07 | Stroma 2 | si:dkey-40c11.2 |
| pah2 | 3.8E-12 | 0.6015275 | 0.405 | 0.348 | 1.22E-07 | Stroma 2 | pah |
| dennd4a1 | 3.8E-12 | 0.5067012 | 0.126 | 0.085 | 1.24E-07 | Stroma 2 | dennd4a |
| abca1a1 | 3.9E-12 | 0.3642932 | 0.253 | 0.195 | 1.28E-07 | Stroma 2 | abca1a |
| rnd3a3 | 4.7E-12 | 0.3365119 | 0.349 | 0.283 | 1.53E-07 | Stroma 2 | rnd3a |
| ahr22 | 5.4E-12 | 0.3377311 | 0.441 | 0.381 | 1.77E-07 | Stroma 2 | ahr2 |
| ms4a17a.93 | 6.5E-12 | 0.4134162 | 0.27 | 0.214 | 2.1E-07 | Stroma 2 | ms4a17a.9 |
| pear12 | 6.5E-12 | 0.5119261 | 0.176 | 0.129 | 2.11E-07 | Stroma 2 | pear1 |
| slc16a102 | 1.4E-11 | 0.4604312 | 0.179 | 0.131 | 4.63E-07 | Stroma 2 | slc16a10 |
| rhogb | 1.6E-11 | 0.5905211 | 0.105 | 0.069 | 5.19E-07 | Stroma 2 | rhogb |
| tnrc182 | 2.3E-11 | 0.2284906 | 0.47 | 0.404 | 7.59E-07 | Stroma 2 | tnrc18 |
| nck1a1 | 3.5E-11 | 0.6407747 | 0.115 | 0.078 | 1.15E-06 | Stroma 2 | nck1a |
| si:dkey-238i5.21 | 3.6E-11 | 0.6115691 | 0.11 | 0.074 | 1.16E-06 | Stroma 2 | si:dkey-238i5.2 |
| dhrs13a.11 | 3.8E-11 | 0.5636382 | 0.142 | 0.101 | 1.24E-06 | Stroma 2 | dhrs13a.1 |
| asap31 | 6.7E-11 | 0.4337209 | 0.17 | 0.124 | 2.18E-06 | Stroma 2 | asap3 |
| svila2 | 2.1E-10 | 0.4840095 | 0.192 | 0.146 | 6.7E-06 | Stroma 2 | svila |
| cspg5a | 2.4E-10 | 0.3767221 | 0.112 | 0.076 | 7.8E-06 | Stroma 2 | cspg5a |
| si:ch1073-392o20.22 | 2.7E-10 | 0.3731983 | 0.294 | 0.24 | 8.63E-06 | Stroma 2 | si:ch1073-392o20.2 |
| cnn3a2 | 2.7E-10 | 0.2044389 | 0.536 | 0.474 | 8.77E-06 | Stroma 2 | cnn3a |
| fam107b1 | 2.7E-10 | 0.3732896 | 0.261 | 0.207 | 8.93E-06 | Stroma 2 | fam107b |
| dhrs93 | 2.9E-10 | 0.4651656 | 0.185 | 0.139 | 9.47E-06 | Stroma 2 | dhrs9 |
| zgc:1103391 | 4E-10 | 0.3644808 | 0.265 | 0.214 | 1.29E-05 | Stroma 2 | zgc:110339 |
| si:dkey-7j14.5 | 4.1E-10 | 0.5105976 | 0.103 | 0.069 | 1.33E-05 | Stroma 2 | si:dkey-7j14.5 |
| cpn12 | 4.1E-10 | 0.4042496 | 0.26 | 0.21 | 1.35E-05 | Stroma 2 | cpn1 |
| prss232 | 5.1E-10 | 0.4582403 | 0.246 | 0.198 | 1.65E-05 | Stroma 2 | prss23 |
| arrdc3a1 | 7.4E-10 | 0.4350329 | 0.209 | 0.163 | 2.39E-05 | Stroma 2 | arrdc3a |
| pepd | 1E-09 | 0.3612555 | 0.261 | 0.211 | 3.34E-05 | Stroma 2 | pepd |
| eif4ebp31 | 1.1E-09 | 0.3303275 | 0.219 | 0.172 | 3.52E-05 | Stroma 2 | eif4ebp3 |
| neu3.22 | 1.3E-09 | 0.212916 | 0.278 | 0.22 | 4.16E-05 | Stroma 2 | neu3.2 |
| ywhag13 | 1.5E-09 | 0.5125565 | 0.148 | 0.109 | 4.79E-05 | Stroma 2 | ywhag1 |
| tuba1b2 | 1.5E-09 | 0.2295263 | 0.365 | 0.306 | 5.03E-05 | Stroma 2 | tuba1b |
| chka3 | 1.7E-09 | 0.5012974 | 0.125 | 0.089 | 5.56E-05 | Stroma 2 | chka |
| scamp41 | 2.1E-09 | 0.4740112 | 0.111 | 0.077 | 6.83E-05 | Stroma 2 | scamp4 |
| thbs4a1 | 2.3E-09 | 0.2028608 | 0.113 | 0.078 | 7.42E-05 | Stroma 2 | thbs4a |
| ifi451 | 2.9E-09 | 0.4023465 | 0.274 | 0.227 | 9.38E-05 | Stroma 2 | ifi45 |
| vwa1 | 3.5E-09 | 0.2833539 | 0.18 | 0.137 | 0.000112 | Stroma 2 | vwa1 |
| stom1 | 3.7E-09 | 0.3391088 | 0.32 | 0.268 | 0.000121 | Stroma 2 | stom |
| arhgdig | 5.1E-09 | 0.5231373 | 0.112 | 0.079 | 0.000167 | Stroma 2 | arhgdig |
| amotl2a | 5.5E-09 | 0.4644127 | 0.199 | 0.155 | 0.00018 | Stroma 2 | amotl2a |
| tgfbr2b2 | 5.8E-09 | 0.6037878 | 0.181 | 0.142 | 0.00019 | Stroma 2 | tgfbr2b |
| si:dkey-188i13.101 | 6.4E-09 | 0.2951683 | 0.298 | 0.246 | 0.000207 | Stroma 2 | si:dkey-188i13.10 |

| | | | | | | | |
|---|---|---|---|---|---|---|---|
| zgc:1533722 | 6.7E-09 | 0.6410959 | 0.119 | 0.086 | 0.000219 | Stroma 2 | zgc:153372 |
| hmcn11 | 7.2E-09 | 0.5438667 | 0.102 | 0.071 | 0.000234 | Stroma 2 | hmcn1 |
| ldhba1 | 7.5E-09 | 0.3322241 | 0.528 | 0.493 | 0.000243 | Stroma 2 | ldhba |
| zgc:173587 | 8.4E-09 | 0.4972927 | 0.17 | 0.13 | 0.000273 | Stroma 2 | zgc:173587 |
| peli1a2 | 8.7E-09 | 0.4703179 | 0.167 | 0.127 | 0.000284 | Stroma 2 | peli1a |
| itm2bb1 | 9.4E-09 | 0.4212915 | 0.123 | 0.088 | 0.000305 | Stroma 2 | itm2bb |
| antxr2a2 | 1E-08 | 0.2465579 | 0.386 | 0.333 | 0.000326 | Stroma 2 | antxr2a |
| asah1b1 | 1E-08 | 0.4150931 | 0.228 | 0.183 | 0.000333 | Stroma 2 | asah1b |
| fxyd6l1 | 1.1E-08 | 0.2863743 | 0.304 | 0.252 | 0.00035 | Stroma 2 | fxyd6l |
| gli3 | 1.2E-08 | 0.4540933 | 0.147 | 0.109 | 0.000393 | Stroma 2 | gli3 |
| mab21l12 | 1.5E-08 | 0.4715283 | 0.178 | 0.138 | 0.000489 | Stroma 2 | mab21l1 |
| nrp1a1 | 2.3E-08 | 0.3361661 | 0.17 | 0.129 | 0.000733 | Stroma 2 | nrp1a |
| sntb21 | 2.3E-08 | 0.3038393 | 0.284 | 0.235 | 0.000743 | Stroma 2 | sntb2 |
| slit31 | 2.4E-08 | 0.3222726 | 0.246 | 0.2 | 0.000795 | Stroma 2 | slit3 |
| gpx1a2 | 2.6E-08 | 0.3772561 | 0.235 | 0.191 | 0.000834 | Stroma 2 | gpx1a |
| tlr5b1 | 3E-08 | 0.5272782 | 0.103 | 0.073 | 0.000965 | Stroma 2 | tlr5b |
| si:ch211-212k18.92 | 3.4E-08 | 0.3146175 | 0.221 | 0.176 | 0.00112 | Stroma 2 | si:ch211-212k18.9 |
| gadd45ga1 | 4.2E-08 | 0.259042 | 0.327 | 0.275 | 0.001377 | Stroma 2 | gadd45ga |
| cebpa2 | 4.4E-08 | 0.3822598 | 0.216 | 0.174 | 0.001427 | Stroma 2 | cebpa |
| spry4 | 5.1E-08 | 0.2830906 | 0.22 | 0.177 | 0.001669 | Stroma 2 | spry4 |
| golim4b1 | 6E-08 | 0.9141524 | 0.14 | 0.108 | 0.001959 | Stroma 2 | golim4b |
| CPNE82 | 6.5E-08 | 0.2094357 | 0.337 | 0.284 | 0.002111 | Stroma 2 | CPNE8 |
| plekho22 | 6.6E-08 | 0.3902214 | 0.224 | 0.181 | 0.002142 | Stroma 2 | plekho2 |
| parp41 | 7.3E-08 | 0.3000799 | 0.115 | 0.083 | 0.002371 | Stroma 2 | parp4 |
| eef1db3 | 8.9E-08 | 0.2162357 | 0.612 | 0.58 | 0.002881 | Stroma 2 | eef1db |
| man2b1 | 9.4E-08 | 0.4353196 | 0.129 | 0.096 | 0.003049 | Stroma 2 | man2b1 |
| lpl2 | 9.8E-08 | 0.2238557 | 0.148 | 0.111 | 0.00318 | Stroma 2 | lpl |
| cilp1 | 1.1E-07 | 1.0324212 | 0.208 | 0.171 | 0.003683 | Stroma 2 | cilp |
| lgals9l31 | 1.1E-07 | 0.3722777 | 0.158 | 0.121 | 0.00369 | Stroma 2 | lgals9l3 |
| sult6b12 | 1.2E-07 | 0.3265274 | 0.217 | 0.176 | 0.00404 | Stroma 2 | sult6b1 |
| h1-102 | 1.3E-07 | 0.2552786 | 0.249 | 0.203 | 0.004076 | Stroma 2 | h1-10 |
| il6st1 | 2E-07 | 0.2119869 | 0.463 | 0.422 | 0.006516 | Stroma 2 | il6st |
| gas1a2 | 2.2E-07 | 0.3313157 | 0.198 | 0.159 | 0.007051 | Stroma 2 | gas1a |
| scarb2a1 | 2.3E-07 | 0.3520917 | 0.227 | 0.187 | 0.007611 | Stroma 2 | scarb2a |
| ddit31 | 2.8E-07 | 0.213262 | 0.61 | 0.581 | 0.009004 | Stroma 2 | ddit3 |
| dlx5a1 | 3.9E-07 | 0.2510324 | 0.341 | 0.297 | 0.012707 | Stroma 2 | dlx5a |
| ms4a17a.83 | 4E-07 | 0.3475246 | 0.146 | 0.112 | 0.012889 | Stroma 2 | ms4a17a.8 |
| polr2a1 | 4.2E-07 | 0.2200527 | 0.409 | 0.361 | 0.013529 | Stroma 2 | polr2a |
| prickle1b2 | 5.2E-07 | 0.2866021 | 0.217 | 0.177 | 0.01687 | Stroma 2 | prickle1b |
| fgl2a2 | 5.8E-07 | 0.2438851 | 0.197 | 0.157 | 0.01889 | Stroma 2 | fgl2a |
| sema4ba | 6.6E-07 | 0.4222647 | 0.112 | 0.083 | 0.021403 | Stroma 2 | sema4ba |
| arhgap35a1 | 7.5E-07 | 0.3376344 | 0.219 | 0.179 | 0.024375 | Stroma 2 | arhgap35a |
| cbr1l1 | 8.6E-07 | 0.3533687 | 0.253 | 0.213 | 0.028097 | Stroma 2 | cbr1l |
| ahcy3 | 8.7E-07 | 0.3855637 | 0.22 | 0.183 | 0.02833 | Stroma 2 | ahcy |
| ccdc85b2 | 9.2E-07 | 0.3127957 | 0.229 | 0.188 | 0.029828 | Stroma 2 | ccdc85b |
| wls1 | 1E-06 | 0.201175 | 0.436 | 0.393 | 0.032564 | Stroma 2 | wls |
| fcho2 | 1.1E-06 | 0.5114712 | 0.192 | 0.158 | 0.036732 | Stroma 2 | fcho2 |
| adprh | 1.2E-06 | 0.4165265 | 0.107 | 0.08 | 0.037652 | Stroma 2 | adprh |

| | | | | | | | |
|---|---|---|---|---|---|---|---|
| irf9 | 1.4E-06 | 0.3248918 | 0.161 | 0.127 | 0.044598 | Stroma 2 | irf9 |
| gstt1b2 | 1.4E-06 | 0.2655979 | 0.154 | 0.121 | 0.046809 | Stroma 2 | gstt1b |
| lhfpl2a2 | 1.5E-06 | 0.3997823 | 0.137 | 0.106 | 0.050395 | Stroma 2 | lhfpl2a |
| rasgef1bb1 | 2E-06 | 0.3204267 | 0.19 | 0.154 | 0.064336 | Stroma 2 | rasgef1bb |
| serpina10a1 | 2.3E-06 | 0.229105 | 0.422 | 0.377 | 0.073574 | Stroma 2 | serpina10a |
| pdlim1 | 2.4E-06 | 0.3068783 | 0.267 | 0.229 | 0.077331 | Stroma 2 | pdlim1 |
| si:ch211-281l24.31 | 2.4E-06 | 0.2507499 | 0.26 | 0.22 | 0.078545 | Stroma 2 | si:ch211-281l24.3 |
| selenbp1 | 2.4E-06 | 0.4201417 | 0.104 | 0.077 | 0.079342 | Stroma 2 | selenbp1 |
| trim3b3 | 2.4E-06 | 0.4045186 | 0.122 | 0.094 | 0.079602 | Stroma 2 | trim3b |
| adam19a1 | 2.5E-06 | 0.4476412 | 0.11 | 0.083 | 0.079939 | Stroma 2 | adam19a |
| si:ch73-86n18.12 | 2.5E-06 | 0.458909 | 0.188 | 0.153 | 0.080468 | Stroma 2 | si:ch73-86n18.1 |
| frem1b1 | 2.9E-06 | 0.4185249 | 0.103 | 0.077 | 0.09407 | Stroma 2 | frem1b |
| pdia41 | 2.9E-06 | 0.2954136 | 0.182 | 0.148 | 0.095164 | Stroma 2 | pdia4 |
| psme11 | 3.3E-06 | 0.3115949 | 0.144 | 0.112 | 0.107366 | Stroma 2 | psme1 |
| lrrc153 | 3.4E-06 | 0.3341812 | 0.168 | 0.133 | 0.109366 | Stroma 2 | lrrc15 |
| sfxn5b3 | 3.7E-06 | 0.3315951 | 0.17 | 0.136 | 0.119851 | Stroma 2 | sfxn5b |
| ATP11B | 3.8E-06 | 0.3975833 | 0.137 | 0.108 | 0.123122 | Stroma 2 | ATP11B |
| sh3bp5a | 4.4E-06 | 0.3746789 | 0.161 | 0.129 | 0.143886 | Stroma 2 | sh3bp5a |
| aldh6a11 | 4.9E-06 | 0.2713933 | 0.31 | 0.272 | 0.15881 | Stroma 2 | aldh6a1 |
| twist1b1 | 4.9E-06 | 0.478546 | 0.19 | 0.158 | 0.160884 | Stroma 2 | twist1b |
| arf2a1 | 5.7E-06 | 0.2338174 | 0.281 | 0.242 | 0.184685 | Stroma 2 | arf2a |
| mob1ba2 | 5.7E-06 | 0.3832059 | 0.128 | 0.1 | 0.186375 | Stroma 2 | mob1ba |
| si:dkey-195m11.81 | 7.4E-06 | 0.2798003 | 0.158 | 0.126 | 0.239094 | Stroma 2 | si:dkey-195m11.8 |
| hopx | 7.9E-06 | 0.2821673 | 0.307 | 0.269 | 0.255714 | Stroma 2 | hopx |
| palm1b | 1E-05 | 0.4362237 | 0.109 | 0.083 | 0.331407 | Stroma 2 | palm1b |
| nhsl1a1 | 1E-05 | 0.440249 | 0.103 | 0.078 | 0.335839 | Stroma 2 | nhsl1a |
| tnrc6a | 1.1E-05 | 0.2514652 | 0.308 | 0.271 | 0.366968 | Stroma 2 | tnrc6a |
| klf12a2 | 1.2E-05 | 0.2430317 | 0.17 | 0.137 | 0.404503 | Stroma 2 | klf12a |
| slc7a10a3 | 1.4E-05 | 0.4015732 | 0.134 | 0.107 | 0.450227 | Stroma 2 | slc7a10a |
| klf152 | 1.6E-05 | 0.2442053 | 0.243 | 0.207 | 0.508452 | Stroma 2 | klf15 |
| cmbl1 | 1.7E-05 | 0.2888537 | 0.207 | 0.174 | 0.553319 | Stroma 2 | cmbl |
| cbsb2 | 1.8E-05 | 0.2112576 | 0.17 | 0.138 | 0.581485 | Stroma 2 | cbsb |
| hdac41 | 1.9E-05 | 0.4087986 | 0.147 | 0.119 | 0.615168 | Stroma 2 | hdac4 |
| bnc22 | 1.9E-05 | 0.4071424 | 0.106 | 0.081 | 0.633551 | Stroma 2 | bnc2 |
| adam8b2 | 2E-05 | 0.3660504 | 0.103 | 0.079 | 0.66648 | Stroma 2 | adam8b |
| kank31 | 2.4E-05 | 0.4229163 | 0.103 | 0.079 | 0.788892 | Stroma 2 | kank3 |
| ndnf3 | 2.7E-05 | 0.2811033 | 0.13 | 0.103 | 0.862229 | Stroma 2 | ndnf |
| fbln51 | 2.7E-05 | 0.3479865 | 0.107 | 0.083 | 0.869514 | Stroma 2 | fbln5 |
| gas1b3 | 2.9E-05 | 0.3839682 | 0.225 | 0.195 | 0.94172 | Stroma 2 | gas1b |
| adh8b2 | 2.9E-05 | 0.2441572 | 0.269 | 0.233 | 0.943636 | Stroma 2 | adh8b |
| tmem119b1 | 3.3E-05 | 0.2374598 | 0.241 | 0.205 | 1 | Stroma 2 | tmem119b |
| si:rp71-36a1.2 | 3.6E-05 | 0.3594448 | 0.111 | 0.087 | 1 | Stroma 2 | si:rp71-36a1.2 |
| bahcc1a1 | 3.8E-05 | 0.2382979 | 0.111 | 0.086 | 1 | Stroma 2 | bahcc1a |
| hspa8b1 | 4.2E-05 | 0.3345479 | 0.117 | 0.092 | 1 | Stroma 2 | hspa8b |
| slmapa | 4.7E-05 | 0.3449479 | 0.124 | 0.099 | 1 | Stroma 2 | slmapa |
| inka1b1 | 4.7E-05 | 0.259503 | 0.117 | 0.091 | 1 | Stroma 2 | inka1b |
| abat | 5.4E-05 | 0.3981202 | 0.124 | 0.099 | 1 | Stroma 2 | abat |
| fbxo342 | 6.6E-05 | 0.4189178 | 0.113 | 0.089 | 1 | Stroma 2 | fbxo34 |

| | | | | | | | |
|---|---|---|---|---|---|---|---|
| atp1b2a2 | 6.8E-05 | 0.3311969 | 0.104 | 0.081 | 1 | Stroma 2 | atp1b2a |
| tcf7l1a | 7E-05 | 0.322905 | 0.143 | 0.116 | 1 | Stroma 2 | tcf7l1a |
| lgmn1 | 7.4E-05 | 0.304955 | 0.201 | 0.171 | 1 | Stroma 2 | lgmn |
| gpc6a3 | 7.5E-05 | 0.2383335 | 0.234 | 0.201 | 1 | Stroma 2 | gpc6a |
| gadd45ab2 | 8.1E-05 | 0.2744123 | 0.205 | 0.175 | 1 | Stroma 2 | gadd45ab |
| cyp3c1 | 8.9E-05 | 0.3771813 | 0.118 | 0.094 | 1 | Stroma 2 | cyp3c1 |
| CABZ01064941.11 | 8.9E-05 | 0.4583477 | 0.107 | 0.085 | 1 | Stroma 2 | CABZ01064941.1 |
| porb2 | 9.5E-05 | 0.3724619 | 0.132 | 0.107 | 1 | Stroma 2 | porb |
| lman2 | 9.5E-05 | 0.2205157 | 0.268 | 0.235 | 1 | Stroma 2 | lman2 |
| pigr1 | 1E-04 | 0.2209866 | 0.219 | 0.188 | 1 | Stroma 2 | pigr |
| hif1al1 | 0.0001 | 0.344922 | 0.136 | 0.111 | 1 | Stroma 2 | hif1al |
| apmap2 | 0.00011 | 0.3473192 | 0.102 | 0.079 | 1 | Stroma 2 | apmap |
| prr12b | 0.00012 | 0.3623689 | 0.141 | 0.116 | 1 | Stroma 2 | prr12b |
| lamc11 | 0.00012 | 0.226414 | 0.155 | 0.128 | 1 | Stroma 2 | lamc1 |
| snx171 | 0.00013 | 0.2430545 | 0.274 | 0.242 | 1 | Stroma 2 | snx17 |
| abhd15a1 | 0.00017 | 0.2808585 | 0.158 | 0.132 | 1 | Stroma 2 | abhd15a |
| bcl9 | 0.00019 | 0.3608744 | 0.106 | 0.085 | 1 | Stroma 2 | bcl9 |
| si:dkeyp-44a8.2 | 0.00019 | 0.3435463 | 0.163 | 0.137 | 1 | Stroma 2 | si:dkeyp-44a8.2 |
| flot2a1 | 0.0002 | 0.2434219 | 0.142 | 0.116 | 1 | Stroma 2 | flot2a |
| limch1b | 0.0002 | 0.5300043 | 0.111 | 0.09 | 1 | Stroma 2 | limch1b |
| rbm4.11 | 0.00021 | 0.3512338 | 0.123 | 0.1 | 1 | Stroma 2 | rbm4.1 |
| pdia73 | 0.00021 | 0.2307477 | 0.171 | 0.144 | 1 | Stroma 2 | pdia7 |
| abcc51 | 0.00021 | 0.297206 | 0.17 | 0.143 | 1 | Stroma 2 | abcc5 |
| mvp2 | 0.00022 | 0.3093416 | 0.341 | 0.316 | 1 | Stroma 2 | mvp |
| slc25a25a1 | 0.00023 | 0.2395238 | 0.25 | 0.218 | 1 | Stroma 2 | slc25a25a |
| trappc51 | 0.00024 | 0.2476924 | 0.138 | 0.113 | 1 | Stroma 2 | trappc5 |
| zgc:77486.12 | 0.00025 | 0.2048287 | 0.197 | 0.168 | 1 | Stroma 2 | zgc:77486.1 |
| klf3 | 0.00027 | 0.3298711 | 0.124 | 0.101 | 1 | Stroma 2 | klf3 |
| map2k61 | 0.00027 | 0.2566832 | 0.217 | 0.189 | 1 | Stroma 2 | map2k6 |
| srsf5b1 | 0.00028 | 0.3441747 | 0.139 | 0.115 | 1 | Stroma 2 | srsf5b |
| myo1ca3 | 0.0003 | 0.2116647 | 0.232 | 0.202 | 1 | Stroma 2 | myo1ca |
| sesn12 | 0.00032 | 0.2938985 | 0.243 | 0.217 | 1 | Stroma 2 | sesn1 |
| mhc1zaa2 | 0.00032 | 0.3492713 | 0.114 | 0.092 | 1 | Stroma 2 | mhc1zaa |
| tgfbr3 | 0.00033 | 0.2273571 | 0.1 | 0.079 | 1 | Stroma 2 | tgfbr3 |
| etv5b1 | 0.00036 | 0.3738168 | 0.121 | 0.1 | 1 | Stroma 2 | etv5b |
| serinc53 | 0.00036 | 0.2424773 | 0.268 | 0.239 | 1 | Stroma 2 | serinc5 |
| grwd12 | 0.00036 | 0.3143331 | 0.166 | 0.141 | 1 | Stroma 2 | grwd1 |
| si:zfos-2326c3.2 | 0.00038 | 0.2620001 | 0.188 | 0.162 | 1 | Stroma 2 | si:zfos-2326c3.2 |
| cmtm6 | 0.00038 | 0.2515048 | 0.165 | 0.14 | 1 | Stroma 2 | cmtm6 |
| lix1l2 | 0.00039 | 0.2363777 | 0.202 | 0.173 | 1 | Stroma 2 | lix1l |
| rgmd1 | 0.00039 | 0.339505 | 0.105 | 0.085 | 1 | Stroma 2 | rgmd |
| limk2 | 0.0004 | 0.371455 | 0.1 | 0.081 | 1 | Stroma 2 | limk2 |
| rnf213a1 | 0.00045 | 0.2250685 | 0.299 | 0.268 | 1 | Stroma 2 | rnf213a |
| smarcc21 | 0.00047 | 0.2348588 | 0.199 | 0.172 | 1 | Stroma 2 | smarcc2 |
| eml31 | 0.00051 | 0.2355843 | 0.144 | 0.12 | 1 | Stroma 2 | eml3 |
| sash1a1 | 0.00057 | 0.3184345 | 0.234 | 0.209 | 1 | Stroma 2 | sash1a |
| rmdn11 | 0.00057 | 0.3193399 | 0.115 | 0.094 | 1 | Stroma 2 | rmdn1 |
| mapkapk2a1 | 0.00068 | 0.2322473 | 0.248 | 0.222 | 1 | Stroma 2 | mapkapk2a |

| | | | | | | | |
|---|---|---|---|---|---|---|---|
| ak3 | 0.0007 | 0.2982164 | 0.128 | 0.107 | 1 | Stroma 2 | ak3 |
| phactr21 | 0.0007 | 0.2857007 | 0.164 | 0.14 | 1 | Stroma 2 | phactr2 |
| bco2l1 | 0.00076 | 0.2851736 | 0.146 | 0.124 | 1 | Stroma 2 | bco2l |
| cnpy32 | 0.0008 | 0.3658859 | 0.107 | 0.088 | 1 | Stroma 2 | cnpy3 |
| grinab1 | 0.00081 | 0.2361957 | 0.237 | 0.212 | 1 | Stroma 2 | grinab |
| zgc:110591 | 0.00082 | 0.3763487 | 0.113 | 0.093 | 1 | Stroma 2 | zgc:110591 |
| rbm281 | 0.00091 | 0.2608963 | 0.266 | 0.24 | 1 | Stroma 2 | rbm28 |
| phyh | 0.00093 | 0.3550292 | 0.111 | 0.091 | 1 | Stroma 2 | phyh |
| zgc:1102392 | 0.00107 | 0.2423531 | 0.256 | 0.232 | 1 | Stroma 2 | zgc:110239 |
| sat1b2 | 0.00107 | 0.2748444 | 0.131 | 0.11 | 1 | Stroma 2 | sat1b |
| gusb1 | 0.00108 | 0.2278067 | 0.152 | 0.129 | 1 | Stroma 2 | gusb |
| thbs4b1 | 0.00116 | 0.3980908 | 0.28 | 0.259 | 1 | Stroma 2 | thbs4b |
| nfe2l1b1 | 0.00125 | 0.2895254 | 0.106 | 0.087 | 1 | Stroma 2 | nfe2l1b |
| cltca1 | 0.00158 | 0.2303457 | 0.221 | 0.197 | 1 | Stroma 2 | cltca |
| ngfb1 | 0.00159 | 0.300041 | 0.1 | 0.081 | 1 | Stroma 2 | ngfb |
| hmgn32 | 0.00163 | 0.2313228 | 0.185 | 0.162 | 1 | Stroma 2 | hmgn3 |
| ppp1r13l1 | 0.00163 | 0.2983983 | 0.106 | 0.087 | 1 | Stroma 2 | ppp1r13l |
| cdc42bpb | 0.00179 | 0.2196201 | 0.158 | 0.136 | 1 | Stroma 2 | cdc42bpb |
| stau1 | 0.00193 | 0.240531 | 0.164 | 0.143 | 1 | Stroma 2 | stau1 |
| ARF51 | 0.00194 | 0.2053454 | 0.175 | 0.152 | 1 | Stroma 2 | ARF5 |
| serpinh2 | 0.00222 | 0.2477928 | 0.138 | 0.117 | 1 | Stroma 2 | serpinh2 |
| ptp4a3a | 0.00301 | 0.2408813 | 0.14 | 0.12 | 1 | Stroma 2 | ptp4a3a |
| rnd1b3 | 0.00355 | 0.2958464 | 0.121 | 0.103 | 1 | Stroma 2 | rnd1b |
| dbpb1 | 0.00376 | 0.3863079 | 0.131 | 0.113 | 1 | Stroma 2 | dbpb |
| znf703 | 0.00383 | 0.2855689 | 0.113 | 0.096 | 1 | Stroma 2 | znf703 |
| proza | 0.00388 | 0.2328179 | 0.191 | 0.17 | 1 | Stroma 2 | proza |
| dipk1c3 | 0.00393 | 0.2362429 | 0.133 | 0.115 | 1 | Stroma 2 | dipk1c |
| zgc:1626121 | 0.00402 | 0.4241369 | 0.134 | 0.116 | 1 | Stroma 2 | zgc:162612 |
| lonrf1l1 | 0.00407 | 0.2216404 | 0.172 | 0.151 | 1 | Stroma 2 | lonrf1l |
| fam171a2a1 | 0.00416 | 0.2471431 | 0.1 | 0.083 | 1 | Stroma 2 | fam171a2a |
| tnfrsf191 | 0.00429 | 0.2343703 | 0.127 | 0.108 | 1 | Stroma 2 | tnfrsf19 |
| gsk3ba1 | 0.00448 | 0.3190239 | 0.163 | 0.145 | 1 | Stroma 2 | gsk3ba |
| arhgef1b | 0.00477 | 0.2621516 | 0.284 | 0.26 | 1 | Stroma 2 | arhgef1b |
| si:ch211-284f22.31 | 0.00489 | 0.2560669 | 0.135 | 0.117 | 1 | Stroma 2 | si:ch211-284f22.3 |
| galnt2 | 0.00505 | 0.3492385 | 0.106 | 0.09 | 1 | Stroma 2 | galnt2 |
| dusp8a2 | 0.00533 | 0.340788 | 0.106 | 0.09 | 1 | Stroma 2 | dusp8a |
| dusp52 | 0.0056 | 0.318817 | 0.218 | 0.198 | 1 | Stroma 2 | dusp5 |
| ugt2a71 | 0.00577 | 0.2950847 | 0.11 | 0.094 | 1 | Stroma 2 | ugt2a7 |
| nr1d2b1 | 0.00712 | 0.3046397 | 0.17 | 0.152 | 1 | Stroma 2 | nr1d2b |
| c1qtnf124 | 0.00784 | 0.3306237 | 0.117 | 0.101 | 1 | Stroma 2 | c1qtnf12 |
| ube2e2 | 0.00789 | 0.2717919 | 0.131 | 0.114 | 1 | Stroma 2 | ube2e2 |
| fam193a1 | 0.00811 | 0.260462 | 0.119 | 0.103 | 1 | Stroma 2 | fam193a |
| ddt | 0.00857 | 0.2318753 | 0.212 | 0.193 | 1 | Stroma 2 | ddt |
| slc29a1a2 | 0.0086 | 0.2298842 | 0.217 | 0.198 | 1 | Stroma 2 | slc29a1a |
| rif12 | 0.00876 | 0.2457809 | 0.101 | 0.086 | 1 | Stroma 2 | rif1 |
| asap1b2 | 0.0093 | 0.2263247 | 0.17 | 0.151 | 1 | Stroma 2 | asap1b |
| zgc:1140461 | 0.00935 | 0.3275604 | 0.115 | 0.1 | 1 | Stroma 2 | zgc:114046 |
| hacd2 | 0.0097 | 0.3187438 | 0.105 | 0.09 | 1 | Stroma 2 | hacd2 |

| | | | | | | | |
|---|---|---|---|---|---|---|---|
| mamdc2b | 0 | 3.9488686 | 0.868 | 0.205 | 0 | DF 4 | mamdc2b |
| ca62 | 0 | 2.9303754 | 0.944 | 0.312 | 0 | DF 4 | ca6 |
| aqp3a3 | 0 | 2.6426098 | 0.862 | 0.288 | 0 | DF 4 | aqp3a |
| ccdc80l2 | 0 | 3.6797059 | 0.654 | 0.082 | 0 | DF 4 | ccdc80l2 |
| col7a11 | 0 | 3.4011129 | 0.672 | 0.125 | 0 | DF 4 | col7a1 |
| ccl441 | 0 | 2.947456 | 0.639 | 0.098 | 0 | DF 4 | ccl44 |
| fmoda1 | 0 | 2.4624965 | 0.642 | 0.156 | 0 | DF 4 | fmoda |
| si:ch211-207e14.4 | 0 | 4.2372794 | 0.499 | 0.044 | 0 | DF 4 | si:ch211-207e14.4 |
| postnb1 | 0 | 2.3755411 | 0.881 | 0.431 | 0 | DF 4 | postnb |
| cyp26b1 | 0 | 1.8840862 | 0.79 | 0.344 | 0 | DF 4 | cyp26b1 |
| col11a1a1 | 0 | 0.4210164 | 0.743 | 0.297 | 0 | DF 4 | col11a1a |
| pah3 | 0 | 1.6729547 | 0.768 | 0.326 | 0 | DF 4 | pah |
| wfdc2 | 0 | 2.261787 | 0.813 | 0.377 | 0 | DF 4 | wfdc2 |
| adma2 | 0 | 2.1858661 | 0.644 | 0.21 | 0 | DF 4 | adma |
| spock33 | 0 | 1.2669185 | 0.805 | 0.39 | 0 | DF 4 | spock3 |
| sfrp23 | 0 | 1.5482728 | 0.784 | 0.375 | 0 | DF 4 | sfrp2 |
| fstb4 | 0 | 1.5839762 | 0.748 | 0.339 | 0 | DF 4 | fstb |
| fgl12 | 0 | 2.007142 | 0.593 | 0.188 | 0 | DF 4 | fgl1 |
| pcbd13 | 0 | 1.7547553 | 0.69 | 0.295 | 0 | DF 4 | pcbd1 |
| qdpra3 | 0 | 1.4655163 | 0.798 | 0.405 | 0 | DF 4 | qdpra |
| zgc:113307 | 0 | 3.2803121 | 0.459 | 0.071 | 0 | DF 4 | zgc:113307 |
| tcf121 | 0 | 1.6342824 | 0.888 | 0.502 | 0 | DF 4 | tcf12 |
| twist24 | 0 | 1.6055174 | 0.706 | 0.32 | 0 | DF 4 | twist2 |
| CABZ01085139.1 | 0 | 3.6634382 | 0.427 | 0.043 | 0 | DF 4 | CABZ01085139.1 |
| hpdb4 | 0 | 1.0560334 | 0.869 | 0.488 | 0 | DF 4 | hpdb |
| col5a3b | 0 | 4.464569 | 0.404 | 0.026 | 0 | DF 4 | col5a3b |
| col2a1b1 | 0 | 0.9625092 | 0.858 | 0.481 | 0 | DF 4 | col2a1b |
| si:dkeyp-44a8.42 | 0 | 2.3022866 | 0.52 | 0.145 | 0 | DF 4 | si:dkeyp-44a8.4 |
| tat3 | 0 | 1.8645966 | 0.585 | 0.226 | 0 | DF 4 | tat |
| twist32 | 0 | 1.677096 | 0.593 | 0.234 | 0 | DF 4 | twist3 |
| alpl | 0 | 3.3827579 | 0.406 | 0.048 | 0 | DF 4 | alpl |
| fsta | 0 | 4.7214205 | 0.382 | 0.025 | 0 | DF 4 | fsta |
| asip13 | 0 | 2.3226965 | 0.509 | 0.158 | 0 | DF 4 | asip1 |
| adamts23 | 0 | 1.493804 | 0.705 | 0.354 | 0 | DF 4 | adamts2 |
| si:ch211-243a20.32 | 0 | 1.5333033 | 0.611 | 0.263 | 0 | DF 4 | si:ch211-243a20.3 |
| smoc12 | 0 | 1.629158 | 0.509 | 0.19 | 0 | DF 4 | smoc1 |
| cpxm1a1 | 0 | 2.1579255 | 0.479 | 0.161 | 0 | DF 4 | cpxm1a |
| col12a1b1 | 0 | 1.3731047 | 0.916 | 0.607 | 0 | DF 4 | col12a1b |
| vstm4a1 | 0 | 2.3497453 | 0.437 | 0.128 | 0 | DF 4 | vstm4a |
| ccl25b3 | 0 | 1.701093 | 0.953 | 0.647 | 0 | DF 4 | ccl25b |
| si:ch211-170d8.82 | 0 | 1.5138834 | 0.769 | 0.468 | 0 | DF 4 | si:ch211-170d8.8 |
| ism11 | 0 | 2.3349818 | 0.404 | 0.108 | 0 | DF 4 | ism1 |
| cx432 | 0 | 2.1853023 | 0.446 | 0.154 | 0 | DF 4 | cx43 |
| lamc3 | 0 | 2.1026431 | 0.391 | 0.105 | 0 | DF 4 | lamc3 |
| ptgdsb.23 | 0 | 1.6566102 | 0.933 | 0.652 | 0 | DF 4 | ptgdsb.2 |
| agtr22 | 0 | 2.2063766 | 0.38 | 0.101 | 0 | DF 4 | agtr2 |
| col12a1a1 | 0 | 1.8875094 | 0.994 | 0.738 | 0 | DF 4 | col12a1a |
| ngfra | 0 | 3.1573903 | 0.293 | 0.04 | 0 | DF 4 | ngfra |

| | | | | | | | |
|---|---|---|---|---|---|---|---|
| paplnb | 0 | 4.7350379 | 0.263 | 0.02 | 0 | DF 4 | paplnb |
| adamtsl31 | 0 | 2.6568902 | 0.297 | 0.056 | 0 | DF 4 | adamtsl3 |
| si:ch211-196i2.11 | 0 | 2.7792491 | 0.28 | 0.04 | 0 | DF 4 | si:ch211-196i2.1 |
| lum2 | 0 | 0.8152182 | 0.98 | 0.746 | 0 | DF 4 | lum |
| megf6a | 0 | 2.796339 | 0.259 | 0.047 | 0 | DF 4 | megf6a |
| si:ch211-252f13.5 | 0 | 3.7772539 | 0.225 | 0.018 | 0 | DF 4 | si:ch211-252f13.5 |
| admb | 0 | 3.0608582 | 0.231 | 0.028 | 0 | DF 4 | admb |
| tnn | 0 | 4.5672595 | 0.224 | 0.022 | 0 | DF 4 | tnn |
| col6a4a | 0 | 2.4627218 | 0.242 | 0.046 | 0 | DF 4 | col6a4a |
| ccn1l1 | 0 | 3.3417589 | 0.222 | 0.027 | 0 | DF 4 | ccn1l1 |
| si:ch211-67e16.11 | 0 | 3.6771875 | 0.206 | 0.018 | 0 | DF 4 | si:ch211-67e16.11 |
| cemip | 0 | 3.213739 | 0.23 | 0.048 | 0 | DF 4 | cemip |
| si:ch211-254n4.3 | 0 | 3.4636062 | 0.194 | 0.019 | 0 | DF 4 | si:ch211-254n4.3 |
| chrdl2 | 0 | 3.1234335 | 0.203 | 0.029 | 0 | DF 4 | chrdl2 |
| si:ch1073-459j12.12 | 0 | 1.0006395 | 0.968 | 0.802 | 0 | DF 4 | si:ch1073-459j12.1 |
| fgf18b | 0 | 3.8614199 | 0.151 | 0.011 | 0 | DF 4 | fgf18b |
| htr2b | 0 | 4.1446003 | 0.147 | 0.008 | 0 | DF 4 | htr2b |
| defbl1 | 0 | 4.4240469 | 0.136 | 0.011 | 0 | DF 4 | defbl1 |
| col6a12 | 0 | 1.1307336 | 0.988 | 0.868 | 0 | DF 4 | col6a1 |
| stc1 | 0 | 4.4463607 | 0.126 | 0.009 | 0 | DF 4 | stc1 |
| col6a22 | 0 | 1.0507811 | 0.987 | 0.871 | 0 | DF 4 | col6a2 |
| selenow12 | 0 | 0.8016321 | 0.97 | 0.893 | 0 | DF 4 | selenow1 |
| col6a3.11 | 0 | 1.130522 | 0.999 | 0.935 | 0 | DF 4 | col6a3.1 |
| BX663503.1 | 6E-307 | 2.304889 | 0.253 | 0.056 | 1.9E-302 | DF 4 | BX663503.1 |
| vmo1a2 | 4E-302 | 0.9403326 | 0.953 | 0.686 | 1.3E-297 | DF 4 | vmo1a |
| tmsb12 | 3E-296 | 1.3190902 | 0.757 | 0.432 | 1.1E-291 | DF 4 | tmsb1 |
| vamp1 | 5E-293 | 3.696384 | 0.107 | 0.01 | 1.8E-288 | DF 4 | vamp1 |
| C18H15orf483 | 1E-289 | 1.3531064 | 0.626 | 0.292 | 3.3E-285 | DF 4 | C18H15orf48 |
| timp4.12 | 4E-287 | 1.6539501 | 0.309 | 0.083 | 1.3E-282 | DF 4 | timp4.1 |
| ucp23 | 2E-286 | 1.5986193 | 0.704 | 0.399 | 5.1E-282 | DF 4 | ucp2 |
| pcdh10b | 6E-286 | 2.5031792 | 0.192 | 0.035 | 1.8E-281 | DF 4 | pcdh10b |
| fstl32 | 5E-282 | 1.44485 | 0.632 | 0.304 | 1.5E-277 | DF 4 | fstl3 |
| gstr1 | 4E-276 | 1.4663815 | 0.699 | 0.4 | 1.2E-271 | DF 4 | gstr |
| htra3a | 2E-271 | 2.3935291 | 0.194 | 0.038 | 5.4E-267 | DF 4 | htra3a |
| col1a1b3 | 1E-266 | 0.6007259 | 1 | 0.986 | 3.3E-262 | DF 4 | col1a1b |
| capn123 | 1E-264 | 1.1766583 | 0.528 | 0.208 | 3.6E-260 | DF 4 | capn12 |
| heyl2 | 7E-264 | 2.0501552 | 0.252 | 0.062 | 2.4E-259 | DF 4 | heyl |
| axin21 | 4E-260 | 1.5013457 | 0.495 | 0.204 | 1.2E-255 | DF 4 | axin2 |
| wnt112 | 1E-257 | 1.5953587 | 0.561 | 0.268 | 3.8E-253 | DF 4 | wnt11 |
| tent5ab | 2E-257 | 2.382775 | 0.219 | 0.049 | 5E-253 | DF 4 | tent5ab |
| sema5a1 | 3E-255 | 2.1099118 | 0.267 | 0.071 | 1E-250 | DF 4 | sema5a |
| gstp13 | 6E-252 | 1.0002675 | 0.864 | 0.631 | 1.9E-247 | DF 4 | gstp1 |
| si:ch1073-291c23.22 | 2E-244 | 1.1551581 | 0.739 | 0.46 | 5.7E-240 | DF 4 | si:ch1073-291c23.2 |
| kera2 | 2E-244 | 1.4584685 | 0.422 | 0.158 | 6.7E-240 | DF 4 | kera |
| prex2 | 1E-242 | 2.1088111 | 0.274 | 0.077 | 3.2E-238 | DF 4 | prex2 |
| FO203432.11 | 2E-237 | 2.6259191 | 0.206 | 0.047 | 6.1E-233 | DF 4 | FO203432.1 |
| gask1a1 | 9E-231 | 1.662099 | 0.438 | 0.181 | 2.9E-226 | DF 4 | gask1a |
| col1a23 | 3E-224 | 0.4840428 | 1 | 0.991 | 1E-219 | DF 4 | col1a2 |

| | | | | | | | |
|---|---|---|---|---|---|---|---|
| fgfbp2a4 | 7E-224 | 0.7378914 | 0.853 | 0.518 | 2.2E-219 | DF 4 | fgfbp2a |
| rbp43 | 1E-220 | 0.6907198 | 0.995 | 0.905 | 4.6E-216 | DF 4 | rbp4 |
| gstm.13 | 2E-218 | 0.9236005 | 0.819 | 0.584 | 6.3E-214 | DF 4 | gstm.1 |
| postna5 | 6E-217 | 0.7270339 | 0.899 | 0.573 | 1.8E-212 | DF 4 | postna |
| ecm1b3 | 8E-217 | 0.9685073 | 0.751 | 0.456 | 2.7E-212 | DF 4 | ecm1b |
| ecrg4a | 1E-214 | 0.9631419 | 0.823 | 0.549 | 3.3E-210 | DF 4 | ecrg4a |
| podn2 | 3E-212 | 1.5001707 | 0.403 | 0.159 | 1.1E-207 | DF 4 | podn |
| phldb2b1 | 7E-212 | 1.6932125 | 0.356 | 0.132 | 2.3E-207 | DF 4 | phldb2b |
| pleca1 | 9E-211 | 0.6633187 | 0.985 | 0.914 | 2.8E-206 | DF 4 | pleca |
| aebp11 | 1E-208 | 0.8905588 | 0.786 | 0.526 | 3.1E-204 | DF 4 | aebp1 |
| itga101 | 2E-207 | 1.1561539 | 0.519 | 0.239 | 6.2E-203 | DF 4 | itga10 |
| slc6a17 | 5E-206 | 2.2204673 | 0.172 | 0.037 | 1.8E-201 | DF 4 | slc6a17 |
| col1a1a3 | 4E-202 | 0.39981 | 1 | 0.979 | 1.2E-197 | DF 4 | col1a1a |
| rtn1b | 1E-200 | 2.3428926 | 0.156 | 0.032 | 3.9E-196 | DF 4 | rtn1b |
| slc51a4 | 4E-200 | 1.3453175 | 0.38 | 0.146 | 1.3E-195 | DF 4 | slc51a |
| ttc364 | 6E-199 | 1.0432439 | 0.634 | 0.338 | 1.9E-194 | DF 4 | ttc36 |
| zgc:565854 | 2E-194 | 1.039075 | 0.602 | 0.313 | 7.2E-190 | DF 4 | zgc:56585 |
| si:ch73-46j18.52 | 9E-192 | 1.0984449 | 0.785 | 0.592 | 3E-187 | DF 4 | si:ch73-46j18.5 |
| itga8 | 1E-190 | 2.537574 | 0.125 | 0.022 | 4.6E-186 | DF 4 | itga8 |
| si:ch211-160j14.33 | 3E-190 | 1.0625482 | 0.584 | 0.299 | 9.3E-186 | DF 4 | si:ch211-160j14.3 |
| cyba3 | 5E-190 | 1.6897063 | 0.372 | 0.152 | 1.7E-185 | DF 4 | cyba |
| vwa11 | 9E-190 | 1.3824271 | 0.34 | 0.127 | 3E-185 | DF 4 | vwa1 |
| col11a1b3 | 6E-187 | 0.8627277 | 0.794 | 0.529 | 2E-182 | DF 4 | col11a1b |
| lix1l3 | 1E-183 | 1.4364294 | 0.388 | 0.162 | 3.7E-179 | DF 4 | lix1l |
| cdo12 | 2E-180 | 1.2409539 | 0.566 | 0.316 | 8.1E-176 | DF 4 | cdo1 |
| ptx3a3 | 6E-179 | 0.6416707 | 0.796 | 0.47 | 1.9E-174 | DF 4 | ptx3a |
| sgtb1 | 1E-177 | 1.5811707 | 0.31 | 0.114 | 3.5E-173 | DF 4 | sgtb |
| abi3bpb1 | 5E-177 | 0.491415 | 0.842 | 0.571 | 1.7E-172 | DF 4 | abi3bpb |
| stk17al1 | 7E-175 | 1.8906478 | 0.224 | 0.068 | 2.2E-170 | DF 4 | stk17al |
| si:dkey-226m8.102 | 8E-175 | 1.6944167 | 0.268 | 0.09 | 2.6E-170 | DF 4 | si:dkey-226m8.10 |
| hgd3 | 8E-173 | 0.9472469 | 0.68 | 0.415 | 2.6E-168 | DF 4 | hgd |
| col18a1a | 3E-172 | 1.1664672 | 0.354 | 0.142 | 9.2E-168 | DF 4 | col18a1a |
| osr11 | 7E-170 | 1.3117855 | 0.381 | 0.161 | 2.4E-165 | DF 4 | osr1 |
| pbxip1b1 | 2E-167 | 1.2233451 | 0.468 | 0.228 | 5.8E-163 | DF 4 | pbxip1b |
| lipf | 3E-166 | 1.3799887 | 0.409 | 0.186 | 8.7E-162 | DF 4 | lipf |
| CABZ01092781.1 | 9E-166 | 1.9413285 | 0.17 | 0.043 | 2.9E-161 | DF 4 | CABZ01092781.1 |
| hhla2a.11 | 2E-164 | 1.6990384 | 0.231 | 0.073 | 5.1E-160 | DF 4 | hhla2a.1 |
| sema3aa2 | 4E-164 | 1.5496683 | 0.279 | 0.099 | 1.2E-159 | DF 4 | sema3aa |
| alx4a1 | 4E-164 | 0.9689993 | 0.621 | 0.356 | 1.4E-159 | DF 4 | alx4a |
| pigrl2.3 | 2E-162 | 2.3573614 | 0.113 | 0.021 | 6.2E-158 | DF 4 | pigrl2.3 |
| gpr1432 | 3E-162 | 1.7144754 | 0.211 | 0.063 | 1E-157 | DF 4 | gpr143 |
| nfe2l2a3 | 4E-162 | 1.0645289 | 0.58 | 0.331 | 1.4E-157 | DF 4 | nfe2l2a |
| sfrp1a3 | 5E-160 | 0.8915091 | 0.681 | 0.424 | 1.5E-155 | DF 4 | sfrp1a |
| neu3.23 | 3E-157 | 1.2158039 | 0.441 | 0.21 | 1.1E-152 | DF 4 | neu3.2 |
| c42 | 1E-156 | 1.0279074 | 0.424 | 0.195 | 4.5E-152 | DF 4 | c4 |
| scinla1 | 6E-156 | 0.9400449 | 0.645 | 0.396 | 1.8E-151 | DF 4 | scinla |
| ackr3a | 5E-151 | 1.9110192 | 0.158 | 0.041 | 1.7E-146 | DF 4 | ackr3a |
| cd99l21 | 1E-150 | 1.2701944 | 0.411 | 0.196 | 3.4E-146 | DF 4 | cd99l2 |

| | | | | | | | |
|---|---|---|---|---|---|---|---|
| FQ323156.13 | 2E-150 | 1.1649838 | 0.401 | 0.184 | 6.7E-146 | DF 4 | FQ323156.1 |
| nfil3-63 | 1E-149 | 1.1594951 | 0.534 | 0.293 | 4.4E-145 | DF 4 | nfil3-6 |
| csf1b | 8E-149 | 1.8824573 | 0.148 | 0.036 | 2.8E-144 | DF 4 | csf1b |
| sparc2 | 5E-148 | 0.2354594 | 0.998 | 0.979 | 1.5E-143 | DF 4 | sparc |
| pigr2 | 5E-147 | 1.3819413 | 0.382 | 0.178 | 1.6E-142 | DF 4 | pigr |
| smarcd21 | 2E-145 | 1.3801155 | 0.35 | 0.155 | 5.8E-141 | DF 4 | smarcd2 |
| ripor3 | 2E-142 | 1.8617128 | 0.157 | 0.042 | 6.7E-138 | DF 4 | ripor3 |
| ism2a1 | 4E-140 | 1.7209828 | 0.166 | 0.046 | 1.5E-135 | DF 4 | ism2a |
| epas1a | 3E-139 | 1.6206932 | 0.189 | 0.057 | 9.2E-135 | DF 4 | epas1a |
| ppfibp1a | 4E-139 | 1.8958622 | 0.17 | 0.049 | 1.2E-134 | DF 4 | ppfibp1a |
| plpp33 | 6E-137 | 0.6641735 | 0.708 | 0.44 | 1.9E-132 | DF 4 | plpp3 |
| adh8a2 | 9E-137 | 1.1659867 | 0.407 | 0.2 | 3E-132 | DF 4 | adh8a |
| calm1a1 | 2E-136 | 0.6884763 | 0.795 | 0.609 | 8E-132 | DF 4 | calm1a |
| tmem119b2 | 4E-134 | 1.1955102 | 0.399 | 0.196 | 1.4E-129 | DF 4 | tmem119b |
| si:ch211-191i18.2 | 2E-133 | 1.5229727 | 0.133 | 0.032 | 5E-129 | DF 4 | si:ch211-191i18.2 |
| entpd13 | 3E-132 | 1.4920038 | 0.217 | 0.074 | 9.5E-128 | DF 4 | entpd1 |
| wnt4 | 8E-132 | 1.9721807 | 0.124 | 0.029 | 2.5E-127 | DF 4 | wnt4 |
| pdgfra2 | 6E-131 | 0.802152 | 0.692 | 0.447 | 1.8E-126 | DF 4 | pdgfra |
| clcf1 | 6E-128 | 1.7934812 | 0.156 | 0.044 | 2E-123 | DF 4 | clcf1 |
| baiap2l1a | 2E-127 | 1.8519433 | 0.13 | 0.033 | 8E-123 | DF 4 | baiap2l1a |
| thbs4b2 | 7E-126 | 0.4460295 | 0.48 | 0.247 | 2.4E-121 | DF 4 | thbs4b |
| appb | 2E-125 | 0.8469877 | 0.665 | 0.457 | 6.9E-121 | DF 4 | appb |
| scube3 | 2E-124 | 1.4271831 | 0.237 | 0.088 | 5.2E-120 | DF 4 | scube3 |
| dcn3 | 5E-124 | 0.389655 | 0.973 | 0.863 | 1.6E-119 | DF 4 | dcn |
| cfp3 | 1E-122 | 0.8669282 | 0.56 | 0.328 | 3.5E-118 | DF 4 | cfp |
| rasl12 | 3E-122 | 0.5162936 | 0.173 | 0.053 | 1E-117 | DF 4 | rasl12 |
| ctsk4 | 1E-120 | 0.7069872 | 0.77 | 0.583 | 3.8E-116 | DF 4 | ctsk |
| nkd1 | 2E-119 | 1.567038 | 0.212 | 0.076 | 5.2E-115 | DF 4 | nkd1 |
| ednraa2 | 3E-118 | 1.0144224 | 0.401 | 0.203 | 1.1E-113 | DF 4 | ednraa |
| btc | 1E-115 | 1.6463053 | 0.147 | 0.042 | 3.2E-111 | DF 4 | btc |
| eef1da1 | 2E-114 | 0.4761605 | 0.939 | 0.872 | 7.6E-110 | DF 4 | eef1da |
| pdgfrl2 | 1E-113 | 0.6444813 | 0.76 | 0.575 | 4.9E-109 | DF 4 | pdgfrl |
| col6a31 | 2E-113 | 1.0723346 | 0.422 | 0.228 | 6E-109 | DF 4 | col6a3 |
| slc1a3a2 | 5E-112 | 1.1974883 | 0.308 | 0.141 | 1.6E-107 | DF 4 | slc1a3a |
| grb10b | 9E-112 | 0.906065 | 0.56 | 0.35 | 3E-107 | DF 4 | grb10b |
| ccdc80l12 | 2E-111 | 1.4297431 | 0.202 | 0.072 | 5.1E-107 | DF 4 | ccdc80l1 |
| bhmt4 | 4E-110 | 0.4929053 | 0.889 | 0.698 | 1.4E-105 | DF 4 | bhmt |
| glulb2 | 6E-110 | 0.6528101 | 0.768 | 0.558 | 2E-105 | DF 4 | glulb |
| mbd3b | 1E-109 | 1.7519421 | 0.154 | 0.048 | 4.6E-105 | DF 4 | mbd3b |
| si:ch211-250c4.41 | 1E-108 | 1.6987377 | 0.137 | 0.039 | 3.3E-104 | DF 4 | si:ch211-250c4.4 |
| sod1 | 1E-107 | 0.8405872 | 0.633 | 0.457 | 3.5E-103 | DF 4 | sod1 |
| kitlga1 | 8E-106 | 1.5023899 | 0.183 | 0.065 | 2.5E-101 | DF 4 | kitlga |
| abi3bpa3 | 1E-105 | 0.6835298 | 0.511 | 0.288 | 4E-101 | DF 4 | abi3bpa |
| stc2a1 | 2E-105 | 1.6549072 | 0.152 | 0.049 | 7.4E-101 | DF 4 | stc2a |
| glula2 | 4E-105 | 0.5815469 | 0.827 | 0.607 | 1.4E-100 | DF 4 | glula |
| hapln1a2 | 1E-104 | 0.834196 | 0.67 | 0.468 | 3.5E-100 | DF 4 | hapln1a |
| hmgb3a | 3E-104 | 1.3615101 | 0.252 | 0.108 | 1.1E-99 | DF 4 | hmgb3a |
| itgb41 | 3E-104 | 0.8069886 | 0.513 | 0.301 | 1.1E-99 | DF 4 | itgb4 |

| | | | | | | | |
|---|---|---|---|---|---|---|---|
| mmp14a2 | 6E-102 | 0.5520284 | 0.801 | 0.639 | 2.04E-97 | DF 4 | mmp14a |
| itgb1b.23 | 4E-101 | 1.3813074 | 0.245 | 0.104 | 1.25E-96 | DF 4 | itgb1b.2 |
| ptgdsb.14 | 1E-100 | 0.8640525 | 0.66 | 0.452 | 3.2E-96 | DF 4 | ptgdsb.1 |
| wls2 | 1E-100 | 0.86057 | 0.573 | 0.385 | 3.66E-96 | DF 4 | wls |
| pnrc21 | 3E-100 | 0.4734578 | 0.94 | 0.878 | 1.1E-95 | DF 4 | pnrc2 |
| timp4.31 | 1.6E-98 | 0.475738 | 0.643 | 0.429 | 5.07E-94 | DF 4 | timp4.3 |
| cdh113 | 2.1E-98 | 0.5226907 | 0.786 | 0.579 | 6.76E-94 | DF 4 | cdh11 |
| gchfr2 | 3.5E-97 | 1.1261307 | 0.316 | 0.154 | 1.15E-92 | DF 4 | gchfr |
| ccdc3b2 | 3.6E-96 | 1.2341139 | 0.26 | 0.117 | 1.16E-91 | DF 4 | ccdc3b |
| rem11 | 7.7E-96 | 1.6468784 | 0.167 | 0.059 | 2.51E-91 | DF 4 | rem1 |
| gpm6bb2 | 7.7E-96 | 0.9262293 | 0.513 | 0.325 | 2.52E-91 | DF 4 | gpm6bb |
| mylka1 | 4E-93 | 1.0643125 | 0.236 | 0.101 | 1.31E-88 | DF 4 | mylka |
| FQ323119.11 | 1.7E-90 | 1.2681339 | 0.227 | 0.097 | 5.59E-86 | DF 4 | FQ323119.1 |
| efemp2a2 | 3E-90 | 0.7872445 | 0.541 | 0.352 | 9.88E-86 | DF 4 | efemp2a |
| apof1 | 3E-90 | 1.4981379 | 0.167 | 0.061 | 9.89E-86 | DF 4 | apof |
| nrn1a1 | 4.1E-90 | 1.4909181 | 0.204 | 0.083 | 1.32E-85 | DF 4 | nrn1a |
| tmeff2b | 2.3E-89 | 1.8068412 | 0.101 | 0.027 | 7.41E-85 | DF 4 | tmeff2b |
| cited1 | 1.5E-88 | 1.7957898 | 0.112 | 0.032 | 4.75E-84 | DF 4 | cited1 |
| cxcl12a4 | 1.5E-88 | 0.311661 | 0.867 | 0.672 | 4.83E-84 | DF 4 | cxcl12a |
| abca1b | 6.7E-87 | 1.4515992 | 0.132 | 0.042 | 2.18E-82 | DF 4 | abca1b |
| crip12 | 9.9E-86 | 0.2883946 | 0.655 | 0.452 | 3.21E-81 | DF 4 | crip1 |
| lmo4b3 | 1.3E-85 | 0.7584473 | 0.497 | 0.309 | 4.36E-81 | DF 4 | lmo4b |
| eya12 | 1.5E-85 | 1.2307129 | 0.268 | 0.129 | 4.99E-81 | DF 4 | eya1 |
| sept12 | 2.5E-85 | 1.5506509 | 0.124 | 0.039 | 8.29E-81 | DF 4 | sept12 |
| cd82a3 | 6.1E-85 | 0.7400747 | 0.534 | 0.345 | 1.98E-80 | DF 4 | cd82a |
| iscub1 | 8.7E-85 | 0.9346288 | 0.345 | 0.184 | 2.83E-80 | DF 4 | iscub |
| gstz13 | 2E-84 | 0.6997679 | 0.52 | 0.335 | 6.45E-80 | DF 4 | gstz1 |
| pcdh18a2 | 2.1E-84 | 1.1556692 | 0.192 | 0.077 | 6.96E-80 | DF 4 | pcdh18a |
| klf153 | 3.1E-84 | 0.9707582 | 0.363 | 0.2 | 1.01E-79 | DF 4 | klf15 |
| daam1a | 7.2E-84 | 0.9850732 | 0.356 | 0.196 | 2.33E-79 | DF 4 | daam1a |
| ppdpfa | 8.7E-84 | 1.6584724 | 0.104 | 0.03 | 2.82E-79 | DF 4 | ppdpfa |
| slc1a53 | 1.2E-83 | 0.9102019 | 0.42 | 0.249 | 3.9E-79 | DF 4 | slc1a5 |
| tpbga | 1.6E-83 | 1.6059981 | 0.119 | 0.037 | 5.26E-79 | DF 4 | tpbga |
| selenop1 | 5.7E-83 | 0.4399198 | 0.87 | 0.734 | 1.84E-78 | DF 4 | selenop |
| tpst12 | 1E-82 | 0.9772854 | 0.323 | 0.168 | 3.34E-78 | DF 4 | tpst1 |
| itm2ba1 | 2.1E-81 | 0.4447195 | 0.845 | 0.736 | 6.89E-77 | DF 4 | itm2ba |
| tob1b2 | 6.4E-81 | 0.4713854 | 0.842 | 0.695 | 2.1E-76 | DF 4 | tob1b |
| ephb3a2 | 1E-80 | 1.215791 | 0.224 | 0.1 | 3.35E-76 | DF 4 | ephb3a |
| fam169aa3 | 1.1E-80 | 0.4805067 | 0.607 | 0.393 | 3.69E-76 | DF 4 | fam169aa |
| znf609b2 | 2.7E-80 | 0.8867705 | 0.431 | 0.265 | 8.9E-76 | DF 4 | znf609b |
| tmem59l2 | 7.3E-80 | 1.1048943 | 0.224 | 0.1 | 2.39E-75 | DF 4 | tmem59l |
| ccbe1 | 2E-78 | 1.6427963 | 0.131 | 0.045 | 6.49E-74 | DF 4 | ccbe1 |
| tnfsf122 | 2.5E-78 | 1.0026196 | 0.279 | 0.139 | 8.13E-74 | DF 4 | tnfsf12 |
| ckbb | 2.9E-78 | 1.1790123 | 0.339 | 0.191 | 9.34E-74 | DF 4 | ckbb |
| tnrc183 | 4.6E-78 | 0.7460347 | 0.573 | 0.398 | 1.5E-73 | DF 4 | tnrc18 |
| dlc | 5.8E-78 | 1.8552972 | 0.101 | 0.03 | 1.88E-73 | DF 4 | dlc |
| si:ch211-136a13.11 | 2.4E-77 | 1.4363652 | 0.189 | 0.08 | 7.76E-73 | DF 4 | si:ch211-136a13.1 |
| arpc1b | 2.8E-77 | 0.2287375 | 0.41 | 0.239 | 9.17E-73 | DF 4 | arpc1b |

| | | | | | | | |
|---|---|---|---|---|---|---|---|
| tbx15 | 3.3E-77 | 1.4267095 | 0.156 | 0.059 | 1.07E-72 | DF 4 | tbx15 |
| ANPEP1 | 1E-76 | 0.9126836 | 0.358 | 0.202 | 3.33E-72 | DF 4 | ANPEP |
| prrx1a2 | 1E-76 | 0.5225005 | 0.747 | 0.578 | 3.36E-72 | DF 4 | prrx1a |
| myl9a2 | 5.9E-76 | 0.7202258 | 0.547 | 0.373 | 1.93E-71 | DF 4 | myl9a |
| si:ch211-251b21.12 | 6.4E-76 | 0.4716619 | 0.65 | 0.44 | 2.1E-71 | DF 4 | si:ch211-251b21.1 |
| lamb1a | 6.6E-76 | 1.4388741 | 0.136 | 0.049 | 2.14E-71 | DF 4 | lamb1a |
| zgc:1943983 | 9E-76 | 0.4327799 | 0.581 | 0.373 | 2.93E-71 | DF 4 | zgc:194398 |
| anos1a | 1.2E-75 | 1.3266906 | 0.11 | 0.035 | 3.95E-71 | DF 4 | anos1a |
| shroom41 | 1.5E-75 | 1.2256699 | 0.255 | 0.127 | 4.76E-71 | DF 4 | shroom4 |
| edil3a2 | 7.9E-75 | 0.5880111 | 0.617 | 0.425 | 2.56E-70 | DF 4 | edil3a |
| hspa12b2 | 9.1E-74 | 1.1007773 | 0.243 | 0.116 | 2.97E-69 | DF 4 | hspa12b |
| si:dkeyp-72e1.9 | 1.4E-73 | 1.6172772 | 0.103 | 0.032 | 4.48E-69 | DF 4 | si:dkeyp-72e1.9 |
| spink2.1 | 2.5E-73 | 1.0490252 | 0.25 | 0.122 | 8.22E-69 | DF 4 | spink2.1 |
| s100v2 | 8.4E-73 | 0.4763868 | 0.792 | 0.673 | 2.72E-68 | DF 4 | s100v2 |
| dse | 3.2E-72 | 1.4507673 | 0.137 | 0.051 | 1.03E-67 | DF 4 | dse |
| agmo2 | 1.1E-70 | 1.1988521 | 0.181 | 0.077 | 3.59E-66 | DF 4 | agmo |
| srpx21 | 1.4E-70 | 0.909286 | 0.401 | 0.247 | 4.64E-66 | DF 4 | srpx2 |
| col5a12 | 9.1E-69 | 0.2544644 | 0.995 | 0.963 | 2.96E-64 | DF 4 | col5a1 |
| si:dkey-240h12.43 | 1.9E-68 | 1.0429368 | 0.232 | 0.112 | 6.07E-64 | DF 4 | si:dkey-240h12.4 |
| pdlim21 | 8E-68 | 0.983452 | 0.281 | 0.15 | 2.6E-63 | DF 4 | pdlim2 |
| mgst1.21 | 1.1E-67 | 0.6806558 | 0.462 | 0.292 | 3.63E-63 | DF 4 | mgst1.2 |
| slc25a483 | 9.8E-67 | 1.0335797 | 0.257 | 0.132 | 3.2E-62 | DF 4 | slc25a48 |
| ccdc85ca | 8.1E-65 | 1.4099613 | 0.129 | 0.048 | 2.62E-60 | DF 4 | ccdc85ca |
| rhoq1 | 1.1E-64 | 1.0020815 | 0.197 | 0.09 | 3.42E-60 | DF 4 | rhoq |
| gch12 | 1.5E-64 | 1.0779719 | 0.251 | 0.13 | 4.94E-60 | DF 4 | gch1 |
| zgc:1530312 | 2.5E-64 | 1.2746905 | 0.145 | 0.058 | 8.13E-60 | DF 4 | zgc:153031 |
| ada1 | 3.5E-64 | 0.851936 | 0.358 | 0.214 | 1.14E-59 | DF 4 | ada |
| zgc:154164 | 8.5E-64 | 1.4318043 | 0.14 | 0.056 | 2.78E-59 | DF 4 | zgc:154164 |
| arhgap361 | 1.9E-63 | 1.1619974 | 0.146 | 0.058 | 6.28E-59 | DF 4 | arhgap36 |
| cd82b | 5.4E-63 | 0.6667284 | 0.523 | 0.367 | 1.74E-58 | DF 4 | cd82b |
| hsd11b23 | 7.9E-63 | 1.2600003 | 0.191 | 0.088 | 2.57E-58 | DF 4 | hsd11b2 |
| net12 | 1.4E-62 | 0.6901786 | 0.52 | 0.365 | 4.47E-58 | DF 4 | net1 |
| tril2 | 1.9E-62 | 0.8675964 | 0.213 | 0.101 | 6.21E-58 | DF 4 | tril |
| fgl2a3 | 7.8E-61 | 0.7061773 | 0.282 | 0.151 | 2.53E-56 | DF 4 | fgl2a |
| sh3bgrl | 2.1E-60 | 0.8219778 | 0.36 | 0.22 | 6.72E-56 | DF 4 | sh3bgrl |
| zbtb16a | 6E-60 | 1.0287164 | 0.239 | 0.125 | 1.94E-55 | DF 4 | zbtb16a |
| apoeb2 | 1.3E-59 | 1.0403096 | 0.385 | 0.241 | 4.3E-55 | DF 4 | apoeb |
| b2ml2 | 1.9E-59 | 0.2906369 | 0.942 | 0.895 | 6.06E-55 | DF 4 | b2ml |
| sod3b3 | 5.2E-59 | 1.1033199 | 0.227 | 0.118 | 1.7E-54 | DF 4 | sod3b |
| bmp52 | 5.5E-59 | 1.0152317 | 0.181 | 0.083 | 1.8E-54 | DF 4 | bmp5 |
| slc16a103 | 8.9E-59 | 0.982003 | 0.242 | 0.127 | 2.9E-54 | DF 4 | slc16a10 |
| zbtb16b1 | 1.2E-58 | 1.1243105 | 0.202 | 0.099 | 3.83E-54 | DF 4 | zbtb16b |
| caskin1 | 1.7E-58 | 1.5379542 | 0.133 | 0.054 | 5.39E-54 | DF 4 | caskin1 |
| gpc4 | 1.2E-57 | 0.8090103 | 0.373 | 0.236 | 4E-53 | DF 4 | gpc4 |
| si:ch211-106h4.122 | 1.3E-57 | 0.5111621 | 0.614 | 0.45 | 4.09E-53 | DF 4 | si:ch211-106h4.12 |
| igfbp31 | 2E-57 | 1.1550414 | 0.207 | 0.104 | 6.63E-53 | DF 4 | igfbp3 |
| fibina3 | 2.3E-57 | 0.5826013 | 0.347 | 0.206 | 7.37E-53 | DF 4 | fibina |
| acsl1a | 3.8E-57 | 1.2969809 | 0.159 | 0.07 | 1.22E-52 | DF 4 | acsl1a |

| | | | | | | | |
|---|---|---|---|---|---|---|---|
| fbxo32 | 4E-57 | 1.527985 | 0.112 | 0.042 | 1.31E-52 | DF 4 | fbxo32 |
| si:dkey-16p21.83 | 4.1E-57 | 0.788044 | 0.34 | 0.203 | 1.33E-52 | DF 4 | si:dkey-16p21.8 |
| tlcd4b2 | 3E-56 | 1.0322953 | 0.172 | 0.078 | 9.79E-52 | DF 4 | tlcd4b |
| pawr3 | 3.2E-56 | 0.7477765 | 0.313 | 0.18 | 1.04E-51 | DF 4 | pawr |
| creg22 | 3.8E-56 | 0.6920861 | 0.461 | 0.318 | 1.23E-51 | DF 4 | creg2 |
| lrrfip22 | 4.7E-56 | 1.1227784 | 0.2 | 0.1 | 1.52E-51 | DF 4 | lrrfip2 |
| CABZ01068358.11 | 1.3E-55 | 0.6820124 | 0.337 | 0.201 | 4.29E-51 | DF 4 | CABZ01068358.1 |
| lrp1aa2 | 2.3E-55 | 0.3834861 | 0.807 | 0.667 | 7.55E-51 | DF 4 | lrp1aa |
| twsg1a1 | 2.8E-55 | 0.7308408 | 0.39 | 0.251 | 9.21E-51 | DF 4 | twsg1a |
| raph1b | 3.1E-55 | 1.1352646 | 0.202 | 0.101 | 1.02E-50 | DF 4 | raph1b |
| ube2e3 | 4.6E-55 | 0.6926945 | 0.462 | 0.322 | 1.49E-50 | DF 4 | ube2e3 |
| runx32 | 7.9E-55 | 0.7620262 | 0.403 | 0.265 | 2.56E-50 | DF 4 | runx3 |
| igf2b2 | 1.5E-54 | 0.5361403 | 0.581 | 0.429 | 4.98E-50 | DF 4 | igf2b |
| gig2d1 | 5.5E-54 | 0.9764465 | 0.124 | 0.049 | 1.78E-49 | DF 4 | gig2d |
| tent5c | 6.3E-54 | 1.1436968 | 0.169 | 0.079 | 2.06E-49 | DF 4 | tent5c |
| frzb3 | 1.8E-53 | 0.250263 | 0.237 | 0.124 | 5.97E-49 | DF 4 | frzb |
| gstm.3 | 7.1E-53 | 1.1880212 | 0.119 | 0.047 | 2.31E-48 | DF 4 | gstm.3 |
| bhlhe401 | 9.7E-53 | 0.5479804 | 0.606 | 0.457 | 3.17E-48 | DF 4 | bhlhe40 |
| pbxip1a1 | 8.4E-52 | 0.8303673 | 0.302 | 0.181 | 2.75E-47 | DF 4 | pbxip1a |
| gpt2l3 | 3.3E-51 | 0.8993113 | 0.231 | 0.124 | 1.07E-46 | DF 4 | gpt2l |
| kremen11 | 4.2E-51 | 1.1676056 | 0.162 | 0.076 | 1.38E-46 | DF 4 | kremen1 |
| enpp12 | 4.7E-51 | 0.8849735 | 0.301 | 0.181 | 1.53E-46 | DF 4 | enpp1 |
| PPM1K1 | 7.6E-51 | 1.0994172 | 0.137 | 0.059 | 2.46E-46 | DF 4 | PPM1K |
| tmem198b1 | 9.3E-51 | 0.7929936 | 0.307 | 0.183 | 3.04E-46 | DF 4 | tmem198b |
| wnt7bb | 1.3E-50 | 1.4946981 | 0.101 | 0.038 | 4.29E-46 | DF 4 | wnt7bb |
| moxd1 | 1.9E-50 | 1.300786 | 0.12 | 0.049 | 6.02E-46 | DF 4 | moxd1 |
| serpina10a2 | 2.8E-50 | 0.5774617 | 0.515 | 0.371 | 9.19E-46 | DF 4 | serpina10a |
| gng13b | 3.6E-50 | 1.1931688 | 0.138 | 0.061 | 1.17E-45 | DF 4 | gng13b |
| pcolcea4 | 3.7E-50 | 0.3376241 | 0.72 | 0.56 | 1.21E-45 | DF 4 | pcolcea |
| si:ch211-288g17.31 | 6.1E-50 | 0.5670127 | 0.534 | 0.397 | 1.98E-45 | DF 4 | si:ch211-288g17.3 |
| sec61a1l1 | 1E-49 | 1.0314405 | 0.239 | 0.135 | 3.38E-45 | DF 4 | sec61a1l |
| zgc:56235 | 1.3E-49 | 0.9821976 | 0.212 | 0.112 | 4.26E-45 | DF 4 | zgc:56235 |
| ttyh3b | 2.4E-49 | 1.1191208 | 0.121 | 0.05 | 7.68E-45 | DF 4 | ttyh3b |
| mao2 | 2.8E-49 | 1.0542218 | 0.15 | 0.068 | 9.07E-45 | DF 4 | mao |
| pcm13 | 7.2E-49 | 0.808678 | 0.258 | 0.147 | 2.33E-44 | DF 4 | pcm1 |
| frem1b2 | 4.4E-48 | 1.0453991 | 0.157 | 0.074 | 1.42E-43 | DF 4 | frem1b |
| rnd3b2 | 8.3E-48 | 0.5755083 | 0.462 | 0.324 | 2.7E-43 | DF 4 | rnd3b |
| zbtb201 | 1.7E-47 | 0.5741024 | 0.506 | 0.368 | 5.54E-43 | DF 4 | zbtb20 |
| atp1b3b2 | 7.2E-47 | 1.168646 | 0.156 | 0.075 | 2.33E-42 | DF 4 | atp1b3b |
| ponzr12 | 9.3E-47 | 0.9113338 | 0.355 | 0.227 | 3.02E-42 | DF 4 | ponzr1 |
| ptgr12 | 1.3E-46 | 1.0050892 | 0.236 | 0.135 | 4.2E-42 | DF 4 | ptgr1 |
| sdc21 | 3.4E-46 | 0.4327093 | 0.705 | 0.582 | 1.1E-41 | DF 4 | sdc2 |
| zfhx42 | 9.6E-45 | 0.4802871 | 0.581 | 0.443 | 3.11E-40 | DF 4 | zfhx4 |
| sfrp1b3 | 9.9E-45 | 0.2960378 | 0.517 | 0.353 | 3.21E-40 | DF 4 | sfrp1b |
| CU638870.12 | 1.5E-44 | 0.5819871 | 0.456 | 0.322 | 5.02E-40 | DF 4 | CU638870.1 |
| per31 | 3E-44 | 0.5887044 | 0.472 | 0.34 | 9.83E-40 | DF 4 | per3 |
| itgb52 | 3.3E-44 | 1.2044969 | 0.161 | 0.081 | 1.06E-39 | DF 4 | itgb5 |
| inka1a1 | 3.6E-44 | 0.8257784 | 0.28 | 0.17 | 1.18E-39 | DF 4 | inka1a |

| | | | | | | | |
|---|---|---|---|---|---|---|---|
| fuca2 | 6.3E-44 | 0.8716382 | 0.243 | 0.141 | 2.06E-39 | DF 4 | fuca2 |
| ssuh2rs1 | 1.7E-43 | 1.1299507 | 0.138 | 0.064 | 5.68E-39 | DF 4 | ssuh2rs1 |
| syt11a2 | 2E-43 | 0.5248769 | 0.275 | 0.159 | 6.57E-39 | DF 4 | syt11a |
| thraa | 4.2E-43 | 1.2417881 | 0.127 | 0.057 | 1.38E-38 | DF 4 | thraa |
| mettl7a.1 | 4.4E-43 | 1.00264 | 0.135 | 0.062 | 1.44E-38 | DF 4 | mettl7a.1 |
| si:ch211-262i1.42 | 5.6E-43 | 0.7766388 | 0.158 | 0.077 | 1.81E-38 | DF 4 | si:ch211-262i1.4 |
| gphna1 | 5.7E-43 | 1.5202943 | 0.117 | 0.052 | 1.85E-38 | DF 4 | gphna |
| dysf | 6E-43 | 1.314445 | 0.103 | 0.042 | 1.94E-38 | DF 4 | dysf |
| olfml3b2 | 7.8E-43 | 0.4074854 | 0.636 | 0.508 | 2.54E-38 | DF 4 | olfml3b |
| slc29a1a3 | 1.8E-42 | 0.6796844 | 0.307 | 0.192 | 5.84E-38 | DF 4 | slc29a1a |
| si:ch211-132b12.71 | 6.2E-42 | 0.8396573 | 0.254 | 0.151 | 2.02E-37 | DF 4 | si:ch211-132b12.7 |
| zgc:1135313 | 7.4E-42 | 0.5482472 | 0.404 | 0.272 | 2.39E-37 | DF 4 | zgc:113531 |
| slc7a23 | 1.8E-41 | 0.606753 | 0.263 | 0.154 | 5.93E-37 | DF 4 | slc7a2 |
| clec11a | 1.8E-41 | 1.191628 | 0.12 | 0.054 | 5.95E-37 | DF 4 | clec11a |
| ccn2a1 | 2.4E-41 | 0.53677 | 0.315 | 0.201 | 7.72E-37 | DF 4 | ccn2a |
| ak43 | 3.2E-41 | 0.7110678 | 0.289 | 0.18 | 1.05E-36 | DF 4 | ak4 |
| cbr1l2 | 4.7E-41 | 0.7515489 | 0.319 | 0.209 | 1.52E-36 | DF 4 | cbr1l |
| thrb1 | 6.6E-41 | 0.7656709 | 0.265 | 0.161 | 2.14E-36 | DF 4 | thrb |
| en1b1 | 7.1E-41 | 1.0654755 | 0.193 | 0.106 | 2.32E-36 | DF 4 | en1b |
| cemip2 | 3.7E-40 | 1.1763082 | 0.141 | 0.069 | 1.2E-35 | DF 4 | cemip2 |
| prrx1b2 | 5.1E-40 | 0.4484674 | 0.513 | 0.373 | 1.66E-35 | DF 4 | prrx1b |
| tpm22 | 7E-40 | 0.7993331 | 0.202 | 0.112 | 2.29E-35 | DF 4 | tpm2 |
| mob1bb | 1.2E-39 | 1.0254195 | 0.155 | 0.079 | 3.8E-35 | DF 4 | mob1bb |
| gulp1a1 | 1.4E-39 | 0.7531122 | 0.287 | 0.183 | 4.62E-35 | DF 4 | gulp1a |
| hmgn33 | 2.5E-39 | 0.7890834 | 0.258 | 0.157 | 8.21E-35 | DF 4 | hmgn3 |
| kif26ba3 | 2.8E-39 | 0.7600679 | 0.191 | 0.104 | 9.03E-35 | DF 4 | kif26ba |
| plpbp | 4E-39 | 1.2368793 | 0.101 | 0.043 | 1.29E-34 | DF 4 | plpbp |
| angptl52 | 5.4E-39 | 0.6390682 | 0.274 | 0.168 | 1.76E-34 | DF 4 | angptl5 |
| qsox11 | 1.6E-38 | 1.1197664 | 0.135 | 0.065 | 5.04E-34 | DF 4 | qsox1 |
| fabp33 | 3.1E-38 | 0.4160786 | 0.643 | 0.525 | 1E-33 | DF 4 | fabp3 |
| myl9b1 | 4.9E-38 | 0.4478028 | 0.542 | 0.419 | 1.6E-33 | DF 4 | myl9b |
| si:ch211-133n4.42 | 6.2E-38 | 0.4737526 | 0.562 | 0.442 | 2.01E-33 | DF 4 | si:ch211-133n4.4 |
| dhrs94 | 1.3E-37 | 0.7746523 | 0.231 | 0.136 | 4.23E-33 | DF 4 | dhrs9 |
| itm2bb2 | 1.4E-37 | 0.8345353 | 0.164 | 0.086 | 4.42E-33 | DF 4 | itm2bb |
| stox2b | 2.1E-37 | 1.1834976 | 0.102 | 0.044 | 6.78E-33 | DF 4 | stox2b |
| zgc:1140462 | 2.3E-37 | 1.0268779 | 0.176 | 0.096 | 7.45E-33 | DF 4 | zgc:114046 |
| scinlb3 | 3.6E-37 | 0.5153787 | 0.425 | 0.302 | 1.16E-32 | DF 4 | scinlb |
| CU927890.1 | 5.1E-37 | 1.0585518 | 0.119 | 0.056 | 1.65E-32 | DF 4 | CU927890.1 |
| tagln21 | 1.5E-36 | 0.3546257 | 0.759 | 0.661 | 4.92E-32 | DF 4 | tagln2 |
| slc7a8a2 | 1.7E-36 | 0.920593 | 0.156 | 0.082 | 5.67E-32 | DF 4 | slc7a8a |
| itga6b2 | 4.8E-36 | 0.4054694 | 0.42 | 0.29 | 1.57E-31 | DF 4 | itga6b |
| adamts15b.11 | 4.8E-36 | 0.7605273 | 0.167 | 0.089 | 1.58E-31 | DF 4 | adamts15b.1 |
| ms4a17a.72 | 6.6E-36 | 0.5188839 | 0.412 | 0.291 | 2.16E-31 | DF 4 | ms4a17a.7 |
| dlx4a5 | 8E-36 | 0.6970302 | 0.29 | 0.188 | 2.61E-31 | DF 4 | dlx4a |
| aoc22 | 8.7E-36 | 0.4795976 | 0.354 | 0.235 | 2.84E-31 | DF 4 | aoc2 |
| lrrc58b1 | 1.7E-35 | 0.8348019 | 0.204 | 0.118 | 5.5E-31 | DF 4 | lrrc58b |
| si:dkey-7j14.61 | 1.9E-35 | 0.2882888 | 0.836 | 0.774 | 6.13E-31 | DF 4 | si:dkey-7j14.6 |
| mmp14b2 | 4.5E-35 | 0.3512859 | 0.666 | 0.544 | 1.45E-30 | DF 4 | mmp14b |

| | | | | | | | |
|---|---|---|---|---|---|---|---|
| hhip1 | 7.1E-35 | 1.0063062 | 0.159 | 0.085 | 2.32E-30 | DF 4 | hhip |
| dbpb2 | 9.5E-35 | 0.8111776 | 0.191 | 0.109 | 3.09E-30 | DF 4 | dbpb |
| capn2b4 | 1E-34 | 0.4698029 | 0.442 | 0.316 | 3.4E-30 | DF 4 | capn2b |
| CT956002.2 | 1.4E-34 | 0.9078105 | 0.141 | 0.072 | 4.64E-30 | DF 4 | CT956002.2 |
| dio12 | 1.8E-34 | 0.7360529 | 0.204 | 0.118 | 5.99E-30 | DF 4 | dio1 |
| limch1a2 | 1.9E-34 | 1.0454727 | 0.143 | 0.075 | 6.06E-30 | DF 4 | limch1a |
| kdm6bb2 | 4.6E-34 | 0.7092639 | 0.226 | 0.136 | 1.48E-29 | DF 4 | kdm6bb |
| nrarpa | 1.8E-33 | 1.0098465 | 0.156 | 0.085 | 5.89E-29 | DF 4 | nrarpa |
| plxdc21 | 1.9E-33 | 0.7068208 | 0.246 | 0.153 | 6.27E-29 | DF 4 | plxdc2 |
| ctsla2 | 3.1E-33 | 0.2907859 | 0.805 | 0.717 | 1.02E-28 | DF 4 | ctsla |
| svila3 | 3.8E-33 | 0.6586992 | 0.235 | 0.144 | 1.25E-28 | DF 4 | svila |
| fosb2 | 4.4E-33 | 0.2709385 | 0.782 | 0.688 | 1.45E-28 | DF 4 | fosb |
| ccnd2a1 | 4.5E-33 | 0.6116688 | 0.316 | 0.213 | 1.48E-28 | DF 4 | ccnd2a |
| wipf1b | 7.1E-33 | 0.8708569 | 0.178 | 0.101 | 2.3E-28 | DF 4 | wipf1b |
| smurf21 | 9.3E-33 | 1.0009794 | 0.126 | 0.063 | 3.01E-28 | DF 4 | smurf2 |
| emp2 | 1E-32 | 0.742979 | 0.236 | 0.148 | 3.31E-28 | DF 4 | emp2 |
| mgst1.12 | 2.1E-32 | 0.5114966 | 0.355 | 0.245 | 6.9E-28 | DF 4 | mgst1.1 |
| gpc1b1 | 4.1E-32 | 0.4115511 | 0.517 | 0.396 | 1.33E-27 | DF 4 | gpc1b |
| irf2bpl | 4.9E-32 | 0.7568847 | 0.288 | 0.194 | 1.58E-27 | DF 4 | irf2bpl |
| dynll2b | 4.9E-32 | 0.7399365 | 0.238 | 0.15 | 1.6E-27 | DF 4 | dynll2b |
| h3f3c1 | 6.5E-32 | 0.451838 | 0.565 | 0.46 | 2.1E-27 | DF 4 | h3f3c |
| f102 | 7E-32 | 0.4569634 | 0.505 | 0.394 | 2.27E-27 | DF 4 | f10 |
| calm3a.1 | 7.5E-32 | 0.4736776 | 0.499 | 0.393 | 2.45E-27 | DF 4 | calm3a.1 |
| C20H1orf1981 | 8.7E-32 | 0.7212806 | 0.272 | 0.18 | 2.83E-27 | DF 4 | C20H1orf198 |
| roraa | 2.1E-31 | 0.7702831 | 0.223 | 0.137 | 6.79E-27 | DF 4 | roraa |
| hist2h2l1 | 2.1E-31 | 0.4728724 | 0.478 | 0.369 | 6.96E-27 | DF 4 | hist2h2l |
| adgrl2a3 | 2.7E-31 | 0.6396462 | 0.282 | 0.186 | 8.68E-27 | DF 4 | adgrl2a |
| dedd12 | 3.4E-31 | 0.3186382 | 0.66 | 0.547 | 1.11E-26 | DF 4 | dedd1 |
| rnf213a2 | 6.4E-31 | 0.5456363 | 0.371 | 0.264 | 2.09E-26 | DF 4 | rnf213a |
| limk21 | 1.1E-30 | 0.9882384 | 0.143 | 0.078 | 3.55E-26 | DF 4 | limk2 |
| ephb2b1 | 1.8E-30 | 0.5954137 | 0.292 | 0.195 | 5.85E-26 | DF 4 | ephb2b |
| CR759885.13 | 2.1E-30 | 0.7281788 | 0.171 | 0.097 | 6.79E-26 | DF 4 | CR759885.1 |
| trim71 | 3.6E-30 | 0.8746735 | 0.172 | 0.099 | 1.17E-25 | DF 4 | trim71 |
| hyal2b3 | 4E-30 | 0.5150114 | 0.291 | 0.192 | 1.31E-25 | DF 4 | hyal2b |
| mxra5b1 | 4.1E-30 | 0.8252307 | 0.115 | 0.057 | 1.35E-25 | DF 4 | mxra5b |
| pde3b1 | 8.8E-30 | 0.7984161 | 0.194 | 0.117 | 2.86E-25 | DF 4 | pde3b |
| ehd1b | 1E-29 | 0.7017456 | 0.259 | 0.171 | 3.32E-25 | DF 4 | ehd1b |
| clstn1 | 1.3E-29 | 0.6705874 | 0.215 | 0.132 | 4.38E-25 | DF 4 | clstn1 |
| fam102ab | 1.5E-29 | 1.0407062 | 0.123 | 0.064 | 4.97E-25 | DF 4 | fam102ab |
| prkab1b | 1.9E-29 | 0.5732264 | 0.37 | 0.273 | 6.23E-25 | DF 4 | prkab1b |
| pbx1a2 | 2E-29 | 0.5820152 | 0.324 | 0.228 | 6.64E-25 | DF 4 | pbx1a |
| uacab2 | 2.5E-29 | 0.9113757 | 0.132 | 0.07 | 8.2E-25 | DF 4 | uacab |
| homer3b | 6.1E-29 | 0.6709449 | 0.226 | 0.143 | 1.98E-24 | DF 4 | homer3b |
| col16a12 | 6.2E-29 | 0.3168668 | 0.573 | 0.449 | 2E-24 | DF 4 | col16a1 |
| cmtm32 | 6.2E-29 | 0.5645492 | 0.283 | 0.19 | 2.01E-24 | DF 4 | cmtm3 |
| hpgd | 6.9E-29 | 0.8345889 | 0.108 | 0.053 | 2.24E-24 | DF 4 | hpgd |
| adamts5 | 6.9E-29 | 0.8832216 | 0.121 | 0.062 | 2.25E-24 | DF 4 | adamts5 |
| cdkn1ba2 | 8.7E-29 | 0.8959073 | 0.187 | 0.114 | 2.84E-24 | DF 4 | cdkn1ba |

| | | | | | | | |
|---|---|---|---|---|---|---|---|
| asip2b1 | 9.7E-29 | 0.2475861 | 0.215 | 0.131 | 3.14E-24 | DF 4 | asip2b |
| serpinf13 | 1.2E-28 | 0.2583615 | 0.669 | 0.548 | 3.81E-24 | DF 4 | serpinf1 |
| dhrs3a1 | 1.6E-28 | 0.5239687 | 0.126 | 0.065 | 5.1E-24 | DF 4 | dhrs3a |
| usp2a | 1.7E-28 | 1.1472893 | 0.127 | 0.068 | 5.61E-24 | DF 4 | usp2a |
| nid1b1 | 2.3E-28 | 0.6653755 | 0.141 | 0.076 | 7.38E-24 | DF 4 | nid1b |
| pbx3b3 | 2.6E-28 | 0.6981043 | 0.22 | 0.139 | 8.42E-24 | DF 4 | pbx3b |
| prdm1a2 | 2.9E-28 | 0.7776347 | 0.137 | 0.074 | 9.38E-24 | DF 4 | prdm1a |
| ap1s2 | 6.2E-28 | 0.6882967 | 0.242 | 0.16 | 2.01E-23 | DF 4 | ap1s2 |
| mdh1aa1 | 7E-28 | 0.5388186 | 0.464 | 0.369 | 2.28E-23 | DF 4 | mdh1aa |
| cpamd8 | 8.6E-28 | 0.7316467 | 0.114 | 0.057 | 2.81E-23 | DF 4 | cpamd8 |
| KIF2A | 1.3E-27 | 0.8375927 | 0.177 | 0.106 | 4.28E-23 | DF 4 | KIF2A |
| ifitm15 | 2.1E-27 | 0.291887 | 0.91 | 0.783 | 6.85E-23 | DF 4 | ifitm1 |
| tnrc6c11 | 2.2E-27 | 0.3793051 | 0.581 | 0.481 | 7.25E-23 | DF 4 | tnrc6c1 |
| rgmd2 | 2.3E-27 | 0.8197011 | 0.146 | 0.082 | 7.53E-23 | DF 4 | rgmd |
| zfp36l1a4 | 3.2E-27 | 0.553979 | 0.295 | 0.202 | 1.04E-22 | DF 4 | zfp36l1a |
| ctdspla1 | 6.3E-27 | 0.8009518 | 0.183 | 0.111 | 2.04E-22 | DF 4 | ctdspla |
| rnf44 | 6.6E-27 | 0.7579984 | 0.189 | 0.117 | 2.16E-22 | DF 4 | rnf44 |
| ifi461 | 9.1E-27 | 0.8440089 | 0.158 | 0.092 | 2.95E-22 | DF 4 | ifi46 |
| aldob4 | 1.4E-26 | 0.3772608 | 0.414 | 0.306 | 4.52E-22 | DF 4 | aldob |
| skilb | 1.6E-26 | 0.9641418 | 0.136 | 0.076 | 5.32E-22 | DF 4 | skilb |
| nucks1b | 1.7E-26 | 0.4806777 | 0.413 | 0.313 | 5.45E-22 | DF 4 | nucks1b |
| jag1b1 | 2.8E-26 | 0.4934945 | 0.305 | 0.211 | 9.25E-22 | DF 4 | jag1b |
| tnfrsf192 | 3.1E-26 | 0.6514155 | 0.176 | 0.105 | 1E-21 | DF 4 | tnfrsf19 |
| napsa1 | 4.3E-26 | 0.5912696 | 0.249 | 0.166 | 1.39E-21 | DF 4 | napsa |
| hist1h4l.10 | 5.3E-26 | 0.8164471 | 0.13 | 0.071 | 1.72E-21 | DF 4 | hist1h4l.10 |
| nsd1b | 5.3E-26 | 0.7770704 | 0.13 | 0.071 | 1.73E-21 | DF 4 | nsd1b |
| tfap2c2 | 8.5E-26 | 0.4797356 | 0.281 | 0.192 | 2.75E-21 | DF 4 | tfap2c |
| ptbp1b1 | 9.2E-26 | 0.5824134 | 0.271 | 0.185 | 2.99E-21 | DF 4 | ptbp1b |
| ndrg21 | 1.8E-25 | 0.4581427 | 0.414 | 0.318 | 5.8E-21 | DF 4 | ndrg2 |
| msx1b2 | 1.8E-25 | 0.4430316 | 0.387 | 0.289 | 5.96E-21 | DF 4 | msx1b |
| spon2b2 | 2.8E-25 | 0.68071 | 0.198 | 0.124 | 9.03E-21 | DF 4 | spon2b |
| chd53 | 3.7E-25 | 0.7301019 | 0.196 | 0.123 | 1.21E-20 | DF 4 | chd5 |
| si:cabz01069012.22 | 7E-25 | 0.5801776 | 0.168 | 0.1 | 2.27E-20 | DF 4 | si:cabz01069012.2 |
| zfhx32 | 8.2E-25 | 0.2912178 | 0.589 | 0.482 | 2.68E-20 | DF 4 | zfhx3 |
| ppm1la | 9.3E-25 | 0.9338042 | 0.126 | 0.07 | 3.02E-20 | DF 4 | ppm1la |
| nr3c1 | 9.6E-25 | 0.5761577 | 0.283 | 0.197 | 3.11E-20 | DF 4 | nr3c1 |
| zgc:1735871 | 1E-24 | 0.8274201 | 0.199 | 0.128 | 3.28E-20 | DF 4 | zgc:173587 |
| si:ch73-1a9.32 | 1E-24 | 0.2149136 | 0.806 | 0.715 | 3.4E-20 | DF 4 | si:ch73-1a9.3 |
| cbsb3 | 1.3E-24 | 0.4827718 | 0.213 | 0.135 | 4.12E-20 | DF 4 | cbsb |
| paip2b | 1.3E-24 | 0.4142028 | 0.465 | 0.373 | 4.12E-20 | DF 4 | paip2b |
| dvl3a | 1.8E-24 | 0.9317482 | 0.136 | 0.078 | 5.9E-20 | DF 4 | dvl3a |
| si:ch211-232b12.5 | 1.8E-24 | 0.7323471 | 0.228 | 0.153 | 5.97E-20 | DF 4 | si:ch211-232b12.5 |
| hmcn12 | 2E-24 | 0.7861927 | 0.126 | 0.07 | 6.48E-20 | DF 4 | hmcn1 |
| ognb3 | 2.1E-24 | 0.3524276 | 0.206 | 0.129 | 6.93E-20 | DF 4 | ognb |
| ISCU | 2.6E-24 | 0.4662972 | 0.353 | 0.261 | 8.59E-20 | DF 4 | ISCU |
| spry21 | 2.6E-24 | 0.8600134 | 0.133 | 0.076 | 8.6E-20 | DF 4 | spry2 |
| rab11fip1a | 4.4E-24 | 0.8957106 | 0.11 | 0.059 | 1.45E-19 | DF 4 | rab11fip1a |
| lpar12 | 5.3E-24 | 0.549725 | 0.26 | 0.178 | 1.71E-19 | DF 4 | lpar1 |

| | | | | | | | |
|---|---|---|---|---|---|---|---|
| ivd | 9.3E-24 | 0.8287407 | 0.146 | 0.086 | 3.02E-19 | DF 4 | ivd |
| ccdc85b3 | 9.7E-24 | 0.5491307 | 0.269 | 0.186 | 3.16E-19 | DF 4 | ccdc85b |
| si:ch211-246m6.5 | 1E-23 | 0.6771282 | 0.101 | 0.052 | 3.25E-19 | DF 4 | si:ch211-246m6.5 |
| tefa | 1.4E-23 | 0.5012253 | 0.301 | 0.215 | 4.65E-19 | DF 4 | tefa |
| rtn4b1 | 1.5E-23 | 0.9230645 | 0.149 | 0.09 | 5.01E-19 | DF 4 | rtn4b |
| col21a12 | 1.8E-23 | 0.7009549 | 0.135 | 0.077 | 5.73E-19 | DF 4 | col21a1 |
| lef11 | 1.8E-23 | 0.7305292 | 0.17 | 0.105 | 5.93E-19 | DF 4 | lef1 |
| tob1a1 | 1.8E-23 | 0.260481 | 0.702 | 0.605 | 6.02E-19 | DF 4 | tob1a |
| pgrmc1 | 1.9E-23 | 0.4428549 | 0.449 | 0.36 | 6.13E-19 | DF 4 | pgrmc1 |
| si:ch73-386h18.13 | 1.9E-23 | 0.5455011 | 0.22 | 0.144 | 6.29E-19 | DF 4 | si:ch73-386h18.1 |
| C24H8orf44 | 2.1E-23 | 0.8727477 | 0.149 | 0.089 | 6.85E-19 | DF 4 | C24H8orf44 |
| smad3a1 | 2.3E-23 | 0.4029172 | 0.397 | 0.301 | 7.41E-19 | DF 4 | smad3a |
| trps12 | 4E-23 | 0.5805722 | 0.249 | 0.17 | 1.3E-18 | DF 4 | trps1 |
| stau2 | 4.1E-23 | 0.3406722 | 0.531 | 0.434 | 1.34E-18 | DF 4 | stau2 |
| h2ax11 | 6.7E-23 | 0.3169852 | 0.697 | 0.625 | 2.18E-18 | DF 4 | h2ax1 |
| ror2 | 1.1E-22 | 0.6986879 | 0.128 | 0.073 | 3.62E-18 | DF 4 | ror2 |
| kazald34 | 1.1E-22 | 0.2008351 | 0.36 | 0.256 | 3.65E-18 | DF 4 | kazald3 |
| prickle1a | 1.4E-22 | 1.0203645 | 0.105 | 0.057 | 4.45E-18 | DF 4 | prickle1a |
| lpp3 | 1.7E-22 | 0.5526583 | 0.334 | 0.249 | 5.45E-18 | DF 4 | lpp |
| si:ch73-237c6.13 | 3.3E-22 | 0.2684717 | 0.471 | 0.36 | 1.07E-17 | DF 4 | si:ch73-237c6.1 |
| si:dkeyp-44a8.21 | 4E-22 | 0.664248 | 0.203 | 0.135 | 1.31E-17 | DF 4 | si:dkeyp-44a8.2 |
| gcdha1 | 5E-22 | 0.7111097 | 0.168 | 0.106 | 1.62E-17 | DF 4 | gcdha |
| efna5b | 5.7E-22 | 0.8139546 | 0.119 | 0.067 | 1.87E-17 | DF 4 | efna5b |
| cebpd3 | 7.6E-22 | 0.2748011 | 0.666 | 0.573 | 2.47E-17 | DF 4 | cebpd |
| gadd45ga2 | 8.7E-22 | 0.5533188 | 0.361 | 0.273 | 2.83E-17 | DF 4 | gadd45ga |
| zbtb18 | 2.2E-21 | 0.9334135 | 0.104 | 0.057 | 7.11E-17 | DF 4 | zbtb18 |
| cry1b | 2.8E-21 | 0.8893791 | 0.1 | 0.054 | 9.15E-17 | DF 4 | cry1b |
| cilp3 | 2.8E-21 | 0.4137851 | 0.245 | 0.169 | 9.16E-17 | DF 4 | cilp |
| ciarta2 | 3E-21 | 0.4336797 | 0.368 | 0.279 | 9.85E-17 | DF 4 | ciarta |
| tsc22d11 | 3.4E-21 | 0.5774704 | 0.177 | 0.112 | 1.12E-16 | DF 4 | tsc22d1 |
| rgn | 5.5E-21 | 0.7502346 | 0.122 | 0.07 | 1.8E-16 | DF 4 | rgn |
| CR847566.12 | 5.7E-21 | 0.6497749 | 0.163 | 0.102 | 1.86E-16 | DF 4 | CR847566.1 |
| tgfb1a1 | 7.4E-21 | 0.3375447 | 0.361 | 0.27 | 2.41E-16 | DF 4 | tgfb1a |
| mxra8b2 | 9.3E-21 | 0.2235623 | 0.502 | 0.404 | 3.01E-16 | DF 4 | mxra8b |
| tmem1763 | 1.1E-20 | 0.2468494 | 0.496 | 0.389 | 3.69E-16 | DF 4 | tmem176 |
| wipi11 | 2.2E-20 | 0.7671769 | 0.18 | 0.119 | 7.05E-16 | DF 4 | wipi1 |
| yap1 | 2.3E-20 | 0.3671717 | 0.444 | 0.355 | 7.52E-16 | DF 4 | yap1 |
| si:dkey-164f24.24 | 2.9E-20 | 0.6214487 | 0.241 | 0.167 | 9.38E-16 | DF 4 | si:dkey-164f24.2 |
| si:dkey-81l17.61 | 3.1E-20 | 0.8610274 | 0.127 | 0.076 | 1.01E-15 | DF 4 | si:dkey-81l17.6 |
| ubtd21 | 3.5E-20 | 0.6410115 | 0.16 | 0.101 | 1.13E-15 | DF 4 | ubtd2 |
| bcat2 | 4.1E-20 | 0.7871044 | 0.141 | 0.086 | 1.32E-15 | DF 4 | bcat2 |
| ddt1 | 4.1E-20 | 0.539125 | 0.264 | 0.19 | 1.33E-15 | DF 4 | ddt |
| ubl3a | 4.6E-20 | 0.360336 | 0.422 | 0.333 | 1.5E-15 | DF 4 | ubl3a |
| pthlha2 | 4.8E-20 | 0.5890227 | 0.121 | 0.07 | 1.55E-15 | DF 4 | pthlha |
| zbtb41 | 7.1E-20 | 0.3393988 | 0.411 | 0.321 | 2.3E-15 | DF 4 | zbtb4 |
| slc12a23 | 8.3E-20 | 0.4385699 | 0.21 | 0.141 | 2.69E-15 | DF 4 | slc12a2 |
| si:ch211-110e21.33 | 8.4E-20 | 0.7699386 | 0.126 | 0.074 | 2.74E-15 | DF 4 | si:ch211-110e21.3 |
| rassf2a1 | 1.9E-19 | 0.5832165 | 0.186 | 0.123 | 6.07E-15 | DF 4 | rassf2a |

| | | | | | | | |
|---|---|---|---|---|---|---|---|
| znf609a2 | 1.9E-19 | 0.503625 | 0.245 | 0.172 | 6.15E-15 | DF 4 | znf609a |
| luzp12 | 3.1E-19 | 0.541402 | 0.241 | 0.172 | 1E-14 | DF 4 | luzp1 |
| smc51 | 6.3E-19 | 0.6371187 | 0.14 | 0.086 | 2.04E-14 | DF 4 | smc5 |
| selenbp11 | 6.4E-19 | 0.7501286 | 0.126 | 0.076 | 2.09E-14 | DF 4 | selenbp1 |
| si:dkey-282h22.52 | 6.5E-19 | 0.5806306 | 0.117 | 0.068 | 2.1E-14 | DF 4 | si:dkey-282h22.5 |
| mocs22 | 7.8E-19 | 0.706369 | 0.138 | 0.085 | 2.54E-14 | DF 4 | mocs2 |
| sfrp52 | 1.6E-18 | 0.2218095 | 0.282 | 0.206 | 5.07E-14 | DF 4 | sfrp5 |
| phactr4b1 | 1.7E-18 | 0.7549766 | 0.119 | 0.071 | 5.49E-14 | DF 4 | phactr4b |
| itgb23 | 2.1E-18 | 0.4314031 | 0.162 | 0.103 | 6.69E-14 | DF 4 | itgb2 |
| pbx4 | 4E-18 | 0.4477391 | 0.332 | 0.256 | 1.31E-13 | DF 4 | pbx4 |
| hopx1 | 5E-18 | 0.343667 | 0.351 | 0.266 | 1.64E-13 | DF 4 | hopx |
| ENSDARG0000010520 | 5.6E-18 | 0.7187921 | 0.103 | 0.059 | 1.81E-13 | DF 4 | ENSDARG00000105 |
| sulf2b2 | 6.5E-18 | 0.3860192 | 0.232 | 0.162 | 2.12E-13 | DF 4 | sulf2b |
| ctnnb23 | 8.5E-18 | 0.446953 | 0.384 | 0.309 | 2.77E-13 | DF 4 | ctnnb2 |
| ecm23 | 9.8E-18 | 0.3318555 | 0.379 | 0.294 | 3.2E-13 | DF 4 | ecm2 |
| wnt5b | 1.1E-17 | 0.5839478 | 0.164 | 0.107 | 3.46E-13 | DF 4 | wnt5b |
| asah1b2 | 1.2E-17 | 0.522166 | 0.25 | 0.182 | 3.96E-13 | DF 4 | asah1b |
| unkl | 1.6E-17 | 0.6942882 | 0.101 | 0.058 | 5.13E-13 | DF 4 | unkl |
| ptenb1 | 1.6E-17 | 0.5272634 | 0.238 | 0.172 | 5.34E-13 | DF 4 | ptenb |
| CABZ01070631.2 | 2.2E-17 | 0.7211738 | 0.103 | 0.06 | 6.99E-13 | DF 4 | CABZ01070631.2 |
| fbxo251 | 2.4E-17 | 0.731434 | 0.125 | 0.077 | 7.76E-13 | DF 4 | fbxo25 |
| c1qtnf13 | 2.7E-17 | 0.6260523 | 0.121 | 0.073 | 8.78E-13 | DF 4 | c1qtnf1 |
| rad54l21 | 2.8E-17 | 0.7800728 | 0.13 | 0.081 | 9.13E-13 | DF 4 | rad54l2 |
| hmgn71 | 2.9E-17 | 0.3013882 | 0.592 | 0.52 | 9.5E-13 | DF 4 | hmgn7 |
| deptor1 | 3.2E-17 | 0.7470957 | 0.115 | 0.069 | 1.05E-12 | DF 4 | deptor |
| plcd1a1 | 3.7E-17 | 0.5918512 | 0.147 | 0.094 | 1.19E-12 | DF 4 | plcd1a |
| adka1 | 4.6E-17 | 0.8788517 | 0.192 | 0.135 | 1.48E-12 | DF 4 | adka |
| mfap23 | 5.1E-17 | 0.2966702 | 0.472 | 0.38 | 1.65E-12 | DF 4 | mfap2 |
| smc31 | 5.6E-17 | 0.5074195 | 0.205 | 0.143 | 1.84E-12 | DF 4 | smc3 |
| dbpa | 5.7E-17 | 0.6858701 | 0.127 | 0.078 | 1.87E-12 | DF 4 | dbpa |
| pgrmc2 | 5.9E-17 | 0.5410924 | 0.198 | 0.137 | 1.92E-12 | DF 4 | pgrmc2 |
| gabarapb | 7.8E-17 | 0.3314797 | 0.496 | 0.421 | 2.54E-12 | DF 4 | gabarapb |
| lamp2 | 8.5E-17 | 0.3649732 | 0.423 | 0.346 | 2.77E-12 | DF 4 | lamp2 |
| rorab | 8.6E-17 | 0.5587484 | 0.181 | 0.123 | 2.81E-12 | DF 4 | rorab |
| hmgb1b1 | 9.7E-17 | 0.4536147 | 0.321 | 0.249 | 3.15E-12 | DF 4 | hmgb1b |
| prdx51 | 9.9E-17 | 0.2306147 | 0.66 | 0.587 | 3.23E-12 | DF 4 | prdx5 |
| zmat5 | 1.1E-16 | 0.7834045 | 0.118 | 0.072 | 3.46E-12 | DF 4 | zmat5 |
| chst14 | 1.1E-16 | 0.7356496 | 0.115 | 0.07 | 3.64E-12 | DF 4 | chst14 |
| emx22 | 1.1E-16 | 0.5083042 | 0.142 | 0.09 | 3.65E-12 | DF 4 | emx2 |
| evla1 | 1.4E-16 | 0.7075897 | 0.134 | 0.085 | 4.42E-12 | DF 4 | evla |
| creg1 | 1.5E-16 | 0.7242642 | 0.116 | 0.07 | 4.78E-12 | DF 4 | creg1 |
| rn7sk4 | 1.8E-16 | 0.3188136 | 0.539 | 0.452 | 5.78E-12 | DF 4 | rn7sk |
| cbx1a1 | 1.9E-16 | 0.4882973 | 0.269 | 0.201 | 6.31E-12 | DF 4 | cbx1a |
| tmtc12 | 3.3E-16 | 0.4653357 | 0.171 | 0.114 | 1.07E-11 | DF 4 | tmtc1 |
| tcima3 | 3.6E-16 | 0.3792041 | 0.295 | 0.223 | 1.18E-11 | DF 4 | tcima |
| ddx62 | 3.9E-16 | 0.3172067 | 0.424 | 0.345 | 1.27E-11 | DF 4 | ddx6 |
| ivns1abpa2 | 4.1E-16 | 0.4705201 | 0.354 | 0.283 | 1.33E-11 | DF 4 | ivns1abpa |
| ahr23 | 4.6E-16 | 0.3374121 | 0.459 | 0.38 | 1.5E-11 | DF 4 | ahr2 |

| | | | | | | | |
|---|---|---|---|---|---|---|---|
| map1lc3a | 5.1E-16 | 0.4658085 | 0.278 | 0.211 | 1.65E-11 | DF 4 | map1lc3a |
| cmbl2 | 5.4E-16 | 0.4806555 | 0.237 | 0.172 | 1.74E-11 | DF 4 | cmbl |
| kctd12 | 7.3E-16 | 0.6438961 | 0.123 | 0.077 | 2.38E-11 | DF 4 | kctd1 |
| tgfbr1b | 7.4E-16 | 0.6027555 | 0.165 | 0.112 | 2.41E-11 | DF 4 | tgfbr1b |
| ip6k2a1 | 7.8E-16 | 0.4874457 | 0.259 | 0.194 | 2.53E-11 | DF 4 | ip6k2a |
| nos1apa2 | 7.9E-16 | 0.5321534 | 0.159 | 0.106 | 2.57E-11 | DF 4 | nos1apa |
| serf22 | 8.1E-16 | 0.2013738 | 0.828 | 0.77 | 2.63E-11 | DF 4 | serf2 |
| CDPF11 | 1.2E-15 | 0.4961125 | 0.238 | 0.176 | 3.84E-11 | DF 4 | CDPF1 |
| syf2 | 1.4E-15 | 0.4069421 | 0.35 | 0.279 | 4.7E-11 | DF 4 | syf2 |
| si:ch211-57n23.13 | 1.6E-15 | 0.4322935 | 0.284 | 0.214 | 5.14E-11 | DF 4 | si:ch211-57n23.1 |
| mbnl3 | 1.8E-15 | 0.5139546 | 0.188 | 0.131 | 6.01E-11 | DF 4 | mbnl3 |
| tgm1l13 | 1.9E-15 | 0.3697446 | 0.184 | 0.125 | 6.07E-11 | DF 4 | tgm1l1 |
| lonrf1 | 1.9E-15 | 0.6761825 | 0.101 | 0.06 | 6.22E-11 | DF 4 | lonrf1 |
| ptk7a2 | 2.2E-15 | 0.3365649 | 0.387 | 0.311 | 7.04E-11 | DF 4 | ptk7a |
| tcea21 | 2.3E-15 | 0.6916402 | 0.121 | 0.076 | 7.36E-11 | DF 4 | tcea2 |
| prickle1b3 | 2.4E-15 | 0.4477809 | 0.239 | 0.175 | 7.94E-11 | DF 4 | prickle1b |
| mcl1b1 | 2.6E-15 | 0.3950475 | 0.545 | 0.48 | 8.33E-11 | DF 4 | mcl1b |
| fgfr1a1 | 3E-15 | 0.5218481 | 0.213 | 0.154 | 9.84E-11 | DF 4 | fgfr1a |
| bco2l2 | 3.4E-15 | 0.4823801 | 0.178 | 0.122 | 1.11E-10 | DF 4 | bco2l |
| lfng2 | 3.6E-15 | 0.2458006 | 0.329 | 0.251 | 1.15E-10 | DF 4 | lfng |
| si:dkey-110g7.8 | 4.2E-15 | 0.5397871 | 0.156 | 0.104 | 1.36E-10 | DF 4 | si:dkey-110g7.8 |
| mylipb1 | 4.2E-15 | 0.5269064 | 0.124 | 0.078 | 1.37E-10 | DF 4 | mylipb |
| pde10a | 5.1E-15 | 0.7512919 | 0.101 | 0.061 | 1.66E-10 | DF 4 | pde10a |
| hs3st3b1b | 5.2E-15 | 0.5396412 | 0.181 | 0.127 | 1.69E-10 | DF 4 | hs3st3b1b |
| cd1641 | 7.5E-15 | 0.3062813 | 0.409 | 0.334 | 2.43E-10 | DF 4 | cd164 |
| fam43a1 | 8.3E-15 | 0.5399384 | 0.109 | 0.067 | 2.7E-10 | DF 4 | fam43a |
| slit21 | 9.4E-15 | 0.6843374 | 0.109 | 0.068 | 3.07E-10 | DF 4 | slit2 |
| arnt2 | 1.1E-14 | 0.7085136 | 0.134 | 0.088 | 3.57E-10 | DF 4 | arnt2 |
| cbx1b | 1.2E-14 | 0.5934434 | 0.155 | 0.105 | 4.03E-10 | DF 4 | cbx1b |
| gng21 | 1.2E-14 | 0.5306254 | 0.158 | 0.107 | 4.03E-10 | DF 4 | gng2 |
| lsm71 | 1.5E-14 | 0.3989208 | 0.324 | 0.257 | 4.74E-10 | DF 4 | lsm7 |
| mob2a1 | 2.1E-14 | 0.543716 | 0.19 | 0.135 | 6.75E-10 | DF 4 | mob2a |
| asap1a1 | 2.3E-14 | 0.6575552 | 0.137 | 0.09 | 7.39E-10 | DF 4 | asap1a |
| ier2a1 | 2.5E-14 | 0.3412638 | 0.544 | 0.483 | 7.98E-10 | DF 4 | ier2a |
| fgfr1b1 | 2.5E-14 | 0.5659416 | 0.148 | 0.099 | 8.26E-10 | DF 4 | fgfr1b |
| pik3r21 | 2.8E-14 | 0.5126623 | 0.26 | 0.198 | 9.06E-10 | DF 4 | pik3r2 |
| tspan4b3 | 3.1E-14 | 0.3437332 | 0.407 | 0.333 | 1.02E-09 | DF 4 | tspan4b |
| gadd45ab3 | 3.3E-14 | 0.4147471 | 0.233 | 0.173 | 1.07E-09 | DF 4 | gadd45ab |
| si:ch211-212k18.93 | 3.4E-14 | 0.4429516 | 0.236 | 0.175 | 1.1E-09 | DF 4 | si:ch211-212k18.9 |
| rbfox2 | 3.8E-14 | 0.3705539 | 0.337 | 0.268 | 1.23E-09 | DF 4 | rbfox2 |
| fam13b | 4.3E-14 | 0.594981 | 0.168 | 0.118 | 1.38E-09 | DF 4 | fam13b |
| tnfsf10l3 | 4.6E-14 | 0.2374683 | 0.291 | 0.218 | 1.48E-09 | DF 4 | tnfsf10l |
| si:ch211-194b1.11 | 4.7E-14 | 0.4041684 | 0.278 | 0.213 | 1.53E-09 | DF 4 | si:ch211-194b1.1 |
| mcl1a1 | 5.8E-14 | 0.3301425 | 0.798 | 0.77 | 1.89E-09 | DF 4 | mcl1a |
| FP236812.1 | 6.5E-14 | 0.820965 | 0.143 | 0.097 | 2.1E-09 | DF 4 | FP236812.1 |
| myca2 | 6.5E-14 | 0.5146818 | 0.212 | 0.155 | 2.1E-09 | DF 4 | myca |
| zbtb33 | 7E-14 | 0.6796467 | 0.11 | 0.069 | 2.27E-09 | DF 4 | zbtb33 |
| mxra8a3 | 7.2E-14 | 0.3049601 | 0.321 | 0.251 | 2.35E-09 | DF 4 | mxra8a |

| | | | | | | | |
|---|---|---|---|---|---|---|---|
| srgap31 | 8.3E-14 | 0.2794012 | 0.269 | 0.2 | 2.69E-09 | DF 4 | srgap3 |
| cybrd12 | 9E-14 | 0.6901612 | 0.124 | 0.081 | 2.93E-09 | DF 4 | cybrd1 |
| thbs2a3 | 9.8E-14 | 0.2202724 | 0.268 | 0.2 | 3.17E-09 | DF 4 | thbs2a |
| cspg5a1 | 1.1E-13 | 0.3880471 | 0.119 | 0.076 | 3.72E-09 | DF 4 | cspg5a |
| fam50a1 | 1.2E-13 | 0.5102114 | 0.215 | 0.16 | 3.79E-09 | DF 4 | fam50a |
| boc2 | 1.4E-13 | 0.2430953 | 0.449 | 0.369 | 4.64E-09 | DF 4 | boc |
| nkx3-23 | 1.6E-13 | 0.321355 | 0.256 | 0.193 | 5.18E-09 | DF 4 | nkx3-2 |
| per1b | 1.7E-13 | 0.4324734 | 0.261 | 0.2 | 5.69E-09 | DF 4 | per1b |
| eef2k | 1.8E-13 | 0.6545079 | 0.141 | 0.095 | 5.9E-09 | DF 4 | eef2k |
| per1a | 2.2E-13 | 0.6296655 | 0.107 | 0.067 | 7.02E-09 | DF 4 | per1a |
| top12 | 3.3E-13 | 0.49951 | 0.123 | 0.08 | 1.07E-08 | DF 4 | top1 |
| myh103 | 3.4E-13 | 0.2669679 | 0.406 | 0.332 | 1.1E-08 | DF 4 | myh10 |
| adkb | 4.5E-13 | 0.6879877 | 0.109 | 0.07 | 1.45E-08 | DF 4 | adkb |
| cpda | 5.5E-13 | 0.6289607 | 0.119 | 0.078 | 1.78E-08 | DF 4 | cpda |
| tgfbr31 | 6.4E-13 | 0.54296 | 0.12 | 0.078 | 2.09E-08 | DF 4 | tgfbr3 |
| dhrs13l11 | 6.8E-13 | 0.4001854 | 0.205 | 0.15 | 2.22E-08 | DF 4 | dhrs13l1 |
| prrc2a | 7.3E-13 | 0.5416111 | 0.186 | 0.135 | 2.37E-08 | DF 4 | prrc2a |
| caly1 | 8.6E-13 | 0.5327991 | 0.111 | 0.071 | 2.8E-08 | DF 4 | caly |
| lrrc154 | 1.1E-12 | 0.3867954 | 0.185 | 0.132 | 3.62E-08 | DF 4 | lrrc15 |
| tnrc6a1 | 1.2E-12 | 0.3362219 | 0.336 | 0.269 | 3.88E-08 | DF 4 | tnrc6a |
| gal3st21 | 1.7E-12 | 0.6318648 | 0.114 | 0.075 | 5.69E-08 | DF 4 | gal3st2 |
| gpx81 | 2.3E-12 | 0.4192294 | 0.227 | 0.172 | 7.45E-08 | DF 4 | gpx8 |
| zgc:175214 | 2.4E-12 | 0.5823114 | 0.151 | 0.106 | 7.8E-08 | DF 4 | zgc:175214 |
| tshz13 | 2.9E-12 | 0.4049152 | 0.185 | 0.134 | 9.36E-08 | DF 4 | tshz1 |
| ldhba2 | 2.9E-12 | 0.2215715 | 0.559 | 0.491 | 9.47E-08 | DF 4 | ldhba |
| gbe1a3 | 3.6E-12 | 0.2919616 | 0.159 | 0.11 | 1.16E-07 | DF 4 | gbe1a |
| si:dkey-222f8.3 | 3.7E-12 | 0.4121864 | 0.279 | 0.219 | 1.22E-07 | DF 4 | si:dkey-222f8.3 |
| mef2d | 5E-12 | 0.3606494 | 0.242 | 0.185 | 1.62E-07 | DF 4 | mef2d |
| ZFP36L1.11 | 5.9E-12 | 0.2024182 | 0.902 | 0.872 | 1.9E-07 | DF 4 | ZFP36L1.1 |
| si:ch73-281n10.21 | 6E-12 | 0.2206777 | 0.507 | 0.439 | 1.96E-07 | DF 4 | si:ch73-281n10.2 |
| ppt11 | 6E-12 | 0.2811614 | 0.409 | 0.342 | 1.96E-07 | DF 4 | ppt1 |
| nrp1a2 | 6.3E-12 | 0.3911896 | 0.179 | 0.129 | 2.05E-07 | DF 4 | nrp1a |
| zgc:1107962 | 7.3E-12 | 0.7611192 | 0.118 | 0.079 | 2.36E-07 | DF 4 | zgc:110796 |
| coq10b1 | 7.4E-12 | 0.447428 | 0.219 | 0.166 | 2.4E-07 | DF 4 | coq10b |
| cacna1g2 | 7.6E-12 | 0.4881926 | 0.138 | 0.095 | 2.46E-07 | DF 4 | cacna1g |
| lrp1ab2 | 8E-12 | 0.2807154 | 0.406 | 0.338 | 2.61E-07 | DF 4 | lrp1ab |
| tmem200b2 | 8.4E-12 | 0.4540916 | 0.136 | 0.093 | 2.74E-07 | DF 4 | tmem200b |
| wbp2nl | 9E-12 | 0.3476754 | 0.35 | 0.288 | 2.92E-07 | DF 4 | wbp2nl |
| stk35 | 9.1E-12 | 0.5756826 | 0.122 | 0.082 | 2.96E-07 | DF 4 | stk35 |
| acox1 | 9.4E-12 | 0.5722573 | 0.144 | 0.101 | 3.06E-07 | DF 4 | acox1 |
| CABZ01032488.1 | 1.4E-11 | 0.4609133 | 0.147 | 0.103 | 4.42E-07 | DF 4 | CABZ01032488.1 |
| pcdh112 | 1.4E-11 | 0.5375919 | 0.112 | 0.074 | 4.58E-07 | DF 4 | pcdh11 |
| aldh6a12 | 1.5E-11 | 0.3274597 | 0.332 | 0.271 | 5E-07 | DF 4 | aldh6a1 |
| fmr1 | 1.7E-11 | 0.4566099 | 0.201 | 0.152 | 5.41E-07 | DF 4 | fmr1 |
| rev3l | 2E-11 | 0.5913897 | 0.135 | 0.094 | 6.44E-07 | DF 4 | rev3l |
| pebp11 | 2.2E-11 | 0.7522884 | 0.107 | 0.071 | 7.3E-07 | DF 4 | pebp1 |
| bhlhe411 | 2.3E-11 | 0.4929871 | 0.174 | 0.127 | 7.54E-07 | DF 4 | bhlhe41 |
| nectin3b1 | 2.5E-11 | 0.4529542 | 0.204 | 0.153 | 8.06E-07 | DF 4 | nectin3b |

| | | | | | | | |
|---|---|---|---|---|---|---|---|
| si:dkeyp-69b9.6 | 3.2E-11 | 0.5866692 | 0.146 | 0.103 | 1.04E-06 | DF 4 | si:dkeyp-69b9.6 |
| dhrs13a.3 | 3.2E-11 | 0.4736931 | 0.169 | 0.123 | 1.05E-06 | DF 4 | dhrs13a.3 |
| smarca21 | 3.4E-11 | 0.369654 | 0.281 | 0.224 | 1.09E-06 | DF 4 | smarca2 |
| si:dkey-199f5.82 | 3.4E-11 | 0.402273 | 0.214 | 0.162 | 1.1E-06 | DF 4 | si:dkey-199f5.8 |
| fh1 | 3.6E-11 | 0.5987969 | 0.104 | 0.069 | 1.17E-06 | DF 4 | fh |
| klf132 | 3.9E-11 | 0.5687312 | 0.171 | 0.126 | 1.28E-06 | DF 4 | klf13 |
| smoc2 | 6.4E-11 | 0.4739873 | 0.129 | 0.089 | 2.07E-06 | DF 4 | smoc2 |
| si:dkey-27i16.21 | 7E-11 | 0.3733181 | 0.161 | 0.116 | 2.27E-06 | DF 4 | si:dkey-27i16.2 |
| ctdsp21 | 7.3E-11 | 0.3715998 | 0.241 | 0.188 | 2.37E-06 | DF 4 | ctdsp2 |
| axin11 | 7.3E-11 | 0.4825514 | 0.155 | 0.111 | 2.39E-06 | DF 4 | axin1 |
| tmem59 | 7.5E-11 | 0.3182199 | 0.371 | 0.311 | 2.45E-06 | DF 4 | tmem59 |
| pros1 | 8.1E-11 | 0.5093685 | 0.158 | 0.115 | 2.64E-06 | DF 4 | pros1 |
| rerea | 8.6E-11 | 0.4223364 | 0.223 | 0.173 | 2.79E-06 | DF 4 | rerea |
| calr3a2 | 1E-10 | 0.2395526 | 0.429 | 0.359 | 3.28E-06 | DF 4 | calr3a |
| wdr26b | 1E-10 | 0.4937183 | 0.13 | 0.091 | 3.31E-06 | DF 4 | wdr26b |
| glis2a1 | 1.1E-10 | 0.3310898 | 0.257 | 0.202 | 3.44E-06 | DF 4 | glis2a |
| mink1 | 1.1E-10 | 0.463422 | 0.142 | 0.1 | 3.62E-06 | DF 4 | mink1 |
| bmpr2b | 1.1E-10 | 0.5521486 | 0.12 | 0.083 | 3.7E-06 | DF 4 | bmpr2b |
| ppp1r3ca2 | 1.2E-10 | 0.2335022 | 0.265 | 0.206 | 3.84E-06 | DF 4 | ppp1r3ca |
| mmp15a2 | 1.4E-10 | 0.4336949 | 0.12 | 0.082 | 4.62E-06 | DF 4 | mmp15a |
| ctdspl2b2 | 1.5E-10 | 0.4366012 | 0.155 | 0.111 | 4.96E-06 | DF 4 | ctdspl2b |
| nr2f21 | 2.1E-10 | 0.4020935 | 0.157 | 0.114 | 6.81E-06 | DF 4 | nr2f2 |
| arhgap5 | 2.1E-10 | 0.279121 | 0.41 | 0.348 | 6.83E-06 | DF 4 | arhgap5 |
| gpx1a3 | 2.5E-10 | 0.2519019 | 0.244 | 0.191 | 8.01E-06 | DF 4 | gpx1a |
| pitpnc1b1 | 2.6E-10 | 0.5201748 | 0.117 | 0.08 | 8.57E-06 | DF 4 | pitpnc1b |
| rpe1 | 2.8E-10 | 0.5211719 | 0.13 | 0.092 | 9.1E-06 | DF 4 | rpe |
| tspan312 | 3.1E-10 | 0.2633363 | 0.281 | 0.224 | 1.02E-05 | DF 4 | tspan31 |
| zgc:1105911 | 3.3E-10 | 0.498243 | 0.13 | 0.092 | 1.09E-05 | DF 4 | zgc:110591 |
| abcc6a1 | 4E-10 | 0.3455097 | 0.11 | 0.075 | 1.3E-05 | DF 4 | abcc6a |
| tspan18b3 | 4E-10 | 0.3667768 | 0.187 | 0.14 | 1.3E-05 | DF 4 | tspan18b |
| notch21 | 4.2E-10 | 0.2201356 | 0.44 | 0.376 | 1.38E-05 | DF 4 | notch2 |
| me3 | 4.8E-10 | 0.5432785 | 0.119 | 0.083 | 1.55E-05 | DF 4 | me3 |
| map2k62 | 5E-10 | 0.3154799 | 0.24 | 0.188 | 1.61E-05 | DF 4 | map2k6 |
| ppp3r1a | 5.1E-10 | 0.4550306 | 0.184 | 0.14 | 1.65E-05 | DF 4 | ppp3r1a |
| gng7 | 5.2E-10 | 0.457031 | 0.21 | 0.163 | 1.68E-05 | DF 4 | gng7 |
| C19H6orf62 | 5.8E-10 | 0.3277247 | 0.454 | 0.4 | 1.9E-05 | DF 4 | C19H6orf62 |
| dnajc5aa | 5.9E-10 | 0.5826914 | 0.119 | 0.083 | 1.92E-05 | DF 4 | dnajc5aa |
| phactr22 | 6.4E-10 | 0.4036047 | 0.185 | 0.139 | 2.09E-05 | DF 4 | phactr2 |
| puraa | 7E-10 | 0.3086298 | 0.305 | 0.251 | 2.27E-05 | DF 4 | puraa |
| tspan4a1 | 7.2E-10 | 0.4080495 | 0.18 | 0.136 | 2.33E-05 | DF 4 | tspan4a |
| C7H4orf48 | 9E-10 | 0.3070573 | 0.297 | 0.241 | 2.93E-05 | DF 4 | C7H4orf48 |
| kank1a3 | 1E-09 | 0.5274234 | 0.151 | 0.111 | 3.28E-05 | DF 4 | kank1a |
| vamp51 | 1.1E-09 | 0.5800498 | 0.114 | 0.079 | 3.63E-05 | DF 4 | vamp5 |
| zgc:1626122 | 1.1E-09 | 0.4146844 | 0.156 | 0.115 | 3.69E-05 | DF 4 | zgc:162612 |
| serinc1 | 1.3E-09 | 0.3261256 | 0.312 | 0.258 | 4.2E-05 | DF 4 | serinc1 |
| prr12b1 | 1.3E-09 | 0.555556 | 0.155 | 0.115 | 4.37E-05 | DF 4 | prr12b |
| eml32 | 1.5E-09 | 0.4910408 | 0.16 | 0.119 | 4.95E-05 | DF 4 | eml3 |
| ddr13 | 1.5E-09 | 0.4698765 | 0.122 | 0.085 | 5.04E-05 | DF 4 | ddr1 |

| | | | | | | | |
|---|---|---|---|---|---|---|---|
| mob1ba3 | 1.7E-09 | 0.3876686 | 0.138 | 0.099 | 5.62E-05 | DF 4 | mob1ba |
| smo1 | 1.7E-09 | 0.3415657 | 0.308 | 0.255 | 5.68E-05 | DF 4 | smo |
| purab | 1.8E-09 | 0.2962752 | 0.228 | 0.178 | 5.96E-05 | DF 4 | purab |
| ctsf | 1.9E-09 | 0.4856984 | 0.104 | 0.071 | 6.27E-05 | DF 4 | ctsf |
| aldh7a1 | 2E-09 | 0.4026835 | 0.232 | 0.186 | 6.39E-05 | DF 4 | aldh7a1 |
| fzd21 | 2E-09 | 0.3695838 | 0.136 | 0.097 | 6.42E-05 | DF 4 | fzd2 |
| tspan7 | 2E-09 | 0.3487844 | 0.263 | 0.213 | 6.62E-05 | DF 4 | tspan7 |
| kdm5ba | 2.7E-09 | 0.4260179 | 0.159 | 0.118 | 8.82E-05 | DF 4 | kdm5ba |
| kdm4b | 2.9E-09 | 0.3585715 | 0.23 | 0.183 | 9.46E-05 | DF 4 | kdm4b |
| p4ha1a2 | 3.1E-09 | 0.5476415 | 0.131 | 0.095 | 0.000102 | DF 4 | p4ha1a |
| picalma | 3.2E-09 | 0.5961514 | 0.106 | 0.074 | 0.000105 | DF 4 | picalma |
| arglu1a2 | 3.3E-09 | 0.3713994 | 0.218 | 0.171 | 0.000106 | DF 4 | arglu1a |
| si:cabz01007802.12 | 3.3E-09 | 0.4502029 | 0.125 | 0.089 | 0.000108 | DF 4 | si:cabz01007802.1 |
| zgc:921401 | 3.3E-09 | 0.3918462 | 0.153 | 0.112 | 0.000108 | DF 4 | zgc:92140 |
| brd3b | 3.3E-09 | 0.3442174 | 0.275 | 0.225 | 0.000109 | DF 4 | brd3b |
| mycbp1 | 3.4E-09 | 0.3185193 | 0.297 | 0.246 | 0.000109 | DF 4 | mycbp |
| zmp:00000007112 | 3.8E-09 | 0.5056696 | 0.115 | 0.081 | 0.000123 | DF 4 | zmp:0000000711 |
| capns1a | 4E-09 | 0.229931 | 0.451 | 0.398 | 0.00013 | DF 4 | capns1a |
| si:ch1073-184j22.11 | 4.3E-09 | 0.4142517 | 0.113 | 0.079 | 0.000139 | DF 4 | si:ch1073-184j22.1 |
| zc4h2 | 4.3E-09 | 0.5697527 | 0.111 | 0.077 | 0.000139 | DF 4 | zc4h2 |
| sept9a | 4.8E-09 | 0.2609481 | 0.266 | 0.214 | 0.000156 | DF 4 | sept9a |
| ngfb2 | 4.8E-09 | 0.3561665 | 0.115 | 0.081 | 0.000157 | DF 4 | ngfb |
| si:dkey-235d18.51 | 5.3E-09 | 0.4222063 | 0.134 | 0.097 | 0.000173 | DF 4 | si:dkey-235d18.5 |
| hspa8b2 | 5.7E-09 | 0.2643238 | 0.129 | 0.092 | 0.000186 | DF 4 | hspa8b |
| bag3 | 6E-09 | 0.6502265 | 0.102 | 0.07 | 0.000194 | DF 4 | bag3 |
| nfs1 | 6.4E-09 | 0.4452047 | 0.121 | 0.086 | 0.000209 | DF 4 | nfs1 |
| mbd3a | 6.5E-09 | 0.5630996 | 0.133 | 0.098 | 0.00021 | DF 4 | mbd3a |
| vegfc2 | 6.5E-09 | 0.3845432 | 0.123 | 0.087 | 0.000211 | DF 4 | vegfc |
| znf532 | 7.9E-09 | 0.4206302 | 0.101 | 0.069 | 0.000258 | DF 4 | znf532 |
| lnpep | 8.9E-09 | 0.6202293 | 0.102 | 0.071 | 0.000288 | DF 4 | lnpep |
| sntb22 | 1.1E-08 | 0.2997538 | 0.286 | 0.235 | 0.000347 | DF 4 | sntb2 |
| si:dkey-51e6.13 | 1.1E-08 | 0.2773727 | 0.308 | 0.256 | 0.000357 | DF 4 | si:dkey-51e6.1 |
| fndc3a | 1.1E-08 | 0.5879355 | 0.114 | 0.082 | 0.000364 | DF 4 | fndc3a |
| c62 | 1.1E-08 | 0.2930911 | 0.104 | 0.072 | 0.000364 | DF 4 | c6 |
| tnrc6c2 | 1.1E-08 | 0.3893793 | 0.121 | 0.087 | 0.000371 | DF 4 | tnrc6c2 |
| fam120c | 1.2E-08 | 0.2605121 | 0.351 | 0.299 | 0.000385 | DF 4 | fam120c |
| cblb | 1.2E-08 | 0.4993167 | 0.13 | 0.095 | 0.00039 | DF 4 | cblb |
| gabarapl2 | 1.4E-08 | 0.3886982 | 0.205 | 0.163 | 0.000463 | DF 4 | gabarapl2 |
| dag1 | 1.4E-08 | 0.334526 | 0.2 | 0.156 | 0.000468 | DF 4 | dag1 |
| oip5-as1 | 1.5E-08 | 0.3510492 | 0.195 | 0.152 | 0.000496 | DF 4 | oip5-as1 |
| mex3b1 | 1.5E-08 | 0.3600064 | 0.215 | 0.17 | 0.000498 | DF 4 | mex3b |
| xirp2a2 | 1.9E-08 | 0.293887 | 0.307 | 0.256 | 0.000627 | DF 4 | xirp2a |
| ttyh3a2 | 2E-08 | 0.4194324 | 0.13 | 0.095 | 0.000635 | DF 4 | ttyh3a |
| ndfip1l1 | 2.1E-08 | 0.2113026 | 0.39 | 0.335 | 0.000683 | DF 4 | ndfip1l |
| pds5b1 | 2.1E-08 | 0.4726888 | 0.108 | 0.076 | 0.000683 | DF 4 | pds5b |
| si:dkeyp-113d7.11 | 2.2E-08 | 0.4934515 | 0.181 | 0.143 | 0.000703 | DF 4 | si:dkeyp-113d7.1 |
| pls31 | 2.3E-08 | 0.2601961 | 0.369 | 0.319 | 0.000747 | DF 4 | pls3 |
| cs1 | 2.4E-08 | 0.4548538 | 0.19 | 0.15 | 0.000794 | DF 4 | cs |

| | | | | | | | |
|---|---|---|---|---|---|---|---|
| col14a1a2 | 2.5E-08 | 0.541319 | 0.104 | 0.073 | 0.000816 | DF 4 | col14a1a |
| gas1b4 | 2.7E-08 | 0.2624349 | 0.242 | 0.194 | 0.000875 | DF 4 | gas1b |
| gsto22 | 2.7E-08 | 0.4062076 | 0.138 | 0.102 | 0.000891 | DF 4 | gsto2 |
| ccdc6b1 | 2.8E-08 | 0.4748992 | 0.111 | 0.079 | 0.000914 | DF 4 | ccdc6b |
| pcdh18b1 | 3E-08 | 0.2845988 | 0.266 | 0.218 | 0.000972 | DF 4 | pcdh18b |
| capza1a1 | 3.5E-08 | 0.289161 | 0.318 | 0.27 | 0.001154 | DF 4 | capza1a |
| ENSDARG00000597( | 3.6E-08 | 0.4673373 | 0.116 | 0.084 | 0.001159 | DF 4 | ENSDARG00000059 |
| ITLN15 | 3.8E-08 | 0.2014079 | 0.26 | 0.209 | 0.00124 | DF 4 | ITLN1 |
| lrrfip1a1 | 3.9E-08 | 0.6362676 | 0.103 | 0.073 | 0.001257 | DF 4 | lrrfip1a |
| cep2901 | 4.2E-08 | 0.4192683 | 0.146 | 0.11 | 0.001359 | DF 4 | cep290 |
| fkbp5 | 4.2E-08 | 0.258045 | 0.242 | 0.195 | 0.001371 | DF 4 | fkbp5 |
| h1-103 | 4.7E-08 | 0.3754522 | 0.247 | 0.203 | 0.001538 | DF 4 | h1-10 |
| sfxn5b4 | 4.8E-08 | 0.2886083 | 0.177 | 0.136 | 0.001575 | DF 4 | sfxn5b |
| itga11a2 | 5.1E-08 | 0.2223746 | 0.146 | 0.109 | 0.001643 | DF 4 | itga11a |
| zgc:101566 | 6.1E-08 | 0.4966237 | 0.119 | 0.087 | 0.001978 | DF 4 | zgc:101566 |
| nedd4a2 | 6.9E-08 | 0.4285199 | 0.123 | 0.09 | 0.002243 | DF 4 | nedd4a |
| cdkn1bb1 | 7.2E-08 | 0.2345549 | 0.347 | 0.296 | 0.002343 | DF 4 | cdkn1bb |
| galnt1 | 7.6E-08 | 0.3273946 | 0.204 | 0.162 | 0.002477 | DF 4 | galnt1 |
| mif1 | 8.2E-08 | 0.2067843 | 0.449 | 0.398 | 0.002666 | DF 4 | mif |
| crim11 | 8.3E-08 | 0.3548664 | 0.181 | 0.142 | 0.002698 | DF 4 | crim1 |
| pepd1 | 9.7E-08 | 0.3645379 | 0.254 | 0.212 | 0.003151 | DF 4 | pepd |
| dab2ipa1 | 9.9E-08 | 0.3880494 | 0.147 | 0.111 | 0.003226 | DF 4 | dab2ipa |
| rnf1691 | 1E-07 | 0.364721 | 0.165 | 0.128 | 0.003319 | DF 4 | rnf169 |
| metrn1 | 1.2E-07 | 0.2026165 | 0.154 | 0.115 | 0.00378 | DF 4 | metrn |
| si:dkey-276j7.11 | 1.2E-07 | 0.395414 | 0.206 | 0.166 | 0.003911 | DF 4 | si:dkey-276j7.1 |
| eif4a21 | 1.3E-07 | 0.2892118 | 0.235 | 0.191 | 0.004096 | DF 4 | eif4a2 |
| lin7c | 1.3E-07 | 0.4662917 | 0.131 | 0.099 | 0.004162 | DF 4 | lin7c |
| psap | 1.4E-07 | 0.2534002 | 0.396 | 0.348 | 0.004445 | DF 4 | psap |
| qser1 | 1.5E-07 | 0.6004755 | 0.1 | 0.072 | 0.004843 | DF 4 | qser1 |
| dnmt3ab | 1.7E-07 | 0.3723969 | 0.155 | 0.119 | 0.005371 | DF 4 | dnmt3ab |
| hsf2 | 1.9E-07 | 0.3087456 | 0.225 | 0.183 | 0.00612 | DF 4 | hsf2 |
| ptpn11a | 1.9E-07 | 0.3779177 | 0.147 | 0.112 | 0.006264 | DF 4 | ptpn11a |
| spred2a | 2E-07 | 0.4445498 | 0.104 | 0.075 | 0.006632 | DF 4 | spred2a |
| bloc1s62 | 2.1E-07 | 0.3044536 | 0.22 | 0.178 | 0.006759 | DF 4 | bloc1s6 |
| lonrf1l2 | 2.3E-07 | 0.3654652 | 0.188 | 0.151 | 0.007435 | DF 4 | lonrf1l |
| rbm39a | 2.4E-07 | 0.2978936 | 0.273 | 0.228 | 0.007717 | DF 4 | rbm39a |
| grb10a1 | 2.4E-07 | 0.375598 | 0.142 | 0.108 | 0.007845 | DF 4 | grb10a |
| fam168a | 2.4E-07 | 0.324258 | 0.26 | 0.219 | 0.00793 | DF 4 | fam168a |
| runx1t11 | 2.7E-07 | 0.3146152 | 0.122 | 0.09 | 0.008912 | DF 4 | runx1t1 |
| phf2 | 2.8E-07 | 0.3399622 | 0.226 | 0.185 | 0.008978 | DF 4 | phf2 |
| smad2 | 2.8E-07 | 0.3275071 | 0.231 | 0.19 | 0.009101 | DF 4 | smad2 |
| dtnbp1a | 2.9E-07 | 0.469435 | 0.144 | 0.111 | 0.009364 | DF 4 | dtnbp1a |
| ube2z | 3E-07 | 0.422299 | 0.107 | 0.078 | 0.009653 | DF 4 | ube2z |
| tmem50a | 3E-07 | 0.2454396 | 0.28 | 0.233 | 0.009776 | DF 4 | tmem50a |
| ENSDARG000000758! | 3.4E-07 | 0.4139755 | 0.114 | 0.084 | 0.010961 | DF 4 | ENSDARG00000075 |
| tbl1xr1a | 3.4E-07 | 0.2940612 | 0.189 | 0.15 | 0.011095 | DF 4 | tbl1xr1a |
| os9 | 3.4E-07 | 0.2875836 | 0.256 | 0.214 | 0.011183 | DF 4 | os9 |
| PLEKHH3 | 3.4E-07 | 0.354712 | 0.201 | 0.162 | 0.011191 | DF 4 | PLEKHH3 |

| | | | | | | | |
|---|---|---|---|---|---|---|---|
| zgc:1949812 | 3.5E-07 | 0.5467329 | 0.109 | 0.08 | 0.011523 | DF 4 | zgc:194981 |
| megf81 | 3.7E-07 | 0.3370969 | 0.163 | 0.127 | 0.011947 | DF 4 | megf8 |
| phip1 | 3.7E-07 | 0.3090735 | 0.195 | 0.156 | 0.01213 | DF 4 | phip |
| prelid3b1 | 3.9E-07 | 0.2670755 | 0.402 | 0.356 | 0.012599 | DF 4 | prelid3b |
| smarcc22 | 4.5E-07 | 0.3230456 | 0.212 | 0.171 | 0.014573 | DF 4 | smarcc2 |
| ndfip1 | 4.6E-07 | 0.2265642 | 0.421 | 0.373 | 0.014922 | DF 4 | ndfip1 |
| phyh1 | 4.6E-07 | 0.3540811 | 0.122 | 0.091 | 0.014952 | DF 4 | phyh |
| sp4 | 4.6E-07 | 0.3424333 | 0.185 | 0.148 | 0.014963 | DF 4 | sp4 |
| ska2 | 5E-07 | 0.3888929 | 0.156 | 0.123 | 0.016139 | DF 4 | ska2 |
| ehbp1l1a2 | 5.1E-07 | 0.2987419 | 0.125 | 0.093 | 0.016724 | DF 4 | ehbp1l1a |
| crebbpa | 5.3E-07 | 0.3082003 | 0.198 | 0.16 | 0.017083 | DF 4 | crebbpa |
| skib2 | 5.3E-07 | 0.2741995 | 0.242 | 0.201 | 0.017158 | DF 4 | skib |
| si:rp71-79p20.2 | 5.6E-07 | 0.2650224 | 0.19 | 0.151 | 0.018079 | DF 4 | si:rp71-79p20.2 |
| cnpy4 | 6.3E-07 | 0.4573053 | 0.156 | 0.123 | 0.020347 | DF 4 | cnpy4 |
| mpc21 | 6.7E-07 | 0.3171623 | 0.218 | 0.18 | 0.021746 | DF 4 | mpc2 |
| sart31 | 6.9E-07 | 0.4855959 | 0.117 | 0.087 | 0.022376 | DF 4 | sart3 |
| fkbp10b2 | 6.9E-07 | 0.234901 | 0.298 | 0.253 | 0.022513 | DF 4 | fkbp10b |
| selenot1b1 | 7.4E-07 | 0.384443 | 0.146 | 0.113 | 0.024136 | DF 4 | selenot1b |
| wasf2 | 8E-07 | 0.2410128 | 0.304 | 0.259 | 0.026091 | DF 4 | wasf2 |
| camk2g1 | 8.3E-07 | 0.2667406 | 0.256 | 0.213 | 0.027013 | DF 4 | camk2g1 |
| si:ch211-212k18.82 | 8.6E-07 | 0.2589137 | 0.178 | 0.14 | 0.027895 | DF 4 | si:ch211-212k18.8 |
| arid5b2 | 9E-07 | 0.3588131 | 0.17 | 0.134 | 0.029322 | DF 4 | arid5b |
| acat1 | 1.1E-06 | 0.377541 | 0.125 | 0.094 | 0.034494 | DF 4 | acat1 |
| zswim51 | 1.1E-06 | 0.2606965 | 0.218 | 0.179 | 0.035559 | DF 4 | zswim5 |
| htatip2 | 1.1E-06 | 0.4433195 | 0.111 | 0.083 | 0.035598 | DF 4 | htatip2 |
| tmem106a | 1.1E-06 | 0.5102871 | 0.103 | 0.076 | 0.036066 | DF 4 | tmem106a |
| plekhb2 | 1.1E-06 | 0.4045026 | 0.132 | 0.101 | 0.036391 | DF 4 | plekhb2 |
| zgc:1103392 | 1.1E-06 | 0.2217184 | 0.257 | 0.214 | 0.037214 | DF 4 | zgc:110339 |
| si:dkey-9i23.15 | 1.2E-06 | 0.3796337 | 0.186 | 0.151 | 0.040064 | DF 4 | si:dkey-9i23.15 |
| dip2ca2 | 1.3E-06 | 0.4361412 | 0.114 | 0.086 | 0.042328 | DF 4 | dip2ca |
| max1 | 1.3E-06 | 0.2367094 | 0.332 | 0.288 | 0.042751 | DF 4 | max |
| CABZ01064941.12 | 1.3E-06 | 0.408608 | 0.113 | 0.084 | 0.043072 | DF 4 | CABZ01064941.1 |
| si:dkey-17m8.11 | 1.4E-06 | 0.3945204 | 0.154 | 0.122 | 0.044744 | DF 4 | si:dkey-17m8.1 |
| chp2 | 1.4E-06 | 0.3836645 | 0.106 | 0.078 | 0.046727 | DF 4 | chp2 |
| farp22 | 1.5E-06 | 0.4186545 | 0.146 | 0.114 | 0.047505 | DF 4 | farp2 |
| CU929418.1 | 1.5E-06 | 0.322395 | 0.157 | 0.124 | 0.048494 | DF 4 | CU929418.1 |
| cry21 | 2E-06 | 0.333406 | 0.202 | 0.165 | 0.065467 | DF 4 | cry2 |
| BX511129.21 | 2.1E-06 | 0.4565374 | 0.116 | 0.088 | 0.067612 | DF 4 | BX511129.2 |
| si:ch211-126j24.11 | 2.4E-06 | 0.4592022 | 0.108 | 0.081 | 0.077305 | DF 4 | si:ch211-126j24.1 |
| cipcb | 2.5E-06 | 0.3948805 | 0.105 | 0.078 | 0.080498 | DF 4 | cipcb |
| AL929435.11 | 2.5E-06 | 0.4367477 | 0.136 | 0.106 | 0.082101 | DF 4 | AL929435.1 |
| irf2bp1 | 2.6E-06 | 0.3816128 | 0.126 | 0.097 | 0.084174 | DF 4 | irf2bp1 |
| igf1rb | 2.8E-06 | 0.2495245 | 0.31 | 0.269 | 0.089851 | DF 4 | igf1rb |
| si:dkey-17o15.2 | 3.2E-06 | 0.4066562 | 0.109 | 0.081 | 0.103682 | DF 4 | si:dkey-17o15.2 |
| foxo3b1 | 3.4E-06 | 0.4676299 | 0.108 | 0.081 | 0.110007 | DF 4 | foxo3b |
| pik3c2a | 3.5E-06 | 0.4161506 | 0.118 | 0.09 | 0.112649 | DF 4 | pik3c2a |
| ppp1r13l2 | 4.1E-06 | 0.3709721 | 0.114 | 0.087 | 0.133441 | DF 4 | ppp1r13l |
| etnk11 | 4.2E-06 | 0.2093802 | 0.226 | 0.187 | 0.137392 | DF 4 | etnk1 |

| | | | | | | | |
|---|---|---|---|---|---|---|---|
| kyat3 | 4.6E-06 | 0.2511801 | 0.154 | 0.121 | 0.148552 | DF 4 | kyat3 |
| ptena | 4.8E-06 | 0.2456997 | 0.295 | 0.255 | 0.157331 | DF 4 | ptena |
| tpd52l2b | 5.2E-06 | 0.3157679 | 0.255 | 0.219 | 0.169586 | DF 4 | tpd52l2b |
| limch1b1 | 5.6E-06 | 0.3852238 | 0.117 | 0.09 | 0.181031 | DF 4 | limch1b |
| aff23 | 5.8E-06 | 0.2121973 | 0.217 | 0.178 | 0.188825 | DF 4 | aff2 |
| hbp1 | 6E-06 | 0.4080434 | 0.141 | 0.111 | 0.19429 | DF 4 | hbp1 |
| map1lc3b | 6E-06 | 0.2501624 | 0.301 | 0.262 | 0.194548 | DF 4 | map1lc3b |
| slc43a3b2 | 6E-06 | 0.3726767 | 0.104 | 0.078 | 0.196715 | DF 4 | slc43a3b |
| tmem106ba | 6.1E-06 | 0.2467583 | 0.226 | 0.189 | 0.198027 | DF 4 | tmem106ba |
| fasn | 6.4E-06 | 0.3632296 | 0.13 | 0.102 | 0.207938 | DF 4 | fasn |
| znf395b | 6.5E-06 | 0.3101532 | 0.17 | 0.137 | 0.212851 | DF 4 | znf395b |
| CR381618.13 | 6.7E-06 | 0.3355844 | 0.264 | 0.226 | 0.217593 | DF 4 | CR381618.1 |
| kmt2ca | 7.1E-06 | 0.2961714 | 0.204 | 0.169 | 0.23232 | DF 4 | kmt2ca |
| ip6k2b1 | 7.2E-06 | 0.208974 | 0.385 | 0.342 | 0.235677 | DF 4 | ip6k2b |
| aldh2.1 | 7.3E-06 | 0.3759896 | 0.14 | 0.11 | 0.236334 | DF 4 | aldh2.1 |
| fam172a | 7.6E-06 | 0.4009021 | 0.105 | 0.08 | 0.248242 | DF 4 | fam172a |
| pcmt1 | 7.7E-06 | 0.4284031 | 0.119 | 0.092 | 0.248824 | DF 4 | pcmt |
| abcc52 | 8.4E-06 | 0.2282847 | 0.177 | 0.143 | 0.273728 | DF 4 | abcc5 |
| ccnd13 | 8.5E-06 | 0.4074181 | 0.101 | 0.076 | 0.276174 | DF 4 | ccnd1 |
| rtf1 | 8.6E-06 | 0.233134 | 0.288 | 0.25 | 0.280625 | DF 4 | rtf1 |
| ranbp9 | 8.8E-06 | 0.3326054 | 0.153 | 0.122 | 0.28643 | DF 4 | ranbp9 |
| gigyf1b | 9.9E-06 | 0.2846893 | 0.172 | 0.139 | 0.320732 | DF 4 | gigyf1b |
| adprh1 | 9.9E-06 | 0.3939148 | 0.105 | 0.08 | 0.321627 | DF 4 | adprh |
| zfx | 1.1E-05 | 0.4659208 | 0.1 | 0.076 | 0.342863 | DF 4 | zfx |
| fkbp41 | 1.1E-05 | 0.2925667 | 0.189 | 0.157 | 0.354566 | DF 4 | fkbp4 |
| zgc:123105 | 1.1E-05 | 0.4933624 | 0.1 | 0.075 | 0.356259 | DF 4 | zgc:123105 |
| fntb | 1.1E-05 | 0.4612215 | 0.133 | 0.105 | 0.362759 | DF 4 | fntb |
| cyp3c11 | 1.2E-05 | 0.3363728 | 0.122 | 0.094 | 0.380221 | DF 4 | cyp3c1 |
| PPP2CA | 1.2E-05 | 0.358516 | 0.165 | 0.134 | 0.380764 | DF 4 | PPP2CA |
| pdcl | 1.2E-05 | 0.2542652 | 0.235 | 0.199 | 0.388499 | DF 4 | pdcl |
| zgc:112962 | 1.2E-05 | 0.2752469 | 0.155 | 0.125 | 0.390899 | DF 4 | zgc:112962 |
| CU464124.11 | 1.2E-05 | 0.3903631 | 0.131 | 0.103 | 0.399673 | DF 4 | CU464124.1 |
| aamp3 | 1.3E-05 | 0.3488004 | 0.184 | 0.153 | 0.413889 | DF 4 | aamp |
| tssc41 | 1.3E-05 | 0.4974419 | 0.117 | 0.091 | 0.42118 | DF 4 | tssc4 |
| hipk3b | 1.4E-05 | 0.3902592 | 0.101 | 0.077 | 0.449008 | DF 4 | hipk3b |
| kmt5c | 1.4E-05 | 0.3642591 | 0.134 | 0.106 | 0.455208 | DF 4 | kmt5c |
| rab7a | 1.5E-05 | 0.3386195 | 0.171 | 0.14 | 0.474331 | DF 4 | rab7a |
| churc1 | 1.6E-05 | 0.2341282 | 0.16 | 0.129 | 0.506978 | DF 4 | churc1 |
| rasl11b2 | 1.6E-05 | 0.2553876 | 0.252 | 0.215 | 0.509682 | DF 4 | rasl11b |
| si:ch211-140m22.7 | 1.6E-05 | 0.3759098 | 0.17 | 0.14 | 0.514897 | DF 4 | si:ch211-140m22.7 |
| auts2a1 | 1.7E-05 | 0.2207911 | 0.26 | 0.221 | 0.539295 | DF 4 | auts2a |
| cwc273 | 1.7E-05 | 0.2979093 | 0.151 | 0.122 | 0.562196 | DF 4 | cwc27 |
| mpp6a1 | 1.7E-05 | 0.3918529 | 0.106 | 0.081 | 0.562909 | DF 4 | mpp6a |
| cdc42ep4b2 | 1.9E-05 | 0.4301502 | 0.109 | 0.085 | 0.630271 | DF 4 | cdc42ep4b |
| waca | 2E-05 | 0.2654937 | 0.149 | 0.119 | 0.647608 | DF 4 | waca |
| napab | 2.1E-05 | 0.2547063 | 0.237 | 0.204 | 0.670551 | DF 4 | napab |
| mnta1 | 2.1E-05 | 0.2460111 | 0.129 | 0.101 | 0.682072 | DF 4 | mnta |
| meis2a.11 | 2.3E-05 | 0.215044 | 0.164 | 0.132 | 0.733802 | DF 4 | meis2a.1 |

| | | | | | | | |
|---|---|---|---|---|---|---|---|
| si:dkeyp-69e1.81 | 2.3E-05 | 0.2933786 | 0.135 | 0.107 | 0.735324 | DF 4 | si:dkeyp-69e1.8 |
| pnisr1 | 2.3E-05 | 0.294963 | 0.219 | 0.185 | 0.753578 | DF 4 | pnisr |
| stag2b | 2.4E-05 | 0.2923589 | 0.199 | 0.166 | 0.767741 | DF 4 | stag2b |
| crlf31 | 2.4E-05 | 0.4176162 | 0.112 | 0.088 | 0.774091 | DF 4 | crlf3 |
| gli31 | 2.4E-05 | 0.3686993 | 0.138 | 0.11 | 0.796503 | DF 4 | gli3 |
| park71 | 2.5E-05 | 0.2798411 | 0.181 | 0.15 | 0.797896 | DF 4 | park7 |
| dlg5a | 2.5E-05 | 0.2719043 | 0.126 | 0.099 | 0.820436 | DF 4 | dlg5a |
| tcf25 | 2.5E-05 | 0.320531 | 0.153 | 0.124 | 0.821693 | DF 4 | tcf25 |
| tent4b1 | 2.7E-05 | 0.2488303 | 0.237 | 0.203 | 0.880189 | DF 4 | tent4b |
| nhsl1a2 | 2.9E-05 | 0.3938001 | 0.102 | 0.078 | 0.930226 | DF 4 | nhsl1a |
| arl4ab2 | 3E-05 | 0.3646018 | 0.137 | 0.109 | 0.974938 | DF 4 | arl4ab |
| stau11 | 3.3E-05 | 0.2629189 | 0.172 | 0.142 | 1 | DF 4 | stau1 |
| nr1d11 | 3.4E-05 | 0.2805856 | 0.164 | 0.134 | 1 | DF 4 | nr1d1 |
| serpinb142 | 3.6E-05 | 0.2725537 | 0.184 | 0.152 | 1 | DF 4 | serpinb14 |
| ppp2r5cb | 3.7E-05 | 0.4525698 | 0.112 | 0.088 | 1 | DF 4 | ppp2r5cb |
| rin31 | 3.9E-05 | 0.2410619 | 0.184 | 0.153 | 1 | DF 4 | rin3 |
| zgc:100918 | 4E-05 | 0.242494 | 0.3 | 0.266 | 1 | DF 4 | zgc:100918 |
| smim19 | 4E-05 | 0.3821547 | 0.102 | 0.079 | 1 | DF 4 | smim19 |
| pnpla6 | 4.1E-05 | 0.222189 | 0.133 | 0.106 | 1 | DF 4 | pnpla6 |
| zmat21 | 4.3E-05 | 0.2452064 | 0.245 | 0.214 | 1 | DF 4 | zmat2 |
| anp32e1 | 4.3E-05 | 0.3060897 | 0.171 | 0.142 | 1 | DF 4 | anp32e |
| znf384l | 4.3E-05 | 0.3675146 | 0.143 | 0.117 | 1 | DF 4 | znf384l |
| setd3 | 4.8E-05 | 0.3094212 | 0.101 | 0.078 | 1 | DF 4 | setd3 |
| sult1st61 | 4.9E-05 | 0.2559276 | 0.143 | 0.115 | 1 | DF 4 | sult1st6 |
| tmbim4 | 5.2E-05 | 0.2765057 | 0.143 | 0.115 | 1 | DF 4 | tmbim4 |
| nr2f1a1 | 5.8E-05 | 0.250388 | 0.182 | 0.152 | 1 | DF 4 | nr2f1a |
| zbtb38 | 5.8E-05 | 0.4227541 | 0.104 | 0.081 | 1 | DF 4 | zbtb38 |
| slc16a1b3 | 5.9E-05 | 0.2070312 | 0.159 | 0.13 | 1 | DF 4 | slc16a1b |
| ezh12 | 6.5E-05 | 0.2856524 | 0.17 | 0.14 | 1 | DF 4 | ezh1 |
| rcor11 | 6.8E-05 | 0.3130288 | 0.161 | 0.133 | 1 | DF 4 | rcor1 |
| uqcc3 | 7.1E-05 | 0.3159141 | 0.106 | 0.083 | 1 | DF 4 | uqcc3 |
| tiparp1 | 7.4E-05 | 0.2108594 | 0.152 | 0.124 | 1 | DF 4 | tiparp |
| trim25 | 8E-05 | 0.3720556 | 0.119 | 0.095 | 1 | DF 4 | trim25 |
| rab18a | 8.5E-05 | 0.2703702 | 0.182 | 0.153 | 1 | DF 4 | rab18a |
| ano5b2 | 8.6E-05 | 0.3657924 | 0.109 | 0.086 | 1 | DF 4 | ano5b |
| abat1 | 8.6E-05 | 0.2806417 | 0.124 | 0.099 | 1 | DF 4 | abat |
| hipk3a | 8.7E-05 | 0.3126329 | 0.131 | 0.106 | 1 | DF 4 | hipk3a |
| si:dkey-56m19.52 | 9.2E-05 | 0.2289447 | 0.108 | 0.085 | 1 | DF 4 | si:dkey-56m19.5 |
| pkig1 | 9.3E-05 | 0.3184246 | 0.143 | 0.117 | 1 | DF 4 | pkig |
| eif3jb1 | 9.7E-05 | 0.2047968 | 0.258 | 0.224 | 1 | DF 4 | eif3jb |
| cnep1r11 | 0.0001 | 0.3248244 | 0.125 | 0.101 | 1 | DF 4 | cnep1r1 |
| foxo4 | 0.00011 | 0.4171791 | 0.13 | 0.106 | 1 | DF 4 | foxo4 |
| ptprsa.12 | 0.00011 | 0.2735493 | 0.151 | 0.124 | 1 | DF 4 | ptprsa.1 |
| kansl1b | 0.00012 | 0.2238086 | 0.2 | 0.169 | 1 | DF 4 | kansl1b |
| snap23.1 | 0.00014 | 0.2290193 | 0.212 | 0.182 | 1 | DF 4 | snap23.1 |
| si:ch211-71n6.41 | 0.00014 | 0.3230103 | 0.138 | 0.113 | 1 | DF 4 | si:ch211-71n6.4 |
| zfand5a1 | 0.00015 | 0.2015994 | 0.219 | 0.188 | 1 | DF 4 | zfand5a |
| cpvl1 | 0.00015 | 0.3197473 | 0.115 | 0.092 | 1 | DF 4 | cpvl |

| | | | | | | | |
|---|---|---|---|---|---|---|---|
| ehmt1b | 0.00015 | 0.2618805 | 0.206 | 0.176 | 1 | DF 4 | ehmt1b |
| smim13 | 0.00015 | 0.2900389 | 0.145 | 0.12 | 1 | DF 4 | smim13 |
| slc44a22 | 0.00017 | 0.2240262 | 0.141 | 0.114 | 1 | DF 4 | slc44a2 |
| prkd3 | 0.00018 | 0.3105728 | 0.101 | 0.079 | 1 | DF 4 | prkd3 |
| zgc:1102393 | 0.00018 | 0.2001078 | 0.263 | 0.231 | 1 | DF 4 | zgc:110239 |
| larp4ab1 | 0.00018 | 0.2197331 | 0.188 | 0.159 | 1 | DF 4 | larp4ab |
| mib1 | 0.00018 | 0.269801 | 0.149 | 0.124 | 1 | DF 4 | mib1 |
| rybpb | 0.00019 | 0.3168609 | 0.11 | 0.088 | 1 | DF 4 | rybpb |
| stk35l | 0.0002 | 0.2666763 | 0.151 | 0.125 | 1 | DF 4 | stk35l |
| mapk3 | 0.00021 | 0.2561903 | 0.141 | 0.117 | 1 | DF 4 | mapk3 |
| slc25a28 | 0.00022 | 0.3801223 | 0.106 | 0.085 | 1 | DF 4 | slc25a28 |
| lrig2 | 0.00022 | 0.3264979 | 0.114 | 0.092 | 1 | DF 4 | lrig2 |
| snx31 | 0.00024 | 0.2652697 | 0.204 | 0.177 | 1 | DF 4 | snx3 |
| zswim81 | 0.00024 | 0.2865202 | 0.103 | 0.081 | 1 | DF 4 | zswim8 |
| si:ch211-191j22.3 | 0.00026 | 0.2272059 | 0.144 | 0.119 | 1 | DF 4 | si:ch211-191j22.3 |
| si:dkey-6e2.2 | 0.00026 | 0.3442411 | 0.112 | 0.091 | 1 | DF 4 | si:dkey-6e2.2 |
| fbxw11b | 0.00026 | 0.325557 | 0.185 | 0.159 | 1 | DF 4 | fbxw11b |
| jade11 | 0.00026 | 0.318123 | 0.125 | 0.102 | 1 | DF 4 | jade1 |
| tcf20 | 0.00026 | 0.2528336 | 0.168 | 0.141 | 1 | DF 4 | tcf20 |
| nsd1a | 0.00027 | 0.4068972 | 0.114 | 0.092 | 1 | DF 4 | nsd1a |
| eif4ebp3l2 | 0.00027 | 0.2433305 | 0.18 | 0.154 | 1 | DF 4 | eif4ebp3l |
| hexim11 | 0.00028 | 0.2055278 | 0.263 | 0.233 | 1 | DF 4 | hexim1 |
| ctsa | 0.00028 | 0.2690568 | 0.141 | 0.117 | 1 | DF 4 | ctsa |
| chuk | 0.00029 | 0.35779 | 0.106 | 0.085 | 1 | DF 4 | chuk |
| arvcfb2 | 0.0003 | 0.3937862 | 0.125 | 0.102 | 1 | DF 4 | arvcfb |
| kank42 | 0.0003 | 0.2802313 | 0.126 | 0.103 | 1 | DF 4 | kank4 |
| rab8b1 | 0.00031 | 0.2465219 | 0.162 | 0.136 | 1 | DF 4 | rab8b |
| nfyba1 | 0.00031 | 0.2544839 | 0.21 | 0.182 | 1 | DF 4 | nfyba |
| man2a2 | 0.00033 | 0.3252623 | 0.154 | 0.129 | 1 | DF 4 | man2a2 |
| aktip | 0.00041 | 0.4200148 | 0.118 | 0.097 | 1 | DF 4 | aktip |
| trip111 | 0.00042 | 0.2812103 | 0.112 | 0.091 | 1 | DF 4 | trip11 |
| rbm39b | 0.00045 | 0.3747984 | 0.126 | 0.104 | 1 | DF 4 | rbm39b |
| scaf4a | 0.00046 | 0.2901516 | 0.111 | 0.09 | 1 | DF 4 | scaf4a |
| fam168b | 0.00046 | 0.3111416 | 0.149 | 0.126 | 1 | DF 4 | fam168b |
| bckdk1 | 0.00047 | 0.275599 | 0.109 | 0.088 | 1 | DF 4 | bckdk |
| lgals9l32 | 0.00049 | 0.2554508 | 0.146 | 0.122 | 1 | DF 4 | lgals9l3 |
| si:dkey-253d23.4 | 0.00053 | 0.2299642 | 0.109 | 0.088 | 1 | DF 4 | si:dkey-253d23.4 |
| rnf241 | 0.00056 | 0.3292605 | 0.113 | 0.092 | 1 | DF 4 | rnf24 |
| cry1a | 0.00057 | 0.2437072 | 0.188 | 0.161 | 1 | DF 4 | cry1a |
| rtraf1 | 0.00063 | 0.3536346 | 0.125 | 0.105 | 1 | DF 4 | rtraf |
| SSBP41 | 0.00065 | 0.2284995 | 0.144 | 0.121 | 1 | DF 4 | SSBP4 |
| si:ch73-308l14.21 | 0.00065 | 0.2740732 | 0.131 | 0.109 | 1 | DF 4 | si:ch73-308l14.2 |
| tbc1d10aa | 0.00071 | 0.2348745 | 0.138 | 0.115 | 1 | DF 4 | tbc1d10aa |
| uri11 | 0.00077 | 0.3602884 | 0.151 | 0.129 | 1 | DF 4 | uri1 |
| mgat1b2 | 0.0008 | 0.324358 | 0.117 | 0.097 | 1 | DF 4 | mgat1b |
| rnf13 | 0.00081 | 0.2092638 | 0.151 | 0.128 | 1 | DF 4 | rnf13 |
| RAPH12 | 0.00084 | 0.2210159 | 0.111 | 0.09 | 1 | DF 4 | RAPH1 |
| ggcx2 | 0.0009 | 0.2281758 | 0.191 | 0.166 | 1 | DF 4 | ggcx |

| | | | | | | | |
|---|---|---|---|---|---|---|---|
| dcaf7 | 0.0009 | 0.3588008 | 0.15 | 0.129 | 1 | DF 4 | dcaf7 |
| sft2d1 | 0.0009 | 0.2082197 | 0.122 | 0.101 | 1 | DF 4 | sft2d1 |
| helz | 0.00094 | 0.3537366 | 0.106 | 0.087 | 1 | DF 4 | helz |
| caska3 | 0.00095 | 0.2389228 | 0.106 | 0.086 | 1 | DF 4 | caska |
| zgc:1527913 | 0.001 | 0.2410022 | 0.127 | 0.105 | 1 | DF 4 | zgc:152791 |
| morf4l11 | 0.00106 | 0.2459948 | 0.202 | 0.177 | 1 | DF 4 | morf4l1 |
| dnase1l1 | 0.00112 | 0.3289508 | 0.135 | 0.114 | 1 | DF 4 | dnase1l1 |
| ldb1a | 0.00113 | 0.3391595 | 0.125 | 0.105 | 1 | DF 4 | ldb1a |
| rab22a1 | 0.00116 | 0.2893943 | 0.122 | 0.102 | 1 | DF 4 | rab22a |
| znf1035 | 0.00124 | 0.2084645 | 0.187 | 0.162 | 1 | DF 4 | znf1035 |
| atrnl1b1 | 0.00126 | 0.2209798 | 0.13 | 0.109 | 1 | DF 4 | atrnl1b |
| gskip2 | 0.00127 | 0.2683997 | 0.184 | 0.161 | 1 | DF 4 | gskip |
| cpn13 | 0.00127 | 0.2149225 | 0.238 | 0.211 | 1 | DF 4 | cpn1 |
| nr1d2b2 | 0.0013 | 0.2391154 | 0.175 | 0.152 | 1 | DF 4 | nr1d2b |
| ppp2r5d | 0.00132 | 0.27488 | 0.135 | 0.114 | 1 | DF 4 | ppp2r5d |
| dchs1b4 | 0.00141 | 0.227419 | 0.11 | 0.091 | 1 | DF 4 | dchs1b |
| dnaja2b1 | 0.00142 | 0.2731348 | 0.131 | 0.111 | 1 | DF 4 | dnaja2b |
| lamtor51 | 0.00145 | 0.3597125 | 0.106 | 0.087 | 1 | DF 4 | lamtor5 |
| lamp1b | 0.00146 | 0.2457811 | 0.108 | 0.089 | 1 | DF 4 | lamp1b |
| cicb2 | 0.00175 | 0.2097097 | 0.149 | 0.127 | 1 | DF 4 | cicb |
| fam199x | 0.00182 | 0.231814 | 0.154 | 0.132 | 1 | DF 4 | fam199x |
| vgll4b1 | 0.00184 | 0.235779 | 0.185 | 0.163 | 1 | DF 4 | vgll4b |
| wasla1 | 0.00196 | 0.2292868 | 0.201 | 0.177 | 1 | DF 4 | wasla |
| trib23 | 0.00207 | 0.3297835 | 0.106 | 0.088 | 1 | DF 4 | trib2 |
| arpp19a | 0.00209 | 0.3278424 | 0.101 | 0.083 | 1 | DF 4 | arpp19a |
| sh2b1 | 0.00222 | 0.2944425 | 0.101 | 0.084 | 1 | DF 4 | sh2b1 |
| BX005309.11 | 0.00222 | 0.2572521 | 0.1 | 0.082 | 1 | DF 4 | BX005309.1 |
| slc48a1a1 | 0.00228 | 0.2841108 | 0.107 | 0.089 | 1 | DF 4 | slc48a1a |
| mgaa | 0.00229 | 0.213104 | 0.163 | 0.14 | 1 | DF 4 | mgaa |
| mbtps1 | 0.00266 | 0.2672665 | 0.128 | 0.109 | 1 | DF 4 | mbtps1 |
| schip11 | 0.00274 | 0.3021214 | 0.165 | 0.145 | 1 | DF 4 | schip1 |
| si:ch211-276i12.9 | 0.00275 | 0.2317389 | 0.109 | 0.091 | 1 | DF 4 | si:ch211-276i12.9 |
| ube2h | 0.00293 | 0.2898493 | 0.129 | 0.11 | 1 | DF 4 | ube2h |
| fcho21 | 0.00326 | 0.2328488 | 0.18 | 0.159 | 1 | DF 4 | fcho2 |
| araf | 0.00366 | 0.291199 | 0.113 | 0.095 | 1 | DF 4 | araf |
| prss161 | 0.00391 | 0.2345049 | 0.107 | 0.09 | 1 | DF 4 | prss16 |
| mdc12 | 0.004 | 0.2886206 | 0.123 | 0.105 | 1 | DF 4 | mdc1 |
| slc9a6a | 0.00412 | 0.2821208 | 0.101 | 0.084 | 1 | DF 4 | slc9a6a |
| sod21 | 0.00419 | 0.2906399 | 0.147 | 0.128 | 1 | DF 4 | sod2 |
| ctbp11 | 0.0042 | 0.2332063 | 0.119 | 0.102 | 1 | DF 4 | ctbp1 |
| nudt3b | 0.00469 | 0.2726226 | 0.113 | 0.096 | 1 | DF 4 | nudt3b |
| arpc5b1 | 0.00482 | 0.2250547 | 0.104 | 0.087 | 1 | DF 4 | arpc5b |
| ythdc11 | 0.00515 | 0.2458717 | 0.149 | 0.129 | 1 | DF 4 | ythdc1 |
| znf654 | 0.00516 | 0.2814071 | 0.111 | 0.094 | 1 | DF 4 | znf654 |
| mta21 | 0.00523 | 0.3470662 | 0.109 | 0.093 | 1 | DF 4 | mta2 |
| ctnnd11 | 0.00565 | 0.2246537 | 0.186 | 0.166 | 1 | DF 4 | ctnnd1 |
| mapk14a2 | 0.00567 | 0.2343537 | 0.1 | 0.084 | 1 | DF 4 | mapk14a |
| pak2b1 | 0.00603 | 0.2158333 | 0.167 | 0.148 | 1 | DF 4 | pak2b |

| | | | | | | | |
|---|---|---|---|---|---|---|---|
| birc61 | 0.0061 | 0.2503647 | 0.131 | 0.113 | 1 | DF 4 | birc6 |
| slc25a331 | 0.00616 | 0.2772759 | 0.134 | 0.117 | 1 | DF 4 | slc25a33 |
| si:ch211-198a12.6 | 0.00617 | 0.2329813 | 0.129 | 0.111 | 1 | DF 4 | si:ch211-198a12.6 |
| zgc:1626981 | 0.00632 | 0.2927811 | 0.131 | 0.114 | 1 | DF 4 | zgc:162698 |
| ap3m11 | 0.00648 | 0.2170964 | 0.1 | 0.085 | 1 | DF 4 | ap3m1 |
| prkar1aa | 0.00655 | 0.2372724 | 0.162 | 0.143 | 1 | DF 4 | prkar1aa |
| sinhcafl1 | 0.00731 | 0.200087 | 0.143 | 0.125 | 1 | DF 4 | sinhcafl |
| sufu | 0.00756 | 0.2215209 | 0.102 | 0.086 | 1 | DF 4 | sufu |
| cry3a | 0.00773 | 0.2385372 | 0.11 | 0.094 | 1 | DF 4 | cry3a |
| tsg101b | 0.00788 | 0.2954429 | 0.106 | 0.09 | 1 | DF 4 | tsg101b |
| srsf5b2 | 0.00871 | 0.2552913 | 0.132 | 0.115 | 1 | DF 4 | srsf5b |
| c6.12 | 0 | 3.3431389 | 0.693 | 0.181 | 0 | Stroma 3 | c6.1 |
| fgl2b2 | 0 | 3.3680597 | 0.579 | 0.1 | 0 | Stroma 3 | fgl2b |
| nova22 | 0 | 2.0459044 | 0.614 | 0.194 | 0 | Stroma 3 | nova2 |
| col4a12 | 0 | 2.9939548 | 0.611 | 0.208 | 0 | Stroma 3 | col4a1 |
| krt943 | 0 | 1.9928655 | 0.679 | 0.277 | 0 | Stroma 3 | krt94 |
| fabp11a1 | 0 | 4.1033205 | 0.453 | 0.054 | 0 | Stroma 3 | fabp11a |
| col4a22 | 0 | 2.6655629 | 0.58 | 0.201 | 0 | Stroma 3 | col4a2 |
| fn1a2 | 0 | 1.7352226 | 0.667 | 0.301 | 0 | Stroma 3 | fn1a |
| CR925728.12 | 0 | 3.1893273 | 0.44 | 0.076 | 0 | Stroma 3 | CR925728.1 |
| fn1b1 | 0 | 1.6108146 | 0.747 | 0.385 | 0 | Stroma 3 | fn1b |
| si:ch211-227m13.11 | 0 | 3.9967816 | 0.393 | 0.037 | 0 | Stroma 3 | si:ch211-227m13.1 |
| si:dkey-261h17.13 | 0 | 1.968408 | 0.624 | 0.275 | 0 | Stroma 3 | si:dkey-261h17.1 |
| neu3.51 | 0 | 3.4261102 | 0.393 | 0.053 | 0 | Stroma 3 | neu3.5 |
| akap12b2 | 0 | 2.480508 | 0.46 | 0.128 | 0 | Stroma 3 | akap12b |
| col5a3a | 0 | 3.9071978 | 0.376 | 0.052 | 0 | Stroma 3 | col5a3a |
| add3a3 | 0 | 1.692349 | 0.75 | 0.434 | 0 | Stroma 3 | add3a |
| phlda22 | 0 | 2.3432585 | 0.517 | 0.205 | 0 | Stroma 3 | phlda2 |
| c43 | 0 | 2.4501212 | 0.503 | 0.193 | 0 | Stroma 3 | c4 |
| zgc:174863 | 0 | 3.6006428 | 0.514 | 0.216 | 0 | Stroma 3 | zgc:174863 |
| ccl19a.1 | 0 | 5.968985 | 0.299 | 0.008 | 0 | Stroma 3 | ccl19a.1 |
| hic11 | 0 | 3.0805244 | 0.297 | 0.053 | 0 | Stroma 3 | hic1 |
| marcksl1a2 | 0 | 1.869009 | 0.935 | 0.696 | 0 | Stroma 3 | marcksl1a |
| itih5 | 0 | 2.1194574 | 0.296 | 0.058 | 0 | Stroma 3 | itih5 |
| pmp22a2 | 0 | 1.6098208 | 0.916 | 0.678 | 0 | Stroma 3 | pmp22a |
| clec19a2 | 0 | 2.010045 | 0.278 | 0.053 | 0 | Stroma 3 | clec19a |
| ebf3a.1 | 0 | 3.0700432 | 0.247 | 0.038 | 0 | Stroma 3 | ebf3a.1 |
| AL954191.2 | 0 | 4.975959 | 0.208 | 0.011 | 0 | Stroma 3 | AL954191.2 |
| vwde | 0 | 3.3532088 | 0.196 | 0.019 | 0 | Stroma 3 | vwde |
| cyp2ad3 | 0 | 4.1309431 | 0.167 | 0.014 | 0 | Stroma 3 | cyp2ad3 |
| flnb | 0 | 4.694316 | 0.15 | 0.01 | 0 | Stroma 3 | flnb |
| socs3a1 | 1E-306 | 1.6056585 | 0.935 | 0.788 | 3.4E-302 | Stroma 3 | socs3a |
| rgs5b4 | 4E-299 | 1.9040822 | 0.812 | 0.509 | 1.2E-294 | Stroma 3 | rgs5b |
| cxcl18b2 | 8E-287 | 1.4843247 | 0.892 | 0.693 | 2.4E-282 | Stroma 3 | cxcl18b |
| thbs1a1 | 1E-284 | 2.216573 | 0.604 | 0.299 | 3.6E-280 | Stroma 3 | thbs1a |
| nt5e1 | 5E-283 | 3.0372622 | 0.25 | 0.055 | 1.6E-278 | Stroma 3 | nt5e |
| cox6a2 | 7E-277 | 2.6449509 | 0.212 | 0.04 | 2.4E-272 | Stroma 3 | cox6a2 |
| htra1b2 | 9E-277 | 1.5737151 | 0.707 | 0.398 | 3.1E-272 | Stroma 3 | htra1b |

| | | | | | | | |
|---|---|---|---|---|---|---|---|
| cilp4 | 1E-275 | 2.16678 | 0.438 | 0.159 | 4E-271 | Stroma 3 | cilp |
| mast22 | 4E-272 | 2.2193338 | 0.424 | 0.152 | 1.4E-267 | Stroma 3 | mast2 |
| ablim21 | 9E-271 | 2.7573145 | 0.208 | 0.04 | 3E-266 | Stroma 3 | ablim2 |
| si:ch211-198c19.32 | 2E-267 | 1.9530778 | 0.499 | 0.198 | 7E-263 | Stroma 3 | si:ch211-198c19.3 |
| ace | 9E-267 | 3.0081432 | 0.214 | 0.043 | 2.8E-262 | Stroma 3 | ace |
| rspo32 | 6E-259 | 2.4812968 | 0.245 | 0.056 | 1.9E-254 | Stroma 3 | rspo3 |
| tfpia4 | 2E-245 | 2.0006641 | 0.434 | 0.167 | 7.4E-241 | Stroma 3 | tfpia |
| plpp1a4 | 1E-239 | 1.6514088 | 0.657 | 0.362 | 4.4E-235 | Stroma 3 | plpp1a |
| hsd11b24 | 1E-232 | 2.5160712 | 0.291 | 0.084 | 3.6E-228 | Stroma 3 | hsd11b2 |
| dpysl32 | 1E-231 | 1.8964907 | 0.406 | 0.15 | 3.8E-227 | Stroma 3 | dpysl3 |
| rgcc1 | 1E-228 | 2.0696314 | 0.454 | 0.186 | 3.8E-224 | Stroma 3 | rgcc |
| dcn4 | 3E-228 | 0.8253795 | 0.968 | 0.864 | 8.4E-224 | Stroma 3 | dcn |
| c63 | 8E-219 | 2.4674197 | 0.246 | 0.064 | 2.7E-214 | Stroma 3 | c6 |
| hspb12 | 9E-217 | 1.7048539 | 0.425 | 0.168 | 2.8E-212 | Stroma 3 | hspb1 |
| prelp2 | 2E-213 | 1.9060475 | 0.353 | 0.123 | 6E-209 | Stroma 3 | prelp |
| mdka3 | 4E-206 | 1.064201 | 0.802 | 0.509 | 1.3E-201 | Stroma 3 | mdka |
| nfia2 | 2E-205 | 1.8513993 | 0.401 | 0.159 | 6.1E-201 | Stroma 3 | nfia |
| nid1a1 | 2E-203 | 1.8892251 | 0.475 | 0.223 | 6.2E-199 | Stroma 3 | nid1a |
| mdkb1 | 3E-198 | 2.6636412 | 0.171 | 0.036 | 1E-193 | Stroma 3 | mdkb |
| cd28 | 2E-193 | 3.3157551 | 0.113 | 0.016 | 6.1E-189 | Stroma 3 | cd28 |
| lama42 | 6E-192 | 1.7855451 | 0.511 | 0.259 | 2E-187 | Stroma 3 | lama4 |
| si:dkey-238i5.22 | 5E-191 | 2.1017829 | 0.242 | 0.067 | 1.6E-186 | Stroma 3 | si:dkey-238i5.2 |
| angptl1a2 | 2E-190 | 1.4543362 | 0.636 | 0.381 | 6.9E-186 | Stroma 3 | angptl1a |
| c1qtnf22 | 6E-187 | 1.9496088 | 0.272 | 0.084 | 1.9E-182 | Stroma 3 | c1qtnf2 |
| adam10a3 | 4E-185 | 1.6471325 | 0.598 | 0.371 | 1.2E-180 | Stroma 3 | adam10a |
| aldh1a22 | 1E-183 | 2.3855203 | 0.195 | 0.048 | 4.4E-179 | Stroma 3 | aldh1a2 |
| foxp43 | 3E-180 | 1.5050975 | 0.438 | 0.192 | 8.7E-176 | Stroma 3 | foxp4 |
| wu:fj16a034 | 2E-178 | 1.2521502 | 0.431 | 0.18 | 5E-174 | Stroma 3 | wu:fj16a03 |
| si:ch211-212k18.71 | 4E-177 | 2.2038508 | 0.212 | 0.057 | 1.3E-172 | Stroma 3 | si:ch211-212k18.7 |
| peli1a3 | 1E-172 | 1.959328 | 0.322 | 0.12 | 4E-168 | Stroma 3 | peli1a |
| fam107b2 | 2E-172 | 1.5761198 | 0.437 | 0.198 | 5.2E-168 | Stroma 3 | fam107b |
| dhrs3a2 | 9E-166 | 2.4963414 | 0.214 | 0.061 | 3E-161 | Stroma 3 | dhrs3a |
| fstl1b1 | 3E-165 | 0.9192715 | 0.862 | 0.752 | 9.1E-161 | Stroma 3 | fstl1b |
| dock9b3 | 1E-160 | 2.0905481 | 0.269 | 0.092 | 3.6E-156 | Stroma 3 | dock9b |
| tns2b | 2E-159 | 2.8954069 | 0.146 | 0.032 | 5.3E-155 | Stroma 3 | tns2b |
| vasna2 | 3E-159 | 1.9844439 | 0.292 | 0.106 | 9.2E-155 | Stroma 3 | vasna |
| c7a4 | 2E-155 | 1.8151176 | 0.375 | 0.165 | 6.8E-151 | Stroma 3 | c7a |
| lamb1b1 | 1E-154 | 1.780668 | 0.4 | 0.186 | 4.5E-150 | Stroma 3 | lamb1b |
| tmem1764 | 4E-149 | 1.2618147 | 0.603 | 0.384 | 1.3E-144 | Stroma 3 | tmem176 |
| plpp34 | 1E-148 | 1.1174249 | 0.681 | 0.444 | 3.4E-144 | Stroma 3 | plpp3 |
| her93 | 2E-148 | 2.1919526 | 0.428 | 0.219 | 5.7E-144 | Stroma 3 | her9 |
| emp21 | 5E-148 | 2.1544448 | 0.335 | 0.144 | 1.7E-143 | Stroma 3 | emp2 |
| ognb4 | 3E-144 | 1.7611339 | 0.311 | 0.124 | 1.1E-139 | Stroma 3 | ognb |
| vat13 | 1E-142 | 1.8604827 | 0.512 | 0.306 | 3.5E-138 | Stroma 3 | vat1 |
| aplp1 | 2E-142 | 2.304339 | 0.153 | 0.037 | 7.6E-138 | Stroma 3 | aplp1 |
| zeb1b2 | 9E-140 | 1.9367641 | 0.238 | 0.081 | 2.9E-135 | Stroma 3 | zeb1b |
| junba1 | 3E-139 | 0.860199 | 0.929 | 0.842 | 1.1E-134 | Stroma 3 | junba |
| fgl2a4 | 1E-138 | 1.7937943 | 0.344 | 0.149 | 3.6E-134 | Stroma 3 | fgl2a |

| | | | | | | | |
|---|---|---|---|---|---|---|---|
| nid1b2 | 3E-133 | 2.2649614 | 0.218 | 0.073 | 9.5E-129 | Stroma 3 | nid1b |
| sptan12 | 5E-131 | 1.5236526 | 0.438 | 0.236 | 1.7E-126 | Stroma 3 | sptan1 |
| pim12 | 2E-130 | 1.2311655 | 0.719 | 0.537 | 7.6E-126 | Stroma 3 | pim1 |
| endouc2 | 1E-129 | 2.0228155 | 0.422 | 0.228 | 4.7E-125 | Stroma 3 | endouc |
| ntn1a3 | 2E-129 | 1.6828577 | 0.288 | 0.113 | 6.7E-125 | Stroma 3 | ntn1a |
| lamc12 | 4E-129 | 1.8664726 | 0.293 | 0.121 | 1.4E-124 | Stroma 3 | lamc1 |
| kirrel1a2 | 8E-129 | 1.4403522 | 0.518 | 0.317 | 2.5E-124 | Stroma 3 | kirrel1a |
| CR383676.24 | 2E-127 | 0.4169 | 1 | 1 | 5.8E-123 | Stroma 3 | CR383676.2 |
| fstl1a2 | 5E-127 | 1.2206677 | 0.269 | 0.099 | 1.6E-122 | Stroma 3 | fstl1a |
| zfp36l1b3 | 3E-126 | 1.0841799 | 0.721 | 0.538 | 1E-121 | Stroma 3 | zfp36l1b |
| f3b3 | 1E-124 | 1.1068483 | 0.591 | 0.37 | 3.3E-120 | Stroma 3 | f3b |
| jun1 | 3E-124 | 0.8110033 | 0.941 | 0.85 | 1.1E-119 | Stroma 3 | jun |
| ablim1b3 | 3E-123 | 1.4191917 | 0.455 | 0.247 | 8.7E-119 | Stroma 3 | ablim1b |
| crabp2a2 | 3E-123 | 1.4904266 | 0.341 | 0.154 | 8.8E-119 | Stroma 3 | crabp2a |
| mfap54 | 1E-122 | 0.9516717 | 0.739 | 0.537 | 3.9E-118 | Stroma 3 | mfap5 |
| mgp | 3E-120 | 0.4499301 | 0.339 | 0.151 | 8.6E-116 | Stroma 3 | mgp |
| si:ch211-105c13.34 | 2E-118 | 0.8507735 | 0.752 | 0.561 | 6.4E-114 | Stroma 3 | si:ch211-105c13.3 |
| tmem178b | 3E-117 | 3.2718578 | 0.101 | 0.021 | 8.2E-113 | Stroma 3 | tmem178b |
| gadd45ba1 | 8E-115 | 0.9857802 | 0.881 | 0.8 | 2.7E-110 | Stroma 3 | gadd45ba |
| arhgef41 | 7E-114 | 1.9449103 | 0.203 | 0.07 | 2.4E-109 | Stroma 3 | arhgef4 |
| CU914776.12 | 1E-113 | 1.6294758 | 0.219 | 0.078 | 3.7E-109 | Stroma 3 | CU914776.1 |
| tgfbr2b3 | 1E-109 | 1.7013656 | 0.301 | 0.136 | 3.3E-105 | Stroma 3 | tgfbr2b |
| CR753876.1 | 5E-107 | 2.8312755 | 0.127 | 0.033 | 1.5E-102 | Stroma 3 | CR753876.1 |
| tgm2b | 2E-106 | 1.8143442 | 0.146 | 0.042 | 5E-102 | Stroma 3 | tgm2b |
| sptbn12 | 1E-105 | 1.3705902 | 0.427 | 0.241 | 4.4E-101 | Stroma 3 | sptbn1 |
| bnc23 | 7E-103 | 1.826254 | 0.207 | 0.076 | 2.29E-98 | Stroma 3 | bnc2 |
| wnt9a1 | 5E-102 | 2.3856542 | 0.108 | 0.026 | 1.78E-97 | Stroma 3 | wnt9a |
| celf24 | 1E-101 | 1.2145271 | 0.374 | 0.19 | 4.5E-97 | Stroma 3 | celf2 |
| sulf12 | 2E-100 | 1.7404065 | 0.298 | 0.14 | 5.38E-96 | Stroma 3 | sulf1 |
| zfp36l21 | 3E-100 | 1.4850513 | 0.436 | 0.26 | 9.9E-96 | Stroma 3 | zfp36l2 |
| arsj | 1E-99 | 2.2248757 | 0.122 | 0.032 | 4.55E-95 | Stroma 3 | arsj |
| CABZ01092746.14 | 4E-99 | 0.632478 | 0.852 | 0.703 | 1.34E-94 | Stroma 3 | CABZ01092746.1 |
| ehbp1l1b1 | 1.2E-96 | 1.5225791 | 0.271 | 0.12 | 3.77E-92 | Stroma 3 | ehbp1l1b |
| mustn1a2 | 2.3E-96 | 1.5357437 | 0.204 | 0.076 | 7.54E-92 | Stroma 3 | mustn1a |
| b2ml3 | 4.8E-96 | 0.5432137 | 0.924 | 0.896 | 1.58E-91 | Stroma 3 | b2ml |
| rdh10a1 | 1.3E-95 | 2.1631728 | 0.126 | 0.035 | 4.07E-91 | Stroma 3 | rdh10a |
| irf8 | 5.2E-91 | 2.8800692 | 0.101 | 0.025 | 1.7E-86 | Stroma 3 | irf8 |
| abca1a2 | 9.7E-91 | 1.4713202 | 0.352 | 0.19 | 3.16E-86 | Stroma 3 | abca1a |
| pcdh191 | 1.3E-89 | 2.0565988 | 0.149 | 0.049 | 4.24E-85 | Stroma 3 | pcdh19 |
| egr12 | 1.4E-89 | 0.8596811 | 0.765 | 0.622 | 4.39E-85 | Stroma 3 | egr1 |
| cyp2p7 | 3.8E-88 | 2.044331 | 0.135 | 0.041 | 1.24E-83 | Stroma 3 | cyp2p7 |
| adcyap1b1 | 5.6E-87 | 1.5074413 | 0.136 | 0.042 | 1.84E-82 | Stroma 3 | adcyap1b |
| meox2a | 7.4E-86 | 2.1539056 | 0.122 | 0.036 | 2.42E-81 | Stroma 3 | meox2a |
| vim3 | 1.6E-83 | 1.057695 | 0.544 | 0.362 | 5.17E-79 | Stroma 3 | vim |
| uap12 | 3E-83 | 1.4103722 | 0.265 | 0.126 | 9.7E-79 | Stroma 3 | uap1 |
| serpinf14 | 8.2E-83 | 0.7849549 | 0.698 | 0.548 | 2.65E-78 | Stroma 3 | serpinf1 |
| ppdpfa1 | 8.5E-83 | 3.023803 | 0.108 | 0.03 | 2.76E-78 | Stroma 3 | ppdpfa |
| scara32 | 3E-82 | 1.3891219 | 0.271 | 0.128 | 9.62E-78 | Stroma 3 | scara3 |

| | | | | | | | |
|---|---|---|---|---|---|---|---|
| adh8a3 | 3E-82 | 0.9239604 | 0.377 | 0.204 | 9.63E-78 | Stroma 3 | adh8a |
| zgc:1583431 | 4.9E-81 | 1.4209193 | 0.529 | 0.371 | 1.6E-76 | Stroma 3 | zgc:158343 |
| thbs3a2 | 5.1E-81 | 0.9303303 | 0.479 | 0.301 | 1.66E-76 | Stroma 3 | thbs3a |
| zcchc244 | 9.2E-80 | 1.049755 | 0.418 | 0.249 | 3.01E-75 | Stroma 3 | zcchc24 |
| si:ch211-195b13.12 | 1.1E-79 | 1.1725419 | 0.398 | 0.236 | 3.72E-75 | Stroma 3 | si:ch211-195b13.1 |
| fosab1 | 8.5E-79 | 0.5642909 | 0.951 | 0.893 | 2.76E-74 | Stroma 3 | fosab |
| cxcl12a5 | 1.2E-77 | 1.2079673 | 0.792 | 0.678 | 3.78E-73 | Stroma 3 | cxcl12a |
| ramp23 | 3.8E-77 | 1.5881217 | 0.274 | 0.136 | 1.23E-72 | Stroma 3 | ramp2 |
| fbln22 | 1.3E-76 | 1.3197005 | 0.25 | 0.117 | 4.23E-72 | Stroma 3 | fbln2 |
| tppp31 | 1.1E-75 | 1.2949685 | 0.154 | 0.055 | 3.62E-71 | Stroma 3 | tppp3 |
| EPB41L21 | 2.5E-75 | 1.5921646 | 0.25 | 0.121 | 8.08E-71 | Stroma 3 | EPB41L2 |
| CABZ01064941.13 | 2.6E-74 | 1.6562407 | 0.192 | 0.08 | 8.56E-70 | Stroma 3 | CABZ01064941.1 |
| ch25hl33 | 4.6E-74 | 1.443058 | 0.224 | 0.102 | 1.48E-69 | Stroma 3 | ch25hl3 |
| klf2a3 | 9.2E-74 | 1.136873 | 0.438 | 0.279 | 3E-69 | Stroma 3 | klf2a |
| c1qtnf51 | 1.3E-73 | 1.3097549 | 0.285 | 0.146 | 4.11E-69 | Stroma 3 | c1qtnf5 |
| specc1la1 | 8.1E-73 | 1.6306777 | 0.183 | 0.075 | 2.64E-68 | Stroma 3 | specc1la |
| clic21 | 2.7E-72 | 0.7542734 | 0.402 | 0.24 | 8.94E-68 | Stroma 3 | clic2 |
| dpt2 | 3.9E-72 | 0.9394628 | 0.291 | 0.148 | 1.28E-67 | Stroma 3 | dpt |
| mgst1.22 | 1E-70 | 1.0143292 | 0.451 | 0.294 | 3.41E-66 | Stroma 3 | mgst1.2 |
| lgmn2 | 5.7E-69 | 1.3340858 | 0.302 | 0.166 | 1.85E-64 | Stroma 3 | lgmn |
| adamts15a4 | 6.5E-69 | 1.1352985 | 0.272 | 0.137 | 2.11E-64 | Stroma 3 | adamts15a |
| tsku3 | 9.9E-69 | 0.9767902 | 0.463 | 0.313 | 3.23E-64 | Stroma 3 | tsku |
| sult1st62 | 2.7E-68 | 1.4320331 | 0.231 | 0.111 | 8.79E-64 | Stroma 3 | sult1st6 |
| CR855311.12 | 3.9E-68 | 0.6963405 | 0.289 | 0.148 | 1.25E-63 | Stroma 3 | CR855311.1 |
| adamts15b1 | 1.5E-67 | 1.5786356 | 0.138 | 0.049 | 4.8E-63 | Stroma 3 | adamts15b |
| ddr2a1 | 1.9E-67 | 1.187893 | 0.343 | 0.202 | 6.09E-63 | Stroma 3 | ddr2a |
| icam32 | 2.1E-67 | 1.6504714 | 0.183 | 0.078 | 6.68E-63 | Stroma 3 | icam3 |
| si:ch1073-268j14.1 | 4.4E-67 | 1.6922756 | 0.12 | 0.04 | 1.42E-62 | Stroma 3 | si:ch1073-268j14.1 |
| angptl42 | 2.1E-66 | 0.9545908 | 0.591 | 0.45 | 6.81E-62 | Stroma 3 | angptl4 |
| loxl13 | 6.8E-66 | 1.056963 | 0.341 | 0.195 | 2.21E-61 | Stroma 3 | loxl1 |
| dennd2b | 7.7E-66 | 1.4727197 | 0.243 | 0.122 | 2.49E-61 | Stroma 3 | dennd2b |
| itga53 | 1E-65 | 1.0435837 | 0.399 | 0.25 | 3.34E-61 | Stroma 3 | itga5 |
| pxdn | 1.5E-65 | 1.742079 | 0.119 | 0.04 | 4.76E-61 | Stroma 3 | pxdn |
| antxr2a3 | 5.7E-65 | 0.9493805 | 0.473 | 0.329 | 1.87E-60 | Stroma 3 | antxr2a |
| arrdc3a2 | 6.2E-65 | 1.401012 | 0.292 | 0.159 | 2.01E-60 | Stroma 3 | arrdc3a |
| rhogb1 | 7.7E-65 | 1.6360989 | 0.163 | 0.066 | 2.5E-60 | Stroma 3 | rhogb |
| crip13 | 8.4E-65 | 0.6820705 | 0.599 | 0.456 | 2.73E-60 | Stroma 3 | crip1 |
| lrp1aa3 | 1.1E-63 | 0.6113038 | 0.765 | 0.67 | 3.66E-59 | Stroma 3 | lrp1aa |
| spry41 | 2.6E-62 | 1.4339648 | 0.301 | 0.173 | 8.59E-58 | Stroma 3 | spry4 |
| b2m4 | 2.7E-61 | 0.5759401 | 0.759 | 0.683 | 8.73E-57 | Stroma 3 | b2m |
| rasl121 | 3.2E-61 | 0.7113978 | 0.144 | 0.055 | 1.04E-56 | Stroma 3 | rasl12 |
| col5a2a2 | 3.3E-61 | 0.4996265 | 0.877 | 0.788 | 1.08E-56 | Stroma 3 | col5a2a |
| lepb | 1E-60 | 2.0551831 | 0.102 | 0.033 | 3.35E-56 | Stroma 3 | lepb |
| adam15 | 3.2E-60 | 1.6272455 | 0.136 | 0.052 | 1.04E-55 | Stroma 3 | adam15 |
| mknk2b1 | 5.2E-60 | 0.79787 | 0.699 | 0.604 | 1.69E-55 | Stroma 3 | mknk2b |
| olfml2bb1 | 6.7E-60 | 1.1539423 | 0.191 | 0.087 | 2.17E-55 | Stroma 3 | olfml2bb |
| ppap2d3 | 6.9E-60 | 0.9851384 | 0.365 | 0.223 | 2.24E-55 | Stroma 3 | ppap2d |
| CR848833.12 | 1.6E-59 | 1.6780314 | 0.161 | 0.068 | 5.14E-55 | Stroma 3 | CR848833.1 |

| | | | | | | | |
|---|---|---|---|---|---|---|---|
| cxcl8b.1 | 2.9E-59 | 2.2588847 | 0.104 | 0.034 | 9.57E-55 | Stroma 3 | cxcl8b.1 |
| krt154 | 3E-59 | 0.4396264 | 0.825 | 0.695 | 9.7E-55 | Stroma 3 | krt15 |
| edil3a3 | 3.3E-59 | 0.8440687 | 0.566 | 0.429 | 1.08E-54 | Stroma 3 | edil3a |
| angptl2a3 | 3.1E-58 | 0.877624 | 0.459 | 0.312 | 1.02E-53 | Stroma 3 | angptl2a |
| plxdc22 | 1.1E-57 | 1.1831799 | 0.276 | 0.153 | 3.7E-53 | Stroma 3 | plxdc2 |
| si:ch211-195o20.7 | 1.2E-57 | 1.7405429 | 0.106 | 0.036 | 3.98E-53 | Stroma 3 | si:ch211-195o20.7 |
| egfra1 | 5E-57 | 1.066218 | 0.387 | 0.253 | 1.63E-52 | Stroma 3 | egfra |
| il15l3 | 4.1E-56 | 1.3256337 | 0.223 | 0.114 | 1.32E-51 | Stroma 3 | il15l |
| sulf2b3 | 1.2E-55 | 1.2114472 | 0.283 | 0.16 | 4.04E-51 | Stroma 3 | sulf2b |
| im:71523482 | 2.4E-55 | 0.9199644 | 0.367 | 0.23 | 7.67E-51 | Stroma 3 | im:7152348 |
| abi3bpa4 | 8.2E-55 | 0.3382416 | 0.462 | 0.292 | 2.67E-50 | Stroma 3 | abi3bpa |
| fndc13 | 1.2E-54 | 1.0109639 | 0.288 | 0.162 | 3.82E-50 | Stroma 3 | fndc1 |
| si:dkey-16l2.201 | 7.8E-54 | 1.5784497 | 0.143 | 0.06 | 2.55E-49 | Stroma 3 | si:dkey-16l2.20 |
| svild1 | 4.1E-53 | 1.0787735 | 0.378 | 0.251 | 1.34E-48 | Stroma 3 | svild |
| nhsl23 | 4.8E-53 | 1.3025525 | 0.2 | 0.098 | 1.55E-48 | Stroma 3 | nhsl2 |
| lsp1b4 | 3.6E-52 | 0.9600296 | 0.388 | 0.252 | 1.17E-47 | Stroma 3 | lsp1b |
| ddit32 | 4.4E-52 | 0.6574872 | 0.682 | 0.578 | 1.44E-47 | Stroma 3 | ddit3 |
| ston2 | 8.1E-52 | 1.9879025 | 0.1 | 0.035 | 2.64E-47 | Stroma 3 | ston2 |
| spon2b3 | 2.8E-51 | 1.7868197 | 0.228 | 0.123 | 9.16E-47 | Stroma 3 | spon2b |
| ier2b1 | 4.4E-51 | 0.7985485 | 0.741 | 0.646 | 1.42E-46 | Stroma 3 | ier2b |
| zgc:1752141 | 8.9E-51 | 1.6013193 | 0.201 | 0.104 | 2.89E-46 | Stroma 3 | zgc:175214 |
| crhbp1 | 7.6E-50 | 1.6613177 | 0.102 | 0.037 | 2.46E-45 | Stroma 3 | crhbp |
| plekho23 | 8E-50 | 1.1776144 | 0.295 | 0.178 | 2.61E-45 | Stroma 3 | plekho2 |
| glipr2l1 | 6.8E-49 | 1.6631634 | 0.102 | 0.037 | 2.22E-44 | Stroma 3 | glipr2l |
| pdgfra3 | 8.7E-49 | 0.6418649 | 0.584 | 0.455 | 2.84E-44 | Stroma 3 | pdgfra |
| stard9 | 2E-48 | 1.6809843 | 0.119 | 0.048 | 6.51E-44 | Stroma 3 | stard9 |
| igfbp5b3 | 4.8E-48 | 0.757422 | 0.679 | 0.567 | 1.56E-43 | Stroma 3 | igfbp5b |
| rbp44 | 3E-47 | 0.3125262 | 0.969 | 0.907 | 9.76E-43 | Stroma 3 | rbp4 |
| tuba1a1 | 3.7E-47 | 1.3255192 | 0.176 | 0.085 | 1.21E-42 | Stroma 3 | tuba1a |
| spon1b1 | 5.5E-47 | 2.0552903 | 0.127 | 0.054 | 1.79E-42 | Stroma 3 | spon1b |
| pak11 | 1.8E-46 | 1.5503924 | 0.144 | 0.065 | 5.78E-42 | Stroma 3 | pak1 |
| sesn13 | 2.6E-46 | 1.1442774 | 0.328 | 0.213 | 8.61E-42 | Stroma 3 | sesn1 |
| dusp8a3 | 3.7E-46 | 1.3689764 | 0.175 | 0.086 | 1.21E-41 | Stroma 3 | dusp8a |
| ddr14 | 6.5E-46 | 1.3651515 | 0.171 | 0.083 | 2.11E-41 | Stroma 3 | ddr1 |
| rpz53 | 1.5E-45 | 0.9484673 | 0.424 | 0.302 | 4.99E-41 | Stroma 3 | rpz5 |
| fzd4 | 1.9E-45 | 1.7714745 | 0.12 | 0.05 | 6.14E-41 | Stroma 3 | fzd4 |
| tnfsf10l4 | 2.3E-45 | 0.9841211 | 0.337 | 0.217 | 7.33E-41 | Stroma 3 | tnfsf10l |
| atf31 | 4.2E-45 | 0.9005344 | 0.464 | 0.341 | 1.36E-40 | Stroma 3 | atf3 |
| sfrp53 | 5E-45 | 1.1563444 | 0.321 | 0.205 | 1.62E-40 | Stroma 3 | sfrp5 |
| junbb1 | 5.6E-45 | 0.5654138 | 0.698 | 0.597 | 1.81E-40 | Stroma 3 | junbb |
| tgfb53 | 1.2E-44 | 1.3938663 | 0.204 | 0.11 | 3.81E-40 | Stroma 3 | tgfb5 |
| svila4 | 2.1E-44 | 1.0954594 | 0.25 | 0.144 | 6.71E-40 | Stroma 3 | svila |
| tsc22d31 | 3.7E-44 | 0.8615481 | 0.507 | 0.398 | 1.22E-39 | Stroma 3 | tsc22d3 |
| ZFP36L1.12 | 5.2E-44 | 0.4878272 | 0.922 | 0.871 | 1.71E-39 | Stroma 3 | ZFP36L1.1 |
| ptprk2 | 6.7E-44 | 1.4797863 | 0.144 | 0.066 | 2.18E-39 | Stroma 3 | ptprk |
| rab32a | 7.4E-44 | 1.6907919 | 0.102 | 0.04 | 2.41E-39 | Stroma 3 | rab32a |
| fgf18a1 | 8.1E-44 | 1.3723874 | 0.106 | 0.042 | 2.64E-39 | Stroma 3 | fgf18a |
| lrp53 | 1.4E-43 | 1.2992868 | 0.203 | 0.108 | 4.65E-39 | Stroma 3 | lrp5 |

| | | | | | | | |
|---|---|---|---|---|---|---|---|
| col5a13 | 2E-43 | 0.3873677 | 0.977 | 0.964 | 6.42E-39 | Stroma 3 | col5a1 |
| dub | 3.4E-43 | 1.3727679 | 0.102 | 0.039 | 1.09E-38 | Stroma 3 | dub |
| col6a13 | 1.1E-42 | 0.4620347 | 0.904 | 0.874 | 3.63E-38 | Stroma 3 | col6a1 |
| meox12 | 2.5E-42 | 1.0143319 | 0.136 | 0.06 | 8.06E-38 | Stroma 3 | meox1 |
| osr12 | 7.4E-42 | 1.0065506 | 0.275 | 0.168 | 2.41E-37 | Stroma 3 | osr1 |
| tenm33 | 1.5E-41 | 1.0613051 | 0.312 | 0.203 | 4.73E-37 | Stroma 3 | tenm3 |
| anxa2a2 | 3.9E-41 | 0.3108976 | 0.93 | 0.889 | 1.26E-36 | Stroma 3 | anxa2a |
| CU914487.11 | 1.4E-40 | 1.1507322 | 0.127 | 0.056 | 4.58E-36 | Stroma 3 | CU914487.1 |
| itgam | 1.7E-40 | 1.5202194 | 0.1 | 0.04 | 5.53E-36 | Stroma 3 | itgam |
| serpinb1 | 6.1E-40 | 1.94895 | 0.136 | 0.064 | 2E-35 | Stroma 3 | serpinb1 |
| ubb1 | 2.4E-39 | 0.5976553 | 0.745 | 0.678 | 7.72E-35 | Stroma 3 | ubb |
| selenop2 | 3.8E-39 | 0.4729451 | 0.809 | 0.738 | 1.22E-34 | Stroma 3 | selenop |
| si:dkey-192g7.31 | 7.6E-39 | 1.4470839 | 0.121 | 0.053 | 2.46E-34 | Stroma 3 | si:dkey-192g7.3 |
| TCIM | 8.8E-39 | 1.2949793 | 0.135 | 0.063 | 2.87E-34 | Stroma 3 | TCIM |
| slco2b14 | 7.1E-38 | 0.9518224 | 0.325 | 0.217 | 2.3E-33 | Stroma 3 | slco2b1 |
| nfkbiab2 | 8.4E-38 | 0.5788523 | 0.665 | 0.58 | 2.74E-33 | Stroma 3 | nfkbiab |
| bahcc1a2 | 1.2E-37 | 1.5291954 | 0.162 | 0.084 | 3.89E-33 | Stroma 3 | bahcc1a |
| si:ch211-250c4.42 | 1.4E-36 | 1.516938 | 0.101 | 0.042 | 4.43E-32 | Stroma 3 | si:ch211-250c4.4 |
| klf6a1 | 4.7E-36 | 0.4795057 | 0.805 | 0.744 | 1.54E-31 | Stroma 3 | klf6a |
| si:ch73-173h19.34 | 6.6E-36 | 0.6894728 | 0.35 | 0.234 | 2.15E-31 | Stroma 3 | si:ch73-173h19.3 |
| si:ch73-335l21.41 | 1.2E-35 | 0.5691259 | 0.667 | 0.566 | 3.91E-31 | Stroma 3 | si:ch73-335l21.4 |
| mxra8a4 | 1.4E-35 | 0.8127143 | 0.354 | 0.25 | 4.4E-31 | Stroma 3 | mxra8a |
| nfixb3 | 4.3E-35 | 0.7705727 | 0.393 | 0.29 | 1.38E-30 | Stroma 3 | nfixb |
| si:dkey-27i16.22 | 6.9E-35 | 0.8500528 | 0.202 | 0.114 | 2.26E-30 | Stroma 3 | si:dkey-27i16.2 |
| sash1a2 | 1.2E-34 | 1.0299033 | 0.302 | 0.206 | 3.77E-30 | Stroma 3 | sash1a |
| cebpa3 | 2.3E-34 | 1.1709607 | 0.267 | 0.172 | 7.33E-30 | Stroma 3 | cebpa |
| lsp1a2 | 4.3E-34 | 0.9975119 | 0.303 | 0.205 | 1.4E-29 | Stroma 3 | lsp1a |
| camk1b1 | 6.2E-34 | 1.3306009 | 0.12 | 0.056 | 2.02E-29 | Stroma 3 | camk1b |
| dnajb1b1 | 6.8E-34 | 0.7302312 | 0.546 | 0.461 | 2.2E-29 | Stroma 3 | dnajb1b |
| tspan141 | 8.7E-34 | 1.2079252 | 0.202 | 0.118 | 2.83E-29 | Stroma 3 | tspan14 |
| srgn2 | 1.8E-33 | 0.6501028 | 0.543 | 0.448 | 5.74E-29 | Stroma 3 | srgn |
| arhgef1a | 1.8E-33 | 1.4316603 | 0.122 | 0.058 | 5.92E-29 | Stroma 3 | arhgef1a |
| lipf1 | 2E-33 | 1.0722197 | 0.291 | 0.194 | 6.57E-29 | Stroma 3 | lipf |
| arhgef10lb | 2.7E-33 | 1.4210621 | 0.119 | 0.056 | 8.75E-29 | Stroma 3 | arhgef10lb |
| si:ch1073-291c23.23 | 3.3E-33 | 0.7036908 | 0.559 | 0.471 | 1.06E-28 | Stroma 3 | si:ch1073-291c23.2 |
| tns1b2 | 4.2E-33 | 0.6174762 | 0.464 | 0.364 | 1.38E-28 | Stroma 3 | tns1b |
| niban1b1 | 1.3E-32 | 1.3052495 | 0.167 | 0.092 | 4.28E-28 | Stroma 3 | niban1b |
| sema3fa2 | 1E-31 | 1.0702888 | 0.125 | 0.061 | 3.32E-27 | Stroma 3 | sema3fa |
| crabp2b2 | 1E-31 | 0.66896 | 0.513 | 0.416 | 3.34E-27 | Stroma 3 | crabp2b |
| mhc1zba2 | 1.8E-31 | 0.4933725 | 0.646 | 0.582 | 5.9E-27 | Stroma 3 | mhc1zba |
| her61 | 2.5E-31 | 0.6706013 | 0.522 | 0.429 | 8.27E-27 | Stroma 3 | her6 |
| mfap24 | 3.3E-31 | 0.5992227 | 0.481 | 0.38 | 1.08E-26 | Stroma 3 | mfap2 |
| grk42 | 4.7E-31 | 0.9474391 | 0.267 | 0.176 | 1.52E-26 | Stroma 3 | grk4 |
| mych2 | 5.4E-31 | 0.7623468 | 0.47 | 0.376 | 1.74E-26 | Stroma 3 | mych |
| pdgfrb2 | 9.9E-31 | 0.7856417 | 0.489 | 0.409 | 3.21E-26 | Stroma 3 | pdgfrb |
| ier5l1 | 1.4E-30 | 0.8758603 | 0.408 | 0.316 | 4.6E-26 | Stroma 3 | ier5l |
| colec123 | 1.6E-30 | 0.6232011 | 0.529 | 0.435 | 5.05E-26 | Stroma 3 | colec12 |
| adam91 | 1.9E-30 | 1.2569217 | 0.128 | 0.065 | 6.09E-26 | Stroma 3 | adam9 |

| | | | | | | | |
|---|---|---|---|---|---|---|---|
| rasgef1bb2 | 2E-30 | 0.9962796 | 0.237 | 0.152 | 6.47E-26 | Stroma 3 | rasgef1bb |
| lpin2 | 2.5E-30 | 1.4689199 | 0.109 | 0.052 | 8.1E-26 | Stroma 3 | lpin2 |
| socs3b2 | 3.9E-30 | 0.8389871 | 0.541 | 0.469 | 1.28E-25 | Stroma 3 | socs3b |
| cgnl12 | 4.8E-30 | 1.2246303 | 0.135 | 0.07 | 1.56E-25 | Stroma 3 | cgnl1 |
| stom2 | 5.2E-30 | 0.7604782 | 0.36 | 0.266 | 1.69E-25 | Stroma 3 | stom |
| arhgap29b1 | 1.3E-29 | 1.4638585 | 0.113 | 0.055 | 4.19E-25 | Stroma 3 | arhgap29b |
| dipk1c4 | 2.7E-29 | 1.1707388 | 0.187 | 0.112 | 8.72E-25 | Stroma 3 | dipk1c |
| tjp1a1 | 2.9E-29 | 0.9976975 | 0.235 | 0.151 | 9.51E-25 | Stroma 3 | tjp1a |
| zgc:1951734 | 5.6E-29 | 0.6967216 | 0.483 | 0.394 | 1.81E-24 | Stroma 3 | zgc:195173 |
| CR936442.1 | 9.4E-29 | 1.5533843 | 0.271 | 0.188 | 3.06E-24 | Stroma 3 | CR936442.1 |
| ENSDARG00000908 | 1.2E-28 | 1.1320924 | 0.157 | 0.088 | 3.82E-24 | Stroma 3 | ENSDARG00000090 |
| btg21 | 1.2E-28 | 0.3424115 | 0.779 | 0.696 | 3.9E-24 | Stroma 3 | btg2 |
| gas1a3 | 1.4E-28 | 0.8314353 | 0.244 | 0.157 | 4.68E-24 | Stroma 3 | gas1a |
| bcl2a1 | 1.6E-28 | 0.9462741 | 0.235 | 0.152 | 5.19E-24 | Stroma 3 | bcl2a |
| si:ch73-380l3.24 | 1.7E-28 | 0.6280524 | 0.504 | 0.413 | 5.44E-24 | Stroma 3 | si:ch73-380l3.2 |
| enc13 | 2.3E-28 | 0.9040575 | 0.201 | 0.121 | 7.34E-24 | Stroma 3 | enc1 |
| lifrb1 | 2.9E-28 | 0.7891467 | 0.286 | 0.195 | 9.33E-24 | Stroma 3 | lifrb |
| si:dkey-11f4.204 | 5.2E-28 | 0.6882012 | 0.556 | 0.486 | 1.69E-23 | Stroma 3 | si:dkey-11f4.20 |
| vamp52 | 5.4E-28 | 1.3463252 | 0.143 | 0.078 | 1.75E-23 | Stroma 3 | vamp5 |
| rab101 | 6.3E-28 | 0.5675351 | 0.528 | 0.455 | 2.05E-23 | Stroma 3 | rab10 |
| cfd1 | 8.3E-28 | 2.1034233 | 0.165 | 0.097 | 2.69E-23 | Stroma 3 | cfd |
| C25H11orf962 | 9.2E-28 | 0.6505537 | 0.475 | 0.38 | 2.98E-23 | Stroma 3 | C25H11orf96 |
| igf2b3 | 1.2E-27 | 0.5761542 | 0.529 | 0.433 | 3.78E-23 | Stroma 3 | igf2b |
| efnb2a | 1.2E-27 | 1.0282308 | 0.227 | 0.147 | 3.95E-23 | Stroma 3 | efnb2a |
| vmp12 | 1.6E-27 | 0.5598618 | 0.409 | 0.315 | 5.1E-23 | Stroma 3 | vmp1 |
| itm2ba2 | 1.9E-27 | 0.3916984 | 0.766 | 0.741 | 6.3E-23 | Stroma 3 | itm2ba |
| dcbld23 | 2.9E-27 | 0.8045252 | 0.201 | 0.122 | 9.36E-23 | Stroma 3 | dcbld2 |
| CR318588.42 | 6.6E-27 | 0.3430342 | 0.992 | 0.986 | 2.14E-22 | Stroma 3 | CR318588.4 |
| scara54 | 8.4E-27 | 0.7828901 | 0.232 | 0.148 | 2.74E-22 | Stroma 3 | scara5 |
| loxl2b2 | 1.5E-26 | 0.7575555 | 0.397 | 0.314 | 4.81E-22 | Stroma 3 | loxl2b |
| znf385a1 | 1.6E-26 | 1.4875392 | 0.102 | 0.049 | 5.08E-22 | Stroma 3 | znf385a |
| col6a23 | 2.1E-26 | 0.3677027 | 0.901 | 0.876 | 6.97E-22 | Stroma 3 | col6a2 |
| gadd45bb2 | 2.5E-26 | 1.0040283 | 0.303 | 0.217 | 8.23E-22 | Stroma 3 | gadd45bb |
| ifi462 | 3E-26 | 1.1310365 | 0.16 | 0.093 | 9.9E-22 | Stroma 3 | ifi46 |
| pcolce2b2 | 4E-26 | 0.6268048 | 0.385 | 0.293 | 1.31E-21 | Stroma 3 | pcolce2b |
| adma3 | 4.3E-26 | 0.7276369 | 0.322 | 0.23 | 1.4E-21 | Stroma 3 | adma |
| loxa4 | 5.8E-26 | 0.6781827 | 0.257 | 0.171 | 1.89E-21 | Stroma 3 | loxa |
| fbln52 | 8.7E-26 | 1.1743132 | 0.145 | 0.081 | 2.83E-21 | Stroma 3 | fbln5 |
| pfkfb31 | 1.1E-25 | 1.3701663 | 0.135 | 0.074 | 3.49E-21 | Stroma 3 | pfkfb3 |
| ywhag14 | 3.9E-25 | 0.9514234 | 0.179 | 0.108 | 1.26E-20 | Stroma 3 | ywhag1 |
| nid2a | 4.4E-25 | 1.3172346 | 0.13 | 0.071 | 1.42E-20 | Stroma 3 | nid2a |
| tcima4 | 8.8E-25 | 1.3148709 | 0.303 | 0.224 | 2.87E-20 | Stroma 3 | tcima |
| si:dkey-172h23.22 | 8.9E-25 | 0.8959545 | 0.22 | 0.144 | 2.89E-20 | Stroma 3 | si:dkey-172h23.2 |
| pde3b2 | 8.9E-25 | 1.0434125 | 0.189 | 0.118 | 2.89E-20 | Stroma 3 | pde3b |
| rfx22 | 9.2E-25 | 1.0642242 | 0.127 | 0.068 | 3E-20 | Stroma 3 | rfx2 |
| rgmb2 | 1.1E-24 | 1.1425304 | 0.186 | 0.117 | 3.67E-20 | Stroma 3 | rgmb |
| lrig1 | 1.4E-24 | 1.0808265 | 0.134 | 0.074 | 4.5E-20 | Stroma 3 | lrig1 |
| si:ch211-156j16.12 | 1.9E-24 | 0.6577172 | 0.23 | 0.149 | 6.11E-20 | Stroma 3 | si:ch211-156j16.1 |

| | | | | | | | |
|---|---|---|---|---|---|---|---|
| qki2 | 2E-24 | 1.2320612 | 0.126 | 0.069 | 6.57E-20 | Stroma 3 | qki2 |
| ponzr13 | 3E-24 | 0.5315521 | 0.323 | 0.23 | 9.62E-20 | Stroma 3 | ponzr1 |
| dennd4a2 | 4.1E-24 | 1.1530383 | 0.145 | 0.084 | 1.33E-19 | Stroma 3 | dennd4a |
| rasl11b3 | 4.7E-24 | 0.9790315 | 0.292 | 0.214 | 1.53E-19 | Stroma 3 | rasl11b |
| cnn3b | 5.1E-24 | 1.401469 | 0.103 | 0.053 | 1.66E-19 | Stroma 3 | cnn3b |
| foxo3b2 | 7.7E-24 | 1.2884513 | 0.139 | 0.08 | 2.5E-19 | Stroma 3 | foxo3b |
| macf1a1 | 9.8E-24 | 0.9278342 | 0.249 | 0.173 | 3.18E-19 | Stroma 3 | macf1a |
| p4ha1b3 | 1.5E-23 | 0.9648418 | 0.21 | 0.138 | 4.88E-19 | Stroma 3 | p4ha1b |
| lrp1ab3 | 1.9E-23 | 0.6597761 | 0.417 | 0.338 | 6.06E-19 | Stroma 3 | lrp1ab |
| cd632 | 2.8E-23 | 0.2506568 | 0.843 | 0.827 | 9.1E-19 | Stroma 3 | cd63 |
| qkia2 | 3.1E-23 | 0.6288121 | 0.524 | 0.466 | 1.01E-18 | Stroma 3 | qkia |
| raph1b1 | 6.7E-23 | 1.0713577 | 0.168 | 0.104 | 2.17E-18 | Stroma 3 | raph1b |
| il341 | 7.4E-23 | 1.149247 | 0.1 | 0.051 | 2.41E-18 | Stroma 3 | il34 |
| ms4a17a.112 | 7.8E-23 | 0.8970587 | 0.324 | 0.249 | 2.55E-18 | Stroma 3 | ms4a17a.11 |
| klf7a | 8.4E-23 | 1.4343919 | 0.105 | 0.056 | 2.73E-18 | Stroma 3 | klf7a |
| si:cabz01069012.23 | 9E-23 | 0.9861166 | 0.165 | 0.1 | 2.94E-18 | Stroma 3 | si:cabz01069012.2 |
| il6st2 | 1.2E-22 | 0.5772815 | 0.495 | 0.421 | 3.89E-18 | Stroma 3 | il6st |
| ppp1r101 | 2E-22 | 0.4628639 | 0.557 | 0.482 | 6.41E-18 | Stroma 3 | ppp1r10 |
| baz2ba1 | 3E-22 | 0.8847944 | 0.24 | 0.167 | 9.64E-18 | Stroma 3 | baz2ba |
| bhlhe402 | 6.3E-22 | 0.6401721 | 0.526 | 0.463 | 2.05E-17 | Stroma 3 | bhlhe40 |
| fzd7a | 2.1E-21 | 1.2356039 | 0.133 | 0.077 | 6.78E-17 | Stroma 3 | fzd7a |
| tmod21 | 2.7E-21 | 0.8812166 | 0.24 | 0.169 | 8.78E-17 | Stroma 3 | tmod2 |
| peli1b1 | 3E-21 | 1.2440133 | 0.145 | 0.088 | 9.72E-17 | Stroma 3 | peli1b |
| txnipa | 5.6E-21 | 0.7793786 | 0.317 | 0.242 | 1.81E-16 | Stroma 3 | txnipa |
| si:ch211-165e15.11 | 6.9E-21 | 1.263135 | 0.116 | 0.065 | 2.23E-16 | Stroma 3 | si:ch211-165e15.1 |
| eml1 | 7.3E-21 | 1.3372582 | 0.101 | 0.054 | 2.38E-16 | Stroma 3 | eml1 |
| podn3 | 8.2E-21 | 0.839749 | 0.242 | 0.169 | 2.65E-16 | Stroma 3 | podn |
| si:rp71-36a1.51 | 8.9E-21 | 0.9578607 | 0.143 | 0.085 | 2.9E-16 | Stroma 3 | si:rp71-36a1.5 |
| txn3 | 9.8E-21 | 0.2770485 | 0.466 | 0.369 | 3.2E-16 | Stroma 3 | txn |
| gulp1a2 | 1.8E-20 | 0.833978 | 0.256 | 0.185 | 6E-16 | Stroma 3 | gulp1a |
| sept9b | 3E-20 | 1.2071659 | 0.109 | 0.06 | 9.79E-16 | Stroma 3 | sept9b |
| gstt1b3 | 3.4E-20 | 0.8993878 | 0.184 | 0.12 | 1.1E-15 | Stroma 3 | gstt1b |
| afdna | 4E-20 | 0.8594772 | 0.262 | 0.191 | 1.29E-15 | Stroma 3 | afdna |
| si:ch211-284f22.32 | 4.2E-20 | 0.9992594 | 0.177 | 0.115 | 1.36E-15 | Stroma 3 | si:ch211-284f22.3 |
| lepr1 | 6E-20 | 1.2058773 | 0.111 | 0.062 | 1.96E-15 | Stroma 3 | lepr |
| arrdc23 | 9.6E-20 | 0.6472692 | 0.318 | 0.24 | 3.13E-15 | Stroma 3 | arrdc2 |
| arhgap42a1 | 1.1E-19 | 1.1648388 | 0.122 | 0.071 | 3.5E-15 | Stroma 3 | arhgap42a |
| twist1b2 | 1.1E-19 | 0.8695999 | 0.225 | 0.157 | 3.54E-15 | Stroma 3 | twist1b |
| mycb1 | 1.2E-19 | 0.3004743 | 0.726 | 0.661 | 3.78E-15 | Stroma 3 | mycb |
| selenbp12 | 2.4E-19 | 1.1218888 | 0.128 | 0.076 | 7.85E-15 | Stroma 3 | selenbp1 |
| rsrp11 | 2.6E-19 | 0.3152739 | 0.611 | 0.54 | 8.32E-15 | Stroma 3 | rsrp1 |
| reep3a1 | 2.6E-19 | 0.7913052 | 0.23 | 0.163 | 8.51E-15 | Stroma 3 | reep3a |
| rnd3a4 | 3.1E-19 | 0.5957716 | 0.36 | 0.283 | 9.95E-15 | Stroma 3 | rnd3a |
| id12 | 3.7E-19 | 0.4580916 | 0.689 | 0.64 | 1.19E-14 | Stroma 3 | id1 |
| sema4ba1 | 5.2E-19 | 1.1686894 | 0.136 | 0.082 | 1.69E-14 | Stroma 3 | sema4ba |
| col6a4a1 | 9.7E-19 | 1.033719 | 0.101 | 0.055 | 3.15E-14 | Stroma 3 | col6a4a |
| zgc:1527914 | 1.2E-18 | 0.9205439 | 0.163 | 0.103 | 3.97E-14 | Stroma 3 | zgc:152791 |
| jpt1b1 | 1.2E-18 | 0.3076035 | 0.718 | 0.691 | 4.06E-14 | Stroma 3 | jpt1b |

| | | | | | | | |
|---|---|---|---|---|---|---|---|
| entpd14 | 1.8E-18 | 0.9250159 | 0.133 | 0.08 | 6.02E-14 | Stroma 3 | entpd1 |
| dab2ipb2 | 2.3E-18 | 0.9978533 | 0.169 | 0.111 | 7.42E-14 | Stroma 3 | dab2ipb |
| kank32 | 2.8E-18 | 1.0080334 | 0.13 | 0.078 | 9.25E-14 | Stroma 3 | kank3 |
| eva1ba1 | 3.1E-18 | 1.1350686 | 0.162 | 0.106 | 1.01E-13 | Stroma 3 | eva1ba |
| pde7a1 | 3.7E-18 | 1.1491532 | 0.143 | 0.09 | 1.21E-13 | Stroma 3 | pde7a |
| amotl2a1 | 5.2E-18 | 0.9229721 | 0.217 | 0.155 | 1.68E-13 | Stroma 3 | amotl2a |
| sik2b2 | 5.5E-18 | 0.7900821 | 0.271 | 0.205 | 1.78E-13 | Stroma 3 | sik2b |
| rgs42 | 1.1E-17 | 0.8873316 | 0.118 | 0.068 | 3.45E-13 | Stroma 3 | rgs4 |
| gna12a | 1.1E-17 | 1.1442436 | 0.122 | 0.074 | 3.6E-13 | Stroma 3 | gna12a |
| fpr11 | 1.1E-17 | 0.9524893 | 0.278 | 0.216 | 3.64E-13 | Stroma 3 | fpr1 |
| furina1 | 1.2E-17 | 0.8909509 | 0.215 | 0.154 | 3.93E-13 | Stroma 3 | furina |
| ppp1r15a1 | 1.3E-17 | 0.7929726 | 0.272 | 0.207 | 4.1E-13 | Stroma 3 | ppp1r15a |
| si:ch73-44m9.3.11 | 1.4E-17 | 0.9430317 | 0.152 | 0.097 | 4.43E-13 | Stroma 3 | si:ch73-44m9.3.1 |
| tmem176l.1 | 1.5E-17 | 0.3085498 | 0.758 | 0.741 | 4.82E-13 | Stroma 3 | tmem176l.1 |
| klf31 | 2.3E-17 | 1.0350816 | 0.153 | 0.099 | 7.59E-13 | Stroma 3 | klf3 |
| zgc:1539111 | 2.6E-17 | 1.2120539 | 0.115 | 0.067 | 8.59E-13 | Stroma 3 | zgc:153911 |
| slc25a25a2 | 4.3E-17 | 0.7390497 | 0.282 | 0.217 | 1.4E-12 | Stroma 3 | slc25a25a |
| zgc:1750882 | 4.5E-17 | 0.6737204 | 0.334 | 0.272 | 1.48E-12 | Stroma 3 | zgc:175088 |
| pros11 | 5.1E-17 | 0.9954304 | 0.169 | 0.114 | 1.67E-12 | Stroma 3 | pros1 |
| plod1a | 6.4E-17 | 0.7635582 | 0.303 | 0.241 | 2.09E-12 | Stroma 3 | plod1a |
| emilin2a2 | 8.3E-17 | 0.7443021 | 0.177 | 0.118 | 2.69E-12 | Stroma 3 | emilin2a |
| soul52 | 8.8E-17 | 0.6743993 | 0.338 | 0.275 | 2.87E-12 | Stroma 3 | soul5 |
| rras2 | 9.5E-17 | 1.0758986 | 0.112 | 0.066 | 3.1E-12 | Stroma 3 | rras2 |
| otud1 | 1.1E-16 | 1.5213516 | 0.1 | 0.057 | 3.69E-12 | Stroma 3 | otud1 |
| rnd3b3 | 1.4E-16 | 0.5653161 | 0.395 | 0.329 | 4.57E-12 | Stroma 3 | rnd3b |
| adam19a2 | 1.9E-16 | 1.1054841 | 0.131 | 0.082 | 6.05E-12 | Stroma 3 | adam19a |
| tspan352 | 2.3E-16 | 0.8246028 | 0.179 | 0.122 | 7.55E-12 | Stroma 3 | tspan35 |
| arhgap35a2 | 2.6E-16 | 0.707815 | 0.241 | 0.178 | 8.32E-12 | Stroma 3 | arhgap35a |
| apcdd1l3 | 3.8E-16 | 0.69325 | 0.165 | 0.108 | 1.24E-11 | Stroma 3 | apcdd1l |
| anxa3b | 4E-16 | 0.6875008 | 0.308 | 0.246 | 1.31E-11 | Stroma 3 | anxa3b |
| bmp1a1 | 5.4E-16 | 0.8387823 | 0.244 | 0.185 | 1.74E-11 | Stroma 3 | bmp1a |
| tbx3a1 | 5.7E-16 | 0.6382109 | 0.182 | 0.122 | 1.86E-11 | Stroma 3 | tbx3a |
| ssh2a1 | 5.9E-16 | 1.0273248 | 0.126 | 0.078 | 1.92E-11 | Stroma 3 | ssh2a |
| adamtsl32 | 6.1E-16 | 1.0100115 | 0.113 | 0.068 | 2E-11 | Stroma 3 | adamtsl3 |
| fam43a2 | 7.2E-16 | 1.0273037 | 0.112 | 0.067 | 2.35E-11 | Stroma 3 | fam43a |
| atf5b1 | 7.4E-16 | 0.6472799 | 0.313 | 0.251 | 2.39E-11 | Stroma 3 | atf5b |
| s100v21 | 8.5E-16 | 0.3362791 | 0.696 | 0.679 | 2.76E-11 | Stroma 3 | s100v2 |
| gstt1a2 | 9E-16 | 1.0646362 | 0.12 | 0.074 | 2.92E-11 | Stroma 3 | gstt1a |
| ccl25b4 | 1.2E-15 | 0.4766084 | 0.761 | 0.66 | 4.06E-11 | Stroma 3 | ccl25b |
| mmp14a3 | 2.4E-15 | 0.3975286 | 0.669 | 0.647 | 7.83E-11 | Stroma 3 | mmp14a |
| adamts15b.12 | 3E-15 | 0.8736634 | 0.142 | 0.091 | 9.81E-11 | Stroma 3 | adamts15b.1 |
| cadm42 | 3.2E-15 | 0.8678278 | 0.121 | 0.074 | 1.03E-10 | Stroma 3 | cadm4 |
| nr2f22 | 4E-15 | 0.8918472 | 0.167 | 0.113 | 1.29E-10 | Stroma 3 | nr2f2 |
| pdgfrl3 | 7.7E-15 | 0.2941576 | 0.623 | 0.584 | 2.5E-10 | Stroma 3 | pdgfrl |
| lhfpl2a3 | 8.6E-15 | 0.8936042 | 0.156 | 0.106 | 2.78E-10 | Stroma 3 | lhfpl2a |
| myoc2 | 1E-14 | 0.7579086 | 0.111 | 0.067 | 3.24E-10 | Stroma 3 | myoc |
| cdh22 | 1E-14 | 0.5062584 | 0.353 | 0.29 | 3.35E-10 | Stroma 3 | cdh2 |
| mmp15a3 | 1E-14 | 0.9236248 | 0.128 | 0.082 | 3.38E-10 | Stroma 3 | mmp15a |

| | | | | | | | |
|---|---|---|---|---|---|---|---|
| si:ch211-212k18.83 | 1.1E-14 | 0.7170495 | 0.197 | 0.14 | 3.57E-10 | Stroma 3 | si:ch211-212k18.8 |
| dedd13 | 1.1E-14 | 0.3612727 | 0.591 | 0.551 | 3.65E-10 | Stroma 3 | dedd1 |
| add11 | 1.2E-14 | 0.778154 | 0.191 | 0.136 | 3.74E-10 | Stroma 3 | add1 |
| sdcbp2 | 1.3E-14 | 0.2458814 | 0.642 | 0.597 | 4.39E-10 | Stroma 3 | sdcbp2 |
| cebpb1 | 2.3E-14 | 0.2995628 | 0.514 | 0.446 | 7.45E-10 | Stroma 3 | cebpb |
| hexb1 | 3.1E-14 | 0.9681723 | 0.136 | 0.09 | 1E-09 | Stroma 3 | hexb |
| pdia52 | 4E-14 | 0.7866529 | 0.219 | 0.164 | 1.29E-09 | Stroma 3 | pdia5 |
| fzd7b1 | 6.4E-14 | 0.8957995 | 0.157 | 0.108 | 2.07E-09 | Stroma 3 | fzd7b |
| efhd11 | 9.3E-14 | 1.2997809 | 0.104 | 0.064 | 3.01E-09 | Stroma 3 | efhd1 |
| mcl1b2 | 1E-13 | 0.4052543 | 0.529 | 0.481 | 3.26E-09 | Stroma 3 | mcl1b |
| arhgap21a1 | 1.1E-13 | 0.9769497 | 0.152 | 0.105 | 3.6E-09 | Stroma 3 | arhgap21a |
| hist2h2l2 | 1.1E-13 | 0.513305 | 0.429 | 0.373 | 3.71E-09 | Stroma 3 | hist2h2l |
| agrn1 | 1.2E-13 | 0.7419661 | 0.238 | 0.183 | 3.93E-09 | Stroma 3 | agrn |
| btg11 | 1.3E-13 | 0.2508775 | 0.831 | 0.814 | 4.26E-09 | Stroma 3 | btg1 |
| pdlim11 | 1.4E-13 | 0.6624543 | 0.285 | 0.228 | 4.62E-09 | Stroma 3 | pdlim1 |
| ezh13 | 1.6E-13 | 0.7626704 | 0.192 | 0.139 | 5.14E-09 | Stroma 3 | ezh1 |
| col18a1a1 | 1.7E-13 | 1.7859245 | 0.201 | 0.152 | 5.48E-09 | Stroma 3 | col18a1a |
| skia3 | 1.9E-13 | 0.4492696 | 0.498 | 0.448 | 6.07E-09 | Stroma 3 | skia |
| jund2 | 3.5E-13 | 0.3207463 | 0.618 | 0.568 | 1.13E-08 | Stroma 3 | jund |
| SBNO21 | 4.6E-13 | 0.6651589 | 0.281 | 0.226 | 1.5E-08 | Stroma 3 | SBNO2 |
| gpc41 | 4.6E-13 | 0.6283978 | 0.296 | 0.241 | 1.5E-08 | Stroma 3 | gpc4 |
| phc2b1 | 7.4E-13 | 1.1916973 | 0.115 | 0.074 | 2.41E-08 | Stroma 3 | phc2b |
| sat1a.22 | 8.2E-13 | 0.4195044 | 0.504 | 0.456 | 2.67E-08 | Stroma 3 | sat1a.2 |
| C7H4orf481 | 8.5E-13 | 0.522443 | 0.297 | 0.241 | 2.78E-08 | Stroma 3 | C7H4orf48 |
| abtb2b | 1.3E-12 | 0.9281971 | 0.114 | 0.073 | 4.38E-08 | Stroma 3 | abtb2b |
| aplp22 | 1.7E-12 | 0.4945917 | 0.371 | 0.322 | 5.42E-08 | Stroma 3 | aplp2 |
| boc3 | 1.9E-12 | 0.4565816 | 0.42 | 0.371 | 6.03E-08 | Stroma 3 | boc |
| zgc:1122341 | 2E-12 | 1.07827 | 0.113 | 0.073 | 6.48E-08 | Stroma 3 | zgc:112234 |
| rhbdf1a1 | 2.4E-12 | 0.8328624 | 0.137 | 0.093 | 7.72E-08 | Stroma 3 | rhbdf1a |
| ier2a2 | 2.8E-12 | 0.4619142 | 0.529 | 0.484 | 9.13E-08 | Stroma 3 | ier2a |
| aif1l2 | 3.2E-12 | 1.0714893 | 0.126 | 0.085 | 1.03E-07 | Stroma 3 | aif1l |
| serpinh21 | 3.4E-12 | 0.7224792 | 0.164 | 0.116 | 1.11E-07 | Stroma 3 | serpinh2 |
| thbs2a4 | 4.2E-12 | 0.7098553 | 0.254 | 0.202 | 1.35E-07 | Stroma 3 | thbs2a |
| sesn32 | 4.5E-12 | 0.7705982 | 0.153 | 0.108 | 1.46E-07 | Stroma 3 | sesn3 |
| emilin1b2 | 6.8E-12 | 0.7646212 | 0.127 | 0.085 | 2.23E-07 | Stroma 3 | emilin1b |
| irf1b2 | 8.7E-12 | 0.9448152 | 0.162 | 0.116 | 2.83E-07 | Stroma 3 | irf1b |
| zgc:1749042 | 9.7E-12 | 1.1735359 | 0.171 | 0.123 | 3.16E-07 | Stroma 3 | zgc:174904 |
| adamts51 | 1E-11 | 1.0129968 | 0.1 | 0.064 | 3.33E-07 | Stroma 3 | adamts5 |
| arl4ab3 | 1.5E-11 | 0.7896908 | 0.153 | 0.109 | 4.8E-07 | Stroma 3 | arl4ab |
| si:ch211-212k18.94 | 1.5E-11 | 0.6158911 | 0.227 | 0.176 | 4.86E-07 | Stroma 3 | si:ch211-212k18.9 |
| prss233 | 1.6E-11 | 0.6114831 | 0.251 | 0.199 | 5.29E-07 | Stroma 3 | prss23 |
| nabp1a1 | 1.6E-11 | 0.7175752 | 0.226 | 0.178 | 5.31E-07 | Stroma 3 | nabp1a |
| si:dkeyp-23e4.33 | 1.7E-11 | 0.715014 | 0.126 | 0.084 | 5.56E-07 | Stroma 3 | si:dkeyp-23e4.3 |
| sertad41 | 1.9E-11 | 0.6577667 | 0.245 | 0.193 | 6.08E-07 | Stroma 3 | sertad4 |
| nck1a2 | 2E-11 | 0.8208005 | 0.118 | 0.078 | 6.52E-07 | Stroma 3 | nck1a |
| ccn12 | 2E-11 | 0.2301914 | 0.687 | 0.641 | 6.66E-07 | Stroma 3 | ccn1 |
| pttg1ipb1 | 2.6E-11 | 0.9752406 | 0.102 | 0.066 | 8.4E-07 | Stroma 3 | pttg1ipb |
| fkbp51 | 3.1E-11 | 0.6766244 | 0.244 | 0.196 | 1.02E-06 | Stroma 3 | fkbp5 |

| | | | | | | | |
|---|---|---|---|---|---|---|---|
| tgfbi1 | 3.2E-11 | 0.4802279 | 0.255 | 0.198 | 1.05E-06 | Stroma 3 | tgfbi |
| ifi452 | 3.7E-11 | 1.0811584 | 0.276 | 0.227 | 1.2E-06 | Stroma 3 | ifi45 |
| si:ch1073-185p12.21 | 6E-11 | 0.8091619 | 0.121 | 0.082 | 1.94E-06 | Stroma 3 | si:ch1073-185p12.2 |
| npc11 | 6E-11 | 0.9226333 | 0.11 | 0.073 | 1.94E-06 | Stroma 3 | npc1 |
| fbxo343 | 7.6E-11 | 0.7941589 | 0.129 | 0.089 | 2.46E-06 | Stroma 3 | fbxo34 |
| glis2a2 | 7.8E-11 | 0.6199066 | 0.25 | 0.203 | 2.53E-06 | Stroma 3 | glis2a |
| sh3bp5a1 | 8.5E-11 | 0.7536343 | 0.174 | 0.129 | 2.77E-06 | Stroma 3 | sh3bp5a |
| zgc:1748881 | 9.6E-11 | 0.5107981 | 0.268 | 0.215 | 3.11E-06 | Stroma 3 | zgc:174888 |
| tril3 | 1E-10 | 0.5862685 | 0.149 | 0.105 | 3.37E-06 | Stroma 3 | tril |
| kank21 | 1.1E-10 | 0.5684315 | 0.26 | 0.211 | 3.71E-06 | Stroma 3 | kank2 |
| arhgap51 | 1.3E-10 | 0.5106622 | 0.392 | 0.35 | 4.16E-06 | Stroma 3 | arhgap5 |
| aoc23 | 1.5E-10 | 0.4048847 | 0.296 | 0.239 | 4.72E-06 | Stroma 3 | aoc2 |
| clasp2 | 1.7E-10 | 0.9120507 | 0.107 | 0.071 | 5.65E-06 | Stroma 3 | clasp2 |
| ATP11B1 | 2.1E-10 | 0.8373664 | 0.148 | 0.107 | 6.76E-06 | Stroma 3 | ATP11B |
| sox4a1 | 2.1E-10 | 0.5946625 | 0.241 | 0.193 | 6.89E-06 | Stroma 3 | sox4a |
| rbpjb2 | 2.1E-10 | 0.747923 | 0.136 | 0.096 | 6.89E-06 | Stroma 3 | rbpjb |
| cxxc5a | 2.2E-10 | 0.7153523 | 0.148 | 0.106 | 7.16E-06 | Stroma 3 | cxxc5a |
| hivep11 | 2.2E-10 | 0.5653036 | 0.201 | 0.152 | 7.16E-06 | Stroma 3 | hivep1 |
| sfrp1b4 | 2.2E-10 | 0.4299266 | 0.414 | 0.36 | 7.17E-06 | Stroma 3 | sfrp1b |
| si:ch73-46j18.53 | 2.5E-10 | 0.2238679 | 0.637 | 0.602 | 8.02E-06 | Stroma 3 | si:ch73-46j18.5 |
| CR383676.33 | 2.5E-10 | 0.3370739 | 0.508 | 0.465 | 8.25E-06 | Stroma 3 | CR383676.3 |
| ugp2b | 3.5E-10 | 0.8020833 | 0.158 | 0.116 | 1.13E-05 | Stroma 3 | ugp2b |
| CU984600.14 | 3.9E-10 | 0.6832145 | 0.214 | 0.169 | 1.26E-05 | Stroma 3 | CU984600.1 |
| ctsh1 | 4.4E-10 | 0.8179545 | 0.149 | 0.109 | 1.42E-05 | Stroma 3 | ctsh |
| kank43 | 4.8E-10 | 0.6385672 | 0.143 | 0.102 | 1.54E-05 | Stroma 3 | kank4 |
| si:ch1073-392o20.23 | 6.3E-10 | 0.5308006 | 0.288 | 0.241 | 2.06E-05 | Stroma 3 | si:ch1073-392o20.2 |
| nrarpa1 | 6.6E-10 | 0.8352636 | 0.125 | 0.087 | 2.16E-05 | Stroma 3 | nrarpa |
| ntd52 | 7.1E-10 | 0.7632235 | 0.186 | 0.141 | 2.32E-05 | Stroma 3 | ntd5 |
| tlr5b2 | 8E-10 | 0.893016 | 0.108 | 0.073 | 2.6E-05 | Stroma 3 | tlr5b |
| twist1a1 | 8.4E-10 | 0.4363084 | 0.478 | 0.437 | 2.72E-05 | Stroma 3 | twist1a |
| jak12 | 8.6E-10 | 0.3804367 | 0.474 | 0.437 | 2.79E-05 | Stroma 3 | jak1 |
| gamt | 9.7E-10 | 0.8740931 | 0.105 | 0.071 | 3.15E-05 | Stroma 3 | gamt |
| tuba1b3 | 9.8E-10 | 0.2951301 | 0.365 | 0.307 | 3.18E-05 | Stroma 3 | tuba1b |
| plk32 | 1E-09 | 0.6688235 | 0.245 | 0.201 | 3.28E-05 | Stroma 3 | plk3 |
| HACD41 | 1E-09 | 0.6631728 | 0.175 | 0.133 | 3.34E-05 | Stroma 3 | HACD4 |
| si:ch211-194e15.52 | 1.1E-09 | 0.5756128 | 0.143 | 0.102 | 3.48E-05 | Stroma 3 | si:ch211-194e15.5 |
| hbegfa3 | 1.2E-09 | 0.4692316 | 0.442 | 0.401 | 3.83E-05 | Stroma 3 | hbegfa |
| adgre101 | 1.4E-09 | 0.7909407 | 0.11 | 0.075 | 4.47E-05 | Stroma 3 | adgre10 |
| pde4ba1 | 1.5E-09 | 0.6929982 | 0.156 | 0.116 | 5.01E-05 | Stroma 3 | pde4ba |
| ctsc | 2E-09 | 0.8451617 | 0.1 | 0.067 | 6.4E-05 | Stroma 3 | ctsc |
| tgm1l14 | 2.1E-09 | 0.8187268 | 0.167 | 0.127 | 6.71E-05 | Stroma 3 | tgm1l1 |
| ahr24 | 2.4E-09 | 0.4431219 | 0.416 | 0.383 | 7.91E-05 | Stroma 3 | ahr2 |
| eif4g3a | 2.6E-09 | 0.8387431 | 0.1 | 0.067 | 8.58E-05 | Stroma 3 | eif4g3a |
| fhl1b2 | 2.8E-09 | 0.8603272 | 0.13 | 0.093 | 9.03E-05 | Stroma 3 | fhl1b |
| b4galt6 | 4E-09 | 0.8216293 | 0.142 | 0.104 | 0.000131 | Stroma 3 | b4galt6 |
| fmnl2a1 | 4.2E-09 | 0.7705691 | 0.147 | 0.109 | 0.000136 | Stroma 3 | fmnl2a |
| tcf122 | 4.8E-09 | 0.2739441 | 0.565 | 0.522 | 0.000155 | Stroma 3 | tcf12 |
| itgb24 | 6.4E-09 | 0.6971274 | 0.143 | 0.105 | 0.000207 | Stroma 3 | itgb2 |

| | | | | | | | |
|---|---|---|---|---|---|---|---|
| ctsba1 | 7.3E-09 | 0.4401673 | 0.415 | 0.383 | 0.000236 | Stroma 3 | ctsba |
| bag31 | 9.4E-09 | 1.1781017 | 0.102 | 0.071 | 0.000305 | Stroma 3 | bag3 |
| RALGDS2 | 9.8E-09 | 0.9605372 | 0.169 | 0.132 | 0.000318 | Stroma 3 | RALGDS |
| skib3 | 1.1E-08 | 0.5925522 | 0.243 | 0.201 | 0.000349 | Stroma 3 | skib |
| FP102018.12 | 1.1E-08 | 0.287784 | 0.43 | 0.39 | 0.000352 | Stroma 3 | FP102018.1 |
| trim3b4 | 1.1E-08 | 0.7958444 | 0.129 | 0.094 | 0.000358 | Stroma 3 | trim3b |
| thbs4b3 | 1.5E-08 | 0.353776 | 0.3 | 0.258 | 0.000489 | Stroma 3 | thbs4b |
| sost3 | 1.6E-08 | 0.7275857 | 0.111 | 0.077 | 0.000528 | Stroma 3 | sost |
| socs1a2 | 1.8E-08 | 0.7201089 | 0.161 | 0.123 | 0.000581 | Stroma 3 | socs1a |
| ngfb3 | 2.2E-08 | 0.8285081 | 0.114 | 0.081 | 0.000721 | Stroma 3 | ngfb |
| CABZ01074397.12 | 2.4E-08 | 0.5933155 | 0.189 | 0.148 | 0.00078 | Stroma 3 | CABZ01074397.1 |
| nucb11 | 2.4E-08 | 0.6042012 | 0.245 | 0.206 | 0.000789 | Stroma 3 | nucb1 |
| jdp2b1 | 2.6E-08 | 0.257308 | 0.61 | 0.57 | 0.000854 | Stroma 3 | jdp2b |
| tmsb23 | 3.2E-08 | 0.8373238 | 0.131 | 0.096 | 0.001031 | Stroma 3 | tmsb2 |
| f103 | 3.2E-08 | 0.3581468 | 0.435 | 0.399 | 0.001055 | Stroma 3 | f10 |
| polr2a2 | 3.3E-08 | 0.5663911 | 0.392 | 0.363 | 0.001071 | Stroma 3 | polr2a |
| psap1 | 3.8E-08 | 0.4583261 | 0.379 | 0.349 | 0.001241 | Stroma 3 | psap |
| il11ra4 | 4.1E-08 | 0.4774698 | 0.282 | 0.239 | 0.001322 | Stroma 3 | il11ra |
| serpina10a3 | 4.1E-08 | 0.3712275 | 0.412 | 0.378 | 0.001334 | Stroma 3 | serpina10a |
| grb10a2 | 5.1E-08 | 0.7289434 | 0.143 | 0.108 | 0.001646 | Stroma 3 | grb10a |
| tnfaip2a2 | 5.9E-08 | 0.7206785 | 0.121 | 0.088 | 0.00192 | Stroma 3 | tnfaip2a |
| CR318588.11 | 6.2E-08 | 0.7507128 | 0.336 | 0.294 | 0.002004 | Stroma 3 | CR318588.1 |
| clu2 | 6.7E-08 | 0.3317616 | 0.433 | 0.382 | 0.002176 | Stroma 3 | clu |
| si:dkey-16l2.161 | 7.3E-08 | 0.6585556 | 0.153 | 0.117 | 0.002368 | Stroma 3 | si:dkey-16l2.16 |
| irf2bp2b | 7.8E-08 | 0.5007302 | 0.248 | 0.209 | 0.00254 | Stroma 3 | irf2bp2b |
| asap1b3 | 8E-08 | 0.6723821 | 0.187 | 0.151 | 0.002601 | Stroma 3 | asap1b |
| gli2a | 9.1E-08 | 0.7505608 | 0.1 | 0.069 | 0.002972 | Stroma 3 | gli2a |
| si:ch211-196l7.42 | 1.1E-07 | 0.3471173 | 0.494 | 0.476 | 0.003552 | Stroma 3 | si:ch211-196l7.4 |
| dusp61 | 1.1E-07 | 0.7575056 | 0.181 | 0.145 | 0.003724 | Stroma 3 | dusp6 |
| si:ch211-132g1.31 | 1.2E-07 | 0.59327 | 0.137 | 0.103 | 0.003831 | Stroma 3 | si:ch211-132g1.3 |
| epn22 | 1.3E-07 | 0.6295855 | 0.143 | 0.108 | 0.004226 | Stroma 3 | epn2 |
| mmp14b3 | 1.3E-07 | 0.2485713 | 0.574 | 0.55 | 0.004249 | Stroma 3 | mmp14b |
| atf4b1 | 1.6E-07 | 0.216138 | 0.641 | 0.634 | 0.005226 | Stroma 3 | atf4b |
| hspg21 | 2.1E-07 | 0.6034509 | 0.182 | 0.145 | 0.006732 | Stroma 3 | hspg2 |
| dtnbp1a1 | 2.2E-07 | 0.6938575 | 0.144 | 0.111 | 0.007078 | Stroma 3 | dtnbp1a |
| MYADM1 | 2.3E-07 | 0.7040222 | 0.129 | 0.096 | 0.007379 | Stroma 3 | MYADM |
| cyp3c12 | 2.9E-07 | 0.698524 | 0.126 | 0.094 | 0.00954 | Stroma 3 | cyp3c1 |
| pnpla61 | 3.9E-07 | 0.612158 | 0.139 | 0.105 | 0.012602 | Stroma 3 | pnpla6 |
| rab5aa | 4E-07 | 0.4786611 | 0.296 | 0.261 | 0.013026 | Stroma 3 | rab5aa |
| rnf242 | 4.3E-07 | 0.7539801 | 0.122 | 0.092 | 0.013904 | Stroma 3 | rnf24 |
| ube2e21 | 5.2E-07 | 0.6647534 | 0.146 | 0.113 | 0.016815 | Stroma 3 | ube2e2 |
| ano63 | 5.6E-07 | 0.6220661 | 0.158 | 0.124 | 0.018133 | Stroma 3 | ano6 |
| si:dkey-42i9.6 | 7.4E-07 | 0.6003211 | 0.228 | 0.195 | 0.023923 | Stroma 3 | si:dkey-42i9.6 |
| sntb23 | 1E-06 | 0.4914002 | 0.27 | 0.236 | 0.033143 | Stroma 3 | sntb2 |
| cdon2 | 1.2E-06 | 0.520263 | 0.224 | 0.187 | 0.038939 | Stroma 3 | cdon |
| ifi30 | 1.3E-06 | 0.6285443 | 0.178 | 0.145 | 0.04133 | Stroma 3 | ifi30 |
| cica1 | 1.5E-06 | 0.5903376 | 0.219 | 0.185 | 0.049237 | Stroma 3 | cica |
| zgc:77486.13 | 1.7E-06 | 0.4992742 | 0.203 | 0.168 | 0.053763 | Stroma 3 | zgc:77486.1 |

| | | | | | | | |
|---|---|---|---|---|---|---|---|
| mical3a.1 | 1.7E-06 | 0.673075 | 0.134 | 0.104 | 0.054044 | Stroma 3 | mical3a.1 |
| asap32 | 1.7E-06 | 0.5453117 | 0.158 | 0.125 | 0.056618 | Stroma 3 | asap3 |
| mvp3 | 1.8E-06 | 0.4141529 | 0.347 | 0.316 | 0.057163 | Stroma 3 | mvp |
| fam49a1 | 2E-06 | 0.8315386 | 0.101 | 0.074 | 0.065342 | Stroma 3 | fam49a |
| rexo11 | 2.3E-06 | 0.754168 | 0.148 | 0.118 | 0.075234 | Stroma 3 | rexo1 |
| hdlbpa3 | 2.5E-06 | 0.2329874 | 0.674 | 0.676 | 0.081367 | Stroma 3 | hdlbpa |
| pgrmc11 | 2.9E-06 | 0.4058372 | 0.385 | 0.364 | 0.093618 | Stroma 3 | pgrmc1 |
| ccnl1a1 | 3.2E-06 | 0.4823287 | 0.295 | 0.262 | 0.105445 | Stroma 3 | ccnl1a |
| arhgef401 | 3.3E-06 | 0.5539056 | 0.223 | 0.191 | 0.10679 | Stroma 3 | arhgef40 |
| tax1bp31 | 3.8E-06 | 0.4067449 | 0.285 | 0.252 | 0.122601 | Stroma 3 | tax1bp3 |
| loxl2a1 | 4.1E-06 | 0.3246121 | 0.26 | 0.223 | 0.132007 | Stroma 3 | loxl2a |
| zyx4 | 4.3E-06 | 0.2277604 | 0.359 | 0.317 | 0.139081 | Stroma 3 | zyx |
| ctsd1 | 4.5E-06 | 0.5152042 | 0.185 | 0.153 | 0.147453 | Stroma 3 | ctsd |
| id2b2 | 5.6E-06 | 0.6990336 | 0.222 | 0.19 | 0.180693 | Stroma 3 | id2b |
| cald1b4 | 5.7E-06 | 0.2022502 | 0.204 | 0.167 | 0.184818 | Stroma 3 | cald1b |
| prss162 | 6.1E-06 | 0.5909743 | 0.117 | 0.09 | 0.197409 | Stroma 3 | prss16 |
| nfkb21 | 6.6E-06 | 0.7772013 | 0.127 | 0.1 | 0.214498 | Stroma 3 | nfkb2 |
| nr1d2a1 | 6.8E-06 | 0.4146029 | 0.283 | 0.251 | 0.222099 | Stroma 3 | nr1d2a |
| pcolcea5 | 7.1E-06 | 0.4023884 | 0.568 | 0.569 | 0.229426 | Stroma 3 | pcolcea |
| klf12a3 | 7.5E-06 | 0.5055696 | 0.169 | 0.138 | 0.244233 | Stroma 3 | klf12a |
| ptges3 | 8.3E-06 | 0.6760347 | 0.138 | 0.108 | 0.268458 | Stroma 3 | ptges |
| zgc:1949891 | 8.5E-06 | 0.6927514 | 0.117 | 0.09 | 0.275985 | Stroma 3 | zgc:194989 |
| marcksl1b3 | 9.2E-06 | 0.4325592 | 0.289 | 0.254 | 0.297867 | Stroma 3 | marcksl1b |
| cbx7a1 | 9.2E-06 | 0.2314406 | 0.38 | 0.346 | 0.299701 | Stroma 3 | cbx7a |
| pora1 | 1.2E-05 | 0.5921116 | 0.138 | 0.11 | 0.375088 | Stroma 3 | pora |
| ctsz | 1.5E-05 | 0.5769489 | 0.191 | 0.162 | 0.478941 | Stroma 3 | ctsz |
| akap132 | 1.9E-05 | 0.3570905 | 0.346 | 0.321 | 0.618452 | Stroma 3 | akap13 |
| sort1a2 | 2.1E-05 | 0.7111245 | 0.14 | 0.113 | 0.69287 | Stroma 3 | sort1a |
| rasgef1ba2 | 2.3E-05 | 0.4773552 | 0.191 | 0.16 | 0.736767 | Stroma 3 | rasgef1ba |
| adprh2 | 2.4E-05 | 0.6674769 | 0.104 | 0.08 | 0.782261 | Stroma 3 | adprh |
| barx12 | 2.8E-05 | 0.4197863 | 0.158 | 0.127 | 0.925742 | Stroma 3 | barx1 |
| sqstm11 | 3.2E-05 | 0.3829756 | 0.329 | 0.305 | 1 | Stroma 3 | sqstm1 |
| rnf213b | 3.5E-05 | 0.6281381 | 0.102 | 0.079 | 1 | Stroma 3 | rnf213b |
| socs5b | 3.7E-05 | 0.7051133 | 0.116 | 0.091 | 1 | Stroma 3 | socs5b |
| nccrp11 | 4.5E-05 | 0.5138834 | 0.297 | 0.276 | 1 | Stroma 3 | nccrp1 |
| amotl2b | 5.3E-05 | 0.7448009 | 0.103 | 0.08 | 1 | Stroma 3 | amotl2b |
| mgst3a1 | 5.5E-05 | 0.4728922 | 0.28 | 0.255 | 1 | Stroma 3 | mgst3a |
| nck1b1 | 5.5E-05 | 0.6148886 | 0.122 | 0.097 | 1 | Stroma 3 | nck1b |
| ebf3a3 | 6.5E-05 | 0.329944 | 0.247 | 0.215 | 1 | Stroma 3 | ebf3a |
| mgst3b | 7.7E-05 | 0.3163838 | 0.399 | 0.385 | 1 | Stroma 3 | mgst3b |
| brd2a1 | 7.7E-05 | 0.3162574 | 0.464 | 0.447 | 1 | Stroma 3 | brd2a |
| tspan17 | 7.9E-05 | 0.5930625 | 0.1 | 0.077 | 1 | Stroma 3 | tspan17 |
| fuca21 | 8.9E-05 | 0.4471365 | 0.173 | 0.146 | 1 | Stroma 3 | fuca2 |
| myo10l31 | 0.0001 | 0.5643966 | 0.123 | 0.099 | 1 | Stroma 3 | myo10l3 |
| sc:d2171 | 0.0001 | 0.5237784 | 0.161 | 0.135 | 1 | Stroma 3 | sc:d217 |
| lpar13 | 0.00011 | 0.4455358 | 0.208 | 0.181 | 1 | Stroma 3 | lpar1 |
| tmem50a1 | 0.00011 | 0.443119 | 0.257 | 0.235 | 1 | Stroma 3 | tmem50a |
| map3k41 | 0.00012 | 0.6337999 | 0.106 | 0.083 | 1 | Stroma 3 | map3k4 |

| | | | | | | | |
|---|---|---|---|---|---|---|---|
| emx23 | 0.00013 | 0.5724243 | 0.115 | 0.092 | 1 | Stroma 3 | emx2 |
| tasor21 | 0.00013 | 0.440247 | 0.174 | 0.147 | 1 | Stroma 3 | tasor2 |
| CTBP11 | 0.00014 | 0.4773373 | 0.194 | 0.168 | 1 | Stroma 3 | CTBP1 |
| rhoaa1 | 0.00016 | 0.3080521 | 0.434 | 0.421 | 1 | Stroma 3 | rhoaa |
| plscr3b2 | 0.00017 | 0.3890422 | 0.265 | 0.24 | 1 | Stroma 3 | plscr3b |
| slc38a21 | 0.00018 | 0.6902855 | 0.1 | 0.079 | 1 | Stroma 3 | slc38a2 |
| zgc:1105912 | 0.00019 | 0.5036369 | 0.116 | 0.093 | 1 | Stroma 3 | zgc:110591 |
| anxa5b | 0.0002 | 0.3364266 | 0.411 | 0.403 | 1 | Stroma 3 | anxa5b |
| ctsa1 | 0.0002 | 0.4482065 | 0.142 | 0.117 | 1 | Stroma 3 | ctsa |
| frzb4 | 0.00023 | 0.2401054 | 0.155 | 0.129 | 1 | Stroma 3 | frzb |
| clk4a3 | 0.00028 | 0.4057789 | 0.253 | 0.229 | 1 | Stroma 3 | clk4a |
| clic11 | 0.00028 | 0.6151046 | 0.126 | 0.104 | 1 | Stroma 3 | clic1 |
| lamp1b1 | 0.00029 | 0.6386952 | 0.11 | 0.089 | 1 | Stroma 3 | lamp1b |
| lrba | 0.00032 | 0.6836053 | 0.199 | 0.177 | 1 | Stroma 3 | lrba |
| cox4i21 | 0.00032 | 0.3593887 | 0.338 | 0.321 | 1 | Stroma 3 | cox4i2 |
| dap1b.12 | 0.00034 | 0.2177335 | 0.515 | 0.508 | 1 | Stroma 3 | dap1b.1 |
| fosl21 | 0.00035 | 0.3012151 | 0.44 | 0.429 | 1 | Stroma 3 | fosl2 |
| rbm39a1 | 0.00035 | 0.387489 | 0.252 | 0.229 | 1 | Stroma 3 | rbm39a |
| fndc3ba1 | 0.00036 | 0.6764004 | 0.16 | 0.139 | 1 | Stroma 3 | fndc3ba |
| fcho22 | 0.00036 | 0.4138004 | 0.183 | 0.159 | 1 | Stroma 3 | fcho2 |
| prex21 | 0.00036 | 0.4270217 | 0.109 | 0.087 | 1 | Stroma 3 | prex2 |
| rbm51 | 0.00037 | 0.3822204 | 0.211 | 0.185 | 1 | Stroma 3 | rbm5 |
| gpx4a | 0.00045 | 0.365341 | 0.322 | 0.303 | 1 | Stroma 3 | gpx4a |
| zgc:1735872 | 0.00046 | 0.5574205 | 0.154 | 0.131 | 1 | Stroma 3 | zgc:173587 |
| rnd1b4 | 0.00049 | 0.5508149 | 0.124 | 0.103 | 1 | Stroma 3 | rnd1b |
| tnrc6a2 | 0.00049 | 0.3508876 | 0.294 | 0.272 | 1 | Stroma 3 | tnrc6a |
| tor11 | 0.0005 | 0.4485402 | 0.13 | 0.108 | 1 | Stroma 3 | tor1 |
| si:dkey-235d18.52 | 0.00055 | 0.5117814 | 0.119 | 0.098 | 1 | Stroma 3 | si:dkey-235d18.5 |
| rbfox21 | 0.00061 | 0.3643125 | 0.293 | 0.271 | 1 | Stroma 3 | rbfox2 |
| mapk14b | 0.00067 | 0.6380142 | 0.106 | 0.086 | 1 | Stroma 3 | mapk14b |
| tmem131 | 0.00067 | 0.3875554 | 0.22 | 0.197 | 1 | Stroma 3 | tmem131 |
| si:dkey-195m11.82 | 0.00069 | 0.5141416 | 0.15 | 0.127 | 1 | Stroma 3 | si:dkey-195m11.8 |
| man2b11 | 0.00072 | 0.4695753 | 0.118 | 0.097 | 1 | Stroma 3 | man2b1 |
| dpp71 | 0.00075 | 0.4273913 | 0.235 | 0.215 | 1 | Stroma 3 | dpp7 |
| il4r.11 | 0.00079 | 0.5162401 | 0.174 | 0.152 | 1 | Stroma 3 | il4r.1 |
| gsta.13 | 0.00087 | 0.2244699 | 0.378 | 0.357 | 1 | Stroma 3 | gsta.1 |
| nfkbiaa3 | 0.00091 | 0.2590588 | 0.44 | 0.423 | 1 | Stroma 3 | nfkbiaa |
| ptp4a3a1 | 0.00091 | 0.5014202 | 0.141 | 0.12 | 1 | Stroma 3 | ptp4a3a |
| ip6k2b2 | 0.00094 | 0.3639482 | 0.359 | 0.344 | 1 | Stroma 3 | ip6k2b |
| tcf7l1a1 | 0.00095 | 0.5788396 | 0.137 | 0.116 | 1 | Stroma 3 | tcf7l1a |
| trim8a1 | 0.00103 | 0.5495515 | 0.102 | 0.083 | 1 | Stroma 3 | trim8a |
| csrnp1b2 | 0.00104 | 0.4999959 | 0.207 | 0.187 | 1 | Stroma 3 | csrnp1b |
| sgk11 | 0.00115 | 0.2346811 | 0.388 | 0.367 | 1 | Stroma 3 | sgk1 |
| midn1 | 0.00117 | 0.2265063 | 0.446 | 0.434 | 1 | Stroma 3 | midn |
| cyb5r11 | 0.00129 | 0.3610776 | 0.224 | 0.201 | 1 | Stroma 3 | cyb5r1 |
| myclb1 | 0.0014 | 0.5365335 | 0.161 | 0.14 | 1 | Stroma 3 | myclb |
| hivep3a2 | 0.00147 | 0.3478376 | 0.13 | 0.109 | 1 | Stroma 3 | hivep3a |
| yaf22 | 0.00149 | 0.496829 | 0.128 | 0.109 | 1 | Stroma 3 | yaf2 |

| | | | | | | | |
|---|---|---|---|---|---|---|---|
| map3k7 | 0.00159 | 0.5177445 | 0.148 | 0.129 | 1 | Stroma 3 | map3k7 |
| zgc:163080 | 0.00177 | 0.5278971 | 0.142 | 0.123 | 1 | Stroma 3 | zgc:163080 |
| hdac42 | 0.00177 | 0.4951596 | 0.14 | 0.12 | 1 | Stroma 3 | hdac4 |
| zfp36l1a5 | 0.00187 | 0.3035858 | 0.229 | 0.206 | 1 | Stroma 3 | zfp36l1a |
| diaph22 | 0.00191 | 0.4890884 | 0.136 | 0.116 | 1 | Stroma 3 | diaph2 |
| dag11 | 0.00191 | 0.5504551 | 0.176 | 0.157 | 1 | Stroma 3 | dag1 |
| sox11a1 | 0.00202 | 0.2688831 | 0.358 | 0.344 | 1 | Stroma 3 | sox11a |
| tnfrsf1a1 | 0.00205 | 0.3615576 | 0.27 | 0.252 | 1 | Stroma 3 | tnfrsf1a |
| tspan4b4 | 0.00207 | 0.251644 | 0.353 | 0.337 | 1 | Stroma 3 | tspan4b |
| si:zfos-2326c3.21 | 0.00208 | 0.4685593 | 0.182 | 0.162 | 1 | Stroma 3 | si:zfos-2326c3.2 |
| gli32 | 0.00216 | 0.4987468 | 0.129 | 0.111 | 1 | Stroma 3 | gli3 |
| cited4a2 | 0.00231 | 0.5417109 | 0.134 | 0.115 | 1 | Stroma 3 | cited4a |
| chmp6b1 | 0.00249 | 0.4084019 | 0.172 | 0.151 | 1 | Stroma 3 | chmp6b |
| per1b1 | 0.0025 | 0.4048323 | 0.222 | 0.203 | 1 | Stroma 3 | per1b |
| ccdc3b3 | 0.00251 | 0.4992501 | 0.143 | 0.125 | 1 | Stroma 3 | ccdc3b |
| pkig2 | 0.00262 | 0.5076467 | 0.136 | 0.118 | 1 | Stroma 3 | pkig |
| dido1 | 0.00265 | 0.254924 | 0.432 | 0.427 | 1 | Stroma 3 | dido1 |
| elmsan1b2 | 0.0028 | 0.2733371 | 0.368 | 0.357 | 1 | Stroma 3 | elmsan1b |
| cyth1a.11 | 0.00283 | 0.5177275 | 0.154 | 0.136 | 1 | Stroma 3 | cyth1a.1 |
| taok1a | 0.00285 | 0.4288164 | 0.166 | 0.147 | 1 | Stroma 3 | taok1a |
| foxo41 | 0.00306 | 0.526145 | 0.124 | 0.106 | 1 | Stroma 3 | foxo4 |
| plxna32 | 0.00315 | 0.5995381 | 0.105 | 0.089 | 1 | Stroma 3 | plxna3 |
| golga1 | 0.00327 | 0.5647372 | 0.108 | 0.091 | 1 | Stroma 3 | golga1 |
| porb3 | 0.00347 | 0.4608899 | 0.127 | 0.108 | 1 | Stroma 3 | porb |
| rmdn12 | 0.00364 | 0.6015304 | 0.111 | 0.094 | 1 | Stroma 3 | rmdn1 |
| hmox1a2 | 0.00378 | 0.4436306 | 0.195 | 0.174 | 1 | Stroma 3 | hmox1a |
| ugt2a72 | 0.00403 | 0.5262438 | 0.11 | 0.094 | 1 | Stroma 3 | ugt2a7 |
| si:dkey-17m8.12 | 0.00406 | 0.4448766 | 0.141 | 0.123 | 1 | Stroma 3 | si:dkey-17m8.1 |
| nt5dc2 | 0.00415 | 0.4223647 | 0.147 | 0.129 | 1 | Stroma 3 | nt5dc2 |
| pepd2 | 0.00424 | 0.4395819 | 0.228 | 0.213 | 1 | Stroma 3 | pepd |
| id2a2 | 0.00425 | 0.2143236 | 0.507 | 0.504 | 1 | Stroma 3 | id2a |
| si:dkey-253d23.41 | 0.00431 | 0.5838718 | 0.105 | 0.088 | 1 | Stroma 3 | si:dkey-253d23.4 |
| napsa2 | 0.00438 | 0.4133779 | 0.188 | 0.169 | 1 | Stroma 3 | napsa |
| coro1ca3 | 0.00469 | 0.3808297 | 0.252 | 0.238 | 1 | Stroma 3 | coro1ca |
| plxnb2a.11 | 0.00502 | 0.4777264 | 0.106 | 0.09 | 1 | Stroma 3 | plxnb2a.1 |
| u2surp2 | 0.00502 | 0.3082758 | 0.323 | 0.314 | 1 | Stroma 3 | u2surp |
| fblim12 | 0.00516 | 0.3797427 | 0.129 | 0.111 | 1 | Stroma 3 | fblim1 |
| cacna1g3 | 0.0052 | 0.466426 | 0.113 | 0.096 | 1 | Stroma 3 | cacna1g |
| sik13 | 0.00536 | 0.2265693 | 0.275 | 0.253 | 1 | Stroma 3 | sik1 |
| atp1a1a.12 | 0.00547 | 0.3187221 | 0.331 | 0.323 | 1 | Stroma 3 | atp1a1a.1 |
| per21 | 0.00578 | 0.4451988 | 0.122 | 0.105 | 1 | Stroma 3 | per2 |
| mbnl31 | 0.00605 | 0.4872675 | 0.15 | 0.133 | 1 | Stroma 3 | mbnl3 |
| cd1642 | 0.00727 | 0.3284273 | 0.344 | 0.338 | 1 | Stroma 3 | cd164 |
| rcan1a2 | 0.00748 | 0.4022523 | 0.244 | 0.23 | 1 | Stroma 3 | rcan1a |
| fam13b1 | 0.0075 | 0.4864572 | 0.136 | 0.12 | 1 | Stroma 3 | fam13b |
| itm2bb3 | 0.0075 | 0.3630821 | 0.105 | 0.089 | 1 | Stroma 3 | itm2bb |
| maff1 | 0.00797 | 0.3614588 | 0.238 | 0.224 | 1 | Stroma 3 | maff |
| tnfsf102 | 0.00802 | 0.3767825 | 0.115 | 0.098 | 1 | Stroma 3 | tnfsf10 |

| | | | | | | | |
|---|---|---|---|---|---|---|---|
| si:dkey-188i13.102 | 0.00818 | 0.3343628 | 0.262 | 0.248 | 1 | Stroma 3 | si:dkey-188i13.10 |
| mcf2l21 | 0.00828 | 0.5370619 | 0.122 | 0.107 | 1 | Stroma 3 | mcf2l2 |
| cmtm61 | 0.00829 | 0.4493787 | 0.156 | 0.14 | 1 | Stroma 3 | cmtm6 |
| tmem1891 | 0.00844 | 0.4345334 | 0.147 | 0.131 | 1 | Stroma 3 | tmem189 |
| dstyk | 0.00853 | 0.5461582 | 0.113 | 0.098 | 1 | Stroma 3 | dstyk |
| rgs12b2 | 0.00889 | 0.4980679 | 0.103 | 0.088 | 1 | Stroma 3 | rgs12b |
| gusb2 | 0.0091 | 0.3298152 | 0.146 | 0.129 | 1 | Stroma 3 | gusb |
| prdx61 | 0.00933 | 0.4555631 | 0.341 | 0.345 | 1 | Stroma 3 | prdx6 |
| zgc:1140463 | 0.0095 | 0.3448138 | 0.116 | 0.1 | 1 | Stroma 3 | zgc:114046 |
| ca63 | 0 | 1.720825 | 0.852 | 0.322 | 0 | DF 5 | ca6 |
| si:ch211-251b21.13 | 0 | 1.3510687 | 0.899 | 0.429 | 0 | DF 5 | si:ch211-251b21.1 |
| fgl13 | 0 | 2.431847 | 0.621 | 0.19 | 0 | DF 5 | fgl1 |
| col7a12 | 0 | 2.2876563 | 0.535 | 0.136 | 0 | DF 5 | col7a1 |
| tfap2c3 | 0 | 2.1734716 | 0.573 | 0.177 | 0 | DF 5 | tfap2c |
| ecrg4a1 | 0 | 2.2796715 | 0.923 | 0.546 | 0 | DF 5 | ecrg4a |
| ckbb1 | 0 | 2.409693 | 0.552 | 0.182 | 0 | DF 5 | ckbb |
| glulb3 | 0 | 1.4327624 | 0.869 | 0.555 | 0 | DF 5 | glulb |
| prdm1a3 | 0 | 2.8401032 | 0.344 | 0.064 | 0 | DF 5 | prdm1a |
| tmeff2b1 | 0 | 3.9473057 | 0.256 | 0.02 | 0 | DF 5 | tmeff2b |
| inhbb | 0 | 2.9876405 | 0.273 | 0.046 | 0 | DF 5 | inhbb |
| CR786577.1 | 0 | 4.3727657 | 0.131 | 0.007 | 0 | DF 5 | CR786577.1 |
| ucp24 | 2E-289 | 1.4067776 | 0.744 | 0.399 | 7.4E-285 | DF 5 | ucp2 |
| qdpra4 | 3E-263 | 1.089045 | 0.78 | 0.409 | 1E-258 | DF 5 | qdpra |
| cyp1c13 | 2E-258 | 2.4261989 | 0.322 | 0.092 | 6.2E-254 | DF 5 | cyp1c1 |
| grem2b | 3E-257 | 3.9187059 | 0.1 | 0.009 | 1E-252 | DF 5 | grem2b |
| igsf102 | 3E-251 | 1.3052885 | 0.605 | 0.262 | 1E-246 | DF 5 | igsf10 |
| arhgef7a | 5E-242 | 2.4342843 | 0.187 | 0.035 | 1.6E-237 | DF 5 | arhgef7a |
| baiap2l1a1 | 5E-241 | 2.4915787 | 0.176 | 0.031 | 1.6E-236 | DF 5 | baiap2l1a |
| eif11 | 6E-240 | 0.9899288 | 0.929 | 0.825 | 2.1E-235 | DF 5 | eif1 |
| col6a4a2 | 2E-236 | 2.3486334 | 0.22 | 0.048 | 6.1E-232 | DF 5 | col6a4a |
| hsp70.11 | 8E-236 | 1.8300221 | 0.628 | 0.328 | 2.5E-231 | DF 5 | hsp70.1 |
| hyal42 | 4E-234 | 2.0851578 | 0.32 | 0.096 | 1.2E-229 | DF 5 | hyal4 |
| net13 | 2E-229 | 1.3852584 | 0.665 | 0.359 | 6.8E-225 | DF 5 | net1 |
| en1b2 | 6E-229 | 1.7842147 | 0.33 | 0.1 | 2E-224 | DF 5 | en1b |
| hsp70.31 | 2E-219 | 1.6110078 | 0.783 | 0.543 | 7.6E-215 | DF 5 | hsp70.3 |
| pcm14 | 2E-219 | 1.7920226 | 0.393 | 0.141 | 7.9E-215 | DF 5 | pcm1 |
| twist33 | 7E-219 | 1.2676087 | 0.551 | 0.239 | 2.4E-214 | DF 5 | twist3 |
| fstb5 | 6E-218 | 1.0347277 | 0.687 | 0.346 | 1.9E-213 | DF 5 | fstb |
| hsp70.21 | 1E-216 | 1.5049968 | 0.756 | 0.481 | 4.3E-212 | DF 5 | hsp70.2 |
| zgc:1626123 | 3E-216 | 1.7470738 | 0.334 | 0.106 | 9.5E-212 | DF 5 | zgc:162612 |
| fmoda2 | 9E-215 | 1.2319937 | 0.457 | 0.17 | 3E-210 | DF 5 | fmoda |
| cyp26b11 | 6E-206 | 1.1972327 | 0.67 | 0.354 | 2E-201 | DF 5 | cyp26b1 |
| kif26ba4 | 1E-200 | 1.7371677 | 0.313 | 0.098 | 3.1E-196 | DF 5 | kif26ba |
| cdh114 | 2E-199 | 0.940802 | 0.848 | 0.578 | 5.9E-195 | DF 5 | cdh11 |
| ndrg41 | 2E-199 | 1.6172373 | 0.388 | 0.143 | 7.9E-195 | DF 5 | ndrg4 |
| hsp70l1 | 8E-196 | 1.7353364 | 0.562 | 0.29 | 2.5E-191 | DF 5 | hsp70l |
| zgc:565855 | 1E-191 | 1.1239821 | 0.625 | 0.314 | 3.6E-187 | DF 5 | zgc:56585 |
| moxd11 | 1E-190 | 2.3730657 | 0.195 | 0.046 | 4.7E-186 | DF 5 | moxd1 |

| | | | | | | | |
|---|---|---|---|---|---|---|---|
| pnrc22 | 4E-190 | 0.8182895 | 0.96 | 0.877 | 1.1E-185 | DF 5 | pnrc2 |
| spock1 | 2E-189 | 2.4941423 | 0.165 | 0.034 | 5.1E-185 | DF 5 | spock1 |
| inka1a2 | 1E-184 | 1.5903587 | 0.407 | 0.164 | 4.4E-180 | DF 5 | inka1a |
| spock34 | 2E-179 | 0.7724913 | 0.741 | 0.396 | 6.2E-175 | DF 5 | spock3 |
| adma4 | 6E-165 | 1.1445559 | 0.488 | 0.222 | 1.8E-160 | DF 5 | adma |
| col5a3b1 | 5E-163 | 1.9969044 | 0.173 | 0.041 | 1.5E-158 | DF 5 | col5a3b |
| cebpb2 | 1E-162 | 1.4571302 | 0.674 | 0.438 | 3.6E-158 | DF 5 | cebpb |
| hsp90aa1.21 | 7E-159 | 1.3181295 | 0.795 | 0.591 | 2.3E-154 | DF 5 | hsp90aa1.2 |
| hhip2 | 3E-157 | 1.7077571 | 0.253 | 0.081 | 8.5E-153 | DF 5 | hhip |
| im:71523483 | 5E-152 | 1.3095002 | 0.466 | 0.225 | 1.6E-147 | DF 5 | im:7152348 |
| ednraa3 | 1E-148 | 1.2550208 | 0.44 | 0.203 | 3.5E-144 | DF 5 | ednraa |
| tnc5 | 2E-147 | 0.897181 | 0.73 | 0.458 | 5.3E-143 | DF 5 | tnc |
| sfrp24 | 1E-146 | 0.7430821 | 0.683 | 0.384 | 3.3E-142 | DF 5 | sfrp2 |
| smarcd22 | 3E-143 | 1.4273151 | 0.367 | 0.155 | 8.2E-139 | DF 5 | smarcd2 |
| tbx3a2 | 1E-138 | 1.3989694 | 0.305 | 0.116 | 3.3E-134 | DF 5 | tbx3a |
| junbb2 | 1E-137 | 0.9700529 | 0.79 | 0.592 | 3.4E-133 | DF 5 | junbb |
| ptmab2 | 4E-137 | 0.7624328 | 0.881 | 0.707 | 1.4E-132 | DF 5 | ptmab |
| mcl1a2 | 7E-137 | 0.8580919 | 0.887 | 0.766 | 2.4E-132 | DF 5 | mcl1a |
| si:dkey-23i12.71 | 1E-136 | 1.680305 | 0.321 | 0.132 | 3.8E-132 | DF 5 | si:dkey-23i12.7 |
| rsrp12 | 1E-135 | 1.1790751 | 0.721 | 0.534 | 3.4E-131 | DF 5 | rsrp1 |
| qsox12 | 2E-134 | 1.6360044 | 0.206 | 0.062 | 5.6E-130 | DF 5 | qsox1 |
| tfap2b | 1E-132 | 2.1673495 | 0.11 | 0.021 | 4.8E-128 | DF 5 | tfap2b |
| fosl1a1 | 2E-132 | 1.0259466 | 0.664 | 0.427 | 5.9E-128 | DF 5 | fosl1a |
| gstm.14 | 3E-131 | 0.8480596 | 0.776 | 0.589 | 9E-127 | DF 5 | gstm.1 |
| twist25 | 2E-130 | 0.9362012 | 0.591 | 0.329 | 5.4E-126 | DF 5 | twist2 |
| zgc:1229791 | 3E-130 | 1.9290397 | 0.221 | 0.073 | 1.1E-125 | DF 5 | zgc:122979 |
| sdc41 | 1E-129 | 0.771948 | 0.853 | 0.697 | 3.5E-125 | DF 5 | sdc4 |
| dlc1 | 1E-129 | 2.1804229 | 0.128 | 0.029 | 3.6E-125 | DF 5 | dlc |
| cald1b5 | 3E-129 | 1.3075879 | 0.364 | 0.158 | 1.1E-124 | DF 5 | cald1b |
| pdgfra4 | 2E-126 | 0.896938 | 0.698 | 0.449 | 5.8E-122 | DF 5 | pdgfra |
| lmo4b4 | 9E-126 | 1.0815381 | 0.545 | 0.308 | 3.1E-121 | DF 5 | lmo4b |
| wnt5b1 | 4E-125 | 1.6048003 | 0.27 | 0.102 | 1.3E-120 | DF 5 | wnt5b |
| fosl22 | 1E-122 | 0.9909587 | 0.642 | 0.419 | 3.6E-118 | DF 5 | fosl2 |
| rpl23a3 | 2E-122 | 0.4331767 | 0.984 | 0.97 | 5.1E-118 | DF 5 | rpl23a |
| id31 | 3E-122 | 1.0195797 | 0.712 | 0.502 | 1E-117 | DF 5 | id3 |
| ptx3a4 | 2E-121 | 0.3344255 | 0.787 | 0.473 | 5.6E-117 | DF 5 | ptx3a |
| lfng3 | 2E-120 | 1.2497533 | 0.462 | 0.244 | 6.4E-116 | DF 5 | lfng |
| rps261 | 1E-118 | 0.4016739 | 0.983 | 0.968 | 4.1E-114 | DF 5 | rps26 |
| her62 | 8E-117 | 1.393119 | 0.622 | 0.424 | 2.6E-112 | DF 5 | her6 |
| ttc365 | 1E-116 | 0.8364826 | 0.6 | 0.342 | 4.6E-112 | DF 5 | ttc36 |
| mycb2 | 2E-116 | 0.8118879 | 0.814 | 0.656 | 7.5E-112 | DF 5 | mycb |
| ubb2 | 2E-115 | 0.8227153 | 0.822 | 0.674 | 5.7E-111 | DF 5 | ubb |
| scinla2 | 2E-115 | 0.9825523 | 0.625 | 0.399 | 6.3E-111 | DF 5 | scinla |
| xbp11 | 1E-114 | 0.6878844 | 0.879 | 0.756 | 3.9E-110 | DF 5 | xbp1 |
| gstp14 | 5E-113 | 0.7108227 | 0.823 | 0.636 | 1.5E-108 | DF 5 | gstp1 |
| sgk12 | 2E-112 | 1.0108742 | 0.576 | 0.357 | 6.5E-108 | DF 5 | sgk1 |
| si:ch73-335l21.42 | 3E-112 | 0.9822824 | 0.743 | 0.563 | 9.2E-108 | DF 5 | si:ch73-335l21.4 |
| glula3 | 1E-110 | 0.6550869 | 0.818 | 0.609 | 3.5E-106 | DF 5 | glula |

| | | | | | | | |
|---|---|---|---|---|---|---|---|
| adamts24 | 6E-108 | 0.826059 | 0.602 | 0.363 | 2E-103 | DF 5 | adamts2 |
| id13 | 1E-106 | 0.8357842 | 0.813 | 0.633 | 4.4E-102 | DF 5 | id1 |
| col2a1b2 | 6E-105 | 0.4180151 | 0.731 | 0.491 | 1.9E-100 | DF 5 | col2a1b |
| prelid3b2 | 2E-104 | 1.0308597 | 0.552 | 0.349 | 5.7E-100 | DF 5 | prelid3b |
| UBB1 | 5E-104 | 0.9328146 | 0.728 | 0.555 | 1.8E-99 | DF 5 | UBB |
| krt82 | 2E-101 | 0.4029309 | 0.995 | 0.984 | 6.85E-97 | DF 5 | krt8 |
| ms4a17a.5.1 | 3E-101 | 2.1389289 | 0.107 | 0.025 | 1.02E-96 | DF 5 | ms4a17a.5.1 |
| dlx3b3 | 4E-100 | 0.9454824 | 0.563 | 0.351 | 1.18E-95 | DF 5 | dlx3b |
| capn2b5 | 3E-99 | 0.858329 | 0.53 | 0.313 | 9.16E-95 | DF 5 | capn2b |
| fosab2 | 1E-98 | 0.6134989 | 0.956 | 0.892 | 3.19E-94 | DF 5 | fosab |
| efna1b1 | 1.1E-97 | 1.2728341 | 0.301 | 0.136 | 3.48E-93 | DF 5 | efna1b |
| jdp2b2 | 1.2E-97 | 0.9014909 | 0.754 | 0.563 | 3.75E-93 | DF 5 | jdp2b |
| rpl312 | 3E-97 | 0.3058664 | 0.995 | 0.993 | 9.85E-93 | DF 5 | rpl31 |
| ppp1r102 | 3.7E-97 | 0.9235607 | 0.667 | 0.476 | 1.21E-92 | DF 5 | ppp1r10 |
| hmgb3a1 | 3.8E-97 | 1.3114796 | 0.262 | 0.109 | 1.22E-92 | DF 5 | hmgb3a |
| tpbga1 | 1.8E-95 | 1.7343044 | 0.132 | 0.037 | 5.79E-91 | DF 5 | tpbga |
| msx1b3 | 1.8E-94 | 1.0481185 | 0.483 | 0.285 | 5.73E-90 | DF 5 | msx1b |
| si:ch211-246m6.51 | 7.7E-93 | 1.6859796 | 0.156 | 0.05 | 2.49E-88 | DF 5 | si:ch211-246m6.5 |
| rpsa3 | 2E-92 | 0.316614 | 0.996 | 0.984 | 6.39E-88 | DF 5 | rpsa |
| rpl18a3 | 5.5E-91 | 0.3131543 | 0.992 | 0.984 | 1.79E-86 | DF 5 | rpl18a |
| lamc31 | 9.5E-91 | 1.3094548 | 0.264 | 0.114 | 3.09E-86 | DF 5 | lamc3 |
| dnajb1b2 | 1.5E-90 | 0.9218189 | 0.642 | 0.456 | 4.91E-86 | DF 5 | dnajb1b |
| cpe1 | 1.9E-90 | 0.9705029 | 0.38 | 0.195 | 6.22E-86 | DF 5 | cpe |
| map1sb | 5.8E-90 | 1.7448636 | 0.163 | 0.055 | 1.9E-85 | DF 5 | map1sb |
| si:ch211-222l21.11 | 2.5E-89 | 0.5095851 | 0.764 | 0.545 | 8.19E-85 | DF 5 | si:ch211-222l21.1 |
| CR936442.11 | 3.5E-88 | 1.1154583 | 0.357 | 0.183 | 1.12E-83 | DF 5 | CR936442.1 |
| tcf123 | 4.9E-88 | 0.6596161 | 0.722 | 0.514 | 1.59E-83 | DF 5 | tcf12 |
| rpl192 | 5.7E-88 | 0.3332215 | 0.987 | 0.983 | 1.85E-83 | DF 5 | rpl19 |
| mamdc2b1 | 8.2E-88 | 0.484844 | 0.438 | 0.233 | 2.66E-83 | DF 5 | mamdc2b |
| mcl1b3 | 1.5E-87 | 0.9129552 | 0.654 | 0.475 | 4.93E-83 | DF 5 | mcl1b |
| ccdc80l13 | 2.6E-87 | 1.2732662 | 0.197 | 0.073 | 8.32E-83 | DF 5 | ccdc80l1 |
| wnt7bb1 | 5.1E-87 | 2.0984817 | 0.126 | 0.037 | 1.65E-82 | DF 5 | wnt7bb |
| btg22 | 1.6E-86 | 0.6898278 | 0.83 | 0.693 | 5.14E-82 | DF 5 | btg2 |
| pim22 | 2.4E-86 | 1.0165703 | 0.396 | 0.218 | 7.96E-82 | DF 5 | pim2 |
| tmem176l.4 | 3.4E-85 | 2.2692647 | 0.122 | 0.036 | 1.11E-80 | DF 5 | tmem176l.4 |
| grb10b1 | 6.9E-85 | 0.8671929 | 0.55 | 0.352 | 2.23E-80 | DF 5 | grb10b |
| rpl7a2 | 1.2E-84 | 0.2987887 | 0.998 | 0.991 | 3.89E-80 | DF 5 | rpl7a |
| socs1a3 | 8.9E-84 | 1.3594487 | 0.262 | 0.117 | 2.89E-79 | DF 5 | socs1a |
| si:ch211-196i2.12 | 2.4E-83 | 1.4997455 | 0.149 | 0.049 | 7.75E-79 | DF 5 | si:ch211-196i2.1 |
| ampd2b | 2.5E-83 | 2.0916753 | 0.105 | 0.028 | 8.12E-79 | DF 5 | ampd2b |
| arl5c2 | 2E-82 | 1.4052595 | 0.255 | 0.116 | 6.46E-78 | DF 5 | arl5c |
| si:dkey-151g10.63 | 4.6E-82 | 0.2549369 | 0.999 | 0.999 | 1.49E-77 | DF 5 | si:dkey-151g10.6 |
| junba2 | 2.1E-81 | 0.5278932 | 0.918 | 0.843 | 6.97E-77 | DF 5 | junba |
| hpdb5 | 4.3E-81 | 0.3715753 | 0.755 | 0.497 | 1.4E-76 | DF 5 | hpdb |
| si:ch1073-184j22.12 | 4.5E-80 | 1.4715478 | 0.192 | 0.075 | 1.46E-75 | DF 5 | si:ch1073-184j22.1 |
| rps27.13 | 5.4E-79 | 0.2958575 | 0.985 | 0.977 | 1.77E-74 | DF 5 | rps27.1 |
| krt913 | 5.5E-79 | 0.5460139 | 0.955 | 0.899 | 1.8E-74 | DF 5 | krt91 |
| rflnb2 | 3.4E-78 | 1.3004932 | 0.238 | 0.106 | 1.1E-73 | DF 5 | rflnb |

| | | | | | | | |
|---|---|---|---|---|---|---|---|
| ms4a17a.73 | 5.2E-78 | 0.9411723 | 0.475 | 0.289 | 1.71E-73 | DF 5 | ms4a17a.7 |
| foxd2 | 5.8E-78 | 1.5641492 | 0.134 | 0.043 | 1.89E-73 | DF 5 | foxd2 |
| tbrg1 | 6.1E-78 | 1.904816 | 0.137 | 0.045 | 1.98E-73 | DF 5 | tbrg1 |
| prickle1a1 | 1.3E-77 | 1.5356323 | 0.155 | 0.055 | 4.38E-73 | DF 5 | prickle1a |
| si:dkey-195m11.83 | 2.6E-77 | 1.1865719 | 0.262 | 0.121 | 8.42E-73 | DF 5 | si:dkey-195m11.8 |
| FQ323156.14 | 3.4E-77 | 0.8788406 | 0.361 | 0.188 | 1.11E-72 | DF 5 | FQ323156.1 |
| ifitm16 | 3.2E-76 | 0.6010614 | 0.908 | 0.785 | 1.05E-71 | DF 5 | ifitm1 |
| irs2b2 | 6.1E-76 | 0.9185573 | 0.396 | 0.223 | 1.99E-71 | DF 5 | irs2b |
| cited4b2 | 2.7E-75 | 1.1854754 | 0.29 | 0.144 | 8.71E-71 | DF 5 | cited4b |
| cfp4 | 2.9E-74 | 0.6604717 | 0.537 | 0.331 | 9.44E-70 | DF 5 | cfp |
| axin22 | 4.9E-74 | 0.9572349 | 0.382 | 0.213 | 1.6E-69 | DF 5 | axin2 |
| rpl222 | 5.6E-74 | 0.2974157 | 0.978 | 0.973 | 1.83E-69 | DF 5 | rpl22 |
| rpl10a2 | 8.8E-74 | 0.2951381 | 0.988 | 0.978 | 2.87E-69 | DF 5 | rpl10a |
| klf11b1 | 9.5E-74 | 0.9756151 | 0.398 | 0.229 | 3.09E-69 | DF 5 | klf11b |
| rpl133 | 2.1E-73 | 0.2529706 | 0.994 | 0.994 | 6.71E-69 | DF 5 | rpl13 |
| rpl283 | 2.7E-73 | 0.2600185 | 0.994 | 0.991 | 8.79E-69 | DF 5 | rpl28 |
| saa | 6.6E-73 | 1.5150005 | 0.131 | 0.044 | 2.14E-68 | DF 5 | saa |
| usp2a1 | 1.3E-72 | 1.552946 | 0.171 | 0.067 | 4.34E-68 | DF 5 | usp2a |
| scube31 | 1.4E-72 | 1.3285105 | 0.212 | 0.091 | 4.56E-68 | DF 5 | scube3 |
| ENSDARG000001152 | 1.6E-72 | 0.2464966 | 0.995 | 0.993 | 5.26E-68 | DF 5 | ENSDARG00000115 |
| sept9b1 | 3.5E-72 | 1.3731402 | 0.157 | 0.058 | 1.12E-67 | DF 5 | sept9b |
| nrn1a2 | 3.9E-71 | 1.2070722 | 0.201 | 0.084 | 1.27E-66 | DF 5 | nrn1a |
| rps27 | 7.4E-71 | 0.279163 | 0.997 | 0.99 | 2.4E-66 | DF 5 | rps2 |
| socs3b3 | 2.2E-70 | 0.7075213 | 0.642 | 0.464 | 7E-66 | DF 5 | socs3b |
| ubc1 | 2.3E-70 | 0.6676411 | 0.783 | 0.657 | 7.44E-66 | DF 5 | ubc |
| abhd15a2 | 3.5E-70 | 1.1339964 | 0.264 | 0.127 | 1.15E-65 | DF 5 | abhd15a |
| rpl13a2 | 9.8E-70 | 0.2743402 | 0.987 | 0.985 | 3.19E-65 | DF 5 | rpl13a |
| rpl72 | 1.2E-69 | 0.300162 | 0.986 | 0.972 | 3.74E-65 | DF 5 | rpl7 |
| wnt113 | 1.3E-69 | 0.665943 | 0.463 | 0.276 | 4.38E-65 | DF 5 | wnt11 |
| tob1a2 | 1.4E-69 | 0.6822108 | 0.746 | 0.603 | 4.47E-65 | DF 5 | tob1a |
| rps8a3 | 3.4E-69 | 0.255119 | 0.994 | 0.991 | 1.11E-64 | DF 5 | rps8a |
| midn2 | 6.3E-69 | 0.7864062 | 0.601 | 0.426 | 2.05E-64 | DF 5 | midn |
| runx33 | 8.9E-69 | 0.7608547 | 0.44 | 0.264 | 2.91E-64 | DF 5 | runx3 |
| ier2a3 | 9.3E-69 | 0.6492878 | 0.654 | 0.478 | 3.04E-64 | DF 5 | ier2a |
| ctsla3 | 1.7E-68 | 0.9566743 | 0.813 | 0.717 | 5.48E-64 | DF 5 | ctsla |
| mych3 | 1.8E-68 | 0.7409272 | 0.554 | 0.372 | 5.75E-64 | DF 5 | mych |
| rpl37 | 3.6E-68 | 0.2801861 | 0.99 | 0.977 | 1.17E-63 | DF 5 | rpl3 |
| serpine12 | 4E-68 | 0.514733 | 0.868 | 0.757 | 1.29E-63 | DF 5 | serpine1 |
| il6st3 | 5.2E-68 | 0.817349 | 0.586 | 0.416 | 1.69E-63 | DF 5 | il6st |
| pcbd14 | 8.8E-68 | 0.622341 | 0.502 | 0.308 | 2.85E-63 | DF 5 | pcbd1 |
| col18a1a2 | 1.5E-67 | 0.7939054 | 0.291 | 0.147 | 4.88E-63 | DF 5 | col18a1a |
| osr13 | 2.2E-67 | 0.8702318 | 0.319 | 0.166 | 7.19E-63 | DF 5 | osr1 |
| cxcl8a2 | 4.1E-67 | 0.79305 | 0.431 | 0.258 | 1.33E-62 | DF 5 | cxcl8a |
| timp4.32 | 4.6E-67 | 0.2553139 | 0.655 | 0.43 | 1.51E-62 | DF 5 | timp4.3 |
| pcdh113 | 6E-67 | 1.1987309 | 0.177 | 0.071 | 1.96E-62 | DF 5 | pcdh11 |
| ivns1abpa3 | 6.5E-67 | 0.8989489 | 0.448 | 0.279 | 2.11E-62 | DF 5 | ivns1abpa |
| slmapb | 1.4E-66 | 1.4235011 | 0.132 | 0.046 | 4.68E-62 | DF 5 | slmapb |
| rps93 | 1.7E-66 | 0.2553149 | 0.988 | 0.987 | 5.49E-62 | DF 5 | rps9 |

| | | | | | | | |
|---|---|---|---|---|---|---|---|
| uba522 | 3E-66 | 0.2283239 | 0.992 | 0.992 | 9.84E-62 | DF 5 | uba52 |
| rpl37.11 | 3.8E-66 | 0.2611052 | 0.989 | 0.991 | 1.25E-61 | DF 5 | rpl37.1 |
| ENSDARG00000075[illegible] | 7.3E-66 | 1.1589932 | 0.19 | 0.08 | 2.37E-61 | DF 5 | ENSDARG00000075 |
| prrx1b3 | 2.6E-65 | 0.6503191 | 0.561 | 0.371 | 8.48E-61 | DF 5 | prrx1b |
| rps27.2 | 2.7E-65 | 0.3039262 | 0.974 | 0.952 | 8.65E-61 | DF 5 | rps27.2 |
| cd99l22 | 3.1E-65 | 0.9496959 | 0.355 | 0.201 | 1E-60 | DF 5 | cd99l2 |
| rpl122 | 9E-65 | 0.2571785 | 0.989 | 0.982 | 2.94E-60 | DF 5 | rpl12 |
| rps182 | 3.4E-64 | 0.3028537 | 0.976 | 0.963 | 1.11E-59 | DF 5 | rps18 |
| rpl352 | 3.5E-64 | 0.2407769 | 0.993 | 0.992 | 1.14E-59 | DF 5 | rpl35 |
| raraa4 | 4.4E-64 | 0.7971647 | 0.413 | 0.246 | 1.43E-59 | DF 5 | raraa |
| ptk7a3 | 1.3E-63 | 0.7291961 | 0.481 | 0.307 | 4.08E-59 | DF 5 | ptk7a |
| eef1da2 | 1.5E-63 | 0.3408595 | 0.938 | 0.873 | 4.8E-59 | DF 5 | eef1da |
| hs3st3b1b1 | 2.7E-63 | 1.1433589 | 0.251 | 0.124 | 8.72E-59 | DF 5 | hs3st3b1b |
| slc7a24 | 4.2E-63 | 0.8130591 | 0.298 | 0.153 | 1.37E-58 | DF 5 | slc7a2 |
| LO018188.14 | 7.6E-63 | 0.6323793 | 0.649 | 0.47 | 2.46E-58 | DF 5 | LO018188.1 |
| f3b4 | 1.7E-62 | 1.0840221 | 0.535 | 0.373 | 5.41E-58 | DF 5 | f3b |
| postnb2 | 1.7E-62 | 0.5227328 | 0.625 | 0.448 | 5.5E-58 | DF 5 | postnb |
| atf4b2 | 1.8E-62 | 0.5519039 | 0.757 | 0.629 | 5.86E-58 | DF 5 | atf4b |
| dop1a1 | 2E-62 | 1.4150909 | 0.173 | 0.072 | 6.52E-58 | DF 5 | dop1a |
| rpl322 | 3.6E-62 | 0.2393349 | 0.99 | 0.989 | 1.16E-57 | DF 5 | rpl32 |
| col12a1a2 | 8.9E-62 | 0.4094903 | 0.905 | 0.745 | 2.9E-57 | DF 5 | col12a1a |
| fabp34 | 5.5E-61 | 0.5688997 | 0.68 | 0.524 | 1.78E-56 | DF 5 | fabp3 |
| rps15a3 | 1.2E-60 | 0.2326262 | 0.991 | 0.99 | 3.86E-56 | DF 5 | rps15a |
| ZFP36L1.13 | 5.6E-60 | 0.4576497 | 0.932 | 0.871 | 1.81E-55 | DF 5 | ZFP36L1.1 |
| fthl272 | 5.7E-60 | 0.8230535 | 0.901 | 0.837 | 1.87E-55 | DF 5 | fthl27 |
| nr1d2a2 | 7.5E-60 | 0.8713439 | 0.398 | 0.245 | 2.45E-55 | DF 5 | nr1d2a |
| ptp4a2b1 | 1.5E-59 | 0.8462521 | 0.455 | 0.297 | 4.97E-55 | DF 5 | ptp4a2b |
| nkd11 | 1.6E-59 | 1.2061806 | 0.182 | 0.079 | 5.21E-55 | DF 5 | nkd1 |
| plk33 | 3.3E-59 | 0.9322961 | 0.34 | 0.196 | 1.07E-54 | DF 5 | plk3 |
| zswim52 | 5.5E-59 | 0.95099 | 0.313 | 0.174 | 1.78E-54 | DF 5 | zswim5 |
| lrrc58b2 | 1.3E-58 | 1.1146223 | 0.237 | 0.117 | 4.24E-54 | DF 5 | lrrc58b |
| hspa81 | 1.8E-58 | 0.3139834 | 0.969 | 0.935 | 5.99E-54 | DF 5 | hspa8 |
| naca2 | 2.5E-58 | 0.2818689 | 0.976 | 0.961 | 8.23E-54 | DF 5 | naca |
| twist1a2 | 2.7E-58 | 0.5493468 | 0.604 | 0.431 | 8.68E-54 | DF 5 | twist1a |
| rpl36a3 | 3.5E-58 | 0.2156929 | 0.995 | 0.994 | 1.13E-53 | DF 5 | rpl36a |
| ip6k2b3 | 3.5E-58 | 0.7509344 | 0.495 | 0.337 | 1.14E-53 | DF 5 | ip6k2b |
| rps73 | 4.4E-58 | 0.2260812 | 0.995 | 0.991 | 1.44E-53 | DF 5 | rps7 |
| rps162 | 1.6E-57 | 0.2388699 | 0.994 | 0.989 | 5.13E-53 | DF 5 | rps16 |
| ctnnb24 | 4.5E-57 | 0.7600628 | 0.461 | 0.305 | 1.47E-52 | DF 5 | ctnnb2 |
| id2a3 | 7.6E-57 | 0.650782 | 0.658 | 0.496 | 2.46E-52 | DF 5 | id2a |
| postna6 | 7.8E-57 | 0.2846033 | 0.797 | 0.581 | 2.55E-52 | DF 5 | postna |
| usp54a1 | 1.5E-56 | 1.5571538 | 0.138 | 0.054 | 4.83E-52 | DF 5 | usp54a |
| gadd45ab4 | 1.6E-56 | 1.0912046 | 0.302 | 0.171 | 5.27E-52 | DF 5 | gadd45ab |
| zgc:1748882 | 8.4E-56 | 0.9634277 | 0.355 | 0.211 | 2.72E-51 | DF 5 | zgc:174888 |
| spry42 | 3.9E-55 | 1.0459105 | 0.303 | 0.173 | 1.27E-50 | DF 5 | spry4 |
| bzw1b1 | 4.3E-55 | 0.7095806 | 0.63 | 0.485 | 1.39E-50 | DF 5 | bzw1b |
| slc44a1b3 | 4.9E-55 | 1.0241243 | 0.231 | 0.115 | 1.61E-50 | DF 5 | slc44a1b |
| jun2 | 7E-55 | 0.4427249 | 0.916 | 0.851 | 2.27E-50 | DF 5 | jun |

| | | | | | | | |
|---|---|---|---|---|---|---|---|
| rps142 | 8E-55 | 0.2103001 | 0.996 | 0.993 | 2.6E-50 | DF 5 | rps14 |
| col27a1b3 | 8.1E-55 | 0.6685626 | 0.499 | 0.33 | 2.63E-50 | DF 5 | col27a1b |
| gstr2 | 8.7E-55 | 0.5384954 | 0.579 | 0.409 | 2.84E-50 | DF 5 | gstr |
| rplp21 | 9.4E-55 | 0.2218918 | 0.985 | 0.988 | 3.06E-50 | DF 5 | rplp2 |
| rps112 | 1.3E-54 | 0.2173979 | 0.995 | 0.989 | 4.27E-50 | DF 5 | rps11 |
| rps3a2 | 2.3E-54 | 0.2392322 | 0.985 | 0.981 | 7.55E-50 | DF 5 | rps3a |
| gnai2b2 | 7.1E-54 | 1.3880952 | 0.145 | 0.06 | 2.31E-49 | DF 5 | gnai2b |
| rpl63 | 7.6E-54 | 0.2804072 | 0.979 | 0.958 | 2.49E-49 | DF 5 | rpl6 |
| neo1b1 | 2.1E-53 | 1.254473 | 0.172 | 0.077 | 6.85E-49 | DF 5 | neo1b |
| ier51 | 2.4E-53 | 0.5485132 | 0.622 | 0.463 | 7.79E-49 | DF 5 | ier5 |
| btg12 | 3.3E-53 | 0.3869394 | 0.898 | 0.811 | 1.08E-48 | DF 5 | btg1 |
| fstl33 | 3.4E-53 | 0.6764393 | 0.471 | 0.315 | 1.12E-48 | DF 5 | fstl3 |
| hpgd1 | 5.4E-53 | 1.3933706 | 0.133 | 0.052 | 1.77E-48 | DF 5 | hpgd |
| rps63 | 1.3E-52 | 0.2531507 | 0.978 | 0.96 | 4.28E-48 | DF 5 | rps6 |
| rpl183 | 1.5E-52 | 0.2286508 | 0.988 | 0.981 | 4.78E-48 | DF 5 | rpl18 |
| inka1b2 | 1.6E-52 | 1.0825717 | 0.188 | 0.088 | 5.05E-48 | DF 5 | inka1b |
| slc25a484 | 1.7E-52 | 1.122094 | 0.251 | 0.133 | 5.58E-48 | DF 5 | slc25a48 |
| prrx1a3 | 4.7E-52 | 0.5349608 | 0.722 | 0.58 | 1.52E-47 | DF 5 | prrx1a |
| rpl392 | 5.8E-52 | 0.2208396 | 0.988 | 0.984 | 1.87E-47 | DF 5 | rpl39 |
| ccnl1a2 | 6.8E-52 | 0.8177313 | 0.4 | 0.256 | 2.2E-47 | DF 5 | ccnl1a |
| nanos12 | 8E-52 | 0.8558035 | 0.335 | 0.199 | 2.59E-47 | DF 5 | nanos1 |
| cspg5a2 | 8.9E-52 | 1.2187487 | 0.166 | 0.074 | 2.88E-47 | DF 5 | cspg5a |
| si:dkey-85k7.73 | 1.6E-51 | 0.547682 | 0.438 | 0.274 | 5.2E-47 | DF 5 | si:dkey-85k7.7 |
| rpl342 | 1.6E-51 | 0.2218171 | 0.99 | 0.985 | 5.35E-47 | DF 5 | rpl34 |
| ier5l2 | 3E-51 | 0.6126808 | 0.469 | 0.313 | 9.79E-47 | DF 5 | ier5l |
| rpl173 | 3.3E-51 | 0.2180192 | 0.994 | 0.989 | 1.06E-46 | DF 5 | rpl17 |
| alpl1 | 5.3E-51 | 1.1577871 | 0.151 | 0.064 | 1.71E-46 | DF 5 | alpl |
| slc25a52 | 7.1E-51 | 0.2559352 | 0.981 | 0.962 | 2.3E-46 | DF 5 | slc25a5 |
| ier2b2 | 8.1E-51 | 0.4698015 | 0.778 | 0.644 | 2.64E-46 | DF 5 | ier2b |
| btf33 | 1.8E-50 | 0.3455661 | 0.924 | 0.873 | 5.92E-46 | DF 5 | btf3 |
| serbp1a2 | 5.5E-50 | 0.2963162 | 0.944 | 0.907 | 1.77E-45 | DF 5 | serbp1a |
| enpp13 | 1.8E-49 | 0.81566 | 0.312 | 0.181 | 5.71E-45 | DF 5 | enpp1 |
| rpl35a3 | 1.9E-49 | 0.2006235 | 0.988 | 0.989 | 6.29E-45 | DF 5 | rpl35a |
| zfhx33 | 2E-49 | 0.513321 | 0.638 | 0.48 | 6.65E-45 | DF 5 | zfhx3 |
| rack13 | 2.2E-49 | 0.2612894 | 0.97 | 0.947 | 7.19E-45 | DF 5 | rack1 |
| abcc6a2 | 2.7E-49 | 1.0822063 | 0.162 | 0.072 | 8.7E-45 | DF 5 | abcc6a |
| rpl5a3 | 2.9E-49 | 0.2602229 | 0.954 | 0.917 | 9.48E-45 | DF 5 | rpl5a |
| zgc:1133071 | 6.4E-49 | 0.9319552 | 0.187 | 0.089 | 2.1E-44 | DF 5 | zgc:113307 |
| id2b3 | 1E-48 | 0.9471537 | 0.312 | 0.185 | 3.24E-44 | DF 5 | id2b |
| C25H11orf963 | 2E-48 | 0.5980072 | 0.538 | 0.377 | 6.65E-44 | DF 5 | C25H11orf96 |
| fosb3 | 2.2E-48 | 0.475145 | 0.813 | 0.687 | 7.06E-44 | DF 5 | fosb |
| rpl243 | 3.5E-48 | 0.2014129 | 0.996 | 0.993 | 1.14E-43 | DF 5 | rpl24 |
| lef12 | 1.4E-47 | 0.9022605 | 0.208 | 0.104 | 4.5E-43 | DF 5 | lef1 |
| cpn14 | 1.8E-47 | 0.6802277 | 0.344 | 0.206 | 5.83E-43 | DF 5 | cpn1 |
| rps33 | 4.4E-47 | 0.257204 | 0.963 | 0.946 | 1.43E-42 | DF 5 | rps3 |
| enpp22 | 4.4E-47 | 0.465475 | 0.599 | 0.438 | 1.43E-42 | DF 5 | enpp2 |
| rps4x3 | 4.8E-47 | 0.2324861 | 0.983 | 0.966 | 1.56E-42 | DF 5 | rps4x |
| dgkza | 5.7E-47 | 1.2061076 | 0.131 | 0.054 | 1.85E-42 | DF 5 | dgkza |

| | | | | | | | |
|---|---|---|---|---|---|---|---|
| cirbpb2 | 7.2E-47 | 0.3206951 | 0.9 | 0.831 | 2.35E-42 | DF 5 | cirbpb |
| si:ch211-207e14.41 | 8.3E-47 | 1.006861 | 0.149 | 0.065 | 2.69E-42 | DF 5 | si:ch211-207e14.4 |
| adgrl2a4 | 1.3E-46 | 0.6549144 | 0.318 | 0.185 | 4.21E-42 | DF 5 | adgrl2a |
| mvp4 | 2.5E-46 | 0.7272984 | 0.451 | 0.311 | 8.05E-42 | DF 5 | mvp |
| brd2a2 | 2.8E-46 | 0.6314929 | 0.583 | 0.441 | 9.21E-42 | DF 5 | brd2a |
| rgl1 | 3.4E-46 | 1.065434 | 0.225 | 0.12 | 1.11E-41 | DF 5 | rgl1 |
| rpl273 | 4.4E-46 | 0.2160458 | 0.984 | 0.978 | 1.45E-41 | DF 5 | rpl27 |
| gch13 | 7.4E-46 | 0.7061281 | 0.246 | 0.131 | 2.4E-41 | DF 5 | gch1 |
| egr13 | 1.7E-45 | 0.507924 | 0.748 | 0.623 | 5.39E-41 | DF 5 | egr1 |
| rps26l3 | 3.5E-45 | 0.2163248 | 0.98 | 0.974 | 1.14E-40 | DF 5 | rps26l |
| si:ch211-170d8.83 | 4.1E-45 | 0.493053 | 0.633 | 0.478 | 1.34E-40 | DF 5 | si:ch211-170d8.8 |
| higd1a2 | 5.9E-45 | 0.5760708 | 0.591 | 0.452 | 1.91E-40 | DF 5 | higd1a |
| iqsec1b | 5.9E-45 | 0.6093284 | 0.484 | 0.338 | 1.92E-40 | DF 5 | iqsec1b |
| metrn2 | 8.2E-45 | 0.8230782 | 0.218 | 0.112 | 2.66E-40 | DF 5 | metrn |
| rpl362 | 1.2E-44 | 0.2126685 | 0.982 | 0.986 | 3.85E-40 | DF 5 | rpl36 |
| mid1ip1l1 | 2E-44 | 0.8305705 | 0.307 | 0.185 | 6.4E-40 | DF 5 | mid1ip1l |
| ddit33 | 3.6E-44 | 0.4681853 | 0.703 | 0.577 | 1.16E-39 | DF 5 | ddit3 |
| rpl83 | 3.9E-44 | 0.2193033 | 0.985 | 0.978 | 1.27E-39 | DF 5 | rpl8 |
| tob1b3 | 4.3E-44 | 0.3720598 | 0.812 | 0.698 | 1.41E-39 | DF 5 | tob1b |
| mknk2b2 | 5.7E-44 | 0.4957689 | 0.73 | 0.602 | 1.86E-39 | DF 5 | mknk2b |
| faua2 | 6.2E-44 | 0.2088134 | 0.982 | 0.974 | 2.02E-39 | DF 5 | faua |
| enpp61 | 6.5E-44 | 1.1813519 | 0.127 | 0.054 | 2.11E-39 | DF 5 | enpp6 |
| smad3a2 | 1.7E-43 | 0.7917329 | 0.432 | 0.3 | 5.56E-39 | DF 5 | smad3a |
| rpl103 | 2E-43 | 0.2196594 | 0.98 | 0.968 | 6.58E-39 | DF 5 | rpl10 |
| hnrnpa0l.12 | 2.6E-43 | 0.4148645 | 0.77 | 0.663 | 8.49E-39 | DF 5 | hnrnpa0l.1 |
| csf1b1 | 4.2E-43 | 1.2439086 | 0.103 | 0.04 | 1.36E-38 | DF 5 | csf1b |
| hyal2b4 | 7E-43 | 0.6812744 | 0.319 | 0.192 | 2.26E-38 | DF 5 | hyal2b |
| zgc:1107963 | 1.2E-42 | 1.2921558 | 0.16 | 0.077 | 3.97E-38 | DF 5 | zgc:110796 |
| cx433 | 1.2E-42 | 0.7410928 | 0.281 | 0.165 | 4.01E-38 | DF 5 | cx43 |
| bmp1a2 | 1.9E-42 | 0.8528428 | 0.301 | 0.182 | 6.25E-38 | DF 5 | bmp1a |
| BX663503.11 | 1.8E-41 | 0.9340357 | 0.141 | 0.063 | 5.94E-37 | DF 5 | BX663503.1 |
| c44 | 2.5E-41 | 0.4205305 | 0.332 | 0.202 | 8.23E-37 | DF 5 | c4 |
| si:ch211-178n15.11 | 2.6E-41 | 0.907847 | 0.275 | 0.164 | 8.38E-37 | DF 5 | si:ch211-178n15.1 |
| slc25a55a2 | 2.8E-41 | 0.8773636 | 0.309 | 0.193 | 8.96E-37 | DF 5 | slc25a55a |
| brdt1 | 4.8E-41 | 0.7334535 | 0.435 | 0.306 | 1.57E-36 | DF 5 | brdt |
| hspa4a2 | 8.7E-41 | 0.9124814 | 0.251 | 0.145 | 2.84E-36 | DF 5 | hspa4a |
| mdh1aa2 | 2E-40 | 0.5639165 | 0.502 | 0.368 | 6.66E-36 | DF 5 | mdh1aa |
| arrdc24 | 3E-40 | 0.5792128 | 0.371 | 0.237 | 9.62E-36 | DF 5 | arrdc2 |
| tgfbi2 | 3.2E-40 | 0.4600724 | 0.321 | 0.195 | 1.03E-35 | DF 5 | tgfbi |
| rgs5b5 | 3.8E-40 | 0.5992572 | 0.641 | 0.518 | 1.24E-35 | DF 5 | rgs5b |
| DAB24 | 5E-40 | 0.5110696 | 0.512 | 0.372 | 1.63E-35 | DF 5 | DAB2 |
| ddx53 | 6.5E-40 | 0.5216506 | 0.703 | 0.593 | 2.12E-35 | DF 5 | ddx5 |
| rcan2 | 1E-39 | 1.0441621 | 0.127 | 0.056 | 3.25E-35 | DF 5 | rcan2 |
| trim711 | 1.2E-39 | 1.0178113 | 0.189 | 0.099 | 3.93E-35 | DF 5 | trim71 |
| trib31 | 1.4E-39 | 0.851393 | 0.281 | 0.171 | 4.68E-35 | DF 5 | trib3 |
| SBNO22 | 1.6E-39 | 0.7380051 | 0.345 | 0.223 | 5.12E-35 | DF 5 | SBNO2 |
| rps53 | 1.6E-39 | 0.2116445 | 0.986 | 0.972 | 5.2E-35 | DF 5 | rps5 |
| mex3b2 | 1.9E-39 | 0.8231606 | 0.278 | 0.167 | 6.31E-35 | DF 5 | mex3b |

| | | | | | | | |
|---|---|---|---|---|---|---|---|
| ephb2b2 | 7.1E-39 | 0.6931648 | 0.313 | 0.194 | 2.3E-34 | DF 5 | ephb2b |
| meis2a.12 | 1E-38 | 0.750437 | 0.232 | 0.129 | 3.34E-34 | DF 5 | meis2a.1 |
| slc25a332 | 1.8E-38 | 0.9415089 | 0.207 | 0.114 | 5.76E-34 | DF 5 | slc25a33 |
| rplp03 | 2.1E-38 | 0.208173 | 0.988 | 0.982 | 6.76E-34 | DF 5 | rplp0 |
| rpl5b2 | 5.4E-38 | 0.2604217 | 0.904 | 0.848 | 1.74E-33 | DF 5 | rpl5b |
| spry22 | 6.6E-38 | 1.0513885 | 0.154 | 0.075 | 2.15E-33 | DF 5 | spry2 |
| pou3f3b | 1.7E-37 | 1.0149587 | 0.121 | 0.053 | 5.56E-33 | DF 5 | pou3f3b |
| rasl11b4 | 2.9E-37 | 0.6744239 | 0.33 | 0.212 | 9.43E-33 | DF 5 | rasl11b |
| rpl153 | 3.2E-37 | 0.2038559 | 0.981 | 0.97 | 1.03E-32 | DF 5 | rpl15 |
| nt5c3a1 | 1.6E-36 | 0.8494926 | 0.271 | 0.165 | 5.13E-32 | DF 5 | nt5c3a |
| pitpnc1b2 | 2E-36 | 0.9577331 | 0.157 | 0.078 | 6.34E-32 | DF 5 | pitpnc1b |
| tbx2a | 2.3E-36 | 1.1360037 | 0.128 | 0.059 | 7.63E-32 | DF 5 | tbx2a |
| serpine22 | 4.6E-36 | 0.70732 | 0.434 | 0.311 | 1.49E-31 | DF 5 | serpine2 |
| etv5b2 | 4.8E-36 | 1.0084932 | 0.182 | 0.097 | 1.58E-31 | DF 5 | etv5b |
| h3f3d2 | 5.5E-36 | 0.3345839 | 0.82 | 0.724 | 1.79E-31 | DF 5 | h3f3d |
| scxa3 | 6E-36 | 0.914571 | 0.159 | 0.08 | 1.96E-31 | DF 5 | scxa |
| eef2b3 | 7.7E-36 | 0.2584286 | 0.954 | 0.907 | 2.5E-31 | DF 5 | eef2b |
| neu3.24 | 1E-35 | 0.5532359 | 0.342 | 0.217 | 3.3E-31 | DF 5 | neu3.2 |
| skib4 | 1.9E-35 | 0.8725784 | 0.304 | 0.198 | 6.32E-31 | DF 5 | skib |
| heyl3 | 4.4E-35 | 0.9679144 | 0.143 | 0.069 | 1.42E-30 | DF 5 | heyl |
| ccdc80l21 | 5.4E-35 | 0.4411819 | 0.205 | 0.11 | 1.76E-30 | DF 5 | ccdc80l2 |
| hmox1a3 | 1E-34 | 1.0222453 | 0.271 | 0.17 | 3.37E-30 | DF 5 | hmox1a |
| fpr12 | 1.1E-34 | 0.4710982 | 0.334 | 0.213 | 3.49E-30 | DF 5 | fpr1 |
| igf1rb1 | 1.1E-34 | 0.6948429 | 0.383 | 0.266 | 3.62E-30 | DF 5 | igf1rb |
| rsl24d13 | 2.2E-34 | 0.3049097 | 0.835 | 0.745 | 7.3E-30 | DF 5 | rsl24d1 |
| fzd10 | 3E-34 | 1.0994039 | 0.1 | 0.042 | 9.84E-30 | DF 5 | fzd10 |
| slc25a25a3 | 4.5E-34 | 0.6704123 | 0.326 | 0.214 | 1.47E-29 | DF 5 | slc25a25a |
| rpl43 | 5.1E-34 | 0.2332954 | 0.94 | 0.894 | 1.67E-29 | DF 5 | rpl4 |
| nabp1a2 | 5.1E-34 | 0.8580705 | 0.278 | 0.176 | 1.67E-29 | DF 5 | nabp1a |
| AL929435.12 | 9.3E-34 | 0.9893374 | 0.187 | 0.103 | 3.01E-29 | DF 5 | AL929435.1 |
| BX908782.21 | 1.8E-33 | 3.16738 | 0.115 | 0.054 | 5.93E-29 | DF 5 | BX908782.2 |
| spink2.11 | 2E-33 | 0.8005415 | 0.217 | 0.125 | 6.6E-29 | DF 5 | spink2.1 |
| unc5c | 2.5E-33 | 1.2110405 | 0.117 | 0.054 | 8.01E-29 | DF 5 | unc5c |
| tent5ab1 | 3.2E-33 | 1.2570269 | 0.12 | 0.056 | 1.03E-28 | DF 5 | tent5ab |
| ccn13 | 3.3E-33 | 0.4988386 | 0.752 | 0.638 | 1.08E-28 | DF 5 | ccn1 |
| ip6k2a2 | 5.1E-33 | 0.6911976 | 0.3 | 0.193 | 1.67E-28 | DF 5 | ip6k2a |
| nfe2l2a4 | 8.5E-33 | 0.4296859 | 0.477 | 0.338 | 2.76E-28 | DF 5 | nfe2l2a |
| dusp11 | 1.2E-32 | 0.5119692 | 0.505 | 0.375 | 4.06E-28 | DF 5 | dusp1 |
| cdo13 | 1.6E-32 | 0.5424695 | 0.447 | 0.324 | 5.29E-28 | DF 5 | cdo1 |
| adgre102 | 2.2E-32 | 0.8950249 | 0.145 | 0.073 | 7.13E-28 | DF 5 | adgre10 |
| jund3 | 2.3E-32 | 0.4337565 | 0.684 | 0.565 | 7.57E-28 | DF 5 | jund |
| hgd4 | 2.9E-32 | 0.244086 | 0.581 | 0.423 | 9.41E-28 | DF 5 | hgd |
| fbp1a2 | 7.6E-32 | 0.5798025 | 0.389 | 0.272 | 2.47E-27 | DF 5 | fbp1a |
| fgfr1b2 | 1.2E-31 | 0.8764103 | 0.179 | 0.098 | 4.04E-27 | DF 5 | fgfr1b |
| bcl6ab | 1.8E-31 | 1.0336252 | 0.122 | 0.059 | 5.91E-27 | DF 5 | bcl6ab |
| C18H15orf484 | 3.6E-31 | 0.3176711 | 0.447 | 0.304 | 1.16E-26 | DF 5 | C18H15orf48 |
| si:ch211-136a13.12 | 4.3E-31 | 1.0113409 | 0.156 | 0.083 | 1.38E-26 | DF 5 | si:ch211-136a13.1 |
| FQ323119.12 | 4.6E-31 | 0.7672696 | 0.182 | 0.1 | 1.49E-26 | DF 5 | FQ323119.1 |

| | | | | | | | |
|---|---|---|---|---|---|---|---|
| pptc7a1 | 5E-31 | 0.6100013 | 0.329 | 0.22 | 1.63E-26 | DF 5 | pptc7a |
| dusp62 | 8.8E-31 | 0.8065389 | 0.235 | 0.143 | 2.87E-26 | DF 5 | dusp6 |
| snai1a1 | 9.1E-31 | 0.6053541 | 0.382 | 0.268 | 2.95E-26 | DF 5 | snai1a |
| fth1a1 | 1.4E-30 | 0.3341488 | 0.796 | 0.715 | 4.39E-26 | DF 5 | fth1a |
| nfkbiab3 | 1.7E-30 | 0.4879677 | 0.682 | 0.579 | 5.5E-26 | DF 5 | nfkbiab |
| fbrsl1 | 1.8E-30 | 1.2517391 | 0.11 | 0.051 | 6.01E-26 | DF 5 | fbrsl1 |
| pdk2b2 | 2.1E-30 | 0.595703 | 0.3 | 0.195 | 6.79E-26 | DF 5 | pdk2b |
| mocs23 | 3E-30 | 0.8779648 | 0.158 | 0.085 | 9.85E-26 | DF 5 | mocs2 |
| marveld11 | 1.8E-29 | 0.6219786 | 0.31 | 0.206 | 5.89E-25 | DF 5 | marveld1 |
| nr2f1a2 | 2.8E-29 | 0.6722682 | 0.242 | 0.149 | 9.2E-25 | DF 5 | nr2f1a |
| si:ch1073-291c23.24 | 3.1E-29 | 0.3242518 | 0.609 | 0.469 | 1.01E-24 | DF 5 | si:ch1073-291c23.2 |
| egfl62 | 3.5E-29 | 0.836571 | 0.112 | 0.053 | 1.13E-24 | DF 5 | egfl6 |
| dusp53 | 4.1E-29 | 0.6384479 | 0.295 | 0.194 | 1.34E-24 | DF 5 | dusp5 |
| tmc6b1 | 6.9E-29 | 0.7335185 | 0.137 | 0.07 | 2.25E-24 | DF 5 | tmc6b |
| tmem198b2 | 9.2E-29 | 0.7382369 | 0.282 | 0.186 | 3.01E-24 | DF 5 | tmem198b |
| trps13 | 1.5E-28 | 0.7095219 | 0.266 | 0.17 | 4.85E-24 | DF 5 | trps1 |
| rpe65b2 | 2.2E-28 | 1.0587978 | 0.131 | 0.067 | 7.16E-24 | DF 5 | rpe65b |
| CABZ01068358.12 | 2.4E-28 | 0.6166975 | 0.307 | 0.204 | 7.81E-24 | DF 5 | CABZ01068358.1 |
| myo10l32 | 3.6E-28 | 0.9391182 | 0.17 | 0.097 | 1.16E-23 | DF 5 | myo10l3 |
| tspan353 | 4.3E-28 | 0.6613871 | 0.205 | 0.12 | 1.41E-23 | DF 5 | tspan35 |
| mylka2 | 4.4E-28 | 0.6818076 | 0.182 | 0.105 | 1.42E-23 | DF 5 | mylka |
| spred2b | 6E-28 | 0.9079848 | 0.181 | 0.104 | 1.96E-23 | DF 5 | spred2b |
| si:dkey-164f24.25 | 6.1E-28 | 0.4886303 | 0.265 | 0.166 | 1.98E-23 | DF 5 | si:dkey-164f24.2 |
| eif3f2 | 7.5E-28 | 0.2798632 | 0.812 | 0.725 | 2.43E-23 | DF 5 | eif3f |
| cbx7a2 | 2.3E-27 | 0.4675222 | 0.46 | 0.342 | 7.58E-23 | DF 5 | cbx7a |
| si:ch211-232b12.51 | 3.3E-27 | 0.6652383 | 0.242 | 0.153 | 1.08E-22 | DF 5 | si:ch211-232b12.5 |
| kctd12.22 | 4.3E-27 | 0.6327864 | 0.321 | 0.222 | 1.4E-22 | DF 5 | kctd12.2 |
| snai1b2 | 6.4E-27 | 0.6697361 | 0.187 | 0.109 | 2.09E-22 | DF 5 | snai1b |
| cebpd4 | 6.5E-27 | 0.3112591 | 0.696 | 0.572 | 2.13E-22 | DF 5 | cebpd |
| barx13 | 7.1E-27 | 0.5190754 | 0.21 | 0.125 | 2.3E-22 | DF 5 | barx1 |
| mepce1 | 7.9E-27 | 0.8283888 | 0.175 | 0.101 | 2.57E-22 | DF 5 | mepce |
| hist2h2l3 | 7.9E-27 | 0.5568658 | 0.48 | 0.37 | 2.57E-22 | DF 5 | hist2h2l |
| smad71 | 1.8E-26 | 0.73743 | 0.236 | 0.15 | 5.83E-22 | DF 5 | smad7 |
| lxn3 | 2.2E-26 | 0.2064891 | 0.818 | 0.696 | 7.2E-22 | DF 5 | lxn |
| ponzr14 | 2.9E-26 | 0.6408497 | 0.333 | 0.23 | 9.44E-22 | DF 5 | ponzr1 |
| gadd45ga3 | 3.3E-26 | 0.5051151 | 0.378 | 0.273 | 1.06E-21 | DF 5 | gadd45ga |
| slc16a104 | 5.4E-26 | 0.6342076 | 0.212 | 0.13 | 1.74E-21 | DF 5 | slc16a10 |
| tuft1a2 | 5.7E-26 | 0.2556849 | 0.708 | 0.586 | 1.87E-21 | DF 5 | tuft1a |
| cx43.43 | 6.5E-26 | 0.5812719 | 0.316 | 0.218 | 2.1E-21 | DF 5 | cx43.4 |
| nfil3-64 | 6.7E-26 | 0.3074375 | 0.424 | 0.301 | 2.17E-21 | DF 5 | nfil3-6 |
| gchfr3 | 8.9E-26 | 0.6281246 | 0.248 | 0.159 | 2.88E-21 | DF 5 | gchfr |
| si:dkey-226m8.103 | 1.1E-25 | 0.7154321 | 0.169 | 0.097 | 3.44E-21 | DF 5 | si:dkey-226m8.10 |
| tnrc6a3 | 1.1E-25 | 0.5923791 | 0.372 | 0.268 | 3.53E-21 | DF 5 | tnrc6a |
| h3f3b.12 | 1.7E-25 | 0.3574088 | 0.598 | 0.491 | 5.44E-21 | DF 5 | h3f3b.1 |
| tat4 | 2.7E-25 | 0.2681195 | 0.354 | 0.24 | 8.86E-21 | DF 5 | tat |
| foxo1a | 4E-25 | 0.6873095 | 0.275 | 0.186 | 1.29E-20 | DF 5 | foxo1a |
| kremen12 | 5.2E-25 | 0.8485844 | 0.142 | 0.078 | 1.68E-20 | DF 5 | kremen1 |
| si:rp71-79p20.21 | 5.5E-25 | 0.6644958 | 0.234 | 0.149 | 1.78E-20 | DF 5 | si:rp71-79p20.2 |

| | | | | | | | |
|---|---|---|---|---|---|---|---|
| gng101 | 5.9E-25 | 0.9472696 | 0.119 | 0.062 | 1.91E-20 | DF 5 | gng10 |
| CU927890.11 | 9.2E-25 | 0.8665828 | 0.113 | 0.057 | 2.98E-20 | DF 5 | CU927890.1 |
| tuba1b4 | 1E-24 | 0.4732154 | 0.411 | 0.305 | 3.41E-20 | DF 5 | tuba1b |
| yap11 | 1.2E-24 | 0.4427173 | 0.467 | 0.355 | 3.88E-20 | DF 5 | yap1 |
| cbsb4 | 1.8E-24 | 0.5550545 | 0.218 | 0.136 | 5.84E-20 | DF 5 | cbsb |
| tln2a1 | 2.8E-24 | 0.868416 | 0.137 | 0.075 | 9.01E-20 | DF 5 | tln2a |
| zbtb181 | 3.1E-24 | 0.9867721 | 0.111 | 0.057 | 9.93E-20 | DF 5 | zbtb18 |
| si:dkey-16p21.84 | 3.4E-24 | 0.4434229 | 0.306 | 0.205 | 1.09E-19 | DF 5 | si:dkey-16p21.8 |
| krt18b2 | 4.9E-24 | 0.2346039 | 0.942 | 0.889 | 1.58E-19 | DF 5 | krt18b |
| glrx1 | 6.9E-24 | 0.4269897 | 0.459 | 0.353 | 2.25E-19 | DF 5 | glrx |
| ppp1r15a2 | 8.1E-24 | 0.50381 | 0.3 | 0.206 | 2.65E-19 | DF 5 | ppp1r15a |
| si:dkeyp-44a8.43 | 1.1E-23 | 0.4650306 | 0.25 | 0.162 | 3.45E-19 | DF 5 | si:dkeyp-44a8.4 |
| hist1h4l.32 | 1.4E-23 | 0.6555519 | 0.22 | 0.14 | 4.51E-19 | DF 5 | hist1h4l.3 |
| pim13 | 1.7E-23 | 0.3228309 | 0.645 | 0.541 | 5.5E-19 | DF 5 | pim1 |
| etf1b1 | 1.9E-23 | 0.4704048 | 0.492 | 0.389 | 6.34E-19 | DF 5 | etf1b |
| gse12 | 2.4E-23 | 0.6592927 | 0.198 | 0.122 | 7.81E-19 | DF 5 | gse1 |
| kctd9a | 2.9E-23 | 0.768027 | 0.162 | 0.095 | 9.3E-19 | DF 5 | kctd9a |
| scinlb4 | 3.9E-23 | 0.3984304 | 0.411 | 0.304 | 1.28E-18 | DF 5 | scinlb |
| si:ch211-225b11.11 | 4.4E-23 | 0.8280426 | 0.132 | 0.072 | 1.44E-18 | DF 5 | si:ch211-225b11.1 |
| si:dkey-23i12.52 | 6E-23 | 0.565112 | 0.306 | 0.215 | 1.95E-18 | DF 5 | si:dkey-23i12.5 |
| ubl3a1 | 8E-23 | 0.4645295 | 0.437 | 0.333 | 2.61E-18 | DF 5 | ubl3a |
| phldb2b2 | 1E-22 | 0.7315034 | 0.218 | 0.141 | 3.4E-18 | DF 5 | phldb2b |
| lhfpl63 | 1.2E-22 | 0.2697203 | 0.536 | 0.412 | 3.84E-18 | DF 5 | lhfpl6 |
| abl2 | 1.4E-22 | 1.058646 | 0.101 | 0.051 | 4.59E-18 | DF 5 | abl2 |
| cbr1l3 | 1.8E-22 | 0.5319152 | 0.3 | 0.211 | 5.84E-18 | DF 5 | cbr1l |
| smoc21 | 2.4E-22 | 0.7533338 | 0.152 | 0.088 | 7.78E-18 | DF 5 | smoc2 |
| rtn4b2 | 2.5E-22 | 0.8356167 | 0.153 | 0.09 | 8.21E-18 | DF 5 | rtn4b |
| ifrd11 | 3E-22 | 0.5677271 | 0.371 | 0.279 | 9.76E-18 | DF 5 | ifrd1 |
| sult6b13 | 3.5E-22 | 0.606 | 0.257 | 0.174 | 1.14E-17 | DF 5 | sult6b1 |
| eif5a3 | 4E-22 | 0.23599 | 0.805 | 0.717 | 1.29E-17 | DF 5 | eif5a |
| ms4a17a.94 | 4.2E-22 | 0.4660697 | 0.309 | 0.212 | 1.35E-17 | DF 5 | ms4a17a.9 |
| prickle1b4 | 6.4E-22 | 0.5562896 | 0.26 | 0.175 | 2.09E-17 | DF 5 | prickle1b |
| suco3 | 1.4E-21 | 0.488304 | 0.255 | 0.17 | 4.71E-17 | DF 5 | suco |
| tmem176l.11 | 1.7E-21 | 0.2918859 | 0.807 | 0.739 | 5.66E-17 | DF 5 | tmem176l.1 |
| abcf2a2 | 2E-21 | 0.5722145 | 0.256 | 0.175 | 6.35E-17 | DF 5 | abcf2a |
| pebp12 | 3.1E-21 | 0.9020697 | 0.126 | 0.071 | 9.92E-17 | DF 5 | pebp1 |
| hspa12b3 | 3.6E-21 | 0.5390911 | 0.193 | 0.12 | 1.17E-16 | DF 5 | hspa12b |
| fxyd6l2 | 4.1E-21 | 0.3246574 | 0.35 | 0.25 | 1.34E-16 | DF 5 | fxyd6l |
| tubb4b2 | 4.4E-21 | 0.2045386 | 0.865 | 0.807 | 1.44E-16 | DF 5 | tubb4b |
| zfand5b1 | 6.3E-21 | 0.4866196 | 0.363 | 0.269 | 2.06E-16 | DF 5 | zfand5b |
| jmjd1cb2 | 6.6E-21 | 0.3926213 | 0.416 | 0.315 | 2.14E-16 | DF 5 | jmjd1cb |
| mknk2a3 | 6.8E-21 | 0.5262415 | 0.314 | 0.226 | 2.21E-16 | DF 5 | mknk2a |
| ncl1 | 8.1E-21 | 0.2550842 | 0.696 | 0.6 | 2.62E-16 | DF 5 | ncl |
| CABZ01068367.13 | 8.3E-21 | 0.2694917 | 0.501 | 0.389 | 2.7E-16 | DF 5 | CABZ01068367.1 |
| srsf5a1 | 8.6E-21 | 0.3032423 | 0.671 | 0.576 | 2.81E-16 | DF 5 | srsf5a |
| ada2 | 1.6E-20 | 0.5377184 | 0.306 | 0.218 | 5.05E-16 | DF 5 | ada |
| swap70b1 | 1.6E-20 | 1.0402692 | 0.106 | 0.057 | 5.17E-16 | DF 5 | swap70b |
| ntd53 | 1.6E-20 | 0.4065641 | 0.217 | 0.139 | 5.31E-16 | DF 5 | ntd5 |

| | | | | | | | |
|---|---|---|---|---|---|---|---|
| si:dkey-51e6.14 | 1.7E-20 | 0.4548742 | 0.345 | 0.255 | 5.44E-16 | DF 5 | si:dkey-51e6.1 |
| tbx151 | 1.9E-20 | 0.6816825 | 0.115 | 0.062 | 6.33E-16 | DF 5 | tbx15 |
| vstm4a2 | 2.1E-20 | 0.7356885 | 0.214 | 0.142 | 6.69E-16 | DF 5 | vstm4a |
| mxra8b3 | 2.3E-20 | 0.2806217 | 0.515 | 0.404 | 7.35E-16 | DF 5 | mxra8b |
| si:ch73-237c6.14 | 3E-20 | 0.2519377 | 0.477 | 0.361 | 9.79E-16 | DF 5 | si:ch73-237c6.1 |
| tanc2a2 | 4.1E-20 | 0.8192142 | 0.133 | 0.077 | 1.34E-15 | DF 5 | tanc2a |
| csrnp1b3 | 4.1E-20 | 0.5172058 | 0.265 | 0.184 | 1.34E-15 | DF 5 | csrnp1b |
| caskin11 | 4.5E-20 | 0.9030049 | 0.105 | 0.056 | 1.47E-15 | DF 5 | caskin1 |
| pmaip11 | 5.1E-20 | 0.8980438 | 0.152 | 0.092 | 1.66E-15 | DF 5 | pmaip1 |
| khdrbs1b | 5.3E-20 | 0.494268 | 0.324 | 0.237 | 1.72E-15 | DF 5 | khdrbs1b |
| si:ch73-386h18.14 | 5.6E-20 | 0.4504086 | 0.222 | 0.144 | 1.81E-15 | DF 5 | si:ch73-386h18.1 |
| fam43a3 | 5.6E-20 | 0.7174397 | 0.12 | 0.066 | 1.81E-15 | DF 5 | fam43a |
| nedd92 | 7.1E-20 | 1.0121434 | 0.124 | 0.071 | 2.32E-15 | DF 5 | nedd9 |
| ppl3 | 7.2E-20 | 0.3368495 | 0.233 | 0.152 | 2.35E-15 | DF 5 | ppl |
| si:ch211-288g17.32 | 1E-19 | 0.4601061 | 0.494 | 0.4 | 3.4E-15 | DF 5 | si:ch211-288g17.3 |
| etv5a1 | 1.3E-19 | 0.7645885 | 0.164 | 0.102 | 4.3E-15 | DF 5 | etv5a |
| ptbp1b2 | 1.3E-19 | 0.5539733 | 0.268 | 0.186 | 4.35E-15 | DF 5 | ptbp1b |
| col6a3.12 | 2.3E-19 | 0.2019249 | 0.979 | 0.936 | 7.52E-15 | DF 5 | col6a3.1 |
| ccng21 | 3.5E-19 | 0.5016255 | 0.302 | 0.22 | 1.14E-14 | DF 5 | ccng2 |
| btbd6b | 4.2E-19 | 0.8427623 | 0.107 | 0.058 | 1.37E-14 | DF 5 | btbd6b |
| ets21 | 4.5E-19 | 0.556489 | 0.232 | 0.157 | 1.47E-14 | DF 5 | ets2 |
| adh51 | 4.9E-19 | 0.3381044 | 0.601 | 0.514 | 1.59E-14 | DF 5 | adh5 |
| calm1a2 | 6.1E-19 | 0.2509016 | 0.723 | 0.614 | 1.99E-14 | DF 5 | calm1a |
| hspa52 | 6.1E-19 | 0.217308 | 0.601 | 0.487 | 1.99E-14 | DF 5 | hspa5 |
| adipor22 | 7.6E-19 | 0.6030863 | 0.25 | 0.174 | 2.47E-14 | DF 5 | adipor2 |
| gng22 | 8.8E-19 | 0.7670351 | 0.169 | 0.107 | 2.87E-14 | DF 5 | gng2 |
| acvrl13 | 9.5E-19 | 0.442118 | 0.329 | 0.243 | 3.09E-14 | DF 5 | acvrl1 |
| phip2 | 1.2E-18 | 0.4747905 | 0.23 | 0.155 | 3.79E-14 | DF 5 | phip |
| gal3st22 | 1.2E-18 | 0.918981 | 0.127 | 0.075 | 4.04E-14 | DF 5 | gal3st2 |
| slc9a3r1a | 2E-18 | 0.7241062 | 0.114 | 0.064 | 6.63E-14 | DF 5 | slc9a3r1a |
| CR381618.14 | 2.3E-18 | 0.5482205 | 0.303 | 0.224 | 7.53E-14 | DF 5 | CR381618.1 |
| irf1b3 | 3.2E-18 | 0.6461234 | 0.179 | 0.115 | 1.05E-13 | DF 5 | irf1b |
| h3f3c2 | 3.8E-18 | 0.3266254 | 0.552 | 0.462 | 1.25E-13 | DF 5 | h3f3c |
| apof2 | 4.1E-18 | 0.6445586 | 0.115 | 0.065 | 1.33E-13 | DF 5 | apof |
| pbx41 | 4.2E-18 | 0.3926207 | 0.344 | 0.256 | 1.37E-13 | DF 5 | pbx4 |
| polr2a3 | 6.1E-18 | 0.3563082 | 0.455 | 0.359 | 2E-13 | DF 5 | polr2a |
| tnfrsf11a2 | 6.2E-18 | 0.5110264 | 0.256 | 0.181 | 2.03E-13 | DF 5 | tnfrsf11a |
| auts2a2 | 1.4E-17 | 0.4542437 | 0.301 | 0.219 | 4.59E-13 | DF 5 | auts2a |
| hist1h4l.182 | 1.6E-17 | 0.4703565 | 0.183 | 0.118 | 5.26E-13 | DF 5 | hist1h4l.18 |
| ENSDARG00000597( | 1.8E-17 | 0.6616422 | 0.137 | 0.083 | 5.97E-13 | DF 5 | ENSDARG00000059 |
| zfhx43 | 2.2E-17 | 0.3453397 | 0.535 | 0.446 | 7.03E-13 | DF 5 | zfhx4 |
| vgll4b2 | 2.4E-17 | 0.5232843 | 0.231 | 0.161 | 7.74E-13 | DF 5 | vgll4b |
| nsa22 | 2.4E-17 | 0.2397766 | 0.704 | 0.629 | 7.75E-13 | DF 5 | nsa2 |
| megf6a1 | 2.8E-17 | 0.9931264 | 0.101 | 0.057 | 9.2E-13 | DF 5 | megf6a |
| si:dkey-102m7.32 | 3E-17 | 0.6639994 | 0.18 | 0.118 | 9.77E-13 | DF 5 | si:dkey-102m7.3 |
| ldb1a1 | 3.6E-17 | 0.6692875 | 0.162 | 0.103 | 1.16E-12 | DF 5 | ldb1a |
| tsc22d32 | 4.6E-17 | 0.3317004 | 0.49 | 0.399 | 1.5E-12 | DF 5 | tsc22d3 |
| hopx2 | 5.6E-17 | 0.3415149 | 0.356 | 0.267 | 1.82E-12 | DF 5 | hopx |

| | | | | | | | |
|---|---|---|---|---|---|---|---|
| wee1 | 5.7E-17 | 0.7799141 | 0.108 | 0.061 | 1.86E-12 | DF 5 | wee1 |
| PLEKHH31 | 7.1E-17 | 0.5992715 | 0.229 | 0.161 | 2.32E-12 | DF 5 | PLEKHH3 |
| eml33 | 7.2E-17 | 0.6164972 | 0.18 | 0.118 | 2.35E-12 | DF 5 | eml3 |
| thrap3a2 | 7.6E-17 | 0.3041052 | 0.478 | 0.38 | 2.46E-12 | DF 5 | thrap3a |
| si:dkey-222f8.31 | 7.7E-17 | 0.4605467 | 0.297 | 0.219 | 2.51E-12 | DF 5 | si:dkey-222f8.3 |
| romo11 | 1.1E-16 | 0.2116852 | 0.774 | 0.704 | 3.61E-12 | DF 5 | romo1 |
| si:dkey-79d12.51 | 2.2E-16 | 0.5987116 | 0.19 | 0.127 | 7.05E-12 | DF 5 | si:dkey-79d12.5 |
| hdac9b1 | 2.3E-16 | 0.7541541 | 0.116 | 0.068 | 7.41E-12 | DF 5 | hdac9b |
| csnk1da2 | 2.4E-16 | 0.4259514 | 0.287 | 0.21 | 7.71E-12 | DF 5 | csnk1da |
| slc22a23 | 2.4E-16 | 0.2756309 | 0.274 | 0.192 | 7.82E-12 | DF 5 | slc22a2 |
| arl4ab4 | 2.6E-16 | 0.6378747 | 0.167 | 0.108 | 8.53E-12 | DF 5 | arl4ab |
| ccni | 2.6E-16 | 0.2672644 | 0.676 | 0.591 | 8.59E-12 | DF 5 | ccni |
| tshz3b | 2.7E-16 | 0.6923019 | 0.106 | 0.061 | 8.67E-12 | DF 5 | tshz3b |
| NPAS32 | 2.7E-16 | 0.450389 | 0.138 | 0.084 | 8.74E-12 | DF 5 | NPAS3 |
| cish1 | 2.8E-16 | 0.6407222 | 0.216 | 0.152 | 9.24E-12 | DF 5 | cish |
| pcdh18b2 | 3.9E-16 | 0.3765303 | 0.297 | 0.216 | 1.27E-11 | DF 5 | pcdh18b |
| tuba8l21 | 4.3E-16 | 0.4785212 | 0.289 | 0.215 | 1.4E-11 | DF 5 | tuba8l2 |
| si:ch73-366l1.53 | 4.8E-16 | 0.6330203 | 0.132 | 0.081 | 1.55E-11 | DF 5 | si:ch73-366l1.5 |
| si:ch211-15b10.61 | 5.3E-16 | 0.6240725 | 0.153 | 0.098 | 1.71E-11 | DF 5 | si:ch211-15b10.6 |
| mmp92 | 6.5E-16 | 2.1677525 | 0.118 | 0.073 | 2.1E-11 | DF 5 | mmp9 |
| dusp24 | 6.7E-16 | 0.4388991 | 0.295 | 0.22 | 2.18E-11 | DF 5 | dusp2 |
| slc25a43 | 7.8E-16 | 0.7668256 | 0.104 | 0.06 | 2.54E-11 | DF 5 | slc25a43 |
| skilb1 | 8.5E-16 | 0.8894624 | 0.126 | 0.077 | 2.77E-11 | DF 5 | skilb |
| zfp36l1b4 | 8.6E-16 | 0.3232574 | 0.622 | 0.543 | 2.8E-11 | DF 5 | zfp36l1b |
| oaz1b2 | 8.7E-16 | 0.3613272 | 0.389 | 0.304 | 2.82E-11 | DF 5 | oaz1b |
| hexim12 | 1E-15 | 0.4966138 | 0.306 | 0.231 | 3.38E-11 | DF 5 | hexim1 |
| zfand2a1 | 1.1E-15 | 0.8526225 | 0.111 | 0.066 | 3.61E-11 | DF 5 | zfand2a |
| nfkbiaa4 | 1.3E-15 | 0.5742439 | 0.491 | 0.421 | 4.17E-11 | DF 5 | nfkbiaa |
| mpp6b | 1.3E-15 | 0.6818497 | 0.101 | 0.057 | 4.35E-11 | DF 5 | mpp6b |
| fscn1a3 | 1.4E-15 | 0.498564 | 0.245 | 0.176 | 4.55E-11 | DF 5 | fscn1a |
| srsf3b1 | 2E-15 | 0.3369992 | 0.356 | 0.271 | 6.5E-11 | DF 5 | srsf3b |
| slc20a1b1 | 2E-15 | 0.4643778 | 0.369 | 0.292 | 6.54E-11 | DF 5 | slc20a1b |
| znf609a3 | 2.1E-15 | 0.4233834 | 0.242 | 0.173 | 6.75E-11 | DF 5 | znf609a |
| col6a32 | 2.1E-15 | 0.4324052 | 0.313 | 0.235 | 6.88E-11 | DF 5 | col6a3 |
| tubb2b2 | 2.3E-15 | 0.3296884 | 0.424 | 0.338 | 7.32E-11 | DF 5 | tubb2b |
| zfand5a2 | 2.3E-15 | 0.4999126 | 0.256 | 0.186 | 7.38E-11 | DF 5 | zfand5a |
| zgc:1530182 | 2.3E-15 | 0.5451064 | 0.3 | 0.23 | 7.42E-11 | DF 5 | zgc:153018 |
| mapk14a3 | 2.3E-15 | 0.5859001 | 0.133 | 0.082 | 7.46E-11 | DF 5 | mapk14a |
| sh3bp5la1 | 2.5E-15 | 0.595147 | 0.159 | 0.104 | 8.15E-11 | DF 5 | sh3bp5la |
| pdcd4b1 | 2.7E-15 | 0.6293612 | 0.162 | 0.106 | 8.86E-11 | DF 5 | pdcd4b |
| ninl | 7.6E-15 | 0.6625434 | 0.113 | 0.067 | 2.48E-10 | DF 5 | ninl |
| mafba3 | 1.2E-14 | 0.3718431 | 0.47 | 0.387 | 3.78E-10 | DF 5 | mafba |
| zc3h41 | 1.2E-14 | 0.5883194 | 0.197 | 0.137 | 3.91E-10 | DF 5 | zc3h4 |
| atoh8 | 1.3E-14 | 0.9093444 | 0.103 | 0.061 | 4.29E-10 | DF 5 | atoh8 |
| rbfox22 | 1.7E-14 | 0.37403 | 0.346 | 0.268 | 5.5E-10 | DF 5 | rbfox2 |
| rarga1 | 1.8E-14 | 0.3991645 | 0.35 | 0.275 | 5.76E-10 | DF 5 | rarga |
| ism12 | 1.8E-14 | 0.559455 | 0.18 | 0.123 | 5.88E-10 | DF 5 | ism1 |
| BX936337.11 | 2.1E-14 | 0.4107257 | 0.466 | 0.386 | 6.74E-10 | DF 5 | BX936337.1 |

| | | | | | | | |
|---|---|---|---|---|---|---|---|
| sqstm12 | 2.1E-14 | 0.349723 | 0.382 | 0.302 | 6.82E-10 | DF 5 | sqstm1 |
| twsg1a2 | 2.5E-14 | 0.3198022 | 0.335 | 0.255 | 8.1E-10 | DF 5 | twsg1a |
| spred2a1 | 2.7E-14 | 0.6941033 | 0.12 | 0.074 | 8.8E-10 | DF 5 | spred2a |
| cmip | 4.3E-14 | 0.5704895 | 0.175 | 0.119 | 1.41E-09 | DF 5 | cmip |
| eif3ea3 | 4.5E-14 | 0.2250013 | 0.644 | 0.557 | 1.45E-09 | DF 5 | eif3ea |
| arg22 | 4.7E-14 | 0.5233359 | 0.225 | 0.162 | 1.53E-09 | DF 5 | arg2 |
| rhoq2 | 4.8E-14 | 0.6241179 | 0.145 | 0.094 | 1.56E-09 | DF 5 | rhoq |
| si:ch73-86n18.13 | 5.7E-14 | 0.7317743 | 0.213 | 0.152 | 1.85E-09 | DF 5 | si:ch73-86n18.1 |
| elmsan1b3 | 6E-14 | 0.3333582 | 0.437 | 0.354 | 1.94E-09 | DF 5 | elmsan1b |
| prmt11 | 6.1E-14 | 0.340066 | 0.255 | 0.186 | 1.99E-09 | DF 5 | prmt1 |
| rad54l22 | 7.8E-14 | 0.5932525 | 0.128 | 0.081 | 2.53E-09 | DF 5 | rad54l2 |
| zgc:1534091 | 8.1E-14 | 0.3502374 | 0.495 | 0.417 | 2.62E-09 | DF 5 | zgc:153409 |
| phactr4b2 | 9.7E-14 | 0.6998949 | 0.116 | 0.071 | 3.15E-09 | DF 5 | phactr4b |
| sox11b2 | 1.2E-13 | 0.4628624 | 0.276 | 0.208 | 3.79E-09 | DF 5 | sox11b |
| gadd45bb3 | 1.3E-13 | 0.394472 | 0.289 | 0.218 | 4.26E-09 | DF 5 | gadd45bb |
| nfil3-51 | 1.4E-13 | 0.4621847 | 0.195 | 0.137 | 4.56E-09 | DF 5 | nfil3-5 |
| rbbp61 | 1.8E-13 | 0.3327984 | 0.393 | 0.314 | 5.74E-09 | DF 5 | rbbp6 |
| cmtm33 | 2.3E-13 | 0.4137928 | 0.259 | 0.192 | 7.56E-09 | DF 5 | cmtm3 |
| sulf2b4 | 2.6E-13 | 0.4231924 | 0.226 | 0.163 | 8.39E-09 | DF 5 | sulf2b |
| si:ch211-195b15.81 | 3.3E-13 | 0.5003988 | 0.138 | 0.089 | 1.07E-08 | DF 5 | si:ch211-195b15.8 |
| smoc13 | 3.3E-13 | 0.2518276 | 0.279 | 0.205 | 1.07E-08 | DF 5 | smoc1 |
| irf2bpl1 | 3.6E-13 | 0.4671207 | 0.261 | 0.197 | 1.16E-08 | DF 5 | irf2bpl |
| si:cabz01007802.13 | 3.6E-13 | 0.4750938 | 0.137 | 0.088 | 1.16E-08 | DF 5 | si:cabz01007802.1 |
| c1qtnf52 | 3.8E-13 | 0.4662664 | 0.21 | 0.15 | 1.24E-08 | DF 5 | c1qtnf5 |
| anp32e2 | 4.2E-13 | 0.5027075 | 0.199 | 0.141 | 1.36E-08 | DF 5 | anp32e |
| si:dkey-7j14.51 | 4.4E-13 | 0.4404248 | 0.112 | 0.069 | 1.44E-08 | DF 5 | si:dkey-7j14.5 |
| nr1d12 | 4.7E-13 | 0.4891543 | 0.19 | 0.133 | 1.52E-08 | DF 5 | nr1d1 |
| si:dkey-7j14.62 | 4.8E-13 | 0.2340531 | 0.823 | 0.775 | 1.55E-08 | DF 5 | si:dkey-7j14.6 |
| atp1b3b3 | 5.2E-13 | 0.6506094 | 0.121 | 0.077 | 1.69E-08 | DF 5 | atp1b3b |
| ndrg22 | 5.5E-13 | 0.2768287 | 0.404 | 0.319 | 1.8E-08 | DF 5 | ndrg2 |
| alyref2 | 7.1E-13 | 0.2962323 | 0.425 | 0.345 | 2.32E-08 | DF 5 | alyref |
| C24H8orf441 | 9.7E-13 | 0.6012137 | 0.137 | 0.09 | 3.17E-08 | DF 5 | C24H8orf44 |
| tbl1xr1a1 | 1.1E-12 | 0.4892903 | 0.207 | 0.149 | 3.69E-08 | DF 5 | tbl1xr1a |
| adamts15b.13 | 1.2E-12 | 0.4183948 | 0.14 | 0.091 | 3.9E-08 | DF 5 | adamts15b.1 |
| sat1a.23 | 1.3E-12 | 0.267405 | 0.536 | 0.454 | 4.12E-08 | DF 5 | sat1a.2 |
| fbxl3a | 1.3E-12 | 0.6782841 | 0.14 | 0.094 | 4.31E-08 | DF 5 | fbxl3a |
| oaz1a2 | 1.3E-12 | 0.3557843 | 0.518 | 0.45 | 4.35E-08 | DF 5 | oaz1a |
| mpp6a2 | 2E-12 | 0.5960345 | 0.124 | 0.08 | 6.56E-08 | DF 5 | mpp6a |
| h2ax12 | 2.1E-12 | 0.2646215 | 0.684 | 0.626 | 6.82E-08 | DF 5 | h2ax1 |
| aebp12 | 2.2E-12 | 0.2152716 | 0.624 | 0.536 | 7.14E-08 | DF 5 | aebp1 |
| gtpbp42 | 2.8E-12 | 0.2472004 | 0.49 | 0.408 | 8.98E-08 | DF 5 | gtpbp4 |
| tmem1765 | 3.1E-12 | 0.2071081 | 0.486 | 0.39 | 1.02E-07 | DF 5 | tmem176 |
| gask1a2 | 3.2E-12 | 0.412942 | 0.256 | 0.193 | 1.04E-07 | DF 5 | gask1a |
| rem12 | 3.5E-12 | 0.3935458 | 0.103 | 0.063 | 1.14E-07 | DF 5 | rem1 |
| emilin1a2 | 3.9E-12 | 0.4394146 | 0.139 | 0.092 | 1.28E-07 | DF 5 | emilin1a |
| sgtb2 | 4.2E-12 | 0.4506169 | 0.177 | 0.123 | 1.36E-07 | DF 5 | sgtb |
| zbtb16b2 | 4.3E-12 | 0.5491016 | 0.151 | 0.102 | 1.4E-07 | DF 5 | zbtb16b |
| ptenb2 | 4.5E-12 | 0.3654573 | 0.234 | 0.173 | 1.46E-07 | DF 5 | ptenb |

| | | | | | | | |
|---|---|---|---|---|---|---|---|
| cux1a2 | 5.3E-12 | 0.707896 | 0.106 | 0.067 | 1.72E-07 | DF 5 | cux1a |
| lima1a2 | 5.6E-12 | 0.4334585 | 0.134 | 0.088 | 1.83E-07 | DF 5 | lima1a |
| mob2a2 | 8.6E-12 | 0.5354096 | 0.188 | 0.136 | 2.8E-07 | DF 5 | mob2a |
| trim8b | 9.1E-12 | 0.472359 | 0.187 | 0.134 | 2.95E-07 | DF 5 | trim8b |
| ddit41 | 9.7E-12 | 0.5400939 | 0.123 | 0.08 | 3.15E-07 | DF 5 | ddit4 |
| zgc:171679 | 1E-11 | 0.5626446 | 0.106 | 0.066 | 3.27E-07 | DF 5 | zgc:171679 |
| tet31 | 1.4E-11 | 0.6167859 | 0.143 | 0.097 | 4.41E-07 | DF 5 | tet3 |
| tnfsf10l5 | 1.4E-11 | 0.226579 | 0.29 | 0.219 | 4.56E-07 | DF 5 | tnfsf10l |
| ciarta3 | 1.5E-11 | 0.2711798 | 0.353 | 0.281 | 4.76E-07 | DF 5 | ciarta |
| rasgef1bb3 | 1.8E-11 | 0.3930564 | 0.21 | 0.153 | 5.84E-07 | DF 5 | rasgef1bb |
| plxnb2a.12 | 2E-11 | 0.5686167 | 0.132 | 0.089 | 6.35E-07 | DF 5 | plxnb2a.1 |
| tnrc6c12 | 2E-11 | 0.22481 | 0.558 | 0.483 | 6.36E-07 | DF 5 | tnrc6c1 |
| zgc:1122342 | 2.1E-11 | 0.6371753 | 0.113 | 0.073 | 6.75E-07 | DF 5 | zgc:112234 |
| ror21 | 2.5E-11 | 0.3890741 | 0.115 | 0.074 | 7.99E-07 | DF 5 | ror2 |
| lrrfip23 | 2.6E-11 | 0.5395587 | 0.15 | 0.103 | 8.32E-07 | DF 5 | lrrfip2 |
| irf2bp2b1 | 2.9E-11 | 0.3878594 | 0.268 | 0.208 | 9.38E-07 | DF 5 | irf2bp2b |
| ssuh2rs11 | 3.1E-11 | 0.6102682 | 0.105 | 0.067 | 1.02E-06 | DF 5 | ssuh2rs1 |
| angptl53 | 3.4E-11 | 0.2174156 | 0.235 | 0.171 | 1.1E-06 | DF 5 | angptl5 |
| neo1a1 | 3.5E-11 | 0.492836 | 0.236 | 0.181 | 1.13E-06 | DF 5 | neo1a |
| pcf111 | 4.1E-11 | 0.2933667 | 0.317 | 0.25 | 1.35E-06 | DF 5 | pcf11 |
| zmp:00000007113 | 4.2E-11 | 0.4758587 | 0.123 | 0.081 | 1.37E-06 | DF 5 | zmp:0000000711 |
| SCARF22 | 4.9E-11 | 0.2988491 | 0.333 | 0.266 | 1.6E-06 | DF 5 | SCARF2 |
| si:ch211-284f22.33 | 5.2E-11 | 0.4991426 | 0.163 | 0.115 | 1.69E-06 | DF 5 | si:ch211-284f22.3 |
| tgif12 | 5.3E-11 | 0.4595984 | 0.261 | 0.204 | 1.73E-06 | DF 5 | tgif1 |
| mbd3a1 | 5.6E-11 | 0.5862431 | 0.142 | 0.097 | 1.81E-06 | DF 5 | mbd3a |
| ube2al1 | 6.4E-11 | 0.3888424 | 0.256 | 0.197 | 2.07E-06 | DF 5 | ube2al |
| btbd6a1 | 7.4E-11 | 0.5901609 | 0.119 | 0.078 | 2.42E-06 | DF 5 | btbd6a |
| eif4a22 | 9.2E-11 | 0.3416533 | 0.251 | 0.191 | 3E-06 | DF 5 | eif4a2 |
| oaz2a | 1E-10 | 0.299236 | 0.117 | 0.077 | 3.3E-06 | DF 5 | oaz2a |
| oser11 | 1E-10 | 0.6374813 | 0.125 | 0.085 | 3.39E-06 | DF 5 | oser1 |
| mob1bb1 | 1.3E-10 | 0.566231 | 0.121 | 0.081 | 4.09E-06 | DF 5 | mob1bb |
| dab2ipa2 | 1.3E-10 | 0.5055935 | 0.157 | 0.111 | 4.38E-06 | DF 5 | dab2ipa |
| eif4ebp32 | 1.5E-10 | 0.3171573 | 0.229 | 0.172 | 4.82E-06 | DF 5 | eif4ebp3 |
| park72 | 1.5E-10 | 0.4743023 | 0.201 | 0.15 | 4.85E-06 | DF 5 | park7 |
| zbtb16a1 | 1.5E-10 | 0.405861 | 0.179 | 0.129 | 4.96E-06 | DF 5 | zbtb16a |
| ccdc3a1 | 1.6E-10 | 0.641766 | 0.103 | 0.067 | 5.16E-06 | DF 5 | ccdc3a |
| maff2 | 1.8E-10 | 0.371234 | 0.281 | 0.222 | 5.74E-06 | DF 5 | maff |
| raph1b2 | 1.9E-10 | 0.4449482 | 0.151 | 0.105 | 6.18E-06 | DF 5 | raph1b |
| ndst1b | 2E-10 | 0.4939556 | 0.102 | 0.066 | 6.52E-06 | DF 5 | ndst1b |
| ptena1 | 2E-10 | 0.263788 | 0.321 | 0.254 | 6.66E-06 | DF 5 | ptena |
| pik3r3a1 | 2.1E-10 | 0.4497618 | 0.183 | 0.133 | 6.72E-06 | DF 5 | pik3r3a |
| fbxw11b1 | 2.9E-10 | 0.4204403 | 0.211 | 0.158 | 9.44E-06 | DF 5 | fbxw11b |
| zgc:921402 | 2.9E-10 | 0.4405717 | 0.158 | 0.112 | 9.54E-06 | DF 5 | zgc:92140 |
| cx35.42 | 3.1E-10 | 0.4896251 | 0.114 | 0.075 | 9.93E-06 | DF 5 | cx35.4 |
| aldh2.11 | 3.3E-10 | 0.5083365 | 0.155 | 0.11 | 1.08E-05 | DF 5 | aldh2.1 |
| bhlhe403 | 3.7E-10 | 0.2727292 | 0.531 | 0.462 | 1.19E-05 | DF 5 | bhlhe40 |
| rapgef2 | 4.5E-10 | 0.5206291 | 0.148 | 0.104 | 1.46E-05 | DF 5 | rapgef2 |
| tmtc13 | 6.2E-10 | 0.3781307 | 0.162 | 0.115 | 2.02E-05 | DF 5 | tmtc1 |

| | | | | | | | |
|---|---|---|---|---|---|---|---|
| xpo1b1 | 6.5E-10 | 0.50199 | 0.143 | 0.1 | 2.1E-05 | DF 5 | xpo1b |
| ak23 | 6.9E-10 | 0.2439193 | 0.449 | 0.377 | 2.24E-05 | DF 5 | ak2 |
| myd881 | 8.6E-10 | 0.6426617 | 0.114 | 0.077 | 2.81E-05 | DF 5 | myd88 |
| nid1b3 | 9E-10 | 0.4572795 | 0.117 | 0.078 | 2.93E-05 | DF 5 | nid1b |
| CU929070.53 | 9.3E-10 | 0.352712 | 0.297 | 0.239 | 3.02E-05 | DF 5 | CU929070.5 |
| sox4a2 | 9.3E-10 | 0.3342978 | 0.249 | 0.192 | 3.04E-05 | DF 5 | sox4a |
| ell1 | 1.3E-09 | 0.5109369 | 0.132 | 0.092 | 4.35E-05 | DF 5 | ell |
| lmo4a1 | 1.4E-09 | 0.4153423 | 0.17 | 0.124 | 4.39E-05 | DF 5 | lmo4a |
| ankrd111 | 1.6E-09 | 0.2602589 | 0.509 | 0.44 | 5.17E-05 | DF 5 | ankrd11 |
| ISCU1 | 1.8E-09 | 0.3933688 | 0.321 | 0.264 | 5.73E-05 | DF 5 | ISCU |
| ano5b3 | 1.8E-09 | 0.5809078 | 0.124 | 0.086 | 5.83E-05 | DF 5 | ano5b |
| CU467861.61 | 1.8E-09 | 0.4557202 | 0.151 | 0.108 | 5.95E-05 | DF 5 | CU467861.6 |
| BX005309.12 | 2E-09 | 0.4661446 | 0.119 | 0.081 | 6.37E-05 | DF 5 | BX005309.1 |
| bco2l3 | 2E-09 | 0.3083688 | 0.171 | 0.123 | 6.51E-05 | DF 5 | bco2l |
| tsc22d12 | 2.3E-09 | 0.4693126 | 0.157 | 0.114 | 7.33E-05 | DF 5 | tsc22d1 |
| ccn2a2 | 2.4E-09 | 0.3066207 | 0.259 | 0.205 | 7.71E-05 | DF 5 | ccn2a |
| CPNE83 | 2.4E-09 | 0.2148418 | 0.353 | 0.284 | 7.76E-05 | DF 5 | CPNE8 |
| slc1a3a3 | 2.4E-09 | 0.3602614 | 0.199 | 0.148 | 7.77E-05 | DF 5 | slc1a3a |
| si:ch73-46j18.54 | 2.7E-09 | 0.33966 | 0.659 | 0.601 | 8.92E-05 | DF 5 | si:ch73-46j18.5 |
| cry22 | 3.4E-09 | 0.3437858 | 0.216 | 0.165 | 0.000109 | DF 5 | cry2 |
| gabarapb1 | 3.6E-09 | 0.2053113 | 0.491 | 0.422 | 0.000117 | DF 5 | gabarapb |
| etnk12 | 4.6E-09 | 0.2759333 | 0.242 | 0.187 | 0.000151 | DF 5 | etnk1 |
| zdhhc8b1 | 4.8E-09 | 0.4851048 | 0.115 | 0.079 | 0.000156 | DF 5 | zdhhc8b |
| iscub2 | 5.4E-09 | 0.2712252 | 0.248 | 0.191 | 0.000176 | DF 5 | iscub |
| si:ch1073-429i10.3 | 5.5E-09 | 0.2172526 | 0.465 | 0.401 | 0.000177 | DF 5 | si:ch1073-429i10.3 |
| BX284696.1 | 5.6E-09 | 0.5623307 | 0.105 | 0.071 | 0.000182 | DF 5 | BX284696.1 |
| sart32 | 6.1E-09 | 0.5379093 | 0.125 | 0.087 | 0.000197 | DF 5 | sart3 |
| cdkn1a2 | 6.2E-09 | 0.474219 | 0.164 | 0.121 | 0.000202 | DF 5 | cdkn1a |
| notch22 | 7E-09 | 0.2291756 | 0.443 | 0.376 | 0.000229 | DF 5 | notch2 |
| rpia1 | 7.2E-09 | 0.5412284 | 0.134 | 0.096 | 0.000234 | DF 5 | rpia |
| roraa1 | 7.9E-09 | 0.4354734 | 0.186 | 0.14 | 0.000255 | DF 5 | roraa |
| zbtb381 | 8.2E-09 | 0.6409002 | 0.117 | 0.081 | 0.000265 | DF 5 | zbtb38 |
| wtap1 | 8.4E-09 | 0.3492187 | 0.17 | 0.125 | 0.000273 | DF 5 | wtap |
| mnta2 | 1.1E-08 | 0.5012795 | 0.14 | 0.101 | 0.000366 | DF 5 | mnta |
| limk22 | 1.1E-08 | 0.5893966 | 0.115 | 0.08 | 0.00037 | DF 5 | limk2 |
| tmem119b3 | 1.3E-08 | 0.2501877 | 0.261 | 0.204 | 0.00041 | DF 5 | tmem119b |
| podn4 | 1.3E-08 | 0.2196929 | 0.224 | 0.17 | 0.000434 | DF 5 | podn |
| dynll2b1 | 1.5E-08 | 0.3706303 | 0.2 | 0.153 | 0.000483 | DF 5 | dynll2b |
| stard13b1 | 1.7E-08 | 0.4771784 | 0.119 | 0.083 | 0.00056 | DF 5 | stard13b |
| ajuba1 | 1.7E-08 | 0.5501661 | 0.111 | 0.077 | 0.000569 | DF 5 | ajuba |
| cct72 | 1.8E-08 | 0.2083257 | 0.435 | 0.37 | 0.000579 | DF 5 | cct7 |
| kdm2ba1 | 1.8E-08 | 0.3715205 | 0.182 | 0.137 | 0.000599 | DF 5 | kdm2ba |
| C20H1orf1982 | 2.3E-08 | 0.3959121 | 0.231 | 0.183 | 0.000764 | DF 5 | C20H1orf198 |
| si:ch211-132b12.72 | 2.4E-08 | 0.3073479 | 0.202 | 0.155 | 0.000774 | DF 5 | si:ch211-132b12.7 |
| ugdh1 | 2.4E-08 | 0.2379696 | 0.225 | 0.176 | 0.00078 | DF 5 | ugdh |
| myclb2 | 2.8E-08 | 0.3165511 | 0.185 | 0.139 | 0.000899 | DF 5 | myclb |
| letm21 | 2.9E-08 | 0.4599985 | 0.233 | 0.187 | 0.000942 | DF 5 | letm2 |
| rock1 | 3E-08 | 0.4125404 | 0.125 | 0.089 | 0.000979 | DF 5 | rock1 |

| | | | | | | | |
|---|---|---|---|---|---|---|---|
| bag32 | 3E-08 | 0.6089837 | 0.103 | 0.071 | 0.00098 | DF 5 | bag3 |
| larp4ab2 | 3E-08 | 0.364944 | 0.205 | 0.158 | 0.00099 | DF 5 | larp4ab |
| klf154 | 3.6E-08 | 0.2320032 | 0.263 | 0.207 | 0.001182 | DF 5 | klf15 |
| chd54 | 4.1E-08 | 0.3387575 | 0.168 | 0.125 | 0.001325 | DF 5 | chd5 |
| ep300b1 | 4.4E-08 | 0.3561867 | 0.173 | 0.13 | 0.001442 | DF 5 | ep300b |
| angptl43 | 4.5E-08 | 0.2735778 | 0.517 | 0.454 | 0.001464 | DF 5 | angptl4 |
| si:dkey-239h2.32 | 4.7E-08 | 0.3052236 | 0.199 | 0.154 | 0.001543 | DF 5 | si:dkey-239h2.3 |
| rassf2a2 | 4.9E-08 | 0.3778441 | 0.166 | 0.124 | 0.001602 | DF 5 | rassf2a |
| efna5b1 | 5E-08 | 0.5274669 | 0.101 | 0.069 | 0.001633 | DF 5 | efna5b |
| ms4a17a.113 | 5.1E-08 | 0.2759392 | 0.305 | 0.25 | 0.001664 | DF 5 | ms4a17a.11 |
| klf133 | 5.3E-08 | 0.50324 | 0.167 | 0.126 | 0.001711 | DF 5 | klf13 |
| tshz14 | 5.9E-08 | 0.268498 | 0.179 | 0.135 | 0.001915 | DF 5 | tshz1 |
| wdr26b1 | 6.1E-08 | 0.4330702 | 0.127 | 0.091 | 0.001968 | DF 5 | wdr26b |
| fam107b3 | 6.1E-08 | 0.3222183 | 0.257 | 0.208 | 0.001999 | DF 5 | fam107b |
| ankrd10b | 6.5E-08 | 0.4478832 | 0.134 | 0.098 | 0.002126 | DF 5 | ankrd10b |
| si:ch211-166a6.51 | 6.8E-08 | 0.3431209 | 0.201 | 0.157 | 0.00221 | DF 5 | si:ch211-166a6.5 |
| tcf3b | 8.5E-08 | 0.3258609 | 0.269 | 0.219 | 0.002764 | DF 5 | tcf3b |
| gng71 | 1E-07 | 0.3901135 | 0.208 | 0.164 | 0.003269 | DF 5 | gng7 |
| cyp3c13 | 1.2E-07 | 0.4975449 | 0.129 | 0.094 | 0.003832 | DF 5 | cyp3c1 |
| si:dkeyp-113d7.12 | 1.3E-07 | 0.4382991 | 0.184 | 0.143 | 0.004116 | DF 5 | si:dkeyp-113d7.1 |
| ulk21 | 1.4E-07 | 0.4271427 | 0.11 | 0.077 | 0.004657 | DF 5 | ulk2 |
| irf2bp11 | 1.5E-07 | 0.376615 | 0.133 | 0.097 | 0.005002 | DF 5 | irf2bp1 |
| uacab3 | 1.8E-07 | 0.5013118 | 0.103 | 0.072 | 0.005858 | DF 5 | uacab |
| ms4a17a.84 | 1.8E-07 | 0.3930193 | 0.15 | 0.112 | 0.005993 | DF 5 | ms4a17a.8 |
| atf4a3 | 2E-07 | 0.2871522 | 0.349 | 0.296 | 0.006519 | DF 5 | atf4a |
| crsp71 | 2.3E-07 | 0.584716 | 0.102 | 0.072 | 0.007408 | DF 5 | crsp7 |
| atf1 | 2.6E-07 | 0.3892563 | 0.149 | 0.112 | 0.008386 | DF 5 | atf1 |
| CU929418.11 | 2.7E-07 | 0.403889 | 0.163 | 0.124 | 0.00865 | DF 5 | CU929418.1 |
| rnf441 | 2.7E-07 | 0.4013253 | 0.157 | 0.119 | 0.008755 | DF 5 | rnf44 |
| nhsl1a3 | 2.7E-07 | 0.4140042 | 0.11 | 0.078 | 0.00879 | DF 5 | nhsl1a |
| sfxn5b5 | 3.1E-07 | 0.2926077 | 0.178 | 0.136 | 0.010003 | DF 5 | sfxn5b |
| sp41 | 3.2E-07 | 0.2950777 | 0.191 | 0.148 | 0.010389 | DF 5 | sp4 |
| il13ra11 | 3.3E-07 | 0.5773841 | 0.116 | 0.084 | 0.010714 | DF 5 | il13ra1 |
| slc7a8a3 | 3.4E-07 | 0.3435558 | 0.118 | 0.084 | 0.010988 | DF 5 | slc7a8a |
| atrnl1b2 | 3.4E-07 | 0.4377062 | 0.145 | 0.109 | 0.011027 | DF 5 | atrnl1b |
| tefa1 | 3.6E-07 | 0.2338871 | 0.268 | 0.218 | 0.011718 | DF 5 | tefa |
| smad11 | 3.7E-07 | 0.3354823 | 0.149 | 0.111 | 0.011901 | DF 5 | smad1 |
| crema1 | 3.9E-07 | 0.4095778 | 0.144 | 0.108 | 0.012843 | DF 5 | crema |
| rusc2 | 4E-07 | 0.3863528 | 0.155 | 0.117 | 0.012973 | DF 5 | rusc2 |
| cct81 | 4E-07 | 0.2640969 | 0.318 | 0.27 | 0.01311 | DF 5 | cct8 |
| setd21 | 4.1E-07 | 0.343599 | 0.16 | 0.122 | 0.013445 | DF 5 | setd2 |
| zgc:110626 | 4.8E-07 | 0.3433068 | 0.126 | 0.092 | 0.015645 | DF 5 | zgc:110626 |
| nav31 | 4.8E-07 | 0.3360865 | 0.114 | 0.082 | 0.015763 | DF 5 | nav3 |
| eml2 | 5E-07 | 0.4566413 | 0.101 | 0.071 | 0.016361 | DF 5 | eml2 |
| BX548028.12 | 5.5E-07 | 0.5289854 | 0.103 | 0.073 | 0.017875 | DF 5 | BX548028.1 |
| rpz21 | 5.5E-07 | 0.4737558 | 0.218 | 0.178 | 0.017887 | DF 5 | rpz2 |
| hnrnpr1 | 5.7E-07 | 0.3805371 | 0.156 | 0.12 | 0.018588 | DF 5 | hnrnpr |
| rorab1 | 5.7E-07 | 0.3850016 | 0.163 | 0.125 | 0.018636 | DF 5 | rorab |

| | | | | | | | |
|---|---|---|---|---|---|---|---|
| ppp1r13l3 | 5.7E-07 | 0.2942133 | 0.12 | 0.086 | 0.018697 | DF 5 | ppp1r13l |
| si:ch211-286b5.5 | 6.7E-07 | 0.2599915 | 0.305 | 0.257 | 0.021681 | DF 5 | si:ch211-286b5.5 |
| FP236812.11 | 7.7E-07 | 0.2656298 | 0.133 | 0.098 | 0.025029 | DF 5 | FP236812.1 |
| waslb1 | 7.7E-07 | 0.3120067 | 0.217 | 0.174 | 0.025109 | DF 5 | waslb |
| ube2e31 | 7.8E-07 | 0.2532528 | 0.38 | 0.327 | 0.025248 | DF 5 | ube2e3 |
| trim251 | 7.9E-07 | 0.3704016 | 0.128 | 0.095 | 0.025583 | DF 5 | trim25 |
| C2H14orf119 | 7.9E-07 | 0.3686308 | 0.121 | 0.089 | 0.025807 | DF 5 | C2H14orf119 |
| fbxo11a | 9.3E-07 | 0.431582 | 0.136 | 0.102 | 0.03009 | DF 5 | fbxo11a |
| gtf2b1 | 1E-06 | 0.5258655 | 0.108 | 0.078 | 0.032846 | DF 5 | gtf2b |
| psd22 | 1.1E-06 | 0.3432049 | 0.13 | 0.096 | 0.036778 | DF 5 | psd2 |
| dlg5a1 | 1.2E-06 | 0.3844749 | 0.132 | 0.099 | 0.039064 | DF 5 | dlg5a |
| si:ch211-110e21.34 | 1.2E-06 | 0.3674302 | 0.106 | 0.076 | 0.039706 | DF 5 | si:ch211-110e21.3 |
| kctd5a2 | 1.3E-06 | 0.3096473 | 0.364 | 0.318 | 0.041955 | DF 5 | kctd5a |
| pim31 | 1.4E-06 | 0.4431818 | 0.14 | 0.106 | 0.045298 | DF 5 | pim3 |
| sept9a1 | 1.4E-06 | 0.3425681 | 0.258 | 0.215 | 0.046999 | DF 5 | sept9a |
| TST1 | 1.9E-06 | 0.3370996 | 0.1 | 0.072 | 0.061213 | DF 5 | TST |
| cry3a1 | 1.9E-06 | 0.3851326 | 0.125 | 0.093 | 0.061547 | DF 5 | cry3a |
| amd11 | 1.9E-06 | 0.3329427 | 0.127 | 0.095 | 0.061856 | DF 5 | amd1 |
| fhl3b2 | 1.9E-06 | 0.3052287 | 0.145 | 0.111 | 0.062437 | DF 5 | fhl3b |
| glud1b | 2E-06 | 0.3612345 | 0.14 | 0.106 | 0.065022 | DF 5 | glud1b |
| brd3b1 | 2E-06 | 0.2647994 | 0.269 | 0.226 | 0.066457 | DF 5 | brd3b |
| ppp3r1a1 | 2.1E-06 | 0.3456149 | 0.178 | 0.141 | 0.067141 | DF 5 | ppp3r1a |
| rps6ka3a | 2.1E-06 | 0.4074345 | 0.103 | 0.074 | 0.067958 | DF 5 | rps6ka3a |
| srsf7a1 | 2.4E-06 | 0.2799299 | 0.226 | 0.185 | 0.077552 | DF 5 | srsf7a |
| histh1l2 | 2.8E-06 | 0.4985964 | 0.251 | 0.21 | 0.092365 | DF 5 | histh1l |
| jak13 | 2.9E-06 | 0.2252849 | 0.489 | 0.436 | 0.094201 | DF 5 | jak1 |
| mta11 | 3E-06 | 0.3201797 | 0.121 | 0.089 | 0.096059 | DF 5 | mta1 |
| rhpn22 | 3.3E-06 | 0.3519488 | 0.179 | 0.142 | 0.1087 | DF 5 | rhpn2 |
| lrrc155 | 3.4E-06 | 0.2158911 | 0.172 | 0.133 | 0.110924 | DF 5 | lrrc15 |
| tmeff1b2 | 3.5E-06 | 0.2868441 | 0.139 | 0.105 | 0.113639 | DF 5 | tmeff1b |
| peli1b2 | 3.5E-06 | 0.3853516 | 0.12 | 0.09 | 0.114164 | DF 5 | peli1b |
| ncoa5 | 4E-06 | 0.2758636 | 0.143 | 0.109 | 0.128499 | DF 5 | ncoa5 |
| stag2b1 | 4E-06 | 0.2820528 | 0.207 | 0.166 | 0.130991 | DF 5 | stag2b |
| asap33 | 4.1E-06 | 0.2538782 | 0.161 | 0.125 | 0.13463 | DF 5 | asap3 |
| ccnl1b2 | 4.1E-06 | 0.4347855 | 0.112 | 0.083 | 0.134909 | DF 5 | ccnl1b |
| rexo12 | 4.6E-06 | 0.4345436 | 0.151 | 0.118 | 0.148373 | DF 5 | rexo1 |
| foxo42 | 4.6E-06 | 0.3800416 | 0.138 | 0.106 | 0.148516 | DF 5 | foxo4 |
| agap11 | 4.8E-06 | 0.3479547 | 0.104 | 0.076 | 0.15572 | DF 5 | agap1 |
| hnrnpl21 | 5.2E-06 | 0.2519327 | 0.185 | 0.147 | 0.170337 | DF 5 | hnrnpl2 |
| tiparp2 | 6E-06 | 0.2900711 | 0.159 | 0.124 | 0.194506 | DF 5 | tiparp |
| si:dkeyp-121d4.3 | 6.5E-06 | 0.2640863 | 0.143 | 0.11 | 0.211864 | DF 5 | si:dkeyp-121d4.3 |
| luc7l31 | 6.9E-06 | 0.3395165 | 0.149 | 0.116 | 0.223611 | DF 5 | luc7l3 |
| kdm6ba1 | 6.9E-06 | 0.2041661 | 0.263 | 0.22 | 0.225235 | DF 5 | kdm6ba |
| tktb1 | 7.1E-06 | 0.3226336 | 0.174 | 0.138 | 0.229748 | DF 5 | tktb |
| per22 | 7.2E-06 | 0.3854544 | 0.135 | 0.104 | 0.234489 | DF 5 | per2 |
| ENSDARG000000713. | 8.1E-06 | 0.3441253 | 0.115 | 0.086 | 0.263411 | DF 5 | ENSDARG00000071 |
| limch1b2 | 8.6E-06 | 0.3853992 | 0.12 | 0.09 | 0.2802 | DF 5 | limch1b |
| rtf21 | 8.6E-06 | 0.2513597 | 0.151 | 0.118 | 0.280888 | DF 5 | rtf2 |

| | | | | | | | |
|---|---|---|---|---|---|---|---|
| si:ch211-140m22.71 | 8.9E-06 | 0.313011 | 0.175 | 0.14 | 0.289605 | DF 5 | si:ch211-140m22.7 |
| ino801 | 9.4E-06 | 0.3526732 | 0.134 | 0.104 | 0.306742 | DF 5 | ino80 |
| waca1 | 1E-05 | 0.2764233 | 0.152 | 0.119 | 0.341026 | DF 5 | waca |
| rnf243 | 1.1E-05 | 0.3958771 | 0.121 | 0.092 | 0.34502 | DF 5 | rnf24 |
| si:ch211-183d21.13 | 1.1E-05 | 0.2306562 | 0.436 | 0.38 | 0.3564 | DF 5 | si:ch211-183d21.1 |
| si:ch211-222k6.32 | 1.2E-05 | 0.2914655 | 0.142 | 0.11 | 0.374785 | DF 5 | si:ch211-222k6.3 |
| asap1a2 | 1.2E-05 | 0.3446686 | 0.121 | 0.092 | 0.388233 | DF 5 | asap1a |
| spoplb | 1.2E-05 | 0.4247147 | 0.127 | 0.098 | 0.406322 | DF 5 | spoplb |
| eef2k1 | 1.3E-05 | 0.2845343 | 0.126 | 0.096 | 0.411114 | DF 5 | eef2k |
| grwd13 | 1.3E-05 | 0.4714158 | 0.174 | 0.141 | 0.422556 | DF 5 | grwd1 |
| ctdspl2b3 | 1.4E-05 | 0.3260974 | 0.145 | 0.112 | 0.45312 | DF 5 | ctdspl2b |
| si:dkey-177p2.61 | 1.4E-05 | 0.2033211 | 0.308 | 0.262 | 0.45585 | DF 5 | si:dkey-177p2.6 |
| bckdhb | 1.4E-05 | 0.307027 | 0.109 | 0.082 | 0.46867 | DF 5 | bckdhb |
| stk351 | 1.5E-05 | 0.3418628 | 0.111 | 0.083 | 0.48995 | DF 5 | stk35 |
| tent5c1 | 1.5E-05 | 0.3453675 | 0.111 | 0.083 | 0.496318 | DF 5 | tent5c |
| zc4h21 | 1.6E-05 | 0.3623799 | 0.105 | 0.078 | 0.505055 | DF 5 | zc4h2 |
| srrt1 | 1.6E-05 | 0.2814197 | 0.16 | 0.126 | 0.516294 | DF 5 | srrt |
| bmpr2b1 | 1.6E-05 | 0.3614208 | 0.111 | 0.083 | 0.519762 | DF 5 | bmpr2b |
| znf384l1 | 1.7E-05 | 0.2312227 | 0.149 | 0.117 | 0.55156 | DF 5 | znf384l |
| crlf32 | 1.7E-05 | 0.3456166 | 0.116 | 0.088 | 0.558403 | DF 5 | crlf3 |
| rin22 | 1.7E-05 | 0.42694 | 0.142 | 0.112 | 0.560573 | DF 5 | rin2 |
| slc25a36a3 | 1.9E-05 | 0.3169308 | 0.136 | 0.106 | 0.616364 | DF 5 | slc25a36a |
| med281 | 2E-05 | 0.4589638 | 0.104 | 0.078 | 0.653343 | DF 5 | med28 |
| grinab2 | 2E-05 | 0.235055 | 0.251 | 0.211 | 0.654681 | DF 5 | grinab |
| cdc42ep4b3 | 2.1E-05 | 0.5350705 | 0.111 | 0.085 | 0.675879 | DF 5 | cdc42ep4b |
| gli33 | 2.1E-05 | 0.3289431 | 0.141 | 0.11 | 0.683196 | DF 5 | gli3 |
| smo2 | 2.1E-05 | 0.2061198 | 0.3 | 0.256 | 0.684306 | DF 5 | smo |
| pfkfb32 | 2.3E-05 | 0.4080404 | 0.101 | 0.076 | 0.743381 | DF 5 | pfkfb3 |
| zgc:1949892 | 2.3E-05 | 0.3446001 | 0.119 | 0.09 | 0.757476 | DF 5 | zgc:194989 |
| hbp11 | 2.4E-05 | 0.3778514 | 0.141 | 0.112 | 0.795282 | DF 5 | hbp1 |
| fzd7b2 | 2.5E-05 | 0.2933377 | 0.139 | 0.109 | 0.824224 | DF 5 | fzd7b |
| nbr1a | 2.7E-05 | 0.3851442 | 0.106 | 0.08 | 0.876082 | DF 5 | nbr1a |
| dnpep1 | 2.7E-05 | 0.3576148 | 0.127 | 0.098 | 0.880166 | DF 5 | dnpep |
| esd1 | 3E-05 | 0.3032786 | 0.167 | 0.135 | 0.975161 | DF 5 | esd |
| mllt101 | 3.1E-05 | 0.4403816 | 0.108 | 0.082 | 1 | DF 5 | mllt10 |
| CU459056.11 | 3.2E-05 | 0.2286675 | 0.205 | 0.168 | 1 | DF 5 | CU459056.1 |
| dstyk1 | 3.4E-05 | 0.3190829 | 0.125 | 0.097 | 1 | DF 5 | dstyk |
| stat31 | 3.4E-05 | 0.2817954 | 0.183 | 0.149 | 1 | DF 5 | stat3 |
| efnb3b2 | 3.4E-05 | 0.242313 | 0.172 | 0.138 | 1 | DF 5 | efnb3b |
| b4galt61 | 3.5E-05 | 0.2535764 | 0.134 | 0.105 | 1 | DF 5 | b4galt6 |
| tet2 | 3.7E-05 | 0.3236246 | 0.101 | 0.075 | 1 | DF 5 | tet2 |
| prex22 | 3.9E-05 | 0.3003931 | 0.115 | 0.087 | 1 | DF 5 | prex2 |
| nid1a2 | 4E-05 | 0.2808937 | 0.272 | 0.234 | 1 | DF 5 | nid1a |
| pxna1 | 4.1E-05 | 0.2829284 | 0.137 | 0.108 | 1 | DF 5 | pxna |
| nr1d2b3 | 4.1E-05 | 0.2344331 | 0.186 | 0.151 | 1 | DF 5 | nr1d2b |
| rbms2a | 4.2E-05 | 0.261183 | 0.176 | 0.143 | 1 | DF 5 | rbms2a |
| cad1 | 4.4E-05 | 0.488351 | 0.105 | 0.08 | 1 | DF 5 | cad |
| cdca4 | 4.6E-05 | 0.4217392 | 0.1 | 0.076 | 1 | DF 5 | cdca4 |

| | | | | | | | |
|---|---|---|---|---|---|---|---|
| polr1a1 | 4.6E-05 | 0.4052954 | 0.101 | 0.077 | 1 | DF 5 | polr1a |
| BX511129.22 | 5.1E-05 | 0.3712184 | 0.114 | 0.088 | 1 | DF 5 | BX511129.2 |
| tmem176l.3a1 | 5.3E-05 | 0.2384493 | 0.182 | 0.149 | 1 | DF 5 | tmem176l.3a |
| sh2b11 | 5.7E-05 | 0.4170934 | 0.109 | 0.083 | 1 | DF 5 | sh2b1 |
| tbx2b2 | 5.8E-05 | 0.304957 | 0.156 | 0.126 | 1 | DF 5 | tbx2b |
| eif4ebp12 | 6E-05 | 0.2975903 | 0.184 | 0.152 | 1 | DF 5 | eif4ebp1 |
| nktr1 | 6E-05 | 0.2515783 | 0.192 | 0.158 | 1 | DF 5 | nktr |
| ap2m1a1 | 6.1E-05 | 0.2076126 | 0.346 | 0.304 | 1 | DF 5 | ap2m1a |
| bmp2b2 | 6.6E-05 | 0.3622487 | 0.128 | 0.101 | 1 | DF 5 | bmp2b |
| zmynd11 | 6.9E-05 | 0.2035493 | 0.171 | 0.138 | 1 | DF 5 | zmynd11 |
| tcf201 | 7.6E-05 | 0.2716338 | 0.173 | 0.141 | 1 | DF 5 | tcf20 |
| arhgap31 | 7.8E-05 | 0.2355929 | 0.142 | 0.112 | 1 | DF 5 | arhgap31 |
| si:dkey-253i9.4 | 7.9E-05 | 0.3718083 | 0.101 | 0.077 | 1 | DF 5 | si:dkey-253i9.4 |
| nocta1 | 8.4E-05 | 0.2852353 | 0.141 | 0.113 | 1 | DF 5 | nocta |
| spen1 | 9.4E-05 | 0.2238367 | 0.232 | 0.197 | 1 | DF 5 | spen |
| kansl1b1 | 0.00011 | 0.2797806 | 0.203 | 0.17 | 1 | DF 5 | kansl1b |
| si:dkey-9i23.151 | 0.00011 | 0.211419 | 0.182 | 0.151 | 1 | DF 5 | si:dkey-9i23.15 |
| rbm4.21 | 0.00012 | 0.2684449 | 0.198 | 0.165 | 1 | DF 5 | rbm4.2 |
| per1b2 | 0.00013 | 0.2405656 | 0.239 | 0.202 | 1 | DF 5 | per1b |
| phf20b | 0.00014 | 0.2504756 | 0.208 | 0.175 | 1 | DF 5 | phf20b |
| clip21 | 0.00014 | 0.3937576 | 0.128 | 0.102 | 1 | DF 5 | clip2 |
| dag12 | 0.00015 | 0.255311 | 0.188 | 0.157 | 1 | DF 5 | dag1 |
| crebbpb | 0.00015 | 0.2224647 | 0.18 | 0.148 | 1 | DF 5 | crebbpb |
| epc2 | 0.00016 | 0.3460073 | 0.107 | 0.084 | 1 | DF 5 | epc2 |
| luc7l | 0.00018 | 0.2022185 | 0.177 | 0.146 | 1 | DF 5 | luc7l |
| tspan9b | 0.00018 | 0.2280418 | 0.162 | 0.133 | 1 | DF 5 | tspan9b |
| mybbp1a1 | 0.00019 | 0.323111 | 0.152 | 0.125 | 1 | DF 5 | mybbp1a |
| kdm4b1 | 0.0002 | 0.2057341 | 0.218 | 0.184 | 1 | DF 5 | kdm4b |
| ccnt2b1 | 0.00022 | 0.2417823 | 0.127 | 0.101 | 1 | DF 5 | ccnt2b |
| rest | 0.00023 | 0.3174946 | 0.106 | 0.083 | 1 | DF 5 | rest |
| nsd1a1 | 0.00024 | 0.2536261 | 0.117 | 0.092 | 1 | DF 5 | nsd1a |
| dhx151 | 0.00024 | 0.2277358 | 0.137 | 0.11 | 1 | DF 5 | dhx15 |
| meaf6 | 0.00028 | 0.2484084 | 0.198 | 0.167 | 1 | DF 5 | meaf6 |
| aamp4 | 0.00028 | 0.258156 | 0.182 | 0.154 | 1 | DF 5 | aamp |
| cicb3 | 0.0003 | 0.2035873 | 0.155 | 0.127 | 1 | DF 5 | cicb |
| dennd4a3 | 0.00033 | 0.2905124 | 0.109 | 0.086 | 1 | DF 5 | dennd4a |
| nfyba2 | 0.00034 | 0.2439069 | 0.212 | 0.182 | 1 | DF 5 | nfyba |
| selenot1a1 | 0.00037 | 0.2729976 | 0.198 | 0.169 | 1 | DF 5 | selenot1a |
| zgc:1752142 | 0.00039 | 0.3229981 | 0.131 | 0.107 | 1 | DF 5 | zgc:175214 |
| kansl1a | 0.00043 | 0.3089206 | 0.103 | 0.081 | 1 | DF 5 | kansl1a |
| si:ch211-213a13.11 | 0.00044 | 0.2458836 | 0.156 | 0.128 | 1 | DF 5 | si:ch211-213a13.1 |
| lonrf1l3 | 0.00044 | 0.2413462 | 0.18 | 0.151 | 1 | DF 5 | lonrf1l |
| terfa | 0.00049 | 0.3851107 | 0.11 | 0.088 | 1 | DF 5 | terfa |
| arid1aa1 | 0.00049 | 0.3034485 | 0.142 | 0.117 | 1 | DF 5 | arid1aa |
| fgfr1a2 | 0.00051 | 0.3176035 | 0.183 | 0.156 | 1 | DF 5 | fgfr1a |
| fam199x1 | 0.00051 | 0.3429919 | 0.158 | 0.132 | 1 | DF 5 | fam199x |
| bbx | 0.00054 | 0.3087222 | 0.103 | 0.081 | 1 | DF 5 | bbx |
| acox11 | 0.00055 | 0.2408676 | 0.126 | 0.102 | 1 | DF 5 | acox1 |

| | | | | | | | |
|---|---|---|---|---|---|---|---|
| prelid1a1 | 0.00058 | 0.2327918 | 0.15 | 0.124 | 1 | DF 5 | prelid1a |
| creb1a1 | 0.00062 | 0.3110993 | 0.19 | 0.162 | 1 | DF 5 | creb1a |
| bach1b1 | 0.00064 | 0.275945 | 0.115 | 0.092 | 1 | DF 5 | bach1b |
| vapal1 | 0.00065 | 0.2436115 | 0.193 | 0.164 | 1 | DF 5 | vapal |
| rassf12 | 0.00067 | 0.3302604 | 0.144 | 0.12 | 1 | DF 5 | rassf1 |
| dzip1 | 0.00068 | 0.2098261 | 0.137 | 0.112 | 1 | DF 5 | dzip1 |
| zgc:922421 | 0.00077 | 0.2473891 | 0.135 | 0.11 | 1 | DF 5 | zgc:92242 |
| wipi12 | 0.0008 | 0.2709892 | 0.146 | 0.121 | 1 | DF 5 | wipi1 |
| CR936321.52 | 0.00084 | 0.3083936 | 0.117 | 0.094 | 1 | DF 5 | CR936321.5 |
| hif1al2 | 0.00085 | 0.3656839 | 0.135 | 0.112 | 1 | DF 5 | hif1al |
| tgfbr1b1 | 0.00097 | 0.3170321 | 0.137 | 0.114 | 1 | DF 5 | tgfbr1b |
| h1-0 | 0.00101 | 0.2746279 | 0.208 | 0.181 | 1 | DF 5 | h1-0 |
| pepd3 | 0.00106 | 0.2077937 | 0.242 | 0.213 | 1 | DF 5 | pepd |
| rnf1111 | 0.00111 | 0.2897521 | 0.114 | 0.093 | 1 | DF 5 | rnf111 |
| fam13b2 | 0.00112 | 0.2303736 | 0.144 | 0.119 | 1 | DF 5 | fam13b |
| timp2a3 | 0.00114 | 0.3692221 | 0.783 | 0.755 | 1 | DF 5 | timp2a |
| uqcc31 | 0.00115 | 0.3110442 | 0.103 | 0.083 | 1 | DF 5 | uqcc3 |
| arih2 | 0.00116 | 0.3647781 | 0.101 | 0.081 | 1 | DF 5 | arih2 |
| pprc12 | 0.00116 | 0.2879308 | 0.139 | 0.116 | 1 | DF 5 | pprc1 |
| bckdk2 | 0.0013 | 0.2702591 | 0.109 | 0.088 | 1 | DF 5 | bckdk |
| pik3r11 | 0.0013 | 0.215873 | 0.22 | 0.191 | 1 | DF 5 | pik3r1 |
| kmt5aa1 | 0.00134 | 0.3323298 | 0.136 | 0.114 | 1 | DF 5 | kmt5aa |
| rif13 | 0.00142 | 0.2354616 | 0.106 | 0.085 | 1 | DF 5 | rif1 |
| ep300a1 | 0.00143 | 0.2239043 | 0.145 | 0.12 | 1 | DF 5 | ep300a |
| dohh2 | 0.00144 | 0.2143078 | 0.175 | 0.149 | 1 | DF 5 | dohh |
| pdcd111 | 0.0015 | 0.3368845 | 0.134 | 0.112 | 1 | DF 5 | pdcd11 |
| ube2d4 | 0.00154 | 0.3557795 | 0.104 | 0.084 | 1 | DF 5 | ube2d4 |
| runx1t12 | 0.00165 | 0.219338 | 0.113 | 0.091 | 1 | DF 5 | runx1t1 |
| taf9 | 0.00168 | 0.2127688 | 0.148 | 0.124 | 1 | DF 5 | taf9 |
| aff41 | 0.00201 | 0.2817951 | 0.105 | 0.085 | 1 | DF 5 | aff4 |
| prr12a | 0.00219 | 0.2384681 | 0.128 | 0.106 | 1 | DF 5 | prr12a |
| lysmd2 | 0.00232 | 0.2454004 | 0.101 | 0.081 | 1 | DF 5 | lysmd2 |
| hk11 | 0.00256 | 0.2365527 | 0.141 | 0.118 | 1 | DF 5 | hk1 |
| mgat1b3 | 0.00263 | 0.2970071 | 0.117 | 0.097 | 1 | DF 5 | mgat1b |
| epc1b | 0.00265 | 0.2644748 | 0.12 | 0.1 | 1 | DF 5 | epc1b |
| srsf10b2 | 0.00272 | 0.2705031 | 0.105 | 0.086 | 1 | DF 5 | srsf10b |
| cbx1b1 | 0.00273 | 0.2718727 | 0.128 | 0.107 | 1 | DF 5 | cbx1b |
| si:dkey-276j7.12 | 0.00288 | 0.2197385 | 0.192 | 0.167 | 1 | DF 5 | si:dkey-276j7.1 |
| farp23 | 0.00296 | 0.2993522 | 0.135 | 0.115 | 1 | DF 5 | farp2 |
| zgc:1531151 | 0.00313 | 0.2194183 | 0.101 | 0.082 | 1 | DF 5 | zgc:153115 |
| ranbp91 | 0.00317 | 0.2062344 | 0.146 | 0.123 | 1 | DF 5 | ranbp9 |
| NDUFB11 | 0.00343 | 0.2499024 | 0.113 | 0.094 | 1 | DF 5 | NDUFB1 |
| hcfc1a1 | 0.00349 | 0.2587039 | 0.115 | 0.095 | 1 | DF 5 | hcfc1a |
| mlf2 | 0.00355 | 0.2049643 | 0.148 | 0.125 | 1 | DF 5 | mlf2 |
| nr4a12 | 0.0037 | 0.3307115 | 0.121 | 0.102 | 1 | DF 5 | nr4a1 |
| nfs11 | 0.00386 | 0.2976178 | 0.106 | 0.087 | 1 | DF 5 | nfs1 |
| osbpl9 | 0.00393 | 0.2734062 | 0.179 | 0.156 | 1 | DF 5 | osbpl9 |
| yy1a | 0.00402 | 0.2027938 | 0.151 | 0.129 | 1 | DF 5 | yy1a |

| | | | | | | | |
|---|---|---|---|---|---|---|---|
| tcf7l1a2 | 0.00419 | 0.2103504 | 0.138 | 0.116 | 1 | DF 5 | tcf7l1a |
| si:dkey-118j18.2 | 0.00429 | 0.2301367 | 0.111 | 0.092 | 1 | DF 5 | si:dkey-118j18.2 |
| cdc731 | 0.00439 | 0.2500267 | 0.1 | 0.082 | 1 | DF 5 | cdc73 |
| spred1 | 0.00456 | 0.2208672 | 0.149 | 0.128 | 1 | DF 5 | spred1 |
| baz1a1 | 0.00512 | 0.2365794 | 0.134 | 0.113 | 1 | DF 5 | baz1a |
| C2H1orf352 | 0.00528 | 0.2348077 | 0.103 | 0.086 | 1 | DF 5 | C2H1orf35 |
| cul4b | 0.00531 | 0.2125372 | 0.165 | 0.143 | 1 | DF 5 | cul4b |
| G3BP21 | 0.00548 | 0.2491291 | 0.13 | 0.11 | 1 | DF 5 | G3BP2 |
| parp42 | 0.00591 | 0.3259965 | 0.101 | 0.084 | 1 | DF 5 | parp4 |
| cnn22 | 0.00609 | 0.2252682 | 0.471 | 0.445 | 1 | DF 5 | cnn2 |
| FOXK22 | 0.00655 | 0.2295794 | 0.157 | 0.137 | 1 | DF 5 | FOXK2 |
| ncbp21 | 0.00705 | 0.2697597 | 0.142 | 0.123 | 1 | DF 5 | ncbp2 |
| si:dkey-112a7.41 | 0.00749 | 0.3150816 | 0.105 | 0.088 | 1 | DF 5 | si:dkey-112a7.4 |
| pank2 | 0.00775 | 0.214119 | 0.118 | 0.099 | 1 | DF 5 | pank2 |
| lrig21 | 0.00804 | 0.2466915 | 0.11 | 0.092 | 1 | DF 5 | lrig2 |
| dnajc5aa1 | 0.00805 | 0.243213 | 0.101 | 0.084 | 1 | DF 5 | dnajc5aa |
| znf395b1 | 0.00858 | 0.2008573 | 0.159 | 0.138 | 1 | DF 5 | znf395b |
| kmt5c1 | 0.00885 | 0.2293502 | 0.125 | 0.107 | 1 | DF 5 | kmt5c |
| si:dkeyp-120h9.12 | 0.00988 | 0.2442894 | 0.1 | 0.084 | 1 | DF 5 | si:dkeyp-120h9.1 |
| spock35 | 0 | 2.4419068 | 0.928 | 0.395 | 0 | DF 6 | spock3 |
| tbx3a3 | 0 | 2.7130143 | 0.529 | 0.111 | 0 | DF 6 | tbx3a |
| foxd11 | 0 | 3.6147301 | 0.439 | 0.055 | 0 | DF 6 | foxd1 |
| dkk21 | 0 | 3.4549332 | 0.365 | 0.045 | 0 | DF 6 | dkk2 |
| foxl2a | 0 | 3.4352726 | 0.192 | 0.021 | 0 | DF 6 | foxl2a |
| sox7 | 0 | 4.6062262 | 0.185 | 0.014 | 0 | DF 6 | sox7 |
| hhipl2 | 0 | 4.2459913 | 0.175 | 0.011 | 0 | DF 6 | hhipl2 |
| wnt6b | 0 | 4.3918215 | 0.169 | 0.01 | 0 | DF 6 | wnt6b |
| si:ch211-161f7.2 | 0 | 4.4387789 | 0.145 | 0.008 | 0 | DF 6 | si:ch211-161f7.2 |
| aplp11 | 4E-291 | 3.1329249 | 0.234 | 0.036 | 1.2E-286 | DF 6 | aplp1 |
| ptn | 1E-282 | 2.7582058 | 0.25 | 0.042 | 4.4E-278 | DF 6 | ptn |
| hpgd2 | 2E-272 | 2.833794 | 0.267 | 0.049 | 6E-268 | DF 6 | hpgd |
| gstr3 | 3E-271 | 1.9075246 | 0.748 | 0.406 | 1.1E-266 | DF 6 | gstr |
| lrp1ba | 4E-255 | 3.3808877 | 0.137 | 0.014 | 1.2E-250 | DF 6 | lrp1ba |
| znf536 | 2E-254 | 3.248516 | 0.165 | 0.02 | 6.1E-250 | DF 6 | znf536 |
| ndrg42 | 4E-249 | 2.0261669 | 0.462 | 0.145 | 1.4E-244 | DF 6 | ndrg4 |
| ntn1a4 | 7E-238 | 2.1327895 | 0.395 | 0.112 | 2.1E-233 | DF 6 | ntn1a |
| kera3 | 4E-234 | 2.078385 | 0.477 | 0.162 | 1.4E-229 | DF 6 | kera |
| bhmt5 | 4E-231 | 1.1471648 | 0.953 | 0.7 | 1.4E-226 | DF 6 | bhmt |
| f13a1b | 4E-229 | 0.7733124 | 0.649 | 0.283 | 1.2E-224 | DF 6 | f13a1b |
| fmoda3 | 3E-221 | 1.7028805 | 0.5 | 0.173 | 9.6E-217 | DF 6 | fmoda |
| krt914 | 8E-213 | 1.0095415 | 0.984 | 0.899 | 2.6E-208 | DF 6 | krt91 |
| tfap2b1 | 1E-211 | 3.0246298 | 0.155 | 0.021 | 4.1E-207 | DF 6 | tfap2b |
| col9a3 | 2E-201 | 0.2592715 | 0.278 | 0.063 | 6.8E-197 | DF 6 | col9a3 |
| FQ323119.13 | 1E-199 | 2.197722 | 0.339 | 0.096 | 4E-195 | DF 6 | FQ323119.1 |
| cxcl14 | 7E-194 | 1.5337326 | 0.56 | 0.239 | 2.4E-189 | DF 6 | cxcl14 |
| hapln1a3 | 2E-192 | 0.9474828 | 0.839 | 0.467 | 5.9E-188 | DF 6 | hapln1a |
| krt83 | 1E-186 | 0.7066351 | 0.995 | 0.984 | 3.7E-182 | DF 6 | krt8 |
| gpm6ab1 | 1E-181 | 2.0841735 | 0.299 | 0.082 | 4.5E-177 | DF 6 | gpm6ab |

| | | | | | | | |
|---|---|---|---|---|---|---|---|
| si:ch211-251b21.14 | 6E-173 | 0.9376827 | 0.799 | 0.44 | 1.9E-168 | DF 6 | si:ch211-251b21.1 |
| pah4 | 4E-169 | 0.9161384 | 0.706 | 0.339 | 1.5E-164 | DF 6 | pah |
| capn124 | 3E-164 | 1.3260612 | 0.525 | 0.216 | 8.7E-160 | DF 6 | capn12 |
| dachd | 4E-158 | 3.4140077 | 0.1 | 0.012 | 1.2E-153 | DF 6 | dachd |
| lmo4b5 | 4E-152 | 1.4684669 | 0.591 | 0.31 | 1.4E-147 | DF 6 | lmo4b |
| cxcl12a6 | 1E-151 | 0.7766314 | 0.921 | 0.675 | 3.4E-147 | DF 6 | cxcl12a |
| zfhx34 | 3E-147 | 1.2460626 | 0.729 | 0.479 | 1E-142 | DF 6 | zfhx3 |
| si:ch211-243a20.33 | 1E-142 | 1.1394867 | 0.578 | 0.273 | 4.8E-138 | DF 6 | si:ch211-243a20.3 |
| alx4a2 | 2E-137 | 1.1996621 | 0.646 | 0.361 | 6.8E-133 | DF 6 | alx4a |
| epas1a1 | 2E-130 | 2.0272591 | 0.22 | 0.059 | 4.9E-126 | DF 6 | epas1a |
| sfrp25 | 3E-128 | 1.1575111 | 0.68 | 0.389 | 1E-123 | DF 6 | sfrp2 |
| twist34 | 4E-125 | 1.3415672 | 0.502 | 0.246 | 1.3E-120 | DF 6 | twist3 |
| id14 | 5E-125 | 1.0899469 | 0.833 | 0.636 | 1.7E-120 | DF 6 | id1 |
| rgs16 | 2E-124 | 2.3464189 | 0.119 | 0.021 | 7.3E-120 | DF 6 | rgs16 |
| ephb2b3 | 4E-123 | 1.6599153 | 0.423 | 0.193 | 1.2E-118 | DF 6 | ephb2b |
| qsox13 | 4E-122 | 2.0650582 | 0.223 | 0.064 | 1.5E-117 | DF 6 | qsox1 |
| hpdb6 | 8E-122 | 0.5823678 | 0.833 | 0.499 | 2.8E-117 | DF 6 | hpdb |
| bco2a1 | 4E-121 | 1.8353557 | 0.354 | 0.143 | 1.4E-116 | DF 6 | bco2a |
| pou3f3b1 | 2E-120 | 2.2883114 | 0.195 | 0.052 | 7E-116 | DF 6 | pou3f3b |
| postna7 | 2E-117 | 0.9541186 | 0.827 | 0.584 | 5.4E-113 | DF 6 | postna |
| tgfbi3 | 2E-115 | 1.1845111 | 0.435 | 0.193 | 6.2E-111 | DF 6 | tgfbi |
| tshz3b1 | 1E-112 | 2.1724485 | 0.203 | 0.058 | 4.8E-108 | DF 6 | tshz3b |
| myh7ba | 6E-111 | 2.3073229 | 0.188 | 0.051 | 2E-106 | DF 6 | myh7ba |
| abi3bpb2 | 4E-108 | 0.7872653 | 0.815 | 0.579 | 1.4E-103 | DF 6 | abi3bpb |
| si:ch211-106h4.123 | 3E-106 | 0.5836536 | 0.761 | 0.449 | 1.1E-101 | DF 6 | si:ch211-106h4.12 |
| BX663503.34 | 2E-105 | 1.0295609 | 0.646 | 0.395 | 5.3E-101 | DF 6 | BX663503.3 |
| enpp23 | 6E-105 | 1.0818913 | 0.666 | 0.438 | 1.9E-100 | DF 6 | enpp2 |
| ttc366 | 1E-104 | 1.0129969 | 0.606 | 0.346 | 3.3E-100 | DF 6 | ttc36 |
| dok4 | 4E-103 | 2.9926764 | 0.101 | 0.018 | 1.38E-98 | DF 6 | dok4 |
| sik14 | 5E-101 | 1.3016352 | 0.473 | 0.247 | 1.61E-96 | DF 6 | sik1 |
| scml2 | 5.6E-98 | 2.6872152 | 0.105 | 0.02 | 1.81E-93 | DF 6 | scml2 |
| adamts25 | 2.2E-95 | 1.1423317 | 0.593 | 0.367 | 7.03E-91 | DF 6 | adamts2 |
| nt5c3a2 | 1E-94 | 1.5568139 | 0.358 | 0.164 | 3.25E-90 | DF 6 | nt5c3a |
| FQ323156.15 | 4.3E-93 | 1.3482012 | 0.395 | 0.189 | 1.38E-88 | DF 6 | FQ323156.1 |
| irx5a2 | 8.7E-93 | 1.8520982 | 0.239 | 0.086 | 2.81E-88 | DF 6 | irx5a |
| mab21l22 | 2.1E-92 | 1.1705389 | 0.545 | 0.325 | 6.71E-88 | DF 6 | mab21l2 |
| emilin1a3 | 5.4E-92 | 1.6852154 | 0.247 | 0.089 | 1.74E-87 | DF 6 | emilin1a |
| en1b3 | 1.8E-91 | 1.5154268 | 0.274 | 0.105 | 5.77E-87 | DF 6 | en1b |
| CABZ01044746.1 | 2.6E-91 | 2.4204488 | 0.125 | 0.029 | 8.54E-87 | DF 6 | CABZ01044746.1 |
| ecrg4a2 | 5.7E-90 | 0.2693833 | 0.814 | 0.556 | 1.87E-85 | DF 6 | ecrg4a |
| arhgef7a1 | 4.3E-89 | 2.2178641 | 0.147 | 0.039 | 1.41E-84 | DF 6 | arhgef7a |
| tob1a3 | 3.9E-87 | 0.9006087 | 0.749 | 0.605 | 1.27E-82 | DF 6 | tob1a |
| cd248a3 | 1.1E-83 | 1.6917139 | 0.274 | 0.112 | 3.69E-79 | DF 6 | cd248a |
| CR383676.25 | 1.3E-82 | 0.3517067 | 1 | 1 | 4.14E-78 | DF 6 | CR383676.2 |
| pnrc23 | 8.3E-82 | 0.6110815 | 0.927 | 0.88 | 2.71E-77 | DF 6 | pnrc2 |
| si:dkey-85k7.74 | 3.1E-81 | 1.0059726 | 0.493 | 0.274 | 9.97E-77 | DF 6 | si:dkey-85k7.7 |
| ckbb2 | 5.9E-81 | 1.2493016 | 0.386 | 0.193 | 1.93E-76 | DF 6 | ckbb |
| clcf11 | 2.1E-80 | 2.056742 | 0.158 | 0.047 | 6.7E-76 | DF 6 | clcf1 |

| | | | | | | | |
|---|---|---|---|---|---|---|---|
| pkdcca | 9.9E-79 | 1.973281 | 0.192 | 0.066 | 3.21E-74 | DF 6 | pkdcca |
| svild2 | 6.1E-78 | 1.2761337 | 0.443 | 0.251 | 1.97E-73 | DF 6 | svild |
| meis2a.13 | 2.5E-76 | 1.5221606 | 0.292 | 0.129 | 8.28E-72 | DF 6 | meis2a.1 |
| vwa12 | 3.3E-73 | 1.2911649 | 0.297 | 0.134 | 1.07E-68 | DF 6 | vwa1 |
| ednraa4 | 8.7E-68 | 1.1570552 | 0.388 | 0.208 | 2.82E-63 | DF 6 | ednraa |
| egfl63 | 2E-67 | 1.7347813 | 0.159 | 0.052 | 6.35E-63 | DF 6 | egfl6 |
| barx14 | 9.7E-67 | 1.3077442 | 0.276 | 0.124 | 3.15E-62 | DF 6 | barx1 |
| junbb3 | 7.3E-66 | 0.8729777 | 0.717 | 0.598 | 2.37E-61 | DF 6 | junbb |
| inhbaa1 | 7.2E-65 | 0.6821422 | 0.463 | 0.267 | 2.34E-60 | DF 6 | inhbaa |
| b2ml4 | 5.4E-63 | 0.4439327 | 0.956 | 0.896 | 1.77E-58 | DF 6 | b2ml |
| tcf124 | 8.3E-63 | 0.7864429 | 0.681 | 0.518 | 2.7E-58 | DF 6 | tcf12 |
| ppp1r1c1 | 5.5E-62 | 0.8252434 | 0.282 | 0.127 | 1.79E-57 | DF 6 | ppp1r1c |
| junba3 | 6.4E-62 | 0.6505099 | 0.876 | 0.845 | 2.08E-57 | DF 6 | junba |
| slc22a24 | 2.5E-61 | 1.291605 | 0.354 | 0.19 | 7.99E-57 | DF 6 | slc22a2 |
| CABZ01092746.15 | 3.6E-61 | 0.4295688 | 0.886 | 0.704 | 1.18E-56 | DF 6 | CABZ01092746.1 |
| cald1b6 | 2.1E-60 | 0.9375548 | 0.326 | 0.163 | 6.82E-56 | DF 6 | cald1b |
| socs1a4 | 1.3E-59 | 1.5181054 | 0.259 | 0.12 | 4.09E-55 | DF 6 | socs1a |
| dll4 | 1.8E-57 | 1.8004605 | 0.127 | 0.04 | 5.74E-53 | DF 6 | dll4 |
| fgfbp2a5 | 2.1E-56 | 0.4009292 | 0.753 | 0.529 | 6.94E-52 | DF 6 | fgfbp2a |
| nanos13 | 3.7E-55 | 1.2609072 | 0.356 | 0.201 | 1.19E-50 | DF 6 | nanos1 |
| ptk7a4 | 1.3E-53 | 1.0445108 | 0.466 | 0.31 | 4.11E-49 | DF 6 | ptk7a |
| sult1st63 | 1.4E-53 | 1.3351162 | 0.242 | 0.113 | 4.71E-49 | DF 6 | sult1st6 |
| cbsb5 | 2.6E-53 | 1.3869737 | 0.271 | 0.135 | 8.35E-49 | DF 6 | cbsb |
| fam169aa4 | 3.1E-53 | 0.6655199 | 0.584 | 0.399 | 9.98E-49 | DF 6 | fam169aa |
| sfrp1b5 | 1.2E-52 | 0.6539141 | 0.546 | 0.356 | 3.78E-48 | DF 6 | sfrp1b |
| mllt3 | 5.5E-51 | 1.8811185 | 0.111 | 0.035 | 1.8E-46 | DF 6 | mllt3 |
| agrn2 | 6.3E-51 | 1.293704 | 0.322 | 0.181 | 2.04E-46 | DF 6 | agrn |
| twist1a3 | 8E-51 | 0.7900489 | 0.589 | 0.434 | 2.59E-46 | DF 6 | twist1a |
| pdgfra5 | 2.3E-50 | 0.7348266 | 0.62 | 0.456 | 7.33E-46 | DF 6 | pdgfra |
| si:ch73-1a9.33 | 7.6E-50 | 0.4987673 | 0.833 | 0.716 | 2.46E-45 | DF 6 | si:ch73-1a9.3 |
| zgc:1626124 | 7.8E-50 | 1.3312775 | 0.238 | 0.113 | 2.53E-45 | DF 6 | zgc:162612 |
| cpe2 | 7.8E-50 | 1.1293844 | 0.345 | 0.199 | 2.55E-45 | DF 6 | cpe |
| csf1b2 | 1.6E-49 | 1.7285982 | 0.12 | 0.04 | 5.05E-45 | DF 6 | csf1b |
| prrx1b4 | 1.8E-49 | 0.7603444 | 0.544 | 0.375 | 5.95E-45 | DF 6 | prrx1b |
| lxn4 | 1.9E-49 | 0.4497523 | 0.83 | 0.698 | 6.25E-45 | DF 6 | lxn |
| CR936442.12 | 5.6E-48 | 1.5976031 | 0.326 | 0.187 | 1.83E-43 | DF 6 | CR936442.1 |
| zgc:1748883 | 1.1E-47 | 0.9912159 | 0.363 | 0.213 | 3.71E-43 | DF 6 | zgc:174888 |
| adma5 | 2.1E-47 | 0.8135342 | 0.393 | 0.229 | 6.83E-43 | DF 6 | adma |
| si:ch211-246m6.52 | 3.8E-47 | 1.5408758 | 0.14 | 0.052 | 1.23E-42 | DF 6 | si:ch211-246m6.5 |
| gfra1a1 | 9.1E-47 | 1.7189678 | 0.141 | 0.053 | 2.97E-42 | DF 6 | gfra1a |
| cebpb3 | 3.3E-46 | 1.14102 | 0.572 | 0.445 | 1.06E-41 | DF 6 | cebpb |
| itgb42 | 4.2E-46 | 0.9021438 | 0.465 | 0.308 | 1.38E-41 | DF 6 | itgb4 |
| auts2a3 | 4.6E-46 | 1.1113258 | 0.36 | 0.219 | 1.5E-41 | DF 6 | auts2a |
| si:dkeyp-72e1.91 | 1.1E-45 | 1.9792587 | 0.104 | 0.033 | 3.63E-41 | DF 6 | si:dkeyp-72e1.9 |
| hmgn34 | 1.5E-45 | 1.2122951 | 0.289 | 0.159 | 4.83E-41 | DF 6 | hmgn3 |
| abca1b1 | 6.6E-44 | 1.9485068 | 0.123 | 0.045 | 2.15E-39 | DF 6 | abca1b |
| eif12 | 1E-43 | 0.632565 | 0.86 | 0.829 | 3.39E-39 | DF 6 | eif1 |
| mycb3 | 1.2E-43 | 0.9510502 | 0.721 | 0.662 | 3.92E-39 | DF 6 | mycb |

| | | | | | | | |
|---|---|---|---|---|---|---|---|
| vmo1a3 | 1.7E-43 | 0.2882289 | 0.895 | 0.695 | 5.67E-39 | DF 6 | vmo1a |
| marcksl1a3 | 3.5E-43 | 0.2729913 | 0.836 | 0.704 | 1.13E-38 | DF 6 | marcksl1a |
| zeb2a3 | 1.7E-42 | 0.904045 | 0.474 | 0.338 | 5.44E-38 | DF 6 | zeb2a |
| nfe2l2a5 | 3.4E-42 | 0.8016865 | 0.484 | 0.34 | 1.09E-37 | DF 6 | nfe2l2a |
| fgfbp1b.1 | 8.8E-42 | 2.0570808 | 0.113 | 0.04 | 2.86E-37 | DF 6 | fgfbp1b.1 |
| mfap55 | 1.3E-41 | 0.4124574 | 0.716 | 0.541 | 4.11E-37 | DF 6 | mfap5 |
| inka1a3 | 1.3E-41 | 1.1846235 | 0.299 | 0.172 | 4.19E-37 | DF 6 | inka1a |
| CABZ01068358.13 | 1.5E-41 | 1.0879459 | 0.337 | 0.204 | 4.9E-37 | DF 6 | CABZ01068358.1 |
| sdc42 | 3.4E-41 | 0.8238868 | 0.762 | 0.703 | 1.12E-36 | DF 6 | sdc4 |
| saa1 | 3.7E-41 | 2.4964415 | 0.122 | 0.045 | 1.19E-36 | DF 6 | saa |
| rgs43 | 7E-41 | 1.3240531 | 0.16 | 0.068 | 2.26E-36 | DF 6 | rgs4 |
| ephb3a3 | 7.6E-41 | 1.3486077 | 0.212 | 0.104 | 2.47E-36 | DF 6 | ephb3a |
| zgc:565856 | 1E-40 | 0.7782805 | 0.477 | 0.324 | 3.33E-36 | DF 6 | zgc:56585 |
| runx34 | 1.1E-40 | 0.8955147 | 0.408 | 0.268 | 3.47E-36 | DF 6 | runx3 |
| lef13 | 2.5E-40 | 1.2999022 | 0.212 | 0.105 | 8.14E-36 | DF 6 | lef1 |
| hspb13 | 4.3E-40 | 1.174295 | 0.304 | 0.176 | 1.39E-35 | DF 6 | hspb1 |
| si:ch211-286o17.13 | 4.9E-40 | 1.140686 | 0.342 | 0.213 | 1.59E-35 | DF 6 | si:ch211-286o17.1 |
| plpp35 | 6.4E-40 | 0.4939248 | 0.619 | 0.45 | 2.09E-35 | DF 6 | plpp3 |
| bcl11ba | 4.2E-39 | 1.6694473 | 0.114 | 0.042 | 1.38E-34 | DF 6 | bcl11ba |
| gsta.14 | 9.8E-39 | 0.858449 | 0.485 | 0.354 | 3.18E-34 | DF 6 | gsta.1 |
| ptgdsb.15 | 1.2E-38 | 0.6941303 | 0.602 | 0.459 | 4.05E-34 | DF 6 | ptgdsb.1 |
| nr2f23 | 1.5E-38 | 1.2238899 | 0.22 | 0.112 | 4.96E-34 | DF 6 | nr2f2 |
| smoc22 | 4.3E-38 | 1.2806488 | 0.185 | 0.088 | 1.39E-33 | DF 6 | smoc2 |
| angptl54 | 5.5E-38 | 0.9669235 | 0.295 | 0.17 | 1.78E-33 | DF 6 | angptl5 |
| aldob5 | 8.3E-38 | 0.8043859 | 0.449 | 0.307 | 2.7E-33 | DF 6 | aldob |
| rem13 | 1.9E-37 | 1.49626 | 0.146 | 0.062 | 6.12E-33 | DF 6 | rem1 |
| zgc:1943984 | 3.8E-37 | 0.6557686 | 0.529 | 0.379 | 1.23E-32 | DF 6 | zgc:194398 |
| tuba8l22 | 6.2E-37 | 1.1059141 | 0.336 | 0.214 | 2.01E-32 | DF 6 | tuba8l2 |
| zbtb16a2 | 7.7E-37 | 1.2521144 | 0.236 | 0.127 | 2.51E-32 | DF 6 | zbtb16a |
| zgc:1135314 | 1.2E-36 | 0.8845483 | 0.404 | 0.275 | 3.82E-32 | DF 6 | zgc:113531 |
| ptx3a5 | 4.3E-36 | 0.440483 | 0.657 | 0.482 | 1.4E-31 | DF 6 | ptx3a |
| prss234 | 4.4E-36 | 1.0935176 | 0.318 | 0.197 | 1.43E-31 | DF 6 | prss23 |
| hsp70.12 | 4.6E-36 | 0.7343136 | 0.475 | 0.338 | 1.49E-31 | DF 6 | hsp70.1 |
| cfp5 | 4.9E-36 | 0.6481388 | 0.486 | 0.336 | 1.58E-31 | DF 6 | cfp |
| snai1a2 | 7.6E-36 | 0.9752161 | 0.39 | 0.269 | 2.48E-31 | DF 6 | snai1a |
| rbp45 | 1.1E-35 | 0.2083878 | 0.986 | 0.907 | 3.42E-31 | DF 6 | rbp4 |
| id2a4 | 1.3E-35 | 0.6718865 | 0.612 | 0.5 | 4.1E-31 | DF 6 | id2a |
| neu3.52 | 2.5E-35 | 1.1859571 | 0.153 | 0.067 | 8.24E-31 | DF 6 | neu3.5 |
| sema3aa3 | 5.1E-35 | 1.4048637 | 0.204 | 0.106 | 1.66E-30 | DF 6 | sema3aa |
| btg13 | 1.2E-34 | 0.4002564 | 0.856 | 0.814 | 3.97E-30 | DF 6 | btg1 |
| btg23 | 1.8E-34 | 0.5167982 | 0.762 | 0.698 | 5.95E-30 | DF 6 | btg2 |
| smoc14 | 2.5E-34 | 0.8709525 | 0.331 | 0.204 | 8.15E-30 | DF 6 | smoc1 |
| ucp25 | 3.3E-34 | 0.3777465 | 0.56 | 0.411 | 1.08E-29 | DF 6 | ucp2 |
| neu3.25 | 6.5E-34 | 0.8974756 | 0.345 | 0.219 | 2.13E-29 | DF 6 | neu3.2 |
| adamts15b.14 | 1.2E-33 | 1.125618 | 0.185 | 0.09 | 3.76E-29 | DF 6 | adamts15b.1 |
| capn2b6 | 1.5E-33 | 0.7556339 | 0.448 | 0.319 | 4.83E-29 | DF 6 | capn2b |
| eef1da3 | 2.1E-33 | 0.3779452 | 0.885 | 0.876 | 6.9E-29 | DF 6 | eef1da |
| dhrs3a3 | 9.5E-33 | 1.2936143 | 0.146 | 0.066 | 3.1E-28 | DF 6 | dhrs3a |

| | | | | | | | |
|---|---|---|---|---|---|---|---|
| hsp90aa1.22 | 1.1E-32 | 0.7149352 | 0.702 | 0.597 | 3.42E-28 | DF 6 | hsp90aa1.2 |
| sgip1a2 | 1.6E-32 | 1.4861122 | 0.121 | 0.05 | 5.17E-28 | DF 6 | sgip1a |
| cdo14 | 3.3E-32 | 0.7539862 | 0.453 | 0.326 | 1.08E-27 | DF 6 | cdo1 |
| hmgb1a1 | 6.1E-32 | 0.6063437 | 0.652 | 0.562 | 1.97E-27 | DF 6 | hmgb1a |
| hmgb3a2 | 6.1E-32 | 1.1379411 | 0.211 | 0.113 | 1.99E-27 | DF 6 | hmgb3a |
| cebpd5 | 7.2E-32 | 0.503357 | 0.689 | 0.574 | 2.34E-27 | DF 6 | cebpd |
| ada3 | 9.1E-32 | 0.9714797 | 0.334 | 0.218 | 2.95E-27 | DF 6 | ada |
| sept121 | 1.4E-31 | 1.5771353 | 0.105 | 0.041 | 4.46E-27 | DF 6 | sept12 |
| hist1h4l.183 | 3E-31 | 0.9486243 | 0.219 | 0.118 | 9.68E-27 | DF 6 | hist1h4l.18 |
| hsp70l2 | 2.7E-30 | 0.9699903 | 0.419 | 0.299 | 8.65E-26 | DF 6 | hsp70l |
| mid1ip1l2 | 3.8E-30 | 1.0208755 | 0.297 | 0.188 | 1.25E-25 | DF 6 | mid1ip1l |
| fosab3 | 8E-30 | 0.5464915 | 0.895 | 0.895 | 2.6E-25 | DF 6 | fosab |
| CABZ01085139.11 | 1.2E-29 | 1.2354578 | 0.138 | 0.062 | 4.04E-25 | DF 6 | CABZ01085139.1 |
| fbp1a3 | 1.7E-29 | 0.7990829 | 0.388 | 0.274 | 5.38E-25 | DF 6 | fbp1a |
| aoc24 | 3.3E-29 | 0.5101054 | 0.368 | 0.238 | 1.07E-24 | DF 6 | aoc2 |
| col6a3.13 | 3.5E-29 | 0.268377 | 0.984 | 0.937 | 1.15E-24 | DF 6 | col6a3.1 |
| pim23 | 6.4E-29 | 0.9248294 | 0.334 | 0.223 | 2.07E-24 | DF 6 | pim2 |
| gstp15 | 1.1E-28 | 0.416361 | 0.742 | 0.641 | 3.74E-24 | DF 6 | gstp1 |
| tnfrsf193 | 1.5E-28 | 1.3050122 | 0.195 | 0.106 | 4.83E-24 | DF 6 | tnfrsf19 |
| hapln1b | 6E-28 | 1.034019 | 0.167 | 0.085 | 1.96E-23 | DF 6 | hapln1b |
| gpc1b2 | 7.9E-28 | 0.7015829 | 0.507 | 0.399 | 2.56E-23 | DF 6 | gpc1b |
| etnk13 | 1.1E-27 | 0.957264 | 0.29 | 0.186 | 3.58E-23 | DF 6 | etnk1 |
| wnt5b2 | 1.2E-27 | 1.2211789 | 0.195 | 0.108 | 3.94E-23 | DF 6 | wnt5b |
| si:ch1073-459j12.13 | 2.3E-27 | 0.2328221 | 0.916 | 0.808 | 7.32E-23 | DF 6 | si:ch1073-459j12.1 |
| gbe1a4 | 4.6E-27 | 0.9628532 | 0.201 | 0.11 | 1.49E-22 | DF 6 | gbe1a |
| qdpra5 | 5.5E-27 | 0.3122395 | 0.576 | 0.422 | 1.78E-22 | DF 6 | qdpra |
| cdh115 | 6.2E-27 | 0.4434037 | 0.703 | 0.587 | 2.03E-22 | DF 6 | cdh11 |
| nfia3 | 1.4E-26 | 0.9725469 | 0.267 | 0.167 | 4.47E-22 | DF 6 | nfia |
| frem1b3 | 2.2E-26 | 1.25338 | 0.151 | 0.076 | 7.32E-22 | DF 6 | frem1b |
| fosl1a2 | 6.3E-26 | 0.7415943 | 0.527 | 0.436 | 2.06E-21 | DF 6 | fosl1a |
| bhlhe404 | 6.7E-26 | 0.6556523 | 0.557 | 0.463 | 2.17E-21 | DF 6 | bhlhe40 |
| cavin1b2 | 3.2E-25 | 0.4136036 | 0.741 | 0.644 | 1.04E-20 | DF 6 | cavin1b |
| ablim1b4 | 5E-25 | 0.5864918 | 0.367 | 0.253 | 1.62E-20 | DF 6 | ablim1b |
| rsrp13 | 8.9E-25 | 0.5996789 | 0.611 | 0.541 | 2.88E-20 | DF 6 | rsrp1 |
| bmp1a3 | 9.2E-25 | 1.083181 | 0.277 | 0.185 | 2.98E-20 | DF 6 | bmp1a |
| zbtb202 | 2.2E-24 | 0.7454677 | 0.467 | 0.373 | 7.07E-20 | DF 6 | zbtb20 |
| slc44a1b4 | 2.5E-24 | 1.1708309 | 0.202 | 0.118 | 8.14E-20 | DF 6 | slc44a1b |
| tob1b4 | 5.4E-24 | 0.3278466 | 0.776 | 0.701 | 1.76E-19 | DF 6 | tob1b |
| zgc:1133072 | 9.9E-24 | 1.0104133 | 0.168 | 0.091 | 3.22E-19 | DF 6 | zgc:113307 |
| eif4ebp33 | 1.6E-23 | 0.9772225 | 0.264 | 0.172 | 5.16E-19 | DF 6 | eif4ebp3 |
| lrba1 | 1.6E-23 | 1.0568596 | 0.267 | 0.175 | 5.27E-19 | DF 6 | lrba |
| mknk2b3 | 1.7E-23 | 0.4552798 | 0.675 | 0.606 | 5.6E-19 | DF 6 | mknk2b |
| gstz14 | 2.2E-23 | 0.5899063 | 0.448 | 0.342 | 7.28E-19 | DF 6 | gstz1 |
| rhoq3 | 4.4E-23 | 1.1489706 | 0.17 | 0.094 | 1.43E-18 | DF 6 | rhoq |
| ism2a2 | 6.8E-23 | 1.3613315 | 0.11 | 0.051 | 2.2E-18 | DF 6 | ism2a |
| ccdc80l14 | 7.9E-23 | 1.1250141 | 0.148 | 0.077 | 2.57E-18 | DF 6 | ccdc80l1 |
| si:dkey-195m11.84 | 5E-22 | 0.9535904 | 0.209 | 0.125 | 1.63E-17 | DF 6 | si:dkey-195m11.8 |
| DDX171 | 6.2E-22 | 0.7331886 | 0.393 | 0.298 | 2.02E-17 | DF 6 | DDX17 |

| | | | | | | | |
|---|---|---|---|---|---|---|---|
| hsp70.22 | 1.7E-21 | 0.7109569 | 0.579 | 0.491 | 5.65E-17 | DF 6 | hsp70.2 |
| fam43a4 | 2.4E-21 | 1.1929099 | 0.13 | 0.067 | 7.74E-17 | DF 6 | fam43a |
| hopx3 | 2.5E-21 | 0.6489195 | 0.369 | 0.268 | 8.26E-17 | DF 6 | hopx |
| crhr12 | 3.1E-21 | 1.1217484 | 0.145 | 0.077 | 1.02E-16 | DF 6 | crhr1 |
| c4b1 | 4.2E-21 | 1.3367307 | 0.111 | 0.054 | 1.37E-16 | DF 6 | c4b |
| gstm.15 | 5.4E-21 | 0.3714207 | 0.677 | 0.595 | 1.76E-16 | DF 6 | gstm.1 |
| hmox1a4 | 8.1E-21 | 1.017267 | 0.259 | 0.172 | 2.64E-16 | DF 6 | hmox1a |
| crabp2b3 | 8.4E-21 | 0.3513578 | 0.527 | 0.417 | 2.73E-16 | DF 6 | crabp2b |
| ak44 | 9.8E-21 | 0.8912294 | 0.27 | 0.183 | 3.19E-16 | DF 6 | ak4 |
| gadd45ga4 | 1.9E-20 | 0.8722347 | 0.366 | 0.275 | 6.3E-16 | DF 6 | gadd45ga |
| mgst1.23 | 2E-20 | 0.5592004 | 0.4 | 0.298 | 6.35E-16 | DF 6 | mgst1.2 |
| bnc24 | 5.4E-20 | 1.2053137 | 0.146 | 0.08 | 1.76E-15 | DF 6 | bnc2 |
| alx4b2 | 7E-20 | 1.3526933 | 0.113 | 0.057 | 2.26E-15 | DF 6 | alx4b |
| bmp41 | 8E-20 | 1.3334486 | 0.124 | 0.064 | 2.59E-15 | DF 6 | bmp4 |
| plecb1 | 8.1E-20 | 0.3735554 | 0.728 | 0.656 | 2.64E-15 | DF 6 | plecb |
| spink2.12 | 1.3E-19 | 0.9055266 | 0.206 | 0.127 | 4.14E-15 | DF 6 | spink2.1 |
| sept9a2 | 1.3E-19 | 0.772056 | 0.3 | 0.214 | 4.35E-15 | DF 6 | sept9a |
| il1rapl1a | 1.7E-19 | 1.1721679 | 0.104 | 0.05 | 5.64E-15 | DF 6 | il1rapl1a |
| tnfsf103 | 2.4E-19 | 0.9276325 | 0.167 | 0.097 | 7.8E-15 | DF 6 | tnfsf10 |
| thrap3a3 | 2.7E-19 | 0.6084028 | 0.464 | 0.382 | 8.79E-15 | DF 6 | thrap3a |
| C25H11orf964 | 2.8E-19 | 0.3964457 | 0.482 | 0.381 | 9.13E-15 | DF 6 | C25H11orf96 |
| hs3st3b1b2 | 4.7E-19 | 1.0338814 | 0.202 | 0.127 | 1.53E-14 | DF 6 | hs3st3b1b |
| aldh9a1a.14 | 1.2E-18 | 0.3619327 | 0.571 | 0.479 | 4.06E-14 | DF 6 | aldh9a1a.1 |
| ehbp1l1a3 | 1.3E-18 | 0.9523037 | 0.16 | 0.092 | 4.23E-14 | DF 6 | ehbp1l1a |
| kat6b | 1.6E-18 | 1.4017187 | 0.103 | 0.051 | 5.1E-14 | DF 6 | kat6b |
| phldb2b3 | 1.6E-18 | 0.9007018 | 0.219 | 0.142 | 5.15E-14 | DF 6 | phldb2b |
| tfap2c4 | 2.2E-18 | 0.6845295 | 0.281 | 0.194 | 7.03E-14 | DF 6 | tfap2c |
| gpx4a1 | 2.2E-18 | 0.6862923 | 0.387 | 0.301 | 7.3E-14 | DF 6 | gpx4a |
| h2ax13 | 3.5E-18 | 0.3976173 | 0.671 | 0.628 | 1.15E-13 | DF 6 | h2ax1 |
| slc40a12 | 4.6E-18 | 0.7268676 | 0.284 | 0.2 | 1.51E-13 | DF 6 | slc40a1 |
| hyal2b5 | 5.3E-18 | 0.9147848 | 0.275 | 0.195 | 1.73E-13 | DF 6 | hyal2b |
| kank44 | 8.9E-18 | 0.9432288 | 0.17 | 0.102 | 2.88E-13 | DF 6 | kank4 |
| cited4b3 | 9E-18 | 0.8200089 | 0.226 | 0.148 | 2.93E-13 | DF 6 | cited4b |
| slc7a25 | 2.3E-17 | 0.8499238 | 0.237 | 0.157 | 7.47E-13 | DF 6 | slc7a2 |
| cbx7a3 | 2.4E-17 | 0.7025728 | 0.428 | 0.345 | 7.84E-13 | DF 6 | cbx7a |
| efnb2a1 | 2.7E-17 | 0.9074796 | 0.224 | 0.149 | 8.67E-13 | DF 6 | efnb2a |
| prelp3 | 4.4E-17 | 0.5208129 | 0.21 | 0.131 | 1.43E-12 | DF 6 | prelp |
| mcl1a3 | 5.2E-17 | 0.4741322 | 0.776 | 0.771 | 1.71E-12 | DF 6 | mcl1a |
| sulf13 | 5.8E-17 | 0.5842511 | 0.224 | 0.145 | 1.89E-12 | DF 6 | sulf1 |
| dap1b.13 | 6.9E-17 | 0.2971547 | 0.597 | 0.505 | 2.25E-12 | DF 6 | dap1b.1 |
| itm2bb4 | 9.3E-17 | 0.978942 | 0.15 | 0.088 | 3.02E-12 | DF 6 | itm2bb |
| dop1a2 | 1.1E-16 | 1.0511087 | 0.134 | 0.075 | 3.7E-12 | DF 6 | dop1a |
| hspa8b3 | 1.6E-16 | 1.0586288 | 0.154 | 0.092 | 5.3E-12 | DF 6 | hspa8b |
| dedd14 | 1.7E-16 | 0.4268981 | 0.607 | 0.551 | 5.5E-12 | DF 6 | dedd1 |
| tmem200b3 | 2.7E-16 | 0.9607821 | 0.155 | 0.093 | 8.63E-12 | DF 6 | tmem200b |
| col12a1b2 | 3.4E-16 | 0.25318 | 0.73 | 0.621 | 1.1E-11 | DF 6 | col12a1b |
| celf25 | 5.3E-16 | 0.5614872 | 0.281 | 0.196 | 1.72E-11 | DF 6 | celf2 |
| zbtb42 | 7.8E-16 | 0.6187253 | 0.399 | 0.323 | 2.55E-11 | DF 6 | zbtb4 |

| | | | | | | | |
|---|---|---|---|---|---|---|---|
| mylipb2 | 9.5E-16 | 1.11259 | 0.135 | 0.078 | 3.07E-11 | DF 6 | mylipb |
| pcdh18b3 | 1.1E-15 | 0.7487465 | 0.294 | 0.218 | 3.56E-11 | DF 6 | pcdh18b |
| pcdh114 | 1.5E-15 | 0.9607755 | 0.13 | 0.074 | 5.03E-11 | DF 6 | pcdh11 |
| cavin2b2 | 1.8E-15 | 0.3537727 | 0.602 | 0.529 | 5.76E-11 | DF 6 | cavin2b |
| adgrl2a5 | 2.1E-15 | 0.7113444 | 0.265 | 0.189 | 6.94E-11 | DF 6 | adgrl2a |
| glula4 | 3.2E-15 | 0.2816154 | 0.713 | 0.616 | 1.04E-10 | DF 6 | glula |
| caskin12 | 4.8E-15 | 1.2947613 | 0.105 | 0.057 | 1.56E-10 | DF 6 | caskin1 |
| ms4a17a.95 | 1.3E-14 | 0.658826 | 0.292 | 0.215 | 4.11E-10 | DF 6 | ms4a17a.9 |
| ntd54 | 1.9E-14 | 0.5694781 | 0.212 | 0.141 | 6.15E-10 | DF 6 | ntd5 |
| adamts15a5 | 2.5E-14 | 0.8611966 | 0.209 | 0.142 | 7.99E-10 | DF 6 | adamts15a |
| wnt114 | 2.5E-14 | 0.2647381 | 0.376 | 0.282 | 8.13E-10 | DF 6 | wnt11 |
| rnf213a3 | 2.8E-14 | 0.5505379 | 0.345 | 0.267 | 9.12E-10 | DF 6 | rnf213a |
| cxcl18b3 | 3.2E-14 | 0.3754133 | 0.751 | 0.701 | 1.05E-09 | DF 6 | cxcl18b |
| ccdc80l22 | 3.3E-14 | 0.6100289 | 0.178 | 0.113 | 1.07E-09 | DF 6 | ccdc80l2 |
| fgfr1b3 | 3.3E-14 | 1.0064622 | 0.159 | 0.1 | 1.07E-09 | DF 6 | fgfr1b |
| hsp70.32 | 3.7E-14 | 0.6453634 | 0.607 | 0.553 | 1.21E-09 | DF 6 | hsp70.3 |
| tnrc184 | 3.8E-14 | 0.5709859 | 0.467 | 0.406 | 1.24E-09 | DF 6 | tnrc18 |
| ecm1b4 | 4.3E-14 | 0.2749467 | 0.565 | 0.469 | 1.41E-09 | DF 6 | ecm1b |
| ninl1 | 6.1E-14 | 1.1324893 | 0.118 | 0.068 | 1.98E-09 | DF 6 | ninl |
| mhc1zba3 | 7E-14 | 0.3044134 | 0.64 | 0.583 | 2.29E-09 | DF 6 | mhc1zba |
| runx1t13 | 7.1E-14 | 0.9479689 | 0.146 | 0.09 | 2.32E-09 | DF 6 | runx1t1 |
| dio3b1 | 8.9E-14 | 1.1631665 | 0.1 | 0.054 | 2.9E-09 | DF 6 | dio3b |
| edil3a4 | 9.1E-14 | 0.2855609 | 0.52 | 0.433 | 2.97E-09 | DF 6 | edil3a |
| tuba8l42 | 1.1E-13 | 0.2559242 | 0.756 | 0.733 | 3.42E-09 | DF 6 | tuba8l4 |
| gpatch82 | 1.2E-13 | 0.5779031 | 0.397 | 0.328 | 3.75E-09 | DF 6 | gpatch8 |
| kif26ba5 | 1.2E-13 | 0.7447069 | 0.168 | 0.107 | 3.97E-09 | DF 6 | kif26ba |
| igsf103 | 1.2E-13 | 0.4388164 | 0.363 | 0.276 | 4.02E-09 | DF 6 | igsf10 |
| map2k63 | 1.3E-13 | 0.7082058 | 0.258 | 0.188 | 4.11E-09 | DF 6 | map2k6 |
| col7a13 | 1.3E-13 | 0.321958 | 0.228 | 0.153 | 4.16E-09 | DF 6 | col7a1 |
| im:71523484 | 1.6E-13 | 0.8046517 | 0.306 | 0.235 | 5.12E-09 | DF 6 | im:7152348 |
| prickle1a2 | 1.9E-13 | 1.0950546 | 0.104 | 0.058 | 6.26E-09 | DF 6 | prickle1a |
| jdp2b3 | 2.3E-13 | 0.6146112 | 0.61 | 0.571 | 7.51E-09 | DF 6 | jdp2b |
| srsf3b2 | 3.2E-13 | 0.6682343 | 0.34 | 0.273 | 1.03E-08 | DF 6 | srsf3b |
| CR318588.12 | 3.2E-13 | 1.4079251 | 0.359 | 0.294 | 1.04E-08 | DF 6 | CR318588.1 |
| sf3b11 | 3.3E-13 | 0.6286486 | 0.401 | 0.337 | 1.06E-08 | DF 6 | sf3b1 |
| si:ch73-335l21.43 | 4.9E-13 | 0.836107 | 0.588 | 0.571 | 1.58E-08 | DF 6 | si:ch73-335l21.4 |
| srsf5a2 | 5.8E-13 | 0.4200301 | 0.609 | 0.579 | 1.87E-08 | DF 6 | srsf5a |
| bmp61 | 5.8E-13 | 0.9546581 | 0.105 | 0.059 | 1.88E-08 | DF 6 | bmp6 |
| tpst13 | 9.4E-13 | 0.7074967 | 0.239 | 0.175 | 3.07E-08 | DF 6 | tpst1 |
| mxra5b2 | 1E-12 | 0.8252479 | 0.105 | 0.059 | 3.34E-08 | DF 6 | mxra5b |
| gpr1433 | 1.3E-12 | 0.8630682 | 0.118 | 0.07 | 4.35E-08 | DF 6 | gpr143 |
| hist2h2l4 | 1.6E-12 | 0.650633 | 0.43 | 0.374 | 5.27E-08 | DF 6 | hist2h2l |
| fgfr23 | 1.8E-12 | 0.4855632 | 0.417 | 0.347 | 5.92E-08 | DF 6 | fgfr2 |
| gse13 | 2.6E-12 | 0.9412662 | 0.181 | 0.124 | 8.47E-08 | DF 6 | gse1 |
| mdh1aa3 | 2.8E-12 | 0.4644223 | 0.431 | 0.373 | 9.05E-08 | DF 6 | mdh1aa |
| ldhba3 | 3.2E-12 | 0.4175017 | 0.536 | 0.493 | 1.04E-07 | DF 6 | ldhba |
| sdc22 | 3.3E-12 | 0.3736199 | 0.632 | 0.587 | 1.09E-07 | DF 6 | sdc2 |
| cd82a4 | 4.2E-12 | 0.4131194 | 0.426 | 0.354 | 1.35E-07 | DF 6 | cd82a |

| | | | | | | | |
|---|---|---|---|---|---|---|---|
| SSBP42 | 4.2E-12 | 0.8545002 | 0.177 | 0.12 | 1.36E-07 | DF 6 | SSBP4 |
| raraa5 | 4.5E-12 | 0.5871912 | 0.321 | 0.252 | 1.47E-07 | DF 6 | raraa |
| rasl11b5 | 5.9E-12 | 0.6808221 | 0.279 | 0.215 | 1.91E-07 | DF 6 | rasl11b |
| ccnl1a3 | 7.3E-12 | 0.7779336 | 0.321 | 0.261 | 2.37E-07 | DF 6 | ccnl1a |
| tent5ab2 | 1.1E-11 | 1.0921579 | 0.1 | 0.058 | 3.71E-07 | DF 6 | tent5ab |
| fgfr1a3 | 1.3E-11 | 0.8725565 | 0.213 | 0.156 | 4.08E-07 | DF 6 | fgfr1a |
| ddx63 | 1.5E-11 | 0.4953389 | 0.408 | 0.347 | 4.77E-07 | DF 6 | ddx6 |
| sox11b3 | 1.5E-11 | 0.6104563 | 0.273 | 0.209 | 4.97E-07 | DF 6 | sox11b |
| plekhg2 | 1.7E-11 | 1.0128421 | 0.118 | 0.072 | 5.6E-07 | DF 6 | plekhg2 |
| psd23 | 1.9E-11 | 0.8854515 | 0.148 | 0.096 | 6.31E-07 | DF 6 | psd2 |
| atrx1 | 2.3E-11 | 0.4349812 | 0.515 | 0.466 | 7.33E-07 | DF 6 | atrx |
| gask1a3 | 2.9E-11 | 0.8469398 | 0.251 | 0.194 | 9.28E-07 | DF 6 | gask1a |
| ANPEP2 | 3.2E-11 | 0.6550775 | 0.271 | 0.209 | 1.03E-06 | DF 6 | ANPEP |
| si:ch73-237c6.15 | 3.9E-11 | 0.3903963 | 0.434 | 0.364 | 1.28E-06 | DF 6 | si:ch73-237c6.1 |
| csnk1da3 | 4.4E-11 | 0.5804594 | 0.274 | 0.211 | 1.42E-06 | DF 6 | csnk1da |
| hhla2a.12 | 7.8E-11 | 0.8352114 | 0.128 | 0.081 | 2.53E-06 | DF 6 | hhla2a.1 |
| chd55 | 1E-10 | 0.7260724 | 0.18 | 0.126 | 3.41E-06 | DF 6 | chd5 |
| zbtb16b3 | 1.5E-10 | 0.7905624 | 0.153 | 0.103 | 4.73E-06 | DF 6 | zbtb16b |
| ENSDARG00000762 | 1.7E-10 | 0.2964327 | 0.646 | 0.624 | 5.52E-06 | DF 6 | ENSDARG00000076 |
| klf7b3 | 1.9E-10 | 0.7063701 | 0.287 | 0.227 | 6.02E-06 | DF 6 | klf7b |
| ctbp2a2 | 2.1E-10 | 0.5140269 | 0.292 | 0.229 | 6.92E-06 | DF 6 | ctbp2a |
| ppl4 | 2.6E-10 | 0.5687746 | 0.213 | 0.154 | 8.55E-06 | DF 6 | ppl |
| thbs2a5 | 3.3E-10 | 0.7222229 | 0.262 | 0.202 | 1.08E-05 | DF 6 | thbs2a |
| BX663503.12 | 3.5E-10 | 0.8021363 | 0.108 | 0.066 | 1.13E-05 | DF 6 | BX663503.1 |
| si:rp71-79p20.22 | 4.3E-10 | 0.7625734 | 0.205 | 0.151 | 1.4E-05 | DF 6 | si:rp71-79p20.2 |
| kremen13 | 5E-10 | 0.8772039 | 0.123 | 0.079 | 1.64E-05 | DF 6 | kremen1 |
| ddx55 | 5.9E-10 | 0.503893 | 0.604 | 0.598 | 1.92E-05 | DF 6 | ddx5 |
| fstb6 | 6.3E-10 | 0.3142178 | 0.435 | 0.36 | 2.06E-05 | DF 6 | fstb |
| pik3ip11 | 6.9E-10 | 1.0114271 | 0.117 | 0.075 | 2.23E-05 | DF 6 | pik3ip1 |
| NPAS33 | 7.9E-10 | 0.8440999 | 0.13 | 0.085 | 2.56E-05 | DF 6 | NPAS3 |
| plekhg5a2 | 1E-09 | 1.0341322 | 0.112 | 0.071 | 3.33E-05 | DF 6 | plekhg5a |
| bzw1b2 | 1.3E-09 | 0.5492834 | 0.52 | 0.491 | 4.24E-05 | DF 6 | bzw1b |
| si:ch73-386h18.15 | 1.5E-09 | 0.708772 | 0.2 | 0.146 | 4.84E-05 | DF 6 | si:ch73-386h18.1 |
| robo11 | 2.1E-09 | 0.9734676 | 0.119 | 0.078 | 6.76E-05 | DF 6 | robo1 |
| tnrc6c13 | 2.2E-09 | 0.3981262 | 0.522 | 0.486 | 7.16E-05 | DF 6 | tnrc6c1 |
| smc32 | 2.7E-09 | 0.7483705 | 0.196 | 0.145 | 8.66E-05 | DF 6 | smc3 |
| fat1a2 | 2.8E-09 | 0.6218306 | 0.321 | 0.272 | 9.1E-05 | DF 6 | fat1a |
| ches12 | 3E-09 | 0.7335057 | 0.232 | 0.182 | 9.81E-05 | DF 6 | ches1 |
| gch14 | 3E-09 | 0.6596627 | 0.186 | 0.135 | 9.91E-05 | DF 6 | gch1 |
| BX322530.1 | 3.9E-09 | 0.4150447 | 0.466 | 0.419 | 0.000127 | DF 6 | BX322530.1 |
| acadm | 4.3E-09 | 0.6487136 | 0.301 | 0.254 | 0.00014 | DF 6 | acadm |
| ciarta4 | 4.9E-09 | 0.5466128 | 0.334 | 0.283 | 0.000158 | DF 6 | ciarta |
| dnajb1b3 | 5.2E-09 | 0.3176031 | 0.513 | 0.464 | 0.00017 | DF 6 | dnajb1b |
| myca3 | 5.5E-09 | 0.6866348 | 0.208 | 0.157 | 0.000178 | DF 6 | myca |
| mmp14b4 | 6.5E-09 | 0.2760899 | 0.595 | 0.55 | 0.000212 | DF 6 | mmp14b |
| pcbd15 | 6.5E-09 | 0.2512723 | 0.388 | 0.315 | 0.000212 | DF 6 | pcbd1 |
| axin23 | 6.7E-09 | 0.5878741 | 0.274 | 0.219 | 0.000219 | DF 6 | axin2 |
| pbx42 | 7E-09 | 0.5816549 | 0.309 | 0.258 | 0.000228 | DF 6 | pbx4 |

| | | | | | | | |
|---|---|---|---|---|---|---|---|
| rsl24d14 | 7.2E-09 | 0.2243183 | 0.752 | 0.749 | 0.000234 | DF 6 | rsl24d1 |
| suco4 | 7.3E-09 | 0.646403 | 0.223 | 0.172 | 0.000238 | DF 6 | suco |
| ncl2 | 7.4E-09 | 0.2204734 | 0.631 | 0.604 | 0.000241 | DF 6 | ncl |
| wls3 | 1E-08 | 0.4460026 | 0.433 | 0.394 | 0.00033 | DF 6 | wls |
| slc16a105 | 1.1E-08 | 0.7277569 | 0.18 | 0.132 | 0.000354 | DF 6 | slc16a10 |
| fosb4 | 1.1E-08 | 0.2739333 | 0.718 | 0.692 | 0.000368 | DF 6 | fosb |
| proza1 | 1.1E-08 | 0.7049512 | 0.219 | 0.169 | 0.000371 | DF 6 | proza |
| tgfbr32 | 1.3E-08 | 0.8909579 | 0.119 | 0.079 | 0.000437 | DF 6 | tgfbr3 |
| serpina10a4 | 1.6E-08 | 0.4701469 | 0.421 | 0.378 | 0.000535 | DF 6 | serpina10a |
| cadm43 | 1.9E-08 | 0.8457395 | 0.114 | 0.075 | 0.00062 | DF 6 | cadm4 |
| polr2a4 | 2E-08 | 0.469554 | 0.409 | 0.363 | 0.000635 | DF 6 | polr2a |
| hist1h4l.33 | 2E-08 | 0.5830838 | 0.192 | 0.142 | 0.000645 | DF 6 | hist1h4l.3 |
| nfil3-65 | 2.2E-08 | 0.2911379 | 0.369 | 0.305 | 0.000714 | DF 6 | nfil3-6 |
| tril4 | 2.2E-08 | 0.6704911 | 0.151 | 0.106 | 0.000721 | DF 6 | tril |
| CABZ01068367.14 | 2.7E-08 | 0.3730171 | 0.441 | 0.393 | 0.000894 | DF 6 | CABZ01068367.1 |
| cmbl3 | 3.8E-08 | 0.6094545 | 0.223 | 0.174 | 0.001227 | DF 6 | cmbl |
| ldha1 | 4.1E-08 | 0.4026569 | 0.282 | 0.229 | 0.001318 | DF 6 | ldha |
| ccni1 | 4.2E-08 | 0.282311 | 0.614 | 0.594 | 0.00136 | DF 6 | ccni |
| ubb3 | 4.5E-08 | 0.3218595 | 0.676 | 0.682 | 0.001467 | DF 6 | ubb |
| hhip3 | 4.7E-08 | 0.6090618 | 0.129 | 0.088 | 0.001544 | DF 6 | hhip |
| raph1b3 | 5.1E-08 | 0.8027346 | 0.148 | 0.106 | 0.001654 | DF 6 | raph1b |
| lrrc58b3 | 5.7E-08 | 0.6889439 | 0.166 | 0.122 | 0.001844 | DF 6 | lrrc58b |
| sh3bp5a2 | 5.8E-08 | 0.7339374 | 0.174 | 0.13 | 0.001878 | DF 6 | sh3bp5a |
| zmiz1a1 | 5.9E-08 | 0.7636678 | 0.163 | 0.119 | 0.001907 | DF 6 | zmiz1a |
| slc12a7b2 | 6.6E-08 | 0.6621736 | 0.197 | 0.151 | 0.002145 | DF 6 | slc12a7b |
| hyal43 | 6.7E-08 | 0.5162723 | 0.149 | 0.105 | 0.002179 | DF 6 | hyal4 |
| ube2e32 | 6.9E-08 | 0.5049194 | 0.365 | 0.328 | 0.002244 | DF 6 | ube2e3 |
| ccdc1252 | 6.9E-08 | 0.806026 | 0.111 | 0.074 | 0.00226 | DF 6 | ccdc125 |
| aff24 | 7.8E-08 | 0.6585444 | 0.225 | 0.178 | 0.002542 | DF 6 | aff2 |
| galnt21 | 8.2E-08 | 0.8142033 | 0.128 | 0.09 | 0.00268 | DF 6 | galnt2 |
| nsd1b1 | 8.3E-08 | 0.9070182 | 0.109 | 0.073 | 0.002686 | DF 6 | nsd1b |
| phip3 | 8.4E-08 | 0.616244 | 0.204 | 0.157 | 0.002732 | DF 6 | phip |
| si:ch211-196l7.43 | 8.7E-08 | 0.3806617 | 0.499 | 0.476 | 0.002844 | DF 6 | si:ch211-196l7.4 |
| CR383676.34 | 9.3E-08 | 0.3757037 | 0.502 | 0.466 | 0.003018 | DF 6 | CR383676.3 |
| map1ab1 | 1.1E-07 | 0.4371285 | 0.246 | 0.196 | 0.003668 | DF 6 | map1ab |
| lpl3 | 1.5E-07 | 0.4674863 | 0.158 | 0.112 | 0.004898 | DF 6 | lpl |
| trim712 | 1.5E-07 | 0.8774297 | 0.141 | 0.102 | 0.004946 | DF 6 | trim71 |
| rbfox23 | 1.7E-07 | 0.600856 | 0.314 | 0.271 | 0.005641 | DF 6 | rbfox2 |
| lamc32 | 1.8E-07 | 0.676517 | 0.163 | 0.12 | 0.005954 | DF 6 | lamc3 |
| cry3a2 | 2.2E-07 | 0.8373549 | 0.132 | 0.094 | 0.007177 | DF 6 | cry3a |
| FAT4 | 2.2E-07 | 0.6207536 | 0.264 | 0.222 | 0.007276 | DF 6 | FAT4 |
| ryr33 | 2.3E-07 | 0.6062195 | 0.128 | 0.089 | 0.007337 | DF 6 | ryr3 |
| si:dkey-23a13.221 | 2.7E-07 | 0.9721534 | 0.103 | 0.07 | 0.008686 | DF 6 | si:dkey-23a13.22 |
| metrn3 | 2.9E-07 | 0.6131665 | 0.158 | 0.116 | 0.009363 | DF 6 | metrn |
| cnbpb1 | 3.3E-07 | 0.3274779 | 0.591 | 0.577 | 0.010848 | DF 6 | cnbpb |
| add12 | 4.1E-07 | 0.6833655 | 0.179 | 0.137 | 0.013379 | DF 6 | add1 |
| irs2b3 | 4.2E-07 | 0.5791104 | 0.274 | 0.23 | 0.013806 | DF 6 | irs2b |
| BX005309.13 | 4.7E-07 | 0.7365647 | 0.118 | 0.082 | 0.015342 | DF 6 | BX005309.1 |

| | | | | | | | |
|---|---|---|---|---|---|---|---|
| sod11 | 5.6E-07 | 0.345071 | 0.49 | 0.466 | 0.018326 | DF 6 | sod1 |
| marcksl1b4 | 5.7E-07 | 0.2240641 | 0.304 | 0.254 | 0.018471 | DF 6 | marcksl1b |
| sltm1 | 5.8E-07 | 0.5305418 | 0.321 | 0.281 | 0.01875 | DF 6 | sltm |
| nkd12 | 7.4E-07 | 0.7806217 | 0.118 | 0.083 | 0.023949 | DF 6 | nkd1 |
| cdr2l1 | 7.4E-07 | 0.7221038 | 0.107 | 0.073 | 0.023957 | DF 6 | cdr2l |
| sqstm13 | 8.3E-07 | 0.4684876 | 0.346 | 0.305 | 0.026904 | DF 6 | sqstm1 |
| notch23 | 8.7E-07 | 0.412329 | 0.412 | 0.379 | 0.028169 | DF 6 | notch2 |
| ehd1b1 | 9.2E-07 | 0.601364 | 0.217 | 0.174 | 0.029788 | DF 6 | ehd1b |
| top13 | 9.5E-07 | 0.6343726 | 0.117 | 0.081 | 0.030739 | DF 6 | top1 |
| id32 | 9.8E-07 | 0.3319532 | 0.54 | 0.512 | 0.031879 | DF 6 | id3 |
| dnajc5aa2 | 9.9E-07 | 0.8698214 | 0.118 | 0.084 | 0.032287 | DF 6 | dnajc5aa |
| MDFI2 | 1.1E-06 | 0.5598005 | 0.279 | 0.238 | 0.036591 | DF 6 | MDFI |
| tmc6b2 | 1.5E-06 | 0.7440876 | 0.105 | 0.072 | 0.048207 | DF 6 | tmc6b |
| ldb1a2 | 1.7E-06 | 0.7479432 | 0.141 | 0.105 | 0.05398 | DF 6 | ldb1a |
| atf32 | 1.8E-06 | 0.5665019 | 0.383 | 0.346 | 0.059368 | DF 6 | atf3 |
| mhc1zaa3 | 1.9E-06 | 0.738766 | 0.128 | 0.093 | 0.061613 | DF 6 | mhc1zaa |
| ppp1r9ba1 | 2E-06 | 0.7086324 | 0.107 | 0.074 | 0.064006 | DF 6 | ppp1r9ba |
| si:dkey-16p21.85 | 2.1E-06 | 0.3066186 | 0.26 | 0.209 | 0.067098 | DF 6 | si:dkey-16p21.8 |
| srsf5b3 | 2.2E-06 | 0.7331366 | 0.152 | 0.115 | 0.072098 | DF 6 | srsf5b |
| midn3 | 2.3E-06 | 0.4403813 | 0.459 | 0.434 | 0.074461 | DF 6 | midn |
| sh3gl1b1 | 2.4E-06 | 0.4875517 | 0.284 | 0.244 | 0.078513 | DF 6 | sh3gl1b |
| zgc:1103393 | 2.6E-06 | 0.485795 | 0.258 | 0.215 | 0.084906 | DF 6 | zgc:110339 |
| mafba4 | 2.7E-06 | 0.2652436 | 0.427 | 0.39 | 0.088598 | DF 6 | mafba |
| sema5a2 | 3.1E-06 | 0.7607472 | 0.114 | 0.081 | 0.101825 | DF 6 | sema5a |
| tbl1xr1a2 | 3.8E-06 | 0.6286284 | 0.19 | 0.151 | 0.123862 | DF 6 | tbl1xr1a |
| rbm25a1 | 4E-06 | 0.7039288 | 0.113 | 0.08 | 0.12929 | DF 6 | rbm25a |
| smarca11 | 4E-06 | 0.7260896 | 0.114 | 0.081 | 0.129342 | DF 6 | smarca1 |
| arglu1a3 | 4.3E-06 | 0.584643 | 0.212 | 0.172 | 0.138732 | DF 6 | arglu1a |
| zgc:921403 | 4.8E-06 | 0.6324128 | 0.15 | 0.113 | 0.156395 | DF 6 | zgc:92140 |
| spry43 | 5E-06 | 0.7088664 | 0.216 | 0.178 | 0.163011 | DF 6 | spry4 |
| fthl273 | 5.1E-06 | 0.2069977 | 0.774 | 0.842 | 0.167221 | DF 6 | fthl27 |
| twsg1a3 | 5.3E-06 | 0.3689232 | 0.301 | 0.257 | 0.173652 | DF 6 | twsg1a |
| hlfa2 | 5.6E-06 | 0.5470329 | 0.17 | 0.131 | 0.182941 | DF 6 | hlfa |
| irf1b4 | 5.7E-06 | 0.9539544 | 0.151 | 0.117 | 0.18472 | DF 6 | irf1b |
| ywhag15 | 5.9E-06 | 0.5477619 | 0.147 | 0.11 | 0.192959 | DF 6 | ywhag1 |
| akt1s11 | 6.3E-06 | 0.7220955 | 0.192 | 0.156 | 0.20649 | DF 6 | akt1s1 |
| si:ch211-166a6.52 | 6.7E-06 | 0.7009524 | 0.194 | 0.157 | 0.218314 | DF 6 | si:ch211-166a6.5 |
| zgc:1735873 | 7E-06 | 0.5330136 | 0.17 | 0.131 | 0.228643 | DF 6 | zgc:173587 |
| ubtf | 7.2E-06 | 0.4129394 | 0.378 | 0.348 | 0.234697 | DF 6 | ubtf |
| aldh2.12 | 9.6E-06 | 0.734424 | 0.144 | 0.111 | 0.31103 | DF 6 | aldh2.1 |
| si:ch1073-185p12.22 | 1.1E-05 | 0.7254123 | 0.114 | 0.083 | 0.348836 | DF 6 | si:ch1073-185p12.2 |
| stag2b2 | 1.1E-05 | 0.5952089 | 0.204 | 0.167 | 0.362378 | DF 6 | stag2b |
| si:ch211-140m22.72 | 1.3E-05 | 0.5171984 | 0.177 | 0.14 | 0.429402 | DF 6 | si:ch211-140m22.7 |
| oga2 | 1.5E-05 | 0.7284287 | 0.129 | 0.097 | 0.480902 | DF 6 | oga |
| tet32 | 1.6E-05 | 0.6162722 | 0.131 | 0.098 | 0.519415 | DF 6 | tet3 |
| gstk11 | 1.6E-05 | 0.5568871 | 0.197 | 0.161 | 0.524155 | DF 6 | gstk1 |
| ecm24 | 1.7E-05 | 0.413699 | 0.336 | 0.298 | 0.544402 | DF 6 | ecm2 |
| tuba8l5 | 1.7E-05 | 0.2996246 | 0.44 | 0.41 | 0.566438 | DF 6 | tuba8l |

| | | | | | | | |
|---|---|---|---|---|---|---|---|
| sgk13 | 1.8E-05 | 0.3990809 | 0.398 | 0.367 | 0.578723 | DF 6 | sgk1 |
| aldh6a13 | 2E-05 | 0.4391012 | 0.306 | 0.274 | 0.651542 | DF 6 | aldh6a1 |
| rsf1b.1.1 | 2.3E-05 | 0.6661058 | 0.186 | 0.151 | 0.738939 | DF 6 | rsf1b.1.1 |
| pcdh18a3 | 2.3E-05 | 0.4533887 | 0.114 | 0.082 | 0.745538 | DF 6 | pcdh18a |
| rac3b2 | 2.5E-05 | 0.6374436 | 0.113 | 0.082 | 0.813622 | DF 6 | rac3b |
| si:ch73-366l1.54 | 2.5E-05 | 0.6252376 | 0.112 | 0.082 | 0.828906 | DF 6 | si:ch73-366l1.5 |
| ENSDARG00000075893 | 2.9E-05 | 0.7498358 | 0.113 | 0.084 | 0.92738 | DF 6 | ENSDARG00000075 |
| pptc7a2 | 3.4E-05 | 0.3396093 | 0.264 | 0.224 | 1 | DF 6 | pptc7a |
| slc1a54 | 3.5E-05 | 0.4783836 | 0.292 | 0.257 | 1 | DF 6 | slc1a5 |
| nid1b4 | 3.6E-05 | 0.6449081 | 0.108 | 0.079 | 1 | DF 6 | nid1b |
| sgtb3 | 3.8E-05 | 0.5473387 | 0.158 | 0.124 | 1 | DF 6 | sgtb |
| srrm11 | 4.3E-05 | 0.3856867 | 0.427 | 0.408 | 1 | DF 6 | srrm1 |
| adh52 | 4.3E-05 | 0.240025 | 0.535 | 0.518 | 1 | DF 6 | adh5 |
| pik3r3a2 | 4.3E-05 | 0.5836842 | 0.167 | 0.134 | 1 | DF 6 | pik3r3a |
| mepce2 | 4.4E-05 | 0.6875487 | 0.133 | 0.103 | 1 | DF 6 | mepce |
| col27a1b4 | 4.5E-05 | 0.2947675 | 0.376 | 0.337 | 1 | DF 6 | col27a1b |
| ddit42 | 4.7E-05 | 0.6238113 | 0.11 | 0.081 | 1 | DF 6 | ddit4 |
| niban1b2 | 4.8E-05 | 0.5421864 | 0.125 | 0.095 | 1 | DF 6 | niban1b |
| bco2l4 | 5.2E-05 | 0.6042425 | 0.157 | 0.124 | 1 | DF 6 | bco2l |
| fubp11 | 5.3E-05 | 0.5552017 | 0.225 | 0.193 | 1 | DF 6 | fubp1 |
| tlcd4b3 | 5.3E-05 | 0.6605638 | 0.111 | 0.083 | 1 | DF 6 | tlcd4b |
| si:ch73-281n10.22 | 5.6E-05 | 0.2515887 | 0.471 | 0.441 | 1 | DF 6 | si:ch73-281n10.2 |
| creg23 | 5.7E-05 | 0.3730613 | 0.354 | 0.325 | 1 | DF 6 | creg2 |
| srgap21 | 6.3E-05 | 0.7306996 | 0.112 | 0.084 | 1 | DF 6 | srgap2 |
| znf7031 | 6.4E-05 | 0.7094433 | 0.125 | 0.096 | 1 | DF 6 | znf703 |
| zgc:562351 | 6.6E-05 | 0.6720694 | 0.148 | 0.117 | 1 | DF 6 | zgc:56235 |
| dhrs11 | 6.8E-05 | 0.5343362 | 0.224 | 0.192 | 1 | DF 6 | dhrs1 |
| etv5a2 | 6.8E-05 | 0.6998478 | 0.133 | 0.104 | 1 | DF 6 | etv5a |
| ehbp1l1b2 | 6.9E-05 | 0.6219226 | 0.158 | 0.127 | 1 | DF 6 | ehbp1l1b |
| fmr11 | 7.3E-05 | 0.5398759 | 0.186 | 0.153 | 1 | DF 6 | fmr1 |
| CU914776.13 | 7.8E-05 | 0.3119467 | 0.113 | 0.084 | 1 | DF 6 | CU914776.1 |
| atf4b3 | 8.8E-05 | 0.2056178 | 0.632 | 0.635 | 1 | DF 6 | atf4b |
| zgc:1949893 | 9.3E-05 | 0.6435943 | 0.119 | 0.09 | 1 | DF 6 | zgc:194989 |
| si:cabz01007802.14 | 9.3E-05 | 0.7158328 | 0.118 | 0.09 | 1 | DF 6 | si:cabz01007802.1 |
| il6st4 | 9.4E-05 | 0.4784989 | 0.435 | 0.424 | 1 | DF 6 | il6st |
| arg23 | 9.6E-05 | 0.7227578 | 0.195 | 0.164 | 1 | DF 6 | arg2 |
| CT956002.21 | 1E-04 | 0.7700626 | 0.102 | 0.075 | 1 | DF 6 | CT956002.2 |
| CABZ01074397.13 | 0.0001 | 0.6173421 | 0.181 | 0.149 | 1 | DF 6 | CABZ01074397.1 |
| UBB2 | 0.0001 | 0.3461724 | 0.57 | 0.563 | 1 | DF 6 | UBB |
| loxl14 | 0.00012 | 0.3833415 | 0.237 | 0.201 | 1 | DF 6 | loxl1 |
| myofl2 | 0.00012 | 0.4904111 | 0.192 | 0.159 | 1 | DF 6 | myofl |
| luc7l32 | 0.00013 | 0.5812994 | 0.147 | 0.117 | 1 | DF 6 | luc7l3 |
| hexim13 | 0.00014 | 0.543106 | 0.264 | 0.234 | 1 | DF 6 | hexim1 |
| si:dkey-276j7.13 | 0.00014 | 0.5817721 | 0.196 | 0.167 | 1 | DF 6 | si:dkey-276j7.1 |
| ankrd12 | 0.00015 | 0.4445039 | 0.282 | 0.25 | 1 | DF 6 | ankrd12 |
| eif4a23 | 0.00018 | 0.4031095 | 0.225 | 0.193 | 1 | DF 6 | eif4a2 |
| ptbp32 | 0.00019 | 0.6396273 | 0.163 | 0.134 | 1 | DF 6 | ptbp3 |
| FP236812.12 | 0.0002 | 0.4812005 | 0.128 | 0.099 | 1 | DF 6 | FP236812.1 |

| | | | | | | | |
|---|---|---|---|---|---|---|---|
| u2surp3 | 0.00022 | 0.3624613 | 0.337 | 0.314 | 1 | DF 6 | u2surp |
| fbxl3a1 | 0.00024 | 0.5915896 | 0.122 | 0.095 | 1 | DF 6 | fbxl3a |
| snai23 | 0.00026 | 0.4689471 | 0.133 | 0.105 | 1 | DF 6 | snai2 |
| her63 | 0.00029 | 0.3824185 | 0.453 | 0.433 | 1 | DF 6 | her6 |
| sf3b61 | 0.00031 | 0.3970084 | 0.308 | 0.282 | 1 | DF 6 | sf3b6 |
| bcat21 | 0.00031 | 0.7449235 | 0.113 | 0.088 | 1 | DF 6 | bcat2 |
| tpm23 | 0.00032 | 0.4778088 | 0.145 | 0.116 | 1 | DF 6 | tpm2 |
| mex3b3 | 0.00033 | 0.5037192 | 0.202 | 0.172 | 1 | DF 6 | mex3b |
| axin12 | 0.00035 | 0.6115689 | 0.14 | 0.113 | 1 | DF 6 | axin1 |
| clstn11 | 0.00036 | 0.510537 | 0.165 | 0.136 | 1 | DF 6 | clstn1 |
| nrn1a3 | 0.00043 | 0.4701913 | 0.115 | 0.089 | 1 | DF 6 | nrn1a |
| rpia2 | 0.00045 | 0.657194 | 0.122 | 0.097 | 1 | DF 6 | rpia |
| brd2a3 | 0.00045 | 0.277617 | 0.464 | 0.448 | 1 | DF 6 | brd2a |
| pabpn11 | 0.00046 | 0.3309222 | 0.421 | 0.411 | 1 | DF 6 | pabpn1 |
| zmynd111 | 0.00047 | 0.5049592 | 0.167 | 0.139 | 1 | DF 6 | zmynd11 |
| scube32 | 0.00049 | 0.4248311 | 0.123 | 0.096 | 1 | DF 6 | scube3 |
| srsf1b | 0.0005 | 0.6677189 | 0.144 | 0.118 | 1 | DF 6 | srsf1b |
| irf2bpl2 | 0.00051 | 0.4535957 | 0.228 | 0.199 | 1 | DF 6 | irf2bpl |
| ubtfl | 0.00051 | 0.6017592 | 0.187 | 0.16 | 1 | DF 6 | ubtfl |
| id2b4 | 0.00051 | 0.4705479 | 0.22 | 0.19 | 1 | DF 6 | id2b |
| pnisr2 | 0.00051 | 0.4939509 | 0.215 | 0.186 | 1 | DF 6 | pnisr |
| tgfb1a2 | 0.00052 | 0.2874107 | 0.306 | 0.274 | 1 | DF 6 | tgfb1a |
| khdrbs1b1 | 0.00053 | 0.4732653 | 0.264 | 0.24 | 1 | DF 6 | khdrbs1b |
| cbr1l4 | 0.00055 | 0.4531298 | 0.243 | 0.214 | 1 | DF 6 | cbr1l |
| rab40b1 | 0.00057 | 0.6151271 | 0.102 | 0.078 | 1 | DF 6 | rab40b |
| ppp1r103 | 0.00059 | 0.2899761 | 0.489 | 0.485 | 1 | DF 6 | ppp1r10 |
| ctdspl2b4 | 0.0006 | 0.6737701 | 0.138 | 0.113 | 1 | DF 6 | ctdspl2b |
| myclb3 | 0.00061 | 0.4821552 | 0.169 | 0.14 | 1 | DF 6 | myclb |
| sox11a2 | 0.00073 | 0.3152996 | 0.371 | 0.343 | 1 | DF 6 | sox11a |
| sat1a.24 | 0.00076 | 0.4049021 | 0.464 | 0.458 | 1 | DF 6 | sat1a.2 |
| ubc2 | 0.00082 | 0.3544159 | 0.634 | 0.664 | 1 | DF 6 | ubc |
| dio13 | 0.00084 | 0.4486836 | 0.149 | 0.122 | 1 | DF 6 | dio1 |
| mllt102 | 0.00099 | 0.5413145 | 0.105 | 0.082 | 1 | DF 6 | mllt10 |
| zgc:1140464 | 0.00101 | 0.3955965 | 0.125 | 0.1 | 1 | DF 6 | zgc:114046 |
| jun3 | 0.00115 | 0.2131532 | 0.814 | 0.856 | 1 | DF 6 | jun |
| lmo4a2 | 0.00117 | 0.5629965 | 0.15 | 0.125 | 1 | DF 6 | lmo4a |
| mapk1 | 0.00126 | 0.6255039 | 0.106 | 0.084 | 1 | DF 6 | mapk1 |
| thbs1a2 | 0.0013 | 0.3537704 | 0.339 | 0.313 | 1 | DF 6 | thbs1a |
| rorab2 | 0.00146 | 0.4976265 | 0.151 | 0.126 | 1 | DF 6 | rorab |
| CPNE84 | 0.0015 | 0.3080131 | 0.314 | 0.287 | 1 | DF 6 | CPNE8 |
| per1b3 | 0.00163 | 0.4438526 | 0.227 | 0.203 | 1 | DF 6 | per1b |
| si:dkey-102m7.33 | 0.00164 | 0.4753117 | 0.145 | 0.12 | 1 | DF 6 | si:dkey-102m7.3 |
| arnt21 | 0.00176 | 0.5470681 | 0.112 | 0.09 | 1 | DF 6 | arnt2 |
| cdc42ep5 | 0.00187 | 0.3304067 | 0.321 | 0.3 | 1 | DF 6 | cdc42ep5 |
| tdh2 | 0.00192 | 0.4348321 | 0.262 | 0.241 | 1 | DF 6 | tdh |
| si:ch211-212k18.4 | 0.002 | 0.4500128 | 0.239 | 0.216 | 1 | DF 6 | si:ch211-212k18.4 |
| zgc:1229792 | 0.00217 | 0.5570913 | 0.101 | 0.08 | 1 | DF 6 | zgc:122979 |
| BX511129.23 | 0.00227 | 0.609043 | 0.11 | 0.089 | 1 | DF 6 | BX511129.2 |

| | | | | | | | |
|---|---|---|---|---|---|---|---|
| klf155 | 0.0023 | 0.3690136 | 0.234 | 0.208 | 1 | DF 6 | klf15 |
| brdt2 | 0.00244 | 0.3018438 | 0.333 | 0.311 | 1 | DF 6 | brdt |
| trip10a | 0.00249 | 0.6682806 | 0.101 | 0.081 | 1 | DF 6 | trip10a |
| ISCU2 | 0.0025 | 0.41619 | 0.284 | 0.266 | 1 | DF 6 | ISCU |
| si:ch211-195b15.82 | 0.00254 | 0.4115808 | 0.113 | 0.091 | 1 | DF 6 | si:ch211-195b15.8 |
| rnf442 | 0.00255 | 0.6396191 | 0.143 | 0.12 | 1 | DF 6 | rnf44 |
| irf91 | 0.00279 | 0.5631593 | 0.15 | 0.128 | 1 | DF 6 | irf9 |
| fbln53 | 0.00281 | 0.561171 | 0.105 | 0.084 | 1 | DF 6 | fbln5 |
| amd12 | 0.00302 | 0.5948533 | 0.117 | 0.096 | 1 | DF 6 | amd1 |
| sh3bgrl1 | 0.00309 | 0.3485139 | 0.249 | 0.227 | 1 | DF 6 | sh3bgrl |
| tnrc6c21 | 0.00321 | 0.6108351 | 0.108 | 0.088 | 1 | DF 6 | tnrc6c2 |
| nr3c11 | 0.00324 | 0.4737757 | 0.224 | 0.201 | 1 | DF 6 | nr3c1 |
| nucks1a1 | 0.0034 | 0.3455482 | 0.375 | 0.363 | 1 | DF 6 | nucks1a |
| asph1 | 0.00343 | 0.2057217 | 0.4 | 0.378 | 1 | DF 6 | asph |
| bcl91 | 0.00389 | 0.6349686 | 0.105 | 0.085 | 1 | DF 6 | bcl9 |
| fasn1 | 0.00393 | 0.5786624 | 0.123 | 0.103 | 1 | DF 6 | fasn |
| vapal2 | 0.00396 | 0.4737278 | 0.186 | 0.165 | 1 | DF 6 | vapal |
| setbp13 | 0.00399 | 0.3389739 | 0.272 | 0.249 | 1 | DF 6 | setbp1 |
| tra2a | 0.00402 | 0.5403845 | 0.209 | 0.188 | 1 | DF 6 | tra2a |
| crlf33 | 0.00426 | 0.6462939 | 0.108 | 0.088 | 1 | DF 6 | crlf3 |
| tnrc6a4 | 0.00431 | 0.3453499 | 0.292 | 0.272 | 1 | DF 6 | tnrc6a |
| sh3bp5la2 | 0.00462 | 0.5476499 | 0.126 | 0.106 | 1 | DF 6 | sh3bp5la |
| setb1 | 0.00482 | 0.2311454 | 0.392 | 0.376 | 1 | DF 6 | setb |
| macroh2a2 | 0.00499 | 0.3189149 | 0.4 | 0.396 | 1 | DF 6 | macroh2a2 |
| pik3r22 | 0.00505 | 0.4201454 | 0.222 | 0.201 | 1 | DF 6 | pik3r2 |
| tmem106ba1 | 0.00506 | 0.4434659 | 0.212 | 0.19 | 1 | DF 6 | tmem106ba |
| mocs24 | 0.00541 | 0.4813216 | 0.108 | 0.088 | 1 | DF 6 | mocs2 |
| nktr2 | 0.00583 | 0.5517082 | 0.18 | 0.159 | 1 | DF 6 | nktr |
| syne31 | 0.00622 | 0.5351125 | 0.131 | 0.111 | 1 | DF 6 | syne3 |
| paics1 | 0.00626 | 0.5552574 | 0.102 | 0.083 | 1 | DF 6 | paics |
| si:dkey-56m19.53 | 0.0064 | 0.5969499 | 0.104 | 0.085 | 1 | DF 6 | si:dkey-56m19.5 |
| mgaa1 | 0.00659 | 0.5125379 | 0.161 | 0.141 | 1 | DF 6 | mgaa |
| caprin1b2 | 0.00665 | 0.2864587 | 0.312 | 0.297 | 1 | DF 6 | caprin1b |
| zyx5 | 0.00732 | 0.2623748 | 0.335 | 0.318 | 1 | DF 6 | zyx |
| rab7a1 | 0.00732 | 0.4992674 | 0.161 | 0.141 | 1 | DF 6 | rab7a |
| homer3b1 | 0.00735 | 0.5710606 | 0.167 | 0.147 | 1 | DF 6 | homer3b |
| gyg1a | 0.00739 | 0.4407126 | 0.202 | 0.183 | 1 | DF 6 | gyg1a |
| prpf391 | 0.00754 | 0.4781972 | 0.15 | 0.13 | 1 | DF 6 | prpf39 |
| srrt2 | 0.00781 | 0.5676793 | 0.146 | 0.127 | 1 | DF 6 | srrt |
| pitpnc1b3 | 0.00785 | 0.5165719 | 0.1 | 0.082 | 1 | DF 6 | pitpnc1b |
| msra | 0.00795 | 0.4969274 | 0.103 | 0.084 | 1 | DF 6 | msra |
| ramp24 | 0.008 | 0.4067317 | 0.163 | 0.142 | 1 | DF 6 | ramp2 |
| CABZ01032488.11 | 0.00803 | 0.4557221 | 0.125 | 0.105 | 1 | DF 6 | CABZ01032488.1 |
| has22 | 0.00807 | 0.2250248 | 0.135 | 0.113 | 1 | DF 6 | has2 |
| ghrb4 | 0.00854 | 0.4390275 | 0.128 | 0.108 | 1 | DF 6 | ghrb |
| pank21 | 0.00909 | 0.5066343 | 0.118 | 0.1 | 1 | DF 6 | pank2 |
| sfpq1 | 0.00925 | 0.4588859 | 0.225 | 0.207 | 1 | DF 6 | sfpq |
| dusp54 | 0.00931 | 0.2259717 | 0.222 | 0.198 | 1 | DF 6 | dusp5 |

| | | | | | | | |
|---|---|---|---|---|---|---|---|
| pcf112 | 0.0094 | 0.3744409 | 0.269 | 0.253 | 1 | DF 6 | pcf11 |
| cbx1a2 | 0.00943 | 0.4461323 | 0.223 | 0.204 | 1 | DF 6 | cbx1a |
| metrnla2 | 0.00959 | 0.2620926 | 0.33 | 0.316 | 1 | DF 6 | metrnla |
| snorc | 0 | 9.4100777 | 0.952 | 0.011 | 0 | Chondrocytes | snorc |
| col9a2 | 0 | 9.039203 | 0.952 | 0.02 | 0 | Chondrocytes | col9a2 |
| cnmd | 0 | 9.1565406 | 0.941 | 0.021 | 0 | Chondrocytes | cnmd |
| col2a1a | 0 | 10.294065 | 0.945 | 0.043 | 0 | Chondrocytes | col2a1a |
| tnxba | 0 | 8.9344104 | 0.915 | 0.021 | 0 | Chondrocytes | tnxba |
| col9a31 | 0 | 7.3632613 | 0.931 | 0.042 | 0 | Chondrocytes | col9a3 |
| col11a21 | 0 | 5.2430182 | 0.969 | 0.084 | 0 | Chondrocytes | col11a2 |
| mia | 0 | 10.686801 | 0.832 | 0.005 | 0 | Chondrocytes | mia |
| epyc | 0 | 7.5521334 | 0.858 | 0.039 | 0 | Chondrocytes | epyc |
| ucmab | 0 | 8.3776732 | 0.823 | 0.015 | 0 | Chondrocytes | ucmab |
| col9a1a | 0 | 6.2729973 | 0.928 | 0.133 | 0 | Chondrocytes | col9a1a |
| acana | 0 | 6.9615693 | 0.78 | 0.012 | 0 | Chondrocytes | acana |
| fgfbp2b | 0 | 10.452401 | 0.767 | 0.009 | 0 | Chondrocytes | fgfbp2b |
| matn1 | 0 | 10.481125 | 0.75 | 0.019 | 0 | Chondrocytes | matn1 |
| col8a1b1 | 0 | 5.8111878 | 0.846 | 0.119 | 0 | Chondrocytes | col8a1b |
| meltf | 0 | 9.0122444 | 0.727 | 0.002 | 0 | Chondrocytes | meltf |
| thbs1b1 | 0 | 4.1587228 | 0.915 | 0.192 | 0 | Chondrocytes | thbs1b |
| hapln1b1 | 0 | 4.6406919 | 0.776 | 0.065 | 0 | Chondrocytes | hapln1b |
| col11a1a2 | 0 | 5.4290632 | 0.98 | 0.301 | 0 | Chondrocytes | col11a1a |
| ccn2a3 | 0 | 3.1158921 | 0.856 | 0.186 | 0 | Chondrocytes | ccn2a |
| acanb | 0 | 9.7823987 | 0.664 | 0.001 | 0 | Chondrocytes | acanb |
| col15a1b | 0 | 4.5510676 | 0.673 | 0.03 | 0 | Chondrocytes | col15a1b |
| wwp2 | 0 | 3.7311403 | 0.723 | 0.085 | 0 | Chondrocytes | wwp2 |
| itih1 | 0 | 5.7524219 | 0.648 | 0.012 | 0 | Chondrocytes | itih1 |
| arpc1b1 | 0 | 4.215977 | 0.852 | 0.229 | 0 | Chondrocytes | arpc1b |
| loxl3b | 0 | 3.1387438 | 0.755 | 0.135 | 0 | Chondrocytes | loxl3b |
| cdc42ep31 | 0 | 4.5698296 | 0.648 | 0.043 | 0 | Chondrocytes | cdc42ep3 |
| sox62 | 0 | 3.4547239 | 0.681 | 0.085 | 0 | Chondrocytes | sox6 |
| si:ch1073-204b8.1 | 0 | 5.3628506 | 0.602 | 0.023 | 0 | Chondrocytes | si:ch1073-204b8.1 |
| crip21 | 0 | 3.5255867 | 0.633 | 0.06 | 0 | Chondrocytes | crip2 |
| tent5ba1 | 0 | 3.1779062 | 0.685 | 0.126 | 0 | Chondrocytes | tent5ba |
| si:dkey-6n6.1 | 0 | 9.4384533 | 0.549 | 0.001 | 0 | Chondrocytes | si:dkey-6n6.1 |
| scin | 0 | 8.9002149 | 0.54 | 0.002 | 0 | Chondrocytes | scin |
| fn1b2 | 0 | 2.9485998 | 0.919 | 0.385 | 0 | Chondrocytes | fn1b |
| fgfr32 | 0 | 1.8252535 | 0.787 | 0.254 | 0 | Chondrocytes | fgfr3 |
| bgnb1 | 0 | 3.2074726 | 0.653 | 0.122 | 0 | Chondrocytes | bgnb |
| eps8l2 | 0 | 7.1271891 | 0.533 | 0.004 | 0 | Chondrocytes | eps8l2 |
| ccdc802 | 0 | 2.1841095 | 0.893 | 0.371 | 0 | Chondrocytes | ccdc80 |
| ccn1l2 | 0 | 7.8365655 | 0.522 | 0.003 | 0 | Chondrocytes | ccn1l2 |
| pcolce2b3 | 0 | 2.3994651 | 0.799 | 0.282 | 0 | Chondrocytes | pcolce2b |
| itga102 | 0 | 1.7319412 | 0.753 | 0.239 | 0 | Chondrocytes | itga10 |
| CLEC3A | 0 | 9.9472003 | 0.508 | 0.001 | 0 | Chondrocytes | CLEC3A |
| dhrs3b2 | 0 | 2.7551755 | 0.607 | 0.101 | 0 | Chondrocytes | dhrs3b |
| pgam1a1 | 0 | 2.6085002 | 0.677 | 0.172 | 0 | Chondrocytes | pgam1a |
| p3h21 | 0 | 3.774066 | 0.547 | 0.047 | 0 | Chondrocytes | p3h2 |

| | | | | | | | |
|---|---|---|---|---|---|---|---|
| alcama2 | 0 | 3.1302523 | 0.544 | 0.061 | 0 | Chondrocytes | alcama |
| si:dkey-262k9.21 | 0 | 3.270208 | 0.571 | 0.088 | 0 | Chondrocytes | si:dkey-262k9.2 |
| slc38a5a2 | 0 | 3.8981691 | 0.537 | 0.056 | 0 | Chondrocytes | slc38a5a |
| foxc1b1 | 0 | 3.4177909 | 0.522 | 0.049 | 0 | Chondrocytes | foxc1b |
| pcolceb2 | 0 | 2.6665899 | 0.632 | 0.159 | 0 | Chondrocytes | pcolceb |
| rock2b.12 | 0 | 2.1308343 | 0.631 | 0.158 | 0 | Chondrocytes | rock2b.1 |
| papss2b1 | 0 | 3.0175939 | 0.555 | 0.084 | 0 | Chondrocytes | papss2b |
| foxa3 | 0 | 8.1587754 | 0.468 | 0.002 | 0 | Chondrocytes | foxa3 |
| elovl1b1 | 0 | 1.9230283 | 0.802 | 0.338 | 0 | Chondrocytes | elovl1b |
| pdgfab | 0 | 4.5811908 | 0.486 | 0.024 | 0 | Chondrocytes | pdgfab |
| ubl3a2 | 0 | 1.8807702 | 0.782 | 0.323 | 0 | Chondrocytes | ubl3a |
| f13a1b1 | 0 | 3.3324023 | 0.74 | 0.281 | 0 | Chondrocytes | f13a1b |
| arhgef12a | 0 | 3.0228274 | 0.53 | 0.074 | 0 | Chondrocytes | arhgef12a |
| BX511021.2 | 0 | 4.1886458 | 0.483 | 0.033 | 0 | Chondrocytes | BX511021.2 |
| tgfbi4 | 0 | 2.7774118 | 0.635 | 0.186 | 0 | Chondrocytes | tgfbi |
| si:dkey-19b23.8 | 0 | 3.7686862 | 0.488 | 0.041 | 0 | Chondrocytes | si:dkey-19b23.8 |
| traf4a1 | 0 | 3.2792189 | 0.506 | 0.062 | 0 | Chondrocytes | traf4a |
| atp1b1a | 0 | 2.2573883 | 0.912 | 0.472 | 0 | Chondrocytes | atp1b1a |
| frzb5 | 0 | 2.6745103 | 0.555 | 0.116 | 0 | Chondrocytes | frzb |
| col2a1b3 | 0 | 3.3653276 | 0.926 | 0.488 | 0 | Chondrocytes | col2a1b |
| six2a2 | 0 | 2.8642191 | 0.525 | 0.089 | 0 | Chondrocytes | six2a |
| loxl2a2 | 0 | 2.3602705 | 0.642 | 0.211 | 0 | Chondrocytes | loxl2a |
| tent5bb3 | 0 | 2.3992464 | 0.558 | 0.13 | 0 | Chondrocytes | tent5bb |
| angptl1b1 | 0 | 3.2338073 | 0.478 | 0.052 | 0 | Chondrocytes | angptl1b |
| tgm2l | 0 | 6.6427238 | 0.432 | 0.006 | 0 | Chondrocytes | tgm2l |
| EMP3 | 0 | 3.2233952 | 0.477 | 0.052 | 0 | Chondrocytes | EMP3 |
| plod1a1 | 0 | 2.0717409 | 0.65 | 0.231 | 0 | Chondrocytes | plod1a |
| rhbdl2 | 0 | 4.4054777 | 0.438 | 0.021 | 0 | Chondrocytes | rhbdl2 |
| ecrg4a3 | 0 | 2.8953604 | 0.967 | 0.551 | 0 | Chondrocytes | ecrg4a |
| otos | 0 | 9.9718889 | 0.418 | 0.004 | 0 | Chondrocytes | otos |
| si:busm1-57f23.12 | 0 | 2.3410234 | 0.971 | 0.562 | 0 | Chondrocytes | si:busm1-57f23.1 |
| si:ch211-156j16.13 | 0 | 1.5277556 | 0.546 | 0.14 | 0 | Chondrocytes | si:ch211-156j16.1 |
| p4ha1b4 | 0 | 2.0132128 | 0.532 | 0.128 | 0 | Chondrocytes | p4ha1b |
| camsap2b1 | 0 | 2.5211936 | 0.494 | 0.096 | 0 | Chondrocytes | camsap2b |
| ccn6 | 0 | 10.794773 | 0.398 | 0 | 0 | Chondrocytes | ccn6 |
| si:ch211-1a19.31 | 0 | 1.7707302 | 0.9 | 0.504 | 0 | Chondrocytes | si:ch211-1a19.3 |
| cthrc1b | 0 | 5.7642323 | 0.401 | 0.008 | 0 | Chondrocytes | cthrc1b |
| fbln71 | 0 | 2.9912491 | 0.482 | 0.089 | 0 | Chondrocytes | fbln7 |
| srpx3 | 0 | 2.0826152 | 0.545 | 0.155 | 0 | Chondrocytes | srpx |
| nid2a1 | 0 | 2.9617618 | 0.451 | 0.062 | 0 | Chondrocytes | nid2a |
| sept8b | 0 | 6.7452022 | 0.39 | 0.004 | 0 | Chondrocytes | sept8b |
| CR855337.1 | 0 | 2.8536996 | 0.465 | 0.083 | 0 | Chondrocytes | CR855337.1 |
| fgfrl1b | 0 | 4.1558173 | 0.402 | 0.021 | 0 | Chondrocytes | fgfrl1b |
| ehd4 | 0 | 2.9737295 | 0.419 | 0.044 | 0 | Chondrocytes | ehd4 |
| etv5b3 | 0 | 2.4870729 | 0.46 | 0.089 | 0 | Chondrocytes | etv5b |
| sox9a | 0 | 2.5355154 | 0.451 | 0.08 | 0 | Chondrocytes | sox9a |
| si:ch211-222l21.12 | 0 | 2.1335876 | 0.915 | 0.544 | 0 | Chondrocytes | si:ch211-222l21.1 |
| tgm2b1 | 0 | 3.9092656 | 0.398 | 0.035 | 0 | Chondrocytes | tgm2b |

| | | | | | | | |
|---|---|---|---|---|---|---|---|
| ninj21 | 0 | 5.9635953 | 0.374 | 0.012 | 0 | Chondrocytes | ninj2 |
| rapgef21 | 0 | 2.3637589 | 0.456 | 0.095 | 0 | Chondrocytes | rapgef2 |
| slc35d1a | 0 | 3.6031828 | 0.385 | 0.03 | 0 | Chondrocytes | slc35d1a |
| mob2b1 | 0 | 2.8548851 | 0.423 | 0.07 | 0 | Chondrocytes | mob2b |
| plod23 | 0 | 2.534519 | 0.414 | 0.061 | 0 | Chondrocytes | plod2 |
| mibp21 | 0 | 2.4277462 | 0.953 | 0.602 | 0 | Chondrocytes | mibp2 |
| ptn1 | 0 | 3.6901752 | 0.384 | 0.038 | 0 | Chondrocytes | ptn |
| sox10 | 0 | 7.2267632 | 0.347 | 0.002 | 0 | Chondrocytes | sox10 |
| fxyd12 | 0 | 1.6514217 | 0.907 | 0.562 | 0 | Chondrocytes | fxyd1 |
| wscd2 | 0 | 4.5508774 | 0.355 | 0.016 | 0 | Chondrocytes | wscd2 |
| cux1a3 | 0 | 2.7855183 | 0.396 | 0.058 | 0 | Chondrocytes | cux1a |
| pim32 | 0 | 2.2178943 | 0.435 | 0.097 | 0 | Chondrocytes | pim3 |
| chad1 | 0 | 3.4442604 | 0.375 | 0.042 | 0 | Chondrocytes | chad |
| sox5 | 0 | 4.4756494 | 0.344 | 0.012 | 0 | Chondrocytes | sox5 |
| rasal2 | 0 | 2.7532208 | 0.393 | 0.062 | 0 | Chondrocytes | rasal2 |
| msi2b | 0 | 2.1256137 | 0.416 | 0.089 | 0 | Chondrocytes | msi2b |
| aldocb | 0 | 2.6960197 | 0.392 | 0.066 | 0 | Chondrocytes | aldocb |
| insig1 | 0 | 3.4983729 | 0.386 | 0.064 | 0 | Chondrocytes | insig1 |
| si:ch211-120e1.7 | 0 | 5.1295042 | 0.333 | 0.011 | 0 | Chondrocytes | si:ch211-120e1.7 |
| bmp2b3 | 0 | 2.6552868 | 0.414 | 0.092 | 0 | Chondrocytes | bmp2b |
| plxnb1a | 0 | 2.8250155 | 0.365 | 0.044 | 0 | Chondrocytes | plxnb1a |
| tspan5a | 0 | 3.852181 | 0.344 | 0.023 | 0 | Chondrocytes | tspan5a |
| aff3 | 0 | 7.6498286 | 0.322 | 0.002 | 0 | Chondrocytes | aff3 |
| mtss1 | 0 | 2.9053461 | 0.363 | 0.046 | 0 | Chondrocytes | mtss1 |
| ccn1l11 | 0 | 3.5116439 | 0.342 | 0.028 | 0 | Chondrocytes | ccn1l1 |
| thbs4a2 | 0 | 1.4399357 | 0.382 | 0.071 | 0 | Chondrocytes | thbs4a |
| slc38a22 | 0 | 2.7187471 | 0.379 | 0.07 | 0 | Chondrocytes | slc38a2 |
| si:ch211-196g2.4 | 0 | 2.1290314 | 0.39 | 0.082 | 0 | Chondrocytes | si:ch211-196g2.4 |
| ptgdsa | 0 | 8.2487311 | 0.309 | 0.001 | 0 | Chondrocytes | ptgdsa |
| matn4 | 0 | 3.3310991 | 0.343 | 0.036 | 0 | Chondrocytes | matn4 |
| pxylp1 | 0 | 3.7868167 | 0.327 | 0.021 | 0 | Chondrocytes | pxylp1 |
| il16 | 0 | 2.9965301 | 0.328 | 0.036 | 0 | Chondrocytes | il16 |
| pnp5a | 0 | 2.770346 | 0.334 | 0.05 | 0 | Chondrocytes | pnp5a |
| itga1 | 0 | 3.3426673 | 0.306 | 0.023 | 0 | Chondrocytes | itga1 |
| sema3d | 0 | 2.6531389 | 0.318 | 0.036 | 0 | Chondrocytes | sema3d |
| tiam2a1 | 0 | 2.7351669 | 0.333 | 0.053 | 0 | Chondrocytes | tiam2a |
| gas6 | 0 | 2.3344408 | 0.336 | 0.058 | 0 | Chondrocytes | gas6 |
| alpi.1 | 0 | 3.9993571 | 0.293 | 0.017 | 0 | Chondrocytes | alpi.1 |
| tmem26b | 0 | 7.7884667 | 0.275 | 0.002 | 0 | Chondrocytes | tmem26b |
| gask1b | 0 | 3.0841131 | 0.299 | 0.031 | 0 | Chondrocytes | gask1b |
| klf2b1 | 0 | 2.4308884 | 0.319 | 0.053 | 0 | Chondrocytes | klf2b |
| stxbp6 | 0 | 2.8568912 | 0.307 | 0.041 | 0 | Chondrocytes | stxbp6 |
| frya | 0 | 3.9267837 | 0.262 | 0.014 | 0 | Chondrocytes | frya |
| mgat5 | 0 | 3.3929576 | 0.269 | 0.022 | 0 | Chondrocytes | mgat5 |
| fthl28 | 0 | 6.0481618 | 0.26 | 0.016 | 0 | Chondrocytes | fthl28 |
| lgr4 | 0 | 3.3251631 | 0.254 | 0.017 | 0 | Chondrocytes | lgr4 |
| chst3a | 0 | 7.7213753 | 0.238 | 0.001 | 0 | Chondrocytes | chst3a |
| klhl21 | 0 | 3.0216318 | 0.266 | 0.033 | 0 | Chondrocytes | klhl21 |

| | | | | | | | |
|---|---|---|---|---|---|---|---|
| si:ch211-247j9.1 | 0 | 3.5280213 | 0.26 | 0.027 | 0 | Chondrocytes | si:ch211-247j9.1 |
| mibp | 0 | 4.4869124 | 0.241 | 0.011 | 0 | Chondrocytes | mibp |
| zgc:165573 | 0 | 3.170131 | 0.255 | 0.03 | 0 | Chondrocytes | zgc:165573 |
| pcolce2a | 0 | 4.9581443 | 0.223 | 0.006 | 0 | Chondrocytes | pcolce2a |
| rasd1 | 0 | 3.6859816 | 0.23 | 0.016 | 0 | Chondrocytes | rasd1 |
| znf385c | 0 | 6.9309474 | 0.215 | 0.001 | 0 | Chondrocytes | znf385c |
| loxl5b | 0 | 6.4649945 | 0.216 | 0.005 | 0 | Chondrocytes | loxl5b |
| dcun1d2b | 0 | 2.6995583 | 0.239 | 0.028 | 0 | Chondrocytes | dcun1d2b |
| CR936300.1 | 0 | 4.7778445 | 0.216 | 0.006 | 0 | Chondrocytes | CR936300.1 |
| slc6a8 | 0 | 3.9308357 | 0.224 | 0.015 | 0 | Chondrocytes | slc6a8 |
| itih6 | 0 | 4.6980444 | 0.214 | 0.007 | 0 | Chondrocytes | itih6 |
| mgat4a | 0 | 3.1478554 | 0.228 | 0.022 | 0 | Chondrocytes | mgat4a |
| VIT1 | 0 | 3.3178503 | 0.226 | 0.021 | 0 | Chondrocytes | VIT |
| stard8 | 0 | 2.7230833 | 0.221 | 0.023 | 0 | Chondrocytes | stard8 |
| hif1aa | 0 | 4.5005749 | 0.205 | 0.008 | 0 | Chondrocytes | hif1aa |
| pip5k1ca | 0 | 5.1909903 | 0.2 | 0.005 | 0 | Chondrocytes | pip5k1ca |
| fryb | 0 | 4.4391535 | 0.199 | 0.007 | 0 | Chondrocytes | fryb |
| loxl5a | 0 | 9.6948119 | 0.183 | 0 | 0 | Chondrocytes | loxl5a |
| kcnq5b | 0 | 5.0446216 | 0.181 | 0.004 | 0 | Chondrocytes | kcnq5b |
| slc44a5b | 0 | 3.1148427 | 0.189 | 0.017 | 0 | Chondrocytes | slc44a5b |
| sept5a | 0 | 3.1644052 | 0.187 | 0.017 | 0 | Chondrocytes | sept5a |
| scube1 | 0 | 4.0181686 | 0.17 | 0.01 | 0 | Chondrocytes | scube1 |
| pde5ab | 0 | 4.7038056 | 0.163 | 0.003 | 0 | Chondrocytes | pde5ab |
| rac2 | 0 | 3.1712753 | 0.168 | 0.015 | 0 | Chondrocytes | rac2 |
| si:dkey-194e6.1 | 0 | 10.866565 | 0.153 | 0 | 0 | Chondrocytes | si:dkey-194e6.1 |
| chadlb | 0 | 7.2395726 | 0.153 | 0.001 | 0 | Chondrocytes | chadlb |
| CABZ01076996.1 | 0 | 5.9181404 | 0.155 | 0.003 | 0 | Chondrocytes | CABZ01076996.1 |
| tfeb | 0 | 4.4779481 | 0.154 | 0.009 | 0 | Chondrocytes | tfeb |
| emp3b | 0 | 4.8698354 | 0.143 | 0.003 | 0 | Chondrocytes | emp3b |
| adamts3 | 0 | 3.3467614 | 0.149 | 0.011 | 0 | Chondrocytes | adamts3 |
| si:dkey-264d12.1 | 0 | 3.725899 | 0.145 | 0.01 | 0 | Chondrocytes | si:dkey-264d12.1 |
| si:ch211-106h11.3 | 0 | 6.4577221 | 0.132 | 0.002 | 0 | Chondrocytes | si:ch211-106h11.3 |
| fads2 | 0 | 4.6474448 | 0.134 | 0.005 | 0 | Chondrocytes | fads2 |
| tspan2b | 0 | 4.419189 | 0.131 | 0.005 | 0 | Chondrocytes | tspan2b |
| FCHSD2 | 0 | 8.6419638 | 0.125 | 0 | 0 | Chondrocytes | FCHSD2 |
| b4galnt4a | 0 | 3.936319 | 0.131 | 0.007 | 0 | Chondrocytes | b4galnt4a |
| sb:cb1058 | 0 | 5.4279643 | 0.126 | 0.004 | 0 | Chondrocytes | sb:cb1058 |
| atcaya | 0 | 7.0952574 | 0.119 | 0.001 | 0 | Chondrocytes | atcaya |
| si:dkeyp-71f10.5 | 0 | 7.2675094 | 0.118 | 0.001 | 0 | Chondrocytes | si:dkeyp-71f10.5 |
| stk38a | 0 | 3.6516249 | 0.123 | 0.007 | 0 | Chondrocytes | stk38a |
| ankrd33ba | 0 | 3.5319603 | 0.124 | 0.008 | 0 | Chondrocytes | ankrd33ba |
| bicdl2 | 0 | 5.9680824 | 0.115 | 0.002 | 0 | Chondrocytes | bicdl2 |
| FP016234.1 | 0 | 6.2769428 | 0.114 | 0.001 | 0 | Chondrocytes | FP016234.1 |
| ddr2b | 0 | 7.0951893 | 0.114 | 0.001 | 0 | Chondrocytes | ddr2b |
| adamtsl2 | 0 | 5.542223 | 0.114 | 0.002 | 0 | Chondrocytes | adamtsl2 |
| csgalnact1a | 0 | 4.1064544 | 0.115 | 0.005 | 0 | Chondrocytes | csgalnact1a |
| sesn2 | 0 | 4.4262482 | 0.113 | 0.004 | 0 | Chondrocytes | sesn2 |
| syt5a | 0 | 4.8789043 | 0.109 | 0.003 | 0 | Chondrocytes | syt5a |

| | | | | | | | |
|---|---|---|---|---|---|---|---|
| si:ch211-153b23.5 | 0 | 4.9627996 | 0.112 | 0.007 | 0 | Chondrocytes | si:ch211-153b23.5 |
| hsc70 | 0 | 4.6395597 | 0.1 | 0.003 | 0 | Chondrocytes | hsc70 |
| mybl1 | 0 | 4.8705178 | 0.1 | 0.003 | 0 | Chondrocytes | mybl1 |
| sparc3 | 0 | 1.418701 | 0.997 | 0.98 | 0 | Chondrocytes | sparc |
| cpne21 | 3E-306 | 2.4017744 | 0.322 | 0.059 | 8.9E-302 | Chondrocytes | cpne2 |
| tgfb1a3 | 9E-304 | 2.0962092 | 0.688 | 0.261 | 3E-299 | Chondrocytes | tgfb1a |
| foxc1a | 2E-302 | 2.9585208 | 0.168 | 0.017 | 6.2E-298 | Chondrocytes | foxc1a |
| copz22 | 1E-298 | 1.9690862 | 0.57 | 0.176 | 3.5E-294 | Chondrocytes | copz2 |
| si:dkey-204f11.64 | 1E-298 | 2.2913127 | 0.353 | 0.071 | 4.2E-294 | Chondrocytes | si:dkey-204f11.64 |
| plcd3b | 2E-298 | 4.0150393 | 0.117 | 0.008 | 6.1E-294 | Chondrocytes | plcd3b |
| cthrc1a2 | 6E-298 | 1.8831477 | 0.515 | 0.143 | 2E-293 | Chondrocytes | cthrc1a |
| sgsm3 | 1E-296 | 3.5442229 | 0.121 | 0.008 | 4.6E-292 | Chondrocytes | sgsm3 |
| nanos14 | 9E-296 | 1.9130818 | 0.598 | 0.193 | 3E-291 | Chondrocytes | nanos1 |
| fibinb2 | 3E-293 | 1.9000709 | 0.683 | 0.262 | 9.5E-289 | Chondrocytes | fibinb |
| eva1bb1 | 1E-291 | 2.2800822 | 0.237 | 0.034 | 4.4E-287 | Chondrocytes | eva1bb |
| ldha2 | 2E-291 | 2.061471 | 0.625 | 0.218 | 5.2E-287 | Chondrocytes | ldha |
| aopep2 | 4E-281 | 2.1562107 | 0.336 | 0.068 | 1.3E-276 | Chondrocytes | aopep |
| gcnt4a | 2E-274 | 3.3588396 | 0.105 | 0.007 | 6.1E-270 | Chondrocytes | gcnt4a |
| cetn4 | 1E-273 | 2.6470236 | 0.281 | 0.05 | 4.7E-269 | Chondrocytes | cetn4 |
| myo5aa | 3E-273 | 2.8393029 | 0.143 | 0.013 | 1.1E-268 | Chondrocytes | myo5aa |
| ACBD3 | 1E-271 | 2.4496766 | 0.223 | 0.032 | 4.8E-267 | Chondrocytes | ACBD3 |
| BCR | 6E-270 | 3.3825226 | 0.112 | 0.008 | 2.1E-265 | Chondrocytes | BCR |
| csrnp1a | 7E-269 | 2.6661464 | 0.202 | 0.027 | 2.2E-264 | Chondrocytes | csrnp1a |
| crispld1a1 | 9E-268 | 3.3768853 | 0.244 | 0.04 | 2.8E-263 | Chondrocytes | crispld1a |
| ace1 | 3E-266 | 2.2370848 | 0.261 | 0.044 | 8.8E-262 | Chondrocytes | ace |
| phlda32 | 4E-264 | 1.7621793 | 0.541 | 0.169 | 1.3E-259 | Chondrocytes | phlda3 |
| AL929222.2 | 3E-259 | 3.5196042 | 0.179 | 0.023 | 1.1E-254 | Chondrocytes | AL929222.2 |
| tnika1 | 4E-259 | 2.7167943 | 0.192 | 0.026 | 1.4E-254 | Chondrocytes | tnika |
| arhgef1b1 | 2E-258 | 1.5416251 | 0.671 | 0.248 | 6.3E-254 | Chondrocytes | arhgef1b |
| plod31 | 6E-256 | 1.9416074 | 0.481 | 0.141 | 2E-251 | Chondrocytes | plod3 |
| sh3kbp11 | 2E-253 | 1.8733093 | 0.449 | 0.123 | 5.1E-249 | Chondrocytes | sh3kbp1 |
| slc37a2 | 2E-252 | 1.906736 | 0.414 | 0.106 | 7E-248 | Chondrocytes | slc37a2 |
| sat1b3 | 6E-252 | 1.9489593 | 0.402 | 0.102 | 2.1E-247 | Chondrocytes | sat1b |
| gpia1 | 4E-248 | 1.894965 | 0.435 | 0.119 | 1.3E-243 | Chondrocytes | gpia |
| inpp4b | 7E-247 | 3.3709649 | 0.105 | 0.008 | 2.2E-242 | Chondrocytes | inpp4b |
| myo6b | 3E-245 | 3.3361392 | 0.106 | 0.008 | 8.1E-241 | Chondrocytes | myo6b |
| vdac32 | 5E-243 | 1.407967 | 0.855 | 0.517 | 1.7E-238 | Chondrocytes | vdac3 |
| bambia2 | 1E-242 | 2.3808032 | 0.457 | 0.139 | 4E-238 | Chondrocytes | bambia |
| timp4.33 | 1E-241 | 2.6606946 | 0.758 | 0.431 | 4.6E-237 | Chondrocytes | timp4.3 |
| rarga2 | 5E-240 | 1.6830038 | 0.663 | 0.266 | 1.7E-235 | Chondrocytes | rarga |
| stc2a2 | 3E-239 | 2.297214 | 0.257 | 0.048 | 1.1E-234 | Chondrocytes | stc2a |
| hivep12 | 2E-237 | 1.7313912 | 0.479 | 0.144 | 7.9E-233 | Chondrocytes | hivep1 |
| sort1a3 | 8E-235 | 2.357869 | 0.393 | 0.106 | 2.6E-230 | Chondrocytes | sort1a |
| eno1a1 | 5E-234 | 1.9670597 | 0.702 | 0.329 | 1.6E-229 | Chondrocytes | eno1a |
| rftn21 | 2E-233 | 2.6606212 | 0.17 | 0.022 | 6.9E-229 | Chondrocytes | rftn2 |
| krt18a.13 | 1E-232 | 1.2613535 | 0.977 | 0.838 | 3.4E-228 | Chondrocytes | krt18a.1 |
| serpine13 | 4E-231 | 1.7561794 | 0.94 | 0.757 | 1.4E-226 | Chondrocytes | serpine1 |
| dkk3b2 | 3E-230 | 1.1141333 | 0.642 | 0.236 | 8.6E-226 | Chondrocytes | dkk3b |

| | | | | | | | |
|---|---|---|---|---|---|---|---|
| prelp4 | 9E-228 | 1.6614644 | 0.44 | 0.124 | 2.8E-223 | Chondrocytes | prelp |
| shisa3 | 1E-227 | 3.2972032 | 0.1 | 0.008 | 3.1E-223 | Chondrocytes | shisa3 |
| aldoaa2 | 1E-220 | 1.4111015 | 0.741 | 0.357 | 4.5E-216 | Chondrocytes | aldoaa |
| slc31a12 | 4E-219 | 1.6477535 | 0.535 | 0.188 | 1.3E-214 | Chondrocytes | slc31a1 |
| abtb2b1 | 1E-218 | 1.8826597 | 0.305 | 0.068 | 4.6E-214 | Chondrocytes | abtb2b |
| plekhg5a3 | 1E-217 | 2.0386474 | 0.297 | 0.065 | 3.6E-213 | Chondrocytes | plekhg5a |
| rab11fip3 | 3E-216 | 2.224931 | 0.232 | 0.042 | 1.1E-211 | Chondrocytes | rab11fip3 |
| lrba2 | 1E-214 | 1.7629298 | 0.503 | 0.167 | 3.9E-210 | Chondrocytes | lrba |
| si:dkey-32e6.61 | 2E-211 | 3.7515344 | 0.186 | 0.029 | 6.6E-207 | Chondrocytes | si:dkey-32e6.6 |
| dap2 | 1E-208 | 1.63745 | 0.524 | 0.185 | 3.4E-204 | Chondrocytes | dap |
| myo1ea2 | 1E-207 | 1.8532412 | 0.315 | 0.075 | 3.5E-203 | Chondrocytes | myo1ea |
| mydgf2 | 2E-207 | 1.9533897 | 0.309 | 0.073 | 5.2E-203 | Chondrocytes | mydgf |
| hexb2 | 2E-207 | 1.8085268 | 0.337 | 0.084 | 7.6E-203 | Chondrocytes | hexb |
| gpc42 | 3E-207 | 1.4715544 | 0.605 | 0.232 | 1.1E-202 | Chondrocytes | gpc4 |
| chst112 | 9E-207 | 2.2550652 | 0.215 | 0.038 | 3E-202 | Chondrocytes | chst11 |
| nupr1b1 | 1E-204 | 2.5971009 | 0.458 | 0.154 | 4.2E-200 | Chondrocytes | nupr1b |
| si:rp71-62i8.1 | 1E-204 | 1.9243716 | 0.247 | 0.049 | 4.6E-200 | Chondrocytes | si:rp71-62i8.1 |
| pde4ba2 | 1E-203 | 1.7728676 | 0.388 | 0.109 | 3.4E-199 | Chondrocytes | pde4ba |
| trpv4 | 5E-203 | 2.2729322 | 0.167 | 0.024 | 1.6E-198 | Chondrocytes | trpv4 |
| sdcbp21 | 4E-201 | 1.311591 | 0.878 | 0.59 | 1.4E-196 | Chondrocytes | sdcbp2 |
| clec11a1 | 6E-201 | 2.4314908 | 0.249 | 0.052 | 1.9E-196 | Chondrocytes | clec11a |
| ndel1b | 4E-197 | 1.8211158 | 0.342 | 0.09 | 1.2E-192 | Chondrocytes | ndel1b |
| slc10a7 | 2E-196 | 2.2793746 | 0.189 | 0.031 | 6E-192 | Chondrocytes | slc10a7 |
| entpd5a2 | 7E-196 | 3.4645117 | 0.234 | 0.048 | 2.3E-191 | Chondrocytes | entpd5a |
| ldlrap1a1 | 8E-194 | 1.9226015 | 0.276 | 0.063 | 2.7E-189 | Chondrocytes | ldlrap1a |
| steap42 | 2E-193 | 1.0878383 | 0.504 | 0.174 | 7.5E-189 | Chondrocytes | steap4 |
| tpi1b1 | 1E-190 | 1.4923452 | 0.696 | 0.348 | 4.7E-186 | Chondrocytes | tpi1b |
| slc38a3a | 4E-189 | 3.0250708 | 0.122 | 0.014 | 1.4E-184 | Chondrocytes | slc38a3a |
| tmem45a2 | 1E-188 | 1.6984911 | 0.448 | 0.149 | 4.4E-184 | Chondrocytes | tmem45a |
| capgb2 | 7E-188 | 1.6632632 | 0.351 | 0.096 | 2.4E-183 | Chondrocytes | capgb |
| peli1b3 | 8E-188 | 1.5515763 | 0.326 | 0.083 | 2.5E-183 | Chondrocytes | peli1b |
| cpamd81 | 6E-187 | 2.6423824 | 0.248 | 0.055 | 2E-182 | Chondrocytes | cpamd8 |
| znf185 | 2E-186 | 3.1157566 | 0.105 | 0.011 | 7.6E-182 | Chondrocytes | znf185 |
| bmp53 | 3E-186 | 1.9534742 | 0.315 | 0.081 | 8.8E-182 | Chondrocytes | bmp5 |
| nog32 | 2E-185 | 2.0121698 | 0.301 | 0.076 | 5E-181 | Chondrocytes | nog3 |
| slc26a2 | 2E-185 | 2.005957 | 0.208 | 0.039 | 6.7E-181 | Chondrocytes | slc26a2 |
| cdc42se1 | 4E-185 | 1.9144462 | 0.291 | 0.071 | 1.1E-180 | Chondrocytes | cdc42se1 |
| crtap1 | 5E-185 | 1.3381302 | 0.667 | 0.309 | 1.7E-180 | Chondrocytes | crtap |
| ehd2b1 | 5E-183 | 1.4585422 | 0.832 | 0.552 | 1.7E-178 | Chondrocytes | ehd2b |
| iqsec1b1 | 3E-182 | 1.5449788 | 0.689 | 0.334 | 8.6E-178 | Chondrocytes | iqsec1b |
| fam114a12 | 8E-182 | 1.6563244 | 0.359 | 0.103 | 2.5E-177 | Chondrocytes | fam114a1 |
| nat16 | 1E-180 | 2.6244776 | 0.125 | 0.015 | 3.4E-176 | Chondrocytes | nat16 |
| nucb2b2 | 2E-179 | 1.5715266 | 0.374 | 0.111 | 6.4E-175 | Chondrocytes | nucb2b |
| rab13 | 5E-179 | 1.4045985 | 0.548 | 0.213 | 1.5E-174 | Chondrocytes | rab13 |
| gclm | 5E-179 | 2.1884516 | 0.255 | 0.058 | 1.7E-174 | Chondrocytes | gclm |
| si:ch1073-184j22.13 | 7E-179 | 1.9260079 | 0.294 | 0.074 | 2.2E-174 | Chondrocytes | si:ch1073-184j22.1 |
| tmem591 | 4E-178 | 1.1950332 | 0.678 | 0.303 | 1.2E-173 | Chondrocytes | tmem59 |
| grasp | 5E-178 | 2.4717049 | 0.169 | 0.028 | 1.6E-173 | Chondrocytes | grasp |

| | | | | | | | |
|---|---|---|---|---|---|---|---|
| tex264a | 4E-177 | 1.7240418 | 0.375 | 0.114 | 1.3E-172 | Chondrocytes | tex264a |
| rpl22l13 | 4E-177 | 0.8383183 | 0.953 | 0.742 | 1.4E-172 | Chondrocytes | rpl22l1 |
| rrbp1b2 | 7E-177 | 1.4540189 | 0.568 | 0.229 | 2.3E-172 | Chondrocytes | rrbp1b |
| ndnf4 | 2E-176 | 1.8107288 | 0.34 | 0.097 | 5.9E-172 | Chondrocytes | ndnf |
| slc16a3 | 2E-176 | 2.4873822 | 0.162 | 0.026 | 6.9E-172 | Chondrocytes | slc16a3 |
| trps14 | 1E-175 | 1.4646111 | 0.475 | 0.165 | 3.8E-171 | Chondrocytes | trps1 |
| cyth1b | 3E-175 | 2.1064415 | 0.211 | 0.042 | 1E-170 | Chondrocytes | cyth1b |
| oaz1b3 | 2E-174 | 1.3641573 | 0.645 | 0.297 | 7.9E-170 | Chondrocytes | oaz1b |
| fthl31 | 1E-173 | 3.2090242 | 0.181 | 0.033 | 3.8E-169 | Chondrocytes | fthl31 |
| atp2b1a | 3E-170 | 1.5340298 | 0.571 | 0.237 | 9.6E-166 | Chondrocytes | atp2b1a |
| C7H16orf872 | 5E-170 | 1.5117178 | 0.384 | 0.119 | 1.6E-165 | Chondrocytes | C7H16orf87 |
| gsna | 4E-169 | 1.2419324 | 0.699 | 0.348 | 1.4E-164 | Chondrocytes | gsna |
| spsb4a | 2E-168 | 2.0551527 | 0.192 | 0.036 | 7.8E-164 | Chondrocytes | spsb4a |
| ackr4b1 | 7E-168 | 1.7702109 | 0.22 | 0.046 | 2.2E-163 | Chondrocytes | ackr4b |
| fam20b1 | 3E-167 | 1.8030812 | 0.254 | 0.06 | 9.2E-163 | Chondrocytes | fam20b |
| golim4b2 | 2E-166 | 1.4147406 | 0.35 | 0.102 | 7.3E-162 | Chondrocytes | golim4b |
| CR381618.15 | 1E-165 | 1.4157916 | 0.542 | 0.218 | 4E-161 | Chondrocytes | CR381618.1 |
| micu3a | 2E-165 | 2.3340797 | 0.151 | 0.024 | 4.9E-161 | Chondrocytes | micu3a |
| prnprs31 | 3E-164 | 1.6172625 | 0.32 | 0.09 | 8.3E-160 | Chondrocytes | prnprs3 |
| ppp1r1c2 | 9E-164 | 2.1204685 | 0.382 | 0.124 | 3E-159 | Chondrocytes | ppp1r1c |
| atp2a3 | 3E-163 | 1.8134689 | 0.192 | 0.037 | 1E-158 | Chondrocytes | atp2a3 |
| ppfibp1a1 | 2E-161 | 1.7179011 | 0.228 | 0.05 | 6.5E-157 | Chondrocytes | ppfibp1a |
| tnfrsfa | 2E-160 | 2.3471292 | 0.125 | 0.017 | 5.9E-156 | Chondrocytes | tnfrsfa |
| rhoab | 3E-160 | 1.3162445 | 0.758 | 0.449 | 9E-156 | Chondrocytes | rhoab |
| mgp1 | 7E-160 | 2.7409224 | 0.416 | 0.152 | 2.4E-155 | Chondrocytes | mgp |
| smoc23 | 1E-158 | 2.1477141 | 0.301 | 0.084 | 4.5E-154 | Chondrocytes | smoc2 |
| mfge8b1 | 3E-157 | 1.9050149 | 0.227 | 0.052 | 1.1E-152 | Chondrocytes | mfge8b |
| arg24 | 3E-156 | 1.7761374 | 0.426 | 0.156 | 8.2E-152 | Chondrocytes | arg2 |
| p4ha22 | 4E-155 | 1.5108625 | 0.397 | 0.135 | 1.3E-150 | Chondrocytes | p4ha2 |
| emilin1a4 | 5E-155 | 1.5885814 | 0.31 | 0.087 | 1.5E-150 | Chondrocytes | emilin1a |
| rassf13 | 6E-155 | 1.6557488 | 0.356 | 0.114 | 1.9E-150 | Chondrocytes | rassf1 |
| xylt1 | 1E-153 | 1.6064847 | 0.205 | 0.043 | 3.7E-149 | Chondrocytes | xylt1 |
| rnf144ab | 1E-152 | 2.5083174 | 0.126 | 0.018 | 4.4E-148 | Chondrocytes | rnf144ab |
| gale1 | 2E-152 | 2.2846152 | 0.214 | 0.048 | 5.3E-148 | Chondrocytes | gale |
| arpc1a1 | 4E-152 | 1.1691883 | 0.648 | 0.308 | 1.3E-147 | Chondrocytes | arpc1a |
| hspg22 | 8E-152 | 1.3303844 | 0.411 | 0.138 | 2.6E-147 | Chondrocytes | hspg2 |
| sdf42 | 6E-150 | 1.2724882 | 0.626 | 0.294 | 2E-145 | Chondrocytes | sdf4 |
| ivns1abpa4 | 9E-150 | 1.3309116 | 0.602 | 0.277 | 2.9E-145 | Chondrocytes | ivns1abpa |
| smad72 | 5E-149 | 1.5858121 | 0.408 | 0.146 | 1.5E-144 | Chondrocytes | smad7 |
| slc39a132 | 1E-148 | 1.440256 | 0.393 | 0.134 | 3.6E-144 | Chondrocytes | slc39a13 |
| mmp22 | 1E-147 | 0.6770703 | 0.966 | 0.738 | 3.5E-143 | Chondrocytes | mmp2 |
| irf2bp2b2 | 1E-147 | 1.5091756 | 0.493 | 0.201 | 3.7E-143 | Chondrocytes | irf2bp2b |
| ppa1b2 | 2E-146 | 1.4360111 | 0.385 | 0.131 | 6.9E-142 | Chondrocytes | ppa1b |
| CR848833.13 | 3E-146 | 1.6139844 | 0.258 | 0.067 | 1.1E-141 | Chondrocytes | CR848833.1 |
| agxta | 3E-145 | 2.6249437 | 0.1 | 0.012 | 1.1E-140 | Chondrocytes | agxta |
| nfixb4 | 5E-145 | 1.1694391 | 0.62 | 0.284 | 1.6E-140 | Chondrocytes | nfixb |
| tmed33 | 5E-145 | 1.5233839 | 0.307 | 0.09 | 1.7E-140 | Chondrocytes | tmed3 |
| cav22 | 9E-145 | 1.3718161 | 0.443 | 0.164 | 2.8E-140 | Chondrocytes | cav2 |

| | | | | | | | |
|---|---|---|---|---|---|---|---|
| lrrfip1a2 | 1E-144 | 1.8355652 | 0.257 | 0.068 | 4.4E-140 | Chondrocytes | lrrfip1a |
| fam83d | 4E-144 | 2.4523359 | 0.116 | 0.017 | 1.4E-139 | Chondrocytes | fam83d |
| sgcg2 | 6E-144 | 1.5071715 | 0.297 | 0.085 | 2.1E-139 | Chondrocytes | sgcg |
| pnp5b | 1E-143 | 1.9183778 | 0.175 | 0.035 | 4.3E-139 | Chondrocytes | pnp5b |
| mfge8a2 | 2E-143 | 1.2386479 | 0.355 | 0.113 | 6.3E-139 | Chondrocytes | mfge8a |
| clstn12 | 3E-142 | 1.1824809 | 0.388 | 0.129 | 8.5E-138 | Chondrocytes | clstn1 |
| igflr1 | 6E-142 | 1.9203294 | 0.183 | 0.038 | 2.1E-137 | Chondrocytes | igflr1 |
| uxs1 | 7E-142 | 1.5070057 | 0.25 | 0.064 | 2.3E-137 | Chondrocytes | uxs1 |
| mthfd2 | 9E-142 | 1.767382 | 0.232 | 0.058 | 3E-137 | Chondrocytes | mthfd2 |
| ucmaa | 2E-141 | 1.835121 | 0.183 | 0.039 | 6.1E-137 | Chondrocytes | ucmaa |
| nkx3-24 | 4E-141 | 1.4952279 | 0.469 | 0.188 | 1.2E-136 | Chondrocytes | nkx3-2 |
| foxp44 | 5E-141 | 1.1904973 | 0.493 | 0.195 | 1.7E-136 | Chondrocytes | foxp4 |
| mfsd2ab | 8E-141 | 1.9368409 | 0.178 | 0.037 | 2.7E-136 | Chondrocytes | mfsd2ab |
| schip12 | 1E-140 | 1.3243112 | 0.397 | 0.138 | 3.9E-136 | Chondrocytes | schip1 |
| eya2 | 2E-140 | 2.1357506 | 0.141 | 0.024 | 7.4E-136 | Chondrocytes | eya2 |
| tcea31 | 2E-139 | 2.2426681 | 0.139 | 0.024 | 6.6E-135 | Chondrocytes | tcea3 |
| slc38a4 | 1E-138 | 2.3832308 | 0.1 | 0.013 | 3.9E-134 | Chondrocytes | slc38a4 |
| usp6nl1 | 2E-138 | 1.9924661 | 0.172 | 0.035 | 6.9E-134 | Chondrocytes | usp6nl |
| jagn1b1 | 5E-138 | 1.9034177 | 0.411 | 0.154 | 1.5E-133 | Chondrocytes | jagn1b |
| stau21 | 9E-138 | 1.0160177 | 0.755 | 0.429 | 2.9E-133 | Chondrocytes | stau2 |
| cntf | 2E-137 | 2.2497318 | 0.146 | 0.026 | 6.7E-133 | Chondrocytes | cntf |
| hdac43 | 2E-137 | 1.2643469 | 0.352 | 0.113 | 7.5E-133 | Chondrocytes | hdac4 |
| nfatc12 | 2E-137 | 1.3979443 | 0.284 | 0.08 | 7.8E-133 | Chondrocytes | nfatc1 |
| cc2d1b | 1E-136 | 1.967479 | 0.171 | 0.035 | 3.9E-132 | Chondrocytes | cc2d1b |
| il4r.12 | 5E-136 | 1.3463359 | 0.402 | 0.145 | 1.7E-131 | Chondrocytes | il4r.1 |
| sytl1 | 4E-135 | 2.3721696 | 0.103 | 0.014 | 1.2E-130 | Chondrocytes | sytl1 |
| calm1a3 | 1E-134 | 1.1316452 | 0.86 | 0.611 | 4.5E-130 | Chondrocytes | calm1a |
| vkorc1 | 2E-134 | 1.6965606 | 0.199 | 0.046 | 7.2E-130 | Chondrocytes | vkorc1 |
| CR376783.1 | 3E-133 | 2.2645283 | 0.127 | 0.021 | 9.5E-129 | Chondrocytes | CR376783.1 |
| slc38a10 | 2E-129 | 1.0414177 | 0.525 | 0.221 | 6.9E-125 | Chondrocytes | slc38a10 |
| timp2a4 | 1E-128 | 0.7537591 | 0.949 | 0.75 | 3.7E-124 | Chondrocytes | timp2a |
| atp1a1a.13 | 1E-128 | 1.044513 | 0.636 | 0.313 | 4.4E-124 | Chondrocytes | atp1a1a.1 |
| calr3b2 | 5E-128 | 1.1317284 | 0.57 | 0.261 | 1.5E-123 | Chondrocytes | calr3b |
| fndc3ba2 | 8E-128 | 1.275702 | 0.374 | 0.132 | 2.6E-123 | Chondrocytes | fndc3ba |
| rrad1 | 9E-128 | 1.7712938 | 0.239 | 0.065 | 2.8E-123 | Chondrocytes | rrad |
| ndrg3a | 6E-126 | 1.5346898 | 0.294 | 0.092 | 1.9E-121 | Chondrocytes | ndrg3a |
| CPNE85 | 6E-126 | 1.1922627 | 0.577 | 0.278 | 2.1E-121 | Chondrocytes | CPNE8 |
| rnf144aa | 2E-125 | 2.1425738 | 0.128 | 0.022 | 7.9E-121 | Chondrocytes | rnf144aa |
| angptl2b2 | 2E-124 | 1.1456128 | 0.374 | 0.134 | 7E-120 | Chondrocytes | angptl2b |
| psph1 | 2E-124 | 1.9568562 | 0.152 | 0.03 | 8.1E-120 | Chondrocytes | psph |
| pgd1 | 1E-123 | 1.4169452 | 0.257 | 0.074 | 3.7E-119 | Chondrocytes | pgd |
| zgc:771581 | 2E-123 | 1.9256268 | 0.217 | 0.057 | 6E-119 | Chondrocytes | zgc:77158 |
| col6a14 | 6E-123 | 0.7551343 | 0.977 | 0.872 | 1.8E-118 | Chondrocytes | col6a1 |
| pdcd61 | 5E-122 | 1.0674515 | 0.752 | 0.469 | 1.5E-117 | Chondrocytes | pdcd6 |
| slc25a3b2 | 6E-122 | 0.8269724 | 0.832 | 0.543 | 1.8E-117 | Chondrocytes | slc25a3b |
| pld73 | 1E-121 | 1.2600025 | 0.389 | 0.146 | 4.5E-117 | Chondrocytes | pld7 |
| olfml3b3 | 7E-121 | 0.9312055 | 0.8 | 0.506 | 2.4E-116 | Chondrocytes | olfml3b |
| zgc:112148 | 1E-120 | 1.8822517 | 0.151 | 0.031 | 3.9E-116 | Chondrocytes | zgc:112148 |

| | | | | | | | |
|---|---|---|---|---|---|---|---|
| impdh1b3 | 4E-120 | 1.5511949 | 0.261 | 0.078 | 1.1E-115 | Chondrocytes | impdh1b |
| rcn33 | 5E-120 | 1.1468533 | 0.545 | 0.255 | 1.7E-115 | Chondrocytes | rcn3 |
| tsc22d13 | 1E-119 | 1.3815511 | 0.324 | 0.109 | 3.3E-115 | Chondrocytes | tsc22d1 |
| MDFIC1 | 2E-119 | 1.0628772 | 0.448 | 0.179 | 7E-115 | Chondrocytes | MDFIC |
| cmpk1 | 3E-119 | 1.1891968 | 0.408 | 0.157 | 9.9E-115 | Chondrocytes | cmpk |
| mroh1 | 4E-119 | 1.5249751 | 0.187 | 0.044 | 1.2E-114 | Chondrocytes | mroh1 |
| lepr2 | 6E-119 | 1.4397885 | 0.222 | 0.059 | 1.9E-114 | Chondrocytes | lepr |
| nupr1a2 | 7E-119 | 1.252339 | 0.543 | 0.248 | 2.4E-114 | Chondrocytes | nupr1a |
| adgre5b.3.1 | 2E-118 | 2.1389108 | 0.117 | 0.02 | 4.9E-114 | Chondrocytes | adgre5b.3.1 |
| pkma1 | 4E-118 | 1.2372197 | 0.476 | 0.205 | 1.2E-113 | Chondrocytes | pkma |
| snap23.11 | 5E-118 | 1.1200207 | 0.439 | 0.175 | 1.7E-113 | Chondrocytes | snap23.1 |
| foxred2 | 7E-118 | 2.3155727 | 0.138 | 0.027 | 2.4E-113 | Chondrocytes | foxred2 |
| tox1 | 1E-117 | 2.0316477 | 0.208 | 0.055 | 3.5E-113 | Chondrocytes | tox |
| stard13b2 | 5E-117 | 1.2753359 | 0.264 | 0.079 | 1.6E-112 | Chondrocytes | stard13b |
| SBNO23 | 8E-116 | 1.0769391 | 0.5 | 0.22 | 2.5E-111 | Chondrocytes | SBNO2 |
| etv5a3 | 1E-115 | 1.2789769 | 0.3 | 0.098 | 4E-111 | Chondrocytes | etv5a |
| cdkn1bb2 | 2E-115 | 1.2219056 | 0.583 | 0.289 | 7.1E-111 | Chondrocytes | cdkn1bb |
| CT030129.1 | 6E-115 | 2.0280579 | 0.122 | 0.022 | 1.8E-110 | Chondrocytes | CT030129.1 |
| pdia61 | 7E-115 | 1.3180964 | 0.449 | 0.192 | 2.2E-110 | Chondrocytes | pdia6 |
| pfkpb | 9E-115 | 1.9522162 | 0.114 | 0.019 | 3E-110 | Chondrocytes | pfkpb |
| neo1a2 | 5E-114 | 1.1187197 | 0.432 | 0.175 | 1.6E-109 | Chondrocytes | neo1a |
| cspg41 | 9E-114 | 1.2724066 | 0.318 | 0.108 | 2.9E-109 | Chondrocytes | cspg4 |
| emx24 | 1E-113 | 1.448793 | 0.275 | 0.087 | 4.6E-109 | Chondrocytes | emx2 |
| papss2a | 9E-113 | 2.0170049 | 0.111 | 0.019 | 2.9E-108 | Chondrocytes | papss2a |
| si:ch211-176g6.2 | 2E-112 | 2.2833924 | 0.113 | 0.02 | 5.5E-108 | Chondrocytes | si:ch211-176g6.2 |
| slc25a431 | 5E-112 | 1.5431181 | 0.212 | 0.057 | 1.6E-107 | Chondrocytes | slc25a43 |
| arhgef1a1 | 6E-112 | 1.6091407 | 0.209 | 0.056 | 1.9E-107 | Chondrocytes | arhgef1a |
| kif13ba1 | 8E-111 | 1.3697718 | 0.294 | 0.097 | 2.7E-106 | Chondrocytes | kif13ba |
| olfml3a2 | 2E-110 | 1.0038176 | 0.536 | 0.25 | 7.1E-106 | Chondrocytes | olfml3a |
| nucb2a2 | 3E-110 | 0.8598221 | 0.802 | 0.516 | 9.8E-106 | Chondrocytes | nucb2a |
| alas12 | 9E-110 | 1.4733088 | 0.297 | 0.101 | 2.9E-105 | Chondrocytes | alas1 |
| slc2a10 | 5E-109 | 1.7741363 | 0.159 | 0.036 | 1.7E-104 | Chondrocytes | slc2a10 |
| lpin11 | 2E-108 | 1.7354815 | 0.21 | 0.058 | 6.7E-104 | Chondrocytes | lpin1 |
| uggt21 | 2E-108 | 1.6181781 | 0.193 | 0.05 | 6.8E-104 | Chondrocytes | uggt2 |
| adcy7 | 3E-108 | 2.0936537 | 0.125 | 0.024 | 8.5E-104 | Chondrocytes | adcy7 |
| ifrd12 | 3E-108 | 1.0595376 | 0.558 | 0.275 | 9E-104 | Chondrocytes | ifrd1 |
| pitpnaa1 | 3E-108 | 1.8429401 | 0.192 | 0.05 | 9E-104 | Chondrocytes | pitpnaa |
| si:ch73-217n20.11 | 4E-108 | 1.4565633 | 0.189 | 0.048 | 1.2E-103 | Chondrocytes | si:ch73-217n20.1 |
| zgc:92606 | 4E-108 | 1.185824 | 0.346 | 0.127 | 1.4E-103 | Chondrocytes | zgc:92606 |
| fam53b | 1E-107 | 1.3604943 | 0.214 | 0.059 | 3.2E-103 | Chondrocytes | fam53b |
| slc66a32 | 2E-107 | 1.155254 | 0.322 | 0.113 | 6.1E-103 | Chondrocytes | slc66a3 |
| degs11 | 3E-107 | 1.2147923 | 0.427 | 0.181 | 9.3E-103 | Chondrocytes | degs1 |
| zgc:921621 | 5E-107 | 1.5641338 | 0.138 | 0.029 | 1.6E-102 | Chondrocytes | zgc:92162 |
| gpx4a2 | 1E-106 | 0.9679546 | 0.587 | 0.295 | 4.8E-102 | Chondrocytes | gpx4a |
| tbrg11 | 2E-106 | 1.8728955 | 0.18 | 0.046 | 7.4E-102 | Chondrocytes | tbrg1 |
| chsy3 | 6E-106 | 1.7830324 | 0.159 | 0.037 | 1.9E-101 | Chondrocytes | chsy3 |
| atp10a2 | 2E-105 | 1.3546668 | 0.278 | 0.092 | 5.3E-101 | Chondrocytes | atp10a |
| gapdhs1 | 2E-105 | 1.0993481 | 0.838 | 0.623 | 6.8E-101 | Chondrocytes | gapdhs |

| | | | | | | | |
|---|---|---|---|---|---|---|---|
| ptgs2a3 | 6E-105 | 1.51867 | 0.331 | 0.124 | 1.9E-100 | Chondrocytes | ptgs2a |
| nt5c2b1 | 1E-104 | 1.4589429 | 0.206 | 0.057 | 3.8E-100 | Chondrocytes | nt5c2b |
| reep3b | 2E-104 | 1.0859575 | 0.405 | 0.165 | 5.8E-100 | Chondrocytes | reep3b |
| pdia53 | 2E-104 | 1.0961403 | 0.396 | 0.159 | 7.5E-100 | Chondrocytes | pdia5 |
| col8a21 | 3E-104 | 2.3541927 | 0.214 | 0.063 | 1.1E-99 | Chondrocytes | col8a2 |
| zgc:565252 | 4E-104 | 0.9953359 | 0.381 | 0.148 | 1.3E-99 | Chondrocytes | zgc:56525 |
| cpn15 | 5E-104 | 1.3365439 | 0.454 | 0.205 | 1.8E-99 | Chondrocytes | cpn1 |
| tgif13 | 6E-104 | 1.564057 | 0.434 | 0.199 | 1.8E-99 | Chondrocytes | tgif1 |
| relt | 6E-104 | 1.759942 | 0.139 | 0.03 | 2E-99 | Chondrocytes | relt |
| atp6v1e1b2 | 8E-104 | 1.244409 | 0.281 | 0.094 | 2.6E-99 | Chondrocytes | atp6v1e1b |
| fzd7a1 | 9E-104 | 1.3600264 | 0.244 | 0.075 | 3.1E-99 | Chondrocytes | fzd7a |
| samd4a | 2E-103 | 1.3434172 | 0.211 | 0.059 | 8.1E-99 | Chondrocytes | samd4a |
| nfatc3a1 | 4E-103 | 1.48752 | 0.268 | 0.088 | 1.29E-98 | Chondrocytes | nfatc3a |
| hephl1a | 7E-103 | 1.7907332 | 0.131 | 0.027 | 2.33E-98 | Chondrocytes | hephl1a |
| ctnnb25 | 1E-102 | 0.9040596 | 0.601 | 0.304 | 4.72E-98 | Chondrocytes | ctnnb2 |
| gadd45bb4 | 2E-102 | 1.252431 | 0.464 | 0.213 | 5.93E-98 | Chondrocytes | gadd45bb |
| selenow2a1 | 2E-102 | 0.9763935 | 0.393 | 0.157 | 7.41E-98 | Chondrocytes | selenow2a |
| MYO9B2 | 2E-102 | 1.2197214 | 0.323 | 0.117 | 8.04E-98 | Chondrocytes | MYO9B |
| slc20a1b2 | 9E-102 | 0.8135481 | 0.581 | 0.286 | 3.02E-97 | Chondrocytes | slc20a1b |
| marveld12 | 1E-101 | 0.9363004 | 0.464 | 0.202 | 3.93E-97 | Chondrocytes | marveld1 |
| prg4b | 5E-101 | 1.5781295 | 0.177 | 0.046 | 1.7E-96 | Chondrocytes | prg4b |
| serpinh22 | 9E-101 | 1.0071654 | 0.316 | 0.112 | 2.8E-96 | Chondrocytes | serpinh2 |
| kdelr33 | 4E-100 | 1.091755 | 0.417 | 0.176 | 1.22E-95 | Chondrocytes | kdelr3 |
| calua2 | 6E-100 | 0.9401007 | 0.605 | 0.318 | 1.9E-95 | Chondrocytes | calua |
| b3gnt2b | 9E-100 | 1.4432293 | 0.225 | 0.067 | 3.01E-95 | Chondrocytes | b3gnt2b |
| triqk2 | 3E-99 | 1.111019 | 0.383 | 0.156 | 1.04E-94 | Chondrocytes | triqk |
| ctsh2 | 6E-99 | 1.2993699 | 0.295 | 0.105 | 1.91E-94 | Chondrocytes | ctsh |
| extl2 | 1.2E-98 | 1.5308626 | 0.161 | 0.039 | 3.94E-94 | Chondrocytes | extl2 |
| fzd7b3 | 1.3E-97 | 1.0693777 | 0.297 | 0.104 | 4.16E-93 | Chondrocytes | fzd7b |
| ifi301 | 3.5E-97 | 0.9946454 | 0.359 | 0.14 | 1.13E-92 | Chondrocytes | ifi30 |
| scocb1 | 1.3E-96 | 1.2521098 | 0.244 | 0.078 | 4.3E-92 | Chondrocytes | scocb |
| mmp13b1 | 5.9E-96 | 1.9796684 | 0.133 | 0.029 | 1.93E-91 | Chondrocytes | mmp13b |
| b3glcta | 1E-95 | 1.256132 | 0.178 | 0.047 | 3.24E-91 | Chondrocytes | b3glcta |
| erf | 1.9E-95 | 1.4832247 | 0.214 | 0.065 | 6.17E-91 | Chondrocytes | erf |
| reep3a2 | 4.4E-95 | 1.006955 | 0.388 | 0.159 | 1.44E-90 | Chondrocytes | reep3a |
| wee11 | 1E-94 | 1.3808253 | 0.203 | 0.059 | 3.36E-90 | Chondrocytes | wee1 |
| ctsd2 | 3.9E-94 | 1.0297804 | 0.367 | 0.148 | 1.28E-89 | Chondrocytes | ctsd |
| tmed71 | 1.3E-93 | 0.8434233 | 0.596 | 0.308 | 4.32E-89 | Chondrocytes | tmed7 |
| gnpda1 | 1.9E-93 | 1.6906771 | 0.135 | 0.031 | 6.06E-89 | Chondrocytes | gnpda1 |
| si:ch211-132b12.73 | 9.4E-93 | 0.9304297 | 0.374 | 0.15 | 3.07E-88 | Chondrocytes | si:ch211-132b12.7 |
| ccng12 | 1.8E-92 | 1.1631434 | 0.748 | 0.498 | 5.84E-88 | Chondrocytes | ccng1 |
| eef2b4 | 5.1E-92 | 0.4911254 | 0.971 | 0.908 | 1.66E-87 | Chondrocytes | eef2b |
| snx172 | 2E-91 | 1.1168817 | 0.488 | 0.235 | 6.64E-87 | Chondrocytes | snx17 |
| pabpc1b2 | 8E-91 | 0.7309403 | 0.82 | 0.579 | 2.59E-86 | Chondrocytes | pabpc1b |
| npc21 | 1.6E-90 | 0.8277866 | 0.626 | 0.338 | 5.3E-86 | Chondrocytes | npc2 |
| cyp27c1 | 1.7E-90 | 1.4695025 | 0.158 | 0.041 | 5.45E-86 | Chondrocytes | cyp27c1 |
| abcf2a3 | 2E-90 | 1.0277918 | 0.397 | 0.172 | 6.54E-86 | Chondrocytes | abcf2a |
| rnasekb | 2.2E-90 | 1.0405309 | 0.284 | 0.101 | 7.14E-86 | Chondrocytes | rnasekb |

| | | | | | | | |
|---|---|---|---|---|---|---|---|
| chac11 | 7.1E-90 | 0.9684027 | 0.611 | 0.347 | 2.32E-85 | Chondrocytes | chac1 |
| ppp1r3ca3 | 8E-90 | 1.2516333 | 0.438 | 0.202 | 2.61E-85 | Chondrocytes | ppp1r3ca |
| prkd2 | 2.3E-89 | 1.0205321 | 0.322 | 0.123 | 7.46E-85 | Chondrocytes | prkd2 |
| neo1b2 | 2.8E-89 | 1.5215616 | 0.232 | 0.077 | 9.24E-85 | Chondrocytes | neo1b |
| ptmab3 | 3.1E-89 | 0.8825372 | 0.882 | 0.71 | 1.01E-84 | Chondrocytes | ptmab |
| BX510657.1 | 3.9E-89 | 1.8700255 | 0.114 | 0.024 | 1.27E-84 | Chondrocytes | BX510657.1 |
| emc102 | 6E-89 | 0.954381 | 0.349 | 0.139 | 1.94E-84 | Chondrocytes | emc10 |
| etv4 | 7.4E-89 | 2.113885 | 0.103 | 0.02 | 2.4E-84 | Chondrocytes | etv4 |
| gnai2a | 9.2E-89 | 0.8642986 | 0.546 | 0.275 | 2.99E-84 | Chondrocytes | gnai2a |
| atp6v1e1a1 | 1.2E-88 | 1.0431349 | 0.278 | 0.099 | 3.88E-84 | Chondrocytes | atp6v1e1a |
| olfml2bb2 | 2.1E-87 | 1.6782043 | 0.247 | 0.087 | 6.98E-83 | Chondrocytes | olfml2bb |
| slc12a41 | 2.2E-87 | 0.956201 | 0.314 | 0.12 | 7.32E-83 | Chondrocytes | slc12a4 |
| chmp2a | 3.5E-87 | 0.9082849 | 0.426 | 0.189 | 1.15E-82 | Chondrocytes | chmp2a |
| slc25a53 | 4.1E-87 | 0.4847327 | 0.986 | 0.962 | 1.32E-82 | Chondrocytes | slc25a5 |
| foxo1a1 | 6.8E-87 | 1.1839094 | 0.403 | 0.183 | 2.2E-82 | Chondrocytes | foxo1a |
| cgref12 | 8E-87 | 0.9187882 | 0.388 | 0.167 | 2.59E-82 | Chondrocytes | cgref1 |
| thbs4b4 | 9.6E-87 | 1.030708 | 0.488 | 0.253 | 3.13E-82 | Chondrocytes | thbs4b |
| efna5a3 | 4.9E-86 | 1.3604397 | 0.224 | 0.074 | 1.58E-81 | Chondrocytes | efna5a |
| antxr1c1 | 5.7E-86 | 0.7005033 | 0.816 | 0.529 | 1.85E-81 | Chondrocytes | antxr1c |
| bcr | 7E-86 | 1.8327602 | 0.119 | 0.026 | 2.29E-81 | Chondrocytes | bcr |
| pdgfrb3 | 1.3E-85 | 0.7732646 | 0.685 | 0.404 | 4.08E-81 | Chondrocytes | pdgfrb |
| chmp2bb1 | 1.4E-85 | 1.0391819 | 0.271 | 0.097 | 4.46E-81 | Chondrocytes | chmp2bb |
| slc25a333 | 1.6E-85 | 1.182503 | 0.296 | 0.112 | 5.23E-81 | Chondrocytes | slc25a33 |
| phactr23 | 5.9E-85 | 0.9717589 | 0.337 | 0.135 | 1.92E-80 | Chondrocytes | phactr2 |
| si:zfos-943e10.1 | 1.1E-84 | 1.224459 | 0.22 | 0.071 | 3.67E-80 | Chondrocytes | si:zfos-943e10.1 |
| tmem30aa | 2.5E-84 | 0.9302222 | 0.368 | 0.155 | 8.25E-80 | Chondrocytes | tmem30aa |
| fkbp112 | 2.8E-84 | 1.1067153 | 0.362 | 0.154 | 9.23E-80 | Chondrocytes | fkbp11 |
| vwa13 | 6.2E-84 | 0.8329929 | 0.332 | 0.133 | 2.03E-79 | Chondrocytes | vwa1 |
| srebf2 | 7.1E-84 | 1.6128966 | 0.12 | 0.027 | 2.32E-79 | Chondrocytes | srebf2 |
| selenot21 | 9.6E-84 | 1.0695939 | 0.272 | 0.099 | 3.12E-79 | Chondrocytes | selenot2 |
| cdkn1a3 | 1.9E-83 | 1.2486374 | 0.298 | 0.117 | 6.05E-79 | Chondrocytes | cdkn1a |
| acsl4b1 | 2.3E-83 | 1.3259364 | 0.313 | 0.127 | 7.62E-79 | Chondrocytes | acsl4b |
| dlc11 | 2.5E-83 | 1.4792677 | 0.153 | 0.041 | 8.16E-79 | Chondrocytes | dlc1 |
| bet11 | 4.3E-83 | 1.2428174 | 0.26 | 0.094 | 1.41E-78 | Chondrocytes | bet1 |
| snx18a | 4.8E-83 | 1.7250471 | 0.117 | 0.026 | 1.58E-78 | Chondrocytes | snx18a |
| ppiab1 | 5.7E-83 | 0.5178096 | 0.974 | 0.894 | 1.85E-78 | Chondrocytes | ppiab |
| atp5pd2 | 5.8E-83 | 0.7326164 | 0.647 | 0.36 | 1.9E-78 | Chondrocytes | atp5pd |
| fam20a1 | 1.9E-82 | 2.1793462 | 0.102 | 0.021 | 6.24E-78 | Chondrocytes | fam20a |
| col16a13 | 2.1E-82 | 0.7818516 | 0.712 | 0.448 | 6.96E-78 | Chondrocytes | col16a1 |
| tnfaip8l31 | 2.2E-82 | 1.5583794 | 0.158 | 0.043 | 7.16E-78 | Chondrocytes | tnfaip8l3 |
| optn | 8E-82 | 1.777324 | 0.205 | 0.066 | 2.61E-77 | Chondrocytes | optn |
| nog21 | 7.3E-81 | 1.474999 | 0.229 | 0.08 | 2.38E-76 | Chondrocytes | nog2 |
| clint1a1 | 1.1E-80 | 0.7780532 | 0.439 | 0.201 | 3.54E-76 | Chondrocytes | clint1a |
| ndfip11 | 1.2E-80 | 0.7275053 | 0.647 | 0.367 | 3.94E-76 | Chondrocytes | ndfip1 |
| niban2a2 | 2.5E-80 | 1.4734285 | 0.168 | 0.048 | 8.23E-76 | Chondrocytes | niban2a |
| si:ch211-242b18.1 | 5E-80 | 1.1643363 | 0.238 | 0.083 | 1.63E-75 | Chondrocytes | si:ch211-242b18.1 |
| phldb1b2 | 9.9E-80 | 0.9970096 | 0.242 | 0.084 | 3.23E-75 | Chondrocytes | phldb1b |
| slc17a9b | 2.6E-79 | 1.4387045 | 0.158 | 0.044 | 8.43E-75 | Chondrocytes | slc17a9b |

| | | | | | | | |
|---|---|---|---|---|---|---|---|
| p3h33 | 3.2E-79 | 0.9273056 | 0.32 | 0.13 | 1.04E-74 | Chondrocytes | p3h3 |
| shroom42 | 3.6E-79 | 0.9351393 | 0.318 | 0.128 | 1.17E-74 | Chondrocytes | shroom4 |
| hmgb3b2 | 7.3E-79 | 1.0726808 | 0.288 | 0.113 | 2.37E-74 | Chondrocytes | hmgb3b |
| htra1b3 | 8.4E-79 | 0.368638 | 0.712 | 0.403 | 2.73E-74 | Chondrocytes | htra1b |
| sdc23 | 1.9E-78 | 0.7508983 | 0.82 | 0.581 | 6.18E-74 | Chondrocytes | sdc2 |
| ABR | 5.7E-78 | 1.635052 | 0.137 | 0.035 | 1.86E-73 | Chondrocytes | ABR |
| pgk11 | 6.7E-78 | 1.074157 | 0.417 | 0.202 | 2.16E-73 | Chondrocytes | pgk1 |
| cox5ab2 | 7.6E-78 | 0.850614 | 0.446 | 0.214 | 2.47E-73 | Chondrocytes | cox5ab |
| nme32 | 1.1E-77 | 0.7435628 | 0.712 | 0.442 | 3.69E-73 | Chondrocytes | nme3 |
| bnip3lb | 1.5E-77 | 0.8624805 | 0.373 | 0.163 | 4.95E-73 | Chondrocytes | bnip3lb |
| nocta2 | 2.3E-77 | 1.3018767 | 0.276 | 0.109 | 7.45E-73 | Chondrocytes | nocta |
| MDFI3 | 3.1E-77 | 1.1277919 | 0.457 | 0.232 | 1.02E-72 | Chondrocytes | MDFI |
| SYDE1 | 2.3E-76 | 1.1821252 | 0.195 | 0.062 | 7.46E-72 | Chondrocytes | SYDE1 |
| ppfibp1b2 | 2.6E-76 | 0.9692117 | 0.329 | 0.138 | 8.53E-72 | Chondrocytes | ppfibp1b |
| med30 | 5.1E-76 | 0.9347027 | 0.29 | 0.113 | 1.65E-71 | Chondrocytes | med30 |
| hivep3a3 | 7.4E-76 | 1.1104543 | 0.272 | 0.105 | 2.41E-71 | Chondrocytes | hivep3a |
| c1qtnf6a2 | 7.4E-76 | 1.4217129 | 0.184 | 0.058 | 2.42E-71 | Chondrocytes | c1qtnf6a |
| clta1 | 1E-75 | 0.8967008 | 0.336 | 0.141 | 3.26E-71 | Chondrocytes | clta |
| hs3st3b1b3 | 1.3E-75 | 0.955841 | 0.308 | 0.124 | 4.32E-71 | Chondrocytes | hs3st3b1b |
| nedd93 | 1.4E-75 | 1.3430375 | 0.206 | 0.069 | 4.4E-71 | Chondrocytes | nedd9 |
| cops92 | 2.7E-75 | 1.0603883 | 0.341 | 0.148 | 8.85E-71 | Chondrocytes | cops9 |
| btbd6a2 | 4.2E-75 | 1.5186247 | 0.217 | 0.076 | 1.36E-70 | Chondrocytes | btbd6a |
| ppib3 | 8E-75 | 0.4853997 | 0.981 | 0.915 | 2.59E-70 | Chondrocytes | ppib |
| ctsz1 | 2.2E-74 | 0.814234 | 0.362 | 0.157 | 7.22E-70 | Chondrocytes | ctsz |
| plp21 | 2.3E-74 | 0.9870652 | 0.285 | 0.113 | 7.42E-70 | Chondrocytes | plp2 |
| golim4a | 2.4E-74 | 0.910557 | 0.289 | 0.113 | 7.71E-70 | Chondrocytes | golim4a |
| cope2 | 2.5E-74 | 0.8632429 | 0.488 | 0.25 | 7.98E-70 | Chondrocytes | cope |
| appb1 | 2.7E-74 | 0.6500895 | 0.738 | 0.46 | 8.76E-70 | Chondrocytes | appb |
| gyg1a1 | 7.2E-74 | 1.4399414 | 0.382 | 0.177 | 2.35E-69 | Chondrocytes | gyg1a |
| sec22ba2 | 1.2E-73 | 0.8854046 | 0.287 | 0.113 | 3.74E-69 | Chondrocytes | sec22ba |
| eef1g2 | 1.2E-73 | 0.4795331 | 0.98 | 0.951 | 3.96E-69 | Chondrocytes | eef1g |
| atp6ap21 | 2E-73 | 0.9508313 | 0.257 | 0.096 | 6.59E-69 | Chondrocytes | atp6ap2 |
| adamtsl33 | 2.8E-73 | 1.1002989 | 0.199 | 0.066 | 9.15E-69 | Chondrocytes | adamtsl3 |
| chmp4ba | 3.4E-73 | 0.9001243 | 0.423 | 0.206 | 1.1E-68 | Chondrocytes | chmp4ba |
| ezrb1 | 7.1E-73 | 1.1151244 | 0.388 | 0.185 | 2.31E-68 | Chondrocytes | ezrb |
| p4hb2 | 7.6E-73 | 0.6986856 | 0.805 | 0.567 | 2.47E-68 | Chondrocytes | p4hb |
| mcfd21 | 1.1E-72 | 1.0328599 | 0.297 | 0.122 | 3.42E-68 | Chondrocytes | mcfd2 |
| alg2 | 2E-72 | 1.709086 | 0.114 | 0.027 | 6.37E-68 | Chondrocytes | alg2 |
| nr4a2b | 2.3E-72 | 1.4445198 | 0.134 | 0.036 | 7.63E-68 | Chondrocytes | nr4a2b |
| h2az2b1 | 3E-72 | 0.5734484 | 0.672 | 0.389 | 9.9E-68 | Chondrocytes | h2az2b |
| nedd4l1 | 3.8E-72 | 1.0414162 | 0.254 | 0.096 | 1.22E-67 | Chondrocytes | nedd4l |
| ptmaa1 | 3.8E-72 | 0.7162905 | 0.946 | 0.833 | 1.24E-67 | Chondrocytes | ptmaa |
| trappc32 | 3.9E-72 | 1.1711192 | 0.262 | 0.102 | 1.27E-67 | Chondrocytes | trappc3 |
| golim4a.1 | 5.8E-72 | 1.0536848 | 0.248 | 0.093 | 1.88E-67 | Chondrocytes | golim4a.1 |
| adgrg6 | 7.9E-72 | 1.4825511 | 0.155 | 0.046 | 2.56E-67 | Chondrocytes | adgrg6 |
| cpxm1b1 | 1.7E-71 | 1.0918839 | 0.303 | 0.128 | 5.59E-67 | Chondrocytes | cpxm1b |
| ell21 | 8.4E-71 | 0.9688407 | 0.314 | 0.134 | 2.72E-66 | Chondrocytes | ell2 |
| col6a24 | 1E-70 | 0.508793 | 0.973 | 0.874 | 3.33E-66 | Chondrocytes | col6a2 |

| | | | | | | | |
|---|---|---|---|---|---|---|---|
| slc3a2b2 | 1.8E-70 | 1.0492726 | 0.312 | 0.134 | 5.76E-66 | Chondrocytes | slc3a2b |
| palm2akap2.13 | 2.8E-70 | 1.1146596 | 0.224 | 0.081 | 9.02E-66 | Chondrocytes | palm2akap2.1 |
| tnfaip8l1 | 6.9E-70 | 1.3432494 | 0.122 | 0.031 | 2.23E-65 | Chondrocytes | tnfaip8l1 |
| trib32 | 9E-70 | 0.9550142 | 0.368 | 0.171 | 2.91E-65 | Chondrocytes | trib3 |
| prkcsh1 | 1.5E-69 | 0.8424161 | 0.31 | 0.131 | 4.96E-65 | Chondrocytes | prkcsh |
| atl32 | 4E-69 | 0.6889047 | 0.574 | 0.315 | 1.3E-64 | Chondrocytes | atl3 |
| chmp1b1 | 5.4E-69 | 0.8646963 | 0.283 | 0.114 | 1.76E-64 | Chondrocytes | chmp1b |
| tp53inp1 | 5.9E-69 | 1.1049968 | 0.177 | 0.056 | 1.92E-64 | Chondrocytes | tp53inp1 |
| atox13 | 7.3E-69 | 0.7636048 | 0.486 | 0.251 | 2.38E-64 | Chondrocytes | atox1 |
| dpm21 | 1.3E-68 | 0.8138512 | 0.317 | 0.134 | 4.19E-64 | Chondrocytes | dpm2 |
| esyt2a | 1.7E-68 | 0.6810281 | 0.368 | 0.164 | 5.38E-64 | Chondrocytes | esyt2a |
| cmtm7 | 2.7E-68 | 0.9278654 | 0.285 | 0.117 | 8.64E-64 | Chondrocytes | cmtm7 |
| lman12 | 3.4E-68 | 1.1002523 | 0.285 | 0.12 | 1.12E-63 | Chondrocytes | lman1 |
| hif1ab1 | 3.7E-68 | 0.6429107 | 0.629 | 0.368 | 1.19E-63 | Chondrocytes | hif1ab |
| git11 | 4.9E-68 | 1.3334281 | 0.186 | 0.063 | 1.59E-63 | Chondrocytes | git1 |
| prkcdb | 1E-67 | 1.5105676 | 0.118 | 0.03 | 3.35E-63 | Chondrocytes | prkcdb |
| akt2l | 1.1E-67 | 1.3706317 | 0.154 | 0.047 | 3.52E-63 | Chondrocytes | akt2l |
| prickle2b | 1.4E-67 | 1.3554612 | 0.116 | 0.029 | 4.49E-63 | Chondrocytes | prickle2b |
| cblb1 | 3.1E-67 | 0.9028602 | 0.244 | 0.093 | 1.02E-62 | Chondrocytes | cblb |
| arhgap21a2 | 4.4E-67 | 0.9985377 | 0.259 | 0.102 | 1.42E-62 | Chondrocytes | arhgap21a |
| pld1a1 | 1.1E-66 | 0.9108607 | 0.236 | 0.089 | 3.55E-62 | Chondrocytes | pld1a |
| rgcc2 | 2.4E-66 | 0.9129837 | 0.394 | 0.193 | 7.81E-62 | Chondrocytes | rgcc |
| pcbp21 | 3E-66 | 0.6953509 | 0.445 | 0.219 | 9.73E-62 | Chondrocytes | pcbp2 |
| golph3 | 5.2E-66 | 0.8652571 | 0.238 | 0.09 | 1.69E-61 | Chondrocytes | golph3 |
| elf2b | 7.7E-66 | 0.8408352 | 0.268 | 0.108 | 2.52E-61 | Chondrocytes | elf2b |
| micall1a | 7.9E-66 | 1.4732061 | 0.106 | 0.026 | 2.57E-61 | Chondrocytes | micall1a |
| nfatc2a | 2.3E-65 | 1.2514051 | 0.125 | 0.034 | 7.56E-61 | Chondrocytes | nfatc2a |
| ptp4a11 | 3.1E-65 | 0.763726 | 0.448 | 0.227 | 9.94E-61 | Chondrocytes | ptp4a1 |
| lrrc8aa1 | 3.2E-65 | 0.8891964 | 0.245 | 0.095 | 1.05E-60 | Chondrocytes | lrrc8aa |
| qrfpr4 | 3.3E-65 | 1.4737717 | 0.107 | 0.026 | 1.07E-60 | Chondrocytes | qrfpr4 |
| fam219aa | 3.9E-65 | 1.2307737 | 0.114 | 0.029 | 1.27E-60 | Chondrocytes | fam219aa |
| h3f3d3 | 4.1E-65 | 0.5830529 | 0.883 | 0.724 | 1.34E-60 | Chondrocytes | h3f3d |
| fli1a | 6.9E-65 | 1.2875482 | 0.152 | 0.046 | 2.23E-60 | Chondrocytes | fli1a |
| kars12 | 7.8E-65 | 0.8679991 | 0.349 | 0.161 | 2.54E-60 | Chondrocytes | kars1 |
| tep1 | 8.4E-65 | 1.229656 | 0.144 | 0.043 | 2.72E-60 | Chondrocytes | tep1 |
| c1qtnf125 | 8.4E-65 | 1.7419892 | 0.24 | 0.097 | 2.73E-60 | Chondrocytes | c1qtnf12 |
| lgals3a1 | 1.3E-64 | 0.7433124 | 0.503 | 0.271 | 4.19E-60 | Chondrocytes | lgals3a |
| arpc5b2 | 1.5E-64 | 1.0953937 | 0.223 | 0.084 | 4.95E-60 | Chondrocytes | arpc5b |
| arf4a.12 | 1.8E-64 | 1.015543 | 0.201 | 0.071 | 5.97E-60 | Chondrocytes | arf4a.1 |
| rnf244 | 1.8E-64 | 1.0967148 | 0.231 | 0.089 | 5.98E-60 | Chondrocytes | rnf24 |
| rgs12b3 | 2.9E-64 | 0.9884143 | 0.225 | 0.085 | 9.41E-60 | Chondrocytes | rgs12b |
| pef11 | 2.9E-64 | 0.7236609 | 0.34 | 0.152 | 9.55E-60 | Chondrocytes | pef1 |
| srpx22 | 5.1E-64 | 0.826674 | 0.469 | 0.248 | 1.65E-59 | Chondrocytes | srpx2 |
| atp5if1b2 | 6.3E-64 | 0.6424981 | 0.583 | 0.325 | 2.06E-59 | Chondrocytes | atp5if1b |
| tnfrsf11a3 | 1.5E-63 | 0.9556752 | 0.368 | 0.179 | 4.88E-59 | Chondrocytes | tnfrsf11a |
| cnpy22 | 1.6E-63 | 1.341755 | 0.142 | 0.042 | 5.26E-59 | Chondrocytes | cnpy2 |
| slc25a55a3 | 2.1E-63 | 0.8113406 | 0.391 | 0.192 | 6.9E-59 | Chondrocytes | slc25a55a |
| ssr43 | 3.6E-63 | 0.582994 | 0.83 | 0.602 | 1.19E-58 | Chondrocytes | ssr4 |

| | | | | | | | |
|---|---|---|---|---|---|---|---|
| fkbp92 | 4E-63 | 0.8047576 | 0.61 | 0.377 | 1.31E-58 | Chondrocytes | fkbp9 |
| aimp22 | 6.4E-62 | 0.8293476 | 0.354 | 0.167 | 2.08E-57 | Chondrocytes | aimp2 |
| six1a2 | 6.5E-62 | 0.8860136 | 0.291 | 0.126 | 2.13E-57 | Chondrocytes | six1a |
| sox4a3 | 6.9E-62 | 0.9595923 | 0.38 | 0.189 | 2.25E-57 | Chondrocytes | sox4a |
| slc39a72 | 9.9E-62 | 1.0994817 | 0.266 | 0.113 | 3.22E-57 | Chondrocytes | slc39a7 |
| h3f3b.13 | 1.1E-61 | 0.5935777 | 0.742 | 0.488 | 3.43E-57 | Chondrocytes | h3f3b.1 |
| hbp12 | 1.1E-61 | 0.8308677 | 0.263 | 0.108 | 3.63E-57 | Chondrocytes | hbp1 |
| zgc:1583432 | 3.1E-61 | 1.601658 | 0.56 | 0.373 | 1.01E-56 | Chondrocytes | zgc:158343 |
| spsb4b | 5.4E-61 | 1.4426068 | 0.111 | 0.029 | 1.74E-56 | Chondrocytes | spsb4b |
| mbnl11 | 1E-60 | 1.0976684 | 0.188 | 0.067 | 3.26E-56 | Chondrocytes | mbnl1 |
| mbnl2 | 1.2E-60 | 0.7038608 | 0.429 | 0.218 | 3.92E-56 | Chondrocytes | mbnl2 |
| fkbp142 | 1.7E-60 | 0.7720854 | 0.549 | 0.326 | 5.4E-56 | Chondrocytes | fkbp14 |
| pfn22 | 1.7E-60 | 0.6217706 | 0.762 | 0.532 | 5.4E-56 | Chondrocytes | pfn2 |
| fosl23 | 1.7E-60 | 0.6883609 | 0.66 | 0.422 | 5.65E-56 | Chondrocytes | fosl2 |
| arfgap3 | 3E-60 | 1.3745692 | 0.114 | 0.031 | 9.71E-56 | Chondrocytes | arfgap3 |
| fuca22 | 3.1E-60 | 0.7132548 | 0.318 | 0.141 | 1E-55 | Chondrocytes | fuca2 |
| copb21 | 3.3E-60 | 0.696638 | 0.487 | 0.264 | 1.07E-55 | Chondrocytes | copb2 |
| cst14a.21 | 9.7E-60 | 0.77756 | 0.585 | 0.357 | 3.14E-55 | Chondrocytes | cst14a.2 |
| fkbp10b3 | 2.1E-59 | 0.6866947 | 0.469 | 0.248 | 6.89E-55 | Chondrocytes | fkbp10b |
| STMP12 | 3.8E-59 | 0.6465271 | 0.476 | 0.251 | 1.24E-54 | Chondrocytes | STMP1 |
| letm22 | 4.4E-59 | 0.8059194 | 0.371 | 0.183 | 1.44E-54 | Chondrocytes | letm2 |
| rhoj | 1.3E-58 | 1.2221353 | 0.122 | 0.035 | 4.12E-54 | Chondrocytes | rhoj |
| klhl24b1 | 1.7E-58 | 1.0237807 | 0.22 | 0.086 | 5.68E-54 | Chondrocytes | klhl24b |
| prrg11 | 2E-58 | 0.8589146 | 0.395 | 0.202 | 6.62E-54 | Chondrocytes | prrg1 |
| yipf4 | 2.3E-58 | 0.6704854 | 0.366 | 0.175 | 7.45E-54 | Chondrocytes | yipf4 |
| osbpl91 | 2.8E-58 | 0.7287953 | 0.329 | 0.151 | 9.13E-54 | Chondrocytes | osbpl9 |
| tomm72 | 3.1E-58 | 0.6698144 | 0.386 | 0.189 | 1.02E-53 | Chondrocytes | tomm7 |
| ift201 | 5.6E-58 | 1.0910036 | 0.167 | 0.057 | 1.81E-53 | Chondrocytes | ift20 |
| lrrc8c | 7.7E-58 | 1.1341403 | 0.113 | 0.031 | 2.5E-53 | Chondrocytes | lrrc8c |
| idh23 | 8.5E-58 | 0.5936291 | 0.58 | 0.337 | 2.78E-53 | Chondrocytes | idh2 |
| sgms1 | 8.6E-58 | 1.0456511 | 0.162 | 0.054 | 2.8E-53 | Chondrocytes | sgms1 |
| lifrb2 | 1.1E-57 | 0.709201 | 0.39 | 0.194 | 3.62E-53 | Chondrocytes | lifrb |
| srp541 | 1.1E-57 | 0.6895504 | 0.357 | 0.17 | 3.74E-53 | Chondrocytes | srp54 |
| f11r.1 | 2.1E-57 | 0.9911692 | 0.196 | 0.073 | 6.75E-53 | Chondrocytes | f11r.1 |
| cx43.44 | 3.8E-57 | 0.8873077 | 0.411 | 0.217 | 1.24E-52 | Chondrocytes | cx43.4 |
| si:dkey-177p2.62 | 4E-57 | 0.862868 | 0.465 | 0.258 | 1.29E-52 | Chondrocytes | si:dkey-177p2.6 |
| tgfbr2b4 | 4E-57 | 0.5886087 | 0.31 | 0.139 | 1.32E-52 | Chondrocytes | tgfbr2b |
| mapk62 | 5.5E-57 | 0.8679937 | 0.216 | 0.084 | 1.8E-52 | Chondrocytes | mapk6 |
| gbp3 | 1.4E-56 | 1.1018883 | 0.151 | 0.049 | 4.63E-52 | Chondrocytes | gbp3 |
| zgc:559431 | 3.2E-56 | 0.9496611 | 0.19 | 0.07 | 1.04E-51 | Chondrocytes | zgc:55943 |
| serf23 | 4.3E-56 | 0.4174491 | 0.931 | 0.768 | 1.41E-51 | Chondrocytes | serf2 |
| fam177a1 | 6.7E-56 | 0.9541155 | 0.18 | 0.065 | 2.19E-51 | Chondrocytes | fam177a1 |
| atp6v1g12 | 8.6E-56 | 0.7588639 | 0.287 | 0.128 | 2.81E-51 | Chondrocytes | atp6v1g1 |
| il13ra12 | 2.3E-55 | 0.9364747 | 0.209 | 0.081 | 7.38E-51 | Chondrocytes | il13ra1 |
| zcchc14 | 3.5E-55 | 1.3186468 | 0.1 | 0.026 | 1.13E-50 | Chondrocytes | zcchc14 |
| CABZ01080375.11 | 3.6E-55 | 0.9881686 | 0.177 | 0.064 | 1.16E-50 | Chondrocytes | CABZ01080375.1 |
| arfgap21 | 6.4E-55 | 0.8065373 | 0.242 | 0.101 | 2.09E-50 | Chondrocytes | arfgap2 |
| slc35e11 | 1.1E-54 | 1.0207844 | 0.158 | 0.054 | 3.42E-50 | Chondrocytes | slc35e1 |

| | | | | | | | |
|---|---|---|---|---|---|---|---|
| kcnk5b | 1.1E-54 | 1.3978811 | 0.141 | 0.046 | 3.65E-50 | Chondrocytes | kcnk5b |
| si:ch211-160o17.21 | 2.1E-54 | 1.2032691 | 0.156 | 0.054 | 6.7E-50 | Chondrocytes | si:ch211-160o17.2 |
| tmed101 | 2.3E-54 | 0.5790626 | 0.569 | 0.331 | 7.38E-50 | Chondrocytes | tmed10 |
| rbm8a1 | 3.2E-54 | 0.5769894 | 0.586 | 0.346 | 1.03E-49 | Chondrocytes | rbm8a |
| vamp31 | 3.8E-54 | 0.5813452 | 0.459 | 0.244 | 1.25E-49 | Chondrocytes | vamp3 |
| elf1 | 5.3E-54 | 0.8076394 | 0.195 | 0.074 | 1.71E-49 | Chondrocytes | elf1 |
| ddost1 | 8.6E-54 | 0.7046828 | 0.417 | 0.219 | 2.81E-49 | Chondrocytes | ddost |
| eml21 | 9.3E-54 | 1.0266889 | 0.184 | 0.069 | 3.03E-49 | Chondrocytes | eml2 |
| plpp2b1 | 9.8E-54 | 0.8803121 | 0.274 | 0.123 | 3.2E-49 | Chondrocytes | plpp2b |
| kank1a4 | 1E-53 | 0.8019452 | 0.253 | 0.108 | 3.38E-49 | Chondrocytes | kank1a |
| mpc12 | 1.1E-53 | 0.8236563 | 0.331 | 0.161 | 3.66E-49 | Chondrocytes | mpc1 |
| nufip21 | 1.5E-53 | 0.7694485 | 0.306 | 0.144 | 4.91E-49 | Chondrocytes | nufip2 |
| tspan3a | 3.2E-53 | 0.5297874 | 0.337 | 0.158 | 1.03E-48 | Chondrocytes | tspan3a |
| CABZ01050249.1 | 4.6E-53 | 1.3526746 | 0.103 | 0.028 | 1.49E-48 | Chondrocytes | CABZ01050249.1 |
| si:ch211-165e15.12 | 5.9E-53 | 0.921832 | 0.175 | 0.064 | 1.91E-48 | Chondrocytes | si:ch211-165e15.1 |
| iqgap11 | 6.4E-53 | 0.6022941 | 0.423 | 0.22 | 2.09E-48 | Chondrocytes | iqgap1 |
| odc12 | 9.6E-53 | 0.669927 | 0.271 | 0.12 | 3.12E-48 | Chondrocytes | odc1 |
| fam3a1 | 2.2E-52 | 0.6940669 | 0.297 | 0.137 | 7.3E-48 | Chondrocytes | fam3a |
| ikbip2 | 2.8E-52 | 0.8757893 | 0.259 | 0.115 | 8.95E-48 | Chondrocytes | ikbip |
| sgk2b1 | 3.2E-52 | 1.4036726 | 0.119 | 0.036 | 1.04E-47 | Chondrocytes | sgk2b |
| acaca | 3.8E-52 | 1.0843511 | 0.111 | 0.032 | 1.23E-47 | Chondrocytes | acaca |
| arhgef28a | 4.9E-52 | 1.2186366 | 0.101 | 0.028 | 1.6E-47 | Chondrocytes | arhgef28a |
| msmo1 | 8.2E-52 | 1.4291155 | 0.169 | 0.062 | 2.68E-47 | Chondrocytes | msmo1 |
| map1lc3a1 | 9.4E-52 | 0.570803 | 0.405 | 0.208 | 3.07E-47 | Chondrocytes | map1lc3a |
| ggcx3 | 1E-51 | 0.6363434 | 0.336 | 0.162 | 3.29E-47 | Chondrocytes | ggcx |
| bach1b2 | 1E-51 | 0.9549792 | 0.217 | 0.089 | 3.35E-47 | Chondrocytes | bach1b |
| fasn2 | 1.4E-51 | 0.8056031 | 0.235 | 0.099 | 4.47E-47 | Chondrocytes | fasn |
| maff3 | 2.5E-51 | 0.9494518 | 0.395 | 0.219 | 8.21E-47 | Chondrocytes | maff |
| ndel1a | 2.8E-51 | 1.2694508 | 0.103 | 0.029 | 9.12E-47 | Chondrocytes | ndel1a |
| cntfr1 | 3E-51 | 1.2852548 | 0.15 | 0.052 | 9.6E-47 | Chondrocytes | cntfr |
| atf4a4 | 3.7E-51 | 0.6077196 | 0.51 | 0.292 | 1.2E-46 | Chondrocytes | atf4a |
| gabarapa | 3.8E-51 | 0.5418372 | 0.557 | 0.325 | 1.23E-46 | Chondrocytes | gabarapa |
| pdap1b2 | 5.2E-51 | 0.9537906 | 0.22 | 0.092 | 1.68E-46 | Chondrocytes | pdap1b |
| aclya | 5.8E-51 | 1.186779 | 0.138 | 0.046 | 1.88E-46 | Chondrocytes | aclya |
| rab5ab | 7.6E-51 | 0.610754 | 0.459 | 0.249 | 2.49E-46 | Chondrocytes | rab5ab |
| phospho11 | 7.8E-51 | 2.2852852 | 0.122 | 0.039 | 2.53E-46 | Chondrocytes | phospho1 |
| aebp13 | 8.4E-51 | 1.057765 | 0.704 | 0.535 | 2.72E-46 | Chondrocytes | aebp1 |
| dynlrb1.11 | 1.3E-50 | 0.5841579 | 0.537 | 0.311 | 4.33E-46 | Chondrocytes | dynlrb1.1 |
| hip1 | 1.4E-50 | 0.9814089 | 0.17 | 0.063 | 4.65E-46 | Chondrocytes | hip1 |
| gorasp1a | 2E-50 | 1.2689893 | 0.108 | 0.031 | 6.4E-46 | Chondrocytes | gorasp1a |
| si:ch211-133n4.43 | 2.2E-50 | 0.4952303 | 0.676 | 0.441 | 7.06E-46 | Chondrocytes | si:ch211-133n4.4 |
| oser12 | 2.2E-50 | 0.9255598 | 0.205 | 0.083 | 7.23E-46 | Chondrocytes | oser1 |
| tomm20b2 | 2.4E-50 | 0.625966 | 0.562 | 0.337 | 7.94E-46 | Chondrocytes | tomm20b |
| atp5f1c2 | 2.4E-50 | 0.5400273 | 0.458 | 0.25 | 7.96E-46 | Chondrocytes | atp5f1c |
| ddx57 | 4E-50 | 0.3998594 | 0.808 | 0.591 | 1.3E-45 | Chondrocytes | ddx5 |
| atp5mc12 | 7.3E-50 | 0.5937567 | 0.516 | 0.305 | 2.36E-45 | Chondrocytes | atp5mc1 |
| selenof2 | 8E-50 | 0.5856839 | 0.698 | 0.46 | 2.6E-45 | Chondrocytes | selenof |
| hint12 | 1E-49 | 0.5521514 | 0.613 | 0.379 | 3.33E-45 | Chondrocytes | hint1 |

| | | | | | | | |
|---|---|---|---|---|---|---|---|
| h1-01 | 1.7E-49 | 0.7483734 | 0.347 | 0.177 | 5.5E-45 | Chondrocytes | h1-0 |
| map1lc3b1 | 1.9E-49 | 0.6023058 | 0.463 | 0.257 | 6.11E-45 | Chondrocytes | map1lc3b |
| abtb2a | 1.9E-49 | 1.3757391 | 0.103 | 0.029 | 6.34E-45 | Chondrocytes | abtb2a |
| mycbp2 | 2.4E-49 | 0.5376011 | 0.451 | 0.243 | 7.89E-45 | Chondrocytes | mycbp |
| tmem231 | 2.9E-49 | 0.9397626 | 0.132 | 0.043 | 9.45E-45 | Chondrocytes | tmem231 |
| lysmd31 | 4.3E-49 | 0.7817208 | 0.235 | 0.102 | 1.39E-44 | Chondrocytes | lysmd3 |
| gas2l1 | 4.4E-49 | 1.1422634 | 0.16 | 0.059 | 1.42E-44 | Chondrocytes | gas2l1 |
| cox7b2 | 4.7E-49 | 0.4877968 | 0.635 | 0.39 | 1.53E-44 | Chondrocytes | cox7b |
| pxk | 6.2E-49 | 0.9501649 | 0.238 | 0.106 | 2.02E-44 | Chondrocytes | pxk |
| ostc1 | 6.5E-49 | 0.6242165 | 0.496 | 0.285 | 2.12E-44 | Chondrocytes | ostc |
| cd151l2 | 7.5E-49 | 0.9048567 | 0.183 | 0.071 | 2.44E-44 | Chondrocytes | cd151l |
| arrdc3a3 | 8E-49 | 0.8542093 | 0.322 | 0.161 | 2.59E-44 | Chondrocytes | arrdc3a |
| taf131 | 8.2E-49 | 0.5582279 | 0.404 | 0.212 | 2.66E-44 | Chondrocytes | taf13 |
| rin23 | 9E-49 | 0.7929883 | 0.245 | 0.109 | 2.92E-44 | Chondrocytes | rin2 |
| nr4a3 | 9.1E-49 | 1.041853 | 0.14 | 0.048 | 2.96E-44 | Chondrocytes | nr4a3 |
| polb | 1.1E-48 | 1.0624664 | 0.155 | 0.056 | 3.42E-44 | Chondrocytes | polb |
| eif4ebp21 | 1.1E-48 | 0.5515134 | 0.402 | 0.211 | 3.65E-44 | Chondrocytes | eif4ebp2 |
| ube2al2 | 1.5E-48 | 0.6752433 | 0.376 | 0.194 | 4.93E-44 | Chondrocytes | ube2al |
| ammecr11 | 1.8E-48 | 0.6665393 | 0.3 | 0.142 | 5.93E-44 | Chondrocytes | ammecr1 |
| piezo2a.2 | 2.3E-48 | 1.2672134 | 0.125 | 0.04 | 7.55E-44 | Chondrocytes | piezo2a.2 |
| sigmar11 | 3.2E-48 | 0.7759239 | 0.22 | 0.093 | 1.04E-43 | Chondrocytes | sigmar1 |
| eif4a1a2 | 4.5E-48 | 0.7399276 | 0.563 | 0.36 | 1.47E-43 | Chondrocytes | eif4a1a |
| cpeb4b1 | 4.9E-48 | 0.7109572 | 0.327 | 0.163 | 1.59E-43 | Chondrocytes | cpeb4b |
| pik3cb | 5.7E-48 | 1.0690745 | 0.116 | 0.036 | 1.85E-43 | Chondrocytes | pik3cb |
| actn41 | 5.9E-48 | 0.8195398 | 0.24 | 0.106 | 1.91E-43 | Chondrocytes | actn4 |
| stat1a2 | 7.5E-48 | 0.5841392 | 0.555 | 0.333 | 2.43E-43 | Chondrocytes | stat1a |
| gabarapb2 | 9.5E-48 | 0.5146998 | 0.664 | 0.418 | 3.08E-43 | Chondrocytes | gabarapb |
| stxbp5a1 | 1.2E-47 | 1.2533119 | 0.118 | 0.037 | 3.94E-43 | Chondrocytes | stxbp5a |
| si:ch211-288g17.33 | 2.3E-47 | 0.5516134 | 0.626 | 0.398 | 7.47E-43 | Chondrocytes | si:ch211-288g17.3 |
| idh11 | 2.6E-47 | 0.8915688 | 0.214 | 0.091 | 8.33E-43 | Chondrocytes | idh1 |
| pdcd10a1 | 3.8E-47 | 0.7980814 | 0.217 | 0.092 | 1.22E-42 | Chondrocytes | pdcd10a |
| map1ab2 | 5.7E-47 | 1.1027007 | 0.361 | 0.192 | 1.86E-42 | Chondrocytes | map1ab |
| si:dkey-42i9.4 | 6E-47 | 0.7824972 | 0.222 | 0.095 | 1.96E-42 | Chondrocytes | si:dkey-42i9.4 |
| rnaseka2 | 6.6E-47 | 0.8033856 | 0.233 | 0.103 | 2.16E-42 | Chondrocytes | rnaseka |
| cnpy41 | 6.9E-47 | 0.6846503 | 0.263 | 0.121 | 2.26E-42 | Chondrocytes | cnpy4 |
| nfkbie | 8.6E-47 | 0.9823189 | 0.182 | 0.072 | 2.78E-42 | Chondrocytes | nfkbie |
| guk1a1 | 1.3E-46 | 0.6178283 | 0.282 | 0.133 | 4.23E-42 | Chondrocytes | guk1a |
| irak3 | 1.8E-46 | 1.811292 | 0.125 | 0.042 | 5.81E-42 | Chondrocytes | irak3 |
| os91 | 2E-46 | 0.5231427 | 0.398 | 0.21 | 6.54E-42 | Chondrocytes | os9 |
| sel1l | 2.3E-46 | 0.8309473 | 0.16 | 0.06 | 7.62E-42 | Chondrocytes | sel1l |
| si:ch73-269m14.2 | 2.4E-46 | 0.5433361 | 0.416 | 0.223 | 7.9E-42 | Chondrocytes | si:ch73-269m14.2 |
| cycsb1 | 2.7E-46 | 0.3556384 | 0.601 | 0.368 | 8.75E-42 | Chondrocytes | cycsb |
| sumo1 | 3E-46 | 0.5811174 | 0.318 | 0.156 | 9.72E-42 | Chondrocytes | sumo1 |
| smox1 | 3E-46 | 1.5700815 | 0.189 | 0.078 | 9.73E-42 | Chondrocytes | smox |
| zfand5b2 | 4.6E-46 | 0.7683002 | 0.452 | 0.267 | 1.49E-41 | Chondrocytes | zfand5b |
| inhbaa2 | 5.1E-46 | 1.4597216 | 0.423 | 0.268 | 1.65E-41 | Chondrocytes | inhbaa |
| CABZ01058650.12 | 6.3E-46 | 0.9616823 | 0.177 | 0.07 | 2.05E-41 | Chondrocytes | CABZ01058650.1 |
| ccdc6b2 | 7.5E-46 | 0.9735367 | 0.189 | 0.077 | 2.44E-41 | Chondrocytes | ccdc6b |

| | | | | | | | |
|---|---|---|---|---|---|---|---|
| ppt12 | 9.2E-46 | 0.5161184 | 0.562 | 0.338 | 2.99E-41 | Chondrocytes | ppt1 |
| tmem165 | 9.7E-46 | 0.8010507 | 0.171 | 0.066 | 3.16E-41 | Chondrocytes | tmem165 |
| irf2bpl3 | 1.2E-45 | 0.5682956 | 0.372 | 0.194 | 3.9E-41 | Chondrocytes | irf2bpl |
| tapbp.1 | 1.3E-45 | 0.6280681 | 0.224 | 0.097 | 4.26E-41 | Chondrocytes | tapbp.1 |
| bmp62 | 2.2E-45 | 1.0540435 | 0.154 | 0.058 | 7.18E-41 | Chondrocytes | bmp6 |
| trappc6bl | 2.4E-45 | 1.3621573 | 0.104 | 0.032 | 7.67E-41 | Chondrocytes | trappc6bl |
| atf4b4 | 3E-45 | 0.6427572 | 0.789 | 0.63 | 9.62E-41 | Chondrocytes | atf4b |
| sdhb1 | 3.5E-45 | 0.5397282 | 0.34 | 0.172 | 1.12E-40 | Chondrocytes | sdhb |
| dad12 | 3.8E-45 | 0.5092457 | 0.615 | 0.393 | 1.23E-40 | Chondrocytes | dad1 |
| hagh | 4E-45 | 0.9031657 | 0.158 | 0.059 | 1.3E-40 | Chondrocytes | hagh |
| rcan1a3 | 4.2E-45 | 0.724267 | 0.406 | 0.225 | 1.37E-40 | Chondrocytes | rcan1a |
| selenok1 | 4.6E-45 | 0.6169297 | 0.344 | 0.176 | 1.48E-40 | Chondrocytes | selenok |
| hdlbpa4 | 4.9E-45 | 0.4109652 | 0.864 | 0.67 | 1.61E-40 | Chondrocytes | hdlbpa |
| kifc31 | 5.9E-45 | 0.9128027 | 0.166 | 0.064 | 1.92E-40 | Chondrocytes | kifc3 |
| pprc13 | 7.4E-45 | 0.977363 | 0.243 | 0.113 | 2.42E-40 | Chondrocytes | pprc1 |
| mpped2a | 1.1E-44 | 1.3196123 | 0.112 | 0.036 | 3.46E-40 | Chondrocytes | mpped2a |
| txndc51 | 1.1E-44 | 0.6623049 | 0.375 | 0.202 | 3.47E-40 | Chondrocytes | txndc5 |
| srprb2 | 1.6E-44 | 0.6370971 | 0.343 | 0.177 | 5.08E-40 | Chondrocytes | srprb |
| slc35b2 | 2.5E-44 | 0.9736113 | 0.13 | 0.045 | 8.12E-40 | Chondrocytes | slc35b2 |
| hivep3b | 3.7E-44 | 1.113037 | 0.143 | 0.052 | 1.19E-39 | Chondrocytes | hivep3b |
| oaz1a3 | 4E-44 | 0.6368654 | 0.66 | 0.446 | 1.32E-39 | Chondrocytes | oaz1a |
| etnk14 | 4.4E-44 | 0.6620638 | 0.352 | 0.184 | 1.42E-39 | Chondrocytes | etnk1 |
| tuba8l23 | 4.6E-44 | 0.6148013 | 0.389 | 0.213 | 1.5E-39 | Chondrocytes | tuba8l2 |
| si:dkey-28b4.71 | 8.3E-44 | 0.5671604 | 0.368 | 0.195 | 2.7E-39 | Chondrocytes | si:dkey-28b4.7 |
| C8H5orf51 | 9.2E-44 | 0.7104374 | 0.247 | 0.114 | 2.99E-39 | Chondrocytes | C8H5orf51 |
| si:dkey-6e2.21 | 9.6E-44 | 1.1534056 | 0.202 | 0.088 | 3.14E-39 | Chondrocytes | si:dkey-6e2.2 |
| h3f3b.1.1 | 1.1E-43 | 0.5592561 | 0.331 | 0.167 | 3.57E-39 | Chondrocytes | h3f3b.1.1 |
| aars12 | 1.9E-43 | 0.6898104 | 0.277 | 0.134 | 6.31E-39 | Chondrocytes | aars1 |
| supt3h | 2.2E-43 | 0.8757853 | 0.151 | 0.057 | 7.08E-39 | Chondrocytes | supt3h |
| akirin11 | 2.2E-43 | 0.7457466 | 0.216 | 0.094 | 7.16E-39 | Chondrocytes | akirin1 |
| nck1b2 | 2.2E-43 | 0.8354277 | 0.214 | 0.094 | 7.17E-39 | Chondrocytes | nck1b |
| zgc:1629442 | 2.5E-43 | 0.5736906 | 0.488 | 0.288 | 8.14E-39 | Chondrocytes | zgc:162944 |
| gpx4b1 | 2.7E-43 | 0.4122055 | 0.507 | 0.29 | 8.85E-39 | Chondrocytes | gpx4b |
| pdap1a2 | 2.8E-43 | 0.4122119 | 0.4 | 0.214 | 9.04E-39 | Chondrocytes | pdap1a |
| grnb | 3.3E-43 | 0.9368399 | 0.133 | 0.047 | 1.07E-38 | Chondrocytes | grnb |
| rab2a1 | 3.8E-43 | 0.4526439 | 0.734 | 0.516 | 1.24E-38 | Chondrocytes | rab2a |
| eef1a1l13 | 5.4E-43 | 0.2297104 | 1 | 0.996 | 1.74E-38 | Chondrocytes | eef1a1l1 |
| zfand5a3 | 6.2E-43 | 0.7424089 | 0.346 | 0.184 | 2.03E-38 | Chondrocytes | zfand5a |
| acbd31 | 8.4E-43 | 0.6100602 | 0.275 | 0.133 | 2.72E-38 | Chondrocytes | acbd3 |
| myca4 | 9.7E-43 | 0.8445187 | 0.303 | 0.154 | 3.15E-38 | Chondrocytes | myca |
| egr2b | 1E-42 | 1.6039593 | 0.124 | 0.043 | 3.39E-38 | Chondrocytes | egr2b |
| eef1da4 | 1.1E-42 | 0.4873663 | 0.948 | 0.873 | 3.66E-38 | Chondrocytes | eef1da |
| ndufb91 | 1.3E-42 | 0.5361284 | 0.507 | 0.304 | 4.34E-38 | Chondrocytes | ndufb9 |
| eif4eb1 | 1.4E-42 | 0.7128754 | 0.226 | 0.102 | 4.62E-38 | Chondrocytes | eif4eb |
| mef2d1 | 1.6E-42 | 0.5765846 | 0.349 | 0.183 | 5.06E-38 | Chondrocytes | mef2d |
| klf9 | 1.9E-42 | 0.8061955 | 0.158 | 0.061 | 6.06E-38 | Chondrocytes | klf9 |
| zfand6 | 1.9E-42 | 1.0227706 | 0.13 | 0.046 | 6.25E-38 | Chondrocytes | zfand6 |
| CYTL1 | 3E-42 | 1.2147729 | 0.103 | 0.032 | 9.77E-38 | Chondrocytes | CYTL1 |

| | | | | | | | |
|---|---|---|---|---|---|---|---|
| ctnnb12 | 3.1E-42 | 0.4714451 | 0.719 | 0.499 | 1E-37 | Chondrocytes | ctnnb1 |
| tfg1 | 4.3E-42 | 0.6030976 | 0.349 | 0.186 | 1.38E-37 | Chondrocytes | tfg |
| BX927308.11 | 4.7E-42 | 0.8983797 | 0.158 | 0.061 | 1.53E-37 | Chondrocytes | BX927308.1 |
| bet1l | 6E-42 | 0.9792966 | 0.137 | 0.05 | 1.94E-37 | Chondrocytes | bet1l |
| arl6 | 6.2E-42 | 1.0709212 | 0.13 | 0.046 | 2.03E-37 | Chondrocytes | arl6 |
| scfd12 | 6.7E-42 | 0.738872 | 0.224 | 0.102 | 2.19E-37 | Chondrocytes | scfd1 |
| atp6v0ca1 | 8.9E-42 | 0.5176202 | 0.55 | 0.333 | 2.88E-37 | Chondrocytes | atp6v0ca |
| col5a2b1 | 9E-42 | 0.2073628 | 0.17 | 0.068 | 2.94E-37 | Chondrocytes | col5a2b |
| dgkza1 | 1.2E-41 | 0.7214149 | 0.147 | 0.055 | 3.78E-37 | Chondrocytes | dgkza |
| elovl6 | 1.2E-41 | 1.6666766 | 0.107 | 0.035 | 4.04E-37 | Chondrocytes | elovl6 |
| decr2 | 1.3E-41 | 1.1636558 | 0.102 | 0.032 | 4.18E-37 | Chondrocytes | decr2 |
| samd11 | 1.3E-41 | 1.263911 | 0.109 | 0.036 | 4.23E-37 | Chondrocytes | samd11 |
| snrnp272 | 1.8E-41 | 0.6120125 | 0.275 | 0.134 | 5.86E-37 | Chondrocytes | snrnp27 |
| gmppb1 | 2E-41 | 0.6157632 | 0.251 | 0.118 | 6.41E-37 | Chondrocytes | gmppb |
| kctd5a3 | 2.4E-41 | 0.4856938 | 0.522 | 0.314 | 7.66E-37 | Chondrocytes | kctd5a |
| bcl6ab1 | 2.6E-41 | 0.947334 | 0.152 | 0.059 | 8.46E-37 | Chondrocytes | bcl6ab |
| sh3pxd2b | 3E-41 | 0.5512778 | 0.471 | 0.277 | 9.9E-37 | Chondrocytes | sh3pxd2b |
| moxd12 | 4E-41 | 2.1785982 | 0.135 | 0.051 | 1.31E-36 | Chondrocytes | moxd1 |
| fhod12 | 4.3E-41 | 0.8412837 | 0.186 | 0.079 | 1.41E-36 | Chondrocytes | fhod1 |
| slc30a72 | 4.6E-41 | 0.8326061 | 0.233 | 0.109 | 1.5E-36 | Chondrocytes | slc30a7 |
| trak1a | 6.8E-41 | 0.8420626 | 0.15 | 0.058 | 2.22E-36 | Chondrocytes | trak1a |
| eif3ba2 | 8.7E-41 | 0.4703727 | 0.638 | 0.423 | 2.83E-36 | Chondrocytes | eif3ba |
| hk12 | 9.6E-41 | 0.6418697 | 0.245 | 0.115 | 3.13E-36 | Chondrocytes | hk1 |
| si:ch211-178n15.12 | 1.2E-40 | 0.9380853 | 0.309 | 0.165 | 3.82E-36 | Chondrocytes | si:ch211-178n15.1 |
| ak31 | 1.2E-40 | 0.6552338 | 0.227 | 0.104 | 3.97E-36 | Chondrocytes | ak3 |
| rargb | 1.2E-40 | 1.094828 | 0.12 | 0.042 | 4E-36 | Chondrocytes | rargb |
| fam162a2 | 1.4E-40 | 0.4845365 | 0.418 | 0.234 | 4.47E-36 | Chondrocytes | fam162a |
| hmgn21 | 1.5E-40 | 0.309941 | 0.775 | 0.56 | 4.88E-36 | Chondrocytes | hmgn2 |
| txndc121 | 1.6E-40 | 0.844857 | 0.16 | 0.064 | 5.33E-36 | Chondrocytes | txndc12 |
| tubb2b3 | 1.8E-40 | 0.5604198 | 0.53 | 0.336 | 5.71E-36 | Chondrocytes | tubb2b |
| tspan9b1 | 1.9E-40 | 0.6612116 | 0.267 | 0.13 | 6.28E-36 | Chondrocytes | tspan9b |
| bmp1a4 | 2.1E-40 | 0.5129467 | 0.343 | 0.183 | 6.81E-36 | Chondrocytes | bmp1a |
| vasnb3 | 2.9E-40 | 0.9305059 | 0.277 | 0.141 | 9.48E-36 | Chondrocytes | vasnb |
| tspan71 | 3.2E-40 | 0.5321765 | 0.385 | 0.211 | 1.03E-35 | Chondrocytes | tspan7 |
| si:ch211-195m9.3 | 3.3E-40 | 1.5251299 | 0.12 | 0.043 | 1.07E-35 | Chondrocytes | si:ch211-195m9.3 |
| LEPROTL1 | 3.4E-40 | 0.6186663 | 0.234 | 0.109 | 1.09E-35 | Chondrocytes | LEPROTL1 |
| pdk2a1 | 4E-40 | 0.8861635 | 0.187 | 0.08 | 1.3E-35 | Chondrocytes | pdk2a |
| ssr23 | 4E-40 | 0.4062063 | 0.872 | 0.686 | 1.31E-35 | Chondrocytes | ssr2 |
| dnajc13 | 5.2E-40 | 0.4387171 | 0.479 | 0.278 | 1.69E-35 | Chondrocytes | dnajc1 |
| ier3ip12 | 7.9E-40 | 0.5194319 | 0.411 | 0.231 | 2.56E-35 | Chondrocytes | ier3ip1 |
| arhgef402 | 8.7E-40 | 0.5191531 | 0.352 | 0.187 | 2.84E-35 | Chondrocytes | arhgef40 |
| limch1b3 | 1E-39 | 0.6880995 | 0.2 | 0.088 | 3.31E-35 | Chondrocytes | limch1b |
| cox8a2 | 1.2E-39 | 0.4327992 | 0.767 | 0.557 | 3.87E-35 | Chondrocytes | cox8a |
| igfbp32 | 1.3E-39 | 0.8789171 | 0.226 | 0.106 | 4.12E-35 | Chondrocytes | igfbp3 |
| slc7a3a | 1.3E-39 | 0.8856284 | 0.131 | 0.048 | 4.39E-35 | Chondrocytes | slc7a3a |
| rabac12 | 1.7E-39 | 0.5060558 | 0.605 | 0.391 | 5.58E-35 | Chondrocytes | rabac1 |
| gfpt11 | 1.7E-39 | 0.867348 | 0.145 | 0.056 | 5.64E-35 | Chondrocytes | gfpt1 |
| snx14 | 3.4E-39 | 0.7252909 | 0.15 | 0.059 | 1.1E-34 | Chondrocytes | snx14 |

| | | | | | | | |
|---|---|---|---|---|---|---|---|
| gmds1 | 3.7E-39 | 0.9062818 | 0.133 | 0.05 | 1.21E-34 | Chondrocytes | gmds |
| upf3b | 4.3E-39 | 0.4399475 | 0.396 | 0.219 | 1.39E-34 | Chondrocytes | upf3b |
| LO018340.1 | 4.7E-39 | 0.935752 | 0.131 | 0.048 | 1.54E-34 | Chondrocytes | LO018340.1 |
| bcl2l1 | 5.2E-39 | 0.5435579 | 0.3 | 0.153 | 1.7E-34 | Chondrocytes | bcl2l1 |
| cacnb2a | 5.2E-39 | 0.9397525 | 0.125 | 0.045 | 1.71E-34 | Chondrocytes | cacnb2a |
| rab6a1 | 5.4E-39 | 0.5347678 | 0.337 | 0.18 | 1.74E-34 | Chondrocytes | rab6a |
| gnpnat12 | 5.8E-39 | 0.8202154 | 0.195 | 0.087 | 1.87E-34 | Chondrocytes | gnpnat1 |
| zgc:153631 | 6.6E-39 | 1.2602277 | 0.103 | 0.034 | 2.15E-34 | Chondrocytes | zgc:153631 |
| anxa5b1 | 9.7E-39 | 0.4699015 | 0.602 | 0.397 | 3.15E-34 | Chondrocytes | anxa5b |
| si:ch73-46j18.55 | 1.4E-38 | 0.2553955 | 0.815 | 0.596 | 4.64E-34 | Chondrocytes | si:ch73-46j18.5 |
| chordc1a | 1.5E-38 | 0.7676869 | 0.165 | 0.068 | 4.86E-34 | Chondrocytes | chordc1a |
| serp13 | 1.6E-38 | 0.5034391 | 0.506 | 0.31 | 5.25E-34 | Chondrocytes | serp1 |
| rhoac1 | 2.6E-38 | 0.452472 | 0.486 | 0.29 | 8.32E-34 | Chondrocytes | rhoac |
| si:ch211-13k12.2 | 2.9E-38 | 0.8777478 | 0.135 | 0.051 | 9.3E-34 | Chondrocytes | si:ch211-13k12.2 |
| plekhg4 | 2.9E-38 | 1.0317844 | 0.1 | 0.032 | 9.46E-34 | Chondrocytes | plekhg4 |
| ufm12 | 3.3E-38 | 0.6382409 | 0.337 | 0.185 | 1.08E-33 | Chondrocytes | ufm1 |
| sars12 | 3.6E-38 | 0.5676923 | 0.282 | 0.143 | 1.18E-33 | Chondrocytes | sars1 |
| herpud1 | 3.9E-38 | 0.8638238 | 0.145 | 0.057 | 1.28E-33 | Chondrocytes | herpud1 |
| mef2cb1 | 4.1E-38 | 1.1140664 | 0.162 | 0.068 | 1.32E-33 | Chondrocytes | mef2cb |
| sumo2b1 | 5E-38 | 0.4345007 | 0.599 | 0.387 | 1.63E-33 | Chondrocytes | sumo2b |
| wbp2nl1 | 5.3E-38 | 0.4419688 | 0.487 | 0.285 | 1.73E-33 | Chondrocytes | wbp2nl |
| macroh2a21 | 6.2E-38 | 0.4185138 | 0.605 | 0.389 | 2.01E-33 | Chondrocytes | macroh2a2 |
| si:ch211-214j24.10 | 6.8E-38 | 0.478773 | 0.405 | 0.23 | 2.21E-33 | Chondrocytes | si:ch211-214j24.10 |
| rcn2 | 8.6E-38 | 0.6919278 | 0.181 | 0.078 | 2.78E-33 | Chondrocytes | rcn2 |
| bcl7bb2 | 1.6E-37 | 0.7596441 | 0.195 | 0.088 | 5.19E-33 | Chondrocytes | bcl7bb |
| rplp04 | 1.9E-37 | 0.2648497 | 0.994 | 0.982 | 6.16E-33 | Chondrocytes | rplp0 |
| selenot1a2 | 2E-37 | 0.4940699 | 0.315 | 0.165 | 6.38E-33 | Chondrocytes | selenot1a |
| sestd11 | 2.1E-37 | 0.9091834 | 0.217 | 0.103 | 6.96E-33 | Chondrocytes | sestd1 |
| sox11a3 | 2.3E-37 | 0.4994097 | 0.535 | 0.338 | 7.35E-33 | Chondrocytes | sox11a |
| srpra1 | 2.6E-37 | 0.4951251 | 0.383 | 0.215 | 8.45E-33 | Chondrocytes | srpra |
| ube2e33 | 2.6E-37 | 0.4118903 | 0.53 | 0.323 | 8.49E-33 | Chondrocytes | ube2e3 |
| stat32 | 2.6E-37 | 0.4161984 | 0.29 | 0.146 | 8.6E-33 | Chondrocytes | stat3 |
| glod51 | 3.3E-37 | 1.030661 | 0.141 | 0.056 | 1.09E-32 | Chondrocytes | glod5 |
| SGMS1 | 5.4E-37 | 0.9207167 | 0.148 | 0.06 | 1.75E-32 | Chondrocytes | SGMS1 |
| ywhaqa1 | 5.8E-37 | 0.36059 | 0.46 | 0.268 | 1.87E-32 | Chondrocytes | ywhaqa |
| mid1ip1b2 | 6E-37 | 0.7988086 | 0.235 | 0.116 | 1.96E-32 | Chondrocytes | mid1ip1b |
| mfap3l1 | 6.9E-37 | 0.6244729 | 0.195 | 0.087 | 2.25E-32 | Chondrocytes | mfap3l |
| tcf7l1a3 | 7E-37 | 0.549007 | 0.237 | 0.113 | 2.27E-32 | Chondrocytes | tcf7l1a |
| anos1b1 | 7E-37 | 0.7483152 | 0.172 | 0.073 | 2.27E-32 | Chondrocytes | anos1b |
| cnpy11 | 1.1E-36 | 0.7427911 | 0.179 | 0.078 | 3.71E-32 | Chondrocytes | cnpy1 |
| rps27l1 | 1.2E-36 | 0.7901087 | 0.464 | 0.283 | 3.75E-32 | Chondrocytes | rps27l |
| si:ch73-91k6.2 | 1.2E-36 | 0.6740361 | 0.216 | 0.102 | 3.9E-32 | Chondrocytes | si:ch73-91k6.2 |
| vps72a | 1.2E-36 | 1.0486238 | 0.1 | 0.033 | 3.99E-32 | Chondrocytes | vps72a |
| glb1l | 1.2E-36 | 1.037629 | 0.106 | 0.037 | 3.99E-32 | Chondrocytes | glb1l |
| capn2a1 | 1.2E-36 | 0.4062359 | 0.565 | 0.357 | 4E-32 | Chondrocytes | capn2a |
| sec61a13 | 1.4E-36 | 0.4091323 | 0.637 | 0.414 | 4.71E-32 | Chondrocytes | sec61a1 |
| sec24d | 1.9E-36 | 0.812844 | 0.132 | 0.051 | 6.16E-32 | Chondrocytes | sec24d |
| eif3ea4 | 2.1E-36 | 0.3980138 | 0.763 | 0.554 | 6.83E-32 | Chondrocytes | eif3ea |

| | | | | | | | |
|---|---|---|---|---|---|---|---|
| rca2.1 | 2.5E-36 | 0.3624727 | 0.474 | 0.28 | 8.1E-32 | Chondrocytes | rca2.1 |
| serinc11 | 3.1E-36 | 0.4369426 | 0.439 | 0.256 | 1.01E-31 | Chondrocytes | serinc1 |
| ugp2b1 | 6.6E-36 | 0.5135785 | 0.236 | 0.115 | 2.14E-31 | Chondrocytes | ugp2b |
| cox5aa2 | 7.2E-36 | 0.3700904 | 0.472 | 0.281 | 2.33E-31 | Chondrocytes | cox5aa |
| slc25a241 | 8.7E-36 | 0.5806693 | 0.376 | 0.215 | 2.84E-31 | Chondrocytes | slc25a24 |
| selenos1 | 1.1E-35 | 0.408957 | 0.302 | 0.157 | 3.61E-31 | Chondrocytes | selenos |
| ndufa51 | 1.1E-35 | 0.477084 | 0.369 | 0.205 | 3.69E-31 | Chondrocytes | ndufa5 |
| psap2 | 1.2E-35 | 0.3748282 | 0.555 | 0.344 | 3.82E-31 | Chondrocytes | psap |
| plekhb21 | 1.7E-35 | 0.5284354 | 0.214 | 0.1 | 5.38E-31 | Chondrocytes | plekhb2 |
| acsl4a2 | 1.7E-35 | 0.7819761 | 0.173 | 0.076 | 5.43E-31 | Chondrocytes | acsl4a |
| ndufs62 | 1.7E-35 | 0.5494393 | 0.27 | 0.138 | 5.58E-31 | Chondrocytes | ndufs6 |
| sec31a1 | 2.4E-35 | 0.7126596 | 0.219 | 0.106 | 7.65E-31 | Chondrocytes | sec31a |
| amotl2b1 | 2.5E-35 | 0.8196487 | 0.176 | 0.078 | 7.99E-31 | Chondrocytes | amotl2b |
| rnf144b | 2.9E-35 | 1.0304863 | 0.109 | 0.039 | 9.28E-31 | Chondrocytes | rnf144b |
| rnf11b | 3.6E-35 | 0.8935184 | 0.116 | 0.043 | 1.18E-30 | Chondrocytes | rnf11b |
| tmbim41 | 5E-35 | 0.4994501 | 0.233 | 0.113 | 1.64E-30 | Chondrocytes | tmbim4 |
| tcp11l2 | 5.1E-35 | 0.7157307 | 0.162 | 0.069 | 1.66E-30 | Chondrocytes | tcp11l2 |
| cct42 | 5.6E-35 | 0.3743349 | 0.505 | 0.307 | 1.83E-30 | Chondrocytes | cct4 |
| si:ch1073-429i10.31 | 5.7E-35 | 0.3748934 | 0.613 | 0.397 | 1.84E-30 | Chondrocytes | si:ch1073-429i10.3 |
| pi4k2b | 5.9E-35 | 0.9872034 | 0.106 | 0.037 | 1.92E-30 | Chondrocytes | pi4k2b |
| dnaja2b2 | 9.9E-35 | 0.5016973 | 0.225 | 0.108 | 3.23E-30 | Chondrocytes | dnaja2b |
| msna1 | 1.1E-34 | 0.3794786 | 0.88 | 0.747 | 3.46E-30 | Chondrocytes | msna |
| hdac9b2 | 1.2E-34 | 1.0490751 | 0.157 | 0.068 | 3.83E-30 | Chondrocytes | hdac9b |
| tdh3 | 1.9E-34 | 0.5071398 | 0.405 | 0.236 | 6.07E-30 | Chondrocytes | tdh |
| pak2a1 | 1.9E-34 | 0.465025 | 0.286 | 0.149 | 6.27E-30 | Chondrocytes | pak2a |
| rab5aa1 | 2E-34 | 0.3892277 | 0.44 | 0.257 | 6.49E-30 | Chondrocytes | rab5aa |
| ppp1caa1 | 2E-34 | 0.427138 | 0.262 | 0.132 | 6.6E-30 | Chondrocytes | ppp1caa |
| COLGALT1 | 2.5E-34 | 0.9393857 | 0.121 | 0.046 | 8.12E-30 | Chondrocytes | COLGALT1 |
| mgat1a | 2.8E-34 | 0.747151 | 0.134 | 0.053 | 8.99E-30 | Chondrocytes | mgat1a |
| cdon3 | 3.2E-34 | 0.4323244 | 0.335 | 0.184 | 1.05E-29 | Chondrocytes | cdon |
| cxcl141 | 3.6E-34 | 0.3242302 | 0.407 | 0.245 | 1.18E-29 | Chondrocytes | cxcl14 |
| errfi1a1 | 5.3E-34 | 0.9142186 | 0.127 | 0.049 | 1.72E-29 | Chondrocytes | errfi1a |
| itpka | 5.3E-34 | 0.8614461 | 0.114 | 0.042 | 1.72E-29 | Chondrocytes | itpka |
| mapk14b1 | 5.4E-34 | 0.5844935 | 0.186 | 0.084 | 1.76E-29 | Chondrocytes | mapk14b |
| rraga1 | 6.8E-34 | 0.839158 | 0.165 | 0.072 | 2.21E-29 | Chondrocytes | rraga |
| arpc31 | 7.1E-34 | 0.4729296 | 0.578 | 0.379 | 2.31E-29 | Chondrocytes | arpc3 |
| si:dkeyp-27c8.21 | 9E-34 | 1.028803 | 0.122 | 0.047 | 2.94E-29 | Chondrocytes | si:dkeyp-27c8.2 |
| chd4b1 | 1.2E-33 | 0.4369226 | 0.428 | 0.255 | 3.95E-29 | Chondrocytes | chd4b |
| tspan362 | 1.4E-33 | 0.9445621 | 0.285 | 0.159 | 4.71E-29 | Chondrocytes | tspan36 |
| tmem50a2 | 1.5E-33 | 0.3745823 | 0.402 | 0.231 | 4.83E-29 | Chondrocytes | tmem50a |
| ddx213 | 1.6E-33 | 0.4573529 | 0.532 | 0.338 | 5.05E-29 | Chondrocytes | ddx21 |
| id2a5 | 3.1E-33 | 0.4425558 | 0.682 | 0.498 | 1.02E-28 | Chondrocytes | id2a |
| rock11 | 3.2E-33 | 0.5521049 | 0.19 | 0.087 | 1.05E-28 | Chondrocytes | rock1 |
| si:dkey-112a7.42 | 5.3E-33 | 0.7508133 | 0.185 | 0.086 | 1.73E-28 | Chondrocytes | si:dkey-112a7.4 |
| ptk7a5 | 5.3E-33 | 0.3223666 | 0.506 | 0.309 | 1.73E-28 | Chondrocytes | ptk7a |
| cspg5a3 | 5.6E-33 | 2.2378821 | 0.162 | 0.075 | 1.83E-28 | Chondrocytes | cspg5a |
| lasp1 | 6.6E-33 | 0.4282666 | 0.666 | 0.466 | 2.14E-28 | Chondrocytes | lasp1 |
| ENSDARG000001018. | 6.6E-33 | 0.9586645 | 0.114 | 0.043 | 2.15E-28 | Chondrocytes | ENSDARG00000101 |

| | | | | | | | |
|---|---|---|---|---|---|---|---|
| CABZ01044281.1 | 6.8E-33 | 0.7976833 | 0.113 | 0.042 | 2.22E-28 | Chondrocytes | CABZ01044281.1 |
| clic41 | 1.2E-32 | 0.3048241 | 0.351 | 0.195 | 3.78E-28 | Chondrocytes | clic4 |
| fkbp21 | 1.2E-32 | 0.4100635 | 0.355 | 0.2 | 3.8E-28 | Chondrocytes | fkbp2 |
| ppid2 | 1.4E-32 | 0.6014002 | 0.174 | 0.079 | 4.44E-28 | Chondrocytes | ppid |
| mesd1 | 1.8E-32 | 0.494729 | 0.276 | 0.147 | 5.7E-28 | Chondrocytes | mesd |
| letm1 | 2E-32 | 0.7515135 | 0.135 | 0.055 | 6.46E-28 | Chondrocytes | letm1 |
| chid13 | 2.3E-32 | 0.67703 | 0.18 | 0.083 | 7.34E-28 | Chondrocytes | chid1 |
| sulf14 | 2.9E-32 | 0.6564763 | 0.266 | 0.144 | 9.4E-28 | Chondrocytes | sulf1 |
| mbd21 | 3E-32 | 0.6931607 | 0.176 | 0.08 | 9.6E-28 | Chondrocytes | mbd2 |
| pfkfb4b | 3.2E-32 | 0.8767582 | 0.125 | 0.05 | 1.04E-27 | Chondrocytes | pfkfb4b |
| si:ch211-195b13.13 | 3.7E-32 | 0.4686589 | 0.398 | 0.239 | 1.19E-27 | Chondrocytes | si:ch211-195b13.1 |
| tlcd2 | 4.6E-32 | 0.8703648 | 0.11 | 0.041 | 1.5E-27 | Chondrocytes | tlcd2 |
| vdac13 | 4.7E-32 | 0.3946587 | 0.509 | 0.318 | 1.54E-27 | Chondrocytes | vdac1 |
| rtca1 | 4.8E-32 | 0.4263327 | 0.331 | 0.184 | 1.58E-27 | Chondrocytes | rtca |
| hvcn11 | 8.2E-32 | 0.6350908 | 0.165 | 0.074 | 2.66E-27 | Chondrocytes | hvcn1 |
| pgm12 | 1.1E-31 | 0.4960183 | 0.204 | 0.097 | 3.65E-27 | Chondrocytes | pgm1 |
| lrpap11 | 1.1E-31 | 0.3941055 | 0.321 | 0.177 | 3.74E-27 | Chondrocytes | lrpap1 |
| atp5f1b2 | 1.2E-31 | 0.372976 | 0.752 | 0.553 | 3.87E-27 | Chondrocytes | atp5f1b |
| eloa | 1.4E-31 | 0.7691343 | 0.162 | 0.072 | 4.47E-27 | Chondrocytes | eloa |
| wdr62 | 1.5E-31 | 0.88184 | 0.113 | 0.043 | 4.77E-27 | Chondrocytes | wdr62 |
| sumo3a1 | 1.6E-31 | 0.345164 | 0.463 | 0.281 | 5.31E-27 | Chondrocytes | sumo3a |
| tfe3a2 | 1.8E-31 | 0.6641011 | 0.192 | 0.092 | 5.97E-27 | Chondrocytes | tfe3a |
| bloc1s63 | 2E-31 | 0.4002742 | 0.32 | 0.176 | 6.4E-27 | Chondrocytes | bloc1s6 |
| si:dkey-33c12.41 | 2.6E-31 | 0.3799128 | 0.533 | 0.336 | 8.49E-27 | Chondrocytes | si:dkey-33c12.4 |
| kdm5c1 | 2.7E-31 | 0.4091747 | 0.279 | 0.149 | 8.76E-27 | Chondrocytes | kdm5c |
| lmcd12 | 2.9E-31 | 0.6833416 | 0.183 | 0.086 | 9.4E-27 | Chondrocytes | lmcd1 |
| sdhaf4 | 3.7E-31 | 0.802022 | 0.131 | 0.054 | 1.19E-26 | Chondrocytes | sdhaf4 |
| gusb3 | 4E-31 | 0.4608789 | 0.245 | 0.127 | 1.31E-26 | Chondrocytes | gusb |
| fgfr1b4 | 4.1E-31 | 0.6037467 | 0.203 | 0.099 | 1.34E-26 | Chondrocytes | fgfr1b |
| eif2s32 | 4.2E-31 | 0.4074748 | 0.694 | 0.49 | 1.35E-26 | Chondrocytes | eif2s3 |
| cpvl2 | 4.8E-31 | 0.6630598 | 0.19 | 0.091 | 1.56E-26 | Chondrocytes | cpvl |
| sat1a.25 | 5E-31 | 0.6430273 | 0.62 | 0.453 | 1.61E-26 | Chondrocytes | sat1a.2 |
| brd1b1 | 5.8E-31 | 0.4417988 | 0.199 | 0.096 | 1.88E-26 | Chondrocytes | brd1b |
| si:ch211-147a11.3 | 6.2E-31 | 0.7538877 | 0.134 | 0.056 | 2.01E-26 | Chondrocytes | si:ch211-147a11.3 |
| GCA1 | 6.7E-31 | 0.4051271 | 0.469 | 0.291 | 2.18E-26 | Chondrocytes | GCA |
| snx9b | 7.9E-31 | 0.4432538 | 0.318 | 0.177 | 2.56E-26 | Chondrocytes | snx9b |
| creb1a2 | 9.4E-31 | 0.5011576 | 0.291 | 0.159 | 3.05E-26 | Chondrocytes | creb1a |
| nucks1b1 | 1E-30 | 0.314439 | 0.504 | 0.312 | 3.27E-26 | Chondrocytes | nucks1b |
| eif2s1b3 | 1.3E-30 | 0.3527385 | 0.583 | 0.382 | 4.09E-26 | Chondrocytes | eif2s1b |
| xpo1b2 | 1.6E-30 | 0.5372492 | 0.204 | 0.099 | 5.04E-26 | Chondrocytes | xpo1b |
| cers2a | 1.6E-30 | 0.9748776 | 0.105 | 0.04 | 5.07E-26 | Chondrocytes | cers2a |
| serpinh1a3 | 1.6E-30 | 0.409803 | 0.392 | 0.234 | 5.25E-26 | Chondrocytes | serpinh1a |
| pdcl1 | 1.7E-30 | 0.2560503 | 0.351 | 0.196 | 5.4E-26 | Chondrocytes | pdcl |
| scara33 | 2.3E-30 | 1.0202116 | 0.241 | 0.132 | 7.46E-26 | Chondrocytes | scara3 |
| golga7 | 2.4E-30 | 0.3494483 | 0.313 | 0.171 | 7.82E-26 | Chondrocytes | golga7 |
| fam120a2 | 2.7E-30 | 0.4283803 | 0.316 | 0.178 | 8.9E-26 | Chondrocytes | fam120a |
| COX5B.12 | 4.2E-30 | 0.2907221 | 0.744 | 0.536 | 1.36E-25 | Chondrocytes | COX5B.1 |
| zgc:85777 | 4.3E-30 | 1.0366079 | 0.104 | 0.04 | 1.4E-25 | Chondrocytes | zgc:85777 |

| | | | | | | | |
|---|---|---|---|---|---|---|---|
| TMEM191 | 4.4E-30 | 0.6758764 | 0.152 | 0.068 | 1.42E-25 | Chondrocytes | TMEM19 |
| ap1s1 | 4.4E-30 | 0.7312492 | 0.113 | 0.044 | 1.43E-25 | Chondrocytes | ap1s1 |
| hmgn72 | 4.8E-30 | 0.3507759 | 0.732 | 0.517 | 1.57E-25 | Chondrocytes | hmgn7 |
| zmiz1a2 | 5.1E-30 | 0.5271976 | 0.229 | 0.117 | 1.67E-25 | Chondrocytes | zmiz1a |
| unm-hu7910 | 5.3E-30 | 0.6755607 | 0.124 | 0.05 | 1.71E-25 | Chondrocytes | unm-hu7910 |
| pdcd10b1 | 5.4E-30 | 0.5479168 | 0.2 | 0.098 | 1.75E-25 | Chondrocytes | pdcd10b |
| usf1l1 | 6.1E-30 | 0.6540732 | 0.154 | 0.069 | 1.97E-25 | Chondrocytes | usf1l |
| fbrsl11 | 6.1E-30 | 0.7555858 | 0.126 | 0.052 | 1.98E-25 | Chondrocytes | fbrsl1 |
| arf11 | 6.5E-30 | 0.534144 | 0.272 | 0.147 | 2.11E-25 | Chondrocytes | arf1 |
| wipi13 | 6.7E-30 | 0.4933218 | 0.232 | 0.119 | 2.17E-25 | Chondrocytes | wipi1 |
| dnajb112 | 7.6E-30 | 0.6304041 | 0.205 | 0.103 | 2.48E-25 | Chondrocytes | dnajb11 |
| erlec11 | 8E-30 | 0.4660852 | 0.228 | 0.116 | 2.6E-25 | Chondrocytes | erlec1 |
| slc35e4 | 8.8E-30 | 0.7005664 | 0.166 | 0.076 | 2.85E-25 | Chondrocytes | slc35e4 |
| capza1a2 | 9.8E-30 | 0.3169655 | 0.444 | 0.267 | 3.17E-25 | Chondrocytes | capza1a |
| gabarapl21 | 1.1E-29 | 0.4214278 | 0.293 | 0.161 | 3.59E-25 | Chondrocytes | gabarapl2 |
| gulp1a3 | 1.1E-29 | 0.6164951 | 0.319 | 0.184 | 3.61E-25 | Chondrocytes | gulp1a |
| otud7b | 1.2E-29 | 0.7709861 | 0.108 | 0.042 | 3.87E-25 | Chondrocytes | otud7b |
| fam114a22 | 1.2E-29 | 0.5278272 | 0.22 | 0.112 | 4E-25 | Chondrocytes | fam114a2 |
| tm9sf31 | 1.6E-29 | 0.2946422 | 0.306 | 0.169 | 5.32E-25 | Chondrocytes | tm9sf3 |
| xbp12 | 1.6E-29 | 0.3987841 | 0.866 | 0.759 | 5.33E-25 | Chondrocytes | xbp1 |
| zgc:1531152 | 1.7E-29 | 0.5231719 | 0.172 | 0.08 | 5.57E-25 | Chondrocytes | zgc:153115 |
| fgfrl1a1 | 1.9E-29 | 0.6109798 | 0.101 | 0.038 | 6.16E-25 | Chondrocytes | fgfrl1a |
| nos1apa3 | 2.1E-29 | 0.7283207 | 0.208 | 0.106 | 6.72E-25 | Chondrocytes | nos1apa |
| yipf61 | 2.7E-29 | 0.6328669 | 0.165 | 0.076 | 8.75E-25 | Chondrocytes | yipf6 |
| PLEKHH32 | 2.9E-29 | 0.5194851 | 0.288 | 0.16 | 9.3E-25 | Chondrocytes | PLEKHH3 |
| laptm4a1 | 3.2E-29 | 0.3045305 | 0.486 | 0.302 | 1.04E-24 | Chondrocytes | laptm4a |
| si:dkey-17o15.21 | 3.6E-29 | 0.5840122 | 0.171 | 0.08 | 1.17E-24 | Chondrocytes | si:dkey-17o15.2 |
| smdt1b1 | 3.8E-29 | 0.3839988 | 0.408 | 0.247 | 1.23E-24 | Chondrocytes | smdt1b |
| tuba8l43 | 4.3E-29 | 0.361571 | 0.857 | 0.73 | 1.41E-24 | Chondrocytes | tuba8l4 |
| cox6c2 | 5.1E-29 | 0.3497737 | 0.607 | 0.41 | 1.66E-24 | Chondrocytes | cox6c |
| si:ch73-234b20.5 | 5.8E-29 | 0.6840043 | 0.119 | 0.048 | 1.9E-24 | Chondrocytes | si:ch73-234b20.5 |
| kctd9a1 | 6.3E-29 | 0.6529335 | 0.192 | 0.095 | 2.05E-24 | Chondrocytes | kctd9a |
| arl6ip11 | 8E-29 | 0.4255364 | 0.286 | 0.158 | 2.61E-24 | Chondrocytes | arl6ip1 |
| mxra8a5 | 8.1E-29 | 0.7154206 | 0.398 | 0.25 | 2.62E-24 | Chondrocytes | mxra8a |
| DAB25 | 8.6E-29 | 0.259571 | 0.563 | 0.373 | 2.79E-24 | Chondrocytes | DAB2 |
| smim131 | 1.1E-28 | 0.4641524 | 0.229 | 0.118 | 3.49E-24 | Chondrocytes | smim13 |
| fgfr1a4 | 1.1E-28 | 0.5155409 | 0.276 | 0.154 | 3.58E-24 | Chondrocytes | fgfr1a |
| ndufa13 | 1.2E-28 | 0.4439736 | 0.346 | 0.201 | 3.85E-24 | Chondrocytes | ndufa1 |
| bhlhe412 | 1.2E-28 | 0.4715592 | 0.24 | 0.126 | 3.94E-24 | Chondrocytes | bhlhe41 |
| aprt1 | 1.2E-28 | 0.8254039 | 0.102 | 0.039 | 3.97E-24 | Chondrocytes | aprt |
| selenou1a | 1.7E-28 | 0.5786681 | 0.148 | 0.066 | 5.61E-24 | Chondrocytes | selenou1a |
| sertad2b | 1.7E-28 | 0.9279016 | 0.112 | 0.045 | 5.66E-24 | Chondrocytes | sertad2b |
| manf1 | 1.9E-28 | 0.4106093 | 0.317 | 0.182 | 6.13E-24 | Chondrocytes | manf |
| ola13 | 1.9E-28 | 0.3689269 | 0.697 | 0.501 | 6.18E-24 | Chondrocytes | ola1 |
| per23 | 2E-28 | 0.4302165 | 0.205 | 0.102 | 6.42E-24 | Chondrocytes | per2 |
| sgsh | 2E-28 | 0.7364973 | 0.106 | 0.041 | 6.62E-24 | Chondrocytes | sgsh |
| tm9sf41 | 2.3E-28 | 0.6088837 | 0.158 | 0.073 | 7.45E-24 | Chondrocytes | tm9sf4 |
| prelid3b3 | 2.7E-28 | 0.5528582 | 0.514 | 0.353 | 8.82E-24 | Chondrocytes | prelid3b |

| | | | | | | | |
|---|---|---|---|---|---|---|---|
| tsg101b1 | 3E-28 | 0.4995933 | 0.183 | 0.088 | 9.68E-24 | Chondrocytes | tsg101b |
| sdha | 3.3E-28 | 0.4310258 | 0.272 | 0.149 | 1.06E-23 | Chondrocytes | sdha |
| mtmr11 | 3.3E-28 | 0.8154945 | 0.107 | 0.042 | 1.07E-23 | Chondrocytes | mtmr11 |
| efnb2a2 | 3.4E-28 | 0.4159315 | 0.269 | 0.147 | 1.11E-23 | Chondrocytes | efnb2a |
| dlx6a | 3.5E-28 | 0.9206194 | 0.12 | 0.05 | 1.13E-23 | Chondrocytes | dlx6a |
| fam83hb | 3.5E-28 | 0.6794341 | 0.141 | 0.062 | 1.15E-23 | Chondrocytes | fam83hb |
| atp5mf1 | 4E-28 | 0.3245298 | 0.566 | 0.366 | 1.31E-23 | Chondrocytes | atp5mf |
| plk34 | 5.1E-28 | 0.5688494 | 0.333 | 0.199 | 1.64E-23 | Chondrocytes | plk3 |
| cox7c1 | 5.4E-28 | 0.3335365 | 0.473 | 0.295 | 1.75E-23 | Chondrocytes | cox7c |
| kdelr2a1 | 5.6E-28 | 0.3554834 | 0.571 | 0.386 | 1.83E-23 | Chondrocytes | kdelr2a |
| syap11 | 5.7E-28 | 0.3782146 | 0.278 | 0.153 | 1.85E-23 | Chondrocytes | syap1 |
| mvb12a | 7.6E-28 | 0.5594479 | 0.151 | 0.068 | 2.47E-23 | Chondrocytes | mvb12a |
| copa1 | 8.5E-28 | 0.3854052 | 0.323 | 0.186 | 2.75E-23 | Chondrocytes | copa |
| rnaset22 | 8.7E-28 | 0.6456606 | 0.237 | 0.127 | 2.82E-23 | Chondrocytes | rnaset2 |
| furina2 | 8.9E-28 | 0.3818006 | 0.278 | 0.153 | 2.89E-23 | Chondrocytes | furina |
| unc5c1 | 9E-28 | 0.6237477 | 0.128 | 0.055 | 2.92E-23 | Chondrocytes | unc5c |
| yipf1 | 1.1E-27 | 0.8012628 | 0.11 | 0.044 | 3.54E-23 | Chondrocytes | yipf1 |
| tpd52l2b1 | 1.2E-27 | 0.3602744 | 0.366 | 0.216 | 3.9E-23 | Chondrocytes | tpd52l2b |
| aspn2 | 1.4E-27 | 1.1699534 | 0.528 | 0.401 | 4.51E-23 | Chondrocytes | aspn |
| itpkcb | 1.5E-27 | 0.8143255 | 0.11 | 0.044 | 4.8E-23 | Chondrocytes | itpkcb |
| slc7a10a4 | 1.6E-27 | 0.606469 | 0.205 | 0.105 | 5.21E-23 | Chondrocytes | slc7a10a |
| pou3f3b2 | 1.8E-27 | 0.4998528 | 0.128 | 0.054 | 6.01E-23 | Chondrocytes | pou3f3b |
| ndufv11 | 2E-27 | 0.424162 | 0.242 | 0.129 | 6.5E-23 | Chondrocytes | ndufv1 |
| tmem39a2 | 2.3E-27 | 0.4055582 | 0.268 | 0.147 | 7.51E-23 | Chondrocytes | tmem39a |
| prdx42 | 2.9E-27 | 0.3435755 | 0.547 | 0.359 | 9.47E-23 | Chondrocytes | prdx4 |
| higd1a3 | 3E-27 | 0.4664444 | 0.629 | 0.453 | 9.83E-23 | Chondrocytes | higd1a |
| zgc:162183 | 3.1E-27 | 0.6509212 | 0.113 | 0.046 | 1.01E-22 | Chondrocytes | zgc:162183 |
| itga11a3 | 3.1E-27 | 0.5089356 | 0.208 | 0.108 | 1.02E-22 | Chondrocytes | itga11a |
| gmpr2 | 3.2E-27 | 0.7538923 | 0.105 | 0.042 | 1.03E-22 | Chondrocytes | gmpr2 |
| ttpal | 3.4E-27 | 0.6966714 | 0.125 | 0.053 | 1.09E-22 | Chondrocytes | ttpal |
| gorasp22 | 3.7E-27 | 0.5251587 | 0.21 | 0.108 | 1.2E-22 | Chondrocytes | gorasp2 |
| rps54 | 3.7E-27 | 0.208958 | 0.991 | 0.972 | 1.21E-22 | Chondrocytes | rps5 |
| phf20b1 | 4.2E-27 | 0.3230686 | 0.303 | 0.172 | 1.38E-22 | Chondrocytes | phf20b |
| tmed21 | 4.3E-27 | 0.3011815 | 0.58 | 0.386 | 1.4E-22 | Chondrocytes | tmed2 |
| sec11a1 | 4.5E-27 | 0.3123615 | 0.517 | 0.338 | 1.46E-22 | Chondrocytes | sec11a |
| lrp10 | 6E-27 | 0.5338516 | 0.186 | 0.093 | 1.94E-22 | Chondrocytes | lrp10 |
| tmem183a | 6.2E-27 | 0.8148147 | 0.105 | 0.042 | 2.02E-22 | Chondrocytes | tmem183a |
| gli2a1 | 7.9E-27 | 0.7005552 | 0.149 | 0.068 | 2.56E-22 | Chondrocytes | gli2a |
| rcn12 | 1E-26 | 0.4747611 | 0.235 | 0.127 | 3.27E-22 | Chondrocytes | rcn1 |
| brd41 | 1.1E-26 | 0.3506242 | 0.417 | 0.258 | 3.5E-22 | Chondrocytes | brd4 |
| isg201 | 1.3E-26 | 0.8272463 | 0.115 | 0.049 | 4.15E-22 | Chondrocytes | isg20 |
| krtcap22 | 1.4E-26 | 0.4047204 | 0.423 | 0.266 | 4.56E-22 | Chondrocytes | krtcap2 |
| eif5a21 | 1.4E-26 | 0.2808224 | 0.663 | 0.462 | 4.64E-22 | Chondrocytes | eif5a2 |
| elocb1 | 1.4E-26 | 0.3452105 | 0.458 | 0.292 | 4.69E-22 | Chondrocytes | elocb |
| phc2a1 | 1.5E-26 | 0.5117402 | 0.148 | 0.068 | 4.73E-22 | Chondrocytes | phc2a |
| sra12 | 1.5E-26 | 0.524742 | 0.14 | 0.063 | 4.91E-22 | Chondrocytes | sra1 |
| TPGS1 | 1.8E-26 | 0.7594753 | 0.114 | 0.048 | 5.81E-22 | Chondrocytes | TPGS1 |
| faub1 | 1.8E-26 | 0.5361524 | 0.173 | 0.084 | 5.98E-22 | Chondrocytes | faub |

| | | | | | | | |
|---|---|---|---|---|---|---|---|
| echs11 | 1.9E-26 | 0.3367897 | 0.252 | 0.137 | 6.31E-22 | Chondrocytes | echs1 |
| tollip | 2E-26 | 0.6573599 | 0.124 | 0.053 | 6.54E-22 | Chondrocytes | tollip |
| fzd41 | 2.3E-26 | 0.5504615 | 0.12 | 0.051 | 7.4E-22 | Chondrocytes | fzd4 |
| spcs31 | 2.4E-26 | 0.282523 | 0.428 | 0.265 | 7.74E-22 | Chondrocytes | spcs3 |
| sec621 | 2.5E-26 | 0.3085491 | 0.385 | 0.233 | 8.23E-22 | Chondrocytes | sec62 |
| si:dkey-79d12.52 | 2.7E-26 | 0.7187528 | 0.232 | 0.127 | 8.92E-22 | Chondrocytes | si:dkey-79d12.5 |
| csnk2b1 | 2.8E-26 | 0.2813853 | 0.41 | 0.249 | 8.98E-22 | Chondrocytes | csnk2b |
| tram22 | 2.9E-26 | 0.4492535 | 0.208 | 0.107 | 9.34E-22 | Chondrocytes | tram2 |
| mia2 | 3E-26 | 0.9466449 | 0.108 | 0.044 | 9.61E-22 | Chondrocytes | mia2 |
| tspan4a2 | 3E-26 | 0.4201634 | 0.248 | 0.135 | 9.62E-22 | Chondrocytes | tspan4a |
| slirp1 | 3.2E-26 | 0.3938319 | 0.256 | 0.14 | 1.03E-21 | Chondrocytes | slirp |
| atp6ap1b | 3.2E-26 | 0.5051429 | 0.146 | 0.067 | 1.05E-21 | Chondrocytes | atp6ap1b |
| slc43a1b1 | 3.3E-26 | 0.7702459 | 0.115 | 0.048 | 1.08E-21 | Chondrocytes | slc43a1b |
| tcta2 | 4.1E-26 | 0.6262473 | 0.177 | 0.088 | 1.34E-21 | Chondrocytes | tcta |
| eef1b23 | 4.2E-26 | 0.2297608 | 0.965 | 0.928 | 1.38E-21 | Chondrocytes | eef1b2 |
| arpin | 4.8E-26 | 0.53208 | 0.143 | 0.066 | 1.55E-21 | Chondrocytes | arpin |
| vma211 | 4.8E-26 | 0.5651985 | 0.211 | 0.111 | 1.55E-21 | Chondrocytes | vma21 |
| skib5 | 4.9E-26 | 0.2886078 | 0.339 | 0.199 | 1.59E-21 | Chondrocytes | skib |
| ufd1l2 | 5E-26 | 0.5083098 | 0.202 | 0.104 | 1.64E-21 | Chondrocytes | ufd1l |
| mdm2 | 5.2E-26 | 1.0496035 | 0.121 | 0.052 | 1.68E-21 | Chondrocytes | mdm2 |
| ube2g21 | 5.6E-26 | 0.4291368 | 0.21 | 0.109 | 1.81E-21 | Chondrocytes | ube2g2 |
| ptp4a2a1 | 5.8E-26 | 0.340016 | 0.25 | 0.135 | 1.87E-21 | Chondrocytes | ptp4a2a |
| dlg1l | 5.9E-26 | 0.5039553 | 0.161 | 0.077 | 1.93E-21 | Chondrocytes | dlg1l |
| ccn2b1 | 6E-26 | 1.1553557 | 0.317 | 0.203 | 1.94E-21 | Chondrocytes | ccn2b |
| ugdh2 | 6.3E-26 | 0.3895312 | 0.302 | 0.175 | 2.04E-21 | Chondrocytes | ugdh |
| bcar11 | 6.4E-26 | 0.5025292 | 0.183 | 0.091 | 2.09E-21 | Chondrocytes | bcar1 |
| RASA2 | 8.4E-26 | 0.6762048 | 0.106 | 0.044 | 2.74E-21 | Chondrocytes | RASA2 |
| loxl41 | 8.6E-26 | 1.2940915 | 0.128 | 0.058 | 2.81E-21 | Chondrocytes | loxl4 |
| zc2hc1a | 8.7E-26 | 0.7566683 | 0.107 | 0.044 | 2.81E-21 | Chondrocytes | zc2hc1a |
| zfyve9a1 | 8.7E-26 | 0.3202015 | 0.275 | 0.153 | 2.84E-21 | Chondrocytes | zfyve9a |
| C20H1orf1983 | 9.2E-26 | 0.4634735 | 0.307 | 0.181 | 3.01E-21 | Chondrocytes | C20H1orf198 |
| zgc:922422 | 9.5E-26 | 0.8030941 | 0.204 | 0.109 | 3.09E-21 | Chondrocytes | zgc:92242 |
| eif61 | 9.9E-26 | 0.3623717 | 0.299 | 0.173 | 3.22E-21 | Chondrocytes | eif6 |
| megf82 | 1.4E-25 | 0.3032732 | 0.235 | 0.126 | 4.45E-21 | Chondrocytes | megf8 |
| piezo12 | 1.4E-25 | 0.4020715 | 0.227 | 0.121 | 4.68E-21 | Chondrocytes | piezo1 |
| sec61g2 | 1.4E-25 | 0.2655783 | 0.968 | 0.923 | 4.69E-21 | Chondrocytes | sec61g |
| pdgfrl4 | 1.5E-25 | 0.8136473 | 0.706 | 0.582 | 4.84E-21 | Chondrocytes | pdgfrl |
| hsd17b12b1 | 1.6E-25 | 0.4555139 | 0.177 | 0.088 | 5.32E-21 | Chondrocytes | hsd17b12b |
| ano10a | 1.9E-25 | 0.5530735 | 0.134 | 0.06 | 6.26E-21 | Chondrocytes | ano10a |
| ssr33 | 1.9E-25 | 0.3648984 | 0.757 | 0.573 | 6.27E-21 | Chondrocytes | ssr3 |
| sema4c | 2.1E-25 | 0.6474767 | 0.109 | 0.045 | 6.94E-21 | Chondrocytes | sema4c |
| tmem184ba1 | 2.2E-25 | 0.5701369 | 0.145 | 0.067 | 7.3E-21 | Chondrocytes | tmem184ba |
| atp5mc3a1 | 3.1E-25 | 0.2789275 | 0.427 | 0.267 | 1E-20 | Chondrocytes | atp5mc3a |
| ccdc471 | 3.3E-25 | 0.2756956 | 0.251 | 0.136 | 1.08E-20 | Chondrocytes | ccdc47 |
| stt3b1 | 5.1E-25 | 0.4756535 | 0.202 | 0.106 | 1.65E-20 | Chondrocytes | stt3b |
| rnf130 | 5.3E-25 | 0.5262666 | 0.126 | 0.056 | 1.71E-20 | Chondrocytes | rnf130 |
| pard3ab | 5.5E-25 | 0.6039623 | 0.125 | 0.055 | 1.79E-20 | Chondrocytes | pard3ab |
| yipf51 | 6.1E-25 | 0.4100955 | 0.226 | 0.121 | 1.99E-20 | Chondrocytes | yipf5 |

| | | | | | | | |
|---|---|---|---|---|---|---|---|
| uqcc11 | 6.9E-25 | 0.6047906 | 0.169 | 0.084 | 2.23E-20 | Chondrocytes | uqcc1 |
| vegfaa2 | 7.2E-25 | 0.9738798 | 0.208 | 0.114 | 2.34E-20 | Chondrocytes | vegfaa |
| mical3a.11 | 7.5E-25 | 0.4179259 | 0.198 | 0.102 | 2.43E-20 | Chondrocytes | mical3a.1 |
| stk35l1 | 8E-25 | 0.4992036 | 0.226 | 0.123 | 2.59E-20 | Chondrocytes | stk35l |
| psmd71 | 8.3E-25 | 0.2780336 | 0.395 | 0.243 | 2.69E-20 | Chondrocytes | psmd7 |
| tm9sf1 | 9.8E-25 | 0.7423958 | 0.113 | 0.049 | 3.2E-20 | Chondrocytes | tm9sf1 |
| blvrb | 1E-24 | 0.7017898 | 0.122 | 0.054 | 3.28E-20 | Chondrocytes | blvrb |
| arcn1a1 | 1.1E-24 | 0.5739449 | 0.186 | 0.096 | 3.49E-20 | Chondrocytes | arcn1a |
| map1sb1 | 1.2E-24 | 0.547465 | 0.129 | 0.058 | 3.91E-20 | Chondrocytes | map1sb |
| tecrb | 1.3E-24 | 0.5941815 | 0.114 | 0.049 | 4.13E-20 | Chondrocytes | tecrb |
| si:ch211-207i1.21 | 1.4E-24 | 0.3944396 | 0.165 | 0.081 | 4.45E-20 | Chondrocytes | si:ch211-207i1.2 |
| vaspb3 | 1.5E-24 | 0.4829674 | 0.189 | 0.098 | 4.91E-20 | Chondrocytes | vaspb |
| tmem14ca1 | 1.9E-24 | 0.3829651 | 0.222 | 0.119 | 6.12E-20 | Chondrocytes | tmem14ca |
| pfkfb33 | 1.9E-24 | 0.6198069 | 0.153 | 0.074 | 6.22E-20 | Chondrocytes | pfkfb3 |
| srp92 | 2.2E-24 | 0.5281674 | 0.2 | 0.106 | 7.25E-20 | Chondrocytes | srp9 |
| map3k3 | 2.2E-24 | 0.6112903 | 0.122 | 0.054 | 7.31E-20 | Chondrocytes | map3k3 |
| rnf71 | 2.3E-24 | 0.4394491 | 0.337 | 0.206 | 7.33E-20 | Chondrocytes | rnf7 |
| arhgef42 | 2.3E-24 | 0.364114 | 0.154 | 0.074 | 7.42E-20 | Chondrocytes | arhgef4 |
| rpn11 | 2.6E-24 | 0.4170918 | 0.343 | 0.209 | 8.49E-20 | Chondrocytes | rpn1 |
| adipor23 | 3.1E-24 | 0.4586305 | 0.294 | 0.174 | 9.96E-20 | Chondrocytes | adipor2 |
| atp5f1d2 | 3.1E-24 | 0.2657008 | 0.504 | 0.329 | 1E-19 | Chondrocytes | atp5f1d |
| ndufb81 | 3.2E-24 | 0.3079254 | 0.503 | 0.332 | 1.04E-19 | Chondrocytes | ndufb8 |
| uck2b2 | 4.2E-24 | 0.6530003 | 0.118 | 0.052 | 1.38E-19 | Chondrocytes | uck2b |
| gtf2b2 | 4.4E-24 | 0.5333828 | 0.158 | 0.077 | 1.43E-19 | Chondrocytes | gtf2b |
| ube2d21 | 4.8E-24 | 0.3045199 | 0.568 | 0.384 | 1.55E-19 | Chondrocytes | ube2d2 |
| efemp2a3 | 5.1E-24 | 0.3484512 | 0.525 | 0.358 | 1.65E-19 | Chondrocytes | efemp2a |
| tmem2141 | 6.2E-24 | 0.458097 | 0.226 | 0.124 | 2.01E-19 | Chondrocytes | tmem214 |
| psma22 | 6.3E-24 | 0.301099 | 0.333 | 0.199 | 2.04E-19 | Chondrocytes | psma2 |
| ywhabl1 | 6.4E-24 | 0.2927697 | 0.607 | 0.413 | 2.07E-19 | Chondrocytes | ywhabl |
| eef1db4 | 6.4E-24 | 0.2985519 | 0.763 | 0.576 | 2.08E-19 | Chondrocytes | eef1db |
| spcs21 | 7.4E-24 | 0.2454634 | 0.448 | 0.282 | 2.41E-19 | Chondrocytes | spcs2 |
| scaf111 | 7.9E-24 | 0.6022888 | 0.131 | 0.061 | 2.57E-19 | Chondrocytes | scaf11 |
| zgc:1534092 | 8E-24 | 0.3353235 | 0.595 | 0.415 | 2.62E-19 | Chondrocytes | zgc:153409 |
| ndufv31 | 8.5E-24 | 0.3162228 | 0.232 | 0.127 | 2.76E-19 | Chondrocytes | ndufv3 |
| rpl5a4 | 8.9E-24 | 0.2265703 | 0.965 | 0.917 | 2.88E-19 | Chondrocytes | rpl5a |
| brdt3 | 9.8E-24 | 0.4339851 | 0.454 | 0.307 | 3.2E-19 | Chondrocytes | brdt |
| C19H6orf621 | 1.1E-23 | 0.2532773 | 0.587 | 0.397 | 3.55E-19 | Chondrocytes | C19H6orf62 |
| minpp1a | 1.2E-23 | 0.7258237 | 0.106 | 0.045 | 3.78E-19 | Chondrocytes | minpp1a |
| rab18b | 1.4E-23 | 0.3256671 | 0.212 | 0.113 | 4.43E-19 | Chondrocytes | rab18b |
| kras1 | 1.6E-23 | 0.450448 | 0.293 | 0.175 | 5.08E-19 | Chondrocytes | kras |
| gskip3 | 1.6E-23 | 0.3894796 | 0.274 | 0.159 | 5.27E-19 | Chondrocytes | gskip |
| hsbp1b1 | 1.7E-23 | 0.296539 | 0.423 | 0.267 | 5.57E-19 | Chondrocytes | hsbp1b |
| gcat1 | 1.8E-23 | 0.5115989 | 0.153 | 0.075 | 5.74E-19 | Chondrocytes | gcat |
| zmp:0000000729 | 1.9E-23 | 0.5933953 | 0.111 | 0.048 | 6.02E-19 | Chondrocytes | zmp:0000000729 |
| pigx | 2.1E-23 | 0.7979234 | 0.108 | 0.047 | 6.76E-19 | Chondrocytes | pigx |
| cdkn1ba3 | 2.1E-23 | 0.4744411 | 0.211 | 0.115 | 6.77E-19 | Chondrocytes | cdkn1ba |
| prmt12 | 2.1E-23 | 0.4231131 | 0.307 | 0.186 | 6.98E-19 | Chondrocytes | prmt1 |
| magt12 | 2.3E-23 | 0.2759776 | 0.41 | 0.257 | 7.39E-19 | Chondrocytes | magt1 |

| | | | | | | | |
|---|---|---|---|---|---|---|---|
| uqcrfs11 | 2.4E-23 | 0.2601549 | 0.355 | 0.217 | 7.81E-19 | Chondrocytes | uqcrfs1 |
| mcl1a4 | 2.5E-23 | 0.6212526 | 0.846 | 0.769 | 7.98E-19 | Chondrocytes | mcl1a |
| dctn31 | 2.6E-23 | 0.3555969 | 0.26 | 0.148 | 8.38E-19 | Chondrocytes | dctn3 |
| dtd11 | 2.7E-23 | 0.8264311 | 0.109 | 0.047 | 8.77E-19 | Chondrocytes | dtd1 |
| g3bp11 | 3.7E-23 | 0.3172277 | 0.286 | 0.166 | 1.22E-18 | Chondrocytes | g3bp1 |
| lsm61 | 3.8E-23 | 0.318876 | 0.289 | 0.169 | 1.22E-18 | Chondrocytes | lsm6 |
| si:dkey-112e7.21 | 4.3E-23 | 0.3337112 | 0.359 | 0.223 | 1.39E-18 | Chondrocytes | si:dkey-112e7.2 |
| cct6a1 | 4.7E-23 | 0.3213271 | 0.665 | 0.474 | 1.53E-18 | Chondrocytes | cct6a |
| smim8 | 5.2E-23 | 0.6312667 | 0.115 | 0.051 | 1.69E-18 | Chondrocytes | smim8 |
| bsdc11 | 5.4E-23 | 0.5524242 | 0.178 | 0.092 | 1.77E-18 | Chondrocytes | bsdc1 |
| atp5meb1 | 5.5E-23 | 0.292185 | 0.734 | 0.532 | 1.78E-18 | Chondrocytes | atp5meb |
| ankrd54 | 6E-23 | 0.7575509 | 0.104 | 0.045 | 1.95E-18 | Chondrocytes | ankrd54 |
| p3h4 | 6.3E-23 | 0.712406 | 0.107 | 0.046 | 2.05E-18 | Chondrocytes | p3h4 |
| ndufs31 | 6.5E-23 | 0.2611129 | 0.232 | 0.128 | 2.12E-18 | Chondrocytes | ndufs3 |
| fbxw11b2 | 6.9E-23 | 0.333116 | 0.271 | 0.157 | 2.24E-18 | Chondrocytes | fbxw11b |
| sar1b1 | 7.2E-23 | 0.3189272 | 0.451 | 0.295 | 2.34E-18 | Chondrocytes | sar1b |
| tpt12 | 8.7E-23 | 0.2366062 | 0.968 | 0.926 | 2.83E-18 | Chondrocytes | tpt1 |
| ube4b1 | 9.2E-23 | 0.4920173 | 0.137 | 0.065 | 2.98E-18 | Chondrocytes | ube4b |
| tia1l1 | 1E-22 | 0.3719304 | 0.242 | 0.136 | 3.36E-18 | Chondrocytes | tia1l |
| ip6k2b4 | 1E-22 | 0.5145412 | 0.491 | 0.34 | 3.36E-18 | Chondrocytes | ip6k2b |
| zgc:162339 | 1.1E-22 | 0.5216751 | 0.146 | 0.071 | 3.53E-18 | Chondrocytes | zgc:162339 |
| MED91 | 1.1E-22 | 0.5628161 | 0.143 | 0.069 | 3.61E-18 | Chondrocytes | MED9 |
| dnajc3b1 | 1.3E-22 | 0.586033 | 0.128 | 0.06 | 4.16E-18 | Chondrocytes | dnajc3b |
| cyth1a.12 | 1.3E-22 | 0.4201737 | 0.237 | 0.133 | 4.17E-18 | Chondrocytes | cyth1a.1 |
| sppl2 | 1.3E-22 | 0.5239874 | 0.128 | 0.059 | 4.19E-18 | Chondrocytes | sppl2 |
| mob2a3 | 1.4E-22 | 0.3978005 | 0.238 | 0.135 | 4.64E-18 | Chondrocytes | mob2a |
| pla2g7 | 1.6E-22 | 0.6951255 | 0.102 | 0.043 | 5.06E-18 | Chondrocytes | pla2g7 |
| gosr1 | 1.6E-22 | 0.5640555 | 0.142 | 0.069 | 5.21E-18 | Chondrocytes | gosr1 |
| thbs2b3 | 1.8E-22 | 0.3291317 | 0.383 | 0.248 | 5.75E-18 | Chondrocytes | thbs2b |
| fbxo252 | 1.8E-22 | 0.568156 | 0.155 | 0.077 | 5.87E-18 | Chondrocytes | fbxo25 |
| ext2 | 1.8E-22 | 0.3524994 | 0.195 | 0.104 | 5.99E-18 | Chondrocytes | ext2 |
| ost42 | 1.9E-22 | 0.4601041 | 0.257 | 0.148 | 6.17E-18 | Chondrocytes | ost4 |
| ppic | 1.9E-22 | 0.6630013 | 0.118 | 0.054 | 6.3E-18 | Chondrocytes | ppic |
| nbr1a1 | 1.9E-22 | 0.5589574 | 0.157 | 0.079 | 6.33E-18 | Chondrocytes | nbr1a |
| lysmd4 | 2.1E-22 | 0.7199901 | 0.105 | 0.045 | 6.77E-18 | Chondrocytes | lysmd4 |
| atp5pb1 | 2.2E-22 | 0.2604987 | 0.649 | 0.459 | 7.02E-18 | Chondrocytes | atp5pb |
| ganab1 | 2.2E-22 | 0.691559 | 0.126 | 0.059 | 7.06E-18 | Chondrocytes | ganab |
| rer11 | 2.7E-22 | 0.3416584 | 0.331 | 0.203 | 8.82E-18 | Chondrocytes | rer1 |
| ggh | 2.9E-22 | 0.4073053 | 0.136 | 0.064 | 9.57E-18 | Chondrocytes | ggh |
| vps28 | 3.1E-22 | 0.3837814 | 0.206 | 0.112 | 1.01E-17 | Chondrocytes | vps28 |
| fam214a1 | 3.4E-22 | 0.5992596 | 0.175 | 0.092 | 1.1E-17 | Chondrocytes | fam214a |
| ndufa4l2 | 4.1E-22 | 0.2486872 | 0.88 | 0.749 | 1.33E-17 | Chondrocytes | ndufa4l |
| irs2b4 | 4.5E-22 | 0.233433 | 0.362 | 0.227 | 1.47E-17 | Chondrocytes | irs2b |
| derl21 | 5.3E-22 | 0.3539159 | 0.175 | 0.091 | 1.73E-17 | Chondrocytes | derl2 |
| hax1 | 5.4E-22 | 0.6876063 | 0.111 | 0.05 | 1.74E-17 | Chondrocytes | hax1 |
| rnf1852 | 6.5E-22 | 0.3309014 | 0.27 | 0.158 | 2.1E-17 | Chondrocytes | rnf185 |
| lrp1aa4 | 6.5E-22 | 0.2437633 | 0.85 | 0.669 | 2.13E-17 | Chondrocytes | lrp1aa |
| nipa21 | 6.7E-22 | 0.3856935 | 0.15 | 0.074 | 2.19E-17 | Chondrocytes | nipa2 |

| | | | | | | | |
|---|---|---|---|---|---|---|---|
| pitpnb1 | 7.5E-22 | 0.6745228 | 0.106 | 0.047 | 2.46E-17 | Chondrocytes | pitpnb |
| csnk1g2b | 7.6E-22 | 0.3486207 | 0.283 | 0.168 | 2.48E-17 | Chondrocytes | csnk1g2b |
| edf11 | 7.7E-22 | 0.2678627 | 0.705 | 0.508 | 2.49E-17 | Chondrocytes | edf1 |
| baz2ba2 | 8.6E-22 | 0.2566692 | 0.282 | 0.167 | 2.78E-17 | Chondrocytes | baz2ba |
| usp361 | 9.7E-22 | 0.3577234 | 0.322 | 0.198 | 3.16E-17 | Chondrocytes | usp36 |
| atp5f1e2 | 1.1E-21 | 0.2246392 | 0.598 | 0.408 | 3.42E-17 | Chondrocytes | atp5f1e |
| eva1ba2 | 1.1E-21 | 0.2045287 | 0.198 | 0.106 | 3.54E-17 | Chondrocytes | eva1ba |
| parp16 | 1.2E-21 | 0.3713452 | 0.168 | 0.086 | 3.8E-17 | Chondrocytes | parp16 |
| CABZ01054965.1 | 1.2E-21 | 0.5566124 | 0.119 | 0.055 | 3.94E-17 | Chondrocytes | CABZ01054965.1 |
| fundc21 | 1.5E-21 | 0.2016462 | 0.46 | 0.298 | 4.72E-17 | Chondrocytes | fundc2 |
| MYO1D1 | 1.7E-21 | 0.4886873 | 0.187 | 0.101 | 5.47E-17 | Chondrocytes | MYO1D |
| prkar1ab | 2E-21 | 0.3464133 | 0.202 | 0.109 | 6.41E-17 | Chondrocytes | prkar1ab |
| si:ch1073-325m22.2 | 2.1E-21 | 0.2059241 | 0.457 | 0.298 | 6.79E-17 | Chondrocytes | si:ch1073-325m22. |
| p4ha1a3 | 2.4E-21 | 0.3613448 | 0.179 | 0.094 | 7.77E-17 | Chondrocytes | p4ha1a |
| si:ch211-150o23.3 | 2.4E-21 | 0.7150401 | 0.1 | 0.043 | 7.88E-17 | Chondrocytes | si:ch211-150o23.3 |
| prss163 | 2.5E-21 | 0.4240995 | 0.17 | 0.088 | 8.11E-17 | Chondrocytes | prss16 |
| cops7a1 | 3E-21 | 0.3930921 | 0.202 | 0.11 | 9.86E-17 | Chondrocytes | cops7a |
| trip112 | 3.1E-21 | 0.4702188 | 0.17 | 0.089 | 1E-16 | Chondrocytes | trip11 |
| tsc22d2 | 3.2E-21 | 0.5756094 | 0.134 | 0.065 | 1.03E-16 | Chondrocytes | tsc22d2 |
| pdlim22 | 3.3E-21 | 0.4348452 | 0.26 | 0.154 | 1.09E-16 | Chondrocytes | pdlim2 |
| FBXO481 | 3.3E-21 | 0.6757927 | 0.125 | 0.059 | 1.09E-16 | Chondrocytes | FBXO48 |
| atp5mc3b2 | 3.4E-21 | 0.2263354 | 0.786 | 0.618 | 1.09E-16 | Chondrocytes | atp5mc3b |
| parp31 | 3.6E-21 | 0.5684744 | 0.165 | 0.086 | 1.16E-16 | Chondrocytes | parp3 |
| bud311 | 3.6E-21 | 0.2658036 | 0.334 | 0.204 | 1.18E-16 | Chondrocytes | bud31 |
| grb2b1 | 3.8E-21 | 0.262745 | 0.272 | 0.16 | 1.25E-16 | Chondrocytes | grb2b |
| chmp4bb1 | 3.9E-21 | 0.3817453 | 0.257 | 0.151 | 1.28E-16 | Chondrocytes | chmp4bb |
| zgc:1536752 | 4.4E-21 | 0.3505328 | 0.397 | 0.259 | 1.43E-16 | Chondrocytes | zgc:153675 |
| tmem2582 | 4.6E-21 | 0.3255763 | 0.757 | 0.583 | 1.48E-16 | Chondrocytes | tmem258 |
| baz1a2 | 4.8E-21 | 0.4334104 | 0.202 | 0.112 | 1.56E-16 | Chondrocytes | baz1a |
| casc4 | 5.3E-21 | 0.3420812 | 0.165 | 0.085 | 1.73E-16 | Chondrocytes | casc4 |
| mbpb1 | 5.5E-21 | 0.3791584 | 0.122 | 0.057 | 1.79E-16 | Chondrocytes | mbpb |
| ccdc1371 | 6.6E-21 | 0.5867467 | 0.102 | 0.045 | 2.16E-16 | Chondrocytes | ccdc137 |
| ube2h1 | 8.7E-21 | 0.3783165 | 0.198 | 0.108 | 2.82E-16 | Chondrocytes | ube2h |
| elmo12 | 8.7E-21 | 0.3142227 | 0.176 | 0.093 | 2.84E-16 | Chondrocytes | elmo1 |
| srgap22 | 1E-20 | 0.4373087 | 0.16 | 0.082 | 3.28E-16 | Chondrocytes | srgap2 |
| nek71 | 1E-20 | 0.3007731 | 0.211 | 0.117 | 3.4E-16 | Chondrocytes | nek7 |
| hikeshi1 | 1.1E-20 | 0.3754781 | 0.188 | 0.101 | 3.61E-16 | Chondrocytes | hikeshi |
| hint2 | 1.1E-20 | 0.5395862 | 0.128 | 0.061 | 3.68E-16 | Chondrocytes | hint2 |
| abhd2a | 1.2E-20 | 0.436653 | 0.163 | 0.085 | 3.79E-16 | Chondrocytes | abhd2a |
| gtpbp43 | 1.3E-20 | 0.3619285 | 0.573 | 0.407 | 4.15E-16 | Chondrocytes | gtpbp4 |
| uqcc32 | 1.4E-20 | 0.4146236 | 0.158 | 0.081 | 4.57E-16 | Chondrocytes | uqcc3 |
| naa201 | 1.6E-20 | 0.3218629 | 0.251 | 0.147 | 5.08E-16 | Chondrocytes | naa20 |
| ndufab1b2 | 1.6E-20 | 0.2381243 | 0.325 | 0.2 | 5.21E-16 | Chondrocytes | ndufab1b |
| dbi1 | 1.7E-20 | 0.2981802 | 0.614 | 0.437 | 5.51E-16 | Chondrocytes | dbi |
| hmgb1a2 | 1.7E-20 | 0.2170559 | 0.754 | 0.559 | 5.52E-16 | Chondrocytes | hmgb1a |
| ptbp1a2 | 1.7E-20 | 0.2635391 | 0.357 | 0.226 | 5.64E-16 | Chondrocytes | ptbp1a |
| suclg11 | 1.8E-20 | 0.3241533 | 0.285 | 0.172 | 5.93E-16 | Chondrocytes | suclg1 |
| si:ch73-147o17.1 | 2E-20 | 0.3644991 | 0.152 | 0.078 | 6.43E-16 | Chondrocytes | si:ch73-147o17.1 |

| | | | | | | | |
|---|---|---|---|---|---|---|---|
| nomo1 | 2.2E-20 | 0.667964 | 0.1 | 0.045 | 7.07E-16 | Chondrocytes | nomo |
| ndufs51 | 2.3E-20 | 0.2860204 | 0.288 | 0.175 | 7.46E-16 | Chondrocytes | ndufs5 |
| tp53bp2b | 2.4E-20 | 0.6747027 | 0.112 | 0.052 | 7.81E-16 | Chondrocytes | tp53bp2b |
| ENSDARG00000055[illegible] | 3.1E-20 | 0.4723463 | 0.15 | 0.077 | 1.02E-15 | Chondrocytes | ENSDARG00000055 |
| spred2b1 | 3.3E-20 | 0.3082238 | 0.192 | 0.105 | 1.09E-15 | Chondrocytes | spred2b |
| nbr1b | 3.7E-20 | 0.52397 | 0.128 | 0.062 | 1.19E-15 | Chondrocytes | nbr1b |
| man2a21 | 3.8E-20 | 0.3216758 | 0.223 | 0.127 | 1.25E-15 | Chondrocytes | man2a2 |
| prickle1a3 | 3.9E-20 | 0.5109323 | 0.121 | 0.058 | 1.26E-15 | Chondrocytes | prickle1a |
| cdca41 | 3.9E-20 | 0.3937907 | 0.148 | 0.075 | 1.28E-15 | Chondrocytes | cdca4 |
| rrbp1a3 | 4.1E-20 | 0.3054509 | 0.597 | 0.426 | 1.34E-15 | Chondrocytes | rrbp1a |
| ccdc186 | 4.4E-20 | 0.4333023 | 0.147 | 0.075 | 1.43E-15 | Chondrocytes | ccdc186 |
| ergic21 | 4.8E-20 | 0.219287 | 0.267 | 0.158 | 1.56E-15 | Chondrocytes | ergic2 |
| aspa | 4.9E-20 | 0.5280476 | 0.122 | 0.058 | 1.58E-15 | Chondrocytes | aspa |
| rab11ba1 | 5.5E-20 | 0.255577 | 0.297 | 0.18 | 1.78E-15 | Chondrocytes | rab11ba |
| prkar1aa1 | 5.6E-20 | 0.2199119 | 0.245 | 0.141 | 1.81E-15 | Chondrocytes | prkar1aa |
| bbip1 | 5.7E-20 | 0.3463098 | 0.217 | 0.124 | 1.86E-15 | Chondrocytes | bbip1 |
| map2k2a | 5.7E-20 | 0.5409952 | 0.105 | 0.048 | 1.86E-15 | Chondrocytes | map2k2a |
| ndufa22 | 6E-20 | 0.3883735 | 0.171 | 0.092 | 1.95E-15 | Chondrocytes | ndufa2 |
| dnajc12 | 6.5E-20 | 0.4466568 | 0.125 | 0.061 | 2.1E-15 | Chondrocytes | dnajc12 |
| calcoco1a1 | 7.8E-20 | 0.3851753 | 0.183 | 0.1 | 2.53E-15 | Chondrocytes | calcoco1a |
| dnajc41 | 9.1E-20 | 0.3881106 | 0.174 | 0.094 | 2.97E-15 | Chondrocytes | dnajc4 |
| lmo7a2 | 9.8E-20 | 0.2912529 | 0.254 | 0.151 | 3.2E-15 | Chondrocytes | lmo7a |
| farsa2 | 1E-19 | 0.3910734 | 0.154 | 0.08 | 3.37E-15 | Chondrocytes | farsa |
| syncripl1 | 1.1E-19 | 0.2937324 | 0.21 | 0.118 | 3.43E-15 | Chondrocytes | syncripl |
| ndufc21 | 1.2E-19 | 0.2178064 | 0.418 | 0.274 | 3.89E-15 | Chondrocytes | ndufc2 |
| psmd81 | 1.2E-19 | 0.2575155 | 0.254 | 0.149 | 3.94E-15 | Chondrocytes | psmd8 |
| zgc:770561 | 1.6E-19 | 0.5594825 | 0.134 | 0.067 | 5.19E-15 | Chondrocytes | zgc:77056 |
| zgc:1935411 | 1.6E-19 | 0.228264 | 0.564 | 0.389 | 5.24E-15 | Chondrocytes | zgc:193541 |
| plgrkt2 | 2E-19 | 0.5859095 | 0.114 | 0.054 | 6.49E-15 | Chondrocytes | plgrkt |
| pam | 2.2E-19 | 0.3972046 | 0.176 | 0.095 | 7.03E-15 | Chondrocytes | pam |
| eif3ha3 | 2.3E-19 | 0.2289867 | 0.602 | 0.416 | 7.33E-15 | Chondrocytes | eif3ha |
| gatad2ab | 2.3E-19 | 0.3766274 | 0.138 | 0.069 | 7.37E-15 | Chondrocytes | gatad2ab |
| arhgap311 | 2.3E-19 | 0.3623929 | 0.198 | 0.111 | 7.4E-15 | Chondrocytes | arhgap31 |
| vps4b | 2.3E-19 | 0.3172158 | 0.389 | 0.255 | 7.59E-15 | Chondrocytes | vps4b |
| rrs12 | 2.4E-19 | 0.3959877 | 0.194 | 0.109 | 7.66E-15 | Chondrocytes | rrs1 |
| timmdc11 | 2.5E-19 | 0.3337818 | 0.165 | 0.087 | 7.97E-15 | Chondrocytes | timmdc1 |
| tmed91 | 2.5E-19 | 0.259504 | 0.351 | 0.224 | 8.14E-15 | Chondrocytes | tmed9 |
| micos131 | 2.5E-19 | 0.348617 | 0.211 | 0.121 | 8.25E-15 | Chondrocytes | micos13 |
| si:dkey-264d12.5 | 2.9E-19 | 0.6037415 | 0.117 | 0.056 | 9.27E-15 | Chondrocytes | si:dkey-264d12.5 |
| dusp71 | 3.1E-19 | 0.4433219 | 0.142 | 0.072 | 1.01E-14 | Chondrocytes | dusp7 |
| prdx32 | 3.2E-19 | 0.2949872 | 0.236 | 0.139 | 1.04E-14 | Chondrocytes | prdx3 |
| CDPF12 | 3.3E-19 | 0.3844116 | 0.286 | 0.176 | 1.07E-14 | Chondrocytes | CDPF1 |
| ahi11 | 3.3E-19 | 0.347683 | 0.198 | 0.111 | 1.08E-14 | Chondrocytes | ahi1 |
| LOXL41 | 3.5E-19 | 0.3193488 | 0.128 | 0.063 | 1.14E-14 | Chondrocytes | LOXL4 |
| adh53 | 3.5E-19 | 0.3293724 | 0.688 | 0.513 | 1.15E-14 | Chondrocytes | adh5 |
| lonrf1l4 | 4.1E-19 | 0.3634418 | 0.25 | 0.149 | 1.33E-14 | Chondrocytes | lonrf1l |
| cnih11 | 4.2E-19 | 0.3807303 | 0.174 | 0.094 | 1.35E-14 | Chondrocytes | cnih1 |
| micos101 | 4.3E-19 | 0.2235093 | 0.278 | 0.168 | 1.4E-14 | Chondrocytes | micos10 |

| | | | | | | | |
|---|---|---|---|---|---|---|---|
| niban1b3 | 6.5E-19 | 0.2534054 | 0.173 | 0.093 | 2.1E-14 | Chondrocytes | niban1b |
| prkcaa | 7.1E-19 | 0.2612853 | 0.188 | 0.104 | 2.31E-14 | Chondrocytes | prkcaa |
| smco42 | 7.2E-19 | 0.2681007 | 0.237 | 0.139 | 2.35E-14 | Chondrocytes | smco4 |
| calr1 | 7.4E-19 | 0.3191283 | 0.31 | 0.195 | 2.4E-14 | Chondrocytes | calr |
| CABZ01026681.11 | 7.5E-19 | 0.3866146 | 0.155 | 0.082 | 2.45E-14 | Chondrocytes | CABZ01026681.1 |
| inpp5l | 7.7E-19 | 0.2250626 | 0.177 | 0.096 | 2.51E-14 | Chondrocytes | inpp5l |
| eif1ad2 | 8.3E-19 | 0.3297116 | 0.233 | 0.138 | 2.7E-14 | Chondrocytes | eif1ad |
| cnot11 | 9.4E-19 | 0.2653524 | 0.19 | 0.106 | 3.05E-14 | Chondrocytes | cnot1 |
| elof12 | 1.3E-18 | 0.3382935 | 0.238 | 0.142 | 4.17E-14 | Chondrocytes | elof1 |
| gtf2a11 | 1.3E-18 | 0.4271825 | 0.153 | 0.081 | 4.18E-14 | Chondrocytes | gtf2a1 |
| hadhb1 | 1.3E-18 | 0.282002 | 0.223 | 0.129 | 4.26E-14 | Chondrocytes | hadhb |
| cyc11 | 1.3E-18 | 0.2880149 | 0.273 | 0.168 | 4.38E-14 | Chondrocytes | cyc1 |
| si:dkey-28b4.8 | 1.6E-18 | 0.2811648 | 0.105 | 0.049 | 5.1E-14 | Chondrocytes | si:dkey-28b4.8 |
| zgc:1626982 | 1.6E-18 | 0.2079652 | 0.2 | 0.112 | 5.12E-14 | Chondrocytes | zgc:162698 |
| cct73 | 1.6E-18 | 0.228423 | 0.53 | 0.368 | 5.3E-14 | Chondrocytes | cct7 |
| pdia74 | 1.7E-18 | 0.3358536 | 0.24 | 0.142 | 5.46E-14 | Chondrocytes | pdia7 |
| prelid1a2 | 1.7E-18 | 0.3101623 | 0.212 | 0.123 | 5.51E-14 | Chondrocytes | prelid1a |
| sema4ba2 | 1.9E-18 | 0.2907428 | 0.156 | 0.083 | 6.3E-14 | Chondrocytes | sema4ba |
| acadsb2 | 2E-18 | 0.5217548 | 0.125 | 0.063 | 6.38E-14 | Chondrocytes | acadsb |
| anxa11a2 | 2.3E-18 | 0.4055607 | 0.557 | 0.407 | 7.42E-14 | Chondrocytes | anxa11a |
| tm2d3 | 2.5E-18 | 0.4378113 | 0.136 | 0.07 | 8.06E-14 | Chondrocytes | tm2d3 |
| calhm2.1 | 2.5E-18 | 0.5368023 | 0.134 | 0.069 | 8.2E-14 | Chondrocytes | calhm2.1 |
| znhit1 | 2.7E-18 | 0.4499249 | 0.118 | 0.058 | 8.66E-14 | Chondrocytes | znhit1 |
| lrig11 | 2.9E-18 | 0.4649722 | 0.143 | 0.075 | 9.42E-14 | Chondrocytes | lrig1 |
| btbd10a | 2.9E-18 | 0.3540427 | 0.141 | 0.073 | 9.57E-14 | Chondrocytes | btbd10a |
| tp531 | 3.1E-18 | 0.3460756 | 0.257 | 0.158 | 9.94E-14 | Chondrocytes | tp53 |
| sec61b2 | 3.1E-18 | 0.2602697 | 0.758 | 0.578 | 1.02E-13 | Chondrocytes | sec61b |
| rerea1 | 3.2E-18 | 0.2422761 | 0.282 | 0.172 | 1.03E-13 | Chondrocytes | rerea |
| smim201 | 3.3E-18 | 0.429703 | 0.149 | 0.079 | 1.08E-13 | Chondrocytes | smim20 |
| dpp72 | 4.1E-18 | 0.2470843 | 0.334 | 0.212 | 1.34E-13 | Chondrocytes | dpp7 |
| ddr2a2 | 4.5E-18 | 0.2838565 | 0.321 | 0.205 | 1.47E-13 | Chondrocytes | ddr2a |
| eif4e3 | 5.4E-18 | 0.4131703 | 0.123 | 0.061 | 1.76E-13 | Chondrocytes | eif4e3 |
| zfr1 | 5.8E-18 | 0.3424368 | 0.194 | 0.111 | 1.88E-13 | Chondrocytes | zfr |
| glrx51 | 6.3E-18 | 0.5660735 | 0.106 | 0.051 | 2.04E-13 | Chondrocytes | glrx5 |
| gnai2b3 | 6.3E-18 | 0.7194614 | 0.122 | 0.062 | 2.06E-13 | Chondrocytes | gnai2b |
| anks1b1 | 6.4E-18 | 0.622536 | 0.118 | 0.059 | 2.08E-13 | Chondrocytes | anks1b |
| ppp1r104 | 6.9E-18 | 0.2057665 | 0.642 | 0.48 | 2.24E-13 | Chondrocytes | ppp1r10 |
| spcs11 | 7.7E-18 | 0.2908078 | 0.602 | 0.432 | 2.5E-13 | Chondrocytes | spcs1 |
| ggps1 | 8.7E-18 | 0.5280872 | 0.115 | 0.057 | 2.82E-13 | Chondrocytes | ggps1 |
| ddx3xa | 8.7E-18 | 0.2126275 | 0.468 | 0.32 | 2.83E-13 | Chondrocytes | ddx3xa |
| hacd21 | 9.5E-18 | 0.2503756 | 0.163 | 0.088 | 3.1E-13 | Chondrocytes | hacd2 |
| uqcrc2b2 | 9.6E-18 | 0.3355504 | 0.195 | 0.112 | 3.12E-13 | Chondrocytes | uqcrc2b |
| zgc:1132081 | 1.1E-17 | 0.2097062 | 0.3 | 0.189 | 3.54E-13 | Chondrocytes | zgc:113208 |
| boc4 | 1.2E-17 | 0.2931981 | 0.522 | 0.369 | 3.95E-13 | Chondrocytes | boc |
| dbpb3 | 1.2E-17 | 0.3236368 | 0.195 | 0.111 | 3.96E-13 | Chondrocytes | dbpb |
| zc3h42 | 1.5E-17 | 0.2319179 | 0.229 | 0.137 | 4.93E-13 | Chondrocytes | zc3h4 |
| ppp3r1b | 1.6E-17 | 0.3399032 | 0.223 | 0.132 | 5.2E-13 | Chondrocytes | ppp3r1b |
| hmgn6 | 1.7E-17 | 0.2505628 | 0.766 | 0.599 | 5.39E-13 | Chondrocytes | hmgn6 |

| | | | | | | | |
|---|---|---|---|---|---|---|---|
| rnf19a | 1.7E-17 | 0.3119965 | 0.263 | 0.162 | 5.5E-13 | Chondrocytes | rnf19a |
| ptges3b1 | 1.8E-17 | 0.2062705 | 0.234 | 0.14 | 5.78E-13 | Chondrocytes | ptges3b |
| glrx31 | 1.8E-17 | 0.2373064 | 0.217 | 0.127 | 5.91E-13 | Chondrocytes | glrx3 |
| tmem1892 | 1.9E-17 | 0.2620974 | 0.22 | 0.129 | 6.15E-13 | Chondrocytes | tmem189 |
| ranbp92 | 2E-17 | 0.3306482 | 0.207 | 0.121 | 6.47E-13 | Chondrocytes | ranbp9 |
| stat6 | 2E-17 | 0.2813289 | 0.14 | 0.073 | 6.52E-13 | Chondrocytes | stat6 |
| tram12 | 2.1E-17 | 0.2774644 | 0.414 | 0.284 | 6.85E-13 | Chondrocytes | tram1 |
| il4r.2 | 2.2E-17 | 0.4297951 | 0.119 | 0.06 | 7.25E-13 | Chondrocytes | il4r.2 |
| si:dkey-27i16.23 | 2.4E-17 | 0.6613172 | 0.194 | 0.116 | 7.74E-13 | Chondrocytes | si:dkey-27i16.2 |
| fntb1 | 2.7E-17 | 0.3080817 | 0.184 | 0.104 | 8.7E-13 | Chondrocytes | fntb |
| lsm12a1 | 2.7E-17 | 0.3200424 | 0.139 | 0.073 | 8.8E-13 | Chondrocytes | lsm12a |
| dixdc1a1 | 2.9E-17 | 0.6437389 | 0.109 | 0.053 | 9.57E-13 | Chondrocytes | dixdc1a |
| arcn1b2 | 3.3E-17 | 0.3280886 | 0.207 | 0.121 | 1.08E-12 | Chondrocytes | arcn1b |
| mtx2 | 3.3E-17 | 0.4356067 | 0.109 | 0.053 | 1.08E-12 | Chondrocytes | mtx2 |
| nudcd31 | 3.4E-17 | 0.4069355 | 0.152 | 0.082 | 1.1E-12 | Chondrocytes | nudcd3 |
| etf1b2 | 3.5E-17 | 0.2718281 | 0.544 | 0.389 | 1.15E-12 | Chondrocytes | etf1b |
| gspt1l1 | 4.2E-17 | 0.2291468 | 0.342 | 0.222 | 1.38E-12 | Chondrocytes | gspt1l |
| ctsf1 | 4.2E-17 | 0.425033 | 0.136 | 0.071 | 1.38E-12 | Chondrocytes | ctsf |
| ppm1ab | 4.7E-17 | 0.2116508 | 0.134 | 0.07 | 1.51E-12 | Chondrocytes | ppm1ab |
| pno12 | 4.9E-17 | 0.2271818 | 0.571 | 0.404 | 1.6E-12 | Chondrocytes | pno1 |
| scamp21 | 5.2E-17 | 0.294536 | 0.205 | 0.12 | 1.68E-12 | Chondrocytes | scamp2 |
| znf3461 | 5.4E-17 | 0.2381021 | 0.164 | 0.09 | 1.74E-12 | Chondrocytes | znf346 |
| ndufb61 | 6.4E-17 | 0.2170672 | 0.285 | 0.179 | 2.07E-12 | Chondrocytes | ndufb6 |
| hectd3 | 6.4E-17 | 0.3620698 | 0.102 | 0.049 | 2.07E-12 | Chondrocytes | hectd3 |
| chchd21 | 6.9E-17 | 0.410513 | 0.165 | 0.092 | 2.25E-12 | Chondrocytes | chchd2 |
| lrrc592 | 7.1E-17 | 0.3242972 | 0.228 | 0.139 | 2.3E-12 | Chondrocytes | lrrc59 |
| tmem2082 | 7.4E-17 | 0.4137168 | 0.125 | 0.064 | 2.41E-12 | Chondrocytes | tmem208 |
| yipf3 | 7.7E-17 | 0.3859769 | 0.134 | 0.07 | 2.51E-12 | Chondrocytes | yipf3 |
| txndc171 | 8.4E-17 | 0.2082036 | 0.201 | 0.117 | 2.74E-12 | Chondrocytes | txndc17 |
| nbas1 | 1.1E-16 | 0.3774611 | 0.171 | 0.097 | 3.43E-12 | Chondrocytes | nbas |
| anapc111 | 1.1E-16 | 0.3630049 | 0.213 | 0.127 | 3.63E-12 | Chondrocytes | anapc11 |
| sod22 | 1.1E-16 | 0.3730463 | 0.212 | 0.127 | 3.71E-12 | Chondrocytes | sod2 |
| smim191 | 1.2E-16 | 0.2937394 | 0.145 | 0.078 | 3.76E-12 | Chondrocytes | smim19 |
| fxr21 | 1.2E-16 | 0.2394918 | 0.163 | 0.09 | 3.94E-12 | Chondrocytes | fxr2 |
| si:ch73-238c9.12 | 1.3E-16 | 0.3494221 | 0.178 | 0.102 | 4.26E-12 | Chondrocytes | si:ch73-238c9.1 |
| hspa4a3 | 1.3E-16 | 0.2573478 | 0.241 | 0.147 | 4.35E-12 | Chondrocytes | hspa4a |
| deptor2 | 1.5E-16 | 0.5163034 | 0.132 | 0.07 | 4.73E-12 | Chondrocytes | deptor |
| zgc:92335 | 1.6E-16 | 0.4261986 | 0.103 | 0.05 | 5.06E-12 | Chondrocytes | zgc:92335 |
| CU638870.13 | 1.7E-16 | 0.2255557 | 0.471 | 0.325 | 5.43E-12 | Chondrocytes | CU638870.1 |
| smarcd11 | 1.7E-16 | 0.456517 | 0.128 | 0.067 | 5.5E-12 | Chondrocytes | smarcd1 |
| rpz22 | 1.7E-16 | 0.365327 | 0.278 | 0.177 | 5.56E-12 | Chondrocytes | rpz2 |
| grb10a3 | 1.9E-16 | 0.2786816 | 0.186 | 0.107 | 6.25E-12 | Chondrocytes | grb10a |
| spag7 | 1.9E-16 | 0.3970479 | 0.117 | 0.059 | 6.28E-12 | Chondrocytes | spag7 |
| ccn14 | 2.2E-16 | 0.4121861 | 0.763 | 0.639 | 7.12E-12 | Chondrocytes | ccn1 |
| wdr431 | 2.4E-16 | 0.3711276 | 0.242 | 0.151 | 7.87E-12 | Chondrocytes | wdr43 |
| plekha3 | 2.6E-16 | 0.4740305 | 0.111 | 0.056 | 8.32E-12 | Chondrocytes | plekha3 |
| snx42 | 2.6E-16 | 0.3970387 | 0.162 | 0.091 | 8.55E-12 | Chondrocytes | snx4 |
| surf4l1 | 2.6E-16 | 0.2563214 | 0.209 | 0.124 | 8.58E-12 | Chondrocytes | surf4l |

| | | | | | | | |
|---|---|---|---|---|---|---|---|
| chaf1a1 | 2.7E-16 | 0.3265518 | 0.106 | 0.052 | 8.73E-12 | Chondrocytes | chaf1a |
| golga3 | 3E-16 | 0.4023599 | 0.128 | 0.067 | 9.77E-12 | Chondrocytes | golga3 |
| ndufb51 | 3.2E-16 | 0.2734573 | 0.289 | 0.186 | 1.06E-11 | Chondrocytes | ndufb5 |
| nfkbib1 | 3.4E-16 | 0.2852906 | 0.192 | 0.112 | 1.1E-11 | Chondrocytes | nfkbib |
| rbbp62 | 4.6E-16 | 0.425573 | 0.438 | 0.314 | 1.48E-11 | Chondrocytes | rbbp6 |
| si:dkeyp-69e1.82 | 4.8E-16 | 0.2815248 | 0.184 | 0.106 | 1.55E-11 | Chondrocytes | si:dkeyp-69e1.8 |
| ist11 | 5.1E-16 | 0.3401612 | 0.139 | 0.075 | 1.67E-11 | Chondrocytes | ist1 |
| etfb1 | 5.2E-16 | 0.2588446 | 0.264 | 0.167 | 1.69E-11 | Chondrocytes | etfb |
| crebrf | 5.2E-16 | 0.4895034 | 0.112 | 0.056 | 1.69E-11 | Chondrocytes | crebrf |
| cdk5rap31 | 5.6E-16 | 0.328304 | 0.107 | 0.053 | 1.83E-11 | Chondrocytes | cdk5rap3 |
| arih21 | 6.6E-16 | 0.3077631 | 0.146 | 0.08 | 2.16E-11 | Chondrocytes | arih2 |
| dlgap4a1 | 7.2E-16 | 0.318811 | 0.162 | 0.091 | 2.35E-11 | Chondrocytes | dlgap4a |
| baxa.11 | 8E-16 | 0.2700376 | 0.175 | 0.1 | 2.59E-11 | Chondrocytes | baxa.1 |
| higd2a1 | 8.1E-16 | 0.3413831 | 0.218 | 0.134 | 2.64E-11 | Chondrocytes | higd2a |
| lsm51 | 9.2E-16 | 0.269386 | 0.196 | 0.116 | 3E-11 | Chondrocytes | lsm5 |
| tlk2 | 9.6E-16 | 0.2826662 | 0.124 | 0.065 | 3.13E-11 | Chondrocytes | tlk2 |
| fryl1 | 1.1E-15 | 0.2374076 | 0.18 | 0.105 | 3.51E-11 | Chondrocytes | fryl |
| uba51 | 1.1E-15 | 0.4492146 | 0.18 | 0.105 | 3.74E-11 | Chondrocytes | uba5 |
| fam3c2 | 1.2E-15 | 0.2131183 | 0.175 | 0.101 | 3.77E-11 | Chondrocytes | fam3c |
| ccdc32 | 1.2E-15 | 0.4648713 | 0.117 | 0.06 | 3.94E-11 | Chondrocytes | ccdc32 |
| fdps | 1.2E-15 | 0.628962 | 0.115 | 0.059 | 3.97E-11 | Chondrocytes | fdps |
| rgs3a2 | 1.2E-15 | 0.3553334 | 0.162 | 0.092 | 4.01E-11 | Chondrocytes | rgs3a |
| otud5b1 | 1.7E-15 | 0.4115023 | 0.149 | 0.083 | 5.45E-11 | Chondrocytes | otud5b |
| zfand2a2 | 1.7E-15 | 0.4491027 | 0.125 | 0.066 | 5.46E-11 | Chondrocytes | zfand2a |
| lamtor52 | 2E-15 | 0.2646531 | 0.154 | 0.086 | 6.59E-11 | Chondrocytes | lamtor5 |
| dusp25 | 2.3E-15 | 0.4243698 | 0.319 | 0.22 | 7.45E-11 | Chondrocytes | dusp2 |
| ilf21 | 2.3E-15 | 0.2589879 | 0.251 | 0.158 | 7.57E-11 | Chondrocytes | ilf2 |
| march6 | 2.4E-15 | 0.255112 | 0.174 | 0.101 | 7.69E-11 | Chondrocytes | march6 |
| rnf126 | 2.5E-15 | 0.3132515 | 0.138 | 0.075 | 8.21E-11 | Chondrocytes | rnf126 |
| abce11 | 3.5E-15 | 0.2544332 | 0.202 | 0.121 | 1.14E-10 | Chondrocytes | abce1 |
| FAM83G | 3.5E-15 | 0.3806342 | 0.115 | 0.06 | 1.15E-10 | Chondrocytes | FAM83G |
| atg13 | 3.6E-15 | 0.3925537 | 0.14 | 0.078 | 1.16E-10 | Chondrocytes | atg13 |
| kif5ba.1 | 3.8E-15 | 0.286362 | 0.169 | 0.097 | 1.23E-10 | Chondrocytes | kif5ba.1 |
| fnta1 | 4.1E-15 | 0.3473168 | 0.126 | 0.067 | 1.35E-10 | Chondrocytes | fnta |
| klf7a1 | 4.2E-15 | 0.2427647 | 0.11 | 0.056 | 1.35E-10 | Chondrocytes | klf7a |
| eif3f3 | 4.5E-15 | 0.2386135 | 0.842 | 0.726 | 1.46E-10 | Chondrocytes | eif3f |
| smap11 | 4.6E-15 | 0.2205014 | 0.245 | 0.153 | 1.49E-10 | Chondrocytes | smap1 |
| polr3glb | 4.8E-15 | 0.2656127 | 0.118 | 0.061 | 1.55E-10 | Chondrocytes | polr3glb |
| ckap42 | 5E-15 | 0.2069666 | 0.675 | 0.515 | 1.61E-10 | Chondrocytes | ckap4 |
| zbtb8os1 | 5E-15 | 0.2876026 | 0.134 | 0.073 | 1.61E-10 | Chondrocytes | zbtb8os |
| lamtor22 | 5E-15 | 0.2844297 | 0.197 | 0.119 | 1.64E-10 | Chondrocytes | lamtor2 |
| nsfl1c1 | 5.6E-15 | 0.2391159 | 0.206 | 0.124 | 1.82E-10 | Chondrocytes | nsfl1c |
| kmt2e | 6E-15 | 0.2760939 | 0.143 | 0.079 | 1.96E-10 | Chondrocytes | kmt2e |
| hyal11 | 6.7E-15 | 0.5024644 | 0.106 | 0.055 | 2.18E-10 | Chondrocytes | hyal1 |
| rab411 | 7.2E-15 | 0.3325303 | 0.144 | 0.081 | 2.34E-10 | Chondrocytes | rab41 |
| CU464124.12 | 7.6E-15 | 0.3034804 | 0.175 | 0.103 | 2.46E-10 | Chondrocytes | CU464124.1 |
| spry23 | 7.8E-15 | 0.2278692 | 0.14 | 0.077 | 2.55E-10 | Chondrocytes | spry2 |
| hk21 | 9E-15 | 0.3036928 | 0.102 | 0.051 | 2.93E-10 | Chondrocytes | hk2 |

| | | | | | | | |
|---|---|---|---|---|---|---|---|
| ip6k2a3 | 9.2E-15 | 0.2658471 | 0.296 | 0.195 | 3E-10 | Chondrocytes | ip6k2a |
| atxn2l1 | 9.4E-15 | 0.2719079 | 0.158 | 0.091 | 3.05E-10 | Chondrocytes | atxn2l |
| atoh81 | 1E-14 | 0.3842191 | 0.116 | 0.061 | 3.4E-10 | Chondrocytes | atoh8 |
| uso11 | 1.1E-14 | 0.2484314 | 0.193 | 0.117 | 3.47E-10 | Chondrocytes | uso1 |
| atg4a | 1.1E-14 | 0.4896317 | 0.115 | 0.061 | 3.67E-10 | Chondrocytes | atg4a |
| sec131 | 1.1E-14 | 0.2338313 | 0.204 | 0.124 | 3.68E-10 | Chondrocytes | sec13 |
| vta11 | 1.3E-14 | 0.4252648 | 0.134 | 0.074 | 4.19E-10 | Chondrocytes | vta1 |
| bcl2l102 | 1.3E-14 | 0.3783899 | 0.271 | 0.178 | 4.25E-10 | Chondrocytes | bcl2l10 |
| zmat51 | 1.5E-14 | 0.3789963 | 0.133 | 0.073 | 4.8E-10 | Chondrocytes | zmat5 |
| kat2b | 1.5E-14 | 0.385545 | 0.101 | 0.051 | 4.82E-10 | Chondrocytes | kat2b |
| cemip21 | 1.6E-14 | 0.3668073 | 0.131 | 0.071 | 5.08E-10 | Chondrocytes | cemip2 |
| pycr1b2 | 1.6E-14 | 0.4350233 | 0.103 | 0.053 | 5.16E-10 | Chondrocytes | pycr1b |
| cul1b | 1.9E-14 | 0.3418266 | 0.104 | 0.053 | 6.07E-10 | Chondrocytes | cul1b |
| btbd6b1 | 1.9E-14 | 0.4038316 | 0.112 | 0.059 | 6.17E-10 | Chondrocytes | btbd6b |
| pebp13 | 2E-14 | 0.353961 | 0.131 | 0.071 | 6.67E-10 | Chondrocytes | pebp1 |
| si:ch211-195b15.83 | 2.5E-14 | 0.3365821 | 0.155 | 0.089 | 8.08E-10 | Chondrocytes | si:ch211-195b15.8 |
| ndufv21 | 2.6E-14 | 0.2574569 | 0.176 | 0.104 | 8.53E-10 | Chondrocytes | ndufv2 |
| ubr71 | 2.8E-14 | 0.3183099 | 0.159 | 0.093 | 9.09E-10 | Chondrocytes | ubr7 |
| zgc:1949813 | 3.1E-14 | 0.3964128 | 0.142 | 0.08 | 9.99E-10 | Chondrocytes | zgc:194981 |
| arhgap11 | 3.1E-14 | 0.288003 | 0.14 | 0.079 | 1.01E-09 | Chondrocytes | arhgap1 |
| cox6b12 | 5.1E-14 | 0.5713637 | 0.125 | 0.069 | 1.65E-09 | Chondrocytes | cox6b1 |
| btg32 | 5.4E-14 | 0.3250645 | 0.201 | 0.125 | 1.75E-09 | Chondrocytes | btg3 |
| derl11 | 5.8E-14 | 0.2592236 | 0.129 | 0.071 | 1.87E-09 | Chondrocytes | derl1 |
| psat11 | 6.8E-14 | 0.4286307 | 0.124 | 0.068 | 2.22E-09 | Chondrocytes | psat1 |
| zgc:1034382 | 7.2E-14 | 0.3215879 | 0.152 | 0.087 | 2.35E-09 | Chondrocytes | zgc:103438 |
| abcc6a3 | 7.5E-14 | 0.9019354 | 0.131 | 0.075 | 2.45E-09 | Chondrocytes | abcc6a |
| rida1 | 9.3E-14 | 0.3452396 | 0.141 | 0.08 | 3.04E-09 | Chondrocytes | rida |
| tgfbr1b2 | 9.4E-14 | 0.2410995 | 0.186 | 0.113 | 3.05E-09 | Chondrocytes | tgfbr1b |
| CIR1 | 1.1E-13 | 0.33505 | 0.143 | 0.082 | 3.61E-09 | Chondrocytes | CIR1 |
| nudt9 | 1.3E-13 | 0.5093909 | 0.1 | 0.052 | 4.11E-09 | Chondrocytes | nudt9 |
| pdzrn3b1 | 1.4E-13 | 0.2313591 | 0.138 | 0.078 | 4.39E-09 | Chondrocytes | pdzrn3b |
| ubl3b | 1.4E-13 | 0.445634 | 0.109 | 0.058 | 4.61E-09 | Chondrocytes | ubl3b |
| lsm81 | 1.5E-13 | 0.2838643 | 0.143 | 0.081 | 4.78E-09 | Chondrocytes | lsm8 |
| p3h12 | 1.8E-13 | 0.2109477 | 0.315 | 0.211 | 5.78E-09 | Chondrocytes | p3h1 |
| ebag91 | 1.8E-13 | 0.2768922 | 0.132 | 0.074 | 6.01E-09 | Chondrocytes | ebag9 |
| senp2 | 1.9E-13 | 0.2137808 | 0.146 | 0.083 | 6.33E-09 | Chondrocytes | senp2 |
| ptpn11b1 | 2.1E-13 | 0.4228707 | 0.116 | 0.063 | 6.8E-09 | Chondrocytes | ptpn11b |
| nnt1 | 2.3E-13 | 0.289999 | 0.11 | 0.059 | 7.51E-09 | Chondrocytes | nnt |
| pcna1 | 2.4E-13 | 0.2421352 | 0.18 | 0.109 | 7.96E-09 | Chondrocytes | pcna |
| blmh1 | 2.8E-13 | 0.3579173 | 0.108 | 0.058 | 9.04E-09 | Chondrocytes | blmh |
| slc40a13 | 2.8E-13 | 0.4526241 | 0.292 | 0.2 | 9.24E-09 | Chondrocytes | slc40a1 |
| eea1 | 2.9E-13 | 0.2303314 | 0.131 | 0.074 | 9.46E-09 | Chondrocytes | eea1 |
| atp6v1h | 3.1E-13 | 0.4001071 | 0.106 | 0.056 | 1E-08 | Chondrocytes | atp6v1h |
| copg21 | 3.1E-13 | 0.2270569 | 0.231 | 0.148 | 1.01E-08 | Chondrocytes | copg2 |
| atp6v0d1 | 3.5E-13 | 0.2075454 | 0.133 | 0.075 | 1.13E-08 | Chondrocytes | atp6v0d1 |
| lnpep1 | 3.5E-13 | 0.2703023 | 0.128 | 0.071 | 1.15E-08 | Chondrocytes | lnpep |
| rab40b2 | 3.6E-13 | 0.3042964 | 0.136 | 0.077 | 1.17E-08 | Chondrocytes | rab40b |
| nck1a3 | 4.5E-13 | 0.2267839 | 0.137 | 0.078 | 1.46E-08 | Chondrocytes | nck1a |

| | | | | | | | |
|---|---|---|---|---|---|---|---|
| zgc:1533723 | 4.7E-13 | 0.2551869 | 0.147 | 0.086 | 1.53E-08 | Chondrocytes | zgc:153372 |
| vps26bl | 5.2E-13 | 0.410357 | 0.113 | 0.062 | 1.69E-08 | Chondrocytes | vps26bl |
| cd2762 | 6E-13 | 0.3703911 | 0.206 | 0.131 | 1.95E-08 | Chondrocytes | cd276 |
| cnot6a1 | 6.7E-13 | 0.2011653 | 0.155 | 0.091 | 2.18E-08 | Chondrocytes | cnot6a |
| eaf1 | 6.9E-13 | 0.3810294 | 0.118 | 0.066 | 2.26E-08 | Chondrocytes | eaf1 |
| nenf1 | 7.3E-13 | 0.2737493 | 0.16 | 0.096 | 2.37E-08 | Chondrocytes | nenf |
| mxd41 | 7.4E-13 | 0.3670596 | 0.152 | 0.09 | 2.41E-08 | Chondrocytes | mxd4 |
| apbb1 | 7.5E-13 | 0.3419698 | 0.103 | 0.055 | 2.44E-08 | Chondrocytes | apbb1 |
| eif3d2 | 9E-13 | 0.203912 | 0.568 | 0.417 | 2.92E-08 | Chondrocytes | eif3d |
| rbms2b | 1.2E-12 | 0.2825621 | 0.112 | 0.062 | 3.79E-08 | Chondrocytes | rbms2b |
| immt1 | 1.3E-12 | 0.2679194 | 0.149 | 0.088 | 4.22E-08 | Chondrocytes | immt |
| eci21 | 1.7E-12 | 0.3931483 | 0.115 | 0.064 | 5.53E-08 | Chondrocytes | eci2 |
| ccdc1672 | 1.7E-12 | 0.456738 | 0.115 | 0.064 | 5.57E-08 | Chondrocytes | ccdc167 |
| ralab | 1.9E-12 | 0.2133847 | 0.15 | 0.089 | 6.25E-08 | Chondrocytes | ralab |
| plekha81 | 1.9E-12 | 0.2289614 | 0.124 | 0.07 | 6.32E-08 | Chondrocytes | plekha8 |
| picalma1 | 2E-12 | 0.2901983 | 0.129 | 0.074 | 6.53E-08 | Chondrocytes | picalma |
| ywhah1 | 2.2E-12 | 0.3154115 | 0.239 | 0.159 | 7.04E-08 | Chondrocytes | ywhah |
| ccdc50 | 2.4E-12 | 0.4182469 | 0.111 | 0.061 | 7.7E-08 | Chondrocytes | ccdc50 |
| mrc23 | 2.7E-12 | 0.2487223 | 0.147 | 0.087 | 8.77E-08 | Chondrocytes | mrc2 |
| tiparp3 | 2.8E-12 | 0.5895783 | 0.192 | 0.124 | 9E-08 | Chondrocytes | tiparp |
| stx181 | 3E-12 | 0.3408313 | 0.1 | 0.054 | 9.64E-08 | Chondrocytes | stx18 |
| cyb5a | 3.1E-12 | 0.5625297 | 0.15 | 0.089 | 9.95E-08 | Chondrocytes | cyb5a |
| cog41 | 3.2E-12 | 0.3275287 | 0.103 | 0.055 | 1.04E-07 | Chondrocytes | cog4 |
| dcun1d51 | 3.4E-12 | 0.3232292 | 0.128 | 0.073 | 1.12E-07 | Chondrocytes | dcun1d5 |
| crsp72 | 4.6E-12 | 0.2883785 | 0.125 | 0.072 | 1.5E-07 | Chondrocytes | crsp7 |
| cnppd1 | 4.6E-12 | 0.2393854 | 0.129 | 0.074 | 1.5E-07 | Chondrocytes | cnppd1 |
| tns2a1 | 4.8E-12 | 0.3693605 | 0.107 | 0.059 | 1.57E-07 | Chondrocytes | tns2a |
| ppdpfb2 | 5.2E-12 | 0.7239131 | 0.857 | 0.776 | 1.7E-07 | Chondrocytes | ppdpfb |
| gosr21 | 6.5E-12 | 0.2610084 | 0.15 | 0.09 | 2.11E-07 | Chondrocytes | gosr2 |
| pgam51 | 7.3E-12 | 0.232231 | 0.127 | 0.073 | 2.36E-07 | Chondrocytes | pgam5 |
| taf11 | 7.3E-12 | 0.3263001 | 0.102 | 0.056 | 2.39E-07 | Chondrocytes | taf11 |
| AL732488.2 | 7.7E-12 | 0.2697286 | 0.112 | 0.063 | 2.51E-07 | Chondrocytes | AL732488.2 |
| ift46 | 8.8E-12 | 0.2738551 | 0.106 | 0.059 | 2.87E-07 | Chondrocytes | ift46 |
| rbbp41 | 8.9E-12 | 0.220082 | 0.152 | 0.092 | 2.91E-07 | Chondrocytes | rbbp4 |
| kmt5aa2 | 1.1E-11 | 0.2136955 | 0.18 | 0.113 | 3.7E-07 | Chondrocytes | kmt5aa |
| adrm11 | 1.4E-11 | 0.3045691 | 0.147 | 0.089 | 4.7E-07 | Chondrocytes | adrm1 |
| actr10 | 1.5E-11 | 0.3276834 | 0.116 | 0.066 | 4.94E-07 | Chondrocytes | actr10 |
| ciapin11 | 2.1E-11 | 0.3925178 | 0.108 | 0.061 | 6.85E-07 | Chondrocytes | ciapin1 |
| tmed5 | 2.2E-11 | 0.2123993 | 0.108 | 0.06 | 7.14E-07 | Chondrocytes | tmed5 |
| mrpl412 | 3.2E-11 | 0.3704923 | 0.138 | 0.084 | 1.04E-06 | Chondrocytes | mrpl41 |
| akirin21 | 3.6E-11 | 0.2079694 | 0.119 | 0.069 | 1.17E-06 | Chondrocytes | akirin2 |
| smarcc1a1 | 4E-11 | 0.2648477 | 0.123 | 0.072 | 1.3E-06 | Chondrocytes | smarcc1a |
| ppardb | 4.4E-11 | 0.3489263 | 0.103 | 0.058 | 1.42E-06 | Chondrocytes | ppardb |
| fndc3a1 | 4.7E-11 | 0.2423102 | 0.137 | 0.082 | 1.54E-06 | Chondrocytes | fndc3a |
| cmc12 | 4.8E-11 | 0.2453126 | 0.155 | 0.096 | 1.55E-06 | Chondrocytes | cmc1 |
| plecb2 | 5.7E-11 | 0.2908752 | 0.762 | 0.655 | 1.86E-06 | Chondrocytes | plecb |
| sec22a | 5.9E-11 | 0.4134321 | 0.1 | 0.056 | 1.93E-06 | Chondrocytes | sec22a |
| aldob6 | 6E-11 | 0.315013 | 0.42 | 0.309 | 1.95E-06 | Chondrocytes | aldob |

| | | | | | | | |
|---|---|---|---|---|---|---|---|
| pik3r3b2 | 8E-11 | 0.293695 | 0.369 | 0.272 | 2.61E-06 | Chondrocytes | pik3r3b |
| itgb53 | 8.6E-11 | 0.2137522 | 0.138 | 0.083 | 2.81E-06 | Chondrocytes | itgb5 |
| ryr34 | 1.2E-10 | 0.928594 | 0.14 | 0.089 | 3.87E-06 | Chondrocytes | ryr3 |
| nfic1 | 1.3E-10 | 0.2410916 | 0.13 | 0.078 | 4.2E-06 | Chondrocytes | nfic |
| tmub1 | 1.4E-10 | 0.2042016 | 0.118 | 0.069 | 4.52E-06 | Chondrocytes | tmub1 |
| tcea22 | 1.5E-10 | 0.288093 | 0.128 | 0.077 | 4.9E-06 | Chondrocytes | tcea2 |
| ypel5 | 1.5E-10 | 0.2792386 | 0.109 | 0.063 | 4.97E-06 | Chondrocytes | ypel5 |
| terfa1 | 1.6E-10 | 0.2345386 | 0.143 | 0.088 | 5.08E-06 | Chondrocytes | terfa |
| EMC41 | 1.6E-10 | 0.2179429 | 0.103 | 0.058 | 5.18E-06 | Chondrocytes | EMC4 |
| map4k5 | 1.7E-10 | 0.3004942 | 0.1 | 0.056 | 5.63E-06 | Chondrocytes | map4k5 |
| acp11 | 2.6E-10 | 0.3189841 | 0.126 | 0.076 | 8.29E-06 | Chondrocytes | acp1 |
| sdf21 | 4.1E-10 | 0.2338426 | 0.101 | 0.058 | 1.33E-05 | Chondrocytes | sdf2 |
| vps26c1 | 4.3E-10 | 0.2135845 | 0.125 | 0.075 | 1.39E-05 | Chondrocytes | vps26c |
| nr4a13 | 5E-10 | 0.2538653 | 0.158 | 0.101 | 1.63E-05 | Chondrocytes | nr4a1 |
| dlx2a | 6E-10 | 0.2472141 | 0.111 | 0.065 | 1.95E-05 | Chondrocytes | dlx2a |
| golt1bb2 | 7.4E-10 | 0.2058671 | 0.156 | 0.1 | 2.4E-05 | Chondrocytes | golt1bb |
| fscn1a4 | 7.7E-10 | 0.2422943 | 0.252 | 0.177 | 2.5E-05 | Chondrocytes | fscn1a |
| pbrm1l | 7.8E-10 | 0.2604638 | 0.101 | 0.058 | 2.55E-05 | Chondrocytes | pbrm1l |
| fgfr42 | 8.1E-10 | 0.5970589 | 0.122 | 0.076 | 2.64E-05 | Chondrocytes | fgfr4 |
| C25H11orf965 | 8.7E-10 | 0.6125911 | 0.457 | 0.382 | 2.83E-05 | Chondrocytes | C25H11orf96 |
| ubl7b1 | 8.8E-10 | 0.2771352 | 0.123 | 0.075 | 2.85E-05 | Chondrocytes | ubl7b |
| pkdcca1 | 9.9E-10 | 0.2426126 | 0.115 | 0.069 | 3.2E-05 | Chondrocytes | pkdcca |
| prr13 | 1.2E-09 | 0.226864 | 0.104 | 0.061 | 3.96E-05 | Chondrocytes | prr13 |
| ap3s21 | 1.3E-09 | 0.2997744 | 0.105 | 0.062 | 4.23E-05 | Chondrocytes | ap3s2 |
| eif2b22 | 1.5E-09 | 0.2795763 | 0.115 | 0.07 | 4.98E-05 | Chondrocytes | eif2b2 |
| c1galt1c11 | 2.4E-09 | 0.2420434 | 0.114 | 0.069 | 7.86E-05 | Chondrocytes | c1galt1c1 |
| pop72 | 3E-09 | 0.3514636 | 0.113 | 0.069 | 9.76E-05 | Chondrocytes | pop7 |
| lum3 | 3.3E-09 | 0.4340341 | 0.683 | 0.762 | 0.000108 | Chondrocytes | lum |
| nat91 | 4.3E-09 | 0.2169792 | 0.137 | 0.087 | 0.000141 | Chondrocytes | nat9 |
| eprs12 | 4.6E-09 | 0.2468511 | 0.144 | 0.093 | 0.000149 | Chondrocytes | eprs1 |
| sgcd | 6.3E-09 | 0.300171 | 0.109 | 0.066 | 0.000204 | Chondrocytes | sgcd |
| clcn3 | 1.7E-08 | 0.2410398 | 0.109 | 0.067 | 0.000558 | Chondrocytes | clcn3 |
| abi3bpb3 | 2.6E-08 | 0.2956615 | 0.475 | 0.59 | 0.00083 | Chondrocytes | abi3bpb |
| egr2a3 | 2.7E-08 | 0.3177184 | 0.143 | 0.094 | 0.00087 | Chondrocytes | egr2a |
| scarb2a2 | 2.8E-08 | 0.2387214 | 0.262 | 0.187 | 0.000919 | Chondrocytes | scarb2a |
| gsc3 | 3.1E-08 | 0.3074716 | 0.169 | 0.116 | 0.000999 | Chondrocytes | gsc |
| ergic31 | 3.3E-08 | 0.2348415 | 0.1 | 0.06 | 0.001085 | Chondrocytes | ergic3 |
| tmem106a1 | 3.4E-08 | 0.2306927 | 0.121 | 0.076 | 0.001112 | Chondrocytes | tmem106a |
| cdc42ep4b4 | 5.6E-08 | 0.2104778 | 0.131 | 0.085 | 0.001818 | Chondrocytes | cdc42ep4b |
| si:dkeyp-55f12.31 | 2E-07 | 0.2377664 | 0.106 | 0.067 | 0.00653 | Chondrocytes | si:dkeyp-55f12.3 |
| mta31 | 2.5E-07 | 0.2287791 | 0.106 | 0.068 | 0.008228 | Chondrocytes | mta3 |
| nfkbiab4 | 1.2E-06 | 0.3714737 | 0.66 | 0.582 | 0.038448 | Chondrocytes | nfkbiab |
| nedd8l1 | 1.9E-06 | 0.2845441 | 0.103 | 0.068 | 0.060291 | Chondrocytes | nedd8l |
| id33 | 3.2E-06 | 0.2087111 | 0.603 | 0.51 | 0.10499 | Chondrocytes | id3 |
| nfkb22 | 8.8E-06 | 0.2076197 | 0.14 | 0.1 | 0.285279 | Chondrocytes | nfkb2 |
| clu3 | 5.7E-05 | 0.4940104 | 0.438 | 0.383 | 1 | Chondrocytes | clu |
| fabp35 | 0.00019 | 0.2518854 | 0.604 | 0.529 | 1 | Chondrocytes | fabp3 |
| ogna2 | 0.00149 | 0.380717 | 0.231 | 0.189 | 1 | Chondrocytes | ogna |

| | | | | | | | |
|---|---|---|---|---|---|---|---|
| dkk3b3 | 0 | 3.5944598 | 0.918 | 0.23 | 0 | Chondrogenic Fibroblasts | dkk3b |
| MYADM.12 | 0 | 3.2087742 | 0.894 | 0.212 | 0 | Chondrogenic Fibroblasts | MYADM.1 |
| bcan2 | 0 | 3.7972584 | 0.722 | 0.085 | 0 | Chondrogenic Fibroblasts | bcan |
| olfml2bb3 | 0 | 4.1183001 | 0.704 | 0.074 | 0 | Chondrogenic Fibroblasts | olfml2bb |
| si:ch211-156j16.14 | 0 | 2.9424092 | 0.721 | 0.137 | 0 | Chondrogenic Fibroblasts | si:ch211-156j16.1 |
| thbs2b4 | 0 | 2.5615869 | 0.819 | 0.236 | 0 | Chondrogenic Fibroblasts | thbs2b |
| sgcg3 | 0 | 3.4282711 | 0.652 | 0.076 | 0 | Chondrogenic Fibroblasts | sgcg |
| aspn3 | 0 | 3.2311861 | 0.956 | 0.389 | 0 | Chondrogenic Fibroblasts | aspn |
| fn1b3 | 0 | 1.7870677 | 0.939 | 0.387 | 0 | Chondrogenic Fibroblasts | fn1b |
| thbs3a3 | 0 | 2.3090091 | 0.829 | 0.295 | 0 | Chondrogenic Fibroblasts | thbs3a |
| crip22 | 0 | 4.2189171 | 0.582 | 0.063 | 0 | Chondrogenic Fibroblasts | crip2 |
| ogna3 | 0 | 2.4674205 | 0.684 | 0.176 | 0 | Chondrogenic Fibroblasts | ogna |
| fibinb3 | 0 | 2.4357335 | 0.766 | 0.261 | 0 | Chondrogenic Fibroblasts | fibinb |
| loxa5 | 0 | 2.5108296 | 0.664 | 0.161 | 0 | Chondrogenic Fibroblasts | loxa |
| arpc1b2 | 0 | 2.3125556 | 0.731 | 0.235 | 0 | Chondrogenic Fibroblasts | arpc1b |
| ccdc803 | 0 | 2.476354 | 0.865 | 0.374 | 0 | Chondrogenic Fibroblasts | ccdc80 |
| col8a1b2 | 0 | 2.2147532 | 0.616 | 0.129 | 0 | Chondrogenic Fibroblasts | col8a1b |
| thbs4b5 | 0 | 2.1561793 | 0.725 | 0.247 | 0 | Chondrogenic Fibroblasts | thbs4b |
| thbs4a3 | 0 | 3.6940684 | 0.543 | 0.067 | 0 | Chondrogenic Fibroblasts | thbs4a |
| itga11a4 | 0 | 2.9215888 | 0.558 | 0.098 | 0 | Chondrogenic Fibroblasts | itga11a |
| hivep13 | 0 | 2.7277803 | 0.597 | 0.142 | 0 | Chondrogenic Fibroblasts | hivep1 |
| col16a14 | 0 | 2.0077503 | 0.897 | 0.444 | 0 | Chondrogenic Fibroblasts | col16a1 |
| si:ch211-1a19.32 | 0 | 2.2854738 | 0.954 | 0.504 | 0 | Chondrogenic Fibroblasts | si:ch211-1a19.3 |
| rabac13 | 0 | 1.880523 | 0.832 | 0.385 | 0 | Chondrogenic Fibroblasts | rabac1 |
| angptl1a3 | 0 | 2.0031462 | 0.827 | 0.381 | 0 | Chondrogenic Fibroblasts | angptl1a |
| angptl2b3 | 0 | 2.5656076 | 0.564 | 0.129 | 0 | Chondrogenic Fibroblasts | angptl2b |
| fndc14 | 0 | 2.1189741 | 0.588 | 0.156 | 0 | Chondrogenic Fibroblasts | fndc1 |
| prss353 | 0 | 3.0439884 | 0.503 | 0.082 | 0 | Chondrogenic Fibroblasts | prss35 |
| si:ch211-222l21.13 | 0 | 2.2126784 | 0.965 | 0.544 | 0 | Chondrogenic Fibroblasts | si:ch211-222l21.1 |
| si:busm1-57f23.13 | 0 | 1.9556268 | 0.98 | 0.564 | 0 | Chondrogenic Fibroblasts | si:busm1-57f23.1 |
| si:dkey-27i16.24 | 0 | 2.5204813 | 0.509 | 0.107 | 0 | Chondrogenic Fibroblasts | si:dkey-27i16.2 |
| tex264a1 | 0 | 2.6938185 | 0.512 | 0.111 | 0 | Chondrogenic Fibroblasts | tex264a |
| col11a1b4 | 0 | 1.9034642 | 0.925 | 0.533 | 0 | Chondrogenic Fibroblasts | col11a1b |
| angptl1b2 | 0 | 3.3369384 | 0.441 | 0.055 | 0 | Chondrogenic Fibroblasts | angptl1b |
| fxyd13 | 0 | 1.9769391 | 0.938 | 0.563 | 0 | Chondrogenic Fibroblasts | fxyd1 |
| abi3bpb4 | 0 | 2.8040929 | 0.947 | 0.576 | 0 | Chondrogenic Fibroblasts | abi3bpb |
| col12a1b3 | 0 | 1.9407732 | 0.964 | 0.615 | 0 | Chondrogenic Fibroblasts | col12a1b |
| sox63 | 0 | 2.1571329 | 0.436 | 0.094 | 0 | Chondrogenic Fibroblasts | sox6 |
| cpne22 | 0 | 3.035056 | 0.394 | 0.058 | 0 | Chondrogenic Fibroblasts | cpne2 |
| ckba2 | 0 | 2.5607211 | 0.967 | 0.632 | 0 | Chondrogenic Fibroblasts | ckba |
| lgals2a3 | 0 | 2.8079512 | 0.96 | 0.633 | 0 | Chondrogenic Fibroblasts | lgals2a |
| fhod13 | 0 | 2.4918044 | 0.391 | 0.074 | 0 | Chondrogenic Fibroblasts | fhod1 |
| col15a1b1 | 0 | 3.2327754 | 0.344 | 0.042 | 0 | Chondrogenic Fibroblasts | col15a1b |
| BX511021.21 | 0 | 2.9386883 | 0.297 | 0.04 | 0 | Chondrogenic Fibroblasts | BX511021.2 |
| TNC | 0 | 3.0030093 | 0.284 | 0.032 | 0 | Chondrogenic Fibroblasts | TNC |
| ccn4b1 | 0 | 2.8695144 | 0.296 | 0.045 | 0 | Chondrogenic Fibroblasts | ccn4b |
| col12a1a3 | 0 | 1.7099694 | 0.992 | 0.746 | 0 | Chondrogenic Fibroblasts | col12a1a |
| tnfaip8l32 | 0 | 2.9900779 | 0.285 | 0.04 | 0 | Chondrogenic Fibroblasts | tnfaip8l3 |

| | | | | | | | |
|---|---|---|---|---|---|---|---|
| mmp23 | 0 | 1.5380724 | 0.979 | 0.738 | 0 | Chondrogenic Fibroblasts | mmp2 |
| timp2a5 | 0 | 1.553198 | 0.981 | 0.749 | 0 | Chondrogenic Fibroblasts | timp2a |
| msna2 | 0 | 1.5612705 | 0.976 | 0.745 | 0 | Chondrogenic Fibroblasts | msna |
| alpi.11 | 0 | 3.7333024 | 0.248 | 0.02 | 0 | Chondrogenic Fibroblasts | alpi.1 |
| col5a2a3 | 0 | 1.5001006 | 0.981 | 0.787 | 0 | Chondrogenic Fibroblasts | col5a2a |
| si:ch1073-459j12.14 | 0 | 1.4383557 | 0.988 | 0.807 | 0 | Chondrogenic Fibroblasts | si:ch1073-459j12.1 |
| pdgfd | 0 | 3.4547153 | 0.174 | 0.015 | 0 | Chondrogenic Fibroblasts | pdgfd |
| krt18a.14 | 0 | 1.6592691 | 0.994 | 0.838 | 0 | Chondrogenic Fibroblasts | krt18a.1 |
| mpp2b | 0 | 4.2196757 | 0.144 | 0.008 | 0 | Chondrogenic Fibroblasts | mpp2b |
| slc8a1a | 0 | 3.832831 | 0.144 | 0.009 | 0 | Chondrogenic Fibroblasts | slc8a1a |
| anxa2a3 | 0 | 1.7640861 | 0.997 | 0.888 | 0 | Chondrogenic Fibroblasts | anxa2a |
| pleca2 | 0 | 1.4006885 | 0.993 | 0.916 | 0 | Chondrogenic Fibroblasts | pleca |
| col5a14 | 0 | 1.2049317 | 0.997 | 0.964 | 0 | Chondrogenic Fibroblasts | col5a1 |
| col1a1b4 | 0 | 1.3121316 | 0.998 | 0.986 | 0 | Chondrogenic Fibroblasts | col1a1b |
| col1a24 | 0 | 1.4544446 | 1 | 0.991 | 0 | Chondrogenic Fibroblasts | col1a2 |
| col1a1a4 | 4E-302 | 1.2735529 | 1 | 0.979 | 1.2E-297 | Chondrogenic Fibroblasts | col1a1a |
| sparc4 | 6E-301 | 1.3606442 | 0.998 | 0.98 | 2.1E-296 | Chondrogenic Fibroblasts | sparc |
| elovl1b2 | 8E-295 | 1.8912208 | 0.779 | 0.34 | 2.5E-290 | Chondrogenic Fibroblasts | elovl1b |
| si:ch211-176g6.21 | 2E-293 | 3.2663033 | 0.18 | 0.018 | 6.6E-289 | Chondrogenic Fibroblasts | si:ch211-176g6.2 |
| gsna1 | 1E-288 | 1.8909063 | 0.784 | 0.347 | 3.1E-284 | Chondrogenic Fibroblasts | gsna |
| ehd2b2 | 9E-287 | 1.6538864 | 0.917 | 0.551 | 3.1E-282 | Chondrogenic Fibroblasts | ehd2b |
| nfatc3a2 | 2E-283 | 2.3994783 | 0.394 | 0.085 | 6.5E-279 | Chondrogenic Fibroblasts | nfatc3a |
| krt43 | 9E-278 | 1.683791 | 0.966 | 0.753 | 2.9E-273 | Chondrogenic Fibroblasts | krt4 |
| CR293509.1 | 3E-276 | 4.2667696 | 0.142 | 0.012 | 1.1E-271 | Chondrogenic Fibroblasts | CR293509.1 |
| hsd3b71 | 7E-276 | 1.7659317 | 0.54 | 0.149 | 2.2E-271 | Chondrogenic Fibroblasts | hsd3b7 |
| pdcd62 | 2E-275 | 1.8498891 | 0.85 | 0.467 | 5.7E-271 | Chondrogenic Fibroblasts | pdcd6 |
| cgref13 | 2E-275 | 1.9086131 | 0.557 | 0.163 | 7.1E-271 | Chondrogenic Fibroblasts | cgref1 |
| cav23 | 3E-275 | 2.0829179 | 0.549 | 0.162 | 8.3E-271 | Chondrogenic Fibroblasts | cav2 |
| alcama3 | 1E-274 | 2.2510506 | 0.348 | 0.068 | 3.9E-270 | Chondrogenic Fibroblasts | alcama |
| crabp2a3 | 2E-268 | 1.7738176 | 0.54 | 0.152 | 5.9E-264 | Chondrogenic Fibroblasts | crabp2a |
| sgsm31 | 2E-266 | 3.5871418 | 0.122 | 0.009 | 7.5E-262 | Chondrogenic Fibroblasts | sgsm3 |
| lum4 | 1E-261 | 1.8329383 | 0.948 | 0.754 | 3.4E-257 | Chondrogenic Fibroblasts | lum |
| col6a15 | 5E-260 | 1.1784719 | 0.991 | 0.872 | 1.7E-255 | Chondrogenic Fibroblasts | col6a1 |
| cd81a2 | 8E-253 | 1.1003024 | 0.981 | 0.901 | 2.5E-248 | Chondrogenic Fibroblasts | cd81a |
| cst14a.22 | 2E-252 | 1.7216938 | 0.771 | 0.353 | 4.9E-248 | Chondrogenic Fibroblasts | cst14a.2 |
| rhoab1 | 6E-252 | 1.7712252 | 0.84 | 0.448 | 2E-247 | Chondrogenic Fibroblasts | rhoab |
| mxra5a2 | 2E-251 | 2.2788491 | 0.34 | 0.07 | 5.6E-247 | Chondrogenic Fibroblasts | mxra5a |
| slc22a211 | 3E-251 | 3.4439398 | 0.182 | 0.022 | 1E-246 | Chondrogenic Fibroblasts | slc22a21 |
| cd9b3 | 3E-248 | 2.2890273 | 0.391 | 0.093 | 8.8E-244 | Chondrogenic Fibroblasts | cd9b |
| synm2 | 1E-245 | 1.5918901 | 0.64 | 0.22 | 3.4E-241 | Chondrogenic Fibroblasts | synm |
| itga11 | 6E-244 | 2.5407789 | 0.202 | 0.027 | 2E-239 | Chondrogenic Fibroblasts | itga1 |
| atp1b1a1 | 7E-244 | 1.8888636 | 0.85 | 0.475 | 2.3E-239 | Chondrogenic Fibroblasts | atp1b1a |
| FP102018.13 | 1E-243 | 1.5821699 | 0.798 | 0.38 | 4.6E-239 | Chondrogenic Fibroblasts | FP102018.1 |
| sdcbp22 | 6E-242 | 1.5900974 | 0.908 | 0.591 | 1.9E-237 | Chondrogenic Fibroblasts | sdcbp2 |
| plecb3 | 2E-239 | 1.5345442 | 0.924 | 0.651 | 5.6E-235 | Chondrogenic Fibroblasts | plecb |
| loxl2a3 | 9E-239 | 2.0453657 | 0.599 | 0.214 | 2.8E-234 | Chondrogenic Fibroblasts | loxl2a |
| foxp45 | 3E-237 | 1.7986682 | 0.581 | 0.193 | 9.3E-233 | Chondrogenic Fibroblasts | foxp4 |
| atp2a31 | 1E-233 | 2.350944 | 0.232 | 0.036 | 4.6E-229 | Chondrogenic Fibroblasts | atp2a3 |

| | | | | | | | |
|---|---|---|---|---|---|---|---|
| col2a1b4 | 4E-232 | 1.2784799 | 0.868 | 0.492 | 1.2E-227 | Chondrogenic Fibroblasts | col2a1b |
| f13a1b2 | 3E-231 | 3.5376014 | 0.641 | 0.286 | 9.1E-227 | Chondrogenic Fibroblasts | f13a1b |
| camsap2b2 | 5E-230 | 2.0082549 | 0.401 | 0.1 | 1.6E-225 | Chondrogenic Fibroblasts | camsap2b |
| vwa14 | 3E-226 | 2.1962068 | 0.455 | 0.13 | 8.9E-222 | Chondrogenic Fibroblasts | vwa1 |
| dcn5 | 5E-226 | 1.13336 | 0.983 | 0.866 | 1.6E-221 | Chondrogenic Fibroblasts | dcn |
| gpx1b | 9E-226 | 3.3936924 | 0.151 | 0.017 | 2.8E-221 | Chondrogenic Fibroblasts | gpx1b |
| lmna3 | 1E-225 | 1.1354843 | 0.971 | 0.776 | 3.5E-221 | Chondrogenic Fibroblasts | lmna |
| tox2 | 1E-225 | 2.756222 | 0.277 | 0.053 | 4.7E-221 | Chondrogenic Fibroblasts | tox |
| ehd41 | 1E-224 | 2.5195718 | 0.269 | 0.05 | 3.6E-220 | Chondrogenic Fibroblasts | ehd4 |
| col6a25 | 1E-224 | 1.0687453 | 0.991 | 0.874 | 4.7E-220 | Chondrogenic Fibroblasts | col6a2 |
| igf1 | 2E-221 | 3.4799166 | 0.151 | 0.017 | 5.7E-217 | Chondrogenic Fibroblasts | igf1 |
| lrp1bb | 9E-220 | 3.7308223 | 0.113 | 0.009 | 3E-215 | Chondrogenic Fibroblasts | lrp1bb |
| antxr1c2 | 1E-216 | 1.5184419 | 0.87 | 0.528 | 3.5E-212 | Chondrogenic Fibroblasts | antxr1c |
| itgbl1 | 1E-215 | 2.5390627 | 0.184 | 0.025 | 4.4E-211 | Chondrogenic Fibroblasts | itgbl1 |
| schip13 | 2E-214 | 1.7753677 | 0.466 | 0.137 | 5.7E-210 | Chondrogenic Fibroblasts | schip1 |
| sdf43 | 4E-214 | 1.5903587 | 0.689 | 0.294 | 1.2E-209 | Chondrogenic Fibroblasts | sdf4 |
| si:dkey-19b23.81 | 1E-213 | 2.8668523 | 0.259 | 0.049 | 3.3E-209 | Chondrogenic Fibroblasts | si:dkey-19b23.8 |
| micu3a1 | 2E-211 | 2.7876443 | 0.176 | 0.024 | 5.2E-207 | Chondrogenic Fibroblasts | micu3a |
| p4ha1b5 | 4E-211 | 1.8295855 | 0.45 | 0.132 | 1.3E-206 | Chondrogenic Fibroblasts | p4ha1b |
| omd | 5E-209 | 2.9734005 | 0.204 | 0.032 | 1.5E-204 | Chondrogenic Fibroblasts | omd |
| pld74 | 5E-209 | 1.8868789 | 0.471 | 0.144 | 1.8E-204 | Chondrogenic Fibroblasts | pld7 |
| copz23 | 3E-208 | 1.721434 | 0.531 | 0.179 | 1E-203 | Chondrogenic Fibroblasts | copz2 |
| stxbp61 | 7E-208 | 2.367889 | 0.243 | 0.044 | 2.1E-203 | Chondrogenic Fibroblasts | stxbp6 |
| acana1 | 7E-208 | 2.2781169 | 0.204 | 0.031 | 2.2E-203 | Chondrogenic Fibroblasts | acana |
| serpinf15 | 5E-202 | 1.2331639 | 0.89 | 0.546 | 1.5E-197 | Chondrogenic Fibroblasts | serpinf1 |
| ppib4 | 2E-201 | 0.8499047 | 0.991 | 0.915 | 8.1E-197 | Chondrogenic Fibroblasts | ppib |
| postnb3 | 3E-201 | 1.4877926 | 0.817 | 0.446 | 9.5E-197 | Chondrogenic Fibroblasts | postnb |
| odc13 | 1E-200 | 1.9332897 | 0.412 | 0.117 | 3.6E-196 | Chondrogenic Fibroblasts | odc1 |
| lgals3a2 | 2E-199 | 1.6657187 | 0.64 | 0.268 | 6E-195 | Chondrogenic Fibroblasts | lgals3a |
| cd1511 | 4E-199 | 2.1796665 | 0.358 | 0.092 | 1.4E-194 | Chondrogenic Fibroblasts | cd151 |
| nfatc13 | 9E-199 | 1.9860864 | 0.333 | 0.08 | 2.9E-194 | Chondrogenic Fibroblasts | nfatc1 |
| sema3d1 | 6E-198 | 2.3066425 | 0.227 | 0.04 | 1.9E-193 | Chondrogenic Fibroblasts | sema3d |
| zeb1b3 | 2E-197 | 1.7938536 | 0.339 | 0.081 | 6.5E-193 | Chondrogenic Fibroblasts | zeb1b |
| cdkn1a4 | 8E-197 | 2.0077082 | 0.403 | 0.115 | 2.5E-192 | Chondrogenic Fibroblasts | cdkn1a |
| CR848833.14 | 1E-192 | 2.0300467 | 0.297 | 0.066 | 3.4E-188 | Chondrogenic Fibroblasts | CR848833.1 |
| tnmd3 | 8E-192 | 2.2577527 | 0.445 | 0.144 | 2.6E-187 | Chondrogenic Fibroblasts | tnmd |
| dhrs3b3 | 2E-191 | 1.7567902 | 0.393 | 0.109 | 5.5E-187 | Chondrogenic Fibroblasts | dhrs3b |
| calua3 | 3E-191 | 1.5119787 | 0.689 | 0.317 | 9.4E-187 | Chondrogenic Fibroblasts | calua |
| loxl3b1 | 1E-190 | 1.6803326 | 0.465 | 0.146 | 3.4E-186 | Chondrogenic Fibroblasts | loxl3b |
| tgfbr2b5 | 4E-190 | 1.5695819 | 0.449 | 0.135 | 1.3E-185 | Chondrogenic Fibroblasts | tgfbr2b |
| cthrc1a3 | 1E-189 | 1.937619 | 0.454 | 0.146 | 3.3E-185 | Chondrogenic Fibroblasts | cthrc1a |
| anxa5b2 | 7E-189 | 1.4361849 | 0.764 | 0.393 | 2.1E-184 | Chondrogenic Fibroblasts | anxa5b |
| olfml3a3 | 1E-188 | 1.8080685 | 0.602 | 0.249 | 4.5E-184 | Chondrogenic Fibroblasts | olfml3a |
| si:ch211-236l14.4 | 2E-187 | 2.9486656 | 0.127 | 0.014 | 7.7E-183 | Chondrogenic Fibroblasts | si:ch211-236l14.4 |
| lman13 | 3E-185 | 1.8691935 | 0.401 | 0.118 | 9.1E-181 | Chondrogenic Fibroblasts | lman1 |
| ddr2a3 | 3E-185 | 1.5200472 | 0.552 | 0.199 | 9.3E-181 | Chondrogenic Fibroblasts | ddr2a |
| pcolce2b4 | 5E-184 | 1.2878517 | 0.679 | 0.287 | 1.6E-179 | Chondrogenic Fibroblasts | pcolce2b |
| pgam1a2 | 6E-183 | 1.5662173 | 0.513 | 0.178 | 1.9E-178 | Chondrogenic Fibroblasts | pgam1a |

| | | | | | | | |
|---|---|---|---|---|---|---|---|
| prnprs32 | 7E-183 | 1.9556472 | 0.344 | 0.09 | 2.2E-178 | Chondrogenic Fibroblasts | prnprs3 |
| tmem45a3 | 8E-183 | 1.7272749 | 0.46 | 0.15 | 2.5E-178 | Chondrogenic Fibroblasts | tmem45a |
| gask1b1 | 5E-182 | 2.4467915 | 0.202 | 0.035 | 1.7E-177 | Chondrogenic Fibroblasts | gask1b |
| triqk3 | 2E-181 | 1.7151392 | 0.466 | 0.154 | 6.8E-177 | Chondrogenic Fibroblasts | triqk |
| phlda33 | 2E-181 | 1.6458247 | 0.498 | 0.172 | 8E-177 | Chondrogenic Fibroblasts | phlda3 |
| six2a3 | 7E-181 | 2.1516566 | 0.351 | 0.095 | 2.1E-176 | Chondrogenic Fibroblasts | six2a |
| aopep3 | 1E-179 | 2.0409891 | 0.296 | 0.07 | 3.6E-175 | Chondrogenic Fibroblasts | aopep |
| slc35e41 | 2E-179 | 2.1985697 | 0.302 | 0.073 | 6E-175 | Chondrogenic Fibroblasts | slc35e4 |
| mxra8a6 | 3E-176 | 1.5499049 | 0.603 | 0.245 | 1E-171 | Chondrogenic Fibroblasts | mxra8a |
| svep13 | 4E-173 | 1.6615258 | 0.434 | 0.137 | 1.4E-168 | Chondrogenic Fibroblasts | svep1 |
| gas61 | 5E-173 | 2.3849502 | 0.268 | 0.061 | 1.5E-168 | Chondrogenic Fibroblasts | gas6 |
| si:ch211-165e15.13 | 6E-173 | 1.9707152 | 0.273 | 0.062 | 1.9E-168 | Chondrogenic Fibroblasts | si:ch211-165e15.1 |
| krt155 | 3E-166 | 1.4932699 | 0.903 | 0.696 | 1E-161 | Chondrogenic Fibroblasts | krt15 |
| igfbp2b1 | 1E-165 | 2.0302021 | 0.276 | 0.065 | 3.3E-161 | Chondrogenic Fibroblasts | igfbp2b |
| mob2b2 | 2E-165 | 2.1177964 | 0.297 | 0.075 | 5.9E-161 | Chondrogenic Fibroblasts | mob2b |
| ppiab2 | 1E-163 | 0.7510452 | 0.982 | 0.894 | 3.4E-159 | Chondrogenic Fibroblasts | ppiab |
| vasnb4 | 4E-163 | 1.7571392 | 0.418 | 0.137 | 1.4E-158 | Chondrogenic Fibroblasts | vasnb |
| fibina4 | 2E-162 | 1.7185034 | 0.525 | 0.205 | 5.1E-158 | Chondrogenic Fibroblasts | fibina |
| sox51 | 7E-162 | 2.8674311 | 0.139 | 0.019 | 2.3E-157 | Chondrogenic Fibroblasts | sox5 |
| selenof3 | 1E-161 | 1.1757079 | 0.796 | 0.459 | 3.9E-157 | Chondrogenic Fibroblasts | selenof |
| sestd12 | 8E-160 | 1.8812433 | 0.345 | 0.099 | 2.7E-155 | Chondrogenic Fibroblasts | sestd1 |
| pard3bb1 | 2E-159 | 1.4761863 | 0.562 | 0.228 | 6.1E-155 | Chondrogenic Fibroblasts | pard3bb |
| plod24 | 7E-157 | 1.9459085 | 0.273 | 0.067 | 2.3E-152 | Chondrogenic Fibroblasts | plod2 |
| C7H16orf873 | 4E-153 | 1.8529959 | 0.378 | 0.12 | 1.2E-148 | Chondrogenic Fibroblasts | C7H16orf87 |
| cavin1b3 | 2E-152 | 1.12674 | 0.898 | 0.64 | 5.6E-148 | Chondrogenic Fibroblasts | cavin1b |
| tgm2l1 | 1E-151 | 2.7156097 | 0.125 | 0.016 | 3.8E-147 | Chondrogenic Fibroblasts | tgm2l |
| col6a3.14 | 2E-151 | 0.7987477 | 0.996 | 0.937 | 6.5E-147 | Chondrogenic Fibroblasts | col6a3.1 |
| fstl1b2 | 4E-151 | 0.8404159 | 0.953 | 0.752 | 1.2E-146 | Chondrogenic Fibroblasts | fstl1b |
| rcan1a4 | 2E-150 | 1.5056403 | 0.54 | 0.222 | 5.6E-146 | Chondrogenic Fibroblasts | rcan1a |
| dpysl2b2 | 3E-150 | 1.5303758 | 0.442 | 0.154 | 1.1E-145 | Chondrogenic Fibroblasts | dpysl2b |
| mmp14a4 | 1E-149 | 1.1031592 | 0.887 | 0.641 | 4.5E-145 | Chondrogenic Fibroblasts | mmp14a |
| mfge8a3 | 2E-149 | 1.6547632 | 0.367 | 0.113 | 5E-145 | Chondrogenic Fibroblasts | mfge8a |
| CR855337.11 | 3E-149 | 1.7922394 | 0.318 | 0.089 | 9.8E-145 | Chondrogenic Fibroblasts | CR855337.1 |
| reep3a3 | 9E-148 | 1.6814998 | 0.438 | 0.158 | 3E-143 | Chondrogenic Fibroblasts | reep3a |
| calr3b3 | 3E-146 | 1.4212325 | 0.595 | 0.262 | 8.5E-142 | Chondrogenic Fibroblasts | calr3b |
| si:ch1073-204b8.11 | 3E-146 | 2.09346 | 0.19 | 0.037 | 8.8E-142 | Chondrogenic Fibroblasts | si:ch1073-204b8.1 |
| anxa11a3 | 6E-146 | 1.3056662 | 0.731 | 0.402 | 1.9E-141 | Chondrogenic Fibroblasts | anxa11a |
| golim4b3 | 7E-146 | 1.7660503 | 0.341 | 0.103 | 2.3E-141 | Chondrogenic Fibroblasts | golim4b |
| traf4a2 | 2E-145 | 2.1603651 | 0.271 | 0.07 | 6.6E-141 | Chondrogenic Fibroblasts | traf4a |
| nucb2a3 | 2E-145 | 1.0561191 | 0.828 | 0.516 | 7.6E-141 | Chondrogenic Fibroblasts | nucb2a |
| eno1a2 | 4E-145 | 1.5315765 | 0.657 | 0.332 | 1.3E-140 | Chondrogenic Fibroblasts | eno1a |
| p3h22 | 4E-145 | 1.9233895 | 0.244 | 0.058 | 1.4E-140 | Chondrogenic Fibroblasts | p3h2 |
| cav13 | 1E-144 | 0.9783592 | 0.904 | 0.626 | 4.5E-140 | Chondrogenic Fibroblasts | cav1 |
| plp22 | 1E-143 | 1.6529508 | 0.357 | 0.112 | 3.7E-139 | Chondrogenic Fibroblasts | plp2 |
| p4ha31 | 4E-143 | 2.03228 | 0.177 | 0.033 | 1.4E-138 | Chondrogenic Fibroblasts | p4ha3 |
| cspg42 | 6E-142 | 1.6474892 | 0.351 | 0.108 | 2E-137 | Chondrogenic Fibroblasts | cspg4 |
| efna5a4 | 2E-141 | 1.9052109 | 0.276 | 0.073 | 5.5E-137 | Chondrogenic Fibroblasts | efna5a |
| smoc24 | 2E-140 | 1.772462 | 0.302 | 0.085 | 5.2E-136 | Chondrogenic Fibroblasts | smoc2 |

| | | | | | | | |
|---|---|---|---|---|---|---|---|
| calm1a4 | 5E-140 | 1.202697 | 0.878 | 0.612 | 1.5E-135 | Chondrogenic Fibroblasts | calm1a |
| sostdc1a1 | 2E-138 | 2.769704 | 0.167 | 0.031 | 5.1E-134 | Chondrogenic Fibroblasts | sostdc1a |
| tiam2a2 | 4E-138 | 1.9813263 | 0.237 | 0.057 | 1.3E-133 | Chondrogenic Fibroblasts | tiam2a |
| hdac44 | 3E-136 | 1.5234207 | 0.36 | 0.114 | 9.5E-132 | Chondrogenic Fibroblasts | hdac4 |
| calm3a3 | 5E-136 | 1.1372637 | 0.806 | 0.494 | 1.7E-131 | Chondrogenic Fibroblasts | calm3a |
| fkbp10b4 | 6E-136 | 1.2835922 | 0.563 | 0.247 | 2E-131 | Chondrogenic Fibroblasts | fkbp10b |
| sgk2b2 | 3E-135 | 2.2118866 | 0.178 | 0.035 | 9.7E-131 | Chondrogenic Fibroblasts | sgk2b |
| nr4a31 | 1E-134 | 2.6246587 | 0.207 | 0.047 | 3.5E-130 | Chondrogenic Fibroblasts | nr4a3 |
| ptmab4 | 2E-134 | 0.8861657 | 0.948 | 0.709 | 5.8E-130 | Chondrogenic Fibroblasts | ptmab |
| fkbp1aa2 | 5E-134 | 1.0197215 | 0.811 | 0.498 | 1.6E-129 | Chondrogenic Fibroblasts | fkbp1aa |
| anxa1a2 | 1E-133 | 0.7638752 | 0.983 | 0.903 | 3.4E-129 | Chondrogenic Fibroblasts | anxa1a |
| arl6ip5b2 | 3E-133 | 1.0161803 | 0.821 | 0.501 | 1.1E-128 | Chondrogenic Fibroblasts | arl6ip5b |
| mamdc2a | 5E-133 | 2.4877815 | 0.132 | 0.021 | 1.7E-128 | Chondrogenic Fibroblasts | mamdc2a |
| angptl2a4 | 2E-132 | 1.3016416 | 0.639 | 0.31 | 4.9E-128 | Chondrogenic Fibroblasts | angptl2a |
| ctsh3 | 4E-132 | 1.8146048 | 0.332 | 0.105 | 1.2E-127 | Chondrogenic Fibroblasts | ctsh |
| igf2a2 | 4E-132 | 2.2780331 | 0.296 | 0.087 | 1.3E-127 | Chondrogenic Fibroblasts | igf2a |
| prelp5 | 4E-132 | 1.3228818 | 0.384 | 0.127 | 1.4E-127 | Chondrogenic Fibroblasts | prelp |
| si:ch211-196g2.41 | 9E-132 | 1.675967 | 0.299 | 0.086 | 2.8E-127 | Chondrogenic Fibroblasts | si:ch211-196g2.4 |
| ywhag21 | 1E-131 | 1.4855709 | 0.431 | 0.161 | 4.4E-127 | Chondrogenic Fibroblasts | ywhag2 |
| meox13 | 4E-131 | 1.6143569 | 0.239 | 0.059 | 1.4E-126 | Chondrogenic Fibroblasts | meox1 |
| gapdhs2 | 5E-131 | 1.0566642 | 0.876 | 0.623 | 1.5E-126 | Chondrogenic Fibroblasts | gapdhs |
| mxra8b4 | 5E-131 | 1.3646615 | 0.712 | 0.4 | 1.6E-126 | Chondrogenic Fibroblasts | mxra8b |
| ngfb4 | 7E-131 | 1.7251442 | 0.277 | 0.077 | 2.2E-126 | Chondrogenic Fibroblasts | ngfb |
| phldb1b3 | 1E-130 | 1.6336811 | 0.292 | 0.083 | 3.6E-126 | Chondrogenic Fibroblasts | phldb1b |
| angptl55 | 4E-130 | 1.6115802 | 0.436 | 0.166 | 1.2E-125 | Chondrogenic Fibroblasts | angptl5 |
| loxl2b3 | 2E-129 | 1.1283489 | 0.644 | 0.309 | 6.2E-125 | Chondrogenic Fibroblasts | loxl2b |
| kdelr34 | 3E-128 | 1.4031069 | 0.453 | 0.176 | 8.2E-124 | Chondrogenic Fibroblasts | kdelr3 |
| nog33 | 4E-128 | 1.9111107 | 0.274 | 0.077 | 1.3E-123 | Chondrogenic Fibroblasts | nog3 |
| sema3h | 4E-128 | 2.1272345 | 0.155 | 0.028 | 1.3E-123 | Chondrogenic Fibroblasts | sema3h |
| pdgfrl5 | 4E-128 | 1.1681751 | 0.843 | 0.578 | 1.4E-123 | Chondrogenic Fibroblasts | pdgfrl |
| rab11fip1b | 7E-128 | 2.2455163 | 0.138 | 0.023 | 2.3E-123 | Chondrogenic Fibroblasts | rab11fip1b |
| klf2b2 | 3E-127 | 1.9144251 | 0.228 | 0.056 | 9.9E-123 | Chondrogenic Fibroblasts | klf2b |
| crip14 | 9E-126 | 1.5280632 | 0.733 | 0.455 | 2.9E-121 | Chondrogenic Fibroblasts | crip1 |
| ostc2 | 2E-124 | 1.1752813 | 0.601 | 0.283 | 7.8E-120 | Chondrogenic Fibroblasts | ostc |
| fam102ba | 3E-124 | 2.2695163 | 0.137 | 0.023 | 9E-120 | Chondrogenic Fibroblasts | fam102ba |
| atl33 | 2E-123 | 1.1756543 | 0.633 | 0.314 | 5.9E-119 | Chondrogenic Fibroblasts | atl3 |
| arhgef403 | 3E-123 | 1.3635087 | 0.464 | 0.184 | 1E-118 | Chondrogenic Fibroblasts | arhgef40 |
| ccn2b2 | 1E-122 | 1.9792989 | 0.466 | 0.2 | 3.1E-118 | Chondrogenic Fibroblasts | ccn2b |
| si:ch211-133n4.44 | 1E-122 | 1.0861531 | 0.753 | 0.44 | 3.6E-118 | Chondrogenic Fibroblasts | si:ch211-133n4.4 |
| esyt2a1 | 1E-122 | 1.2838756 | 0.434 | 0.163 | 4E-118 | Chondrogenic Fibroblasts | esyt2a |
| mtss11 | 1E-121 | 2.1075831 | 0.211 | 0.051 | 4.1E-117 | Chondrogenic Fibroblasts | mtss1 |
| cd992 | 4E-121 | 0.7861035 | 0.924 | 0.721 | 1.4E-116 | Chondrogenic Fibroblasts | cd99 |
| CU638870.14 | 6E-121 | 1.1080724 | 0.646 | 0.321 | 1.9E-116 | Chondrogenic Fibroblasts | CU638870.1 |
| arpc1a2 | 3E-120 | 1.1736583 | 0.619 | 0.31 | 1.1E-115 | Chondrogenic Fibroblasts | arpc1a |
| spaca4l1 | 8E-120 | 1.9283917 | 0.269 | 0.078 | 2.6E-115 | Chondrogenic Fibroblasts | spaca4l |
| fam114a13 | 9E-120 | 1.551343 | 0.323 | 0.105 | 2.9E-115 | Chondrogenic Fibroblasts | fam114a1 |
| serpinh1a4 | 1E-119 | 1.2717453 | 0.521 | 0.231 | 3.1E-115 | Chondrogenic Fibroblasts | serpinh1a |
| myo7ab1 | 2E-119 | 1.9153204 | 0.188 | 0.042 | 5.4E-115 | Chondrogenic Fibroblasts | myo7ab |

| | | | | | | | |
|---|---|---|---|---|---|---|---|
| nfixb5 | 2E-119 | 1.2503687 | 0.591 | 0.287 | 7.1E-115 | Chondrogenic Fibroblasts | nfixb |
| olfml3b4 | 1E-118 | 1.0902106 | 0.791 | 0.507 | 3.6E-114 | Chondrogenic Fibroblasts | olfml3b |
| rbfox12 | 2E-118 | 1.1263333 | 0.704 | 0.398 | 5.1E-114 | Chondrogenic Fibroblasts | rbfox1 |
| dap4 | 2E-118 | 1.2953391 | 0.463 | 0.189 | 5.2E-114 | Chondrogenic Fibroblasts | dap |
| si:dkey-12l12.12 | 2E-118 | 0.8974789 | 0.859 | 0.573 | 7.4E-114 | Chondrogenic Fibroblasts | si:dkey-12l12.1 |
| si:dkey-238c7.162 | 2E-118 | 1.3871186 | 0.416 | 0.159 | 7.7E-114 | Chondrogenic Fibroblasts | si:dkey-238c7.16 |
| ctsd3 | 3E-118 | 1.4240197 | 0.398 | 0.148 | 8.7E-114 | Chondrogenic Fibroblasts | ctsd |
| rgs3a3 | 2E-117 | 1.8900117 | 0.288 | 0.089 | 6E-113 | Chondrogenic Fibroblasts | rgs3a |
| mcf2l22 | 3E-117 | 1.4499245 | 0.32 | 0.102 | 9.3E-113 | Chondrogenic Fibroblasts | mcf2l2 |
| mid1ip1b3 | 3E-117 | 1.5698107 | 0.338 | 0.114 | 1E-112 | Chondrogenic Fibroblasts | mid1ip1b |
| scinla3 | 1E-115 | 1.269601 | 0.706 | 0.402 | 4.5E-111 | Chondrogenic Fibroblasts | scinla |
| plekhg5a4 | 2E-115 | 1.9600156 | 0.243 | 0.068 | 5.9E-111 | Chondrogenic Fibroblasts | plekhg5a |
| ucmab1 | 1E-114 | 1.0505532 | 0.173 | 0.037 | 4.2E-110 | Chondrogenic Fibroblasts | ucmab |
| ucmaa1 | 3E-114 | 2.4910401 | 0.176 | 0.039 | 8.9E-110 | Chondrogenic Fibroblasts | ucmaa |
| pdlim23 | 1E-113 | 1.4085779 | 0.399 | 0.15 | 4E-109 | Chondrogenic Fibroblasts | pdlim2 |
| roraa2 | 1E-113 | 1.3280657 | 0.377 | 0.136 | 4.9E-109 | Chondrogenic Fibroblasts | roraa |
| cd2763 | 2E-113 | 1.5690484 | 0.356 | 0.127 | 7.2E-109 | Chondrogenic Fibroblasts | cd276 |
| fn1a3 | 9E-113 | 1.0755669 | 0.631 | 0.31 | 2.8E-108 | Chondrogenic Fibroblasts | fn1a |
| rab2a2 | 1E-112 | 0.8843636 | 0.802 | 0.514 | 3.1E-108 | Chondrogenic Fibroblasts | rab2a |
| aldoaa3 | 1E-111 | 1.1082369 | 0.668 | 0.361 | 3.2E-107 | Chondrogenic Fibroblasts | aldoaa |
| pef12 | 2E-111 | 1.3231607 | 0.396 | 0.151 | 5.2E-107 | Chondrogenic Fibroblasts | pef1 |
| arhgap21a3 | 2E-111 | 1.4792954 | 0.311 | 0.101 | 6.9E-107 | Chondrogenic Fibroblasts | arhgap21a |
| rock2b.13 | 2E-111 | 1.4654749 | 0.416 | 0.166 | 7.6E-107 | Chondrogenic Fibroblasts | rock2b.1 |
| TCIM1 | 3E-110 | 2.0302166 | 0.226 | 0.062 | 8.7E-106 | Chondrogenic Fibroblasts | TCIM |
| srpx4 | 1E-109 | 1.3339294 | 0.412 | 0.161 | 3.1E-105 | Chondrogenic Fibroblasts | srpx |
| pdlim41 | 5E-109 | 2.0670411 | 0.168 | 0.037 | 1.5E-104 | Chondrogenic Fibroblasts | pdlim4 |
| kctd9a2 | 6E-109 | 1.6198776 | 0.29 | 0.093 | 2E-104 | Chondrogenic Fibroblasts | kctd9a |
| akap133 | 9E-109 | 1.0539032 | 0.617 | 0.313 | 3E-104 | Chondrogenic Fibroblasts | akap13 |
| ACBD31 | 2E-108 | 1.971369 | 0.162 | 0.035 | 8.1E-104 | Chondrogenic Fibroblasts | ACBD3 |
| pitpnaa2 | 3E-108 | 2.336913 | 0.199 | 0.051 | 8.5E-104 | Chondrogenic Fibroblasts | pitpnaa |
| rrbp1a4 | 3E-108 | 1.0430821 | 0.72 | 0.423 | 1E-103 | Chondrogenic Fibroblasts | rrbp1a |
| foxc1b2 | 5E-108 | 1.8990427 | 0.22 | 0.059 | 1.5E-103 | Chondrogenic Fibroblasts | foxc1b |
| AL732488.21 | 5E-108 | 1.745014 | 0.223 | 0.06 | 1.7E-103 | Chondrogenic Fibroblasts | AL732488.2 |
| tmem2583 | 6E-108 | 0.8747875 | 0.832 | 0.582 | 1.8E-103 | Chondrogenic Fibroblasts | tmem258 |
| tns1b3 | 6E-108 | 1.0287329 | 0.67 | 0.36 | 2E-103 | Chondrogenic Fibroblasts | tns1b |
| aebp14 | 2E-107 | 1.0906515 | 0.802 | 0.533 | 7E-103 | Chondrogenic Fibroblasts | aebp1 |
| col8a22 | 9E-106 | 1.8256453 | 0.227 | 0.063 | 2.8E-101 | Chondrogenic Fibroblasts | col8a2 |
| ndel1b1 | 7E-105 | 1.7066935 | 0.284 | 0.093 | 2.4E-100 | Chondrogenic Fibroblasts | ndel1b |
| tfpia5 | 1E-104 | 1.3806448 | 0.422 | 0.173 | 3.4E-100 | Chondrogenic Fibroblasts | tfpia |
| olfm1b | 1E-104 | 2.3915619 | 0.123 | 0.022 | 3.5E-100 | Chondrogenic Fibroblasts | olfm1b |
| fkbp93 | 3E-104 | 1.0042092 | 0.673 | 0.377 | 1.1E-99 | Chondrogenic Fibroblasts | fkbp9 |
| ppp3r1b1 | 8E-104 | 1.4729335 | 0.349 | 0.129 | 2.7E-99 | Chondrogenic Fibroblasts | ppp3r1b |
| plpp2b2 | 3E-103 | 1.5940497 | 0.334 | 0.122 | 9E-99 | Chondrogenic Fibroblasts | plpp2b |
| tent5bb4 | 3E-103 | 1.3107101 | 0.365 | 0.137 | 9.6E-99 | Chondrogenic Fibroblasts | tent5bb |
| pitx2 | 8E-103 | 2.4213009 | 0.142 | 0.029 | 2.58E-98 | Chondrogenic Fibroblasts | pitx2 |
| p4ha23 | 1E-102 | 1.3553353 | 0.364 | 0.137 | 4.71E-98 | Chondrogenic Fibroblasts | p4ha2 |
| mical2a | 2E-102 | 1.8684395 | 0.162 | 0.036 | 6.53E-98 | Chondrogenic Fibroblasts | mical2a |
| atp2b1a1 | 2E-102 | 1.1858663 | 0.518 | 0.24 | 8.03E-98 | Chondrogenic Fibroblasts | atp2b1a |

| | | | | | | | |
|---|---|---|---|---|---|---|---|
| ches13 | 1E-101 | 1.1228786 | 0.433 | 0.176 | 3.77E-97 | Chondrogenic Fibroblasts | ches1 |
| specc11 | 2E-101 | 1.9756469 | 0.158 | 0.035 | 5.21E-97 | Chondrogenic Fibroblasts | specc1 |
| si:ch211-57n23.14 | 3E-101 | 1.2179509 | 0.475 | 0.211 | 1.02E-96 | Chondrogenic Fibroblasts | si:ch211-57n23.1 |
| plod1a2 | 5E-100 | 1.0865561 | 0.514 | 0.236 | 1.66E-95 | Chondrogenic Fibroblasts | plod1a |
| zgc:110366 | 1E-99 | 2.1838313 | 0.133 | 0.026 | 3.68E-95 | Chondrogenic Fibroblasts | zgc:110366 |
| praf21 | 3E-99 | 1.2873132 | 0.356 | 0.133 | 9.32E-95 | Chondrogenic Fibroblasts | praf2 |
| selenom3 | 1.6E-98 | 1.3172678 | 0.401 | 0.166 | 5.26E-94 | Chondrogenic Fibroblasts | selenom |
| arpc32 | 2.4E-98 | 0.9901039 | 0.669 | 0.377 | 7.93E-94 | Chondrogenic Fibroblasts | arpc3 |
| thbs1b2 | 1.3E-97 | 1.2939748 | 0.464 | 0.207 | 4.23E-93 | Chondrogenic Fibroblasts | thbs1b |
| scocb2 | 2.2E-97 | 1.4736435 | 0.253 | 0.078 | 7.05E-93 | Chondrogenic Fibroblasts | scocb |
| stmn1b3 | 2.3E-97 | 1.5681782 | 0.325 | 0.119 | 7.5E-93 | Chondrogenic Fibroblasts | stmn1b |
| rarga3 | 3.3E-97 | 1.1943267 | 0.54 | 0.271 | 1.08E-92 | Chondrogenic Fibroblasts | rarga |
| tspan9b2 | 3.4E-97 | 1.4384261 | 0.342 | 0.129 | 1.09E-92 | Chondrogenic Fibroblasts | tspan9b |
| itga54 | 2.7E-96 | 1.029318 | 0.527 | 0.25 | 8.88E-92 | Chondrogenic Fibroblasts | itga5 |
| six1a3 | 5.6E-96 | 1.3122988 | 0.338 | 0.125 | 1.84E-91 | Chondrogenic Fibroblasts | six1a |
| uchl31 | 1.5E-95 | 1.3528969 | 0.3 | 0.104 | 4.91E-91 | Chondrogenic Fibroblasts | uchl3 |
| dpp73 | 1.6E-95 | 1.4180235 | 0.458 | 0.209 | 5.17E-91 | Chondrogenic Fibroblasts | dpp7 |
| cmpk2 | 2.1E-95 | 1.2037802 | 0.393 | 0.159 | 6.67E-91 | Chondrogenic Fibroblasts | cmpk |
| arhgef12a1 | 2.1E-95 | 1.5975388 | 0.259 | 0.083 | 6.98E-91 | Chondrogenic Fibroblasts | arhgef12a |
| rab131 | 2.9E-95 | 0.9934768 | 0.486 | 0.216 | 9.33E-91 | Chondrogenic Fibroblasts | rab13 |
| txndc52 | 6.3E-95 | 1.1715305 | 0.451 | 0.2 | 2.06E-90 | Chondrogenic Fibroblasts | txndc5 |
| ctsz2 | 6.6E-94 | 1.2391723 | 0.387 | 0.157 | 2.16E-89 | Chondrogenic Fibroblasts | ctsz |
| col6a33 | 7.9E-94 | 1.0674145 | 0.503 | 0.231 | 2.57E-89 | Chondrogenic Fibroblasts | col6a3 |
| si:ch73-138n13.12 | 5.2E-93 | 0.9753265 | 0.66 | 0.37 | 1.68E-88 | Chondrogenic Fibroblasts | si:ch73-138n13.1 |
| sox9a1 | 6E-93 | 1.5932163 | 0.265 | 0.087 | 1.95E-88 | Chondrogenic Fibroblasts | sox9a |
| itgb1a2 | 1.9E-92 | 0.9883162 | 0.538 | 0.258 | 6.14E-88 | Chondrogenic Fibroblasts | itgb1a |
| ecrg4a4 | 2.3E-92 | 0.2106259 | 0.857 | 0.556 | 7.36E-88 | Chondrogenic Fibroblasts | ecrg4a |
| kif13ba2 | 4.5E-92 | 1.4679903 | 0.284 | 0.098 | 1.45E-87 | Chondrogenic Fibroblasts | kif13ba |
| sept151 | 6.2E-92 | 1.1164729 | 0.446 | 0.198 | 2.01E-87 | Chondrogenic Fibroblasts | sept15 |
| rab1ab1 | 9E-92 | 0.863221 | 0.702 | 0.413 | 2.91E-87 | Chondrogenic Fibroblasts | rab1ab |
| si:ch211-191c10.1 | 3.2E-91 | 1.948271 | 0.146 | 0.033 | 1.04E-86 | Chondrogenic Fibroblasts | si:ch211-191c10.1 |
| atp6v0ca2 | 6.9E-91 | 0.9700148 | 0.619 | 0.332 | 2.24E-86 | Chondrogenic Fibroblasts | atp6v0ca |
| ezrb2 | 1.1E-90 | 1.4754837 | 0.419 | 0.185 | 3.62E-86 | Chondrogenic Fibroblasts | ezrb |
| rhoac2 | 1.2E-90 | 1.0140773 | 0.567 | 0.289 | 3.95E-86 | Chondrogenic Fibroblasts | rhoac |
| serf24 | 1.3E-90 | 0.6353414 | 0.94 | 0.768 | 4.11E-86 | Chondrogenic Fibroblasts | serf2 |
| EMP31 | 1.7E-90 | 1.7668578 | 0.21 | 0.062 | 5.42E-86 | Chondrogenic Fibroblasts | EMP3 |
| arg25 | 2.6E-90 | 1.2003047 | 0.382 | 0.158 | 8.59E-86 | Chondrogenic Fibroblasts | arg2 |
| myof2 | 3.7E-90 | 1.0342444 | 0.351 | 0.133 | 1.2E-85 | Chondrogenic Fibroblasts | myof |
| si:dkey-222f8.32 | 1.1E-89 | 1.1094558 | 0.468 | 0.215 | 3.62E-85 | Chondrogenic Fibroblasts | si:dkey-222f8.3 |
| ppa1b3 | 1.6E-89 | 1.2912551 | 0.342 | 0.133 | 5.05E-85 | Chondrogenic Fibroblasts | ppa1b |
| map1ab3 | 1.7E-89 | 1.6217444 | 0.422 | 0.191 | 5.51E-85 | Chondrogenic Fibroblasts | map1ab |
| kdelr2b2 | 5E-89 | 1.0207504 | 0.586 | 0.314 | 1.61E-84 | Chondrogenic Fibroblasts | kdelr2b |
| iqsec1b2 | 6.4E-89 | 1.0687094 | 0.617 | 0.337 | 2.08E-84 | Chondrogenic Fibroblasts | iqsec1b |
| lrp1aa5 | 9.4E-89 | 0.7386829 | 0.882 | 0.669 | 3.05E-84 | Chondrogenic Fibroblasts | lrp1aa |
| gabarapa1 | 1.2E-88 | 0.9262555 | 0.611 | 0.325 | 3.8E-84 | Chondrogenic Fibroblasts | gabarapa |
| fam83hb1 | 1.8E-88 | 1.5848197 | 0.209 | 0.061 | 5.93E-84 | Chondrogenic Fibroblasts | fam83hb |
| xirp2a3 | 3.6E-88 | 1.0333774 | 0.515 | 0.251 | 1.18E-83 | Chondrogenic Fibroblasts | xirp2a |
| ccn2a4 | 4E-88 | 1.3123074 | 0.438 | 0.201 | 1.31E-83 | Chondrogenic Fibroblasts | ccn2a |

| | | | | | | | |
|---|---|---|---|---|---|---|---|
| cfl11 | 5.1E-88 | 0.6067167 | 0.956 | 0.841 | 1.65E-83 | Chondrogenic Fibroblasts | cfl1 |
| dipk1c5 | 1.2E-87 | 1.4682351 | 0.299 | 0.11 | 3.84E-83 | Chondrogenic Fibroblasts | dipk1c |
| sgms11 | 3.1E-86 | 1.7044346 | 0.192 | 0.054 | 1.01E-81 | Chondrogenic Fibroblasts | sgms1 |
| clec11a2 | 4.2E-86 | 1.9908111 | 0.19 | 0.054 | 1.36E-81 | Chondrogenic Fibroblasts | clec11a |
| chmp2a1 | 5.8E-86 | 1.1551574 | 0.424 | 0.19 | 1.89E-81 | Chondrogenic Fibroblasts | chmp2a |
| pkma2 | 6.5E-86 | 1.1395818 | 0.449 | 0.207 | 2.11E-81 | Chondrogenic Fibroblasts | pkma |
| ccn5 | 1.3E-85 | 2.0225398 | 0.119 | 0.024 | 4.36E-81 | Chondrogenic Fibroblasts | ccn5 |
| cavin2b3 | 3.9E-85 | 0.7966382 | 0.799 | 0.524 | 1.25E-80 | Chondrogenic Fibroblasts | cavin2b |
| adamtsl34 | 1.2E-84 | 1.5127581 | 0.216 | 0.066 | 4.03E-80 | Chondrogenic Fibroblasts | adamtsl3 |
| atp6v1e1b3 | 1.3E-84 | 1.3509303 | 0.271 | 0.095 | 4.37E-80 | Chondrogenic Fibroblasts | atp6v1e1b |
| zgc:1538671 | 1.6E-84 | 0.6842161 | 0.902 | 0.749 | 5.19E-80 | Chondrogenic Fibroblasts | zgc:153867 |
| tmed34 | 6.3E-84 | 1.3466438 | 0.265 | 0.092 | 2.05E-79 | Chondrogenic Fibroblasts | tmed3 |
| cdc42se11 | 1.9E-83 | 1.4246564 | 0.23 | 0.073 | 6.25E-79 | Chondrogenic Fibroblasts | cdc42se1 |
| snap23.12 | 1.3E-82 | 1.1381369 | 0.403 | 0.177 | 4.15E-78 | Chondrogenic Fibroblasts | snap23.1 |
| c1qtnf6a3 | 1.4E-82 | 1.5654282 | 0.198 | 0.058 | 4.39E-78 | Chondrogenic Fibroblasts | c1qtnf6a |
| snx9b1 | 1.8E-82 | 1.1041879 | 0.402 | 0.175 | 5.84E-78 | Chondrogenic Fibroblasts | snx9b |
| vkorc11 | 2.7E-82 | 1.7459908 | 0.173 | 0.047 | 8.92E-78 | Chondrogenic Fibroblasts | vkorc1 |
| nectin3b2 | 3.6E-82 | 1.0135243 | 0.366 | 0.15 | 1.18E-77 | Chondrogenic Fibroblasts | nectin3b |
| copb22 | 4.1E-82 | 0.9434033 | 0.525 | 0.264 | 1.33E-77 | Chondrogenic Fibroblasts | copb2 |
| wwp21 | 1.2E-81 | 1.347959 | 0.277 | 0.1 | 3.83E-77 | Chondrogenic Fibroblasts | wwp2 |
| slc39a133 | 1.8E-81 | 1.1865621 | 0.339 | 0.137 | 5.84E-77 | Chondrogenic Fibroblasts | slc39a13 |
| nupr1a3 | 2.9E-81 | 1.1144075 | 0.499 | 0.251 | 9.33E-77 | Chondrogenic Fibroblasts | nupr1a |
| kdelr2a2 | 6.7E-81 | 0.84428 | 0.659 | 0.384 | 2.17E-76 | Chondrogenic Fibroblasts | kdelr2a |
| lamb1b2 | 1.2E-80 | 1.005646 | 0.42 | 0.19 | 4E-76 | Chondrogenic Fibroblasts | lamb1b |
| adgre5a | 1.7E-80 | 2.107031 | 0.107 | 0.021 | 5.6E-76 | Chondrogenic Fibroblasts | adgre5a |
| map7d1b2 | 2.4E-80 | 1.0660142 | 0.412 | 0.184 | 7.76E-76 | Chondrogenic Fibroblasts | map7d1b |
| c1qtnf53 | 2.8E-80 | 1.6710004 | 0.344 | 0.147 | 9.2E-76 | Chondrogenic Fibroblasts | c1qtnf5 |
| ddost2 | 3.5E-80 | 1.0050816 | 0.459 | 0.219 | 1.15E-75 | Chondrogenic Fibroblasts | ddost |
| dad13 | 5E-80 | 0.8495429 | 0.664 | 0.393 | 1.62E-75 | Chondrogenic Fibroblasts | dad1 |
| ano64 | 1.2E-79 | 1.2340852 | 0.312 | 0.12 | 3.79E-75 | Chondrogenic Fibroblasts | ano6 |
| mmp14b5 | 1.7E-79 | 0.8850825 | 0.797 | 0.544 | 5.6E-75 | Chondrogenic Fibroblasts | mmp14b |
| arf4a.13 | 2.9E-79 | 1.3449785 | 0.222 | 0.071 | 9.38E-75 | Chondrogenic Fibroblasts | arf4a.1 |
| atp5pd3 | 4.1E-79 | 0.8370266 | 0.638 | 0.361 | 1.34E-74 | Chondrogenic Fibroblasts | atp5pd |
| s100w2 | 8.6E-79 | 0.8882796 | 0.693 | 0.427 | 2.79E-74 | Chondrogenic Fibroblasts | s100w |
| col11a1a3 | 1.8E-78 | 0.4000782 | 0.557 | 0.316 | 5.74E-74 | Chondrogenic Fibroblasts | col11a1a |
| MYO9B3 | 2.4E-78 | 1.2645811 | 0.304 | 0.119 | 7.64E-74 | Chondrogenic Fibroblasts | MYO9B |
| clu4 | 2.4E-78 | 0.9327435 | 0.648 | 0.377 | 7.89E-74 | Chondrogenic Fibroblasts | clu |
| phldb1a1 | 2.8E-78 | 1.8304343 | 0.153 | 0.04 | 9.13E-74 | Chondrogenic Fibroblasts | phldb1a |
| nenf2 | 2.9E-78 | 1.3313802 | 0.26 | 0.093 | 9.38E-74 | Chondrogenic Fibroblasts | nenf |
| aldocb1 | 3.6E-78 | 1.4874695 | 0.222 | 0.072 | 1.17E-73 | Chondrogenic Fibroblasts | aldocb |
| skia4 | 8.6E-78 | 0.8617679 | 0.703 | 0.443 | 2.8E-73 | Chondrogenic Fibroblasts | skia |
| sgce1 | 1.2E-77 | 0.9227205 | 0.534 | 0.276 | 3.78E-73 | Chondrogenic Fibroblasts | sgce |
| ndfip12 | 1.5E-77 | 0.8608087 | 0.631 | 0.368 | 4.85E-73 | Chondrogenic Fibroblasts | ndfip1 |
| cope3 | 6.5E-77 | 1.0504407 | 0.491 | 0.251 | 2.12E-72 | Chondrogenic Fibroblasts | cope |
| cnnm2b | 8.6E-77 | 1.9348268 | 0.106 | 0.021 | 2.79E-72 | Chondrogenic Fibroblasts | cnnm2b |
| fkbp72 | 1.3E-76 | 0.7933012 | 0.802 | 0.608 | 4.28E-72 | Chondrogenic Fibroblasts | fkbp7 |
| hint13 | 2.6E-76 | 0.8633923 | 0.637 | 0.38 | 8.38E-72 | Chondrogenic Fibroblasts | hint1 |
| pdia54 | 3.2E-76 | 1.0891947 | 0.37 | 0.161 | 1.04E-71 | Chondrogenic Fibroblasts | pdia5 |

| | | | | | | | |
|---|---|---|---|---|---|---|---|
| C20H1orf1984 | 6.6E-76 | 1.0216753 | 0.399 | 0.179 | 2.15E-71 | Chondrogenic Fibroblasts | C20H1orf198 |
| glis2a3 | 9.2E-76 | 1.073185 | 0.425 | 0.199 | 2.99E-71 | Chondrogenic Fibroblasts | glis2a |
| hexb3 | 3.2E-75 | 1.4503268 | 0.246 | 0.087 | 1.04E-70 | Chondrogenic Fibroblasts | hexb |
| ldlrad4a | 4.2E-75 | 1.6731095 | 0.141 | 0.035 | 1.38E-70 | Chondrogenic Fibroblasts | ldlrad4a |
| slc31a13 | 5.2E-75 | 0.985092 | 0.417 | 0.192 | 1.69E-70 | Chondrogenic Fibroblasts | slc31a1 |
| pld1a2 | 6.2E-75 | 1.3182069 | 0.251 | 0.089 | 2.01E-70 | Chondrogenic Fibroblasts | pld1a |
| rcn34 | 2.1E-74 | 0.9565411 | 0.497 | 0.257 | 6.81E-70 | Chondrogenic Fibroblasts | rcn3 |
| prg4b1 | 3.9E-74 | 1.5942137 | 0.165 | 0.047 | 1.26E-69 | Chondrogenic Fibroblasts | prg4b |
| manf2 | 5.3E-74 | 0.9960764 | 0.399 | 0.18 | 1.74E-69 | Chondrogenic Fibroblasts | manf |
| ssr44 | 5.4E-74 | 0.7115528 | 0.834 | 0.602 | 1.75E-69 | Chondrogenic Fibroblasts | ssr4 |
| ywhah2 | 6.1E-74 | 1.1157815 | 0.361 | 0.156 | 2E-69 | Chondrogenic Fibroblasts | ywhah |
| srgn3 | 6.6E-74 | 0.9213019 | 0.712 | 0.445 | 2.15E-69 | Chondrogenic Fibroblasts | srgn |
| git12 | 1.2E-73 | 1.5679376 | 0.198 | 0.063 | 3.93E-69 | Chondrogenic Fibroblasts | git1 |
| CABZ01044281.11 | 9.1E-73 | 1.4881057 | 0.154 | 0.042 | 2.96E-68 | Chondrogenic Fibroblasts | CABZ01044281.1 |
| arpc5a1 | 1.7E-72 | 1.1077015 | 0.347 | 0.15 | 5.63E-68 | Chondrogenic Fibroblasts | arpc5a |
| fap1 | 1.8E-72 | 1.3800731 | 0.198 | 0.062 | 6.01E-68 | Chondrogenic Fibroblasts | fap |
| wipi14 | 2.5E-72 | 1.2727105 | 0.296 | 0.117 | 8.29E-68 | Chondrogenic Fibroblasts | wipi1 |
| fstl1a3 | 3.2E-72 | 1.609061 | 0.268 | 0.103 | 1.03E-67 | Chondrogenic Fibroblasts | fstl1a |
| sdc24 | 3.4E-72 | 0.7333158 | 0.814 | 0.582 | 1.1E-67 | Chondrogenic Fibroblasts | sdc2 |
| nedd4l2 | 7.1E-72 | 1.3686349 | 0.259 | 0.096 | 2.3E-67 | Chondrogenic Fibroblasts | nedd4l |
| uba53 | 9.4E-72 | 1.2498088 | 0.271 | 0.103 | 3.07E-67 | Chondrogenic Fibroblasts | uba5 |
| plod32 | 1E-71 | 1.1108427 | 0.341 | 0.146 | 3.32E-67 | Chondrogenic Fibroblasts | plod3 |
| prss164 | 1.7E-71 | 1.3741994 | 0.241 | 0.087 | 5.52E-67 | Chondrogenic Fibroblasts | prss16 |
| trps15 | 4E-71 | 1.0604142 | 0.379 | 0.169 | 1.29E-66 | Chondrogenic Fibroblasts | trps1 |
| arhgef12b | 5.7E-71 | 0.9772722 | 0.413 | 0.193 | 1.86E-66 | Chondrogenic Fibroblasts | arhgef12b |
| tspan5a1 | 8.5E-71 | 2.1256716 | 0.125 | 0.031 | 2.76E-66 | Chondrogenic Fibroblasts | tspan5a |
| cox6a12 | 8.6E-71 | 0.7207588 | 0.742 | 0.485 | 2.79E-66 | Chondrogenic Fibroblasts | cox6a1 |
| BX927333.11 | 1.3E-70 | 1.4587575 | 0.162 | 0.046 | 4.34E-66 | Chondrogenic Fibroblasts | BX927333.1 |
| reep51 | 1.6E-70 | 0.7760278 | 0.772 | 0.535 | 5.12E-66 | Chondrogenic Fibroblasts | reep5 |
| rpz23 | 3.5E-70 | 1.0607625 | 0.381 | 0.174 | 1.12E-65 | Chondrogenic Fibroblasts | rpz2 |
| map1lc3a2 | 4.6E-70 | 0.9442935 | 0.432 | 0.208 | 1.49E-65 | Chondrogenic Fibroblasts | map1lc3a |
| ext1a | 8.1E-70 | 1.5287457 | 0.166 | 0.048 | 2.63E-65 | Chondrogenic Fibroblasts | ext1a |
| clic42 | 9E-70 | 0.9573354 | 0.412 | 0.193 | 2.92E-65 | Chondrogenic Fibroblasts | clic4 |
| pdia62 | 9.8E-70 | 0.9696271 | 0.412 | 0.195 | 3.17E-65 | Chondrogenic Fibroblasts | pdia6 |
| zgc:1750883 | 1.5E-69 | 0.9310945 | 0.508 | 0.268 | 4.72E-65 | Chondrogenic Fibroblasts | zgc:175088 |
| ldha3 | 2.1E-69 | 1.0321418 | 0.454 | 0.225 | 6.73E-65 | Chondrogenic Fibroblasts | ldha |
| ndufb92 | 6.9E-69 | 0.8358261 | 0.555 | 0.303 | 2.25E-64 | Chondrogenic Fibroblasts | ndufb9 |
| foxn31 | 9.2E-69 | 1.2522898 | 0.215 | 0.073 | 3E-64 | Chondrogenic Fibroblasts | foxn3 |
| fthl281 | 1.4E-68 | 2.4124009 | 0.101 | 0.022 | 4.69E-64 | Chondrogenic Fibroblasts | fthl28 |
| rca2.11 | 2.2E-68 | 0.9132209 | 0.521 | 0.279 | 7.17E-64 | Chondrogenic Fibroblasts | rca2.1 |
| nfic2 | 1.2E-67 | 1.2331922 | 0.218 | 0.075 | 3.98E-63 | Chondrogenic Fibroblasts | nfic |
| sec31a2 | 3E-67 | 1.1540283 | 0.271 | 0.105 | 9.85E-63 | Chondrogenic Fibroblasts | sec31a |
| cox5ab3 | 3.9E-67 | 0.8623055 | 0.439 | 0.215 | 1.28E-62 | Chondrogenic Fibroblasts | cox5ab |
| ackr4b2 | 4.7E-67 | 1.5740571 | 0.163 | 0.049 | 1.54E-62 | Chondrogenic Fibroblasts | ackr4b |
| im:7151449 | 2.1E-66 | 1.1225998 | 0.156 | 0.045 | 6.7E-62 | Chondrogenic Fibroblasts | im:7151449 |
| ppt13 | 2.3E-66 | 0.8051629 | 0.587 | 0.339 | 7.35E-62 | Chondrogenic Fibroblasts | ppt1 |
| coa3a1 | 2.6E-66 | 1.0546886 | 0.302 | 0.125 | 8.61E-62 | Chondrogenic Fibroblasts | coa3a |
| ptpn131 | 3E-66 | 0.8323271 | 0.54 | 0.296 | 9.73E-62 | Chondrogenic Fibroblasts | ptpn13 |

| | | | | | | | |
|---|---|---|---|---|---|---|---|
| tpi1b2 | 1E-65 | 0.8969086 | 0.595 | 0.352 | 3.24E-61 | Chondrogenic Fibroblasts | tpi1b |
| arpc5b3 | 1.1E-65 | 1.4306815 | 0.229 | 0.084 | 3.42E-61 | Chondrogenic Fibroblasts | arpc5b |
| txndc122 | 1.4E-65 | 1.2996833 | 0.192 | 0.063 | 4.48E-61 | Chondrogenic Fibroblasts | txndc12 |
| si:dkey-188i13.111 | 1.6E-65 | 1.0023627 | 0.406 | 0.198 | 5.24E-61 | Chondrogenic Fibroblasts | si:dkey-188i13.11 |
| cops93 | 2.8E-65 | 1.0369727 | 0.338 | 0.149 | 8.95E-61 | Chondrogenic Fibroblasts | cops9 |
| nupr1b2 | 2.9E-65 | 1.5687273 | 0.347 | 0.159 | 9.3E-61 | Chondrogenic Fibroblasts | nupr1b |
| ccser2a | 5.4E-65 | 1.693878 | 0.132 | 0.035 | 1.75E-60 | Chondrogenic Fibroblasts | ccser2a |
| vma212 | 5.9E-65 | 1.0505693 | 0.276 | 0.11 | 1.92E-60 | Chondrogenic Fibroblasts | vma21 |
| csrp1a3 | 7.4E-65 | 0.7979192 | 0.484 | 0.251 | 2.39E-60 | Chondrogenic Fibroblasts | csrp1a |
| pdia75 | 1.3E-64 | 1.0342415 | 0.324 | 0.14 | 4.13E-60 | Chondrogenic Fibroblasts | pdia7 |
| si:ch1073-385f13.3 | 1.4E-64 | 1.3123859 | 0.183 | 0.059 | 4.66E-60 | Chondrogenic Fibroblasts | si:ch1073-385f13.3 |
| COX5B.13 | 1.7E-64 | 0.598585 | 0.778 | 0.535 | 5.52E-60 | Chondrogenic Fibroblasts | COX5B.1 |
| cremb | 4.3E-64 | 1.7566768 | 0.145 | 0.042 | 1.41E-59 | Chondrogenic Fibroblasts | cremb |
| cilp22 | 6E-64 | 1.3739537 | 0.182 | 0.06 | 1.95E-59 | Chondrogenic Fibroblasts | cilp2 |
| inhbaa3 | 7.5E-64 | 1.4505659 | 0.476 | 0.267 | 2.45E-59 | Chondrogenic Fibroblasts | inhbaa |
| dynlt31 | 2.1E-63 | 0.9096383 | 0.43 | 0.215 | 6.92E-59 | Chondrogenic Fibroblasts | dynlt3 |
| mibp22 | 2.7E-63 | 0.8024549 | 0.824 | 0.607 | 8.82E-59 | Chondrogenic Fibroblasts | mibp2 |
| anxa43 | 3.2E-63 | 0.70985 | 0.869 | 0.7 | 1.04E-58 | Chondrogenic Fibroblasts | anxa4 |
| kif5ba.11 | 4.7E-63 | 1.1536924 | 0.248 | 0.095 | 1.53E-58 | Chondrogenic Fibroblasts | kif5ba.1 |
| gpm6ab2 | 8.7E-63 | 1.1270787 | 0.229 | 0.085 | 2.81E-58 | Chondrogenic Fibroblasts | gpm6ab |
| pls32 | 8.7E-63 | 0.7658832 | 0.556 | 0.315 | 2.82E-58 | Chondrogenic Fibroblasts | pls3 |
| myo1cb | 9.2E-63 | 0.8103123 | 0.479 | 0.248 | 2.99E-58 | Chondrogenic Fibroblasts | myo1cb |
| cyb5r12 | 1.6E-62 | 0.9710389 | 0.401 | 0.196 | 5.27E-58 | Chondrogenic Fibroblasts | cyb5r1 |
| fkbp1ab1 | 2.1E-62 | 1.5754196 | 0.134 | 0.037 | 6.96E-58 | Chondrogenic Fibroblasts | fkbp1ab |
| capns1b1 | 4.4E-62 | 0.7186104 | 0.729 | 0.49 | 1.42E-57 | Chondrogenic Fibroblasts | capns1b |
| ackr3b2 | 4.4E-62 | 1.1037104 | 0.278 | 0.115 | 1.42E-57 | Chondrogenic Fibroblasts | ackr3b |
| cox7a2a2 | 5.5E-62 | 0.7568966 | 0.496 | 0.265 | 1.8E-57 | Chondrogenic Fibroblasts | cox7a2a |
| tmed92 | 7.4E-62 | 0.8696595 | 0.436 | 0.222 | 2.41E-57 | Chondrogenic Fibroblasts | tmed9 |
| tmed72 | 1.2E-61 | 0.8020815 | 0.548 | 0.311 | 4.05E-57 | Chondrogenic Fibroblasts | tmed7 |
| cdc42ep51 | 2.3E-61 | 0.8743738 | 0.523 | 0.295 | 7.47E-57 | Chondrogenic Fibroblasts | cdc42ep5 |
| dynlrb1.12 | 2.4E-61 | 0.8508884 | 0.546 | 0.312 | 7.77E-57 | Chondrogenic Fibroblasts | dynlrb1.1 |
| dnajb113 | 2.8E-61 | 1.1588289 | 0.256 | 0.102 | 9.11E-57 | Chondrogenic Fibroblasts | dnajb11 |
| rab6a2 | 4.2E-61 | 0.9369277 | 0.375 | 0.18 | 1.36E-56 | Chondrogenic Fibroblasts | rab6a |
| sar1b2 | 4.4E-61 | 0.7917192 | 0.527 | 0.294 | 1.44E-56 | Chondrogenic Fibroblasts | sar1b |
| sh3pxd2b1 | 7.2E-61 | 0.8194058 | 0.507 | 0.276 | 2.34E-56 | Chondrogenic Fibroblasts | sh3pxd2b |
| si:ch211-243a20.34 | 7.6E-61 | 1.2375341 | 0.486 | 0.277 | 2.47E-56 | Chondrogenic Fibroblasts | si:ch211-243a20.3 |
| eif4eb2 | 1E-60 | 1.0931519 | 0.255 | 0.101 | 3.32E-56 | Chondrogenic Fibroblasts | eif4eb |
| znf385b | 1.3E-60 | 1.895462 | 0.1 | 0.023 | 4.34E-56 | Chondrogenic Fibroblasts | znf385b |
| prkd21 | 1.5E-60 | 1.0692726 | 0.294 | 0.125 | 4.74E-56 | Chondrogenic Fibroblasts | prkd2 |
| ppfibp1a2 | 5.7E-60 | 1.3172122 | 0.166 | 0.053 | 1.85E-55 | Chondrogenic Fibroblasts | ppfibp1a |
| tacc12 | 7.1E-60 | 0.9109024 | 0.418 | 0.211 | 2.3E-55 | Chondrogenic Fibroblasts | tacc1 |
| thbs2a6 | 8E-60 | 0.9360153 | 0.404 | 0.198 | 2.59E-55 | Chondrogenic Fibroblasts | thbs2a |
| hhla2a.13 | 8E-60 | 1.3997867 | 0.213 | 0.079 | 2.6E-55 | Chondrogenic Fibroblasts | hhla2a.1 |
| scara34 | 9.3E-60 | 1.3161345 | 0.297 | 0.13 | 3.03E-55 | Chondrogenic Fibroblasts | scara3 |
| chmp4bb2 | 1.2E-59 | 0.9168985 | 0.332 | 0.149 | 3.93E-55 | Chondrogenic Fibroblasts | chmp4bb |
| golim4a.11 | 3.2E-59 | 1.0826349 | 0.241 | 0.094 | 1.02E-54 | Chondrogenic Fibroblasts | golim4a.1 |
| vim4 | 4.4E-59 | 0.7166349 | 0.61 | 0.364 | 1.44E-54 | Chondrogenic Fibroblasts | vim |
| arhgef1a2 | 8.8E-59 | 1.5756516 | 0.174 | 0.058 | 2.86E-54 | Chondrogenic Fibroblasts | arhgef1a |

| | | | | | | | |
|---|---|---|---|---|---|---|---|
| chmp4ba1 | 1.4E-58 | 0.8390021 | 0.412 | 0.207 | 4.61E-54 | Chondrogenic Fibroblasts | chmp4ba |
| ENSDARG000001018 | 1.8E-58 | 1.4432007 | 0.143 | 0.043 | 5.94E-54 | Chondrogenic Fibroblasts | ENSDARG00000101 |
| ivns1abpb | 3.7E-58 | 1.6272808 | 0.129 | 0.036 | 1.2E-53 | Chondrogenic Fibroblasts | ivns1abpb |
| RAPH13 | 3.8E-58 | 1.1608733 | 0.229 | 0.087 | 1.24E-53 | Chondrogenic Fibroblasts | RAPH1 |
| rab18a1 | 4.2E-58 | 0.9370229 | 0.329 | 0.15 | 1.37E-53 | Chondrogenic Fibroblasts | rab18a |
| arfgap22 | 8E-58 | 1.0633855 | 0.251 | 0.101 | 2.6E-53 | Chondrogenic Fibroblasts | arfgap2 |
| col27a1b5 | 2.3E-57 | 0.7130967 | 0.565 | 0.332 | 7.6E-53 | Chondrogenic Fibroblasts | col27a1b |
| trak1a1 | 4.2E-57 | 1.2845246 | 0.173 | 0.058 | 1.36E-52 | Chondrogenic Fibroblasts | trak1a |
| scxa4 | 5.7E-57 | 1.6216325 | 0.21 | 0.08 | 1.86E-52 | Chondrogenic Fibroblasts | scxa |
| zfyve21 | 5.9E-57 | 0.8743258 | 0.424 | 0.219 | 1.91E-52 | Chondrogenic Fibroblasts | zfyve21 |
| cycsb2 | 1.1E-56 | 0.5041792 | 0.619 | 0.368 | 3.45E-52 | Chondrogenic Fibroblasts | cycsb |
| fermt21 | 1.2E-56 | 0.9970984 | 0.24 | 0.094 | 3.79E-52 | Chondrogenic Fibroblasts | fermt2 |
| h2az2b2 | 1.2E-56 | 0.7154304 | 0.627 | 0.392 | 3.8E-52 | Chondrogenic Fibroblasts | h2az2b |
| vaspb4 | 1.4E-56 | 1.0752916 | 0.241 | 0.096 | 4.42E-52 | Chondrogenic Fibroblasts | vaspb |
| peli1a4 | 2.6E-56 | 0.8681878 | 0.29 | 0.125 | 8.34E-52 | Chondrogenic Fibroblasts | peli1a |
| h3f3d4 | 3.7E-56 | 0.5277723 | 0.903 | 0.724 | 1.21E-51 | Chondrogenic Fibroblasts | h3f3d |
| clic22 | 4.2E-56 | 1.1317443 | 0.441 | 0.243 | 1.36E-51 | Chondrogenic Fibroblasts | clic2 |
| ppp1cb1 | 1.1E-55 | 0.9015855 | 0.383 | 0.191 | 3.63E-51 | Chondrogenic Fibroblasts | ppp1cb |
| rbm8a2 | 1.4E-55 | 0.6509072 | 0.592 | 0.347 | 4.51E-51 | Chondrogenic Fibroblasts | rbm8a |
| ecm25 | 1.6E-55 | 0.8382236 | 0.517 | 0.293 | 5.31E-51 | Chondrogenic Fibroblasts | ecm2 |
| sec11a2 | 2E-55 | 0.7497545 | 0.566 | 0.337 | 6.56E-51 | Chondrogenic Fibroblasts | sec11a |
| rnaseka3 | 2.4E-55 | 1.0929201 | 0.249 | 0.103 | 7.78E-51 | Chondrogenic Fibroblasts | rnaseka |
| calr3a3 | 4.8E-55 | 0.7369919 | 0.585 | 0.357 | 1.55E-50 | Chondrogenic Fibroblasts | calr3a |
| chmp5b1 | 5.1E-55 | 0.8257524 | 0.431 | 0.225 | 1.65E-50 | Chondrogenic Fibroblasts | chmp5b |
| atp5f1b3 | 5.4E-55 | 0.5630584 | 0.773 | 0.553 | 1.77E-50 | Chondrogenic Fibroblasts | atp5f1b |
| si:ch1073-303k11.2 | 6.7E-55 | 1.5581485 | 0.111 | 0.029 | 2.17E-50 | Chondrogenic Fibroblasts | si:ch1073-303k11.2 |
| sgcd1 | 1.6E-54 | 1.3167095 | 0.182 | 0.065 | 5.17E-50 | Chondrogenic Fibroblasts | sgcd |
| elf11 | 1.6E-54 | 1.1138236 | 0.201 | 0.074 | 5.2E-50 | Chondrogenic Fibroblasts | elf1 |
| cpne31 | 1.8E-54 | 1.0334853 | 0.253 | 0.104 | 5.99E-50 | Chondrogenic Fibroblasts | cpne3 |
| ube2na1 | 2E-54 | 0.7423549 | 0.499 | 0.278 | 6.59E-50 | Chondrogenic Fibroblasts | ube2na |
| hnrnpaba1 | 2.7E-54 | 0.5769443 | 0.811 | 0.6 | 8.69E-50 | Chondrogenic Fibroblasts | hnrnpaba |
| ywhaqa2 | 4E-54 | 0.8368018 | 0.477 | 0.268 | 1.29E-49 | Chondrogenic Fibroblasts | ywhaqa |
| rtn31 | 5.8E-54 | 0.7599894 | 0.43 | 0.222 | 1.87E-49 | Chondrogenic Fibroblasts | rtn3 |
| iqgap12 | 6.4E-54 | 0.7852429 | 0.429 | 0.221 | 2.09E-49 | Chondrogenic Fibroblasts | iqgap1 |
| sept10 | 7.1E-54 | 0.7319964 | 0.523 | 0.297 | 2.32E-49 | Chondrogenic Fibroblasts | sept10 |
| fkbp143 | 1.2E-53 | 0.7450546 | 0.547 | 0.327 | 4.05E-49 | Chondrogenic Fibroblasts | fkbp14 |
| bzw1a1 | 1.3E-53 | 0.6346605 | 0.543 | 0.311 | 4.29E-49 | Chondrogenic Fibroblasts | bzw1a |
| ugdh3 | 1.3E-53 | 1.1354407 | 0.35 | 0.174 | 4.33E-49 | Chondrogenic Fibroblasts | ugdh |
| tmem50a3 | 2.1E-53 | 0.7879346 | 0.438 | 0.23 | 6.89E-49 | Chondrogenic Fibroblasts | tmem50a |
| elocb2 | 2.4E-53 | 0.7537697 | 0.509 | 0.291 | 7.71E-49 | Chondrogenic Fibroblasts | elocb |
| rin24 | 2.6E-53 | 1.1097222 | 0.257 | 0.109 | 8.31E-49 | Chondrogenic Fibroblasts | rin2 |
| pfn2l1 | 6E-53 | 0.6064546 | 0.701 | 0.472 | 1.94E-48 | Chondrogenic Fibroblasts | pfn2l |
| fam3a2 | 7.3E-53 | 0.9892148 | 0.3 | 0.137 | 2.37E-48 | Chondrogenic Fibroblasts | fam3a |
| spcs22 | 7.7E-53 | 0.6904826 | 0.499 | 0.281 | 2.5E-48 | Chondrogenic Fibroblasts | spcs2 |
| rpn12 | 9E-53 | 0.8925208 | 0.401 | 0.208 | 2.92E-48 | Chondrogenic Fibroblasts | rpn1 |
| tmsb24 | 2E-52 | 1.1824931 | 0.231 | 0.094 | 6.64E-48 | Chondrogenic Fibroblasts | tmsb2 |
| eif5a22 | 2.4E-52 | 0.6012219 | 0.699 | 0.462 | 7.74E-48 | Chondrogenic Fibroblasts | eif5a2 |
| GCA2 | 2.7E-52 | 0.7488302 | 0.51 | 0.291 | 8.92E-48 | Chondrogenic Fibroblasts | GCA |

| | | | | | | | |
|---|---|---|---|---|---|---|---|
| chsy31 | 3.4E-52 | 1.41075 | 0.129 | 0.038 | 1.1E-47 | Chondrogenic Fibroblasts | chsy3 |
| sec132 | 4.8E-52 | 1.0138089 | 0.276 | 0.122 | 1.57E-47 | Chondrogenic Fibroblasts | sec13 |
| ssr34 | 5.4E-52 | 0.5638785 | 0.801 | 0.573 | 1.75E-47 | Chondrogenic Fibroblasts | ssr3 |
| nucb2b3 | 7.4E-52 | 0.9973912 | 0.265 | 0.115 | 2.39E-47 | Chondrogenic Fibroblasts | nucb2b |
| glrx2 | 8.1E-52 | 0.6305468 | 0.588 | 0.352 | 2.62E-47 | Chondrogenic Fibroblasts | glrx |
| tmed22 | 8.6E-52 | 0.6959111 | 0.615 | 0.386 | 2.8E-47 | Chondrogenic Fibroblasts | tmed2 |
| gng51 | 1E-51 | 0.7614671 | 0.47 | 0.259 | 3.35E-47 | Chondrogenic Fibroblasts | gng5 |
| dctn32 | 1.1E-51 | 0.9024134 | 0.314 | 0.147 | 3.48E-47 | Chondrogenic Fibroblasts | dctn3 |
| agpat41 | 1.6E-51 | 1.0120129 | 0.177 | 0.062 | 5.27E-47 | Chondrogenic Fibroblasts | agpat4 |
| hsbp1b2 | 1.7E-51 | 0.7320173 | 0.477 | 0.266 | 5.66E-47 | Chondrogenic Fibroblasts | hsbp1b |
| cd82b1 | 2.2E-51 | 0.6688631 | 0.604 | 0.37 | 7.22E-47 | Chondrogenic Fibroblasts | cd82b |
| fuca23 | 2.5E-51 | 0.8836713 | 0.308 | 0.142 | 8.07E-47 | Chondrogenic Fibroblasts | fuca2 |
| dpysl33 | 3.1E-51 | 0.7238676 | 0.333 | 0.158 | 1.02E-46 | Chondrogenic Fibroblasts | dpysl3 |
| tmem167a1 | 3.2E-51 | 0.7414637 | 0.457 | 0.249 | 1.04E-46 | Chondrogenic Fibroblasts | tmem167a |
| yipf41 | 3.3E-51 | 0.9258412 | 0.352 | 0.176 | 1.06E-46 | Chondrogenic Fibroblasts | yipf4 |
| atox14 | 3.7E-51 | 0.8566818 | 0.451 | 0.253 | 1.19E-46 | Chondrogenic Fibroblasts | atox1 |
| hpcal1 | 3.8E-51 | 1.6149859 | 0.104 | 0.028 | 1.24E-46 | Chondrogenic Fibroblasts | hpcal1 |
| abracl1 | 3.9E-51 | 0.8989398 | 0.251 | 0.106 | 1.26E-46 | Chondrogenic Fibroblasts | abracl |
| tagln22 | 4.1E-51 | 0.5864723 | 0.842 | 0.661 | 1.32E-46 | Chondrogenic Fibroblasts | tagln2 |
| prkcsh2 | 6.7E-51 | 0.905535 | 0.29 | 0.132 | 2.19E-46 | Chondrogenic Fibroblasts | prkcsh |
| gnpnat13 | 8.6E-51 | 1.1749044 | 0.216 | 0.087 | 2.8E-46 | Chondrogenic Fibroblasts | gnpnat1 |
| chmp1b2 | 1.1E-50 | 0.9632146 | 0.265 | 0.115 | 3.63E-46 | Chondrogenic Fibroblasts | chmp1b |
| tmem30aa1 | 2E-50 | 0.8345446 | 0.328 | 0.157 | 6.45E-46 | Chondrogenic Fibroblasts | tmem30aa |
| actr11 | 2E-50 | 0.8811345 | 0.304 | 0.141 | 6.6E-46 | Chondrogenic Fibroblasts | actr1 |
| zmat22 | 2.2E-50 | 0.8548293 | 0.398 | 0.21 | 7.1E-46 | Chondrogenic Fibroblasts | zmat2 |
| glb1l1 | 2.6E-50 | 1.4908101 | 0.123 | 0.037 | 8.38E-46 | Chondrogenic Fibroblasts | glb1l |
| TXN1 | 2.8E-50 | 0.4026612 | 0.907 | 0.743 | 9.14E-46 | Chondrogenic Fibroblasts | TXN |
| tgfb32 | 3.3E-50 | 1.0629901 | 0.202 | 0.078 | 1.08E-45 | Chondrogenic Fibroblasts | tgfb3 |
| taf132 | 4.1E-50 | 0.7926535 | 0.405 | 0.213 | 1.34E-45 | Chondrogenic Fibroblasts | taf13 |
| serp14 | 4.2E-50 | 0.6944061 | 0.527 | 0.31 | 1.36E-45 | Chondrogenic Fibroblasts | serp1 |
| dynll12 | 4.8E-50 | 0.8898147 | 0.615 | 0.422 | 1.55E-45 | Chondrogenic Fibroblasts | dynll1 |
| cox7c2 | 1.1E-49 | 0.6984028 | 0.511 | 0.295 | 3.63E-45 | Chondrogenic Fibroblasts | cox7c |
| relt1 | 1.1E-49 | 1.5997158 | 0.11 | 0.031 | 3.68E-45 | Chondrogenic Fibroblasts | relt |
| hagh1 | 1.3E-49 | 1.312812 | 0.167 | 0.059 | 4.16E-45 | Chondrogenic Fibroblasts | hagh |
| ost43 | 1.5E-49 | 0.9538869 | 0.309 | 0.147 | 5.02E-45 | Chondrogenic Fibroblasts | ost4 |
| bloc1s64 | 1.6E-49 | 0.979609 | 0.348 | 0.176 | 5.09E-45 | Chondrogenic Fibroblasts | bloc1s6 |
| sorbs32 | 1.7E-49 | 0.7475095 | 0.371 | 0.186 | 5.39E-45 | Chondrogenic Fibroblasts | sorbs3 |
| si:dkey-28b4.72 | 1.9E-49 | 0.8618612 | 0.376 | 0.195 | 6.2E-45 | Chondrogenic Fibroblasts | si:dkey-28b4.7 |
| capza1a3 | 2E-49 | 0.6967569 | 0.477 | 0.267 | 6.56E-45 | Chondrogenic Fibroblasts | capza1a |
| ufm13 | 2.1E-49 | 0.7681596 | 0.366 | 0.185 | 6.8E-45 | Chondrogenic Fibroblasts | ufm1 |
| mfge8b2 | 3.1E-49 | 1.1076675 | 0.158 | 0.054 | 9.95E-45 | Chondrogenic Fibroblasts | mfge8b |
| irf2bp2b3 | 3.1E-49 | 0.7834821 | 0.394 | 0.205 | 1.02E-44 | Chondrogenic Fibroblasts | irf2bp2b |
| scpep12 | 3.2E-49 | 0.9247536 | 0.282 | 0.128 | 1.05E-44 | Chondrogenic Fibroblasts | scpep1 |
| nfatc2a1 | 3.9E-49 | 1.4589357 | 0.118 | 0.034 | 1.27E-44 | Chondrogenic Fibroblasts | nfatc2a |
| si:ch73-46j18.56 | 4.8E-49 | 0.4026387 | 0.814 | 0.597 | 1.55E-44 | Chondrogenic Fibroblasts | si:ch73-46j18.5 |
| glud1a2 | 5.2E-49 | 0.586937 | 0.669 | 0.435 | 1.7E-44 | Chondrogenic Fibroblasts | glud1a |
| arpc41 | 5.3E-49 | 0.7902891 | 0.35 | 0.174 | 1.71E-44 | Chondrogenic Fibroblasts | arpc4 |
| rab34a2 | 7.7E-49 | 0.9727738 | 0.261 | 0.115 | 2.51E-44 | Chondrogenic Fibroblasts | rab34a |

| | | | | | | | |
|---|---|---|---|---|---|---|---|
| sumo3a2 | 8.1E-49 | 0.7069312 | 0.495 | 0.281 | 2.63E-44 | Chondrogenic Fibroblasts | sumo3a |
| prdx43 | 9.5E-49 | 0.6676074 | 0.582 | 0.359 | 3.1E-44 | Chondrogenic Fibroblasts | prdx4 |
| chst113 | 1.1E-48 | 1.6912969 | 0.131 | 0.042 | 3.72E-44 | Chondrogenic Fibroblasts | chst11 |
| dync1li21 | 1.9E-48 | 0.7391893 | 0.333 | 0.161 | 6.08E-44 | Chondrogenic Fibroblasts | dync1li2 |
| angptl72 | 2.4E-48 | 1.1578758 | 0.367 | 0.199 | 7.82E-44 | Chondrogenic Fibroblasts | angptl7 |
| tcp11l21 | 2.4E-48 | 1.2238462 | 0.183 | 0.069 | 7.92E-44 | Chondrogenic Fibroblasts | tcp11l2 |
| actr2a1 | 2.8E-48 | 0.7015611 | 0.39 | 0.201 | 9.21E-44 | Chondrogenic Fibroblasts | actr2a |
| cxcl142 | 3.7E-48 | 0.964113 | 0.425 | 0.245 | 1.21E-43 | Chondrogenic Fibroblasts | cxcl14 |
| ndufc22 | 4.1E-48 | 0.6755458 | 0.483 | 0.273 | 1.32E-43 | Chondrogenic Fibroblasts | ndufc2 |
| rhoca1 | 4.3E-48 | 0.7139729 | 0.455 | 0.253 | 1.39E-43 | Chondrogenic Fibroblasts | rhoca |
| dgkaa | 6.9E-48 | 1.6050184 | 0.114 | 0.033 | 2.23E-43 | Chondrogenic Fibroblasts | dgkaa |
| fmnl3 | 7.5E-48 | 0.8592204 | 0.292 | 0.135 | 2.45E-43 | Chondrogenic Fibroblasts | fmnl3 |
| sumo11 | 7.7E-48 | 0.8579427 | 0.322 | 0.157 | 2.5E-43 | Chondrogenic Fibroblasts | sumo1 |
| PTPRD | 7.8E-48 | 1.3108875 | 0.134 | 0.043 | 2.54E-43 | Chondrogenic Fibroblasts | PTPRD |
| STMP13 | 8.2E-48 | 0.7001412 | 0.455 | 0.253 | 2.68E-43 | Chondrogenic Fibroblasts | STMP1 |
| nid2a2 | 1.1E-47 | 1.3045342 | 0.186 | 0.071 | 3.55E-43 | Chondrogenic Fibroblasts | nid2a |
| zgc:154058 | 1.3E-47 | 1.4670749 | 0.109 | 0.031 | 4.14E-43 | Chondrogenic Fibroblasts | zgc:154058 |
| CABZ01068367.15 | 1.5E-47 | 0.6659896 | 0.605 | 0.388 | 4.78E-43 | Chondrogenic Fibroblasts | CABZ01068367.1 |
| fhl3a1 | 1.6E-47 | 1.0293695 | 0.178 | 0.066 | 5.31E-43 | Chondrogenic Fibroblasts | fhl3a |
| pafah1b1b1 | 2.7E-47 | 0.7083443 | 0.39 | 0.203 | 8.94E-43 | Chondrogenic Fibroblasts | pafah1b1b |
| pfn23 | 3.5E-47 | 0.5649365 | 0.752 | 0.534 | 1.13E-42 | Chondrogenic Fibroblasts | pfn2 |
| ctsba2 | 3.9E-47 | 0.7812711 | 0.587 | 0.379 | 1.28E-42 | Chondrogenic Fibroblasts | ctsba |
| gmfb1 | 4.4E-47 | 0.880245 | 0.298 | 0.141 | 1.42E-42 | Chondrogenic Fibroblasts | gmfb |
| zgc:1102394 | 4.4E-47 | 0.8231529 | 0.414 | 0.228 | 1.44E-42 | Chondrogenic Fibroblasts | zgc:110239 |
| atp5mc13 | 4.7E-47 | 0.640905 | 0.515 | 0.306 | 1.54E-42 | Chondrogenic Fibroblasts | atp5mc1 |
| zgc:1629443 | 5E-47 | 0.7007195 | 0.496 | 0.289 | 1.64E-42 | Chondrogenic Fibroblasts | zgc:162944 |
| zgc:559432 | 6.1E-47 | 1.1581574 | 0.185 | 0.071 | 1.98E-42 | Chondrogenic Fibroblasts | zgc:55943 |
| si:dkey-17o15.22 | 7.5E-47 | 1.0019776 | 0.201 | 0.08 | 2.43E-42 | Chondrogenic Fibroblasts | si:dkey-17o15.2 |
| atp5f1c3 | 1.1E-46 | 0.7249666 | 0.446 | 0.252 | 3.5E-42 | Chondrogenic Fibroblasts | atp5f1c |
| gpia2 | 1.1E-46 | 1.1295033 | 0.268 | 0.125 | 3.73E-42 | Chondrogenic Fibroblasts | gpia |
| wbp2nl2 | 1.3E-46 | 0.6353639 | 0.499 | 0.286 | 4.31E-42 | Chondrogenic Fibroblasts | wbp2nl |
| LO018188.15 | 1.6E-46 | 0.9139183 | 0.68 | 0.473 | 5.16E-42 | Chondrogenic Fibroblasts | LO018188.1 |
| crtap2 | 2.7E-46 | 0.7027723 | 0.522 | 0.315 | 8.89E-42 | Chondrogenic Fibroblasts | crtap |
| zgc:565253 | 2.8E-46 | 0.8192067 | 0.311 | 0.151 | 9.18E-42 | Chondrogenic Fibroblasts | zgc:56525 |
| vldlr | 3.9E-46 | 1.4393837 | 0.116 | 0.035 | 1.27E-41 | Chondrogenic Fibroblasts | vldlr |
| ndufs63 | 1.4E-45 | 0.8738439 | 0.291 | 0.138 | 4.47E-41 | Chondrogenic Fibroblasts | ndufs6 |
| serinc54 | 1.4E-45 | 0.681163 | 0.43 | 0.235 | 4.54E-41 | Chondrogenic Fibroblasts | serinc5 |
| fkbp10a2 | 1.4E-45 | 0.8791724 | 0.283 | 0.131 | 4.65E-41 | Chondrogenic Fibroblasts | fkbp10a |
| si:ch211-132b12.74 | 1.7E-45 | 0.7643525 | 0.316 | 0.152 | 5.52E-41 | Chondrogenic Fibroblasts | si:ch211-132b12.7 |
| hmgn61 | 2E-45 | 0.5445773 | 0.79 | 0.599 | 6.4E-41 | Chondrogenic Fibroblasts | hmgn6 |
| net14 | 2.9E-45 | 0.7754434 | 0.57 | 0.368 | 9.5E-41 | Chondrogenic Fibroblasts | net1 |
| pdia32 | 3.8E-45 | 0.4959892 | 0.861 | 0.682 | 1.25E-40 | Chondrogenic Fibroblasts | pdia3 |
| fthl311 | 4.8E-45 | 2.0645913 | 0.114 | 0.035 | 1.55E-40 | Chondrogenic Fibroblasts | fthl31 |
| mgp2 | 6.5E-45 | 1.4862724 | 0.299 | 0.156 | 2.11E-40 | Chondrogenic Fibroblasts | mgp |
| PAMR13 | 7.8E-45 | 0.8314264 | 0.259 | 0.117 | 2.54E-40 | Chondrogenic Fibroblasts | PAMR1 |
| si:ch211-76l23.42 | 9.5E-45 | 0.7801998 | 0.354 | 0.183 | 3.09E-40 | Chondrogenic Fibroblasts | si:ch211-76l23.4 |
| psmb71 | 1.3E-44 | 0.6918821 | 0.444 | 0.25 | 4.37E-40 | Chondrogenic Fibroblasts | psmb7 |
| acbd32 | 2.4E-44 | 0.9072954 | 0.279 | 0.133 | 7.92E-40 | Chondrogenic Fibroblasts | acbd3 |

| | | | | | | | |
|---|---|---|---|---|---|---|---|
| ndufb31 | 2.5E-44 | 0.7010717 | 0.471 | 0.274 | 8.11E-40 | Chondrogenic Fibroblasts | ndufb3 |
| tuba8l31 | 2.6E-44 | 0.6035002 | 0.627 | 0.42 | 8.45E-40 | Chondrogenic Fibroblasts | tuba8l3 |
| dnajc14 | 3.7E-44 | 0.6674762 | 0.477 | 0.278 | 1.19E-39 | Chondrogenic Fibroblasts | dnajc1 |
| cnn3a3 | 5.6E-44 | 0.5710184 | 0.689 | 0.471 | 1.83E-39 | Chondrogenic Fibroblasts | cnn3a |
| matn41 | 7.6E-44 | 1.6813086 | 0.129 | 0.043 | 2.48E-39 | Chondrogenic Fibroblasts | matn4 |
| emx25 | 9.1E-44 | 1.0417268 | 0.212 | 0.09 | 2.98E-39 | Chondrogenic Fibroblasts | emx2 |
| eif4g2a1 | 1.2E-43 | 0.7301047 | 0.351 | 0.181 | 3.98E-39 | Chondrogenic Fibroblasts | eif4g2a |
| ssr24 | 1.3E-43 | 0.5130463 | 0.853 | 0.687 | 4.13E-39 | Chondrogenic Fibroblasts | ssr2 |
| sem11 | 1.4E-43 | 0.5727215 | 0.631 | 0.413 | 4.61E-39 | Chondrogenic Fibroblasts | sem1 |
| cnih12 | 1.4E-43 | 0.9417012 | 0.218 | 0.093 | 4.7E-39 | Chondrogenic Fibroblasts | cnih1 |
| yif1a1 | 2.8E-43 | 0.8536718 | 0.262 | 0.121 | 9.21E-39 | Chondrogenic Fibroblasts | yif1a |
| fbxo253 | 3.1E-43 | 1.0117035 | 0.19 | 0.076 | 1.02E-38 | Chondrogenic Fibroblasts | fbxo25 |
| tuba1a2 | 3.3E-43 | 1.0242156 | 0.206 | 0.086 | 1.07E-38 | Chondrogenic Fibroblasts | tuba1a |
| pamr12 | 3.3E-43 | 1.0098748 | 0.239 | 0.107 | 1.09E-38 | Chondrogenic Fibroblasts | pamr1 |
| txn4 | 3.4E-43 | 0.8971331 | 0.549 | 0.369 | 1.09E-38 | Chondrogenic Fibroblasts | txn |
| nos1apa4 | 7.1E-43 | 1.0235842 | 0.236 | 0.106 | 2.32E-38 | Chondrogenic Fibroblasts | nos1apa |
| ptgs2a4 | 7.8E-43 | 1.3035679 | 0.266 | 0.127 | 2.52E-38 | Chondrogenic Fibroblasts | ptgs2a |
| utrn2 | 8.7E-43 | 0.6304889 | 0.358 | 0.184 | 2.84E-38 | Chondrogenic Fibroblasts | utrn |
| snrnp273 | 9.5E-43 | 0.9547395 | 0.278 | 0.135 | 3.09E-38 | Chondrogenic Fibroblasts | snrnp27 |
| ralba1 | 1.2E-42 | 0.9812128 | 0.208 | 0.088 | 3.77E-38 | Chondrogenic Fibroblasts | ralba |
| laptm4a2 | 1.2E-42 | 0.6560504 | 0.503 | 0.302 | 3.87E-38 | Chondrogenic Fibroblasts | laptm4a |
| grb2b2 | 1.3E-42 | 0.7567429 | 0.316 | 0.159 | 4.08E-38 | Chondrogenic Fibroblasts | grb2b |
| golim4a1 | 1.7E-42 | 0.8849766 | 0.251 | 0.115 | 5.37E-38 | Chondrogenic Fibroblasts | golim4a |
| rbx11 | 1.7E-42 | 0.516852 | 0.674 | 0.448 | 5.39E-38 | Chondrogenic Fibroblasts | rbx1 |
| col4a23 | 2.7E-42 | 0.5062726 | 0.392 | 0.215 | 8.82E-38 | Chondrogenic Fibroblasts | col4a2 |
| ergic22 | 2.7E-42 | 0.7901581 | 0.314 | 0.157 | 8.92E-38 | Chondrogenic Fibroblasts | ergic2 |
| oaz1b4 | 2.8E-42 | 0.6694811 | 0.502 | 0.303 | 9.15E-38 | Chondrogenic Fibroblasts | oaz1b |
| bud312 | 5.8E-42 | 0.683353 | 0.378 | 0.204 | 1.9E-37 | Chondrogenic Fibroblasts | bud31 |
| osbpl92 | 5.9E-42 | 0.8151306 | 0.306 | 0.153 | 1.93E-37 | Chondrogenic Fibroblasts | osbpl9 |
| slc25a3b3 | 7.4E-42 | 0.5031914 | 0.747 | 0.546 | 2.42E-37 | Chondrogenic Fibroblasts | slc25a3b |
| rasal21 | 1.1E-41 | 0.9743996 | 0.176 | 0.069 | 3.64E-37 | Chondrogenic Fibroblasts | rasal2 |
| chchd22 | 1.9E-41 | 0.942107 | 0.212 | 0.091 | 6.15E-37 | Chondrogenic Fibroblasts | chchd2 |
| mpc13 | 2.3E-41 | 0.7651026 | 0.319 | 0.162 | 7.54E-37 | Chondrogenic Fibroblasts | mpc1 |
| srpra2 | 2.3E-41 | 0.69301 | 0.391 | 0.216 | 7.62E-37 | Chondrogenic Fibroblasts | srpra |
| med301 | 2.7E-41 | 1.0272937 | 0.247 | 0.115 | 8.94E-37 | Chondrogenic Fibroblasts | med30 |
| zfyve9a2 | 3.1E-41 | 0.8287277 | 0.303 | 0.152 | 1.02E-36 | Chondrogenic Fibroblasts | zfyve9a |
| acsl4b2 | 3.6E-41 | 0.9667143 | 0.268 | 0.129 | 1.17E-36 | Chondrogenic Fibroblasts | acsl4b |
| slc38a5a3 | 3.8E-41 | 1.2432921 | 0.172 | 0.068 | 1.23E-36 | Chondrogenic Fibroblasts | slc38a5a |
| vamp32 | 3.9E-41 | 0.7232468 | 0.428 | 0.246 | 1.26E-36 | Chondrogenic Fibroblasts | vamp3 |
| pkd1a | 4.1E-41 | 1.0396877 | 0.177 | 0.07 | 1.35E-36 | Chondrogenic Fibroblasts | pkd1a |
| atp5pf2 | 4.3E-41 | 0.5037776 | 0.62 | 0.4 | 1.4E-36 | Chondrogenic Fibroblasts | atp5pf |
| snx43 | 6.4E-41 | 1.0006493 | 0.208 | 0.09 | 2.07E-36 | Chondrogenic Fibroblasts | snx4 |
| arpc21 | 6.7E-41 | 0.6093869 | 0.643 | 0.439 | 2.19E-36 | Chondrogenic Fibroblasts | arpc2 |
| tmed102 | 8.3E-41 | 0.5730647 | 0.54 | 0.333 | 2.69E-36 | Chondrogenic Fibroblasts | tmed10 |
| ckap43 | 8.4E-41 | 0.5406241 | 0.713 | 0.514 | 2.73E-36 | Chondrogenic Fibroblasts | ckap4 |
| map1lc3b2 | 9.9E-41 | 0.6820699 | 0.445 | 0.259 | 3.22E-36 | Chondrogenic Fibroblasts | map1lc3b |
| rab18b1 | 1E-40 | 0.9023917 | 0.243 | 0.113 | 3.29E-36 | Chondrogenic Fibroblasts | rab18b |
| gpx4b2 | 1E-40 | 0.6644528 | 0.48 | 0.291 | 3.33E-36 | Chondrogenic Fibroblasts | gpx4b |

| | | | | | | | |
|---|---|---|---|---|---|---|---|
| psmb31 | 1.3E-40 | 0.5385045 | 0.445 | 0.252 | 4.17E-36 | Chondrogenic Fibroblasts | psmb3 |
| gng23 | 1.6E-40 | 0.8867258 | 0.235 | 0.106 | 5.3E-36 | Chondrogenic Fibroblasts | gng2 |
| zgc:1935412 | 2.5E-40 | 0.5440117 | 0.596 | 0.389 | 8.16E-36 | Chondrogenic Fibroblasts | zgc:193541 |
| arl11 | 3.1E-40 | 0.7204321 | 0.317 | 0.162 | 1.01E-35 | Chondrogenic Fibroblasts | arl1 |
| p4hb3 | 3.8E-40 | 0.5378558 | 0.762 | 0.569 | 1.23E-35 | Chondrogenic Fibroblasts | p4hb |
| spcs12 | 6.1E-40 | 0.6159825 | 0.624 | 0.432 | 1.98E-35 | Chondrogenic Fibroblasts | spcs1 |
| si:ch211-207i1.22 | 6.1E-40 | 1.0392356 | 0.192 | 0.08 | 1.99E-35 | Chondrogenic Fibroblasts | si:ch211-207i1.2 |
| gas1a4 | 6.3E-40 | 0.8222321 | 0.308 | 0.157 | 2.05E-35 | Chondrogenic Fibroblasts | gas1a |
| hivep3a4 | 6.5E-40 | 0.8774628 | 0.234 | 0.107 | 2.13E-35 | Chondrogenic Fibroblasts | hivep3a |
| si:dkey-16l2.162 | 7.6E-40 | 0.9181238 | 0.246 | 0.115 | 2.47E-35 | Chondrogenic Fibroblasts | si:dkey-16l2.16 |
| csnk1g2b1 | 8.7E-40 | 0.6731147 | 0.326 | 0.167 | 2.83E-35 | Chondrogenic Fibroblasts | csnk1g2b |
| fbln23 | 9.6E-40 | 0.7915102 | 0.255 | 0.12 | 3.14E-35 | Chondrogenic Fibroblasts | fbln2 |
| uggt22 | 1E-39 | 1.2189684 | 0.143 | 0.052 | 3.25E-35 | Chondrogenic Fibroblasts | uggt2 |
| rer12 | 1.3E-39 | 0.7335989 | 0.371 | 0.202 | 4.07E-35 | Chondrogenic Fibroblasts | rer1 |
| rhocb1 | 1.4E-39 | 0.7958501 | 0.289 | 0.144 | 4.59E-35 | Chondrogenic Fibroblasts | rhocb |
| itgb43 | 1.6E-39 | 0.70628 | 0.499 | 0.308 | 5.23E-35 | Chondrogenic Fibroblasts | itgb4 |
| trappc33 | 1.6E-39 | 1.0236133 | 0.227 | 0.104 | 5.26E-35 | Chondrogenic Fibroblasts | trappc3 |
| ier3ip13 | 1.9E-39 | 0.7383836 | 0.404 | 0.232 | 6.1E-35 | Chondrogenic Fibroblasts | ier3ip1 |
| ubl3a3 | 2E-39 | 0.5930133 | 0.534 | 0.332 | 6.63E-35 | Chondrogenic Fibroblasts | ubl3a |
| uqcrfs12 | 3.1E-39 | 0.6453063 | 0.39 | 0.217 | 1E-34 | Chondrogenic Fibroblasts | uqcrfs1 |
| ndufa61 | 3.6E-39 | 0.6040261 | 0.406 | 0.227 | 1.17E-34 | Chondrogenic Fibroblasts | ndufa6 |
| ywhabb1 | 4E-39 | 0.5109794 | 0.603 | 0.392 | 1.29E-34 | Chondrogenic Fibroblasts | ywhabb |
| hif1ab2 | 4E-39 | 0.5848974 | 0.578 | 0.37 | 1.31E-34 | Chondrogenic Fibroblasts | hif1ab |
| si:ch211-288g17.34 | 4.2E-39 | 0.6355059 | 0.601 | 0.399 | 1.38E-34 | Chondrogenic Fibroblasts | si:ch211-288g17.3 |
| chmp2bb2 | 4.4E-39 | 0.8897284 | 0.221 | 0.099 | 1.43E-34 | Chondrogenic Fibroblasts | chmp2bb |
| fli1a1 | 4.6E-39 | 1.1492246 | 0.133 | 0.047 | 1.5E-34 | Chondrogenic Fibroblasts | fli1a |
| ywhae11 | 4.6E-39 | 0.5623354 | 0.523 | 0.325 | 1.5E-34 | Chondrogenic Fibroblasts | ywhae1 |
| krtcap23 | 6.4E-39 | 0.6898897 | 0.45 | 0.266 | 2.08E-34 | Chondrogenic Fibroblasts | krtcap2 |
| sptssa1 | 8.9E-39 | 0.9731196 | 0.238 | 0.112 | 2.91E-34 | Chondrogenic Fibroblasts | sptssa |
| srpx23 | 9.6E-39 | 0.6753444 | 0.43 | 0.25 | 3.12E-34 | Chondrogenic Fibroblasts | srpx2 |
| srp542 | 9.6E-39 | 0.769305 | 0.325 | 0.172 | 3.13E-34 | Chondrogenic Fibroblasts | srp54 |
| il4r.13 | 9.9E-39 | 0.8439712 | 0.291 | 0.149 | 3.21E-34 | Chondrogenic Fibroblasts | il4r.1 |
| tfg2 | 1E-38 | 0.7802131 | 0.344 | 0.186 | 3.28E-34 | Chondrogenic Fibroblasts | tfg |
| slc39a73 | 1.3E-38 | 0.8555078 | 0.242 | 0.114 | 4.24E-34 | Chondrogenic Fibroblasts | slc39a7 |
| ppfibp1b3 | 1.4E-38 | 0.7968949 | 0.281 | 0.14 | 4.65E-34 | Chondrogenic Fibroblasts | ppfibp1b |
| hyou11 | 1.9E-38 | 0.79599 | 0.294 | 0.149 | 6.34E-34 | Chondrogenic Fibroblasts | hyou1 |
| KCTD153 | 2E-38 | 0.492531 | 0.546 | 0.337 | 6.63E-34 | Chondrogenic Fibroblasts | KCTD15 |
| bcl7bb3 | 2.3E-38 | 1.0556577 | 0.2 | 0.088 | 7.61E-34 | Chondrogenic Fibroblasts | bcl7bb |
| cdc42l1 | 3E-38 | 0.5260803 | 0.616 | 0.399 | 9.67E-34 | Chondrogenic Fibroblasts | cdc42l |
| ptp4a2a2 | 3.1E-38 | 0.7588636 | 0.274 | 0.135 | 1.01E-33 | Chondrogenic Fibroblasts | ptp4a2a |
| psph2 | 3.2E-38 | 1.443272 | 0.102 | 0.032 | 1.04E-33 | Chondrogenic Fibroblasts | psph |
| fundc22 | 3.3E-38 | 0.6122949 | 0.49 | 0.298 | 1.07E-33 | Chondrogenic Fibroblasts | fundc2 |
| si:ch211-288d18.11 | 3.7E-38 | 1.0015988 | 0.187 | 0.079 | 1.21E-33 | Chondrogenic Fibroblasts | si:ch211-288d18.1 |
| scamp22 | 3.9E-38 | 0.8406755 | 0.249 | 0.119 | 1.28E-33 | Chondrogenic Fibroblasts | scamp2 |
| scfd13 | 4.8E-38 | 0.7569921 | 0.225 | 0.102 | 1.55E-33 | Chondrogenic Fibroblasts | scfd1 |
| soga12 | 5.6E-38 | 0.6948299 | 0.259 | 0.123 | 1.83E-33 | Chondrogenic Fibroblasts | soga1 |
| fam162a3 | 6.2E-38 | 0.5921329 | 0.414 | 0.235 | 2.03E-33 | Chondrogenic Fibroblasts | fam162a |
| golt1bb3 | 7.2E-38 | 0.9491506 | 0.216 | 0.099 | 2.33E-33 | Chondrogenic Fibroblasts | golt1bb |

| | | | | | | | |
|---|---|---|---|---|---|---|---|
| MED92 | 7.2E-38 | 1.0690183 | 0.168 | 0.068 | 2.33E-33 | Chondrogenic Fibroblasts | MED9 |
| si:ch211-160o17.22 | 7.2E-38 | 1.1566005 | 0.144 | 0.055 | 2.36E-33 | Chondrogenic Fibroblasts | si:ch211-160o17.2 |
| cdc42ep32 | 7.4E-38 | 1.1862234 | 0.153 | 0.06 | 2.4E-33 | Chondrogenic Fibroblasts | cdc42ep3 |
| pdgfab1 | 9.2E-38 | 1.4457339 | 0.11 | 0.036 | 2.99E-33 | Chondrogenic Fibroblasts | pdgfab |
| bcl2l11 | 1.1E-37 | 0.7301001 | 0.301 | 0.154 | 3.47E-33 | Chondrogenic Fibroblasts | bcl2l1 |
| abhd2a1 | 1.3E-37 | 1.0664241 | 0.194 | 0.084 | 4.32E-33 | Chondrogenic Fibroblasts | abhd2a |
| ywhaqb1 | 1.7E-37 | 0.5541905 | 0.724 | 0.533 | 5.57E-33 | Chondrogenic Fibroblasts | ywhaqb |
| parp32 | 1.7E-37 | 0.9660449 | 0.195 | 0.085 | 5.68E-33 | Chondrogenic Fibroblasts | parp3 |
| mif2 | 2.1E-37 | 0.5760246 | 0.588 | 0.396 | 6.91E-33 | Chondrogenic Fibroblasts | mif |
| rhbdl21 | 2.5E-37 | 1.5839418 | 0.102 | 0.033 | 7.97E-33 | Chondrogenic Fibroblasts | rhbdl2 |
| pmp22b2 | 3.9E-37 | 0.5636233 | 0.552 | 0.358 | 1.28E-32 | Chondrogenic Fibroblasts | pmp22b |
| s100a10b2 | 4.1E-37 | 0.4221754 | 0.999 | 0.994 | 1.33E-32 | Chondrogenic Fibroblasts | s100a10b |
| ube2g22 | 4.7E-37 | 0.8838054 | 0.231 | 0.109 | 1.53E-32 | Chondrogenic Fibroblasts | ube2g2 |
| gabarapb3 | 5.1E-37 | 0.5842269 | 0.617 | 0.42 | 1.67E-32 | Chondrogenic Fibroblasts | gabarapb |
| sertad42 | 5.7E-37 | 0.7221852 | 0.35 | 0.192 | 1.87E-32 | Chondrogenic Fibroblasts | sertad4 |
| niban1b4 | 6.8E-37 | 0.7202039 | 0.21 | 0.092 | 2.22E-32 | Chondrogenic Fibroblasts | niban1b |
| reep3b1 | 7.4E-37 | 0.70736 | 0.319 | 0.168 | 2.4E-32 | Chondrogenic Fibroblasts | reep3b |
| CABZ01054965.11 | 9E-37 | 1.0455217 | 0.143 | 0.054 | 2.91E-32 | Chondrogenic Fibroblasts | CABZ01054965.1 |
| chmp6b2 | 9.2E-37 | 0.6537269 | 0.293 | 0.148 | 2.98E-32 | Chondrogenic Fibroblasts | chmp6b |
| eef2b5 | 1.2E-36 | 0.2881514 | 0.977 | 0.908 | 3.79E-32 | Chondrogenic Fibroblasts | eef2b |
| clta2 | 1.3E-36 | 0.8085516 | 0.28 | 0.143 | 4.23E-32 | Chondrogenic Fibroblasts | clta |
| gnpda11 | 1.8E-36 | 1.6516526 | 0.1 | 0.032 | 5.81E-32 | Chondrogenic Fibroblasts | gnpda1 |
| ndufb82 | 2.4E-36 | 0.4978008 | 0.528 | 0.332 | 7.74E-32 | Chondrogenic Fibroblasts | ndufb8 |
| lrpap12 | 3.9E-36 | 0.6482161 | 0.33 | 0.177 | 1.26E-31 | Chondrogenic Fibroblasts | lrpap1 |
| sec23a1 | 4.9E-36 | 0.7544304 | 0.283 | 0.144 | 1.58E-31 | Chondrogenic Fibroblasts | sec23a |
| serpine23 | 6.2E-36 | 0.5977051 | 0.499 | 0.312 | 2.03E-31 | Chondrogenic Fibroblasts | serpine2 |
| mrps361 | 9.8E-36 | 0.7902195 | 0.293 | 0.153 | 3.2E-31 | Chondrogenic Fibroblasts | mrps36 |
| pfkpa | 1.5E-35 | 1.1664517 | 0.118 | 0.041 | 4.92E-31 | Chondrogenic Fibroblasts | pfkpa |
| itga92 | 1.6E-35 | 0.7677388 | 0.254 | 0.124 | 5.19E-31 | Chondrogenic Fibroblasts | itga9 |
| echs12 | 2.1E-35 | 0.7539318 | 0.269 | 0.137 | 6.88E-31 | Chondrogenic Fibroblasts | echs1 |
| bloc1s21 | 2.7E-35 | 1.0769097 | 0.168 | 0.071 | 8.77E-31 | Chondrogenic Fibroblasts | bloc1s2 |
| zgc:1133073 | 2.8E-35 | 0.8954285 | 0.201 | 0.09 | 9.21E-31 | Chondrogenic Fibroblasts | zgc:113307 |
| raph1b4 | 3.8E-35 | 0.9917491 | 0.22 | 0.104 | 1.25E-30 | Chondrogenic Fibroblasts | raph1b |
| skp11 | 3.9E-35 | 0.5100639 | 0.693 | 0.494 | 1.28E-30 | Chondrogenic Fibroblasts | skp1 |
| tomm73 | 4.8E-35 | 0.667662 | 0.345 | 0.191 | 1.55E-30 | Chondrogenic Fibroblasts | tomm7 |
| gas2l11 | 5E-35 | 1.0597573 | 0.149 | 0.059 | 1.64E-30 | Chondrogenic Fibroblasts | gas2l1 |
| ctnnb13 | 7.3E-35 | 0.4721113 | 0.711 | 0.5 | 2.38E-30 | Chondrogenic Fibroblasts | ctnnb1 |
| dbn11 | 7.5E-35 | 0.5878393 | 0.356 | 0.197 | 2.44E-30 | Chondrogenic Fibroblasts | dbn1 |
| chid14 | 8E-35 | 0.9145253 | 0.188 | 0.083 | 2.59E-30 | Chondrogenic Fibroblasts | chid1 |
| rhoaa2 | 8.8E-35 | 0.4767349 | 0.615 | 0.416 | 2.85E-30 | Chondrogenic Fibroblasts | rhoaa |
| smdt1b2 | 1.1E-34 | 0.6041174 | 0.418 | 0.248 | 3.65E-30 | Chondrogenic Fibroblasts | smdt1b |
| atp1a1a.14 | 1.8E-34 | 0.5486023 | 0.505 | 0.318 | 5.79E-30 | Chondrogenic Fibroblasts | atp1a1a.1 |
| serinc12 | 1.8E-34 | 0.551061 | 0.434 | 0.257 | 5.8E-30 | Chondrogenic Fibroblasts | serinc1 |
| uqcrh2 | 1.8E-34 | 0.4500116 | 0.587 | 0.38 | 5.97E-30 | Chondrogenic Fibroblasts | uqcrh |
| ndufa14 | 2.3E-34 | 0.7069756 | 0.356 | 0.201 | 7.38E-30 | Chondrogenic Fibroblasts | ndufa1 |
| sat1b4 | 2.7E-34 | 0.8006264 | 0.225 | 0.108 | 8.63E-30 | Chondrogenic Fibroblasts | sat1b |
| tuba8l44 | 2.7E-34 | 0.4444947 | 0.875 | 0.73 | 8.94E-30 | Chondrogenic Fibroblasts | tuba8l4 |
| baxa.12 | 3.1E-34 | 0.8085304 | 0.213 | 0.099 | 1E-29 | Chondrogenic Fibroblasts | baxa.1 |

| | | | | | | | |
|---|---|---|---|---|---|---|---|
| cox5aa3 | 3.4E-34 | 0.4990159 | 0.463 | 0.282 | 1.12E-29 | Chondrogenic Fibroblasts | cox5aa |
| gmppb2 | 3.6E-34 | 0.8300913 | 0.241 | 0.119 | 1.16E-29 | Chondrogenic Fibroblasts | gmppb |
| mtpn1 | 4E-34 | 0.5084965 | 0.497 | 0.308 | 1.31E-29 | Chondrogenic Fibroblasts | mtpn |
| tmem2142 | 4.3E-34 | 0.7594986 | 0.249 | 0.124 | 1.41E-29 | Chondrogenic Fibroblasts | tmem214 |
| calm2a2 | 5.3E-34 | 0.5287804 | 0.497 | 0.318 | 1.74E-29 | Chondrogenic Fibroblasts | calm2a |
| ppp1r14bb3 | 5.9E-34 | 0.7725982 | 0.28 | 0.147 | 1.91E-29 | Chondrogenic Fibroblasts | ppp1r14bb |
| smco43 | 5.9E-34 | 0.8297753 | 0.268 | 0.139 | 1.93E-29 | Chondrogenic Fibroblasts | smco4 |
| tlcd21 | 6.2E-34 | 1.2226707 | 0.116 | 0.041 | 2.03E-29 | Chondrogenic Fibroblasts | tlcd2 |
| psma11 | 6.3E-34 | 0.5352224 | 0.494 | 0.307 | 2.05E-29 | Chondrogenic Fibroblasts | psma1 |
| hmgn35 | 6.4E-34 | 0.7357868 | 0.301 | 0.159 | 2.07E-29 | Chondrogenic Fibroblasts | hmgn3 |
| mbnl12 | 7.5E-34 | 1.0901261 | 0.162 | 0.068 | 2.43E-29 | Chondrogenic Fibroblasts | mbnl1 |
| sipa1l2 | 7.7E-34 | 1.084873 | 0.135 | 0.052 | 2.51E-29 | Chondrogenic Fibroblasts | sipa1l2 |
| lrrfip1b | 7.8E-34 | 1.3777117 | 0.107 | 0.037 | 2.52E-29 | Chondrogenic Fibroblasts | lrrfip1b |
| uqcr101 | 8.3E-34 | 0.5215553 | 0.403 | 0.235 | 2.69E-29 | Chondrogenic Fibroblasts | uqcr10 |
| pkd21 | 8.6E-34 | 0.7834201 | 0.223 | 0.105 | 2.8E-29 | Chondrogenic Fibroblasts | pkd2 |
| b3gnt2b1 | 1.5E-33 | 1.0727028 | 0.164 | 0.07 | 4.82E-29 | Chondrogenic Fibroblasts | b3gnt2b |
| jam3b3 | 1.6E-33 | 0.6637524 | 0.377 | 0.218 | 5.24E-29 | Chondrogenic Fibroblasts | jam3b |
| psmb11 | 1.8E-33 | 0.480916 | 0.438 | 0.259 | 5.99E-29 | Chondrogenic Fibroblasts | psmb1 |
| si:ch1073-184j22.14 | 2E-33 | 1.0645876 | 0.178 | 0.078 | 6.59E-29 | Chondrogenic Fibroblasts | si:ch1073-184j22.1 |
| ccdc1872 | 3.4E-33 | 0.5361357 | 0.283 | 0.146 | 1.12E-28 | Chondrogenic Fibroblasts | ccdc187 |
| guk1a2 | 3.6E-33 | 0.7018408 | 0.262 | 0.134 | 1.15E-28 | Chondrogenic Fibroblasts | guk1a |
| dlg1l1 | 4.1E-33 | 0.8400403 | 0.176 | 0.076 | 1.32E-28 | Chondrogenic Fibroblasts | dlg1l |
| psmb41 | 4.4E-33 | 0.5486136 | 0.402 | 0.234 | 1.43E-28 | Chondrogenic Fibroblasts | psmb4 |
| ppm1la1 | 4.6E-33 | 1.0281302 | 0.165 | 0.071 | 1.49E-28 | Chondrogenic Fibroblasts | ppm1la |
| gnai2a1 | 4.9E-33 | 0.5560321 | 0.455 | 0.279 | 1.6E-28 | Chondrogenic Fibroblasts | gnai2a |
| rab102 | 5.4E-33 | 0.5145577 | 0.641 | 0.453 | 1.75E-28 | Chondrogenic Fibroblasts | rab10 |
| cux1a4 | 5.5E-33 | 1.2543235 | 0.157 | 0.067 | 1.79E-28 | Chondrogenic Fibroblasts | cux1a |
| calr2 | 7.2E-33 | 0.6924159 | 0.344 | 0.195 | 2.36E-28 | Chondrogenic Fibroblasts | calr |
| myl61 | 8.4E-33 | 0.5009295 | 0.373 | 0.212 | 2.73E-28 | Chondrogenic Fibroblasts | myl6 |
| fam214a2 | 9.6E-33 | 0.9928531 | 0.198 | 0.092 | 3.11E-28 | Chondrogenic Fibroblasts | fam214a |
| tuft1a3 | 1E-32 | 0.4859323 | 0.777 | 0.587 | 3.35E-28 | Chondrogenic Fibroblasts | tuft1a |
| pear13 | 1.4E-32 | 0.6258249 | 0.255 | 0.128 | 4.41E-28 | Chondrogenic Fibroblasts | pear1 |
| tmem183a1 | 1.4E-32 | 1.1604586 | 0.115 | 0.042 | 4.43E-28 | Chondrogenic Fibroblasts | tmem183a |
| ahi12 | 1.4E-32 | 0.8028978 | 0.228 | 0.11 | 4.6E-28 | Chondrogenic Fibroblasts | ahi1 |
| hsp90b11 | 1.8E-32 | 0.5605654 | 0.526 | 0.341 | 5.77E-28 | Chondrogenic Fibroblasts | hsp90b1 |
| dlg3 | 1.8E-32 | 1.0158246 | 0.131 | 0.051 | 5.96E-28 | Chondrogenic Fibroblasts | dlg3 |
| selenot22 | 2E-32 | 0.7562491 | 0.213 | 0.101 | 6.42E-28 | Chondrogenic Fibroblasts | selenot2 |
| slc38a23 | 2E-32 | 0.8218803 | 0.175 | 0.077 | 6.6E-28 | Chondrogenic Fibroblasts | slc38a2 |
| ufc12 | 2.6E-32 | 0.7987187 | 0.253 | 0.13 | 8.44E-28 | Chondrogenic Fibroblasts | ufc1 |
| cyc12 | 4E-32 | 0.5990631 | 0.308 | 0.167 | 1.3E-27 | Chondrogenic Fibroblasts | cyc1 |
| BX936337.12 | 5.2E-32 | 0.5704171 | 0.575 | 0.384 | 1.7E-27 | Chondrogenic Fibroblasts | BX936337.1 |
| atp5mf2 | 5.3E-32 | 0.4418402 | 0.558 | 0.368 | 1.73E-27 | Chondrogenic Fibroblasts | atp5mf |
| tes1 | 5.4E-32 | 0.7834588 | 0.168 | 0.073 | 1.76E-27 | Chondrogenic Fibroblasts | tes |
| hivep2a1 | 5.5E-32 | 0.6033668 | 0.367 | 0.212 | 1.78E-27 | Chondrogenic Fibroblasts | hivep2a |
| cops7a2 | 5.5E-32 | 0.8373919 | 0.223 | 0.11 | 1.79E-27 | Chondrogenic Fibroblasts | cops7a |
| atp6v1aa | 5.5E-32 | 1.0725364 | 0.118 | 0.044 | 1.8E-27 | Chondrogenic Fibroblasts | atp6v1aa |
| lrrc8aa2 | 8.4E-32 | 0.7718872 | 0.205 | 0.097 | 2.72E-27 | Chondrogenic Fibroblasts | lrrc8aa |
| srpk1a | 9.2E-32 | 0.7940612 | 0.24 | 0.12 | 3E-27 | Chondrogenic Fibroblasts | srpk1a |

| | | | | | | | |
|---|---|---|---|---|---|---|---|
| fkbp22 | 9.8E-32 | 0.57814 | 0.351 | 0.2 | 3.19E-27 | Chondrogenic Fibroblasts | fkbp2 |
| letm11 | 1E-31 | 1.0733236 | 0.137 | 0.055 | 3.37E-27 | Chondrogenic Fibroblasts | letm1 |
| creb3l3l1 | 1.2E-31 | 0.47509 | 0.417 | 0.245 | 4.01E-27 | Chondrogenic Fibroblasts | creb3l3l |
| kif5ba1 | 1.5E-31 | 0.7342183 | 0.201 | 0.095 | 4.99E-27 | Chondrogenic Fibroblasts | kif5ba |
| ube2l3a1 | 1.7E-31 | 0.6554489 | 0.281 | 0.149 | 5.53E-27 | Chondrogenic Fibroblasts | ube2l3a |
| stx12 | 1.8E-31 | 0.6451109 | 0.29 | 0.157 | 5.95E-27 | Chondrogenic Fibroblasts | stx12 |
| atp5mc3a2 | 2E-31 | 0.4877552 | 0.44 | 0.267 | 6.48E-27 | Chondrogenic Fibroblasts | atp5mc3a |
| ctsa2 | 2.3E-31 | 0.6984688 | 0.233 | 0.115 | 7.54E-27 | Chondrogenic Fibroblasts | ctsa |
| u2af2b1 | 2.9E-31 | 0.4794685 | 0.483 | 0.302 | 9.31E-27 | Chondrogenic Fibroblasts | u2af2b |
| pdap1b3 | 3.1E-31 | 0.7935822 | 0.198 | 0.093 | 9.95E-27 | Chondrogenic Fibroblasts | pdap1b |
| prps1a1 | 3.6E-31 | 0.699911 | 0.23 | 0.114 | 1.18E-26 | Chondrogenic Fibroblasts | prps1a |
| atp2b41 | 3.9E-31 | 0.5089954 | 0.535 | 0.352 | 1.27E-26 | Chondrogenic Fibroblasts | atp2b4 |
| dnajc42 | 4.2E-31 | 0.7528916 | 0.198 | 0.093 | 1.36E-26 | Chondrogenic Fibroblasts | dnajc4 |
| ppp2r1bb1 | 4.3E-31 | 0.7671372 | 0.197 | 0.092 | 1.39E-26 | Chondrogenic Fibroblasts | ppp2r1bb |
| clic12 | 5.2E-31 | 0.7563676 | 0.212 | 0.102 | 1.69E-26 | Chondrogenic Fibroblasts | clic1 |
| tmem1893 | 6.4E-31 | 0.7051227 | 0.25 | 0.129 | 2.08E-26 | Chondrogenic Fibroblasts | tmem189 |
| lzts2a1 | 6.4E-31 | 0.9431973 | 0.152 | 0.064 | 2.09E-26 | Chondrogenic Fibroblasts | lzts2a |
| uap13 | 1.1E-30 | 0.6572452 | 0.253 | 0.129 | 3.55E-26 | Chondrogenic Fibroblasts | uap1 |
| slc2a101 | 1.2E-30 | 1.3701315 | 0.106 | 0.038 | 3.84E-26 | Chondrogenic Fibroblasts | slc2a10 |
| lamtor31 | 1.2E-30 | 0.839638 | 0.175 | 0.079 | 3.92E-26 | Chondrogenic Fibroblasts | lamtor3 |
| ndufa121 | 1.3E-30 | 0.5179593 | 0.372 | 0.216 | 4.2E-26 | Chondrogenic Fibroblasts | ndufa12 |
| atp6ap22 | 1.5E-30 | 0.7229224 | 0.205 | 0.098 | 4.86E-26 | Chondrogenic Fibroblasts | atp6ap2 |
| upf3b1 | 1.5E-30 | 0.6347606 | 0.372 | 0.221 | 4.95E-26 | Chondrogenic Fibroblasts | upf3b |
| ist12 | 1.7E-30 | 0.8799255 | 0.168 | 0.074 | 5.47E-26 | Chondrogenic Fibroblasts | ist1 |
| magt13 | 1.8E-30 | 0.5169232 | 0.423 | 0.258 | 5.7E-26 | Chondrogenic Fibroblasts | magt1 |
| ift202 | 2.4E-30 | 0.9214113 | 0.141 | 0.058 | 7.74E-26 | Chondrogenic Fibroblasts | ift20 |
| tmem119b4 | 2.6E-30 | 0.5745044 | 0.354 | 0.203 | 8.37E-26 | Chondrogenic Fibroblasts | tmem119b |
| rab14 | 2.6E-30 | 0.5129544 | 0.381 | 0.223 | 8.5E-26 | Chondrogenic Fibroblasts | rab14 |
| emc103 | 3.4E-30 | 0.6720038 | 0.268 | 0.142 | 1.11E-25 | Chondrogenic Fibroblasts | emc10 |
| pdap1a3 | 3.6E-30 | 0.4968244 | 0.368 | 0.216 | 1.18E-25 | Chondrogenic Fibroblasts | pdap1a |
| rab22a2 | 4E-30 | 0.7252398 | 0.207 | 0.1 | 1.31E-25 | Chondrogenic Fibroblasts | rab22a |
| pdcd10a2 | 4.1E-30 | 0.838853 | 0.196 | 0.093 | 1.34E-25 | Chondrogenic Fibroblasts | pdcd10a |
| mesd2 | 4.6E-30 | 0.7050182 | 0.274 | 0.148 | 1.48E-25 | Chondrogenic Fibroblasts | mesd |
| mt-nd21 | 4.6E-30 | 0.3885368 | 0.805 | 0.627 | 1.51E-25 | Chondrogenic Fibroblasts | mt-nd2 |
| atp6v0d11 | 5.2E-30 | 0.9026191 | 0.166 | 0.074 | 1.69E-25 | Chondrogenic Fibroblasts | atp6v0d1 |
| mthfd21 | 5.3E-30 | 0.8326885 | 0.145 | 0.061 | 1.72E-25 | Chondrogenic Fibroblasts | mthfd2 |
| tram13 | 6.5E-30 | 0.5182991 | 0.451 | 0.284 | 2.13E-25 | Chondrogenic Fibroblasts | tram1 |
| ndufab1a1 | 6.6E-30 | 0.6095362 | 0.245 | 0.126 | 2.15E-25 | Chondrogenic Fibroblasts | ndufab1a |
| pitpnb2 | 7.3E-30 | 1.2097251 | 0.12 | 0.047 | 2.36E-25 | Chondrogenic Fibroblasts | pitpnb |
| fam114a23 | 7.6E-30 | 0.807409 | 0.223 | 0.112 | 2.47E-25 | Chondrogenic Fibroblasts | fam114a2 |
| selenok2 | 8.1E-30 | 0.6620668 | 0.314 | 0.178 | 2.64E-25 | Chondrogenic Fibroblasts | selenok |
| s100v22 | 9.3E-30 | 0.4337998 | 0.82 | 0.676 | 3.04E-25 | Chondrogenic Fibroblasts | s100v2 |
| eif3jb2 | 9.6E-30 | 0.4912664 | 0.377 | 0.222 | 3.14E-25 | Chondrogenic Fibroblasts | eif3jb |
| ifrd13 | 1E-29 | 0.5134797 | 0.448 | 0.279 | 3.34E-25 | Chondrogenic Fibroblasts | ifrd1 |
| tbc1d8b | 1.6E-29 | 1.1654585 | 0.112 | 0.042 | 5.1E-25 | Chondrogenic Fibroblasts | tbc1d8b |
| pim33 | 1.9E-29 | 0.8872703 | 0.211 | 0.105 | 6.1E-25 | Chondrogenic Fibroblasts | pim3 |
| psmd72 | 1.9E-29 | 0.4771915 | 0.405 | 0.243 | 6.19E-25 | Chondrogenic Fibroblasts | psmd7 |
| bmp2k | 2E-29 | 1.0551665 | 0.106 | 0.039 | 6.54E-25 | Chondrogenic Fibroblasts | bmp2k |

| | | | | | | | |
|---|---|---|---|---|---|---|---|
| lrp12 | 2.4E-29 | 1.0739765 | 0.117 | 0.045 | 7.97E-25 | Chondrogenic Fibroblasts | lrp12 |
| ammecr12 | 2.9E-29 | 0.6761908 | 0.267 | 0.143 | 9.5E-25 | Chondrogenic Fibroblasts | ammecr1 |
| tubb51 | 3E-29 | 0.671574 | 0.272 | 0.147 | 9.88E-25 | Chondrogenic Fibroblasts | tubb5 |
| si:ch211-242b18.11 | 3.7E-29 | 0.7888977 | 0.183 | 0.085 | 1.2E-24 | Chondrogenic Fibroblasts | si:ch211-242b18.1 |
| atp5pb2 | 3.8E-29 | 0.4117358 | 0.649 | 0.46 | 1.25E-24 | Chondrogenic Fibroblasts | atp5pb |
| mapkapk2a2 | 4.7E-29 | 0.5455586 | 0.369 | 0.219 | 1.52E-24 | Chondrogenic Fibroblasts | mapkapk2a |
| ccng13 | 5.1E-29 | 0.6331855 | 0.672 | 0.501 | 1.64E-24 | Chondrogenic Fibroblasts | ccng1 |
| calm3b1 | 5.1E-29 | 0.4360353 | 0.601 | 0.412 | 1.66E-24 | Chondrogenic Fibroblasts | calm3b |
| golga31 | 5.2E-29 | 0.7731301 | 0.154 | 0.067 | 1.68E-24 | Chondrogenic Fibroblasts | golga3 |
| si:dkey-25e12.31 | 5.4E-29 | 0.8531377 | 0.16 | 0.071 | 1.76E-24 | Chondrogenic Fibroblasts | si:dkey-25e12.3 |
| ndufa71 | 5.6E-29 | 1.0221781 | 0.122 | 0.048 | 1.83E-24 | Chondrogenic Fibroblasts | ndufa7 |
| ywhaba1 | 6.8E-29 | 0.4223268 | 0.697 | 0.502 | 2.22E-24 | Chondrogenic Fibroblasts | ywhaba |
| npc22 | 7.3E-29 | 0.6011083 | 0.499 | 0.343 | 2.38E-24 | Chondrogenic Fibroblasts | npc2 |
| atp5if1b3 | 7.6E-29 | 0.4312373 | 0.509 | 0.328 | 2.48E-24 | Chondrogenic Fibroblasts | atp5if1b |
| atp6v1h1 | 8E-29 | 1.0896098 | 0.133 | 0.055 | 2.59E-24 | Chondrogenic Fibroblasts | atp6v1h |
| cox7b3 | 8.9E-29 | 0.3849342 | 0.583 | 0.392 | 2.91E-24 | Chondrogenic Fibroblasts | cox7b |
| fam133b1 | 9E-29 | 0.5085012 | 0.408 | 0.248 | 2.94E-24 | Chondrogenic Fibroblasts | fam133b |
| stx5a1 | 9.4E-29 | 0.653057 | 0.2 | 0.097 | 3.05E-24 | Chondrogenic Fibroblasts | stx5a |
| ncam1a2 | 1.1E-28 | 0.8183802 | 0.157 | 0.07 | 3.7E-24 | Chondrogenic Fibroblasts | ncam1a |
| zgc:1132082 | 1.3E-28 | 0.6022649 | 0.327 | 0.188 | 4.12E-24 | Chondrogenic Fibroblasts | zgc:113208 |
| TPGS11 | 1.3E-28 | 1.0178648 | 0.12 | 0.048 | 4.31E-24 | Chondrogenic Fibroblasts | TPGS1 |
| cfl22 | 1.4E-28 | 0.9701253 | 0.133 | 0.055 | 4.64E-24 | Chondrogenic Fibroblasts | cfl2 |
| tenm34 | 1.5E-28 | 0.5372029 | 0.348 | 0.204 | 5.04E-24 | Chondrogenic Fibroblasts | tenm3 |
| dnajc3b2 | 1.6E-28 | 0.9345233 | 0.141 | 0.06 | 5.07E-24 | Chondrogenic Fibroblasts | dnajc3b |
| os92 | 1.7E-28 | 0.4952448 | 0.359 | 0.212 | 5.59E-24 | Chondrogenic Fibroblasts | os9 |
| dbi2 | 1.7E-28 | 0.4322633 | 0.624 | 0.438 | 5.63E-24 | Chondrogenic Fibroblasts | dbi |
| pgk12 | 1.9E-28 | 0.6634604 | 0.343 | 0.205 | 6.27E-24 | Chondrogenic Fibroblasts | pgk1 |
| selenow13 | 2.2E-28 | 0.2675297 | 0.975 | 0.895 | 7.02E-24 | Chondrogenic Fibroblasts | selenow1 |
| fkbp113 | 2.4E-28 | 0.7046804 | 0.281 | 0.157 | 7.91E-24 | Chondrogenic Fibroblasts | fkbp11 |
| mtmr111 | 2.8E-28 | 1.0453507 | 0.111 | 0.042 | 9.16E-24 | Chondrogenic Fibroblasts | mtmr11 |
| slc66a33 | 3.5E-28 | 0.6085263 | 0.229 | 0.117 | 1.13E-23 | Chondrogenic Fibroblasts | slc66a3 |
| ATP11B2 | 3.5E-28 | 0.7007575 | 0.213 | 0.106 | 1.14E-23 | Chondrogenic Fibroblasts | ATP11B |
| sec61a14 | 3.5E-28 | 0.4195316 | 0.609 | 0.416 | 1.15E-23 | Chondrogenic Fibroblasts | sec61a1 |
| cst31 | 3.6E-28 | 0.3468562 | 0.856 | 0.721 | 1.16E-23 | Chondrogenic Fibroblasts | cst3 |
| rcn21 | 3.7E-28 | 0.825398 | 0.17 | 0.079 | 1.22E-23 | Chondrogenic Fibroblasts | rcn2 |
| ube2a1 | 4.2E-28 | 0.7736647 | 0.175 | 0.082 | 1.36E-23 | Chondrogenic Fibroblasts | ube2a |
| rcn13 | 5.8E-28 | 0.7307947 | 0.241 | 0.127 | 1.87E-23 | Chondrogenic Fibroblasts | rcn1 |
| psmb61 | 6.8E-28 | 0.4423666 | 0.486 | 0.311 | 2.21E-23 | Chondrogenic Fibroblasts | psmb6 |
| ptbp1a3 | 7.1E-28 | 0.559224 | 0.372 | 0.226 | 2.3E-23 | Chondrogenic Fibroblasts | ptbp1a |
| surf4l2 | 7.2E-28 | 0.594989 | 0.238 | 0.124 | 2.35E-23 | Chondrogenic Fibroblasts | surf4l |
| nrp2b3 | 7.7E-28 | 0.7008133 | 0.236 | 0.123 | 2.52E-23 | Chondrogenic Fibroblasts | nrp2b |
| pdcd10b2 | 8.1E-28 | 0.5955371 | 0.201 | 0.098 | 2.62E-23 | Chondrogenic Fibroblasts | pdcd10b |
| eif4ebp22 | 8.2E-28 | 0.5650613 | 0.356 | 0.213 | 2.65E-23 | Chondrogenic Fibroblasts | eif4ebp2 |
| zbtb8os2 | 9.4E-28 | 0.9361975 | 0.16 | 0.073 | 3.05E-23 | Chondrogenic Fibroblasts | zbtb8os |
| BX284638.21 | 9.8E-28 | 0.7643254 | 0.103 | 0.038 | 3.17E-23 | Chondrogenic Fibroblasts | BX284638.2 |
| eif4e1c1 | 1.2E-27 | 0.5111135 | 0.265 | 0.142 | 3.77E-23 | Chondrogenic Fibroblasts | eif4e1c |
| syap12 | 1.2E-27 | 0.614831 | 0.278 | 0.154 | 3.8E-23 | Chondrogenic Fibroblasts | syap1 |
| dapk3 | 1.3E-27 | 1.0989151 | 0.101 | 0.037 | 4.17E-23 | Chondrogenic Fibroblasts | dapk3 |

| | | | | | | | |
|---|---|---|---|---|---|---|---|
| lurap11 | 1.3E-27 | 1.0482045 | 0.122 | 0.049 | 4.2E-23 | Chondrogenic Fibroblasts | lurap1 |
| si:ch1073-429i10.32 | 1.4E-27 | 0.4242838 | 0.587 | 0.399 | 4.59E-23 | Chondrogenic Fibroblasts | si:ch1073-429i10.3 |
| prrc12 | 1.5E-27 | 0.8916951 | 0.166 | 0.077 | 4.83E-23 | Chondrogenic Fibroblasts | prrc1 |
| suclg12 | 1.5E-27 | 0.5795884 | 0.303 | 0.172 | 4.93E-23 | Chondrogenic Fibroblasts | suclg1 |
| nudcd32 | 1.6E-27 | 0.8486786 | 0.174 | 0.082 | 5.1E-23 | Chondrogenic Fibroblasts | nudcd3 |
| MDFIC2 | 1.8E-27 | 0.5879809 | 0.32 | 0.183 | 5.74E-23 | Chondrogenic Fibroblasts | MDFIC |
| tram23 | 1.9E-27 | 0.7135789 | 0.213 | 0.108 | 6.02E-23 | Chondrogenic Fibroblasts | tram2 |
| vcp1 | 2E-27 | 0.4441091 | 0.484 | 0.313 | 6.44E-23 | Chondrogenic Fibroblasts | vcp |
| edf12 | 2E-27 | 0.4236441 | 0.696 | 0.509 | 6.46E-23 | Chondrogenic Fibroblasts | edf1 |
| gnb1a | 2.1E-27 | 0.4153229 | 0.662 | 0.485 | 6.92E-23 | Chondrogenic Fibroblasts | gnb1a |
| ugt2a73 | 2.4E-27 | 0.6687177 | 0.191 | 0.092 | 7.71E-23 | Chondrogenic Fibroblasts | ugt2a7 |
| slc25a242 | 2.4E-27 | 0.5079778 | 0.363 | 0.216 | 7.73E-23 | Chondrogenic Fibroblasts | slc25a24 |
| mpc22 | 2.6E-27 | 0.5963406 | 0.309 | 0.179 | 8.35E-23 | Chondrogenic Fibroblasts | mpc2 |
| bri3 | 2.6E-27 | 0.8512501 | 0.137 | 0.058 | 8.39E-23 | Chondrogenic Fibroblasts | bri3 |
| serinc21 | 2.7E-27 | 0.4665947 | 0.478 | 0.314 | 8.82E-23 | Chondrogenic Fibroblasts | serinc2 |
| tmbim42 | 2.8E-27 | 0.6457943 | 0.223 | 0.114 | 9.21E-23 | Chondrogenic Fibroblasts | tmbim4 |
| arcn1b3 | 2.8E-27 | 0.6657399 | 0.231 | 0.121 | 9.23E-23 | Chondrogenic Fibroblasts | arcn1b |
| appb2 | 3E-27 | 0.400797 | 0.651 | 0.464 | 9.76E-23 | Chondrogenic Fibroblasts | appb |
| sdhb2 | 3.5E-27 | 0.6077687 | 0.302 | 0.173 | 1.14E-22 | Chondrogenic Fibroblasts | sdhb |
| capn2a2 | 3.6E-27 | 0.4209055 | 0.533 | 0.358 | 1.17E-22 | Chondrogenic Fibroblasts | capn2a |
| rgs12b4 | 4.1E-27 | 0.8196315 | 0.18 | 0.087 | 1.35E-22 | Chondrogenic Fibroblasts | rgs12b |
| cpne11 | 4.3E-27 | 0.3675793 | 0.657 | 0.466 | 1.39E-22 | Chondrogenic Fibroblasts | cpne1 |
| exoc1 | 4.3E-27 | 0.7672956 | 0.184 | 0.089 | 1.39E-22 | Chondrogenic Fibroblasts | exoc1 |
| ostm1 | 4.7E-27 | 0.8361276 | 0.152 | 0.068 | 1.54E-22 | Chondrogenic Fibroblasts | ostm1 |
| nedd81 | 6.7E-27 | 0.7059872 | 0.209 | 0.106 | 2.19E-22 | Chondrogenic Fibroblasts | nedd8 |
| zc3h151 | 8.9E-27 | 0.5875397 | 0.256 | 0.138 | 2.9E-22 | Chondrogenic Fibroblasts | zc3h15 |
| tmem39a3 | 9.1E-27 | 0.6553793 | 0.266 | 0.148 | 2.96E-22 | Chondrogenic Fibroblasts | tmem39a |
| h3f3b.1.11 | 9.7E-27 | 0.6700651 | 0.295 | 0.168 | 3.16E-22 | Chondrogenic Fibroblasts | h3f3b.1.1 |
| tpd52l2b2 | 1E-26 | 0.5869637 | 0.357 | 0.217 | 3.34E-22 | Chondrogenic Fibroblasts | tpd52l2b |
| si:ch211-235e9.62 | 1.1E-26 | 0.7142652 | 0.187 | 0.091 | 3.57E-22 | Chondrogenic Fibroblasts | si:ch211-235e9.6 |
| sumo2b2 | 1.1E-26 | 0.4883055 | 0.563 | 0.389 | 3.63E-22 | Chondrogenic Fibroblasts | sumo2b |
| si:dkey-33c12.42 | 1.1E-26 | 0.4454167 | 0.513 | 0.338 | 3.73E-22 | Chondrogenic Fibroblasts | si:dkey-33c12.4 |
| psmc51 | 1.4E-26 | 0.5550097 | 0.298 | 0.171 | 4.48E-22 | Chondrogenic Fibroblasts | psmc5 |
| psmd141 | 1.5E-26 | 0.6630332 | 0.246 | 0.132 | 4.75E-22 | Chondrogenic Fibroblasts | psmd14 |
| med19a1 | 1.6E-26 | 0.577955 | 0.243 | 0.13 | 5.12E-22 | Chondrogenic Fibroblasts | med19a |
| timmdc12 | 1.8E-26 | 0.7663749 | 0.18 | 0.087 | 5.81E-22 | Chondrogenic Fibroblasts | timmdc1 |
| pin11 | 1.9E-26 | 0.5053323 | 0.414 | 0.259 | 6.11E-22 | Chondrogenic Fibroblasts | pin1 |
| anxa131 | 2E-26 | 0.4330213 | 0.515 | 0.341 | 6.4E-22 | Chondrogenic Fibroblasts | anxa13 |
| yipf31 | 2E-26 | 0.7980833 | 0.155 | 0.07 | 6.59E-22 | Chondrogenic Fibroblasts | yipf3 |
| ddah2 | 2.3E-26 | 0.6642996 | 0.236 | 0.126 | 7.6E-22 | Chondrogenic Fibroblasts | ddah2 |
| tcea11 | 2.4E-26 | 0.6415461 | 0.206 | 0.104 | 7.71E-22 | Chondrogenic Fibroblasts | tcea1 |
| psma41 | 2.8E-26 | 0.4785696 | 0.446 | 0.286 | 8.97E-22 | Chondrogenic Fibroblasts | psma4 |
| josd1 | 2.8E-26 | 1.0699508 | 0.101 | 0.038 | 9.1E-22 | Chondrogenic Fibroblasts | josd1 |
| yars11 | 2.9E-26 | 1.0027951 | 0.142 | 0.063 | 9.53E-22 | Chondrogenic Fibroblasts | yars1 |
| uck2b3 | 3.2E-26 | 1.0082743 | 0.125 | 0.052 | 1.03E-21 | Chondrogenic Fibroblasts | uck2b |
| lamtor23 | 3.2E-26 | 0.7610278 | 0.224 | 0.118 | 1.04E-21 | Chondrogenic Fibroblasts | lamtor2 |
| glsb1 | 3.5E-26 | 0.6169782 | 0.221 | 0.114 | 1.15E-21 | Chondrogenic Fibroblasts | glsb |
| lasp11 | 3.6E-26 | 0.4109916 | 0.644 | 0.468 | 1.18E-21 | Chondrogenic Fibroblasts | lasp1 |

| | | | | | | | |
|---|---|---|---|---|---|---|---|
| chordc1b1 | 3.9E-26 | 0.6271965 | 0.199 | 0.1 | 1.26E-21 | Chondrogenic Fibroblasts | chordc1b |
| ubl7b2 | 3.9E-26 | 0.7398741 | 0.16 | 0.074 | 1.28E-21 | Chondrogenic Fibroblasts | ubl7b |
| papss2b2 | 4.1E-26 | 0.8323104 | 0.192 | 0.096 | 1.32E-21 | Chondrogenic Fibroblasts | papss2b |
| ppdpfb3 | 4.1E-26 | 0.351939 | 0.899 | 0.775 | 1.33E-21 | Chondrogenic Fibroblasts | ppdpfb |
| frzb6 | 5.8E-26 | 0.910863 | 0.233 | 0.127 | 1.88E-21 | Chondrogenic Fibroblasts | frzb |
| parvaa | 5.8E-26 | 0.5259361 | 0.341 | 0.203 | 1.89E-21 | Chondrogenic Fibroblasts | parvaa |
| slc20a21 | 7.1E-26 | 0.5206991 | 0.421 | 0.269 | 2.31E-21 | Chondrogenic Fibroblasts | slc20a2 |
| slc12a42 | 7.9E-26 | 0.5036742 | 0.234 | 0.123 | 2.57E-21 | Chondrogenic Fibroblasts | slc12a4 |
| niban2a3 | 8.3E-26 | 0.9339473 | 0.12 | 0.05 | 2.7E-21 | Chondrogenic Fibroblasts | niban2a |
| atp6v1g13 | 8.6E-26 | 0.7189471 | 0.24 | 0.13 | 2.81E-21 | Chondrogenic Fibroblasts | atp6v1g1 |
| atp5l1 | 1.2E-25 | 0.3460818 | 0.771 | 0.59 | 4.04E-21 | Chondrogenic Fibroblasts | atp5l |
| nucb12 | 1.3E-25 | 0.4486663 | 0.346 | 0.204 | 4.21E-21 | Chondrogenic Fibroblasts | nucb1 |
| unc502 | 1.3E-25 | 0.9187956 | 0.166 | 0.079 | 4.25E-21 | Chondrogenic Fibroblasts | unc50 |
| cpvl3 | 1.5E-25 | 0.787638 | 0.185 | 0.091 | 4.8E-21 | Chondrogenic Fibroblasts | cpvl |
| hm131 | 1.6E-25 | 0.573786 | 0.253 | 0.139 | 5.2E-21 | Chondrogenic Fibroblasts | hm13 |
| si:ch1073-325m22.23 | 1.9E-25 | 0.4464417 | 0.459 | 0.299 | 6.21E-21 | Chondrogenic Fibroblasts | si:ch1073-325m22. |
| ifi302 | 1.9E-25 | 0.5697207 | 0.259 | 0.144 | 6.34E-21 | Chondrogenic Fibroblasts | ifi30 |
| CABZ01080375.12 | 2E-25 | 0.9178078 | 0.144 | 0.065 | 6.38E-21 | Chondrogenic Fibroblasts | CABZ01080375.1 |
| kifc32 | 2.1E-25 | 0.9238381 | 0.144 | 0.065 | 6.74E-21 | Chondrogenic Fibroblasts | kifc3 |
| dixdc1a2 | 2.2E-25 | 0.9860129 | 0.125 | 0.053 | 7.22E-21 | Chondrogenic Fibroblasts | dixdc1a |
| znhit11 | 2.4E-25 | 0.8969915 | 0.132 | 0.058 | 7.65E-21 | Chondrogenic Fibroblasts | znhit1 |
| tegt1 | 2.4E-25 | 0.536495 | 0.274 | 0.154 | 7.84E-21 | Chondrogenic Fibroblasts | tegt |
| si:dkey-172h23.23 | 2.4E-25 | 0.4795137 | 0.262 | 0.144 | 7.85E-21 | Chondrogenic Fibroblasts | si:dkey-172h23.2 |
| lrrc8da | 2.4E-25 | 0.8851682 | 0.13 | 0.056 | 7.95E-21 | Chondrogenic Fibroblasts | lrrc8da |
| map7d1a | 3E-25 | 0.7060366 | 0.228 | 0.122 | 9.82E-21 | Chondrogenic Fibroblasts | map7d1a |
| cmtm71 | 3E-25 | 0.6865939 | 0.224 | 0.119 | 9.85E-21 | Chondrogenic Fibroblasts | cmtm7 |
| srprb3 | 3.1E-25 | 0.5337206 | 0.306 | 0.179 | 9.94E-21 | Chondrogenic Fibroblasts | srprb |
| mbnl21 | 3.5E-25 | 0.4334564 | 0.364 | 0.22 | 1.12E-20 | Chondrogenic Fibroblasts | mbnl2 |
| calhm2.11 | 3.5E-25 | 0.7425938 | 0.15 | 0.069 | 1.14E-20 | Chondrogenic Fibroblasts | calhm2.1 |
| abcf11 | 3.5E-25 | 0.3442447 | 0.474 | 0.305 | 1.15E-20 | Chondrogenic Fibroblasts | abcf1 |
| lamp21 | 4.4E-25 | 0.4313659 | 0.513 | 0.346 | 1.44E-20 | Chondrogenic Fibroblasts | lamp2 |
| ppp1r37 | 4.5E-25 | 0.8529073 | 0.134 | 0.059 | 1.46E-20 | Chondrogenic Fibroblasts | ppp1r37 |
| si:dkey-21a6.5 | 4.9E-25 | 0.9706277 | 0.11 | 0.044 | 1.6E-20 | Chondrogenic Fibroblasts | si:dkey-21a6.5 |
| cdc42ep4b5 | 5E-25 | 0.7181654 | 0.173 | 0.084 | 1.63E-20 | Chondrogenic Fibroblasts | cdc42ep4b |
| sars13 | 5.1E-25 | 0.5444446 | 0.259 | 0.144 | 1.67E-20 | Chondrogenic Fibroblasts | sars1 |
| rab121 | 5.6E-25 | 0.8664864 | 0.146 | 0.067 | 1.82E-20 | Chondrogenic Fibroblasts | rab12 |
| mt-nd32 | 6.8E-25 | 0.3580211 | 0.534 | 0.357 | 2.21E-20 | Chondrogenic Fibroblasts | mt-nd3 |
| rab412 | 6.8E-25 | 0.6537964 | 0.168 | 0.08 | 2.21E-20 | Chondrogenic Fibroblasts | rab41 |
| txndc172 | 6.9E-25 | 0.5811053 | 0.221 | 0.116 | 2.23E-20 | Chondrogenic Fibroblasts | txndc17 |
| gcsha1 | 7.3E-25 | 0.6749534 | 0.196 | 0.1 | 2.38E-20 | Chondrogenic Fibroblasts | gcsha |
| p3h34 | 7.6E-25 | 0.6433276 | 0.242 | 0.133 | 2.46E-20 | Chondrogenic Fibroblasts | p3h3 |
| ano10a1 | 7.6E-25 | 0.881195 | 0.136 | 0.06 | 2.47E-20 | Chondrogenic Fibroblasts | ano10a |
| CABZ01079281.1 | 7.9E-25 | 1.0304774 | 0.101 | 0.04 | 2.55E-20 | Chondrogenic Fibroblasts | CABZ01079281.1 |
| csnk2b2 | 8.5E-25 | 0.4483565 | 0.399 | 0.25 | 2.75E-20 | Chondrogenic Fibroblasts | csnk2b |
| tnc6 | 9.1E-25 | 0.4062325 | 0.64 | 0.466 | 2.96E-20 | Chondrogenic Fibroblasts | tnc |
| smox2 | 1.2E-24 | 0.6531008 | 0.166 | 0.079 | 3.87E-20 | Chondrogenic Fibroblasts | smox |
| arf51 | 1.2E-24 | 0.3957904 | 0.47 | 0.306 | 3.89E-20 | Chondrogenic Fibroblasts | arf5 |
| ndufa23 | 1.3E-24 | 0.7356156 | 0.183 | 0.092 | 4.08E-20 | Chondrogenic Fibroblasts | ndufa2 |

| | | | | | | | |
|---|---|---|---|---|---|---|---|
| glrx32 | 1.4E-24 | 0.6064871 | 0.235 | 0.127 | 4.59E-20 | Chondrogenic Fibroblasts | glrx3 |
| PLBD2 | 1.4E-24 | 1.1075319 | 0.1 | 0.039 | 4.65E-20 | Chondrogenic Fibroblasts | PLBD2 |
| eprs13 | 1.4E-24 | 0.7270503 | 0.184 | 0.092 | 4.7E-20 | Chondrogenic Fibroblasts | eprs1 |
| uqcrq1 | 1.7E-24 | 0.4779563 | 0.416 | 0.267 | 5.41E-20 | Chondrogenic Fibroblasts | uqcrq |
| lamtor41 | 1.8E-24 | 0.7455984 | 0.197 | 0.101 | 5.82E-20 | Chondrogenic Fibroblasts | lamtor4 |
| slirp2 | 2E-24 | 0.5495283 | 0.253 | 0.141 | 6.64E-20 | Chondrogenic Fibroblasts | slirp |
| rab5aa2 | 2.1E-24 | 0.498557 | 0.404 | 0.259 | 6.88E-20 | Chondrogenic Fibroblasts | rab5aa |
| ndufa4l3 | 2.1E-24 | 0.3048112 | 0.865 | 0.75 | 6.94E-20 | Chondrogenic Fibroblasts | ndufa4l |
| tmod22 | 2.4E-24 | 0.3581368 | 0.298 | 0.169 | 7.73E-20 | Chondrogenic Fibroblasts | tmod2 |
| hmgn73 | 2.4E-24 | 0.3796626 | 0.709 | 0.519 | 7.79E-20 | Chondrogenic Fibroblasts | hmgn7 |
| yipf52 | 2.7E-24 | 0.6351148 | 0.225 | 0.121 | 8.91E-20 | Chondrogenic Fibroblasts | yipf5 |
| tmco12 | 2.9E-24 | 0.6681511 | 0.166 | 0.08 | 9.4E-20 | Chondrogenic Fibroblasts | tmco1 |
| fmnl2a2 | 3.4E-24 | 0.7091059 | 0.206 | 0.108 | 1.11E-19 | Chondrogenic Fibroblasts | fmnl2a |
| hnrnpa1a1 | 3.7E-24 | 0.5655696 | 0.244 | 0.134 | 1.2E-19 | Chondrogenic Fibroblasts | hnrnpa1a |
| arhgef1b2 | 4.1E-24 | 0.4021803 | 0.406 | 0.257 | 1.33E-19 | Chondrogenic Fibroblasts | arhgef1b |
| tmem1651 | 5.1E-24 | 0.7769513 | 0.145 | 0.067 | 1.67E-19 | Chondrogenic Fibroblasts | tmem165 |
| erh1 | 5.4E-24 | 0.3372112 | 0.584 | 0.405 | 1.74E-19 | Chondrogenic Fibroblasts | erh |
| zgc:1531153 | 5.8E-24 | 0.6607876 | 0.167 | 0.081 | 1.88E-19 | Chondrogenic Fibroblasts | zgc:153115 |
| syncripl2 | 6.1E-24 | 0.6645318 | 0.22 | 0.118 | 1.98E-19 | Chondrogenic Fibroblasts | syncripl |
| ndufv32 | 7E-24 | 0.5503033 | 0.233 | 0.127 | 2.27E-19 | Chondrogenic Fibroblasts | ndufv3 |
| pcmt2 | 7E-24 | 0.6310916 | 0.182 | 0.091 | 2.29E-19 | Chondrogenic Fibroblasts | pcmt |
| glmp | 7.1E-24 | 0.8750112 | 0.117 | 0.05 | 2.32E-19 | Chondrogenic Fibroblasts | glmp |
| sorbs12 | 7.3E-24 | 0.8232835 | 0.239 | 0.132 | 2.37E-19 | Chondrogenic Fibroblasts | sorbs1 |
| psmd4a1 | 7.5E-24 | 0.5384438 | 0.193 | 0.098 | 2.44E-19 | Chondrogenic Fibroblasts | psmd4a |
| phex1 | 7.8E-24 | 1.4079207 | 0.155 | 0.076 | 2.55E-19 | Chondrogenic Fibroblasts | phex |
| smap12 | 8.7E-24 | 0.5245843 | 0.27 | 0.153 | 2.82E-19 | Chondrogenic Fibroblasts | smap1 |
| rap1b | 8.7E-24 | 0.3815143 | 0.527 | 0.362 | 2.83E-19 | Chondrogenic Fibroblasts | rap1b |
| rab11a1 | 9.7E-24 | 0.3986825 | 0.332 | 0.198 | 3.15E-19 | Chondrogenic Fibroblasts | rab11a |
| dazap21 | 9.9E-24 | 0.6059261 | 0.256 | 0.144 | 3.21E-19 | Chondrogenic Fibroblasts | dazap2 |
| gnai2b4 | 1E-23 | 0.9335328 | 0.136 | 0.062 | 3.27E-19 | Chondrogenic Fibroblasts | gnai2b |
| cyp20a11 | 1.2E-23 | 0.6747143 | 0.21 | 0.111 | 3.86E-19 | Chondrogenic Fibroblasts | cyp20a1 |
| commd71 | 1.2E-23 | 0.7451416 | 0.169 | 0.083 | 3.96E-19 | Chondrogenic Fibroblasts | commd7 |
| tia1l2 | 1.2E-23 | 0.6064465 | 0.246 | 0.137 | 4E-19 | Chondrogenic Fibroblasts | tia1l |
| ndufs52 | 1.3E-23 | 0.5051757 | 0.298 | 0.175 | 4.11E-19 | Chondrogenic Fibroblasts | ndufs5 |
| dctn5 | 1.3E-23 | 0.5904585 | 0.217 | 0.116 | 4.3E-19 | Chondrogenic Fibroblasts | dctn5 |
| ddx231 | 1.4E-23 | 0.505121 | 0.278 | 0.16 | 4.39E-19 | Chondrogenic Fibroblasts | ddx23 |
| lrp101 | 1.4E-23 | 0.586488 | 0.185 | 0.093 | 4.58E-19 | Chondrogenic Fibroblasts | lrp10 |
| hdlbpa5 | 1.5E-23 | 0.3357916 | 0.824 | 0.672 | 4.74E-19 | Chondrogenic Fibroblasts | hdlbpa |
| mt-atp61 | 1.5E-23 | 0.2579642 | 0.995 | 0.964 | 4.96E-19 | Chondrogenic Fibroblasts | mt-atp6 |
| gdi11 | 1.8E-23 | 0.568482 | 0.198 | 0.103 | 5.7E-19 | Chondrogenic Fibroblasts | gdi1 |
| nsfl1c2 | 1.9E-23 | 0.5576657 | 0.229 | 0.124 | 6.24E-19 | Chondrogenic Fibroblasts | nsfl1c |
| bmpr2a1 | 1.9E-23 | 0.4875386 | 0.299 | 0.175 | 6.34E-19 | Chondrogenic Fibroblasts | bmpr2a |
| lmo7a3 | 2E-23 | 0.7099854 | 0.261 | 0.151 | 6.37E-19 | Chondrogenic Fibroblasts | lmo7a |
| ap1m11 | 2E-23 | 0.689404 | 0.223 | 0.123 | 6.38E-19 | Chondrogenic Fibroblasts | ap1m1 |
| vdac33 | 2E-23 | 0.384425 | 0.698 | 0.523 | 6.5E-19 | Chondrogenic Fibroblasts | vdac3 |
| ralaa1 | 2.3E-23 | 0.9522775 | 0.104 | 0.042 | 7.41E-19 | Chondrogenic Fibroblasts | ralaa |
| rab5c1 | 2.9E-23 | 0.4204333 | 0.366 | 0.228 | 9.27E-19 | Chondrogenic Fibroblasts | rab5c |
| ppp1r1c3 | 3E-23 | 2.1055868 | 0.223 | 0.13 | 9.68E-19 | Chondrogenic Fibroblasts | ppp1r1c |

| | | | | | | | |
|---|---|---|---|---|---|---|---|
| bsg1 | 3.8E-23 | 0.3659534 | 0.732 | 0.57 | 1.24E-18 | Chondrogenic Fibroblasts | bsg |
| ubtf1 | 4E-23 | 0.4211074 | 0.508 | 0.345 | 1.31E-18 | Chondrogenic Fibroblasts | ubtf |
| dnase1l11 | 4.2E-23 | 0.6462605 | 0.21 | 0.112 | 1.36E-18 | Chondrogenic Fibroblasts | dnase1l1 |
| dtnbp1a2 | 4.3E-23 | 0.7389893 | 0.206 | 0.11 | 1.4E-18 | Chondrogenic Fibroblasts | dtnbp1a |
| p4ha1a4 | 4.6E-23 | 0.5841862 | 0.186 | 0.095 | 1.49E-18 | Chondrogenic Fibroblasts | p4ha1a |
| eif2s21 | 4.7E-23 | 0.3176533 | 0.644 | 0.46 | 1.52E-18 | Chondrogenic Fibroblasts | eif2s2 |
| pros12 | 5.2E-23 | 0.5201566 | 0.215 | 0.114 | 1.68E-18 | Chondrogenic Fibroblasts | pros1 |
| sh3pxd2aa1 | 5.2E-23 | 0.637515 | 0.249 | 0.141 | 1.68E-18 | Chondrogenic Fibroblasts | sh3pxd2aa |
| arpc4l1 | 5.2E-23 | 0.343699 | 0.531 | 0.359 | 1.69E-18 | Chondrogenic Fibroblasts | arpc4l |
| nkap1 | 5.6E-23 | 0.4981198 | 0.308 | 0.185 | 1.82E-18 | Chondrogenic Fibroblasts | nkap |
| rida2 | 5.6E-23 | 0.8633904 | 0.162 | 0.08 | 1.84E-18 | Chondrogenic Fibroblasts | rida |
| ptpn2b | 5.9E-23 | 0.5974494 | 0.21 | 0.112 | 1.92E-18 | Chondrogenic Fibroblasts | ptpn2b |
| psap3 | 6.3E-23 | 0.5112454 | 0.497 | 0.346 | 2.04E-18 | Chondrogenic Fibroblasts | psap |
| slc38a101 | 6.7E-23 | 0.3860609 | 0.366 | 0.226 | 2.17E-18 | Chondrogenic Fibroblasts | slc38a10 |
| elof13 | 6.8E-23 | 0.6256647 | 0.25 | 0.142 | 2.21E-18 | Chondrogenic Fibroblasts | elof1 |
| tmem471 | 6.9E-23 | 0.5900647 | 0.273 | 0.158 | 2.25E-18 | Chondrogenic Fibroblasts | tmem47 |
| hmgb1b2 | 7E-23 | 0.3770186 | 0.396 | 0.249 | 2.27E-18 | Chondrogenic Fibroblasts | hmgb1b |
| gnb1b1 | 7.1E-23 | 0.3286405 | 0.733 | 0.564 | 2.3E-18 | Chondrogenic Fibroblasts | gnb1b |
| psmd82 | 7.1E-23 | 0.5485555 | 0.261 | 0.15 | 2.32E-18 | Chondrogenic Fibroblasts | psmd8 |
| dbpb4 | 7.3E-23 | 0.4935629 | 0.21 | 0.111 | 2.39E-18 | Chondrogenic Fibroblasts | dbpb |
| cnep1r12 | 8.5E-23 | 0.6289471 | 0.192 | 0.099 | 2.77E-18 | Chondrogenic Fibroblasts | cnep1r1 |
| bcl2a2 | 8.5E-23 | 0.4828849 | 0.267 | 0.153 | 2.77E-18 | Chondrogenic Fibroblasts | bcl2a |
| rbbp92 | 8.8E-23 | 0.6745898 | 0.223 | 0.123 | 2.86E-18 | Chondrogenic Fibroblasts | rbbp9 |
| cars11 | 9.8E-23 | 0.7094901 | 0.162 | 0.079 | 3.19E-18 | Chondrogenic Fibroblasts | cars1 |
| ENSDARG00000356 | 1E-22 | 0.3967123 | 0.35 | 0.214 | 3.36E-18 | Chondrogenic Fibroblasts | ENSDARG00000035 |
| atp6v0b | 1E-22 | 0.5435019 | 0.222 | 0.121 | 3.38E-18 | Chondrogenic Fibroblasts | atp6v0b |
| bmp63 | 1.4E-22 | 1.0582814 | 0.129 | 0.059 | 4.61E-18 | Chondrogenic Fibroblasts | bmp6 |
| coro1ca4 | 1.5E-22 | 0.4082823 | 0.373 | 0.235 | 4.81E-18 | Chondrogenic Fibroblasts | coro1ca |
| ap1s11 | 1.5E-22 | 1.0113126 | 0.107 | 0.045 | 4.87E-18 | Chondrogenic Fibroblasts | ap1s1 |
| flrt22 | 1.5E-22 | 0.6885277 | 0.154 | 0.074 | 4.94E-18 | Chondrogenic Fibroblasts | flrt2 |
| si:ch73-269m14.21 | 1.5E-22 | 0.3689602 | 0.366 | 0.225 | 5.03E-18 | Chondrogenic Fibroblasts | si:ch73-269m14.2 |
| hnrnpd | 1.6E-22 | 0.4607727 | 0.304 | 0.18 | 5.28E-18 | Chondrogenic Fibroblasts | hnrnpd |
| ciao2b1 | 1.6E-22 | 0.5889528 | 0.234 | 0.131 | 5.36E-18 | Chondrogenic Fibroblasts | ciao2b |
| smim72 | 1.8E-22 | 0.537469 | 0.238 | 0.132 | 5.94E-18 | Chondrogenic Fibroblasts | smim7 |
| higd1a4 | 2E-22 | 0.3932351 | 0.62 | 0.454 | 6.51E-18 | Chondrogenic Fibroblasts | higd1a |
| zgc:1135315 | 2.1E-22 | 0.53819 | 0.418 | 0.275 | 6.96E-18 | Chondrogenic Fibroblasts | zgc:113531 |
| cox8a3 | 2.3E-22 | 0.3129386 | 0.727 | 0.559 | 7.36E-18 | Chondrogenic Fibroblasts | cox8a |
| kctd3 | 2.4E-22 | 0.7414091 | 0.186 | 0.097 | 7.91E-18 | Chondrogenic Fibroblasts | kctd3 |
| cox4i11 | 2.5E-22 | 0.2997766 | 0.893 | 0.775 | 8.2E-18 | Chondrogenic Fibroblasts | cox4i1 |
| sel1l1 | 2.5E-22 | 0.8232185 | 0.132 | 0.061 | 8.26E-18 | Chondrogenic Fibroblasts | sel1l |
| sypl2b | 2.6E-22 | 0.6066662 | 0.207 | 0.111 | 8.43E-18 | Chondrogenic Fibroblasts | sypl2b |
| hypk1 | 2.8E-22 | 0.3873233 | 0.431 | 0.282 | 9.27E-18 | Chondrogenic Fibroblasts | hypk |
| zgc:1231051 | 2.9E-22 | 0.6414827 | 0.155 | 0.075 | 9.32E-18 | Chondrogenic Fibroblasts | zgc:123105 |
| rabif1 | 2.9E-22 | 0.7549218 | 0.135 | 0.062 | 9.5E-18 | Chondrogenic Fibroblasts | rabif |
| ndufa101 | 4.1E-22 | 0.4892901 | 0.216 | 0.117 | 1.32E-17 | Chondrogenic Fibroblasts | ndufa10 |
| nudt51 | 4.2E-22 | 0.9117609 | 0.11 | 0.047 | 1.37E-17 | Chondrogenic Fibroblasts | nudt5 |
| psmc41 | 4.8E-22 | 0.5197364 | 0.277 | 0.163 | 1.57E-17 | Chondrogenic Fibroblasts | psmc4 |
| uqcrc2b3 | 5.4E-22 | 0.587082 | 0.207 | 0.112 | 1.75E-17 | Chondrogenic Fibroblasts | uqcrc2b |

| | | | | | | | |
|---|---|---|---|---|---|---|---|
| hadhb2 | 6.2E-22 | 0.5964834 | 0.231 | 0.13 | 2.02E-17 | Chondrogenic Fibroblasts | hadhb |
| ndufa9a1 | 6.8E-22 | 0.6884881 | 0.193 | 0.103 | 2.2E-17 | Chondrogenic Fibroblasts | ndufa9a |
| si:dkey-204f11.641 | 7.1E-22 | 0.7449457 | 0.158 | 0.078 | 2.3E-17 | Chondrogenic Fibroblasts | si:dkey-204f11.64 |
| arhgef101 | 7.1E-22 | 0.8703974 | 0.107 | 0.046 | 2.32E-17 | Chondrogenic Fibroblasts | arhgef10 |
| eif1ad3 | 7.4E-22 | 0.5093914 | 0.244 | 0.138 | 2.41E-17 | Chondrogenic Fibroblasts | eif1ad |
| kif1b2 | 7.8E-22 | 0.5680815 | 0.189 | 0.099 | 2.55E-17 | Chondrogenic Fibroblasts | kif1b |
| idh24 | 8.4E-22 | 0.3863614 | 0.499 | 0.341 | 2.73E-17 | Chondrogenic Fibroblasts | idh2 |
| perp1 | 8.6E-22 | 0.4198658 | 0.369 | 0.234 | 2.78E-17 | Chondrogenic Fibroblasts | perp |
| calcoco1a2 | 9.1E-22 | 0.7553947 | 0.189 | 0.1 | 2.97E-17 | Chondrogenic Fibroblasts | calcoco1a |
| skib6 | 9.4E-22 | 0.3999062 | 0.327 | 0.2 | 3.05E-17 | Chondrogenic Fibroblasts | skib |
| spoplb1 | 9.7E-22 | 0.6527994 | 0.185 | 0.097 | 3.17E-17 | Chondrogenic Fibroblasts | spoplb |
| mafa2 | 1E-21 | 1.237187 | 0.109 | 0.048 | 3.33E-17 | Chondrogenic Fibroblasts | mafa |
| dda11 | 1.1E-21 | 0.4799659 | 0.265 | 0.153 | 3.69E-17 | Chondrogenic Fibroblasts | dda1 |
| eny21 | 1.2E-21 | 0.6771463 | 0.198 | 0.106 | 3.97E-17 | Chondrogenic Fibroblasts | eny2 |
| CR381618.16 | 1.5E-21 | 0.4519157 | 0.354 | 0.225 | 4.78E-17 | Chondrogenic Fibroblasts | CR381618.1 |
| ctsk5 | 1.5E-21 | 0.3640484 | 0.741 | 0.589 | 4.94E-17 | Chondrogenic Fibroblasts | ctsk |
| faf21 | 1.8E-21 | 0.4838635 | 0.243 | 0.138 | 5.74E-17 | Chondrogenic Fibroblasts | faf2 |
| ilf22 | 1.8E-21 | 0.4648773 | 0.27 | 0.158 | 5.91E-17 | Chondrogenic Fibroblasts | ilf2 |
| clint1a2 | 1.8E-21 | 0.3799721 | 0.334 | 0.205 | 5.94E-17 | Chondrogenic Fibroblasts | clint1a |
| sgcb | 1.8E-21 | 0.7751765 | 0.15 | 0.074 | 6.01E-17 | Chondrogenic Fibroblasts | sgcb |
| gnai11 | 1.9E-21 | 0.47067 | 0.268 | 0.156 | 6.03E-17 | Chondrogenic Fibroblasts | gnai1 |
| p3h13 | 2E-21 | 0.4539254 | 0.338 | 0.211 | 6.53E-17 | Chondrogenic Fibroblasts | p3h1 |
| pik3r3b3 | 2.2E-21 | 0.4877909 | 0.409 | 0.271 | 7.24E-17 | Chondrogenic Fibroblasts | pik3r3b |
| syf21 | 2.4E-21 | 0.386292 | 0.423 | 0.279 | 7.73E-17 | Chondrogenic Fibroblasts | syf2 |
| ndrg43 | 2.6E-21 | 0.5944394 | 0.26 | 0.152 | 8.53E-17 | Chondrogenic Fibroblasts | ndrg4 |
| myo1ea3 | 3.3E-21 | 0.8447151 | 0.159 | 0.08 | 1.07E-16 | Chondrogenic Fibroblasts | myo1ea |
| ccdc251 | 3.4E-21 | 0.4555401 | 0.293 | 0.176 | 1.11E-16 | Chondrogenic Fibroblasts | ccdc25 |
| srsf42 | 3.4E-21 | 0.4966581 | 0.272 | 0.16 | 1.12E-16 | Chondrogenic Fibroblasts | srsf4 |
| hmcn13 | 3.6E-21 | 0.5936919 | 0.147 | 0.071 | 1.17E-16 | Chondrogenic Fibroblasts | hmcn1 |
| psmc1a1 | 3.6E-21 | 0.5210911 | 0.207 | 0.112 | 1.19E-16 | Chondrogenic Fibroblasts | psmc1a |
| arf12 | 3.9E-21 | 0.4641402 | 0.257 | 0.148 | 1.27E-16 | Chondrogenic Fibroblasts | arf1 |
| micos102 | 3.9E-21 | 0.4824127 | 0.282 | 0.169 | 1.28E-16 | Chondrogenic Fibroblasts | micos10 |
| timm17a1 | 4.4E-21 | 0.5533785 | 0.195 | 0.104 | 1.43E-16 | Chondrogenic Fibroblasts | timm17a |
| serpinh23 | 4.6E-21 | 0.5855388 | 0.211 | 0.116 | 1.51E-16 | Chondrogenic Fibroblasts | serpinh2 |
| cald1b7 | 4.7E-21 | 1.1517843 | 0.268 | 0.166 | 1.53E-16 | Chondrogenic Fibroblasts | cald1b |
| marveld13 | 4.9E-21 | 0.5260156 | 0.33 | 0.207 | 1.6E-16 | Chondrogenic Fibroblasts | marveld1 |
| rtn4a1 | 5.6E-21 | 0.2946892 | 0.623 | 0.446 | 1.81E-16 | Chondrogenic Fibroblasts | rtn4a |
| calub1 | 6.1E-21 | 0.659182 | 0.137 | 0.065 | 2E-16 | Chondrogenic Fibroblasts | calub |
| sec24d1 | 6.5E-21 | 0.8075325 | 0.115 | 0.051 | 2.11E-16 | Chondrogenic Fibroblasts | sec24d |
| dctn21 | 6.7E-21 | 0.5353445 | 0.223 | 0.125 | 2.17E-16 | Chondrogenic Fibroblasts | dctn2 |
| zgc:1536753 | 7E-21 | 0.4329335 | 0.398 | 0.259 | 2.27E-16 | Chondrogenic Fibroblasts | zgc:153675 |
| ARF52 | 7.5E-21 | 0.5599147 | 0.258 | 0.15 | 2.43E-16 | Chondrogenic Fibroblasts | ARF5 |
| tbcb1 | 8.6E-21 | 0.3875737 | 0.344 | 0.215 | 2.8E-16 | Chondrogenic Fibroblasts | tbcb |
| ywhabl2 | 8.7E-21 | 0.3300889 | 0.584 | 0.414 | 2.84E-16 | Chondrogenic Fibroblasts | ywhabl |
| eif4a1a3 | 1E-20 | 0.3396883 | 0.519 | 0.362 | 3.36E-16 | Chondrogenic Fibroblasts | eif4a1a |
| mfap25 | 1.1E-20 | 0.3290058 | 0.543 | 0.38 | 3.58E-16 | Chondrogenic Fibroblasts | mfap2 |
| zgc:1034383 | 1.1E-20 | 0.657934 | 0.168 | 0.087 | 3.61E-16 | Chondrogenic Fibroblasts | zgc:103438 |
| tomm20b3 | 1.2E-20 | 0.3723671 | 0.498 | 0.34 | 3.9E-16 | Chondrogenic Fibroblasts | tomm20b |

| | | | | | | | |
|---|---|---|---|---|---|---|---|
| pla2g15 | 1.2E-20 | 0.9020818 | 0.118 | 0.053 | 4E-16 | Chondrogenic Fibroblasts | pla2g15 |
| CTBP12 | 1.2E-20 | 0.4868958 | 0.279 | 0.166 | 4.01E-16 | Chondrogenic Fibroblasts | CTBP1 |
| vps4b1 | 1.6E-20 | 0.3225141 | 0.396 | 0.256 | 5.11E-16 | Chondrogenic Fibroblasts | vps4b |
| slc44a23 | 1.8E-20 | 0.4413449 | 0.208 | 0.113 | 5.73E-16 | Chondrogenic Fibroblasts | slc44a2 |
| lsm72 | 1.8E-20 | 0.3766829 | 0.395 | 0.257 | 5.87E-16 | Chondrogenic Fibroblasts | lsm7 |
| hectd2 | 1.9E-20 | 0.9844992 | 0.1 | 0.042 | 6.11E-16 | Chondrogenic Fibroblasts | hectd2 |
| sox4a4 | 2E-20 | 0.5225544 | 0.308 | 0.192 | 6.43E-16 | Chondrogenic Fibroblasts | sox4a |
| ing4 | 2E-20 | 0.8193448 | 0.107 | 0.047 | 6.66E-16 | Chondrogenic Fibroblasts | ing4 |
| napab1 | 2.1E-20 | 0.4002157 | 0.326 | 0.202 | 6.89E-16 | Chondrogenic Fibroblasts | napab |
| ikbip3 | 2.3E-20 | 0.5870511 | 0.21 | 0.117 | 7.36E-16 | Chondrogenic Fibroblasts | ikbip |
| atp5f1d3 | 2.3E-20 | 0.3404048 | 0.479 | 0.331 | 7.58E-16 | Chondrogenic Fibroblasts | atp5f1d |
| hmgb2b1 | 2.4E-20 | 0.3113933 | 0.519 | 0.36 | 7.74E-16 | Chondrogenic Fibroblasts | hmgb2b |
| mt-nd11 | 2.5E-20 | 0.2445153 | 0.866 | 0.715 | 7.98E-16 | Chondrogenic Fibroblasts | mt-nd1 |
| CU694197.11 | 2.8E-20 | 0.4826411 | 0.325 | 0.204 | 9.09E-16 | Chondrogenic Fibroblasts | CU694197.1 |
| lmcd13 | 2.8E-20 | 0.6660887 | 0.168 | 0.087 | 9.15E-16 | Chondrogenic Fibroblasts | lmcd1 |
| atp6ap1b1 | 2.9E-20 | 0.7386533 | 0.139 | 0.068 | 9.29E-16 | Chondrogenic Fibroblasts | atp6ap1b |
| ubqln41 | 3E-20 | 0.5106139 | 0.239 | 0.138 | 9.89E-16 | Chondrogenic Fibroblasts | ubqln4 |
| ext21 | 3.2E-20 | 0.6869728 | 0.192 | 0.104 | 1.06E-15 | Chondrogenic Fibroblasts | ext2 |
| tmem167b | 3.3E-20 | 0.9568933 | 0.101 | 0.044 | 1.06E-15 | Chondrogenic Fibroblasts | tmem167b |
| cdv31 | 3.3E-20 | 0.3113704 | 0.372 | 0.235 | 1.06E-15 | Chondrogenic Fibroblasts | cdv3 |
| lrch31 | 3.3E-20 | 0.7682032 | 0.153 | 0.077 | 1.09E-15 | Chondrogenic Fibroblasts | lrch3 |
| kras2 | 3.8E-20 | 0.4692325 | 0.29 | 0.176 | 1.25E-15 | Chondrogenic Fibroblasts | kras |
| ube2l3b1 | 3.9E-20 | 0.3721759 | 0.332 | 0.207 | 1.25E-15 | Chondrogenic Fibroblasts | ube2l3b |
| lepr3 | 4.4E-20 | 0.7764499 | 0.131 | 0.062 | 1.42E-15 | Chondrogenic Fibroblasts | lepr |
| gtf2h51 | 4.5E-20 | 0.7101153 | 0.18 | 0.096 | 1.45E-15 | Chondrogenic Fibroblasts | gtf2h5 |
| tcta3 | 4.5E-20 | 0.7219325 | 0.168 | 0.088 | 1.47E-15 | Chondrogenic Fibroblasts | tcta |
| otud5b2 | 4.6E-20 | 0.6868399 | 0.162 | 0.083 | 1.49E-15 | Chondrogenic Fibroblasts | otud5b |
| antxr1b1 | 4.8E-20 | 0.8121626 | 0.112 | 0.05 | 1.57E-15 | Chondrogenic Fibroblasts | antxr1b |
| emc21 | 5.3E-20 | 0.4763088 | 0.319 | 0.2 | 1.71E-15 | Chondrogenic Fibroblasts | emc2 |
| mfsd13a | 5.5E-20 | 0.6748763 | 0.122 | 0.056 | 1.79E-15 | Chondrogenic Fibroblasts | mfsd13a |
| sertad2b1 | 5.9E-20 | 0.801845 | 0.104 | 0.045 | 1.92E-15 | Chondrogenic Fibroblasts | sertad2b |
| pacsin21 | 6.4E-20 | 0.7478976 | 0.131 | 0.062 | 2.1E-15 | Chondrogenic Fibroblasts | pacsin2 |
| ndufb41 | 6.6E-20 | 0.3285371 | 0.472 | 0.322 | 2.14E-15 | Chondrogenic Fibroblasts | ndufb4 |
| higd2a2 | 6.7E-20 | 0.5327389 | 0.231 | 0.134 | 2.18E-15 | Chondrogenic Fibroblasts | higd2a |
| pam1 | 6.7E-20 | 0.6224064 | 0.18 | 0.096 | 2.19E-15 | Chondrogenic Fibroblasts | pam |
| gars12 | 7E-20 | 0.4930054 | 0.25 | 0.147 | 2.29E-15 | Chondrogenic Fibroblasts | gars1 |
| rgcc3 | 7.4E-20 | 0.4062128 | 0.313 | 0.196 | 2.39E-15 | Chondrogenic Fibroblasts | rgcc |
| c1qtnf6b2 | 7.6E-20 | 0.7653559 | 0.149 | 0.075 | 2.49E-15 | Chondrogenic Fibroblasts | c1qtnf6b |
| cndp22 | 8E-20 | 0.4613184 | 0.199 | 0.109 | 2.6E-15 | Chondrogenic Fibroblasts | cndp2 |
| chd56 | 8.4E-20 | 0.460829 | 0.223 | 0.125 | 2.73E-15 | Chondrogenic Fibroblasts | chd5 |
| cblb2 | 8.4E-20 | 0.6887837 | 0.178 | 0.095 | 2.73E-15 | Chondrogenic Fibroblasts | cblb |
| yipf62 | 8.6E-20 | 0.6202627 | 0.151 | 0.076 | 2.8E-15 | Chondrogenic Fibroblasts | yipf6 |
| cnpy12 | 8.7E-20 | 0.8631759 | 0.154 | 0.079 | 2.81E-15 | Chondrogenic Fibroblasts | cnpy1 |
| trappc11 | 9.3E-20 | 0.4105006 | 0.247 | 0.143 | 3.01E-15 | Chondrogenic Fibroblasts | trappc1 |
| gabarapl22 | 1E-19 | 0.4465276 | 0.271 | 0.162 | 3.34E-15 | Chondrogenic Fibroblasts | gabarapl2 |
| erp441 | 1E-19 | 0.4393852 | 0.297 | 0.184 | 3.38E-15 | Chondrogenic Fibroblasts | erp44 |
| zgc:926061 | 1.1E-19 | 0.4892696 | 0.229 | 0.131 | 3.63E-15 | Chondrogenic Fibroblasts | zgc:92606 |
| cox6c3 | 1.2E-19 | 0.3129802 | 0.578 | 0.411 | 3.74E-15 | Chondrogenic Fibroblasts | cox6c |

| | | | | | | | |
|---|---|---|---|---|---|---|---|
| smndc11 | 1.2E-19 | 0.3984864 | 0.329 | 0.205 | 3.76E-15 | Chondrogenic Fibroblasts | smndc1 |
| emilin1a5 | 1.3E-19 | 0.6652647 | 0.174 | 0.092 | 4.22E-15 | Chondrogenic Fibroblasts | emilin1a |
| rnf25 | 1.3E-19 | 0.8023359 | 0.105 | 0.046 | 4.26E-15 | Chondrogenic Fibroblasts | rnf25 |
| sri1 | 1.4E-19 | 0.3600062 | 0.405 | 0.268 | 4.64E-15 | Chondrogenic Fibroblasts | sri |
| thrap3a4 | 1.5E-19 | 0.3007714 | 0.537 | 0.38 | 4.8E-15 | Chondrogenic Fibroblasts | thrap3a |
| syngr2a | 1.5E-19 | 0.6318507 | 0.131 | 0.062 | 4.99E-15 | Chondrogenic Fibroblasts | syngr2a |
| cpda1 | 1.6E-19 | 0.6284306 | 0.154 | 0.078 | 5.19E-15 | Chondrogenic Fibroblasts | cpda |
| cd1643 | 1.6E-19 | 0.301004 | 0.486 | 0.334 | 5.2E-15 | Chondrogenic Fibroblasts | cd164 |
| tmem14ca2 | 1.8E-19 | 0.4897417 | 0.213 | 0.12 | 6.01E-15 | Chondrogenic Fibroblasts | tmem14ca |
| eif4e31 | 1.9E-19 | 0.6340504 | 0.129 | 0.062 | 6.07E-15 | Chondrogenic Fibroblasts | eif4e3 |
| ankrd13c | 2E-19 | 0.7127425 | 0.116 | 0.053 | 6.37E-15 | Chondrogenic Fibroblasts | ankrd13c |
| prkacab | 2.1E-19 | 0.7736256 | 0.123 | 0.058 | 6.93E-15 | Chondrogenic Fibroblasts | prkacab |
| mbd22 | 2.1E-19 | 0.6080228 | 0.158 | 0.081 | 6.95E-15 | Chondrogenic Fibroblasts | mbd2 |
| dtd12 | 2.5E-19 | 0.7904429 | 0.107 | 0.048 | 8.14E-15 | Chondrogenic Fibroblasts | dtd1 |
| stau22 | 2.5E-19 | 0.2962962 | 0.604 | 0.435 | 8.18E-15 | Chondrogenic Fibroblasts | stau2 |
| ptmaa2 | 3E-19 | 0.3333893 | 0.913 | 0.834 | 9.76E-15 | Chondrogenic Fibroblasts | ptmaa |
| stxbp31 | 3.1E-19 | 0.3747526 | 0.265 | 0.157 | 1.02E-14 | Chondrogenic Fibroblasts | stxbp3 |
| sh3bgrl31 | 3.2E-19 | 0.4600406 | 0.229 | 0.132 | 1.03E-14 | Chondrogenic Fibroblasts | sh3bgrl3 |
| arf6a1 | 3.2E-19 | 0.4947097 | 0.196 | 0.108 | 1.04E-14 | Chondrogenic Fibroblasts | arf6a |
| smarce11 | 3.2E-19 | 0.5494208 | 0.249 | 0.149 | 1.05E-14 | Chondrogenic Fibroblasts | smarce1 |
| lysmd41 | 3.3E-19 | 0.9055037 | 0.103 | 0.046 | 1.07E-14 | Chondrogenic Fibroblasts | lysmd4 |
| mycbp3 | 3.4E-19 | 0.3815377 | 0.375 | 0.246 | 1.11E-14 | Chondrogenic Fibroblasts | mycbp |
| stx182 | 3.8E-19 | 0.7429329 | 0.115 | 0.053 | 1.23E-14 | Chondrogenic Fibroblasts | stx18 |
| bmp54 | 3.9E-19 | 0.8523943 | 0.163 | 0.087 | 1.26E-14 | Chondrogenic Fibroblasts | bmp5 |
| mapk63 | 4.4E-19 | 0.5847475 | 0.164 | 0.086 | 1.43E-14 | Chondrogenic Fibroblasts | mapk6 |
| chac12 | 4.4E-19 | 0.4038632 | 0.497 | 0.351 | 1.45E-14 | Chondrogenic Fibroblasts | chac1 |
| si:ch211-276i12.111 | 4.5E-19 | 0.5488396 | 0.187 | 0.102 | 1.45E-14 | Chondrogenic Fibroblasts | si:ch211-276i12.11 |
| fis1 | 4.5E-19 | 0.4923892 | 0.209 | 0.118 | 1.47E-14 | Chondrogenic Fibroblasts | fis1 |
| cers2b1 | 4.5E-19 | 0.7221997 | 0.135 | 0.067 | 1.48E-14 | Chondrogenic Fibroblasts | cers2b |
| tmem592 | 4.6E-19 | 0.3442961 | 0.454 | 0.311 | 1.5E-14 | Chondrogenic Fibroblasts | tmem59 |
| vps292 | 4.6E-19 | 0.4438418 | 0.314 | 0.199 | 1.5E-14 | Chondrogenic Fibroblasts | vps29 |
| pnp5a1 | 5E-19 | 0.6053609 | 0.121 | 0.057 | 1.62E-14 | Chondrogenic Fibroblasts | pnp5a |
| sdha1 | 5.4E-19 | 0.5123069 | 0.252 | 0.151 | 1.77E-14 | Chondrogenic Fibroblasts | sdha |
| sec61b3 | 5.6E-19 | 0.3787838 | 0.739 | 0.579 | 1.83E-14 | Chondrogenic Fibroblasts | sec61b |
| adtrp11 | 5.9E-19 | 0.598563 | 0.155 | 0.08 | 1.92E-14 | Chondrogenic Fibroblasts | adtrp1 |
| p3h41 | 6.4E-19 | 0.8753366 | 0.104 | 0.047 | 2.07E-14 | Chondrogenic Fibroblasts | p3h4 |
| btbd10b1 | 6.4E-19 | 0.5691983 | 0.189 | 0.104 | 2.09E-14 | Chondrogenic Fibroblasts | btbd10b |
| aamdc | 6.5E-19 | 0.7173922 | 0.115 | 0.053 | 2.12E-14 | Chondrogenic Fibroblasts | aamdc |
| atp5f1e3 | 6.7E-19 | 0.3762972 | 0.564 | 0.409 | 2.17E-14 | Chondrogenic Fibroblasts | atp5f1e |
| ube2v11 | 7.3E-19 | 0.3847711 | 0.395 | 0.263 | 2.37E-14 | Chondrogenic Fibroblasts | ube2v1 |
| rbbp42 | 7.4E-19 | 0.5324302 | 0.172 | 0.091 | 2.4E-14 | Chondrogenic Fibroblasts | rbbp4 |
| smarca51 | 8.2E-19 | 0.7958813 | 0.104 | 0.047 | 2.68E-14 | Chondrogenic Fibroblasts | smarca5 |
| selenot1a3 | 8.3E-19 | 0.3526662 | 0.277 | 0.167 | 2.69E-14 | Chondrogenic Fibroblasts | selenot1a |
| htra1a3 | 8.4E-19 | 0.4070122 | 0.346 | 0.225 | 2.74E-14 | Chondrogenic Fibroblasts | htra1a |
| psma31 | 8.7E-19 | 0.2605286 | 0.382 | 0.246 | 2.81E-14 | Chondrogenic Fibroblasts | psma3 |
| abce12 | 8.7E-19 | 0.5362396 | 0.212 | 0.121 | 2.84E-14 | Chondrogenic Fibroblasts | abce1 |
| ndufa82 | 9.4E-19 | 0.3578541 | 0.355 | 0.231 | 3.06E-14 | Chondrogenic Fibroblasts | ndufa8 |
| snx32 | 9.8E-19 | 0.3142598 | 0.29 | 0.176 | 3.2E-14 | Chondrogenic Fibroblasts | snx3 |

| | | | | | | | |
|---|---|---|---|---|---|---|---|
| CABZ01026681.12 | 9.9E-19 | 0.5419894 | 0.158 | 0.082 | 3.21E-14 | Chondrogenic Fibroblasts | CABZ01026681.1 |
| eif1axb1 | 1E-18 | 0.2858387 | 0.447 | 0.301 | 3.29E-14 | Chondrogenic Fibroblasts | eif1axb |
| CABZ01058650.13 | 1.1E-18 | 0.6647918 | 0.142 | 0.071 | 3.61E-14 | Chondrogenic Fibroblasts | CABZ01058650.1 |
| sec22bb1 | 1.1E-18 | 0.8654888 | 0.109 | 0.05 | 3.71E-14 | Chondrogenic Fibroblasts | sec22bb |
| fkbp1b2 | 1.1E-18 | 0.6734277 | 0.143 | 0.073 | 3.72E-14 | Chondrogenic Fibroblasts | fkbp1b |
| lmf2b1 | 1.2E-18 | 0.4999519 | 0.233 | 0.135 | 3.85E-14 | Chondrogenic Fibroblasts | lmf2b |
| emd1 | 1.3E-18 | 0.3320594 | 0.512 | 0.358 | 4.15E-14 | Chondrogenic Fibroblasts | emd |
| ddx39aa1 | 1.3E-18 | 0.3876868 | 0.293 | 0.18 | 4.25E-14 | Chondrogenic Fibroblasts | ddx39aa |
| trir1 | 1.3E-18 | 0.7017494 | 0.142 | 0.072 | 4.35E-14 | Chondrogenic Fibroblasts | trir |
| mex3b4 | 1.5E-18 | 0.4003455 | 0.279 | 0.17 | 4.86E-14 | Chondrogenic Fibroblasts | mex3b |
| CR847953.1 | 1.6E-18 | 0.9690191 | 0.103 | 0.047 | 5.22E-14 | Chondrogenic Fibroblasts | CR847953.1 |
| sdhaf41 | 1.9E-18 | 0.9197963 | 0.115 | 0.055 | 6.09E-14 | Chondrogenic Fibroblasts | sdhaf4 |
| tmem30ab1 | 2E-18 | 0.4551149 | 0.238 | 0.14 | 6.37E-14 | Chondrogenic Fibroblasts | tmem30ab |
| jagn1b2 | 2.1E-18 | 0.5011433 | 0.261 | 0.16 | 6.68E-14 | Chondrogenic Fibroblasts | jagn1b |
| ube2h2 | 2.1E-18 | 0.585978 | 0.194 | 0.109 | 6.9E-14 | Chondrogenic Fibroblasts | ube2h |
| gale2 | 2.1E-18 | 0.7506872 | 0.112 | 0.052 | 6.91E-14 | Chondrogenic Fibroblasts | gale |
| cog42 | 2.1E-18 | 0.6672094 | 0.117 | 0.055 | 6.96E-14 | Chondrogenic Fibroblasts | cog4 |
| lsm52 | 2.2E-18 | 0.4692024 | 0.205 | 0.116 | 7.03E-14 | Chondrogenic Fibroblasts | lsm5 |
| si:ch73-238c9.13 | 2.2E-18 | 0.6363338 | 0.184 | 0.102 | 7.14E-14 | Chondrogenic Fibroblasts | si:ch73-238c9.1 |
| kalrnb1 | 2.2E-18 | 0.6390529 | 0.168 | 0.091 | 7.18E-14 | Chondrogenic Fibroblasts | kalrnb |
| ccdc1241 | 2.3E-18 | 0.4437997 | 0.274 | 0.168 | 7.36E-14 | Chondrogenic Fibroblasts | ccdc124 |
| pdcd6ip1 | 2.3E-18 | 0.5185811 | 0.224 | 0.13 | 7.53E-14 | Chondrogenic Fibroblasts | pdcd6ip |
| sra13 | 2.4E-18 | 0.7876821 | 0.129 | 0.063 | 7.75E-14 | Chondrogenic Fibroblasts | sra1 |
| sdhdb1 | 2.4E-18 | 0.5316463 | 0.217 | 0.126 | 7.91E-14 | Chondrogenic Fibroblasts | sdhdb |
| trmt61a | 2.7E-18 | 0.7921377 | 0.115 | 0.054 | 8.87E-14 | Chondrogenic Fibroblasts | trmt61a |
| ddx39ab1 | 2.7E-18 | 0.2856293 | 0.276 | 0.166 | 8.9E-14 | Chondrogenic Fibroblasts | ddx39ab |
| tmem1471 | 3E-18 | 0.5237846 | 0.249 | 0.15 | 9.84E-14 | Chondrogenic Fibroblasts | tmem147 |
| calm3a.11 | 3.3E-18 | 0.3991064 | 0.539 | 0.395 | 1.07E-13 | Chondrogenic Fibroblasts | calm3a.1 |
| srp192 | 3.5E-18 | 0.446341 | 0.235 | 0.139 | 1.13E-13 | Chondrogenic Fibroblasts | srp19 |
| lsm41 | 3.5E-18 | 0.3708661 | 0.335 | 0.215 | 1.15E-13 | Chondrogenic Fibroblasts | lsm4 |
| uggt11 | 3.6E-18 | 0.675615 | 0.112 | 0.052 | 1.16E-13 | Chondrogenic Fibroblasts | uggt1 |
| cebpg1 | 3.6E-18 | 0.388645 | 0.234 | 0.136 | 1.17E-13 | Chondrogenic Fibroblasts | cebpg |
| mtdhb1 | 3.7E-18 | 0.3994428 | 0.253 | 0.151 | 1.2E-13 | Chondrogenic Fibroblasts | mtdhb |
| stx4 | 3.8E-18 | 0.6781563 | 0.131 | 0.065 | 1.23E-13 | Chondrogenic Fibroblasts | stx4 |
| elob.1 | 3.9E-18 | 0.3311853 | 0.35 | 0.226 | 1.26E-13 | Chondrogenic Fibroblasts | elob.1 |
| ciapin12 | 3.9E-18 | 0.7269503 | 0.125 | 0.061 | 1.27E-13 | Chondrogenic Fibroblasts | ciapin1 |
| txnl4a2 | 4.1E-18 | 0.5736577 | 0.152 | 0.079 | 1.33E-13 | Chondrogenic Fibroblasts | txnl4a |
| top1l1 | 4.1E-18 | 0.2844656 | 0.552 | 0.39 | 1.35E-13 | Chondrogenic Fibroblasts | top1l |
| ube2d22 | 4.3E-18 | 0.3633287 | 0.531 | 0.386 | 1.39E-13 | Chondrogenic Fibroblasts | ube2d2 |
| pdha1a1 | 4.4E-18 | 0.4997154 | 0.228 | 0.134 | 1.44E-13 | Chondrogenic Fibroblasts | pdha1a |
| anapc161 | 4.9E-18 | 0.5122571 | 0.241 | 0.145 | 1.59E-13 | Chondrogenic Fibroblasts | anapc16 |
| polr2l1 | 5E-18 | 0.3601186 | 0.307 | 0.192 | 1.63E-13 | Chondrogenic Fibroblasts | polr2l |
| arpin1 | 5E-18 | 0.6257765 | 0.133 | 0.066 | 1.64E-13 | Chondrogenic Fibroblasts | arpin |
| gulp1a4 | 5.2E-18 | 0.4536114 | 0.296 | 0.186 | 1.7E-13 | Chondrogenic Fibroblasts | gulp1a |
| aco21 | 5.4E-18 | 0.447243 | 0.154 | 0.08 | 1.75E-13 | Chondrogenic Fibroblasts | aco2 |
| si:ch73-147o17.11 | 5.6E-18 | 0.6252626 | 0.15 | 0.078 | 1.81E-13 | Chondrogenic Fibroblasts | si:ch73-147o17.1 |
| neo1a3 | 5.9E-18 | 0.3666214 | 0.29 | 0.18 | 1.92E-13 | Chondrogenic Fibroblasts | neo1a |
| zgc:635871 | 6E-18 | 0.4444376 | 0.159 | 0.083 | 1.94E-13 | Chondrogenic Fibroblasts | zgc:63587 |

| | | | | | | | |
|---|---|---|---|---|---|---|---|
| si:ch211-284f22.34 | 6.3E-18 | 0.4210112 | 0.204 | 0.115 | 2.06E-13 | Chondrogenic Fibroblasts | si:ch211-284f22.3 |
| sat1a.26 | 6.5E-18 | 0.2622614 | 0.615 | 0.454 | 2.11E-13 | Chondrogenic Fibroblasts | sat1a.2 |
| rab5ab1 | 6.5E-18 | 0.3732012 | 0.379 | 0.252 | 2.11E-13 | Chondrogenic Fibroblasts | rab5ab |
| tln11 | 6.6E-18 | 0.232234 | 0.766 | 0.607 | 2.15E-13 | Chondrogenic Fibroblasts | tln1 |
| ndufb101 | 7E-18 | 0.3234667 | 0.36 | 0.233 | 2.29E-13 | Chondrogenic Fibroblasts | ndufb10 |
| capns1a1 | 7.3E-18 | 0.3163133 | 0.55 | 0.397 | 2.38E-13 | Chondrogenic Fibroblasts | capns1a |
| plcd1a2 | 7.6E-18 | 0.5346494 | 0.174 | 0.095 | 2.48E-13 | Chondrogenic Fibroblasts | plcd1a |
| psma6a1 | 7.7E-18 | 0.4944621 | 0.22 | 0.129 | 2.51E-13 | Chondrogenic Fibroblasts | psma6a |
| aspa1 | 7.8E-18 | 0.6900616 | 0.121 | 0.059 | 2.55E-13 | Chondrogenic Fibroblasts | aspa |
| ilf3b1 | 7.8E-18 | 0.4504609 | 0.295 | 0.185 | 2.55E-13 | Chondrogenic Fibroblasts | ilf3b |
| aldh1a23 | 8.1E-18 | 0.9069575 | 0.113 | 0.054 | 2.63E-13 | Chondrogenic Fibroblasts | aldh1a2 |
| atp6v1ba1 | 8.3E-18 | 0.9307497 | 0.116 | 0.056 | 2.71E-13 | Chondrogenic Fibroblasts | atp6v1ba |
| si:ch211-214j24.101 | 9.5E-18 | 0.4228453 | 0.351 | 0.232 | 3.08E-13 | Chondrogenic Fibroblasts | si:ch211-214j24.10 |
| sub1a1 | 9.9E-18 | 0.3225144 | 0.47 | 0.33 | 3.23E-13 | Chondrogenic Fibroblasts | sub1a |
| supt3h1 | 1.1E-17 | 0.7487321 | 0.119 | 0.058 | 3.53E-13 | Chondrogenic Fibroblasts | supt3h |
| kctd5a4 | 1.3E-17 | 0.3453642 | 0.454 | 0.317 | 4.24E-13 | Chondrogenic Fibroblasts | kctd5a |
| si:dkey-100n23.5 | 1.4E-17 | 0.7557251 | 0.108 | 0.051 | 4.41E-13 | Chondrogenic Fibroblasts | si:dkey-100n23.5 |
| hyal12 | 1.4E-17 | 0.7447294 | 0.114 | 0.055 | 4.46E-13 | Chondrogenic Fibroblasts | hyal1 |
| tgfbi5 | 1.4E-17 | 0.4876262 | 0.307 | 0.198 | 4.61E-13 | Chondrogenic Fibroblasts | tgfbi |
| max2 | 1.5E-17 | 0.3574705 | 0.418 | 0.287 | 4.78E-13 | Chondrogenic Fibroblasts | max |
| tpm32 | 1.5E-17 | 0.3111807 | 0.674 | 0.523 | 4.85E-13 | Chondrogenic Fibroblasts | tpm3 |
| tbc1d10aa1 | 1.5E-17 | 0.4364899 | 0.201 | 0.114 | 4.91E-13 | Chondrogenic Fibroblasts | tbc1d10aa |
| serpinb143 | 1.6E-17 | 0.3709666 | 0.252 | 0.151 | 5.19E-13 | Chondrogenic Fibroblasts | serpinb14 |
| rnasekb1 | 1.6E-17 | 0.5210816 | 0.186 | 0.104 | 5.35E-13 | Chondrogenic Fibroblasts | rnasekb |
| si:dkeyp-27c8.22 | 1.7E-17 | 0.7813132 | 0.104 | 0.048 | 5.45E-13 | Chondrogenic Fibroblasts | si:dkeyp-27c8.2 |
| sf3b52 | 1.7E-17 | 0.5502705 | 0.179 | 0.099 | 5.53E-13 | Chondrogenic Fibroblasts | sf3b5 |
| arl8a | 1.8E-17 | 0.6235718 | 0.149 | 0.078 | 5.9E-13 | Chondrogenic Fibroblasts | arl8a |
| cib11 | 2E-17 | 0.4809195 | 0.207 | 0.119 | 6.57E-13 | Chondrogenic Fibroblasts | cib1 |
| anxa11b | 2E-17 | 0.2906429 | 0.533 | 0.384 | 6.58E-13 | Chondrogenic Fibroblasts | anxa11b |
| gorasp23 | 2.1E-17 | 0.5993681 | 0.191 | 0.109 | 6.75E-13 | Chondrogenic Fibroblasts | gorasp2 |
| zgc:114174 | 2.1E-17 | 0.5369173 | 0.175 | 0.097 | 6.85E-13 | Chondrogenic Fibroblasts | zgc:114174 |
| cox6b22 | 2.3E-17 | 0.2208942 | 0.923 | 0.812 | 7.41E-13 | Chondrogenic Fibroblasts | cox6b2 |
| rhogb2 | 2.3E-17 | 0.6122014 | 0.136 | 0.069 | 7.62E-13 | Chondrogenic Fibroblasts | rhogb |
| furina3 | 2.4E-17 | 0.4880481 | 0.251 | 0.154 | 7.68E-13 | Chondrogenic Fibroblasts | furina |
| stat33 | 2.4E-17 | 0.4676554 | 0.245 | 0.148 | 7.8E-13 | Chondrogenic Fibroblasts | stat3 |
| si:dkey-9i23.152 | 2.4E-17 | 0.3542953 | 0.25 | 0.15 | 7.86E-13 | Chondrogenic Fibroblasts | si:dkey-9i23.15 |
| hspa4b1 | 2.5E-17 | 0.3815377 | 0.236 | 0.141 | 8.2E-13 | Chondrogenic Fibroblasts | hspa4b |
| mcfd22 | 2.5E-17 | 0.5569729 | 0.213 | 0.125 | 8.29E-13 | Chondrogenic Fibroblasts | mcfd2 |
| uqcrc11 | 2.6E-17 | 0.3923354 | 0.284 | 0.179 | 8.42E-13 | Chondrogenic Fibroblasts | uqcrc1 |
| rnf2 | 2.6E-17 | 0.5360037 | 0.202 | 0.117 | 8.51E-13 | Chondrogenic Fibroblasts | rnf2 |
| rras1 | 2.7E-17 | 0.4623017 | 0.259 | 0.159 | 8.64E-13 | Chondrogenic Fibroblasts | rras |
| nek72 | 3E-17 | 0.40434 | 0.204 | 0.118 | 9.86E-13 | Chondrogenic Fibroblasts | nek7 |
| copa2 | 3.1E-17 | 0.3010608 | 0.299 | 0.187 | 1.01E-12 | Chondrogenic Fibroblasts | copa |
| sec61a1l2 | 3.3E-17 | 0.51054 | 0.23 | 0.138 | 1.06E-12 | Chondrogenic Fibroblasts | sec61a1l |
| sfrp54 | 3.3E-17 | 0.84179 | 0.308 | 0.208 | 1.06E-12 | Chondrogenic Fibroblasts | sfrp5 |
| adamts12 | 3.4E-17 | 0.6790472 | 0.111 | 0.053 | 1.1E-12 | Chondrogenic Fibroblasts | adamts12 |
| baz1b1 | 3.6E-17 | 0.2817584 | 0.28 | 0.172 | 1.17E-12 | Chondrogenic Fibroblasts | baz1b |
| cacybp1 | 3.8E-17 | 0.4647725 | 0.247 | 0.15 | 1.22E-12 | Chondrogenic Fibroblasts | cacybp |

| | | | | | | | |
|---|---|---|---|---|---|---|---|
| srp93 | 3.8E-17 | 0.4434515 | 0.189 | 0.106 | 1.25E-12 | Chondrogenic Fibroblasts | srp9 |
| ndufs8a2 | 4E-17 | 0.5236811 | 0.166 | 0.091 | 1.3E-12 | Chondrogenic Fibroblasts | ndufs8a |
| aktip1 | 4.9E-17 | 0.6128013 | 0.173 | 0.096 | 1.61E-12 | Chondrogenic Fibroblasts | aktip |
| ppp3r1a2 | 5E-17 | 0.4612245 | 0.233 | 0.14 | 1.63E-12 | Chondrogenic Fibroblasts | ppp3r1a |
| gpc1b3 | 5.2E-17 | 0.2483429 | 0.551 | 0.398 | 1.7E-12 | Chondrogenic Fibroblasts | gpc1b |
| ube2d1b1 | 5.4E-17 | 0.440982 | 0.23 | 0.137 | 1.76E-12 | Chondrogenic Fibroblasts | ube2d1b |
| plce1 | 5.9E-17 | 0.6675124 | 0.101 | 0.046 | 1.92E-12 | Chondrogenic Fibroblasts | plce1 |
| pdcl2 | 6E-17 | 0.3683851 | 0.31 | 0.198 | 1.94E-12 | Chondrogenic Fibroblasts | pdcl |
| bckdhbl | 6.3E-17 | 0.4905684 | 0.274 | 0.173 | 2.04E-12 | Chondrogenic Fibroblasts | bckdhbl |
| gpm6bb3 | 6.4E-17 | 0.2719038 | 0.477 | 0.332 | 2.09E-12 | Chondrogenic Fibroblasts | gpm6bb |
| mt-cyb1 | 6.5E-17 | 0.2030554 | 0.997 | 0.974 | 2.13E-12 | Chondrogenic Fibroblasts | mt-cyb |
| capza1b1 | 7.1E-17 | 0.3018216 | 0.507 | 0.364 | 2.32E-12 | Chondrogenic Fibroblasts | capza1b |
| trip113 | 7.7E-17 | 0.6375606 | 0.163 | 0.09 | 2.49E-12 | Chondrogenic Fibroblasts | trip11 |
| psmc31 | 7.7E-17 | 0.5825288 | 0.172 | 0.096 | 2.51E-12 | Chondrogenic Fibroblasts | psmc3 |
| micos132 | 8.1E-17 | 0.466652 | 0.207 | 0.121 | 2.65E-12 | Chondrogenic Fibroblasts | micos13 |
| nipa22 | 8.6E-17 | 0.5996406 | 0.143 | 0.075 | 2.78E-12 | Chondrogenic Fibroblasts | nipa2 |
| atp5fa11 | 8.7E-17 | 0.2372489 | 0.617 | 0.458 | 2.83E-12 | Chondrogenic Fibroblasts | atp5fa1 |
| vps281 | 9.1E-17 | 0.5365196 | 0.195 | 0.113 | 2.97E-12 | Chondrogenic Fibroblasts | vps28 |
| brk11 | 9.2E-17 | 0.363394 | 0.313 | 0.202 | 3E-12 | Chondrogenic Fibroblasts | brk1 |
| vapal3 | 9.8E-17 | 0.4462401 | 0.261 | 0.163 | 3.2E-12 | Chondrogenic Fibroblasts | vapal |
| ndufs32 | 9.9E-17 | 0.4614162 | 0.217 | 0.129 | 3.2E-12 | Chondrogenic Fibroblasts | ndufs3 |
| ndufb52 | 1E-16 | 0.3948572 | 0.291 | 0.186 | 3.35E-12 | Chondrogenic Fibroblasts | ndufb5 |
| zgc:919101 | 1.2E-16 | 0.3330627 | 0.372 | 0.25 | 3.95E-12 | Chondrogenic Fibroblasts | zgc:91910 |
| sept7b2 | 1.2E-16 | 0.4079194 | 0.304 | 0.196 | 4E-12 | Chondrogenic Fibroblasts | sept7b |
| cap11 | 1.3E-16 | 0.7262359 | 0.103 | 0.049 | 4.25E-12 | Chondrogenic Fibroblasts | cap1 |
| ccdc1673 | 1.3E-16 | 0.6269858 | 0.127 | 0.064 | 4.36E-12 | Chondrogenic Fibroblasts | ccdc167 |
| vps251 | 1.5E-16 | 0.4683685 | 0.17 | 0.094 | 4.95E-12 | Chondrogenic Fibroblasts | vps25 |
| alas13 | 1.6E-16 | 0.4690387 | 0.186 | 0.105 | 5.1E-12 | Chondrogenic Fibroblasts | alas1 |
| blcap1 | 1.6E-16 | 0.45428 | 0.234 | 0.142 | 5.3E-12 | Chondrogenic Fibroblasts | blcap |
| immt2 | 1.7E-16 | 0.5375427 | 0.161 | 0.088 | 5.65E-12 | Chondrogenic Fibroblasts | immt |
| tspan72 | 1.8E-16 | 0.3769233 | 0.325 | 0.213 | 5.77E-12 | Chondrogenic Fibroblasts | tspan7 |
| CU929259.11 | 1.8E-16 | 0.2590254 | 0.455 | 0.315 | 5.81E-12 | Chondrogenic Fibroblasts | CU929259.1 |
| prkcaa1 | 1.8E-16 | 0.4413475 | 0.184 | 0.104 | 5.87E-12 | Chondrogenic Fibroblasts | prkcaa |
| tpm4b3 | 1.8E-16 | 0.5557868 | 0.152 | 0.083 | 5.91E-12 | Chondrogenic Fibroblasts | tpm4b |
| mapk14b2 | 2E-16 | 0.4486236 | 0.158 | 0.085 | 6.39E-12 | Chondrogenic Fibroblasts | mapk14b |
| btg33 | 2E-16 | 0.4994976 | 0.21 | 0.125 | 6.42E-12 | Chondrogenic Fibroblasts | btg3 |
| porb4 | 2E-16 | 0.3403129 | 0.189 | 0.107 | 6.66E-12 | Chondrogenic Fibroblasts | porb |
| tbrg12 | 2.2E-16 | 0.7815694 | 0.102 | 0.048 | 7.22E-12 | Chondrogenic Fibroblasts | tbrg1 |
| cnn3b1 | 2.6E-16 | 0.6275138 | 0.111 | 0.054 | 8.45E-12 | Chondrogenic Fibroblasts | cnn3b |
| zmp:00000007291 | 2.7E-16 | 0.8577979 | 0.102 | 0.049 | 8.64E-12 | Chondrogenic Fibroblasts | zmp:0000000729 |
| bcl2l103 | 2.8E-16 | 0.4937555 | 0.278 | 0.178 | 9.05E-12 | Chondrogenic Fibroblasts | bcl2l10 |
| snx61 | 3.2E-16 | 0.4568016 | 0.199 | 0.115 | 1.05E-11 | Chondrogenic Fibroblasts | snx6 |
| naa202 | 3.6E-16 | 0.4585625 | 0.24 | 0.148 | 1.18E-11 | Chondrogenic Fibroblasts | naa20 |
| mfap11 | 3.9E-16 | 0.2422199 | 0.23 | 0.136 | 1.28E-11 | Chondrogenic Fibroblasts | mfap1 |
| irf2a1 | 4.5E-16 | 0.6303582 | 0.153 | 0.083 | 1.47E-11 | Chondrogenic Fibroblasts | irf2a |
| ptp4a12 | 4.6E-16 | 0.3320174 | 0.347 | 0.231 | 1.49E-11 | Chondrogenic Fibroblasts | ptp4a1 |
| ik | 4.9E-16 | 0.3396978 | 0.273 | 0.172 | 1.6E-11 | Chondrogenic Fibroblasts | ik |
| suds3 | 5E-16 | 0.429332 | 0.179 | 0.101 | 1.64E-11 | Chondrogenic Fibroblasts | suds3 |

| | | | | | | | |
|---|---|---|---|---|---|---|---|
| rnd1b5 | 5.6E-16 | 0.6133403 | 0.179 | 0.102 | 1.84E-11 | Chondrogenic Fibroblasts | rnd1b |
| sgta1 | 5.7E-16 | 0.5165186 | 0.222 | 0.135 | 1.84E-11 | Chondrogenic Fibroblasts | sgta |
| snw11 | 6.3E-16 | 0.3067041 | 0.301 | 0.192 | 2.06E-11 | Chondrogenic Fibroblasts | snw1 |
| fam20b2 | 6.9E-16 | 0.7612306 | 0.125 | 0.064 | 2.25E-11 | Chondrogenic Fibroblasts | fam20b |
| cybrd13 | 7.4E-16 | 0.562225 | 0.15 | 0.082 | 2.41E-11 | Chondrogenic Fibroblasts | cybrd1 |
| vps37c | 7.4E-16 | 0.6461894 | 0.126 | 0.065 | 2.42E-11 | Chondrogenic Fibroblasts | vps37c |
| qars11 | 8.4E-16 | 0.5869381 | 0.149 | 0.081 | 2.73E-11 | Chondrogenic Fibroblasts | qars1 |
| nufip22 | 9.3E-16 | 0.3418624 | 0.238 | 0.146 | 3.03E-11 | Chondrogenic Fibroblasts | nufip2 |
| ndufab1b3 | 9.7E-16 | 0.3675287 | 0.307 | 0.201 | 3.15E-11 | Chondrogenic Fibroblasts | ndufab1b |
| si:dkey-42i9.61 | 9.8E-16 | 0.4112377 | 0.296 | 0.193 | 3.19E-11 | Chondrogenic Fibroblasts | si:dkey-42i9.6 |
| mapre1b1 | 1E-15 | 0.3947083 | 0.205 | 0.121 | 3.24E-11 | Chondrogenic Fibroblasts | mapre1b |
| snrpg2 | 1E-15 | 0.2503073 | 0.285 | 0.181 | 3.32E-11 | Chondrogenic Fibroblasts | snrpg |
| nav1b1 | 1.1E-15 | 0.5319965 | 0.134 | 0.07 | 3.46E-11 | Chondrogenic Fibroblasts | nav1b |
| gskip4 | 1.1E-15 | 0.4124198 | 0.254 | 0.16 | 3.48E-11 | Chondrogenic Fibroblasts | gskip |
| riok31 | 1.1E-15 | 0.863821 | 0.101 | 0.049 | 3.49E-11 | Chondrogenic Fibroblasts | riok3 |
| sap30l1 | 1.1E-15 | 0.3201908 | 0.326 | 0.214 | 3.58E-11 | Chondrogenic Fibroblasts | sap30l |
| myl12.22 | 1.1E-15 | 0.436885 | 0.261 | 0.166 | 3.65E-11 | Chondrogenic Fibroblasts | myl12.2 |
| arpp19b1 | 1.1E-15 | 0.5083196 | 0.186 | 0.109 | 3.73E-11 | Chondrogenic Fibroblasts | arpp19b |
| arcn1a2 | 1.2E-15 | 0.5677076 | 0.17 | 0.097 | 3.75E-11 | Chondrogenic Fibroblasts | arcn1a |
| atp6v1e1a2 | 1.2E-15 | 0.5690934 | 0.178 | 0.103 | 3.75E-11 | Chondrogenic Fibroblasts | atp6v1e1a |
| hnrnpk1 | 1.2E-15 | 0.3060559 | 0.242 | 0.148 | 4.06E-11 | Chondrogenic Fibroblasts | hnrnpk |
| eif1b1 | 1.5E-15 | 0.2678641 | 0.318 | 0.207 | 4.95E-11 | Chondrogenic Fibroblasts | eif1b |
| mrpl331 | 1.5E-15 | 0.4111491 | 0.244 | 0.152 | 4.95E-11 | Chondrogenic Fibroblasts | mrpl33 |
| mrpl471 | 1.5E-15 | 0.4748936 | 0.125 | 0.064 | 5.03E-11 | Chondrogenic Fibroblasts | mrpl47 |
| TMEM192 | 1.6E-15 | 0.6755332 | 0.131 | 0.069 | 5.29E-11 | Chondrogenic Fibroblasts | TMEM19 |
| ppme11 | 1.7E-15 | 0.5410737 | 0.179 | 0.103 | 5.41E-11 | Chondrogenic Fibroblasts | ppme1 |
| si:dkey-6n21.121 | 1.7E-15 | 0.3798999 | 0.214 | 0.128 | 5.52E-11 | Chondrogenic Fibroblasts | si:dkey-6n21.12 |
| tm9sf32 | 1.8E-15 | 0.3718952 | 0.27 | 0.171 | 5.74E-11 | Chondrogenic Fibroblasts | tm9sf3 |
| herpud2 | 1.8E-15 | 0.5250598 | 0.137 | 0.073 | 5.93E-11 | Chondrogenic Fibroblasts | herpud2 |
| tspan2a | 1.9E-15 | 0.6140059 | 0.109 | 0.054 | 6.26E-11 | Chondrogenic Fibroblasts | tspan2a |
| fhl1b3 | 1.9E-15 | 0.6822337 | 0.164 | 0.093 | 6.32E-11 | Chondrogenic Fibroblasts | fhl1b |
| slc43a1a2 | 2.1E-15 | 0.5461176 | 0.149 | 0.081 | 6.97E-11 | Chondrogenic Fibroblasts | slc43a1a |
| gadd45bb5 | 2.2E-15 | 0.4850472 | 0.324 | 0.218 | 6.99E-11 | Chondrogenic Fibroblasts | gadd45bb |
| cobll1b2 | 2.2E-15 | 0.3677151 | 0.123 | 0.062 | 7.14E-11 | Chondrogenic Fibroblasts | cobll1b |
| cttn1 | 2.3E-15 | 0.5280908 | 0.131 | 0.069 | 7.33E-11 | Chondrogenic Fibroblasts | cttn |
| ndufb21 | 2.3E-15 | 0.3396841 | 0.366 | 0.253 | 7.52E-11 | Chondrogenic Fibroblasts | ndufb2 |
| ctsc1 | 2.4E-15 | 0.5664849 | 0.128 | 0.067 | 7.73E-11 | Chondrogenic Fibroblasts | ctsc |
| bcar12 | 2.4E-15 | 0.5135396 | 0.163 | 0.092 | 7.76E-11 | Chondrogenic Fibroblasts | bcar1 |
| pora2 | 2.4E-15 | 0.4042927 | 0.188 | 0.109 | 7.87E-11 | Chondrogenic Fibroblasts | pora |
| m6pr1 | 2.5E-15 | 0.4198475 | 0.205 | 0.122 | 8E-11 | Chondrogenic Fibroblasts | m6pr |
| chmp71 | 2.5E-15 | 0.7527081 | 0.138 | 0.075 | 8.07E-11 | Chondrogenic Fibroblasts | chmp7 |
| megf6a2 | 2.5E-15 | 0.6252208 | 0.114 | 0.057 | 8.12E-11 | Chondrogenic Fibroblasts | megf6a |
| tardbp1 | 2.6E-15 | 0.3461279 | 0.253 | 0.158 | 8.48E-11 | Chondrogenic Fibroblasts | tardbp |
| hp1bp3 | 2.6E-15 | 0.302234 | 0.348 | 0.232 | 8.53E-11 | Chondrogenic Fibroblasts | hp1bp3 |
| rdh10b1 | 2.6E-15 | 0.6619834 | 0.137 | 0.074 | 8.59E-11 | Chondrogenic Fibroblasts | rdh10b |
| nck1b3 | 2.7E-15 | 0.4930789 | 0.169 | 0.096 | 8.63E-11 | Chondrogenic Fibroblasts | nck1b |
| oaz1a4 | 2.7E-15 | 0.2179105 | 0.606 | 0.449 | 8.86E-11 | Chondrogenic Fibroblasts | oaz1a |
| akt2l1 | 2.8E-15 | 0.8647294 | 0.101 | 0.049 | 8.98E-11 | Chondrogenic Fibroblasts | akt2l |

| | | | | | | | |
|---|---|---|---|---|---|---|---|
| ndufa52 | 2.8E-15 | 0.3426123 | 0.314 | 0.207 | 9.12E-11 | Chondrogenic Fibroblasts | ndufa5 |
| lzic | 2.8E-15 | 0.519076 | 0.108 | 0.053 | 9.12E-11 | Chondrogenic Fibroblasts | lzic |
| rad21a1 | 2.9E-15 | 0.3697295 | 0.303 | 0.199 | 9.57E-11 | Chondrogenic Fibroblasts | rad21a |
| pole41 | 3.1E-15 | 0.6485201 | 0.115 | 0.058 | 1.02E-10 | Chondrogenic Fibroblasts | pole4 |
| psmc61 | 3.3E-15 | 0.4208275 | 0.219 | 0.133 | 1.09E-10 | Chondrogenic Fibroblasts | psmc6 |
| atg4a1 | 3.3E-15 | 0.7050015 | 0.119 | 0.061 | 1.09E-10 | Chondrogenic Fibroblasts | atg4a |
| cdk9 | 3.5E-15 | 0.6847726 | 0.14 | 0.076 | 1.14E-10 | Chondrogenic Fibroblasts | cdk9 |
| si:ch211-11k18.41 | 3.7E-15 | 0.6547148 | 0.114 | 0.058 | 1.21E-10 | Chondrogenic Fibroblasts | si:ch211-11k18.4 |
| ndufs22 | 3.8E-15 | 0.4500281 | 0.19 | 0.112 | 1.24E-10 | Chondrogenic Fibroblasts | ndufs2 |
| rasgef1bb4 | 3.9E-15 | 0.4329193 | 0.244 | 0.153 | 1.27E-10 | Chondrogenic Fibroblasts | rasgef1bb |
| tuba1b5 | 4E-15 | 0.2199801 | 0.442 | 0.306 | 1.31E-10 | Chondrogenic Fibroblasts | tuba1b |
| pfdn21 | 4.3E-15 | 0.5901466 | 0.131 | 0.07 | 1.4E-10 | Chondrogenic Fibroblasts | pfdn2 |
| tars11 | 4.5E-15 | 0.5387849 | 0.136 | 0.072 | 1.45E-10 | Chondrogenic Fibroblasts | tars1 |
| stt3b2 | 4.5E-15 | 0.3741521 | 0.184 | 0.107 | 1.47E-10 | Chondrogenic Fibroblasts | stt3b |
| ubtd22 | 5.2E-15 | 0.4623967 | 0.177 | 0.102 | 1.69E-10 | Chondrogenic Fibroblasts | ubtd2 |
| tbca1 | 5.5E-15 | 0.2733604 | 0.444 | 0.313 | 1.8E-10 | Chondrogenic Fibroblasts | tbca |
| baz2ba3 | 5.8E-15 | 0.3304501 | 0.265 | 0.168 | 1.87E-10 | Chondrogenic Fibroblasts | baz2ba |
| syncrip1 | 5.9E-15 | 0.2175486 | 0.654 | 0.492 | 1.92E-10 | Chondrogenic Fibroblasts | syncrip |
| bnip3lb1 | 5.9E-15 | 0.5449232 | 0.259 | 0.167 | 1.93E-10 | Chondrogenic Fibroblasts | bnip3lb |
| kmt5aa3 | 6.1E-15 | 0.4190629 | 0.192 | 0.113 | 1.98E-10 | Chondrogenic Fibroblasts | kmt5aa |
| limch1b4 | 6.5E-15 | 0.6856097 | 0.158 | 0.09 | 2.1E-10 | Chondrogenic Fibroblasts | limch1b |
| znf395b2 | 6.9E-15 | 0.3945911 | 0.223 | 0.137 | 2.25E-10 | Chondrogenic Fibroblasts | znf395b |
| cbx51 | 8.1E-15 | 0.293321 | 0.335 | 0.224 | 2.64E-10 | Chondrogenic Fibroblasts | cbx5 |
| cmtm62 | 8.3E-15 | 0.3101461 | 0.228 | 0.139 | 2.69E-10 | Chondrogenic Fibroblasts | cmtm6 |
| copb11 | 9.3E-15 | 0.3421538 | 0.235 | 0.145 | 3.03E-10 | Chondrogenic Fibroblasts | copb1 |
| cltca2 | 9.6E-15 | 0.2263848 | 0.302 | 0.195 | 3.13E-10 | Chondrogenic Fibroblasts | cltca |
| cmc13 | 9.8E-15 | 0.496709 | 0.167 | 0.096 | 3.19E-10 | Chondrogenic Fibroblasts | cmc1 |
| tp53bp2b1 | 1E-14 | 0.7354278 | 0.106 | 0.053 | 3.27E-10 | Chondrogenic Fibroblasts | tp53bp2b |
| tspan363 | 1E-14 | 0.6935166 | 0.245 | 0.16 | 3.4E-10 | Chondrogenic Fibroblasts | tspan36 |
| fuom | 1.1E-14 | 0.6188833 | 0.124 | 0.065 | 3.49E-10 | Chondrogenic Fibroblasts | fuom |
| rnf401 | 1.1E-14 | 0.4598639 | 0.192 | 0.114 | 3.6E-10 | Chondrogenic Fibroblasts | rnf40 |
| wasf21 | 1.2E-14 | 0.2941922 | 0.378 | 0.258 | 3.79E-10 | Chondrogenic Fibroblasts | wasf2 |
| uri12 | 1.2E-14 | 0.4486765 | 0.21 | 0.128 | 3.97E-10 | Chondrogenic Fibroblasts | uri1 |
| serpinh1b1 | 1.2E-14 | 0.2731494 | 0.715 | 0.579 | 3.98E-10 | Chondrogenic Fibroblasts | serpinh1b |
| mcts11 | 1.2E-14 | 0.3385025 | 0.265 | 0.169 | 4.02E-10 | Chondrogenic Fibroblasts | mcts1 |
| psmb51 | 1.4E-14 | 0.4449006 | 0.219 | 0.136 | 4.69E-10 | Chondrogenic Fibroblasts | psmb5 |
| gorab1 | 1.6E-14 | 0.6427796 | 0.129 | 0.069 | 5.1E-10 | Chondrogenic Fibroblasts | gorab |
| lpin21 | 1.6E-14 | 0.4512814 | 0.107 | 0.053 | 5.26E-10 | Chondrogenic Fibroblasts | lpin2 |
| si:dkey-17m8.13 | 1.6E-14 | 0.3847336 | 0.202 | 0.121 | 5.3E-10 | Chondrogenic Fibroblasts | si:dkey-17m8.1 |
| cd151l3 | 1.6E-14 | 0.6279001 | 0.134 | 0.073 | 5.34E-10 | Chondrogenic Fibroblasts | cd151l |
| adi12 | 1.7E-14 | 0.5108547 | 0.157 | 0.089 | 5.39E-10 | Chondrogenic Fibroblasts | adi1 |
| snd12 | 1.8E-14 | 0.2732279 | 0.351 | 0.238 | 5.8E-10 | Chondrogenic Fibroblasts | snd1 |
| ap3s22 | 1.8E-14 | 0.6967885 | 0.118 | 0.061 | 5.94E-10 | Chondrogenic Fibroblasts | ap3s2 |
| il4r.21 | 1.8E-14 | 0.6268537 | 0.116 | 0.06 | 6E-10 | Chondrogenic Fibroblasts | il4r.2 |
| sptlc2b2 | 1.9E-14 | 0.432061 | 0.196 | 0.118 | 6.13E-10 | Chondrogenic Fibroblasts | sptlc2b |
| lhfpl2a4 | 2E-14 | 0.3883309 | 0.181 | 0.106 | 6.42E-10 | Chondrogenic Fibroblasts | lhfpl2a |
| fam50a2 | 2E-14 | 0.3123365 | 0.253 | 0.16 | 6.43E-10 | Chondrogenic Fibroblasts | fam50a |
| abhd16a1 | 2E-14 | 0.5728471 | 0.128 | 0.068 | 6.58E-10 | Chondrogenic Fibroblasts | abhd16a |

| | | | | | | | |
|---|---|---|---|---|---|---|---|
| erlec12 | 2.1E-14 | 0.3777785 | 0.197 | 0.118 | 6.93E-10 | Chondrogenic Fibroblasts | erlec1 |
| pak2a2 | 2.2E-14 | 0.2728652 | 0.242 | 0.151 | 7E-10 | Chondrogenic Fibroblasts | pak2a |
| tax1bp32 | 2.2E-14 | 0.2706829 | 0.367 | 0.251 | 7.01E-10 | Chondrogenic Fibroblasts | tax1bp3 |
| polr2j1 | 2.2E-14 | 0.4102259 | 0.166 | 0.095 | 7.01E-10 | Chondrogenic Fibroblasts | polr2j |
| bod1 | 2.5E-14 | 0.5823145 | 0.113 | 0.058 | 8E-10 | Chondrogenic Fibroblasts | bod1 |
| erf1 | 2.5E-14 | 0.713053 | 0.126 | 0.068 | 8.05E-10 | Chondrogenic Fibroblasts | erf |
| tm9sf42 | 2.5E-14 | 0.4593852 | 0.136 | 0.074 | 8.25E-10 | Chondrogenic Fibroblasts | tm9sf4 |
| kdm5c2 | 2.7E-14 | 0.3385166 | 0.24 | 0.15 | 8.86E-10 | Chondrogenic Fibroblasts | kdm5c |
| usp51 | 3E-14 | 0.4189223 | 0.179 | 0.105 | 9.62E-10 | Chondrogenic Fibroblasts | usp5 |
| coq10b2 | 3.3E-14 | 0.3108578 | 0.261 | 0.167 | 1.06E-09 | Chondrogenic Fibroblasts | coq10b |
| si:dkeyp-23e4.34 | 3.4E-14 | 0.4096313 | 0.152 | 0.084 | 1.1E-09 | Chondrogenic Fibroblasts | si:dkeyp-23e4.3 |
| psmd31 | 3.4E-14 | 0.3115203 | 0.241 | 0.151 | 1.11E-09 | Chondrogenic Fibroblasts | psmd3 |
| ppp2r5eb1 | 3.8E-14 | 0.3494855 | 0.243 | 0.154 | 1.24E-09 | Chondrogenic Fibroblasts | ppp2r5eb |
| ndufs41 | 3.9E-14 | 0.3951749 | 0.214 | 0.131 | 1.28E-09 | Chondrogenic Fibroblasts | ndufs4 |
| ap2s11 | 4E-14 | 0.261446 | 0.267 | 0.171 | 1.3E-09 | Chondrogenic Fibroblasts | ap2s1 |
| gpx82 | 4E-14 | 0.4087863 | 0.265 | 0.173 | 1.31E-09 | Chondrogenic Fibroblasts | gpx8 |
| tp53inp11 | 4E-14 | 0.6602398 | 0.113 | 0.059 | 1.31E-09 | Chondrogenic Fibroblasts | tp53inp1 |
| mia31 | 4.4E-14 | 0.2970578 | 0.295 | 0.194 | 1.42E-09 | Chondrogenic Fibroblasts | mia3 |
| ilk1 | 4.5E-14 | 0.2210854 | 0.277 | 0.178 | 1.45E-09 | Chondrogenic Fibroblasts | ilk |
| C19H6orf622 | 4.8E-14 | 0.2214868 | 0.543 | 0.4 | 1.56E-09 | Chondrogenic Fibroblasts | C19H6orf62 |
| hdac11 | 5E-14 | 0.2416824 | 0.387 | 0.269 | 1.63E-09 | Chondrogenic Fibroblasts | hdac1 |
| efna5b2 | 5E-14 | 0.4970222 | 0.129 | 0.069 | 1.63E-09 | Chondrogenic Fibroblasts | efna5b |
| gnaia | 5.2E-14 | 0.3251614 | 0.233 | 0.146 | 1.7E-09 | Chondrogenic Fibroblasts | gnaia |
| zgc:1623391 | 5.8E-14 | 0.5796464 | 0.131 | 0.071 | 1.88E-09 | Chondrogenic Fibroblasts | zgc:162339 |
| slc30a73 | 6E-14 | 0.4945395 | 0.185 | 0.111 | 1.95E-09 | Chondrogenic Fibroblasts | slc30a7 |
| mical3a.12 | 6.1E-14 | 0.4283164 | 0.175 | 0.103 | 1.98E-09 | Chondrogenic Fibroblasts | mical3a.1 |
| mob41 | 6.4E-14 | 0.4295556 | 0.204 | 0.125 | 2.08E-09 | Chondrogenic Fibroblasts | mob4 |
| csnk1a1 | 7.3E-14 | 0.2300221 | 0.519 | 0.377 | 2.37E-09 | Chondrogenic Fibroblasts | csnk1a1 |
| phf5a1 | 7.8E-14 | 0.4783268 | 0.182 | 0.109 | 2.54E-09 | Chondrogenic Fibroblasts | phf5a |
| copg22 | 7.9E-14 | 0.3726069 | 0.234 | 0.148 | 2.57E-09 | Chondrogenic Fibroblasts | copg2 |
| rnf10 | 8.3E-14 | 0.2041556 | 0.476 | 0.338 | 2.69E-09 | Chondrogenic Fibroblasts | rnf10 |
| nbas2 | 8.8E-14 | 0.5763802 | 0.166 | 0.098 | 2.87E-09 | Chondrogenic Fibroblasts | nbas |
| igfbp5b4 | 9.3E-14 | 0.231409 | 0.71 | 0.568 | 3.03E-09 | Chondrogenic Fibroblasts | igfbp5b |
| lrba3 | 9.5E-14 | 0.2410392 | 0.271 | 0.175 | 3.09E-09 | Chondrogenic Fibroblasts | lrba |
| casc41 | 9.8E-14 | 0.5178591 | 0.15 | 0.086 | 3.17E-09 | Chondrogenic Fibroblasts | casc4 |
| dync1i2a1 | 1E-13 | 0.3291858 | 0.187 | 0.112 | 3.3E-09 | Chondrogenic Fibroblasts | dync1i2a |
| hnrnph1l1 | 1E-13 | 0.3379919 | 0.213 | 0.132 | 3.32E-09 | Chondrogenic Fibroblasts | hnrnph1l |
| si:dkey-112a7.43 | 1E-13 | 0.7857287 | 0.152 | 0.087 | 3.38E-09 | Chondrogenic Fibroblasts | si:dkey-112a7.4 |
| nt5e2 | 1.1E-13 | 0.4660593 | 0.119 | 0.063 | 3.58E-09 | Chondrogenic Fibroblasts | nt5e |
| txlna1 | 1.1E-13 | 0.3714683 | 0.173 | 0.101 | 3.66E-09 | Chondrogenic Fibroblasts | txlna |
| cnpy33 | 1.2E-13 | 0.5459294 | 0.152 | 0.087 | 3.96E-09 | Chondrogenic Fibroblasts | cnpy3 |
| nkx3-25 | 1.2E-13 | 0.4538494 | 0.289 | 0.194 | 4.05E-09 | Chondrogenic Fibroblasts | nkx3-2 |
| eif4a31 | 1.3E-13 | 0.3091311 | 0.278 | 0.183 | 4.12E-09 | Chondrogenic Fibroblasts | eif4a3 |
| pitpnc1b4 | 1.3E-13 | 0.4979048 | 0.143 | 0.081 | 4.25E-09 | Chondrogenic Fibroblasts | pitpnc1b |
| picalma2 | 1.4E-13 | 0.5848759 | 0.134 | 0.074 | 4.61E-09 | Chondrogenic Fibroblasts | picalma |
| tmem182 | 1.6E-13 | 0.6177942 | 0.151 | 0.087 | 5.09E-09 | Chondrogenic Fibroblasts | tmem18 |
| psmd11 | 1.6E-13 | 0.2949451 | 0.273 | 0.179 | 5.12E-09 | Chondrogenic Fibroblasts | psmd1 |
| rtca2 | 1.6E-13 | 0.3315129 | 0.282 | 0.186 | 5.21E-09 | Chondrogenic Fibroblasts | rtca |

| | | | | | | | |
|---|---|---|---|---|---|---|---|
| ndufv12 | 1.7E-13 | 0.3646011 | 0.21 | 0.13 | 5.48E-09 | Chondrogenic Fibroblasts | ndufv1 |
| psmd131 | 1.7E-13 | 0.3263704 | 0.223 | 0.141 | 5.62E-09 | Chondrogenic Fibroblasts | psmd13 |
| rn7sk5 | 1.7E-13 | 0.2670504 | 0.578 | 0.453 | 5.65E-09 | Chondrogenic Fibroblasts | rn7sk |
| slc7a10a5 | 1.8E-13 | 0.5264513 | 0.178 | 0.106 | 5.91E-09 | Chondrogenic Fibroblasts | slc7a10a |
| pcdh2g282 | 1.9E-13 | 0.2084281 | 0.381 | 0.26 | 6.16E-09 | Chondrogenic Fibroblasts | pcdh2g28 |
| abi1a1 | 1.9E-13 | 0.5531684 | 0.124 | 0.067 | 6.22E-09 | Chondrogenic Fibroblasts | abi1a |
| zgc:114200 | 2E-13 | 0.4248954 | 0.146 | 0.082 | 6.45E-09 | Chondrogenic Fibroblasts | zgc:114200 |
| lsm12b1 | 2E-13 | 0.2124681 | 0.465 | 0.332 | 6.58E-09 | Chondrogenic Fibroblasts | lsm12b |
| cux1b | 2.2E-13 | 0.4268837 | 0.165 | 0.097 | 7.05E-09 | Chondrogenic Fibroblasts | cux1b |
| chchd6a | 2.2E-13 | 0.4378016 | 0.138 | 0.077 | 7.07E-09 | Chondrogenic Fibroblasts | chchd6a |
| aup11 | 2.5E-13 | 0.4501502 | 0.151 | 0.087 | 8E-09 | Chondrogenic Fibroblasts | aup1 |
| bet1l1 | 2.5E-13 | 0.5889608 | 0.101 | 0.052 | 8.16E-09 | Chondrogenic Fibroblasts | bet1l |
| ssrp1b | 2.5E-13 | 0.4165591 | 0.146 | 0.082 | 8.29E-09 | Chondrogenic Fibroblasts | ssrp1b |
| zfand3 | 2.8E-13 | 0.28921 | 0.284 | 0.187 | 9.18E-09 | Chondrogenic Fibroblasts | zfand3 |
| ZNF512B | 2.9E-13 | 0.6215686 | 0.102 | 0.052 | 9.55E-09 | Chondrogenic Fibroblasts | ZNF512B |
| arrdc3a4 | 3E-13 | 0.4903622 | 0.25 | 0.163 | 9.61E-09 | Chondrogenic Fibroblasts | arrdc3a |
| dnajc71 | 3E-13 | 0.2370439 | 0.295 | 0.196 | 9.8E-09 | Chondrogenic Fibroblasts | dnajc7 |
| cyfip1 | 3E-13 | 0.2571435 | 0.308 | 0.208 | 9.81E-09 | Chondrogenic Fibroblasts | cyfip1 |
| elf2b1 | 3E-13 | 0.517276 | 0.182 | 0.111 | 9.83E-09 | Chondrogenic Fibroblasts | elf2b |
| sec22ba3 | 3.2E-13 | 0.4278325 | 0.19 | 0.116 | 1.06E-08 | Chondrogenic Fibroblasts | sec22ba |
| kif3a | 3.3E-13 | 0.3108352 | 0.115 | 0.06 | 1.08E-08 | Chondrogenic Fibroblasts | kif3a |
| CABZ01115113.11 | 3.5E-13 | 0.3887696 | 0.185 | 0.112 | 1.13E-08 | Chondrogenic Fibroblasts | CABZ01115113.1 |
| emc81 | 3.6E-13 | 0.8076369 | 0.101 | 0.052 | 1.16E-08 | Chondrogenic Fibroblasts | emc8 |
| eloa1 | 3.7E-13 | 0.5814345 | 0.132 | 0.073 | 1.21E-08 | Chondrogenic Fibroblasts | eloa |
| zgc:1008294 | 3.9E-13 | 0.4944146 | 0.191 | 0.118 | 1.26E-08 | Chondrogenic Fibroblasts | zgc:100829 |
| selenou1a1 | 3.9E-13 | 0.4650581 | 0.124 | 0.067 | 1.26E-08 | Chondrogenic Fibroblasts | selenou1a |
| lsm62 | 4E-13 | 0.292134 | 0.259 | 0.171 | 1.31E-08 | Chondrogenic Fibroblasts | lsm6 |
| kank22 | 4.1E-13 | 0.31599 | 0.31 | 0.21 | 1.34E-08 | Chondrogenic Fibroblasts | kank2 |
| ap3d11 | 4.3E-13 | 0.3294095 | 0.2 | 0.123 | 1.41E-08 | Chondrogenic Fibroblasts | ap3d1 |
| bhlhe413 | 4.4E-13 | 0.4082013 | 0.205 | 0.128 | 1.44E-08 | Chondrogenic Fibroblasts | bhlhe41 |
| psma6b1 | 4.4E-13 | 0.3192937 | 0.236 | 0.151 | 1.45E-08 | Chondrogenic Fibroblasts | psma6b |
| C20H12orf571 | 4.7E-13 | 0.4547702 | 0.155 | 0.09 | 1.53E-08 | Chondrogenic Fibroblasts | C20H12orf57 |
| bsdc12 | 4.7E-13 | 0.4000935 | 0.16 | 0.093 | 1.54E-08 | Chondrogenic Fibroblasts | bsdc1 |
| sec622 | 4.8E-13 | 0.3245724 | 0.338 | 0.235 | 1.55E-08 | Chondrogenic Fibroblasts | sec62 |
| taf12 | 4.9E-13 | 0.3586203 | 0.301 | 0.206 | 1.58E-08 | Chondrogenic Fibroblasts | taf12 |
| szrd11 | 4.9E-13 | 0.2625942 | 0.247 | 0.158 | 1.61E-08 | Chondrogenic Fibroblasts | szrd1 |
| tjp1b1 | 5.4E-13 | 0.2671995 | 0.224 | 0.141 | 1.77E-08 | Chondrogenic Fibroblasts | tjp1b |
| cct43 | 5.4E-13 | 0.2238762 | 0.433 | 0.31 | 1.77E-08 | Chondrogenic Fibroblasts | cct4 |
| wdr83os1 | 5.5E-13 | 0.259047 | 0.276 | 0.182 | 1.79E-08 | Chondrogenic Fibroblasts | wdr83os |
| dpy301 | 5.5E-13 | 0.3471112 | 0.25 | 0.163 | 1.8E-08 | Chondrogenic Fibroblasts | dpy30 |
| plekhb22 | 5.8E-13 | 0.3950017 | 0.17 | 0.101 | 1.9E-08 | Chondrogenic Fibroblasts | plekhb2 |
| emc61 | 6.1E-13 | 0.343744 | 0.171 | 0.102 | 2E-08 | Chondrogenic Fibroblasts | emc6 |
| gfpt12 | 6.1E-13 | 0.5003148 | 0.109 | 0.057 | 2E-08 | Chondrogenic Fibroblasts | gfpt1 |
| cbx1b2 | 6.3E-13 | 0.3745388 | 0.177 | 0.106 | 2.06E-08 | Chondrogenic Fibroblasts | cbx1b |
| gclm1 | 6.5E-13 | 0.8229131 | 0.117 | 0.063 | 2.11E-08 | Chondrogenic Fibroblasts | gclm |
| ppp2cb1 | 6.5E-13 | 0.2161588 | 0.409 | 0.29 | 2.12E-08 | Chondrogenic Fibroblasts | ppp2cb |
| ttpal1 | 6.7E-13 | 0.6094797 | 0.104 | 0.054 | 2.17E-08 | Chondrogenic Fibroblasts | ttpal |
| hspa53 | 7.5E-13 | 0.2853563 | 0.607 | 0.49 | 2.43E-08 | Chondrogenic Fibroblasts | hspa5 |

| | | | | | | | |
|---|---|---|---|---|---|---|---|
| rerea2 | 8E-13 | 0.2686109 | 0.265 | 0.173 | 2.59E-08 | Chondrogenic Fibroblasts | rerea |
| ap1ar1 | 8.1E-13 | 0.5333431 | 0.143 | 0.082 | 2.65E-08 | Chondrogenic Fibroblasts | ap1ar |
| idh3g1 | 8.4E-13 | 0.6331867 | 0.112 | 0.06 | 2.73E-08 | Chondrogenic Fibroblasts | idh3g |
| adrm12 | 8.9E-13 | 0.4132581 | 0.153 | 0.089 | 2.88E-08 | Chondrogenic Fibroblasts | adrm1 |
| zgc:1949814 | 9.8E-13 | 0.5039162 | 0.141 | 0.08 | 3.18E-08 | Chondrogenic Fibroblasts | zgc:194981 |
| piezo13 | 9.9E-13 | 0.3867527 | 0.198 | 0.123 | 3.21E-08 | Chondrogenic Fibroblasts | piezo1 |
| scyl11 | 1E-12 | 0.4196056 | 0.137 | 0.077 | 3.27E-08 | Chondrogenic Fibroblasts | scyl1 |
| ormdl1 | 1.1E-12 | 0.6346347 | 0.1 | 0.051 | 3.43E-08 | Chondrogenic Fibroblasts | ormdl1 |
| rorab3 | 1.1E-12 | 0.2393118 | 0.202 | 0.124 | 3.48E-08 | Chondrogenic Fibroblasts | rorab |
| gosr11 | 1.1E-12 | 0.5609768 | 0.125 | 0.069 | 3.54E-08 | Chondrogenic Fibroblasts | gosr1 |
| gkap1 | 1.1E-12 | 0.3500257 | 0.18 | 0.109 | 3.69E-08 | Chondrogenic Fibroblasts | gkap1 |
| mtap1 | 1.1E-12 | 0.2737814 | 0.298 | 0.201 | 3.69E-08 | Chondrogenic Fibroblasts | mtap |
| man2b12 | 1.1E-12 | 0.4104238 | 0.162 | 0.096 | 3.71E-08 | Chondrogenic Fibroblasts | man2b1 |
| inppl1a2 | 1.1E-12 | 0.4740316 | 0.149 | 0.087 | 3.72E-08 | Chondrogenic Fibroblasts | inppl1a |
| puraa1 | 1.2E-12 | 0.2190879 | 0.366 | 0.251 | 3.84E-08 | Chondrogenic Fibroblasts | puraa |
| ppp1r7 | 1.2E-12 | 0.5107498 | 0.109 | 0.058 | 3.86E-08 | Chondrogenic Fibroblasts | ppp1r7 |
| mrpl342 | 1.2E-12 | 0.2972953 | 0.204 | 0.128 | 3.92E-08 | Chondrogenic Fibroblasts | mrpl34 |
| sh3bgrl2 | 1.2E-12 | 0.2706324 | 0.328 | 0.225 | 3.98E-08 | Chondrogenic Fibroblasts | sh3bgrl |
| jdp2a | 1.3E-12 | 0.415224 | 0.123 | 0.067 | 4.07E-08 | Chondrogenic Fibroblasts | jdp2a |
| ralbp11 | 1.3E-12 | 0.3500616 | 0.244 | 0.159 | 4.28E-08 | Chondrogenic Fibroblasts | ralbp1 |
| ddx3xa1 | 1.3E-12 | 0.2010488 | 0.448 | 0.321 | 4.36E-08 | Chondrogenic Fibroblasts | ddx3xa |
| phb1 | 1.3E-12 | 0.2663535 | 0.201 | 0.125 | 4.38E-08 | Chondrogenic Fibroblasts | phb |
| cyb5r3 | 1.4E-12 | 0.3188369 | 0.221 | 0.14 | 4.43E-08 | Chondrogenic Fibroblasts | cyb5r3 |
| sap181 | 1.5E-12 | 0.2873935 | 0.297 | 0.201 | 4.79E-08 | Chondrogenic Fibroblasts | sap18 |
| ndnf5 | 1.5E-12 | 0.3746381 | 0.171 | 0.103 | 4.8E-08 | Chondrogenic Fibroblasts | ndnf |
| h1-02 | 1.5E-12 | 0.323121 | 0.27 | 0.18 | 4.87E-08 | Chondrogenic Fibroblasts | h1-0 |
| tsg101b2 | 1.5E-12 | 0.4008411 | 0.153 | 0.089 | 4.87E-08 | Chondrogenic Fibroblasts | tsg101b |
| sirt2 | 1.6E-12 | 0.2995691 | 0.232 | 0.149 | 5.12E-08 | Chondrogenic Fibroblasts | sirt2 |
| bbip11 | 1.6E-12 | 0.3577416 | 0.199 | 0.125 | 5.19E-08 | Chondrogenic Fibroblasts | bbip1 |
| adam19a3 | 1.6E-12 | 0.435579 | 0.144 | 0.083 | 5.36E-08 | Chondrogenic Fibroblasts | adam19a |
| tmem230a | 1.7E-12 | 0.5151596 | 0.101 | 0.053 | 5.38E-08 | Chondrogenic Fibroblasts | tmem230a |
| atf61 | 1.7E-12 | 0.2728516 | 0.189 | 0.116 | 5.44E-08 | Chondrogenic Fibroblasts | atf6 |
| fam76b1 | 1.7E-12 | 0.3327967 | 0.146 | 0.084 | 5.54E-08 | Chondrogenic Fibroblasts | fam76b |
| wipf1b1 | 1.7E-12 | 0.4198339 | 0.172 | 0.104 | 5.54E-08 | Chondrogenic Fibroblasts | wipf1b |
| lamtor53 | 1.7E-12 | 0.4879433 | 0.149 | 0.087 | 5.63E-08 | Chondrogenic Fibroblasts | lamtor5 |
| idh3a1 | 1.7E-12 | 0.4786326 | 0.137 | 0.078 | 5.69E-08 | Chondrogenic Fibroblasts | idh3a |
| xrn1 | 1.8E-12 | 0.6115688 | 0.103 | 0.054 | 5.71E-08 | Chondrogenic Fibroblasts | xrn1 |
| ehd1a1 | 1.8E-12 | 0.4564744 | 0.118 | 0.064 | 5.73E-08 | Chondrogenic Fibroblasts | ehd1a |
| psma51 | 2E-12 | 0.2672403 | 0.287 | 0.192 | 6.38E-08 | Chondrogenic Fibroblasts | psma5 |
| mob3a1 | 2E-12 | 0.5049966 | 0.137 | 0.079 | 6.42E-08 | Chondrogenic Fibroblasts | mob3a |
| maff4 | 2E-12 | 0.3930124 | 0.316 | 0.222 | 6.48E-08 | Chondrogenic Fibroblasts | maff |
| si:dkey-253i9.41 | 2E-12 | 0.5971428 | 0.134 | 0.076 | 6.48E-08 | Chondrogenic Fibroblasts | si:dkey-253i9.4 |
| tollip1 | 2E-12 | 0.6529231 | 0.103 | 0.054 | 6.59E-08 | Chondrogenic Fibroblasts | tollip |
| map3k42 | 2.2E-12 | 0.4358312 | 0.143 | 0.083 | 7.06E-08 | Chondrogenic Fibroblasts | map3k4 |
| api51 | 2.2E-12 | 0.2884836 | 0.216 | 0.137 | 7.25E-08 | Chondrogenic Fibroblasts | api5 |
| rab3il1 | 2.2E-12 | 0.4329198 | 0.119 | 0.065 | 7.25E-08 | Chondrogenic Fibroblasts | rab3il1 |
| psmb21 | 2.2E-12 | 0.2371011 | 0.376 | 0.264 | 7.31E-08 | Chondrogenic Fibroblasts | psmb2 |
| msi2b1 | 2.3E-12 | 0.4301636 | 0.163 | 0.097 | 7.45E-08 | Chondrogenic Fibroblasts | msi2b |

| | | | | | | | |
|---|---|---|---|---|---|---|---|
| fam32a1 | 2.5E-12 | 0.3162635 | 0.298 | 0.204 | 7.99E-08 | Chondrogenic Fibroblasts | fam32a |
| f104 | 2.6E-12 | 0.2439558 | 0.524 | 0.397 | 8.49E-08 | Chondrogenic Fibroblasts | f10 |
| eaf11 | 2.6E-12 | 0.5509044 | 0.119 | 0.066 | 8.55E-08 | Chondrogenic Fibroblasts | eaf1 |
| nedd8l2 | 2.7E-12 | 0.6329686 | 0.121 | 0.068 | 8.7E-08 | Chondrogenic Fibroblasts | nedd8l |
| dpm11 | 2.7E-12 | 0.4542078 | 0.147 | 0.086 | 8.93E-08 | Chondrogenic Fibroblasts | dpm1 |
| fam177a11 | 2.8E-12 | 0.4532127 | 0.122 | 0.067 | 9.2E-08 | Chondrogenic Fibroblasts | fam177a1 |
| zmat52 | 3E-12 | 0.5012074 | 0.13 | 0.073 | 9.83E-08 | Chondrogenic Fibroblasts | zmat5 |
| sec631 | 3.1E-12 | 0.2242583 | 0.318 | 0.216 | 9.94E-08 | Chondrogenic Fibroblasts | sec63 |
| dlgap4a2 | 3.1E-12 | 0.4596994 | 0.155 | 0.092 | 1.01E-07 | Chondrogenic Fibroblasts | dlgap4a |
| supt4h11 | 3.2E-12 | 0.4136092 | 0.209 | 0.134 | 1.05E-07 | Chondrogenic Fibroblasts | supt4h1 |
| rtn4b3 | 3.3E-12 | 0.3131346 | 0.155 | 0.091 | 1.08E-07 | Chondrogenic Fibroblasts | rtn4b |
| med19b | 3.4E-12 | 0.4516654 | 0.12 | 0.066 | 1.12E-07 | Chondrogenic Fibroblasts | med19b |
| tpst1l | 3.5E-12 | 0.3212645 | 0.117 | 0.063 | 1.13E-07 | Chondrogenic Fibroblasts | tpst1l |
| sdhc1 | 3.5E-12 | 0.3360632 | 0.284 | 0.194 | 1.15E-07 | Chondrogenic Fibroblasts | sdhc |
| phc2b2 | 3.5E-12 | 0.7887609 | 0.131 | 0.075 | 1.15E-07 | Chondrogenic Fibroblasts | phc2b |
| spcs32 | 3.6E-12 | 0.2313183 | 0.379 | 0.268 | 1.17E-07 | Chondrogenic Fibroblasts | spcs3 |
| creb3l21 | 3.6E-12 | 0.2349757 | 0.564 | 0.423 | 1.18E-07 | Chondrogenic Fibroblasts | creb3l2 |
| ccdc472 | 3.7E-12 | 0.2631307 | 0.216 | 0.138 | 1.19E-07 | Chondrogenic Fibroblasts | ccdc47 |
| ppp1r21 | 3.8E-12 | 0.4021144 | 0.107 | 0.057 | 1.24E-07 | Chondrogenic Fibroblasts | ppp1r2 |
| psenen1 | 3.9E-12 | 0.4705045 | 0.197 | 0.124 | 1.26E-07 | Chondrogenic Fibroblasts | psenen |
| atp5mea1 | 4.2E-12 | 0.2434762 | 0.357 | 0.252 | 1.38E-07 | Chondrogenic Fibroblasts | atp5mea |
| mrpl271 | 4.4E-12 | 0.5402435 | 0.108 | 0.058 | 1.43E-07 | Chondrogenic Fibroblasts | mrpl27 |
| rab5if2 | 4.4E-12 | 0.4473199 | 0.19 | 0.12 | 1.44E-07 | Chondrogenic Fibroblasts | rab5if |
| degs12 | 4.7E-12 | 0.2346936 | 0.277 | 0.186 | 1.54E-07 | Chondrogenic Fibroblasts | degs1 |
| tomm401 | 5E-12 | 0.3551891 | 0.162 | 0.097 | 1.62E-07 | Chondrogenic Fibroblasts | tomm40 |
| tmem176l.12 | 5.1E-12 | 0.211417 | 0.86 | 0.739 | 1.65E-07 | Chondrogenic Fibroblasts | tmem176l.1 |
| nxn | 5.6E-12 | 0.5513455 | 0.119 | 0.066 | 1.81E-07 | Chondrogenic Fibroblasts | nxn |
| btf3l42 | 5.6E-12 | 0.3151648 | 0.301 | 0.206 | 1.82E-07 | Chondrogenic Fibroblasts | btf3l4 |
| rbm4.31 | 5.7E-12 | 0.2408045 | 0.316 | 0.216 | 1.85E-07 | Chondrogenic Fibroblasts | rbm4.3 |
| sh3gl1b2 | 5.8E-12 | 0.2179643 | 0.346 | 0.242 | 1.89E-07 | Chondrogenic Fibroblasts | sh3gl1b |
| baxb | 5.9E-12 | 0.6474402 | 0.1 | 0.052 | 1.92E-07 | Chondrogenic Fibroblasts | baxb |
| rbm6 | 6.4E-12 | 0.3685406 | 0.15 | 0.088 | 2.09E-07 | Chondrogenic Fibroblasts | rbm6 |
| acp12 | 6.7E-12 | 0.3706815 | 0.133 | 0.076 | 2.17E-07 | Chondrogenic Fibroblasts | acp1 |
| nap1l4b2 | 6.7E-12 | 0.3496492 | 0.183 | 0.114 | 2.18E-07 | Chondrogenic Fibroblasts | nap1l4b |
| ndufb62 | 6.9E-12 | 0.2676475 | 0.266 | 0.18 | 2.25E-07 | Chondrogenic Fibroblasts | ndufb6 |
| bet12 | 7.1E-12 | 0.4020658 | 0.162 | 0.098 | 2.31E-07 | Chondrogenic Fibroblasts | bet1 |
| mcrip11 | 7.1E-12 | 0.4295185 | 0.13 | 0.074 | 2.32E-07 | Chondrogenic Fibroblasts | mcrip1 |
| sdr16c5b1 | 7.4E-12 | 0.3414001 | 0.217 | 0.141 | 2.41E-07 | Chondrogenic Fibroblasts | sdr16c5b |
| nr1d13 | 7.4E-12 | 0.3705509 | 0.21 | 0.134 | 2.41E-07 | Chondrogenic Fibroblasts | nr1d1 |
| si:dkey-118j18.21 | 7.7E-12 | 0.3082058 | 0.154 | 0.091 | 2.52E-07 | Chondrogenic Fibroblasts | si:dkey-118j18.2 |
| kifbp1 | 7.8E-12 | 0.3502653 | 0.149 | 0.088 | 2.52E-07 | Chondrogenic Fibroblasts | kifbp |
| tmem176l.3a2 | 8E-12 | 0.2487864 | 0.229 | 0.148 | 2.59E-07 | Chondrogenic Fibroblasts | tmem176l.3a |
| rab11ba2 | 8.8E-12 | 0.2699508 | 0.27 | 0.182 | 2.86E-07 | Chondrogenic Fibroblasts | rab11ba |
| g3bp12 | 8.8E-12 | 0.2470082 | 0.253 | 0.168 | 2.87E-07 | Chondrogenic Fibroblasts | g3bp1 |
| rbmx21 | 8.9E-12 | 0.5084784 | 0.126 | 0.072 | 2.88E-07 | Chondrogenic Fibroblasts | rbmx2 |
| golga71 | 8.9E-12 | 0.3168936 | 0.259 | 0.174 | 2.9E-07 | Chondrogenic Fibroblasts | golga7 |
| arf6b1 | 9E-12 | 0.5094591 | 0.116 | 0.064 | 2.92E-07 | Chondrogenic Fibroblasts | arf6b |
| rimkla1 | 9.5E-12 | 0.5688375 | 0.119 | 0.067 | 3.1E-07 | Chondrogenic Fibroblasts | rimkla |

| | | | | | | | |
|---|---|---|---|---|---|---|---|
| snx141 | 9.7E-12 | 0.5840611 | 0.11 | 0.06 | 3.16E-07 | Chondrogenic Fibroblasts | snx14 |
| med282 | 9.8E-12 | 0.5429782 | 0.134 | 0.078 | 3.18E-07 | Chondrogenic Fibroblasts | med28 |
| tgif14 | 1E-11 | 0.3696638 | 0.295 | 0.204 | 3.25E-07 | Chondrogenic Fibroblasts | tgif1 |
| tcea23 | 1E-11 | 0.530007 | 0.133 | 0.077 | 3.27E-07 | Chondrogenic Fibroblasts | tcea2 |
| ccdc6b3 | 1E-11 | 0.6119263 | 0.136 | 0.079 | 3.3E-07 | Chondrogenic Fibroblasts | ccdc6b |
| zgc:865981 | 1E-11 | 0.2881143 | 0.242 | 0.16 | 3.32E-07 | Chondrogenic Fibroblasts | zgc:86598 |
| FBXO482 | 1.1E-11 | 0.6442772 | 0.109 | 0.06 | 3.47E-07 | Chondrogenic Fibroblasts | FBXO48 |
| nutf2l2 | 1.1E-11 | 0.3934792 | 0.164 | 0.1 | 3.64E-07 | Chondrogenic Fibroblasts | nutf2l |
| ssr11 | 1.2E-11 | 0.5477721 | 0.103 | 0.055 | 3.74E-07 | Chondrogenic Fibroblasts | ssr1 |
| sdf22 | 1.2E-11 | 0.5191384 | 0.107 | 0.058 | 3.8E-07 | Chondrogenic Fibroblasts | sdf2 |
| glod52 | 1.2E-11 | 0.5431399 | 0.106 | 0.057 | 3.85E-07 | Chondrogenic Fibroblasts | glod5 |
| atf4b5 | 1.2E-11 | 0.2191113 | 0.764 | 0.631 | 3.87E-07 | Chondrogenic Fibroblasts | atf4b |
| stk35l2 | 1.2E-11 | 0.3636683 | 0.196 | 0.124 | 4.06E-07 | Chondrogenic Fibroblasts | stk35l |
| abl1 | 1.3E-11 | 0.612829 | 0.11 | 0.06 | 4.12E-07 | Chondrogenic Fibroblasts | abl1 |
| etv5b4 | 1.3E-11 | 0.4244413 | 0.162 | 0.099 | 4.34E-07 | Chondrogenic Fibroblasts | etv5b |
| tex2611 | 1.3E-11 | 0.3794345 | 0.184 | 0.115 | 4.34E-07 | Chondrogenic Fibroblasts | tex261 |
| trappc2l1 | 1.4E-11 | 0.4859383 | 0.121 | 0.068 | 4.5E-07 | Chondrogenic Fibroblasts | trappc2l |
| C8H5orf511 | 1.4E-11 | 0.330072 | 0.185 | 0.116 | 4.5E-07 | Chondrogenic Fibroblasts | C8H5orf51 |
| s1pr12 | 1.4E-11 | 0.6270234 | 0.151 | 0.092 | 4.63E-07 | Chondrogenic Fibroblasts | s1pr1 |
| ralab1 | 1.4E-11 | 0.3331159 | 0.15 | 0.089 | 4.63E-07 | Chondrogenic Fibroblasts | ralab |
| usp41 | 1.5E-11 | 0.4285782 | 0.156 | 0.094 | 4.78E-07 | Chondrogenic Fibroblasts | usp4 |
| mxd42 | 1.5E-11 | 0.3606854 | 0.151 | 0.09 | 4.86E-07 | Chondrogenic Fibroblasts | mxd4 |
| czib1 | 1.5E-11 | 0.4630179 | 0.156 | 0.095 | 4.98E-07 | Chondrogenic Fibroblasts | czib |
| gng72 | 1.6E-11 | 0.2252738 | 0.248 | 0.164 | 5.25E-07 | Chondrogenic Fibroblasts | gng7 |
| ap1s21 | 1.6E-11 | 0.2544839 | 0.246 | 0.162 | 5.28E-07 | Chondrogenic Fibroblasts | ap1s2 |
| ccdc321 | 1.7E-11 | 0.5885284 | 0.11 | 0.061 | 5.58E-07 | Chondrogenic Fibroblasts | ccdc32 |
| hnrnpua1 | 1.8E-11 | 0.2455429 | 0.28 | 0.191 | 5.71E-07 | Chondrogenic Fibroblasts | hnrnpua |
| akirin12 | 1.9E-11 | 0.5309292 | 0.157 | 0.096 | 6.2E-07 | Chondrogenic Fibroblasts | akirin1 |
| mtx21 | 1.9E-11 | 0.6058183 | 0.1 | 0.053 | 6.3E-07 | Chondrogenic Fibroblasts | mtx2 |
| ufsp21 | 2E-11 | 0.4865003 | 0.123 | 0.07 | 6.4E-07 | Chondrogenic Fibroblasts | ufsp2 |
| rac1a1 | 2.2E-11 | 0.2172202 | 0.273 | 0.186 | 7.05E-07 | Chondrogenic Fibroblasts | rac1a |
| mef2d2 | 2.5E-11 | 0.2267601 | 0.276 | 0.186 | 8.2E-07 | Chondrogenic Fibroblasts | mef2d |
| zmiz1a3 | 2.7E-11 | 0.4132643 | 0.187 | 0.119 | 8.66E-07 | Chondrogenic Fibroblasts | zmiz1a |
| mgst3a2 | 2.7E-11 | 0.2800987 | 0.355 | 0.254 | 8.84E-07 | Chondrogenic Fibroblasts | mgst3a |
| derl12 | 2.8E-11 | 0.4807192 | 0.125 | 0.071 | 9.14E-07 | Chondrogenic Fibroblasts | derl1 |
| si:ch211-194k22.81 | 3E-11 | 0.3701349 | 0.155 | 0.094 | 9.66E-07 | Chondrogenic Fibroblasts | si:ch211-194k22.8 |
| srgap23 | 3.1E-11 | 0.4830599 | 0.141 | 0.083 | 9.94E-07 | Chondrogenic Fibroblasts | srgap2 |
| tent5ba2 | 3.1E-11 | 0.3839653 | 0.214 | 0.142 | 9.99E-07 | Chondrogenic Fibroblasts | tent5ba |
| wasla2 | 3.1E-11 | 0.3835095 | 0.257 | 0.176 | 1.02E-06 | Chondrogenic Fibroblasts | wasla |
| rrad2 | 3.4E-11 | 1.0852149 | 0.12 | 0.069 | 1.11E-06 | Chondrogenic Fibroblasts | rrad |
| prpf38b1 | 3.5E-11 | 0.2868541 | 0.183 | 0.115 | 1.13E-06 | Chondrogenic Fibroblasts | prpf38b |
| mrps141 | 3.5E-11 | 0.3623985 | 0.202 | 0.13 | 1.13E-06 | Chondrogenic Fibroblasts | mrps14 |
| rbm14a | 3.5E-11 | 0.493634 | 0.119 | 0.067 | 1.14E-06 | Chondrogenic Fibroblasts | rbm14a |
| nutf21 | 3.5E-11 | 0.311982 | 0.205 | 0.132 | 1.14E-06 | Chondrogenic Fibroblasts | nutf2 |
| SCARF23 | 3.6E-11 | 0.3181902 | 0.364 | 0.266 | 1.19E-06 | Chondrogenic Fibroblasts | SCARF2 |
| cox141 | 3.8E-11 | 0.3905164 | 0.141 | 0.084 | 1.23E-06 | Chondrogenic Fibroblasts | cox14 |
| gemin81 | 3.8E-11 | 0.374298 | 0.137 | 0.08 | 1.24E-06 | Chondrogenic Fibroblasts | gemin8 |
| ube2d2l2 | 3.8E-11 | 0.2181703 | 0.226 | 0.147 | 1.24E-06 | Chondrogenic Fibroblasts | ube2d2l |

| | | | | | | | |
|---|---|---|---|---|---|---|---|
| ENSDARG000000553 | 4E-11 | 0.3789576 | 0.133 | 0.078 | 1.29E-06 | Chondrogenic Fibroblasts | ENSDARG00000055 |
| rock12 | 4E-11 | 0.3250725 | 0.149 | 0.089 | 1.3E-06 | Chondrogenic Fibroblasts | rock1 |
| rgs12a1 | 4.1E-11 | 0.6579046 | 0.107 | 0.059 | 1.33E-06 | Chondrogenic Fibroblasts | rgs12a |
| znf3301 | 4.1E-11 | 0.5557853 | 0.1 | 0.054 | 1.34E-06 | Chondrogenic Fibroblasts | znf330 |
| pitpnbl1 | 5E-11 | 0.2602357 | 0.182 | 0.114 | 1.62E-06 | Chondrogenic Fibroblasts | pitpnbl |
| eea11 | 5.2E-11 | 0.3975229 | 0.128 | 0.074 | 1.7E-06 | Chondrogenic Fibroblasts | eea1 |
| tm9sf2 | 5.6E-11 | 0.2579674 | 0.252 | 0.17 | 1.81E-06 | Chondrogenic Fibroblasts | tm9sf2 |
| sart11 | 5.9E-11 | 0.2079539 | 0.249 | 0.166 | 1.9E-06 | Chondrogenic Fibroblasts | sart1 |
| snrnp701 | 5.9E-11 | 0.269698 | 0.246 | 0.164 | 1.92E-06 | Chondrogenic Fibroblasts | snrnp70 |
| etfa1 | 6E-11 | 0.2890143 | 0.236 | 0.158 | 1.96E-06 | Chondrogenic Fibroblasts | etfa |
| cnpy42 | 6.1E-11 | 0.3435038 | 0.192 | 0.123 | 1.98E-06 | Chondrogenic Fibroblasts | cnpy4 |
| mrps241 | 6.1E-11 | 0.4689776 | 0.124 | 0.071 | 1.99E-06 | Chondrogenic Fibroblasts | mrps24 |
| rap2c | 6.8E-11 | 0.318498 | 0.205 | 0.133 | 2.2E-06 | Chondrogenic Fibroblasts | rap2c |
| kars13 | 6.8E-11 | 0.2983847 | 0.243 | 0.165 | 2.21E-06 | Chondrogenic Fibroblasts | kars1 |
| dnaja2b3 | 7E-11 | 0.3104816 | 0.175 | 0.11 | 2.28E-06 | Chondrogenic Fibroblasts | dnaja2b |
| BX572619.1 | 7.4E-11 | 0.4014653 | 0.109 | 0.061 | 2.4E-06 | Chondrogenic Fibroblasts | BX572619.1 |
| drap11 | 7.4E-11 | 0.3614969 | 0.2 | 0.13 | 2.42E-06 | Chondrogenic Fibroblasts | drap1 |
| rassf14 | 7.5E-11 | 0.3260589 | 0.187 | 0.119 | 2.43E-06 | Chondrogenic Fibroblasts | rassf1 |
| ccdc91 | 7.7E-11 | 0.4155166 | 0.132 | 0.078 | 2.5E-06 | Chondrogenic Fibroblasts | ccdc9 |
| chd4b2 | 8E-11 | 0.2438469 | 0.358 | 0.257 | 2.62E-06 | Chondrogenic Fibroblasts | chd4b |
| ppm1ab1 | 8.3E-11 | 0.4651848 | 0.122 | 0.07 | 2.68E-06 | Chondrogenic Fibroblasts | ppm1ab |
| phrf1 | 8.6E-11 | 0.2400261 | 0.214 | 0.14 | 2.79E-06 | Chondrogenic Fibroblasts | phrf1 |
| si:ch73-91k6.21 | 9.3E-11 | 0.3615154 | 0.166 | 0.103 | 3.01E-06 | Chondrogenic Fibroblasts | si:ch73-91k6.2 |
| kpna1 | 9.3E-11 | 0.4884514 | 0.104 | 0.057 | 3.01E-06 | Chondrogenic Fibroblasts | kpna1 |
| nat92 | 9.3E-11 | 0.5569189 | 0.143 | 0.087 | 3.04E-06 | Chondrogenic Fibroblasts | nat9 |
| plpp52 | 9.4E-11 | 0.3772092 | 0.114 | 0.065 | 3.07E-06 | Chondrogenic Fibroblasts | plpp5 |
| otud5a | 9.5E-11 | 0.4003687 | 0.132 | 0.078 | 3.08E-06 | Chondrogenic Fibroblasts | otud5a |
| psme31 | 1E-10 | 0.2457378 | 0.15 | 0.091 | 3.24E-06 | Chondrogenic Fibroblasts | psme3 |
| lsm31 | 1E-10 | 0.263733 | 0.295 | 0.205 | 3.35E-06 | Chondrogenic Fibroblasts | lsm3 |
| riok11 | 1E-10 | 0.413365 | 0.117 | 0.067 | 3.37E-06 | Chondrogenic Fibroblasts | riok1 |
| nme33 | 1E-10 | 0.2192948 | 0.578 | 0.447 | 3.39E-06 | Chondrogenic Fibroblasts | nme3 |
| herpud11 | 1.1E-10 | 0.4780085 | 0.105 | 0.058 | 3.72E-06 | Chondrogenic Fibroblasts | herpud1 |
| gabpa | 1.1E-10 | 0.32884 | 0.123 | 0.071 | 3.73E-06 | Chondrogenic Fibroblasts | gabpa |
| leo11 | 1.2E-10 | 0.2868163 | 0.229 | 0.153 | 3.85E-06 | Chondrogenic Fibroblasts | leo1 |
| pik3c2a1 | 1.3E-10 | 0.382056 | 0.148 | 0.09 | 4.22E-06 | Chondrogenic Fibroblasts | pik3c2a |
| dlst1 | 1.3E-10 | 0.2533209 | 0.273 | 0.189 | 4.24E-06 | Chondrogenic Fibroblasts | dlst |
| kpna41 | 1.3E-10 | 0.2032441 | 0.218 | 0.143 | 4.31E-06 | Chondrogenic Fibroblasts | kpna4 |
| scaf1 | 1.3E-10 | 0.2079727 | 0.328 | 0.233 | 4.34E-06 | Chondrogenic Fibroblasts | scaf1 |
| PKN2 | 1.4E-10 | 0.368822 | 0.11 | 0.062 | 4.49E-06 | Chondrogenic Fibroblasts | PKN2 |
| pde4ba3 | 1.4E-10 | 0.329866 | 0.182 | 0.116 | 4.5E-06 | Chondrogenic Fibroblasts | pde4ba |
| si:ch211-107o10.32 | 1.4E-10 | 0.3688935 | 0.152 | 0.093 | 4.57E-06 | Chondrogenic Fibroblasts | si:ch211-107o10.3 |
| acot9.1 | 1.5E-10 | 0.5141695 | 0.106 | 0.059 | 4.91E-06 | Chondrogenic Fibroblasts | acot9.1 |
| si:dkey-235d18.53 | 1.5E-10 | 0.2079112 | 0.159 | 0.097 | 4.94E-06 | Chondrogenic Fibroblasts | si:dkey-235d18.5 |
| rnf19a1 | 1.5E-10 | 0.2987732 | 0.241 | 0.163 | 4.99E-06 | Chondrogenic Fibroblasts | rnf19a |
| hmgb1a3 | 1.6E-10 | 0.2425589 | 0.675 | 0.562 | 5.08E-06 | Chondrogenic Fibroblasts | hmgb1a |
| ostf11 | 1.6E-10 | 0.4537054 | 0.135 | 0.08 | 5.19E-06 | Chondrogenic Fibroblasts | ostf1 |
| abtb2b2 | 1.7E-10 | 0.4061372 | 0.126 | 0.074 | 5.65E-06 | Chondrogenic Fibroblasts | abtb2b |
| jpt2 | 1.8E-10 | 0.2653134 | 0.282 | 0.197 | 5.82E-06 | Chondrogenic Fibroblasts | jpt2 |

| | | | | | | | |
|---|---|---|---|---|---|---|---|
| rars11 | 1.8E-10 | 0.4750499 | 0.112 | 0.063 | 5.89E-06 | Chondrogenic Fibroblasts | rars1 |
| si:dkey-282h22.53 | 1.8E-10 | 0.9251004 | 0.119 | 0.07 | 5.96E-06 | Chondrogenic Fibroblasts | si:dkey-282h22.5 |
| ssu721 | 1.9E-10 | 0.3651515 | 0.152 | 0.094 | 6.08E-06 | Chondrogenic Fibroblasts | ssu72 |
| tm2d31 | 1.9E-10 | 0.3859586 | 0.121 | 0.07 | 6.3E-06 | Chondrogenic Fibroblasts | tm2d3 |
| zgc:921404 | 2E-10 | 0.4074352 | 0.176 | 0.113 | 6.53E-06 | Chondrogenic Fibroblasts | zgc:92140 |
| mrpl142 | 2E-10 | 0.4194478 | 0.122 | 0.071 | 6.64E-06 | Chondrogenic Fibroblasts | mrpl14 |
| nucks1b2 | 2E-10 | 0.2132804 | 0.423 | 0.315 | 6.65E-06 | Chondrogenic Fibroblasts | nucks1b |
| epn23 | 2.2E-10 | 0.4047809 | 0.17 | 0.108 | 7.18E-06 | Chondrogenic Fibroblasts | epn2 |
| cnot91 | 2.3E-10 | 0.4275657 | 0.13 | 0.077 | 7.51E-06 | Chondrogenic Fibroblasts | cnot9 |
| tbl1xr1b | 2.5E-10 | 0.2749368 | 0.117 | 0.067 | 8.16E-06 | Chondrogenic Fibroblasts | tbl1xr1b |
| snx1a | 2.6E-10 | 0.2682787 | 0.15 | 0.092 | 8.45E-06 | Chondrogenic Fibroblasts | snx1a |
| zgc:1626983 | 2.6E-10 | 0.2660261 | 0.178 | 0.113 | 8.45E-06 | Chondrogenic Fibroblasts | zgc:162698 |
| nbr1b1 | 2.7E-10 | 0.454432 | 0.111 | 0.063 | 8.77E-06 | Chondrogenic Fibroblasts | nbr1b |
| si:ch211-22i13.21 | 3E-10 | 0.443389 | 0.125 | 0.074 | 9.76E-06 | Chondrogenic Fibroblasts | si:ch211-22i13.2 |
| cry1a1 | 3.1E-10 | 0.2080283 | 0.24 | 0.161 | 1.01E-05 | Chondrogenic Fibroblasts | cry1a |
| adam92 | 3.3E-10 | 0.4292275 | 0.115 | 0.066 | 1.06E-05 | Chondrogenic Fibroblasts | adam9 |
| pcnp | 3.4E-10 | 0.252904 | 0.231 | 0.156 | 1.11E-05 | Chondrogenic Fibroblasts | pcnp |
| phactr24 | 3.6E-10 | 0.2720675 | 0.212 | 0.139 | 1.16E-05 | Chondrogenic Fibroblasts | phactr2 |
| pgd2 | 4.1E-10 | 0.5174515 | 0.13 | 0.078 | 1.35E-05 | Chondrogenic Fibroblasts | pgd |
| znf3462 | 4.2E-10 | 0.327494 | 0.148 | 0.091 | 1.35E-05 | Chondrogenic Fibroblasts | znf346 |
| mmp15a4 | 4.2E-10 | 0.3030484 | 0.137 | 0.083 | 1.36E-05 | Chondrogenic Fibroblasts | mmp15a |
| anapc112 | 4.3E-10 | 0.2115818 | 0.197 | 0.128 | 1.4E-05 | Chondrogenic Fibroblasts | anapc11 |
| ubfd11 | 4.4E-10 | 0.3618054 | 0.167 | 0.106 | 1.45E-05 | Chondrogenic Fibroblasts | ubfd1 |
| smarcd12 | 4.6E-10 | 0.3731538 | 0.117 | 0.068 | 1.5E-05 | Chondrogenic Fibroblasts | smarcd1 |
| si:ch73-37h15.21 | 4.7E-10 | 0.2455958 | 0.17 | 0.107 | 1.52E-05 | Chondrogenic Fibroblasts | si:ch73-37h15.2 |
| phf20b2 | 4.7E-10 | 0.2917906 | 0.253 | 0.174 | 1.52E-05 | Chondrogenic Fibroblasts | phf20b |
| armh31 | 4.8E-10 | 0.3150027 | 0.14 | 0.085 | 1.55E-05 | Chondrogenic Fibroblasts | armh3 |
| clip22 | 5.2E-10 | 0.3030593 | 0.162 | 0.102 | 1.69E-05 | Chondrogenic Fibroblasts | clip2 |
| slc35e12 | 5.5E-10 | 0.476652 | 0.101 | 0.056 | 1.8E-05 | Chondrogenic Fibroblasts | slc35e1 |
| FO704772.31 | 5.6E-10 | 0.2070593 | 0.436 | 0.327 | 1.81E-05 | Chondrogenic Fibroblasts | FO704772.3 |
| ndufa112 | 5.6E-10 | 0.3108661 | 0.201 | 0.133 | 1.81E-05 | Chondrogenic Fibroblasts | ndufa11 |
| tgfb54 | 5.6E-10 | 0.2482321 | 0.175 | 0.113 | 1.81E-05 | Chondrogenic Fibroblasts | tgfb5 |
| CABZ01044053.11 | 5.7E-10 | 0.2586515 | 0.134 | 0.08 | 1.85E-05 | Chondrogenic Fibroblasts | CABZ01044053.1 |
| med211 | 5.8E-10 | 0.4773239 | 0.119 | 0.07 | 1.89E-05 | Chondrogenic Fibroblasts | med21 |
| farsa3 | 6.2E-10 | 0.3954047 | 0.133 | 0.081 | 2.03E-05 | Chondrogenic Fibroblasts | farsa |
| cpxm1a2 | 6.5E-10 | 0.2867073 | 0.255 | 0.177 | 2.13E-05 | Chondrogenic Fibroblasts | cpxm1a |
| atp5md1 | 6.8E-10 | 0.2884461 | 0.213 | 0.143 | 2.23E-05 | Chondrogenic Fibroblasts | atp5md |
| nfkbie1 | 8E-10 | 0.2896419 | 0.125 | 0.074 | 2.6E-05 | Chondrogenic Fibroblasts | nfkbie |
| rab1ba1 | 8.4E-10 | 0.2418594 | 0.209 | 0.139 | 2.72E-05 | Chondrogenic Fibroblasts | rab1ba |
| brd71 | 8.4E-10 | 0.3847771 | 0.105 | 0.06 | 2.73E-05 | Chondrogenic Fibroblasts | brd7 |
| selenoh1 | 1E-09 | 0.3020073 | 0.186 | 0.122 | 3.33E-05 | Chondrogenic Fibroblasts | selenoh |
| fcf11 | 1.1E-09 | 0.4766724 | 0.103 | 0.059 | 3.44E-05 | Chondrogenic Fibroblasts | fcf1 |
| actl6a1 | 1.1E-09 | 0.4694207 | 0.103 | 0.059 | 3.6E-05 | Chondrogenic Fibroblasts | actl6a |
| ube2e12 | 1.1E-09 | 0.252385 | 0.179 | 0.116 | 3.69E-05 | Chondrogenic Fibroblasts | ube2e1 |
| actr101 | 1.2E-09 | 0.4260191 | 0.113 | 0.067 | 3.78E-05 | Chondrogenic Fibroblasts | actr10 |
| si:ch73-376l24.2 | 1.2E-09 | 0.219882 | 0.133 | 0.08 | 3.84E-05 | Chondrogenic Fibroblasts | si:ch73-376l24.2 |
| gal3st23 | 1.2E-09 | 0.5360825 | 0.125 | 0.076 | 3.93E-05 | Chondrogenic Fibroblasts | gal3st2 |
| AL935174.61 | 1.2E-09 | 0.3865768 | 0.132 | 0.08 | 3.96E-05 | Chondrogenic Fibroblasts | AL935174.6 |

| | | | | | | | |
|---|---|---|---|---|---|---|---|
| galnt22 | 1.2E-09 | 0.226089 | 0.145 | 0.089 | 3.99E-05 | Chondrogenic Fibroblasts | galnt2 |
| vkorc1l11 | 1.3E-09 | 0.2356738 | 0.145 | 0.09 | 4.09E-05 | Chondrogenic Fibroblasts | vkorc1l1 |
| chp21 | 1.3E-09 | 0.4958167 | 0.129 | 0.078 | 4.29E-05 | Chondrogenic Fibroblasts | chp2 |
| ubr72 | 1.4E-09 | 0.4578079 | 0.148 | 0.093 | 4.47E-05 | Chondrogenic Fibroblasts | ubr7 |
| blmh2 | 1.4E-09 | 0.5397683 | 0.101 | 0.058 | 4.49E-05 | Chondrogenic Fibroblasts | blmh |
| klf6a2 | 1.5E-09 | 0.2685204 | 0.809 | 0.745 | 4.72E-05 | Chondrogenic Fibroblasts | klf6a |
| hmgb3b3 | 1.5E-09 | 0.3571243 | 0.178 | 0.116 | 4.81E-05 | Chondrogenic Fibroblasts | hmgb3b |
| clasp21 | 1.6E-09 | 0.3415656 | 0.12 | 0.071 | 5.13E-05 | Chondrogenic Fibroblasts | clasp2 |
| mrpl362 | 1.6E-09 | 0.2951744 | 0.204 | 0.137 | 5.18E-05 | Chondrogenic Fibroblasts | mrpl36 |
| C23H1orf52 | 1.8E-09 | 0.2454097 | 0.168 | 0.108 | 5.93E-05 | Chondrogenic Fibroblasts | C23H1orf52 |
| selenon3 | 1.8E-09 | 0.2002115 | 0.229 | 0.155 | 5.99E-05 | Chondrogenic Fibroblasts | selenon |
| cox6b13 | 1.9E-09 | 0.603709 | 0.116 | 0.07 | 6.34E-05 | Chondrogenic Fibroblasts | cox6b1 |
| purba1 | 2E-09 | 0.2099522 | 0.192 | 0.127 | 6.35E-05 | Chondrogenic Fibroblasts | purba |
| atp10a3 | 2.1E-09 | 0.3218317 | 0.152 | 0.096 | 6.87E-05 | Chondrogenic Fibroblasts | atp10a |
| tmem9b1 | 2.2E-09 | 0.294277 | 0.16 | 0.102 | 7.13E-05 | Chondrogenic Fibroblasts | tmem9b |
| hars2 | 2.2E-09 | 0.2803796 | 0.142 | 0.088 | 7.18E-05 | Chondrogenic Fibroblasts | hars |
| mtus1b | 2.3E-09 | 0.5855932 | 0.114 | 0.068 | 7.55E-05 | Chondrogenic Fibroblasts | mtus1b |
| C5H9orf781 | 2.4E-09 | 0.2625932 | 0.139 | 0.086 | 7.88E-05 | Chondrogenic Fibroblasts | C5H9orf78 |
| churc11 | 2.5E-09 | 0.3009398 | 0.192 | 0.129 | 7.98E-05 | Chondrogenic Fibroblasts | churc1 |
| COX7A21 | 2.6E-09 | 0.2880731 | 0.194 | 0.13 | 8.53E-05 | Chondrogenic Fibroblasts | COX7A2 |
| ppm1bb | 2.7E-09 | 0.2795754 | 0.132 | 0.081 | 8.69E-05 | Chondrogenic Fibroblasts | ppm1bb |
| rbm71 | 2.8E-09 | 0.3025312 | 0.139 | 0.086 | 9.13E-05 | Chondrogenic Fibroblasts | rbm7 |
| uchl51 | 2.9E-09 | 0.4326693 | 0.125 | 0.076 | 9.39E-05 | Chondrogenic Fibroblasts | uchl5 |
| zgc:77486.14 | 2.9E-09 | 0.2602166 | 0.242 | 0.168 | 9.42E-05 | Chondrogenic Fibroblasts | zgc:77486.1 |
| usf1l2 | 3E-09 | 0.4174967 | 0.117 | 0.07 | 9.76E-05 | Chondrogenic Fibroblasts | usf1l |
| lamp1b2 | 3E-09 | 0.3125423 | 0.142 | 0.088 | 9.9E-05 | Chondrogenic Fibroblasts | lamp1b |
| ndufv22 | 3.2E-09 | 0.3069154 | 0.162 | 0.105 | 0.000103 | Chondrogenic Fibroblasts | ndufv2 |
| FOXK23 | 3.3E-09 | 0.2039023 | 0.202 | 0.136 | 0.000107 | Chondrogenic Fibroblasts | FOXK2 |
| eci22 | 3.3E-09 | 0.366328 | 0.109 | 0.064 | 0.000108 | Chondrogenic Fibroblasts | eci2 |
| dpm22 | 3.4E-09 | 0.3629544 | 0.203 | 0.138 | 0.000109 | Chondrogenic Fibroblasts | dpm2 |
| fam168a1 | 3.8E-09 | 0.2198975 | 0.303 | 0.219 | 0.000124 | Chondrogenic Fibroblasts | fam168a |
| samd4a1 | 4E-09 | 0.43561 | 0.107 | 0.063 | 0.000129 | Chondrogenic Fibroblasts | samd4a |
| col21a13 | 4.1E-09 | 0.4494032 | 0.129 | 0.079 | 0.000134 | Chondrogenic Fibroblasts | col21a1 |
| mafba5 | 4.4E-09 | 0.3674154 | 0.486 | 0.389 | 0.000142 | Chondrogenic Fibroblasts | mafba |
| ube2d1a1 | 4.4E-09 | 0.2848751 | 0.158 | 0.101 | 0.000142 | Chondrogenic Fibroblasts | ube2d1a |
| per1a1 | 4.6E-09 | 0.3720615 | 0.115 | 0.068 | 0.00015 | Chondrogenic Fibroblasts | per1a |
| twsg1a4 | 4.7E-09 | 0.2261526 | 0.347 | 0.256 | 0.000151 | Chondrogenic Fibroblasts | twsg1a |
| map7d31 | 4.8E-09 | 0.3048404 | 0.129 | 0.079 | 0.000156 | Chondrogenic Fibroblasts | map7d3 |
| ypel51 | 4.9E-09 | 0.4866868 | 0.107 | 0.063 | 0.000159 | Chondrogenic Fibroblasts | ypel5 |
| cops41 | 4.9E-09 | 0.3358205 | 0.169 | 0.111 | 0.00016 | Chondrogenic Fibroblasts | cops4 |
| acadm1 | 4.9E-09 | 0.2031738 | 0.343 | 0.253 | 0.00016 | Chondrogenic Fibroblasts | acadm |
| isca1 | 5.1E-09 | 0.5876562 | 0.106 | 0.062 | 0.000167 | Chondrogenic Fibroblasts | isca1 |
| fbn2b1 | 5.3E-09 | 0.2224601 | 0.116 | 0.069 | 0.000171 | Chondrogenic Fibroblasts | fbn2b |
| pdia42 | 5.3E-09 | 0.2188299 | 0.216 | 0.148 | 0.000171 | Chondrogenic Fibroblasts | pdia4 |
| nfyba3 | 5.4E-09 | 0.2057993 | 0.259 | 0.181 | 0.000176 | Chondrogenic Fibroblasts | nfyba |
| twist1b3 | 5.4E-09 | 0.2784541 | 0.229 | 0.158 | 0.000176 | Chondrogenic Fibroblasts | twist1b |
| march5l | 5.5E-09 | 0.2521415 | 0.108 | 0.064 | 0.000178 | Chondrogenic Fibroblasts | march5l |
| eif2b23 | 5.7E-09 | 0.521781 | 0.115 | 0.07 | 0.000184 | Chondrogenic Fibroblasts | eif2b2 |

| | | | | | | | |
|---|---|---|---|---|---|---|---|
| me31 | 5.8E-09 | 0.3219183 | 0.135 | 0.084 | 0.000187 | Chondrogenic Fibroblasts | me3 |
| eif4ba1 | 5.8E-09 | 0.2370839 | 0.183 | 0.121 | 0.000189 | Chondrogenic Fibroblasts | eif4ba |
| med31 | 5.8E-09 | 0.5575807 | 0.112 | 0.067 | 0.00019 | Chondrogenic Fibroblasts | med31 |
| pebp14 | 6E-09 | 0.4653354 | 0.119 | 0.072 | 0.000196 | Chondrogenic Fibroblasts | pebp1 |
| etv5a4 | 6.2E-09 | 0.310169 | 0.159 | 0.103 | 0.0002 | Chondrogenic Fibroblasts | etv5a |
| carhsp1 | 6.2E-09 | 0.5524639 | 0.105 | 0.062 | 0.000203 | Chondrogenic Fibroblasts | carhsp1 |
| eif3ja1 | 6.5E-09 | 0.2586783 | 0.171 | 0.112 | 0.000212 | Chondrogenic Fibroblasts | eif3ja |
| pcyt21 | 6.6E-09 | 0.2115557 | 0.15 | 0.095 | 0.000214 | Chondrogenic Fibroblasts | pcyt2 |
| dlat1 | 6.9E-09 | 0.4701234 | 0.124 | 0.076 | 0.000223 | Chondrogenic Fibroblasts | dlat |
| ube2g1a1 | 6.9E-09 | 0.2556706 | 0.134 | 0.083 | 0.000223 | Chondrogenic Fibroblasts | ube2g1a |
| ncbp22 | 7.2E-09 | 0.2297563 | 0.183 | 0.122 | 0.000233 | Chondrogenic Fibroblasts | ncbp2 |
| amot | 8E-09 | 0.2646924 | 0.112 | 0.066 | 0.000261 | Chondrogenic Fibroblasts | amot |
| cdkn1ba4 | 8.2E-09 | 0.4439088 | 0.174 | 0.116 | 0.000267 | Chondrogenic Fibroblasts | cdkn1ba |
| psmd4b1 | 8.4E-09 | 0.2527023 | 0.213 | 0.146 | 0.000274 | Chondrogenic Fibroblasts | psmd4b |
| dr11 | 8.8E-09 | 0.405308 | 0.116 | 0.07 | 0.000286 | Chondrogenic Fibroblasts | dr1 |
| dpp9 | 8.8E-09 | 0.4971961 | 0.101 | 0.059 | 0.000287 | Chondrogenic Fibroblasts | dpp9 |
| inpp5l1 | 9.2E-09 | 0.2179824 | 0.151 | 0.097 | 0.000301 | Chondrogenic Fibroblasts | inpp5l |
| dhx152 | 9.3E-09 | 0.3820117 | 0.167 | 0.11 | 0.000301 | Chondrogenic Fibroblasts | dhx15 |
| cfap201 | 9.4E-09 | 0.5438678 | 0.117 | 0.071 | 0.000304 | Chondrogenic Fibroblasts | cfap20 |
| gtf2f2a1 | 1.2E-08 | 0.2017756 | 0.196 | 0.132 | 0.000383 | Chondrogenic Fibroblasts | gtf2f2a |
| rbmx1 | 1.2E-08 | 0.2517482 | 0.188 | 0.127 | 0.000387 | Chondrogenic Fibroblasts | rbmx |
| ube2ia1 | 1.2E-08 | 0.2124057 | 0.214 | 0.147 | 0.000395 | Chondrogenic Fibroblasts | ube2ia |
| naa501 | 1.3E-08 | 0.3264993 | 0.198 | 0.135 | 0.000409 | Chondrogenic Fibroblasts | naa50 |
| vta12 | 1.3E-08 | 0.3853277 | 0.121 | 0.075 | 0.000415 | Chondrogenic Fibroblasts | vta1 |
| rtraf2 | 1.3E-08 | 0.3288414 | 0.16 | 0.104 | 0.000416 | Chondrogenic Fibroblasts | rtraf |
| arih22 | 1.4E-08 | 0.3283116 | 0.129 | 0.08 | 0.000441 | Chondrogenic Fibroblasts | arih2 |
| stard13b3 | 1.4E-08 | 0.3632028 | 0.132 | 0.083 | 0.000449 | Chondrogenic Fibroblasts | stard13b |
| commd81 | 1.4E-08 | 0.38938 | 0.118 | 0.072 | 0.00046 | Chondrogenic Fibroblasts | commd8 |
| adamts52 | 1.5E-08 | 0.4644875 | 0.107 | 0.064 | 0.000472 | Chondrogenic Fibroblasts | adamts5 |
| faub2 | 1.5E-08 | 0.2527226 | 0.136 | 0.085 | 0.000492 | Chondrogenic Fibroblasts | faub |
| kank33 | 1.7E-08 | 0.2587524 | 0.128 | 0.079 | 0.00054 | Chondrogenic Fibroblasts | kank3 |
| map1sa1 | 1.7E-08 | 0.370929 | 0.151 | 0.098 | 0.000548 | Chondrogenic Fibroblasts | map1sa |
| tmem184ba2 | 1.8E-08 | 0.3952858 | 0.113 | 0.069 | 0.000573 | Chondrogenic Fibroblasts | tmem184ba |
| lgals3b2 | 1.8E-08 | 0.236411 | 0.505 | 0.406 | 0.000577 | Chondrogenic Fibroblasts | lgals3b |
| btbd6a3 | 1.8E-08 | 0.5418326 | 0.125 | 0.079 | 0.000591 | Chondrogenic Fibroblasts | btbd6a |
| BOLA21 | 2E-08 | 0.3879716 | 0.147 | 0.096 | 0.000635 | Chondrogenic Fibroblasts | BOLA2 |
| parp161 | 2E-08 | 0.3156391 | 0.137 | 0.088 | 0.00066 | Chondrogenic Fibroblasts | parp16 |
| ergic32 | 2.1E-08 | 0.287625 | 0.102 | 0.06 | 0.000668 | Chondrogenic Fibroblasts | ergic3 |
| rhogc | 2.1E-08 | 0.404446 | 0.107 | 0.064 | 0.000686 | Chondrogenic Fibroblasts | rhogc |
| c1galt1c12 | 2.1E-08 | 0.2956225 | 0.113 | 0.069 | 0.000697 | Chondrogenic Fibroblasts | c1galt1c1 |
| mff1 | 2.2E-08 | 0.4623288 | 0.113 | 0.07 | 0.000715 | Chondrogenic Fibroblasts | mff |
| exosc32 | 2.5E-08 | 0.4232904 | 0.116 | 0.072 | 0.000829 | Chondrogenic Fibroblasts | exosc3 |
| robo12 | 2.7E-08 | 0.282943 | 0.125 | 0.078 | 0.000875 | Chondrogenic Fibroblasts | robo1 |
| rps27l2 | 2.8E-08 | 0.4277578 | 0.373 | 0.287 | 0.000921 | Chondrogenic Fibroblasts | rps27l |
| tfe3a3 | 2.9E-08 | 0.5457014 | 0.144 | 0.094 | 0.000935 | Chondrogenic Fibroblasts | tfe3a |
| ugp2b2 | 3E-08 | 0.2511182 | 0.174 | 0.117 | 0.000984 | Chondrogenic Fibroblasts | ugp2b |
| btbd10a1 | 3.2E-08 | 0.2822392 | 0.119 | 0.074 | 0.00104 | Chondrogenic Fibroblasts | btbd10a |
| zgc:771582 | 3.7E-08 | 0.6339767 | 0.101 | 0.061 | 0.001204 | Chondrogenic Fibroblasts | zgc:77158 |

| | | | | | | | |
|---|---|---|---|---|---|---|---|
| atp6v0cb1 | 3.8E-08 | 0.3516263 | 0.13 | 0.082 | 0.001225 | Chondrogenic Fibroblasts | atp6v0cb |
| npepl11 | 3.9E-08 | 0.2812697 | 0.167 | 0.112 | 0.001258 | Chondrogenic Fibroblasts | npepl1 |
| zgc:55558 | 4E-08 | 0.2985307 | 0.103 | 0.062 | 0.001292 | Chondrogenic Fibroblasts | zgc:55558 |
| ganab2 | 4.2E-08 | 0.3079416 | 0.101 | 0.06 | 0.001365 | Chondrogenic Fibroblasts | ganab |
| slu7 | 4.3E-08 | 0.2560475 | 0.157 | 0.104 | 0.001404 | Chondrogenic Fibroblasts | slu7 |
| pde10a1 | 4.6E-08 | 0.4212198 | 0.103 | 0.062 | 0.001488 | Chondrogenic Fibroblasts | pde10a |
| dcun1d52 | 5E-08 | 0.4720277 | 0.118 | 0.074 | 0.001618 | Chondrogenic Fibroblasts | dcun1d5 |
| zgc:152830 | 5.5E-08 | 0.3319768 | 0.11 | 0.068 | 0.001787 | Chondrogenic Fibroblasts | zgc:152830 |
| ufd1l3 | 5.7E-08 | 0.3244358 | 0.158 | 0.106 | 0.001856 | Chondrogenic Fibroblasts | ufd1l |
| aff42 | 5.7E-08 | 0.279209 | 0.133 | 0.085 | 0.001857 | Chondrogenic Fibroblasts | aff4 |
| gosr22 | 5.9E-08 | 0.3124447 | 0.14 | 0.091 | 0.001921 | Chondrogenic Fibroblasts | gosr2 |
| adssl1 | 6.1E-08 | 0.3091379 | 0.128 | 0.081 | 0.001987 | Chondrogenic Fibroblasts | adssl |
| mtrr1 | 6.3E-08 | 0.3036323 | 0.141 | 0.092 | 0.002049 | Chondrogenic Fibroblasts | mtrr |
| golga41 | 6.5E-08 | 0.2741308 | 0.227 | 0.161 | 0.002118 | Chondrogenic Fibroblasts | golga4 |
| mob1bb2 | 7E-08 | 0.342866 | 0.128 | 0.082 | 0.002266 | Chondrogenic Fibroblasts | mob1bb |
| pop73 | 7.8E-08 | 0.3932242 | 0.111 | 0.069 | 0.00252 | Chondrogenic Fibroblasts | pop7 |
| serpinb1l31 | 8.3E-08 | 0.3743283 | 0.136 | 0.088 | 0.002708 | Chondrogenic Fibroblasts | serpinb1l3 |
| acadl1 | 1E-07 | 0.2329105 | 0.14 | 0.091 | 0.003292 | Chondrogenic Fibroblasts | acadl |
| polr2eb1 | 1E-07 | 0.2560998 | 0.211 | 0.15 | 0.003318 | Chondrogenic Fibroblasts | polr2eb |
| fnta2 | 1.1E-07 | 0.4069498 | 0.109 | 0.068 | 0.003717 | Chondrogenic Fibroblasts | fnta |
| thoc71 | 1.2E-07 | 0.5022208 | 0.104 | 0.064 | 0.003894 | Chondrogenic Fibroblasts | thoc7 |
| b3gnt5a3 | 1.3E-07 | 0.302785 | 0.164 | 0.111 | 0.00412 | Chondrogenic Fibroblasts | b3gnt5a |
| morf4l12 | 1.3E-07 | 0.2211558 | 0.244 | 0.177 | 0.004263 | Chondrogenic Fibroblasts | morf4l1 |
| nudcd21 | 1.4E-07 | 0.345919 | 0.119 | 0.075 | 0.004468 | Chondrogenic Fibroblasts | nudcd2 |
| gtf2e21 | 1.4E-07 | 0.2971883 | 0.141 | 0.092 | 0.004484 | Chondrogenic Fibroblasts | gtf2e2 |
| dpm32 | 1.4E-07 | 0.2313159 | 0.143 | 0.093 | 0.004554 | Chondrogenic Fibroblasts | dpm3 |
| plekha51 | 1.5E-07 | 0.3162583 | 0.113 | 0.071 | 0.004762 | Chondrogenic Fibroblasts | plekha5 |
| psat12 | 1.5E-07 | 0.3393206 | 0.11 | 0.068 | 0.004877 | Chondrogenic Fibroblasts | psat1 |
| sept9a3 | 1.5E-07 | 0.2390379 | 0.29 | 0.215 | 0.004889 | Chondrogenic Fibroblasts | sept9a |
| gtf2a21 | 1.6E-07 | 0.2809821 | 0.162 | 0.11 | 0.005241 | Chondrogenic Fibroblasts | gtf2a2 |
| snapin | 1.6E-07 | 0.3186076 | 0.101 | 0.062 | 0.005292 | Chondrogenic Fibroblasts | snapin |
| arih1 | 1.6E-07 | 0.2174404 | 0.125 | 0.08 | 0.005294 | Chondrogenic Fibroblasts | arih1 |
| crema2 | 1.7E-07 | 0.5177912 | 0.159 | 0.108 | 0.005396 | Chondrogenic Fibroblasts | crema |
| gpkow | 2E-07 | 0.3073912 | 0.101 | 0.062 | 0.006399 | Chondrogenic Fibroblasts | gpkow |
| gsc4 | 2E-07 | 0.4074756 | 0.17 | 0.116 | 0.006545 | Chondrogenic Fibroblasts | gsc |
| arl21 | 2.1E-07 | 0.5070634 | 0.117 | 0.075 | 0.006775 | Chondrogenic Fibroblasts | arl2 |
| tmem111 | 2.1E-07 | 0.2290174 | 0.153 | 0.102 | 0.006822 | Chondrogenic Fibroblasts | tmem11 |
| ppp1r3ca4 | 2.2E-07 | 0.2367089 | 0.28 | 0.207 | 0.006997 | Chondrogenic Fibroblasts | ppp1r3ca |
| timm131 | 2.2E-07 | 0.3652954 | 0.104 | 0.064 | 0.007169 | Chondrogenic Fibroblasts | timm13 |
| zgc:1105913 | 2.5E-07 | 0.2235609 | 0.141 | 0.093 | 0.008143 | Chondrogenic Fibroblasts | zgc:110591 |
| vwa5a | 2.7E-07 | 0.2008618 | 0.108 | 0.067 | 0.00894 | Chondrogenic Fibroblasts | vwa5a |
| krt18b3 | 3.2E-07 | 0.2118305 | 0.923 | 0.891 | 0.010322 | Chondrogenic Fibroblasts | krt18b |
| acsl4a3 | 3.3E-07 | 0.356611 | 0.12 | 0.077 | 0.010884 | Chondrogenic Fibroblasts | acsl4a |
| rbm221 | 3.5E-07 | 0.2476773 | 0.153 | 0.103 | 0.011234 | Chondrogenic Fibroblasts | rbm22 |
| uqcrb1 | 4.4E-07 | 0.4443969 | 0.101 | 0.063 | 0.014223 | Chondrogenic Fibroblasts | uqcrb |
| rpe2 | 4.4E-07 | 0.3210439 | 0.139 | 0.093 | 0.014296 | Chondrogenic Fibroblasts | rpe |
| tmf1 | 4.6E-07 | 0.2700781 | 0.119 | 0.077 | 0.014846 | Chondrogenic Fibroblasts | tmf1 |
| hikeshi2 | 4.6E-07 | 0.3080723 | 0.151 | 0.103 | 0.014881 | Chondrogenic Fibroblasts | hikeshi |

| | | | | | | | |
|---|---|---|---|---|---|---|---|
| MYO1D2 | 4.6E-07 | 0.2757418 | 0.15 | 0.102 | 0.014893 | Chondrogenic Fibroblasts | MYO1D |
| mrc24 | 5.1E-07 | 0.2431035 | 0.133 | 0.088 | 0.016469 | Chondrogenic Fibroblasts | mrc2 |
| rap1ab1 | 5.2E-07 | 0.3940076 | 0.107 | 0.068 | 0.016855 | Chondrogenic Fibroblasts | rap1ab |
| terfa2 | 5.3E-07 | 0.3537649 | 0.132 | 0.088 | 0.017153 | Chondrogenic Fibroblasts | terfa |
| prelid1a3 | 5.8E-07 | 0.2476339 | 0.178 | 0.124 | 0.01902 | Chondrogenic Fibroblasts | prelid1a |
| prpf31 | 6.2E-07 | 0.2117079 | 0.115 | 0.074 | 0.020083 | Chondrogenic Fibroblasts | prpf3 |
| si:dkey-42i9.41 | 6.3E-07 | 0.3061328 | 0.144 | 0.098 | 0.020529 | Chondrogenic Fibroblasts | si:dkey-42i9.4 |
| wdr52 | 7.1E-07 | 0.3061258 | 0.106 | 0.067 | 0.023107 | Chondrogenic Fibroblasts | wdr5 |
| aimp23 | 7.2E-07 | 0.229697 | 0.233 | 0.171 | 0.023566 | Chondrogenic Fibroblasts | aimp2 |
| ccdc1861 | 7.5E-07 | 0.2537715 | 0.118 | 0.076 | 0.024515 | Chondrogenic Fibroblasts | ccdc186 |
| med12 | 8E-07 | 0.2585869 | 0.131 | 0.087 | 0.025859 | Chondrogenic Fibroblasts | med12 |
| cops81 | 8.2E-07 | 0.2336574 | 0.131 | 0.088 | 0.026729 | Chondrogenic Fibroblasts | cops8 |
| si:ch211-195b15.84 | 8.5E-07 | 0.3303767 | 0.135 | 0.09 | 0.027485 | Chondrogenic Fibroblasts | si:ch211-195b15.8 |
| fntb2 | 8.7E-07 | 0.3292747 | 0.153 | 0.106 | 0.02823 | Chondrogenic Fibroblasts | fntb |
| ercc6 | 8.9E-07 | 0.3207775 | 0.103 | 0.065 | 0.028912 | Chondrogenic Fibroblasts | ercc6 |
| snx72 | 9.9E-07 | 0.4490009 | 0.115 | 0.076 | 0.032112 | Chondrogenic Fibroblasts | snx7 |
| abhd17b | 1.1E-06 | 0.3765398 | 0.103 | 0.066 | 0.03447 | Chondrogenic Fibroblasts | abhd17b |
| psmd11a1 | 1.1E-06 | 0.3067896 | 0.137 | 0.093 | 0.034735 | Chondrogenic Fibroblasts | psmd11a |
| sccpdhb | 1.1E-06 | 0.2941054 | 0.104 | 0.066 | 0.035127 | Chondrogenic Fibroblasts | sccpdhb |
| uqcc12 | 1.1E-06 | 0.2200106 | 0.129 | 0.085 | 0.035294 | Chondrogenic Fibroblasts | uqcc1 |
| tusc2b1 | 1.1E-06 | 0.217658 | 0.119 | 0.078 | 0.035445 | Chondrogenic Fibroblasts | tusc2b |
| ocrl | 1.2E-06 | 0.3135942 | 0.107 | 0.068 | 0.039606 | Chondrogenic Fibroblasts | ocrl |
| phb2b1 | 1.2E-06 | 0.3330088 | 0.114 | 0.075 | 0.040422 | Chondrogenic Fibroblasts | phb2b |
| phlda23 | 1.5E-06 | 0.2458563 | 0.284 | 0.219 | 0.04726 | Chondrogenic Fibroblasts | phlda2 |
| CDPF13 | 1.5E-06 | 0.2047471 | 0.241 | 0.178 | 0.049407 | Chondrogenic Fibroblasts | CDPF1 |
| naa381 | 1.6E-06 | 0.2933968 | 0.1 | 0.064 | 0.052892 | Chondrogenic Fibroblasts | naa38 |
| zgc:770562 | 1.8E-06 | 0.3596588 | 0.106 | 0.069 | 0.059056 | Chondrogenic Fibroblasts | zgc:77056 |
| CIR11 | 1.9E-06 | 0.2314326 | 0.124 | 0.082 | 0.060243 | Chondrogenic Fibroblasts | CIR1 |
| fam3c3 | 2E-06 | 0.2450246 | 0.147 | 0.102 | 0.064963 | Chondrogenic Fibroblasts | fam3c |
| dnajb1a | 2.1E-06 | 0.2672267 | 0.1 | 0.063 | 0.068324 | Chondrogenic Fibroblasts | dnajb1a |
| mrps101 | 2.4E-06 | 0.2283875 | 0.157 | 0.11 | 0.079281 | Chondrogenic Fibroblasts | mrps10 |
| hgs | 2.4E-06 | 0.2646544 | 0.131 | 0.088 | 0.079628 | Chondrogenic Fibroblasts | hgs |
| serpine14 | 2.6E-06 | 0.2486682 | 0.843 | 0.76 | 0.083766 | Chondrogenic Fibroblasts | serpine1 |
| ankrd46a | 3.1E-06 | 0.3359474 | 0.109 | 0.072 | 0.10127 | Chondrogenic Fibroblasts | ankrd46a |
| zfand5a4 | 3.5E-06 | 0.3130391 | 0.247 | 0.188 | 0.113025 | Chondrogenic Fibroblasts | zfand5a |
| psmd91 | 3.6E-06 | 0.2457577 | 0.118 | 0.079 | 0.117132 | Chondrogenic Fibroblasts | psmd9 |
| trim8a2 | 3.7E-06 | 0.3260394 | 0.122 | 0.082 | 0.120177 | Chondrogenic Fibroblasts | trim8a |
| dele1 | 3.8E-06 | 0.3650519 | 0.106 | 0.069 | 0.123694 | Chondrogenic Fibroblasts | dele1 |
| mrpl542 | 3.9E-06 | 0.322895 | 0.134 | 0.092 | 0.125857 | Chondrogenic Fibroblasts | mrpl54 |
| mrps71 | 3.9E-06 | 0.3483769 | 0.116 | 0.078 | 0.126924 | Chondrogenic Fibroblasts | mrps7 |
| gtf2b3 | 4.7E-06 | 0.3133127 | 0.117 | 0.079 | 0.153487 | Chondrogenic Fibroblasts | gtf2b |
| dctn4 | 4.8E-06 | 0.2627595 | 0.111 | 0.073 | 0.154589 | Chondrogenic Fibroblasts | dctn4 |
| pdzrn3b2 | 5.3E-06 | 0.2039148 | 0.118 | 0.079 | 0.173451 | Chondrogenic Fibroblasts | pdzrn3b |
| arhgap42a2 | 5.5E-06 | 0.2095898 | 0.109 | 0.072 | 0.177671 | Chondrogenic Fibroblasts | arhgap42a |
| sephs21 | 6.6E-06 | 0.2725559 | 0.113 | 0.075 | 0.21593 | Chondrogenic Fibroblasts | sephs2 |
| shroom43 | 7.3E-06 | 0.2966503 | 0.182 | 0.133 | 0.235976 | Chondrogenic Fibroblasts | shroom4 |
| NDUFB12 | 7.3E-06 | 0.3685934 | 0.134 | 0.094 | 0.238724 | Chondrogenic Fibroblasts | NDUFB1 |
| zgc:560951 | 7.4E-06 | 0.2384091 | 0.134 | 0.093 | 0.240494 | Chondrogenic Fibroblasts | zgc:56095 |

| | | | | | | | |
|---|---|---|---|---|---|---|---|
| gpr108 | 7.9E-06 | 0.2188525 | 0.155 | 0.11 | 0.257796 | Chondrogenic Fibroblasts | gpr108 |
| itgb54 | 8E-06 | 0.2543221 | 0.123 | 0.084 | 0.260748 | Chondrogenic Fibroblasts | itgb5 |
| abcc6a4 | 8.1E-06 | 0.7532598 | 0.111 | 0.076 | 0.261876 | Chondrogenic Fibroblasts | abcc6a |
| ncln2 | 8.1E-06 | 0.3253295 | 0.108 | 0.072 | 0.262129 | Chondrogenic Fibroblasts | ncln |
| plekha82 | 8.5E-06 | 0.2854129 | 0.106 | 0.07 | 0.277965 | Chondrogenic Fibroblasts | plekha8 |
| hsdl22 | 1E-05 | 0.2023097 | 0.111 | 0.074 | 0.329803 | Chondrogenic Fibroblasts | hsdl2 |
| mydgf3 | 1.1E-05 | 0.2875959 | 0.117 | 0.08 | 0.364337 | Chondrogenic Fibroblasts | mydgf |
| derl22 | 1.2E-05 | 0.2654659 | 0.132 | 0.092 | 0.406088 | Chondrogenic Fibroblasts | derl2 |
| rcc21 | 1.3E-05 | 0.2347571 | 0.11 | 0.074 | 0.408619 | Chondrogenic Fibroblasts | rcc2 |
| taf102 | 1.4E-05 | 0.2819648 | 0.111 | 0.075 | 0.471334 | Chondrogenic Fibroblasts | taf10 |
| hbp13 | 1.5E-05 | 0.2123969 | 0.155 | 0.112 | 0.481136 | Chondrogenic Fibroblasts | hbp1 |
| tmub11 | 1.6E-05 | 0.3563126 | 0.103 | 0.069 | 0.507279 | Chondrogenic Fibroblasts | tmub1 |
| pin42 | 1.7E-05 | 0.2098164 | 0.197 | 0.148 | 0.541668 | Chondrogenic Fibroblasts | pin4 |
| eif2a1 | 1.8E-05 | 0.2979493 | 0.119 | 0.083 | 0.573737 | Chondrogenic Fibroblasts | eif2a |
| rai143 | 1.8E-05 | 0.2487747 | 0.124 | 0.086 | 0.595696 | Chondrogenic Fibroblasts | rai14 |
| tprb1 | 2.1E-05 | 0.2474471 | 0.143 | 0.103 | 0.688976 | Chondrogenic Fibroblasts | tprb |
| afap12 | 2.4E-05 | 0.2424148 | 0.105 | 0.071 | 0.767215 | Chondrogenic Fibroblasts | afap1 |
| plxna33 | 2.7E-05 | 0.2115866 | 0.126 | 0.088 | 0.888583 | Chondrogenic Fibroblasts | plxna3 |
| gatad2ab1 | 3.3E-05 | 0.2423455 | 0.104 | 0.071 | 1 | Chondrogenic Fibroblasts | gatad2ab |
| uqcrc2a2 | 3.4E-05 | 0.2128126 | 0.143 | 0.102 | 1 | Chondrogenic Fibroblasts | uqcrc2a |
| zdhhc8b2 | 3.6E-05 | 0.2071357 | 0.115 | 0.08 | 1 | Chondrogenic Fibroblasts | zdhhc8b |
| gpalpp1 | 3.6E-05 | 0.2911002 | 0.107 | 0.074 | 1 | Chondrogenic Fibroblasts | gpalpp1 |
| eif4g3a1 | 3.7E-05 | 0.2225599 | 0.1 | 0.067 | 1 | Chondrogenic Fibroblasts | eif4g3a |
| si:dkey-88p24.111 | 4.1E-05 | 0.2428559 | 0.116 | 0.081 | 1 | Chondrogenic Fibroblasts | si:dkey-88p24.11 |
| tbc1d231 | 4.7E-05 | 0.2432202 | 0.107 | 0.073 | 1 | Chondrogenic Fibroblasts | tbc1d23 |
| pxna2 | 4.9E-05 | 0.2292346 | 0.149 | 0.108 | 1 | Chondrogenic Fibroblasts | pxna |
| pkig3 | 5.1E-05 | 0.230344 | 0.16 | 0.118 | 1 | Chondrogenic Fibroblasts | pkig |
| rrp362 | 5.3E-05 | 0.2495961 | 0.121 | 0.086 | 1 | Chondrogenic Fibroblasts | rrp36 |
| smim152 | 6.5E-05 | 0.2022074 | 0.218 | 0.169 | 1 | Chondrogenic Fibroblasts | smim15 |
| svbp | 7.7E-05 | 0.2415896 | 0.101 | 0.07 | 1 | Chondrogenic Fibroblasts | svbp |
| snx51 | 8.2E-05 | 0.2515164 | 0.119 | 0.084 | 1 | Chondrogenic Fibroblasts | snx5 |
| clns1a1 | 8.3E-05 | 0.2836236 | 0.11 | 0.077 | 1 | Chondrogenic Fibroblasts | clns1a |
| il15l4 | 8.4E-05 | 0.4992514 | 0.159 | 0.118 | 1 | Chondrogenic Fibroblasts | il15l |
| oser13 | 8.5E-05 | 0.2702596 | 0.12 | 0.086 | 1 | Chondrogenic Fibroblasts | oser1 |
| pgam52 | 8.7E-05 | 0.2455395 | 0.106 | 0.074 | 1 | Chondrogenic Fibroblasts | pgam5 |
| fam204a | 0.00017 | 0.2416384 | 0.101 | 0.071 | 1 | Chondrogenic Fibroblasts | fam204a |
| sort1a4 | 0.00017 | 0.2499088 | 0.151 | 0.114 | 1 | Chondrogenic Fibroblasts | sort1a |
| luzp13 | 0.00056 | 0.2288617 | 0.218 | 0.174 | 1 | Chondrogenic Fibroblasts | luzp1 |
| fbln72 | 0.00086 | 0.7676739 | 0.13 | 0.1 | 1 | Chondrogenic Fibroblasts | fbln7 |
| vegfaa3 | 0.00887 | 0.2219806 | 0.143 | 0.117 | 1 | Chondrogenic Fibroblasts | vegfaa |
| nrn1a4 | 0 | 3.6091266 | 0.648 | 0.074 | 0 | DF 7 | nrn1a |
| cxcl143 | 0 | 3.0963259 | 0.798 | 0.235 | 0 | DF 7 | cxcl14 |
| si:ch211-106h4.124 | 0 | 1.5779964 | 0.963 | 0.445 | 0 | DF 7 | si:ch211-106h4.12 |
| pax3a | 0 | 5.3118852 | 0.492 | 0.017 | 0 | DF 7 | pax3a |
| rgs5a2 | 0 | 3.3633058 | 0.585 | 0.141 | 0 | DF 7 | rgs5a |
| barx15 | 0 | 2.6501028 | 0.553 | 0.117 | 0 | DF 7 | barx1 |
| rspo4 | 0 | 6.004486 | 0.413 | 0.008 | 0 | DF 7 | rspo4 |
| itm2ca | 0 | 4.4194549 | 0.157 | 0.009 | 0 | DF 7 | itm2ca |

| | | | | | | | |
|---|---|---|---|---|---|---|---|
| spock2 | 0 | 3.8917723 | 0.151 | 0.009 | 0 | DF 7 | spock2 |
| msrb2 | 0 | 5.9737913 | 0.111 | 0.002 | 0 | DF 7 | msrb2 |
| ldlrad2 | 0 | 4.9656512 | 0.106 | 0.004 | 0 | DF 7 | ldlrad2 |
| krt915 | 0 | 1.4677222 | 1 | 0.899 | 0 | DF 7 | krt91 |
| asip2b2 | 4E-301 | 2.7177839 | 0.49 | 0.126 | 1.4E-296 | DF 7 | asip2b |
| cpamd82 | 2E-285 | 2.7141174 | 0.311 | 0.054 | 7.4E-281 | DF 7 | cpamd8 |
| postna8 | 2E-275 | 1.357386 | 0.978 | 0.581 | 7E-271 | DF 7 | postna |
| si:ch211-251b21.15 | 2E-273 | 1.3417987 | 0.919 | 0.439 | 5.6E-269 | DF 7 | si:ch211-251b21.1 |
| s100b | 2E-254 | 4.3123933 | 0.108 | 0.007 | 6.4E-250 | DF 7 | s100b |
| irx1a | 5E-251 | 3.6087333 | 0.16 | 0.016 | 1.8E-246 | DF 7 | irx1a |
| tagln4 | 6E-240 | 1.9378739 | 0.857 | 0.531 | 2.1E-235 | DF 7 | tagln |
| krt84 | 1E-234 | 0.8142736 | 0.997 | 0.984 | 3.3E-230 | DF 7 | krt8 |
| lpl4 | 6E-229 | 2.2651297 | 0.407 | 0.105 | 1.9E-224 | DF 7 | lpl |
| slc22a25 | 1E-228 | 1.8481984 | 0.555 | 0.186 | 3.4E-224 | DF 7 | slc22a2 |
| tbx3a4 | 7E-222 | 1.883159 | 0.433 | 0.116 | 2.3E-217 | DF 7 | tbx3a |
| rbp46 | 8E-219 | 1.1280432 | 0.999 | 0.908 | 2.5E-214 | DF 7 | rbp4 |
| ntd55 | 1E-209 | 2.0722975 | 0.449 | 0.134 | 4.8E-205 | DF 7 | ntd5 |
| ccl25b5 | 2E-208 | 0.8449484 | 0.976 | 0.656 | 7.8E-204 | DF 7 | ccl25b |
| pln | 4E-208 | 2.1710739 | 0.228 | 0.039 | 1.2E-203 | DF 7 | pln |
| pnrc24 | 1E-206 | 1.0691228 | 0.978 | 0.879 | 3.8E-202 | DF 7 | pnrc2 |
| tbx152 | 7E-202 | 2.5749087 | 0.281 | 0.059 | 2.2E-197 | DF 7 | tbx15 |
| spock36 | 5E-195 | 1.0987178 | 0.832 | 0.402 | 1.6E-190 | DF 7 | spock3 |
| eif4ebp34 | 7E-195 | 2.1950209 | 0.477 | 0.166 | 2.2E-190 | DF 7 | eif4ebp3 |
| CABZ01092746.16 | 2E-191 | 1.0607304 | 0.962 | 0.703 | 5.5E-187 | DF 7 | CABZ01092746.1 |
| apof3 | 7E-185 | 2.2341198 | 0.279 | 0.061 | 2.4E-180 | DF 7 | apof |
| kera4 | 1E-184 | 1.6805762 | 0.488 | 0.164 | 3.5E-180 | DF 7 | kera |
| ptx3a6 | 2E-183 | 0.9949139 | 0.883 | 0.477 | 5E-179 | DF 7 | ptx3a |
| fgl14 | 1E-179 | 1.4950246 | 0.545 | 0.202 | 4.3E-175 | DF 7 | fgl1 |
| c4b2 | 1E-178 | 2.4217413 | 0.246 | 0.05 | 4.6E-174 | DF 7 | c4b |
| BX663503.35 | 7E-175 | 1.3310175 | 0.763 | 0.394 | 2.3E-170 | DF 7 | BX663503.3 |
| fhl2a2 | 8E-174 | 2.1394468 | 0.406 | 0.128 | 2.5E-169 | DF 7 | fhl2a |
| pcdh10b1 | 2E-172 | 2.5547377 | 0.212 | 0.039 | 5.1E-168 | DF 7 | pcdh10b |
| NPAS34 | 1E-171 | 2.3321894 | 0.312 | 0.08 | 3.5E-167 | DF 7 | NPAS3 |
| si:dkey-85k7.75 | 1E-169 | 1.4584751 | 0.63 | 0.272 | 3.7E-165 | DF 7 | si:dkey-85k7.7 |
| adgrl2a6 | 1E-164 | 1.788465 | 0.489 | 0.183 | 4E-160 | DF 7 | adgrl2a |
| pcdh17 | 3E-161 | 3.2465526 | 0.119 | 0.014 | 1.1E-156 | DF 7 | pcdh17 |
| fbln73 | 3E-157 | 1.6105432 | 0.338 | 0.094 | 1.1E-152 | DF 7 | fbln7 |
| ramp25 | 2E-153 | 1.9328103 | 0.402 | 0.135 | 6.7E-149 | DF 7 | ramp2 |
| hsp70l3 | 3E-153 | 1.7914253 | 0.613 | 0.294 | 1E-148 | DF 7 | hsp70l |
| sik15 | 2E-146 | 1.7168051 | 0.553 | 0.246 | 7.4E-142 | DF 7 | sik1 |
| FQ323156.16 | 1E-145 | 1.6212794 | 0.481 | 0.188 | 3.3E-141 | DF 7 | FQ323156.1 |
| CR383676.26 | 1E-143 | 0.5182568 | 1 | 1 | 3.2E-139 | DF 7 | CR383676.2 |
| ptgdsb.24 | 2E-142 | 0.7549021 | 0.962 | 0.66 | 6.7E-138 | DF 7 | ptgdsb.2 |
| ifitm17 | 6E-141 | 0.699728 | 0.989 | 0.785 | 1.9E-136 | DF 7 | ifitm1 |
| junba4 | 1E-139 | 1.0905348 | 0.943 | 0.844 | 3.2E-135 | DF 7 | junba |
| ckbb3 | 3E-139 | 1.6878244 | 0.477 | 0.192 | 9E-135 | DF 7 | ckbb |
| nfil3-66 | 3E-136 | 1.2467347 | 0.628 | 0.298 | 9.7E-132 | DF 7 | nfil3-6 |
| bhmt6 | 3E-135 | 0.8794643 | 0.945 | 0.702 | 1E-130 | DF 7 | bhmt |

| | | | | | | | |
|---|---|---|---|---|---|---|---|
| dll41 | 1E-133 | 2.371136 | 0.188 | 0.039 | 3.5E-129 | DF 7 | dll4 |
| vcanb3 | 3E-131 | 0.7694539 | 0.812 | 0.463 | 1.1E-126 | DF 7 | vcanb |
| f13a1b3 | 5E-128 | 0.3946189 | 0.607 | 0.287 | 1.5E-123 | DF 7 | f13a1b |
| hsp70.13 | 7E-128 | 1.6736306 | 0.631 | 0.334 | 2.2E-123 | DF 7 | hsp70.1 |
| glula5 | 8E-128 | 0.9428992 | 0.879 | 0.612 | 2.7E-123 | DF 7 | glula |
| actb23 | 6E-126 | 0.5731397 | 1 | 0.991 | 1.9E-121 | DF 7 | actb2 |
| ntn1a5 | 1E-125 | 1.6042689 | 0.348 | 0.115 | 3.8E-121 | DF 7 | ntn1a |
| si:dkey-164f24.26 | 2E-124 | 1.3192241 | 0.434 | 0.164 | 6.2E-120 | DF 7 | si:dkey-164f24.2 |
| wnt115 | 6E-124 | 1.3831334 | 0.579 | 0.277 | 2.1E-119 | DF 7 | wnt11 |
| hsp70.23 | 3E-123 | 1.5629141 | 0.75 | 0.487 | 1E-118 | DF 7 | hsp70.2 |
| plpp36 | 9E-122 | 1.0253746 | 0.756 | 0.447 | 2.8E-117 | DF 7 | plpp3 |
| mycb4 | 2E-120 | 1.2930817 | 0.836 | 0.659 | 7.6E-116 | DF 7 | mycb |
| cebpd6 | 1E-118 | 1.0546305 | 0.825 | 0.571 | 3.4E-114 | DF 7 | cebpd |
| lxn5 | 2E-115 | 0.8318217 | 0.905 | 0.697 | 5E-111 | DF 7 | lxn |
| FO818704.2 | 2E-115 | 2.8350695 | 0.108 | 0.016 | 5.3E-111 | DF 7 | FO818704.2 |
| vmo1a4 | 6E-115 | 0.6441784 | 0.967 | 0.694 | 2E-110 | DF 7 | vmo1a |
| sdc43 | 3E-114 | 1.2171065 | 0.861 | 0.7 | 1.1E-109 | DF 7 | sdc4 |
| c45 | 2E-110 | 1.1337212 | 0.473 | 0.201 | 7.3E-106 | DF 7 | c4 |
| im:71523485 | 2E-110 | 1.5893711 | 0.489 | 0.23 | 7.5E-106 | DF 7 | im:7152348 |
| cxcl18b4 | 6E-110 | 1.0310261 | 0.894 | 0.697 | 2.1E-105 | DF 7 | cxcl18b |
| twist35 | 6E-109 | 1.3867732 | 0.519 | 0.247 | 2E-104 | DF 7 | twist3 |
| pcdh115 | 2E-104 | 1.8047063 | 0.244 | 0.071 | 4.9E-100 | DF 7 | pcdh11 |
| twist1a4 | 2E-103 | 1.0761926 | 0.703 | 0.432 | 8E-99 | DF 7 | twist1a |
| tmsb4x2 | 6E-103 | 0.5612542 | 0.999 | 0.992 | 2.03E-98 | DF 7 | tmsb4x |
| ttyh3a3 | 2E-102 | 1.9196218 | 0.279 | 0.092 | 5.42E-98 | DF 7 | ttyh3a |
| tob1a4 | 3E-102 | 0.9710183 | 0.807 | 0.605 | 9.93E-98 | DF 7 | tob1a |
| bco2a2 | 2E-101 | 1.5892367 | 0.366 | 0.144 | 5.96E-97 | DF 7 | bco2a |
| eif13 | 1E-100 | 0.9432323 | 0.905 | 0.828 | 3.85E-96 | DF 7 | eif1 |
| cavin2b4 | 2E-100 | 0.9980507 | 0.764 | 0.525 | 6.32E-96 | DF 7 | cavin2b |
| efnb2a3 | 8E-100 | 1.6441315 | 0.363 | 0.145 | 2.46E-95 | DF 7 | efnb2a |
| cxcl12a7 | 8E-100 | 0.5740975 | 0.947 | 0.676 | 2.56E-95 | DF 7 | cxcl12a |
| ccni2 | 4E-99 | 0.9685208 | 0.776 | 0.59 | 1.29E-94 | DF 7 | ccni |
| actb12 | 7.3E-98 | 0.7466682 | 0.949 | 0.865 | 2.38E-93 | DF 7 | actb1 |
| nsmfb | 5.2E-97 | 2.6590885 | 0.102 | 0.016 | 1.69E-92 | DF 7 | nsmfb |
| tgfbi6 | 2E-96 | 1.0908589 | 0.442 | 0.194 | 6.53E-92 | DF 7 | tgfbi |
| pcdh11.1 | 2.5E-96 | 2.164232 | 0.171 | 0.042 | 8.21E-92 | DF 7 | pcdh11.1 |
| tbx2a1 | 8.9E-96 | 2.240554 | 0.207 | 0.058 | 2.91E-91 | DF 7 | tbx2a |
| zgc:1626125 | 7.1E-95 | 1.6262317 | 0.306 | 0.112 | 2.32E-90 | DF 7 | zgc:162612 |
| tmem200b4 | 1.4E-94 | 1.8136172 | 0.268 | 0.09 | 4.56E-90 | DF 7 | tmem200b |
| efna1a2 | 2.9E-94 | 1.5766785 | 0.395 | 0.172 | 9.4E-90 | DF 7 | efna1a |
| egfl64 | 1.7E-90 | 1.8460624 | 0.192 | 0.052 | 5.52E-86 | DF 7 | egfl6 |
| spink2.13 | 3.4E-90 | 1.5147901 | 0.324 | 0.124 | 1.12E-85 | DF 7 | spink2.1 |
| hsp70.33 | 6.7E-90 | 1.3742232 | 0.749 | 0.55 | 2.17E-85 | DF 7 | hsp70.3 |
| aoc25 | 1.2E-89 | 1.1297458 | 0.487 | 0.235 | 3.86E-85 | DF 7 | aoc2 |
| rsrp14 | 5.2E-89 | 1.0594421 | 0.73 | 0.538 | 1.7E-84 | DF 7 | rsrp1 |
| atp1b4 | 2.8E-88 | 2.1456049 | 0.129 | 0.027 | 8.98E-84 | DF 7 | atp1b4 |
| timp4.34 | 1.5E-87 | 0.4879572 | 0.745 | 0.433 | 4.87E-83 | DF 7 | timp4.3 |
| emilin2b1 | 7E-87 | 1.6622936 | 0.217 | 0.065 | 2.27E-82 | DF 7 | emilin2b |

| | | | | | | | |
|---|---|---|---|---|---|---|---|
| fstb7 | 1E-86 | 0.8881736 | 0.637 | 0.355 | 3.33E-82 | DF 7 | fstb |
| tob1b5 | 2.5E-83 | 0.7142731 | 0.874 | 0.698 | 8.13E-79 | DF 7 | tob1b |
| fmoda4 | 2.8E-83 | 1.0996553 | 0.407 | 0.178 | 9.11E-79 | DF 7 | fmoda |
| fosl1a3 | 6E-82 | 1.2021904 | 0.642 | 0.433 | 1.95E-77 | DF 7 | fosl1a |
| boc5 | 2.3E-81 | 1.209849 | 0.585 | 0.368 | 7.42E-77 | DF 7 | boc |
| csrp22 | 7.1E-80 | 1.7789135 | 0.291 | 0.115 | 2.31E-75 | DF 7 | csrp2 |
| fam169aa5 | 7.9E-80 | 0.8162445 | 0.67 | 0.397 | 2.56E-75 | DF 7 | fam169aa |
| slc1a3a4 | 1.2E-79 | 1.5097107 | 0.339 | 0.146 | 4E-75 | DF 7 | slc1a3a |
| hsp90aa1.23 | 3.1E-79 | 1.2751473 | 0.777 | 0.596 | 1.01E-74 | DF 7 | hsp90aa1.2 |
| zgc:565857 | 3E-78 | 0.9775597 | 0.575 | 0.322 | 9.79E-74 | DF 7 | zgc:56585 |
| lmod1b2 | 3.7E-78 | 1.9052858 | 0.255 | 0.093 | 1.2E-73 | DF 7 | lmod1b |
| igsf21a2 | 1.3E-77 | 1.891574 | 0.188 | 0.056 | 4.22E-73 | DF 7 | igsf21a |
| ndrg44 | 2.4E-77 | 1.2877835 | 0.35 | 0.15 | 7.82E-73 | DF 7 | ndrg4 |
| cyp26b12 | 3.8E-76 | 1.1295663 | 0.607 | 0.363 | 1.24E-71 | DF 7 | cyp26b1 |
| lhfpl64 | 5.2E-75 | 0.9896494 | 0.63 | 0.412 | 1.68E-70 | DF 7 | lhfpl6 |
| emilin1a6 | 3.3E-74 | 1.4574276 | 0.251 | 0.09 | 1.07E-69 | DF 7 | emilin1a |
| BX323087.1 | 7.1E-74 | 2.3772953 | 0.101 | 0.02 | 2.31E-69 | DF 7 | BX323087.1 |
| gpm6ab3 | 8.8E-74 | 1.6487374 | 0.238 | 0.085 | 2.87E-69 | DF 7 | gpm6ab |
| btg24 | 3.6E-73 | 1.0180857 | 0.826 | 0.696 | 1.18E-68 | DF 7 | btg2 |
| si:ch211-195b13.14 | 6E-73 | 1.2054313 | 0.455 | 0.238 | 1.94E-68 | DF 7 | si:ch211-195b13.1 |
| tmem198b3 | 1.3E-72 | 1.3904092 | 0.384 | 0.185 | 4.27E-68 | DF 7 | tmem198b |
| btg14 | 1.9E-72 | 0.6982691 | 0.895 | 0.813 | 6.3E-68 | DF 7 | btg1 |
| hhip4 | 1.2E-71 | 1.6124691 | 0.237 | 0.085 | 3.81E-67 | DF 7 | hhip |
| mcl1a5 | 1.4E-71 | 0.7853067 | 0.875 | 0.769 | 4.69E-67 | DF 7 | mcl1a |
| rhoq4 | 3E-70 | 1.6306042 | 0.245 | 0.092 | 9.83E-66 | DF 7 | rhoq |
| b2ml5 | 1.7E-69 | 0.4770885 | 0.969 | 0.896 | 5.56E-65 | DF 7 | b2ml |
| ZFP36L1.14 | 4.9E-69 | 0.6584808 | 0.953 | 0.872 | 1.6E-64 | DF 7 | ZFP36L1.1 |
| myl9b2 | 4.8E-68 | 1.1008605 | 0.617 | 0.421 | 1.55E-63 | DF 7 | myl9b |
| metrn4 | 6.5E-68 | 1.5274441 | 0.277 | 0.113 | 2.11E-63 | DF 7 | metrn |
| zeb2a4 | 8.7E-68 | 1.0367523 | 0.556 | 0.336 | 2.82E-63 | DF 7 | zeb2a |
| vim5 | 3.1E-67 | 0.6281745 | 0.618 | 0.364 | 1.01E-62 | DF 7 | vim |
| snai1a3 | 4.3E-67 | 1.1847118 | 0.477 | 0.268 | 1.4E-62 | DF 7 | snai1a |
| ephb2b4 | 1.2E-66 | 1.2908819 | 0.388 | 0.195 | 3.78E-62 | DF 7 | ephb2b |
| tpst14 | 6.1E-66 | 1.3247665 | 0.357 | 0.172 | 1.98E-61 | DF 7 | tpst1 |
| tpm4a2 | 2.1E-65 | 0.763184 | 0.744 | 0.549 | 6.82E-61 | DF 7 | tpm4a |
| si:ch73-386h18.16 | 2.7E-64 | 1.3688641 | 0.318 | 0.144 | 8.63E-60 | DF 7 | si:ch73-386h18.1 |
| id34 | 3.5E-63 | 0.9429723 | 0.69 | 0.508 | 1.13E-58 | DF 7 | id3 |
| hspa8b4 | 4.5E-63 | 1.531665 | 0.236 | 0.09 | 1.45E-58 | DF 7 | hspa8b |
| efna1b2 | 4.9E-63 | 1.3304709 | 0.312 | 0.139 | 1.59E-58 | DF 7 | efna1b |
| fgfr1b5 | 5E-63 | 1.6497021 | 0.246 | 0.098 | 1.62E-58 | DF 7 | fgfr1b |
| agrn3 | 1.3E-62 | 1.2327905 | 0.368 | 0.18 | 4.13E-58 | DF 7 | agrn |
| ptn2 | 1.9E-62 | 1.5310938 | 0.154 | 0.046 | 6.23E-58 | DF 7 | ptn |
| zgc:1943985 | 5.4E-62 | 0.7004862 | 0.623 | 0.378 | 1.77E-57 | DF 7 | zgc:194398 |
| BX663503.13 | 7.4E-62 | 1.6456917 | 0.188 | 0.064 | 2.42E-57 | DF 7 | BX663503.1 |
| cnn23 | 7.5E-62 | 1.071407 | 0.628 | 0.441 | 2.43E-57 | DF 7 | cnn2 |
| mgst1.24 | 9.2E-62 | 0.9274875 | 0.515 | 0.296 | 2.98E-57 | DF 7 | mgst1.2 |
| qdpra6 | 1.1E-61 | 0.6047669 | 0.677 | 0.42 | 3.71E-57 | DF 7 | qdpra |
| tnfsf104 | 2.4E-61 | 1.4307645 | 0.243 | 0.095 | 7.71E-57 | DF 7 | tnfsf10 |

| | | | | | | | |
|---|---|---|---|---|---|---|---|
| slc6a171 | 5.4E-61 | 1.8575001 | 0.145 | 0.042 | 1.77E-56 | DF 7 | slc6a17 |
| rgs44 | 5.7E-61 | 1.5044989 | 0.194 | 0.067 | 1.85E-56 | DF 7 | rgs4 |
| slc1a55 | 1.7E-60 | 1.0864463 | 0.453 | 0.253 | 5.38E-56 | DF 7 | slc1a5 |
| junbb4 | 2.1E-60 | 0.7863033 | 0.761 | 0.598 | 6.7E-56 | DF 7 | junbb |
| meis1b4 | 3.4E-60 | 0.9616822 | 0.531 | 0.319 | 1.09E-55 | DF 7 | meis1b |
| ca64 | 4.2E-59 | 0.4411979 | 0.606 | 0.341 | 1.37E-54 | DF 7 | ca6 |
| sik2b3 | 8.8E-59 | 1.1983132 | 0.389 | 0.203 | 2.87E-54 | DF 7 | sik2b |
| smarcd23 | 1.6E-58 | 1.2230997 | 0.335 | 0.161 | 5.32E-54 | DF 7 | smarcd2 |
| gbe1a5 | 2.1E-57 | 1.2919968 | 0.26 | 0.109 | 6.83E-53 | DF 7 | gbe1a |
| CABZ01068358.14 | 3.6E-57 | 1.2153514 | 0.385 | 0.204 | 1.17E-52 | DF 7 | CABZ01068358.1 |
| lamc33 | 6.4E-57 | 1.2970808 | 0.272 | 0.117 | 2.08E-52 | DF 7 | lamc3 |
| ptgdsb.16 | 7.3E-57 | 0.9651248 | 0.65 | 0.459 | 2.37E-52 | DF 7 | ptgdsb.1 |
| mab21l23 | 1.4E-56 | 0.9515692 | 0.53 | 0.327 | 4.4E-52 | DF 7 | mab21l2 |
| tgfb1a4 | 1.6E-56 | 1.0520733 | 0.465 | 0.269 | 5.34E-52 | DF 7 | tgfb1a |
| irs2b5 | 4E-56 | 1.1401437 | 0.413 | 0.226 | 1.3E-51 | DF 7 | irs2b |
| hpdb7 | 4.2E-56 | 0.2734041 | 0.804 | 0.502 | 1.36E-51 | DF 7 | hpdb |
| tmem59l3 | 8.5E-56 | 1.3638164 | 0.247 | 0.103 | 2.75E-51 | DF 7 | tmem59l |
| lhx61 | 1E-55 | 1.7583287 | 0.146 | 0.045 | 3.41E-51 | DF 7 | lhx6 |
| mknk2b4 | 2.5E-55 | 0.6891142 | 0.764 | 0.604 | 8.04E-51 | DF 7 | mknk2b |
| hist1h4l.184 | 3.7E-55 | 1.2256882 | 0.27 | 0.117 | 1.21E-50 | DF 7 | hist1h4l.18 |
| cited4b4 | 1.2E-54 | 1.1381635 | 0.31 | 0.146 | 3.78E-50 | DF 7 | cited4b |
| slco2b15 | 1.2E-53 | 0.8976879 | 0.41 | 0.217 | 3.77E-49 | DF 7 | slco2b1 |
| ms4a17a.96 | 1.5E-53 | 1.0739093 | 0.394 | 0.212 | 4.79E-49 | DF 7 | ms4a17a.9 |
| epas1a2 | 7.7E-53 | 1.3245611 | 0.176 | 0.062 | 2.51E-48 | DF 7 | epas1a |
| cfp6 | 1.6E-52 | 0.7426984 | 0.553 | 0.335 | 5.12E-48 | DF 7 | cfp |
| fstl34 | 2E-52 | 1.2878297 | 0.495 | 0.318 | 6.4E-48 | DF 7 | fstl3 |
| enpp24 | 8.5E-52 | 0.6313642 | 0.658 | 0.44 | 2.76E-47 | DF 7 | enpp2 |
| si:ch73-237c6.16 | 1.4E-51 | 0.7637406 | 0.571 | 0.361 | 4.62E-47 | DF 7 | si:ch73-237c6.1 |
| sept9b2 | 2.8E-51 | 1.6307725 | 0.169 | 0.06 | 9.06E-47 | DF 7 | sept9b |
| hyal2b6 | 3.2E-51 | 1.1762616 | 0.364 | 0.193 | 1.03E-46 | DF 7 | hyal2b |
| ubb4 | 1.8E-50 | 0.7409803 | 0.784 | 0.679 | 5.98E-46 | DF 7 | ubb |
| si:ch1073-291c23.25 | 2.6E-50 | 0.7150443 | 0.658 | 0.47 | 8.35E-46 | DF 7 | si:ch1073-291c23.2 |
| tmsb13 | 3.1E-50 | 0.5634076 | 0.657 | 0.445 | 1.01E-45 | DF 7 | tmsb1 |
| gadd45ga5 | 4.2E-50 | 1.471927 | 0.442 | 0.273 | 1.37E-45 | DF 7 | gadd45ga |
| bzw1b3 | 1.3E-49 | 1.3512357 | 0.625 | 0.488 | 4.24E-45 | DF 7 | bzw1b |
| rps27.21 | 1.5E-49 | 0.3837353 | 0.955 | 0.953 | 4.72E-45 | DF 7 | rps27.2 |
| ackr3b3 | 2.2E-49 | 0.8604334 | 0.262 | 0.115 | 7.03E-45 | DF 7 | ackr3b |
| gstr4 | 4.3E-49 | 0.7714706 | 0.595 | 0.412 | 1.4E-44 | DF 7 | gstr |
| cbsb6 | 1.4E-48 | 1.2656573 | 0.283 | 0.136 | 4.53E-44 | DF 7 | cbsb |
| gpm6aa1 | 1.5E-48 | 1.7534156 | 0.157 | 0.055 | 4.8E-44 | DF 7 | gpm6aa |
| tubb4b3 | 1.5E-48 | 0.4839395 | 0.883 | 0.808 | 4.9E-44 | DF 7 | tubb4b |
| ptmaa3 | 2.8E-48 | 0.5635159 | 0.91 | 0.835 | 9.03E-44 | DF 7 | ptmaa |
| tcf125 | 3.4E-48 | 0.5876735 | 0.711 | 0.518 | 1.1E-43 | DF 7 | tcf12 |
| adamts15b.15 | 5E-48 | 1.292262 | 0.218 | 0.09 | 1.63E-43 | DF 7 | adamts15b.1 |
| sgk14 | 1.6E-47 | 0.8880098 | 0.545 | 0.363 | 5.17E-43 | DF 7 | sgk1 |
| glulb4 | 2.4E-47 | 0.6186888 | 0.735 | 0.566 | 7.84E-43 | DF 7 | glulb |
| slc40a14 | 2.9E-47 | 0.9763202 | 0.369 | 0.198 | 9.32E-43 | DF 7 | slc40a1 |
| cotl11 | 3.2E-47 | 1.3792447 | 0.227 | 0.099 | 1.05E-42 | DF 7 | cotl1 |

| | | | | | | | |
|---|---|---|---|---|---|---|---|
| ENSDARG00000007589 | 4E-47 | 1.3758828 | 0.202 | 0.082 | 1.29E-42 | DF 7 | ENSDARG00000075 |
| ppp1r1c4 | 8.7E-47 | 0.8508617 | 0.274 | 0.128 | 2.81E-42 | DF 7 | ppp1r1c |
| suco5 | 9.3E-47 | 1.2482129 | 0.323 | 0.17 | 3.03E-42 | DF 7 | suco |
| slc20a1b3 | 1.7E-46 | 1.1996719 | 0.456 | 0.291 | 5.54E-42 | DF 7 | slc20a1b |
| sh3bgrl4 | 2E-46 | 1.1311136 | 0.387 | 0.223 | 6.51E-42 | DF 7 | sh3bgrl |
| ccdc1253 | 2E-46 | 1.4267234 | 0.185 | 0.072 | 6.58E-42 | DF 7 | ccdc125 |
| ddx58 | 4.1E-46 | 1.0326565 | 0.693 | 0.596 | 1.32E-41 | DF 7 | ddx5 |
| ddx64 | 4.8E-46 | 0.8518715 | 0.518 | 0.344 | 1.57E-41 | DF 7 | ddx6 |
| rbfox3a | 8.7E-46 | 1.8775985 | 0.116 | 0.035 | 2.82E-41 | DF 7 | rbfox3a |
| efna2a | 3.5E-45 | 1.6069919 | 0.132 | 0.043 | 1.13E-40 | DF 7 | efna2a |
| col18a1a3 | 4.3E-45 | 0.6946869 | 0.304 | 0.15 | 1.39E-40 | DF 7 | col18a1a |
| ephb3a4 | 5.1E-45 | 1.3468905 | 0.232 | 0.104 | 1.65E-40 | DF 7 | ephb3a |
| selenop3 | 7.9E-45 | 0.4008342 | 0.899 | 0.737 | 2.58E-40 | DF 7 | selenop |
| usp2a2 | 2.1E-44 | 1.5161595 | 0.177 | 0.069 | 6.91E-40 | DF 7 | usp2a |
| gstm.16 | 5.6E-44 | 0.5839088 | 0.745 | 0.594 | 1.82E-39 | DF 7 | gstm.1 |
| hpgd3 | 1.4E-43 | 1.5303953 | 0.149 | 0.054 | 4.67E-39 | DF 7 | hpgd |
| si:ch211-232b12.52 | 1.8E-43 | 1.1906875 | 0.297 | 0.153 | 5.78E-39 | DF 7 | si:ch211-232b12.5 |
| socs1a5 | 2.4E-43 | 1.2337782 | 0.254 | 0.121 | 7.78E-39 | DF 7 | socs1a |
| cbx7a4 | 3.2E-43 | 1.0029573 | 0.511 | 0.343 | 1.03E-38 | DF 7 | cbx7a |
| LO018188.16 | 3.2E-43 | 0.6335776 | 0.649 | 0.474 | 1.05E-38 | DF 7 | LO018188.1 |
| xylt11 | 3.3E-42 | 1.6845955 | 0.133 | 0.046 | 1.08E-37 | DF 7 | xylt1 |
| pou3f3b3 | 1.4E-41 | 1.4870134 | 0.147 | 0.054 | 4.42E-37 | DF 7 | pou3f3b |
| fam43a5 | 1.4E-41 | 1.4500508 | 0.169 | 0.066 | 4.56E-37 | DF 7 | fam43a |
| tie1 | 1.7E-41 | 1.7655179 | 0.106 | 0.032 | 5.62E-37 | DF 7 | tie1 |
| loxl15 | 2.2E-41 | 1.0006891 | 0.355 | 0.198 | 7.06E-37 | DF 7 | loxl1 |
| foxd12 | 7.1E-41 | 1.3516188 | 0.167 | 0.065 | 2.31E-36 | DF 7 | foxd1 |
| hist2h2l5 | 1E-40 | 0.8214783 | 0.539 | 0.371 | 3.31E-36 | DF 7 | hist2h2l |
| higd1a5 | 1.2E-40 | 0.8490565 | 0.598 | 0.455 | 3.84E-36 | DF 7 | higd1a |
| gpr1434 | 2.6E-40 | 1.4442757 | 0.17 | 0.068 | 8.32E-36 | DF 7 | gpr143 |
| hgd5 | 3E-40 | 0.4573838 | 0.636 | 0.425 | 9.78E-36 | DF 7 | hgd |
| ism2a3 | 3.3E-40 | 1.3215154 | 0.14 | 0.05 | 1.07E-35 | DF 7 | ism2a |
| flna3 | 3.8E-40 | 0.8083551 | 0.682 | 0.54 | 1.23E-35 | DF 7 | flna |
| atf33 | 8.1E-40 | 0.875367 | 0.504 | 0.342 | 2.65E-35 | DF 7 | atf3 |
| si:ch211-262i1.43 | 8.7E-40 | 1.0341295 | 0.189 | 0.078 | 2.82E-35 | DF 7 | si:ch211-262i1.4 |
| tiam1a1 | 8.7E-40 | 1.6315836 | 0.154 | 0.06 | 2.82E-35 | DF 7 | tiam1a |
| meis2a.14 | 1.1E-39 | 1.0362344 | 0.265 | 0.131 | 3.69E-35 | DF 7 | meis2a.1 |
| tfap2c5 | 1.2E-39 | 0.9834472 | 0.342 | 0.193 | 3.93E-35 | DF 7 | tfap2c |
| sept9a4 | 5.2E-39 | 1.0127606 | 0.363 | 0.213 | 1.68E-34 | DF 7 | sept9a |
| fosab4 | 6.3E-39 | 0.6037383 | 0.948 | 0.894 | 2.05E-34 | DF 7 | fosab |
| tshz3b2 | 3.4E-38 | 1.397523 | 0.154 | 0.06 | 1.11E-33 | DF 7 | tshz3b |
| camk2n1a3 | 6.1E-38 | 1.3522733 | 0.187 | 0.081 | 1.99E-33 | DF 7 | camk2n1a |
| ednraa5 | 5.3E-37 | 0.9038982 | 0.361 | 0.21 | 1.72E-32 | DF 7 | ednraa |
| pim24 | 5.3E-37 | 0.9522632 | 0.372 | 0.223 | 1.73E-32 | DF 7 | pim2 |
| gstp16 | 6.3E-37 | 0.4791398 | 0.781 | 0.641 | 2.04E-32 | DF 7 | gstp1 |
| aplp12 | 1E-36 | 1.4351538 | 0.118 | 0.041 | 3.38E-32 | DF 7 | aplp1 |
| rbpms2a1 | 2.5E-36 | 1.07802 | 0.256 | 0.131 | 8.19E-32 | DF 7 | rbpms2a |
| agmo3 | 3E-36 | 1.2977968 | 0.184 | 0.08 | 9.75E-32 | DF 7 | agmo |
| tagln23 | 4.2E-36 | 0.5133712 | 0.793 | 0.663 | 1.36E-31 | DF 7 | tagln2 |

| | | | | | | | |
|---|---|---|---|---|---|---|---|
| klf11b2 | 4.6E-36 | 0.9285992 | 0.383 | 0.234 | 1.51E-31 | DF 7 | klf11b |
| hmgb3a3 | 7.3E-36 | 1.0432364 | 0.233 | 0.113 | 2.36E-31 | DF 7 | hmgb3a |
| tspan354 | 2E-35 | 1.2591455 | 0.24 | 0.121 | 6.48E-31 | DF 7 | tspan35 |
| mhc1zba4 | 2.6E-35 | 0.5583004 | 0.711 | 0.581 | 8.43E-31 | DF 7 | mhc1zba |
| iqsec1b3 | 2.6E-35 | 0.6921789 | 0.5 | 0.341 | 8.58E-31 | DF 7 | iqsec1b |
| si:ch211-191i18.21 | 3.1E-35 | 1.2793171 | 0.108 | 0.036 | 1.01E-30 | DF 7 | si:ch211-191i18.2 |
| gadd45ba2 | 3.4E-35 | 0.668344 | 0.886 | 0.802 | 1.11E-30 | DF 7 | gadd45ba |
| dnajb1b4 | 5E-35 | 0.6540565 | 0.615 | 0.461 | 1.64E-30 | DF 7 | dnajb1b |
| sema3aa4 | 8E-35 | 1.1178522 | 0.221 | 0.106 | 2.61E-30 | DF 7 | sema3aa |
| si:ch211-63p21.12 | 1.8E-34 | 1.4308236 | 0.13 | 0.049 | 5.79E-30 | DF 7 | si:ch211-63p21.1 |
| si:dkey-195m11.85 | 1.8E-34 | 1.1164466 | 0.245 | 0.125 | 5.85E-30 | DF 7 | si:dkey-195m11.8 |
| arhgap101 | 2E-34 | 1.0474527 | 0.272 | 0.145 | 6.42E-30 | DF 7 | arhgap10 |
| atn1 | 7.5E-34 | 1.381716 | 0.12 | 0.044 | 2.43E-29 | DF 7 | atn1 |
| gch15 | 1.3E-33 | 0.93908 | 0.258 | 0.133 | 4.33E-29 | DF 7 | gch1 |
| nr1d2a3 | 1.5E-33 | 0.915372 | 0.389 | 0.249 | 4.91E-29 | DF 7 | nr1d2a |
| nfil3-52 | 4.1E-33 | 1.1584865 | 0.256 | 0.136 | 1.34E-28 | DF 7 | nfil3-5 |
| h2ax14 | 4.3E-33 | 0.5674731 | 0.742 | 0.626 | 1.4E-28 | DF 7 | h2ax1 |
| fgfr24 | 4.6E-33 | 0.660015 | 0.502 | 0.345 | 1.49E-28 | DF 7 | fgfr2 |
| srsf5a3 | 4.9E-33 | 0.556378 | 0.703 | 0.577 | 1.59E-28 | DF 7 | srsf5a |
| fgfr1a5 | 6.6E-33 | 1.2306468 | 0.274 | 0.154 | 2.15E-28 | DF 7 | fgfr1a |
| nfe2l2a6 | 6.7E-33 | 0.650843 | 0.499 | 0.341 | 2.18E-28 | DF 7 | nfe2l2a |
| si:dkey-51e6.15 | 7.4E-33 | 0.8173551 | 0.4 | 0.255 | 2.42E-28 | DF 7 | si:dkey-51e6.1 |
| ablim1b5 | 8.9E-33 | 0.6291327 | 0.411 | 0.253 | 2.89E-28 | DF 7 | ablim1b |
| crabp2b4 | 2.1E-32 | 0.4484667 | 0.595 | 0.416 | 6.97E-28 | DF 7 | crabp2b |
| cadm44 | 2.8E-32 | 1.2181989 | 0.168 | 0.074 | 9.25E-28 | DF 7 | cadm4 |
| tnfrsf194 | 7.2E-32 | 1.1711327 | 0.214 | 0.106 | 2.36E-27 | DF 7 | tnfrsf19 |
| clcf12 | 1.1E-31 | 1.3144504 | 0.125 | 0.048 | 3.73E-27 | DF 7 | clcf1 |
| gpatch83 | 1.1E-31 | 0.6805117 | 0.48 | 0.326 | 3.74E-27 | DF 7 | gpatch8 |
| wnt5b3 | 1.7E-31 | 1.1146962 | 0.214 | 0.108 | 5.41E-27 | DF 7 | wnt5b |
| rtn32 | 4.8E-31 | 0.8651471 | 0.36 | 0.224 | 1.55E-26 | DF 7 | rtn3 |
| ehbp1l1a4 | 7E-31 | 0.9544755 | 0.195 | 0.092 | 2.28E-26 | DF 7 | ehbp1l1a |
| atf4b6 | 7.1E-31 | 0.5077833 | 0.746 | 0.632 | 2.31E-26 | DF 7 | atf4b |
| mlip2 | 8.1E-31 | 1.3495988 | 0.162 | 0.072 | 2.62E-26 | DF 7 | mlip |
| etv5a5 | 8.8E-31 | 1.3763408 | 0.203 | 0.102 | 2.88E-26 | DF 7 | etv5a |
| rplp22 | 9.2E-31 | 0.2487927 | 0.995 | 0.987 | 2.99E-26 | DF 7 | rplp2 |
| hmgb1a4 | 1.3E-30 | 0.5428812 | 0.678 | 0.562 | 4.25E-26 | DF 7 | hmgb1a |
| btf34 | 1.5E-30 | 0.3761578 | 0.909 | 0.875 | 4.98E-26 | DF 7 | btf3 |
| si:ch211-286o17.14 | 6.1E-30 | 0.984416 | 0.342 | 0.214 | 1.99E-25 | DF 7 | si:ch211-286o17.1 |
| gas1b5 | 1.3E-29 | 0.997846 | 0.318 | 0.193 | 4.28E-25 | DF 7 | gas1b |
| slc16a106 | 1.5E-29 | 1.127972 | 0.241 | 0.131 | 4.78E-25 | DF 7 | slc16a10 |
| ak45 | 2.1E-29 | 0.8218408 | 0.31 | 0.182 | 6.74E-25 | DF 7 | ak4 |
| hhla2a.14 | 2.2E-29 | 1.2567944 | 0.171 | 0.08 | 7.26E-25 | DF 7 | hhla2a.1 |
| prrx1b5 | 2.3E-29 | 0.6283676 | 0.523 | 0.377 | 7.56E-25 | DF 7 | prrx1b |
| ccnl1a4 | 2.3E-29 | 0.9002906 | 0.388 | 0.26 | 7.59E-25 | DF 7 | ccnl1a |
| dedd15 | 2.9E-29 | 0.570224 | 0.683 | 0.549 | 9.27E-25 | DF 7 | dedd1 |
| heyl4 | 5.2E-29 | 1.310369 | 0.156 | 0.07 | 1.69E-24 | DF 7 | heyl |
| marcksl1b5 | 9.5E-29 | 0.5090285 | 0.396 | 0.251 | 3.09E-24 | DF 7 | marcksl1b |
| ptk7a6 | 1.2E-28 | 0.6824486 | 0.451 | 0.312 | 3.99E-24 | DF 7 | ptk7a |

| | | | | | | | |
|---|---|---|---|---|---|---|---|
| hspb14 | 1.3E-28 | 1.104393 | 0.3 | 0.177 | 4.18E-24 | DF 7 | hspb1 |
| UBB3 | 1.4E-28 | 0.6604531 | 0.667 | 0.56 | 4.63E-24 | DF 7 | UBB |
| id15 | 1.4E-28 | 0.5290254 | 0.771 | 0.639 | 4.67E-24 | DF 7 | id1 |
| slmapb1 | 2E-28 | 1.2797632 | 0.121 | 0.048 | 6.67E-24 | DF 7 | slmapb |
| romo12 | 2.4E-28 | 0.4569205 | 0.788 | 0.705 | 7.74E-24 | DF 7 | romo1 |
| irx5a3 | 3.2E-28 | 1.2431048 | 0.181 | 0.089 | 1.05E-23 | DF 7 | irx5a |
| arrdc3b | 3.9E-28 | 1.5843084 | 0.106 | 0.04 | 1.28E-23 | DF 7 | arrdc3b |
| pde3b3 | 4E-28 | 1.0017068 | 0.224 | 0.119 | 1.29E-23 | DF 7 | pde3b |
| metrnla3 | 4.4E-28 | 0.8079104 | 0.447 | 0.313 | 1.43E-23 | DF 7 | metrnla |
| aldob7 | 6.4E-28 | 0.5966141 | 0.455 | 0.308 | 2.07E-23 | DF 7 | aldob |
| dop1a3 | 1.2E-27 | 1.1488274 | 0.162 | 0.075 | 3.76E-23 | DF 7 | dop1a |
| mylka3 | 1.3E-27 | 1.01473 | 0.206 | 0.106 | 4.33E-23 | DF 7 | mylka |
| crhr13 | 1.5E-27 | 1.0883742 | 0.165 | 0.077 | 4.88E-23 | DF 7 | crhr1 |
| sept7a2 | 1.6E-27 | 0.7005678 | 0.489 | 0.359 | 5.29E-23 | DF 7 | sept7a |
| nr4a14 | 1.7E-27 | 1.4093906 | 0.194 | 0.1 | 5.63E-23 | DF 7 | nr4a1 |
| tmtc14 | 1.8E-27 | 0.9524703 | 0.219 | 0.114 | 5.73E-23 | DF 7 | tmtc1 |
| hist1h4l.34 | 1.8E-27 | 0.9454113 | 0.253 | 0.141 | 5.87E-23 | DF 7 | hist1h4l.3 |
| si:ch211-195b15.85 | 2.3E-27 | 1.2717724 | 0.18 | 0.089 | 7.61E-23 | DF 7 | si:ch211-195b15.8 |
| frmd62 | 5.2E-27 | 0.9696327 | 0.303 | 0.187 | 1.69E-22 | DF 7 | frmd6 |
| rrad3 | 5.5E-27 | 1.6046233 | 0.149 | 0.069 | 1.8E-22 | DF 7 | rrad |
| ciarta5 | 9.1E-27 | 0.7774113 | 0.408 | 0.281 | 2.97E-22 | DF 7 | ciarta |
| dap1b.14 | 1.6E-26 | 0.4523361 | 0.653 | 0.504 | 5.29E-22 | DF 7 | dap1b.1 |
| zfp36l22 | 2E-26 | 0.7277799 | 0.395 | 0.266 | 6.63E-22 | DF 7 | zfp36l2 |
| sinhcaf | 2.4E-26 | 1.3074953 | 0.131 | 0.056 | 7.65E-22 | DF 7 | sinhcaf |
| tlcd4b4 | 3.2E-26 | 1.2479408 | 0.168 | 0.081 | 1.05E-21 | DF 7 | tlcd4b |
| fzd101 | 3.6E-26 | 1.3214538 | 0.109 | 0.043 | 1.18E-21 | DF 7 | fzd10 |
| mid1ip1l3 | 3.9E-26 | 0.8969062 | 0.309 | 0.188 | 1.25E-21 | DF 7 | mid1ip1l |
| fth1a2 | 4.5E-26 | 0.4576019 | 0.769 | 0.718 | 1.46E-21 | DF 7 | fth1a |
| slc12a7b3 | 4.6E-26 | 0.9000992 | 0.26 | 0.149 | 1.5E-21 | DF 7 | slc12a7b |
| sgtb4 | 5.6E-26 | 1.0624292 | 0.224 | 0.123 | 1.83E-21 | DF 7 | sgtb |
| FQ323119.14 | 7.1E-26 | 0.9723392 | 0.198 | 0.102 | 2.3E-21 | DF 7 | FQ323119.1 |
| sf3b12 | 8.6E-26 | 0.6866358 | 0.462 | 0.335 | 2.79E-21 | DF 7 | sf3b1 |
| synm3 | 1E-25 | 0.708461 | 0.357 | 0.228 | 3.27E-21 | DF 7 | synm |
| sat1a.27 | 1.2E-25 | 0.8445915 | 0.557 | 0.455 | 3.77E-21 | DF 7 | sat1a.2 |
| etnk15 | 1.8E-25 | 0.8398346 | 0.303 | 0.186 | 5.92E-21 | DF 7 | etnk1 |
| creg24 | 2.7E-25 | 0.6767272 | 0.45 | 0.323 | 8.73E-21 | DF 7 | creg2 |
| zbtb16b4 | 4.6E-25 | 1.0317503 | 0.196 | 0.102 | 1.48E-20 | DF 7 | zbtb16b |
| ccdc80l15 | 5.6E-25 | 1.1245875 | 0.161 | 0.077 | 1.81E-20 | DF 7 | ccdc80l1 |
| csnk1da4 | 6.5E-25 | 0.8441486 | 0.328 | 0.21 | 2.12E-20 | DF 7 | csnk1da |
| ldhba4 | 1E-24 | 0.503184 | 0.611 | 0.492 | 3.31E-20 | DF 7 | ldhba |
| dusp12 | 1.2E-24 | 0.5644838 | 0.51 | 0.378 | 3.77E-20 | DF 7 | dusp1 |
| zgc:1538672 | 1.2E-24 | 0.4480805 | 0.835 | 0.751 | 3.86E-20 | DF 7 | zgc:153867 |
| runx35 | 1.2E-24 | 0.6967505 | 0.397 | 0.269 | 4.03E-20 | DF 7 | runx3 |
| mdh1aa4 | 1.7E-24 | 0.6249953 | 0.485 | 0.372 | 5.58E-20 | DF 7 | mdh1aa |
| trib33 | 2.7E-24 | 1.0657373 | 0.281 | 0.174 | 8.81E-20 | DF 7 | trib3 |
| zbtb16a3 | 2.9E-24 | 0.9401229 | 0.229 | 0.128 | 9.29E-20 | DF 7 | zbtb16a |
| rcan21 | 3E-24 | 1.3707731 | 0.128 | 0.057 | 9.76E-20 | DF 7 | rcan2 |
| prelid3b4 | 4.2E-24 | 0.9023018 | 0.459 | 0.356 | 1.38E-19 | DF 7 | prelid3b |

| | | | | | | | |
|---|---|---|---|---|---|---|---|
| aldocb2 | 5E-24 | 1.2073377 | 0.154 | 0.074 | 1.64E-19 | DF 7 | aldocb |
| zbtb203 | 5.4E-24 | 0.5924768 | 0.499 | 0.372 | 1.76E-19 | DF 7 | zbtb20 |
| twist1b4 | 5.8E-24 | 0.9615197 | 0.263 | 0.157 | 1.88E-19 | DF 7 | twist1b |
| cicb4 | 6.1E-24 | 0.886215 | 0.225 | 0.125 | 1.98E-19 | DF 7 | cicb |
| arl5c3 | 8E-24 | 1.1206305 | 0.214 | 0.12 | 2.6E-19 | DF 7 | arl5c |
| prickle1a4 | 1E-23 | 1.1283596 | 0.129 | 0.058 | 3.31E-19 | DF 7 | prickle1a |
| mhc1zaa4 | 1.5E-23 | 0.8650928 | 0.18 | 0.091 | 4.88E-19 | DF 7 | mhc1zaa |
| lonrf1l5 | 1.5E-23 | 0.9767688 | 0.252 | 0.15 | 5.03E-19 | DF 7 | lonrf1l |
| cavin1b4 | 2E-23 | 0.3352169 | 0.784 | 0.643 | 6.38E-19 | DF 7 | cavin1b |
| hopx4 | 2E-23 | 0.6707615 | 0.395 | 0.268 | 6.57E-19 | DF 7 | hopx |
| has23 | 2.3E-23 | 0.6986805 | 0.208 | 0.111 | 7.52E-19 | DF 7 | has2 |
| slc7a26 | 2.7E-23 | 0.6224664 | 0.269 | 0.157 | 8.73E-19 | DF 7 | slc7a2 |
| si:ch211-225b11.12 | 3.1E-23 | 1.1430393 | 0.151 | 0.073 | 1.02E-18 | DF 7 | si:ch211-225b11.1 |
| rem14 | 3.4E-23 | 1.229329 | 0.136 | 0.063 | 1.12E-18 | DF 7 | rem1 |
| rab11fip1a1 | 5.1E-23 | 1.3821883 | 0.13 | 0.06 | 1.65E-18 | DF 7 | rab11fip1a |
| si:ch211-284f22.35 | 1.1E-22 | 1.0390114 | 0.207 | 0.115 | 3.7E-18 | DF 7 | si:ch211-284f22.3 |
| DDX172 | 1.3E-22 | 0.6088826 | 0.421 | 0.297 | 4.27E-18 | DF 7 | DDX17 |
| siah1 | 1.3E-22 | 1.302534 | 0.129 | 0.06 | 4.3E-18 | DF 7 | siah1 |
| her64 | 1.6E-22 | 0.662286 | 0.547 | 0.43 | 5.21E-18 | DF 7 | her6 |
| map2k64 | 1.7E-22 | 0.8211123 | 0.297 | 0.188 | 5.47E-18 | DF 7 | map2k6 |
| midn4 | 1.9E-22 | 0.7852738 | 0.533 | 0.432 | 6.09E-18 | DF 7 | midn |
| nocta3 | 2.2E-22 | 1.4182997 | 0.199 | 0.112 | 7.26E-18 | DF 7 | nocta |
| ror22 | 2.5E-22 | 1.1405018 | 0.15 | 0.074 | 8.23E-18 | DF 7 | ror2 |
| smox3 | 2.7E-22 | 1.3920995 | 0.156 | 0.08 | 8.75E-18 | DF 7 | smox |
| irs12 | 4.4E-22 | 0.9449645 | 0.226 | 0.131 | 1.42E-17 | DF 7 | irs1 |
| cish2 | 7.2E-22 | 1.0932149 | 0.247 | 0.152 | 2.33E-17 | DF 7 | cish |
| tgfbr33 | 7.7E-22 | 1.0501427 | 0.156 | 0.078 | 2.51E-17 | DF 7 | tgfbr3 |
| capn2b7 | 7.8E-22 | 0.5057627 | 0.45 | 0.32 | 2.52E-17 | DF 7 | capn2b |
| zcchc245 | 9.9E-22 | 0.6076141 | 0.38 | 0.254 | 3.23E-17 | DF 7 | zcchc24 |
| myl9a3 | 1E-21 | 0.7363175 | 0.485 | 0.38 | 3.38E-17 | DF 7 | myl9a |
| ppp1r3cb | 1.1E-21 | 1.2274561 | 0.11 | 0.048 | 3.54E-17 | DF 7 | ppp1r3cb |
| fbxl72 | 1.4E-21 | 1.3057287 | 0.149 | 0.075 | 4.47E-17 | DF 7 | fbxl7 |
| mao3 | 1.5E-21 | 1.2039825 | 0.145 | 0.071 | 4.79E-17 | DF 7 | mao |
| niban1b5 | 1.8E-21 | 0.9804516 | 0.177 | 0.093 | 5.93E-17 | DF 7 | niban1b |
| ier2b3 | 2.3E-21 | 0.4242856 | 0.747 | 0.648 | 7.48E-17 | DF 7 | ier2b |
| vegfc3 | 2.5E-21 | 0.9759224 | 0.168 | 0.087 | 8.09E-17 | DF 7 | vegfc |
| dchs1b5 | 2.6E-21 | 1.0997227 | 0.17 | 0.09 | 8.52E-17 | DF 7 | dchs1b |
| si:dkey-23i12.72 | 2.9E-21 | 1.1117768 | 0.231 | 0.139 | 9.42E-17 | DF 7 | si:dkey-23i12.7 |
| rps262 | 3.5E-21 | 0.2130645 | 0.986 | 0.968 | 1.14E-16 | DF 7 | rps26 |
| c1qtnf23 | 3.5E-21 | 1.0339312 | 0.172 | 0.091 | 1.15E-16 | DF 7 | c1qtnf2 |
| scube33 | 4.4E-21 | 1.0262537 | 0.177 | 0.094 | 1.43E-16 | DF 7 | scube3 |
| tnfaip2a3 | 4.4E-21 | 1.1443891 | 0.166 | 0.087 | 1.43E-16 | DF 7 | tnfaip2a |
| pptc7a3 | 5.3E-21 | 0.717027 | 0.333 | 0.222 | 1.71E-16 | DF 7 | pptc7a |
| tuba8l24 | 5.8E-21 | 0.8747692 | 0.321 | 0.215 | 1.89E-16 | DF 7 | tuba8l2 |
| jun4 | 6.6E-21 | 0.3318611 | 0.911 | 0.853 | 2.13E-16 | DF 7 | jun |
| oaz2a1 | 7.7E-21 | 0.828811 | 0.152 | 0.077 | 2.49E-16 | DF 7 | oaz2a |
| bhlhe405 | 8E-21 | 0.5614874 | 0.568 | 0.463 | 2.59E-16 | DF 7 | bhlhe40 |
| flrt3 | 9.4E-21 | 1.3869352 | 0.12 | 0.056 | 3.06E-16 | DF 7 | flrt3 |

| | | | | | | | |
|---|---|---|---|---|---|---|---|
| dhcr7 | 9.4E-21 | 1.1872537 | 0.109 | 0.048 | 3.07E-16 | DF 7 | dhcr7 |
| zgc:1229793 | 1.4E-20 | 1.4173755 | 0.153 | 0.078 | 4.45E-16 | DF 7 | zgc:122979 |
| gpr146 | 1.4E-20 | 1.1752104 | 0.112 | 0.05 | 4.48E-16 | DF 7 | gpr146 |
| tnrc185 | 1.4E-20 | 0.6238009 | 0.515 | 0.405 | 4.68E-16 | DF 7 | tnrc18 |
| jund4 | 1.5E-20 | 0.4509214 | 0.66 | 0.568 | 4.75E-16 | DF 7 | jund |
| wls4 | 1.6E-20 | 0.5735748 | 0.508 | 0.393 | 5.05E-16 | DF 7 | wls |
| ppp1r105 | 1.9E-20 | 0.6174713 | 0.571 | 0.483 | 6.12E-16 | DF 7 | ppp1r10 |
| tuba8l45 | 2E-20 | 0.3196319 | 0.804 | 0.732 | 6.54E-16 | DF 7 | tuba8l4 |
| bsg2 | 2.2E-20 | 0.4023348 | 0.675 | 0.572 | 7.23E-16 | DF 7 | bsg |
| ackr3a1 | 2.2E-20 | 1.4557442 | 0.104 | 0.046 | 7.28E-16 | DF 7 | ackr3a |
| ctdsp22 | 2.4E-20 | 0.7600771 | 0.292 | 0.189 | 7.83E-16 | DF 7 | ctdsp2 |
| il1rapl1a1 | 2.5E-20 | 1.2278981 | 0.112 | 0.05 | 8.06E-16 | DF 7 | il1rapl1a |
| gse14 | 2.6E-20 | 0.9363335 | 0.213 | 0.124 | 8.45E-16 | DF 7 | gse1 |
| brdt4 | 3.3E-20 | 0.7331859 | 0.414 | 0.309 | 1.06E-15 | DF 7 | brdt |
| abtb2b3 | 6.4E-20 | 1.1535639 | 0.145 | 0.073 | 2.07E-15 | DF 7 | abtb2b |
| sept152 | 7.5E-20 | 0.7441228 | 0.301 | 0.202 | 2.45E-15 | DF 7 | sept15 |
| rsl24d15 | 7.8E-20 | 0.3494444 | 0.817 | 0.747 | 2.53E-15 | DF 7 | rsl24d1 |
| psd24 | 8.4E-20 | 1.0072283 | 0.176 | 0.096 | 2.74E-15 | DF 7 | psd2 |
| kdm6ba2 | 1.2E-19 | 0.9187273 | 0.319 | 0.22 | 3.76E-15 | DF 7 | kdm6ba |
| adamts15a6 | 1.2E-19 | 0.7211697 | 0.237 | 0.142 | 3.89E-15 | DF 7 | adamts15a |
| tuba1b6 | 1.5E-19 | 0.5982364 | 0.425 | 0.307 | 4.84E-15 | DF 7 | tuba1b |
| eef1da5 | 1.5E-19 | 0.3089781 | 0.91 | 0.875 | 4.86E-15 | DF 7 | eef1da |
| pbxip1b2 | 1.7E-19 | 0.5989978 | 0.351 | 0.239 | 5.64E-15 | DF 7 | pbxip1b |
| sec61g3 | 1.8E-19 | 0.256162 | 0.946 | 0.924 | 5.73E-15 | DF 7 | sec61g |
| mrtfbb1 | 1.9E-19 | 1.0923068 | 0.14 | 0.071 | 6.24E-15 | DF 7 | mrtfbb |
| btc1 | 2.8E-19 | 1.3745105 | 0.104 | 0.047 | 9.1E-15 | DF 7 | btc |
| nabp1a3 | 4.6E-19 | 0.8601598 | 0.272 | 0.178 | 1.48E-14 | DF 7 | nabp1a |
| myh9b2 | 5.7E-19 | 0.7054782 | 0.409 | 0.3 | 1.86E-14 | DF 7 | myh9b |
| anos1b2 | 7E-19 | 1.1580634 | 0.144 | 0.075 | 2.28E-14 | DF 7 | anos1b |
| itm2bb5 | 7.3E-19 | 0.8493746 | 0.165 | 0.088 | 2.38E-14 | DF 7 | itm2bb |
| celf26 | 7.4E-19 | 0.5783189 | 0.305 | 0.196 | 2.4E-14 | DF 7 | celf2 |
| slc25a54 | 9.5E-19 | 0.2177562 | 0.976 | 0.962 | 3.08E-14 | DF 7 | slc25a5 |
| vegfaa4 | 1.6E-18 | 0.95538 | 0.197 | 0.115 | 5.31E-14 | DF 7 | vegfaa |
| slit32 | 1.7E-18 | 0.7973365 | 0.297 | 0.2 | 5.44E-14 | DF 7 | slit3 |
| arg26 | 1.7E-18 | 0.8843917 | 0.255 | 0.162 | 5.52E-14 | DF 7 | arg2 |
| en1b4 | 2E-18 | 0.6559408 | 0.192 | 0.108 | 6.53E-14 | DF 7 | en1b |
| itgb44 | 2.1E-18 | 0.5025089 | 0.425 | 0.31 | 6.98E-14 | DF 7 | itgb4 |
| sorbs33 | 2.2E-18 | 0.8351921 | 0.282 | 0.189 | 7E-14 | DF 7 | sorbs3 |
| sh3bp5la3 | 3E-18 | 1.0561722 | 0.181 | 0.104 | 9.76E-14 | DF 7 | sh3bp5la |
| gsta.15 | 3.5E-18 | 0.4998825 | 0.468 | 0.355 | 1.12E-13 | DF 7 | gsta.1 |
| ENSDARG00000762 | 5E-18 | 0.3981553 | 0.698 | 0.622 | 1.63E-13 | DF 7 | ENSDARG00000076 |
| klf7b4 | 5.4E-18 | 0.7853628 | 0.324 | 0.226 | 1.75E-13 | DF 7 | klf7b |
| si:dkey-79d12.53 | 7.2E-18 | 0.9670852 | 0.21 | 0.128 | 2.34E-13 | DF 7 | si:dkey-79d12.5 |
| iscub3 | 9.3E-18 | 0.7162729 | 0.286 | 0.191 | 3.02E-13 | DF 7 | iscub |
| lrp1ab4 | 1E-17 | 0.5095791 | 0.45 | 0.339 | 3.27E-13 | DF 7 | lrp1ab |
| fscn1a5 | 1.1E-17 | 0.7192245 | 0.27 | 0.177 | 3.63E-13 | DF 7 | fscn1a |
| proza2 | 1.2E-17 | 0.8075878 | 0.259 | 0.168 | 3.96E-13 | DF 7 | proza |
| aldh9a1a.15 | 1.3E-17 | 0.3059304 | 0.599 | 0.479 | 4.2E-13 | DF 7 | aldh9a1a.1 |

| | | | | | | | |
|---|---|---|---|---|---|---|---|
| hmgn36 | 1.3E-17 | 0.7719039 | 0.251 | 0.161 | 4.26E-13 | DF 7 | hmgn3 |
| col6a4a3 | 1.6E-17 | 0.6049464 | 0.116 | 0.055 | 5.2E-13 | DF 7 | col6a4a |
| phldb2b4 | 2E-17 | 0.8056865 | 0.229 | 0.142 | 6.54E-13 | DF 7 | phldb2b |
| ankrd121 | 2.1E-17 | 0.5990213 | 0.35 | 0.248 | 6.87E-13 | DF 7 | ankrd12 |
| ier52 | 2.3E-17 | 0.4864755 | 0.572 | 0.468 | 7.35E-13 | DF 7 | ier5 |
| rbm10 | 2.3E-17 | 1.2393971 | 0.101 | 0.046 | 7.51E-13 | DF 7 | rbm10 |
| gstk12 | 2.5E-17 | 0.8614573 | 0.247 | 0.16 | 7.98E-13 | DF 7 | gstk1 |
| chac13 | 2.9E-17 | 0.7118762 | 0.444 | 0.353 | 9.43E-13 | DF 7 | chac1 |
| xbp13 | 3E-17 | 0.3714094 | 0.821 | 0.76 | 9.67E-13 | DF 7 | xbp1 |
| igsf104 | 3.1E-17 | 0.3507797 | 0.396 | 0.276 | 1.02E-12 | DF 7 | igsf10 |
| prrx1a4 | 3.9E-17 | 0.4165753 | 0.677 | 0.585 | 1.27E-12 | DF 7 | prrx1a |
| cd99l23 | 4.1E-17 | 0.6945695 | 0.302 | 0.206 | 1.35E-12 | DF 7 | cd99l2 |
| si:dkey-178e17.3 | 5.2E-17 | 1.2370025 | 0.102 | 0.048 | 1.7E-12 | DF 7 | si:dkey-178e17.3 |
| tpm33 | 5.3E-17 | 0.4398784 | 0.615 | 0.525 | 1.73E-12 | DF 7 | tpm3 |
| ttc367 | 6.7E-17 | 0.2668661 | 0.491 | 0.351 | 2.17E-12 | DF 7 | ttc36 |
| ugt5c2 | 7.3E-17 | 1.1702161 | 0.1 | 0.046 | 2.36E-12 | DF 7 | ugt5c2 |
| il11ra5 | 9.4E-17 | 0.5731049 | 0.341 | 0.238 | 3.06E-12 | DF 7 | il11ra |
| cirbpb3 | 1E-16 | 0.2647332 | 0.868 | 0.833 | 3.25E-12 | DF 7 | cirbpb |
| spen2 | 1.5E-16 | 0.7613774 | 0.288 | 0.196 | 4.97E-12 | DF 7 | spen |
| ncalda | 1.9E-16 | 1.2460375 | 0.111 | 0.055 | 6.22E-12 | DF 7 | ncalda |
| fbp1a4 | 2E-16 | 0.5407949 | 0.381 | 0.275 | 6.63E-12 | DF 7 | fbp1a |
| SSBP43 | 3.4E-16 | 0.8720374 | 0.198 | 0.12 | 1.1E-11 | DF 7 | SSBP4 |
| si:rp71-79p20.23 | 5.2E-16 | 0.80106 | 0.235 | 0.151 | 1.69E-11 | DF 7 | si:rp71-79p20.2 |
| lpp4 | 5.4E-16 | 0.6644152 | 0.346 | 0.252 | 1.74E-11 | DF 7 | lpp |
| zgc:158564 | 5.4E-16 | 0.7299093 | 0.32 | 0.23 | 1.75E-11 | DF 7 | zgc:158564 |
| capn125 | 5.6E-16 | 0.2932153 | 0.337 | 0.224 | 1.82E-11 | DF 7 | capn12 |
| naca3 | 6.6E-16 | 0.202976 | 0.979 | 0.961 | 2.16E-11 | DF 7 | naca |
| ulk22 | 7.3E-16 | 1.0691784 | 0.141 | 0.077 | 2.38E-11 | DF 7 | ulk2 |
| ctbp2a3 | 8.1E-16 | 0.5724715 | 0.327 | 0.229 | 2.62E-11 | DF 7 | ctbp2a |
| si:dkey-240h12.44 | 8.6E-16 | 0.7637789 | 0.194 | 0.116 | 2.81E-11 | DF 7 | si:dkey-240h12.4 |
| stat1a3 | 8.7E-16 | 0.6548438 | 0.428 | 0.338 | 2.81E-11 | DF 7 | stat1a |
| rock2a1 | 8.8E-16 | 0.7715289 | 0.282 | 0.195 | 2.87E-11 | DF 7 | rock2a |
| si:dkey-16p21.86 | 1.3E-15 | 0.4622513 | 0.309 | 0.208 | 4.38E-11 | DF 7 | si:dkey-16p21.8 |
| fosb5 | 1.5E-15 | 0.4270789 | 0.74 | 0.692 | 4.87E-11 | DF 7 | fosb |
| ARHGAP61 | 1.9E-15 | 0.9323338 | 0.122 | 0.063 | 6.31E-11 | DF 7 | ARHGAP6 |
| abat2 | 2.1E-15 | 0.9832889 | 0.169 | 0.098 | 6.7E-11 | DF 7 | abat |
| mpp6b1 | 2.6E-15 | 0.9449838 | 0.114 | 0.058 | 8.39E-11 | DF 7 | mpp6b |
| CU929418.12 | 3.5E-15 | 0.8334833 | 0.199 | 0.124 | 1.13E-10 | DF 7 | CU929418.1 |
| robo13 | 3.7E-15 | 1.1104089 | 0.139 | 0.077 | 1.2E-10 | DF 7 | robo1 |
| ssbp4 | 4.4E-15 | 0.9545469 | 0.136 | 0.074 | 1.42E-10 | DF 7 | ssbp4 |
| slc25a334 | 4.4E-15 | 0.9786746 | 0.188 | 0.116 | 1.44E-10 | DF 7 | slc25a33 |
| kcne42 | 4.5E-15 | 0.8283736 | 0.117 | 0.06 | 1.46E-10 | DF 7 | kcne4 |
| serbp1a3 | 4.9E-15 | 0.2245461 | 0.93 | 0.908 | 1.61E-10 | DF 7 | serbp1a |
| ca10a1 | 5.5E-15 | 1.1947863 | 0.102 | 0.051 | 1.78E-10 | DF 7 | ca10a |
| si:dkeyp-23e4.35 | 6.4E-15 | 0.7680175 | 0.151 | 0.085 | 2.07E-10 | DF 7 | si:dkeyp-23e4.3 |
| BX322530.11 | 6.6E-15 | 0.4186742 | 0.52 | 0.418 | 2.16E-10 | DF 7 | BX322530.1 |
| myh7ba1 | 1E-14 | 0.9674172 | 0.108 | 0.054 | 3.35E-10 | DF 7 | myh7ba |
| rab40b3 | 1.4E-14 | 0.9405114 | 0.139 | 0.077 | 4.45E-10 | DF 7 | rab40b |

| | | | | | | | |
|---|---|---|---|---|---|---|---|
| CR383676.35 | 1.5E-14 | 0.3896994 | 0.561 | 0.464 | 4.86E-10 | DF 7 | CR383676.3 |
| hlfa3 | 1.7E-14 | 0.8550373 | 0.205 | 0.131 | 5.58E-10 | DF 7 | hlfa |
| zgc:1583433 | 1.8E-14 | 0.5640277 | 0.475 | 0.376 | 5.73E-10 | DF 7 | zgc:158343 |
| kirrel1a3 | 1.9E-14 | 0.4362622 | 0.423 | 0.325 | 6.21E-10 | DF 7 | kirrel1a |
| pik3r3a3 | 2E-14 | 0.8224829 | 0.208 | 0.134 | 6.41E-10 | DF 7 | pik3r3a |
| cd248a4 | 2.6E-14 | 0.6006159 | 0.19 | 0.115 | 8.29E-10 | DF 7 | cd248a |
| zgc:1135316 | 3.7E-14 | 0.4303473 | 0.378 | 0.277 | 1.22E-09 | DF 7 | zgc:113531 |
| klf11a | 4.1E-14 | 1.0645394 | 0.123 | 0.066 | 1.33E-09 | DF 7 | klf11a |
| cebpb4 | 4.1E-14 | 0.7039124 | 0.527 | 0.447 | 1.35E-09 | DF 7 | cebpb |
| pdlim12 | 4.3E-14 | 0.7038296 | 0.312 | 0.228 | 1.39E-09 | DF 7 | pdlim1 |
| jag1b2 | 6.2E-14 | 0.6505054 | 0.3 | 0.214 | 2.02E-09 | DF 7 | jag1b |
| mmp15a5 | 7.8E-14 | 1.079421 | 0.143 | 0.083 | 2.55E-09 | DF 7 | mmp15a |
| foxo3b3 | 1.1E-13 | 0.9485862 | 0.141 | 0.081 | 3.62E-09 | DF 7 | foxo3b |
| ell3 | 1.2E-13 | 0.8697591 | 0.155 | 0.092 | 3.92E-09 | DF 7 | ell |
| enpp62 | 1.2E-13 | 1.1003799 | 0.107 | 0.056 | 3.99E-09 | DF 7 | enpp6 |
| sesn14 | 1.4E-13 | 0.5752668 | 0.305 | 0.216 | 4.41E-09 | DF 7 | sesn1 |
| mertka1 | 1.5E-13 | 0.9818531 | 0.188 | 0.12 | 4.97E-09 | DF 7 | mertka |
| zbtb43 | 1.7E-13 | 0.5496994 | 0.41 | 0.324 | 5.58E-09 | DF 7 | zbtb4 |
| akt1s12 | 1.8E-13 | 0.6888302 | 0.231 | 0.155 | 5.86E-09 | DF 7 | akt1s1 |
| id2a6 | 1.9E-13 | 0.3874544 | 0.589 | 0.502 | 6.06E-09 | DF 7 | id2a |
| klf156 | 2E-13 | 0.4991093 | 0.295 | 0.207 | 6.66E-09 | DF 7 | klf15 |
| grb10b2 | 3E-13 | 0.3837882 | 0.455 | 0.359 | 9.74E-09 | DF 7 | grb10b |
| ppl5 | 3E-13 | 0.5506184 | 0.235 | 0.153 | 9.75E-09 | DF 7 | ppl |
| ucp26 | 3.1E-13 | 0.2545286 | 0.522 | 0.413 | 1E-08 | DF 7 | ucp2 |
| nck1a4 | 3.1E-13 | 0.917517 | 0.137 | 0.078 | 1.01E-08 | DF 7 | nck1a |
| qsox14 | 3.5E-13 | 0.728358 | 0.124 | 0.068 | 1.15E-08 | DF 7 | qsox1 |
| jdp2b4 | 4E-13 | 0.4615251 | 0.667 | 0.569 | 1.31E-08 | DF 7 | jdp2b |
| csrnp1b4 | 4.2E-13 | 0.7042312 | 0.264 | 0.186 | 1.36E-08 | DF 7 | csrnp1b |
| srsf3b3 | 4.6E-13 | 0.5392137 | 0.36 | 0.273 | 1.49E-08 | DF 7 | srsf3b |
| ptp4a3a2 | 6.5E-13 | 0.7378203 | 0.187 | 0.119 | 2.12E-08 | DF 7 | ptp4a3a |
| appa1 | 6.7E-13 | 0.554922 | 0.365 | 0.281 | 2.18E-08 | DF 7 | appa |
| lrrc58b4 | 7.7E-13 | 0.8384833 | 0.189 | 0.121 | 2.52E-08 | DF 7 | lrrc58b |
| efnb3b3 | 8.5E-13 | 0.7206846 | 0.208 | 0.138 | 2.78E-08 | DF 7 | efnb3b |
| kank45 | 9.8E-13 | 0.6619764 | 0.168 | 0.102 | 3.17E-08 | DF 7 | kank4 |
| pdk2b3 | 1.2E-12 | 0.6295041 | 0.278 | 0.198 | 3.8E-08 | DF 7 | pdk2b |
| hmox1a5 | 1.2E-12 | 0.4868834 | 0.252 | 0.173 | 3.93E-08 | DF 7 | hmox1a |
| zgc:1748884 | 1.3E-12 | 0.7469793 | 0.296 | 0.216 | 4.28E-08 | DF 7 | zgc:174888 |
| osr14 | 1.3E-12 | 0.3675933 | 0.256 | 0.171 | 4.35E-08 | DF 7 | osr1 |
| ddit43 | 1.4E-12 | 0.7804229 | 0.139 | 0.081 | 4.4E-08 | DF 7 | ddit4 |
| bcat22 | 1.4E-12 | 1.0093268 | 0.146 | 0.088 | 4.44E-08 | DF 7 | bcat2 |
| camk2b1 | 1.4E-12 | 0.9223558 | 0.113 | 0.061 | 4.52E-08 | DF 7 | camk2b1 |
| gfra1a2 | 1.4E-12 | 0.792611 | 0.104 | 0.055 | 4.56E-08 | DF 7 | gfra1a |
| per1b4 | 1.5E-12 | 0.5540676 | 0.283 | 0.202 | 4.72E-08 | DF 7 | per1b |
| znf219 | 1.6E-12 | 1.0662457 | 0.102 | 0.054 | 5.09E-08 | DF 7 | znf219 |
| sned12 | 1.6E-12 | 0.5969818 | 0.115 | 0.062 | 5.13E-08 | DF 7 | sned1 |
| ier2a4 | 1.6E-12 | 0.4942919 | 0.552 | 0.485 | 5.33E-08 | DF 7 | ier2a |
| clec3ba3 | 1.7E-12 | 0.4780265 | 0.198 | 0.125 | 5.55E-08 | DF 7 | clec3ba |
| lef14 | 2.7E-12 | 0.6616318 | 0.172 | 0.107 | 8.79E-08 | DF 7 | lef1 |

| | | | | | | | |
|---|---|---|---|---|---|---|---|
| cst32 | 2.8E-12 | 0.248218 | 0.801 | 0.722 | 9.18E-08 | DF 7 | cst3 |
| gstz15 | 2.9E-12 | 0.3348771 | 0.44 | 0.343 | 9.37E-08 | DF 7 | gstz1 |
| smad12 | 3.5E-12 | 0.7814675 | 0.175 | 0.111 | 1.15E-07 | DF 7 | smad1 |
| hspa4a4 | 4.1E-12 | 0.7743651 | 0.218 | 0.148 | 1.32E-07 | DF 7 | hspa4a |
| rad54l23 | 4.1E-12 | 0.8297969 | 0.139 | 0.082 | 1.34E-07 | DF 7 | rad54l2 |
| zyx6 | 4.1E-12 | 0.4659241 | 0.402 | 0.316 | 1.35E-07 | DF 7 | zyx |
| adipor24 | 5.2E-12 | 0.7281865 | 0.249 | 0.176 | 1.69E-07 | DF 7 | adipor2 |
| hist1h4l.101 | 5.5E-12 | 0.8029188 | 0.127 | 0.073 | 1.78E-07 | DF 7 | hist1h4l.10 |
| acvrl14 | 6E-12 | 0.6933901 | 0.32 | 0.245 | 1.96E-07 | DF 7 | acvrl1 |
| pkdcca2 | 6.3E-12 | 1.0668448 | 0.12 | 0.069 | 2.06E-07 | DF 7 | pkdcca |
| soul53 | 7.8E-12 | 0.241203 | 0.372 | 0.275 | 2.53E-07 | DF 7 | soul5 |
| lmo4b6 | 9.1E-12 | 0.3501404 | 0.413 | 0.317 | 2.96E-07 | DF 7 | lmo4b |
| pcdh18a4 | 9.6E-12 | 0.5855866 | 0.139 | 0.082 | 3.12E-07 | DF 7 | pcdh18a |
| rbfox24 | 9.9E-12 | 0.5471977 | 0.348 | 0.27 | 3.22E-07 | DF 7 | rbfox2 |
| ubl3a4 | 1.3E-11 | 0.4169193 | 0.419 | 0.336 | 4.38E-07 | DF 7 | ubl3a |
| ddit34 | 1.6E-11 | 0.3263032 | 0.648 | 0.581 | 5.21E-07 | DF 7 | ddit3 |
| nova23 | 1.7E-11 | 0.2386194 | 0.305 | 0.212 | 5.68E-07 | DF 7 | nova2 |
| ENSDARG000000713. | 2.5E-11 | 0.7721413 | 0.141 | 0.086 | 8.2E-07 | DF 7 | ENSDARG00000071 |
| il6st5 | 2.6E-11 | 0.5776602 | 0.496 | 0.422 | 8.46E-07 | DF 7 | il6st |
| pik3ip12 | 2.7E-11 | 0.6872805 | 0.128 | 0.075 | 8.89E-07 | DF 7 | pik3ip1 |
| fxyd6l3 | 3.7E-11 | 0.398829 | 0.335 | 0.253 | 1.21E-06 | DF 7 | fxyd6l |
| coq10b3 | 4E-11 | 0.7148473 | 0.236 | 0.167 | 1.29E-06 | DF 7 | coq10b |
| ptp4a2b2 | 4.2E-11 | 0.5699348 | 0.373 | 0.303 | 1.36E-06 | DF 7 | ptp4a2b |
| twist26 | 4.3E-11 | 0.2601431 | 0.44 | 0.339 | 1.4E-06 | DF 7 | twist2 |
| spry24 | 4.7E-11 | 0.8862778 | 0.13 | 0.078 | 1.52E-06 | DF 7 | spry2 |
| enah3 | 5.6E-11 | 0.664086 | 0.177 | 0.115 | 1.82E-06 | DF 7 | enah |
| cdr2l2 | 5.7E-11 | 0.7277369 | 0.124 | 0.073 | 1.84E-06 | DF 7 | cdr2l |
| ubc3 | 6.9E-11 | 0.5818209 | 0.688 | 0.663 | 2.25E-06 | DF 7 | ubc |
| CR318588.13 | 8.1E-11 | 1.5896899 | 0.353 | 0.295 | 2.63E-06 | DF 7 | CR318588.1 |
| amd13 | 8.7E-11 | 0.813747 | 0.151 | 0.095 | 2.83E-06 | DF 7 | amd1 |
| cry1b1 | 9E-11 | 0.8421876 | 0.1 | 0.055 | 2.93E-06 | DF 7 | cry1b |
| eif3g2 | 9.7E-11 | 0.3875898 | 0.595 | 0.525 | 3.14E-06 | DF 7 | eif3g |
| nt5c3a3 | 1.1E-10 | 0.6677773 | 0.236 | 0.169 | 3.42E-06 | DF 7 | nt5c3a |
| fhl3b3 | 1.1E-10 | 0.924438 | 0.168 | 0.111 | 3.74E-06 | DF 7 | fhl3b |
| gyg1a2 | 1.2E-10 | 0.5976773 | 0.251 | 0.182 | 3.84E-06 | DF 7 | gyg1a |
| nsd1b2 | 1.2E-10 | 0.8524868 | 0.123 | 0.073 | 3.94E-06 | DF 7 | nsd1b |
| creb3l22 | 1.2E-10 | 0.4076599 | 0.502 | 0.425 | 3.99E-06 | DF 7 | creb3l2 |
| hnrnpa0l.13 | 1.3E-10 | 0.3953576 | 0.702 | 0.667 | 4.25E-06 | DF 7 | hnrnpa0l.1 |
| ncl3 | 1.3E-10 | 0.3505469 | 0.665 | 0.603 | 4.29E-06 | DF 7 | ncl |
| myclb4 | 1.5E-10 | 0.6483608 | 0.205 | 0.14 | 4.76E-06 | DF 7 | myclb |
| atp2b1a2 | 1.5E-10 | 0.5480788 | 0.318 | 0.245 | 4.9E-06 | DF 7 | atp2b1a |
| kank23 | 1.5E-10 | 0.6054603 | 0.282 | 0.211 | 4.97E-06 | DF 7 | kank2 |
| skia5 | 1.6E-10 | 0.3193493 | 0.527 | 0.448 | 5.15E-06 | DF 7 | skia |
| eif4a24 | 1.7E-10 | 0.5917757 | 0.261 | 0.192 | 5.56E-06 | DF 7 | eif4a2 |
| tspan3a1 | 1.7E-10 | 0.5938222 | 0.229 | 0.161 | 5.69E-06 | DF 7 | tspan3a |
| myca5 | 1.8E-10 | 0.6257414 | 0.223 | 0.157 | 5.95E-06 | DF 7 | myca |
| efna5b3 | 1.9E-10 | 0.8124357 | 0.117 | 0.069 | 6.03E-06 | DF 7 | efna5b |
| mgaa2 | 2.2E-10 | 0.6695855 | 0.204 | 0.14 | 7.09E-06 | DF 7 | mgaa |

| | | | | | | | |
|---|---|---|---|---|---|---|---|
| klf32 | 2.4E-10 | 0.7222052 | 0.157 | 0.1 | 7.72E-06 | DF 7 | klf3 |
| thbs1a3 | 2.6E-10 | 0.3062294 | 0.394 | 0.312 | 8.3E-06 | DF 7 | thbs1a |
| cry23 | 2.6E-10 | 0.5181376 | 0.234 | 0.165 | 8.61E-06 | DF 7 | cry2 |
| si:ch211-286b5.51 | 2.8E-10 | 0.4797969 | 0.33 | 0.257 | 9.01E-06 | DF 7 | si:ch211-286b5.5 |
| SERF2 | 3.1E-10 | 0.6834771 | 0.152 | 0.097 | 1E-05 | DF 7 | SERF2 |
| zgc:1140465 | 3.2E-10 | 0.6831482 | 0.155 | 0.099 | 1.04E-05 | DF 7 | zgc:114046 |
| ldb2a | 3.9E-10 | 0.7956178 | 0.109 | 0.063 | 1.27E-05 | DF 7 | ldb2a |
| ctdsp11 | 4.2E-10 | 0.701656 | 0.169 | 0.112 | 1.36E-05 | DF 7 | ctdsp1 |
| actn14 | 4.2E-10 | 0.5909935 | 0.386 | 0.316 | 1.38E-05 | DF 7 | actn1 |
| gstm.2 | 4.5E-10 | 0.9206036 | 0.117 | 0.07 | 1.46E-05 | DF 7 | gstm.2 |
| arl4ab5 | 5E-10 | 0.6911936 | 0.167 | 0.109 | 1.61E-05 | DF 7 | arl4ab |
| brd2a4 | 5E-10 | 0.5134855 | 0.51 | 0.446 | 1.64E-05 | DF 7 | brd2a |
| zgc:921405 | 5.2E-10 | 0.5888336 | 0.171 | 0.113 | 1.7E-05 | DF 7 | zgc:92140 |
| mcl1b4 | 5.3E-10 | 0.313216 | 0.555 | 0.482 | 1.71E-05 | DF 7 | mcl1b |
| caskin13 | 5.3E-10 | 0.7910281 | 0.102 | 0.058 | 1.73E-05 | DF 7 | caskin1 |
| prkar2aa | 7.2E-10 | 0.5708363 | 0.226 | 0.162 | 2.36E-05 | DF 7 | prkar2aa |
| tnrc6c14 | 7.3E-10 | 0.3464644 | 0.553 | 0.485 | 2.38E-05 | DF 7 | tnrc6c1 |
| sltm2 | 7.8E-10 | 0.4349206 | 0.356 | 0.28 | 2.54E-05 | DF 7 | sltm |
| zgc:1530313 | 8.1E-10 | 0.8326433 | 0.106 | 0.061 | 2.64E-05 | DF 7 | zgc:153031 |
| arid5b3 | 8.7E-10 | 0.5515904 | 0.196 | 0.135 | 2.83E-05 | DF 7 | arid5b |
| tnrc6c22 | 1.1E-09 | 0.8163944 | 0.138 | 0.087 | 3.5E-05 | DF 7 | tnrc6c2 |
| CPNE86 | 1.3E-09 | 0.3835703 | 0.361 | 0.285 | 4.15E-05 | DF 7 | CPNE8 |
| pcbd16 | 1.4E-09 | 0.2751674 | 0.407 | 0.315 | 4.5E-05 | DF 7 | pcbd1 |
| pgm13 | 1.5E-09 | 0.7569414 | 0.152 | 0.099 | 4.86E-05 | DF 7 | pgm1 |
| sh3gl1b3 | 1.5E-09 | 0.584239 | 0.309 | 0.243 | 4.99E-05 | DF 7 | sh3gl1b |
| aff25 | 1.6E-09 | 0.5402244 | 0.244 | 0.178 | 5.34E-05 | DF 7 | aff2 |
| zgc:1949894 | 1.7E-09 | 0.724635 | 0.142 | 0.09 | 5.47E-05 | DF 7 | zgc:194989 |
| si:ch211-196l7.44 | 1.7E-09 | 0.3508453 | 0.542 | 0.475 | 5.52E-05 | DF 7 | si:ch211-196l7.4 |
| mych4 | 1.8E-09 | 0.4209138 | 0.455 | 0.378 | 5.88E-05 | DF 7 | mych |
| map1ab4 | 2E-09 | 0.4352337 | 0.264 | 0.196 | 6.41E-05 | DF 7 | map1ab |
| pdcd4b2 | 2.2E-09 | 0.8199563 | 0.161 | 0.107 | 7.05E-05 | DF 7 | pdcd4b |
| adh54 | 2.5E-09 | 0.2963987 | 0.585 | 0.517 | 8.03E-05 | DF 7 | adh5 |
| kif26ba6 | 2.6E-09 | 0.4735705 | 0.164 | 0.107 | 8.54E-05 | DF 7 | kif26ba |
| fkbp52 | 2.6E-09 | 0.4739697 | 0.265 | 0.196 | 8.54E-05 | DF 7 | fkbp5 |
| ndrg3a1 | 2.8E-09 | 0.7530775 | 0.148 | 0.097 | 8.95E-05 | DF 7 | ndrg3a |
| PPM1K2 | 3.1E-09 | 0.6905623 | 0.106 | 0.062 | 0.000101 | DF 7 | PPM1K |
| sox11a4 | 3.3E-09 | 0.4174965 | 0.421 | 0.342 | 0.000106 | DF 7 | sox11a |
| srrm12 | 3.4E-09 | 0.3500347 | 0.476 | 0.407 | 0.000111 | DF 7 | srrm1 |
| ada4 | 3.5E-09 | 0.4835993 | 0.289 | 0.22 | 0.000113 | DF 7 | ada |
| pear14 | 3.8E-09 | 0.6323354 | 0.187 | 0.13 | 0.000123 | DF 7 | pear1 |
| myo10l33 | 4.8E-09 | 0.71853 | 0.15 | 0.099 | 0.000155 | DF 7 | myo10l3 |
| auts2a4 | 5E-09 | 0.596521 | 0.285 | 0.222 | 0.000164 | DF 7 | auts2a |
| smarcb1b | 5.1E-09 | 0.855893 | 0.104 | 0.062 | 0.000164 | DF 7 | smarcb1b |
| rap1b1 | 5.2E-09 | 0.4429658 | 0.425 | 0.365 | 0.000169 | DF 7 | rap1b |
| dusp55 | 5.2E-09 | 0.6510514 | 0.26 | 0.197 | 0.00017 | DF 7 | dusp5 |
| bag33 | 5.5E-09 | 0.8106003 | 0.116 | 0.071 | 0.000178 | DF 7 | bag3 |
| pawr4 | 5.5E-09 | 0.3093148 | 0.257 | 0.186 | 0.000179 | DF 7 | pawr |
| slc7a8a4 | 5.5E-09 | 0.8629148 | 0.132 | 0.085 | 0.00018 | DF 7 | slc7a8a |

| | | | | | | | |
|---|---|---|---|---|---|---|---|
| crema3 | 5.8E-09 | 0.7464143 | 0.161 | 0.108 | 0.000188 | DF 7 | crema |
| cmbl4 | 5.9E-09 | 0.5548552 | 0.237 | 0.174 | 0.000192 | DF 7 | cmbl |
| pcf113 | 7.2E-09 | 0.4945149 | 0.316 | 0.251 | 0.000234 | DF 7 | pcf11 |
| dip2ca3 | 7.7E-09 | 0.6995069 | 0.134 | 0.086 | 0.000249 | DF 7 | dip2ca |
| cpe3 | 8.3E-09 | 0.4463732 | 0.269 | 0.202 | 0.000269 | DF 7 | cpe |
| rbms2a1 | 8.3E-09 | 0.5476522 | 0.201 | 0.143 | 0.000269 | DF 7 | rbms2a |
| prx2 | 1E-08 | 0.2364391 | 0.109 | 0.065 | 0.000329 | DF 7 | prx |
| cgnl13 | 1.1E-08 | 0.68926 | 0.117 | 0.072 | 0.000349 | DF 7 | cgnl1 |
| srpx24 | 1.1E-08 | 0.4459463 | 0.322 | 0.253 | 0.000361 | DF 7 | srpx2 |
| zfand2a3 | 1.1E-08 | 0.9174999 | 0.109 | 0.067 | 0.000362 | DF 7 | zfand2a |
| sp42 | 1.2E-08 | 0.6125319 | 0.207 | 0.149 | 0.000375 | DF 7 | sp4 |
| inka1b3 | 1.4E-08 | 0.7485652 | 0.139 | 0.091 | 0.000455 | DF 7 | inka1b |
| nktr3 | 1.4E-08 | 0.6241333 | 0.216 | 0.158 | 0.000455 | DF 7 | nktr |
| pebp15 | 1.4E-08 | 0.8784857 | 0.115 | 0.072 | 0.00046 | DF 7 | pebp1 |
| pbx43 | 1.4E-08 | 0.4111547 | 0.327 | 0.258 | 0.000461 | DF 7 | pbx4 |
| tnfsf10l6 | 1.7E-08 | 0.3602791 | 0.29 | 0.221 | 0.000545 | DF 7 | tnfsf10l |
| C7H4orf482 | 2E-08 | 0.5046015 | 0.305 | 0.242 | 0.000637 | DF 7 | C7H4orf48 |
| zmp:00000007114 | 2E-08 | 0.7226731 | 0.127 | 0.082 | 0.00066 | DF 7 | zmp:0000000711 |
| plekhj12 | 2.2E-08 | 0.5384618 | 0.175 | 0.121 | 0.000708 | DF 7 | plekhj1 |
| lims1 | 2.2E-08 | 0.5146606 | 0.354 | 0.294 | 0.000729 | DF 7 | lims1 |
| cnbpb2 | 2.7E-08 | 0.264446 | 0.637 | 0.576 | 0.00088 | DF 7 | cnbpb |
| CT956002.22 | 2.8E-08 | 0.785009 | 0.119 | 0.075 | 0.000901 | DF 7 | CT956002.2 |
| tent5c2 | 2.9E-08 | 0.7710772 | 0.128 | 0.083 | 0.000951 | DF 7 | tent5c |
| sod3b4 | 3E-08 | 0.3847412 | 0.178 | 0.123 | 0.000968 | DF 7 | sod3b |
| dhrs12 | 3.8E-08 | 0.5644736 | 0.249 | 0.191 | 0.001235 | DF 7 | dhrs1 |
| lrba4 | 4E-08 | 0.5358875 | 0.236 | 0.176 | 0.001294 | DF 7 | lrba |
| ddx3xa2 | 4E-08 | 0.4234899 | 0.384 | 0.323 | 0.001298 | DF 7 | ddx3xa |
| ube2al3 | 4.2E-08 | 0.5842454 | 0.257 | 0.198 | 0.001379 | DF 7 | ube2al |
| serpina10a5 | 4.6E-08 | 0.3759673 | 0.438 | 0.378 | 0.001486 | DF 7 | serpina10a |
| msmo11 | 4.7E-08 | 0.7923353 | 0.104 | 0.064 | 0.001513 | DF 7 | msmo1 |
| znf5321 | 4.9E-08 | 0.6534202 | 0.112 | 0.07 | 0.0016 | DF 7 | znf532 |
| acadm2 | 5.2E-08 | 0.5028196 | 0.313 | 0.254 | 0.001676 | DF 7 | acadm |
| fbxl3a2 | 5.5E-08 | 0.7493248 | 0.14 | 0.095 | 0.001802 | DF 7 | fbxl3a |
| dusp8a4 | 5.7E-08 | 0.6157676 | 0.136 | 0.089 | 0.001843 | DF 7 | dusp8a |
| rfc11 | 5.7E-08 | 0.7766613 | 0.105 | 0.065 | 0.001868 | DF 7 | rfc1 |
| gadd45ab5 | 6.4E-08 | 0.5252495 | 0.234 | 0.175 | 0.00208 | DF 7 | gadd45ab |
| slc48a1b2 | 6.4E-08 | 0.5515952 | 0.195 | 0.14 | 0.002082 | DF 7 | slc48a1b |
| ghrb5 | 6.4E-08 | 0.678262 | 0.156 | 0.108 | 0.002086 | DF 7 | ghrb |
| pde10a2 | 6.5E-08 | 0.7442851 | 0.102 | 0.062 | 0.002116 | DF 7 | pde10a |
| cpeb4b2 | 6.5E-08 | 0.5941829 | 0.222 | 0.167 | 0.002126 | DF 7 | cpeb4b |
| cdo15 | 6.6E-08 | 0.366726 | 0.4 | 0.328 | 0.002158 | DF 7 | cdo1 |
| tmem1766 | 7.1E-08 | 0.2026776 | 0.477 | 0.393 | 0.0023 | DF 7 | tmem176 |
| ptpn11b2 | 7.2E-08 | 0.8317258 | 0.103 | 0.064 | 0.002357 | DF 7 | ptpn11b |
| mprip | 8.9E-08 | 0.7691944 | 0.117 | 0.076 | 0.002883 | DF 7 | mprip |
| si:ch73-86n18.14 | 9.1E-08 | 0.4677902 | 0.212 | 0.153 | 0.002957 | DF 7 | si:ch73-86n18.1 |
| slc25a55a4 | 9.9E-08 | 0.6822036 | 0.25 | 0.197 | 0.003231 | DF 7 | slc25a55a |
| arhgdia2 | 1.1E-07 | 0.3441485 | 0.474 | 0.411 | 0.003648 | DF 7 | arhgdia |
| nos1apa5 | 1.1E-07 | 0.8440482 | 0.154 | 0.108 | 0.003715 | DF 7 | nos1apa |

| | | | | | | | |
|---|---|---|---|---|---|---|---|
| polr2a5 | 1.2E-07 | 0.3547508 | 0.426 | 0.362 | 0.003825 | DF 7 | polr2a |
| spry44 | 1.2E-07 | 0.6989251 | 0.23 | 0.178 | 0.003995 | DF 7 | spry4 |
| trim3b5 | 1.3E-07 | 0.4878027 | 0.141 | 0.094 | 0.004186 | DF 7 | trim3b |
| cox4i22 | 1.3E-07 | 0.3876461 | 0.382 | 0.32 | 0.004264 | DF 7 | cox4i2 |
| nectin1b1 | 1.3E-07 | 0.7385343 | 0.114 | 0.073 | 0.004292 | DF 7 | nectin1b |
| hspa12b4 | 1.4E-07 | 0.4284958 | 0.175 | 0.122 | 0.004607 | DF 7 | hspa12b |
| ppp2r2d1 | 1.5E-07 | 0.5529832 | 0.194 | 0.142 | 0.004892 | DF 7 | ppp2r2d |
| ccdc3a2 | 1.6E-07 | 0.9445808 | 0.106 | 0.067 | 0.005136 | DF 7 | ccdc3a |
| adgre103 | 1.9E-07 | 0.4350542 | 0.117 | 0.075 | 0.006273 | DF 7 | adgre10 |
| MDFI4 | 1.9E-07 | 0.4419591 | 0.298 | 0.238 | 0.006302 | DF 7 | MDFI |
| pmaip12 | 2.1E-07 | 0.6345808 | 0.139 | 0.094 | 0.006825 | DF 7 | pmaip1 |
| mdh21 | 2.3E-07 | 0.3253955 | 0.492 | 0.439 | 0.007317 | DF 7 | mdh2 |
| hexim14 | 2.3E-07 | 0.5107619 | 0.291 | 0.233 | 0.00732 | DF 7 | hexim1 |
| akap8l1 | 2.3E-07 | 0.4454716 | 0.252 | 0.195 | 0.007351 | DF 7 | akap8l |
| mapk11 | 2.7E-07 | 0.8641063 | 0.124 | 0.084 | 0.008628 | DF 7 | mapk1 |
| zgc:1533724 | 3.2E-07 | 0.8389474 | 0.128 | 0.086 | 0.010267 | DF 7 | zgc:153372 |
| stk352 | 3.3E-07 | 0.7831699 | 0.124 | 0.084 | 0.010693 | DF 7 | stk35 |
| fbxw11b3 | 3.3E-07 | 0.5063694 | 0.211 | 0.159 | 0.010821 | DF 7 | fbxw11b |
| znf395b3 | 3.8E-07 | 0.5073838 | 0.188 | 0.138 | 0.012374 | DF 7 | znf395b |
| smc33 | 4.3E-07 | 0.5642459 | 0.196 | 0.145 | 0.014116 | DF 7 | smc3 |
| snai1b3 | 4.6E-07 | 0.5700414 | 0.158 | 0.112 | 0.014805 | DF 7 | snai1b |
| smarca12 | 4.7E-07 | 0.6155942 | 0.123 | 0.081 | 0.015384 | DF 7 | smarca1 |
| f11r.11 | 5E-07 | 0.703196 | 0.116 | 0.076 | 0.016144 | DF 7 | f11r.1 |
| oaz1a5 | 6.4E-07 | 0.3025606 | 0.505 | 0.452 | 0.020727 | DF 7 | oaz1a |
| txn21 | 6.4E-07 | 0.6264913 | 0.156 | 0.111 | 0.020855 | DF 7 | txn2 |
| tefa2 | 6.5E-07 | 0.3926179 | 0.275 | 0.219 | 0.021024 | DF 7 | tefa |
| tmem472 | 6.5E-07 | 0.62944 | 0.208 | 0.16 | 0.021068 | DF 7 | tmem47 |
| tspan313 | 7.1E-07 | 0.4847698 | 0.28 | 0.226 | 0.023195 | DF 7 | tspan31 |
| abcc53 | 7.3E-07 | 0.5257822 | 0.193 | 0.143 | 0.023673 | DF 7 | abcc5 |
| slc25a25a4 | 7.4E-07 | 0.48749 | 0.271 | 0.218 | 0.024134 | DF 7 | slc25a25a |
| fosl24 | 7.9E-07 | 0.4044942 | 0.471 | 0.428 | 0.025781 | DF 7 | fosl2 |
| znf106a | 8.1E-07 | 0.8006267 | 0.101 | 0.065 | 0.026417 | DF 7 | znf106a |
| si:dkey-7j14.52 | 8.3E-07 | 0.5967479 | 0.108 | 0.07 | 0.027077 | DF 7 | si:dkey-7j14.5 |
| sult1st64 | 8.7E-07 | 0.4522785 | 0.163 | 0.116 | 0.028239 | DF 7 | sult1st6 |
| rnf1112 | 8.9E-07 | 0.5241832 | 0.135 | 0.093 | 0.028951 | DF 7 | rnf111 |
| ANPEP3 | 9E-07 | 0.4253658 | 0.266 | 0.21 | 0.029393 | DF 7 | ANPEP |
| tmem119b5 | 9.8E-07 | 0.3688135 | 0.265 | 0.206 | 0.031779 | DF 7 | tmem119b |
| ehbp13 | 1E-06 | 0.5910862 | 0.154 | 0.11 | 0.032849 | DF 7 | ehbp1 |
| clstn13 | 1.1E-06 | 0.4904374 | 0.185 | 0.136 | 0.034406 | DF 7 | clstn1 |
| per24 | 1.1E-06 | 0.6563788 | 0.147 | 0.104 | 0.034867 | DF 7 | per2 |
| tardbpl1 | 1.1E-06 | 0.502885 | 0.204 | 0.155 | 0.036145 | DF 7 | tardbpl |
| si:ch1073-185p12.23 | 1.1E-06 | 0.7687796 | 0.121 | 0.083 | 0.0367 | DF 7 | si:ch1073-185p12.2 |
| cipcb1 | 1.2E-06 | 0.6400212 | 0.117 | 0.079 | 0.038389 | DF 7 | cipcb |
| irf1b5 | 1.3E-06 | 0.9081632 | 0.159 | 0.117 | 0.041485 | DF 7 | irf1b |
| si:ch211-281l24.32 | 1.4E-06 | 0.3982664 | 0.277 | 0.221 | 0.044761 | DF 7 | si:ch211-281l24.3 |
| fubp12 | 1.4E-06 | 0.4846015 | 0.244 | 0.193 | 0.045439 | DF 7 | fubp1 |
| FP236812.13 | 1.4E-06 | 0.7409368 | 0.14 | 0.099 | 0.047031 | DF 7 | FP236812.1 |
| foxj32 | 1.4E-06 | 0.5185676 | 0.188 | 0.14 | 0.047116 | DF 7 | foxj3 |

| | | | | | | | |
|---|---|---|---|---|---|---|---|
| zfand5a5 | 1.5E-06 | 0.5194333 | 0.238 | 0.188 | 0.048424 | DF 7 | zfand5a |
| tsc22d14 | 1.5E-06 | 0.6824113 | 0.159 | 0.114 | 0.049398 | DF 7 | tsc22d1 |
| gamt1 | 1.6E-06 | 0.6510156 | 0.109 | 0.072 | 0.051202 | DF 7 | gamt |
| arrb2b | 1.6E-06 | 0.7358614 | 0.1 | 0.065 | 0.053622 | DF 7 | arrb2b |
| tril5 | 1.7E-06 | 0.4275618 | 0.151 | 0.106 | 0.05447 | DF 7 | tril |
| rnd3a5 | 1.9E-06 | 0.2537952 | 0.348 | 0.285 | 0.061634 | DF 7 | rnd3a |
| atrnl1b3 | 1.9E-06 | 0.501571 | 0.154 | 0.109 | 0.06176 | DF 7 | atrnl1b |
| camk2g21 | 2E-06 | 0.6282077 | 0.17 | 0.126 | 0.063674 | DF 7 | camk2g2 |
| dnajc5aa3 | 2.1E-06 | 0.6671981 | 0.123 | 0.084 | 0.067858 | DF 7 | dnajc5aa |
| chka4 | 2.1E-06 | 0.5165381 | 0.131 | 0.09 | 0.069737 | DF 7 | chka |
| si:ch211-212k18.95 | 2.3E-06 | 0.3779192 | 0.23 | 0.177 | 0.073217 | DF 7 | si:ch211-212k18.9 |
| mbnl32 | 2.4E-06 | 0.5138164 | 0.179 | 0.133 | 0.07721 | DF 7 | mbnl3 |
| BX284696.11 | 2.4E-06 | 0.6705607 | 0.108 | 0.071 | 0.077283 | DF 7 | BX284696.1 |
| pcyt22 | 2.5E-06 | 0.5968437 | 0.136 | 0.096 | 0.081554 | DF 7 | pcyt2 |
| slc35c23 | 2.6E-06 | 0.5288929 | 0.198 | 0.151 | 0.085199 | DF 7 | slc35c2 |
| sept8a | 2.7E-06 | 0.4688778 | 0.177 | 0.131 | 0.089013 | DF 7 | sept8a |
| rac3b3 | 2.8E-06 | 0.6333045 | 0.121 | 0.082 | 0.08943 | DF 7 | rac3b |
| selenbp13 | 2.8E-06 | 0.6776745 | 0.115 | 0.078 | 0.090204 | DF 7 | selenbp1 |
| zfp36l1b5 | 2.9E-06 | 0.4142652 | 0.577 | 0.546 | 0.095355 | DF 7 | zfp36l1b |
| roraa3 | 3.1E-06 | 0.5785547 | 0.188 | 0.141 | 0.10073 | DF 7 | roraa |
| cdc42bpb1 | 3.1E-06 | 0.5867166 | 0.181 | 0.136 | 0.101713 | DF 7 | cdc42bpb |
| dcbld24 | 3.6E-06 | 0.5934166 | 0.168 | 0.124 | 0.116429 | DF 7 | dcbld2 |
| ywhag16 | 3.7E-06 | 0.5664799 | 0.154 | 0.11 | 0.120847 | DF 7 | ywhag1 |
| thrb2 | 3.7E-06 | 0.4638851 | 0.216 | 0.165 | 0.121326 | DF 7 | thrb |
| rorab4 | 3.7E-06 | 0.5190648 | 0.17 | 0.125 | 0.12185 | DF 7 | rorab |
| setb2 | 3.9E-06 | 0.2971637 | 0.426 | 0.376 | 0.12811 | DF 7 | setb |
| sort1a5 | 4E-06 | 0.5543719 | 0.155 | 0.114 | 0.129862 | DF 7 | sort1a |
| zgc:1122343 | 4.6E-06 | 0.9068605 | 0.109 | 0.074 | 0.149641 | DF 7 | zgc:112234 |
| ube2e34 | 5.1E-06 | 0.3290872 | 0.378 | 0.328 | 0.164543 | DF 7 | ube2e3 |
| grk43 | 5.1E-06 | 0.3956722 | 0.23 | 0.18 | 0.166434 | DF 7 | grk4 |
| pitpnc1b5 | 5.2E-06 | 0.6329055 | 0.118 | 0.081 | 0.168704 | DF 7 | pitpnc1b |
| ifi463 | 5.2E-06 | 0.4631657 | 0.135 | 0.095 | 0.16989 | DF 7 | ifi46 |
| eno3 | 6.7E-06 | 0.3522214 | 0.35 | 0.301 | 0.217542 | DF 7 | eno3 |
| cul4b1 | 6.9E-06 | 0.5980501 | 0.186 | 0.143 | 0.223743 | DF 7 | cul4b |
| angptl44 | 7.7E-06 | 0.2534949 | 0.514 | 0.456 | 0.250833 | DF 7 | angptl4 |
| gpc1b4 | 7.8E-06 | 0.2421568 | 0.46 | 0.401 | 0.255091 | DF 7 | gpc1b |
| ifrd14 | 7.9E-06 | 0.4887329 | 0.327 | 0.282 | 0.255352 | DF 7 | ifrd1 |
| mnta3 | 8.9E-06 | 0.6419755 | 0.14 | 0.102 | 0.290737 | DF 7 | mnta |
| atp1b2a3 | 9.6E-06 | 0.600057 | 0.117 | 0.081 | 0.313688 | DF 7 | atp1b2a |
| vdac14 | 9.7E-06 | 0.3810452 | 0.366 | 0.323 | 0.315737 | DF 7 | vdac1 |
| bcl2l12 | 9.9E-06 | 0.5115226 | 0.2 | 0.157 | 0.322165 | DF 7 | bcl2l1 |
| snai24 | 1.1E-05 | 0.4722036 | 0.145 | 0.105 | 0.345118 | DF 7 | snai2 |
| wipf1b2 | 1.1E-05 | 0.5128697 | 0.144 | 0.105 | 0.366497 | DF 7 | wipf1b |
| ptenb3 | 1.2E-05 | 0.5559236 | 0.218 | 0.175 | 0.392707 | DF 7 | ptenb |
| prrg12 | 1.2E-05 | 0.436826 | 0.252 | 0.207 | 0.401032 | DF 7 | prrg1 |
| srsf2a1 | 1.3E-05 | 0.4018128 | 0.274 | 0.225 | 0.409294 | DF 7 | srsf2a |
| gnas | 1.5E-05 | 0.5336307 | 0.198 | 0.155 | 0.502252 | DF 7 | gnas |
| ccng22 | 1.6E-05 | 0.4709983 | 0.27 | 0.222 | 0.511235 | DF 7 | ccng2 |

| | | | | | | | |
|---|---|---|---|---|---|---|---|
| sh3pxd2aa2 | 1.7E-05 | 0.5476732 | 0.184 | 0.143 | 0.555905 | DF 7 | sh3pxd2aa |
| slc44a1b5 | 1.8E-05 | 0.42277 | 0.16 | 0.119 | 0.5819 | DF 7 | slc44a1b |
| si:dkey-226m8.104 | 1.8E-05 | 0.3459132 | 0.139 | 0.099 | 0.590433 | DF 7 | si:dkey-226m8.10 |
| dennd4a4 | 1.8E-05 | 0.5491394 | 0.122 | 0.086 | 0.591937 | DF 7 | dennd4a |
| prex23 | 1.9E-05 | 0.4765075 | 0.124 | 0.087 | 0.632052 | DF 7 | prex2 |
| rnd3b4 | 2.2E-05 | 0.326155 | 0.38 | 0.331 | 0.711326 | DF 7 | rnd3b |
| arl3b | 2.3E-05 | 0.7356904 | 0.117 | 0.083 | 0.757066 | DF 7 | arl3b |
| zmynd112 | 2.4E-05 | 0.4113324 | 0.181 | 0.139 | 0.770584 | DF 7 | zmynd11 |
| lsp1b5 | 2.5E-05 | 0.2075143 | 0.314 | 0.258 | 0.800964 | DF 7 | lsp1b |
| ninl2 | 2.5E-05 | 0.5461809 | 0.101 | 0.068 | 0.811971 | DF 7 | ninl |
| tcf7l1a4 | 2.5E-05 | 0.4993359 | 0.155 | 0.116 | 0.816129 | DF 7 | tcf7l1a |
| rbbp63 | 2.6E-05 | 0.2809052 | 0.369 | 0.316 | 0.83819 | DF 7 | rbbp6 |
| ppm1k | 2.8E-05 | 0.6419763 | 0.102 | 0.07 | 0.90685 | DF 7 | ppm1k |
| h3f3b.14 | 2.9E-05 | 0.2327953 | 0.533 | 0.495 | 0.944027 | DF 7 | h3f3b.1 |
| sqstm14 | 3E-05 | 0.2482412 | 0.356 | 0.305 | 0.970462 | DF 7 | sqstm1 |
| xpo1b3 | 3.1E-05 | 0.5735886 | 0.138 | 0.102 | 1 | DF 7 | xpo1b |
| BX511129.24 | 3.1E-05 | 0.7574795 | 0.122 | 0.089 | 1 | DF 7 | BX511129.2 |
| itgb25 | 3.4E-05 | 0.375537 | 0.144 | 0.105 | 1 | DF 7 | itgb2 |
| glo1 | 3.7E-05 | 0.2803333 | 0.415 | 0.373 | 1 | DF 7 | glo1 |
| sorl11 | 3.7E-05 | 0.6813551 | 0.103 | 0.072 | 1 | DF 7 | sorl1 |
| top14 | 3.7E-05 | 0.474212 | 0.116 | 0.082 | 1 | DF 7 | top1 |
| isoc2 | 4E-05 | 0.4847141 | 0.144 | 0.107 | 1 | DF 7 | isoc2 |
| phc2a2 | 4E-05 | 0.7885439 | 0.1 | 0.07 | 1 | DF 7 | phc2a |
| msi2b2 | 4.1E-05 | 0.5659808 | 0.133 | 0.098 | 1 | DF 7 | msi2b |
| uchl12 | 4.1E-05 | 0.4807825 | 0.167 | 0.128 | 1 | DF 7 | uchl1 |
| cdc421 | 4.2E-05 | 0.3082555 | 0.418 | 0.376 | 1 | DF 7 | cdc42 |
| b4galt62 | 4.4E-05 | 0.6179887 | 0.141 | 0.105 | 1 | DF 7 | b4galt6 |
| laptm4b | 4.4E-05 | 0.555504 | 0.181 | 0.142 | 1 | DF 7 | laptm4b |
| rap2c1 | 4.4E-05 | 0.4391852 | 0.173 | 0.134 | 1 | DF 7 | rap2c |
| phf21 | 4.4E-05 | 0.3841564 | 0.23 | 0.186 | 1 | DF 7 | phf2 |
| gpx4a3 | 4.5E-05 | 0.2647294 | 0.356 | 0.303 | 1 | DF 7 | gpx4a |
| prickle1b5 | 5.5E-05 | 0.3130849 | 0.222 | 0.178 | 1 | DF 7 | prickle1b |
| nkd13 | 6E-05 | 0.4368971 | 0.117 | 0.083 | 1 | DF 7 | nkd1 |
| fam168b1 | 6.1E-05 | 0.4813553 | 0.163 | 0.126 | 1 | DF 7 | fam168b |
| adam8b3 | 6.2E-05 | 0.4749363 | 0.111 | 0.079 | 1 | DF 7 | adam8b |
| KIF2A1 | 6.6E-05 | 0.4804466 | 0.146 | 0.109 | 1 | DF 7 | KIF2A |
| zmiz1a4 | 6.8E-05 | 0.4149414 | 0.157 | 0.12 | 1 | DF 7 | zmiz1a |
| ptbp1b3 | 7E-05 | 0.478862 | 0.229 | 0.189 | 1 | DF 7 | ptbp1b |
| fam13b3 | 7E-05 | 0.4979196 | 0.156 | 0.119 | 1 | DF 7 | fam13b |
| waca2 | 7.3E-05 | 0.4992567 | 0.157 | 0.12 | 1 | DF 7 | waca |
| lysmd21 | 7.8E-05 | 0.6334589 | 0.113 | 0.082 | 1 | DF 7 | lysmd2 |
| pgrmc21 | 8E-05 | 0.4338375 | 0.178 | 0.139 | 1 | DF 7 | pgrmc2 |
| per32 | 8.3E-05 | 0.2726062 | 0.396 | 0.347 | 1 | DF 7 | per3 |
| ube2d41 | 8.4E-05 | 0.6026184 | 0.117 | 0.084 | 1 | DF 7 | ube2d4 |
| axin13 | 9.1E-05 | 0.4455716 | 0.149 | 0.113 | 1 | DF 7 | axin1 |
| srsf7a2 | 9.3E-05 | 0.410603 | 0.226 | 0.186 | 1 | DF 7 | srsf7a |
| BX005309.14 | 9.8E-05 | 0.5584625 | 0.114 | 0.082 | 1 | DF 7 | BX005309.1 |
| nr1d14 | 0.0001 | 0.5068847 | 0.172 | 0.135 | 1 | DF 7 | nr1d1 |

| | | | | | | | |
|---|---|---|---|---|---|---|---|
| acsl4a4 | 0.0001 | 0.7212224 | 0.108 | 0.078 | 1 | DF 7 | acsl4a |
| mta12 | 0.0001 | 0.5872569 | 0.122 | 0.09 | 1 | DF 7 | mta1 |
| fam120c1 | 0.0001 | 0.3778864 | 0.339 | 0.301 | 1 | DF 7 | fam120c |
| cry1a2 | 0.00011 | 0.3957982 | 0.201 | 0.162 | 1 | DF 7 | cry1a |
| fblim13 | 0.00012 | 0.4449865 | 0.146 | 0.11 | 1 | DF 7 | fblim1 |
| nudt3a | 0.00012 | 0.5452085 | 0.12 | 0.088 | 1 | DF 7 | nudt3a |
| raraa6 | 0.00012 | 0.2057511 | 0.304 | 0.253 | 1 | DF 7 | raraa |
| gdi22 | 0.00012 | 0.2390332 | 0.508 | 0.474 | 1 | DF 7 | gdi2 |
| tmem106ba2 | 0.00012 | 0.3906087 | 0.231 | 0.19 | 1 | DF 7 | tmem106ba |
| gabarapl23 | 0.00013 | 0.4358821 | 0.202 | 0.164 | 1 | DF 7 | gabarapl2 |
| rbm4.12 | 0.00014 | 0.5370347 | 0.133 | 0.1 | 1 | DF 7 | rbm4.1 |
| zgc:77486.15 | 0.00014 | 0.4202723 | 0.209 | 0.169 | 1 | DF 7 | zgc:77486.1 |
| abcf2a4 | 0.00015 | 0.481664 | 0.216 | 0.178 | 1 | DF 7 | abcf2a |
| lasp12 | 0.00016 | 0.2568239 | 0.504 | 0.472 | 1 | DF 7 | lasp1 |
| neu3.26 | 0.00016 | 0.2157304 | 0.271 | 0.222 | 1 | DF 7 | neu3.2 |
| si:dkey-199f5.83 | 0.00016 | 0.4249046 | 0.201 | 0.164 | 1 | DF 7 | si:dkey-199f5.8 |
| dnpep2 | 0.00016 | 0.6025202 | 0.131 | 0.099 | 1 | DF 7 | dnpep |
| ptbp33 | 0.00016 | 0.3955137 | 0.171 | 0.134 | 1 | DF 7 | ptbp3 |
| tmc6b3 | 0.00017 | 0.4890434 | 0.102 | 0.072 | 1 | DF 7 | tmc6b |
| atrx2 | 0.00017 | 0.2125295 | 0.502 | 0.467 | 1 | DF 7 | atrx |
| cblb3 | 0.00018 | 0.5312575 | 0.129 | 0.097 | 1 | DF 7 | cblb |
| sema3e2 | 0.00019 | 0.3439484 | 0.128 | 0.095 | 1 | DF 7 | sema3e |
| palld1 | 0.00019 | 0.4988155 | 0.182 | 0.147 | 1 | DF 7 | palld |
| axin24 | 0.0002 | 0.304993 | 0.264 | 0.22 | 1 | DF 7 | axin2 |
| arpc4l2 | 0.00021 | 0.2770239 | 0.4 | 0.363 | 1 | DF 7 | arpc4l |
| oga3 | 0.00024 | 0.6065393 | 0.128 | 0.097 | 1 | DF 7 | oga |
| sf3b62 | 0.00024 | 0.350811 | 0.319 | 0.282 | 1 | DF 7 | sf3b6 |
| lmcd14 | 0.00024 | 0.568142 | 0.118 | 0.088 | 1 | DF 7 | lmcd1 |
| glsb2 | 0.00025 | 0.4849493 | 0.15 | 0.116 | 1 | DF 7 | glsb |
| kmt2cb1 | 0.00029 | 0.336672 | 0.316 | 0.274 | 1 | DF 7 | kmt2cb |
| mepce3 | 0.00029 | 0.4919818 | 0.135 | 0.103 | 1 | DF 7 | mepce |
| slc3a2b3 | 0.00033 | 0.5477788 | 0.171 | 0.138 | 1 | DF 7 | slc3a2b |
| rusc21 | 0.00033 | 0.5210947 | 0.151 | 0.118 | 1 | DF 7 | rusc2 |
| fkbp32 | 0.00033 | 0.2774761 | 0.396 | 0.358 | 1 | DF 7 | fkbp3 |
| arl6ip12 | 0.00035 | 0.4944769 | 0.196 | 0.161 | 1 | DF 7 | arl6ip1 |
| grb10a4 | 0.00035 | 0.3552375 | 0.141 | 0.109 | 1 | DF 7 | grb10a |
| pik3r23 | 0.00035 | 0.3541762 | 0.239 | 0.201 | 1 | DF 7 | pik3r2 |
| prkar1ab1 | 0.00037 | 0.4857855 | 0.143 | 0.111 | 1 | DF 7 | prkar1ab |
| maff5 | 0.00041 | 0.4511778 | 0.257 | 0.224 | 1 | DF 7 | maff |
| sesn33 | 0.00041 | 0.3341481 | 0.142 | 0.109 | 1 | DF 7 | sesn3 |
| dio14 | 0.00041 | 0.4031492 | 0.156 | 0.122 | 1 | DF 7 | dio1 |
| trnau1apb | 0.00041 | 0.3268856 | 0.199 | 0.162 | 1 | DF 7 | trnau1apb |
| ncoa51 | 0.00043 | 0.524526 | 0.141 | 0.11 | 1 | DF 7 | ncoa5 |
| ehbp1l1b3 | 0.00045 | 0.2729989 | 0.162 | 0.127 | 1 | DF 7 | ehbp1l1b |
| rin32 | 0.00046 | 0.4326595 | 0.189 | 0.154 | 1 | DF 7 | rin3 |
| oaz1b5 | 0.00047 | 0.3182284 | 0.341 | 0.308 | 1 | DF 7 | oaz1b |
| luc7l33 | 0.00048 | 0.450433 | 0.149 | 0.117 | 1 | DF 7 | luc7l3 |
| rif14 | 0.00048 | 0.5511287 | 0.114 | 0.086 | 1 | DF 7 | rif1 |

| | | | | | | | |
|---|---|---|---|---|---|---|---|
| srgap32 | 0.0005 | 0.2241064 | 0.244 | 0.203 | 1 | DF 7 | srgap3 |
| rnf131 | 0.00051 | 0.4736738 | 0.16 | 0.128 | 1 | DF 7 | rnf13 |
| rest1 | 0.00052 | 0.5771183 | 0.111 | 0.083 | 1 | DF 7 | rest |
| plekhg21 | 0.00054 | 0.5418164 | 0.1 | 0.073 | 1 | DF 7 | plekhg2 |
| ndel1b2 | 0.00054 | 0.565252 | 0.126 | 0.098 | 1 | DF 7 | ndel1b |
| ece2b2 | 0.00058 | 0.4325898 | 0.131 | 0.1 | 1 | DF 7 | ece2b |
| gnsa1 | 0.00062 | 0.4791879 | 0.194 | 0.162 | 1 | DF 7 | gnsa |
| clk4a4 | 0.00064 | 0.3204025 | 0.267 | 0.23 | 1 | DF 7 | clk4a |
| vdrb | 0.00065 | 0.4488609 | 0.18 | 0.148 | 1 | DF 7 | vdrb |
| mrtfab2 | 0.00067 | 0.4852147 | 0.114 | 0.086 | 1 | DF 7 | mrtfab |
| srsf5b4 | 0.00067 | 0.4034641 | 0.147 | 0.115 | 1 | DF 7 | srsf5b |
| tcf7l21 | 0.00067 | 0.3934313 | 0.198 | 0.164 | 1 | DF 7 | tcf7l2 |
| lamp22 | 0.00068 | 0.2973253 | 0.383 | 0.349 | 1 | DF 7 | lamp2 |
| kmt2ca1 | 0.0007 | 0.4037994 | 0.205 | 0.17 | 1 | DF 7 | kmt2ca |
| fam168a2 | 0.00073 | 0.3488642 | 0.255 | 0.22 | 1 | DF 7 | fam168a |
| gng73 | 0.00075 | 0.3175751 | 0.199 | 0.165 | 1 | DF 7 | gng7 |
| srsf1b1 | 0.00075 | 0.3364866 | 0.15 | 0.118 | 1 | DF 7 | srsf1b |
| brd3b2 | 0.00076 | 0.3475958 | 0.261 | 0.227 | 1 | DF 7 | brd3b |
| kctd31 | 0.00077 | 0.3905577 | 0.128 | 0.098 | 1 | DF 7 | kctd3 |
| unc119.11 | 0.00078 | 0.3721024 | 0.105 | 0.078 | 1 | DF 7 | unc119.1 |
| prkar1aa2 | 0.00079 | 0.4617805 | 0.175 | 0.144 | 1 | DF 7 | prkar1aa |
| capza1b2 | 0.0008 | 0.3597361 | 0.399 | 0.367 | 1 | DF 7 | capza1b |
| chchd3a | 0.00082 | 0.3508224 | 0.133 | 0.103 | 1 | DF 7 | chchd3a |
| pik3r12 | 0.00083 | 0.3773853 | 0.226 | 0.191 | 1 | DF 7 | pik3r1 |
| mllt103 | 0.00083 | 0.4772889 | 0.109 | 0.082 | 1 | DF 7 | mllt10 |
| ldha4 | 0.00089 | 0.2473275 | 0.268 | 0.23 | 1 | DF 7 | ldha |
| dusp63 | 0.00094 | 0.3753343 | 0.178 | 0.146 | 1 | DF 7 | dusp6 |
| idh12 | 0.00094 | 0.6428768 | 0.121 | 0.094 | 1 | DF 7 | idh1 |
| pabpn12 | 0.00096 | 0.2341869 | 0.439 | 0.411 | 1 | DF 7 | pabpn1 |
| hnrnpm1 | 0.00096 | 0.3122255 | 0.218 | 0.184 | 1 | DF 7 | hnrnpm |
| kmt2a1 | 0.00098 | 0.3471176 | 0.195 | 0.162 | 1 | DF 7 | kmt2a |
| pbxip1a2 | 0.001 | 0.3160093 | 0.223 | 0.187 | 1 | DF 7 | pbxip1a |
| hs3st3b1b4 | 0.00101 | 0.4291416 | 0.16 | 0.129 | 1 | DF 7 | hs3st3b1b |
| phip4 | 0.00102 | 0.3239749 | 0.192 | 0.157 | 1 | DF 7 | phip |
| runx1t14 | 0.00103 | 0.4965897 | 0.118 | 0.091 | 1 | DF 7 | runx1t1 |
| si:ch211-194b1.12 | 0.00106 | 0.345358 | 0.251 | 0.216 | 1 | DF 7 | si:ch211-194b1.1 |
| ENSDARG00000597( | 0.00108 | 0.3567208 | 0.112 | 0.085 | 1 | DF 7 | ENSDARG00000059 |
| gpt2l4 | 0.00113 | 0.334821 | 0.161 | 0.129 | 1 | DF 7 | gpt2l |
| pfkfb34 | 0.00113 | 0.592964 | 0.102 | 0.076 | 1 | DF 7 | pfkfb3 |
| zgc:1368641 | 0.00122 | 0.3986848 | 0.101 | 0.075 | 1 | DF 7 | zgc:136864 |
| sorbs13 | 0.00127 | 0.3645545 | 0.166 | 0.134 | 1 | DF 7 | sorbs1 |
| tmod23 | 0.00127 | 0.3863616 | 0.205 | 0.172 | 1 | DF 7 | tmod2 |
| zgc:1231052 | 0.00139 | 0.5334081 | 0.101 | 0.076 | 1 | DF 7 | zgc:123105 |
| nrp2a2 | 0.00141 | 0.3119153 | 0.16 | 0.128 | 1 | DF 7 | nrp2a |
| arhgap312 | 0.00143 | 0.3646979 | 0.142 | 0.113 | 1 | DF 7 | arhgap31 |
| specc1la2 | 0.00152 | 0.4255453 | 0.105 | 0.08 | 1 | DF 7 | specc1la |
| mpp1 | 0.00158 | 0.5276632 | 0.101 | 0.076 | 1 | DF 7 | mpp1 |
| pcbp22 | 0.00165 | 0.3426112 | 0.254 | 0.225 | 1 | DF 7 | pcbp2 |

| | | | | | | | |
|---|---|---|---|---|---|---|---|
| tp53bp2a | 0.00166 | 0.5156881 | 0.104 | 0.079 | 1 | DF 7 | tp53bp2a |
| tra2a1 | 0.00169 | 0.3384809 | 0.22 | 0.188 | 1 | DF 7 | tra2a |
| phactr25 | 0.00172 | 0.4498821 | 0.17 | 0.141 | 1 | DF 7 | phactr2 |
| sema5a3 | 0.00176 | 0.4766124 | 0.107 | 0.082 | 1 | DF 7 | sema5a |
| ets22 | 0.00179 | 0.4568379 | 0.19 | 0.16 | 1 | DF 7 | ets2 |
| emilin1b3 | 0.00182 | 0.4288489 | 0.112 | 0.086 | 1 | DF 7 | emilin1b |
| add13 | 0.00193 | 0.429053 | 0.167 | 0.138 | 1 | DF 7 | add1 |
| CABZ01074397.14 | 0.00201 | 0.2583158 | 0.182 | 0.149 | 1 | DF 7 | CABZ01074397.1 |
| oser14 | 0.00201 | 0.5860067 | 0.11 | 0.086 | 1 | DF 7 | oser1 |
| nr1d2b4 | 0.00203 | 0.406442 | 0.182 | 0.152 | 1 | DF 7 | nr1d2b |
| ptprsa.13 | 0.00234 | 0.4461048 | 0.152 | 0.125 | 1 | DF 7 | ptprsa.1 |
| eif4g2b1 | 0.00244 | 0.4011943 | 0.221 | 0.191 | 1 | DF 7 | eif4g2b |
| mocs25 | 0.00247 | 0.4900649 | 0.112 | 0.088 | 1 | DF 7 | mocs2 |
| slc29a1a4 | 0.00257 | 0.3183153 | 0.229 | 0.198 | 1 | DF 7 | slc29a1a |
| cad2 | 0.00259 | 0.3785562 | 0.105 | 0.081 | 1 | DF 7 | cad |
| cab39l1 | 0.0026 | 0.4093405 | 0.148 | 0.122 | 1 | DF 7 | cab39l |
| si:ch211-178n15.13 | 0.00263 | 0.337188 | 0.199 | 0.169 | 1 | DF 7 | si:ch211-178n15.1 |
| apcdd1l4 | 0.00273 | 0.3281584 | 0.139 | 0.11 | 1 | DF 7 | apcdd1l |
| nr3c12 | 0.00274 | 0.3421409 | 0.232 | 0.201 | 1 | DF 7 | nr3c1 |
| aldh7a11 | 0.00274 | 0.3155229 | 0.217 | 0.187 | 1 | DF 7 | aldh7a1 |
| srsf111 | 0.00275 | 0.2336318 | 0.277 | 0.244 | 1 | DF 7 | srsf11 |
| fbxo344 | 0.00281 | 0.3213748 | 0.116 | 0.09 | 1 | DF 7 | fbxo34 |
| porb5 | 0.00287 | 0.3705786 | 0.135 | 0.108 | 1 | DF 7 | porb |
| znf292a | 0.00288 | 0.3858693 | 0.184 | 0.155 | 1 | DF 7 | znf292a |
| tet33 | 0.00288 | 0.388306 | 0.124 | 0.099 | 1 | DF 7 | tet3 |
| ube2d2l3 | 0.00293 | 0.3601652 | 0.177 | 0.149 | 1 | DF 7 | ube2d2l |
| zc3h11a | 0.00293 | 0.359112 | 0.172 | 0.143 | 1 | DF 7 | zc3h11a |
| si:ch211-276i12.91 | 0.00294 | 0.5433086 | 0.116 | 0.091 | 1 | DF 7 | si:ch211-276i12.9 |
| nanos15 | 0.00296 | 0.3287626 | 0.237 | 0.205 | 1 | DF 7 | nanos1 |
| rbm4.22 | 0.00297 | 0.3425953 | 0.195 | 0.166 | 1 | DF 7 | rbm4.2 |
| prr12b2 | 0.00299 | 0.3446535 | 0.144 | 0.117 | 1 | DF 7 | prr12b |
| pxk1 | 0.00308 | 0.5046386 | 0.135 | 0.109 | 1 | DF 7 | pxk |
| macroh2a22 | 0.00313 | 0.2123802 | 0.425 | 0.395 | 1 | DF 7 | macroh2a2 |
| rbm52 | 0.00313 | 0.3528632 | 0.217 | 0.186 | 1 | DF 7 | rbm5 |
| tnrc6a5 | 0.00316 | 0.2675168 | 0.305 | 0.272 | 1 | DF 7 | tnrc6a |
| srgap24 | 0.0032 | 0.5116476 | 0.108 | 0.084 | 1 | DF 7 | srgap2 |
| crlf34 | 0.00321 | 0.4712179 | 0.112 | 0.088 | 1 | DF 7 | crlf3 |
| letm23 | 0.00326 | 0.2766604 | 0.221 | 0.188 | 1 | DF 7 | letm2 |
| AL929435.13 | 0.00332 | 0.4291925 | 0.132 | 0.107 | 1 | DF 7 | AL929435.1 |
| nedd4a3 | 0.00338 | 0.412793 | 0.116 | 0.091 | 1 | DF 7 | nedd4a |
| mgst1.13 | 0.00345 | 0.2667234 | 0.284 | 0.25 | 1 | DF 7 | mgst1.1 |
| si:dkey-276j7.14 | 0.00345 | 0.3020009 | 0.197 | 0.168 | 1 | DF 7 | si:dkey-276j7.1 |
| mpp6a3 | 0.00377 | 0.512121 | 0.104 | 0.082 | 1 | DF 7 | mpp6a |
| ugdh4 | 0.00381 | 0.3497035 | 0.207 | 0.178 | 1 | DF 7 | ugdh |
| si:dkey-177p2.63 | 0.00384 | 0.3296985 | 0.29 | 0.263 | 1 | DF 7 | si:dkey-177p2.6 |
| arhgef12b1 | 0.00393 | 0.2533528 | 0.229 | 0.198 | 1 | DF 7 | arhgef12b |
| arglu1a4 | 0.00398 | 0.2852338 | 0.202 | 0.173 | 1 | DF 7 | arglu1a |
| zgc:1735874 | 0.00401 | 0.2983009 | 0.161 | 0.132 | 1 | DF 7 | zgc:173587 |

| | | | | | | | |
|---|---|---|---|---|---|---|---|
| arhgap35a3 | 0.00419 | 0.2682808 | 0.211 | 0.18 | 1 | DF 7 | arhgap35a |
| pip4k2ca | 0.00421 | 0.354888 | 0.175 | 0.147 | 1 | DF 7 | pip4k2ca |
| lmo4a3 | 0.00428 | 0.3916166 | 0.152 | 0.126 | 1 | DF 7 | lmo4a |
| tcf3b1 | 0.0045 | 0.3264022 | 0.249 | 0.221 | 1 | DF 7 | tcf3b |
| igfbp33 | 0.00451 | 0.2772886 | 0.136 | 0.11 | 1 | DF 7 | igfbp3 |
| hspg23 | 0.00468 | 0.3959789 | 0.173 | 0.146 | 1 | DF 7 | hspg2 |
| acox12 | 0.00483 | 0.3184206 | 0.128 | 0.103 | 1 | DF 7 | acox1 |
| cdkn1a5 | 0.00484 | 0.5849107 | 0.146 | 0.122 | 1 | DF 7 | cdkn1a |
| abhd15a3 | 0.0049 | 0.2336862 | 0.161 | 0.132 | 1 | DF 7 | abhd15a |
| ap1s22 | 0.00501 | 0.3893344 | 0.189 | 0.164 | 1 | DF 7 | ap1s2 |
| mef2d3 | 0.00514 | 0.4044855 | 0.214 | 0.188 | 1 | DF 7 | mef2d |
| acin1b | 0.00515 | 0.3465331 | 0.239 | 0.212 | 1 | DF 7 | acin1b |
| cry3a3 | 0.00524 | 0.4634431 | 0.117 | 0.094 | 1 | DF 7 | cry3a |
| tbl1xr1a3 | 0.00534 | 0.294018 | 0.18 | 0.151 | 1 | DF 7 | tbl1xr1a |
| myofl3 | 0.00559 | 0.3708898 | 0.187 | 0.16 | 1 | DF 7 | myofl |
| fam171a2a2 | 0.006 | 0.3854869 | 0.105 | 0.083 | 1 | DF 7 | fam171a2a |
| si:ch211-214j24.102 | 0.006 | 0.3045651 | 0.261 | 0.235 | 1 | DF 7 | si:ch211-214j24.10 |
| ip6k2a4 | 0.00613 | 0.2850009 | 0.226 | 0.197 | 1 | DF 7 | ip6k2a |
| ccnl1b3 | 0.00625 | 0.4337291 | 0.106 | 0.084 | 1 | DF 7 | ccnl1b |
| zgc:1132083 | 0.0066 | 0.3305486 | 0.219 | 0.191 | 1 | DF 7 | zgc:113208 |
| si:ch73-335l21.44 | 0.00671 | 0.3170389 | 0.572 | 0.571 | 1 | DF 7 | si:ch73-335l21.4 |
| sult6b14 | 0.00673 | 0.3349695 | 0.205 | 0.177 | 1 | DF 7 | sult6b1 |
| anxa3b1 | 0.00673 | 0.2735127 | 0.275 | 0.248 | 1 | DF 7 | anxa3b |
| eif4a1a4 | 0.00689 | 0.2489074 | 0.39 | 0.366 | 1 | DF 7 | eif4a1a |
| prr12a1 | 0.00692 | 0.4085882 | 0.13 | 0.107 | 1 | DF 7 | prr12a |
| limk23 | 0.00694 | 0.4723647 | 0.102 | 0.081 | 1 | DF 7 | limk2 |
| fzd7a2 | 0.00709 | 0.4281633 | 0.101 | 0.08 | 1 | DF 7 | fzd7a |
| wdr33 | 0.00728 | 0.385637 | 0.111 | 0.089 | 1 | DF 7 | wdr33 |
| helz1 | 0.00765 | 0.4867671 | 0.109 | 0.087 | 1 | DF 7 | helz |
| mbnl22 | 0.00778 | 0.2616046 | 0.251 | 0.224 | 1 | DF 7 | mbnl2 |
| foxo43 | 0.00814 | 0.3686708 | 0.13 | 0.107 | 1 | DF 7 | foxo4 |
| zc3h43 | 0.00823 | 0.3623116 | 0.163 | 0.139 | 1 | DF 7 | zc3h4 |
| il13ra13 | 0.00833 | 0.3577282 | 0.106 | 0.085 | 1 | DF 7 | il13ra1 |
| zgc:922423 | 0.00848 | 0.3808322 | 0.134 | 0.111 | 1 | DF 7 | zgc:92242 |
| crebbpb1 | 0.00853 | 0.3237687 | 0.174 | 0.149 | 1 | DF 7 | crebbpb |
| kpna31 | 0.00867 | 0.3736819 | 0.176 | 0.152 | 1 | DF 7 | kpna3 |
| st131 | 0.0091 | 0.2596596 | 0.239 | 0.213 | 1 | DF 7 | st13 |
| ENSDARG000000124. | 0.00914 | 0.3098708 | 0.308 | 0.286 | 1 | DF 7 | ENSDARG00000012 |
| atp2b42 | 0.0092 | 0.20008 | 0.383 | 0.356 | 1 | DF 7 | atp2b4 |
| march61 | 0.00928 | 0.4622866 | 0.124 | 0.103 | 1 | DF 7 | march6 |
| paics2 | 0.00929 | 0.4815971 | 0.103 | 0.083 | 1 | DF 7 | paics |
| rnasekb2 | 0.00977 | 0.4246873 | 0.128 | 0.106 | 1 | DF 7 | rnasekb |
| fmr12 | 0.00993 | 0.3233594 | 0.179 | 0.154 | 1 | DF 7 | fmr1 |
| bckdhb1 | 0.00997 | 0.5536035 | 0.102 | 0.082 | 1 | DF 7 | bckdhb |
| clec19a3 | 0 | 5.2393389 | 0.732 | 0.047 | 0 | Synoviocytes | clec19a |
| itih51 | 0 | 4.8973819 | 0.705 | 0.053 | 0 | Synoviocytes | itih5 |
| crabp2a4 | 0 | 3.6234813 | 0.771 | 0.147 | 0 | Synoviocytes | crabp2a |
| krt944 | 0 | 3.4414327 | 0.904 | 0.282 | 0 | Synoviocytes | krt94 |

| | | | | | | | |
|---|---|---|---|---|---|---|---|
| si:dkey-261h17.14 | 0 | 3.3964542 | 0.898 | 0.276 | 0 | Synoviocytes | si:dkey-261h17.1 |
| clic23 | 0 | 4.0701496 | 0.852 | 0.233 | 0 | Synoviocytes | clic2 |
| CU914487.12 | 0 | 4.6141121 | 0.619 | 0.045 | 0 | Synoviocytes | CU914487.1 |
| MYADM.13 | 0 | 3.0248268 | 0.781 | 0.217 | 0 | Synoviocytes | MYADM.1 |
| si:ch211-156j16.15 | 0 | 3.4351939 | 0.693 | 0.14 | 0 | Synoviocytes | si:ch211-156j16.1 |
| dpysl34 | 0 | 3.0697593 | 0.7 | 0.149 | 0 | Synoviocytes | dpysl3 |
| adcyap1b2 | 0 | 6.0218338 | 0.582 | 0.033 | 0 | Synoviocytes | adcyap1b |
| cd9b4 | 0 | 4.1335982 | 0.628 | 0.087 | 0 | Synoviocytes | cd9b |
| im:71514491 | 0 | 4.9844341 | 0.572 | 0.034 | 0 | Synoviocytes | im:7151449 |
| prg4b2 | 0 | 6.7127528 | 0.571 | 0.037 | 0 | Synoviocytes | prg4b |
| hsd3b72 | 0 | 3.125923 | 0.672 | 0.147 | 0 | Synoviocytes | hsd3b7 |
| dub1 | 0 | 4.9930929 | 0.54 | 0.03 | 0 | Synoviocytes | dub |
| tppp32 | 0 | 4.7850781 | 0.549 | 0.047 | 0 | Synoviocytes | tppp3 |
| txn5 | 0 | 2.0950824 | 0.86 | 0.361 | 0 | Synoviocytes | txn |
| si:dkey-172h23.24 | 0 | 3.2702149 | 0.631 | 0.135 | 0 | Synoviocytes | si:dkey-172h23.2 |
| mgp3 | 0 | 4.9445354 | 0.641 | 0.148 | 0 | Synoviocytes | mgp |
| nova24 | 0 | 2.2099247 | 0.693 | 0.203 | 0 | Synoviocytes | nova2 |
| fbln24 | 0 | 3.1924051 | 0.595 | 0.112 | 0 | Synoviocytes | fbln2 |
| fgl2b3 | 0 | 2.9879794 | 0.589 | 0.112 | 0 | Synoviocytes | fgl2b |
| spaca4l2 | 0 | 4.0803951 | 0.54 | 0.071 | 0 | Synoviocytes | spaca4l |
| ackr3b4 | 0 | 3.4817264 | 0.571 | 0.108 | 0 | Synoviocytes | ackr3b |
| fstl1a4 | 0 | 3.1512821 | 0.557 | 0.096 | 0 | Synoviocytes | fstl1a |
| sptbn13 | 0 | 2.665192 | 0.698 | 0.239 | 0 | Synoviocytes | sptbn1 |
| pmp22b3 | 0 | 2.6363101 | 0.806 | 0.352 | 0 | Synoviocytes | pmp22b |
| htra1b4 | 0 | 2.7945254 | 0.85 | 0.402 | 0 | Synoviocytes | htra1b |
| anxa5b3 | 0 | 2.5249519 | 0.834 | 0.393 | 0 | Synoviocytes | anxa5b |
| lamb1b3 | 0 | 2.3419477 | 0.622 | 0.186 | 0 | Synoviocytes | lamb1b |
| ccn2b3 | 0 | 3.2111981 | 0.618 | 0.196 | 0 | Synoviocytes | ccn2b |
| vcana1 | 0 | 5.2248982 | 0.431 | 0.021 | 0 | Synoviocytes | vcana |
| si:ch211-105c13.35 | 0 | 2.2512902 | 0.954 | 0.56 | 0 | Synoviocytes | si:ch211-105c13.3 |
| calm3a4 | 0 | 2.4109402 | 0.871 | 0.493 | 0 | Synoviocytes | calm3a |
| dock9b4 | 0 | 2.6417288 | 0.439 | 0.092 | 0 | Synoviocytes | dock9b |
| gfpt21 | 0 | 3.2808528 | 0.397 | 0.061 | 0 | Synoviocytes | gfpt2 |
| klf2b3 | 0 | 3.5651496 | 0.362 | 0.053 | 0 | Synoviocytes | klf2b |
| BX284638.22 | 0 | 3.8293899 | 0.313 | 0.033 | 0 | Synoviocytes | BX284638.2 |
| mfge8b3 | 0 | 3.2448468 | 0.321 | 0.051 | 0 | Synoviocytes | mfge8b |
| itih11 | 0 | 4.2410247 | 0.289 | 0.025 | 0 | Synoviocytes | itih1 |
| kif1aa | 0 | 4.2080825 | 0.263 | 0.018 | 0 | Synoviocytes | kif1aa |
| zmp:0000000991 | 0 | 3.6169356 | 0.261 | 0.026 | 0 | Synoviocytes | zmp:0000000991 |
| gap43 | 0 | 6.51731 | 0.233 | 0.005 | 0 | Synoviocytes | gap43 |
| specc12 | 0 | 3.593016 | 0.26 | 0.033 | 0 | Synoviocytes | specc1 |
| wnt9a2 | 0 | 3.8886766 | 0.251 | 0.024 | 0 | Synoviocytes | wnt9a |
| sst6 | 0 | 5.7193742 | 0.229 | 0.007 | 0 | Synoviocytes | sst6 |
| sncga | 0 | 4.1299258 | 0.231 | 0.022 | 0 | Synoviocytes | sncga |
| hbegfb | 0 | 5.212005 | 0.206 | 0.012 | 0 | Synoviocytes | hbegfb |
| syt4 | 0 | 5.3326704 | 0.181 | 0.005 | 0 | Synoviocytes | syt4 |
| pdgfd1 | 0 | 3.8514897 | 0.183 | 0.015 | 0 | Synoviocytes | pdgfd |
| sema7a | 0 | 4.5714484 | 0.12 | 0.006 | 0 | Synoviocytes | sema7a |

| | | | | | | | |
|---|---|---|---|---|---|---|---|
| anxa2a4 | 0 | 1.7988205 | 0.991 | 0.889 | 0 | Synoviocytes | anxa2a |
| exoc3l2b | 0 | 4.861015 | 0.106 | 0.005 | 0 | Synoviocytes | exoc3l2b |
| xkr9 | 0 | 5.5756522 | 0.102 | 0.002 | 0 | Synoviocytes | xkr9 |
| b3gnt3.4 | 0 | 5.4460721 | 0.1 | 0.003 | 0 | Synoviocytes | b3gnt3.4 |
| s100a10b3 | 0 | 1.4623044 | 1 | 0.994 | 0 | Synoviocytes | s100a10b |
| ezrb3 | 3E-298 | 2.7275337 | 0.591 | 0.181 | 1E-293 | Synoviocytes | ezrb |
| uap14 | 3E-297 | 2.6338344 | 0.494 | 0.123 | 1E-292 | Synoviocytes | uap1 |
| adam151 | 9E-294 | 2.9464051 | 0.311 | 0.05 | 2.8E-289 | Synoviocytes | adam15 |
| cd1512 | 6E-293 | 2.8108307 | 0.422 | 0.092 | 1.9E-288 | Synoviocytes | cd151 |
| has11 | 8E-290 | 3.3647272 | 0.238 | 0.03 | 2.6E-285 | Synoviocytes | has1 |
| fn1b4 | 8E-289 | 1.6419765 | 0.874 | 0.39 | 2.7E-284 | Synoviocytes | fn1b |
| phlda24 | 2E-285 | 2.7682745 | 0.632 | 0.21 | 7.8E-281 | Synoviocytes | phlda2 |
| crabp2b5 | 7E-284 | 2.081435 | 0.828 | 0.41 | 2.2E-279 | Synoviocytes | crabp2b |
| crip15 | 2E-282 | 2.5284671 | 0.841 | 0.453 | 5.3E-278 | Synoviocytes | crip1 |
| sema3d2 | 2E-277 | 3.1480253 | 0.269 | 0.04 | 6.2E-273 | Synoviocytes | sema3d |
| myof3 | 4E-276 | 2.3256547 | 0.501 | 0.13 | 1.4E-271 | Synoviocytes | myof |
| ehd42 | 1E-275 | 2.9536539 | 0.303 | 0.05 | 3.3E-271 | Synoviocytes | ehd4 |
| crip23 | 4E-259 | 2.6867227 | 0.355 | 0.071 | 1.2E-254 | Synoviocytes | crip2 |
| itga55 | 4E-258 | 2.0967351 | 0.662 | 0.247 | 1.3E-253 | Synoviocytes | itga5 |
| sema3h1 | 9E-258 | 3.2003734 | 0.215 | 0.027 | 2.8E-253 | Synoviocytes | sema3h |
| olfml2ba3 | 2E-256 | 2.4081709 | 0.466 | 0.118 | 4.9E-252 | Synoviocytes | olfml2ba |
| sost4 | 9E-256 | 3.3473962 | 0.351 | 0.072 | 3E-251 | Synoviocytes | sost |
| add3a4 | 4E-255 | 2.0413383 | 0.822 | 0.44 | 1.2E-250 | Synoviocytes | add3a |
| ablim1b6 | 8E-251 | 2.094033 | 0.659 | 0.247 | 2.6E-246 | Synoviocytes | ablim1b |
| rpz54 | 2E-250 | 1.8877527 | 0.725 | 0.297 | 4.9E-246 | Synoviocytes | rpz5 |
| tgfbr2b6 | 2E-247 | 2.4131909 | 0.485 | 0.135 | 7.7E-243 | Synoviocytes | tgfbr2b |
| ckba3 | 7E-245 | 1.9407084 | 0.901 | 0.635 | 2.3E-240 | Synoviocytes | ckba |
| pmp22a3 | 1E-243 | 1.7933791 | 0.918 | 0.684 | 3.6E-239 | Synoviocytes | pmp22a |
| fn1a4 | 3E-241 | 1.9344068 | 0.738 | 0.308 | 9.4E-237 | Synoviocytes | fn1a |
| akap134 | 3E-241 | 2.1045745 | 0.709 | 0.312 | 1.1E-236 | Synoviocytes | akap13 |
| krt44 | 3E-237 | 1.5284859 | 0.961 | 0.754 | 1.1E-232 | Synoviocytes | krt4 |
| emilin2b2 | 5E-237 | 3.0325477 | 0.318 | 0.063 | 1.5E-232 | Synoviocytes | emilin2b |
| cfap45 | 1E-235 | 3.2727642 | 0.174 | 0.019 | 3.4E-231 | Synoviocytes | cfap45 |
| AL732488.22 | 4E-235 | 2.7166738 | 0.307 | 0.058 | 1.2E-230 | Synoviocytes | AL732488.2 |
| bmp64 | 6E-235 | 3.258306 | 0.294 | 0.055 | 2E-230 | Synoviocytes | bmp6 |
| niban1b6 | 9E-235 | 2.4798868 | 0.383 | 0.088 | 3.1E-230 | Synoviocytes | niban1b |
| si:dkeyp-27c8.23 | 7E-226 | 3.2394003 | 0.259 | 0.044 | 2.1E-221 | Synoviocytes | si:dkeyp-27c8.2 |
| fstl1b3 | 2E-222 | 1.5383444 | 0.925 | 0.753 | 7.4E-218 | Synoviocytes | fstl1b |
| anxa1a3 | 1E-218 | 1.2599725 | 0.979 | 0.903 | 3.4E-214 | Synoviocytes | anxa1a |
| rhogb3 | 1E-218 | 2.5465879 | 0.315 | 0.065 | 4E-214 | Synoviocytes | rhogb |
| scara35 | 6E-213 | 2.1210559 | 0.451 | 0.127 | 2E-208 | Synoviocytes | scara3 |
| fxyd14 | 6E-212 | 1.7004752 | 0.847 | 0.566 | 1.8E-207 | Synoviocytes | fxyd1 |
| angptl2a5 | 3E-211 | 1.8958216 | 0.701 | 0.31 | 9.2E-207 | Synoviocytes | angptl2a |
| cd993 | 6E-211 | 1.4138036 | 0.915 | 0.722 | 2.1E-206 | Synoviocytes | cd99 |
| rgcc4 | 1E-206 | 2.1618944 | 0.547 | 0.191 | 3.3E-202 | Synoviocytes | rgcc |
| cd81a3 | 1E-201 | 1.146771 | 0.962 | 0.902 | 4.7E-197 | Synoviocytes | cd81a |
| synm4 | 9E-199 | 1.7907309 | 0.593 | 0.222 | 2.9E-194 | Synoviocytes | synm |
| cthrc1a4 | 3E-196 | 2.035483 | 0.472 | 0.146 | 1.1E-191 | Synoviocytes | cthrc1a |

| | | | | | | | |
|---|---|---|---|---|---|---|---|
| vim6 | 8E-193 | 1.7058794 | 0.739 | 0.362 | 2.7E-188 | Synoviocytes | vim |
| kalrnb2 | 4E-192 | 2.3953172 | 0.349 | 0.086 | 1.2E-187 | Synoviocytes | kalrnb |
| aldh1a24 | 8E-192 | 3.1203046 | 0.258 | 0.05 | 2.7E-187 | Synoviocytes | aldh1a2 |
| adamts15b2 | 8E-191 | 2.6164297 | 0.256 | 0.049 | 2.4E-186 | Synoviocytes | adamts15b |
| fam107b4 | 1E-186 | 1.8448411 | 0.551 | 0.201 | 4.8E-182 | Synoviocytes | fam107b |
| tns2b1 | 5E-186 | 2.8653924 | 0.206 | 0.033 | 1.7E-181 | Synoviocytes | tns2b |
| dpysl2b3 | 1E-183 | 2.0360949 | 0.47 | 0.154 | 4.8E-179 | Synoviocytes | dpysl2b |
| bgna1 | 2E-183 | 1.4759174 | 0.409 | 0.112 | 5.2E-179 | Synoviocytes | bgna |
| CR925728.13 | 4E-181 | 2.5371063 | 0.344 | 0.088 | 1.4E-176 | Synoviocytes | CR925728.1 |
| nccrp12 | 5E-181 | 2.4937758 | 0.595 | 0.268 | 1.5E-176 | Synoviocytes | nccrp1 |
| zbtb7c1 | 1E-176 | 3.6631452 | 0.126 | 0.014 | 3.2E-172 | Synoviocytes | zbtb7c |
| stmn1b4 | 7E-176 | 2.3918619 | 0.398 | 0.117 | 2.2E-171 | Synoviocytes | stmn1b |
| endouc3 | 2E-173 | 2.9438126 | 0.54 | 0.23 | 5.5E-169 | Synoviocytes | endouc |
| ugdh5 | 7E-173 | 2.2265239 | 0.48 | 0.171 | 2.2E-168 | Synoviocytes | ugdh |
| frya1 | 5E-172 | 3.5709869 | 0.145 | 0.018 | 1.5E-167 | Synoviocytes | frya |
| capns1b2 | 2E-170 | 1.613578 | 0.761 | 0.489 | 7E-166 | Synoviocytes | capns1b |
| tns1b4 | 1E-165 | 1.6150988 | 0.694 | 0.36 | 4.4E-161 | Synoviocytes | tns1b |
| tagln24 | 1E-165 | 1.4844326 | 0.881 | 0.661 | 4.6E-161 | Synoviocytes | tagln2 |
| si:ch211-157j23.3 | 3E-165 | 2.922053 | 0.17 | 0.026 | 9.9E-161 | Synoviocytes | si:ch211-157j23.3 |
| si:dkey-238i5.23 | 2E-164 | 2.192014 | 0.297 | 0.07 | 6E-160 | Synoviocytes | si:dkey-238i5.2 |
| qki21 | 8E-162 | 2.3194213 | 0.284 | 0.066 | 2.6E-157 | Synoviocytes | qki2 |
| nav1b2 | 2E-160 | 2.5029247 | 0.281 | 0.067 | 5.9E-156 | Synoviocytes | nav1b |
| peli1a5 | 4E-159 | 1.986715 | 0.399 | 0.123 | 1.2E-154 | Synoviocytes | peli1a |
| pdcd63 | 7E-159 | 1.8017553 | 0.749 | 0.471 | 2.4E-154 | Synoviocytes | pdcd6 |
| map7d1a1 | 2E-158 | 2.1482096 | 0.384 | 0.118 | 5.6E-154 | Synoviocytes | map7d1a |
| capn2a3 | 7E-156 | 1.6475327 | 0.675 | 0.355 | 2.3E-151 | Synoviocytes | capn2a |
| arhgef10lb1 | 1E-155 | 2.3671965 | 0.25 | 0.054 | 4.6E-151 | Synoviocytes | arhgef10lb |
| ndufaf4 | 2E-155 | 3.3386065 | 0.17 | 0.027 | 6.2E-151 | Synoviocytes | ndufaf4 |
| kif5ba.12 | 8E-154 | 2.2042377 | 0.334 | 0.094 | 2.5E-149 | Synoviocytes | kif5ba.1 |
| cox6a21 | 4E-153 | 2.5449098 | 0.221 | 0.044 | 1.3E-148 | Synoviocytes | cox6a2 |
| krt156 | 5E-152 | 1.3532104 | 0.906 | 0.696 | 1.6E-147 | Synoviocytes | krt15 |
| emilin2a3 | 2E-151 | 2.6405578 | 0.363 | 0.114 | 5.3E-147 | Synoviocytes | emilin2a |
| gsna2 | 1E-148 | 1.9870215 | 0.644 | 0.352 | 4.8E-144 | Synoviocytes | gsna |
| agpat42 | 2E-147 | 2.2969475 | 0.259 | 0.061 | 6.7E-143 | Synoviocytes | agpat4 |
| rhoab2 | 7E-147 | 1.5218642 | 0.741 | 0.451 | 2.2E-142 | Synoviocytes | rhoab |
| lhfpl2a5 | 2E-145 | 2.2328478 | 0.342 | 0.102 | 5.8E-141 | Synoviocytes | lhfpl2a |
| klf4 | 4E-145 | 3.1474207 | 0.117 | 0.014 | 1.1E-140 | Synoviocytes | klf4 |
| lpin22 | 6E-145 | 2.6370006 | 0.23 | 0.05 | 1.8E-140 | Synoviocytes | lpin2 |
| calr3b4 | 5E-143 | 1.863881 | 0.573 | 0.263 | 1.8E-138 | Synoviocytes | calr3b |
| gpc1a | 1E-142 | 2.7947197 | 0.173 | 0.03 | 4.4E-138 | Synoviocytes | gpc1a |
| thbs2b5 | 1E-142 | 1.7314911 | 0.562 | 0.244 | 4.9E-138 | Synoviocytes | thbs2b |
| mgll | 1E-141 | 3.0561775 | 0.147 | 0.022 | 3.3E-137 | Synoviocytes | mgll |
| cobll1b3 | 2E-139 | 2.154619 | 0.251 | 0.059 | 7.5E-135 | Synoviocytes | cobll1b |
| abi3bpa5 | 1E-138 | 1.6127891 | 0.61 | 0.293 | 4E-134 | Synoviocytes | abi3bpa |
| hivep14 | 2E-137 | 1.8482683 | 0.416 | 0.148 | 5E-133 | Synoviocytes | hivep1 |
| celf27 | 8E-137 | 1.6768871 | 0.489 | 0.191 | 2.7E-132 | Synoviocytes | celf2 |
| ugp2b3 | 6E-136 | 2.1327101 | 0.352 | 0.112 | 1.9E-131 | Synoviocytes | ugp2b |
| CR318588.43 | 1E-135 | 0.844535 | 0.999 | 0.986 | 4.9E-131 | Synoviocytes | CR318588.4 |

| | | | | | | | |
|---|---|---|---|---|---|---|---|
| tax1bp33 | 2E-135 | 1.6885185 | 0.545 | 0.246 | 7E-131 | Synoviocytes | tax1bp3 |
| myoc3 | 6E-135 | 2.8820992 | 0.257 | 0.065 | 1.8E-130 | Synoviocytes | myoc |
| rabac14 | 1E-134 | 1.5676464 | 0.668 | 0.391 | 3.8E-130 | Synoviocytes | rabac1 |
| si:ch211-1a19.33 | 2E-132 | 1.2621452 | 0.788 | 0.509 | 6.4E-128 | Synoviocytes | si:ch211-1a19.3 |
| ccl442 | 2E-132 | 3.1494344 | 0.361 | 0.123 | 7.9E-128 | Synoviocytes | ccl44 |
| si:ch73-217n20.12 | 4E-131 | 3.2825702 | 0.215 | 0.049 | 1.2E-126 | Synoviocytes | si:ch73-217n20.1 |
| tmsb25 | 1E-130 | 2.2300008 | 0.312 | 0.092 | 4.1E-126 | Synoviocytes | tmsb2 |
| zgc:193505 | 2E-130 | 2.98176 | 0.167 | 0.03 | 4.9E-126 | Synoviocytes | zgc:193505 |
| EPB41L22 | 4E-130 | 1.9404025 | 0.366 | 0.121 | 1.2E-125 | Synoviocytes | EPB41L2 |
| s100w3 | 3E-129 | 1.4364548 | 0.718 | 0.427 | 1.1E-124 | Synoviocytes | s100w |
| s100v23 | 3E-129 | 1.1483482 | 0.853 | 0.675 | 1.1E-124 | Synoviocytes | s100v2 |
| rnd3a6 | 3E-129 | 1.6955665 | 0.576 | 0.279 | 1.1E-124 | Synoviocytes | rnd3a |
| lsp1b6 | 3E-128 | 1.5879402 | 0.553 | 0.251 | 1.1E-123 | Synoviocytes | lsp1b |
| si:busm1-57f23.14 | 4E-127 | 1.2802327 | 0.822 | 0.569 | 1.4E-122 | Synoviocytes | si:busm1-57f23.1 |
| fgf18a2 | 1E-125 | 2.4804715 | 0.194 | 0.041 | 4.6E-121 | Synoviocytes | fgf18a |
| ltbp1 | 2E-125 | 2.6185908 | 0.195 | 0.042 | 5.2E-121 | Synoviocytes | ltbp1 |
| ACBD32 | 7E-125 | 2.8502419 | 0.176 | 0.035 | 2.2E-120 | Synoviocytes | ACBD3 |
| zeb1b4 | 2E-124 | 2.0070678 | 0.292 | 0.083 | 6E-120 | Synoviocytes | zeb1b |
| kank24 | 8E-124 | 1.6516081 | 0.489 | 0.206 | 2.7E-119 | Synoviocytes | kank2 |
| glud1a3 | 3E-122 | 1.3749148 | 0.695 | 0.435 | 1.1E-117 | Synoviocytes | glud1a |
| arsj1 | 7E-122 | 2.4270207 | 0.171 | 0.033 | 2.2E-117 | Synoviocytes | arsj |
| FP102018.14 | 3E-121 | 1.3640021 | 0.673 | 0.385 | 9.4E-117 | Synoviocytes | FP102018.1 |
| sptan13 | 2E-119 | 1.6834824 | 0.518 | 0.239 | 5.8E-115 | Synoviocytes | sptan1 |
| krt18b4 | 9E-119 | 0.8272124 | 0.97 | 0.89 | 2.9E-114 | Synoviocytes | krt18b |
| fam219aa1 | 1E-118 | 3.0816191 | 0.154 | 0.029 | 3.8E-114 | Synoviocytes | fam219aa |
| pamr13 | 2E-118 | 2.0518594 | 0.325 | 0.105 | 7.5E-114 | Synoviocytes | pamr1 |
| mcf2l23 | 3E-118 | 1.9507557 | 0.322 | 0.102 | 8.3E-114 | Synoviocytes | mcf2l2 |
| tuba1a3 | 8E-117 | 1.9027149 | 0.287 | 0.085 | 2.5E-112 | Synoviocytes | tuba1a |
| lrp1aa6 | 9E-117 | 1.0269898 | 0.861 | 0.67 | 2.8E-112 | Synoviocytes | lrp1aa |
| msna3 | 5E-116 | 1.0260449 | 0.885 | 0.748 | 1.7E-111 | Synoviocytes | msna |
| enc14 | 2E-115 | 1.8814586 | 0.351 | 0.119 | 6.8E-111 | Synoviocytes | enc1 |
| si:dkey-27i16.25 | 3E-115 | 2.0204273 | 0.336 | 0.113 | 1.1E-110 | Synoviocytes | si:dkey-27i16.2 |
| si:ch211-227m13.12 | 9E-115 | 2.0252209 | 0.212 | 0.05 | 2.8E-110 | Synoviocytes | si:ch211-227m13.1 |
| igfbp5b5 | 7E-114 | 1.599933 | 0.808 | 0.566 | 2.2E-109 | Synoviocytes | igfbp5b |
| efna5a5 | 4E-112 | 2.1215257 | 0.26 | 0.074 | 1.3E-107 | Synoviocytes | efna5a |
| ahnak2 | 5E-112 | 0.6965306 | 0.994 | 0.961 | 1.7E-107 | Synoviocytes | ahnak |
| elovl1b3 | 3E-111 | 1.3950535 | 0.62 | 0.346 | 8.8E-107 | Synoviocytes | elovl1b |
| tln12 | 6E-110 | 1.0623842 | 0.816 | 0.606 | 2.1E-105 | Synoviocytes | tln1 |
| reep3a4 | 1E-109 | 1.8828142 | 0.401 | 0.16 | 3.7E-105 | Synoviocytes | reep3a |
| ywhaqb2 | 2E-108 | 1.3091729 | 0.743 | 0.533 | 5.6E-104 | Synoviocytes | ywhaqb |
| hapln3 | 5E-108 | 3.3162077 | 0.101 | 0.014 | 1.6E-103 | Synoviocytes | hapln3 |
| si:dkey-282h22.54 | 1E-107 | 1.8071414 | 0.244 | 0.067 | 3.6E-103 | Synoviocytes | si:dkey-282h22.5 |
| mast23 | 3E-107 | 1.6055576 | 0.406 | 0.159 | 1E-102 | Synoviocytes | mast2 |
| jag1b3 | 8E-107 | 1.6839574 | 0.469 | 0.21 | 2.6E-102 | Synoviocytes | jag1b |
| perp2 | 2E-105 | 1.8316126 | 0.487 | 0.232 | 5.9E-101 | Synoviocytes | perp |
| klf6a3 | 1E-104 | 1.0465089 | 0.891 | 0.743 | 4.6E-100 | Synoviocytes | klf6a |
| raph1b5 | 3E-104 | 1.9075939 | 0.307 | 0.102 | 1.1E-99 | Synoviocytes | raph1b |
| cast2 | 8E-104 | 1.0509495 | 0.793 | 0.587 | 2.7E-99 | Synoviocytes | cast |

| | | | | | | | |
|---|---|---|---|---|---|---|---|
| anxa11a4 | 2E-103 | 1.3021014 | 0.662 | 0.405 | 7.9E-99 | Synoviocytes | anxa11a |
| angptl73 | 3E-102 | 2.2095063 | 0.442 | 0.198 | 9.59E-98 | Synoviocytes | angptl7 |
| rasal22 | 2E-101 | 2.050062 | 0.239 | 0.068 | 5.95E-97 | Synoviocytes | rasal2 |
| atp2b43 | 8E-101 | 1.2858167 | 0.619 | 0.35 | 2.62E-96 | Synoviocytes | atp2b4 |
| cfl1l2 | 8E-100 | 1.9752844 | 0.365 | 0.143 | 2.45E-95 | Synoviocytes | cfl1l |
| tmod24 | 1E-99 | 1.5312762 | 0.41 | 0.166 | 3.82E-95 | Synoviocytes | tmod2 |
| htra1a4 | 2E-99 | 1.8863723 | 0.469 | 0.222 | 6.99E-95 | Synoviocytes | htra1a |
| fam102bb | 1E-98 | 2.2003931 | 0.172 | 0.039 | 3.41E-94 | Synoviocytes | fam102bb |
| ackr4b3 | 2.1E-98 | 2.2697845 | 0.193 | 0.048 | 6.9E-94 | Synoviocytes | ackr4b |
| tgm1l15 | 3E-98 | 2.0218429 | 0.333 | 0.123 | 9.76E-94 | Synoviocytes | tgm1l1 |
| kras3 | 5.9E-98 | 1.5664207 | 0.412 | 0.173 | 1.91E-93 | Synoviocytes | kras |
| emp22 | 1.3E-97 | 1.6239097 | 0.375 | 0.148 | 4.11E-93 | Synoviocytes | emp2 |
| TCIM2 | 1.6E-97 | 1.8183417 | 0.224 | 0.063 | 5.28E-93 | Synoviocytes | TCIM |
| CABZ01044281.12 | 4.7E-97 | 2.1726436 | 0.176 | 0.041 | 1.53E-92 | Synoviocytes | CABZ01044281.1 |
| mustn1a3 | 7.5E-97 | 1.853028 | 0.258 | 0.078 | 2.43E-92 | Synoviocytes | mustn1a |
| akap12b3 | 2.7E-96 | 1.9824373 | 0.355 | 0.138 | 8.69E-92 | Synoviocytes | akap12b |
| inhbaa4 | 1.7E-95 | 1.669789 | 0.522 | 0.267 | 5.39E-91 | Synoviocytes | inhbaa |
| itga93 | 5.9E-95 | 1.7232545 | 0.333 | 0.123 | 1.91E-90 | Synoviocytes | itga9 |
| arhgap21a4 | 6.5E-95 | 1.783611 | 0.298 | 0.102 | 2.11E-90 | Synoviocytes | arhgap21a |
| zgc:1750884 | 6.2E-93 | 1.3829425 | 0.524 | 0.268 | 2.01E-88 | Synoviocytes | zgc:175088 |
| aatkb | 6.4E-93 | 2.532373 | 0.12 | 0.022 | 2.09E-88 | Synoviocytes | aatkb |
| ston21 | 6.6E-93 | 2.3006992 | 0.158 | 0.035 | 2.16E-88 | Synoviocytes | ston2 |
| si:ch211-165e15.14 | 1.4E-92 | 2.0845307 | 0.222 | 0.064 | 4.45E-88 | Synoviocytes | si:ch211-165e15.1 |
| ace2 | 1.5E-92 | 1.9159766 | 0.189 | 0.048 | 4.82E-88 | Synoviocytes | ace |
| hbegfa4 | 2.6E-92 | 1.4702744 | 0.653 | 0.397 | 8.48E-88 | Synoviocytes | hbegfa |
| rhoaa3 | 9E-92 | 1.2322981 | 0.646 | 0.416 | 2.94E-87 | Synoviocytes | rhoaa |
| map1ab5 | 1.5E-91 | 2.0490995 | 0.421 | 0.192 | 5.01E-87 | Synoviocytes | map1ab |
| pkig4 | 2.7E-91 | 1.8204002 | 0.311 | 0.114 | 8.68E-87 | Synoviocytes | pkig |
| bag2 | 1.9E-90 | 2.7778837 | 0.112 | 0.02 | 6.02E-86 | Synoviocytes | bag2 |
| si:ch73-119p20.1 | 6.8E-90 | 2.5789046 | 0.142 | 0.03 | 2.21E-85 | Synoviocytes | si:ch73-119p20.1 |
| mid1ip1b4 | 7.4E-90 | 1.684738 | 0.315 | 0.115 | 2.39E-85 | Synoviocytes | mid1ip1b |
| hspb8 | 1.2E-88 | 2.3664265 | 0.157 | 0.036 | 3.91E-84 | Synoviocytes | hspb8 |
| mt-atp62 | 2.2E-88 | 0.6060938 | 0.994 | 0.965 | 7.22E-84 | Synoviocytes | mt-atp6 |
| ano65 | 2.5E-88 | 1.6581829 | 0.322 | 0.121 | 8.02E-84 | Synoviocytes | ano6 |
| cst14a.23 | 5.7E-88 | 1.2644763 | 0.604 | 0.358 | 1.84E-83 | Synoviocytes | cst14a.2 |
| anxa132 | 4.7E-87 | 1.3715108 | 0.58 | 0.34 | 1.52E-82 | Synoviocytes | anxa13 |
| zgc:77486.16 | 6E-87 | 1.6542612 | 0.382 | 0.165 | 1.97E-82 | Synoviocytes | zgc:77486.1 |
| cyb5r13 | 4.4E-86 | 1.5104871 | 0.423 | 0.196 | 1.43E-81 | Synoviocytes | cyb5r1 |
| scara55 | 7.1E-86 | 1.5624936 | 0.362 | 0.147 | 2.32E-81 | Synoviocytes | scara5 |
| scinla4 | 1.4E-84 | 1.5545708 | 0.635 | 0.405 | 4.54E-80 | Synoviocytes | scinla |
| tuba1c | 3.5E-84 | 2.3416616 | 0.16 | 0.039 | 1.14E-79 | Synoviocytes | tuba1c |
| tmem198a | 6.3E-84 | 2.4909965 | 0.122 | 0.024 | 2.05E-79 | Synoviocytes | tmem198a |
| sulf15 | 1.1E-83 | 1.8085388 | 0.346 | 0.143 | 3.69E-79 | Synoviocytes | sulf1 |
| mfap56 | 1.4E-83 | 0.9148261 | 0.804 | 0.54 | 4.5E-79 | Synoviocytes | mfap5 |
| hspb15 | 2.5E-82 | 1.549472 | 0.39 | 0.175 | 8.08E-78 | Synoviocytes | hspb1 |
| arl6ip5b3 | 3.1E-82 | 1.0191374 | 0.709 | 0.505 | 1.02E-77 | Synoviocytes | arl6ip5b |
| glis2a4 | 5.4E-82 | 1.4723766 | 0.426 | 0.199 | 1.76E-77 | Synoviocytes | glis2a |
| icn2 | 1.4E-81 | 0.6017114 | 0.999 | 0.996 | 4.63E-77 | Synoviocytes | icn |

| | | | | | | | |
|---|---|---|---|---|---|---|---|
| cyth1a.13 | 3.2E-81 | 1.6115089 | 0.33 | 0.132 | 1.05E-76 | Synoviocytes | cyth1a.1 |
| june | 6.2E-81 | 2.1133266 | 0.132 | 0.028 | 2.01E-76 | Synoviocytes | june |
| afdna1 | 7.9E-81 | 1.4612267 | 0.414 | 0.189 | 2.58E-76 | Synoviocytes | afdna |
| csnk1e | 2.2E-80 | 2.5306247 | 0.118 | 0.024 | 7.26E-76 | Synoviocytes | csnk1e |
| dcbld25 | 8.7E-78 | 1.6106249 | 0.309 | 0.121 | 2.82E-73 | Synoviocytes | dcbld2 |
| znf385a2 | 5.2E-77 | 2.2720143 | 0.175 | 0.049 | 1.68E-72 | Synoviocytes | znf385a |
| cremb1 | 9.9E-77 | 2.2639499 | 0.16 | 0.041 | 3.22E-72 | Synoviocytes | cremb |
| gpr1 | 1E-76 | 2.4240985 | 0.109 | 0.021 | 3.29E-72 | Synoviocytes | gpr1 |
| iqsec1b4 | 1.2E-76 | 1.1545534 | 0.577 | 0.339 | 3.96E-72 | Synoviocytes | iqsec1b |
| si:ch73-138n13.13 | 1.7E-75 | 1.1167518 | 0.613 | 0.372 | 5.55E-71 | Synoviocytes | si:ch73-138n13.1 |
| rab103 | 1.3E-73 | 0.9806307 | 0.667 | 0.453 | 4.38E-69 | Synoviocytes | rab10 |
| clip3 | 2.8E-73 | 2.2837064 | 0.114 | 0.024 | 9.18E-69 | Synoviocytes | clip3 |
| foxc1b3 | 2.9E-73 | 1.890485 | 0.197 | 0.06 | 9.27E-69 | Synoviocytes | foxc1b |
| sema4ba3 | 3.7E-73 | 1.9511424 | 0.235 | 0.081 | 1.2E-68 | Synoviocytes | sema4ba |
| slit22 | 6.4E-73 | 1.8921721 | 0.209 | 0.066 | 2.07E-68 | Synoviocytes | slit2 |
| klf12b | 5.4E-72 | 2.6921433 | 0.1 | 0.019 | 1.75E-67 | Synoviocytes | klf12b |
| ywhaqa3 | 8.1E-72 | 1.4173239 | 0.477 | 0.269 | 2.62E-67 | Synoviocytes | ywhaqa |
| sdcbp23 | 1.2E-71 | 0.8276138 | 0.771 | 0.595 | 3.92E-67 | Synoviocytes | sdcbp2 |
| cdc42l2 | 1.5E-71 | 1.0324127 | 0.62 | 0.4 | 4.74E-67 | Synoviocytes | cdc42l |
| klf8 | 2.9E-71 | 2.3833514 | 0.114 | 0.024 | 9.57E-67 | Synoviocytes | klf8 |
| lgals3b3 | 3.3E-71 | 1.3410101 | 0.586 | 0.404 | 1.07E-66 | Synoviocytes | lgals3b |
| abi1b1 | 7.2E-71 | 2.1075805 | 0.13 | 0.031 | 2.35E-66 | Synoviocytes | abi1b |
| atf5b2 | 1.3E-70 | 1.4465792 | 0.457 | 0.248 | 4.28E-66 | Synoviocytes | atf5b |
| pfn24 | 4.2E-70 | 0.8094515 | 0.74 | 0.534 | 1.36E-65 | Synoviocytes | pfn2 |
| cyfip11 | 9E-70 | 1.3032383 | 0.416 | 0.205 | 2.92E-65 | Synoviocytes | cyfip1 |
| si:dkey-19b23.82 | 9.8E-70 | 2.220374 | 0.176 | 0.052 | 3.18E-65 | Synoviocytes | si:dkey-19b23.8 |
| hdac45 | 1.2E-69 | 1.6071964 | 0.293 | 0.117 | 3.8E-65 | Synoviocytes | hdac4 |
| myo1ca4 | 1.5E-69 | 1.2444454 | 0.414 | 0.198 | 4.91E-65 | Synoviocytes | myo1ca |
| ccdc804 | 5E-69 | 1.0882149 | 0.61 | 0.382 | 1.62E-64 | Synoviocytes | ccdc80 |
| c1qtnf24 | 1.2E-68 | 1.5000458 | 0.248 | 0.089 | 4.01E-64 | Synoviocytes | c1qtnf2 |
| cpne12 | 1.2E-68 | 0.9379244 | 0.669 | 0.466 | 4.06E-64 | Synoviocytes | cpne1 |
| vwa15 | 2.8E-68 | 2.0133771 | 0.31 | 0.135 | 9.1E-64 | Synoviocytes | vwa1 |
| pld75 | 6.8E-68 | 1.6528643 | 0.333 | 0.149 | 2.2E-63 | Synoviocytes | pld7 |
| foxp46 | 6.9E-68 | 1.1714426 | 0.412 | 0.199 | 2.26E-63 | Synoviocytes | foxp4 |
| rras21 | 9.6E-68 | 1.8033434 | 0.201 | 0.065 | 3.13E-63 | Synoviocytes | rras2 |
| myo9aa1 | 1.6E-67 | 1.8181072 | 0.21 | 0.07 | 5.08E-63 | Synoviocytes | myo9aa |
| cpne23 | 5.9E-67 | 1.8464567 | 0.198 | 0.064 | 1.92E-62 | Synoviocytes | cpne2 |
| osr15 | 1.3E-66 | 1.5334664 | 0.361 | 0.168 | 4.17E-62 | Synoviocytes | osr1 |
| rhocb2 | 1.4E-66 | 1.4494725 | 0.329 | 0.143 | 4.49E-62 | Synoviocytes | rhocb |
| ppfibp2b | 1.8E-66 | 1.7293807 | 0.129 | 0.031 | 5.94E-62 | Synoviocytes | ppfibp2b |
| ablim22 | 4.4E-66 | 1.7733931 | 0.161 | 0.045 | 1.43E-61 | Synoviocytes | ablim2 |
| angptl1a4 | 7.6E-66 | 1.0761178 | 0.609 | 0.388 | 2.46E-61 | Synoviocytes | angptl1a |
| pdia55 | 8.1E-66 | 1.4555054 | 0.351 | 0.162 | 2.65E-61 | Synoviocytes | pdia5 |
| cst33 | 1.3E-65 | 0.7771956 | 0.835 | 0.722 | 4.07E-61 | Synoviocytes | cst3 |
| prnprs33 | 4.6E-65 | 1.7097793 | 0.247 | 0.093 | 1.51E-60 | Synoviocytes | prnprs3 |
| cdc42se12 | 8.2E-65 | 1.7017003 | 0.216 | 0.074 | 2.67E-60 | Synoviocytes | cdc42se1 |
| ctnnbip11 | 9E-65 | 2.0481037 | 0.148 | 0.041 | 2.91E-60 | Synoviocytes | ctnnbip1 |
| HACD42 | 1.5E-64 | 1.42395 | 0.309 | 0.131 | 4.92E-60 | Synoviocytes | HACD4 |

| | | | | | | | |
|---|---|---|---|---|---|---|---|
| lgals2a4 | 6.6E-64 | 1.1306726 | 0.782 | 0.638 | 2.16E-59 | Synoviocytes | lgals2a |
| inavaa | 9.2E-64 | 2.7181286 | 0.101 | 0.021 | 2.98E-59 | Synoviocytes | inavaa |
| ehbp1l1b4 | 1.2E-63 | 1.4316479 | 0.297 | 0.123 | 4.04E-59 | Synoviocytes | ehbp1l1b |
| klf2a4 | 3.6E-63 | 1.2543596 | 0.493 | 0.281 | 1.16E-58 | Synoviocytes | klf2a |
| pam2 | 1.5E-62 | 1.610299 | 0.247 | 0.094 | 4.79E-58 | Synoviocytes | pam |
| esyt2a2 | 8.7E-62 | 1.3229319 | 0.352 | 0.166 | 2.83E-57 | Synoviocytes | esyt2a |
| plecb4 | 1.2E-61 | 0.788834 | 0.8 | 0.655 | 3.76E-57 | Synoviocytes | plecb |
| sema3fb | 1.6E-61 | 2.3390132 | 0.102 | 0.022 | 5.23E-57 | Synoviocytes | sema3fb |
| clic13 | 3.3E-61 | 1.4402275 | 0.258 | 0.101 | 1.08E-56 | Synoviocytes | clic1 |
| schip14 | 6.9E-61 | 1.3982042 | 0.316 | 0.142 | 2.26E-56 | Synoviocytes | schip1 |
| adam93 | 1E-60 | 1.729914 | 0.193 | 0.065 | 3.32E-56 | Synoviocytes | adam9 |
| rhoca2 | 1.8E-60 | 1.1523621 | 0.458 | 0.253 | 5.7E-56 | Synoviocytes | rhoca |
| dynlt32 | 1E-59 | 1.2498822 | 0.407 | 0.217 | 3.4E-55 | Synoviocytes | dynlt3 |
| gpc1b5 | 1.4E-59 | 0.9020326 | 0.613 | 0.397 | 4.59E-55 | Synoviocytes | gpc1b |
| si:dkey-25e12.32 | 1.5E-59 | 1.8426603 | 0.2 | 0.07 | 4.75E-55 | Synoviocytes | si:dkey-25e12.3 |
| lama43 | 3.6E-59 | 1.1518389 | 0.469 | 0.267 | 1.18E-54 | Synoviocytes | lama4 |
| ywhaz1 | 4.4E-59 | 0.8698291 | 0.693 | 0.527 | 1.44E-54 | Synoviocytes | ywhaz |
| dync1li22 | 6.2E-59 | 1.3727047 | 0.34 | 0.161 | 2.02E-54 | Synoviocytes | dync1li2 |
| tnfaip63 | 2.3E-58 | 1.7344588 | 0.607 | 0.428 | 7.58E-54 | Synoviocytes | tnfaip6 |
| thbs3a4 | 7.2E-58 | 0.916966 | 0.524 | 0.304 | 2.33E-53 | Synoviocytes | thbs3a |
| rtn1a1 | 1E-57 | 1.2516948 | 0.412 | 0.219 | 3.4E-53 | Synoviocytes | rtn1a |
| asap1b4 | 1.4E-57 | 1.4340729 | 0.321 | 0.148 | 4.42E-53 | Synoviocytes | asap1b |
| tlr5b3 | 1.6E-57 | 1.7233724 | 0.201 | 0.071 | 5.13E-53 | Synoviocytes | tlr5b |
| klf33 | 1.7E-57 | 1.7807782 | 0.243 | 0.098 | 5.39E-53 | Synoviocytes | klf3 |
| slc25a19 | 2E-57 | 1.8878764 | 0.141 | 0.04 | 6.62E-53 | Synoviocytes | slc25a19 |
| siae | 2.2E-57 | 2.1625346 | 0.126 | 0.034 | 7.19E-53 | Synoviocytes | siae |
| prelp6 | 2.8E-57 | 1.0773223 | 0.303 | 0.13 | 9.14E-53 | Synoviocytes | prelp |
| iqgap13 | 7.3E-57 | 1.1424304 | 0.42 | 0.222 | 2.37E-52 | Synoviocytes | iqgap1 |
| sgcg4 | 1.2E-56 | 1.4357238 | 0.231 | 0.089 | 4.02E-52 | Synoviocytes | sgcg |
| col5a3a1 | 1.4E-56 | 1.2390106 | 0.191 | 0.065 | 4.47E-52 | Synoviocytes | col5a3a |
| dgkaa1 | 7.8E-56 | 2.1014661 | 0.125 | 0.033 | 2.53E-51 | Synoviocytes | dgkaa |
| si:ch211-195m9.31 | 2.2E-55 | 2.7101605 | 0.143 | 0.043 | 7.22E-51 | Synoviocytes | si:ch211-195m9.3 |
| fpr13 | 3.7E-55 | 1.1443902 | 0.406 | 0.214 | 1.22E-50 | Synoviocytes | fpr1 |
| chmp5b2 | 7.6E-55 | 1.1391021 | 0.417 | 0.226 | 2.48E-50 | Synoviocytes | chmp5b |
| tmem1894 | 1.7E-54 | 1.4118612 | 0.288 | 0.128 | 5.42E-50 | Synoviocytes | tmem189 |
| si:ch73-173h19.35 | 2.2E-54 | 1.158901 | 0.432 | 0.235 | 7.06E-50 | Synoviocytes | si:ch73-173h19.3 |
| ebf3a.11 | 2.3E-54 | 1.8008202 | 0.15 | 0.046 | 7.36E-50 | Synoviocytes | ebf3a.1 |
| fundc23 | 2.7E-54 | 1.0843693 | 0.489 | 0.298 | 8.7E-50 | Synoviocytes | fundc2 |
| fhod14 | 3.1E-54 | 1.6239713 | 0.211 | 0.079 | 9.93E-50 | Synoviocytes | fhod1 |
| anxa3b2 | 3.7E-54 | 1.1088158 | 0.435 | 0.244 | 1.22E-49 | Synoviocytes | anxa3b |
| pim14 | 9E-54 | 0.9216368 | 0.732 | 0.541 | 2.92E-49 | Synoviocytes | pim1 |
| arhgef404 | 9.7E-54 | 1.1770589 | 0.37 | 0.188 | 3.15E-49 | Synoviocytes | arhgef40 |
| c1qtnf54 | 1.3E-53 | 1.7721731 | 0.309 | 0.149 | 4.14E-49 | Synoviocytes | c1qtnf5 |
| fbxo345 | 4.5E-53 | 1.4972781 | 0.224 | 0.087 | 1.48E-48 | Synoviocytes | fbxo34 |
| pafah1b1b2 | 5.2E-53 | 1.1487293 | 0.386 | 0.204 | 1.7E-48 | Synoviocytes | pafah1b1b |
| si:dkey-112a7.44 | 8.9E-53 | 1.8663133 | 0.216 | 0.086 | 2.89E-48 | Synoviocytes | si:dkey-112a7.4 |
| nucb13 | 1E-52 | 1.1276281 | 0.386 | 0.204 | 3.24E-48 | Synoviocytes | nucb1 |
| map7d1b3 | 1.1E-52 | 1.2319204 | 0.363 | 0.186 | 3.45E-48 | Synoviocytes | map7d1b |

| | | | | | | | |
|---|---|---|---|---|---|---|---|
| si:dkeyp-23e4.36 | 7.3E-52 | 1.4168314 | 0.216 | 0.083 | 2.37E-47 | Synoviocytes | si:dkeyp-23e4.3 |
| tuba8l46 | 2E-51 | 0.7218639 | 0.823 | 0.732 | 6.61E-47 | Synoviocytes | tuba8l4 |
| ywhah3 | 4.2E-51 | 1.2061469 | 0.327 | 0.157 | 1.35E-46 | Synoviocytes | ywhah |
| itgb1a3 | 2.7E-50 | 1.1678144 | 0.443 | 0.261 | 8.77E-46 | Synoviocytes | itgb1a |
| ehd2b3 | 3.4E-50 | 0.7106704 | 0.727 | 0.557 | 1.11E-45 | Synoviocytes | ehd2b |
| atl34 | 3.8E-50 | 1.0729093 | 0.499 | 0.319 | 1.24E-45 | Synoviocytes | atl3 |
| slc35e42 | 4.9E-50 | 1.5915637 | 0.2 | 0.076 | 1.6E-45 | Synoviocytes | slc35e4 |
| rca2.12 | 9.5E-50 | 1.0501444 | 0.466 | 0.281 | 3.08E-45 | Synoviocytes | rca2.1 |
| tuba8l32 | 1.2E-49 | 1.0219325 | 0.585 | 0.422 | 3.81E-45 | Synoviocytes | tuba8l3 |
| vasna3 | 1.7E-49 | 1.3539876 | 0.258 | 0.112 | 5.37E-45 | Synoviocytes | vasna |
| cavin2b5 | 2.6E-49 | 0.6854887 | 0.714 | 0.527 | 8.49E-45 | Synoviocytes | cavin2b |
| jpt1b2 | 6E-49 | 0.5961183 | 0.824 | 0.689 | 1.94E-44 | Synoviocytes | jpt1b |
| otud11 | 3E-48 | 1.8623528 | 0.163 | 0.056 | 9.8E-44 | Synoviocytes | otud1 |
| arhgef43 | 1E-47 | 1.6101477 | 0.193 | 0.073 | 3.39E-43 | Synoviocytes | arhgef4 |
| ENSDARG000001018 | 2.8E-47 | 1.6992485 | 0.137 | 0.043 | 9.11E-43 | Synoviocytes | ENSDARG00000101 |
| plscr3b3 | 3.2E-47 | 1.0681063 | 0.417 | 0.237 | 1.03E-42 | Synoviocytes | plscr3b |
| kank34 | 5.4E-47 | 1.4849731 | 0.199 | 0.078 | 1.75E-42 | Synoviocytes | kank3 |
| trak1a2 | 9.8E-47 | 1.532316 | 0.165 | 0.058 | 3.19E-42 | Synoviocytes | trak1a |
| CU929259.12 | 1.9E-46 | 1.0703641 | 0.489 | 0.314 | 6.09E-42 | Synoviocytes | CU929259.1 |
| camsap2b3 | 2.2E-46 | 1.363333 | 0.241 | 0.105 | 7.15E-42 | Synoviocytes | camsap2b |
| plekho24 | 2.5E-46 | 1.1992269 | 0.344 | 0.179 | 8.04E-42 | Synoviocytes | plekho2 |
| rras3 | 5.9E-46 | 1.2494342 | 0.315 | 0.158 | 1.91E-41 | Synoviocytes | rras |
| plod25 | 8.8E-46 | 1.5787906 | 0.183 | 0.07 | 2.86E-41 | Synoviocytes | plod2 |
| ddr2a4 | 9.4E-46 | 1.1644766 | 0.374 | 0.204 | 3.06E-41 | Synoviocytes | ddr2a |
| s100t | 1.6E-45 | 1.9996041 | 0.102 | 0.027 | 5.17E-41 | Synoviocytes | s100t |
| dip2ba1 | 2.1E-45 | 1.4950027 | 0.189 | 0.073 | 6.91E-41 | Synoviocytes | dip2ba |
| nfixb6 | 3E-45 | 1.0148371 | 0.467 | 0.291 | 9.87E-41 | Synoviocytes | nfixb |
| cdc42ep52 | 6.2E-45 | 1.0026178 | 0.479 | 0.297 | 2.01E-40 | Synoviocytes | cdc42ep5 |
| stxbp62 | 6.4E-45 | 1.8268578 | 0.142 | 0.047 | 2.07E-40 | Synoviocytes | stxbp6 |
| rhoac3 | 6.4E-45 | 0.9593701 | 0.468 | 0.292 | 2.08E-40 | Synoviocytes | rhoac |
| syngr2a1 | 6.9E-45 | 1.6032811 | 0.169 | 0.062 | 2.25E-40 | Synoviocytes | syngr2a |
| numbl | 7.5E-45 | 2.0101396 | 0.1 | 0.027 | 2.45E-40 | Synoviocytes | numbl |
| arpc33 | 2E-44 | 0.9010152 | 0.552 | 0.381 | 6.57E-40 | Synoviocytes | arpc3 |
| igfbp2b2 | 2.2E-44 | 1.4924514 | 0.18 | 0.068 | 7.03E-40 | Synoviocytes | igfbp2b |
| anxa44 | 3.1E-44 | 0.6831245 | 0.801 | 0.703 | 1.02E-39 | Synoviocytes | anxa4 |
| ehd1a2 | 3.5E-44 | 1.5292714 | 0.17 | 0.063 | 1.14E-39 | Synoviocytes | ehd1a |
| gas1a5 | 3.6E-44 | 1.2796623 | 0.309 | 0.157 | 1.17E-39 | Synoviocytes | gas1a |
| myl12.11 | 7.1E-44 | 0.7287394 | 0.68 | 0.532 | 2.29E-39 | Synoviocytes | myl12.1 |
| epn24 | 7.6E-44 | 1.3500373 | 0.239 | 0.107 | 2.48E-39 | Synoviocytes | epn2 |
| vat14 | 8E-44 | 1.0024201 | 0.493 | 0.312 | 2.61E-39 | Synoviocytes | vat1 |
| pfn2l2 | 1.6E-43 | 0.7359861 | 0.642 | 0.474 | 5.17E-39 | Synoviocytes | pfn2l |
| smim14 | 2E-43 | 1.8327958 | 0.141 | 0.048 | 6.44E-39 | Synoviocytes | smim14 |
| TNC1 | 2E-43 | 2.0345972 | 0.12 | 0.037 | 6.5E-39 | Synoviocytes | TNC |
| zcchc246 | 2.4E-43 | 1.1438473 | 0.422 | 0.253 | 7.91E-39 | Synoviocytes | zcchc24 |
| dynlrb1.13 | 3.6E-43 | 0.9531174 | 0.482 | 0.314 | 1.17E-38 | Synoviocytes | dynlrb1.1 |
| mt-nd12 | 8.2E-43 | 0.6551975 | 0.839 | 0.716 | 2.68E-38 | Synoviocytes | mt-nd1 |
| cd633 | 1.2E-42 | 0.4110266 | 0.898 | 0.826 | 4.01E-38 | Synoviocytes | cd63 |
| zgc:154093 | 3.5E-42 | 1.6011441 | 0.137 | 0.046 | 1.13E-37 | Synoviocytes | zgc:154093 |

| | | | | | | | |
|---|---|---|---|---|---|---|---|
| sh3pxd2aa3 | 4.4E-42 | 1.2599261 | 0.284 | 0.14 | 1.42E-37 | Synoviocytes | sh3pxd2aa |
| si:dkey-42i9.62 | 7.3E-42 | 1.1902274 | 0.348 | 0.192 | 2.37E-37 | Synoviocytes | si:dkey-42i9.6 |
| rapgef1a | 8.2E-42 | 1.6778048 | 0.125 | 0.04 | 2.67E-37 | Synoviocytes | rapgef1a |
| serpinb144 | 1.5E-41 | 1.0718913 | 0.302 | 0.15 | 4.81E-37 | Synoviocytes | serpinb14 |
| rasgef1bb5 | 1.9E-41 | 1.1159737 | 0.301 | 0.152 | 6.31E-37 | Synoviocytes | rasgef1bb |
| atf4b7 | 2.5E-41 | 0.6688103 | 0.757 | 0.632 | 8.01E-37 | Synoviocytes | atf4b |
| ramp26 | 5.2E-41 | 1.3084767 | 0.282 | 0.139 | 1.68E-36 | Synoviocytes | ramp2 |
| serinc22 | 1.5E-40 | 0.8526947 | 0.488 | 0.314 | 4.84E-36 | Synoviocytes | serinc2 |
| arpc5a2 | 1.6E-40 | 1.1226321 | 0.298 | 0.152 | 5.06E-36 | Synoviocytes | arpc5a |
| rab1ab2 | 1.6E-40 | 0.78548 | 0.581 | 0.417 | 5.07E-36 | Synoviocytes | rab1ab |
| reep52 | 1.6E-40 | 0.6753791 | 0.69 | 0.538 | 5.18E-36 | Synoviocytes | reep5 |
| palm1b1 | 2.2E-40 | 1.4397902 | 0.196 | 0.082 | 7.21E-36 | Synoviocytes | palm1b |
| dagla | 3.1E-40 | 1.8142017 | 0.102 | 0.03 | 9.99E-36 | Synoviocytes | dagla |
| cfl12 | 5.8E-40 | 0.4752816 | 0.898 | 0.842 | 1.9E-35 | Synoviocytes | cfl1 |
| rac1a2 | 6.9E-40 | 1.0789632 | 0.337 | 0.184 | 2.25E-35 | Synoviocytes | rac1a |
| mt-nd22 | 8.8E-40 | 0.6408719 | 0.77 | 0.628 | 2.85E-35 | Synoviocytes | mt-nd2 |
| sri2 | 1.1E-39 | 0.9261181 | 0.437 | 0.268 | 3.56E-35 | Synoviocytes | sri |
| serpinb1l32 | 1.1E-39 | 1.3410969 | 0.203 | 0.087 | 3.67E-35 | Synoviocytes | serpinb1l3 |
| atp2a32 | 1.2E-39 | 1.5610508 | 0.123 | 0.04 | 3.79E-35 | Synoviocytes | atp2a3 |
| dynll13 | 1.6E-39 | 1.0725114 | 0.567 | 0.424 | 5.07E-35 | Synoviocytes | dynll1 |
| lgals3a3 | 3.4E-39 | 1.0417656 | 0.434 | 0.274 | 1.12E-34 | Synoviocytes | lgals3a |
| cd2764 | 3.7E-39 | 1.141307 | 0.265 | 0.13 | 1.2E-34 | Synoviocytes | cd276 |
| lepr4 | 5.8E-39 | 1.4956874 | 0.161 | 0.062 | 1.88E-34 | Synoviocytes | lepr |
| scocb3 | 7.6E-39 | 1.4945647 | 0.191 | 0.08 | 2.46E-34 | Synoviocytes | scocb |
| MYADM2 | 1.3E-38 | 1.3044917 | 0.215 | 0.095 | 4.37E-34 | Synoviocytes | MYADM |
| abi1a2 | 1.6E-38 | 1.4520418 | 0.168 | 0.066 | 5.23E-34 | Synoviocytes | abi1a |
| tmem60 | 3E-38 | 1.6082076 | 0.133 | 0.046 | 9.72E-34 | Synoviocytes | tmem60 |
| si:dkey-6n21.122 | 5.1E-38 | 1.2349458 | 0.259 | 0.127 | 1.65E-33 | Synoviocytes | si:dkey-6n21.12 |
| BX936337.13 | 5.9E-38 | 0.8386339 | 0.541 | 0.386 | 1.93E-33 | Synoviocytes | BX936337.1 |
| CR848833.15 | 7.4E-38 | 1.3735481 | 0.174 | 0.07 | 2.42E-33 | Synoviocytes | CR848833.1 |
| map3k43 | 1.4E-37 | 1.3814521 | 0.192 | 0.082 | 4.51E-33 | Synoviocytes | map3k4 |
| sept101 | 1.4E-37 | 0.8578994 | 0.464 | 0.299 | 4.66E-33 | Synoviocytes | sept10 |
| hopx5 | 2.5E-37 | 1.0462575 | 0.422 | 0.267 | 8.15E-33 | Synoviocytes | hopx |
| neu3.53 | 2.7E-37 | 1.6160426 | 0.168 | 0.067 | 8.73E-33 | Synoviocytes | neu3.5 |
| tubb52 | 4.1E-37 | 1.0923919 | 0.285 | 0.147 | 1.34E-32 | Synoviocytes | tubb5 |
| fkbp1aa3 | 7.6E-37 | 0.6928383 | 0.649 | 0.503 | 2.48E-32 | Synoviocytes | fkbp1aa |
| chmp1b3 | 1.2E-36 | 1.2931533 | 0.24 | 0.116 | 3.74E-32 | Synoviocytes | chmp1b |
| phldb1a2 | 1.6E-36 | 1.5116311 | 0.121 | 0.041 | 5.19E-32 | Synoviocytes | phldb1a |
| clasp22 | 2.1E-36 | 1.4235999 | 0.171 | 0.07 | 6.81E-32 | Synoviocytes | clasp2 |
| jund5 | 2.5E-36 | 0.568559 | 0.725 | 0.567 | 8.11E-32 | Synoviocytes | jund |
| fam114a14 | 2.7E-36 | 1.3214995 | 0.227 | 0.108 | 8.69E-32 | Synoviocytes | fam114a1 |
| col4a13 | 3.3E-36 | 0.68831 | 0.383 | 0.224 | 1.06E-31 | Synoviocytes | col4a1 |
| h3f3d5 | 3.4E-36 | 0.4555569 | 0.839 | 0.726 | 1.09E-31 | Synoviocytes | h3f3d |
| dsc2l2 | 4.4E-36 | 1.3600023 | 0.166 | 0.066 | 1.43E-31 | Synoviocytes | dsc2l |
| dspa2 | 4.8E-36 | 1.1977561 | 0.249 | 0.123 | 1.57E-31 | Synoviocytes | dspa |
| nckap12 | 6E-36 | 0.9461521 | 0.391 | 0.235 | 1.96E-31 | Synoviocytes | nckap1 |
| CU638870.15 | 7.2E-36 | 0.8981237 | 0.484 | 0.326 | 2.34E-31 | Synoviocytes | CU638870.1 |
| fam83hb2 | 8.2E-36 | 1.4783637 | 0.158 | 0.063 | 2.67E-31 | Synoviocytes | fam83hb |

| | | | | | | | |
|---|---|---|---|---|---|---|---|
| rimkla2 | 1.1E-35 | 1.5896758 | 0.163 | 0.066 | 3.43E-31 | Synoviocytes | rimkla |
| skia6 | 1.3E-35 | 0.676292 | 0.601 | 0.446 | 4.07E-31 | Synoviocytes | skia |
| fgl2a5 | 1.4E-35 | 1.1388693 | 0.293 | 0.155 | 4.53E-31 | Synoviocytes | fgl2a |
| zfyve9a3 | 1.9E-35 | 1.1514294 | 0.289 | 0.153 | 6.02E-31 | Synoviocytes | zfyve9a |
| ches14 | 1.9E-35 | 1.0798928 | 0.322 | 0.18 | 6.23E-31 | Synoviocytes | ches1 |
| cib12 | 2.4E-35 | 1.2196599 | 0.241 | 0.118 | 7.88E-31 | Synoviocytes | cib1 |
| mical3a.13 | 2.7E-35 | 1.3397247 | 0.218 | 0.102 | 8.7E-31 | Synoviocytes | mical3a.1 |
| cpne32 | 4.7E-35 | 1.15786 | 0.225 | 0.106 | 1.52E-30 | Synoviocytes | cpne3 |
| pkn1a | 5.5E-35 | 1.3936112 | 0.162 | 0.065 | 1.8E-30 | Synoviocytes | pkn1a |
| si:dkey-238c7.163 | 6.4E-35 | 1.0789463 | 0.3 | 0.163 | 2.08E-30 | Synoviocytes | si:dkey-238c7.16 |
| afap13 | 7.6E-35 | 1.5155653 | 0.168 | 0.069 | 2.46E-30 | Synoviocytes | afap1 |
| ywhae12 | 1.3E-34 | 0.812784 | 0.482 | 0.327 | 4.35E-30 | Synoviocytes | ywhae1 |
| serpine24 | 1.5E-34 | 1.7647865 | 0.456 | 0.314 | 4.98E-30 | Synoviocytes | serpine2 |
| fndc15 | 1.8E-34 | 0.7029779 | 0.312 | 0.165 | 5.79E-30 | Synoviocytes | fndc1 |
| glsb3 | 2.1E-34 | 1.2562859 | 0.235 | 0.114 | 6.74E-30 | Synoviocytes | glsb |
| mafk | 3.9E-34 | 1.69109 | 0.133 | 0.05 | 1.26E-29 | Synoviocytes | mafk |
| myo1cb1 | 4.4E-34 | 0.9327816 | 0.4 | 0.251 | 1.43E-29 | Synoviocytes | myo1cb |
| KCTD154 | 4.6E-34 | 0.8779772 | 0.495 | 0.339 | 1.5E-29 | Synoviocytes | KCTD15 |
| pak12 | 6.2E-34 | 1.5699724 | 0.161 | 0.066 | 2.01E-29 | Synoviocytes | pak1 |
| afap1l21 | 6.2E-34 | 2.0570579 | 0.107 | 0.036 | 2.03E-29 | Synoviocytes | afap1l2 |
| emilin1b4 | 1.1E-33 | 1.1718137 | 0.192 | 0.084 | 3.42E-29 | Synoviocytes | emilin1b |
| sptlc2b3 | 1.1E-33 | 1.2047263 | 0.237 | 0.117 | 3.64E-29 | Synoviocytes | sptlc2b |
| pitpnaa3 | 1.6E-33 | 1.4984225 | 0.138 | 0.053 | 5.34E-29 | Synoviocytes | pitpnaa |
| prkd22 | 1.8E-33 | 1.1489452 | 0.25 | 0.126 | 5.84E-29 | Synoviocytes | prkd2 |
| stx121 | 1.8E-33 | 1.1574525 | 0.288 | 0.157 | 6E-29 | Synoviocytes | stx12 |
| GCA3 | 2.1E-33 | 0.8490939 | 0.445 | 0.293 | 6.9E-29 | Synoviocytes | GCA |
| sema3e3 | 5.3E-33 | 1.2168936 | 0.202 | 0.093 | 1.72E-28 | Synoviocytes | sema3e |
| mt-nd42 | 5.5E-33 | 0.4307608 | 0.925 | 0.855 | 1.8E-28 | Synoviocytes | mt-nd4 |
| ubc4 | 6.6E-33 | 0.4953149 | 0.78 | 0.66 | 2.16E-28 | Synoviocytes | ubc |
| pard3bb2 | 6.9E-33 | 0.9047848 | 0.381 | 0.234 | 2.24E-28 | Synoviocytes | pard3bb |
| phex2 | 6.9E-33 | 1.6823297 | 0.173 | 0.076 | 2.26E-28 | Synoviocytes | phex |
| gask1b2 | 9.3E-33 | 1.6271708 | 0.11 | 0.038 | 3.02E-28 | Synoviocytes | gask1b |
| zfyve211 | 1.1E-32 | 0.9509702 | 0.363 | 0.221 | 3.57E-28 | Synoviocytes | zfyve21 |
| camk2n1a4 | 1.1E-32 | 1.3596555 | 0.185 | 0.081 | 3.63E-28 | Synoviocytes | camk2n1a |
| ddah21 | 1.4E-32 | 1.0792864 | 0.248 | 0.126 | 4.42E-28 | Synoviocytes | ddah2 |
| il4r.22 | 1.6E-32 | 1.5118278 | 0.148 | 0.059 | 5.31E-28 | Synoviocytes | il4r.2 |
| rhbdf1a2 | 1.9E-32 | 1.2558359 | 0.2 | 0.093 | 6.09E-28 | Synoviocytes | rhbdf1a |
| si:dkey-204f11.642 | 2.2E-32 | 1.2075317 | 0.178 | 0.078 | 7.13E-28 | Synoviocytes | si:dkey-204f11.64 |
| itgav2 | 2.5E-32 | 1.2134165 | 0.199 | 0.092 | 8.01E-28 | Synoviocytes | itgav |
| itgb45 | 3.1E-32 | 0.764824 | 0.471 | 0.309 | 1.01E-27 | Synoviocytes | itgb4 |
| bcl2a3 | 3.5E-32 | 1.143806 | 0.282 | 0.153 | 1.14E-27 | Synoviocytes | bcl2a |
| mbd23 | 4.6E-32 | 1.346904 | 0.181 | 0.081 | 1.49E-27 | Synoviocytes | mbd2 |
| cyb5r31 | 5.1E-32 | 1.1030546 | 0.264 | 0.14 | 1.65E-27 | Synoviocytes | cyb5r3 |
| srgn4 | 5.3E-32 | 0.6866465 | 0.61 | 0.449 | 1.73E-27 | Synoviocytes | srgn |
| map1lc3a3 | 5.5E-32 | 0.9964462 | 0.35 | 0.211 | 1.8E-27 | Synoviocytes | map1lc3a |
| calua4 | 6.1E-32 | 0.8064147 | 0.48 | 0.324 | 1.98E-27 | Synoviocytes | calua |
| ywhag22 | 7.1E-32 | 1.0726442 | 0.297 | 0.166 | 2.31E-27 | Synoviocytes | ywhag2 |
| lamc13 | 7.3E-32 | 1.0235944 | 0.249 | 0.126 | 2.38E-27 | Synoviocytes | lamc1 |

| | | | | | | | |
|---|---|---|---|---|---|---|---|
| uchl32 | 7.9E-32 | 1.1519301 | 0.218 | 0.106 | 2.56E-27 | Synoviocytes | uchl3 |
| ctsz3 | 1.1E-31 | 1.0320101 | 0.292 | 0.16 | 3.44E-27 | Synoviocytes | ctsz |
| anxa11b1 | 1.4E-31 | 0.7325032 | 0.526 | 0.385 | 4.59E-27 | Synoviocytes | anxa11b |
| tbrg13 | 1.6E-31 | 1.6826345 | 0.126 | 0.048 | 5.27E-27 | Synoviocytes | tbrg1 |
| ywhaba2 | 1.7E-31 | 0.641757 | 0.635 | 0.504 | 5.68E-27 | Synoviocytes | ywhaba |
| fmnl2a3 | 1.9E-31 | 1.1104522 | 0.221 | 0.108 | 6.03E-27 | Synoviocytes | fmnl2a |
| slka1 | 2.4E-31 | 0.785598 | 0.455 | 0.301 | 7.71E-27 | Synoviocytes | slka |
| actr2a2 | 5.3E-31 | 0.9129108 | 0.343 | 0.203 | 1.72E-26 | Synoviocytes | actr2a |
| loxl16 | 6.7E-31 | 0.9467983 | 0.339 | 0.198 | 2.19E-26 | Synoviocytes | loxl1 |
| gnaia1 | 7.5E-31 | 1.0278137 | 0.27 | 0.145 | 2.44E-26 | Synoviocytes | gnaia |
| diaph23 | 8.6E-31 | 1.0737118 | 0.229 | 0.114 | 2.81E-26 | Synoviocytes | diaph2 |
| nck1a5 | 8.7E-31 | 1.4894051 | 0.173 | 0.078 | 2.82E-26 | Synoviocytes | nck1a |
| zyx7 | 9.9E-31 | 0.7284305 | 0.473 | 0.315 | 3.22E-26 | Synoviocytes | zyx |
| psmb72 | 1.1E-30 | 0.8660027 | 0.394 | 0.252 | 3.46E-26 | Synoviocytes | psmb7 |
| rfx23 | 1.1E-30 | 1.1818251 | 0.162 | 0.069 | 3.49E-26 | Synoviocytes | rfx2 |
| daam1b | 1.3E-30 | 1.4485319 | 0.115 | 0.041 | 4.22E-26 | Synoviocytes | daam1b |
| anos1a1 | 1.4E-30 | 1.8841021 | 0.106 | 0.037 | 4.55E-26 | Synoviocytes | anos1a |
| ralab2 | 1.5E-30 | 1.2538201 | 0.19 | 0.088 | 4.87E-26 | Synoviocytes | ralab |
| wwp22 | 2E-30 | 1.0396397 | 0.213 | 0.103 | 6.61E-26 | Synoviocytes | wwp2 |
| actr12 | 2.1E-30 | 1.0271307 | 0.265 | 0.143 | 6.67E-26 | Synoviocytes | actr1 |
| si:ch73-335l21.45 | 2.4E-30 | 0.733868 | 0.713 | 0.568 | 7.76E-26 | Synoviocytes | si:ch73-335l21.4 |
| itga6b3 | 2.5E-30 | 0.8573259 | 0.441 | 0.294 | 8.27E-26 | Synoviocytes | itga6b |
| si:ch1073-385f13.31 | 2.7E-30 | 1.4439006 | 0.146 | 0.06 | 8.62E-26 | Synoviocytes | si:ch1073-385f13.3 |
| tpm34 | 4.7E-30 | 0.5853418 | 0.648 | 0.524 | 1.53E-25 | Synoviocytes | tpm3 |
| sat1a.28 | 5.3E-30 | 0.6501688 | 0.602 | 0.454 | 1.71E-25 | Synoviocytes | sat1a.2 |
| gdi12 | 7.2E-30 | 1.0257953 | 0.212 | 0.103 | 2.35E-25 | Synoviocytes | gdi1 |
| rab141 | 8E-30 | 0.8532191 | 0.363 | 0.224 | 2.61E-25 | Synoviocytes | rab14 |
| sdf44 | 8.1E-30 | 0.7712874 | 0.452 | 0.301 | 2.62E-25 | Synoviocytes | sdf4 |
| gnb1b2 | 1E-29 | 0.569579 | 0.682 | 0.566 | 3.26E-25 | Synoviocytes | gnb1b |
| skib7 | 1.3E-29 | 0.9283758 | 0.335 | 0.2 | 4.23E-25 | Synoviocytes | skib |
| plekhh1 | 2.5E-29 | 1.6664538 | 0.118 | 0.045 | 8.07E-25 | Synoviocytes | plekhh1 |
| itgb55 | 2.5E-29 | 1.3168334 | 0.179 | 0.083 | 8.25E-25 | Synoviocytes | itgb5 |
| il4r.14 | 3.8E-29 | 1.0874986 | 0.272 | 0.15 | 1.23E-24 | Synoviocytes | il4r.1 |
| aif1l3 | 4.4E-29 | 1.3394705 | 0.181 | 0.084 | 1.42E-24 | Synoviocytes | aif1l |
| psmd73 | 4.7E-29 | 0.870729 | 0.383 | 0.244 | 1.52E-24 | Synoviocytes | psmd7 |
| si:dkey-253i9.42 | 7.8E-29 | 1.3709192 | 0.168 | 0.076 | 2.54E-24 | Synoviocytes | si:dkey-253i9.4 |
| daam1a1 | 8E-29 | 0.9516493 | 0.335 | 0.202 | 2.59E-24 | Synoviocytes | daam1a |
| plp23 | 9E-29 | 1.1000444 | 0.225 | 0.116 | 2.93E-24 | Synoviocytes | plp2 |
| limch1b5 | 1.1E-28 | 1.4947427 | 0.185 | 0.089 | 3.55E-24 | Synoviocytes | limch1b |
| rab5aa3 | 1.4E-28 | 0.8155022 | 0.398 | 0.26 | 4.55E-24 | Synoviocytes | rab5aa |
| tex264a2 | 1.5E-28 | 1.1908063 | 0.229 | 0.119 | 4.91E-24 | Synoviocytes | tex264a |
| si:dkey-16l2.163 | 1.6E-28 | 1.1216379 | 0.225 | 0.116 | 5.06E-24 | Synoviocytes | si:dkey-16l2.16 |
| C25H11orf966 | 2E-28 | 1.0627741 | 0.516 | 0.381 | 6.47E-24 | Synoviocytes | C25H11orf96 |
| socs3a2 | 2.6E-28 | 0.5645103 | 0.883 | 0.793 | 8.36E-24 | Synoviocytes | socs3a |
| slc12a43 | 3.8E-28 | 1.0783449 | 0.234 | 0.123 | 1.25E-23 | Synoviocytes | slc12a4 |
| zgc:1539112 | 4.1E-28 | 1.7748989 | 0.153 | 0.067 | 1.33E-23 | Synoviocytes | zgc:153911 |
| niban2b1 | 5E-28 | 1.0245965 | 0.265 | 0.147 | 1.64E-23 | Synoviocytes | niban2b |
| hspg24 | 6.8E-28 | 1.0073948 | 0.262 | 0.144 | 2.22E-23 | Synoviocytes | hspg2 |

| | | | | | | | |
|---|---|---|---|---|---|---|---|
| si:dkey-222f8.33 | 1.5E-27 | 0.8455304 | 0.352 | 0.219 | 4.72E-23 | Synoviocytes | si:dkey-222f8.3 |
| sptssa2 | 1.5E-27 | 1.0554 | 0.219 | 0.113 | 5.04E-23 | Synoviocytes | sptssa |
| ube2na2 | 1.9E-27 | 0.8008216 | 0.419 | 0.281 | 6.03E-23 | Synoviocytes | ube2na |
| tbc1d2 | 2E-27 | 1.4612959 | 0.115 | 0.044 | 6.51E-23 | Synoviocytes | tbc1d2 |
| ywhabb2 | 2.4E-27 | 0.665306 | 0.535 | 0.394 | 7.69E-23 | Synoviocytes | ywhabb |
| fhl1b4 | 2.7E-27 | 1.2591837 | 0.19 | 0.093 | 8.82E-23 | Synoviocytes | fhl1b |
| myl62 | 3.4E-27 | 0.9179221 | 0.342 | 0.213 | 1.1E-22 | Synoviocytes | myl6 |
| ap1ar2 | 3.5E-27 | 1.1004714 | 0.174 | 0.081 | 1.15E-22 | Synoviocytes | ap1ar |
| tmem50a4 | 3.6E-27 | 0.8672546 | 0.369 | 0.233 | 1.16E-22 | Synoviocytes | tmem50a |
| hdlbpa6 | 4.3E-27 | 0.4455269 | 0.779 | 0.674 | 1.4E-22 | Synoviocytes | hdlbpa |
| si:ch211-207i1.23 | 4.9E-27 | 1.1603433 | 0.173 | 0.081 | 1.58E-22 | Synoviocytes | si:ch211-207i1.2 |
| lmna4 | 5.8E-27 | 0.4226512 | 0.848 | 0.78 | 1.88E-22 | Synoviocytes | lmna |
| mef2aa1 | 7.5E-27 | 0.7481491 | 0.352 | 0.219 | 2.45E-22 | Synoviocytes | mef2aa |
| wbp2nl3 | 8.6E-27 | 0.8382895 | 0.421 | 0.288 | 2.78E-22 | Synoviocytes | wbp2nl |
| plpp1a5 | 1E-26 | 0.3378442 | 0.553 | 0.372 | 3.27E-22 | Synoviocytes | plpp1a |
| mxra5a3 | 1E-26 | 1.3218757 | 0.165 | 0.075 | 3.33E-22 | Synoviocytes | mxra5a |
| rab2a3 | 1E-26 | 0.60548 | 0.631 | 0.52 | 3.41E-22 | Synoviocytes | rab2a |
| nudt91 | 1.3E-26 | 1.3367701 | 0.125 | 0.051 | 4.33E-22 | Synoviocytes | nudt9 |
| nupr1b3 | 1.4E-26 | 1.3847381 | 0.276 | 0.161 | 4.47E-22 | Synoviocytes | nupr1b |
| rab11a2 | 2.4E-26 | 0.9343925 | 0.324 | 0.198 | 7.76E-22 | Synoviocytes | rab11a |
| col5a15 | 2.8E-26 | 0.3665092 | 0.978 | 0.964 | 9.05E-22 | Synoviocytes | col5a1 |
| si:ch211-132g1.32 | 3.2E-26 | 1.1284411 | 0.201 | 0.102 | 1.04E-21 | Synoviocytes | si:ch211-132g1.3 |
| atp6v0d12 | 3.6E-26 | 1.2917987 | 0.162 | 0.074 | 1.16E-21 | Synoviocytes | atp6v0d1 |
| sash1a3 | 3.6E-26 | 0.9431084 | 0.332 | 0.207 | 1.16E-21 | Synoviocytes | sash1a |
| cdc42bpb2 | 4E-26 | 0.985273 | 0.246 | 0.134 | 1.29E-21 | Synoviocytes | cdc42bpb |
| gabarapa2 | 5E-26 | 0.7786181 | 0.461 | 0.329 | 1.61E-21 | Synoviocytes | gabarapa |
| lmcd15 | 5.3E-26 | 1.2121646 | 0.179 | 0.087 | 1.72E-21 | Synoviocytes | lmcd1 |
| zgc:1141741 | 6.5E-26 | 1.2309653 | 0.193 | 0.097 | 2.11E-21 | Synoviocytes | zgc:114174 |
| tsku4 | 1.9E-25 | 0.6539328 | 0.459 | 0.316 | 6.06E-21 | Synoviocytes | tsku |
| ostc3 | 2.2E-25 | 0.7421639 | 0.423 | 0.288 | 7.24E-21 | Synoviocytes | ostc |
| capza1a4 | 2.2E-25 | 0.7532588 | 0.403 | 0.27 | 7.27E-21 | Synoviocytes | capza1a |
| nck1b4 | 2.2E-25 | 1.0759703 | 0.191 | 0.096 | 7.28E-21 | Synoviocytes | nck1b |
| fermt22 | 2.6E-25 | 1.0610487 | 0.192 | 0.096 | 8.46E-21 | Synoviocytes | fermt2 |
| cnih13 | 2.7E-25 | 1.1842271 | 0.188 | 0.094 | 8.93E-21 | Synoviocytes | cnih1 |
| baxb1 | 3.6E-25 | 1.4072963 | 0.125 | 0.052 | 1.18E-20 | Synoviocytes | baxb |
| spon2b4 | 4.6E-25 | 0.6581638 | 0.234 | 0.126 | 1.5E-20 | Synoviocytes | spon2b |
| slc38a24 | 5.8E-25 | 1.1859724 | 0.165 | 0.078 | 1.87E-20 | Synoviocytes | slc38a2 |
| psmd11b1 | 9.8E-25 | 1.3884831 | 0.113 | 0.045 | 3.17E-20 | Synoviocytes | psmd11b |
| pacsin22 | 1.3E-24 | 1.0995521 | 0.141 | 0.062 | 4.31E-20 | Synoviocytes | pacsin2 |
| acsbg2 | 1.8E-24 | 1.4210481 | 0.113 | 0.046 | 5.78E-20 | Synoviocytes | acsbg2 |
| chmp2a2 | 2.5E-24 | 0.7620851 | 0.316 | 0.193 | 8.09E-20 | Synoviocytes | chmp2a |
| pef13 | 2.6E-24 | 0.9950897 | 0.264 | 0.155 | 8.34E-20 | Synoviocytes | pef1 |
| eif4e1c2 | 4.2E-24 | 0.9334598 | 0.251 | 0.142 | 1.37E-19 | Synoviocytes | eif4e1c |
| psmb12 | 4.4E-24 | 0.7976726 | 0.386 | 0.261 | 1.44E-19 | Synoviocytes | psmb1 |
| sgce2 | 5.5E-24 | 0.7419348 | 0.408 | 0.28 | 1.8E-19 | Synoviocytes | sgce |
| pdia63 | 6.3E-24 | 0.8051538 | 0.319 | 0.198 | 2.06E-19 | Synoviocytes | pdia6 |
| elf12 | 7E-24 | 1.3653337 | 0.159 | 0.076 | 2.28E-19 | Synoviocytes | elf1 |
| abracl2 | 7.8E-24 | 0.9019974 | 0.207 | 0.108 | 2.54E-19 | Synoviocytes | abracl |

| | | | | | | | |
|---|---|---|---|---|---|---|---|
| col15a1b2 | 8E-24 | 1.2281085 | 0.118 | 0.049 | 2.61E-19 | Synoviocytes | col15a1b |
| arf13 | 8.5E-24 | 0.9363909 | 0.259 | 0.148 | 2.78E-19 | Synoviocytes | arf1 |
| ppt14 | 9.6E-24 | 0.7106749 | 0.468 | 0.342 | 3.12E-19 | Synoviocytes | ppt1 |
| hivep2a2 | 9.7E-24 | 0.8665321 | 0.333 | 0.213 | 3.17E-19 | Synoviocytes | hivep2a |
| cmpk3 | 1.1E-23 | 0.9471107 | 0.273 | 0.163 | 3.56E-19 | Synoviocytes | cmpk |
| ddx59 | 1.3E-23 | 0.3928302 | 0.735 | 0.595 | 4.15E-19 | Synoviocytes | ddx5 |
| hsp90b12 | 1.8E-23 | 0.713576 | 0.477 | 0.343 | 5.76E-19 | Synoviocytes | hsp90b1 |
| selenof4 | 2.5E-23 | 0.5036713 | 0.596 | 0.465 | 8.27E-19 | Synoviocytes | selenof |
| rap1b2 | 5.4E-23 | 0.6730614 | 0.486 | 0.363 | 1.75E-18 | Synoviocytes | rap1b |
| arl8a1 | 5.4E-23 | 1.057321 | 0.162 | 0.078 | 1.77E-18 | Synoviocytes | arl8a |
| tfg3 | 7.2E-23 | 0.8519694 | 0.303 | 0.188 | 2.36E-18 | Synoviocytes | tfg |
| mtpn2 | 1.1E-22 | 0.6236816 | 0.442 | 0.31 | 3.69E-18 | Synoviocytes | mtpn |
| chmp4bb3 | 1.2E-22 | 0.8968511 | 0.257 | 0.151 | 3.97E-18 | Synoviocytes | chmp4bb |
| prss354 | 1.5E-22 | 0.7682847 | 0.18 | 0.092 | 4.93E-18 | Synoviocytes | prss35 |
| kifc33 | 2.5E-22 | 1.288609 | 0.141 | 0.066 | 7.98E-18 | Synoviocytes | kifc3 |
| arhgdig1 | 2.5E-22 | 1.1331646 | 0.161 | 0.078 | 8.01E-18 | Synoviocytes | arhgdig |
| ppp2r1bb2 | 3.3E-22 | 1.0381231 | 0.18 | 0.093 | 1.06E-17 | Synoviocytes | ppp2r1bb |
| sgms12 | 3.8E-22 | 1.3106947 | 0.126 | 0.056 | 1.25E-17 | Synoviocytes | sgms1 |
| smdt1b3 | 4.4E-22 | 0.7730151 | 0.366 | 0.25 | 1.44E-17 | Synoviocytes | smdt1b |
| hyou12 | 4.7E-22 | 0.8560139 | 0.256 | 0.151 | 1.52E-17 | Synoviocytes | hyou1 |
| hsbp1b3 | 6.2E-22 | 0.7252862 | 0.392 | 0.269 | 2.02E-17 | Synoviocytes | hsbp1b |
| nfic3 | 6.4E-22 | 1.1663914 | 0.157 | 0.077 | 2.07E-17 | Synoviocytes | nfic |
| ndfip13 | 7.2E-22 | 0.6346234 | 0.496 | 0.372 | 2.34E-17 | Synoviocytes | ndfip1 |
| dazap22 | 8E-22 | 0.8478111 | 0.248 | 0.144 | 2.6E-17 | Synoviocytes | dazap2 |
| vegfaa5 | 1.1E-21 | 1.2333831 | 0.206 | 0.115 | 3.42E-17 | Synoviocytes | vegfaa |
| tspan3a2 | 1.1E-21 | 0.9925301 | 0.265 | 0.161 | 3.47E-17 | Synoviocytes | tspan3a |
| TXN2 | 1.3E-21 | 0.3435347 | 0.818 | 0.746 | 4.21E-17 | Synoviocytes | TXN |
| ahi13 | 1.4E-21 | 1.0218146 | 0.204 | 0.111 | 4.62E-17 | Synoviocytes | ahi1 |
| arhgef1a3 | 1.6E-21 | 1.254204 | 0.13 | 0.059 | 5.05E-17 | Synoviocytes | arhgef1a |
| mfge8a4 | 1.8E-21 | 1.2442922 | 0.209 | 0.118 | 5.9E-17 | Synoviocytes | mfge8a |
| cox7c3 | 2E-21 | 0.6949235 | 0.422 | 0.297 | 6.38E-17 | Synoviocytes | cox7c |
| utrn3 | 2.1E-21 | 0.8487459 | 0.297 | 0.186 | 6.79E-17 | Synoviocytes | utrn |
| ralbp12 | 2.3E-21 | 0.7770232 | 0.267 | 0.159 | 7.59E-17 | Synoviocytes | ralbp1 |
| zgc:635872 | 2.5E-21 | 1.0277028 | 0.166 | 0.083 | 8.24E-17 | Synoviocytes | zgc:63587 |
| baxa.13 | 2.7E-21 | 0.9283136 | 0.19 | 0.1 | 8.8E-17 | Synoviocytes | baxa.1 |
| rtn33 | 3.2E-21 | 0.6931589 | 0.345 | 0.225 | 1.05E-16 | Synoviocytes | rtn3 |
| uba54 | 3.5E-21 | 1.166302 | 0.193 | 0.105 | 1.15E-16 | Synoviocytes | uba5 |
| maff6 | 3.9E-21 | 0.8903943 | 0.332 | 0.222 | 1.27E-16 | Synoviocytes | maff |
| b3gnt2b2 | 4.2E-21 | 1.1123247 | 0.147 | 0.071 | 1.37E-16 | Synoviocytes | b3gnt2b |
| hk22 | 4.6E-21 | 1.3788203 | 0.117 | 0.051 | 1.49E-16 | Synoviocytes | hk2 |
| mapkapk2a3 | 4.9E-21 | 0.8041706 | 0.334 | 0.22 | 1.6E-16 | Synoviocytes | mapkapk2a |
| fis11 | 5.3E-21 | 0.9618809 | 0.211 | 0.118 | 1.71E-16 | Synoviocytes | fis1 |
| dnajb114 | 6E-21 | 1.0228138 | 0.193 | 0.104 | 1.94E-16 | Synoviocytes | dnajb11 |
| tcp11l22 | 6.3E-21 | 1.1244109 | 0.146 | 0.07 | 2.04E-16 | Synoviocytes | tcp11l2 |
| arpc5b4 | 7.1E-21 | 1.2052174 | 0.168 | 0.086 | 2.31E-16 | Synoviocytes | arpc5b |
| atp5mc3a3 | 7.8E-21 | 0.7189337 | 0.384 | 0.269 | 2.54E-16 | Synoviocytes | atp5mc3a |
| dbi3 | 8.3E-21 | 0.6195126 | 0.549 | 0.44 | 2.72E-16 | Synoviocytes | dbi |
| ralba2 | 9.5E-21 | 1.0302131 | 0.172 | 0.089 | 3.09E-16 | Synoviocytes | ralba |

| | | | | | | | |
|---|---|---|---|---|---|---|---|
| mt-cyb2 | 1E-20 | 0.277762 | 0.987 | 0.975 | 3.33E-16 | Synoviocytes | mt-cyb |
| appa2 | 1E-20 | 0.7115404 | 0.399 | 0.28 | 3.34E-16 | Synoviocytes | appa |
| nog34 | 1.3E-20 | 1.213793 | 0.16 | 0.081 | 4.3E-16 | Synoviocytes | nog3 |
| psmb32 | 1.4E-20 | 0.7771119 | 0.369 | 0.254 | 4.56E-16 | Synoviocytes | psmb3 |
| capns1a2 | 1.9E-20 | 0.6060983 | 0.517 | 0.398 | 6.18E-16 | Synoviocytes | capns1a |
| gmppb3 | 1.9E-20 | 1.0300063 | 0.211 | 0.12 | 6.27E-16 | Synoviocytes | gmppb |
| col6a16 | 2E-20 | 0.3464464 | 0.899 | 0.874 | 6.36E-16 | Synoviocytes | col6a1 |
| stx41 | 2.6E-20 | 1.1353855 | 0.136 | 0.065 | 8.47E-16 | Synoviocytes | stx4 |
| pdcd6ip2 | 2.8E-20 | 0.9065139 | 0.226 | 0.131 | 8.98E-16 | Synoviocytes | pdcd6ip |
| sirt21 | 3.4E-20 | 0.8544454 | 0.249 | 0.149 | 1.12E-15 | Synoviocytes | sirt2 |
| six1a4 | 3.6E-20 | 0.9558757 | 0.222 | 0.129 | 1.19E-15 | Synoviocytes | six1a |
| eif4eb3 | 4.2E-20 | 0.9995189 | 0.19 | 0.103 | 1.38E-15 | Synoviocytes | eif4eb |
| rab122 | 4.4E-20 | 1.0700317 | 0.14 | 0.067 | 1.42E-15 | Synoviocytes | rab12 |
| casp8 | 4.6E-20 | 1.370698 | 0.106 | 0.046 | 1.5E-15 | Synoviocytes | casp8 |
| psmd83 | 5E-20 | 0.9364148 | 0.248 | 0.15 | 1.61E-15 | Synoviocytes | psmd8 |
| lzic1 | 5.1E-20 | 1.2526166 | 0.118 | 0.053 | 1.66E-15 | Synoviocytes | lzic |
| C7H4orf483 | 6E-20 | 0.7088194 | 0.354 | 0.241 | 1.95E-15 | Synoviocytes | C7H4orf48 |
| rab8b2 | 6.1E-20 | 0.8877427 | 0.231 | 0.135 | 1.97E-15 | Synoviocytes | rab8b |
| arf2a2 | 7.1E-20 | 0.6969454 | 0.355 | 0.241 | 2.3E-15 | Synoviocytes | arf2a |
| notch24 | 8.5E-20 | 0.658309 | 0.489 | 0.377 | 2.75E-15 | Synoviocytes | notch2 |
| col4a24 | 1E-19 | 0.6109873 | 0.33 | 0.217 | 3.35E-15 | Synoviocytes | col4a2 |
| itm2ba3 | 1.1E-19 | 0.3592361 | 0.813 | 0.74 | 3.61E-15 | Synoviocytes | itm2ba |
| ktn12 | 1.5E-19 | 0.6526044 | 0.389 | 0.271 | 5.03E-15 | Synoviocytes | ktn1 |
| fhl3a2 | 1.7E-19 | 1.1555523 | 0.138 | 0.067 | 5.66E-15 | Synoviocytes | fhl3a |
| dda12 | 2.1E-19 | 0.8245481 | 0.254 | 0.154 | 6.71E-15 | Synoviocytes | dda1 |
| hnrnpaba2 | 2.2E-19 | 0.4126569 | 0.695 | 0.603 | 7.25E-15 | Synoviocytes | hnrnpaba |
| eif3jb3 | 2.9E-19 | 0.7300415 | 0.333 | 0.223 | 9.27E-15 | Synoviocytes | eif3jb |
| atp6v0cb2 | 3.5E-19 | 1.0880057 | 0.158 | 0.082 | 1.12E-14 | Synoviocytes | atp6v0cb |
| si:dkey-16l2.202 | 3.6E-19 | 1.1061084 | 0.13 | 0.062 | 1.16E-14 | Synoviocytes | si:dkey-16l2.20 |
| nr4a32 | 4.1E-19 | 1.6862028 | 0.11 | 0.05 | 1.33E-14 | Synoviocytes | nr4a3 |
| ift57 | 4.2E-19 | 1.1311876 | 0.124 | 0.058 | 1.35E-14 | Synoviocytes | ift57 |
| gng52 | 4.9E-19 | 0.7256118 | 0.375 | 0.262 | 1.6E-14 | Synoviocytes | gng5 |
| ocrl1 | 5.1E-19 | 1.1159371 | 0.138 | 0.068 | 1.65E-14 | Synoviocytes | ocrl |
| sem12 | 5.8E-19 | 0.6024936 | 0.521 | 0.416 | 1.88E-14 | Synoviocytes | sem1 |
| stxbp32 | 6E-19 | 0.8373376 | 0.257 | 0.158 | 1.94E-14 | Synoviocytes | stxbp3 |
| jcada2 | 7.6E-19 | 1.0634343 | 0.207 | 0.12 | 2.47E-14 | Synoviocytes | jcada |
| snx9b2 | 9.6E-19 | 0.8250931 | 0.279 | 0.179 | 3.13E-14 | Synoviocytes | snx9b |
| glrx4 | 1E-18 | 0.536602 | 0.479 | 0.355 | 3.41E-14 | Synoviocytes | glrx |
| atp6v0ca3 | 1.1E-18 | 0.6809061 | 0.454 | 0.337 | 3.49E-14 | Synoviocytes | atp6v0ca |
| tbcb2 | 1.1E-18 | 0.6751687 | 0.327 | 0.216 | 3.69E-14 | Synoviocytes | tbcb |
| ctsh4 | 1.6E-18 | 1.0323775 | 0.193 | 0.109 | 5.09E-14 | Synoviocytes | ctsh |
| zc2hc1a1 | 1.6E-18 | 1.1249564 | 0.102 | 0.045 | 5.33E-14 | Synoviocytes | zc2hc1a |
| creb3l3l2 | 1.9E-18 | 0.7025342 | 0.357 | 0.247 | 6.08E-14 | Synoviocytes | creb3l3l |
| pcyt23 | 1.9E-18 | 0.9716694 | 0.175 | 0.095 | 6.32E-14 | Synoviocytes | pcyt2 |
| kif5ba2 | 2E-18 | 0.9477363 | 0.176 | 0.096 | 6.49E-14 | Synoviocytes | kif5ba |
| gapdhs3 | 2E-18 | 0.5477316 | 0.709 | 0.628 | 6.62E-14 | Synoviocytes | gapdhs |
| prkcaa2 | 2.4E-18 | 1.0509118 | 0.186 | 0.104 | 7.96E-14 | Synoviocytes | prkcaa |
| timp2b2 | 2.5E-18 | 0.8070101 | 0.365 | 0.25 | 8.22E-14 | Synoviocytes | timp2b |

| | | | | | | | |
|---|---|---|---|---|---|---|---|
| eva1ba3 | 2.8E-18 | 1.2052696 | 0.187 | 0.107 | 9.1E-14 | Synoviocytes | eva1ba |
| tmem2584 | 2.8E-18 | 0.4125462 | 0.675 | 0.587 | 9.12E-14 | Synoviocytes | tmem258 |
| jam3b4 | 4.3E-18 | 0.6887094 | 0.328 | 0.219 | 1.41E-13 | Synoviocytes | jam3b |
| fkbp10a3 | 4.4E-18 | 0.8724128 | 0.223 | 0.133 | 1.42E-13 | Synoviocytes | fkbp10a |
| eif4g2a2 | 4.4E-18 | 0.745153 | 0.284 | 0.183 | 1.44E-13 | Synoviocytes | eif4g2a |
| abhd2a2 | 5E-18 | 1.0386023 | 0.161 | 0.085 | 1.63E-13 | Synoviocytes | abhd2a |
| puraa2 | 5.2E-18 | 0.7077738 | 0.358 | 0.252 | 1.68E-13 | Synoviocytes | puraa |
| gnpnat14 | 5.2E-18 | 1.0064362 | 0.165 | 0.088 | 1.71E-13 | Synoviocytes | gnpnat1 |
| timmdc13 | 5.9E-18 | 0.9910669 | 0.164 | 0.088 | 1.93E-13 | Synoviocytes | timmdc1 |
| nipa23 | 6.6E-18 | 0.938255 | 0.147 | 0.075 | 2.16E-13 | Synoviocytes | nipa2 |
| tmed73 | 7.5E-18 | 0.6210899 | 0.423 | 0.315 | 2.42E-13 | Synoviocytes | tmed7 |
| zbtb7a | 7.5E-18 | 1.2512514 | 0.125 | 0.061 | 2.44E-13 | Synoviocytes | zbtb7a |
| lix1l4 | 7.7E-18 | 0.8057447 | 0.271 | 0.173 | 2.5E-13 | Synoviocytes | lix1l |
| psma32 | 8.9E-18 | 0.6836886 | 0.354 | 0.247 | 2.89E-13 | Synoviocytes | psma3 |
| commd41 | 8.9E-18 | 1.0553428 | 0.138 | 0.069 | 2.91E-13 | Synoviocytes | commd4 |
| rpz24 | 9.6E-18 | 0.802766 | 0.277 | 0.177 | 3.11E-13 | Synoviocytes | rpz2 |
| abcf12 | 9.6E-18 | 0.5655063 | 0.424 | 0.307 | 3.12E-13 | Synoviocytes | abcf1 |
| zbtb8os3 | 9.7E-18 | 1.135538 | 0.143 | 0.073 | 3.15E-13 | Synoviocytes | zbtb8os |
| prss235 | 1E-17 | 0.7821734 | 0.299 | 0.199 | 3.3E-13 | Synoviocytes | prss23 |
| rgs12b5 | 1E-17 | 0.965325 | 0.163 | 0.087 | 3.37E-13 | Synoviocytes | rgs12b |
| rcan1a5 | 1.1E-17 | 0.8653678 | 0.332 | 0.228 | 3.66E-13 | Synoviocytes | rcan1a |
| cdv32 | 1.3E-17 | 0.6715841 | 0.343 | 0.236 | 4.27E-13 | Synoviocytes | cdv3 |
| ppiab3 | 1.6E-17 | 0.2669692 | 0.919 | 0.896 | 5.3E-13 | Synoviocytes | ppiab |
| inppl1a3 | 1.6E-17 | 1.1087964 | 0.161 | 0.087 | 5.36E-13 | Synoviocytes | inppl1a |
| spcs23 | 2.1E-17 | 0.6319509 | 0.394 | 0.284 | 6.81E-13 | Synoviocytes | spcs2 |
| grk44 | 2.4E-17 | 0.8747903 | 0.274 | 0.178 | 7.68E-13 | Synoviocytes | grk4 |
| ist13 | 2.4E-17 | 1.0621885 | 0.145 | 0.075 | 7.83E-13 | Synoviocytes | ist1 |
| pls33 | 2.5E-17 | 0.6188775 | 0.425 | 0.319 | 8.23E-13 | Synoviocytes | pls3 |
| tor12 | 2.8E-17 | 0.9396318 | 0.188 | 0.107 | 9.14E-13 | Synoviocytes | tor1 |
| hspa54 | 3.1E-17 | 0.6531387 | 0.584 | 0.491 | 1.02E-12 | Synoviocytes | hspa5 |
| ube2l3b2 | 3.7E-17 | 0.724822 | 0.31 | 0.208 | 1.19E-12 | Synoviocytes | ube2l3b |
| psme21 | 4.4E-17 | 1.0520406 | 0.145 | 0.076 | 1.44E-12 | Synoviocytes | psme2 |
| hspa82 | 4.6E-17 | 0.2574 | 0.955 | 0.936 | 1.5E-12 | Synoviocytes | hspa8 |
| ywhabl3 | 5E-17 | 0.5335212 | 0.524 | 0.416 | 1.61E-12 | Synoviocytes | ywhabl |
| gnai12 | 5E-17 | 0.7227 | 0.253 | 0.157 | 1.61E-12 | Synoviocytes | gnai1 |
| plcd1a3 | 5.1E-17 | 0.8579204 | 0.172 | 0.095 | 1.67E-12 | Synoviocytes | plcd1a |
| ptpn132 | 5.4E-17 | 0.5969342 | 0.409 | 0.3 | 1.75E-12 | Synoviocytes | ptpn13 |
| cdon4 | 5.6E-17 | 0.8782594 | 0.284 | 0.187 | 1.81E-12 | Synoviocytes | cdon |
| snx173 | 5.8E-17 | 0.75765 | 0.341 | 0.241 | 1.87E-12 | Synoviocytes | snx17 |
| adamts53 | 5.8E-17 | 1.2564115 | 0.127 | 0.064 | 1.88E-12 | Synoviocytes | adamts5 |
| si:dkeyp-115e12.6 | 6E-17 | 1.2360852 | 0.105 | 0.049 | 1.95E-12 | Synoviocytes | si:dkeyp-115e12.6 |
| rab1ba2 | 6.4E-17 | 0.8023943 | 0.227 | 0.138 | 2.07E-12 | Synoviocytes | rab1ba |
| bzw1a2 | 6.5E-17 | 0.5375713 | 0.425 | 0.315 | 2.11E-12 | Synoviocytes | bzw1a |
| fnbp1l1 | 6.8E-17 | 0.9943091 | 0.132 | 0.066 | 2.22E-12 | Synoviocytes | fnbp1l |
| arpc1a3 | 7.4E-17 | 0.5502295 | 0.425 | 0.316 | 2.41E-12 | Synoviocytes | arpc1a |
| tuba1b7 | 8.3E-17 | 0.4736718 | 0.425 | 0.307 | 2.69E-12 | Synoviocytes | tuba1b |
| cltca3 | 9.1E-17 | 0.7865514 | 0.293 | 0.196 | 2.97E-12 | Synoviocytes | cltca |
| trim3b6 | 1E-16 | 0.9544173 | 0.169 | 0.093 | 3.32E-12 | Synoviocytes | trim3b |

| | | | | | | | |
|---|---|---|---|---|---|---|---|
| arpc42 | 1.1E-16 | 0.7699163 | 0.271 | 0.177 | 3.59E-12 | Synoviocytes | arpc4 |
| si:dkeyp-69e1.83 | 1.2E-16 | 0.921842 | 0.186 | 0.107 | 3.74E-12 | Synoviocytes | si:dkeyp-69e1.8 |
| golt1bb4 | 1.5E-16 | 0.9056573 | 0.176 | 0.1 | 4.81E-12 | Synoviocytes | golt1bb |
| pora3 | 1.6E-16 | 0.877199 | 0.19 | 0.109 | 5.1E-12 | Synoviocytes | pora |
| edf13 | 1.8E-16 | 0.4525213 | 0.605 | 0.511 | 5.9E-12 | Synoviocytes | edf1 |
| irf92 | 1.9E-16 | 0.8499021 | 0.212 | 0.127 | 6.04E-12 | Synoviocytes | irf9 |
| wasf22 | 2.2E-16 | 0.6206791 | 0.362 | 0.259 | 7.1E-12 | Synoviocytes | wasf2 |
| abtb2b4 | 2.3E-16 | 0.9780121 | 0.141 | 0.074 | 7.37E-12 | Synoviocytes | abtb2b |
| sec632 | 2.3E-16 | 0.7253553 | 0.314 | 0.216 | 7.55E-12 | Synoviocytes | sec63 |
| arf6b2 | 2.4E-16 | 1.0779123 | 0.127 | 0.064 | 7.66E-12 | Synoviocytes | arf6b |
| emd2 | 2.6E-16 | 0.4966591 | 0.468 | 0.36 | 8.32E-12 | Synoviocytes | emd |
| oxsr1a1 | 2.6E-16 | 0.5812815 | 0.312 | 0.208 | 8.45E-12 | Synoviocytes | oxsr1a |
| arhgap35a4 | 2.6E-16 | 0.785884 | 0.272 | 0.179 | 8.47E-12 | Synoviocytes | arhgap35a |
| fcho23 | 2.8E-16 | 0.7615734 | 0.248 | 0.158 | 9.19E-12 | Synoviocytes | fcho2 |
| ntn1a6 | 3.4E-16 | 0.61594 | 0.204 | 0.12 | 1.1E-11 | Synoviocytes | ntn1a |
| fkbp23 | 3.5E-16 | 0.6525828 | 0.302 | 0.202 | 1.14E-11 | Synoviocytes | fkbp2 |
| selenot1a4 | 3.6E-16 | 0.7333631 | 0.26 | 0.168 | 1.18E-11 | Synoviocytes | selenot1a |
| klf7b5 | 3.6E-16 | 0.5438304 | 0.333 | 0.226 | 1.19E-11 | Synoviocytes | klf7b |
| CTBP13 | 3.9E-16 | 0.9099953 | 0.256 | 0.167 | 1.26E-11 | Synoviocytes | CTBP1 |
| nfkbie2 | 3.9E-16 | 1.177292 | 0.14 | 0.074 | 1.26E-11 | Synoviocytes | nfkbie |
| sc:d2172 | 4.7E-16 | 0.7776858 | 0.22 | 0.134 | 1.54E-11 | Synoviocytes | sc:d217 |
| coro1ca5 | 6.3E-16 | 0.5747215 | 0.339 | 0.236 | 2.05E-11 | Synoviocytes | coro1ca |
| mapre1b2 | 6.4E-16 | 0.8492513 | 0.202 | 0.121 | 2.08E-11 | Synoviocytes | mapre1b |
| arf6a2 | 6.4E-16 | 0.9402894 | 0.186 | 0.108 | 2.09E-11 | Synoviocytes | arf6a |
| tubb4b4 | 7.3E-16 | 0.3776003 | 0.846 | 0.809 | 2.37E-11 | Synoviocytes | tubb4b |
| nectin3b3 | 7.6E-16 | 0.8003512 | 0.242 | 0.154 | 2.48E-11 | Synoviocytes | nectin3b |
| dctn33 | 9E-16 | 0.7891007 | 0.236 | 0.15 | 2.94E-11 | Synoviocytes | dctn3 |
| pdcl3 | 9.3E-16 | 0.7187471 | 0.293 | 0.199 | 3.02E-11 | Synoviocytes | pdcl |
| selenow14 | 9.3E-16 | 0.2593228 | 0.921 | 0.897 | 3.02E-11 | Synoviocytes | selenow1 |
| higd1a6 | 9.8E-16 | 0.5394723 | 0.553 | 0.456 | 3.17E-11 | Synoviocytes | higd1a |
| mt-co32 | 1E-15 | 0.2255173 | 0.997 | 0.997 | 3.36E-11 | Synoviocytes | mt-co3 |
| dut1 | 1.1E-15 | 0.6704876 | 0.109 | 0.052 | 3.62E-11 | Synoviocytes | dut |
| pitpnb3 | 1.2E-15 | 1.0012087 | 0.102 | 0.047 | 3.86E-11 | Synoviocytes | pitpnb |
| vcp2 | 1.2E-15 | 0.5389519 | 0.42 | 0.316 | 3.91E-11 | Synoviocytes | vcp |
| serinc13 | 1.4E-15 | 0.6689613 | 0.359 | 0.259 | 4.68E-11 | Synoviocytes | serinc1 |
| atf62 | 1.5E-15 | 0.8196985 | 0.195 | 0.116 | 4.82E-11 | Synoviocytes | atf6 |
| ddr15 | 1.6E-15 | 1.1046315 | 0.154 | 0.085 | 5.04E-11 | Synoviocytes | ddr1 |
| eif4a1a5 | 1.7E-15 | 0.5341125 | 0.466 | 0.364 | 5.41E-11 | Synoviocytes | eif4a1a |
| galnt11 | 2E-15 | 0.7561703 | 0.251 | 0.162 | 6.54E-11 | Synoviocytes | galnt1 |
| angptl45 | 2.1E-15 | 0.5725631 | 0.561 | 0.455 | 6.72E-11 | Synoviocytes | angptl4 |
| aopep4 | 2.1E-15 | 1.0275894 | 0.139 | 0.074 | 6.85E-11 | Synoviocytes | aopep |
| clstn14 | 2.4E-15 | 0.8073726 | 0.218 | 0.135 | 7.9E-11 | Synoviocytes | clstn1 |
| CABZ01026681.13 | 2.7E-15 | 0.989266 | 0.15 | 0.082 | 8.83E-11 | Synoviocytes | CABZ01026681.1 |
| si:dkey-17o15.23 | 2.9E-15 | 0.8632155 | 0.149 | 0.081 | 9.39E-11 | Synoviocytes | si:dkey-17o15.2 |
| loxa6 | 3.6E-15 | 0.6697053 | 0.265 | 0.172 | 1.18E-10 | Synoviocytes | loxa |
| ada5 | 3.8E-15 | 0.6375855 | 0.318 | 0.22 | 1.24E-10 | Synoviocytes | ada |
| zgc:1629444 | 3.9E-15 | 0.6464056 | 0.388 | 0.292 | 1.28E-10 | Synoviocytes | zgc:162944 |
| fam83ha | 4.1E-15 | 1.0558482 | 0.107 | 0.052 | 1.33E-10 | Synoviocytes | fam83ha |

| | | | | | | | |
|---|---|---|---|---|---|---|---|
| ugt2a74 | 4.1E-15 | 0.9894585 | 0.163 | 0.093 | 1.35E-10 | Synoviocytes | ugt2a7 |
| clip23 | 4.6E-15 | 0.7926483 | 0.176 | 0.102 | 1.5E-10 | Synoviocytes | clip2 |
| acsl4a5 | 4.8E-15 | 1.1389382 | 0.141 | 0.077 | 1.55E-10 | Synoviocytes | acsl4a |
| vamp4 | 5.1E-15 | 1.1213536 | 0.11 | 0.054 | 1.66E-10 | Synoviocytes | vamp4 |
| gabarapb4 | 5.2E-15 | 0.5314946 | 0.516 | 0.423 | 1.7E-10 | Synoviocytes | gabarapb |
| nenf3 | 5.6E-15 | 0.9023202 | 0.168 | 0.096 | 1.83E-10 | Synoviocytes | nenf |
| si:ch73-269m14.22 | 5.7E-15 | 0.7075726 | 0.32 | 0.226 | 1.85E-10 | Synoviocytes | si:ch73-269m14.2 |
| vmp13 | 6.3E-15 | 0.7388254 | 0.425 | 0.317 | 2.04E-10 | Synoviocytes | vmp1 |
| snap23.13 | 7.7E-15 | 0.7489663 | 0.27 | 0.181 | 2.5E-10 | Synoviocytes | snap23.1 |
| cisd1 | 8.1E-15 | 0.9018537 | 0.124 | 0.063 | 2.63E-10 | Synoviocytes | cisd1 |
| pdap1b4 | 8.4E-15 | 0.9192708 | 0.165 | 0.094 | 2.74E-10 | Synoviocytes | pdap1b |
| dennd4a5 | 8.9E-15 | 0.7809921 | 0.154 | 0.085 | 2.89E-10 | Synoviocytes | dennd4a |
| nt5e3 | 1.1E-14 | 0.9378143 | 0.123 | 0.063 | 3.57E-10 | Synoviocytes | nt5e |
| txnrd31 | 1.2E-14 | 0.8405063 | 0.168 | 0.096 | 3.75E-10 | Synoviocytes | txnrd3 |
| ATP11B3 | 1.2E-14 | 0.856293 | 0.182 | 0.107 | 3.87E-10 | Synoviocytes | ATP11B |
| ccnd2b | 1.3E-14 | 1.0129041 | 0.103 | 0.05 | 4.29E-10 | Synoviocytes | ccnd2b |
| ccn2a5 | 1.4E-14 | 0.668475 | 0.298 | 0.205 | 4.4E-10 | Synoviocytes | ccn2a |
| s100u2 | 1.4E-14 | 0.3283471 | 0.62 | 0.517 | 4.67E-10 | Synoviocytes | s100u |
| baz1a3 | 1.5E-14 | 0.8872425 | 0.188 | 0.113 | 4.74E-10 | Synoviocytes | baz1a |
| ppp2cb2 | 1.5E-14 | 0.5920306 | 0.391 | 0.29 | 4.86E-10 | Synoviocytes | ppp2cb |
| khdrbs1a1 | 1.5E-14 | 0.3675379 | 0.711 | 0.652 | 4.97E-10 | Synoviocytes | khdrbs1a |
| chac14 | 1.5E-14 | 0.4913804 | 0.46 | 0.353 | 5.01E-10 | Synoviocytes | chac1 |
| sult1st65 | 1.9E-14 | 0.8969733 | 0.19 | 0.115 | 6.29E-10 | Synoviocytes | sult1st6 |
| mt-nd33 | 1.9E-14 | 0.542196 | 0.461 | 0.359 | 6.32E-10 | Synoviocytes | mt-nd3 |
| chp22 | 2E-14 | 0.8889275 | 0.143 | 0.078 | 6.41E-10 | Synoviocytes | chp2 |
| cers2b2 | 2E-14 | 1.152171 | 0.126 | 0.067 | 6.59E-10 | Synoviocytes | cers2b |
| ppp2r5eb2 | 2.3E-14 | 0.7824679 | 0.237 | 0.155 | 7.49E-10 | Synoviocytes | ppp2r5eb |
| ybx12 | 2.3E-14 | 0.2056568 | 0.979 | 0.97 | 7.58E-10 | Synoviocytes | ybx1 |
| tfpia6 | 2.4E-14 | 0.6861718 | 0.265 | 0.178 | 7.72E-10 | Synoviocytes | tfpia |
| bnip2 | 2.5E-14 | 0.9469431 | 0.116 | 0.059 | 8.03E-10 | Synoviocytes | bnip2 |
| scamp23 | 2.5E-14 | 0.779122 | 0.197 | 0.12 | 8.16E-10 | Synoviocytes | scamp2 |
| sox64 | 2.5E-14 | 0.8026683 | 0.173 | 0.102 | 8.25E-10 | Synoviocytes | sox6 |
| psma12 | 2.8E-14 | 0.5371494 | 0.407 | 0.31 | 9.18E-10 | Synoviocytes | psma1 |
| lamp1b3 | 2.9E-14 | 0.9910211 | 0.154 | 0.088 | 9.44E-10 | Synoviocytes | lamp1b |
| hyal13 | 3E-14 | 1.1044944 | 0.109 | 0.055 | 9.81E-10 | Synoviocytes | hyal1 |
| pafah1b1a | 3E-14 | 0.734132 | 0.261 | 0.175 | 9.88E-10 | Synoviocytes | pafah1b1a |
| degs13 | 3.1E-14 | 0.6631033 | 0.275 | 0.187 | 1.01E-09 | Synoviocytes | degs1 |
| kif1b3 | 3.2E-14 | 0.8905586 | 0.17 | 0.099 | 1.04E-09 | Synoviocytes | kif1b |
| ppp1r22 | 3.6E-14 | 0.9314769 | 0.113 | 0.057 | 1.17E-09 | Synoviocytes | ppp1r2 |
| napsa3 | 3.6E-14 | 0.8051901 | 0.253 | 0.168 | 1.17E-09 | Synoviocytes | napsa |
| psmd4b2 | 4E-14 | 0.723905 | 0.227 | 0.146 | 1.29E-09 | Synoviocytes | psmd4b |
| gnb1a1 | 4.4E-14 | 0.4446181 | 0.572 | 0.488 | 1.42E-09 | Synoviocytes | gnb1a |
| rab413 | 4.4E-14 | 0.9130763 | 0.145 | 0.081 | 1.43E-09 | Synoviocytes | rab41 |
| tspan142 | 4.5E-14 | 0.7734943 | 0.196 | 0.12 | 1.46E-09 | Synoviocytes | tspan14 |
| pole42 | 4.8E-14 | 1.0623813 | 0.114 | 0.058 | 1.56E-09 | Synoviocytes | pole4 |
| pitpnbl2 | 4.9E-14 | 0.6819037 | 0.19 | 0.114 | 1.59E-09 | Synoviocytes | pitpnbl |
| trir2 | 5.1E-14 | 0.9928083 | 0.133 | 0.072 | 1.67E-09 | Synoviocytes | trir |
| manf3 | 6.1E-14 | 0.7163087 | 0.271 | 0.184 | 1.99E-09 | Synoviocytes | manf |

| | | | | | | | |
|---|---|---|---|---|---|---|---|
| ilf3b2 | 6.9E-14 | 0.6657415 | 0.275 | 0.186 | 2.23E-09 | Synoviocytes | ilf3b |
| ndufc23 | 7.2E-14 | 0.6103067 | 0.37 | 0.276 | 2.35E-09 | Synoviocytes | ndufc2 |
| fbxo45 | 7.6E-14 | 1.0001151 | 0.103 | 0.051 | 2.49E-09 | Synoviocytes | fbxo45 |
| tmem30aa2 | 7.9E-14 | 0.6860475 | 0.243 | 0.159 | 2.56E-09 | Synoviocytes | tmem30aa |
| tp53bp2b2 | 8.7E-14 | 1.0999566 | 0.105 | 0.053 | 2.82E-09 | Synoviocytes | tp53bp2b |
| atp5pb3 | 8.8E-14 | 0.3954539 | 0.561 | 0.463 | 2.85E-09 | Synoviocytes | atp5pb |
| atp1b1a2 | 9.6E-14 | 0.5144256 | 0.574 | 0.484 | 3.11E-09 | Synoviocytes | atp1b1a |
| prkci | 9.6E-14 | 0.8534483 | 0.18 | 0.108 | 3.12E-09 | Synoviocytes | prkci |
| dnajc72 | 9.8E-14 | 0.6401198 | 0.286 | 0.196 | 3.19E-09 | Synoviocytes | dnajc7 |
| npc12 | 1.1E-13 | 0.8824093 | 0.134 | 0.073 | 3.64E-09 | Synoviocytes | npc1 |
| rab5c2 | 1.2E-13 | 0.6157512 | 0.319 | 0.229 | 3.92E-09 | Synoviocytes | rab5c |
| pkma3 | 1.5E-13 | 0.7101825 | 0.299 | 0.212 | 4.86E-09 | Synoviocytes | pkma |
| ndfip1l2 | 1.6E-13 | 0.4935759 | 0.437 | 0.336 | 5.13E-09 | Synoviocytes | ndfip1l |
| oaz1b6 | 1.7E-13 | 0.5065272 | 0.402 | 0.306 | 5.49E-09 | Synoviocytes | oaz1b |
| rgs3a4 | 1.7E-13 | 1.1442562 | 0.157 | 0.093 | 5.64E-09 | Synoviocytes | rgs3a |
| kirrel1a4 | 1.8E-13 | 0.6172772 | 0.427 | 0.325 | 5.9E-09 | Synoviocytes | kirrel1a |
| cox6a13 | 1.8E-13 | 0.4046077 | 0.587 | 0.49 | 5.99E-09 | Synoviocytes | cox6a1 |
| laptm4a3 | 2.1E-13 | 0.5366925 | 0.402 | 0.305 | 6.75E-09 | Synoviocytes | laptm4a |
| col6a26 | 2.1E-13 | 0.2829639 | 0.891 | 0.877 | 6.88E-09 | Synoviocytes | col6a2 |
| bmpr2a2 | 2.4E-13 | 0.6077873 | 0.262 | 0.176 | 7.85E-09 | Synoviocytes | bmpr2a |
| tcea12 | 2.6E-13 | 0.76171 | 0.175 | 0.105 | 8.3E-09 | Synoviocytes | tcea1 |
| wnk1a1 | 2.6E-13 | 1.0594842 | 0.114 | 0.06 | 8.33E-09 | Synoviocytes | wnk1a |
| psmc52 | 2.8E-13 | 0.6915024 | 0.255 | 0.172 | 9.1E-09 | Synoviocytes | psmc5 |
| ppp4cb1 | 2.9E-13 | 0.9470749 | 0.148 | 0.085 | 9.42E-09 | Synoviocytes | ppp4cb |
| rpz4 | 3E-13 | 0.9194876 | 0.141 | 0.079 | 9.82E-09 | Synoviocytes | rpz4 |
| eif5a23 | 3.1E-13 | 0.3853773 | 0.557 | 0.466 | 9.95E-09 | Synoviocytes | eif5a2 |
| m6pr2 | 3.1E-13 | 0.7879703 | 0.196 | 0.123 | 1.01E-08 | Synoviocytes | m6pr |
| letm12 | 3.3E-13 | 0.9497548 | 0.109 | 0.056 | 1.08E-08 | Synoviocytes | letm1 |
| ptpn2b1 | 3.3E-13 | 0.7618198 | 0.184 | 0.113 | 1.08E-08 | Synoviocytes | ptpn2b |
| MDFIC3 | 3.4E-13 | 0.6974742 | 0.269 | 0.185 | 1.1E-08 | Synoviocytes | MDFIC |
| selenok3 | 3.5E-13 | 0.6967502 | 0.261 | 0.179 | 1.15E-08 | Synoviocytes | selenok |
| rnasekb3 | 3.6E-13 | 0.8242559 | 0.174 | 0.105 | 1.16E-08 | Synoviocytes | rnasekb |
| taf133 | 3.8E-13 | 0.6128951 | 0.305 | 0.216 | 1.25E-08 | Synoviocytes | taf13 |
| mtor | 5E-13 | 1.0317072 | 0.148 | 0.086 | 1.64E-08 | Synoviocytes | mtor |
| tjp1a2 | 6.1E-13 | 0.6986964 | 0.233 | 0.153 | 1.97E-08 | Synoviocytes | tjp1a |
| vps37c1 | 6.8E-13 | 0.9545717 | 0.121 | 0.065 | 2.22E-08 | Synoviocytes | vps37c |
| hnrnpub1 | 6.8E-13 | 0.4993509 | 0.415 | 0.32 | 2.22E-08 | Synoviocytes | hnrnpub |
| vps4b2 | 7.7E-13 | 0.5868502 | 0.35 | 0.257 | 2.49E-08 | Synoviocytes | vps4b |
| aldh1a32 | 7.7E-13 | 0.7345765 | 0.181 | 0.11 | 2.52E-08 | Synoviocytes | aldh1a3 |
| ext1a1 | 9.1E-13 | 1.0204103 | 0.1 | 0.05 | 2.95E-08 | Synoviocytes | ext1a |
| gpx4b3 | 9.6E-13 | 0.4986107 | 0.387 | 0.294 | 3.11E-08 | Synoviocytes | gpx4b |
| vapal4 | 1.1E-12 | 0.7084796 | 0.241 | 0.163 | 3.62E-08 | Synoviocytes | vapal |
| rab18a2 | 1.1E-12 | 0.654403 | 0.232 | 0.153 | 3.66E-08 | Synoviocytes | rab18a |
| pbx3b4 | 1.1E-12 | 0.7482912 | 0.218 | 0.142 | 3.73E-08 | Synoviocytes | pbx3b |
| il15l5 | 1.2E-12 | 0.5646675 | 0.192 | 0.117 | 3.83E-08 | Synoviocytes | il15l |
| tnfrsf1a2 | 1.2E-12 | 0.5774651 | 0.341 | 0.251 | 4.03E-08 | Synoviocytes | tnfrsf1a |
| abcf2a5 | 1.3E-12 | 0.6515634 | 0.259 | 0.177 | 4.29E-08 | Synoviocytes | abcf2a |
| nutf22 | 1.3E-12 | 0.7467923 | 0.206 | 0.132 | 4.36E-08 | Synoviocytes | nutf2 |

| | | | | | | | |
|---|---|---|---|---|---|---|---|
| snw12 | 1.4E-12 | 0.5808441 | 0.278 | 0.193 | 4.46E-08 | Synoviocytes | snw1 |
| bud313 | 1.4E-12 | 0.5880229 | 0.292 | 0.206 | 4.5E-08 | Synoviocytes | bud31 |
| psmd15 | 1.4E-12 | 0.7202971 | 0.261 | 0.18 | 4.51E-08 | Synoviocytes | psmd1 |
| ogdha1 | 1.5E-12 | 0.5632775 | 0.3 | 0.212 | 4.74E-08 | Synoviocytes | ogdha |
| zgc:1009181 | 1.5E-12 | 0.571515 | 0.359 | 0.266 | 4.87E-08 | Synoviocytes | zgc:100918 |
| chmp3 | 1.5E-12 | 0.9720792 | 0.101 | 0.051 | 5.02E-08 | Synoviocytes | chmp3 |
| jupa1 | 1.7E-12 | 0.8044229 | 0.2 | 0.128 | 5.64E-08 | Synoviocytes | jupa |
| snx73 | 1.8E-12 | 0.962557 | 0.133 | 0.075 | 5.74E-08 | Synoviocytes | snx7 |
| cmtm63 | 1.8E-12 | 0.6836536 | 0.215 | 0.139 | 5.91E-08 | Synoviocytes | cmtm6 |
| bahcc1a3 | 1.8E-12 | 0.8223222 | 0.148 | 0.086 | 5.93E-08 | Synoviocytes | bahcc1a |
| b3gnt5a4 | 1.9E-12 | 0.9116986 | 0.178 | 0.111 | 6.1E-08 | Synoviocytes | b3gnt5a |
| rorab5 | 2.1E-12 | 0.8364138 | 0.194 | 0.125 | 6.72E-08 | Synoviocytes | rorab |
| si:dkey-118j18.22 | 2.1E-12 | 0.8391279 | 0.154 | 0.091 | 6.91E-08 | Synoviocytes | si:dkey-118j18.2 |
| serinc55 | 2.4E-12 | 0.6574872 | 0.325 | 0.239 | 7.81E-08 | Synoviocytes | serinc5 |
| snrpg3 | 2.4E-12 | 0.6251932 | 0.265 | 0.182 | 7.96E-08 | Synoviocytes | snrpg |
| pleca3 | 2.5E-12 | 0.2160238 | 0.943 | 0.917 | 8.21E-08 | Synoviocytes | pleca |
| mvp5 | 2.7E-12 | 0.3603802 | 0.42 | 0.315 | 8.9E-08 | Synoviocytes | mvp |
| psmc42 | 2.9E-12 | 0.6202938 | 0.243 | 0.164 | 9.28E-08 | Synoviocytes | psmc4 |
| acbd33 | 3E-12 | 0.739466 | 0.209 | 0.135 | 9.68E-08 | Synoviocytes | acbd3 |
| sar1b3 | 3E-12 | 0.5207609 | 0.388 | 0.298 | 9.77E-08 | Synoviocytes | sar1b |
| arhgdia3 | 3.1E-12 | 0.4023445 | 0.5 | 0.41 | 9.96E-08 | Synoviocytes | arhgdia |
| nucb2a4 | 3.1E-12 | 0.3843373 | 0.599 | 0.523 | 1.01E-07 | Synoviocytes | nucb2a |
| ap2a12 | 3.2E-12 | 0.6624618 | 0.265 | 0.183 | 1.04E-07 | Synoviocytes | ap2a1 |
| atf34 | 3.3E-12 | 1.2180677 | 0.416 | 0.345 | 1.06E-07 | Synoviocytes | atf3 |
| adrm13 | 3.7E-12 | 0.9059699 | 0.15 | 0.089 | 1.2E-07 | Synoviocytes | adrm1 |
| slc36a4 | 3.8E-12 | 1.0256392 | 0.103 | 0.054 | 1.25E-07 | Synoviocytes | slc36a4 |
| reep3b2 | 3.9E-12 | 0.7031154 | 0.247 | 0.17 | 1.25E-07 | Synoviocytes | reep3b |
| hmga1a1 | 4E-12 | 0.425582 | 0.272 | 0.187 | 1.31E-07 | Synoviocytes | hmga1a |
| brk12 | 4.1E-12 | 0.5880541 | 0.285 | 0.203 | 1.34E-07 | Synoviocytes | brk1 |
| fbn2b2 | 4.2E-12 | 0.8436605 | 0.125 | 0.069 | 1.36E-07 | Synoviocytes | fbn2b |
| zdhhc5a | 4.3E-12 | 0.6938873 | 0.192 | 0.12 | 1.39E-07 | Synoviocytes | zdhhc5a |
| psmb42 | 4.3E-12 | 0.5772708 | 0.323 | 0.237 | 1.4E-07 | Synoviocytes | psmb4 |
| prrg13 | 4.4E-12 | 0.536218 | 0.291 | 0.206 | 1.42E-07 | Synoviocytes | prrg1 |
| selenot23 | 4.5E-12 | 0.7462447 | 0.168 | 0.103 | 1.46E-07 | Synoviocytes | selenot2 |
| syncrip2 | 4.5E-12 | 0.3926624 | 0.571 | 0.494 | 1.47E-07 | Synoviocytes | syncrip |
| chmp6b3 | 4.5E-12 | 0.7553914 | 0.223 | 0.151 | 1.48E-07 | Synoviocytes | chmp6b |
| ptbp1a4 | 4.6E-12 | 0.583027 | 0.31 | 0.228 | 1.49E-07 | Synoviocytes | ptbp1a |
| atp6v1e1b4 | 4.9E-12 | 1.0160716 | 0.162 | 0.098 | 1.61E-07 | Synoviocytes | atp6v1e1b |
| aldocb3 | 5.2E-12 | 0.9838421 | 0.131 | 0.075 | 1.68E-07 | Synoviocytes | aldocb |
| hif1ab3 | 5.4E-12 | 0.4653798 | 0.466 | 0.374 | 1.75E-07 | Synoviocytes | hif1ab |
| ptpn21 | 5.5E-12 | 0.5720843 | 0.304 | 0.219 | 1.79E-07 | Synoviocytes | ptpn21 |
| her94 | 5.7E-12 | 0.5730621 | 0.315 | 0.227 | 1.84E-07 | Synoviocytes | her9 |
| cbx3a1 | 5.7E-12 | 0.4157775 | 0.413 | 0.313 | 1.85E-07 | Synoviocytes | cbx3a |
| clta3 | 5.8E-12 | 0.6696753 | 0.22 | 0.145 | 1.88E-07 | Synoviocytes | clta |
| psmd61 | 5.9E-12 | 0.787675 | 0.187 | 0.119 | 1.92E-07 | Synoviocytes | psmd6 |
| si:ch211-71m22.1 | 5.9E-12 | 1.132758 | 0.105 | 0.056 | 1.92E-07 | Synoviocytes | si:ch211-71m22.1 |
| kdm5c3 | 6E-12 | 0.66382 | 0.227 | 0.151 | 1.96E-07 | Synoviocytes | kdm5c |
| rdh10b2 | 6.1E-12 | 1.2489014 | 0.129 | 0.074 | 1.97E-07 | Synoviocytes | rdh10b |

| | | | | | | | |
|---|---|---|---|---|---|---|---|
| zgc:1538673 | 6.1E-12 | 0.2885113 | 0.784 | 0.753 | 1.99E-07 | Synoviocytes | zgc:153867 |
| si:ch211-57n23.15 | 6.3E-12 | 0.6156821 | 0.302 | 0.216 | 2.03E-07 | Synoviocytes | si:ch211-57n23.1 |
| si:ch73-44m9.3.12 | 6.4E-12 | 0.8798684 | 0.16 | 0.098 | 2.07E-07 | Synoviocytes | si:ch73-44m9.3.1 |
| mtap2 | 6.4E-12 | 0.5903256 | 0.284 | 0.201 | 2.08E-07 | Synoviocytes | mtap |
| gpm6bb4 | 7.7E-12 | 0.480166 | 0.425 | 0.334 | 2.5E-07 | Synoviocytes | gpm6bb |
| ubtd23 | 8.4E-12 | 0.888643 | 0.166 | 0.103 | 2.73E-07 | Synoviocytes | ubtd2 |
| xirp2a4 | 8.4E-12 | 0.5918361 | 0.346 | 0.256 | 2.74E-07 | Synoviocytes | xirp2a |
| ctnnb14 | 8.5E-12 | 0.3814092 | 0.584 | 0.504 | 2.75E-07 | Synoviocytes | ctnnb1 |
| ubqln42 | 8.7E-12 | 0.8037809 | 0.209 | 0.139 | 2.82E-07 | Synoviocytes | ubqln4 |
| nfkb23 | 8.8E-12 | 0.8659164 | 0.163 | 0.1 | 2.86E-07 | Synoviocytes | nfkb2 |
| sulf2b5 | 9E-12 | 0.6035585 | 0.244 | 0.164 | 2.93E-07 | Synoviocytes | sulf2b |
| MYO1D3 | 9.5E-12 | 0.8661895 | 0.165 | 0.102 | 3.08E-07 | Synoviocytes | MYO1D |
| cmtm72 | 9.5E-12 | 0.6906467 | 0.189 | 0.12 | 3.09E-07 | Synoviocytes | cmtm7 |
| u2af2b2 | 1E-11 | 0.510713 | 0.396 | 0.305 | 3.26E-07 | Synoviocytes | u2af2b |
| rer13 | 1.1E-11 | 0.5780791 | 0.286 | 0.205 | 3.56E-07 | Synoviocytes | rer1 |
| ostm11 | 1.1E-11 | 0.940987 | 0.123 | 0.069 | 3.58E-07 | Synoviocytes | ostm1 |
| jpt21 | 1.1E-11 | 0.6642146 | 0.275 | 0.197 | 3.73E-07 | Synoviocytes | jpt2 |
| fam114a24 | 1.2E-11 | 0.7594868 | 0.18 | 0.114 | 4.01E-07 | Synoviocytes | fam114a2 |
| vps282 | 1.4E-11 | 0.7716323 | 0.179 | 0.113 | 4.48E-07 | Synoviocytes | vps28 |
| kctd32 | 1.4E-11 | 0.8211779 | 0.16 | 0.097 | 4.63E-07 | Synoviocytes | kctd3 |
| csde11 | 1.4E-11 | 0.4741502 | 0.475 | 0.391 | 4.69E-07 | Synoviocytes | csde1 |
| waslb2 | 1.4E-11 | 0.6806778 | 0.249 | 0.175 | 4.71E-07 | Synoviocytes | waslb |
| antxr2a4 | 1.6E-11 | 0.5004635 | 0.422 | 0.334 | 5.28E-07 | Synoviocytes | antxr2a |
| prps1a2 | 1.6E-11 | 0.7504126 | 0.181 | 0.115 | 5.32E-07 | Synoviocytes | prps1a |
| lrpap13 | 1.6E-11 | 0.6085366 | 0.257 | 0.179 | 5.34E-07 | Synoviocytes | lrpap1 |
| trps16 | 1.8E-11 | 0.5315072 | 0.254 | 0.173 | 5.71E-07 | Synoviocytes | trps1 |
| si:zfos-2326c3.22 | 1.8E-11 | 0.7254439 | 0.235 | 0.161 | 5.79E-07 | Synoviocytes | si:zfos-2326c3.2 |
| apcdd1l5 | 1.8E-11 | 0.7850182 | 0.175 | 0.109 | 5.88E-07 | Synoviocytes | apcdd1l |
| copb23 | 1.9E-11 | 0.5616923 | 0.351 | 0.269 | 6.05E-07 | Synoviocytes | copb2 |
| si:ch211-160o17.23 | 1.9E-11 | 0.9847164 | 0.104 | 0.056 | 6.13E-07 | Synoviocytes | si:ch211-160o17.2 |
| nfe2l1b2 | 1.9E-11 | 0.7866718 | 0.146 | 0.086 | 6.19E-07 | Synoviocytes | nfe2l1b |
| yipf42 | 1.9E-11 | 0.6637229 | 0.256 | 0.179 | 6.25E-07 | Synoviocytes | yipf4 |
| kif13ba3 | 1.9E-11 | 0.7681857 | 0.165 | 0.102 | 6.26E-07 | Synoviocytes | kif13ba |
| psmc1b1 | 2E-11 | 0.8656074 | 0.145 | 0.086 | 6.62E-07 | Synoviocytes | psmc1b |
| nsfl1c3 | 2.1E-11 | 0.6014112 | 0.194 | 0.125 | 6.71E-07 | Synoviocytes | nsfl1c |
| dbn12 | 2.3E-11 | 0.6109973 | 0.279 | 0.199 | 7.52E-07 | Synoviocytes | dbn1 |
| clic43 | 2.3E-11 | 0.5747919 | 0.277 | 0.197 | 7.53E-07 | Synoviocytes | clic4 |
| hnrnpua2 | 2.4E-11 | 0.5028556 | 0.274 | 0.191 | 7.66E-07 | Synoviocytes | hnrnpua |
| slc25a55a5 | 2.5E-11 | 0.9232236 | 0.269 | 0.196 | 8.07E-07 | Synoviocytes | slc25a55a |
| lsp1a3 | 2.6E-11 | 0.5785624 | 0.291 | 0.208 | 8.47E-07 | Synoviocytes | lsp1a |
| gmfb2 | 2.6E-11 | 0.614377 | 0.216 | 0.143 | 8.52E-07 | Synoviocytes | gmfb |
| atp5fa12 | 2.7E-11 | 0.3760514 | 0.547 | 0.46 | 8.9E-07 | Synoviocytes | atp5fa1 |
| csnk1g2b2 | 2.8E-11 | 0.4639279 | 0.249 | 0.169 | 9.07E-07 | Synoviocytes | csnk1g2b |
| hmcn14 | 2.8E-11 | 0.864869 | 0.125 | 0.072 | 9.07E-07 | Synoviocytes | hmcn1 |
| tgfb33 | 2.9E-11 | 1.0303139 | 0.135 | 0.08 | 9.56E-07 | Synoviocytes | tgfb3 |
| coa3a2 | 3E-11 | 0.6765595 | 0.196 | 0.128 | 9.6E-07 | Synoviocytes | coa3a |
| txndc123 | 3.1E-11 | 0.8775872 | 0.117 | 0.065 | 1E-06 | Synoviocytes | txndc12 |
| si:ch73-375g18.1 | 3.1E-11 | 0.9326518 | 0.113 | 0.063 | 1.01E-06 | Synoviocytes | si:ch73-375g18.1 |

| | | | | | | | |
|---|---|---|---|---|---|---|---|
| qkia3 | 3.3E-11 | 0.4220723 | 0.548 | 0.467 | 1.06E-06 | Synoviocytes | qkia |
| idh3a2 | 3.4E-11 | 0.7319324 | 0.135 | 0.079 | 1.1E-06 | Synoviocytes | idh3a |
| ddost3 | 3.4E-11 | 0.5213763 | 0.307 | 0.223 | 1.1E-06 | Synoviocytes | ddost |
| cdc422 | 3.4E-11 | 0.4553994 | 0.459 | 0.375 | 1.11E-06 | Synoviocytes | cdc42 |
| EMP32 | 3.4E-11 | 0.7743816 | 0.116 | 0.064 | 1.11E-06 | Synoviocytes | EMP3 |
| pdcd10a3 | 3.6E-11 | 0.7680293 | 0.155 | 0.095 | 1.17E-06 | Synoviocytes | pdcd10a |
| march5l1 | 3.6E-11 | 0.8901748 | 0.114 | 0.064 | 1.18E-06 | Synoviocytes | march5l |
| strn31 | 3.7E-11 | 0.5800367 | 0.257 | 0.18 | 1.2E-06 | Synoviocytes | strn3 |
| ube2d1b2 | 4.1E-11 | 0.655629 | 0.208 | 0.138 | 1.32E-06 | Synoviocytes | ube2d1b |
| pik3c2a2 | 4.1E-11 | 0.8646852 | 0.148 | 0.09 | 1.32E-06 | Synoviocytes | pik3c2a |
| gpc6a4 | 4.2E-11 | 0.6289597 | 0.278 | 0.201 | 1.35E-06 | Synoviocytes | gpc6a |
| ENSDARG00000012[illegible] | 4.3E-11 | 0.5742809 | 0.364 | 0.284 | 1.4E-06 | Synoviocytes | ENSDARG00000012 |
| psmd121 | 4.8E-11 | 0.6327153 | 0.18 | 0.115 | 1.57E-06 | Synoviocytes | psmd12 |
| eea12 | 5E-11 | 0.7736316 | 0.128 | 0.074 | 1.61E-06 | Synoviocytes | eea1 |
| tmx11 | 5.5E-11 | 0.5408481 | 0.186 | 0.118 | 1.8E-06 | Synoviocytes | tmx1 |
| ppa1b4 | 5.9E-11 | 0.6941127 | 0.207 | 0.137 | 1.9E-06 | Synoviocytes | ppa1b |
| zgc:1129621 | 6.1E-11 | 0.6440938 | 0.193 | 0.125 | 2E-06 | Synoviocytes | zgc:112962 |
| ube2h3 | 6.1E-11 | 0.7782148 | 0.172 | 0.11 | 2E-06 | Synoviocytes | ube2h |
| hnrnpabb1 | 6.2E-11 | 0.3248802 | 0.566 | 0.478 | 2.01E-06 | Synoviocytes | hnrnpabb |
| atp5f1b4 | 6.2E-11 | 0.3292115 | 0.634 | 0.557 | 2.02E-06 | Synoviocytes | atp5f1b |
| EMC42 | 6.6E-11 | 0.8706862 | 0.106 | 0.058 | 2.14E-06 | Synoviocytes | EMC4 |
| pfdn22 | 6.6E-11 | 0.846408 | 0.123 | 0.07 | 2.16E-06 | Synoviocytes | pfdn2 |
| ilf23 | 6.8E-11 | 0.5987567 | 0.232 | 0.159 | 2.2E-06 | Synoviocytes | ilf2 |
| zfp36l1b6 | 7.2E-11 | 0.4611658 | 0.617 | 0.545 | 2.35E-06 | Synoviocytes | zfp36l1b |
| cdkn1a6 | 7.5E-11 | 0.7326264 | 0.186 | 0.121 | 2.44E-06 | Synoviocytes | cdkn1a |
| cgref14 | 8.1E-11 | 0.4265037 | 0.249 | 0.172 | 2.64E-06 | Synoviocytes | cgref1 |
| march8 | 8.5E-11 | 1.0257081 | 0.108 | 0.06 | 2.76E-06 | Synoviocytes | march8 |
| tmed93 | 8.5E-11 | 0.5677954 | 0.305 | 0.226 | 2.77E-06 | Synoviocytes | tmed9 |
| fam177a12 | 8.9E-11 | 0.7701742 | 0.119 | 0.067 | 2.9E-06 | Synoviocytes | fam177a1 |
| psme12 | 9.5E-11 | 0.7113214 | 0.175 | 0.112 | 3.07E-06 | Synoviocytes | psme1 |
| npc23 | 9.6E-11 | 0.4281392 | 0.435 | 0.345 | 3.11E-06 | Synoviocytes | npc2 |
| psma52 | 1.1E-10 | 0.6593523 | 0.265 | 0.193 | 3.49E-06 | Synoviocytes | psma5 |
| tardbp2 | 1.1E-10 | 0.5878037 | 0.232 | 0.159 | 3.69E-06 | Synoviocytes | tardbp |
| golga72 | 1.2E-10 | 0.7309996 | 0.244 | 0.174 | 3.92E-06 | Synoviocytes | golga7 |
| tbca2 | 1.3E-10 | 0.47223 | 0.402 | 0.315 | 4.16E-06 | Synoviocytes | tbca |
| pnp5a2 | 1.3E-10 | 0.7580128 | 0.105 | 0.058 | 4.18E-06 | Synoviocytes | pnp5a |
| psmc21 | 1.3E-10 | 0.7092287 | 0.19 | 0.125 | 4.31E-06 | Synoviocytes | psmc2 |
| exoc11 | 1.3E-10 | 0.7489867 | 0.148 | 0.09 | 4.35E-06 | Synoviocytes | exoc1 |
| smarca4a1 | 1.3E-10 | 0.7519384 | 0.153 | 0.095 | 4.35E-06 | Synoviocytes | smarca4a |
| sugt11 | 1.3E-10 | 0.714596 | 0.171 | 0.109 | 4.38E-06 | Synoviocytes | sugt1 |
| pak2a3 | 1.4E-10 | 0.6388465 | 0.222 | 0.152 | 4.55E-06 | Synoviocytes | pak2a |
| yars12 | 1.4E-10 | 0.8823616 | 0.113 | 0.064 | 4.58E-06 | Synoviocytes | yars1 |
| add14 | 1.5E-10 | 0.65601 | 0.204 | 0.137 | 4.83E-06 | Synoviocytes | add1 |
| fam3a3 | 1.5E-10 | 0.6933325 | 0.207 | 0.14 | 4.93E-06 | Synoviocytes | fam3a |
| plekha1a1 | 1.5E-10 | 1.0743554 | 0.104 | 0.058 | 4.99E-06 | Synoviocytes | plekha1a |
| si:ch211-196l7.45 | 1.5E-10 | 0.4024746 | 0.565 | 0.474 | 5E-06 | Synoviocytes | si:ch211-196l7.4 |
| ubl7b3 | 1.6E-10 | 0.8008537 | 0.127 | 0.075 | 5.34E-06 | Synoviocytes | ubl7b |
| stom3 | 1.7E-10 | 0.5108353 | 0.352 | 0.269 | 5.39E-06 | Synoviocytes | stom |

| | | | | | | | |
|---|---|---|---|---|---|---|---|
| rab6a3 | 1.7E-10 | 0.6038464 | 0.257 | 0.183 | 5.49E-06 | Synoviocytes | rab6a |
| ap2s12 | 1.7E-10 | 0.5821168 | 0.244 | 0.172 | 5.55E-06 | Synoviocytes | ap2s1 |
| ndufa122 | 2E-10 | 0.6372587 | 0.293 | 0.219 | 6.4E-06 | Synoviocytes | ndufa12 |
| psmb52 | 2.2E-10 | 0.6747863 | 0.203 | 0.137 | 7.1E-06 | Synoviocytes | psmb5 |
| otud5a1 | 2.2E-10 | 0.7349352 | 0.132 | 0.078 | 7.22E-06 | Synoviocytes | otud5a |
| eif4ebp23 | 2.3E-10 | 0.5372788 | 0.291 | 0.215 | 7.49E-06 | Synoviocytes | eif4ebp2 |
| eml11 | 2.6E-10 | 1.1510802 | 0.1 | 0.055 | 8.41E-06 | Synoviocytes | eml1 |
| hagh2 | 2.6E-10 | 0.9705388 | 0.108 | 0.061 | 8.47E-06 | Synoviocytes | hagh |
| pgam1a3 | 2.7E-10 | 0.5569906 | 0.259 | 0.186 | 8.66E-06 | Synoviocytes | pgam1a |
| cab39l11 | 3.1E-10 | 0.6080533 | 0.223 | 0.155 | 9.99E-06 | Synoviocytes | cab39l1 |
| itsn2b1 | 3.1E-10 | 0.6136188 | 0.208 | 0.141 | 1E-05 | Synoviocytes | itsn2b |
| vamp33 | 3.1E-10 | 0.503334 | 0.328 | 0.249 | 1.01E-05 | Synoviocytes | vamp3 |
| ube2v12 | 3.2E-10 | 0.4399381 | 0.348 | 0.265 | 1.03E-05 | Synoviocytes | ube2v1 |
| si:rp71-36a1.21 | 3.2E-10 | 0.7575276 | 0.142 | 0.087 | 1.04E-05 | Synoviocytes | si:rp71-36a1.2 |
| STMP14 | 3.2E-10 | 0.5216226 | 0.335 | 0.256 | 1.05E-05 | Synoviocytes | STMP1 |
| tma71 | 3.4E-10 | 0.3178409 | 0.463 | 0.374 | 1.1E-05 | Synoviocytes | tma7 |
| kif3a1 | 3.7E-10 | 0.815436 | 0.108 | 0.061 | 1.19E-05 | Synoviocytes | kif3a |
| rusc22 | 3.7E-10 | 0.7157176 | 0.18 | 0.118 | 1.2E-05 | Synoviocytes | rusc2 |
| alas14 | 3.7E-10 | 0.7088468 | 0.166 | 0.106 | 1.21E-05 | Synoviocytes | alas1 |
| sesn15 | 3.8E-10 | 0.8209672 | 0.285 | 0.217 | 1.24E-05 | Synoviocytes | sesn1 |
| kpna42 | 3.9E-10 | 0.5563893 | 0.212 | 0.143 | 1.26E-05 | Synoviocytes | kpna4 |
| htatsf11 | 3.9E-10 | 0.6454073 | 0.177 | 0.115 | 1.27E-05 | Synoviocytes | htatsf1 |
| mrpl171 | 4E-10 | 0.8186639 | 0.117 | 0.068 | 1.3E-05 | Synoviocytes | mrpl17 |
| ostf12 | 4.1E-10 | 0.7500308 | 0.134 | 0.08 | 1.33E-05 | Synoviocytes | ostf1 |
| cycsb3 | 4.3E-10 | 0.3842136 | 0.46 | 0.373 | 1.39E-05 | Synoviocytes | cycsb |
| eif5b1 | 4.3E-10 | 0.4148793 | 0.435 | 0.347 | 1.4E-05 | Synoviocytes | eif5b |
| atp6v1e1a3 | 4.4E-10 | 0.7763722 | 0.161 | 0.103 | 1.42E-05 | Synoviocytes | atp6v1e1a |
| ndel1b3 | 4.5E-10 | 0.7852304 | 0.153 | 0.097 | 1.48E-05 | Synoviocytes | ndel1b |
| plpp2b3 | 4.8E-10 | 0.6659175 | 0.19 | 0.126 | 1.57E-05 | Synoviocytes | plpp2b |
| mfsd13a1 | 5.1E-10 | 0.8829062 | 0.102 | 0.057 | 1.66E-05 | Synoviocytes | mfsd13a |
| herpud21 | 5.3E-10 | 0.6811016 | 0.125 | 0.073 | 1.73E-05 | Synoviocytes | herpud2 |
| kdm6ba3 | 5.5E-10 | 0.5169715 | 0.297 | 0.22 | 1.8E-05 | Synoviocytes | kdm6ba |
| rtca3 | 5.8E-10 | 0.5879883 | 0.259 | 0.187 | 1.89E-05 | Synoviocytes | rtca |
| lims11 | 5.9E-10 | 0.4474382 | 0.375 | 0.294 | 1.92E-05 | Synoviocytes | lims1 |
| abce13 | 6E-10 | 0.6118763 | 0.185 | 0.122 | 1.95E-05 | Synoviocytes | abce1 |
| mbnl23 | 6.2E-10 | 0.5799649 | 0.298 | 0.223 | 2E-05 | Synoviocytes | mbnl2 |
| tsg101b3 | 6.3E-10 | 0.7048899 | 0.145 | 0.09 | 2.04E-05 | Synoviocytes | tsg101b |
| tmem9b2 | 6.4E-10 | 0.6827942 | 0.16 | 0.102 | 2.1E-05 | Synoviocytes | tmem9b |
| psmc1a2 | 6.9E-10 | 0.6540365 | 0.174 | 0.113 | 2.24E-05 | Synoviocytes | psmc1a |
| arhgap42a3 | 6.9E-10 | 0.7582868 | 0.122 | 0.072 | 2.25E-05 | Synoviocytes | arhgap42a |
| tcta4 | 7E-10 | 0.7518759 | 0.144 | 0.089 | 2.29E-05 | Synoviocytes | tcta |
| kdelr2b3 | 7.2E-10 | 0.4177695 | 0.403 | 0.319 | 2.35E-05 | Synoviocytes | kdelr2b |
| med19a2 | 7.3E-10 | 0.5978838 | 0.195 | 0.131 | 2.38E-05 | Synoviocytes | med19a |
| klf7a2 | 7.5E-10 | 0.9038719 | 0.102 | 0.057 | 2.43E-05 | Synoviocytes | klf7a |
| b2m5 | 7.5E-10 | 0.4713311 | 0.716 | 0.686 | 2.45E-05 | Synoviocytes | b2m |
| atrnl1b4 | 7.9E-10 | 0.6779205 | 0.169 | 0.109 | 2.58E-05 | Synoviocytes | atrnl1b |
| atp6v1g14 | 8.6E-10 | 1.0116748 | 0.194 | 0.132 | 2.8E-05 | Synoviocytes | atp6v1g1 |
| csnk2a11 | 9E-10 | 0.5143431 | 0.331 | 0.255 | 2.91E-05 | Synoviocytes | csnk2a1 |

| | | | | | | | |
|---|---|---|---|---|---|---|---|
| scaf12 | 9.3E-10 | 0.4146756 | 0.313 | 0.233 | 3.02E-05 | Synoviocytes | scaf1 |
| suds31 | 9.8E-10 | 0.6989381 | 0.159 | 0.102 | 3.19E-05 | Synoviocytes | suds3 |
| per25 | 1E-09 | 0.6806307 | 0.162 | 0.104 | 3.32E-05 | Synoviocytes | per2 |
| ube2a2 | 1.1E-09 | 0.7707936 | 0.135 | 0.083 | 3.5E-05 | Synoviocytes | ube2a |
| ptges3b2 | 1.1E-09 | 0.5542633 | 0.208 | 0.142 | 3.59E-05 | Synoviocytes | ptges3b |
| ube2d23 | 1.1E-09 | 0.3996011 | 0.463 | 0.388 | 3.66E-05 | Synoviocytes | ube2d2 |
| FBXO483 | 1.1E-09 | 0.9360258 | 0.105 | 0.06 | 3.67E-05 | Synoviocytes | FBXO48 |
| fam49a2 | 1.2E-09 | 0.916209 | 0.124 | 0.074 | 3.81E-05 | Synoviocytes | fam49a |
| tnip11 | 1.2E-09 | 0.6170044 | 0.254 | 0.182 | 3.83E-05 | Synoviocytes | tnip1 |
| nap1l4b3 | 1.2E-09 | 0.5599239 | 0.175 | 0.114 | 3.93E-05 | Synoviocytes | nap1l4b |
| tmem176l.13 | 1.2E-09 | 0.2406065 | 0.791 | 0.741 | 3.93E-05 | Synoviocytes | tmem176l.1 |
| kifbp2 | 1.2E-09 | 0.6993761 | 0.143 | 0.089 | 4.03E-05 | Synoviocytes | kifbp |
| col5a2a4 | 1.3E-09 | 0.2514428 | 0.852 | 0.791 | 4.07E-05 | Synoviocytes | col5a2a |
| tmtc31 | 1.3E-09 | 0.7318846 | 0.125 | 0.074 | 4.3E-05 | Synoviocytes | tmtc3 |
| SGMS11 | 1.4E-09 | 0.9244719 | 0.106 | 0.061 | 4.42E-05 | Synoviocytes | SGMS1 |
| prdx62 | 1.4E-09 | 0.4191815 | 0.419 | 0.343 | 4.49E-05 | Synoviocytes | prdx6 |
| psmd32 | 1.4E-09 | 0.6169296 | 0.216 | 0.152 | 4.55E-05 | Synoviocytes | psmd3 |
| cnp2 | 1.4E-09 | 0.4821606 | 0.37 | 0.291 | 4.67E-05 | Synoviocytes | cnp |
| bcl2l13 | 1.4E-09 | 0.615881 | 0.222 | 0.156 | 4.69E-05 | Synoviocytes | bcl2l1 |
| nek73 | 1.4E-09 | 0.7232703 | 0.178 | 0.118 | 4.71E-05 | Synoviocytes | nek7 |
| ap2m1a2 | 1.6E-09 | 0.4656076 | 0.383 | 0.304 | 5.08E-05 | Synoviocytes | ap2m1a |
| pum11 | 1.6E-09 | 0.9116312 | 0.15 | 0.097 | 5.14E-05 | Synoviocytes | pum1 |
| ccdc1862 | 1.6E-09 | 0.7644879 | 0.126 | 0.076 | 5.35E-05 | Synoviocytes | ccdc186 |
| skp12 | 1.7E-09 | 0.3344601 | 0.562 | 0.498 | 5.6E-05 | Synoviocytes | skp1 |
| asap1a3 | 1.8E-09 | 0.6636095 | 0.147 | 0.092 | 5.82E-05 | Synoviocytes | asap1a |
| si:ch211-243a20.35 | 1.8E-09 | 0.3808825 | 0.371 | 0.281 | 5.82E-05 | Synoviocytes | si:ch211-243a20.3 |
| c6.13 | 1.8E-09 | 0.3724877 | 0.279 | 0.205 | 5.89E-05 | Synoviocytes | c6.1 |
| sypl2a1 | 1.8E-09 | 0.6622597 | 0.231 | 0.164 | 5.94E-05 | Synoviocytes | sypl2a |
| tp532 | 1.8E-09 | 0.5950248 | 0.226 | 0.159 | 6E-05 | Synoviocytes | tp53 |
| atp6v0b1 | 1.9E-09 | 0.7459361 | 0.181 | 0.122 | 6.04E-05 | Synoviocytes | atp6v0b |
| rab5if3 | 1.9E-09 | 0.7040224 | 0.179 | 0.12 | 6.2E-05 | Synoviocytes | rab5if |
| psmd4a2 | 2E-09 | 0.7289335 | 0.155 | 0.1 | 6.62E-05 | Synoviocytes | psmd4a |
| fxr22 | 2E-09 | 0.7311379 | 0.145 | 0.091 | 6.63E-05 | Synoviocytes | fxr2 |
| tbc1d17 | 2.5E-09 | 0.8304493 | 0.121 | 0.073 | 8.21E-05 | Synoviocytes | tbc1d17 |
| tenm35 | 2.7E-09 | 0.5741241 | 0.276 | 0.206 | 8.68E-05 | Synoviocytes | tenm3 |
| ube2d2l4 | 2.7E-09 | 0.682016 | 0.212 | 0.148 | 8.79E-05 | Synoviocytes | ube2d2l |
| ilk2 | 2.7E-09 | 0.5714192 | 0.248 | 0.179 | 8.9E-05 | Synoviocytes | ilk |
| git13 | 2.9E-09 | 0.9273095 | 0.111 | 0.066 | 9.35E-05 | Synoviocytes | git1 |
| ddx39ab2 | 3E-09 | 0.5287176 | 0.235 | 0.168 | 9.63E-05 | Synoviocytes | ddx39ab |
| ctsc2 | 3.3E-09 | 0.9457573 | 0.113 | 0.067 | 0.000108 | Synoviocytes | ctsc |
| si:ch73-147o17.12 | 3.4E-09 | 0.8427342 | 0.128 | 0.079 | 0.00011 | Synoviocytes | si:ch73-147o17.1 |
| dctn22 | 3.4E-09 | 0.5986791 | 0.187 | 0.126 | 0.000112 | Synoviocytes | dctn2 |
| dctn51 | 3.5E-09 | 0.6558912 | 0.175 | 0.117 | 0.000113 | Synoviocytes | dctn5 |
| dlst2 | 3.5E-09 | 0.5634963 | 0.258 | 0.19 | 0.000113 | Synoviocytes | dlst |
| rbm72 | 3.5E-09 | 0.6784876 | 0.139 | 0.087 | 0.000115 | Synoviocytes | rbm7 |
| psap4 | 3.6E-09 | 0.4424935 | 0.421 | 0.348 | 0.000116 | Synoviocytes | psap |
| shroom44 | 3.9E-09 | 0.8436005 | 0.192 | 0.133 | 0.000126 | Synoviocytes | shroom4 |
| cdh23 | 4E-09 | 0.3867645 | 0.375 | 0.291 | 0.000129 | Synoviocytes | cdh2 |

| | | | | | | | |
|---|---|---|---|---|---|---|---|
| peli1b4 | 4.1E-09 | 0.8405261 | 0.141 | 0.09 | 0.000134 | Synoviocytes | peli1b |
| rock2b.14 | 4.1E-09 | 0.6090704 | 0.238 | 0.171 | 0.000134 | Synoviocytes | rock2b.1 |
| elob.11 | 4.1E-09 | 0.5072619 | 0.298 | 0.228 | 0.000134 | Synoviocytes | elob.1 |
| rnaseka4 | 4.2E-09 | 0.7521056 | 0.161 | 0.105 | 0.000135 | Synoviocytes | rnaseka |
| ptp4a3a3 | 4.3E-09 | 0.5859149 | 0.179 | 0.119 | 0.000139 | Synoviocytes | ptp4a3a |
| eno1a3 | 4.9E-09 | 0.7369304 | 0.398 | 0.34 | 0.000158 | Synoviocytes | eno1a |
| dap5 | 4.9E-09 | 0.5324088 | 0.261 | 0.195 | 0.000159 | Synoviocytes | dap |
| atp5mc14 | 4.9E-09 | 0.4681302 | 0.384 | 0.31 | 0.000159 | Synoviocytes | atp5mc1 |
| elavl1a1 | 5E-09 | 0.7263584 | 0.174 | 0.117 | 0.000162 | Synoviocytes | elavl1a |
| susd6 | 5.1E-09 | 0.5482346 | 0.284 | 0.214 | 0.000165 | Synoviocytes | susd6 |
| snx52 | 5.3E-09 | 0.7886066 | 0.134 | 0.084 | 0.000172 | Synoviocytes | snx5 |
| arhgef12b2 | 5.4E-09 | 0.4645475 | 0.269 | 0.197 | 0.000174 | Synoviocytes | arhgef12b |
| rnf146 | 5.5E-09 | 0.7031072 | 0.138 | 0.087 | 0.000179 | Synoviocytes | rnf146 |
| sypl2b1 | 5.6E-09 | 0.7158919 | 0.169 | 0.113 | 0.000182 | Synoviocytes | sypl2b |
| C8H5orf512 | 5.6E-09 | 0.6146918 | 0.174 | 0.116 | 0.000183 | Synoviocytes | C8H5orf51 |
| cpe4 | 5.6E-09 | 0.4036218 | 0.277 | 0.202 | 0.000183 | Synoviocytes | cpe |
| clu5 | 5.6E-09 | 0.5695356 | 0.46 | 0.382 | 0.000183 | Synoviocytes | clu |
| chmp2bb3 | 5.7E-09 | 0.7256119 | 0.155 | 0.101 | 0.000184 | Synoviocytes | chmp2bb |
| ccdc85b4 | 5.8E-09 | 0.4520553 | 0.26 | 0.189 | 0.000187 | Synoviocytes | ccdc85b |
| zgc:1531154 | 5.8E-09 | 0.8549219 | 0.131 | 0.082 | 0.000188 | Synoviocytes | zgc:153115 |
| si:ch1073-392o20.24 | 5.8E-09 | 0.4210651 | 0.317 | 0.242 | 0.000189 | Synoviocytes | si:ch1073-392o20.2 |
| pld1a3 | 5.9E-09 | 0.7858166 | 0.144 | 0.092 | 0.00019 | Synoviocytes | pld1a |
| hnrnpd1 | 5.9E-09 | 0.555603 | 0.249 | 0.181 | 0.000193 | Synoviocytes | hnrnpd |
| bsg3 | 6E-09 | 0.3002799 | 0.631 | 0.573 | 0.000195 | Synoviocytes | bsg |
| rab5ab2 | 6.1E-09 | 0.5155071 | 0.328 | 0.254 | 0.000198 | Synoviocytes | rab5ab |
| sntb24 | 6.2E-09 | 0.5457509 | 0.307 | 0.236 | 0.000201 | Synoviocytes | sntb2 |
| immt3 | 6.3E-09 | 0.7212568 | 0.14 | 0.089 | 0.000204 | Synoviocytes | immt |
| flna4 | 6.3E-09 | 0.235931 | 0.61 | 0.543 | 0.000205 | Synoviocytes | flna |
| ubtf2 | 6.8E-09 | 0.4343255 | 0.423 | 0.348 | 0.000222 | Synoviocytes | ubtf |
| ube2v21 | 6.9E-09 | 0.2850886 | 0.601 | 0.529 | 0.000225 | Synoviocytes | ube2v2 |
| dusp8a5 | 7.3E-09 | 0.8554967 | 0.14 | 0.089 | 0.000237 | Synoviocytes | dusp8a |
| ndufv33 | 7.4E-09 | 0.663985 | 0.187 | 0.128 | 0.000239 | Synoviocytes | ndufv3 |
| faf22 | 7.6E-09 | 0.5552836 | 0.201 | 0.14 | 0.000247 | Synoviocytes | faf2 |
| acsl4b3 | 8E-09 | 0.6843544 | 0.19 | 0.132 | 0.00026 | Synoviocytes | acsl4b |
| dad14 | 8.2E-09 | 0.3587443 | 0.471 | 0.398 | 0.000267 | Synoviocytes | dad1 |
| ergic23 | 9.3E-09 | 0.4990662 | 0.224 | 0.16 | 0.000301 | Synoviocytes | ergic2 |
| cse1l1 | 9.7E-09 | 0.7469104 | 0.105 | 0.062 | 0.000314 | Synoviocytes | cse1l |
| rwdd11 | 9.8E-09 | 0.4381331 | 0.336 | 0.262 | 0.000318 | Synoviocytes | rwdd1 |
| hnrpkl1 | 1.1E-08 | 0.5737576 | 0.203 | 0.142 | 0.000355 | Synoviocytes | hnrpkl |
| psmd11a2 | 1.1E-08 | 0.539333 | 0.146 | 0.093 | 0.000358 | Synoviocytes | psmd11a |
| rab132 | 1.1E-08 | 0.4949028 | 0.292 | 0.222 | 0.000359 | Synoviocytes | rab13 |
| tegt2 | 1.1E-08 | 0.5349398 | 0.219 | 0.155 | 0.000372 | Synoviocytes | tegt |
| psma6a2 | 1.2E-08 | 0.6349898 | 0.189 | 0.13 | 0.000396 | Synoviocytes | psma6a |
| rab4b | 1.2E-08 | 0.8151159 | 0.113 | 0.068 | 0.0004 | Synoviocytes | rab4b |
| cpda2 | 1.2E-08 | 0.8292822 | 0.126 | 0.079 | 0.000401 | Synoviocytes | cpda |
| ndufs42 | 1.3E-08 | 0.5901538 | 0.192 | 0.132 | 0.000407 | Synoviocytes | ndufs4 |
| golga42 | 1.3E-08 | 0.5341041 | 0.225 | 0.161 | 0.000439 | Synoviocytes | golga4 |
| psmc62 | 1.4E-08 | 0.6330324 | 0.193 | 0.134 | 0.000449 | Synoviocytes | psmc6 |

| | | | | | | | |
|---|---|---|---|---|---|---|---|
| si:ch211-242b18.12 | 1.4E-08 | 0.6915922 | 0.137 | 0.087 | 0.000454 | Synoviocytes | si:ch211-242b18.1 |
| tmem1652 | 1.4E-08 | 0.6830501 | 0.113 | 0.068 | 0.000461 | Synoviocytes | tmem165 |
| chchd23 | 1.4E-08 | 0.7497744 | 0.145 | 0.093 | 0.000464 | Synoviocytes | chchd2 |
| atp2a2b1 | 1.4E-08 | 0.5518631 | 0.155 | 0.101 | 0.00047 | Synoviocytes | atp2a2b |
| sertad43 | 1.4E-08 | 0.5408662 | 0.261 | 0.194 | 0.000471 | Synoviocytes | sertad4 |
| denr1 | 1.5E-08 | 0.4494416 | 0.292 | 0.222 | 0.000488 | Synoviocytes | denr |
| dolpp11 | 1.6E-08 | 0.5590146 | 0.151 | 0.098 | 0.000516 | Synoviocytes | dolpp1 |
| vwa5a1 | 1.6E-08 | 0.6692862 | 0.112 | 0.067 | 0.000521 | Synoviocytes | vwa5a |
| ptp4a13 | 1.6E-08 | 0.4467753 | 0.306 | 0.232 | 0.000522 | Synoviocytes | ptp4a1 |
| pdap1a4 | 1.6E-08 | 0.5110102 | 0.288 | 0.218 | 0.000526 | Synoviocytes | pdap1a |
| emc62 | 1.6E-08 | 0.6615057 | 0.155 | 0.102 | 0.000528 | Synoviocytes | emc6 |
| actn42 | 1.7E-08 | 0.6793615 | 0.164 | 0.109 | 0.000557 | Synoviocytes | actn4 |
| frmd8 | 1.7E-08 | 0.7301151 | 0.106 | 0.063 | 0.000564 | Synoviocytes | frmd8 |
| cope4 | 1.7E-08 | 0.4755061 | 0.327 | 0.256 | 0.000568 | Synoviocytes | cope |
| polr2j2 | 1.8E-08 | 0.7190954 | 0.147 | 0.096 | 0.000572 | Synoviocytes | polr2j |
| elocb3 | 1.9E-08 | 0.4328069 | 0.368 | 0.296 | 0.000605 | Synoviocytes | elocb |
| rbbp43 | 1.9E-08 | 0.553433 | 0.144 | 0.092 | 0.000614 | Synoviocytes | rbbp4 |
| ptpra | 1.9E-08 | 0.8799071 | 0.101 | 0.059 | 0.00062 | Synoviocytes | ptpra |
| srp543 | 1.9E-08 | 0.5999019 | 0.238 | 0.175 | 0.000622 | Synoviocytes | srp54 |
| ngfb5 | 1.9E-08 | 0.6995627 | 0.129 | 0.081 | 0.00063 | Synoviocytes | ngfb |
| cops7a3 | 2E-08 | 0.6588585 | 0.167 | 0.112 | 0.000641 | Synoviocytes | cops7a |
| taf71 | 2E-08 | 0.5935921 | 0.176 | 0.12 | 0.000658 | Synoviocytes | taf7 |
| brd3a | 2E-08 | 0.5534415 | 0.234 | 0.169 | 0.000664 | Synoviocytes | brd3a |
| bri31 | 2.1E-08 | 0.9807201 | 0.101 | 0.059 | 0.000675 | Synoviocytes | bri3 |
| txnl4a3 | 2.2E-08 | 0.6875857 | 0.127 | 0.08 | 0.0007 | Synoviocytes | txnl4a |
| taok1a1 | 2.2E-08 | 0.5359567 | 0.208 | 0.147 | 0.000709 | Synoviocytes | taok1a |
| vapb | 2.3E-08 | 0.4213354 | 0.313 | 0.24 | 0.00075 | Synoviocytes | vapb |
| arpc22 | 2.4E-08 | 0.3649155 | 0.503 | 0.443 | 0.000791 | Synoviocytes | arpc2 |
| ifrd15 | 2.6E-08 | 0.3927336 | 0.354 | 0.282 | 0.000832 | Synoviocytes | ifrd1 |
| hip11 | 2.6E-08 | 0.7286874 | 0.108 | 0.065 | 0.000859 | Synoviocytes | hip1 |
| ccdc1242 | 2.7E-08 | 0.5798126 | 0.231 | 0.169 | 0.000883 | Synoviocytes | ccdc124 |
| psma42 | 2.8E-08 | 0.4352726 | 0.361 | 0.288 | 0.000916 | Synoviocytes | psma4 |
| fuca24 | 3E-08 | 0.4923217 | 0.207 | 0.146 | 0.000963 | Synoviocytes | fuca2 |
| rbms2b1 | 3.1E-08 | 1.0461878 | 0.103 | 0.062 | 0.001012 | Synoviocytes | rbms2b |
| ENSDARG000000356 | 3.1E-08 | 0.5281803 | 0.282 | 0.216 | 0.001018 | Synoviocytes | ENSDARG00000035 |
| psmd142 | 3.1E-08 | 0.5370556 | 0.193 | 0.134 | 0.001021 | Synoviocytes | psmd14 |
| mvb12a1 | 3.2E-08 | 0.7698401 | 0.114 | 0.07 | 0.001043 | Synoviocytes | mvb12a |
| smndc12 | 3.5E-08 | 0.4323148 | 0.277 | 0.207 | 0.001153 | Synoviocytes | smndc1 |
| arhgap23a2 | 3.6E-08 | 0.7240066 | 0.135 | 0.087 | 0.001158 | Synoviocytes | arhgap23a |
| BX572619.11 | 3.7E-08 | 0.7385265 | 0.102 | 0.061 | 0.001197 | Synoviocytes | BX572619.1 |
| piezo14 | 3.8E-08 | 0.6162278 | 0.178 | 0.123 | 0.001235 | Synoviocytes | piezo1 |
| scfd14 | 3.9E-08 | 0.6980487 | 0.155 | 0.104 | 0.001272 | Synoviocytes | scfd1 |
| si:dkey-188i13.112 | 4E-08 | 0.5325279 | 0.265 | 0.202 | 0.001295 | Synoviocytes | si:dkey-188i13.11 |
| abi3bpb5 | 4.1E-08 | 0.4354988 | 0.639 | 0.585 | 0.001322 | Synoviocytes | abi3bpb |
| mphosph81 | 4.2E-08 | 0.4382194 | 0.349 | 0.275 | 0.001352 | Synoviocytes | mphosph8 |
| ccng14 | 4.2E-08 | 0.3612088 | 0.563 | 0.505 | 0.001363 | Synoviocytes | ccng1 |
| hnrnpa0a1 | 4.2E-08 | 0.4906054 | 0.3 | 0.231 | 0.001371 | Synoviocytes | hnrnpa0a |
| h2ax15 | 4.3E-08 | 0.2475584 | 0.689 | 0.628 | 0.001414 | Synoviocytes | h2ax1 |

| | | | | | | | |
|---|---|---|---|---|---|---|---|
| vps252 | 5E-08 | 0.7027245 | 0.145 | 0.095 | 0.001633 | Synoviocytes | vps25 |
| atp5mf3 | 5.3E-08 | 0.3841809 | 0.443 | 0.371 | 0.001714 | Synoviocytes | atp5mf |
| zgc:1034384 | 5.3E-08 | 1.0632845 | 0.136 | 0.088 | 0.001715 | Synoviocytes | zgc:103438 |
| tbc1d10aa2 | 5.4E-08 | 0.6117099 | 0.169 | 0.115 | 0.001752 | Synoviocytes | tbc1d10aa |
| taf151 | 5.5E-08 | 0.4483535 | 0.28 | 0.214 | 0.001784 | Synoviocytes | taf15 |
| snx44 | 5.6E-08 | 0.7218574 | 0.14 | 0.092 | 0.001807 | Synoviocytes | snx4 |
| b2ml6 | 6E-08 | 0.2167622 | 0.92 | 0.897 | 0.001939 | Synoviocytes | b2ml |
| epha4a2 | 6E-08 | 0.851864 | 0.109 | 0.067 | 0.001957 | Synoviocytes | epha4a |
| mgst3b1 | 6.1E-08 | 0.3793008 | 0.449 | 0.384 | 0.001972 | Synoviocytes | mgst3b |
| dusp56 | 6.1E-08 | 0.8028039 | 0.26 | 0.197 | 0.001982 | Synoviocytes | dusp5 |
| tacc13 | 6.3E-08 | 0.5454704 | 0.277 | 0.216 | 0.002044 | Synoviocytes | tacc1 |
| uchl13 | 6.7E-08 | 0.5530982 | 0.184 | 0.127 | 0.002175 | Synoviocytes | uchl1 |
| txndc173 | 6.7E-08 | 0.5835279 | 0.171 | 0.118 | 0.002187 | Synoviocytes | txndc17 |
| kdelr2a3 | 6.8E-08 | 0.331384 | 0.466 | 0.389 | 0.002196 | Synoviocytes | kdelr2a |
| ywhag17 | 7.4E-08 | 0.674467 | 0.162 | 0.11 | 0.002394 | Synoviocytes | ywhag1 |
| ppp1cb2 | 7.6E-08 | 0.4847972 | 0.258 | 0.195 | 0.002475 | Synoviocytes | ppp1cb |
| sars14 | 7.6E-08 | 0.574075 | 0.203 | 0.146 | 0.002483 | Synoviocytes | sars1 |
| atp5if1b4 | 7.8E-08 | 0.4357207 | 0.4 | 0.332 | 0.002544 | Synoviocytes | atp5if1b |
| ARF53 | 7.9E-08 | 0.5220715 | 0.211 | 0.152 | 0.002555 | Synoviocytes | ARF5 |
| sec623 | 8.2E-08 | 0.4357088 | 0.303 | 0.236 | 0.002659 | Synoviocytes | sec62 |
| usp52 | 8.4E-08 | 0.5384015 | 0.158 | 0.106 | 0.002722 | Synoviocytes | usp5 |
| MYO9B4 | 8.4E-08 | 0.6907746 | 0.175 | 0.123 | 0.002723 | Synoviocytes | MYO9B |
| psmc32 | 8.5E-08 | 0.6262516 | 0.146 | 0.096 | 0.002761 | Synoviocytes | psmc3 |
| CU459056.12 | 8.5E-08 | 0.5944878 | 0.228 | 0.169 | 0.002769 | Synoviocytes | CU459056.1 |
| MED93 | 9.1E-08 | 0.826138 | 0.112 | 0.07 | 0.002943 | Synoviocytes | MED9 |
| ube2l3a2 | 9.4E-08 | 0.6592682 | 0.208 | 0.151 | 0.003045 | Synoviocytes | ube2l3a |
| sel1l2 | 9.9E-08 | 0.8967662 | 0.102 | 0.062 | 0.003212 | Synoviocytes | sel1l |
| farsa4 | 1.1E-07 | 0.7306881 | 0.125 | 0.081 | 0.003471 | Synoviocytes | farsa |
| mrpl332 | 1.1E-07 | 0.4880814 | 0.212 | 0.153 | 0.003622 | Synoviocytes | mrpl33 |
| zfand5a6 | 1.1E-07 | 0.6186959 | 0.246 | 0.188 | 0.003723 | Synoviocytes | zfand5a |
| zmat23 | 1.2E-07 | 0.4170544 | 0.279 | 0.214 | 0.003833 | Synoviocytes | zmat2 |
| pdlim13 | 1.2E-07 | 0.5691363 | 0.291 | 0.229 | 0.003878 | Synoviocytes | pdlim1 |
| ccn15 | 1.3E-07 | 0.298908 | 0.7 | 0.642 | 0.004303 | Synoviocytes | ccn1 |
| calm3b2 | 1.3E-07 | 0.3479171 | 0.479 | 0.416 | 0.004375 | Synoviocytes | calm3b |
| colec124 | 1.4E-07 | 0.4950155 | 0.493 | 0.438 | 0.004424 | Synoviocytes | colec12 |
| ifngr2 | 1.4E-07 | 0.7135167 | 0.11 | 0.068 | 0.004447 | Synoviocytes | ifngr2 |
| cacybp2 | 1.4E-07 | 0.4512654 | 0.211 | 0.152 | 0.004447 | Synoviocytes | cacybp |
| txnl11 | 1.4E-07 | 0.6212897 | 0.126 | 0.081 | 0.004471 | Synoviocytes | txnl1 |
| aip1 | 1.4E-07 | 0.476318 | 0.263 | 0.2 | 0.004715 | Synoviocytes | aip |
| taf121 | 1.5E-07 | 0.454842 | 0.272 | 0.207 | 0.004724 | Synoviocytes | taf12 |
| trip114 | 1.5E-07 | 0.6076868 | 0.138 | 0.091 | 0.004846 | Synoviocytes | trip11 |
| kpnb11 | 1.6E-07 | 0.5456637 | 0.199 | 0.143 | 0.005126 | Synoviocytes | kpnb1 |
| tial1 | 1.6E-07 | 0.6872725 | 0.126 | 0.082 | 0.005232 | Synoviocytes | tial1 |
| pkd1a1 | 1.6E-07 | 0.6639186 | 0.115 | 0.072 | 0.005322 | Synoviocytes | pkd1a |
| nedd82 | 1.7E-07 | 0.5644544 | 0.157 | 0.107 | 0.005583 | Synoviocytes | nedd8 |
| taf3 | 1.7E-07 | 0.5813763 | 0.226 | 0.169 | 0.005588 | Synoviocytes | taf3 |
| uqcrh3 | 1.9E-07 | 0.3729191 | 0.45 | 0.384 | 0.006284 | Synoviocytes | uqcrh |
| sumo3a3 | 2E-07 | 0.433275 | 0.35 | 0.286 | 0.006414 | Synoviocytes | sumo3a |

| | | | | | | | |
|---|---|---|---|---|---|---|---|
| arhgap52 | 2E-07 | 0.3658723 | 0.421 | 0.35 | 0.006415 | Synoviocytes | arhgap5 |
| adi13 | 2.1E-07 | 0.6596025 | 0.136 | 0.09 | 0.006721 | Synoviocytes | adi1 |
| smap13 | 2.1E-07 | 0.6236157 | 0.21 | 0.154 | 0.006722 | Synoviocytes | smap1 |
| anapc162 | 2.2E-07 | 0.7028993 | 0.198 | 0.146 | 0.007162 | Synoviocytes | anapc16 |
| ammecr13 | 2.3E-07 | 0.5056713 | 0.202 | 0.145 | 0.007369 | Synoviocytes | ammecr1 |
| uqcrq2 | 2.3E-07 | 0.4570313 | 0.331 | 0.269 | 0.007546 | Synoviocytes | uqcrq |
| slain22 | 2.3E-07 | 0.6188896 | 0.159 | 0.109 | 0.007566 | Synoviocytes | slain2 |
| ddx42 | 2.4E-07 | 0.5440299 | 0.192 | 0.136 | 0.007683 | Synoviocytes | ddx42 |
| lman14 | 2.4E-07 | 0.564647 | 0.176 | 0.124 | 0.007803 | Synoviocytes | lman1 |
| ddit35 | 2.5E-07 | 0.2052259 | 0.654 | 0.581 | 0.007976 | Synoviocytes | ddit3 |
| pfdn41 | 2.5E-07 | 0.3895145 | 0.324 | 0.257 | 0.00817 | Synoviocytes | pfdn4 |
| mrps362 | 2.5E-07 | 0.5753127 | 0.212 | 0.156 | 0.008268 | Synoviocytes | mrps36 |
| otud5b3 | 2.6E-07 | 0.5927514 | 0.129 | 0.084 | 0.008393 | Synoviocytes | otud5b |
| dpp74 | 2.6E-07 | 0.542227 | 0.274 | 0.215 | 0.008523 | Synoviocytes | dpp7 |
| znf7061 | 2.7E-07 | 0.6192765 | 0.143 | 0.096 | 0.00865 | Synoviocytes | znf706 |
| tsc22d33 | 2.7E-07 | 0.4355974 | 0.46 | 0.402 | 0.008907 | Synoviocytes | tsc22d3 |
| tufm1 | 2.8E-07 | 0.576885 | 0.181 | 0.129 | 0.009031 | Synoviocytes | tufm |
| pde7a2 | 2.9E-07 | 0.6401473 | 0.138 | 0.092 | 0.009366 | Synoviocytes | pde7a |
| b4galt63 | 2.9E-07 | 0.5567306 | 0.154 | 0.105 | 0.009519 | Synoviocytes | b4galt6 |
| arhgap313 | 3.1E-07 | 0.5368849 | 0.163 | 0.112 | 0.010037 | Synoviocytes | arhgap31 |
| rrad4 | 3.1E-07 | 1.1237749 | 0.11 | 0.07 | 0.010177 | Synoviocytes | rrad |
| lgmn3 | 3.2E-07 | 0.4325445 | 0.231 | 0.171 | 0.010441 | Synoviocytes | lgmn |
| cdkn1ba5 | 3.3E-07 | 0.6967859 | 0.166 | 0.117 | 0.010683 | Synoviocytes | cdkn1ba |
| spred11 | 3.3E-07 | 0.579724 | 0.179 | 0.128 | 0.010757 | Synoviocytes | spred1 |
| ppfibp1b4 | 3.3E-07 | 0.5128894 | 0.197 | 0.143 | 0.01076 | Synoviocytes | ppfibp1b |
| CU694197.12 | 3.4E-07 | 0.5039567 | 0.266 | 0.206 | 0.01113 | Synoviocytes | CU694197.1 |
| crfb41 | 3.6E-07 | 0.7147885 | 0.152 | 0.105 | 0.011841 | Synoviocytes | crfb4 |
| suclg13 | 3.7E-07 | 0.3850604 | 0.235 | 0.174 | 0.011954 | Synoviocytes | suclg1 |
| mycbp21 | 4E-07 | 0.4808891 | 0.291 | 0.23 | 0.012856 | Synoviocytes | mycbp2 |
| praf22 | 4E-07 | 0.5539841 | 0.192 | 0.138 | 0.013138 | Synoviocytes | praf2 |
| snx1a1 | 4.2E-07 | 0.7475543 | 0.137 | 0.092 | 0.013717 | Synoviocytes | snx1a |
| erp442 | 4.3E-07 | 0.5141052 | 0.243 | 0.186 | 0.013849 | Synoviocytes | erp44 |
| palm2akap2.14 | 4.3E-07 | 0.6595091 | 0.128 | 0.085 | 0.013997 | Synoviocytes | palm2akap2.1 |
| baz1b2 | 4.3E-07 | 0.4221957 | 0.233 | 0.174 | 0.014116 | Synoviocytes | baz1b |
| cnpy34 | 4.4E-07 | 0.7008798 | 0.132 | 0.088 | 0.014413 | Synoviocytes | cnpy3 |
| zgc:1626984 | 4.6E-07 | 0.6115184 | 0.163 | 0.114 | 0.014949 | Synoviocytes | zgc:162698 |
| snap29 | 4.7E-07 | 0.6429964 | 0.112 | 0.071 | 0.015268 | Synoviocytes | snap29 |
| srsf43 | 4.8E-07 | 0.4185635 | 0.22 | 0.162 | 0.015701 | Synoviocytes | srsf4 |
| pet1001 | 4.9E-07 | 0.6651476 | 0.126 | 0.083 | 0.015856 | Synoviocytes | pet100 |
| ube2nb1 | 5.1E-07 | 0.5615893 | 0.192 | 0.139 | 0.016534 | Synoviocytes | ube2nb |
| ufm14 | 5.3E-07 | 0.4691686 | 0.247 | 0.189 | 0.017289 | Synoviocytes | ufm1 |
| tomm70a1 | 5.7E-07 | 0.4746465 | 0.232 | 0.174 | 0.018464 | Synoviocytes | tomm70a |
| zgc:926062 | 5.8E-07 | 0.6118306 | 0.183 | 0.132 | 0.01888 | Synoviocytes | zgc:92606 |
| lasp13 | 5.9E-07 | 0.2768677 | 0.526 | 0.471 | 0.019187 | Synoviocytes | lasp1 |
| sec31a3 | 5.9E-07 | 0.5930991 | 0.156 | 0.108 | 0.019221 | Synoviocytes | sec31a |
| tmem30ab2 | 6.1E-07 | 0.4748278 | 0.195 | 0.141 | 0.019948 | Synoviocytes | tmem30ab |
| bmp42 | 6.2E-07 | 0.864422 | 0.103 | 0.065 | 0.020066 | Synoviocytes | bmp4 |
| SBNO24 | 6.3E-07 | 0.4949165 | 0.286 | 0.227 | 0.020599 | Synoviocytes | SBNO2 |

| | | | | | | | |
|---|---|---|---|---|---|---|---|
| ebf3a4 | 6.6E-07 | 0.399689 | 0.279 | 0.215 | 0.021363 | Synoviocytes | ebf3a |
| cnep1r13 | 6.6E-07 | 0.6120596 | 0.147 | 0.101 | 0.021396 | Synoviocytes | cnep1r1 |
| chmp4ba2 | 6.9E-07 | 0.4920189 | 0.271 | 0.211 | 0.022481 | Synoviocytes | chmp4ba |
| tomm402 | 6.9E-07 | 0.6336052 | 0.143 | 0.097 | 0.022484 | Synoviocytes | tomm40 |
| tmed23 | 6.9E-07 | 0.310836 | 0.458 | 0.391 | 0.022582 | Synoviocytes | tmed2 |
| pkd22 | 7.1E-07 | 0.5450242 | 0.155 | 0.107 | 0.023096 | Synoviocytes | pkd2 |
| atp5f1d4 | 7.2E-07 | 0.3709597 | 0.397 | 0.333 | 0.023308 | Synoviocytes | atp5f1d |
| cct52 | 7.5E-07 | 0.3418563 | 0.44 | 0.381 | 0.024506 | Synoviocytes | cct5 |
| ptpn11a1 | 8E-07 | 0.5680601 | 0.161 | 0.113 | 0.025944 | Synoviocytes | ptpn11a |
| bccip2 | 8.1E-07 | 0.6542542 | 0.246 | 0.194 | 0.026284 | Synoviocytes | bccip |
| ik1 | 8.1E-07 | 0.5160289 | 0.227 | 0.173 | 0.026453 | Synoviocytes | ik |
| gigyf1b1 | 8.3E-07 | 0.5315264 | 0.193 | 0.14 | 0.026972 | Synoviocytes | gigyf1b |
| srrm21 | 8.6E-07 | 0.2786857 | 0.381 | 0.311 | 0.028003 | Synoviocytes | srrm2 |
| gng24 | 8.7E-07 | 0.6035801 | 0.156 | 0.109 | 0.028317 | Synoviocytes | gng2 |
| ARHGAP62 | 8.7E-07 | 0.7073148 | 0.102 | 0.064 | 0.028344 | Synoviocytes | ARHGAP6 |
| zfp36l23 | 8.9E-07 | 0.403941 | 0.33 | 0.267 | 0.028931 | Synoviocytes | zfp36l2 |
| rgmb3 | 8.9E-07 | 0.6093773 | 0.168 | 0.119 | 0.029065 | Synoviocytes | rgmb |
| zgc:919102 | 9.1E-07 | 0.4330794 | 0.312 | 0.251 | 0.029437 | Synoviocytes | zgc:91910 |
| uba11 | 9.2E-07 | 0.4778241 | 0.268 | 0.211 | 0.030041 | Synoviocytes | uba1 |
| fam199x2 | 9.3E-07 | 0.4902559 | 0.184 | 0.132 | 0.030145 | Synoviocytes | fam199x |
| sf3b63 | 9.6E-07 | 0.453003 | 0.342 | 0.282 | 0.031063 | Synoviocytes | sf3b6 |
| COX5B.14 | 9.7E-07 | 0.2328545 | 0.605 | 0.541 | 0.031675 | Synoviocytes | COX5B.1 |
| kifap3a2 | 1E-06 | 0.5037869 | 0.137 | 0.092 | 0.032422 | Synoviocytes | kifap3a |
| sgcb1 | 1E-06 | 0.599942 | 0.115 | 0.075 | 0.034072 | Synoviocytes | sgcb |
| map7d32 | 1.1E-06 | 0.5121651 | 0.122 | 0.079 | 0.034574 | Synoviocytes | map7d3 |
| ppm1ba1 | 1.1E-06 | 0.807946 | 0.116 | 0.076 | 0.035735 | Synoviocytes | ppm1ba |
| snrpd21 | 1.1E-06 | 0.3772755 | 0.262 | 0.201 | 0.035743 | Synoviocytes | snrpd2 |
| ciao2b2 | 1.1E-06 | 0.5447997 | 0.183 | 0.132 | 0.035751 | Synoviocytes | ciao2b |
| emc104 | 1.1E-06 | 0.5245776 | 0.195 | 0.144 | 0.036836 | Synoviocytes | emc10 |
| hnrnph1l2 | 1.1E-06 | 0.442184 | 0.185 | 0.133 | 0.036933 | Synoviocytes | hnrnph1l |
| spcs13 | 1.1E-06 | 0.3180529 | 0.495 | 0.436 | 0.036942 | Synoviocytes | spcs1 |
| pcmt3 | 1.1E-06 | 0.6379926 | 0.136 | 0.093 | 0.037357 | Synoviocytes | pcmt |
| arih11 | 1.2E-06 | 0.6514237 | 0.121 | 0.08 | 0.040469 | Synoviocytes | arih1 |
| sdhdb2 | 1.3E-06 | 0.6263627 | 0.176 | 0.127 | 0.040981 | Synoviocytes | sdhdb |
| nbr1b2 | 1.3E-06 | 0.7358892 | 0.101 | 0.064 | 0.041786 | Synoviocytes | nbr1b |
| dnajb1a1 | 1.3E-06 | 0.6148446 | 0.101 | 0.064 | 0.04195 | Synoviocytes | dnajb1a |
| cops94 | 1.3E-06 | 0.5651224 | 0.204 | 0.153 | 0.042088 | Synoviocytes | cops9 |
| antxr1c3 | 1.3E-06 | 0.3147598 | 0.579 | 0.537 | 0.042339 | Synoviocytes | antxr1c |
| gusb4 | 1.4E-06 | 0.5691932 | 0.178 | 0.129 | 0.044413 | Synoviocytes | gusb |
| aplp23 | 1.4E-06 | 0.3595128 | 0.386 | 0.322 | 0.045918 | Synoviocytes | aplp2 |
| nos1apa6 | 1.4E-06 | 0.733517 | 0.153 | 0.108 | 0.046275 | Synoviocytes | nos1apa |
| aldoaa4 | 1.4E-06 | 0.3228435 | 0.433 | 0.368 | 0.046519 | Synoviocytes | aldoaa |
| spty2d1 | 1.5E-06 | 0.4330224 | 0.183 | 0.131 | 0.047633 | Synoviocytes | spty2d1 |
| acap12 | 1.5E-06 | 0.6610908 | 0.123 | 0.082 | 0.047711 | Synoviocytes | acap1 |
| ssh2a2 | 1.5E-06 | 0.8990071 | 0.119 | 0.08 | 0.048137 | Synoviocytes | ssh2a |
| rbx12 | 1.5E-06 | 0.2531757 | 0.516 | 0.453 | 0.049252 | Synoviocytes | rbx1 |
| hmgb2b2 | 1.6E-06 | 0.2169917 | 0.43 | 0.363 | 0.052398 | Synoviocytes | hmgb2b |
| litaf | 1.7E-06 | 0.5111623 | 0.165 | 0.117 | 0.0562 | Synoviocytes | litaf |

| | | | | | | | |
|---|---|---|---|---|---|---|---|
| pgk13 | 1.8E-06 | 0.5363862 | 0.261 | 0.207 | 0.057779 | Synoviocytes | pgk1 |
| rbmx3 | 1.8E-06 | 0.5267592 | 0.176 | 0.127 | 0.059087 | Synoviocytes | rbmx |
| ubfd12 | 1.8E-06 | 0.5468243 | 0.152 | 0.107 | 0.059951 | Synoviocytes | ubfd1 |
| ctbp12 | 1.9E-06 | 0.6622661 | 0.146 | 0.101 | 0.061273 | Synoviocytes | ctbp1 |
| uqcr102 | 1.9E-06 | 0.3799214 | 0.298 | 0.238 | 0.062396 | Synoviocytes | uqcr10 |
| eif4g1a1 | 2E-06 | 0.3219233 | 0.42 | 0.355 | 0.063617 | Synoviocytes | eif4g1a |
| ahr25 | 2E-06 | 0.31821 | 0.445 | 0.383 | 0.064343 | Synoviocytes | ahr2 |
| usp11 | 2E-06 | 0.7321035 | 0.125 | 0.085 | 0.065214 | Synoviocytes | usp1 |
| prex24 | 2E-06 | 0.4552674 | 0.13 | 0.087 | 0.065851 | Synoviocytes | prex2 |
| szrd12 | 2E-06 | 0.5043796 | 0.211 | 0.159 | 0.065947 | Synoviocytes | szrd1 |
| adprh3 | 2E-06 | 0.7522811 | 0.12 | 0.08 | 0.066341 | Synoviocytes | adprh |
| grb2b3 | 2.1E-06 | 0.5337588 | 0.214 | 0.162 | 0.067582 | Synoviocytes | grb2b |
| si:ch211-191j22.31 | 2.1E-06 | 0.5637798 | 0.167 | 0.119 | 0.068011 | Synoviocytes | si:ch211-191j22.3 |
| uqcrc12 | 2.1E-06 | 0.4292788 | 0.236 | 0.181 | 0.068541 | Synoviocytes | uqcrc1 |
| sorbs14 | 2.2E-06 | 0.5790541 | 0.184 | 0.134 | 0.072859 | Synoviocytes | sorbs1 |
| ino802 | 2.3E-06 | 0.6290706 | 0.148 | 0.104 | 0.074147 | Synoviocytes | ino80 |
| micos133 | 2.3E-06 | 0.578755 | 0.169 | 0.122 | 0.074942 | Synoviocytes | micos13 |
| sec133 | 2.4E-06 | 0.4462892 | 0.175 | 0.126 | 0.077877 | Synoviocytes | sec13 |
| stip11 | 2.4E-06 | 0.455383 | 0.212 | 0.16 | 0.078436 | Synoviocytes | stip1 |
| pomp1 | 2.5E-06 | 0.3678621 | 0.399 | 0.341 | 0.080187 | Synoviocytes | pomp |
| smim73 | 2.5E-06 | 0.5750985 | 0.183 | 0.134 | 0.080542 | Synoviocytes | smim7 |
| usp19 | 2.5E-06 | 0.5566531 | 0.124 | 0.083 | 0.082091 | Synoviocytes | usp19 |
| zmiz1a5 | 2.5E-06 | 0.589233 | 0.166 | 0.119 | 0.082165 | Synoviocytes | zmiz1a |
| lzts2a2 | 2.5E-06 | 0.601473 | 0.102 | 0.066 | 0.082925 | Synoviocytes | lzts2a |
| RALGDS3 | 2.6E-06 | 0.6228087 | 0.18 | 0.133 | 0.083544 | Synoviocytes | RALGDS |
| mbnl13 | 2.6E-06 | 0.593573 | 0.107 | 0.07 | 0.083632 | Synoviocytes | mbnl1 |
| naa502 | 2.6E-06 | 0.4824414 | 0.185 | 0.136 | 0.083832 | Synoviocytes | naa50 |
| bnip3lb2 | 2.6E-06 | 0.6159385 | 0.217 | 0.168 | 0.084063 | Synoviocytes | bnip3lb |
| waplb | 2.8E-06 | 0.6653107 | 0.112 | 0.074 | 0.090454 | Synoviocytes | waplb |
| ccdc473 | 2.9E-06 | 0.524801 | 0.187 | 0.139 | 0.092797 | Synoviocytes | ccdc47 |
| mapk14b3 | 3.2E-06 | 0.6584296 | 0.126 | 0.086 | 0.104691 | Synoviocytes | mapk14b |
| thbs2a7 | 3.3E-06 | 0.3415733 | 0.261 | 0.203 | 0.107152 | Synoviocytes | thbs2a |
| ccdc92 | 3.3E-06 | 0.5558772 | 0.118 | 0.078 | 0.107417 | Synoviocytes | ccdc9 |
| elf2b2 | 3.3E-06 | 0.6209834 | 0.156 | 0.112 | 0.107525 | Synoviocytes | elf2b |
| cct44 | 3.4E-06 | 0.3486011 | 0.373 | 0.312 | 0.11116 | Synoviocytes | cct4 |
| CABZ01074397.15 | 3.4E-06 | 0.3788522 | 0.201 | 0.149 | 0.111232 | Synoviocytes | CABZ01074397.1 |
| rif15 | 3.6E-06 | 0.6345435 | 0.125 | 0.085 | 0.118169 | Synoviocytes | rif1 |
| hmgb1b3 | 3.7E-06 | 0.3402051 | 0.314 | 0.252 | 0.12004 | Synoviocytes | hmgb1b |
| seta1 | 3.8E-06 | 0.3820472 | 0.375 | 0.318 | 0.124204 | Synoviocytes | seta |
| phf5a2 | 3.8E-06 | 0.5751002 | 0.154 | 0.11 | 0.124929 | Synoviocytes | phf5a |
| drap12 | 3.9E-06 | 0.4942431 | 0.178 | 0.131 | 0.125472 | Synoviocytes | drap1 |
| crk1 | 3.9E-06 | 0.4868097 | 0.136 | 0.093 | 0.126554 | Synoviocytes | crk |
| rab3il11 | 4.1E-06 | 0.6323464 | 0.101 | 0.065 | 0.134908 | Synoviocytes | rab3il1 |
| btg15 | 4.3E-06 | 0.2159422 | 0.854 | 0.814 | 0.14111 | Synoviocytes | btg1 |
| txndc53 | 4.4E-06 | 0.4470217 | 0.261 | 0.206 | 0.141489 | Synoviocytes | txndc5 |
| dlat2 | 4.4E-06 | 0.5544755 | 0.115 | 0.077 | 0.142636 | Synoviocytes | dlat |
| hdac12 | 4.4E-06 | 0.3350101 | 0.332 | 0.271 | 0.143594 | Synoviocytes | hdac1 |
| pgd3 | 4.4E-06 | 0.7570669 | 0.116 | 0.079 | 0.143602 | Synoviocytes | pgd |

| | | | | | | | |
|---|---|---|---|---|---|---|---|
| syap13 | 4.5E-06 | 0.4548139 | 0.207 | 0.156 | 0.14631 | Synoviocytes | syap1 |
| sorbs34 | 4.6E-06 | 0.3579128 | 0.248 | 0.19 | 0.148334 | Synoviocytes | sorbs3 |
| rbm8a3 | 4.6E-06 | 0.3606205 | 0.404 | 0.353 | 0.148419 | Synoviocytes | rbm8a |
| ndufab1a2 | 4.6E-06 | 0.5465328 | 0.174 | 0.128 | 0.149156 | Synoviocytes | ndufab1a |
| ubl3a5 | 4.7E-06 | 0.3061876 | 0.397 | 0.336 | 0.151234 | Synoviocytes | ubl3a |
| selenoh2 | 4.8E-06 | 0.5119009 | 0.17 | 0.123 | 0.156465 | Synoviocytes | selenoh |
| banf12 | 5.1E-06 | 0.4741276 | 0.174 | 0.127 | 0.165166 | Synoviocytes | banf1 |
| naa203 | 5.2E-06 | 0.4057626 | 0.199 | 0.149 | 0.170076 | Synoviocytes | naa20 |
| wu:fj16a035 | 5.3E-06 | 0.9034195 | 0.243 | 0.191 | 0.171442 | Synoviocytes | wu:fj16a03 |
| ppp6c | 5.5E-06 | 0.6669949 | 0.143 | 0.101 | 0.178882 | Synoviocytes | ppp6c |
| lrrc8aa3 | 5.6E-06 | 0.6537919 | 0.14 | 0.099 | 0.183132 | Synoviocytes | lrrc8aa |
| mob42 | 5.7E-06 | 0.5448631 | 0.171 | 0.126 | 0.186021 | Synoviocytes | mob4 |
| sav11 | 5.8E-06 | 0.6572386 | 0.168 | 0.124 | 0.188898 | Synoviocytes | sav1 |
| psma81 | 5.8E-06 | 0.4359508 | 0.311 | 0.256 | 0.189008 | Synoviocytes | psma8 |
| hectd1 | 5.9E-06 | 0.5076168 | 0.263 | 0.208 | 0.193008 | Synoviocytes | hectd1 |
| chchd6a1 | 6.1E-06 | 0.5204047 | 0.116 | 0.078 | 0.197854 | Synoviocytes | chchd6a |
| stk25b | 6.3E-06 | 0.7117585 | 0.109 | 0.073 | 0.205039 | Synoviocytes | stk25b |
| idh25 | 6.4E-06 | 0.3304473 | 0.406 | 0.343 | 0.208156 | Synoviocytes | idh2 |
| sft2d11 | 6.6E-06 | 0.5949497 | 0.143 | 0.101 | 0.214513 | Synoviocytes | sft2d1 |
| dlgap4a3 | 6.6E-06 | 0.6020049 | 0.133 | 0.093 | 0.214898 | Synoviocytes | dlgap4a |
| zeb2a5 | 6.8E-06 | 0.2972933 | 0.404 | 0.341 | 0.22017 | Synoviocytes | zeb2a |
| gtf2h52 | 6.9E-06 | 0.4901646 | 0.139 | 0.097 | 0.223539 | Synoviocytes | gtf2h5 |
| lamtor42 | 6.9E-06 | 0.590769 | 0.145 | 0.103 | 0.22439 | Synoviocytes | lamtor4 |
| ppp1r26 | 7.1E-06 | 0.5324494 | 0.128 | 0.088 | 0.230221 | Synoviocytes | ppp1r26 |
| micos103 | 7.4E-06 | 0.4194373 | 0.221 | 0.171 | 0.242138 | Synoviocytes | micos10 |
| mtrr2 | 7.5E-06 | 0.6365816 | 0.132 | 0.092 | 0.243802 | Synoviocytes | mtrr |
| atp5pf3 | 7.7E-06 | 0.2559141 | 0.467 | 0.404 | 0.249126 | Synoviocytes | atp5pf |
| zfr2 | 7.8E-06 | 0.6329037 | 0.154 | 0.112 | 0.254389 | Synoviocytes | zfr |
| stau12 | 8E-06 | 0.5515713 | 0.19 | 0.143 | 0.260399 | Synoviocytes | stau1 |
| eaf12 | 8.2E-06 | 0.7447682 | 0.101 | 0.066 | 0.265199 | Synoviocytes | eaf1 |
| guk1a3 | 8.4E-06 | 0.5325657 | 0.182 | 0.136 | 0.273713 | Synoviocytes | guk1a |
| ndufs53 | 8.5E-06 | 0.4989152 | 0.226 | 0.177 | 0.274877 | Synoviocytes | ndufs5 |
| arid5b4 | 8.9E-06 | 0.4790215 | 0.183 | 0.135 | 0.288481 | Synoviocytes | arid5b |
| ppm1bb1 | 9E-06 | 0.5349312 | 0.12 | 0.081 | 0.291088 | Synoviocytes | ppm1bb |
| stat34 | 9E-06 | 0.4675422 | 0.197 | 0.15 | 0.291991 | Synoviocytes | stat3 |
| sept7b3 | 9E-06 | 0.4114696 | 0.25 | 0.198 | 0.293723 | Synoviocytes | sept7b |
| dnajc81 | 9.2E-06 | 0.6838986 | 0.126 | 0.088 | 0.298542 | Synoviocytes | dnajc8 |
| eif4a32 | 9.4E-06 | 0.4073448 | 0.235 | 0.184 | 0.304915 | Synoviocytes | eif4a3 |
| si:ch211-276i12.112 | 9.5E-06 | 0.5259377 | 0.145 | 0.103 | 0.309679 | Synoviocytes | si:ch211-276i12.11 |
| ppp2r5d1 | 9.6E-06 | 0.463634 | 0.158 | 0.114 | 0.312749 | Synoviocytes | ppp2r5d |
| mob3a2 | 9.8E-06 | 0.5969427 | 0.117 | 0.079 | 0.318341 | Synoviocytes | mob3a |
| supt6h2 | 1E-05 | 0.3715474 | 0.25 | 0.196 | 0.332327 | Synoviocytes | supt6h |
| dusp13 | 1E-05 | 0.4396803 | 0.429 | 0.38 | 0.332856 | Synoviocytes | dusp1 |
| glrx33 | 1E-05 | 0.446041 | 0.175 | 0.129 | 0.335017 | Synoviocytes | glrx3 |
| fbxo11a.1 | 1E-05 | 0.473516 | 0.132 | 0.092 | 0.335231 | Synoviocytes | fbxo11a.1 |
| c1galt1b | 1.1E-05 | 0.5485885 | 0.102 | 0.068 | 0.345605 | Synoviocytes | c1galt1b |
| trappc34 | 1.1E-05 | 0.5447262 | 0.148 | 0.106 | 0.366002 | Synoviocytes | trappc3 |
| trim8a3 | 1.1E-05 | 0.6368986 | 0.12 | 0.083 | 0.366788 | Synoviocytes | trim8a |

| | | | | | | | |
|---|---|---|---|---|---|---|---|
| ap1m12 | 1.2E-05 | 0.5942522 | 0.167 | 0.124 | 0.388127 | Synoviocytes | ap1m1 |
| ggh1 | 1.3E-05 | 0.589007 | 0.1 | 0.066 | 0.417787 | Synoviocytes | ggh |
| pum2 | 1.3E-05 | 0.4501764 | 0.116 | 0.079 | 0.422633 | Synoviocytes | pum2 |
| eif1b2 | 1.4E-05 | 0.4608604 | 0.26 | 0.209 | 0.441941 | Synoviocytes | eif1b |
| arfgap23 | 1.4E-05 | 0.5292305 | 0.146 | 0.105 | 0.445831 | Synoviocytes | arfgap2 |
| copb12 | 1.4E-05 | 0.5821496 | 0.19 | 0.146 | 0.447763 | Synoviocytes | copb1 |
| gatad2ab2 | 1.4E-05 | 0.693844 | 0.105 | 0.071 | 0.451786 | Synoviocytes | gatad2ab |
| C19H6orf623 | 1.4E-05 | 0.3017336 | 0.46 | 0.402 | 0.457648 | Synoviocytes | C19H6orf62 |
| g3bp13 | 1.4E-05 | 0.4326463 | 0.216 | 0.169 | 0.458007 | Synoviocytes | g3bp1 |
| csnk2b3 | 1.4E-05 | 0.3761346 | 0.307 | 0.253 | 0.45822 | Synoviocytes | csnk2b |
| vdac23 | 1.4E-05 | 0.2528957 | 0.562 | 0.519 | 0.464424 | Synoviocytes | vdac2 |
| man2b13 | 1.5E-05 | 0.5855154 | 0.136 | 0.097 | 0.471864 | Synoviocytes | man2b1 |
| efna1b3 | 1.5E-05 | 0.4812534 | 0.19 | 0.143 | 0.475487 | Synoviocytes | efna1b |
| ptprk3 | 1.5E-05 | 0.6618207 | 0.103 | 0.069 | 0.492227 | Synoviocytes | ptprk |
| tspan171 | 1.5E-05 | 0.7154405 | 0.113 | 0.077 | 0.49497 | Synoviocytes | tspan17 |
| hm132 | 1.5E-05 | 0.3892431 | 0.188 | 0.141 | 0.501785 | Synoviocytes | hm13 |
| ppp1cab1 | 1.6E-05 | 0.6045501 | 0.101 | 0.067 | 0.527555 | Synoviocytes | ppp1cab |
| eny22 | 1.6E-05 | 0.6408039 | 0.148 | 0.108 | 0.529566 | Synoviocytes | eny2 |
| atp6ap1b2 | 1.7E-05 | 0.7567781 | 0.102 | 0.069 | 0.548122 | Synoviocytes | atp6ap1b |
| nudc1 | 1.7E-05 | 0.2986291 | 0.236 | 0.183 | 0.553518 | Synoviocytes | nudc |
| macf1a2 | 1.7E-05 | 0.3648767 | 0.226 | 0.175 | 0.567838 | Synoviocytes | macf1a |
| rab35b1 | 1.8E-05 | 0.4888013 | 0.13 | 0.092 | 0.572678 | Synoviocytes | rab35b |
| marcksl1a4 | 1.8E-05 | 0.3588806 | 0.586 | 0.711 | 0.586422 | Synoviocytes | marcksl1a |
| uqcrfs13 | 1.8E-05 | 0.3694263 | 0.271 | 0.221 | 0.58995 | Synoviocytes | uqcrfs1 |
| yipf63 | 1.8E-05 | 0.687924 | 0.112 | 0.077 | 0.595198 | Synoviocytes | yipf6 |
| med212 | 1.9E-05 | 0.6378352 | 0.104 | 0.07 | 0.602641 | Synoviocytes | med21 |
| ap2b11 | 2E-05 | 0.3984842 | 0.157 | 0.114 | 0.648953 | Synoviocytes | ap2b1 |
| si:ch73-25f10.6 | 2.1E-05 | 0.5623782 | 0.117 | 0.081 | 0.668108 | Synoviocytes | si:ch73-25f10.6 |
| slirp3 | 2.1E-05 | 0.4341198 | 0.188 | 0.143 | 0.682179 | Synoviocytes | slirp |
| srpra3 | 2.2E-05 | 0.3255981 | 0.274 | 0.219 | 0.702357 | Synoviocytes | srpra |
| slc44a24 | 2.2E-05 | 0.4473309 | 0.157 | 0.115 | 0.724735 | Synoviocytes | slc44a2 |
| cxxc5a1 | 2.3E-05 | 0.5308707 | 0.148 | 0.107 | 0.733855 | Synoviocytes | cxxc5a |
| rnf101 | 2.4E-05 | 0.2781485 | 0.395 | 0.341 | 0.768747 | Synoviocytes | rnf10 |
| eif1axb2 | 2.4E-05 | 0.2835258 | 0.36 | 0.303 | 0.775122 | Synoviocytes | eif1axb |
| ywhae21 | 2.4E-05 | 0.5088207 | 0.187 | 0.143 | 0.786354 | Synoviocytes | ywhae2 |
| ubap2l1 | 2.5E-05 | 0.5011329 | 0.196 | 0.152 | 0.802543 | Synoviocytes | ubap2l |
| pip4k2aa | 2.6E-05 | 0.5848947 | 0.118 | 0.082 | 0.843468 | Synoviocytes | pip4k2aa |
| rbms2a2 | 2.6E-05 | 0.4268654 | 0.189 | 0.143 | 0.854187 | Synoviocytes | rbms2a |
| ENSDARG00000908 | 2.7E-05 | 0.893359 | 0.125 | 0.09 | 0.880839 | Synoviocytes | ENSDARG00000090 |
| si:ch211-22i13.22 | 2.7E-05 | 0.6970776 | 0.108 | 0.074 | 0.890197 | Synoviocytes | si:ch211-22i13.2 |
| eprs14 | 2.8E-05 | 0.6336955 | 0.13 | 0.093 | 0.905769 | Synoviocytes | eprs1 |
| tent4b2 | 2.8E-05 | 0.517783 | 0.25 | 0.204 | 0.916969 | Synoviocytes | tent4b |
| irf2bp2b4 | 2.8E-05 | 0.4017763 | 0.259 | 0.209 | 0.921176 | Synoviocytes | irf2bp2b |
| arl12 | 2.8E-05 | 0.4530876 | 0.211 | 0.165 | 0.921726 | Synoviocytes | arl1 |
| usp42 | 2.9E-05 | 0.5445738 | 0.133 | 0.095 | 0.932906 | Synoviocytes | usp4 |
| tars12 | 2.9E-05 | 0.5306325 | 0.107 | 0.073 | 0.939191 | Synoviocytes | tars1 |
| arglu1b1 | 3E-05 | 0.4723552 | 0.151 | 0.111 | 0.96356 | Synoviocytes | arglu1b |
| odc14 | 3E-05 | 0.5111058 | 0.166 | 0.124 | 0.972764 | Synoviocytes | odc1 |

| | | | | | | | |
|---|---|---|---|---|---|---|---|
| top1l2 | 3E-05 | 0.2547025 | 0.448 | 0.393 | 0.9766 | Synoviocytes | top1l |
| eppk13 | 3E-05 | 0.7843537 | 0.259 | 0.211 | 0.989041 | Synoviocytes | eppk1 |
| nedd4l3 | 3E-05 | 0.5712709 | 0.138 | 0.1 | 0.990087 | Synoviocytes | nedd4l |
| cebpz1 | 3.1E-05 | 0.6780945 | 0.1 | 0.067 | 0.998441 | Synoviocytes | cebpz |
| tomm74 | 3.1E-05 | 0.4226369 | 0.242 | 0.195 | 1 | Synoviocytes | tomm7 |
| nfatc3a3 | 3.2E-05 | 0.4496192 | 0.131 | 0.093 | 1 | Synoviocytes | nfatc3a |
| smg11 | 3.2E-05 | 0.5145586 | 0.1 | 0.067 | 1 | Synoviocytes | smg1 |
| qars12 | 3.2E-05 | 0.5629579 | 0.118 | 0.082 | 1 | Synoviocytes | qars1 |
| cox7a2a3 | 3.2E-05 | 0.2666734 | 0.325 | 0.27 | 1 | Synoviocytes | cox7a2a |
| tram14 | 3.3E-05 | 0.3676097 | 0.34 | 0.287 | 1 | Synoviocytes | tram1 |
| smarcc23 | 3.3E-05 | 0.4260985 | 0.219 | 0.172 | 1 | Synoviocytes | smarcc2 |
| gnai2a2 | 3.4E-05 | 0.3709161 | 0.333 | 0.282 | 1 | Synoviocytes | gnai2a |
| yipf32 | 3.4E-05 | 0.6754313 | 0.104 | 0.072 | 1 | Synoviocytes | yipf3 |
| ube2z1 | 3.5E-05 | 0.6153624 | 0.113 | 0.079 | 1 | Synoviocytes | ube2z |
| lamtor32 | 3.5E-05 | 0.5135904 | 0.116 | 0.081 | 1 | Synoviocytes | lamtor3 |
| fzd22 | 3.5E-05 | 0.4695821 | 0.137 | 0.098 | 1 | Synoviocytes | fzd2 |
| sept153 | 3.5E-05 | 0.3364014 | 0.253 | 0.203 | 1 | Synoviocytes | sept15 |
| gtf2b4 | 3.5E-05 | 0.6015319 | 0.113 | 0.079 | 1 | Synoviocytes | gtf2b |
| ankrd46a1 | 3.6E-05 | 0.5928622 | 0.105 | 0.072 | 1 | Synoviocytes | ankrd46a |
| nbas3 | 3.7E-05 | 0.5288277 | 0.136 | 0.099 | 1 | Synoviocytes | nbas |
| sap30bp1 | 3.8E-05 | 0.6444886 | 0.122 | 0.086 | 1 | Synoviocytes | sap30bp |
| canx1 | 3.9E-05 | 0.2808771 | 0.304 | 0.251 | 1 | Synoviocytes | canx |
| mesd3 | 3.9E-05 | 0.4969057 | 0.193 | 0.15 | 1 | Synoviocytes | mesd |
| si:dkey-177p2.64 | 4E-05 | 0.3467121 | 0.315 | 0.263 | 1 | Synoviocytes | si:dkey-177p2.6 |
| med302 | 4E-05 | 0.5614903 | 0.158 | 0.118 | 1 | Synoviocytes | med30 |
| eif3ja2 | 4.1E-05 | 0.4698861 | 0.152 | 0.112 | 1 | Synoviocytes | eif3ja |
| pa2g4b2 | 4.1E-05 | 0.3478452 | 0.304 | 0.25 | 1 | Synoviocytes | pa2g4b |
| snx62 | 4.2E-05 | 0.3905941 | 0.158 | 0.117 | 1 | Synoviocytes | snx6 |
| tcp12 | 4.3E-05 | 0.3263163 | 0.375 | 0.322 | 1 | Synoviocytes | tcp1 |
| epn1 | 4.4E-05 | 0.5276989 | 0.121 | 0.085 | 1 | Synoviocytes | epn1 |
| rtf11 | 4.6E-05 | 0.340494 | 0.303 | 0.251 | 1 | Synoviocytes | rtf1 |
| rbm341 | 4.6E-05 | 0.7644798 | 0.103 | 0.071 | 1 | Synoviocytes | rbm34 |
| pgrmc12 | 4.7E-05 | 0.2471589 | 0.419 | 0.363 | 1 | Synoviocytes | pgrmc1 |
| gigyf2 | 4.8E-05 | 0.5259323 | 0.147 | 0.108 | 1 | Synoviocytes | gigyf2 |
| pthlha3 | 4.8E-05 | 0.5694967 | 0.105 | 0.072 | 1 | Synoviocytes | pthlha |
| phc2b3 | 5E-05 | 0.9232986 | 0.107 | 0.075 | 1 | Synoviocytes | phc2b |
| rpl7l11 | 5.1E-05 | 0.3962184 | 0.176 | 0.134 | 1 | Synoviocytes | rpl7l1 |
| stau23 | 5.2E-05 | 0.2671351 | 0.487 | 0.439 | 1 | Synoviocytes | stau2 |
| rad21a2 | 5.3E-05 | 0.4122248 | 0.247 | 0.201 | 1 | Synoviocytes | rad21a |
| sik2b4 | 5.3E-05 | 0.4505504 | 0.253 | 0.207 | 1 | Synoviocytes | sik2b |
| hnrnpa1a2 | 5.3E-05 | 0.4133239 | 0.179 | 0.136 | 1 | Synoviocytes | hnrnpa1a |
| cops42 | 5.4E-05 | 0.5065528 | 0.149 | 0.111 | 1 | Synoviocytes | cops4 |
| prkag11 | 5.4E-05 | 0.4994292 | 0.133 | 0.096 | 1 | Synoviocytes | prkag1 |
| ctnnd12 | 5.5E-05 | 0.4830751 | 0.211 | 0.166 | 1 | Synoviocytes | ctnnd1 |
| rnf19a2 | 5.5E-05 | 0.3601725 | 0.21 | 0.164 | 1 | Synoviocytes | rnf19a |
| hgs1 | 5.6E-05 | 0.5210821 | 0.125 | 0.089 | 1 | Synoviocytes | hgs |
| lin7c1 | 5.7E-05 | 0.4783475 | 0.137 | 0.099 | 1 | Synoviocytes | lin7c |
| C23H1orf521 | 5.9E-05 | 0.4201183 | 0.147 | 0.108 | 1 | Synoviocytes | C23H1orf52 |

| | | | | | | | |
|---|---|---|---|---|---|---|---|
| golim4a.12 | 6E-05 | 0.4937685 | 0.134 | 0.097 | 1 | Synoviocytes | golim4a.1 |
| ppp3r1b2 | 6.1E-05 | 0.4605809 | 0.175 | 0.134 | 1 | Synoviocytes | ppp3r1b |
| flii2 | 6.2E-05 | 0.4722556 | 0.148 | 0.109 | 1 | Synoviocytes | flii |
| opa11 | 6.2E-05 | 0.4711444 | 0.112 | 0.078 | 1 | Synoviocytes | opa1 |
| pak2b2 | 6.4E-05 | 0.4441217 | 0.19 | 0.148 | 1 | Synoviocytes | pak2b |
| CABZ01061894.11 | 6.5E-05 | 0.5454408 | 0.135 | 0.098 | 1 | Synoviocytes | CABZ01061894.1 |
| twist1b5 | 6.5E-05 | 0.4435075 | 0.203 | 0.159 | 1 | Synoviocytes | twist1b |
| cnot92 | 6.6E-05 | 0.4556167 | 0.111 | 0.078 | 1 | Synoviocytes | cnot9 |
| kmt5aa4 | 6.7E-05 | 0.5350549 | 0.152 | 0.114 | 1 | Synoviocytes | kmt5aa |
| tmem14ca3 | 6.7E-05 | 0.5232698 | 0.161 | 0.122 | 1 | Synoviocytes | tmem14ca |
| jak14 | 7E-05 | 0.2363805 | 0.493 | 0.437 | 1 | Synoviocytes | jak1 |
| knop12 | 7.1E-05 | 0.4288075 | 0.207 | 0.163 | 1 | Synoviocytes | knop1 |
| cnppd11 | 7.1E-05 | 0.5719697 | 0.107 | 0.075 | 1 | Synoviocytes | cnppd1 |
| snx12 | 7.1E-05 | 0.3074083 | 0.318 | 0.27 | 1 | Synoviocytes | snx12 |
| ezh14 | 7.2E-05 | 0.4894404 | 0.182 | 0.141 | 1 | Synoviocytes | ezh1 |
| myo1b1 | 7.5E-05 | 0.3004071 | 0.157 | 0.117 | 1 | Synoviocytes | myo1b |
| smarcd13 | 7.6E-05 | 0.5045358 | 0.1 | 0.068 | 1 | Synoviocytes | smarcd1 |
| snrpe1 | 7.7E-05 | 0.3033113 | 0.297 | 0.247 | 1 | Synoviocytes | snrpe |
| dpm23 | 8.1E-05 | 0.4773769 | 0.18 | 0.139 | 1 | Synoviocytes | dpm2 |
| cpvl4 | 8.2E-05 | 0.6143819 | 0.127 | 0.093 | 1 | Synoviocytes | cpvl |
| stk35l3 | 8.2E-05 | 0.682705 | 0.163 | 0.126 | 1 | Synoviocytes | stk35l |
| atp5md2 | 8.3E-05 | 0.3741507 | 0.187 | 0.143 | 1 | Synoviocytes | atp5md |
| nansa1 | 8.4E-05 | 0.6575643 | 0.104 | 0.073 | 1 | Synoviocytes | nansa |
| gnas1 | 8.5E-05 | 0.416004 | 0.198 | 0.155 | 1 | Synoviocytes | gnas |
| tgfb55 | 8.5E-05 | 0.4881025 | 0.151 | 0.114 | 1 | Synoviocytes | tgfb5 |
| stx5a2 | 8.6E-05 | 0.658689 | 0.133 | 0.099 | 1 | Synoviocytes | stx5a |
| tpd52l2b3 | 8.9E-05 | 0.397574 | 0.265 | 0.219 | 1 | Synoviocytes | tpd52l2b |
| sart12 | 9E-05 | 0.3001022 | 0.213 | 0.167 | 1 | Synoviocytes | sart1 |
| irf2a2 | 9E-05 | 0.6002168 | 0.118 | 0.084 | 1 | Synoviocytes | irf2a |
| pcnp1 | 9.3E-05 | 0.3647366 | 0.2 | 0.157 | 1 | Synoviocytes | pcnp |
| gripap11 | 9.3E-05 | 0.3901774 | 0.159 | 0.119 | 1 | Synoviocytes | gripap1 |
| usf1l3 | 9.6E-05 | 0.4508419 | 0.102 | 0.071 | 1 | Synoviocytes | usf1l |
| C20H12orf572 | 0.0001 | 0.5680741 | 0.125 | 0.091 | 1 | Synoviocytes | C20H12orf57 |
| rbms1a1 | 0.0001 | 0.611176 | 0.107 | 0.076 | 1 | Synoviocytes | rbms1a |
| nupr1a4 | 0.0001 | 0.4250567 | 0.3 | 0.256 | 1 | Synoviocytes | nupr1a |
| txlna2 | 0.00011 | 0.5073428 | 0.138 | 0.102 | 1 | Synoviocytes | txlna |
| phf3 | 0.00011 | 0.542901 | 0.161 | 0.123 | 1 | Synoviocytes | phf3 |
| zgc:1585641 | 0.00011 | 0.3086005 | 0.281 | 0.232 | 1 | Synoviocytes | zgc:158564 |
| sdhb3 | 0.00011 | 0.3705899 | 0.219 | 0.176 | 1 | Synoviocytes | sdhb |
| prpf40a1 | 0.00011 | 0.5093772 | 0.162 | 0.124 | 1 | Synoviocytes | prpf40a |
| ube2g23 | 0.00011 | 0.4747385 | 0.148 | 0.111 | 1 | Synoviocytes | ube2g2 |
| fam76b2 | 0.00011 | 0.54378 | 0.118 | 0.085 | 1 | Synoviocytes | fam76b |
| luc7l1 | 0.00012 | 0.4622824 | 0.188 | 0.146 | 1 | Synoviocytes | luc7l |
| zc3h152 | 0.00012 | 0.3578607 | 0.182 | 0.141 | 1 | Synoviocytes | zc3h15 |
| cep350 | 0.00012 | 0.4500869 | 0.148 | 0.11 | 1 | Synoviocytes | cep350 |
| polr2eb2 | 0.00012 | 0.331664 | 0.193 | 0.151 | 1 | Synoviocytes | polr2eb |
| caprin1a1 | 0.00012 | 0.4294948 | 0.139 | 0.102 | 1 | Synoviocytes | caprin1a |
| tm2d32 | 0.00012 | 0.5772237 | 0.102 | 0.071 | 1 | Synoviocytes | tm2d3 |

| | | | | | | | |
|---|---|---|---|---|---|---|---|
| nphp3 | 0.00012 | 0.4455759 | 0.152 | 0.113 | 1 | Synoviocytes | nphp3 |
| gorasp24 | 0.00012 | 0.5621684 | 0.147 | 0.111 | 1 | Synoviocytes | gorasp2 |
| usp9 | 0.00013 | 0.367361 | 0.284 | 0.237 | 1 | Synoviocytes | usp9 |
| atg131 | 0.00013 | 0.5199104 | 0.111 | 0.079 | 1 | Synoviocytes | atg13 |
| dync1i2a2 | 0.00013 | 0.3993805 | 0.151 | 0.113 | 1 | Synoviocytes | dync1i2a |
| mapk12 | 0.00013 | 0.5262727 | 0.117 | 0.084 | 1 | Synoviocytes | mapk1 |
| nkap2 | 0.00013 | 0.3848729 | 0.233 | 0.188 | 1 | Synoviocytes | nkap |
| phldb1b4 | 0.00013 | 0.3820945 | 0.123 | 0.088 | 1 | Synoviocytes | phldb1b |
| selenon4 | 0.00014 | 0.3949373 | 0.197 | 0.157 | 1 | Synoviocytes | selenon |
| slc20a22 | 0.00014 | 0.3927609 | 0.319 | 0.272 | 1 | Synoviocytes | slc20a2 |
| yif1a2 | 0.00014 | 0.4775513 | 0.162 | 0.124 | 1 | Synoviocytes | yif1a |
| wipf2a | 0.00014 | 0.5731425 | 0.119 | 0.086 | 1 | Synoviocytes | wipf2a |
| serpinh24 | 0.00014 | 0.5184214 | 0.155 | 0.118 | 1 | Synoviocytes | serpinh2 |
| cfap202 | 0.00014 | 0.6424616 | 0.102 | 0.072 | 1 | Synoviocytes | cfap20 |
| rnf21 | 0.00014 | 0.3991189 | 0.156 | 0.118 | 1 | Synoviocytes | rnf2 |
| mfap12 | 0.00014 | 0.4738875 | 0.177 | 0.138 | 1 | Synoviocytes | mfap1 |
| snrpb21 | 0.00014 | 0.5246051 | 0.102 | 0.072 | 1 | Synoviocytes | snrpb2 |
| egfra2 | 0.00014 | 0.3569603 | 0.307 | 0.258 | 1 | Synoviocytes | egfra |
| rsrc2 | 0.00015 | 0.4400966 | 0.2 | 0.159 | 1 | Synoviocytes | rsrc2 |
| kctd101 | 0.00015 | 0.5699567 | 0.104 | 0.073 | 1 | Synoviocytes | kctd10 |
| usp25 | 0.00015 | 0.6983307 | 0.109 | 0.078 | 1 | Synoviocytes | usp25 |
| ubr41 | 0.00016 | 0.4406597 | 0.135 | 0.1 | 1 | Synoviocytes | ubr4 |
| rab22a3 | 0.00017 | 0.4759568 | 0.137 | 0.102 | 1 | Synoviocytes | rab22a |
| rab34a3 | 0.00017 | 0.4723722 | 0.155 | 0.118 | 1 | Synoviocytes | rab34a |
| ppp2r1ba1 | 0.00017 | 0.3704205 | 0.197 | 0.156 | 1 | Synoviocytes | ppp2r1ba |
| zgc:1935413 | 0.00017 | 0.2497561 | 0.443 | 0.393 | 1 | Synoviocytes | zgc:193541 |
| slc43a1a3 | 0.00018 | 0.5542209 | 0.114 | 0.082 | 1 | Synoviocytes | slc43a1a |
| dido11 | 0.00018 | 0.2275118 | 0.471 | 0.426 | 1 | Synoviocytes | dido1 |
| sf3a21 | 0.00018 | 0.3817046 | 0.2 | 0.159 | 1 | Synoviocytes | sf3a2 |
| acin1a1 | 0.00018 | 0.2622328 | 0.621 | 0.593 | 1 | Synoviocytes | acin1a |
| ctdsp12 | 0.00018 | 0.4284536 | 0.149 | 0.113 | 1 | Synoviocytes | ctdsp1 |
| hnrnpr2 | 0.00019 | 0.4894309 | 0.157 | 0.12 | 1 | Synoviocytes | hnrnpr |
| hsd17b12b2 | 0.00019 | 0.6565329 | 0.122 | 0.09 | 1 | Synoviocytes | hsd17b12b |
| bloc1s22 | 0.00019 | 0.6330631 | 0.102 | 0.073 | 1 | Synoviocytes | bloc1s2 |
| cilp5 | 0.00019 | 0.6451895 | 0.212 | 0.172 | 1 | Synoviocytes | cilp |
| si:ch211-235e9.63 | 0.00019 | 0.4950995 | 0.126 | 0.093 | 1 | Synoviocytes | si:ch211-235e9.6 |
| dnajc15 | 0.00019 | 0.3068121 | 0.33 | 0.283 | 1 | Synoviocytes | dnajc1 |
| ndufa102 | 0.0002 | 0.335962 | 0.156 | 0.119 | 1 | Synoviocytes | ndufa10 |
| cox5ab4 | 0.0002 | 0.3384496 | 0.268 | 0.22 | 1 | Synoviocytes | cox5ab |
| ddx3xb1 | 0.00021 | 0.2690255 | 0.199 | 0.157 | 1 | Synoviocytes | ddx3xb |
| cars12 | 0.00021 | 0.3702108 | 0.113 | 0.081 | 1 | Synoviocytes | cars1 |
| faub3 | 0.00021 | 0.5314295 | 0.118 | 0.086 | 1 | Synoviocytes | faub |
| CIR12 | 0.00022 | 0.4160754 | 0.115 | 0.083 | 1 | Synoviocytes | CIR1 |
| tnrc6b2 | 0.00022 | 0.3735051 | 0.177 | 0.139 | 1 | Synoviocytes | tnrc6b |
| ccdc6a1 | 0.00023 | 0.446668 | 0.15 | 0.114 | 1 | Synoviocytes | ccdc6a |
| hdgfl21 | 0.00023 | 0.436508 | 0.22 | 0.178 | 1 | Synoviocytes | hdgfl2 |
| cyc13 | 0.00023 | 0.4430401 | 0.21 | 0.17 | 1 | Synoviocytes | cyc1 |
| atxn2l2 | 0.00023 | 0.4507229 | 0.125 | 0.092 | 1 | Synoviocytes | atxn2l |

| | | | | | | | |
|---|---|---|---|---|---|---|---|
| srp193 | 0.00023 | 0.3847982 | 0.18 | 0.141 | 1 | Synoviocytes | srp19 |
| nrd1a1 | 0.00024 | 0.507911 | 0.122 | 0.089 | 1 | Synoviocytes | nrd1a |
| uso12 | 0.00024 | 0.4042046 | 0.155 | 0.118 | 1 | Synoviocytes | uso1 |
| fam193a2 | 0.00025 | 0.4353908 | 0.138 | 0.103 | 1 | Synoviocytes | fam193a |
| tax1bp1b1 | 0.00025 | 0.2617323 | 0.332 | 0.281 | 1 | Synoviocytes | tax1bp1b |
| pam161 | 0.00025 | 0.4279575 | 0.145 | 0.11 | 1 | Synoviocytes | pam16 |
| acadvl1 | 0.00025 | 0.6043237 | 0.131 | 0.099 | 1 | Synoviocytes | acadvl |
| kansl1b2 | 0.00025 | 0.4357888 | 0.211 | 0.17 | 1 | Synoviocytes | kansl1b |
| rtn4a2 | 0.00026 | 0.2527931 | 0.494 | 0.45 | 1 | Synoviocytes | rtn4a |
| si:dkey-33c12.43 | 0.00026 | 0.2872147 | 0.391 | 0.342 | 1 | Synoviocytes | si:dkey-33c12.4 |
| gnl32 | 0.00026 | 0.4866815 | 0.177 | 0.14 | 1 | Synoviocytes | gnl3 |
| cd151l4 | 0.00027 | 0.5367619 | 0.103 | 0.074 | 1 | Synoviocytes | cd151l |
| tpra1 | 0.00027 | 0.8009784 | 0.101 | 0.072 | 1 | Synoviocytes | tpra |
| pdgfrb4 | 0.00027 | 0.2317133 | 0.461 | 0.411 | 1 | Synoviocytes | pdgfrb |
| rpn13 | 0.00027 | 0.2945796 | 0.257 | 0.212 | 1 | Synoviocytes | rpn1 |
| wipi15 | 0.00028 | 0.4862458 | 0.157 | 0.121 | 1 | Synoviocytes | wipi1 |
| plxdc23 | 0.0003 | 0.283877 | 0.2 | 0.158 | 1 | Synoviocytes | plxdc2 |
| psenen2 | 0.0003 | 0.4071929 | 0.162 | 0.125 | 1 | Synoviocytes | psenen |
| si:dkey-188i13.103 | 0.0003 | 0.2604139 | 0.296 | 0.248 | 1 | Synoviocytes | si:dkey-188i13.10 |
| rnd3b5 | 0.00031 | 0.342711 | 0.375 | 0.331 | 1 | Synoviocytes | rnd3b |
| btg34 | 0.00031 | 0.3941285 | 0.163 | 0.126 | 1 | Synoviocytes | btg3 |
| zgc:1623392 | 0.00033 | 0.4311191 | 0.102 | 0.072 | 1 | Synoviocytes | zgc:162339 |
| ube2kb1 | 0.00033 | 0.6138601 | 0.112 | 0.082 | 1 | Synoviocytes | ube2kb |
| tmed35 | 0.00033 | 0.4789385 | 0.129 | 0.096 | 1 | Synoviocytes | tmed3 |
| parvaa1 | 0.00033 | 0.2862228 | 0.251 | 0.206 | 1 | Synoviocytes | parvaa |
| zfand31 | 0.00035 | 0.3298171 | 0.23 | 0.188 | 1 | Synoviocytes | zfand3 |
| birc21 | 0.00035 | 0.4262664 | 0.172 | 0.136 | 1 | Synoviocytes | birc2 |
| ndufb102 | 0.00035 | 0.3013476 | 0.279 | 0.236 | 1 | Synoviocytes | ndufb10 |
| rbbp93 | 0.00037 | 0.4603151 | 0.16 | 0.125 | 1 | Synoviocytes | rbbp9 |
| pdcd10b3 | 0.00037 | 0.5309112 | 0.132 | 0.1 | 1 | Synoviocytes | pdcd10b |
| ndufb32 | 0.00037 | 0.3355928 | 0.323 | 0.278 | 1 | Synoviocytes | ndufb3 |
| slmapa1 | 0.00038 | 0.461123 | 0.132 | 0.099 | 1 | Synoviocytes | slmapa |
| nufip23 | 0.00038 | 0.402597 | 0.187 | 0.148 | 1 | Synoviocytes | nufip2 |
| chordc1b2 | 0.00039 | 0.3923965 | 0.135 | 0.102 | 1 | Synoviocytes | chordc1b |
| vma213 | 0.0004 | 0.46599 | 0.148 | 0.114 | 1 | Synoviocytes | vma21 |
| cyp20a12 | 0.00041 | 0.3789369 | 0.147 | 0.113 | 1 | Synoviocytes | cyp20a1 |
| tmed103 | 0.00041 | 0.2643863 | 0.385 | 0.337 | 1 | Synoviocytes | tmed10 |
| lamtor54 | 0.00041 | 0.4838455 | 0.119 | 0.088 | 1 | Synoviocytes | lamtor5 |
| copg23 | 0.00041 | 0.4512851 | 0.186 | 0.149 | 1 | Synoviocytes | copg2 |
| hbp14 | 0.00041 | 0.4825071 | 0.147 | 0.112 | 1 | Synoviocytes | hbp1 |
| tm9sf33 | 0.00042 | 0.2919024 | 0.215 | 0.172 | 1 | Synoviocytes | tm9sf3 |
| lsm53 | 0.00042 | 0.4955941 | 0.151 | 0.118 | 1 | Synoviocytes | lsm5 |
| sh3glb1a | 0.00043 | 0.5825964 | 0.106 | 0.077 | 1 | Synoviocytes | sh3glb1a |
| cyb5a1 | 0.00043 | 0.4852273 | 0.122 | 0.09 | 1 | Synoviocytes | cyb5a |
| lsm12b2 | 0.00044 | 0.3274474 | 0.375 | 0.335 | 1 | Synoviocytes | lsm12b |
| prss165 | 0.00045 | 0.586167 | 0.121 | 0.09 | 1 | Synoviocytes | prss16 |
| myo18ab | 0.00045 | 0.3097908 | 0.229 | 0.187 | 1 | Synoviocytes | myo18ab |
| aktip2 | 0.00046 | 0.6208506 | 0.128 | 0.097 | 1 | Synoviocytes | aktip |

| | | | | | | | |
|---|---|---|---|---|---|---|---|
| creb3l23 | 0.00047 | 0.2837204 | 0.465 | 0.426 | 1 | Synoviocytes | creb3l2 |
| snx33 | 0.00047 | 0.3094832 | 0.219 | 0.178 | 1 | Synoviocytes | snx3 |
| atp6ap23 | 0.00048 | 0.4404852 | 0.133 | 0.101 | 1 | Synoviocytes | atp6ap2 |
| arid4a | 0.00049 | 0.3921714 | 0.207 | 0.169 | 1 | Synoviocytes | arid4a |
| nhsl24 | 0.00049 | 0.4908474 | 0.134 | 0.102 | 1 | Synoviocytes | nhsl2 |
| mtch21 | 0.00049 | 0.4088467 | 0.134 | 0.101 | 1 | Synoviocytes | mtch2 |
| si:ch211-194e15.53 | 0.00049 | 0.5014393 | 0.136 | 0.103 | 1 | Synoviocytes | si:ch211-194e15.5 |
| MZT11 | 0.0005 | 0.5581402 | 0.104 | 0.076 | 1 | Synoviocytes | MZT1 |
| tes2 | 0.0005 | 0.4136153 | 0.103 | 0.075 | 1 | Synoviocytes | tes |
| dhx381 | 0.00051 | 0.5267333 | 0.13 | 0.099 | 1 | Synoviocytes | dhx38 |
| llph2 | 0.00052 | 0.5616668 | 0.113 | 0.084 | 1 | Synoviocytes | llph |
| socs5b1 | 0.00053 | 0.3636525 | 0.124 | 0.092 | 1 | Synoviocytes | socs5b |
| lman21 | 0.00053 | 0.3571985 | 0.276 | 0.236 | 1 | Synoviocytes | lman2 |
| irf22 | 0.00053 | 0.3357919 | 0.186 | 0.149 | 1 | Synoviocytes | irf2 |
| arl6ip13 | 0.00054 | 0.3383806 | 0.2 | 0.161 | 1 | Synoviocytes | arl6ip1 |
| rbm25b1 | 0.00054 | 0.4044641 | 0.249 | 0.209 | 1 | Synoviocytes | rbm25b |
| sf3b21 | 0.00055 | 0.3285518 | 0.244 | 0.203 | 1 | Synoviocytes | sf3b2 |
| slc66a34 | 0.00055 | 0.4116143 | 0.152 | 0.119 | 1 | Synoviocytes | slc66a3 |
| psmb22 | 0.00055 | 0.3031161 | 0.307 | 0.266 | 1 | Synoviocytes | psmb2 |
| cavin1b5 | 0.00056 | 0.3238176 | 0.648 | 0.647 | 1 | Synoviocytes | cavin1b |
| sdha2 | 0.00059 | 0.3751863 | 0.189 | 0.152 | 1 | Synoviocytes | sdha |
| snrpd11 | 0.0006 | 0.257919 | 0.238 | 0.197 | 1 | Synoviocytes | snrpd1 |
| dhx91 | 0.0006 | 0.4462581 | 0.102 | 0.074 | 1 | Synoviocytes | dhx9 |
| ndufb63 | 0.00061 | 0.4002253 | 0.22 | 0.181 | 1 | Synoviocytes | ndufb6 |
| magoh1 | 0.00062 | 0.4033099 | 0.166 | 0.131 | 1 | Synoviocytes | magoh |
| prrc2c1 | 0.00062 | 0.3239159 | 0.151 | 0.117 | 1 | Synoviocytes | prrc2c |
| creg11 | 0.00063 | 0.5290015 | 0.1 | 0.072 | 1 | Synoviocytes | creg1 |
| snrnp2001 | 0.00064 | 0.5227896 | 0.145 | 0.113 | 1 | Synoviocytes | snrnp200 |
| psmb62 | 0.00065 | 0.3290867 | 0.355 | 0.315 | 1 | Synoviocytes | psmb6 |
| phb2 | 0.00065 | 0.4277161 | 0.161 | 0.126 | 1 | Synoviocytes | phb |
| zgc:565254 | 0.00065 | 0.3223059 | 0.193 | 0.154 | 1 | Synoviocytes | zgc:56525 |
| syf22 | 0.00067 | 0.2717856 | 0.328 | 0.282 | 1 | Synoviocytes | syf2 |
| dpm12 | 0.00068 | 0.4395853 | 0.116 | 0.087 | 1 | Synoviocytes | dpm1 |
| mtdhb2 | 0.00071 | 0.4147606 | 0.19 | 0.153 | 1 | Synoviocytes | mtdhb |
| krt18a.15 | 0.00071 | 0.2017862 | 0.842 | 0.842 | 1 | Synoviocytes | krt18a.1 |
| mycbp4 | 0.00072 | 0.3177903 | 0.291 | 0.248 | 1 | Synoviocytes | mycbp |
| trappc12 | 0.00072 | 0.3119313 | 0.182 | 0.145 | 1 | Synoviocytes | trappc1 |
| hspa4b2 | 0.00073 | 0.4354127 | 0.178 | 0.143 | 1 | Synoviocytes | hspa4b |
| six2a4 | 0.00074 | 0.6126016 | 0.132 | 0.102 | 1 | Synoviocytes | six2a |
| lmnb21 | 0.00077 | 0.3123986 | 0.131 | 0.099 | 1 | Synoviocytes | lmnb2 |
| srsf1a1 | 0.00077 | 0.3340215 | 0.208 | 0.17 | 1 | Synoviocytes | srsf1a |
| hikeshi3 | 0.00078 | 0.4506078 | 0.134 | 0.103 | 1 | Synoviocytes | hikeshi |
| phf141 | 0.00078 | 0.4372211 | 0.15 | 0.117 | 1 | Synoviocytes | phf14 |
| kctd5a5 | 0.00078 | 0.2629075 | 0.364 | 0.319 | 1 | Synoviocytes | kctd5a |
| slc25a243 | 0.00079 | 0.2752659 | 0.259 | 0.219 | 1 | Synoviocytes | slc25a24 |
| gpia3 | 0.00079 | 0.5392667 | 0.16 | 0.128 | 1 | Synoviocytes | gpia |
| ERC11 | 0.0008 | 0.4206711 | 0.105 | 0.077 | 1 | Synoviocytes | ERC1 |
| glyr12 | 0.0008 | 0.4549961 | 0.133 | 0.102 | 1 | Synoviocytes | glyr1 |

| | | | | | | | |
|---|---|---|---|---|---|---|---|
| mpc23 | 0.0008 | 0.2768892 | 0.221 | 0.181 | 1 | Synoviocytes | mpc2 |
| cblb4 | 0.00081 | 0.4494566 | 0.127 | 0.097 | 1 | Synoviocytes | cblb |
| vkorc1l12 | 0.00081 | 0.458913 | 0.12 | 0.09 | 1 | Synoviocytes | vkorc1l1 |
| snrpc1 | 0.00083 | 0.4344406 | 0.134 | 0.102 | 1 | Synoviocytes | snrpc |
| cul3b | 0.00083 | 0.5963838 | 0.102 | 0.076 | 1 | Synoviocytes | cul3b |
| capza1b3 | 0.00085 | 0.215884 | 0.408 | 0.367 | 1 | Synoviocytes | capza1b |
| myl12.23 | 0.00086 | 0.4370105 | 0.203 | 0.168 | 1 | Synoviocytes | myl12.2 |
| si:ch211-286b5.52 | 0.00088 | 0.3703788 | 0.295 | 0.258 | 1 | Synoviocytes | si:ch211-286b5.5 |
| pfdn11 | 0.00088 | 0.3520421 | 0.233 | 0.195 | 1 | Synoviocytes | pfdn1 |
| sec23a2 | 0.0009 | 0.2554097 | 0.185 | 0.147 | 1 | Synoviocytes | sec23a |
| nipblb1 | 0.00091 | 0.2808618 | 0.231 | 0.191 | 1 | Synoviocytes | nipblb |
| si:dkey-67c22.21 | 0.00093 | 0.2771253 | 0.269 | 0.228 | 1 | Synoviocytes | si:dkey-67c22.2 |
| prkcsh3 | 0.00094 | 0.3124159 | 0.17 | 0.135 | 1 | Synoviocytes | prkcsh |
| taf1 | 0.00094 | 0.4645225 | 0.125 | 0.095 | 1 | Synoviocytes | taf1 |
| kctd9a3 | 0.00095 | 0.3215594 | 0.128 | 0.097 | 1 | Synoviocytes | kctd9a |
| tusc2b2 | 0.00098 | 0.6014203 | 0.105 | 0.078 | 1 | Synoviocytes | tusc2b |
| si:dkey-286j15.11 | 0.001 | 0.273791 | 0.193 | 0.157 | 1 | Synoviocytes | si:dkey-286j15.1 |
| cct31 | 0.001 | 0.319083 | 0.364 | 0.331 | 1 | Synoviocytes | cct3 |
| parp33 | 0.00101 | 0.4848131 | 0.116 | 0.087 | 1 | Synoviocytes | parp3 |
| grinab3 | 0.00102 | 0.339966 | 0.249 | 0.212 | 1 | Synoviocytes | grinab |
| tmem112 | 0.00104 | 0.3831074 | 0.134 | 0.103 | 1 | Synoviocytes | tmem11 |
| si:dkey-19a16.2 | 0.00106 | 0.5301032 | 0.103 | 0.076 | 1 | Synoviocytes | si:dkey-19a16.2 |
| tcima5 | 0.00106 | 0.2829388 | 0.271 | 0.226 | 1 | Synoviocytes | tcima |
| RAPH14 | 0.00108 | 0.3700891 | 0.12 | 0.09 | 1 | Synoviocytes | RAPH1 |
| tia11 | 0.00108 | 0.4661238 | 0.122 | 0.093 | 1 | Synoviocytes | tia1 |
| tmbim43 | 0.0011 | 0.3654741 | 0.148 | 0.116 | 1 | Synoviocytes | tmbim4 |
| azin1b | 0.00115 | 0.4965343 | 0.129 | 0.1 | 1 | Synoviocytes | azin1b |
| ggnbp2 | 0.00115 | 0.4566789 | 0.104 | 0.078 | 1 | Synoviocytes | ggnbp2 |
| KIF2A2 | 0.00117 | 0.3042512 | 0.142 | 0.109 | 1 | Synoviocytes | KIF2A |
| flot1b | 0.00118 | 0.3803676 | 0.219 | 0.183 | 1 | Synoviocytes | flot1b |
| adam17a1 | 0.00119 | 0.4321551 | 0.128 | 0.098 | 1 | Synoviocytes | adam17a |
| CR383676.36 | 0.00119 | 0.2630211 | 0.511 | 0.466 | 1 | Synoviocytes | CR383676.3 |
| fbxo11a1 | 0.00123 | 0.3148717 | 0.134 | 0.103 | 1 | Synoviocytes | fbxo11a |
| pdha1a2 | 0.00124 | 0.391143 | 0.17 | 0.136 | 1 | Synoviocytes | pdha1a |
| si:ch73-376l24.21 | 0.00125 | 0.3304043 | 0.109 | 0.081 | 1 | Synoviocytes | si:ch73-376l24.2 |
| tomm221 | 0.00126 | 0.3809699 | 0.129 | 0.099 | 1 | Synoviocytes | tomm22 |
| hspa4a5 | 0.00128 | 0.2290816 | 0.185 | 0.149 | 1 | Synoviocytes | hspa4a |
| zgc:1752143 | 0.00129 | 0.3670956 | 0.138 | 0.108 | 1 | Synoviocytes | zgc:175214 |
| yipf53 | 0.00135 | 0.4128065 | 0.155 | 0.123 | 1 | Synoviocytes | yipf5 |
| si:dkey-88p24.112 | 0.00136 | 0.4543601 | 0.108 | 0.081 | 1 | Synoviocytes | si:dkey-88p24.11 |
| mcts12 | 0.00136 | 0.2828431 | 0.208 | 0.171 | 1 | Synoviocytes | mcts1 |
| rhoq5 | 0.00136 | 0.4581042 | 0.125 | 0.096 | 1 | Synoviocytes | rhoq |
| ppm1g1 | 0.00138 | 0.3274916 | 0.191 | 0.156 | 1 | Synoviocytes | ppm1g |
| spoplb2 | 0.00139 | 0.4250932 | 0.127 | 0.098 | 1 | Synoviocytes | spoplb |
| phactr26 | 0.0014 | 0.2705809 | 0.177 | 0.141 | 1 | Synoviocytes | phactr2 |
| mpc14 | 0.00141 | 0.4516082 | 0.199 | 0.166 | 1 | Synoviocytes | mpc1 |
| srsf6b | 0.0015 | 0.3659512 | 0.148 | 0.116 | 1 | Synoviocytes | srsf6b |
| cd1644 | 0.00153 | 0.2745468 | 0.375 | 0.338 | 1 | Synoviocytes | cd164 |

| | | | | | | | |
|---|---|---|---|---|---|---|---|
| uri13 | 0.00154 | 0.3629009 | 0.162 | 0.13 | 1 | Synoviocytes | uri1 |
| csnk1a11 | 0.00154 | 0.2384246 | 0.418 | 0.381 | 1 | Synoviocytes | csnk1a1 |
| ndufb93 | 0.00155 | 0.2529373 | 0.35 | 0.309 | 1 | Synoviocytes | ndufb9 |
| ppp6r3 | 0.00156 | 0.2075055 | 0.21 | 0.171 | 1 | Synoviocytes | ppp6r3 |
| nop581 | 0.00156 | 0.2563321 | 0.329 | 0.288 | 1 | Synoviocytes | nop58 |
| pa2g4a1 | 0.00157 | 0.3189702 | 0.222 | 0.185 | 1 | Synoviocytes | pa2g4a |
| smarca22 | 0.00158 | 0.2716107 | 0.265 | 0.226 | 1 | Synoviocytes | smarca2 |
| atp5if1a1 | 0.0016 | 0.2200228 | 0.371 | 0.326 | 1 | Synoviocytes | atp5if1a |
| foxo1a2 | 0.00163 | 0.4095724 | 0.223 | 0.189 | 1 | Synoviocytes | foxo1a |
| api52 | 0.00171 | 0.3015772 | 0.172 | 0.139 | 1 | Synoviocytes | api5 |
| eloa2 | 0.00174 | 0.5298723 | 0.1 | 0.074 | 1 | Synoviocytes | eloa |
| gna12a1 | 0.00175 | 0.5183189 | 0.101 | 0.075 | 1 | Synoviocytes | gna12a |
| sf3a11 | 0.00176 | 0.3738621 | 0.122 | 0.093 | 1 | Synoviocytes | sf3a1 |
| vps293 | 0.00181 | 0.2380197 | 0.239 | 0.201 | 1 | Synoviocytes | vps29 |
| CABZ01044053.12 | 0.00182 | 0.352129 | 0.108 | 0.081 | 1 | Synoviocytes | CABZ01044053.1 |
| tgif15 | 0.00182 | 0.3403146 | 0.241 | 0.205 | 1 | Synoviocytes | tgif1 |
| snrnp702 | 0.00184 | 0.2542891 | 0.202 | 0.166 | 1 | Synoviocytes | snrnp70 |
| calr3a4 | 0.00186 | 0.2035759 | 0.403 | 0.362 | 1 | Synoviocytes | calr3a |
| krtcap24 | 0.00187 | 0.3385162 | 0.305 | 0.27 | 1 | Synoviocytes | krtcap2 |
| clns1a2 | 0.00187 | 0.3718762 | 0.103 | 0.077 | 1 | Synoviocytes | clns1a |
| upf1 | 0.00187 | 0.3222532 | 0.172 | 0.139 | 1 | Synoviocytes | upf1 |
| csrp1a4 | 0.0019 | 0.2406486 | 0.293 | 0.256 | 1 | Synoviocytes | csrp1a |
| emc7 | 0.00192 | 0.3120891 | 0.163 | 0.131 | 1 | Synoviocytes | emc7 |
| sept62 | 0.00198 | 0.390006 | 0.179 | 0.147 | 1 | Synoviocytes | sept6 |
| dip2ca4 | 0.00199 | 0.4018444 | 0.114 | 0.087 | 1 | Synoviocytes | dip2ca |
| hint14 | 0.00199 | 0.2053015 | 0.426 | 0.386 | 1 | Synoviocytes | hint1 |
| ctcf1 | 0.00201 | 0.400338 | 0.152 | 0.122 | 1 | Synoviocytes | ctcf |
| acadl2 | 0.00201 | 0.3851062 | 0.12 | 0.092 | 1 | Synoviocytes | acadl |
| ewsr1b1 | 0.00201 | 0.2957082 | 0.269 | 0.232 | 1 | Synoviocytes | ewsr1b |
| gtf2a22 | 0.00209 | 0.319197 | 0.141 | 0.11 | 1 | Synoviocytes | gtf2a2 |
| cica2 | 0.00211 | 0.2868141 | 0.223 | 0.186 | 1 | Synoviocytes | cica |
| ehd1b2 | 0.00212 | 0.3348783 | 0.21 | 0.175 | 1 | Synoviocytes | ehd1b |
| zswim53 | 0.00216 | 0.5243495 | 0.211 | 0.18 | 1 | Synoviocytes | zswim5 |
| ndufs23 | 0.00217 | 0.4591135 | 0.142 | 0.113 | 1 | Synoviocytes | ndufs2 |
| emc31 | 0.00221 | 0.4810975 | 0.104 | 0.079 | 1 | Synoviocytes | emc3 |
| puf60a1 | 0.00222 | 0.3997936 | 0.184 | 0.152 | 1 | Synoviocytes | puf60a |
| itgb26 | 0.00229 | 0.4016808 | 0.135 | 0.106 | 1 | Synoviocytes | itgb2 |
| cybrd14 | 0.00231 | 0.5085936 | 0.109 | 0.083 | 1 | Synoviocytes | cybrd1 |
| top2b2 | 0.00235 | 0.2113052 | 0.306 | 0.265 | 1 | Synoviocytes | top2b |
| wnk1b2 | 0.00238 | 0.4555255 | 0.157 | 0.127 | 1 | Synoviocytes | wnk1b |
| eif2a2 | 0.0024 | 0.3444328 | 0.109 | 0.083 | 1 | Synoviocytes | eif2a |
| snrpf1 | 0.00242 | 0.2802577 | 0.352 | 0.314 | 1 | Synoviocytes | snrpf |
| cops82 | 0.00243 | 0.4107043 | 0.115 | 0.088 | 1 | Synoviocytes | cops8 |
| rarga4 | 0.00246 | 0.236169 | 0.315 | 0.278 | 1 | Synoviocytes | rarga |
| chd32 | 0.00246 | 0.2756831 | 0.21 | 0.174 | 1 | Synoviocytes | chd3 |
| svila5 | 0.00252 | 0.3654478 | 0.181 | 0.148 | 1 | Synoviocytes | svila |
| rbm53 | 0.00256 | 0.2618205 | 0.222 | 0.186 | 1 | Synoviocytes | rbm5 |
| cdipt1 | 0.00256 | 0.3867459 | 0.144 | 0.114 | 1 | Synoviocytes | cdipt |

| | | | | | | | |
|---|---|---|---|---|---|---|---|
| asap34 | 0.00261 | 0.4069235 | 0.156 | 0.126 | 1 | Synoviocytes | asap3 |
| fryl2 | 0.00262 | 0.4207809 | 0.135 | 0.106 | 1 | Synoviocytes | fryl |
| ppp1caa2 | 0.00266 | 0.3509009 | 0.168 | 0.136 | 1 | Synoviocytes | ppp1caa |
| thrap3b1 | 0.00267 | 0.2065594 | 0.254 | 0.214 | 1 | Synoviocytes | thrap3b |
| mia32 | 0.00267 | 0.2742055 | 0.232 | 0.196 | 1 | Synoviocytes | mia3 |
| emc22 | 0.00268 | 0.2800012 | 0.238 | 0.203 | 1 | Synoviocytes | emc2 |
| golga11 | 0.00268 | 0.4085425 | 0.118 | 0.091 | 1 | Synoviocytes | golga1 |
| tmem1472 | 0.00271 | 0.2666932 | 0.185 | 0.152 | 1 | Synoviocytes | tmem147 |
| crebzf | 0.00272 | 0.3616058 | 0.162 | 0.13 | 1 | Synoviocytes | crebzf |
| cux1b1 | 0.00278 | 0.5091375 | 0.125 | 0.098 | 1 | Synoviocytes | cux1b |
| ctsd4 | 0.00282 | 0.3951415 | 0.187 | 0.154 | 1 | Synoviocytes | ctsd |
| dipk1c6 | 0.00282 | 0.445753 | 0.143 | 0.115 | 1 | Synoviocytes | dipk1c |
| sf3b31 | 0.00283 | 0.4292092 | 0.11 | 0.085 | 1 | Synoviocytes | sf3b3 |
| smco44 | 0.00283 | 0.4227124 | 0.17 | 0.141 | 1 | Synoviocytes | smco4 |
| u2surp4 | 0.00287 | 0.2230116 | 0.351 | 0.313 | 1 | Synoviocytes | u2surp |
| gspt1l2 | 0.00288 | 0.3337609 | 0.26 | 0.225 | 1 | Synoviocytes | gspt1l |
| lpgat1 | 0.00288 | 0.3558745 | 0.134 | 0.106 | 1 | Synoviocytes | lpgat1 |
| psmd92 | 0.00295 | 0.4425204 | 0.103 | 0.079 | 1 | Synoviocytes | psmd9 |
| tmem183 | 0.00296 | 0.5320396 | 0.113 | 0.088 | 1 | Synoviocytes | tmem18 |
| adam19a4 | 0.00299 | 0.6740322 | 0.108 | 0.084 | 1 | Synoviocytes | adam19a |
| f3b5 | 0.00309 | 0.2156662 | 0.42 | 0.38 | 1 | Synoviocytes | f3b |
| gnb21 | 0.00311 | 0.3729716 | 0.134 | 0.106 | 1 | Synoviocytes | gnb2 |
| tmem2143 | 0.00329 | 0.3683831 | 0.156 | 0.126 | 1 | Synoviocytes | tmem214 |
| sumo12 | 0.00336 | 0.3469226 | 0.192 | 0.161 | 1 | Synoviocytes | sumo1 |
| usp101 | 0.00339 | 0.255789 | 0.286 | 0.249 | 1 | Synoviocytes | usp10 |
| birc62 | 0.00341 | 0.4759637 | 0.141 | 0.114 | 1 | Synoviocytes | birc6 |
| mink11 | 0.00345 | 0.41696 | 0.129 | 0.102 | 1 | Synoviocytes | mink1 |
| prkar2aa1 | 0.00348 | 0.2642313 | 0.195 | 0.162 | 1 | Synoviocytes | prkar2aa |
| icam33 | 0.00352 | 0.2895305 | 0.108 | 0.083 | 1 | Synoviocytes | icam3 |
| cltcb1 | 0.00352 | 0.3938827 | 0.137 | 0.109 | 1 | Synoviocytes | cltcb |
| hnrnpk2 | 0.00353 | 0.3502621 | 0.181 | 0.15 | 1 | Synoviocytes | hnrnpk |
| larp71 | 0.00357 | 0.4073468 | 0.106 | 0.082 | 1 | Synoviocytes | larp7 |
| si:dkey-9i23.153 | 0.00361 | 0.3346968 | 0.184 | 0.152 | 1 | Synoviocytes | si:dkey-9i23.15 |
| tcf251 | 0.00362 | 0.2613612 | 0.155 | 0.125 | 1 | Synoviocytes | tcf25 |
| slc7a6os | 0.00367 | 0.4733569 | 0.103 | 0.079 | 1 | Synoviocytes | slc7a6os |
| ptges3a1 | 0.00369 | 0.334242 | 0.152 | 0.123 | 1 | Synoviocytes | ptges3a |
| klf134 | 0.00369 | 0.3158701 | 0.157 | 0.128 | 1 | Synoviocytes | klf13 |
| fbxw11b4 | 0.0039 | 0.2905977 | 0.193 | 0.159 | 1 | Synoviocytes | fbxw11b |
| rbm222 | 0.00392 | 0.3793773 | 0.13 | 0.104 | 1 | Synoviocytes | rbm22 |
| higd2a3 | 0.00398 | 0.3536403 | 0.164 | 0.136 | 1 | Synoviocytes | higd2a |
| upf3b2 | 0.00398 | 0.2972078 | 0.257 | 0.224 | 1 | Synoviocytes | upf3b |
| supt4h12 | 0.00401 | 0.3217884 | 0.165 | 0.135 | 1 | Synoviocytes | supt4h1 |
| inpp5l2 | 0.00409 | 0.3989379 | 0.124 | 0.098 | 1 | Synoviocytes | inpp5l |
| parp162 | 0.00414 | 0.4685309 | 0.113 | 0.088 | 1 | Synoviocytes | parp16 |
| gars13 | 0.00414 | 0.4516246 | 0.178 | 0.149 | 1 | Synoviocytes | gars1 |
| ptp4a2b3 | 0.00414 | 0.2445655 | 0.337 | 0.304 | 1 | Synoviocytes | ptp4a2b |
| per33 | 0.00415 | 0.2389644 | 0.385 | 0.347 | 1 | Synoviocytes | per3 |
| bsdc13 | 0.00416 | 0.3413251 | 0.121 | 0.095 | 1 | Synoviocytes | bsdc1 |

| | | | | | | | |
|---|---|---|---|---|---|---|---|
| tomm20a.12 | 0.00418 | 0.342062 | 0.192 | 0.16 | 1 | Synoviocytes | tomm20a.1 |
| ckap51 | 0.00423 | 0.4233107 | 0.113 | 0.088 | 1 | Synoviocytes | ckap5 |
| smad21 | 0.00426 | 0.2672999 | 0.225 | 0.191 | 1 | Synoviocytes | smad2 |
| ndufa24 | 0.00426 | 0.501009 | 0.119 | 0.094 | 1 | Synoviocytes | ndufa2 |
| crim12 | 0.00438 | 0.3134736 | 0.174 | 0.143 | 1 | Synoviocytes | crim1 |
| ndufa15 | 0.0044 | 0.2654453 | 0.239 | 0.205 | 1 | Synoviocytes | ndufa1 |
| safb1 | 0.00446 | 0.2403073 | 0.263 | 0.228 | 1 | Synoviocytes | safb |
| snrpa11 | 0.0045 | 0.3533786 | 0.125 | 0.098 | 1 | Synoviocytes | snrpa1 |
| zgc:112982 | 0.00459 | 0.2927309 | 0.166 | 0.136 | 1 | Synoviocytes | zgc:112982 |
| smarcb1a1 | 0.00464 | 0.4114945 | 0.138 | 0.111 | 1 | Synoviocytes | smarcb1a |
| hadhb3 | 0.00475 | 0.3161962 | 0.161 | 0.132 | 1 | Synoviocytes | hadhb |
| ebna1bp21 | 0.00475 | 0.4434364 | 0.17 | 0.142 | 1 | Synoviocytes | ebna1bp2 |
| ssrp1b1 | 0.00478 | 0.4445551 | 0.107 | 0.083 | 1 | Synoviocytes | ssrp1b |
| kpna32 | 0.00487 | 0.273573 | 0.183 | 0.152 | 1 | Synoviocytes | kpna3 |
| LEPROTL11 | 0.00488 | 0.3458243 | 0.139 | 0.112 | 1 | Synoviocytes | LEPROTL1 |
| CABZ01115113.12 | 0.00494 | 0.3541979 | 0.14 | 0.113 | 1 | Synoviocytes | CABZ01115113.1 |
| clint1a3 | 0.00494 | 0.2283856 | 0.242 | 0.208 | 1 | Synoviocytes | clint1a |
| wdr331 | 0.00506 | 0.5203955 | 0.113 | 0.089 | 1 | Synoviocytes | wdr33 |
| exoc61 | 0.00507 | 0.4140117 | 0.125 | 0.099 | 1 | Synoviocytes | exoc6 |
| cnot4b | 0.00508 | 0.4191185 | 0.108 | 0.084 | 1 | Synoviocytes | cnot4b |
| baz2ba4 | 0.00508 | 0.2914549 | 0.202 | 0.17 | 1 | Synoviocytes | baz2ba |
| ube2g1a2 | 0.00509 | 0.4205076 | 0.108 | 0.084 | 1 | Synoviocytes | ube2g1a |
| dstyk2 | 0.0051 | 0.413513 | 0.124 | 0.098 | 1 | Synoviocytes | dstyk |
| si:ch211-132b12.75 | 0.00511 | 0.2659641 | 0.188 | 0.156 | 1 | Synoviocytes | si:ch211-132b12.7 |
| arhgef1b3 | 0.00512 | 0.4547054 | 0.291 | 0.261 | 1 | Synoviocytes | arhgef1b |
| erlin11 | 0.0052 | 0.338768 | 0.119 | 0.093 | 1 | Synoviocytes | erlin1 |
| socs3b4 | 0.00527 | 0.2760033 | 0.502 | 0.471 | 1 | Synoviocytes | socs3b |
| med13a | 0.00534 | 0.5272991 | 0.122 | 0.097 | 1 | Synoviocytes | med13a |
| rbm4.23 | 0.00538 | 0.2880366 | 0.197 | 0.166 | 1 | Synoviocytes | rbm4.2 |
| golim4a2 | 0.00538 | 0.23913 | 0.147 | 0.118 | 1 | Synoviocytes | golim4a |
| ier3ip14 | 0.0054 | 0.2788457 | 0.269 | 0.236 | 1 | Synoviocytes | ier3ip1 |
| mbtps11 | 0.00549 | 0.593949 | 0.133 | 0.109 | 1 | Synoviocytes | mbtps1 |
| gga1 | 0.00551 | 0.4177856 | 0.118 | 0.093 | 1 | Synoviocytes | gga1 |
| cwc25 | 0.00555 | 0.3771673 | 0.162 | 0.134 | 1 | Synoviocytes | cwc25 |
| ch25hl34 | 0.00557 | 0.2576146 | 0.134 | 0.107 | 1 | Synoviocytes | ch25hl3 |
| acp13 | 0.0056 | 0.4773364 | 0.1 | 0.077 | 1 | Synoviocytes | acp1 |
| tnpo21 | 0.00566 | 0.4991165 | 0.11 | 0.087 | 1 | Synoviocytes | tnpo2 |
| wdr26b2 | 0.00567 | 0.4038153 | 0.117 | 0.092 | 1 | Synoviocytes | wdr26b |
| prelid1a4 | 0.00568 | 0.3952623 | 0.152 | 0.125 | 1 | Synoviocytes | prelid1a |
| trappc52 | 0.00577 | 0.4130891 | 0.14 | 0.114 | 1 | Synoviocytes | trappc5 |
| med101 | 0.00585 | 0.2679974 | 0.11 | 0.086 | 1 | Synoviocytes | med10 |
| pin43 | 0.00592 | 0.297298 | 0.178 | 0.148 | 1 | Synoviocytes | pin4 |
| bbip12 | 0.00601 | 0.4806848 | 0.151 | 0.126 | 1 | Synoviocytes | bbip1 |
| sod23 | 0.00607 | 0.2862862 | 0.157 | 0.129 | 1 | Synoviocytes | sod2 |
| lrp102 | 0.00607 | 0.3833173 | 0.12 | 0.095 | 1 | Synoviocytes | lrp10 |
| scyl12 | 0.0062 | 0.4295758 | 0.101 | 0.078 | 1 | Synoviocytes | scyl1 |
| lsm42 | 0.00636 | 0.3232219 | 0.246 | 0.217 | 1 | Synoviocytes | lsm4 |
| dock7 | 0.00672 | 0.2422141 | 0.265 | 0.232 | 1 | Synoviocytes | dock7 |

| | | | | | | | |
|---|---|---|---|---|---|---|---|
| sfrp55 | 0.00673 | 0.4414064 | 0.239 | 0.21 | 1 | Synoviocytes | sfrp5 |
| net15 | 0.00695 | 0.2416628 | 0.402 | 0.373 | 1 | Synoviocytes | net1 |
| si:ch211-196g2.42 | 0.00699 | 0.4720304 | 0.115 | 0.092 | 1 | Synoviocytes | si:ch211-196g2.4 |
| hypk2 | 0.00708 | 0.2198059 | 0.319 | 0.285 | 1 | Synoviocytes | hypk |
| polr2l2 | 0.00719 | 0.2440515 | 0.225 | 0.195 | 1 | Synoviocytes | polr2l |
| dhx161 | 0.00722 | 0.3364647 | 0.135 | 0.109 | 1 | Synoviocytes | dhx16 |
| gsk3ba2 | 0.00724 | 0.2814505 | 0.174 | 0.145 | 1 | Synoviocytes | gsk3ba |
| ddx232 | 0.0073 | 0.2693254 | 0.193 | 0.163 | 1 | Synoviocytes | ddx23 |
| rab18b2 | 0.00739 | 0.335594 | 0.142 | 0.115 | 1 | Synoviocytes | rab18b |
| xrn21 | 0.00747 | 0.2100432 | 0.257 | 0.223 | 1 | Synoviocytes | xrn2 |
| dcaf71 | 0.00753 | 0.3621214 | 0.156 | 0.13 | 1 | Synoviocytes | dcaf7 |
| fmnl31 | 0.00758 | 0.4519562 | 0.166 | 0.139 | 1 | Synoviocytes | fmnl3 |
| pdhb1 | 0.00765 | 0.3557189 | 0.135 | 0.11 | 1 | Synoviocytes | pdhb |
| nono1 | 0.00766 | 0.2665784 | 0.226 | 0.197 | 1 | Synoviocytes | nono |
| ndufs71 | 0.00774 | 0.2957759 | 0.208 | 0.178 | 1 | Synoviocytes | ndufs7 |
| sf11 | 0.00783 | 0.2881867 | 0.267 | 0.237 | 1 | Synoviocytes | sf1 |
| ankrd10b1 | 0.0079 | 0.4457276 | 0.123 | 0.099 | 1 | Synoviocytes | ankrd10b |
| furina4 | 0.00813 | 0.3207439 | 0.183 | 0.156 | 1 | Synoviocytes | furina |
| gcsha2 | 0.00813 | 0.2861607 | 0.126 | 0.102 | 1 | Synoviocytes | gcsha |
| zgc:1008295 | 0.00825 | 0.3955549 | 0.146 | 0.119 | 1 | Synoviocytes | zgc:100829 |
| bms11 | 0.00837 | 0.4770845 | 0.127 | 0.104 | 1 | Synoviocytes | bms1 |
| ndufb83 | 0.00845 | 0.2253025 | 0.369 | 0.337 | 1 | Synoviocytes | ndufb8 |
| med121 | 0.00858 | 0.4130057 | 0.111 | 0.088 | 1 | Synoviocytes | med12 |
| rcn22 | 0.00863 | 0.398292 | 0.102 | 0.081 | 1 | Synoviocytes | rcn2 |
| yaf23 | 0.00881 | 0.3399309 | 0.134 | 0.109 | 1 | Synoviocytes | yaf2 |
| tnfsf10l7 | 0.00884 | 0.3151795 | 0.251 | 0.222 | 1 | Synoviocytes | tnfsf10l |
| copa3 | 0.00889 | 0.3201236 | 0.218 | 0.19 | 1 | Synoviocytes | copa |
| nars11 | 0.00915 | 0.3164235 | 0.176 | 0.148 | 1 | Synoviocytes | nars1 |
| cdkn1bb3 | 0.00923 | 0.2806083 | 0.328 | 0.298 | 1 | Synoviocytes | cdkn1bb |
| C5H9orf782 | 0.00931 | 0.4192971 | 0.109 | 0.087 | 1 | Synoviocytes | C5H9orf78 |
| C2H1orf353 | 0.00957 | 0.5260431 | 0.107 | 0.086 | 1 | Synoviocytes | C2H1orf35 |
| lsm63 | 0.00993 | 0.3333064 | 0.199 | 0.172 | 1 | Synoviocytes | lsm6 |
| coch | 0 | 8.5418499 | 0.639 | 0.015 | 0 | Stroma 4 | coch |
| cxcl8a3 | 0 | 2.8905746 | 0.828 | 0.255 | 0 | Stroma 4 | cxcl8a |
| ndnfl1 | 0 | 4.1228364 | 0.363 | 0.029 | 0 | Stroma 4 | ndnfl |
| tnn1 | 0 | 3.1897846 | 0.272 | 0.028 | 0 | Stroma 4 | tnn |
| cd59 | 0 | 4.8502819 | 0.123 | 0.006 | 0 | Stroma 4 | cd59 |
| cygb14 | 9E-256 | 2.0745426 | 0.882 | 0.413 | 2.8E-251 | Stroma 4 | cygb1 |
| pcsk6 | 5E-219 | 4.2956907 | 0.167 | 0.017 | 1.7E-214 | Stroma 4 | pcsk6 |
| c1qtnf55 | 1E-216 | 2.0633701 | 0.525 | 0.145 | 3.8E-212 | Stroma 4 | c1qtnf5 |
| chad2 | 2E-213 | 2.9245326 | 0.283 | 0.048 | 8E-209 | Stroma 4 | chad |
| col5a3a2 | 1E-201 | 2.3025831 | 0.321 | 0.062 | 3.4E-197 | Stroma 4 | col5a3a |
| spock11 | 1E-191 | 2.8822446 | 0.232 | 0.036 | 3.2E-187 | Stroma 4 | spock1 |
| igf2b4 | 2E-187 | 1.8119102 | 0.834 | 0.429 | 6.1E-183 | Stroma 4 | igf2b |
| inhbaa5 | 1E-159 | 2.2209078 | 0.635 | 0.265 | 4.8E-155 | Stroma 4 | inhbaa |
| rpe65b3 | 6E-159 | 2.3243347 | 0.297 | 0.065 | 2.1E-154 | Stroma 4 | rpe65b |
| dhrs3b4 | 1E-155 | 2.8550958 | 0.387 | 0.112 | 4.6E-151 | Stroma 4 | dhrs3b |
| epha4l | 2E-142 | 3.4161025 | 0.111 | 0.011 | 6.2E-138 | Stroma 4 | epha4l |

| | | | | | | | |
|---|---|---|---|---|---|---|---|
| cald1b8 | 8E-139 | 2.2808217 | 0.466 | 0.162 | 2.5E-134 | Stroma 4 | cald1b |
| cyp26b13 | 1E-136 | 1.3189221 | 0.747 | 0.361 | 3.3E-132 | Stroma 4 | cyp26b1 |
| si:ch211-251b21.16 | 9E-136 | 1.4657271 | 0.818 | 0.444 | 2.9E-131 | Stroma 4 | si:ch211-251b21.1 |
| timp4.13 | 1E-135 | 1.7029795 | 0.341 | 0.091 | 4.2E-131 | Stroma 4 | timp4.1 |
| ncam1a3 | 8E-133 | 2.5157343 | 0.279 | 0.068 | 2.7E-128 | Stroma 4 | ncam1a |
| rgs5b6 | 5E-127 | 1.3424038 | 0.876 | 0.516 | 1.5E-122 | Stroma 4 | rgs5b |
| sema3fa3 | 8E-126 | 2.2872338 | 0.257 | 0.06 | 2.8E-121 | Stroma 4 | sema3fa |
| rpl134 | 4E-122 | 0.5535796 | 0.997 | 0.994 | 1.2E-117 | Stroma 4 | rpl13 |
| rps153 | 3E-110 | 0.4889512 | 0.998 | 0.989 | 1.1E-105 | Stroma 4 | rps15 |
| wnt7bb2 | 3E-109 | 2.5618053 | 0.188 | 0.038 | 8.9E-105 | Stroma 4 | wnt7bb |
| ednraa6 | 1E-106 | 1.6177964 | 0.504 | 0.208 | 4.6E-102 | Stroma 4 | ednraa |
| rplp2l3 | 2E-104 | 0.550545 | 0.994 | 0.987 | 6E-100 | Stroma 4 | rplp2l |
| osr2 | 9E-104 | 2.9036661 | 0.156 | 0.029 | 3.1E-99 | Stroma 4 | osr2 |
| nme2b.13 | 9E-103 | 0.7101342 | 0.974 | 0.918 | 2.79E-98 | Stroma 4 | nme2b.1 |
| clu6 | 2E-102 | 1.2823303 | 0.714 | 0.377 | 7.74E-98 | Stroma 4 | clu |
| rps27.14 | 2E-101 | 0.5080555 | 0.99 | 0.977 | 5.17E-97 | Stroma 4 | rps27.1 |
| mgp4 | 2E-99 | 0.8457619 | 0.408 | 0.155 | 5.56E-95 | Stroma 4 | mgp |
| cx43.45 | 2E-99 | 1.5187595 | 0.499 | 0.217 | 6.12E-95 | Stroma 4 | cx43.4 |
| tcima6 | 1.4E-98 | 1.5394976 | 0.51 | 0.221 | 4.69E-94 | Stroma 4 | tcima |
| ppiaa2 | 3.8E-98 | 0.5882091 | 0.982 | 0.96 | 1.23E-93 | Stroma 4 | ppiaa |
| lepa2 | 2.4E-97 | 2.060737 | 0.228 | 0.058 | 7.78E-93 | Stroma 4 | lepa |
| rps203 | 6.4E-96 | 0.5499412 | 0.99 | 0.986 | 2.07E-91 | Stroma 4 | rps20 |
| nr2f1a3 | 3.8E-95 | 1.9334178 | 0.391 | 0.148 | 1.23E-90 | Stroma 4 | nr2f1a |
| rpl93 | 2.1E-92 | 0.4769979 | 0.993 | 0.988 | 6.98E-88 | Stroma 4 | rpl9 |
| si:dkeyp-44a8.44 | 2.5E-92 | 1.753292 | 0.413 | 0.161 | 8.22E-88 | Stroma 4 | si:dkeyp-44a8.4 |
| rpl18a4 | 2.3E-91 | 0.4956529 | 0.99 | 0.984 | 7.32E-87 | Stroma 4 | rpl18a |
| flrt31 | 1.1E-89 | 2.2175659 | 0.212 | 0.054 | 3.42E-85 | Stroma 4 | flrt3 |
| mmp24 | 4E-89 | 0.8817564 | 0.938 | 0.741 | 1.29E-84 | Stroma 4 | mmp2 |
| rpl64 | 6.5E-89 | 0.55646 | 0.992 | 0.958 | 2.12E-84 | Stroma 4 | rpl6 |
| rpl143 | 1E-88 | 0.3785593 | 0.996 | 0.996 | 3.41E-84 | Stroma 4 | rpl14 |
| fgfr43 | 8.5E-88 | 2.0298829 | 0.251 | 0.073 | 2.75E-83 | Stroma 4 | fgfr4 |
| apcdd1l6 | 7.2E-87 | 1.7159026 | 0.316 | 0.107 | 2.35E-82 | Stroma 4 | apcdd1l |
| cxcl12b | 1.7E-86 | 0.823266 | 0.107 | 0.016 | 5.53E-82 | Stroma 4 | cxcl12b |
| sgk15 | 1.8E-85 | 1.2466875 | 0.651 | 0.362 | 5.92E-81 | Stroma 4 | sgk1 |
| ca10b | 1.9E-85 | 2.8261894 | 0.102 | 0.016 | 6.12E-81 | Stroma 4 | ca10b |
| rpsa4 | 1.1E-83 | 0.4541994 | 0.999 | 0.985 | 3.59E-79 | Stroma 4 | rpsa |
| rpl274 | 2.5E-81 | 0.4559226 | 0.996 | 0.978 | 8.04E-77 | Stroma 4 | rpl27 |
| rpl113 | 5.4E-80 | 0.4434361 | 0.992 | 0.989 | 1.75E-75 | Stroma 4 | rpl11 |
| rps193 | 7.4E-79 | 0.388747 | 0.997 | 0.996 | 2.41E-74 | Stroma 4 | rps19 |
| rpl343 | 1.1E-78 | 0.4409617 | 0.992 | 0.985 | 3.42E-74 | Stroma 4 | rpl34 |
| rps94 | 1.5E-78 | 0.4480736 | 0.994 | 0.987 | 4.96E-74 | Stroma 4 | rps9 |
| CR556696.1 | 6.8E-78 | 2.7383114 | 0.101 | 0.016 | 2.2E-73 | Stroma 4 | CR556696.1 |
| eef1a1l14 | 3.2E-77 | 0.4229182 | 1 | 0.996 | 1.04E-72 | Stroma 4 | eef1a1l1 |
| rplp13 | 1.1E-76 | 0.3767942 | 0.999 | 0.999 | 3.48E-72 | Stroma 4 | rplp1 |
| BX927333.12 | 1.2E-76 | 2.1857516 | 0.182 | 0.047 | 4E-72 | Stroma 4 | BX927333.1 |
| angptl46 | 5.4E-76 | 1.0430846 | 0.732 | 0.451 | 1.75E-71 | Stroma 4 | angptl4 |
| si:dkey-151g10.64 | 3.2E-75 | 0.3684721 | 1 | 0.999 | 1.04E-70 | Stroma 4 | si:dkey-151g10.6 |
| rps64 | 4E-75 | 0.5013105 | 0.987 | 0.96 | 1.3E-70 | Stroma 4 | rps6 |

| | | | | | | | |
|---|---|---|---|---|---|---|---|
| nfkbiaa5 | 1.4E-74 | 1.3191551 | 0.669 | 0.419 | 4.63E-70 | Stroma 4 | nfkbiaa |
| prdm1a4 | 2.9E-74 | 1.485537 | 0.242 | 0.074 | 9.32E-70 | Stroma 4 | prdm1a |
| rpl104 | 2.5E-73 | 0.4844301 | 0.985 | 0.968 | 8.01E-69 | Stroma 4 | rpl10 |
| si:ch211-197l9.2 | 2.1E-72 | 2.5378173 | 0.1 | 0.017 | 6.85E-68 | Stroma 4 | si:ch211-197l9.2 |
| fthl274 | 3E-72 | 0.8289665 | 0.942 | 0.838 | 9.91E-68 | Stroma 4 | fthl27 |
| cdh116 | 9.5E-72 | 0.8713355 | 0.852 | 0.585 | 3.09E-67 | Stroma 4 | cdh11 |
| rps8a4 | 1.7E-71 | 0.4182825 | 0.997 | 0.991 | 5.58E-67 | Stroma 4 | rps8a |
| rps15a4 | 2.1E-71 | 0.4077517 | 1 | 0.989 | 6.7E-67 | Stroma 4 | rps15a |
| rpl310 | 3.6E-71 | 0.4657687 | 0.991 | 0.977 | 1.18E-66 | Stroma 4 | rpl3 |
| rpl7a3 | 5.7E-71 | 0.4327337 | 0.996 | 0.991 | 1.85E-66 | Stroma 4 | rpl7a |
| rps4x4 | 6.1E-71 | 0.4963424 | 0.984 | 0.967 | 1.99E-66 | Stroma 4 | rps4x |
| rplp05 | 2.1E-70 | 0.4860319 | 0.993 | 0.982 | 6.75E-66 | Stroma 4 | rplp0 |
| rpl184 | 2.4E-70 | 0.4379819 | 0.991 | 0.982 | 7.84E-66 | Stroma 4 | rpl18 |
| rpl302 | 6.6E-70 | 0.3992892 | 0.991 | 0.983 | 2.15E-65 | Stroma 4 | rpl30 |
| col7a14 | 2.1E-69 | 1.0973026 | 0.373 | 0.151 | 6.75E-65 | Stroma 4 | col7a1 |
| rpl5a5 | 2.2E-69 | 0.5089887 | 0.963 | 0.918 | 7.29E-65 | Stroma 4 | rpl5a |
| rps34 | 2.3E-69 | 0.5046125 | 0.976 | 0.947 | 7.47E-65 | Stroma 4 | rps3 |
| rps103 | 1.6E-68 | 0.390011 | 0.993 | 0.994 | 5.33E-64 | Stroma 4 | rps10 |
| igfbp1a3 | 3.3E-67 | 2.3703173 | 0.217 | 0.069 | 1.08E-62 | Stroma 4 | igfbp1a |
| rpl353 | 6.1E-67 | 0.3778966 | 0.997 | 0.992 | 1.97E-62 | Stroma 4 | rpl35 |
| rps243 | 7.3E-67 | 0.363198 | 1 | 0.996 | 2.36E-62 | Stroma 4 | rps24 |
| rpl244 | 1.3E-66 | 0.3826255 | 0.999 | 0.993 | 4.39E-62 | Stroma 4 | rpl24 |
| rack14 | 2.6E-66 | 0.4784397 | 0.974 | 0.948 | 8.54E-62 | Stroma 4 | rack1 |
| uba523 | 1.1E-65 | 0.3607904 | 0.992 | 0.992 | 3.52E-61 | Stroma 4 | uba52 |
| rpl233 | 1.8E-65 | 0.3829859 | 0.99 | 0.992 | 5.93E-61 | Stroma 4 | rpl23 |
| rpl174 | 1.9E-65 | 0.408305 | 0.993 | 0.989 | 6.16E-61 | Stroma 4 | rpl17 |
| ctsla4 | 1.9E-65 | 0.8704707 | 0.865 | 0.719 | 6.32E-61 | Stroma 4 | ctsla |
| rps143 | 3.2E-65 | 0.3652607 | 0.994 | 0.993 | 1.04E-60 | Stroma 4 | rps14 |
| rps183 | 4.8E-65 | 0.4769874 | 0.983 | 0.963 | 1.56E-60 | Stroma 4 | rps18 |
| rps113 | 7.2E-65 | 0.3823974 | 0.993 | 0.989 | 2.35E-60 | Stroma 4 | rps11 |
| hsp70.34 | 2E-64 | 0.869956 | 0.809 | 0.549 | 6.44E-60 | Stroma 4 | hsp70.3 |
| rpl123 | 1.2E-62 | 0.4307563 | 0.988 | 0.982 | 3.79E-58 | Stroma 4 | rpl12 |
| rps133 | 1.3E-62 | 0.357581 | 0.997 | 0.99 | 4.08E-58 | Stroma 4 | rps13 |
| rpl154 | 1.3E-62 | 0.4662655 | 0.991 | 0.97 | 4.37E-58 | Stroma 4 | rpl15 |
| rbp53 | 2E-62 | 1.4407345 | 0.298 | 0.115 | 6.54E-58 | Stroma 4 | rbp5 |
| rps173 | 3.6E-62 | 0.4533731 | 0.971 | 0.945 | 1.16E-57 | Stroma 4 | rps17 |
| rpl23a4 | 7.1E-62 | 0.4329062 | 0.983 | 0.971 | 2.32E-57 | Stroma 4 | rpl23a |
| rpl323 | 2.6E-61 | 0.3963 | 0.994 | 0.989 | 8.59E-57 | Stroma 4 | rpl32 |
| nid1a3 | 1.2E-60 | 1.2346017 | 0.46 | 0.231 | 3.88E-56 | Stroma 4 | nid1a |
| plpp37 | 1.9E-60 | 0.8848222 | 0.714 | 0.45 | 6.29E-56 | Stroma 4 | plpp3 |
| rps3a3 | 2.5E-60 | 0.4323956 | 0.988 | 0.981 | 8.27E-56 | Stroma 4 | rps3a |
| rps253 | 5.8E-60 | 0.3287342 | 1 | 0.994 | 1.88E-55 | Stroma 4 | rps25 |
| faua3 | 9.4E-60 | 0.4056175 | 0.984 | 0.974 | 3.07E-55 | Stroma 4 | faua |
| f3b6 | 1.3E-59 | 1.0761091 | 0.619 | 0.376 | 4.2E-55 | Stroma 4 | f3b |
| rps123 | 2.9E-59 | 0.4396959 | 0.989 | 0.978 | 9.31E-55 | Stroma 4 | rps12 |
| rps27a2 | 4E-59 | 0.3659976 | 0.997 | 0.992 | 1.28E-54 | Stroma 4 | rps27a |
| spry45 | 4.9E-59 | 1.0786829 | 0.387 | 0.175 | 1.61E-54 | Stroma 4 | spry4 |
| mmp93 | 1.1E-58 | 1.2771144 | 0.216 | 0.072 | 3.58E-54 | Stroma 4 | mmp9 |

| | | | | | | | |
|---|---|---|---|---|---|---|---|
| enpp63 | 1.3E-58 | 1.6747176 | 0.183 | 0.055 | 4.07E-54 | Stroma 4 | enpp6 |
| rpl22l14 | 1.8E-58 | 0.7408809 | 0.865 | 0.746 | 5.87E-54 | Stroma 4 | rpl22l1 |
| CR855311.13 | 1.6E-57 | 0.8343952 | 0.35 | 0.15 | 5.18E-53 | Stroma 4 | CR855311.1 |
| hsp90ab11 | 6.7E-57 | 0.3645493 | 0.997 | 0.993 | 2.17E-52 | Stroma 4 | hsp90ab1 |
| RPS173 | 1.6E-56 | 0.3606595 | 0.997 | 0.987 | 5.05E-52 | Stroma 4 | RPS17 |
| rpl284 | 2.8E-56 | 0.3530904 | 0.996 | 0.991 | 9.02E-52 | Stroma 4 | rpl28 |
| rpl10a3 | 3.3E-56 | 0.4264481 | 0.988 | 0.978 | 1.08E-51 | Stroma 4 | rpl10a |
| dll42 | 1.1E-55 | 1.8956473 | 0.148 | 0.04 | 3.52E-51 | Stroma 4 | dll4 |
| rpl313 | 1.1E-55 | 0.3704394 | 0.997 | 0.993 | 3.56E-51 | Stroma 4 | rpl31 |
| slc1a3a5 | 1.2E-55 | 1.4288829 | 0.333 | 0.147 | 3.89E-51 | Stroma 4 | slc1a3a |
| nav32 | 6.7E-55 | 1.3419974 | 0.229 | 0.08 | 2.17E-50 | Stroma 4 | nav3 |
| hic12 | 3.3E-54 | 1.8154495 | 0.191 | 0.062 | 1.07E-49 | Stroma 4 | hic1 |
| si:ch211-222l21.14 | 1.3E-53 | 0.4282817 | 0.809 | 0.55 | 4.2E-49 | Stroma 4 | si:ch211-222l21.1 |
| rps74 | 1.5E-53 | 0.3748587 | 0.998 | 0.991 | 4.76E-49 | Stroma 4 | rps7 |
| rps26l4 | 1.7E-53 | 0.3734345 | 0.982 | 0.974 | 5.45E-49 | Stroma 4 | rps26l |
| rps55 | 1.9E-53 | 0.4070375 | 0.988 | 0.972 | 6.23E-49 | Stroma 4 | rps5 |
| mdka4 | 2.5E-53 | 0.8670659 | 0.734 | 0.518 | 8.08E-49 | Stroma 4 | mdka |
| hyal44 | 1.2E-52 | 1.1978185 | 0.266 | 0.103 | 3.88E-48 | Stroma 4 | hyal4 |
| adamts54 | 1.8E-52 | 1.5449841 | 0.192 | 0.063 | 5.75E-48 | Stroma 4 | adamts5 |
| rpl44 | 2.2E-52 | 0.4567317 | 0.955 | 0.895 | 7.11E-48 | Stroma 4 | rpl4 |
| C25H11orf967 | 3.5E-52 | 0.8822484 | 0.629 | 0.379 | 1.13E-47 | Stroma 4 | C25H11orf96 |
| eef1g3 | 2.6E-51 | 0.4471415 | 0.979 | 0.952 | 8.54E-47 | Stroma 4 | eef1g |
| efemp2a4 | 7.1E-51 | 0.88507 | 0.594 | 0.358 | 2.32E-46 | Stroma 4 | efemp2a |
| rps233 | 2E-50 | 0.3526372 | 0.985 | 0.982 | 6.36E-46 | Stroma 4 | rps23 |
| plk35 | 2.2E-50 | 1.2134705 | 0.401 | 0.199 | 7.3E-46 | Stroma 4 | plk3 |
| quo1 | 3.4E-50 | 1.4707697 | 0.194 | 0.066 | 1.1E-45 | Stroma 4 | quo |
| si:ch211-198c19.33 | 5.9E-50 | 1.0544624 | 0.409 | 0.208 | 1.91E-45 | Stroma 4 | si:ch211-198c19.3 |
| id2b5 | 6.7E-50 | 1.2523041 | 0.379 | 0.187 | 2.17E-45 | Stroma 4 | id2b |
| pim15 | 1.2E-49 | 0.6588715 | 0.765 | 0.541 | 4.03E-45 | Stroma 4 | pim1 |
| rpl36a4 | 1.3E-49 | 0.3048929 | 0.997 | 0.994 | 4.17E-45 | Stroma 4 | rpl36a |
| rpl84 | 1.6E-49 | 0.3950558 | 0.99 | 0.978 | 5.12E-45 | Stroma 4 | rpl8 |
| furina5 | 7E-49 | 1.4253308 | 0.328 | 0.153 | 2.28E-44 | Stroma 4 | furina |
| cebpb5 | 3.1E-48 | 0.8895183 | 0.677 | 0.445 | 9.97E-44 | Stroma 4 | cebpb |
| lgals9l33 | 6.5E-48 | 1.2713378 | 0.283 | 0.12 | 2.12E-43 | Stroma 4 | lgals9l3 |
| phlda25 | 6.2E-47 | 0.8663447 | 0.418 | 0.216 | 2.02E-42 | Stroma 4 | phlda2 |
| adam152 | 9.4E-47 | 1.4251949 | 0.167 | 0.054 | 3.07E-42 | Stroma 4 | adam15 |
| sdc44 | 1.4E-46 | 0.6194615 | 0.865 | 0.701 | 4.42E-42 | Stroma 4 | sdc4 |
| msx1b4 | 2.3E-46 | 0.7875865 | 0.52 | 0.29 | 7.57E-42 | Stroma 4 | msx1b |
| hsp70.24 | 7.2E-46 | 0.614809 | 0.738 | 0.489 | 2.33E-41 | Stroma 4 | hsp70.2 |
| slc25a55 | 7.5E-46 | 0.397358 | 0.982 | 0.962 | 2.43E-41 | Stroma 4 | slc25a5 |
| rgcc5 | 6.3E-45 | 0.8085126 | 0.39 | 0.195 | 2.04E-40 | Stroma 4 | rgcc |
| tmem1767 | 1.8E-44 | 0.8065377 | 0.611 | 0.39 | 6.02E-40 | Stroma 4 | tmem176 |
| si:dkey-34d22.11 | 2E-44 | 1.8178164 | 0.127 | 0.036 | 6.54E-40 | Stroma 4 | si:dkey-34d22.1 |
| fhl2a3 | 3E-44 | 1.1747393 | 0.294 | 0.132 | 9.69E-40 | Stroma 4 | fhl2a |
| trabd2b | 3.2E-44 | 1.6742838 | 0.12 | 0.033 | 1.03E-39 | Stroma 4 | trabd2b |
| id35 | 6.2E-44 | 0.7372038 | 0.716 | 0.508 | 2.02E-39 | Stroma 4 | id3 |
| eef1b24 | 1.2E-43 | 0.4258728 | 0.974 | 0.928 | 3.74E-39 | Stroma 4 | eef1b2 |
| htra3a1 | 1.3E-43 | 1.4032906 | 0.145 | 0.045 | 4.28E-39 | Stroma 4 | htra3a |

| | | | | | | | |
|---|---|---|---|---|---|---|---|
| ENSDARG000001152 | 1.5E-43 | 0.3046499 | 0.998 | 0.993 | 4.73E-39 | Stroma 4 | ENSDARG00000115 |
| rpl363 | 1.5E-43 | 0.3078682 | 0.994 | 0.985 | 4.76E-39 | Stroma 4 | rpl36 |
| rps163 | 2.2E-43 | 0.3164213 | 0.988 | 0.989 | 7.26E-39 | Stroma 4 | rps16 |
| rpl212 | 2.3E-43 | 0.3342056 | 0.99 | 0.985 | 7.46E-39 | Stroma 4 | rpl21 |
| arhgap362 | 4.7E-43 | 1.4466115 | 0.176 | 0.061 | 1.51E-38 | Stroma 4 | arhgap36 |
| gstm.31 | 7.3E-43 | 1.6749709 | 0.151 | 0.049 | 2.36E-38 | Stroma 4 | gstm.3 |
| id2a7 | 1.1E-42 | 0.7809609 | 0.705 | 0.5 | 3.73E-38 | Stroma 4 | id2a |
| rpl35a4 | 2E-42 | 0.2892417 | 0.99 | 0.989 | 6.52E-38 | Stroma 4 | rpl35a |
| dusp26 | 2E-42 | 1.1980028 | 0.407 | 0.219 | 6.57E-38 | Stroma 4 | dusp2 |
| pdgfra6 | 2.2E-42 | 0.7446702 | 0.672 | 0.457 | 7.03E-38 | Stroma 4 | pdgfra |
| phex3 | 2.4E-42 | 1.2943179 | 0.201 | 0.076 | 7.81E-38 | Stroma 4 | phex |
| cyp1c14 | 4.1E-41 | 1.5742546 | 0.235 | 0.101 | 1.32E-36 | Stroma 4 | cyp1c1 |
| zfand5b3 | 9.8E-41 | 1.0034087 | 0.459 | 0.269 | 3.18E-36 | Stroma 4 | zfand5b |
| sat1a.29 | 1.7E-40 | 0.7917716 | 0.649 | 0.454 | 5.53E-36 | Stroma 4 | sat1a.2 |
| rpl73 | 2E-40 | 0.3684569 | 0.98 | 0.973 | 6.54E-36 | Stroma 4 | rpl7 |
| rpl13a3 | 6.4E-40 | 0.3392107 | 0.996 | 0.985 | 2.07E-35 | Stroma 4 | rpl13a |
| rpl393 | 9.5E-40 | 0.3070826 | 0.993 | 0.984 | 3.09E-35 | Stroma 4 | rpl39 |
| wu:fj16a036 | 1.1E-39 | 1.0614415 | 0.363 | 0.189 | 3.6E-35 | Stroma 4 | wu:fj16a03 |
| tpt13 | 2.9E-39 | 0.3914719 | 0.975 | 0.926 | 9.43E-35 | Stroma 4 | tpt1 |
| zfhx44 | 6.2E-39 | 0.8338372 | 0.66 | 0.446 | 2.03E-34 | Stroma 4 | zfhx4 |
| rplp23 | 6.8E-39 | 0.302917 | 0.994 | 0.987 | 2.2E-34 | Stroma 4 | rplp2 |
| npm1a2 | 2.3E-38 | 0.7542937 | 0.559 | 0.357 | 7.61E-34 | Stroma 4 | npm1a |
| marcksl1b6 | 3.3E-38 | 0.8505734 | 0.445 | 0.251 | 1.07E-33 | Stroma 4 | marcksl1b |
| gstt1a3 | 1.2E-37 | 1.3531587 | 0.189 | 0.074 | 3.85E-33 | Stroma 4 | gstt1a |
| ppfibp2b1 | 1.8E-37 | 2.1793695 | 0.11 | 0.032 | 5.82E-33 | Stroma 4 | ppfibp2b |
| tnc7 | 3.3E-37 | 0.4145864 | 0.713 | 0.466 | 1.07E-32 | Stroma 4 | tnc |
| tspan364 | 4.1E-37 | 1.102502 | 0.316 | 0.159 | 1.33E-32 | Stroma 4 | tspan36 |
| rpl223 | 5.8E-37 | 0.3327846 | 0.985 | 0.973 | 1.89E-32 | Stroma 4 | rpl22 |
| zgc:1748885 | 8.7E-37 | 0.99151 | 0.389 | 0.214 | 2.82E-32 | Stroma 4 | zgc:174888 |
| serpine25 | 1.8E-36 | 1.1421312 | 0.484 | 0.314 | 5.7E-32 | Stroma 4 | serpine2 |
| cxxc5b | 2.6E-36 | 1.8874497 | 0.1 | 0.028 | 8.34E-32 | Stroma 4 | cxxc5b |
| ch25hl35 | 3.4E-36 | 1.0233169 | 0.24 | 0.105 | 1.12E-31 | Stroma 4 | ch25hl3 |
| eif14 | 4.7E-36 | 0.4315266 | 0.914 | 0.828 | 1.53E-31 | Stroma 4 | eif1 |
| ankha1 | 9.6E-36 | 1.8043849 | 0.115 | 0.036 | 3.11E-31 | Stroma 4 | ankha |
| si:dkey-12l12.13 | 9.7E-36 | 0.3700428 | 0.786 | 0.576 | 3.16E-31 | Stroma 4 | si:dkey-12l12.1 |
| fbln1 | 3.1E-35 | 1.7277606 | 0.109 | 0.032 | 1E-30 | Stroma 4 | fbln1 |
| fosl1a4 | 3.6E-35 | 0.713355 | 0.643 | 0.434 | 1.17E-30 | Stroma 4 | fosl1a |
| ms4a17a.74 | 4.7E-35 | 0.7927786 | 0.482 | 0.294 | 1.53E-30 | Stroma 4 | ms4a17a.7 |
| rpl5b3 | 4.8E-35 | 0.4087932 | 0.917 | 0.849 | 1.57E-30 | Stroma 4 | rpl5b |
| her65 | 8.2E-35 | 0.5959418 | 0.64 | 0.429 | 2.67E-30 | Stroma 4 | her6 |
| arl5c4 | 8.9E-35 | 1.1188008 | 0.256 | 0.119 | 2.9E-30 | Stroma 4 | arl5c |
| cd634 | 1.1E-34 | 0.6177922 | 0.889 | 0.826 | 3.69E-30 | Stroma 4 | cd63 |
| znf710a | 1.3E-34 | 1.7150631 | 0.115 | 0.036 | 4.08E-30 | Stroma 4 | znf710a |
| dnmbp1 | 3E-34 | 1.2182641 | 0.226 | 0.1 | 9.84E-30 | Stroma 4 | dnmbp |
| hsp90aa1.24 | 5.8E-34 | 0.6161294 | 0.765 | 0.597 | 1.89E-29 | Stroma 4 | hsp90aa1.2 |
| rps282 | 6E-34 | 0.3070002 | 0.994 | 0.978 | 1.96E-29 | Stroma 4 | rps28 |
| adamtsl71 | 7E-34 | 0.9510896 | 0.289 | 0.143 | 2.26E-29 | Stroma 4 | adamtsl7 |
| hbegfa5 | 1.6E-33 | 0.8515423 | 0.58 | 0.399 | 5.3E-29 | Stroma 4 | hbegfa |

| | | | | | | | |
|---|---|---|---|---|---|---|---|
| hspa83 | 1.9E-33 | 0.3773136 | 0.964 | 0.936 | 6.2E-29 | Stroma 4 | hspa8 |
| ivns1abpa5 | 2E-33 | 0.9351835 | 0.459 | 0.284 | 6.53E-29 | Stroma 4 | ivns1abpa |
| en1b5 | 3.4E-33 | 1.2783337 | 0.234 | 0.108 | 1.1E-28 | Stroma 4 | en1b |
| pfkfb35 | 3.8E-33 | 1.4532568 | 0.182 | 0.075 | 1.24E-28 | Stroma 4 | pfkfb3 |
| im:71523486 | 7.1E-33 | 0.9470461 | 0.404 | 0.233 | 2.32E-28 | Stroma 4 | im:7152348 |
| smad73 | 8.7E-33 | 0.9732774 | 0.298 | 0.151 | 2.81E-28 | Stroma 4 | smad7 |
| mych5 | 1.2E-32 | 0.815804 | 0.559 | 0.377 | 3.8E-28 | Stroma 4 | mych |
| hapln1b2 | 3E-32 | 0.5820587 | 0.203 | 0.085 | 9.86E-28 | Stroma 4 | hapln1b |
| zfp36l1b7 | 3.4E-32 | 0.574621 | 0.722 | 0.543 | 1.11E-27 | Stroma 4 | zfp36l1b |
| rps263 | 4.4E-32 | 0.3027456 | 0.972 | 0.969 | 1.44E-27 | Stroma 4 | rps26 |
| rps210 | 5.3E-32 | 0.2983072 | 0.991 | 0.991 | 1.74E-27 | Stroma 4 | rps2 |
| ets23 | 9.2E-32 | 1.0741152 | 0.304 | 0.158 | 3E-27 | Stroma 4 | ets2 |
| si:ch211-160j14.34 | 9.5E-32 | 0.6879243 | 0.504 | 0.311 | 3.08E-27 | Stroma 4 | si:ch211-160j14.3 |
| ptmab5 | 1.4E-31 | 0.5075752 | 0.862 | 0.713 | 4.46E-27 | Stroma 4 | ptmab |
| rpl193 | 2.1E-31 | 0.3330895 | 0.989 | 0.983 | 6.67E-27 | Stroma 4 | rpl19 |
| irx5a4 | 5.6E-31 | 1.2955148 | 0.202 | 0.089 | 1.83E-26 | Stroma 4 | irx5a |
| clcn5b | 8.7E-31 | 1.4758221 | 0.139 | 0.051 | 2.82E-26 | Stroma 4 | clcn5b |
| ip6k2b5 | 1.5E-30 | 0.7506202 | 0.524 | 0.341 | 5E-26 | Stroma 4 | ip6k2b |
| inhbb1 | 2E-30 | 1.2847634 | 0.146 | 0.055 | 6.66E-26 | Stroma 4 | inhbb |
| pprc14 | 2.7E-30 | 1.156424 | 0.239 | 0.115 | 8.86E-26 | Stroma 4 | pprc1 |
| bzw1b4 | 7.2E-29 | 0.6634091 | 0.638 | 0.489 | 2.34E-24 | Stroma 4 | bzw1b |
| si:dkey-184p18.21 | 9.4E-29 | 1.6649047 | 0.145 | 0.057 | 3.07E-24 | Stroma 4 | si:dkey-184p18.2 |
| abca1a3 | 1.6E-28 | 0.8212058 | 0.35 | 0.195 | 5.21E-24 | Stroma 4 | abca1a |
| ptp4a2b4 | 1.8E-28 | 0.7637665 | 0.469 | 0.302 | 5.84E-24 | Stroma 4 | ptp4a2b |
| UBB4 | 2E-28 | 0.5311555 | 0.731 | 0.559 | 6.46E-24 | Stroma 4 | UBB |
| rsrp15 | 2.2E-28 | 0.5690814 | 0.706 | 0.539 | 7.31E-24 | Stroma 4 | rsrp1 |
| spon2a | 8E-28 | 1.6702292 | 0.1 | 0.033 | 2.62E-23 | Stroma 4 | spon2a |
| dusp57 | 9.9E-28 | 0.8921379 | 0.339 | 0.196 | 3.22E-23 | Stroma 4 | dusp5 |
| igflr11 | 1.3E-27 | 1.5591571 | 0.115 | 0.041 | 4.36E-23 | Stroma 4 | igflr1 |
| fstl35 | 1.8E-27 | 0.6231689 | 0.497 | 0.319 | 5.81E-23 | Stroma 4 | fstl3 |
| usp54a2 | 2.5E-27 | 1.4732348 | 0.142 | 0.057 | 8.09E-23 | Stroma 4 | usp54a |
| hsp70l4 | 3E-27 | 0.6633816 | 0.466 | 0.3 | 9.69E-23 | Stroma 4 | hsp70l |
| hif1ab4 | 5.2E-27 | 0.6837504 | 0.543 | 0.372 | 1.68E-22 | Stroma 4 | hif1ab |
| zgc:922424 | 5.5E-27 | 0.9177247 | 0.226 | 0.109 | 1.78E-22 | Stroma 4 | zgc:92242 |
| oaz1a6 | 1.2E-26 | 0.6548936 | 0.605 | 0.45 | 3.86E-22 | Stroma 4 | oaz1a |
| ldb2a1 | 1.5E-26 | 1.1382836 | 0.152 | 0.062 | 4.82E-22 | Stroma 4 | ldb2a |
| rhpn23 | 4E-26 | 0.8689009 | 0.27 | 0.141 | 1.3E-21 | Stroma 4 | rhpn2 |
| grwd14 | 6.1E-26 | 1.0922054 | 0.263 | 0.14 | 1.97E-21 | Stroma 4 | grwd1 |
| hmox1a6 | 8.1E-26 | 0.8385461 | 0.308 | 0.172 | 2.64E-21 | Stroma 4 | hmox1a |
| rps293 | 1.2E-25 | 0.2758974 | 0.976 | 0.97 | 3.81E-21 | Stroma 4 | rps29 |
| arhgef7a2 | 1.4E-25 | 1.366978 | 0.112 | 0.041 | 4.41E-21 | Stroma 4 | arhgef7a |
| snai1b4 | 2.1E-25 | 0.827962 | 0.226 | 0.111 | 6.8E-21 | Stroma 4 | snai1b |
| kng13 | 2.2E-25 | 0.9531287 | 0.177 | 0.079 | 7E-21 | Stroma 4 | kng1 |
| ahcy4 | 3.3E-25 | 0.7763057 | 0.322 | 0.183 | 1.08E-20 | Stroma 4 | ahcy |
| eef1db5 | 6.6E-25 | 0.45601 | 0.721 | 0.579 | 2.14E-20 | Stroma 4 | eef1db |
| fabp36 | 7E-25 | 0.5279887 | 0.685 | 0.528 | 2.28E-20 | Stroma 4 | fabp3 |
| prdx63 | 8.1E-25 | 0.6492568 | 0.501 | 0.342 | 2.62E-20 | Stroma 4 | prdx6 |
| rpl37.12 | 1.3E-24 | 0.2141347 | 0.994 | 0.991 | 4.11E-20 | Stroma 4 | rpl37.1 |

| | | | | | | | |
|---|---|---|---|---|---|---|---|
| ugdh6 | 1.3E-24 | 0.9117303 | 0.304 | 0.176 | 4.24E-20 | Stroma 4 | ugdh |
| pmaip13 | 1.4E-24 | 1.0854668 | 0.195 | 0.093 | 4.71E-20 | Stroma 4 | pmaip1 |
| letm24 | 2.9E-24 | 0.7630715 | 0.325 | 0.186 | 9.35E-20 | Stroma 4 | letm2 |
| ier2a5 | 3.2E-24 | 0.5824652 | 0.664 | 0.483 | 1.05E-19 | Stroma 4 | ier2a |
| cd82a5 | 3.3E-24 | 0.6975838 | 0.516 | 0.353 | 1.08E-19 | Stroma 4 | cd82a |
| sqstm15 | 3.5E-24 | 0.7851338 | 0.451 | 0.303 | 1.13E-19 | Stroma 4 | sqstm1 |
| ubb5 | 3.5E-24 | 0.4513754 | 0.82 | 0.679 | 1.14E-19 | Stroma 4 | ubb |
| prelid3b5 | 5.2E-24 | 0.6892005 | 0.507 | 0.355 | 1.7E-19 | Stroma 4 | prelid3b |
| timp2a6 | 5.8E-24 | 0.6481837 | 0.859 | 0.754 | 1.88E-19 | Stroma 4 | timp2a |
| gfpt22 | 6.3E-24 | 0.8714513 | 0.156 | 0.067 | 2.04E-19 | Stroma 4 | gfpt2 |
| slc25a55a6 | 9.4E-24 | 0.7175163 | 0.335 | 0.195 | 3.07E-19 | Stroma 4 | slc25a55a |
| her95 | 9.6E-24 | 0.4159214 | 0.372 | 0.226 | 3.14E-19 | Stroma 4 | her9 |
| polr2a6 | 9.9E-24 | 0.6439351 | 0.525 | 0.361 | 3.2E-19 | Stroma 4 | polr2a |
| selenon5 | 1.1E-23 | 0.9668276 | 0.278 | 0.155 | 3.68E-19 | Stroma 4 | selenon |
| adam8a1 | 1.3E-23 | 0.644807 | 0.254 | 0.134 | 4.23E-19 | Stroma 4 | adam8a |
| dio3b2 | 1.7E-23 | 0.961627 | 0.133 | 0.054 | 5.66E-19 | Stroma 4 | dio3b |
| thbs4b6 | 1.8E-23 | 0.2972396 | 0.419 | 0.257 | 5.82E-19 | Stroma 4 | thbs4b |
| ppiab4 | 3.8E-23 | 0.2989699 | 0.955 | 0.895 | 1.23E-18 | Stroma 4 | ppiab |
| brd2a5 | 3.8E-23 | 0.6127019 | 0.592 | 0.445 | 1.23E-18 | Stroma 4 | brd2a |
| atp1b1a3 | 4E-23 | 0.3762148 | 0.645 | 0.482 | 1.31E-18 | Stroma 4 | atp1b1a |
| neo1b3 | 4.2E-23 | 1.0078129 | 0.173 | 0.08 | 1.36E-18 | Stroma 4 | neo1b |
| prdx12 | 4.6E-23 | 1.0913356 | 0.15 | 0.066 | 1.48E-18 | Stroma 4 | prdx1 |
| marcksl1a5 | 6.5E-23 | 0.2559887 | 0.853 | 0.705 | 2.1E-18 | Stroma 4 | marcksl1a |
| zgc:921613 | 8.9E-23 | 0.6903995 | 0.325 | 0.192 | 2.89E-18 | Stroma 4 | zgc:92161 |
| amotl2a2 | 1.2E-22 | 0.9344265 | 0.275 | 0.155 | 3.82E-18 | Stroma 4 | amotl2a |
| cited4b5 | 1.3E-22 | 0.9013413 | 0.267 | 0.148 | 4.22E-18 | Stroma 4 | cited4b |
| naca4 | 1.5E-22 | 0.2769505 | 0.97 | 0.961 | 4.8E-18 | Stroma 4 | naca |
| rps211 | 2E-22 | 0.2115464 | 0.992 | 0.978 | 6.35E-18 | Stroma 4 | rps21 |
| ssh2a3 | 2.2E-22 | 1.1271338 | 0.169 | 0.079 | 7.18E-18 | Stroma 4 | ssh2a |
| si:dkey-195m11.86 | 3.2E-22 | 0.766895 | 0.239 | 0.126 | 1.04E-17 | Stroma 4 | si:dkey-195m11.8 |
| ccn16 | 4.5E-22 | 0.4892307 | 0.775 | 0.64 | 1.47E-17 | Stroma 4 | ccn1 |
| ppp1r106 | 1E-21 | 0.4584107 | 0.647 | 0.482 | 3.36E-17 | Stroma 4 | ppp1r10 |
| suco6 | 1.1E-21 | 0.6419137 | 0.299 | 0.171 | 3.51E-17 | Stroma 4 | suco |
| bach1b3 | 1.4E-21 | 1.0276878 | 0.186 | 0.091 | 4.49E-17 | Stroma 4 | bach1b |
| nr4a33 | 1.5E-21 | 0.697525 | 0.122 | 0.05 | 4.98E-17 | Stroma 4 | nr4a3 |
| BX908782.22 | 1.8E-21 | 1.4593272 | 0.13 | 0.056 | 5.74E-17 | Stroma 4 | BX908782.2 |
| egr2a4 | 3.6E-21 | 1.0382742 | 0.188 | 0.094 | 1.17E-16 | Stroma 4 | egr2a |
| zgc:1749043 | 3.9E-21 | 0.7240164 | 0.232 | 0.123 | 1.27E-16 | Stroma 4 | zgc:174904 |
| crema4 | 4.8E-21 | 0.9954441 | 0.209 | 0.108 | 1.56E-16 | Stroma 4 | crema |
| tgif16 | 5.4E-21 | 0.7406982 | 0.332 | 0.204 | 1.75E-16 | Stroma 4 | tgif1 |
| mxra5a4 | 6.6E-21 | 0.9297309 | 0.163 | 0.076 | 2.16E-16 | Stroma 4 | mxra5a |
| sox4a5 | 7.1E-21 | 0.6261804 | 0.322 | 0.192 | 2.33E-16 | Stroma 4 | sox4a |
| twist27 | 8.4E-21 | 0.517713 | 0.512 | 0.338 | 2.73E-16 | Stroma 4 | twist2 |
| tbx2b3 | 1.3E-20 | 0.891502 | 0.231 | 0.125 | 4.2E-16 | Stroma 4 | tbx2b |
| eva1a2 | 1.6E-20 | 0.8482499 | 0.131 | 0.056 | 5.12E-16 | Stroma 4 | eva1a |
| ms4a17a.85 | 2E-20 | 0.8099354 | 0.214 | 0.112 | 6.66E-16 | Stroma 4 | ms4a17a.8 |
| pdia33 | 2.7E-20 | 0.3853564 | 0.786 | 0.685 | 8.87E-16 | Stroma 4 | pdia3 |
| tenm36 | 2.7E-20 | 0.6687745 | 0.334 | 0.205 | 8.88E-16 | Stroma 4 | tenm3 |

| | | | | | | | |
|---|---|---|---|---|---|---|---|
| prdx21 | 2.8E-20 | 0.5934825 | 0.478 | 0.328 | 9.06E-16 | Stroma 4 | prdx2 |
| pcolcea6 | 2.8E-20 | 0.4339807 | 0.732 | 0.566 | 9.09E-16 | Stroma 4 | pcolcea |
| stk17al2 | 3.5E-20 | 1.1263532 | 0.159 | 0.075 | 1.13E-15 | Stroma 4 | stk17al |
| lifrb3 | 3.9E-20 | 0.6381072 | 0.324 | 0.197 | 1.27E-15 | Stroma 4 | lifrb |
| camk1b2 | 9.4E-20 | 1.2165265 | 0.13 | 0.057 | 3.05E-15 | Stroma 4 | camk1b |
| eef2b6 | 9.6E-20 | 0.2749169 | 0.95 | 0.909 | 3.12E-15 | Stroma 4 | eef2b |
| homeza | 1.1E-19 | 0.9597612 | 0.158 | 0.075 | 3.7E-15 | Stroma 4 | homeza |
| maff7 | 1.2E-19 | 0.6229741 | 0.357 | 0.222 | 3.93E-15 | Stroma 4 | maff |
| s1pr13 | 1.3E-19 | 0.8160373 | 0.183 | 0.091 | 4.23E-15 | Stroma 4 | s1pr1 |
| atoh82 | 1.4E-19 | 1.0544802 | 0.137 | 0.061 | 4.63E-15 | Stroma 4 | atoh8 |
| fhl3a3 | 2.2E-19 | 1.1662588 | 0.145 | 0.067 | 7E-15 | Stroma 4 | fhl3a |
| igfbp6b1 | 2.2E-19 | 1.467968 | 0.131 | 0.059 | 7.02E-15 | Stroma 4 | igfbp6b |
| hspd11 | 2.5E-19 | 0.7051105 | 0.34 | 0.212 | 8.09E-15 | Stroma 4 | hspd1 |
| ldb2b | 2.9E-19 | 0.9373376 | 0.149 | 0.069 | 9.44E-15 | Stroma 4 | ldb2b |
| gadd45ab6 | 3E-19 | 0.7056748 | 0.293 | 0.174 | 9.9E-15 | Stroma 4 | gadd45ab |
| efhd2 | 3.7E-19 | 1.1672206 | 0.107 | 0.043 | 1.22E-14 | Stroma 4 | efhd2 |
| ubc5 | 4.4E-19 | 0.3981602 | 0.785 | 0.661 | 1.42E-14 | Stroma 4 | ubc |
| SBNO25 | 4.9E-19 | 0.5525374 | 0.36 | 0.226 | 1.58E-14 | Stroma 4 | SBNO2 |
| zgc:771583 | 8.9E-19 | 1.2559816 | 0.132 | 0.061 | 2.89E-14 | Stroma 4 | zgc:77158 |
| zswim54 | 1E-18 | 0.5590522 | 0.303 | 0.178 | 3.3E-14 | Stroma 4 | zswim5 |
| c7a5 | 1.5E-18 | 0.8388716 | 0.284 | 0.173 | 4.98E-14 | Stroma 4 | c7a |
| mt-cyb3 | 1.6E-18 | 0.2881275 | 0.981 | 0.975 | 5.08E-14 | Stroma 4 | mt-cyb |
| acsl4a6 | 1.9E-18 | 0.8746763 | 0.158 | 0.077 | 6.26E-14 | Stroma 4 | acsl4a |
| pgam1a4 | 2.1E-18 | 0.4909848 | 0.311 | 0.185 | 6.9E-14 | Stroma 4 | pgam1a |
| cycsb4 | 2.2E-18 | 0.4565073 | 0.516 | 0.372 | 7.04E-14 | Stroma 4 | cycsb |
| ptges4 | 2.3E-18 | 0.8484352 | 0.201 | 0.107 | 7.59E-14 | Stroma 4 | ptges |
| si:dkey-192g7.32 | 2.6E-18 | 0.7857025 | 0.126 | 0.055 | 8.59E-14 | Stroma 4 | si:dkey-192g7.3 |
| trib34 | 4.1E-18 | 0.7251044 | 0.289 | 0.174 | 1.35E-13 | Stroma 4 | trib3 |
| mknk2a4 | 7.1E-18 | 0.6572998 | 0.352 | 0.227 | 2.31E-13 | Stroma 4 | mknk2a |
| slc25a25a5 | 7.6E-18 | 0.6250829 | 0.345 | 0.217 | 2.49E-13 | Stroma 4 | slc25a25a |
| eif51 | 7.9E-18 | 0.4933181 | 0.614 | 0.478 | 2.58E-13 | Stroma 4 | eif5 |
| hsp70.14 | 9.4E-18 | 0.3251248 | 0.488 | 0.339 | 3.07E-13 | Stroma 4 | hsp70.1 |
| lamc34 | 1.1E-17 | 0.8672823 | 0.216 | 0.119 | 3.53E-13 | Stroma 4 | lamc3 |
| plxdc24 | 1.5E-17 | 0.4936685 | 0.271 | 0.156 | 4.77E-13 | Stroma 4 | plxdc2 |
| ldha5 | 1.6E-17 | 0.4765701 | 0.358 | 0.228 | 5.32E-13 | Stroma 4 | ldha |
| rrad5 | 2.2E-17 | 0.9281468 | 0.143 | 0.069 | 7.18E-13 | Stroma 4 | rrad |
| mvp6 | 2.3E-17 | 0.4861399 | 0.45 | 0.315 | 7.45E-13 | Stroma 4 | mvp |
| brdt5 | 2.5E-17 | 0.5946308 | 0.445 | 0.309 | 8.27E-13 | Stroma 4 | brdt |
| tuba8l6 | 2.7E-17 | 0.4863023 | 0.546 | 0.408 | 8.7E-13 | Stroma 4 | tuba8l |
| dlx5a2 | 3.2E-17 | 0.6118563 | 0.432 | 0.297 | 1.05E-12 | Stroma 4 | dlx5a |
| rsl24d16 | 3.8E-17 | 0.381397 | 0.83 | 0.747 | 1.24E-12 | Stroma 4 | rsl24d1 |
| yap12 | 4E-17 | 0.5111984 | 0.504 | 0.357 | 1.29E-12 | Stroma 4 | yap1 |
| si:dkey-7j14.63 | 4.4E-17 | 0.3334031 | 0.85 | 0.776 | 1.44E-12 | Stroma 4 | si:dkey-7j14.6 |
| TXN3 | 4.9E-17 | 0.332427 | 0.837 | 0.746 | 1.58E-12 | Stroma 4 | TXN |
| vmp14 | 5.3E-17 | 0.4226584 | 0.451 | 0.317 | 1.71E-12 | Stroma 4 | vmp1 |
| crabp2b6 | 5.9E-17 | 0.4974684 | 0.558 | 0.418 | 1.92E-12 | Stroma 4 | crabp2b |
| midn5 | 5.9E-17 | 0.4733336 | 0.573 | 0.432 | 1.93E-12 | Stroma 4 | midn |
| scfd15 | 6.7E-17 | 1.0654096 | 0.189 | 0.104 | 2.19E-12 | Stroma 4 | scfd1 |

| | | | | | | | |
|---|---|---|---|---|---|---|---|
| rbbp64 | 7E-17 | 0.5251819 | 0.451 | 0.315 | 2.28E-12 | Stroma 4 | rbbp6 |
| nocta4 | 7.4E-17 | 0.6518196 | 0.205 | 0.112 | 2.39E-12 | Stroma 4 | nocta |
| mcl1a6 | 7.7E-17 | 0.3224464 | 0.872 | 0.769 | 2.49E-12 | Stroma 4 | mcl1a |
| dnajb1b5 | 9.5E-17 | 0.253283 | 0.618 | 0.462 | 3.08E-12 | Stroma 4 | dnajb1b |
| ssr35 | 9.5E-17 | 0.3752978 | 0.704 | 0.576 | 3.09E-12 | Stroma 4 | ssr3 |
| dhrs3a4 | 9.7E-17 | 1.2388353 | 0.138 | 0.067 | 3.14E-12 | Stroma 4 | dhrs3a |
| inka1b4 | 9.7E-17 | 0.8669392 | 0.174 | 0.091 | 3.16E-12 | Stroma 4 | inka1b |
| cited4a3 | 1.5E-16 | 0.9012579 | 0.205 | 0.114 | 4.77E-12 | Stroma 4 | cited4a |
| gpx1a4 | 1.5E-16 | 0.7825768 | 0.302 | 0.191 | 4.86E-12 | Stroma 4 | gpx1a |
| bag34 | 1.5E-16 | 0.9724265 | 0.143 | 0.071 | 4.89E-12 | Stroma 4 | bag3 |
| tubb2b4 | 1.6E-16 | 0.4160403 | 0.485 | 0.339 | 5.15E-12 | Stroma 4 | tubb2b |
| si:dkey-79d12.54 | 1.8E-16 | 0.7149907 | 0.225 | 0.128 | 5.81E-12 | Stroma 4 | si:dkey-79d12.5 |
| tent5c3 | 1.8E-16 | 0.9704049 | 0.161 | 0.083 | 5.87E-12 | Stroma 4 | tent5c |
| DAB26 | 2.4E-16 | 0.4252792 | 0.506 | 0.376 | 7.75E-12 | Stroma 4 | DAB2 |
| hspa55 | 2.5E-16 | 0.4333384 | 0.627 | 0.49 | 8.07E-12 | Stroma 4 | hspa5 |
| gadd45bb6 | 2.6E-16 | 0.5704872 | 0.335 | 0.219 | 8.58E-12 | Stroma 4 | gadd45bb |
| cfp7 | 3.5E-16 | 0.3483343 | 0.497 | 0.338 | 1.13E-11 | Stroma 4 | cfp |
| cct2.13 | 4.5E-16 | 0.4398275 | 0.553 | 0.422 | 1.46E-11 | Stroma 4 | cct2.1 |
| eif4a1a6 | 5.4E-16 | 0.4476657 | 0.506 | 0.364 | 1.75E-11 | Stroma 4 | eif4a1a |
| rspo33 | 5.6E-16 | 0.5542802 | 0.133 | 0.064 | 1.82E-11 | Stroma 4 | rspo3 |
| eif2s1b4 | 6.1E-16 | 0.4507651 | 0.521 | 0.385 | 1.98E-11 | Stroma 4 | eif2s1b |
| irf23 | 6.4E-16 | 0.7149855 | 0.248 | 0.148 | 2.09E-11 | Stroma 4 | irf2 |
| pcxa2 | 6.8E-16 | 0.6917118 | 0.206 | 0.115 | 2.21E-11 | Stroma 4 | pcxa |
| slc3a2b4 | 8.8E-16 | 0.6292268 | 0.235 | 0.137 | 2.88E-11 | Stroma 4 | slc3a2b |
| ano5b4 | 1.4E-15 | 0.8567825 | 0.164 | 0.086 | 4.6E-11 | Stroma 4 | ano5b |
| dbi4 | 1.5E-15 | 0.4551229 | 0.57 | 0.44 | 4.94E-11 | Stroma 4 | dbi |
| kctd12.23 | 1.7E-15 | 0.7437376 | 0.333 | 0.225 | 5.6E-11 | Stroma 4 | kctd12.2 |
| prmt13 | 1.9E-15 | 0.6286965 | 0.296 | 0.187 | 6.12E-11 | Stroma 4 | prmt1 |
| ak24 | 1.9E-15 | 0.5046423 | 0.506 | 0.378 | 6.34E-11 | Stroma 4 | ak2 |
| cox6b23 | 3.1E-15 | 0.2614945 | 0.869 | 0.814 | 1.02E-10 | Stroma 4 | cox6b2 |
| svild3 | 3.2E-15 | 0.5697268 | 0.379 | 0.254 | 1.04E-10 | Stroma 4 | svild |
| arg27 | 3.9E-15 | 0.5524558 | 0.263 | 0.163 | 1.28E-10 | Stroma 4 | arg2 |
| dipk1c7 | 4.5E-15 | 0.6376773 | 0.201 | 0.114 | 1.46E-10 | Stroma 4 | dipk1c |
| cpxm1a3 | 5.3E-15 | 0.3866168 | 0.287 | 0.177 | 1.73E-10 | Stroma 4 | cpxm1a |
| si:zfos-943e10.11 | 5.5E-15 | 0.8504608 | 0.146 | 0.075 | 1.79E-10 | Stroma 4 | si:zfos-943e10.1 |
| zfand5a7 | 5.9E-15 | 0.5260284 | 0.297 | 0.187 | 1.91E-10 | Stroma 4 | zfand5a |
| nfkbiab5 | 6.3E-15 | 0.3549305 | 0.703 | 0.582 | 2.06E-10 | Stroma 4 | nfkbiab |
| atf35 | 8.1E-15 | 0.2063526 | 0.482 | 0.344 | 2.65E-10 | Stroma 4 | atf3 |
| eif3i3 | 1.2E-14 | 0.4519264 | 0.473 | 0.343 | 3.98E-10 | Stroma 4 | eif3i |
| rpl292 | 1.3E-14 | 0.2010161 | 0.966 | 0.931 | 4.13E-10 | Stroma 4 | rpl29 |
| sec61b4 | 1.4E-14 | 0.3319724 | 0.701 | 0.581 | 4.7E-10 | Stroma 4 | sec61b |
| egr14 | 1.7E-14 | 0.2692699 | 0.779 | 0.626 | 5.52E-10 | Stroma 4 | egr1 |
| fap2 | 1.7E-14 | 0.9184477 | 0.13 | 0.065 | 5.57E-10 | Stroma 4 | fap |
| eif3k1 | 1.8E-14 | 0.3782106 | 0.612 | 0.472 | 5.8E-10 | Stroma 4 | eif3k |
| tmem176l.14 | 1.9E-14 | 0.29211 | 0.831 | 0.74 | 6.04E-10 | Stroma 4 | tmem176l.1 |
| eif3ea5 | 2.3E-14 | 0.3576426 | 0.673 | 0.558 | 7.49E-10 | Stroma 4 | eif3ea |
| oaz2a2 | 2.6E-14 | 0.6705496 | 0.148 | 0.077 | 8.59E-10 | Stroma 4 | oaz2a |
| si:ch211-286o17.15 | 2.7E-14 | 0.5105007 | 0.328 | 0.215 | 8.87E-10 | Stroma 4 | si:ch211-286o17.1 |

| | | | | | | | |
|---|---|---|---|---|---|---|---|
| nedd94 | 3.1E-14 | 0.8331403 | 0.14 | 0.072 | 1.02E-09 | Stroma 4 | nedd9 |
| serp15 | 3.4E-14 | 0.4889482 | 0.438 | 0.314 | 1.12E-09 | Stroma 4 | serp1 |
| fscn1a6 | 3.5E-14 | 0.5885675 | 0.279 | 0.177 | 1.12E-09 | Stroma 4 | fscn1a |
| glulb5 | 3.7E-14 | 0.2243145 | 0.719 | 0.567 | 1.2E-09 | Stroma 4 | glulb |
| ptges3b3 | 4.8E-14 | 0.4687175 | 0.237 | 0.141 | 1.58E-09 | Stroma 4 | ptges3b |
| si:ch211-15b10.62 | 7.5E-14 | 0.6398103 | 0.178 | 0.099 | 2.45E-09 | Stroma 4 | si:ch211-15b10.6 |
| rpl383 | 7.7E-14 | 0.2124309 | 0.937 | 0.91 | 2.49E-09 | Stroma 4 | rpl38 |
| rasl11b6 | 7.8E-14 | 0.7614591 | 0.317 | 0.215 | 2.54E-09 | Stroma 4 | rasl11b |
| lrrc174 | 1.8E-13 | 0.3103518 | 0.536 | 0.394 | 5.82E-09 | Stroma 4 | lrrc17 |
| lygl12 | 1.9E-13 | 0.5006164 | 0.261 | 0.164 | 6.17E-09 | Stroma 4 | lygl1 |
| ponzr15 | 2E-13 | 0.4144324 | 0.348 | 0.232 | 6.52E-09 | Stroma 4 | ponzr1 |
| ube2al4 | 2E-13 | 0.6086351 | 0.3 | 0.198 | 6.53E-09 | Stroma 4 | ube2al |
| CABZ01061591.1 | 2.1E-13 | 0.9462433 | 0.137 | 0.072 | 6.85E-09 | Stroma 4 | CABZ01061591.1 |
| zgc:1107964 | 2.3E-13 | 0.9073766 | 0.148 | 0.08 | 7.61E-09 | Stroma 4 | zgc:110796 |
| lfng4 | 2.8E-13 | 0.4365678 | 0.371 | 0.252 | 9.15E-09 | Stroma 4 | lfng |
| aebp15 | 3.5E-13 | 0.2930964 | 0.664 | 0.538 | 1.15E-08 | Stroma 4 | aebp1 |
| adam8b4 | 3.9E-13 | 0.7459824 | 0.147 | 0.079 | 1.28E-08 | Stroma 4 | adam8b |
| pcdh192 | 4.2E-13 | 0.8187565 | 0.109 | 0.052 | 1.35E-08 | Stroma 4 | pcdh19 |
| htra1b5 | 4.2E-13 | 0.2681419 | 0.53 | 0.41 | 1.36E-08 | Stroma 4 | htra1b |
| rbms1a2 | 4.3E-13 | 0.8461755 | 0.141 | 0.075 | 1.4E-08 | Stroma 4 | rbms1a |
| rarga5 | 4.9E-13 | 0.636611 | 0.387 | 0.276 | 1.58E-08 | Stroma 4 | rarga |
| cct6a2 | 5E-13 | 0.3599678 | 0.599 | 0.478 | 1.63E-08 | Stroma 4 | cct6a |
| eif3ha4 | 5.1E-13 | 0.4113045 | 0.535 | 0.42 | 1.66E-08 | Stroma 4 | eif3ha |
| socs1a6 | 5.1E-13 | 0.4427998 | 0.207 | 0.123 | 1.67E-08 | Stroma 4 | socs1a |
| si:ch73-86n18.15 | 5.2E-13 | 0.5299753 | 0.243 | 0.153 | 1.71E-08 | Stroma 4 | si:ch73-86n18.1 |
| irf1b6 | 6.3E-13 | 0.5575265 | 0.197 | 0.117 | 2.04E-08 | Stroma 4 | irf1b |
| bmp1a5 | 6.3E-13 | 0.5729738 | 0.285 | 0.186 | 2.04E-08 | Stroma 4 | bmp1a |
| ism13 | 6.3E-13 | 0.5361053 | 0.209 | 0.124 | 2.05E-08 | Stroma 4 | ism1 |
| rgl11 | 1.4E-12 | 0.5345668 | 0.207 | 0.124 | 4.53E-08 | Stroma 4 | rgl1 |
| bambia3 | 1.4E-12 | 0.5676112 | 0.234 | 0.147 | 4.59E-08 | Stroma 4 | bambia |
| eif3m2 | 1.5E-12 | 0.2804107 | 0.735 | 0.625 | 4.98E-08 | Stroma 4 | eif3m |
| egfl65 | 1.5E-12 | 0.9022869 | 0.11 | 0.054 | 5.02E-08 | Stroma 4 | egfl6 |
| btf35 | 1.6E-12 | 0.2737904 | 0.898 | 0.875 | 5.2E-08 | Stroma 4 | btf3 |
| enosf1 | 1.8E-12 | 0.6737484 | 0.102 | 0.049 | 5.84E-08 | Stroma 4 | enosf1 |
| fgfr25 | 2.2E-12 | 0.4179842 | 0.466 | 0.347 | 7.24E-08 | Stroma 4 | fgfr2 |
| tgfbr2a | 2.4E-12 | 0.8976156 | 0.105 | 0.052 | 7.94E-08 | Stroma 4 | tgfbr2a |
| pde4ba4 | 2.5E-12 | 0.5722229 | 0.196 | 0.116 | 8.22E-08 | Stroma 4 | pde4ba |
| nop532 | 2.8E-12 | 0.3345068 | 0.584 | 0.456 | 8.99E-08 | Stroma 4 | nop53 |
| fhl3b4 | 2.8E-12 | 0.5820935 | 0.187 | 0.111 | 9.2E-08 | Stroma 4 | fhl3b |
| slc25a485 | 2.9E-12 | 0.5735131 | 0.222 | 0.137 | 9.35E-08 | Stroma 4 | slc25a48 |
| ptenb4 | 2.9E-12 | 0.6492118 | 0.265 | 0.174 | 9.46E-08 | Stroma 4 | ptenb |
| eif5a4 | 3.3E-12 | 0.2588998 | 0.812 | 0.719 | 1.09E-07 | Stroma 4 | eif5a |
| hspa4a6 | 4.4E-12 | 0.5272752 | 0.237 | 0.148 | 1.42E-07 | Stroma 4 | hspa4a |
| cldni2 | 4.5E-12 | 0.7689263 | 0.119 | 0.061 | 1.47E-07 | Stroma 4 | cldni |
| h3f3b.15 | 4.7E-12 | 0.3956607 | 0.602 | 0.494 | 1.54E-07 | Stroma 4 | h3f3b.1 |
| dab2ipb3 | 5E-12 | 0.6195611 | 0.188 | 0.112 | 1.63E-07 | Stroma 4 | dab2ipb |
| abcf2a6 | 5.1E-12 | 0.4420384 | 0.274 | 0.177 | 1.66E-07 | Stroma 4 | abcf2a |
| tshz15 | 5.2E-12 | 0.5756223 | 0.22 | 0.135 | 1.7E-07 | Stroma 4 | tshz1 |

| | | | | | | | |
|---|---|---|---|---|---|---|---|
| hnrnpa0b1 | 5.7E-12 | 0.3647949 | 0.515 | 0.401 | 1.87E-07 | Stroma 4 | hnrnpa0b |
| spen3 | 6.7E-12 | 0.5791636 | 0.294 | 0.196 | 2.19E-07 | Stroma 4 | spen |
| oaz1b7 | 8E-12 | 0.4279174 | 0.414 | 0.306 | 2.62E-07 | Stroma 4 | oaz1b |
| ddx510 | 8.3E-12 | 0.2938594 | 0.704 | 0.596 | 2.71E-07 | Stroma 4 | ddx5 |
| fblim14 | 9.2E-12 | 0.5520822 | 0.186 | 0.11 | 3E-07 | Stroma 4 | fblim1 |
| plxnb2a.13 | 9.5E-12 | 0.7043535 | 0.157 | 0.089 | 3.1E-07 | Stroma 4 | plxnb2a.1 |
| mt-co22 | 1E-11 | 0.2124137 | 1 | 0.998 | 3.41E-07 | Stroma 4 | mt-co2 |
| psmg22 | 1.1E-11 | 0.6767996 | 0.178 | 0.105 | 3.52E-07 | Stroma 4 | psmg2 |
| ctnnb26 | 1.2E-11 | 0.4826662 | 0.411 | 0.311 | 3.89E-07 | Stroma 4 | ctnnb2 |
| rps27.22 | 1.2E-11 | 0.2043816 | 0.96 | 0.953 | 4.01E-07 | Stroma 4 | rps27.2 |
| slc38a25 | 1.3E-11 | 0.6622651 | 0.142 | 0.079 | 4.09E-07 | Stroma 4 | slc38a2 |
| fosl25 | 1.3E-11 | 0.3432838 | 0.552 | 0.427 | 4.38E-07 | Stroma 4 | fosl2 |
| pcm15 | 1.5E-11 | 0.6093993 | 0.233 | 0.152 | 4.85E-07 | Stroma 4 | pcm1 |
| sema4ba4 | 1.6E-11 | 0.6754532 | 0.149 | 0.084 | 5.23E-07 | Stroma 4 | sema4ba |
| steap43 | 1.7E-11 | 0.418812 | 0.275 | 0.183 | 5.47E-07 | Stroma 4 | steap4 |
| ncl4 | 2E-11 | 0.3508257 | 0.675 | 0.604 | 6.63E-07 | Stroma 4 | ncl |
| myo10l34 | 2.1E-11 | 0.6331389 | 0.169 | 0.099 | 6.69E-07 | Stroma 4 | myo10l3 |
| klf135 | 2.2E-11 | 0.6563142 | 0.205 | 0.127 | 7.26E-07 | Stroma 4 | klf13 |
| adipor25 | 3.1E-11 | 0.5325965 | 0.269 | 0.176 | 1.01E-06 | Stroma 4 | adipor2 |
| nabp1a4 | 3.1E-11 | 0.5546243 | 0.267 | 0.179 | 1.02E-06 | Stroma 4 | nabp1a |
| sox65 | 3.2E-11 | 0.4039479 | 0.174 | 0.102 | 1.05E-06 | Stroma 4 | sox6 |
| si:dkey-23i12.73 | 3.3E-11 | 0.5677338 | 0.221 | 0.14 | 1.08E-06 | Stroma 4 | si:dkey-23i12.7 |
| eif3f4 | 3.4E-11 | 0.2499712 | 0.804 | 0.728 | 1.12E-06 | Stroma 4 | eif3f |
| si:ch1073-392o20.25 | 3.5E-11 | 0.4558377 | 0.345 | 0.241 | 1.13E-06 | Stroma 4 | si:ch1073-392o20.2 |
| sik16 | 3.6E-11 | 0.2951444 | 0.363 | 0.252 | 1.16E-06 | Stroma 4 | sik1 |
| bcl6ab2 | 3.7E-11 | 0.7479637 | 0.115 | 0.06 | 1.19E-06 | Stroma 4 | bcl6ab |
| eif2s33 | 4.1E-11 | 0.2885506 | 0.615 | 0.494 | 1.34E-06 | Stroma 4 | eif2s3 |
| col14a1a3 | 4.6E-11 | 0.5069139 | 0.133 | 0.073 | 1.51E-06 | Stroma 4 | col14a1a |
| si:dkey-248g15.32 | 5.2E-11 | 0.3862135 | 0.295 | 0.199 | 1.71E-06 | Stroma 4 | si:dkey-248g15.3 |
| snu13b2 | 5.3E-11 | 0.5420748 | 0.267 | 0.18 | 1.71E-06 | Stroma 4 | snu13b |
| hmgb3a4 | 6.4E-11 | 0.5555482 | 0.188 | 0.115 | 2.09E-06 | Stroma 4 | hmgb3a |
| atf4a5 | 6.8E-11 | 0.4552514 | 0.4 | 0.297 | 2.23E-06 | Stroma 4 | atf4a |
| ifrd16 | 7.2E-11 | 0.432219 | 0.39 | 0.281 | 2.35E-06 | Stroma 4 | ifrd1 |
| scarb2a3 | 8.7E-11 | 0.3618386 | 0.284 | 0.187 | 2.83E-06 | Stroma 4 | scarb2a |
| cad3 | 9.1E-11 | 0.6388363 | 0.141 | 0.08 | 2.97E-06 | Stroma 4 | cad |
| tbx2a2 | 9.2E-11 | 0.5673131 | 0.115 | 0.061 | 3E-06 | Stroma 4 | tbx2a |
| ap2m1a3 | 9.6E-11 | 0.3876866 | 0.41 | 0.304 | 3.11E-06 | Stroma 4 | ap2m1a |
| nrarpa2 | 9.8E-11 | 0.6762733 | 0.151 | 0.088 | 3.18E-06 | Stroma 4 | nrarpa |
| zc3h44 | 9.9E-11 | 0.6587415 | 0.215 | 0.138 | 3.21E-06 | Stroma 4 | zc3h4 |
| hspe11 | 1.1E-10 | 0.3056398 | 0.399 | 0.293 | 3.5E-06 | Stroma 4 | hspe1 |
| bahcc1b | 1.1E-10 | 0.5065267 | 0.195 | 0.12 | 3.63E-06 | Stroma 4 | bahcc1b |
| relch2 | 1.2E-10 | 0.7844073 | 0.126 | 0.069 | 3.91E-06 | Stroma 4 | relch |
| tspan4b5 | 1.3E-10 | 0.2857323 | 0.461 | 0.335 | 4.39E-06 | Stroma 4 | tspan4b |
| parp43 | 1.5E-10 | 0.5570719 | 0.146 | 0.083 | 4.81E-06 | Stroma 4 | parp4 |
| tiparp4 | 1.7E-10 | 0.5973606 | 0.198 | 0.124 | 5.48E-06 | Stroma 4 | tiparp |
| si:ch211-195b13.15 | 1.9E-10 | 0.2870078 | 0.344 | 0.242 | 6.03E-06 | Stroma 4 | si:ch211-195b13.1 |
| uqcrh4 | 2.1E-10 | 0.296205 | 0.492 | 0.383 | 6.77E-06 | Stroma 4 | uqcrh |
| jak15 | 2.2E-10 | 0.3075467 | 0.546 | 0.436 | 7.01E-06 | Stroma 4 | jak1 |

| | | | | | | | |
|---|---|---|---|---|---|---|---|
| mafba6 | 2.3E-10 | 0.2854258 | 0.508 | 0.389 | 7.37E-06 | Stroma 4 | mafba |
| b3gnt5a5 | 2.5E-10 | 0.5366367 | 0.182 | 0.111 | 8.04E-06 | Stroma 4 | b3gnt5a |
| nap1l11 | 2.7E-10 | 0.2715852 | 0.63 | 0.525 | 8.79E-06 | Stroma 4 | nap1l1 |
| tspan355 | 2.9E-10 | 0.265502 | 0.201 | 0.123 | 9.36E-06 | Stroma 4 | tspan35 |
| cxcl18b5 | 3.3E-10 | 0.265048 | 0.79 | 0.701 | 1.08E-05 | Stroma 4 | cxcl18b |
| ephb2b5 | 3.5E-10 | 0.4274218 | 0.29 | 0.198 | 1.13E-05 | Stroma 4 | ephb2b |
| ptgs2a5 | 3.6E-10 | 0.4765407 | 0.204 | 0.129 | 1.17E-05 | Stroma 4 | ptgs2a |
| moxd13 | 3.7E-10 | 0.6509003 | 0.101 | 0.052 | 1.19E-05 | Stroma 4 | moxd1 |
| kras4 | 4E-10 | 0.4387453 | 0.263 | 0.177 | 1.3E-05 | Stroma 4 | kras |
| zgc:1530183 | 4.1E-10 | 0.5079234 | 0.326 | 0.231 | 1.34E-05 | Stroma 4 | zgc:153018 |
| romo13 | 4.8E-10 | 0.2889191 | 0.787 | 0.705 | 1.56E-05 | Stroma 4 | romo1 |
| napsa4 | 5.4E-10 | 0.4774187 | 0.253 | 0.168 | 1.76E-05 | Stroma 4 | napsa |
| klf12a4 | 5.4E-10 | 0.2988325 | 0.217 | 0.137 | 1.77E-05 | Stroma 4 | klf12a |
| chordc1a1 | 5.9E-10 | 0.6965729 | 0.126 | 0.07 | 1.91E-05 | Stroma 4 | chordc1a |
| nfil3-53 | 6.4E-10 | 0.4179049 | 0.215 | 0.138 | 2.07E-05 | Stroma 4 | nfil3-5 |
| hk13 | 7.1E-10 | 0.4702806 | 0.188 | 0.118 | 2.31E-05 | Stroma 4 | hk1 |
| ube2d2l5 | 7.6E-10 | 0.2873622 | 0.23 | 0.148 | 2.49E-05 | Stroma 4 | ube2d2l |
| sec61a15 | 8.8E-10 | 0.3403175 | 0.531 | 0.419 | 2.85E-05 | Stroma 4 | sec61a1 |
| smim153 | 1E-09 | 0.4821906 | 0.249 | 0.169 | 3.31E-05 | Stroma 4 | smim15 |
| snrpf2 | 1.1E-09 | 0.2809381 | 0.42 | 0.313 | 3.52E-05 | Stroma 4 | snrpf |
| PLEKHH33 | 1.1E-09 | 0.4478727 | 0.243 | 0.163 | 3.56E-05 | Stroma 4 | PLEKHH3 |
| tuba1b8 | 1.3E-09 | 0.3314735 | 0.407 | 0.308 | 4.3E-05 | Stroma 4 | tuba1b |
| hnrnpa0l.14 | 1.4E-09 | 0.2830419 | 0.739 | 0.666 | 4.53E-05 | Stroma 4 | hnrnpa0l.1 |
| si:ch211-225b11.13 | 1.4E-09 | 0.5564681 | 0.13 | 0.074 | 4.7E-05 | Stroma 4 | si:ch211-225b11.1 |
| pepd4 | 1.5E-09 | 0.3973307 | 0.304 | 0.212 | 4.88E-05 | Stroma 4 | pepd |
| hexim15 | 1.5E-09 | 0.3888535 | 0.325 | 0.233 | 4.98E-05 | Stroma 4 | hexim1 |
| cox5aa4 | 1.6E-09 | 0.3084142 | 0.386 | 0.285 | 5.32E-05 | Stroma 4 | cox5aa |
| psma23 | 1.9E-09 | 0.4232999 | 0.287 | 0.201 | 6.05E-05 | Stroma 4 | psma2 |
| tfap2c6 | 1.9E-09 | 0.3971101 | 0.285 | 0.195 | 6.17E-05 | Stroma 4 | tfap2c |
| pptc7a4 | 1.9E-09 | 0.3493739 | 0.318 | 0.223 | 6.24E-05 | Stroma 4 | pptc7a |
| si:dkey-25e12.33 | 2E-09 | 0.5746081 | 0.127 | 0.072 | 6.5E-05 | Stroma 4 | si:dkey-25e12.3 |
| icam34 | 2.2E-09 | 0.3399201 | 0.141 | 0.082 | 7.24E-05 | Stroma 4 | icam3 |
| dock71 | 2.3E-09 | 0.3940599 | 0.326 | 0.23 | 7.45E-05 | Stroma 4 | dock7 |
| ddit36 | 2.3E-09 | 0.2448215 | 0.689 | 0.58 | 7.58E-05 | Stroma 4 | ddit3 |
| socs3b5 | 2.5E-09 | 0.2159461 | 0.601 | 0.469 | 8.15E-05 | Stroma 4 | socs3b |
| atp5fa13 | 2.7E-09 | 0.2961804 | 0.566 | 0.46 | 8.82E-05 | Stroma 4 | atp5fa1 |
| si:dkey-177p2.65 | 2.9E-09 | 0.2968912 | 0.362 | 0.262 | 9.56E-05 | Stroma 4 | si:dkey-177p2.6 |
| btbd6a4 | 3E-09 | 0.6753824 | 0.135 | 0.079 | 9.88E-05 | Stroma 4 | btbd6a |
| ENSDARG0000007621 | 3E-09 | 0.2818693 | 0.719 | 0.622 | 9.9E-05 | Stroma 4 | ENSDARG00000076 |
| irs2b6 | 3.2E-09 | 0.4242898 | 0.321 | 0.229 | 0.000103 | Stroma 4 | irs2b |
| pfn11 | 3.2E-09 | 0.2806706 | 0.411 | 0.307 | 0.000106 | Stroma 4 | pfn1 |
| flot1b1 | 3.5E-09 | 0.4767497 | 0.262 | 0.182 | 0.000114 | Stroma 4 | flot1b |
| csrnp1b5 | 3.6E-09 | 0.4575267 | 0.269 | 0.186 | 0.000118 | Stroma 4 | csrnp1b |
| rasgef1ba3 | 4.4E-09 | 0.3739828 | 0.237 | 0.16 | 0.000142 | Stroma 4 | rasgef1ba |
| ecm1b5 | 4.5E-09 | 0.2484173 | 0.591 | 0.47 | 0.000148 | Stroma 4 | ecm1b |
| gstt1b4 | 5E-09 | 0.4887403 | 0.189 | 0.121 | 0.000162 | Stroma 4 | gstt1b |
| il4r.15 | 5.1E-09 | 0.4518499 | 0.226 | 0.152 | 0.000164 | Stroma 4 | il4r.1 |
| net16 | 5.4E-09 | 0.2637933 | 0.478 | 0.372 | 0.000175 | Stroma 4 | net1 |

| | | | | | | | |
|---|---|---|---|---|---|---|---|
| f105 | 5.7E-09 | 0.4147235 | 0.501 | 0.398 | 0.000184 | Stroma 4 | f10 |
| il13ra14 | 6.2E-09 | 0.5960713 | 0.14 | 0.084 | 0.0002 | Stroma 4 | il13ra1 |
| ier5l3 | 6.4E-09 | 0.3854767 | 0.419 | 0.318 | 0.000209 | Stroma 4 | ier5l |
| asap35 | 6.7E-09 | 0.4025343 | 0.195 | 0.126 | 0.000218 | Stroma 4 | asap3 |
| hnrnpabb2 | 8.7E-09 | 0.3420856 | 0.575 | 0.478 | 0.000283 | Stroma 4 | hnrnpabb |
| slc25a3b4 | 9.1E-09 | 0.261114 | 0.649 | 0.55 | 0.000296 | Stroma 4 | slc25a3b |
| tcp13 | 9.8E-09 | 0.3326474 | 0.415 | 0.321 | 0.000318 | Stroma 4 | tcp1 |
| prr12b3 | 9.9E-09 | 0.4851244 | 0.182 | 0.116 | 0.000323 | Stroma 4 | prr12b |
| tcf7l1a5 | 1E-08 | 0.5375616 | 0.18 | 0.116 | 0.000326 | Stroma 4 | tcf7l1a |
| ppp1r15a3 | 1E-08 | 0.463088 | 0.294 | 0.209 | 0.000336 | Stroma 4 | ppp1r15a |
| rdh10b3 | 1.1E-08 | 0.682353 | 0.127 | 0.074 | 0.000365 | Stroma 4 | rdh10b |
| marveld14 | 1.2E-08 | 0.3625744 | 0.294 | 0.209 | 0.000383 | Stroma 4 | marveld1 |
| plekha52 | 1.2E-08 | 0.8122839 | 0.121 | 0.071 | 0.000404 | Stroma 4 | plekha5 |
| peli1b5 | 1.3E-08 | 0.5789073 | 0.147 | 0.09 | 0.000423 | Stroma 4 | peli1b |
| ebf3a5 | 1.7E-08 | 0.213749 | 0.308 | 0.215 | 0.000544 | Stroma 4 | ebf3a |
| meis2a.15 | 1.8E-08 | 0.3732998 | 0.202 | 0.133 | 0.000585 | Stroma 4 | meis2a.1 |
| hivep3a5 | 1.9E-08 | 0.421775 | 0.172 | 0.109 | 0.000622 | Stroma 4 | hivep3a |
| tomm20b4 | 2E-08 | 0.34116 | 0.443 | 0.342 | 0.000653 | Stroma 4 | tomm20b |
| rbm4.24 | 2.1E-08 | 0.4128399 | 0.241 | 0.165 | 0.000667 | Stroma 4 | rbm4.2 |
| nutf2l3 | 2.2E-08 | 0.398086 | 0.16 | 0.101 | 0.000726 | Stroma 4 | nutf2l |
| isoc21 | 2.2E-08 | 0.5237638 | 0.168 | 0.107 | 0.000727 | Stroma 4 | isoc2 |
| txn6 | 2.3E-08 | 0.288873 | 0.451 | 0.372 | 0.000736 | Stroma 4 | txn |
| cpeb4b3 | 2.3E-08 | 0.4451342 | 0.242 | 0.167 | 0.000744 | Stroma 4 | cpeb4b |
| limch1b6 | 2.3E-08 | 0.4620696 | 0.147 | 0.09 | 0.000753 | Stroma 4 | limch1b |
| etv5b5 | 2.4E-08 | 0.4282756 | 0.159 | 0.099 | 0.000774 | Stroma 4 | etv5b |
| ptena2 | 2.5E-08 | 0.3216139 | 0.348 | 0.256 | 0.000813 | Stroma 4 | ptena |
| acadvl2 | 2.5E-08 | 0.5826318 | 0.156 | 0.098 | 0.000817 | Stroma 4 | acadvl |
| slc25a335 | 2.7E-08 | 0.4427156 | 0.182 | 0.117 | 0.000894 | Stroma 4 | slc25a33 |
| cox7a2a4 | 2.8E-08 | 0.3940779 | 0.358 | 0.269 | 0.000904 | Stroma 4 | cox7a2a |
| creb1a3 | 3E-08 | 0.4048873 | 0.237 | 0.162 | 0.000989 | Stroma 4 | creb1a |
| zgc:1103394 | 3.2E-08 | 0.2967037 | 0.302 | 0.215 | 0.001029 | Stroma 4 | zgc:110339 |
| vdac24 | 3.7E-08 | 0.272816 | 0.613 | 0.518 | 0.001187 | Stroma 4 | vdac2 |
| etv5a6 | 3.7E-08 | 0.6541477 | 0.161 | 0.104 | 0.001194 | Stroma 4 | etv5a |
| uqcrc2a3 | 4.3E-08 | 0.3854182 | 0.163 | 0.102 | 0.001412 | Stroma 4 | uqcrc2a |
| crsp73 | 4.6E-08 | 0.353607 | 0.123 | 0.072 | 0.001486 | Stroma 4 | crsp7 |
| cbfb3 | 4.6E-08 | 0.5290953 | 0.186 | 0.123 | 0.001507 | Stroma 4 | cbfb |
| si:ch211-114n24.61 | 4.9E-08 | 0.4846724 | 0.122 | 0.073 | 0.001579 | Stroma 4 | si:ch211-114n24.6 |
| cyp3c14 | 5E-08 | 0.5675564 | 0.15 | 0.095 | 0.001624 | Stroma 4 | cyp3c1 |
| ddx214 | 5.5E-08 | 0.3083861 | 0.437 | 0.342 | 0.001783 | Stroma 4 | ddx21 |
| rgmb4 | 5.5E-08 | 0.3927169 | 0.182 | 0.119 | 0.001784 | Stroma 4 | rgmb |
| snrpd12 | 7.2E-08 | 0.2871878 | 0.275 | 0.196 | 0.002347 | Stroma 4 | snrpd1 |
| pno13 | 7.5E-08 | 0.2954129 | 0.506 | 0.407 | 0.002436 | Stroma 4 | pno1 |
| tax1bp1b2 | 7.7E-08 | 0.2456232 | 0.374 | 0.28 | 0.002518 | Stroma 4 | tax1bp1b |
| cdh24 | 8.9E-08 | 0.2252624 | 0.39 | 0.291 | 0.002897 | Stroma 4 | cdh2 |
| atp5mc3b3 | 9E-08 | 0.2290527 | 0.696 | 0.622 | 0.002939 | Stroma 4 | atp5mc3b |
| CU927890.12 | 9.7E-08 | 0.6794078 | 0.102 | 0.059 | 0.003162 | Stroma 4 | CU927890.1 |
| pgk14 | 1.1E-07 | 0.3156832 | 0.287 | 0.207 | 0.003618 | Stroma 4 | pgk1 |
| snrpb1 | 1.2E-07 | 0.3008278 | 0.39 | 0.299 | 0.003763 | Stroma 4 | snrpb |

| | | | | | | | |
|---|---|---|---|---|---|---|---|
| ip6k2a5 | 1.2E-07 | 0.3883953 | 0.275 | 0.196 | 0.003974 | Stroma 4 | ip6k2a |
| sri3 | 1.2E-07 | 0.3231916 | 0.357 | 0.27 | 0.004011 | Stroma 4 | sri |
| pfdn42 | 1.2E-07 | 0.2779511 | 0.344 | 0.257 | 0.004027 | Stroma 4 | pfdn4 |
| ckbb4 | 1.3E-07 | 0.2794465 | 0.274 | 0.198 | 0.004078 | Stroma 4 | ckbb |
| tpi1b3 | 1.3E-07 | 0.2592054 | 0.455 | 0.356 | 0.00424 | Stroma 4 | tpi1b |
| zgc:1629445 | 1.4E-07 | 0.3330492 | 0.381 | 0.293 | 0.004518 | Stroma 4 | zgc:162944 |
| oser15 | 1.5E-07 | 0.5780699 | 0.138 | 0.086 | 0.004818 | Stroma 4 | oser1 |
| pdk2a2 | 1.5E-07 | 0.5617216 | 0.133 | 0.082 | 0.004889 | Stroma 4 | pdk2a |
| eif3s6ip3 | 1.5E-07 | 0.3680235 | 0.404 | 0.319 | 0.004894 | Stroma 4 | eif3s6ip |
| agpat31 | 1.5E-07 | 0.4459392 | 0.13 | 0.079 | 0.004897 | Stroma 4 | agpat3 |
| hexb4 | 1.5E-07 | 0.6719881 | 0.142 | 0.091 | 0.004995 | Stroma 4 | hexb |
| man2b14 | 1.6E-07 | 0.3755571 | 0.152 | 0.097 | 0.005229 | Stroma 4 | man2b1 |
| snrpd22 | 1.7E-07 | 0.3969696 | 0.277 | 0.201 | 0.005457 | Stroma 4 | snrpd2 |
| STMP15 | 1.7E-07 | 0.3501079 | 0.339 | 0.256 | 0.00555 | Stroma 4 | STMP1 |
| cct82 | 1.8E-07 | 0.3066081 | 0.359 | 0.27 | 0.005788 | Stroma 4 | cct8 |
| htra1a5 | 1.8E-07 | 0.3067137 | 0.306 | 0.227 | 0.005946 | Stroma 4 | htra1a |
| eif62 | 1.9E-07 | 0.3791823 | 0.247 | 0.175 | 0.006055 | Stroma 4 | eif6 |
| enc15 | 1.9E-07 | 0.2270425 | 0.187 | 0.124 | 0.006244 | Stroma 4 | enc1 |
| jpt1b3 | 2.1E-07 | 0.2539833 | 0.747 | 0.691 | 0.006864 | Stroma 4 | jpt1b |
| nufip24 | 2.1E-07 | 0.4569753 | 0.213 | 0.148 | 0.006923 | Stroma 4 | nufip2 |
| mt-nd13 | 2.1E-07 | 0.2211849 | 0.789 | 0.717 | 0.006986 | Stroma 4 | mt-nd1 |
| srp721 | 2.2E-07 | 0.4293114 | 0.217 | 0.15 | 0.007062 | Stroma 4 | srp72 |
| eif3d3 | 2.4E-07 | 0.2421861 | 0.525 | 0.42 | 0.007761 | Stroma 4 | eif3d |
| hmga1a2 | 2.4E-07 | 0.2580063 | 0.261 | 0.187 | 0.007892 | Stroma 4 | hmga1a |
| pim25 | 2.5E-07 | 0.3739907 | 0.305 | 0.225 | 0.008162 | Stroma 4 | pim2 |
| ehbp1l1b5 | 2.5E-07 | 0.4002146 | 0.188 | 0.126 | 0.008265 | Stroma 4 | ehbp1l1b |
| commd91 | 2.7E-07 | 0.4030965 | 0.109 | 0.064 | 0.008874 | Stroma 4 | commd9 |
| si:dkey-110g7.81 | 2.8E-07 | 0.3673041 | 0.164 | 0.106 | 0.009263 | Stroma 4 | si:dkey-110g7.8 |
| pfdn61 | 3.1E-07 | 0.3422927 | 0.203 | 0.139 | 0.009982 | Stroma 4 | pfdn6 |
| arhgef44 | 3.2E-07 | 0.723491 | 0.121 | 0.075 | 0.010433 | Stroma 4 | arhgef4 |
| mbd24 | 3.5E-07 | 0.7351097 | 0.13 | 0.082 | 0.011419 | Stroma 4 | mbd2 |
| kdm6ba4 | 3.5E-07 | 0.3932585 | 0.296 | 0.221 | 0.011448 | Stroma 4 | kdm6ba |
| crebzf1 | 3.5E-07 | 0.3076807 | 0.194 | 0.13 | 0.011492 | Stroma 4 | crebzf |
| bzw1a3 | 3.6E-07 | 0.2910016 | 0.401 | 0.316 | 0.011857 | Stroma 4 | bzw1a |
| si:ch211-232b12.53 | 3.8E-07 | 0.4399972 | 0.222 | 0.156 | 0.012211 | Stroma 4 | si:ch211-232b12.5 |
| mrpl111 | 4E-07 | 0.3285143 | 0.178 | 0.119 | 0.012897 | Stroma 4 | mrpl11 |
| nt5c3a4 | 4.3E-07 | 0.4619233 | 0.235 | 0.169 | 0.014003 | Stroma 4 | nt5c3a |
| si:ch1073-291c23.26 | 4.4E-07 | 0.2244353 | 0.587 | 0.473 | 0.014242 | Stroma 4 | si:ch1073-291c23.2 |
| gmps2 | 4.4E-07 | 0.6710042 | 0.195 | 0.136 | 0.014358 | Stroma 4 | gmps |
| igfbp34 | 4.7E-07 | 0.3787417 | 0.165 | 0.109 | 0.015424 | Stroma 4 | igfbp3 |
| rin25 | 4.8E-07 | 0.569651 | 0.168 | 0.112 | 0.015581 | Stroma 4 | rin2 |
| CU459056.13 | 4.8E-07 | 0.3858519 | 0.238 | 0.169 | 0.015692 | Stroma 4 | CU459056.1 |
| pak13 | 4.9E-07 | 0.7477944 | 0.111 | 0.068 | 0.015848 | Stroma 4 | pak1 |
| cbx7a5 | 4.9E-07 | 0.2133834 | 0.443 | 0.345 | 0.015853 | Stroma 4 | cbx7a |
| cers2b3 | 5.2E-07 | 0.5126926 | 0.112 | 0.068 | 0.016799 | Stroma 4 | cers2b |
| ngdn4 | 5.2E-07 | 0.3219013 | 0.224 | 0.157 | 0.017027 | Stroma 4 | ngdn |
| nuak1b4 | 5.2E-07 | 0.26271 | 0.37 | 0.281 | 0.017067 | Stroma 4 | nuak1b |
| pik3r3b4 | 5.4E-07 | 0.3046434 | 0.358 | 0.273 | 0.017664 | Stroma 4 | pik3r3b |

| | | | | | | | |
|---|---|---|---|---|---|---|---|
| slit33 | 5.6E-07 | 0.2630443 | 0.278 | 0.201 | 0.01813 | Stroma 4 | slit3 |
| mat2aa | 6.1E-07 | 0.5088442 | 0.105 | 0.063 | 0.019685 | Stroma 4 | mat2aa |
| mdh22 | 7.3E-07 | 0.2749045 | 0.526 | 0.439 | 0.023625 | Stroma 4 | mdh2 |
| cspg5a4 | 8.1E-07 | 0.5900214 | 0.122 | 0.077 | 0.026347 | Stroma 4 | cspg5a |
| xpo1b4 | 8.6E-07 | 0.4597499 | 0.155 | 0.101 | 0.028091 | Stroma 4 | xpo1b |
| skilb2 | 9.2E-07 | 0.551417 | 0.124 | 0.079 | 0.030042 | Stroma 4 | skilb |
| atp1a1a.15 | 9.6E-07 | 0.3284325 | 0.408 | 0.322 | 0.03109 | Stroma 4 | atp1a1a.1 |
| BX284696.12 | 1.1E-06 | 0.572553 | 0.115 | 0.071 | 0.036666 | Stroma 4 | BX284696.1 |
| kdm5c4 | 1.3E-06 | 0.3817861 | 0.214 | 0.152 | 0.041414 | Stroma 4 | kdm5c |
| mrpl232 | 1.3E-06 | 0.6270842 | 0.123 | 0.079 | 0.043673 | Stroma 4 | mrpl23 |
| foxo1a3 | 1.4E-06 | 0.267242 | 0.26 | 0.188 | 0.044638 | Stroma 4 | foxo1a |
| gadd45ga6 | 1.6E-06 | 0.2859568 | 0.36 | 0.276 | 0.051683 | Stroma 4 | gadd45ga |
| grinab4 | 1.8E-06 | 0.2185096 | 0.288 | 0.212 | 0.057791 | Stroma 4 | grinab |
| snrpg4 | 1.8E-06 | 0.2019175 | 0.25 | 0.183 | 0.059243 | Stroma 4 | snrpg |
| gse15 | 1.9E-06 | 0.3691819 | 0.182 | 0.125 | 0.060733 | Stroma 4 | gse1 |
| si:ch211-22i13.23 | 1.9E-06 | 0.5323113 | 0.118 | 0.074 | 0.061397 | Stroma 4 | si:ch211-22i13.2 |
| acss1 | 2.1E-06 | 0.4483924 | 0.109 | 0.067 | 0.068428 | Stroma 4 | acss1 |
| si:ch211-178n15.14 | 2.2E-06 | 0.2723353 | 0.233 | 0.168 | 0.07074 | Stroma 4 | si:ch211-178n15.1 |
| ddx39ab3 | 2.2E-06 | 0.3515012 | 0.231 | 0.168 | 0.072586 | Stroma 4 | ddx39ab |
| arvcfb3 | 2.3E-06 | 0.751625 | 0.15 | 0.103 | 0.075713 | Stroma 4 | arvcfb |
| calr3 | 2.4E-06 | 0.4193901 | 0.263 | 0.198 | 0.077245 | Stroma 4 | calr |
| eif4a1b2 | 2.7E-06 | 0.2096297 | 0.7 | 0.61 | 0.086881 | Stroma 4 | eif4a1b |
| ehd1a3 | 2.7E-06 | 0.4398971 | 0.105 | 0.064 | 0.088087 | Stroma 4 | ehd1a |
| si:ch1073-325m22.24 | 2.9E-06 | 0.3174885 | 0.378 | 0.302 | 0.094743 | Stroma 4 | si:ch1073-325m22. |
| dlx3b4 | 2.9E-06 | 0.2117796 | 0.453 | 0.359 | 0.095522 | Stroma 4 | dlx3b |
| rmdn13 | 3E-06 | 0.2148004 | 0.143 | 0.094 | 0.096971 | Stroma 4 | rmdn1 |
| fkbp24 | 3.1E-06 | 0.326041 | 0.272 | 0.203 | 0.10145 | Stroma 4 | fkbp2 |
| si:dkey-188i13.113 | 3.2E-06 | 0.3250534 | 0.272 | 0.202 | 0.102597 | Stroma 4 | si:dkey-188i13.11 |
| cct74 | 3.4E-06 | 0.2372423 | 0.461 | 0.371 | 0.109813 | Stroma 4 | cct7 |
| id16 | 3.7E-06 | 0.2381275 | 0.729 | 0.64 | 0.119274 | Stroma 4 | id1 |
| eif4g2b2 | 3.7E-06 | 0.2660408 | 0.259 | 0.191 | 0.121787 | Stroma 4 | eif4g2b |
| znf395a | 3.9E-06 | 0.5780733 | 0.108 | 0.067 | 0.126409 | Stroma 4 | znf395a |
| abracl3 | 3.9E-06 | 0.3257856 | 0.16 | 0.109 | 0.126545 | Stroma 4 | abracl |
| phb2a1 | 3.9E-06 | 0.4836157 | 0.163 | 0.111 | 0.126854 | Stroma 4 | phb2a |
| emc71 | 4.1E-06 | 0.452737 | 0.185 | 0.13 | 0.13367 | Stroma 4 | emc7 |
| tdh4 | 4.3E-06 | 0.4214848 | 0.312 | 0.24 | 0.138871 | Stroma 4 | tdh |
| tcf3b2 | 4.6E-06 | 0.304399 | 0.29 | 0.22 | 0.151002 | Stroma 4 | tcf3b |
| pbx44 | 4.6E-06 | 0.3753362 | 0.331 | 0.259 | 0.151126 | Stroma 4 | pbx4 |
| nanos16 | 4.7E-06 | 0.522711 | 0.267 | 0.205 | 0.153903 | Stroma 4 | nanos1 |
| srgap33 | 4.9E-06 | 0.3911514 | 0.27 | 0.202 | 0.158191 | Stroma 4 | srgap3 |
| ppp1caa3 | 4.9E-06 | 0.2197657 | 0.194 | 0.135 | 0.159389 | Stroma 4 | ppp1caa |
| ncor2 | 5E-06 | 0.2617075 | 0.256 | 0.188 | 0.162484 | Stroma 4 | ncor2 |
| larp4ab3 | 5.1E-06 | 0.343997 | 0.22 | 0.159 | 0.166221 | Stroma 4 | larp4ab |
| akirin22 | 5.2E-06 | 0.3900916 | 0.111 | 0.07 | 0.168025 | Stroma 4 | akirin2 |
| rassf15 | 5.3E-06 | 0.3801553 | 0.173 | 0.12 | 0.171801 | Stroma 4 | rassf1 |
| gtpbp44 | 5.4E-06 | 0.2445509 | 0.502 | 0.41 | 0.176813 | Stroma 4 | gtpbp4 |
| fgfr1a6 | 5.5E-06 | 0.399062 | 0.215 | 0.156 | 0.177344 | Stroma 4 | fgfr1a |
| glrx6 | 5.5E-06 | 0.2358174 | 0.442 | 0.357 | 0.179329 | Stroma 4 | glrx |

| | | | | | | | |
|---|---|---|---|---|---|---|---|
| dop1a4 | 6.1E-06 | 0.4082911 | 0.119 | 0.076 | 0.199899 | Stroma 4 | dop1a |
| hint15 | 6.2E-06 | 0.2100834 | 0.473 | 0.385 | 0.201291 | Stroma 4 | hint1 |
| rflnb3 | 6.7E-06 | 0.373527 | 0.161 | 0.111 | 0.218031 | Stroma 4 | rflnb |
| fbl2 | 7.5E-06 | 0.2905114 | 0.287 | 0.22 | 0.24525 | Stroma 4 | fbl |
| sulf2b6 | 8E-06 | 0.3293017 | 0.225 | 0.165 | 0.260911 | Stroma 4 | sulf2b |
| tomm20a.13 | 8.2E-06 | 0.3647927 | 0.217 | 0.16 | 0.265662 | Stroma 4 | tomm20a.1 |
| ube2e13 | 8.4E-06 | 0.3265924 | 0.168 | 0.116 | 0.271667 | Stroma 4 | ube2e1 |
| optn1 | 8.6E-06 | 0.5809661 | 0.11 | 0.07 | 0.280506 | Stroma 4 | optn |
| sigmar12 | 9.2E-06 | 0.4399648 | 0.142 | 0.096 | 0.2983 | Stroma 4 | sigmar1 |
| mt-nd34 | 9.5E-06 | 0.2554466 | 0.443 | 0.36 | 0.310053 | Stroma 4 | mt-nd3 |
| dusp72 | 9.8E-06 | 0.411051 | 0.114 | 0.074 | 0.318449 | Stroma 4 | dusp7 |
| chd22 | 9.8E-06 | 0.3016554 | 0.24 | 0.178 | 0.319685 | Stroma 4 | chd2 |
| dip2ba2 | 1E-05 | 0.4954085 | 0.115 | 0.075 | 0.332359 | Stroma 4 | dip2ba |
| bcar13 | 1.2E-05 | 0.5044852 | 0.138 | 0.093 | 0.37586 | Stroma 4 | bcar1 |
| MYO9B5 | 1.3E-05 | 0.3132135 | 0.175 | 0.123 | 0.421339 | Stroma 4 | MYO9B |
| naa102 | 1.3E-05 | 0.2235917 | 0.354 | 0.277 | 0.421876 | Stroma 4 | naa10 |
| alas15 | 1.3E-05 | 0.408637 | 0.155 | 0.107 | 0.429488 | Stroma 4 | alas1 |
| eif1b3 | 1.4E-05 | 0.2572183 | 0.277 | 0.209 | 0.444977 | Stroma 4 | eif1b |
| gpx4a4 | 1.5E-05 | 0.2246285 | 0.385 | 0.303 | 0.474832 | Stroma 4 | gpx4a |
| mt-nd23 | 1.5E-05 | 0.2252614 | 0.694 | 0.631 | 0.476644 | Stroma 4 | mt-nd2 |
| cox5ab5 | 1.6E-05 | 0.373252 | 0.281 | 0.22 | 0.510697 | Stroma 4 | cox5ab |
| ndufb22 | 1.6E-05 | 0.3256935 | 0.321 | 0.255 | 0.520812 | Stroma 4 | ndufb2 |
| atp2a2b2 | 1.6E-05 | 0.4289478 | 0.148 | 0.102 | 0.523143 | Stroma 4 | atp2a2b |
| fkbp33 | 1.6E-05 | 0.2708097 | 0.443 | 0.357 | 0.53214 | Stroma 4 | fkbp3 |
| cct45 | 1.7E-05 | 0.2777065 | 0.382 | 0.312 | 0.539347 | Stroma 4 | cct4 |
| phb3 | 1.7E-05 | 0.3440316 | 0.177 | 0.126 | 0.553366 | Stroma 4 | phb |
| plekhg22 | 1.7E-05 | 0.4972687 | 0.112 | 0.073 | 0.564 | Stroma 4 | plekhg2 |
| rasgef1bb6 | 1.8E-05 | 0.3028696 | 0.213 | 0.155 | 0.601611 | Stroma 4 | rasgef1bb |
| cox7b4 | 2E-05 | 0.2060688 | 0.479 | 0.396 | 0.647487 | Stroma 4 | cox7b |
| kdm5bb | 2E-05 | 0.390712 | 0.186 | 0.135 | 0.652882 | Stroma 4 | kdm5bb |
| ccng23 | 2E-05 | 0.2050242 | 0.29 | 0.222 | 0.660699 | Stroma 4 | ccng2 |
| tomm52 | 2.1E-05 | 0.2286331 | 0.526 | 0.447 | 0.670155 | Stroma 4 | tomm5 |
| g3bp14 | 2.2E-05 | 0.3354553 | 0.226 | 0.169 | 0.70411 | Stroma 4 | g3bp1 |
| gna12a2 | 2.2E-05 | 0.4303281 | 0.114 | 0.075 | 0.713489 | Stroma 4 | gna12a |
| metrn5 | 2.2E-05 | 0.2333151 | 0.167 | 0.117 | 0.713925 | Stroma 4 | metrn |
| pfdn51 | 2.3E-05 | 0.2552013 | 0.237 | 0.176 | 0.741137 | Stroma 4 | pfdn5 |
| cct53 | 2.4E-05 | 0.2061999 | 0.463 | 0.381 | 0.771608 | Stroma 4 | cct5 |
| psmb23 | 2.4E-05 | 0.2324551 | 0.336 | 0.265 | 0.774598 | Stroma 4 | psmb2 |
| ilf24 | 2.5E-05 | 0.2262554 | 0.217 | 0.16 | 0.814077 | Stroma 4 | ilf2 |
| ndufs43 | 2.7E-05 | 0.2542007 | 0.185 | 0.133 | 0.869968 | Stroma 4 | ndufs4 |
| tnfrsf11a4 | 2.7E-05 | 0.2355108 | 0.244 | 0.183 | 0.878756 | Stroma 4 | tnfrsf11a |
| myo18ab1 | 2.7E-05 | 0.325645 | 0.249 | 0.187 | 0.884506 | Stroma 4 | myo18ab |
| mapk64 | 3.1E-05 | 0.4787565 | 0.128 | 0.087 | 1 | Stroma 4 | mapk6 |
| mepce4 | 3.2E-05 | 0.5148683 | 0.147 | 0.103 | 1 | Stroma 4 | mepce |
| stip12 | 3.2E-05 | 0.2611423 | 0.217 | 0.16 | 1 | Stroma 4 | stip1 |
| zfand2a4 | 3.6E-05 | 0.4723301 | 0.103 | 0.067 | 1 | Stroma 4 | zfand2a |
| efnb3b4 | 3.7E-05 | 0.3331568 | 0.191 | 0.139 | 1 | Stroma 4 | efnb3b |
| COX7A22 | 3.7E-05 | 0.2771969 | 0.182 | 0.131 | 1 | Stroma 4 | COX7A2 |

| | | | | | | | |
|---|---|---|---|---|---|---|---|
| cmip1 | 4.4E-05 | 0.3111528 | 0.169 | 0.121 | 1 | Stroma 4 | cmip |
| hnrnpl22 | 4.4E-05 | 0.2783302 | 0.202 | 0.148 | 1 | Stroma 4 | hnrnpl2 |
| si:ch211-284f22.36 | 4.6E-05 | 0.2994889 | 0.165 | 0.117 | 1 | Stroma 4 | si:ch211-284f22.3 |
| dhx382 | 4.6E-05 | 0.3131586 | 0.142 | 0.098 | 1 | Stroma 4 | dhx38 |
| tet34 | 4.7E-05 | 0.3909112 | 0.141 | 0.099 | 1 | Stroma 4 | tet3 |
| baz2a1 | 5E-05 | 0.40014 | 0.148 | 0.104 | 1 | Stroma 4 | baz2a |
| nfil32 | 5.5E-05 | 0.3510433 | 0.12 | 0.081 | 1 | Stroma 4 | nfil3 |
| rnf1113 | 6.3E-05 | 0.433465 | 0.135 | 0.093 | 1 | Stroma 4 | rnf111 |
| prdx33 | 6.6E-05 | 0.2210363 | 0.193 | 0.141 | 1 | Stroma 4 | prdx3 |
| guk1a4 | 7.1E-05 | 0.2836655 | 0.185 | 0.136 | 1 | Stroma 4 | guk1a |
| rrs13 | 7.2E-05 | 0.3360566 | 0.156 | 0.111 | 1 | Stroma 4 | rrs1 |
| zgc:1620251 | 7.3E-05 | 0.328939 | 0.123 | 0.084 | 1 | Stroma 4 | zgc:162025 |
| eif3g3 | 7.9E-05 | 0.2102179 | 0.599 | 0.526 | 1 | Stroma 4 | eif3g |
| npm31 | 8.5E-05 | 0.3031586 | 0.17 | 0.123 | 1 | Stroma 4 | npm3 |
| khdrbs1b2 | 8.6E-05 | 0.2076125 | 0.306 | 0.239 | 1 | Stroma 4 | khdrbs1b |
| dkc12 | 8.8E-05 | 0.2423579 | 0.25 | 0.19 | 1 | Stroma 4 | dkc1 |
| eml34 | 9E-05 | 0.3667312 | 0.166 | 0.12 | 1 | Stroma 4 | eml3 |
| tp533 | 9.2E-05 | 0.274637 | 0.212 | 0.16 | 1 | Stroma 4 | tp53 |
| glsb4 | 9.3E-05 | 0.2960708 | 0.161 | 0.116 | 1 | Stroma 4 | glsb |
| zgc:1122344 | 9.4E-05 | 0.3432575 | 0.11 | 0.074 | 1 | Stroma 4 | zgc:112234 |
| sept9a5 | 9.6E-05 | 0.2392541 | 0.275 | 0.216 | 1 | Stroma 4 | sept9a |
| cfl1l3 | 9.9E-05 | 0.2947585 | 0.196 | 0.148 | 1 | Stroma 4 | cfl1l |
| rnaset23 | 0.0001 | 0.2509496 | 0.177 | 0.13 | 1 | Stroma 4 | rnaset2 |
| alyref3 | 0.00011 | 0.2399359 | 0.411 | 0.347 | 1 | Stroma 4 | alyref |
| czib2 | 0.00011 | 0.21334 | 0.137 | 0.095 | 1 | Stroma 4 | czib |
| psmb13 | 0.00012 | 0.2103109 | 0.326 | 0.262 | 1 | Stroma 4 | psmb1 |
| myd882 | 0.00012 | 0.3828994 | 0.114 | 0.078 | 1 | Stroma 4 | myd88 |
| atp5if1a2 | 0.00012 | 0.2153679 | 0.395 | 0.326 | 1 | Stroma 4 | atp5if1a |
| dusp64 | 0.00012 | 0.2540717 | 0.195 | 0.146 | 1 | Stroma 4 | dusp6 |
| zgc:1588031 | 0.00012 | 0.3119805 | 0.133 | 0.093 | 1 | Stroma 4 | zgc:158803 |
| galnt72 | 0.00012 | 0.3643775 | 0.161 | 0.117 | 1 | Stroma 4 | galnt7 |
| cox6a14 | 0.00012 | 0.2127944 | 0.561 | 0.491 | 1 | Stroma 4 | cox6a1 |
| phactr4b3 | 0.00012 | 0.460983 | 0.108 | 0.073 | 1 | Stroma 4 | phactr4b |
| nr1d2b5 | 0.00013 | 0.2773797 | 0.203 | 0.152 | 1 | Stroma 4 | nr1d2b |
| psma13 | 0.00013 | 0.2816397 | 0.376 | 0.311 | 1 | Stroma 4 | psma1 |
| yy1a1 | 0.00013 | 0.262322 | 0.177 | 0.129 | 1 | Stroma 4 | yy1a |
| cd151l5 | 0.00013 | 0.4282108 | 0.109 | 0.074 | 1 | Stroma 4 | cd151l |
| CU467861.62 | 0.00014 | 0.3523719 | 0.151 | 0.109 | 1 | Stroma 4 | CU467861.6 |
| aars13 | 0.00014 | 0.2103105 | 0.187 | 0.138 | 1 | Stroma 4 | aars1 |
| jcada3 | 0.00015 | 0.4992313 | 0.163 | 0.121 | 1 | Stroma 4 | jcada |
| srcap1 | 0.00016 | 0.2224334 | 0.196 | 0.146 | 1 | Stroma 4 | srcap |
| si:ch211-191j22.32 | 0.00016 | 0.3449272 | 0.164 | 0.12 | 1 | Stroma 4 | si:ch211-191j22.3 |
| rab5if4 | 0.00016 | 0.2858276 | 0.165 | 0.121 | 1 | Stroma 4 | rab5if |
| u2af11 | 0.00016 | 0.2413009 | 0.228 | 0.174 | 1 | Stroma 4 | u2af1 |
| gtf2a12 | 0.00017 | 0.4087349 | 0.119 | 0.083 | 1 | Stroma 4 | gtf2a1 |
| acp14 | 0.00017 | 0.3430867 | 0.112 | 0.077 | 1 | Stroma 4 | acp1 |
| taf91 | 0.00017 | 0.2866605 | 0.169 | 0.125 | 1 | Stroma 4 | taf9 |
| prelid1a5 | 0.00017 | 0.239452 | 0.169 | 0.124 | 1 | Stroma 4 | prelid1a |

| | | | | | | | |
|---|---|---|---|---|---|---|---|
| tuba8l25 | 0.00017 | 0.3107902 | 0.272 | 0.217 | 1 | Stroma 4 | tuba8l2 |
| amot1 | 0.00018 | 0.372832 | 0.1 | 0.067 | 1 | Stroma 4 | amot |
| sh3bp5la4 | 0.00018 | 0.2461852 | 0.147 | 0.106 | 1 | Stroma 4 | sh3bp5la |
| lrp1aa7 | 0.00019 | 0.2455614 | 0.747 | 0.673 | 1 | Stroma 4 | lrp1aa |
| oxsr1a2 | 0.0002 | 0.2044189 | 0.268 | 0.21 | 1 | Stroma 4 | oxsr1a |
| nucb14 | 0.00022 | 0.2422898 | 0.262 | 0.207 | 1 | Stroma 4 | nucb1 |
| suclg14 | 0.00023 | 0.2342319 | 0.226 | 0.175 | 1 | Stroma 4 | suclg1 |
| pum31 | 0.00024 | 0.2791395 | 0.117 | 0.081 | 1 | Stroma 4 | pum3 |
| cyb5a2 | 0.00024 | 0.5822828 | 0.126 | 0.09 | 1 | Stroma 4 | cyb5a |
| bcl2l104 | 0.00025 | 0.2229898 | 0.229 | 0.179 | 1 | Stroma 4 | bcl2l10 |
| tsr22 | 0.00025 | 0.4301862 | 0.103 | 0.07 | 1 | Stroma 4 | tsr2 |
| asah1b3 | 0.00026 | 0.2252419 | 0.239 | 0.185 | 1 | Stroma 4 | asah1b |
| wapla | 0.00027 | 0.4542313 | 0.128 | 0.091 | 1 | Stroma 4 | wapla |
| C2H14orf1191 | 0.00028 | 0.4166672 | 0.127 | 0.09 | 1 | Stroma 4 | C2H14orf119 |
| chd4a1 | 0.00029 | 0.2274047 | 0.285 | 0.228 | 1 | Stroma 4 | chd4a |
| ppp1r13l4 | 0.0003 | 0.2051664 | 0.124 | 0.087 | 1 | Stroma 4 | ppp1r13l |
| rnf102 | 0.00031 | 0.2140072 | 0.41 | 0.341 | 1 | Stroma 4 | rnf10 |
| scamp42 | 0.00031 | 0.3588709 | 0.112 | 0.078 | 1 | Stroma 4 | scamp4 |
| ANPEP4 | 0.00032 | 0.205591 | 0.268 | 0.21 | 1 | Stroma 4 | ANPEP |
| arfgap24 | 0.00033 | 0.3585526 | 0.143 | 0.105 | 1 | Stroma 4 | arfgap2 |
| degs14 | 0.00034 | 0.3331763 | 0.24 | 0.188 | 1 | Stroma 4 | degs1 |
| ENSDARG000000758 | 0.00034 | 0.3724937 | 0.12 | 0.085 | 1 | Stroma 4 | ENSDARG00000075 |
| mrpl302 | 0.00035 | 0.2194738 | 0.148 | 0.108 | 1 | Stroma 4 | mrpl30 |
| vbp12 | 0.00037 | 0.2025459 | 0.172 | 0.127 | 1 | Stroma 4 | vbp1 |
| gli34 | 0.00037 | 0.2501315 | 0.151 | 0.111 | 1 | Stroma 4 | gli3 |
| fasn3 | 0.00038 | 0.359105 | 0.141 | 0.103 | 1 | Stroma 4 | fasn |
| timm8b1 | 0.00038 | 0.2450107 | 0.143 | 0.104 | 1 | Stroma 4 | timm8b |
| atp5if1b5 | 0.00038 | 0.2157241 | 0.395 | 0.332 | 1 | Stroma 4 | atp5if1b |
| znf7032 | 0.00038 | 0.26657 | 0.133 | 0.096 | 1 | Stroma 4 | znf703 |
| tsc22d34 | 0.0004 | 0.2163935 | 0.478 | 0.402 | 1 | Stroma 4 | tsc22d3 |
| btg35 | 0.00042 | 0.2704488 | 0.168 | 0.126 | 1 | Stroma 4 | btg3 |
| gal3st24 | 0.00043 | 0.3030384 | 0.11 | 0.076 | 1 | Stroma 4 | gal3st2 |
| ssb1 | 0.0005 | 0.2125573 | 0.177 | 0.134 | 1 | Stroma 4 | ssb |
| rcan1a6 | 0.00051 | 0.2357625 | 0.286 | 0.23 | 1 | Stroma 4 | rcan1a |
| psme13 | 0.00056 | 0.2060385 | 0.154 | 0.113 | 1 | Stroma 4 | psme1 |
| mbnl24 | 0.00057 | 0.3077294 | 0.279 | 0.223 | 1 | Stroma 4 | mbnl2 |
| rif16 | 0.00059 | 0.3098357 | 0.12 | 0.086 | 1 | Stroma 4 | rif1 |
| cdc42ep4b6 | 0.00059 | 0.4607849 | 0.119 | 0.085 | 1 | Stroma 4 | cdc42ep4b |
| arid1aa2 | 0.0006 | 0.3415292 | 0.157 | 0.117 | 1 | Stroma 4 | arid1aa |
| ddx3xb2 | 0.00065 | 0.256619 | 0.202 | 0.157 | 1 | Stroma 4 | ddx3xb |
| sod24 | 0.00065 | 0.2175308 | 0.17 | 0.129 | 1 | Stroma 4 | sod2 |
| si:dkeyp-120h9.13 | 0.0007 | 0.2385645 | 0.118 | 0.084 | 1 | Stroma 4 | si:dkeyp-120h9.1 |
| ISCU3 | 0.00071 | 0.2351815 | 0.32 | 0.266 | 1 | Stroma 4 | ISCU |
| rexo13 | 0.00071 | 0.203427 | 0.158 | 0.119 | 1 | Stroma 4 | rexo1 |
| bahcc1a4 | 0.00073 | 0.2514917 | 0.121 | 0.087 | 1 | Stroma 4 | bahcc1a |
| fkbp10a4 | 0.00078 | 0.2008962 | 0.177 | 0.135 | 1 | Stroma 4 | fkbp10a |
| gtf2a23 | 0.0008 | 0.3636656 | 0.148 | 0.11 | 1 | Stroma 4 | gtf2a2 |
| snai25 | 0.00081 | 0.2188215 | 0.142 | 0.105 | 1 | Stroma 4 | snai2 |

| | | | | | | | |
|---|---|---|---|---|---|---|---|
| arl13 | 0.00082 | 0.3100148 | 0.209 | 0.165 | 1 | Stroma 4 | arl1 |
| smg7 | 0.00082 | 0.2498134 | 0.141 | 0.104 | 1 | Stroma 4 | smg7 |
| hacd22 | 0.00086 | 0.2517484 | 0.124 | 0.09 | 1 | Stroma 4 | hacd2 |
| cab39l12 | 0.00095 | 0.2282104 | 0.201 | 0.156 | 1 | Stroma 4 | cab39l1 |
| sf3a22 | 0.00095 | 0.2387352 | 0.204 | 0.16 | 1 | Stroma 4 | sf3a2 |
| lsm82 | 0.00097 | 0.4342678 | 0.114 | 0.083 | 1 | Stroma 4 | lsm8 |
| cbx1a3 | 0.0011 | 0.2212053 | 0.253 | 0.204 | 1 | Stroma 4 | cbx1a |
| polr2j3 | 0.00115 | 0.348944 | 0.13 | 0.096 | 1 | Stroma 4 | polr2j |
| antxr2a5 | 0.00116 | 0.2312826 | 0.392 | 0.335 | 1 | Stroma 4 | antxr2a |
| rnf1853 | 0.00117 | 0.2474854 | 0.204 | 0.16 | 1 | Stroma 4 | rnf185 |
| cmpk4 | 0.00127 | 0.2033338 | 0.209 | 0.165 | 1 | Stroma 4 | cmpk |
| ctsba3 | 0.00127 | 0.2929826 | 0.436 | 0.384 | 1 | Stroma 4 | ctsba |
| atf11 | 0.00132 | 0.2671557 | 0.149 | 0.113 | 1 | Stroma 4 | atf1 |
| pora4 | 0.00143 | 0.2975243 | 0.146 | 0.11 | 1 | Stroma 4 | pora |
| phf20b3 | 0.00145 | 0.2338694 | 0.22 | 0.175 | 1 | Stroma 4 | phf20b |
| npepl12 | 0.00147 | 0.237933 | 0.148 | 0.112 | 1 | Stroma 4 | npepl1 |
| uqcc21 | 0.00148 | 0.3109259 | 0.159 | 0.121 | 1 | Stroma 4 | uqcc2 |
| vamp53 | 0.00153 | 0.2823985 | 0.111 | 0.081 | 1 | Stroma 4 | vamp5 |
| map1lc3a4 | 0.00162 | 0.2122886 | 0.261 | 0.214 | 1 | Stroma 4 | map1lc3a |
| txnrd32 | 0.00162 | 0.2904485 | 0.131 | 0.097 | 1 | Stroma 4 | txnrd3 |
| nfe2l1b3 | 0.00163 | 0.2896435 | 0.119 | 0.087 | 1 | Stroma 4 | nfe2l1b |
| psmc1b2 | 0.00166 | 0.3534242 | 0.119 | 0.087 | 1 | Stroma 4 | psmc1b |
| ythdf22 | 0.00178 | 0.3479093 | 0.139 | 0.105 | 1 | Stroma 4 | ythdf2 |
| zswim82 | 0.00195 | 0.354695 | 0.112 | 0.082 | 1 | Stroma 4 | zswim8 |
| baz2ba5 | 0.00207 | 0.2391636 | 0.212 | 0.17 | 1 | Stroma 4 | baz2ba |
| cxxc5a2 | 0.00217 | 0.2474804 | 0.141 | 0.108 | 1 | Stroma 4 | cxxc5a |
| hcfc1b1 | 0.00224 | 0.2172952 | 0.161 | 0.124 | 1 | Stroma 4 | hcfc1b |
| tmtc32 | 0.0025 | 0.2726769 | 0.103 | 0.075 | 1 | Stroma 4 | tmtc3 |
| cars13 | 0.00261 | 0.4003075 | 0.11 | 0.081 | 1 | Stroma 4 | cars1 |
| tars13 | 0.00266 | 0.371816 | 0.101 | 0.073 | 1 | Stroma 4 | tars1 |
| fdx11 | 0.00282 | 0.2208401 | 0.142 | 0.109 | 1 | Stroma 4 | fdx1 |
| dspa3 | 0.00286 | 0.320942 | 0.16 | 0.126 | 1 | Stroma 4 | dspa |
| cdipt2 | 0.00299 | 0.2086851 | 0.149 | 0.114 | 1 | Stroma 4 | cdipt |
| atxn2l3 | 0.00326 | 0.2521 | 0.123 | 0.093 | 1 | Stroma 4 | atxn2l |
| baz1a4 | 0.0034 | 0.2282758 | 0.148 | 0.114 | 1 | Stroma 4 | baz1a |
| epc1b1 | 0.00346 | 0.4028198 | 0.131 | 0.1 | 1 | Stroma 4 | epc1b |
| rab35b2 | 0.0035 | 0.2732983 | 0.122 | 0.092 | 1 | Stroma 4 | rab35b |
| mapk14a4 | 0.00362 | 0.2637169 | 0.113 | 0.084 | 1 | Stroma 4 | mapk14a |
| stat22 | 0.00364 | 0.2086606 | 0.156 | 0.122 | 1 | Stroma 4 | stat2 |
| CU464124.13 | 0.00365 | 0.2627893 | 0.136 | 0.104 | 1 | Stroma 4 | CU464124.1 |
| zgc:1949895 | 0.00365 | 0.2796227 | 0.121 | 0.091 | 1 | Stroma 4 | zgc:194989 |
| gtf2b5 | 0.00374 | 0.313428 | 0.107 | 0.079 | 1 | Stroma 4 | gtf2b |
| tsg101b4 | 0.00376 | 0.286374 | 0.12 | 0.09 | 1 | Stroma 4 | tsg101b |
| cnot12 | 0.00384 | 0.2533517 | 0.14 | 0.108 | 1 | Stroma 4 | cnot1 |
| rad21a3 | 0.0042 | 0.2737435 | 0.241 | 0.201 | 1 | Stroma 4 | rad21a |
| frmd63 | 0.00457 | 0.2377981 | 0.231 | 0.189 | 1 | Stroma 4 | frmd6 |
| adgre104 | 0.00458 | 0.5381131 | 0.101 | 0.076 | 1 | Stroma 4 | adgre10 |
| smoc25 | 0.0058 | 0.3822164 | 0.119 | 0.091 | 1 | Stroma 4 | smoc2 |

| | | | | | | | |
|---|---|---|---|---|---|---|---|
| lamtor24 | 0.00595 | 0.2025105 | 0.154 | 0.121 | 1 | Stroma 4 | lamtor2 |
| dhrs13a.31 | 0.00658 | 0.2363646 | 0.158 | 0.125 | 1 | Stroma 4 | dhrs13a.3 |
| prkcsh4 | 0.00692 | 0.2345785 | 0.169 | 0.136 | 1 | Stroma 4 | prkcsh |
| irf2bp12 | 0.00737 | 0.2610365 | 0.127 | 0.098 | 1 | Stroma 4 | irf2bp1 |
| trim8b1 | 0.00806 | 0.2816742 | 0.168 | 0.136 | 1 | Stroma 4 | trim8b |
| zgc:1752144 | 0.00848 | 0.2396321 | 0.137 | 0.108 | 1 | Stroma 4 | zgc:175214 |
| pxk2 | 0.00863 | 0.2385532 | 0.139 | 0.109 | 1 | Stroma 4 | pxk |
| tgfb56 | 0.00866 | 0.2567217 | 0.143 | 0.114 | 1 | Stroma 4 | tgfb5 |
| ube2kb2 | 0.00882 | 0.3541832 | 0.108 | 0.083 | 1 | Stroma 4 | ube2kb |
| fbxo91 | 0.00904 | 0.3240686 | 0.105 | 0.081 | 1 | Stroma 4 | fbxo9 |
| ahcyl2b | 0.00962 | 0.2437983 | 0.112 | 0.086 | 1 | Stroma 4 | ahcyl2b |
| rapgef22 | 0.00983 | 0.2066197 | 0.136 | 0.106 | 1 | Stroma 4 | rapgef2 |
| timm92 | 0.00995 | 0.2355194 | 0.124 | 0.097 | 1 | Stroma 4 | timm9 |
| apoeb3 | 0 | 2.9049814 | 0.817 | 0.237 | 0 | DF 8 | apoeb |
| gyg1b4 | 0 | 2.3334476 | 0.834 | 0.256 | 0 | DF 8 | gyg1b |
| capn126 | 0 | 2.209623 | 0.727 | 0.216 | 0 | DF 8 | capn12 |
| hapln1a4 | 0 | 2.7206738 | 0.97 | 0.468 | 0 | DF 8 | hapln1a |
| aldh1a33 | 0 | 2.9382 | 0.585 | 0.101 | 0 | DF 8 | aldh1a3 |
| ptx3a7 | 0 | 1.990339 | 0.954 | 0.478 | 0 | DF 8 | ptx3a |
| si:ch211-260p9.3 | 0 | 6.2113392 | 0.398 | 0.009 | 0 | DF 8 | si:ch211-260p9.3 |
| sned13 | 0 | 3.1282589 | 0.389 | 0.056 | 0 | DF 8 | sned1 |
| FO203432.12 | 0 | 2.9161159 | 0.358 | 0.05 | 0 | DF 8 | FO203432.1 |
| crabp1b | 0 | 3.4086568 | 0.286 | 0.028 | 0 | DF 8 | crabp1b |
| rbp47 | 0 | 1.6042988 | 1 | 0.908 | 0 | DF 8 | rbp4 |
| vcanb4 | 5E-290 | 1.9834877 | 0.933 | 0.463 | 1.8E-285 | DF 8 | vcanb |
| cxcl12a8 | 3E-276 | 1.5649007 | 0.99 | 0.677 | 1E-271 | DF 8 | cxcl12a |
| ptgdsb.25 | 3E-275 | 1.6319666 | 0.992 | 0.661 | 9E-271 | DF 8 | ptgdsb.2 |
| slc51a5 | 8E-271 | 2.3472598 | 0.567 | 0.151 | 2.5E-266 | DF 8 | slc51a |
| plpp1a6 | 3E-233 | 1.695929 | 0.816 | 0.367 | 1E-228 | DF 8 | plpp1a |
| CR383676.27 | 9E-210 | 0.7276436 | 1 | 1 | 2.9E-205 | DF 8 | CR383676.2 |
| clec3ba4 | 1E-207 | 2.5455536 | 0.45 | 0.12 | 3.8E-203 | DF 8 | clec3ba |
| si:ch211-213d14.2 | 4E-207 | 3.3667229 | 0.185 | 0.022 | 1.2E-202 | DF 8 | si:ch211-213d14.2 |
| icn3 | 6E-201 | 0.8881542 | 1 | 0.996 | 2E-196 | DF 8 | icn |
| ccl25b6 | 4E-198 | 0.9718172 | 0.997 | 0.658 | 1.3E-193 | DF 8 | ccl25b |
| si:dkey-164f24.27 | 2E-185 | 2.0037286 | 0.517 | 0.164 | 5E-181 | DF 8 | si:dkey-164f24.2 |
| thy12 | 1E-180 | 2.0366847 | 0.595 | 0.224 | 4.7E-176 | DF 8 | thy1 |
| zgc:1943986 | 2E-179 | 1.5359561 | 0.772 | 0.376 | 5.7E-175 | DF 8 | zgc:194398 |
| fam169aa6 | 9E-179 | 1.4865879 | 0.789 | 0.397 | 3.1E-174 | DF 8 | fam169aa |
| postna9 | 4E-178 | 1.2350139 | 0.972 | 0.584 | 1.3E-173 | DF 8 | postna |
| ptgdsb.17 | 1E-169 | 1.3070291 | 0.855 | 0.456 | 3.1E-165 | DF 8 | ptgdsb.1 |
| hpdb8 | 1E-167 | 1.0995068 | 0.911 | 0.501 | 4.3E-163 | DF 8 | hpdb |
| has24 | 2E-166 | 2.2311887 | 0.395 | 0.108 | 8.1E-162 | DF 8 | has2 |
| ahnak3 | 2E-165 | 0.9713833 | 0.998 | 0.961 | 7.7E-161 | DF 8 | ahnak |
| bhmt7 | 4E-163 | 1.1402338 | 0.972 | 0.703 | 1.3E-158 | DF 8 | bhmt |
| CABZ01080042.1 | 5E-161 | 2.9705794 | 0.16 | 0.021 | 1.5E-156 | DF 8 | CABZ01080042.1 |
| kera5 | 4E-158 | 1.9443502 | 0.492 | 0.166 | 1.2E-153 | DF 8 | kera |
| hgd6 | 1E-150 | 1.2334116 | 0.807 | 0.422 | 3.5E-146 | DF 8 | hgd |
| vmo1a5 | 2E-150 | 1.0445052 | 0.979 | 0.695 | 5.5E-146 | DF 8 | vmo1a |

| | | | | | | | |
|---|---|---|---|---|---|---|---|
| lxn6 | 2E-148 | 1.0708425 | 0.952 | 0.697 | 8E-144 | DF 8 | lxn |
| shox1 | 6E-138 | 2.4729296 | 0.245 | 0.052 | 2E-133 | DF 8 | shox |
| si:ch211-243a20.36 | 3E-137 | 1.259472 | 0.652 | 0.275 | 9.3E-133 | DF 8 | si:ch211-243a20.3 |
| aoc26 | 4E-137 | 1.7434143 | 0.563 | 0.235 | 1.3E-132 | DF 8 | aoc2 |
| pcolcea7 | 4E-133 | 1.0537972 | 0.884 | 0.562 | 1.4E-128 | DF 8 | pcolcea |
| pck1 | 4E-132 | 3.2598932 | 0.102 | 0.01 | 1.4E-127 | DF 8 | pck1 |
| si:dkey-11f4.205 | 6E-131 | 1.2868414 | 0.8 | 0.483 | 1.9E-126 | DF 8 | si:dkey-11f4.20 |
| krt157 | 1E-130 | 0.9959628 | 0.951 | 0.696 | 4E-126 | DF 8 | krt15 |
| mfap57 | 4E-130 | 1.0978729 | 0.872 | 0.54 | 1.4E-125 | DF 8 | mfap5 |
| si:dkey-85k7.76 | 9E-130 | 1.4137322 | 0.616 | 0.274 | 2.9E-125 | DF 8 | si:dkey-85k7.7 |
| pawr5 | 1E-128 | 1.7634008 | 0.481 | 0.182 | 4.1E-124 | DF 8 | pawr |
| igfbp5b6 | 4E-126 | 1.1971736 | 0.856 | 0.566 | 1.5E-121 | DF 8 | igfbp5b |
| ecm1b6 | 1E-122 | 1.2731784 | 0.779 | 0.466 | 4.3E-118 | DF 8 | ecm1b |
| si:ch73-237c6.17 | 4E-122 | 1.3194347 | 0.699 | 0.359 | 1.4E-117 | DF 8 | si:ch73-237c6.1 |
| sema3e4 | 1E-121 | 2.1127216 | 0.319 | 0.091 | 4E-117 | DF 8 | sema3e |
| fibinb4 | 2E-120 | 1.4745692 | 0.59 | 0.269 | 6.6E-116 | DF 8 | fibinb |
| si:ch211-106h4.125 | 3E-118 | 0.8555052 | 0.809 | 0.452 | 9.7E-114 | DF 8 | si:ch211-106h4.12 |
| prrx1a5 | 5E-118 | 1.0727989 | 0.84 | 0.582 | 1.5E-113 | DF 8 | prrx1a |
| pah5 | 4E-117 | 1.1184386 | 0.703 | 0.344 | 1.3E-112 | DF 8 | pah |
| spock37 | 4E-117 | 0.8734695 | 0.802 | 0.405 | 1.4E-112 | DF 8 | spock3 |
| kctd15a | 1E-115 | 2.6633326 | 0.19 | 0.038 | 3.3E-111 | DF 8 | kctd15a |
| mlip3 | 1E-115 | 2.1843001 | 0.269 | 0.07 | 4.5E-111 | DF 8 | mlip |
| syt11a3 | 2E-115 | 1.6841856 | 0.437 | 0.16 | 4.9E-111 | DF 8 | syt11a |
| ccl443 | 2E-115 | 1.6772552 | 0.378 | 0.123 | 6.8E-111 | DF 8 | ccl44 |
| fstb8 | 5E-115 | 1.3161936 | 0.695 | 0.355 | 1.6E-110 | DF 8 | fstb |
| igsf21a3 | 3E-113 | 2.1615391 | 0.236 | 0.056 | 9.4E-109 | DF 8 | igsf21a |
| tnfsf105 | 4E-113 | 1.9778921 | 0.319 | 0.094 | 1.5E-108 | DF 8 | tnfsf10 |
| si:ch211-63p21.13 | 6E-112 | 2.1623675 | 0.214 | 0.047 | 1.8E-107 | DF 8 | si:ch211-63p21.1 |
| slc7a27 | 5E-110 | 1.5820702 | 0.425 | 0.154 | 1.8E-105 | DF 8 | slc7a2 |
| fgfbp2a6 | 2E-108 | 0.8834997 | 0.861 | 0.53 | 5.2E-104 | DF 8 | fgfbp2a |
| si:ch211-262i1.44 | 2E-106 | 1.8653471 | 0.279 | 0.077 | 6.2E-102 | DF 8 | si:ch211-262i1.4 |
| cfp8 | 3E-104 | 1.208383 | 0.652 | 0.334 | 9.4E-100 | DF 8 | cfp |
| tagln5 | 3E-103 | 0.7188917 | 0.855 | 0.533 | 8.3E-99 | DF 8 | tagln |
| herc12 | 3E-103 | 2.1645506 | 0.304 | 0.095 | 1.05E-98 | DF 8 | herc1 |
| C18H15orf485 | 4E-102 | 1.3039386 | 0.608 | 0.304 | 1.15E-97 | DF 8 | C18H15orf48 |
| sfrp1a4 | 9E-102 | 1.0360341 | 0.756 | 0.432 | 2.91E-97 | DF 8 | sfrp1a |
| si:dkey-285b23.4 | 1E-101 | 2.8384312 | 0.112 | 0.016 | 4.36E-97 | DF 8 | si:dkey-285b23.4 |
| ryr35 | 8E-99 | 1.8705976 | 0.288 | 0.086 | 2.75E-94 | DF 8 | ryr3 |
| gstz16 | 4E-97 | 1.3349986 | 0.63 | 0.339 | 1.29E-92 | DF 8 | gstz1 |
| krt916 | 2.7E-95 | 0.5940178 | 0.994 | 0.9 | 8.82E-91 | DF 8 | krt91 |
| nfil3-67 | 4.4E-95 | 1.3062699 | 0.595 | 0.301 | 1.44E-90 | DF 8 | nfil3-6 |
| itga6b4 | 4.8E-92 | 1.2807299 | 0.578 | 0.291 | 1.55E-87 | DF 8 | itga6b |
| tcf126 | 6.3E-91 | 0.9824867 | 0.78 | 0.518 | 2.06E-86 | DF 8 | tcf12 |
| cdon5 | 1.5E-90 | 1.5807231 | 0.432 | 0.184 | 4.73E-86 | DF 8 | cdon |
| ca65 | 4.4E-87 | 0.4749942 | 0.7 | 0.341 | 1.43E-82 | DF 8 | ca6 |
| lhfpl65 | 8.5E-87 | 1.2248765 | 0.664 | 0.413 | 2.75E-82 | DF 8 | lhfpl6 |
| asip14 | 1.6E-84 | 1.3867524 | 0.416 | 0.173 | 5.17E-80 | DF 8 | asip1 |
| krt45 | 2.4E-84 | 0.7512974 | 0.935 | 0.756 | 7.74E-80 | DF 8 | krt4 |

| | | | | | | | |
|---|---|---|---|---|---|---|---|
| tob1b6 | 1.1E-83 | 0.8101282 | 0.895 | 0.699 | 3.65E-79 | DF 8 | tob1b |
| wfdc21 | 7.4E-83 | 1.0389326 | 0.675 | 0.396 | 2.41E-78 | DF 8 | wfdc2 |
| BX663503.36 | 2.9E-82 | 0.9540498 | 0.686 | 0.398 | 9.46E-78 | DF 8 | BX663503.3 |
| ppap2d4 | 1.8E-80 | 1.2606325 | 0.481 | 0.225 | 5.81E-76 | DF 8 | ppap2d |
| aqp3a4 | 1.2E-79 | 0.895681 | 0.607 | 0.314 | 3.96E-75 | DF 8 | aqp3a |
| tnfaip64 | 2.5E-79 | 0.989586 | 0.707 | 0.426 | 8.1E-75 | DF 8 | tnfaip6 |
| si:ch211-170d8.84 | 8E-79 | 0.9792666 | 0.732 | 0.48 | 2.59E-74 | DF 8 | si:ch211-170d8.8 |
| zgc:1951735 | 8.5E-79 | 1.2333775 | 0.649 | 0.392 | 2.76E-74 | DF 8 | zgc:195173 |
| si:dkey-261h17.15 | 2.8E-78 | 0.7119086 | 0.576 | 0.285 | 9E-74 | DF 8 | si:dkey-261h17.1 |
| emid12 | 4.1E-76 | 1.2822036 | 0.35 | 0.136 | 1.32E-71 | DF 8 | emid1 |
| si:ch73-173h19.36 | 1.1E-75 | 1.1348912 | 0.496 | 0.234 | 3.44E-71 | DF 8 | si:ch73-173h19.3 |
| enpp25 | 4.1E-75 | 0.931857 | 0.71 | 0.44 | 1.33E-70 | DF 8 | enpp2 |
| fmoda5 | 1.6E-74 | 1.1045752 | 0.424 | 0.179 | 5.06E-70 | DF 8 | fmoda |
| gbe1a6 | 1.2E-72 | 1.5296126 | 0.3 | 0.109 | 3.85E-68 | DF 8 | gbe1a |
| olfml2a1 | 9.2E-71 | 2.2950416 | 0.154 | 0.037 | 3E-66 | DF 8 | olfml2a |
| tspan314 | 1.2E-70 | 1.3502106 | 0.451 | 0.222 | 4.04E-66 | DF 8 | tspan31 |
| gfra2a | 5.9E-70 | 2.078056 | 0.157 | 0.039 | 1.91E-65 | DF 8 | gfra2a |
| plpp38 | 3.1E-69 | 0.8462658 | 0.724 | 0.45 | 9.97E-65 | DF 8 | plpp3 |
| ccn1l12 | 2.4E-67 | 2.1310893 | 0.147 | 0.036 | 7.84E-63 | DF 8 | ccn1l1 |
| mao4 | 1.5E-63 | 1.8029472 | 0.215 | 0.07 | 4.74E-59 | DF 8 | mao |
| prrx1b6 | 1.9E-63 | 1.0550401 | 0.609 | 0.376 | 6.16E-59 | DF 8 | prrx1b |
| si:ch211-105c13.36 | 8.5E-63 | 0.677077 | 0.801 | 0.565 | 2.76E-58 | DF 8 | si:ch211-105c13.3 |
| metrnla4 | 8.9E-63 | 1.1846392 | 0.538 | 0.311 | 2.9E-58 | DF 8 | metrnla |
| fzd102 | 2.2E-62 | 1.9706919 | 0.158 | 0.042 | 7.05E-58 | DF 8 | fzd10 |
| kazald35 | 1E-61 | 1.1198778 | 0.489 | 0.257 | 3.35E-57 | DF 8 | kazald3 |
| qdpra7 | 1.3E-61 | 0.6913776 | 0.699 | 0.421 | 4.21E-57 | DF 8 | qdpra |
| tat5 | 7.5E-59 | 1.2911511 | 0.46 | 0.241 | 2.43E-54 | DF 8 | tat |
| elnb1 | 3.2E-58 | 1.4226989 | 0.167 | 0.048 | 1.04E-53 | DF 8 | elnb |
| CABZ01085139.12 | 3.9E-58 | 1.5670571 | 0.196 | 0.062 | 1.25E-53 | DF 8 | CABZ01085139.1 |
| serpinb145 | 7.8E-58 | 1.4431723 | 0.334 | 0.15 | 2.53E-53 | DF 8 | serpinb14 |
| cebpa4 | 6.4E-57 | 1.31108 | 0.368 | 0.172 | 2.07E-52 | DF 8 | cebpa |
| gig2d2 | 2.1E-56 | 1.71336 | 0.171 | 0.051 | 6.74E-52 | DF 8 | gig2d |
| efemp2a5 | 6E-56 | 0.969884 | 0.581 | 0.358 | 1.94E-51 | DF 8 | efemp2a |
| capn2b8 | 6.3E-55 | 1.0212879 | 0.538 | 0.318 | 2.06E-50 | DF 8 | capn2b |
| tmsb4x3 | 1.1E-54 | 0.3576678 | 0.998 | 0.992 | 3.5E-50 | DF 8 | tmsb4x |
| cdh7a | 1.3E-54 | 2.4202002 | 0.1 | 0.021 | 4.26E-50 | DF 8 | cdh7a |
| gsta.16 | 4.3E-54 | 1.0241337 | 0.562 | 0.354 | 1.4E-49 | DF 8 | gsta.1 |
| slc12a7b4 | 4.5E-54 | 1.410116 | 0.328 | 0.149 | 1.45E-49 | DF 8 | slc12a7b |
| mxra5b3 | 5.7E-54 | 1.7042124 | 0.182 | 0.058 | 1.87E-49 | DF 8 | mxra5b |
| timp4.35 | 2.8E-53 | 0.4903672 | 0.697 | 0.436 | 9.22E-49 | DF 8 | timp4.3 |
| tnrc186 | 5.2E-53 | 0.9748014 | 0.604 | 0.404 | 1.68E-48 | DF 8 | tnrc18 |
| soul54 | 5.6E-53 | 0.9500742 | 0.492 | 0.273 | 1.84E-48 | DF 8 | soul5 |
| dio15 | 3.2E-52 | 1.5396019 | 0.282 | 0.12 | 1.04E-47 | DF 8 | dio1 |
| mfap26 | 3.5E-52 | 1.0437317 | 0.586 | 0.381 | 1.13E-47 | DF 8 | mfap2 |
| itm2ba4 | 1.6E-51 | 0.6167522 | 0.856 | 0.74 | 5.16E-47 | DF 8 | itm2ba |
| b2ml7 | 3.6E-51 | 0.4459639 | 0.971 | 0.896 | 1.16E-46 | DF 8 | b2ml |
| smoc15 | 4.3E-51 | 1.1438324 | 0.402 | 0.204 | 1.38E-46 | DF 8 | smoc1 |
| lix1l5 | 5.6E-51 | 1.2857936 | 0.354 | 0.171 | 1.82E-46 | DF 8 | lix1l |

| | | | | | | | |
|---|---|---|---|---|---|---|---|
| eppk14 | 2.2E-50 | 1.1696905 | 0.401 | 0.208 | 7.18E-46 | DF 8 | eppk1 |
| aldh9a1a.16 | 4.4E-50 | 0.9591367 | 0.668 | 0.478 | 1.42E-45 | DF 8 | aldh9a1a.1 |
| selenop4 | 4.4E-50 | 0.5659928 | 0.907 | 0.738 | 1.44E-45 | DF 8 | selenop |
| igfbp2b3 | 5E-50 | 1.6114972 | 0.196 | 0.068 | 1.63E-45 | DF 8 | igfbp2b |
| ebf3a6 | 6.1E-50 | 1.1709967 | 0.405 | 0.213 | 1.98E-45 | DF 8 | ebf3a |
| anxa1a4 | 1.1E-49 | 0.4506898 | 0.98 | 0.904 | 3.57E-45 | DF 8 | anxa1a |
| adcy6b | 6E-49 | 1.9707238 | 0.113 | 0.028 | 1.95E-44 | DF 8 | adcy6b |
| cavin1b6 | 6.4E-49 | 0.555435 | 0.838 | 0.643 | 2.08E-44 | DF 8 | cavin1b |
| cebpd7 | 1.8E-48 | 0.8079333 | 0.754 | 0.574 | 5.81E-44 | DF 8 | cebpd |
| slc1a56 | 2.4E-48 | 1.0651906 | 0.45 | 0.254 | 7.96E-44 | DF 8 | slc1a5 |
| efna2a1 | 1.3E-47 | 1.9060786 | 0.145 | 0.043 | 4.15E-43 | DF 8 | efna2a |
| ism2a4 | 2.2E-47 | 1.7120137 | 0.159 | 0.05 | 7.2E-43 | DF 8 | ism2a |
| gstp17 | 2.5E-47 | 0.6711405 | 0.8 | 0.641 | 8.15E-43 | DF 8 | gstp1 |
| skia7 | 2.6E-47 | 0.8416792 | 0.635 | 0.446 | 8.33E-43 | DF 8 | skia |
| ifitm18 | 1.2E-46 | 0.3807524 | 0.983 | 0.786 | 3.95E-42 | DF 8 | ifitm1 |
| si:ch73-380l3.25 | 3.4E-46 | 0.8277761 | 0.615 | 0.413 | 1.11E-41 | DF 8 | si:ch73-380l3.2 |
| pcdh18a5 | 5E-46 | 1.5766734 | 0.212 | 0.081 | 1.62E-41 | DF 8 | pcdh18a |
| slc22a26 | 8E-46 | 1.0805776 | 0.377 | 0.192 | 2.62E-41 | DF 8 | slc22a2 |
| ttc368 | 9.5E-45 | 0.7368732 | 0.577 | 0.35 | 3.1E-40 | DF 8 | ttc36 |
| tsku5 | 1.5E-44 | 0.9542462 | 0.511 | 0.316 | 4.76E-40 | DF 8 | tsku |
| neu3.27 | 5.5E-44 | 0.9936725 | 0.409 | 0.219 | 1.78E-39 | DF 8 | neu3.2 |
| cav14 | 8.6E-44 | 0.5807919 | 0.809 | 0.63 | 2.79E-39 | DF 8 | cav1 |
| pcbd17 | 1.2E-43 | 0.8170496 | 0.524 | 0.313 | 4.06E-39 | DF 8 | pcbd1 |
| slc40a15 | 1.3E-43 | 1.0880646 | 0.377 | 0.199 | 4.2E-39 | DF 8 | slc40a1 |
| crybg1b3 | 3.6E-43 | 0.7162056 | 0.664 | 0.477 | 1.16E-38 | DF 8 | crybg1b |
| sema3aa5 | 1E-42 | 1.2784324 | 0.249 | 0.106 | 3.34E-38 | DF 8 | sema3aa |
| agmo4 | 2.7E-42 | 1.619217 | 0.204 | 0.08 | 8.68E-38 | DF 8 | agmo |
| ramp27 | 3.3E-42 | 1.5410357 | 0.29 | 0.139 | 1.07E-37 | DF 8 | ramp2 |
| dcbld26 | 1.3E-41 | 1.2693854 | 0.27 | 0.122 | 4.33E-37 | DF 8 | dcbld2 |
| madd2 | 1.8E-41 | 1.6059974 | 0.172 | 0.062 | 5.89E-37 | DF 8 | madd |
| FQ323156.17 | 2.5E-41 | 1.0692534 | 0.365 | 0.192 | 8.16E-37 | DF 8 | FQ323156.1 |
| serpinh1b2 | 3E-41 | 0.8844123 | 0.724 | 0.58 | 9.85E-37 | DF 8 | serpinh1b |
| cd248a5 | 1E-40 | 1.2195247 | 0.257 | 0.114 | 3.24E-36 | DF 8 | cd248a |
| slc48a1b3 | 1.4E-40 | 1.2503116 | 0.288 | 0.139 | 4.56E-36 | DF 8 | slc48a1b |
| col6a27 | 1.4E-40 | 0.3837561 | 0.967 | 0.875 | 4.66E-36 | DF 8 | col6a2 |
| gpc1b6 | 1.6E-40 | 0.8502156 | 0.578 | 0.399 | 5.36E-36 | DF 8 | gpc1b |
| si:dkey-40c11.24 | 2E-40 | 0.9777362 | 0.493 | 0.314 | 6.57E-36 | DF 8 | si:dkey-40c11.2 |
| vegfc4 | 3.5E-40 | 1.4985191 | 0.212 | 0.086 | 1.15E-35 | DF 8 | vegfc |
| nfe2l2a7 | 5.7E-40 | 0.8496622 | 0.53 | 0.341 | 1.84E-35 | DF 8 | nfe2l2a |
| zfhx35 | 3.8E-39 | 0.7631669 | 0.652 | 0.484 | 1.25E-34 | DF 8 | zfhx3 |
| il1rapl1a2 | 9.3E-39 | 1.7340049 | 0.147 | 0.05 | 3.03E-34 | DF 8 | il1rapl1a |
| s100a10b4 | 1E-38 | 0.2311771 | 0.998 | 0.994 | 3.27E-34 | DF 8 | s100a10b |
| lpar14 | 1.5E-38 | 1.1044088 | 0.339 | 0.179 | 4.85E-34 | DF 8 | lpar1 |
| mhc1zaa5 | 2.3E-38 | 1.3538861 | 0.217 | 0.091 | 7.56E-34 | DF 8 | mhc1zaa |
| bsg4 | 3.3E-38 | 0.7215244 | 0.706 | 0.572 | 1.08E-33 | DF 8 | bsg |
| zgc:565858 | 3.9E-38 | 0.6973144 | 0.527 | 0.325 | 1.26E-33 | DF 8 | zgc:56585 |
| zcchc247 | 6.1E-38 | 1.0268206 | 0.428 | 0.254 | 1.98E-33 | DF 8 | zcchc24 |
| gstm.17 | 6.9E-38 | 0.633468 | 0.736 | 0.595 | 2.24E-33 | DF 8 | gstm.1 |

| | | | | | | | |
|---|---|---|---|---|---|---|---|
| ephb3a5 | 7.3E-38 | 1.4136335 | 0.235 | 0.104 | 2.39E-33 | DF 8 | ephb3a |
| cxcl18b6 | 1.2E-37 | 0.4347541 | 0.863 | 0.699 | 3.98E-33 | DF 8 | cxcl18b |
| notch32 | 2.9E-37 | 0.7134174 | 0.529 | 0.343 | 9.47E-33 | DF 8 | notch3 |
| si:dkey-240h12.45 | 3.6E-37 | 1.2347456 | 0.253 | 0.116 | 1.17E-32 | DF 8 | si:dkey-240h12.4 |
| fstl1b4 | 6.7E-37 | 0.4514676 | 0.868 | 0.755 | 2.17E-32 | DF 8 | fstl1b |
| slc35c24 | 8.1E-37 | 1.2087872 | 0.296 | 0.149 | 2.65E-32 | DF 8 | slc35c2 |
| entpd15 | 1.1E-36 | 1.3651762 | 0.196 | 0.08 | 3.42E-32 | DF 8 | entpd1 |
| mamdc2b2 | 1.5E-36 | 0.2426869 | 0.43 | 0.239 | 4.98E-32 | DF 8 | mamdc2b |
| rgmd3 | 1.5E-36 | 1.5241479 | 0.2 | 0.083 | 4.99E-32 | DF 8 | rgmd |
| CR383676.37 | 1.9E-36 | 0.8842076 | 0.627 | 0.463 | 6.19E-32 | DF 8 | CR383676.3 |
| nkx3-26 | 3.2E-36 | 0.9976889 | 0.354 | 0.193 | 1.04E-31 | DF 8 | nkx3-2 |
| ppl6 | 5.9E-36 | 1.1562636 | 0.301 | 0.152 | 1.91E-31 | DF 8 | ppl |
| pcdh116 | 6E-36 | 1.4059315 | 0.185 | 0.074 | 1.95E-31 | DF 8 | pcdh11 |
| col6a3.15 | 1.2E-35 | 0.3394481 | 0.99 | 0.937 | 3.87E-31 | DF 8 | col6a3.1 |
| tmem59l4 | 2.6E-35 | 1.3467661 | 0.229 | 0.104 | 8.53E-31 | DF 8 | tmem59l |
| pik3ip13 | 6.2E-35 | 1.5562309 | 0.182 | 0.074 | 2.03E-30 | DF 8 | pik3ip1 |
| glud1a4 | 1.6E-34 | 0.7191956 | 0.594 | 0.438 | 5.06E-30 | DF 8 | glud1a |
| zbtb182 | 1.6E-34 | 1.6407375 | 0.154 | 0.057 | 5.09E-30 | DF 8 | zbtb18 |
| lmna5 | 2.5E-34 | 0.381487 | 0.891 | 0.779 | 8.13E-30 | DF 8 | lmna |
| myh9b3 | 2.7E-34 | 1.0295083 | 0.462 | 0.3 | 8.66E-30 | DF 8 | myh9b |
| ptprua | 3.3E-34 | 2.1137992 | 0.101 | 0.03 | 1.08E-29 | DF 8 | ptprua |
| gstk13 | 3.5E-34 | 1.220219 | 0.3 | 0.159 | 1.14E-29 | DF 8 | gstk1 |
| ppp1r1c5 | 9E-34 | 0.9916802 | 0.265 | 0.129 | 2.94E-29 | DF 8 | ppp1r1c |
| agtr23 | 1E-33 | 1.2518809 | 0.242 | 0.114 | 3.27E-29 | DF 8 | agtr2 |
| twist36 | 1.7E-33 | 0.8094624 | 0.421 | 0.251 | 5.48E-29 | DF 8 | twist3 |
| tlcd4b5 | 1.9E-33 | 1.3664506 | 0.192 | 0.081 | 6.33E-29 | DF 8 | tlcd4b |
| CU914776.14 | 2.4E-33 | 0.9896228 | 0.196 | 0.082 | 7.96E-29 | DF 8 | CU914776.1 |
| slco2b16 | 2.8E-33 | 0.7848327 | 0.385 | 0.219 | 9.17E-29 | DF 8 | slco2b1 |
| prx3 | 3.2E-33 | 1.3035775 | 0.165 | 0.064 | 1.05E-28 | DF 8 | prx |
| slc12a24 | 3.9E-33 | 1.2185143 | 0.278 | 0.142 | 1.25E-28 | DF 8 | slc12a2 |
| loxl17 | 4.5E-33 | 0.8618692 | 0.359 | 0.198 | 1.46E-28 | DF 8 | loxl1 |
| hspa12b5 | 1.1E-32 | 1.1834202 | 0.249 | 0.121 | 3.49E-28 | DF 8 | hspa12b |
| angptl56 | 2.4E-32 | 1.0158749 | 0.316 | 0.171 | 7.78E-28 | DF 8 | angptl5 |
| serpina10a6 | 3E-32 | 0.8173972 | 0.529 | 0.376 | 9.62E-28 | DF 8 | serpina10a |
| angptl2a6 | 3.1E-32 | 0.639204 | 0.499 | 0.316 | 1.01E-27 | DF 8 | angptl2a |
| tbx3a5 | 4.1E-32 | 0.9776534 | 0.254 | 0.122 | 1.32E-27 | DF 8 | tbx3a |
| emilin2b3 | 4.1E-32 | 1.2698916 | 0.168 | 0.067 | 1.33E-27 | DF 8 | emilin2b |
| creg25 | 4.7E-32 | 0.8710855 | 0.48 | 0.323 | 1.51E-27 | DF 8 | creg2 |
| plxna41 | 5.8E-32 | 1.4682656 | 0.152 | 0.058 | 1.87E-27 | DF 8 | plxna4 |
| olfml2ba4 | 6.4E-32 | 0.9643202 | 0.257 | 0.124 | 2.08E-27 | DF 8 | olfml2ba |
| ccnd2a2 | 1.1E-31 | 1.0650732 | 0.364 | 0.216 | 3.45E-27 | DF 8 | ccnd2a |
| ppp1r3ca5 | 2.5E-31 | 0.9839833 | 0.356 | 0.206 | 8.02E-27 | DF 8 | ppp1r3ca |
| igfbp6b2 | 2.8E-31 | 1.4659726 | 0.152 | 0.059 | 9.08E-27 | DF 8 | igfbp6b |
| cdr2l3 | 4.7E-31 | 1.4137194 | 0.173 | 0.072 | 1.52E-26 | DF 8 | cdr2l |
| aldob8 | 6.1E-31 | 0.7959036 | 0.476 | 0.309 | 1.98E-26 | DF 8 | aldob |
| mustn1a4 | 9.2E-31 | 1.2134071 | 0.187 | 0.08 | 2.99E-26 | DF 8 | mustn1a |
| pigr3 | 1.6E-30 | 1.1956734 | 0.324 | 0.186 | 5.26E-26 | DF 8 | pigr |
| sema5a4 | 1.8E-30 | 1.4297437 | 0.184 | 0.08 | 5.91E-26 | DF 8 | sema5a |

| | | | | | | | |
|---|---|---|---|---|---|---|---|
| tgfb1a5 | 2.3E-30 | 0.8039596 | 0.428 | 0.271 | 7.46E-26 | DF 8 | tgfb1a |
| emx26 | 4.6E-30 | 1.353732 | 0.2 | 0.091 | 1.5E-25 | DF 8 | emx2 |
| fosb6 | 7.4E-30 | 0.6779701 | 0.776 | 0.691 | 2.39E-25 | DF 8 | fosb |
| dlx4a6 | 7.6E-30 | 1.0927361 | 0.329 | 0.191 | 2.47E-25 | DF 8 | dlx4a |
| tnc8 | 1E-29 | 0.5731082 | 0.646 | 0.467 | 3.37E-25 | DF 8 | tnc |
| tmtc15 | 1.2E-29 | 1.1896237 | 0.233 | 0.115 | 4E-25 | DF 8 | tmtc1 |
| CABZ01092746.17 | 1.3E-29 | 0.3741387 | 0.863 | 0.707 | 4.27E-25 | DF 8 | CABZ01092746.1 |
| rac3b4 | 2.2E-29 | 1.3819621 | 0.184 | 0.081 | 7.06E-25 | DF 8 | rac3b |
| asip2b3 | 2.3E-29 | 0.6605894 | 0.265 | 0.133 | 7.63E-25 | DF 8 | asip2b |
| pbxip1a3 | 4.2E-29 | 1.0749698 | 0.322 | 0.185 | 1.38E-24 | DF 8 | pbxip1a |
| ccdc1254 | 4.3E-29 | 1.3025817 | 0.173 | 0.073 | 1.39E-24 | DF 8 | ccdc125 |
| CABZ01068358.15 | 5.5E-29 | 1.0054728 | 0.347 | 0.206 | 1.8E-24 | DF 8 | CABZ01068358.1 |
| tmem200b5 | 9.1E-29 | 1.2347768 | 0.201 | 0.093 | 2.96E-24 | DF 8 | tmem200b |
| dhcr71 | 1.5E-28 | 1.5776449 | 0.128 | 0.048 | 4.95E-24 | DF 8 | dhcr7 |
| myzap | 2.2E-28 | 1.6437094 | 0.103 | 0.034 | 7.16E-24 | DF 8 | myzap |
| map2k65 | 3E-28 | 1.0366147 | 0.321 | 0.188 | 9.66E-24 | DF 8 | map2k6 |
| cgnl14 | 3E-28 | 1.2517998 | 0.167 | 0.071 | 9.8E-24 | DF 8 | cgnl1 |
| cxcl8a4 | 3.5E-28 | 0.4142016 | 0.432 | 0.263 | 1.14E-23 | DF 8 | cxcl8a |
| ildr21 | 4E-28 | 1.5101223 | 0.148 | 0.06 | 1.29E-23 | DF 8 | ildr2 |
| ITLN16 | 1.6E-27 | 0.9447985 | 0.348 | 0.209 | 5.33E-23 | DF 8 | ITLN1 |
| plpp2a | 1.8E-27 | 1.5951812 | 0.123 | 0.046 | 5.88E-23 | DF 8 | plpp2a |
| pln1 | 2.2E-27 | 0.6665657 | 0.118 | 0.042 | 7.23E-23 | DF 8 | pln |
| phyhd1 | 3.6E-27 | 1.5864735 | 0.122 | 0.046 | 1.17E-22 | DF 8 | phyhd1 |
| suco7 | 7.6E-27 | 1.0248509 | 0.3 | 0.171 | 2.46E-22 | DF 8 | suco |
| sc:d2173 | 8.7E-27 | 1.1484728 | 0.253 | 0.134 | 2.82E-22 | DF 8 | sc:d217 |
| zgc:101744 | 9.5E-27 | 1.572332 | 0.128 | 0.049 | 3.09E-22 | DF 8 | zgc:101744 |
| si:ch1073-291c23.27 | 1.2E-26 | 0.6114434 | 0.614 | 0.472 | 3.82E-22 | DF 8 | si:ch1073-291c23.2 |
| dock9b5 | 3.1E-26 | 1.0600824 | 0.204 | 0.098 | 9.95E-22 | DF 8 | dock9b |
| b2m6 | 3.7E-26 | 0.5505464 | 0.78 | 0.685 | 1.2E-21 | DF 8 | b2m |
| gpc6a5 | 4.9E-26 | 0.9704208 | 0.332 | 0.201 | 1.58E-21 | DF 8 | gpc6a |
| NPAS35 | 7.8E-26 | 1.2231256 | 0.183 | 0.084 | 2.53E-21 | DF 8 | NPAS3 |
| si:ch73-86n18.16 | 1.1E-25 | 0.6582571 | 0.281 | 0.152 | 3.58E-21 | DF 8 | si:ch73-86n18.1 |
| axin25 | 1.6E-25 | 0.8422047 | 0.358 | 0.218 | 5.23E-21 | DF 8 | axin2 |
| ucp27 | 2.3E-25 | 0.7330477 | 0.559 | 0.413 | 7.33E-21 | DF 8 | ucp2 |
| si:dkey-178e17.31 | 3.3E-25 | 1.5073879 | 0.123 | 0.048 | 1.07E-20 | DF 8 | si:dkey-178e17.3 |
| ccdc80l23 | 3.7E-25 | 0.5238556 | 0.229 | 0.112 | 1.19E-20 | DF 8 | ccdc80l2 |
| hyal2b7 | 3.9E-25 | 0.9888697 | 0.322 | 0.195 | 1.28E-20 | DF 8 | hyal2b |
| twsg1a5 | 5.2E-25 | 0.8649676 | 0.389 | 0.256 | 1.69E-20 | DF 8 | twsg1a |
| alx4a3 | 5.6E-25 | 0.5076213 | 0.531 | 0.368 | 1.81E-20 | DF 8 | alx4a |
| creb3l24 | 8.2E-25 | 0.6919909 | 0.549 | 0.425 | 2.68E-20 | DF 8 | creb3l2 |
| si:ch73-366l1.55 | 1.5E-24 | 1.309625 | 0.174 | 0.081 | 4.99E-20 | DF 8 | si:ch73-366l1.5 |
| si:ch211-57n23.16 | 2.3E-24 | 0.9271876 | 0.346 | 0.216 | 7.4E-20 | DF 8 | si:ch211-57n23.1 |
| fbp1a5 | 3E-24 | 0.7988991 | 0.411 | 0.275 | 9.68E-20 | DF 8 | fbp1a |
| lpl5 | 6.7E-24 | 1.0016355 | 0.218 | 0.111 | 2.18E-19 | DF 8 | lpl |
| psd25 | 9.2E-24 | 1.1820331 | 0.194 | 0.096 | 2.98E-19 | DF 8 | psd2 |
| gas1a6 | 9.8E-24 | 0.9110506 | 0.278 | 0.158 | 3.17E-19 | DF 8 | gas1a |
| arl6ip5b4 | 9.9E-24 | 0.6027343 | 0.627 | 0.508 | 3.22E-19 | DF 8 | arl6ip5b |
| tpm24 | 1.7E-23 | 1.1062286 | 0.22 | 0.115 | 5.47E-19 | DF 8 | tpm2 |

| | | | | | | | |
|---|---|---|---|---|---|---|---|
| porb6 | 2E-23 | 1.2723701 | 0.207 | 0.107 | 6.66E-19 | DF 8 | porb |
| ghrb6 | 3.4E-23 | 1.1495873 | 0.208 | 0.107 | 1.11E-18 | DF 8 | ghrb |
| sfxn5b6 | 3.8E-23 | 1.0754042 | 0.246 | 0.136 | 1.23E-18 | DF 8 | sfxn5b |
| CU984600.15 | 6.5E-23 | 0.9934399 | 0.284 | 0.168 | 2.12E-18 | DF 8 | CU984600.1 |
| cast3 | 7.1E-23 | 0.4676568 | 0.698 | 0.589 | 2.32E-18 | DF 8 | cast |
| tpst15 | 7.4E-23 | 0.9803188 | 0.293 | 0.174 | 2.41E-18 | DF 8 | tpst1 |
| xylt12 | 1E-22 | 1.7964393 | 0.117 | 0.047 | 3.39E-18 | DF 8 | xylt1 |
| ccdc80l16 | 1.5E-22 | 1.1437056 | 0.166 | 0.078 | 4.96E-18 | DF 8 | ccdc80l1 |
| slc43a3b3 | 1.9E-22 | 1.2522842 | 0.165 | 0.078 | 6.25E-18 | DF 8 | slc43a3b |
| adam10a4 | 2E-22 | 0.5829298 | 0.515 | 0.379 | 6.66E-18 | DF 8 | adam10a |
| cldn19 | 2.1E-22 | 1.52214 | 0.116 | 0.046 | 6.72E-18 | DF 8 | cldn19 |
| hmgb1a5 | 2.7E-22 | 0.5458621 | 0.668 | 0.563 | 8.91E-18 | DF 8 | hmgb1a |
| actb24 | 9.9E-22 | 0.2001948 | 1 | 0.991 | 3.22E-17 | DF 8 | actb2 |
| KCTD155 | 1.3E-21 | 0.6799021 | 0.473 | 0.34 | 4.31E-17 | DF 8 | KCTD15 |
| ENSDARG000000713. | 1.4E-21 | 1.2931932 | 0.173 | 0.085 | 4.67E-17 | DF 8 | ENSDARG00000071 |
| itgb27 | 1.4E-21 | 1.1569573 | 0.201 | 0.104 | 4.68E-17 | DF 8 | itgb2 |
| ugt5c21 | 1.8E-21 | 1.4302566 | 0.114 | 0.046 | 5.83E-17 | DF 8 | ugt5c2 |
| rpz55 | 3E-21 | 0.7336997 | 0.434 | 0.305 | 9.68E-17 | DF 8 | rpz5 |
| svila6 | 3.3E-21 | 0.9667101 | 0.257 | 0.147 | 1.08E-16 | DF 8 | svila |
| snap23.2 | 4.1E-21 | 1.5578583 | 0.103 | 0.04 | 1.34E-16 | DF 8 | snap23.2 |
| pomt1 | 4.1E-21 | 1.3766568 | 0.132 | 0.058 | 1.35E-16 | DF 8 | pomt1 |
| mgst1.25 | 4.7E-21 | 0.6944223 | 0.428 | 0.299 | 1.53E-16 | DF 8 | mgst1.2 |
| hspa8b5 | 4.9E-21 | 1.1629845 | 0.183 | 0.092 | 1.6E-16 | DF 8 | hspa8b |
| ywhag18 | 5.8E-21 | 1.0623033 | 0.207 | 0.109 | 1.87E-16 | DF 8 | ywhag1 |
| crhbp2 | 7.2E-21 | 1.1384536 | 0.101 | 0.039 | 2.36E-16 | DF 8 | crhbp |
| gch16 | 7.3E-21 | 0.8958551 | 0.24 | 0.134 | 2.36E-16 | DF 8 | gch1 |
| wls5 | 7.4E-21 | 0.5957911 | 0.515 | 0.393 | 2.4E-16 | DF 8 | wls |
| mgst1.14 | 8E-21 | 0.7578044 | 0.373 | 0.249 | 2.62E-16 | DF 8 | mgst1.1 |
| ppdpfb4 | 1.7E-20 | 0.346772 | 0.842 | 0.777 | 5.64E-16 | DF 8 | ppdpfb |
| add3a5 | 2.6E-20 | 0.3529271 | 0.589 | 0.446 | 8.47E-16 | DF 8 | add3a |
| atp1b3b4 | 3.6E-20 | 1.4092302 | 0.158 | 0.078 | 1.17E-15 | DF 8 | atp1b3b |
| rnd3a7 | 4.2E-20 | 0.7064521 | 0.411 | 0.284 | 1.36E-15 | DF 8 | rnd3a |
| thrb3 | 6.1E-20 | 0.9201398 | 0.273 | 0.164 | 1.98E-15 | DF 8 | thrb |
| abcc54 | 7.9E-20 | 0.9855239 | 0.245 | 0.143 | 2.58E-15 | DF 8 | abcc5 |
| rbm54 | 9.6E-20 | 0.8494731 | 0.297 | 0.184 | 3.13E-15 | DF 8 | rbm5 |
| alpl2 | 9.9E-20 | 1.0048428 | 0.144 | 0.066 | 3.22E-15 | DF 8 | alpl |
| gpatch84 | 1E-19 | 0.6500187 | 0.453 | 0.328 | 3.39E-15 | DF 8 | gpatch8 |
| gxylt2 | 1.1E-19 | 1.3753337 | 0.121 | 0.053 | 3.43E-15 | DF 8 | gxylt2 |
| ncor11 | 1.3E-19 | 0.8054782 | 0.393 | 0.28 | 4.12E-15 | DF 8 | ncor1 |
| crhr14 | 2E-19 | 1.0370115 | 0.159 | 0.077 | 6.44E-15 | DF 8 | crhr1 |
| sfrp1b6 | 2.1E-19 | 0.3606545 | 0.514 | 0.359 | 6.83E-15 | DF 8 | sfrp1b |
| tspan18b4 | 2.6E-19 | 0.8970235 | 0.242 | 0.14 | 8.49E-15 | DF 8 | tspan18b |
| tmeff1b3 | 3E-19 | 1.175422 | 0.194 | 0.105 | 9.76E-15 | DF 8 | tmeff1b |
| ppp1r9alb1 | 3.3E-19 | 1.5043516 | 0.102 | 0.041 | 1.08E-14 | DF 8 | ppp1r9alb |
| PPM1K3 | 3.6E-19 | 1.2178893 | 0.135 | 0.062 | 1.19E-14 | DF 8 | PPM1K |
| ehd2b4 | 6E-19 | 0.2865994 | 0.692 | 0.558 | 1.96E-14 | DF 8 | ehd2b |
| aplp24 | 6.3E-19 | 0.6299695 | 0.441 | 0.321 | 2.04E-14 | DF 8 | aplp2 |
| si:dkey-7j14.53 | 7.3E-19 | 1.2006457 | 0.145 | 0.069 | 2.39E-14 | DF 8 | si:dkey-7j14.5 |

| | | | | | | | |
|---|---|---|---|---|---|---|---|
| il161 | 9.7E-19 | 1.5540175 | 0.104 | 0.044 | 3.15E-14 | DF 8 | il16 |
| itih52 | 1.3E-18 | 0.6899247 | 0.145 | 0.068 | 4.16E-14 | DF 8 | itih5 |
| tmem1768 | 1.3E-18 | 0.5235193 | 0.521 | 0.392 | 4.22E-14 | DF 8 | tmem176 |
| si:ch211-212k18.96 | 1.6E-18 | 0.8340721 | 0.283 | 0.176 | 5.22E-14 | DF 8 | si:ch211-212k18.9 |
| pamr14 | 2.3E-18 | 0.9981691 | 0.199 | 0.108 | 7.54E-14 | DF 8 | pamr1 |
| LO018340.11 | 2.3E-18 | 1.2549478 | 0.114 | 0.05 | 7.58E-14 | DF 8 | LO018340.1 |
| ttyh3b1 | 2.6E-18 | 1.1919066 | 0.119 | 0.053 | 8.5E-14 | DF 8 | ttyh3b |
| si:dkey-16p21.87 | 2.9E-18 | 0.8542333 | 0.318 | 0.208 | 9.55E-14 | DF 8 | si:dkey-16p21.8 |
| rn7sk6 | 3.2E-18 | 0.5828463 | 0.571 | 0.454 | 1.04E-13 | DF 8 | rn7sk |
| sf3b13 | 1E-17 | 0.6145693 | 0.453 | 0.336 | 3.34E-13 | DF 8 | sf3b1 |
| gas1b6 | 1.1E-17 | 0.8448879 | 0.3 | 0.194 | 3.66E-13 | DF 8 | gas1b |
| setbp14 | 1.9E-17 | 0.8241438 | 0.352 | 0.247 | 6.09E-13 | DF 8 | setbp1 |
| sesn16 | 2.8E-17 | 0.6608511 | 0.325 | 0.216 | 9.07E-13 | DF 8 | sesn1 |
| edil3a5 | 3.4E-17 | 0.4454457 | 0.56 | 0.433 | 1.1E-12 | DF 8 | edil3a |
| rnf213a4 | 4.7E-17 | 0.6787027 | 0.38 | 0.268 | 1.52E-12 | DF 8 | rnf213a |
| pthlha4 | 5.2E-17 | 1.106298 | 0.144 | 0.071 | 1.69E-12 | DF 8 | pthlha |
| gpr1435 | 5.8E-17 | 1.0945809 | 0.141 | 0.07 | 1.88E-12 | DF 8 | gpr143 |
| pgrmc13 | 5.8E-17 | 0.6477118 | 0.464 | 0.363 | 1.88E-12 | DF 8 | pgrmc1 |
| adamts15b3 | 6.8E-17 | 0.9178258 | 0.117 | 0.052 | 2.2E-12 | DF 8 | adamts15b |
| ognb5 | 7.6E-17 | 0.5414908 | 0.229 | 0.131 | 2.47E-12 | DF 8 | ognb |
| tuft1a4 | 8.5E-17 | 0.4401177 | 0.686 | 0.59 | 2.78E-12 | DF 8 | tuft1a |
| tmsb14 | 1.1E-16 | 0.2843469 | 0.581 | 0.448 | 3.63E-12 | DF 8 | tmsb1 |
| srpx25 | 1.3E-16 | 0.6962104 | 0.36 | 0.253 | 4.26E-12 | DF 8 | srpx2 |
| C7H4orf484 | 1.3E-16 | 0.7383178 | 0.345 | 0.242 | 4.27E-12 | DF 8 | C7H4orf48 |
| itgb1b.24 | 1.4E-16 | 0.9787684 | 0.195 | 0.11 | 4.41E-12 | DF 8 | itgb1b.2 |
| ebf1b1 | 1.4E-16 | 1.2221069 | 0.103 | 0.045 | 4.63E-12 | DF 8 | ebf1b |
| ARHGAP63 | 1.6E-16 | 1.0525808 | 0.131 | 0.063 | 5.24E-12 | DF 8 | ARHGAP6 |
| ablim1b7 | 1.7E-16 | 0.533139 | 0.371 | 0.254 | 5.45E-12 | DF 8 | ablim1b |
| fgl2a6 | 2E-16 | 0.6440854 | 0.258 | 0.157 | 6.61E-12 | DF 8 | fgl2a |
| cbsb7 | 2.1E-16 | 0.8658292 | 0.23 | 0.138 | 6.68E-12 | DF 8 | cbsb |
| cwc274 | 2.4E-16 | 1.0048216 | 0.208 | 0.122 | 7.93E-12 | DF 8 | cwc27 |
| myh104 | 3.3E-16 | 0.567984 | 0.443 | 0.334 | 1.07E-11 | DF 8 | myh10 |
| arid5b5 | 3.4E-16 | 0.8561728 | 0.224 | 0.134 | 1.1E-11 | DF 8 | arid5b |
| asah1b4 | 4.2E-16 | 0.8157941 | 0.282 | 0.184 | 1.38E-11 | DF 8 | asah1b |
| laptm4a4 | 5.9E-16 | 0.6735606 | 0.407 | 0.305 | 1.92E-11 | DF 8 | laptm4a |
| ece2b3 | 6.7E-16 | 1.1305943 | 0.178 | 0.1 | 2.16E-11 | DF 8 | ece2b |
| rfng | 8.9E-16 | 1.2596266 | 0.126 | 0.061 | 2.89E-11 | DF 8 | rfng |
| fstl36 | 1E-15 | 0.483117 | 0.441 | 0.32 | 3.41E-11 | DF 8 | fstl3 |
| scara56 | 1.2E-15 | 0.7745627 | 0.245 | 0.151 | 4.02E-11 | DF 8 | scara5 |
| cst34 | 1.4E-15 | 0.3141214 | 0.796 | 0.723 | 4.62E-11 | DF 8 | cst3 |
| c1qtnf14 | 1.9E-15 | 1.0501116 | 0.145 | 0.074 | 6.2E-11 | DF 8 | c1qtnf1 |
| pdlim5a1 | 1.9E-15 | 1.2084112 | 0.113 | 0.053 | 6.21E-11 | DF 8 | pdlim5a |
| ak46 | 2.1E-15 | 0.8439674 | 0.278 | 0.184 | 6.73E-11 | DF 8 | ak4 |
| smc34 | 2.6E-15 | 0.8758216 | 0.235 | 0.145 | 8.61E-11 | DF 8 | smc3 |
| tnfrsf195 | 2.8E-15 | 0.8236818 | 0.19 | 0.107 | 9.25E-11 | DF 8 | tnfrsf19 |
| BX663503.14 | 3.6E-15 | 1.1236239 | 0.131 | 0.066 | 1.18E-10 | DF 8 | BX663503.1 |
| calm3a.12 | 4.1E-15 | 0.6458986 | 0.486 | 0.397 | 1.34E-10 | DF 8 | calm3a.1 |
| CABZ01112387.1 | 4.9E-15 | 1.2549176 | 0.109 | 0.051 | 1.59E-10 | DF 8 | CABZ01112387.1 |

| | | | | | | | |
|---|---|---|---|---|---|---|---|
| podn5 | 6.5E-15 | 0.6283482 | 0.268 | 0.171 | 2.1E-10 | DF 8 | podn |
| h2ax16 | 6.6E-15 | 0.3787076 | 0.71 | 0.627 | 2.15E-10 | DF 8 | h2ax1 |
| reep53 | 6.9E-15 | 0.4517509 | 0.629 | 0.54 | 2.24E-10 | DF 8 | reep5 |
| zfp36l1a6 | 7.4E-15 | 0.6365554 | 0.306 | 0.205 | 2.41E-10 | DF 8 | zfp36l1a |
| zbtb204 | 7.8E-15 | 0.53168 | 0.475 | 0.374 | 2.54E-10 | DF 8 | zbtb20 |
| si:dkey-9i23.16 | 8.3E-15 | 1.2947893 | 0.111 | 0.053 | 2.69E-10 | DF 8 | si:dkey-9i23.16 |
| cyp2aa71 | 8.7E-15 | 1.0967173 | 0.116 | 0.056 | 2.83E-10 | DF 8 | cyp2aa7 |
| gnsa2 | 9.8E-15 | 0.9149891 | 0.248 | 0.161 | 3.18E-10 | DF 8 | gnsa |
| cdk61 | 1.1E-14 | 1.1506771 | 0.129 | 0.065 | 3.47E-10 | DF 8 | cdk6 |
| nfia4 | 1.1E-14 | 0.6647596 | 0.265 | 0.169 | 3.72E-10 | DF 8 | nfia |
| tob1a5 | 1.8E-14 | 0.3844805 | 0.703 | 0.608 | 5.7E-10 | DF 8 | tob1a |
| CR759885.14 | 3.6E-14 | 0.9221165 | 0.175 | 0.1 | 1.16E-09 | DF 8 | CR759885.1 |
| ehbp14 | 3.6E-14 | 0.993695 | 0.186 | 0.11 | 1.17E-09 | DF 8 | ehbp1 |
| dhrs13a.12 | 5.3E-14 | 1.1212306 | 0.174 | 0.102 | 1.71E-09 | DF 8 | dhrs13a.1 |
| atf5b3 | 5.3E-14 | 0.6208969 | 0.35 | 0.252 | 1.74E-09 | DF 8 | atf5b |
| ly752 | 5.6E-14 | 1.0676772 | 0.134 | 0.069 | 1.83E-09 | DF 8 | ly75 |
| heyl5 | 8.3E-14 | 1.0785631 | 0.136 | 0.071 | 2.71E-09 | DF 8 | heyl |
| wsb12 | 9.6E-14 | 0.8798981 | 0.272 | 0.187 | 3.11E-09 | DF 8 | wsb1 |
| celf28 | 1.2E-13 | 0.5127829 | 0.297 | 0.197 | 3.96E-09 | DF 8 | celf2 |
| sh3pxd2ab1 | 1.3E-13 | 1.0437722 | 0.144 | 0.078 | 4.16E-09 | DF 8 | sh3pxd2ab |
| zgc:1530314 | 1.3E-13 | 1.0879714 | 0.121 | 0.061 | 4.29E-09 | DF 8 | zgc:153031 |
| si:dkeyp-23e4.37 | 1.6E-13 | 0.8254203 | 0.155 | 0.085 | 5.24E-09 | DF 8 | si:dkeyp-23e4.3 |
| mhc1zba5 | 1.7E-13 | 0.3489042 | 0.666 | 0.583 | 5.52E-09 | DF 8 | mhc1zba |
| sept7a3 | 1.9E-13 | 0.606837 | 0.446 | 0.36 | 6.09E-09 | DF 8 | sept7a |
| scinlb5 | 2.1E-13 | 0.6041152 | 0.402 | 0.307 | 6.79E-09 | DF 8 | scinlb |
| ubap2a2 | 2.2E-13 | 0.8023309 | 0.282 | 0.193 | 7.1E-09 | DF 8 | ubap2a |
| si:ch211-212k18.84 | 2.6E-13 | 0.8313577 | 0.222 | 0.141 | 8.36E-09 | DF 8 | si:ch211-212k18.8 |
| csnk1da5 | 2.6E-13 | 0.6808844 | 0.303 | 0.211 | 8.57E-09 | DF 8 | csnk1da |
| clstn15 | 2.9E-13 | 0.7665189 | 0.217 | 0.135 | 9.57E-09 | DF 8 | clstn1 |
| adgrl2a7 | 3.1E-13 | 0.6058374 | 0.282 | 0.189 | 9.99E-09 | DF 8 | adgrl2a |
| hpgd4 | 3.2E-13 | 0.8772317 | 0.112 | 0.055 | 1.03E-08 | DF 8 | hpgd |
| rmdn14 | 3.9E-13 | 1.0306839 | 0.162 | 0.093 | 1.28E-08 | DF 8 | rmdn1 |
| hist2h2l6 | 4.1E-13 | 0.5387659 | 0.466 | 0.374 | 1.32E-08 | DF 8 | hist2h2l |
| park73 | 4.1E-13 | 0.9479141 | 0.229 | 0.15 | 1.34E-08 | DF 8 | park7 |
| emilin2a4 | 4.3E-13 | 0.6664991 | 0.198 | 0.119 | 1.39E-08 | DF 8 | emilin2a |
| btg25 | 4.6E-13 | 0.4110658 | 0.756 | 0.699 | 1.49E-08 | DF 8 | btg2 |
| glula6 | 4.9E-13 | 0.2157405 | 0.751 | 0.617 | 1.61E-08 | DF 8 | glula |
| galnt12 | 5.3E-13 | 0.7338701 | 0.247 | 0.163 | 1.72E-08 | DF 8 | galnt1 |
| adamts121 | 5.4E-13 | 1.3778782 | 0.107 | 0.053 | 1.77E-08 | DF 8 | adamts12 |
| ubtf3 | 5.7E-13 | 0.5863383 | 0.438 | 0.348 | 1.87E-08 | DF 8 | ubtf |
| bco2a3 | 5.8E-13 | 0.6416381 | 0.233 | 0.148 | 1.89E-08 | DF 8 | bco2a |
| nova25 | 6.6E-13 | 0.3240576 | 0.32 | 0.213 | 2.13E-08 | DF 8 | nova2 |
| atp2b44 | 7.3E-13 | 0.5260249 | 0.443 | 0.355 | 2.39E-08 | DF 8 | atp2b4 |
| zgc:1135317 | 7.6E-13 | 0.553237 | 0.376 | 0.277 | 2.46E-08 | DF 8 | zgc:113531 |
| stat5a | 8.1E-13 | 1.1864496 | 0.116 | 0.059 | 2.64E-08 | DF 8 | stat5a |
| DDX173 | 8.3E-13 | 0.5641947 | 0.395 | 0.299 | 2.7E-08 | DF 8 | DDX17 |
| sypl2a2 | 8.3E-13 | 0.7874183 | 0.246 | 0.164 | 2.7E-08 | DF 8 | sypl2a |
| zbtb44 | 9E-13 | 0.6065878 | 0.415 | 0.324 | 2.94E-08 | DF 8 | zbtb4 |

| | | | | | | | |
|---|---|---|---|---|---|---|---|
| rgn1 | 1.2E-12 | 1.1004697 | 0.132 | 0.072 | 3.84E-08 | DF 8 | rgn |
| top15 | 1.3E-12 | 0.9794829 | 0.146 | 0.081 | 4.39E-08 | DF 8 | top1 |
| actb13 | 1.4E-12 | 0.2251164 | 0.916 | 0.866 | 4.58E-08 | DF 8 | actb1 |
| gpt2l5 | 1.4E-12 | 0.7998993 | 0.207 | 0.129 | 4.62E-08 | DF 8 | gpt2l |
| ndrg23 | 2.4E-12 | 0.656806 | 0.409 | 0.322 | 7.77E-08 | DF 8 | ndrg2 |
| dbt | 2.9E-12 | 1.1728471 | 0.102 | 0.051 | 9.37E-08 | DF 8 | dbt |
| thraa1 | 3.4E-12 | 1.1551439 | 0.116 | 0.06 | 1.12E-07 | DF 8 | thraa |
| arhgdia4 | 3.5E-12 | 0.5324839 | 0.493 | 0.411 | 1.13E-07 | DF 8 | arhgdia |
| dhrs95 | 3.6E-12 | 0.8587702 | 0.215 | 0.14 | 1.18E-07 | DF 8 | dhrs9 |
| fgfr26 | 3.8E-12 | 0.4534949 | 0.442 | 0.348 | 1.25E-07 | DF 8 | fgfr2 |
| dpysl2b4 | 4.4E-12 | 0.6742401 | 0.242 | 0.161 | 1.42E-07 | DF 8 | dpysl2b |
| tmem106ba3 | 5.4E-12 | 0.7344614 | 0.27 | 0.189 | 1.74E-07 | DF 8 | tmem106ba |
| ddx65 | 6.1E-12 | 0.5502636 | 0.434 | 0.347 | 1.99E-07 | DF 8 | ddx6 |
| kctd13 | 6.4E-12 | 1.0133388 | 0.139 | 0.079 | 2.07E-07 | DF 8 | kctd1 |
| si:ch73-138n13.14 | 7.7E-12 | 0.4398762 | 0.473 | 0.376 | 2.52E-07 | DF 8 | si:ch73-138n13.1 |
| zgc:921406 | 8.4E-12 | 0.9881437 | 0.183 | 0.113 | 2.73E-07 | DF 8 | zgc:92140 |
| gcdha2 | 1.2E-11 | 0.9280279 | 0.175 | 0.108 | 4.01E-07 | DF 8 | gcdha |
| klf7b6 | 1.4E-11 | 0.5413855 | 0.318 | 0.227 | 4.56E-07 | DF 8 | klf7b |
| cx35.43 | 1.5E-11 | 1.0894589 | 0.135 | 0.076 | 4.75E-07 | DF 8 | cx35.4 |
| si:ch211-196l7.46 | 1.7E-11 | 0.4353754 | 0.557 | 0.475 | 5.45E-07 | DF 8 | si:ch211-196l7.4 |
| selenbp14 | 1.9E-11 | 0.9837732 | 0.137 | 0.077 | 6.15E-07 | DF 8 | selenbp1 |
| enah4 | 2.1E-11 | 0.7527436 | 0.185 | 0.115 | 6.82E-07 | DF 8 | enah |
| plxna34 | 2.3E-11 | 1.0037039 | 0.149 | 0.088 | 7.48E-07 | DF 8 | plxna3 |
| synm5 | 2.4E-11 | 0.4701736 | 0.322 | 0.229 | 7.7E-07 | DF 8 | synm |
| ktn13 | 2.7E-11 | 0.5980879 | 0.356 | 0.272 | 8.63E-07 | DF 8 | ktn1 |
| flna5 | 3.2E-11 | 0.3531574 | 0.633 | 0.542 | 1.04E-06 | DF 8 | flna |
| klf157 | 3.3E-11 | 0.6196551 | 0.293 | 0.207 | 1.07E-06 | DF 8 | klf15 |
| GNAZ3 | 4.2E-11 | 0.6937508 | 0.209 | 0.134 | 1.36E-06 | DF 8 | GNAZ |
| znf609a4 | 5.8E-11 | 0.7231036 | 0.251 | 0.174 | 1.9E-06 | DF 8 | znf609a |
| kyat31 | 7.5E-11 | 0.8174821 | 0.191 | 0.122 | 2.44E-06 | DF 8 | kyat3 |
| nsd1b3 | 8.3E-11 | 0.8192527 | 0.13 | 0.073 | 2.69E-06 | DF 8 | nsd1b |
| iscub4 | 8.6E-11 | 0.6183811 | 0.272 | 0.192 | 2.79E-06 | DF 8 | iscub |
| opcml | 1.1E-10 | 0.975584 | 0.114 | 0.062 | 3.58E-06 | DF 8 | opcml |
| ldhba5 | 1.1E-10 | 0.4038774 | 0.563 | 0.493 | 3.65E-06 | DF 8 | ldhba |
| efnb3b5 | 1.1E-10 | 0.8062542 | 0.208 | 0.138 | 3.66E-06 | DF 8 | efnb3b |
| sik17 | 1.1E-10 | 0.4345242 | 0.34 | 0.252 | 3.69E-06 | DF 8 | sik1 |
| myo1ca5 | 1.2E-10 | 0.6136684 | 0.284 | 0.202 | 3.93E-06 | DF 8 | myo1ca |
| lrrc156 | 1.3E-10 | 0.660624 | 0.205 | 0.134 | 4.25E-06 | DF 8 | lrrc15 |
| MDFI5 | 1.6E-10 | 0.5410906 | 0.321 | 0.238 | 5.31E-06 | DF 8 | MDFI |
| HACD43 | 1.8E-10 | 0.7940082 | 0.202 | 0.134 | 5.74E-06 | DF 8 | HACD4 |
| si:ch211-194b1.13 | 1.9E-10 | 0.6771365 | 0.294 | 0.215 | 6.28E-06 | DF 8 | si:ch211-194b1.1 |
| rtn1a2 | 1.9E-10 | 0.5844093 | 0.302 | 0.222 | 6.32E-06 | DF 8 | rtn1a |
| inka1b5 | 2E-10 | 0.876555 | 0.152 | 0.091 | 6.43E-06 | DF 8 | inka1b |
| boc6 | 2.6E-10 | 0.5242966 | 0.448 | 0.372 | 8.57E-06 | DF 8 | boc |
| col6a34 | 2.7E-10 | 0.5547963 | 0.318 | 0.237 | 8.94E-06 | DF 8 | col6a3 |
| abat3 | 3.2E-10 | 0.8358833 | 0.16 | 0.099 | 1.04E-05 | DF 8 | abat |
| mab21l24 | 3.3E-10 | 0.4446089 | 0.414 | 0.33 | 1.09E-05 | DF 8 | mab21l2 |
| egfra3 | 3.4E-10 | 0.5154543 | 0.339 | 0.258 | 1.09E-05 | DF 8 | egfra |

| | | | | | | | |
|---|---|---|---|---|---|---|---|
| ehbp1l1a5 | 3.4E-10 | 0.8712666 | 0.154 | 0.093 | 1.1E-05 | DF 8 | ehbp1l1a |
| zgc:1140466 | 3.8E-10 | 0.8855568 | 0.162 | 0.099 | 1.25E-05 | DF 8 | zgc:114046 |
| arl6ip14 | 4E-10 | 0.7502451 | 0.23 | 0.16 | 1.31E-05 | DF 8 | arl6ip1 |
| tbx153 | 4.1E-10 | 0.8314148 | 0.116 | 0.064 | 1.32E-05 | DF 8 | tbx15 |
| actn15 | 4.1E-10 | 0.4531944 | 0.402 | 0.317 | 1.34E-05 | DF 8 | actn1 |
| phldb2b5 | 4.2E-10 | 0.6865918 | 0.214 | 0.143 | 1.35E-05 | DF 8 | phldb2b |
| jmjd1cb3 | 4.9E-10 | 0.5835941 | 0.397 | 0.318 | 1.58E-05 | DF 8 | jmjd1cb |
| dolpp12 | 5E-10 | 0.8903554 | 0.157 | 0.098 | 1.63E-05 | DF 8 | dolpp1 |
| glulb6 | 5.3E-10 | 0.2110358 | 0.669 | 0.568 | 1.73E-05 | DF 8 | glulb |
| colec125 | 6.5E-10 | 0.3540494 | 0.522 | 0.438 | 2.13E-05 | DF 8 | colec12 |
| bco2l5 | 7.6E-10 | 0.7563645 | 0.189 | 0.124 | 2.48E-05 | DF 8 | bco2l |
| slc48a1a2 | 8.2E-10 | 0.8551189 | 0.146 | 0.089 | 2.68E-05 | DF 8 | slc48a1a |
| ash1l1 | 9.7E-10 | 0.4355531 | 0.48 | 0.407 | 3.16E-05 | DF 8 | ash1l |
| napepld | 1E-09 | 1.1237026 | 0.108 | 0.06 | 3.3E-05 | DF 8 | napepld |
| metrn6 | 1.3E-09 | 0.7599144 | 0.18 | 0.116 | 4.19E-05 | DF 8 | metrn |
| kdm5ba1 | 1.3E-09 | 0.6789335 | 0.184 | 0.119 | 4.29E-05 | DF 8 | kdm5ba |
| dchs1a2 | 1.5E-09 | 0.9287163 | 0.134 | 0.08 | 4.81E-05 | DF 8 | dchs1a |
| ptbp34 | 1.5E-09 | 0.7807585 | 0.199 | 0.134 | 4.95E-05 | DF 8 | ptbp3 |
| vasna4 | 1.5E-09 | 0.5801123 | 0.178 | 0.114 | 5.01E-05 | DF 8 | vasna |
| cavin2b6 | 1.7E-09 | 0.3074788 | 0.603 | 0.53 | 5.37E-05 | DF 8 | cavin2b |
| slc44a25 | 1.7E-09 | 0.8015041 | 0.176 | 0.114 | 5.45E-05 | DF 8 | slc44a2 |
| degs15 | 1.8E-09 | 0.7881274 | 0.255 | 0.187 | 5.83E-05 | DF 8 | degs1 |
| ddr16 | 1.8E-09 | 0.7262094 | 0.143 | 0.086 | 5.87E-05 | DF 8 | ddr1 |
| cdh25 | 1.8E-09 | 0.4109607 | 0.375 | 0.292 | 5.9E-05 | DF 8 | cdh2 |
| gli2a2 | 1.9E-09 | 1.0311091 | 0.12 | 0.07 | 6.26E-05 | DF 8 | gli2a |
| rin33 | 2E-09 | 0.800137 | 0.22 | 0.153 | 6.55E-05 | DF 8 | rin3 |
| mhc1uba2 | 2.3E-09 | 0.5290377 | 0.292 | 0.214 | 7.42E-05 | DF 8 | mhc1uba |
| lgmn4 | 2.6E-09 | 0.6039341 | 0.24 | 0.171 | 8.51E-05 | DF 8 | lgmn |
| srgap34 | 2.7E-09 | 0.6198519 | 0.276 | 0.202 | 8.93E-05 | DF 8 | srgap3 |
| dlx2a1 | 2.9E-09 | 0.9609001 | 0.114 | 0.065 | 9.49E-05 | DF 8 | dlx2a |
| lsp1b7 | 2.9E-09 | 0.3920062 | 0.34 | 0.257 | 9.55E-05 | DF 8 | lsp1b |
| aff26 | 3E-09 | 0.6204163 | 0.249 | 0.178 | 9.74E-05 | DF 8 | aff2 |
| lamc35 | 3.3E-09 | 0.6156829 | 0.184 | 0.12 | 0.000107 | DF 8 | lamc3 |
| snai26 | 3.7E-09 | 0.699284 | 0.165 | 0.105 | 0.000119 | DF 8 | snai2 |
| qki22 | 4.5E-09 | 0.8388968 | 0.121 | 0.071 | 0.000145 | DF 8 | qki2 |
| map3k14a2 | 4.5E-09 | 0.8810931 | 0.127 | 0.076 | 0.000148 | DF 8 | map3k14a |
| bcat23 | 5.6E-09 | 0.9233727 | 0.141 | 0.088 | 0.000184 | DF 8 | bcat2 |
| si:ch211-110e21.35 | 5.7E-09 | 0.8840614 | 0.127 | 0.076 | 0.000186 | DF 8 | si:ch211-110e21.3 |
| top1l3 | 6.6E-09 | 0.4603912 | 0.459 | 0.393 | 0.000215 | DF 8 | top1l |
| pou3f3b4 | 6.9E-09 | 1.0529799 | 0.1 | 0.056 | 0.000224 | DF 8 | pou3f3b |
| CABZ01068367.16 | 8.3E-09 | 0.2487178 | 0.48 | 0.393 | 0.00027 | DF 8 | CABZ01068367.1 |
| pcolce2b5 | 8.4E-09 | 0.2130142 | 0.392 | 0.296 | 0.000272 | DF 8 | pcolce2b |
| en1b6 | 9.1E-09 | 0.4888083 | 0.171 | 0.109 | 0.000295 | DF 8 | en1b |
| snx174 | 1E-08 | 0.4963724 | 0.316 | 0.242 | 0.000325 | DF 8 | snx17 |
| dip2ca5 | 1.2E-08 | 0.8228937 | 0.139 | 0.086 | 0.00038 | DF 8 | dip2ca |
| cmtm34 | 1.4E-08 | 0.5682919 | 0.262 | 0.194 | 0.000448 | DF 8 | cmtm3 |
| si:ch73-25f10.61 | 1.4E-08 | 0.8390766 | 0.131 | 0.08 | 0.000454 | DF 8 | si:ch73-25f10.6 |
| cd82a6 | 1.6E-08 | 0.4113426 | 0.43 | 0.354 | 0.000528 | DF 8 | cd82a |

| | | | | | | | |
|---|---|---|---|---|---|---|---|
| ier53 | 1.7E-08 | 0.5001301 | 0.524 | 0.469 | 0.000543 | DF 8 | ier5 |
| caska4 | 1.7E-08 | 0.8201039 | 0.138 | 0.086 | 0.000556 | DF 8 | caska |
| si:ch211-140m22.73 | 1.9E-08 | 0.7831333 | 0.2 | 0.14 | 0.000625 | DF 8 | si:ch211-140m22.7 |
| uchl14 | 2.3E-08 | 0.7523067 | 0.186 | 0.127 | 0.000757 | DF 8 | uchl1 |
| si:rp71-36a1.52 | 2.4E-08 | 0.8042974 | 0.138 | 0.087 | 0.00077 | DF 8 | si:rp71-36a1.5 |
| slc1a3a6 | 2.4E-08 | 0.6502941 | 0.212 | 0.149 | 0.000787 | DF 8 | slc1a3a |
| pacsin23 | 2.5E-08 | 1.0829739 | 0.108 | 0.063 | 0.000811 | DF 8 | pacsin2 |
| CU459056.14 | 3.5E-08 | 0.6354312 | 0.233 | 0.169 | 0.001151 | DF 8 | CU459056.1 |
| srrm13 | 3.7E-08 | 0.4464333 | 0.469 | 0.407 | 0.001198 | DF 8 | srrm1 |
| FQ323119.15 | 4.2E-08 | 0.6556005 | 0.159 | 0.103 | 0.001379 | DF 8 | FQ323119.1 |
| pbxip1b3 | 4.4E-08 | 0.6019819 | 0.305 | 0.241 | 0.001438 | DF 8 | pbxip1b |
| znf7062 | 4.6E-08 | 0.7981957 | 0.148 | 0.096 | 0.00148 | DF 8 | znf706 |
| antxr1c4 | 4.6E-08 | 0.2817164 | 0.6 | 0.536 | 0.001496 | DF 8 | antxr1c |
| aldh7a12 | 4.6E-08 | 0.719733 | 0.246 | 0.187 | 0.001497 | DF 8 | aldh7a1 |
| si:ch211-194e15.54 | 5.1E-08 | 0.7534471 | 0.157 | 0.103 | 0.001648 | DF 8 | si:ch211-194e15.5 |
| mta13 | 5.6E-08 | 0.8613272 | 0.14 | 0.09 | 0.001812 | DF 8 | mta1 |
| ndfip22 | 6E-08 | 0.5784399 | 0.35 | 0.286 | 0.001963 | DF 8 | ndfip2 |
| prickle1b6 | 6.1E-08 | 0.538805 | 0.242 | 0.178 | 0.001983 | DF 8 | prickle1b |
| dpyda.13 | 6.6E-08 | 0.67031 | 0.166 | 0.111 | 0.002161 | DF 8 | dpyda.1 |
| nr2f24 | 6.8E-08 | 0.6445767 | 0.172 | 0.115 | 0.002196 | DF 8 | nr2f2 |
| myca6 | 6.9E-08 | 0.7633 | 0.218 | 0.157 | 0.002238 | DF 8 | myca |
| glo11 | 6.9E-08 | 0.4555281 | 0.432 | 0.373 | 0.002241 | DF 8 | glo1 |
| lama44 | 7.3E-08 | 0.4116352 | 0.341 | 0.27 | 0.002374 | DF 8 | lama4 |
| slka2 | 7.5E-08 | 0.4658348 | 0.368 | 0.303 | 0.002443 | DF 8 | slka |
| lamp23 | 8.6E-08 | 0.4918338 | 0.409 | 0.349 | 0.002793 | DF 8 | lamp2 |
| si:ch73-386h18.17 | 8.9E-08 | 0.5426041 | 0.209 | 0.147 | 0.002882 | DF 8 | si:ch73-386h18.1 |
| sod3b5 | 9.2E-08 | 0.5803039 | 0.182 | 0.123 | 0.002979 | DF 8 | sod3b |
| u2surp5 | 9.4E-08 | 0.405631 | 0.385 | 0.313 | 0.003045 | DF 8 | u2surp |
| prkar2aa2 | 9.8E-08 | 0.5948016 | 0.222 | 0.162 | 0.003202 | DF 8 | prkar2aa |
| ctbp13 | 1E-07 | 0.7399738 | 0.154 | 0.101 | 0.003322 | DF 8 | ctbp1 |
| zgc:114045 | 1E-07 | 0.6883735 | 0.102 | 0.059 | 0.003385 | DF 8 | zgc:114045 |
| pgrmc22 | 1.1E-07 | 0.6701721 | 0.196 | 0.139 | 0.00351 | DF 8 | pgrmc2 |
| apbb2b | 1.1E-07 | 0.8535232 | 0.119 | 0.073 | 0.003665 | DF 8 | apbb2b |
| si:ch211-281l24.33 | 1.2E-07 | 0.5284934 | 0.286 | 0.221 | 0.003813 | DF 8 | si:ch211-281l24.3 |
| znf5322 | 1.2E-07 | 0.7969437 | 0.116 | 0.07 | 0.003982 | DF 8 | znf532 |
| mpp6b2 | 1.3E-07 | 0.8609342 | 0.1 | 0.058 | 0.004184 | DF 8 | mpp6b |
| sesn34 | 1.3E-07 | 0.8788137 | 0.16 | 0.109 | 0.004227 | DF 8 | sesn3 |
| trnau1apb1 | 1.3E-07 | 0.6890242 | 0.22 | 0.162 | 0.004256 | DF 8 | trnau1apb |
| dynll2b2 | 1.3E-07 | 0.6458522 | 0.212 | 0.154 | 0.004374 | DF 8 | dynll2b |
| fmr13 | 1.7E-07 | 0.6431227 | 0.212 | 0.153 | 0.005409 | DF 8 | fmr1 |
| ctbp2a4 | 2.1E-07 | 0.5259307 | 0.294 | 0.23 | 0.006714 | DF 8 | ctbp2a |
| mylipb3 | 2.1E-07 | 0.8582001 | 0.126 | 0.079 | 0.006737 | DF 8 | mylipb |
| si:ch211-207e14.42 | 2.3E-07 | 0.2798143 | 0.114 | 0.069 | 0.007551 | DF 8 | si:ch211-207e14.4 |
| mettl7a.11 | 2.4E-07 | 0.931803 | 0.108 | 0.066 | 0.007944 | DF 8 | mettl7a.1 |
| dyrk1b1 | 2.5E-07 | 0.7479177 | 0.141 | 0.093 | 0.008276 | DF 8 | dyrk1b |
| pde3b4 | 2.6E-07 | 0.7764508 | 0.173 | 0.12 | 0.008399 | DF 8 | pde3b |
| apmap3 | 2.6E-07 | 0.9834073 | 0.125 | 0.08 | 0.008571 | DF 8 | apmap |
| sult6b15 | 2.7E-07 | 0.5811616 | 0.238 | 0.177 | 0.008683 | DF 8 | sult6b1 |

| | | | | | | | |
|---|---|---|---|---|---|---|---|
| ptprfb2 | 2.7E-07 | 0.4745314 | 0.318 | 0.254 | 0.008722 | DF 8 | ptprfb |
| kank46 | 2.8E-07 | 0.7263161 | 0.154 | 0.103 | 0.009157 | DF 8 | kank4 |
| fabp37 | 3.1E-07 | 0.304592 | 0.602 | 0.53 | 0.00999 | DF 8 | fabp3 |
| rab11fip1a2 | 3.2E-07 | 0.9661321 | 0.101 | 0.061 | 0.010272 | DF 8 | rab11fip1a |
| zmp:00000007115 | 3.2E-07 | 0.7589757 | 0.128 | 0.082 | 0.010501 | DF 8 | zmp:0000000711 |
| fbxo346 | 3.3E-07 | 0.6918799 | 0.138 | 0.09 | 0.010575 | DF 8 | fbxo34 |
| anp32a1 | 3.3E-07 | 0.3812001 | 0.42 | 0.356 | 0.010803 | DF 8 | anp32a |
| peli32 | 3.6E-07 | 0.7385952 | 0.109 | 0.066 | 0.011783 | DF 8 | peli3 |
| cd1645 | 3.7E-07 | 0.437737 | 0.398 | 0.337 | 0.012185 | DF 8 | cd164 |
| pgm14 | 3.8E-07 | 0.8248598 | 0.148 | 0.1 | 0.012304 | DF 8 | pgm1 |
| lpp5 | 3.8E-07 | 0.4424705 | 0.316 | 0.253 | 0.012329 | DF 8 | lpp |
| cpamd83 | 3.9E-07 | 0.6055353 | 0.101 | 0.06 | 0.012771 | DF 8 | cpamd8 |
| kank25 | 3.9E-07 | 0.4868261 | 0.275 | 0.212 | 0.012781 | DF 8 | kank2 |
| gpx4a5 | 4.1E-07 | 0.3992758 | 0.367 | 0.303 | 0.013175 | DF 8 | gpx4a |
| ylpm11 | 4.3E-07 | 0.6161288 | 0.173 | 0.12 | 0.013963 | DF 8 | ylpm1 |
| plcd4b2 | 4.4E-07 | 0.902558 | 0.126 | 0.081 | 0.014288 | DF 8 | plcd4b |
| mnta4 | 4.7E-07 | 0.6870364 | 0.152 | 0.102 | 0.015371 | DF 8 | mnta |
| CT956002.23 | 5.1E-07 | 0.7429814 | 0.12 | 0.075 | 0.016554 | DF 8 | CT956002.2 |
| rca2.13 | 5.4E-07 | 0.4890904 | 0.345 | 0.284 | 0.017448 | DF 8 | rca2.1 |
| mknk2b5 | 5.5E-07 | 0.286256 | 0.662 | 0.608 | 0.017977 | DF 8 | mknk2b |
| cemip22 | 5.9E-07 | 0.8452832 | 0.114 | 0.072 | 0.019069 | DF 8 | cemip2 |
| si:cabz01069012.24 | 6.6E-07 | 0.6068898 | 0.153 | 0.103 | 0.02135 | DF 8 | si:cabz01069012.2 |
| CABZ01074397.16 | 6.6E-07 | 0.562673 | 0.205 | 0.149 | 0.02137 | DF 8 | CABZ01074397.1 |
| tiam1a2 | 6.8E-07 | 0.855599 | 0.101 | 0.061 | 0.021959 | DF 8 | tiam1a |
| si:ch211-276i12.92 | 7.5E-07 | 0.8626053 | 0.137 | 0.091 | 0.024304 | DF 8 | si:ch211-276i12.9 |
| scarb2a4 | 7.5E-07 | 0.6085481 | 0.246 | 0.188 | 0.024317 | DF 8 | scarb2a |
| frem1b4 | 7.8E-07 | 0.7781351 | 0.121 | 0.078 | 0.025346 | DF 8 | frem1b |
| dedd16 | 7.8E-07 | 0.2916872 | 0.607 | 0.552 | 0.025447 | DF 8 | dedd1 |
| mgaa3 | 8.8E-07 | 0.5439836 | 0.195 | 0.14 | 0.028696 | DF 8 | mgaa |
| sgtb5 | 8.9E-07 | 0.6404924 | 0.176 | 0.124 | 0.028925 | DF 8 | sgtb |
| tmem119b6 | 9.7E-07 | 0.5749079 | 0.264 | 0.206 | 0.031497 | DF 8 | tmem119b |
| ptp4a3a4 | 1E-06 | 0.6860774 | 0.171 | 0.12 | 0.033061 | DF 8 | ptp4a3a |
| zgc:1585642 | 1E-06 | 0.5504958 | 0.288 | 0.232 | 0.033308 | DF 8 | zgc:158564 |
| kcne43 | 1.1E-06 | 0.555042 | 0.101 | 0.061 | 0.035393 | DF 8 | kcne4 |
| camk2b11 | 1.2E-06 | 0.8421821 | 0.101 | 0.062 | 0.038491 | DF 8 | camk2b1 |
| laptm4b1 | 1.2E-06 | 0.561981 | 0.195 | 0.142 | 0.039862 | DF 8 | laptm4b |
| axin14 | 1.2E-06 | 0.7942966 | 0.16 | 0.113 | 0.040383 | DF 8 | axin1 |
| snai1a4 | 1.3E-06 | 0.4825315 | 0.332 | 0.272 | 0.043462 | DF 8 | snai1a |
| fbn2b3 | 1.4E-06 | 0.8282425 | 0.11 | 0.069 | 0.044593 | DF 8 | fbn2b |
| auts2a5 | 1.5E-06 | 0.4493007 | 0.283 | 0.222 | 0.04752 | DF 8 | auts2a |
| acap13 | 1.6E-06 | 0.7944554 | 0.125 | 0.082 | 0.050816 | DF 8 | acap1 |
| pnn2 | 1.6E-06 | 0.4966711 | 0.294 | 0.235 | 0.052147 | DF 8 | pnn |
| lef15 | 1.7E-06 | 0.5655305 | 0.157 | 0.108 | 0.055666 | DF 8 | lef1 |
| anxa11a5 | 1.7E-06 | 0.3187084 | 0.47 | 0.41 | 0.056904 | DF 8 | anxa11a |
| ttyh3a4 | 1.9E-06 | 0.6726419 | 0.143 | 0.096 | 0.061791 | DF 8 | ttyh3a |
| adprh4 | 2E-06 | 0.8295679 | 0.122 | 0.08 | 0.063854 | DF 8 | adprh |
| prkab1b1 | 2.1E-06 | 0.4910332 | 0.332 | 0.277 | 0.068218 | DF 8 | prkab1b |
| arpp19a1 | 2.4E-06 | 0.7354193 | 0.126 | 0.083 | 0.076622 | DF 8 | arpp19a |

| | | | | | | | |
|---|---|---|---|---|---|---|---|
| grk45 | 2.8E-06 | 0.4364781 | 0.236 | 0.18 | 0.091808 | DF 8 | grk4 |
| kdm6bb3 | 3.3E-06 | 0.6016094 | 0.191 | 0.14 | 0.1065 | DF 8 | kdm6bb |
| puraa3 | 3.4E-06 | 0.4383789 | 0.309 | 0.253 | 0.1091 | DF 8 | puraa |
| diaph24 | 3.7E-06 | 0.6813729 | 0.164 | 0.116 | 0.118983 | DF 8 | diaph2 |
| appb3 | 3.9E-06 | 0.316927 | 0.521 | 0.468 | 0.126052 | DF 8 | appb |
| ENSDARG0000005970 | 3.9E-06 | 0.7091896 | 0.127 | 0.085 | 0.126327 | DF 8 | ENSDARG00000059 |
| CPNE87 | 4.1E-06 | 0.3804671 | 0.348 | 0.286 | 0.132931 | DF 8 | CPNE8 |
| sorbs15 | 4.2E-06 | 0.6177606 | 0.183 | 0.134 | 0.136273 | DF 8 | sorbs1 |
| slc16a107 | 4.2E-06 | 0.6065084 | 0.181 | 0.133 | 0.137995 | DF 8 | slc16a10 |
| mrtfab3 | 4.5E-06 | 0.5640004 | 0.129 | 0.086 | 0.145948 | DF 8 | mrtfab |
| rab8b3 | 4.5E-06 | 0.7301092 | 0.184 | 0.136 | 0.146712 | DF 8 | rab8b |
| mab21l13 | 4.5E-06 | 0.6421492 | 0.187 | 0.139 | 0.147904 | DF 8 | mab21l1 |
| si:ch211-160j14.35 | 4.9E-06 | 0.2199173 | 0.386 | 0.314 | 0.160234 | DF 8 | si:ch211-160j14.3 |
| cd82b2 | 5.6E-06 | 0.3772514 | 0.424 | 0.375 | 0.182982 | DF 8 | cd82b |
| znf609b3 | 5.7E-06 | 0.4660558 | 0.328 | 0.273 | 0.185038 | DF 8 | znf609b |
| calhm2.12 | 6.3E-06 | 0.7941268 | 0.108 | 0.07 | 0.203293 | DF 8 | calhm2.1 |
| ciarta6 | 6.4E-06 | 0.4514959 | 0.338 | 0.283 | 0.207377 | DF 8 | ciarta |
| itga94 | 6.5E-06 | 0.5579069 | 0.175 | 0.127 | 0.210265 | DF 8 | itga9 |
| gadd45ba3 | 7.2E-06 | 0.2138734 | 0.822 | 0.804 | 0.234264 | DF 8 | gadd45ba |
| rtn34 | 7.6E-06 | 0.5028651 | 0.279 | 0.227 | 0.24605 | DF 8 | rtn3 |
| rnf132 | 8.3E-06 | 0.6800218 | 0.174 | 0.128 | 0.26852 | DF 8 | rnf13 |
| top2b3 | 8.4E-06 | 0.4604809 | 0.318 | 0.265 | 0.273806 | DF 8 | top2b |
| cpxm1b2 | 8.9E-06 | 0.7104703 | 0.178 | 0.132 | 0.288398 | DF 8 | cpxm1b |
| itm2bb6 | 9.6E-06 | 0.5987018 | 0.131 | 0.089 | 0.312087 | DF 8 | itm2bb |
| si:ch211-71n6.42 | 9.7E-06 | 0.7326394 | 0.157 | 0.114 | 0.314467 | DF 8 | si:ch211-71n6.4 |
| ndfip1l3 | 1E-05 | 0.385633 | 0.386 | 0.337 | 0.331509 | DF 8 | ndfip1l |
| PAMR14 | 1E-05 | 0.5838985 | 0.165 | 0.12 | 0.33199 | DF 8 | PAMR1 |
| myofl4 | 1.1E-05 | 0.5389444 | 0.21 | 0.159 | 0.357526 | DF 8 | myofl |
| emilin1b5 | 1.3E-05 | 0.6640818 | 0.127 | 0.086 | 0.406514 | DF 8 | emilin1b |
| twist28 | 1.3E-05 | 0.2203155 | 0.409 | 0.34 | 0.409833 | DF 8 | twist2 |
| si:ch73-44m9.3.13 | 1.3E-05 | 0.6560684 | 0.14 | 0.099 | 0.417547 | DF 8 | si:ch73-44m9.3.1 |
| lman22 | 1.3E-05 | 0.4462837 | 0.287 | 0.236 | 0.421209 | DF 8 | lman2 |
| cmbl5 | 1.4E-05 | 0.5049002 | 0.226 | 0.175 | 0.446979 | DF 8 | cmbl |
| si:dkey-253d23.42 | 1.4E-05 | 0.5913199 | 0.129 | 0.088 | 0.448441 | DF 8 | si:dkey-253d23.4 |
| aldh6a14 | 1.6E-05 | 0.4634643 | 0.32 | 0.274 | 0.525506 | DF 8 | aldh6a1 |
| tmem198b4 | 1.6E-05 | 0.4579928 | 0.239 | 0.189 | 0.529914 | DF 8 | tmem198b |
| stom4 | 1.7E-05 | 0.4311768 | 0.322 | 0.27 | 0.537563 | DF 8 | stom |
| purab1 | 1.7E-05 | 0.4818631 | 0.231 | 0.179 | 0.552369 | DF 8 | purab |
| prss236 | 1.7E-05 | 0.4854391 | 0.253 | 0.2 | 0.562905 | DF 8 | prss23 |
| srsf5a4 | 1.7E-05 | 0.2302473 | 0.623 | 0.579 | 0.564941 | DF 8 | srsf5a |
| nedd4a4 | 1.8E-05 | 0.608313 | 0.131 | 0.091 | 0.591404 | DF 8 | nedd4a |
| pbx1a3 | 1.9E-05 | 0.4261513 | 0.286 | 0.232 | 0.612827 | DF 8 | pbx1a |
| fryl3 | 1.9E-05 | 0.6504266 | 0.148 | 0.106 | 0.614689 | DF 8 | fryl |
| si:ch211-271b14.1 | 2E-05 | 0.6169916 | 0.148 | 0.106 | 0.651699 | DF 8 | si:ch211-271b14.1 |
| tmem30ab3 | 2E-05 | 0.6543855 | 0.185 | 0.141 | 0.665109 | DF 8 | tmem30ab |
| nxn1 | 2.2E-05 | 0.7734215 | 0.101 | 0.066 | 0.71887 | DF 8 | nxn |
| rab133 | 2.2E-05 | 0.4459071 | 0.273 | 0.222 | 0.727906 | DF 8 | rab13 |
| slc44a1b6 | 2.2E-05 | 0.514929 | 0.165 | 0.119 | 0.731065 | DF 8 | slc44a1b |

| | | | | | | | |
|---|---|---|---|---|---|---|---|
| ap2a13 | 2.4E-05 | 0.5074069 | 0.231 | 0.184 | 0.772597 | DF 8 | ap2a1 |
| osr16 | 2.4E-05 | 0.3840492 | 0.224 | 0.172 | 0.796339 | DF 8 | osr1 |
| myo1cb2 | 2.8E-05 | 0.3733872 | 0.305 | 0.253 | 0.89686 | DF 8 | myo1cb |
| ifngr21 | 2.8E-05 | 0.7601548 | 0.103 | 0.069 | 0.91905 | DF 8 | ifngr2 |
| smarcd24 | 2.9E-05 | 0.523869 | 0.212 | 0.165 | 0.948995 | DF 8 | smarcd2 |
| nrp2a3 | 3E-05 | 0.5755627 | 0.172 | 0.128 | 0.979357 | DF 8 | nrp2a |
| acin1a2 | 3.1E-05 | 0.2021913 | 0.622 | 0.593 | 1 | DF 8 | acin1a |
| hist1h4l.185 | 3.1E-05 | 0.3772792 | 0.166 | 0.12 | 1 | DF 8 | hist1h4l.18 |
| ginm1 | 3.2E-05 | 0.4134134 | 0.382 | 0.334 | 1 | DF 8 | ginm1 |
| cab39l13 | 3.4E-05 | 0.540671 | 0.201 | 0.156 | 1 | DF 8 | cab39l1 |
| ckap52 | 3.4E-05 | 0.6624188 | 0.126 | 0.088 | 1 | DF 8 | ckap5 |
| acin1b1 | 3.5E-05 | 0.4840578 | 0.259 | 0.212 | 1 | DF 8 | acin1b |
| zc3h131 | 3.5E-05 | 0.4075444 | 0.281 | 0.234 | 1 | DF 8 | zc3h13 |
| ephx2 | 3.6E-05 | 0.7497893 | 0.111 | 0.076 | 1 | DF 8 | ephx2 |
| cacna1g4 | 3.7E-05 | 0.5686437 | 0.137 | 0.096 | 1 | DF 8 | cacna1g |
| trim3b7 | 3.9E-05 | 0.5509686 | 0.135 | 0.094 | 1 | DF 8 | trim3b |
| gdi23 | 4.1E-05 | 0.2558397 | 0.516 | 0.474 | 1 | DF 8 | gdi2 |
| si:ch211-262i1.6 | 4.1E-05 | 0.7664237 | 0.135 | 0.097 | 1 | DF 8 | si:ch211-262i1.6 |
| ubtfl1 | 4.1E-05 | 0.5614616 | 0.204 | 0.16 | 1 | DF 8 | ubtfl |
| zranb22 | 4.4E-05 | 0.5695063 | 0.173 | 0.13 | 1 | DF 8 | zranb2 |
| tln13 | 4.5E-05 | 0.2086379 | 0.655 | 0.611 | 1 | DF 8 | tln1 |
| soga13 | 4.8E-05 | 0.5054031 | 0.169 | 0.126 | 1 | DF 8 | soga1 |
| dchs1b6 | 4.9E-05 | 0.6366382 | 0.129 | 0.091 | 1 | DF 8 | dchs1b |
| appa3 | 5E-05 | 0.4249031 | 0.329 | 0.282 | 1 | DF 8 | appa |
| msra1 | 5.1E-05 | 0.7256092 | 0.12 | 0.084 | 1 | DF 8 | msra |
| atl35 | 6.1E-05 | 0.4117809 | 0.364 | 0.322 | 1 | DF 8 | atl3 |
| tspan3a3 | 6.3E-05 | 0.547776 | 0.205 | 0.162 | 1 | DF 8 | tspan3a |
| ep300b2 | 6.3E-05 | 0.4925267 | 0.174 | 0.131 | 1 | DF 8 | ep300b |
| lrig12 | 6.4E-05 | 0.6261602 | 0.111 | 0.076 | 1 | DF 8 | lrig1 |
| hhip5 | 6.7E-05 | 0.2819277 | 0.127 | 0.088 | 1 | DF 8 | hhip |
| plscr3b4 | 6.8E-05 | 0.418206 | 0.288 | 0.24 | 1 | DF 8 | plscr3b |
| spink2.14 | 6.9E-05 | 0.4666526 | 0.172 | 0.129 | 1 | DF 8 | spink2.1 |
| vegfaa6 | 7.5E-05 | 0.6746003 | 0.155 | 0.116 | 1 | DF 8 | vegfaa |
| ntd56 | 7.8E-05 | 0.5747772 | 0.187 | 0.142 | 1 | DF 8 | ntd5 |
| smek1 | 8E-05 | 0.6028629 | 0.134 | 0.096 | 1 | DF 8 | smek1 |
| ddt2 | 8.2E-05 | 0.4783479 | 0.237 | 0.193 | 1 | DF 8 | ddt |
| fkbp53 | 8.2E-05 | 0.580224 | 0.239 | 0.197 | 1 | DF 8 | fkbp5 |
| runx1t15 | 8.4E-05 | 0.6233388 | 0.128 | 0.091 | 1 | DF 8 | runx1t1 |
| ctnna11 | 8.5E-05 | 0.4829317 | 0.22 | 0.178 | 1 | DF 8 | ctnna1 |
| ada6 | 9.9E-05 | 0.3812186 | 0.268 | 0.221 | 1 | DF 8 | ada |
| slit23 | 1E-04 | 0.6504472 | 0.102 | 0.069 | 1 | DF 8 | slit2 |
| ssbp41 | 0.0001 | 0.6790228 | 0.109 | 0.075 | 1 | DF 8 | ssbp4 |
| ptprsa.14 | 0.00011 | 0.6007873 | 0.164 | 0.125 | 1 | DF 8 | ptprsa.1 |
| oga4 | 0.00012 | 0.6070599 | 0.134 | 0.097 | 1 | DF 8 | oga |
| trps17 | 0.00012 | 0.3859584 | 0.218 | 0.174 | 1 | DF 8 | trps1 |
| adh8b3 | 0.00012 | 0.366764 | 0.281 | 0.234 | 1 | DF 8 | adh8b |
| nudt3b1 | 0.00012 | 0.5783032 | 0.132 | 0.096 | 1 | DF 8 | nudt3b |
| fstl1a5 | 0.00013 | 0.2138033 | 0.148 | 0.107 | 1 | DF 8 | fstl1a |

| | | | | | | | |
|---|---|---|---|---|---|---|---|
| tmod25 | 0.00014 | 0.4453894 | 0.215 | 0.172 | 1 | DF 8 | tmod2 |
| kng14 | 0.00014 | 0.7010515 | 0.114 | 0.08 | 1 | DF 8 | kng1 |
| me32 | 0.00014 | 0.8164749 | 0.118 | 0.085 | 1 | DF 8 | me3 |
| CU457778.9 | 0.00014 | 0.6717873 | 0.104 | 0.072 | 1 | DF 8 | CU457778.9 |
| spryd3 | 0.00015 | 0.5948372 | 0.139 | 0.103 | 1 | DF 8 | spryd3 |
| rassf2a3 | 0.00016 | 0.6033964 | 0.164 | 0.126 | 1 | DF 8 | rassf2a |
| rab7a2 | 0.00018 | 0.6344431 | 0.178 | 0.141 | 1 | DF 8 | rab7a |
| sfpq2 | 0.00018 | 0.5451269 | 0.247 | 0.207 | 1 | DF 8 | sfpq |
| smarca23 | 0.00018 | 0.3567112 | 0.272 | 0.226 | 1 | DF 8 | smarca2 |
| srsf6b1 | 0.00018 | 0.5164961 | 0.154 | 0.116 | 1 | DF 8 | srsf6b |
| vkorc1l13 | 0.00018 | 0.6598831 | 0.125 | 0.09 | 1 | DF 8 | vkorc1l1 |
| bckdhbl1 | 0.00018 | 0.6136043 | 0.213 | 0.175 | 1 | DF 8 | bckdhbl |
| zeb2a6 | 0.00018 | 0.2038453 | 0.395 | 0.341 | 1 | DF 8 | zeb2a |
| cyb5r32 | 0.00018 | 0.6717499 | 0.18 | 0.142 | 1 | DF 8 | cyb5r3 |
| igf2a3 | 0.00018 | 0.268127 | 0.129 | 0.092 | 1 | DF 8 | igf2a |
| FAT41 | 0.00019 | 0.3921846 | 0.267 | 0.222 | 1 | DF 8 | FAT4 |
| tead1b2 | 0.00019 | 0.5476961 | 0.332 | 0.293 | 1 | DF 8 | tead1b |
| atrx3 | 0.00021 | 0.2997116 | 0.498 | 0.467 | 1 | DF 8 | atrx |
| c64 | 0.00021 | 0.4114912 | 0.105 | 0.073 | 1 | DF 8 | c6 |
| slc16a1b4 | 0.00021 | 0.6922378 | 0.168 | 0.131 | 1 | DF 8 | slc16a1b |
| nuak1b5 | 0.00022 | 0.3425265 | 0.325 | 0.282 | 1 | DF 8 | nuak1b |
| tril6 | 0.00022 | 0.4177792 | 0.145 | 0.107 | 1 | DF 8 | tril |
| kifap3a3 | 0.00024 | 0.5937103 | 0.127 | 0.092 | 1 | DF 8 | kifap3a |
| tcf202 | 0.00024 | 0.5577021 | 0.181 | 0.142 | 1 | DF 8 | tcf20 |
| mia33 | 0.00025 | 0.4417868 | 0.238 | 0.196 | 1 | DF 8 | mia3 |
| tmem50a5 | 0.00025 | 0.4973937 | 0.273 | 0.235 | 1 | DF 8 | tmem50a |
| sntb25 | 0.00025 | 0.3539272 | 0.282 | 0.237 | 1 | DF 8 | sntb2 |
| chd23 | 0.00028 | 0.5491821 | 0.219 | 0.179 | 1 | DF 8 | chd2 |
| erlin12 | 0.00028 | 0.62528 | 0.127 | 0.093 | 1 | DF 8 | erlin1 |
| si:ch211-286o17.16 | 0.00029 | 0.4149798 | 0.258 | 0.217 | 1 | DF 8 | si:ch211-286o17.1 |
| si:ch211-225b11.14 | 0.00029 | 0.6985188 | 0.105 | 0.075 | 1 | DF 8 | si:ch211-225b11.1 |
| ptpn31 | 0.0003 | 0.5846034 | 0.1 | 0.069 | 1 | DF 8 | ptpn3 |
| BX005309.15 | 0.00031 | 0.801626 | 0.113 | 0.082 | 1 | DF 8 | BX005309.1 |
| trioa1 | 0.00033 | 0.550823 | 0.127 | 0.093 | 1 | DF 8 | trioa |
| tspan143 | 0.00033 | 0.4886955 | 0.158 | 0.121 | 1 | DF 8 | tspan14 |
| hspg25 | 0.00034 | 0.5402406 | 0.184 | 0.146 | 1 | DF 8 | hspg2 |
| bptf1 | 0.00034 | 0.314367 | 0.223 | 0.18 | 1 | DF 8 | bptf |
| rrbp1a5 | 0.00034 | 0.3021868 | 0.46 | 0.431 | 1 | DF 8 | rrbp1a |
| fzd23 | 0.00036 | 0.6465392 | 0.132 | 0.099 | 1 | DF 8 | fzd2 |
| ms4a17a.97 | 0.00036 | 0.4154294 | 0.259 | 0.216 | 1 | DF 8 | ms4a17a.9 |
| nphp31 | 0.00037 | 0.4294502 | 0.15 | 0.114 | 1 | DF 8 | nphp3 |
| myclb5 | 0.00041 | 0.4525333 | 0.18 | 0.141 | 1 | DF 8 | myclb |
| setd31 | 0.00044 | 0.6813041 | 0.109 | 0.079 | 1 | DF 8 | setd3 |
| si:ch211-166a6.53 | 0.00048 | 0.4396146 | 0.198 | 0.158 | 1 | DF 8 | si:ch211-166a6.5 |
| crim13 | 0.00049 | 0.5086551 | 0.18 | 0.143 | 1 | DF 8 | crim1 |
| st132 | 0.0005 | 0.4994338 | 0.25 | 0.212 | 1 | DF 8 | st13 |
| tmx12 | 0.00053 | 0.6378358 | 0.153 | 0.119 | 1 | DF 8 | tmx1 |
| slc29a1a5 | 0.00057 | 0.3860628 | 0.238 | 0.198 | 1 | DF 8 | slc29a1a |

| | | | | | | | |
|---|---|---|---|---|---|---|---|
| ppp2r2d2 | 0.00058 | 0.5327618 | 0.178 | 0.142 | 1 | DF 8 | ppp2r2d |
| znf10351 | 0.00058 | 0.49143 | 0.201 | 0.163 | 1 | DF 8 | znf1035 |
| zgc:562352 | 0.00063 | 0.6003333 | 0.15 | 0.117 | 1 | DF 8 | zgc:56235 |
| hlfa4 | 0.00066 | 0.4918153 | 0.168 | 0.132 | 1 | DF 8 | hlfa |
| rhoq6 | 0.00069 | 0.3979564 | 0.129 | 0.096 | 1 | DF 8 | rhoq |
| zgc:1231053 | 0.00069 | 0.629092 | 0.105 | 0.076 | 1 | DF 8 | zgc:123105 |
| acox13 | 0.00069 | 0.5000882 | 0.136 | 0.103 | 1 | DF 8 | acox1 |
| tbcb3 | 0.00073 | 0.5208675 | 0.251 | 0.218 | 1 | DF 8 | tbcb |
| adamts15a7 | 0.00076 | 0.4358606 | 0.181 | 0.143 | 1 | DF 8 | adamts15a |
| birc22 | 0.00077 | 0.4470747 | 0.172 | 0.136 | 1 | DF 8 | birc2 |
| plekhj13 | 0.00081 | 0.4680571 | 0.156 | 0.122 | 1 | DF 8 | plekhj1 |
| PPP2CA1 | 0.00082 | 0.5721926 | 0.168 | 0.135 | 1 | DF 8 | PPP2CA |
| rnd3b6 | 0.00085 | 0.2661537 | 0.373 | 0.331 | 1 | DF 8 | rnd3b |
| ctsa3 | 0.00088 | 0.6155876 | 0.149 | 0.118 | 1 | DF 8 | ctsa |
| supt6h3 | 0.00094 | 0.3261054 | 0.236 | 0.197 | 1 | DF 8 | supt6h |
| kdm4b2 | 0.00099 | 0.453768 | 0.221 | 0.185 | 1 | DF 8 | kdm4b |
| capns1a3 | 0.00102 | 0.2982322 | 0.428 | 0.401 | 1 | DF 8 | capns1a |
| lrp1ab5 | 0.00108 | 0.2848585 | 0.382 | 0.341 | 1 | DF 8 | lrp1ab |
| kremen14 | 0.00108 | 0.5611894 | 0.109 | 0.08 | 1 | DF 8 | kremen1 |
| ogt.1.11 | 0.00108 | 0.5867033 | 0.146 | 0.113 | 1 | DF 8 | ogt.1.1 |
| kmt2ca2 | 0.00109 | 0.3868727 | 0.207 | 0.17 | 1 | DF 8 | kmt2ca |
| ddx3xa3 | 0.00111 | 0.3860022 | 0.351 | 0.324 | 1 | DF 8 | ddx3xa |
| rsrc21 | 0.00117 | 0.3655811 | 0.196 | 0.159 | 1 | DF 8 | rsrc2 |
| ehbp1l1b6 | 0.00126 | 0.4809966 | 0.16 | 0.127 | 1 | DF 8 | ehbp1l1b |
| lpgat11 | 0.00126 | 0.4386703 | 0.138 | 0.106 | 1 | DF 8 | lpgat1 |
| si:ch211-108d22.21 | 0.0013 | 0.3574987 | 0.251 | 0.213 | 1 | DF 8 | si:ch211-108d22.2 |
| lmf2b2 | 0.00132 | 0.5323557 | 0.169 | 0.137 | 1 | DF 8 | lmf2b |
| cand11 | 0.00132 | 0.4977917 | 0.147 | 0.115 | 1 | DF 8 | cand1 |
| cep1311 | 0.00136 | 0.5540651 | 0.127 | 0.097 | 1 | DF 8 | cep131 |
| cmtm64 | 0.0015 | 0.4922124 | 0.173 | 0.14 | 1 | DF 8 | cmtm6 |
| slc20a1b4 | 0.00153 | 0.5044189 | 0.324 | 0.295 | 1 | DF 8 | slc20a1b |
| mpp6a4 | 0.00154 | 0.7417089 | 0.109 | 0.082 | 1 | DF 8 | mpp6a |
| rbm25a2 | 0.00169 | 0.4861845 | 0.109 | 0.081 | 1 | DF 8 | rbm25a |
| trip10a1 | 0.00171 | 0.5010606 | 0.109 | 0.081 | 1 | DF 8 | trip10a |
| tdh5 | 0.00172 | 0.3757372 | 0.276 | 0.241 | 1 | DF 8 | tdh |
| fam120c2 | 0.00191 | 0.3332143 | 0.33 | 0.301 | 1 | DF 8 | fam120c |
| trappc53 | 0.00195 | 0.5415128 | 0.144 | 0.114 | 1 | DF 8 | trappc5 |
| cdc42l3 | 0.00199 | 0.2372273 | 0.433 | 0.405 | 1 | DF 8 | cdc42l |
| cyb5a3 | 0.00199 | 0.5462089 | 0.119 | 0.091 | 1 | DF 8 | cyb5a |
| gmfb3 | 0.002 | 0.453958 | 0.176 | 0.144 | 1 | DF 8 | gmfb |
| ppp1r12a1 | 0.00204 | 0.489331 | 0.158 | 0.127 | 1 | DF 8 | ppp1r12a |
| akr1b1.2 | 0.00206 | 0.366074 | 0.299 | 0.266 | 1 | DF 8 | akr1b1.2 |
| ankhd11 | 0.00206 | 0.4954926 | 0.192 | 0.16 | 1 | DF 8 | ankhd1 |
| si:dkey-235d18.54 | 0.00207 | 0.5387906 | 0.127 | 0.098 | 1 | DF 8 | si:dkey-235d18.5 |
| wapla1 | 0.00207 | 0.5608846 | 0.119 | 0.091 | 1 | DF 8 | wapla |
| pnisr3 | 0.00208 | 0.3551267 | 0.221 | 0.186 | 1 | DF 8 | pnisr |
| aqp11 | 0.00211 | 0.6037752 | 0.104 | 0.078 | 1 | DF 8 | aqp11 |
| zbtb16a4 | 0.00211 | 0.4903117 | 0.163 | 0.13 | 1 | DF 8 | zbtb16a |

| | | | | | | | |
|---|---|---|---|---|---|---|---|
| lrrc58b5 | 0.00216 | 0.5173883 | 0.153 | 0.122 | 1 | DF 8 | lrrc58b |
| rab104 | 0.00219 | 0.2232819 | 0.481 | 0.458 | 1 | DF 8 | rab10 |
| KIF2A3 | 0.00224 | 0.5208505 | 0.139 | 0.11 | 1 | DF 8 | KIF2A |
| wnk1b3 | 0.00226 | 0.3838743 | 0.159 | 0.127 | 1 | DF 8 | wnk1b |
| foxj33 | 0.00233 | 0.521381 | 0.173 | 0.14 | 1 | DF 8 | foxj3 |
| setd22 | 0.00237 | 0.5781995 | 0.153 | 0.123 | 1 | DF 8 | setd2 |
| gnas2 | 0.00244 | 0.3983238 | 0.187 | 0.155 | 1 | DF 8 | gnas |
| camk2g22 | 0.00245 | 0.5115455 | 0.157 | 0.127 | 1 | DF 8 | camk2g2 |
| specc1la3 | 0.00247 | 0.4734138 | 0.107 | 0.08 | 1 | DF 8 | specc1la |
| lrp54 | 0.00252 | 0.4274144 | 0.143 | 0.112 | 1 | DF 8 | lrp5 |
| flii3 | 0.00254 | 0.6176633 | 0.138 | 0.11 | 1 | DF 8 | flii |
| selenot1b2 | 0.00257 | 0.5011248 | 0.144 | 0.115 | 1 | DF 8 | selenot1b |
| hectd11 | 0.00278 | 0.3819296 | 0.241 | 0.209 | 1 | DF 8 | hectd1 |
| pdlim24 | 0.00281 | 0.4089185 | 0.189 | 0.157 | 1 | DF 8 | pdlim2 |
| foxo3b4 | 0.00281 | 0.5159256 | 0.109 | 0.082 | 1 | DF 8 | foxo3b |
| fblim15 | 0.00282 | 0.5290032 | 0.14 | 0.111 | 1 | DF 8 | fblim1 |
| rnf213b1 | 0.00283 | 0.5224067 | 0.105 | 0.079 | 1 | DF 8 | rnf213b |
| eif4a25 | 0.00287 | 0.3481616 | 0.228 | 0.193 | 1 | DF 8 | eif4a2 |
| tmem1311 | 0.00303 | 0.4419544 | 0.228 | 0.197 | 1 | DF 8 | tmem131 |
| mllt104 | 0.00304 | 0.4655308 | 0.109 | 0.082 | 1 | DF 8 | mllt10 |
| rprd2b2 | 0.00307 | 0.4594425 | 0.153 | 0.123 | 1 | DF 8 | rprd2b |
| npepps1 | 0.00309 | 0.4453613 | 0.175 | 0.144 | 1 | DF 8 | npepps |
| cmtm41 | 0.00314 | 0.4929476 | 0.145 | 0.116 | 1 | DF 8 | cmtm4 |
| tm9sf21 | 0.00317 | 0.4163123 | 0.203 | 0.171 | 1 | DF 8 | tm9sf2 |
| chd33 | 0.00332 | 0.3760974 | 0.207 | 0.174 | 1 | DF 8 | chd3 |
| rhbdf1a3 | 0.00333 | 0.6595103 | 0.121 | 0.095 | 1 | DF 8 | rhbdf1a |
| dag13 | 0.00341 | 0.4418913 | 0.189 | 0.158 | 1 | DF 8 | dag1 |
| zgc:1105914 | 0.00348 | 0.6228742 | 0.12 | 0.094 | 1 | DF 8 | zgc:110591 |
| prkd31 | 0.00357 | 0.5497434 | 0.105 | 0.08 | 1 | DF 8 | prkd3 |
| ppp2r5d2 | 0.00367 | 0.4548856 | 0.143 | 0.114 | 1 | DF 8 | ppp2r5d |
| nfe2l1b4 | 0.00379 | 0.5576031 | 0.113 | 0.087 | 1 | DF 8 | nfe2l1b |
| tjp1a3 | 0.0039 | 0.35346 | 0.186 | 0.154 | 1 | DF 8 | tjp1a |
| CABZ01113192.11 | 0.00393 | 0.4077477 | 0.164 | 0.134 | 1 | DF 8 | CABZ01113192.1 |
| sfswap1 | 0.00396 | 0.5168864 | 0.137 | 0.109 | 1 | DF 8 | sfswap |
| tcf7l1a6 | 0.00396 | 0.513767 | 0.145 | 0.117 | 1 | DF 8 | tcf7l1a |
| bmpr2a3 | 0.00408 | 0.2986275 | 0.212 | 0.178 | 1 | DF 8 | bmpr2a |
| tmem593 | 0.00409 | 0.3332395 | 0.338 | 0.314 | 1 | DF 8 | tmem59 |
| ANPEP5 | 0.0041 | 0.2533679 | 0.245 | 0.21 | 1 | DF 8 | ANPEP |
| zc3h11a1 | 0.00421 | 0.4285565 | 0.173 | 0.143 | 1 | DF 8 | zc3h11a |
| ctdsp23 | 0.00421 | 0.3259776 | 0.226 | 0.191 | 1 | DF 8 | ctdsp2 |
| usp22 | 0.00457 | 0.4677682 | 0.114 | 0.088 | 1 | DF 8 | usp22 |
| zmynd113 | 0.00471 | 0.3278645 | 0.171 | 0.139 | 1 | DF 8 | zmynd11 |
| washc2c1 | 0.00472 | 0.5288631 | 0.15 | 0.123 | 1 | DF 8 | washc2c |
| map3k71 | 0.00472 | 0.4720135 | 0.157 | 0.129 | 1 | DF 8 | map3k7 |
| phyh2 | 0.00472 | 0.5866058 | 0.117 | 0.092 | 1 | DF 8 | phyh |
| rybpb1 | 0.00477 | 0.3792952 | 0.114 | 0.088 | 1 | DF 8 | rybpb |
| proza3 | 0.00486 | 0.4145365 | 0.201 | 0.17 | 1 | DF 8 | proza |
| mmp15a6 | 0.00491 | 0.4779236 | 0.109 | 0.084 | 1 | DF 8 | mmp15a |

| | | | | | | | |
|---|---|---|---|---|---|---|---|
| mob1a | 0.00494 | 0.601541 | 0.117 | 0.092 | 1 | DF 8 | mob1a |
| med102 | 0.005 | 0.626913 | 0.11 | 0.086 | 1 | DF 8 | med10 |
| AL935174.62 | 0.00503 | 0.5792965 | 0.105 | 0.081 | 1 | DF 8 | AL935174.6 |
| ppp2r5cb1 | 0.00526 | 0.4689045 | 0.114 | 0.089 | 1 | DF 8 | ppp2r5cb |
| ccdc474 | 0.00531 | 0.6343093 | 0.164 | 0.139 | 1 | DF 8 | ccdc47 |
| c1qtnf25 | 0.00531 | 0.4560486 | 0.119 | 0.092 | 1 | DF 8 | c1qtnf2 |
| zgc:1748886 | 0.00533 | 0.2527287 | 0.253 | 0.217 | 1 | DF 8 | zgc:174888 |
| sufu1 | 0.00545 | 0.4645489 | 0.111 | 0.086 | 1 | DF 8 | sufu |
| id17 | 0.00572 | 0.2243921 | 0.645 | 0.642 | 1 | DF 8 | id1 |
| rbm61 | 0.00576 | 0.5924699 | 0.114 | 0.089 | 1 | DF 8 | rbm6 |
| CABZ01061894.12 | 0.00582 | 0.5036386 | 0.123 | 0.098 | 1 | DF 8 | CABZ01061894.1 |
| cmip2 | 0.00588 | 0.4641419 | 0.148 | 0.121 | 1 | DF 8 | cmip |
| znf3181 | 0.006 | 0.4646267 | 0.185 | 0.157 | 1 | DF 8 | znf318 |
| smo3 | 0.0061 | 0.3711488 | 0.284 | 0.258 | 1 | DF 8 | smo |
| lamc14 | 0.0062 | 0.3251579 | 0.158 | 0.129 | 1 | DF 8 | lamc1 |
| si:rp71-36a1.22 | 0.00647 | 0.6245916 | 0.111 | 0.087 | 1 | DF 8 | si:rp71-36a1.2 |
| tdrd3 | 0.00648 | 0.4869152 | 0.128 | 0.103 | 1 | DF 8 | tdrd3 |
| jag1b4 | 0.0065 | 0.261248 | 0.249 | 0.216 | 1 | DF 8 | jag1b |
| wdr332 | 0.00713 | 0.4266047 | 0.113 | 0.089 | 1 | DF 8 | wdr33 |
| rbm25b2 | 0.00731 | 0.4066822 | 0.238 | 0.21 | 1 | DF 8 | rbm25b |
| pbx45 | 0.00749 | 0.261468 | 0.291 | 0.259 | 1 | DF 8 | pbx4 |
| irf93 | 0.00772 | 0.3867594 | 0.156 | 0.129 | 1 | DF 8 | irf9 |
| mink12 | 0.00779 | 0.3831524 | 0.128 | 0.102 | 1 | DF 8 | mink1 |
| cdc42ep53 | 0.00781 | 0.2071436 | 0.33 | 0.3 | 1 | DF 8 | cdc42ep5 |
| camk2g11 | 0.00786 | 0.3327166 | 0.242 | 0.215 | 1 | DF 8 | camk2g1 |
| piezo15 | 0.00858 | 0.4972354 | 0.149 | 0.124 | 1 | DF 8 | piezo1 |
| cep2902 | 0.00863 | 0.4307973 | 0.137 | 0.112 | 1 | DF 8 | cep290 |
| usp7 | 0.00872 | 0.4352303 | 0.183 | 0.157 | 1 | DF 8 | usp7 |
| tspan73 | 0.00878 | 0.3535342 | 0.242 | 0.216 | 1 | DF 8 | tspan7 |
| paxbp11 | 0.00878 | 0.3199838 | 0.145 | 0.118 | 1 | DF 8 | paxbp1 |
| maco1a | 0.00897 | 0.569753 | 0.109 | 0.086 | 1 | DF 8 | maco1a |
| pias4a | 0.00952 | 0.4728372 | 0.103 | 0.081 | 1 | DF 8 | pias4a |
| msx1b5 | 0.00958 | 0.2427594 | 0.328 | 0.294 | 1 | DF 8 | msx1b |
| ubb6 | 0.00963 | 0.2025989 | 0.68 | 0.682 | 1 | DF 8 | ubb |
| pik3r24 | 0.00981 | 0.2892981 | 0.231 | 0.201 | 1 | DF 8 | pik3r2 |
| vwde1 | 0 | 6.5030351 | 0.814 | 0.019 | 0 | Lig/Ten | vwde |
| ucmaa2 | 0 | 6.0138196 | 0.743 | 0.036 | 0 | Lig/Ten | ucmaa |
| thbs4a4 | 0 | 4.1063742 | 0.768 | 0.073 | 0 | Lig/Ten | thbs4a |
| thbs4b7 | 0 | 3.748841 | 0.931 | 0.253 | 0 | Lig/Ten | thbs4b |
| gpm6ab4 | 0 | 3.9362352 | 0.748 | 0.082 | 0 | Lig/Ten | gpm6ab |
| prelp7 | 0 | 3.3836986 | 0.763 | 0.127 | 0 | Lig/Ten | prelp |
| tnmd4 | 0 | 3.8525259 | 0.761 | 0.146 | 0 | Lig/Ten | tnmd |
| olfml2bb4 | 0 | 3.3994717 | 0.697 | 0.085 | 0 | Lig/Ten | olfml2bb |
| ognb6 | 0 | 3.1330604 | 0.721 | 0.127 | 0 | Lig/Ten | ognb |
| matn42 | 0 | 4.8837507 | 0.6 | 0.039 | 0 | Lig/Ten | matn4 |
| col22a11 | 0 | 3.9259561 | 0.602 | 0.056 | 0 | Lig/Ten | col22a1 |
| comp | 0 | 7.8337244 | 0.535 | 0.005 | 0 | Lig/Ten | comp |
| meox14 | 0 | 3.6422836 | 0.577 | 0.058 | 0 | Lig/Ten | meox1 |

| | | | | | | | |
|---|---|---|---|---|---|---|---|
| scxa5 | 0 | 3.6857634 | 0.573 | 0.078 | 0 | Lig/Ten | scxa |
| TNC2 | 0 | 4.2711607 | 0.498 | 0.034 | 0 | Lig/Ten | TNC |
| cilp23 | 0 | 4.3108327 | 0.507 | 0.058 | 0 | Lig/Ten | cilp2 |
| chad3 | 0 | 3.7910594 | 0.48 | 0.048 | 0 | Lig/Ten | chad |
| col28a1b | 0 | 5.8622022 | 0.438 | 0.013 | 0 | Lig/Ten | col28a1b |
| si:ch211-159i8.4 | 0 | 5.0756016 | 0.425 | 0.014 | 0 | Lig/Ten | si:ch211-159i8.4 |
| itga11b | 0 | 5.3281783 | 0.361 | 0.011 | 0 | Lig/Ten | itga11b |
| CR753876.11 | 0 | 3.6042417 | 0.381 | 0.034 | 0 | Lig/Ten | CR753876.1 |
| si:dkey-30e9.7 | 0 | 5.8705476 | 0.352 | 0.006 | 0 | Lig/Ten | si:dkey-30e9.7 |
| si:ch1073-303k11.21 | 0 | 3.8584347 | 0.358 | 0.028 | 0 | Lig/Ten | si:ch1073-303k11.2 |
| col4a6 | 0 | 6.4696112 | 0.25 | 0.004 | 0 | Lig/Ten | col4a6 |
| vgf | 0 | 5.0837731 | 0.237 | 0.012 | 0 | Lig/Ten | vgf |
| col4a5 | 0 | 5.5320466 | 0.215 | 0.006 | 0 | Lig/Ten | col4a5 |
| ptx4 | 0 | 5.2173103 | 0.192 | 0.004 | 0 | Lig/Ten | ptx4 |
| enpep | 0 | 5.5992208 | 0.186 | 0.003 | 0 | Lig/Ten | enpep |
| olfm2a | 0 | 5.5605615 | 0.181 | 0.005 | 0 | Lig/Ten | olfm2a |
| bmp1b | 0 | 5.0515926 | 0.157 | 0.006 | 0 | Lig/Ten | bmp1b |
| mmp17b | 0 | 5.5781894 | 0.124 | 0.003 | 0 | Lig/Ten | mmp17b |
| gdf10a | 0 | 5.2995452 | 0.117 | 0.003 | 0 | Lig/Ten | gdf10a |
| irf81 | 5E-283 | 4.0648125 | 0.308 | 0.026 | 1.5E-278 | Lig/Ten | irf8 |
| ogna4 | 1E-271 | 2.9311581 | 0.785 | 0.184 | 3.4E-267 | Lig/Ten | ogna |
| si:dkey-27i16.26 | 6E-248 | 3.1441947 | 0.606 | 0.113 | 1.9E-243 | Lig/Ten | si:dkey-27i16.2 |
| meox2a1 | 8E-247 | 3.1786261 | 0.35 | 0.037 | 2.7E-242 | Lig/Ten | meox2a |
| pcolce2b6 | 2E-233 | 2.6162575 | 0.878 | 0.292 | 5.7E-229 | Lig/Ten | pcolce2b |
| egr3 | 4E-232 | 3.5987777 | 0.316 | 0.033 | 1.3E-227 | Lig/Ten | egr3 |
| tgfbi7 | 5E-231 | 2.684341 | 0.759 | 0.195 | 1.5E-226 | Lig/Ten | tgfbi |
| col4a14 | 1E-225 | 2.6161192 | 0.803 | 0.222 | 3.9E-221 | Lig/Ten | col4a1 |
| cthrc1b1 | 1E-210 | 3.6678317 | 0.223 | 0.018 | 3.2E-206 | Lig/Ten | cthrc1b |
| f81 | 2E-209 | 4.2739015 | 0.19 | 0.013 | 5.3E-205 | Lig/Ten | f8 |
| crip16 | 3E-209 | 2.4241971 | 0.971 | 0.458 | 8.8E-205 | Lig/Ten | crip1 |
| bmp16 | 2E-202 | 4.1568402 | 0.21 | 0.017 | 6.1E-198 | Lig/Ten | bmp16 |
| cd248b | 9E-195 | 3.7360882 | 0.21 | 0.017 | 2.9E-190 | Lig/Ten | cd248b |
| zgc:1748631 | 7E-187 | 3.0954235 | 0.726 | 0.225 | 2.3E-182 | Lig/Ten | zgc:174863 |
| col4a25 | 8E-185 | 2.5102396 | 0.732 | 0.214 | 2.7E-180 | Lig/Ten | col4a2 |
| fndc16 | 3E-177 | 2.3082505 | 0.653 | 0.163 | 8.4E-173 | Lig/Ten | fndc1 |
| myct1a | 4E-170 | 3.7865035 | 0.166 | 0.012 | 1.3E-165 | Lig/Ten | myct1a |
| nid1a4 | 7E-168 | 2.4918683 | 0.726 | 0.23 | 2.3E-163 | Lig/Ten | nid1a |
| p4ha1b6 | 3E-158 | 2.4007036 | 0.562 | 0.137 | 1E-153 | Lig/Ten | p4ha1b |
| ecrg4a5 | 5E-157 | 2.2320039 | 0.951 | 0.56 | 1.6E-152 | Lig/Ten | ecrg4a |
| col2a1b5 | 6E-156 | 1.8967745 | 0.925 | 0.498 | 1.9E-151 | Lig/Ten | col2a1b |
| col11a1b5 | 1E-151 | 1.8391733 | 0.956 | 0.54 | 3.6E-147 | Lig/Ten | col11a1b |
| postnb4 | 1E-142 | 1.685283 | 0.918 | 0.451 | 4E-138 | Lig/Ten | postnb |
| dcn6 | 1E-141 | 1.4514995 | 1 | 0.867 | 3.8E-137 | Lig/Ten | dcn |
| foxp47 | 2E-140 | 2.1424874 | 0.657 | 0.199 | 6.5E-136 | Lig/Ten | foxp4 |
| mkxa | 6E-139 | 3.5763338 | 0.201 | 0.022 | 2E-134 | Lig/Ten | mkxa |
| dachb | 3E-138 | 3.8552478 | 0.108 | 0.006 | 8.8E-134 | Lig/Ten | dachb |
| fgf16 | 1E-137 | 4.0883933 | 0.111 | 0.007 | 4.3E-133 | Lig/Ten | fgf16 |
| fn1b5 | 5E-137 | 2.5508876 | 0.885 | 0.397 | 1.8E-132 | Lig/Ten | fn1b |

| | | | | | | | |
|---|---|---|---|---|---|---|---|
| col18a1a4 | 6E-135 | 3.0747774 | 0.538 | 0.15 | 2.1E-130 | Lig/Ten | col18a1a |
| FP102018.15 | 3E-134 | 1.8668721 | 0.834 | 0.387 | 9.1E-130 | Lig/Ten | FP102018.1 |
| fam107b5 | 3E-134 | 2.407186 | 0.642 | 0.205 | 1.1E-129 | Lig/Ten | fam107b |
| col12a1a4 | 8E-134 | 1.5106687 | 0.998 | 0.75 | 2.5E-129 | Lig/Ten | col12a1a |
| loxa7 | 3E-133 | 2.3109048 | 0.591 | 0.17 | 8.7E-129 | Lig/Ten | loxa |
| npnta | 2E-132 | 3.0050293 | 0.206 | 0.023 | 7.5E-128 | Lig/Ten | npnta |
| mmp14a5 | 1E-126 | 1.6506529 | 0.94 | 0.645 | 3.4E-122 | Lig/Ten | mmp14a |
| wu:fc23c09 | 6E-121 | 3.093465 | 0.184 | 0.02 | 2E-116 | Lig/Ten | wu:fc23c09 |
| hsd17b3 | 1E-119 | 3.6957961 | 0.157 | 0.015 | 3.9E-115 | Lig/Ten | hsd17b3 |
| col1a25 | 3E-118 | 1.2427544 | 1 | 0.991 | 9E-114 | Lig/Ten | col1a2 |
| loxl2b4 | 4E-117 | 1.7581082 | 0.763 | 0.313 | 1.4E-112 | Lig/Ten | loxl2b |
| ebf3a.12 | 1E-116 | 2.6019224 | 0.281 | 0.046 | 3.9E-112 | Lig/Ten | ebf3a.1 |
| hlx1 | 3E-116 | 3.9476925 | 0.1 | 0.006 | 8.9E-112 | Lig/Ten | hlx1 |
| vwa16 | 6E-115 | 2.4313108 | 0.493 | 0.135 | 2.1E-110 | Lig/Ten | vwa1 |
| omd1 | 8E-113 | 3.4464444 | 0.235 | 0.035 | 2.6E-108 | Lig/Ten | omd |
| plod1a3 | 3E-112 | 2.0260813 | 0.648 | 0.24 | 1.1E-107 | Lig/Ten | plod1a |
| col1a1a5 | 2E-111 | 1.2567209 | 1 | 0.98 | 6.2E-107 | Lig/Ten | col1a1a |
| hhipl1 | 3E-111 | 2.9136565 | 0.177 | 0.02 | 1.1E-106 | Lig/Ten | hhipl1 |
| ecel1 | 9E-111 | 3.8846336 | 0.1 | 0.007 | 3E-106 | Lig/Ten | ecel1 |
| tgfb57 | 3E-109 | 2.103588 | 0.442 | 0.111 | 8.4E-105 | Lig/Ten | tgfb5 |
| angptl2b4 | 4E-109 | 1.8249711 | 0.502 | 0.138 | 1.2E-104 | Lig/Ten | angptl2b |
| egfra4 | 1E-108 | 2.08394 | 0.675 | 0.255 | 3.4E-104 | Lig/Ten | egfra |
| angptl1a5 | 4E-102 | 1.6863961 | 0.803 | 0.389 | 1.33E-97 | Lig/Ten | angptl1a |
| si:ch211-243a20.37 | 4E-102 | 1.5598644 | 0.715 | 0.278 | 1.4E-97 | Lig/Ten | si:ch211-243a20.3 |
| sparc5 | 7.2E-98 | 1.0121049 | 1 | 0.98 | 2.35E-93 | Lig/Ten | sparc |
| tfpia7 | 1.1E-97 | 1.8593617 | 0.544 | 0.176 | 3.65E-93 | Lig/Ten | tfpia |
| col5a16 | 1.5E-94 | 1.0214515 | 1 | 0.964 | 4.98E-90 | Lig/Ten | col5a1 |
| mmp25 | 1E-89 | 1.277361 | 0.954 | 0.743 | 3.31E-85 | Lig/Ten | mmp2 |
| hic13 | 2.7E-86 | 2.2647266 | 0.292 | 0.063 | 8.73E-82 | Lig/Ten | hic1 |
| tent5bb5 | 3.4E-86 | 1.9075075 | 0.46 | 0.14 | 1.1E-81 | Lig/Ten | tent5bb |
| prss355 | 4.4E-85 | 1.5674661 | 0.363 | 0.091 | 1.43E-80 | Lig/Ten | prss35 |
| col1a1b5 | 1.6E-78 | 0.8261097 | 0.998 | 0.987 | 5.18E-74 | Lig/Ten | col1a1b |
| nt5e4 | 4.5E-76 | 1.9887971 | 0.279 | 0.062 | 1.46E-71 | Lig/Ten | nt5e |
| col16a15 | 1.2E-75 | 1.2889098 | 0.812 | 0.452 | 3.79E-71 | Lig/Ten | col16a1 |
| p4ha24 | 2.6E-74 | 1.9316927 | 0.431 | 0.14 | 8.6E-70 | Lig/Ten | p4ha2 |
| tgfb34 | 4.1E-73 | 2.0366486 | 0.312 | 0.079 | 1.35E-68 | Lig/Ten | tgfb3 |
| loxl2a4 | 2.7E-72 | 1.6975513 | 0.558 | 0.221 | 8.89E-68 | Lig/Ten | loxl2a |
| nfixb7 | 3.1E-72 | 1.5572138 | 0.635 | 0.291 | 9.93E-68 | Lig/Ten | nfixb |
| angptl57 | 5E-72 | 1.7347359 | 0.482 | 0.17 | 1.63E-67 | Lig/Ten | angptl5 |
| si:ch1073-459j12.15 | 6.1E-72 | 0.9809756 | 0.978 | 0.81 | 2E-67 | Lig/Ten | si:ch1073-459j12.1 |
| si:ch211-195b13.16 | 8.8E-71 | 1.5434199 | 0.588 | 0.24 | 2.85E-66 | Lig/Ten | si:ch211-195b13.1 |
| dkk3b4 | 2.1E-70 | 1.3048808 | 0.602 | 0.245 | 6.8E-66 | Lig/Ten | dkk3b |
| cilp6 | 7.1E-69 | 1.783984 | 0.471 | 0.17 | 2.31E-64 | Lig/Ten | cilp |
| f3b7 | 2.3E-68 | 1.7624153 | 0.715 | 0.377 | 7.57E-64 | Lig/Ten | f3b |
| gadd45bb7 | 9E-68 | 2.099342 | 0.529 | 0.218 | 2.93E-63 | Lig/Ten | gadd45bb |
| pmp22a4 | 1.4E-66 | 0.9315418 | 0.923 | 0.688 | 4.68E-62 | Lig/Ten | pmp22a |
| ccn17 | 9.7E-66 | 1.4514297 | 0.878 | 0.641 | 3.15E-61 | Lig/Ten | ccn1 |
| PAMR15 | 1.8E-65 | 1.6466272 | 0.383 | 0.118 | 5.7E-61 | Lig/Ten | PAMR1 |

| | | | | | | | |
|---|---|---|---|---|---|---|---|
| aplp13 | 7.5E-65 | 1.9732381 | 0.206 | 0.041 | 2.44E-60 | Lig/Ten | aplp1 |
| adamts26 | 7.9E-65 | 1.2006458 | 0.737 | 0.37 | 2.55E-60 | Lig/Ten | adamts2 |
| col5a3a3 | 5.1E-64 | 1.5419527 | 0.27 | 0.066 | 1.65E-59 | Lig/Ten | col5a3a |
| si:ch211-37e10.2 | 2E-62 | 2.4652583 | 0.15 | 0.025 | 6.38E-58 | Lig/Ten | si:ch211-37e10.2 |
| thbs1a4 | 2.5E-62 | 1.3438731 | 0.644 | 0.31 | 8.11E-58 | Lig/Ten | thbs1a |
| hexb5 | 3.9E-61 | 1.8859257 | 0.314 | 0.089 | 1.27E-56 | Lig/Ten | hexb |
| si:ch211-236l14.41 | 8.4E-61 | 2.8135664 | 0.117 | 0.016 | 2.74E-56 | Lig/Ten | si:ch211-236l14.4 |
| serpinf16 | 1E-60 | 1.1144656 | 0.85 | 0.552 | 3.32E-56 | Lig/Ten | serpinf1 |
| mxra8a7 | 4.2E-60 | 1.4572231 | 0.569 | 0.251 | 1.35E-55 | Lig/Ten | mxra8a |
| fibinb5 | 1.2E-59 | 1.3261045 | 0.602 | 0.272 | 3.77E-55 | Lig/Ten | fibinb |
| phlda26 | 8.9E-59 | 1.7961787 | 0.507 | 0.217 | 2.88E-54 | Lig/Ten | phlda2 |
| rnd1b6 | 6.8E-58 | 1.7291594 | 0.334 | 0.101 | 2.21E-53 | Lig/Ten | rnd1b |
| dpt3 | 5.2E-57 | 0.9387104 | 0.434 | 0.152 | 1.7E-52 | Lig/Ten | dpt |
| angptl2a7 | 2E-56 | 1.2600501 | 0.646 | 0.316 | 6.38E-52 | Lig/Ten | angptl2a |
| pknox2 | 2.8E-55 | 2.9557814 | 0.113 | 0.016 | 9.09E-51 | Lig/Ten | pknox2 |
| mgst1.26 | 4.7E-55 | 1.2355962 | 0.624 | 0.298 | 1.54E-50 | Lig/Ten | mgst1.2 |
| xirp2a5 | 1.1E-53 | 1.5286364 | 0.546 | 0.255 | 3.56E-49 | Lig/Ten | xirp2a |
| tmem45a4 | 1.5E-53 | 1.6168289 | 0.414 | 0.156 | 4.79E-49 | Lig/Ten | tmem45a |
| timp2a7 | 9.4E-53 | 0.81266 | 0.931 | 0.754 | 3.04E-48 | Lig/Ten | timp2a |
| myoc4 | 5.3E-52 | 2.6430902 | 0.248 | 0.068 | 1.72E-47 | Lig/Ten | myoc |
| lipf2 | 2.1E-51 | 1.5066885 | 0.476 | 0.196 | 6.97E-47 | Lig/Ten | lipf |
| si:ch211-133n4.45 | 5.5E-51 | 1.2088095 | 0.739 | 0.446 | 1.78E-46 | Lig/Ten | si:ch211-133n4.4 |
| clec11a3 | 9.1E-51 | 1.9538598 | 0.221 | 0.056 | 2.96E-46 | Lig/Ten | clec11a |
| srgn5 | 1.3E-49 | 1.0663347 | 0.765 | 0.45 | 4.15E-45 | Lig/Ten | srgn |
| srpx5 | 1.4E-49 | 1.5029615 | 0.425 | 0.165 | 4.49E-45 | Lig/Ten | srpx |
| pim16 | 1.4E-49 | 1.1802873 | 0.794 | 0.543 | 4.66E-45 | Lig/Ten | pim1 |
| mast24 | 6E-49 | 1.4806119 | 0.418 | 0.162 | 1.95E-44 | Lig/Ten | mast2 |
| zgc:1583434 | 1.3E-48 | 1.5864624 | 0.659 | 0.376 | 4.1E-44 | Lig/Ten | zgc:158343 |
| il342 | 1.5E-47 | 1.7151269 | 0.206 | 0.051 | 4.94E-43 | Lig/Ten | il34 |
| jdp2b5 | 7.2E-47 | 0.9884189 | 0.838 | 0.569 | 2.33E-42 | Lig/Ten | jdp2b |
| zfp36l24 | 5.3E-46 | 1.5193976 | 0.549 | 0.266 | 1.71E-41 | Lig/Ten | zfp36l2 |
| abi3bpb6 | 1.6E-45 | 0.8387895 | 0.838 | 0.584 | 5.13E-41 | Lig/Ten | abi3bpb |
| si:dkey-282h22.55 | 2.5E-45 | 1.980775 | 0.241 | 0.069 | 8.26E-41 | Lig/Ten | si:dkey-282h22.5 |
| mycn | 3.5E-45 | 2.3423603 | 0.184 | 0.045 | 1.15E-40 | Lig/Ten | mycn |
| tent5ba3 | 8.2E-45 | 1.5521179 | 0.369 | 0.142 | 2.68E-40 | Lig/Ten | tent5ba |
| fabp11a2 | 1.4E-44 | 1.0496848 | 0.248 | 0.072 | 4.52E-40 | Lig/Ten | fabp11a |
| col12a1b4 | 2E-44 | 0.7654712 | 0.903 | 0.622 | 6.41E-40 | Lig/Ten | col12a1b |
| rasd11 | 2E-44 | 2.0709477 | 0.119 | 0.021 | 6.42E-40 | Lig/Ten | rasd1 |
| igsf105 | 2.3E-44 | 1.0806631 | 0.58 | 0.276 | 7.35E-40 | Lig/Ten | igsf10 |
| mdka5 | 1.5E-42 | 0.9838059 | 0.803 | 0.52 | 4.98E-38 | Lig/Ten | mdka |
| rgs5b7 | 2.2E-41 | 1.0955087 | 0.777 | 0.521 | 7.09E-37 | Lig/Ten | rgs5b |
| fndc3ba3 | 6.6E-41 | 1.4976604 | 0.354 | 0.137 | 2.14E-36 | Lig/Ten | fndc3ba |
| zgc:1752145 | 2.1E-38 | 1.4466573 | 0.296 | 0.106 | 6.72E-34 | Lig/Ten | zgc:175214 |
| rgcc6 | 1.1E-37 | 1.1103547 | 0.445 | 0.197 | 3.59E-33 | Lig/Ten | rgcc |
| crip24 | 1.3E-37 | 0.9027655 | 0.241 | 0.076 | 4.37E-33 | Lig/Ten | crip2 |
| dclk1a | 1.5E-37 | 2.7262745 | 0.102 | 0.019 | 4.76E-33 | Lig/Ten | dclk1a |
| CR356233.1 | 3.3E-37 | 1.808315 | 0.166 | 0.042 | 1.08E-32 | Lig/Ten | CR356233.1 |
| timp2b3 | 7.8E-37 | 2.3729284 | 0.491 | 0.25 | 2.55E-32 | Lig/Ten | timp2b |

| | | | | | | | |
|---|---|---|---|---|---|---|---|
| itga103 | 6.9E-36 | 1.4004472 | 0.498 | 0.252 | 2.23E-31 | Lig/Ten | itga10 |
| CR381618.17 | 4E-35 | 1.1049656 | 0.467 | 0.226 | 1.3E-30 | Lig/Ten | CR381618.1 |
| tgfbr2b7 | 1.7E-34 | 1.2684105 | 0.345 | 0.142 | 5.56E-30 | Lig/Ten | tgfbr2b |
| igf11 | 1.9E-34 | 2.7919212 | 0.102 | 0.02 | 6.09E-30 | Lig/Ten | igf1 |
| marcksl1b7 | 2.7E-34 | 0.7046868 | 0.515 | 0.252 | 8.92E-30 | Lig/Ten | marcksl1b |
| marcksl1a6 | 5.9E-34 | 0.616448 | 0.912 | 0.706 | 1.9E-29 | Lig/Ten | marcksl1a |
| rbfox13 | 2.1E-33 | 0.9919216 | 0.653 | 0.404 | 6.84E-29 | Lig/Ten | rbfox1 |
| pdgfrb5 | 2.6E-33 | 0.9923368 | 0.668 | 0.41 | 8.41E-29 | Lig/Ten | pdgfrb |
| epha71 | 1.7E-32 | 1.6231458 | 0.177 | 0.051 | 5.52E-28 | Lig/Ten | epha7 |
| c1qtnf56 | 2.9E-32 | 1.3861468 | 0.35 | 0.151 | 9.39E-28 | Lig/Ten | c1qtnf5 |
| pabpc1b3 | 3.8E-32 | 0.7998787 | 0.799 | 0.585 | 1.24E-27 | Lig/Ten | pabpc1b |
| thbs3a5 | 6E-32 | 1.0247273 | 0.56 | 0.307 | 1.94E-27 | Lig/Ten | thbs3a |
| fzd8a1 | 8.9E-32 | 2.0840625 | 0.131 | 0.032 | 2.9E-27 | Lig/Ten | fzd8a |
| her96 | 1E-31 | 1.8284941 | 0.44 | 0.227 | 3.26E-27 | Lig/Ten | her9 |
| cd248a6 | 1.6E-31 | 1.0044125 | 0.301 | 0.115 | 5.33E-27 | Lig/Ten | cd248a |
| kdelr35 | 2.5E-31 | 1.3528148 | 0.383 | 0.181 | 8.12E-27 | Lig/Ten | kdelr3 |
| lgals2a5 | 5.6E-31 | 0.6609243 | 0.867 | 0.639 | 1.83E-26 | Lig/Ten | lgals2a |
| rrbp1b3 | 8.2E-31 | 1.0367623 | 0.469 | 0.237 | 2.65E-26 | Lig/Ten | rrbp1b |
| CR339041.11 | 1.1E-30 | 1.6370518 | 0.257 | 0.097 | 3.44E-26 | Lig/Ten | CR339041.1 |
| ccn2a6 | 1.4E-30 | 1.7469495 | 0.405 | 0.205 | 4.47E-26 | Lig/Ten | ccn2a |
| aebp16 | 2.6E-30 | 0.7808261 | 0.792 | 0.538 | 8.59E-26 | Lig/Ten | aebp1 |
| podn6 | 4.1E-30 | 1.1597731 | 0.381 | 0.171 | 1.33E-25 | Lig/Ten | podn |
| icam35 | 1.1E-29 | 1.4381329 | 0.23 | 0.082 | 3.49E-25 | Lig/Ten | icam3 |
| plekhg5a5 | 1.3E-29 | 1.5255906 | 0.21 | 0.071 | 4.12E-25 | Lig/Ten | plekhg5a |
| baz2ba6 | 2E-29 | 1.2266366 | 0.367 | 0.169 | 6.61E-25 | Lig/Ten | baz2ba |
| itgam1 | 4.8E-29 | 1.3924605 | 0.15 | 0.041 | 1.56E-24 | Lig/Ten | itgam |
| ddr2a5 | 6.2E-29 | 1.1082728 | 0.42 | 0.206 | 2.01E-24 | Lig/Ten | ddr2a |
| phldb2a | 1.3E-28 | 1.5253703 | 0.186 | 0.059 | 4.17E-24 | Lig/Ten | phldb2a |
| hmcn15 | 1.5E-28 | 1.3744927 | 0.21 | 0.071 | 5.02E-24 | Lig/Ten | hmcn1 |
| lrp1aa8 | 3.5E-28 | 0.6570962 | 0.858 | 0.673 | 1.15E-23 | Lig/Ten | lrp1aa |
| mknk2a5 | 4.9E-28 | 1.0398134 | 0.442 | 0.228 | 1.58E-23 | Lig/Ten | mknk2a |
| copz24 | 6.6E-28 | 1.0108471 | 0.392 | 0.186 | 2.15E-23 | Lig/Ten | copz2 |
| tmem1769 | 2.6E-27 | 0.7928293 | 0.637 | 0.392 | 8.36E-23 | Lig/Ten | tmem176 |
| pdgfrl6 | 3E-27 | 0.6956542 | 0.77 | 0.584 | 9.62E-23 | Lig/Ten | pdgfrl |
| ifi303 | 6E-27 | 1.1521097 | 0.325 | 0.145 | 1.94E-22 | Lig/Ten | ifi30 |
| rasgef1bb7 | 1.3E-26 | 1.2009335 | 0.336 | 0.154 | 4.31E-22 | Lig/Ten | rasgef1bb |
| bhlhe414 | 9.7E-26 | 1.1856121 | 0.296 | 0.128 | 3.14E-21 | Lig/Ten | bhlhe41 |
| cnn3a4 | 1.2E-25 | 0.7953907 | 0.684 | 0.475 | 3.96E-21 | Lig/Ten | cnn3a |
| col5a2a5 | 3.7E-25 | 0.7606715 | 0.869 | 0.792 | 1.2E-20 | Lig/Ten | col5a2a |
| lima1a3 | 5.6E-25 | 1.338953 | 0.23 | 0.089 | 1.84E-20 | Lig/Ten | lima1a |
| f106 | 6E-25 | 0.8394632 | 0.617 | 0.398 | 1.96E-20 | Lig/Ten | f10 |
| plxdc25 | 7.2E-25 | 1.0805019 | 0.338 | 0.157 | 2.35E-20 | Lig/Ten | plxdc2 |
| adam10a5 | 8E-25 | 0.8189181 | 0.604 | 0.38 | 2.6E-20 | Lig/Ten | adam10a |
| plk36 | 1.2E-24 | 1.1551432 | 0.389 | 0.201 | 3.83E-20 | Lig/Ten | plk3 |
| rab20 | 1.8E-24 | 1.6015037 | 0.135 | 0.039 | 5.71E-20 | Lig/Ten | rab20 |
| klf7a3 | 2.1E-24 | 1.3873177 | 0.17 | 0.057 | 6.77E-20 | Lig/Ten | klf7a |
| fkbp73 | 2.9E-24 | 0.6924126 | 0.794 | 0.611 | 9.52E-20 | Lig/Ten | fkbp7 |
| tgfbr2a1 | 3E-24 | 1.5509461 | 0.159 | 0.052 | 9.71E-20 | Lig/Ten | tgfbr2a |

| | | | | | | | |
|---|---|---|---|---|---|---|---|
| vasna5 | 3.2E-24 | 1.0227399 | 0.272 | 0.114 | 1.05E-19 | Lig/Ten | vasna |
| nfia5 | 4.1E-24 | 0.9818654 | 0.352 | 0.169 | 1.32E-19 | Lig/Ten | nfia |
| sgk2b3 | 5.5E-24 | 1.6544842 | 0.131 | 0.038 | 1.79E-19 | Lig/Ten | sgk2b |
| efcc1 | 6E-24 | 1.7952089 | 0.124 | 0.035 | 1.95E-19 | Lig/Ten | efcc1 |
| dennd2b1 | 1E-23 | 1.0819869 | 0.288 | 0.126 | 3.31E-19 | Lig/Ten | dennd2b |
| CR936442.13 | 1.2E-23 | 0.9736192 | 0.374 | 0.19 | 3.93E-19 | Lig/Ten | CR936442.1 |
| ext1a2 | 1.6E-23 | 1.6865804 | 0.155 | 0.051 | 5.11E-19 | Lig/Ten | ext1a |
| cgref15 | 1.6E-23 | 0.8696229 | 0.356 | 0.172 | 5.16E-19 | Lig/Ten | cgref1 |
| tnn2 | 2.5E-23 | 1.4139645 | 0.117 | 0.032 | 8.13E-19 | Lig/Ten | tnn |
| selenop5 | 2.6E-23 | 0.832837 | 0.843 | 0.74 | 8.4E-19 | Lig/Ten | selenop |
| dusp65 | 3.1E-23 | 1.3698922 | 0.308 | 0.145 | 1E-18 | Lig/Ten | dusp6 |
| zmp:00000007292 | 4.8E-23 | 1.7882506 | 0.15 | 0.049 | 1.57E-18 | Lig/Ten | zmp:0000000729 |
| spon2b5 | 8.3E-23 | 1.6051563 | 0.277 | 0.127 | 2.69E-18 | Lig/Ten | spon2b |
| nucb2a5 | 1E-22 | 0.6808003 | 0.723 | 0.523 | 3.32E-18 | Lig/Ten | nucb2a |
| rrbp1a6 | 1.8E-22 | 0.6858977 | 0.657 | 0.429 | 5.93E-18 | Lig/Ten | rrbp1a |
| il17rd | 2E-22 | 1.572264 | 0.104 | 0.027 | 6.61E-18 | Lig/Ten | il17rd |
| creb3l13 | 2.4E-22 | 1.1732158 | 0.259 | 0.112 | 7.74E-18 | Lig/Ten | creb3l1 |
| fkbp94 | 5E-22 | 0.7189556 | 0.595 | 0.383 | 1.62E-17 | Lig/Ten | fkbp9 |
| cpn16 | 6.6E-22 | 0.9019931 | 0.403 | 0.211 | 2.15E-17 | Lig/Ten | cpn1 |
| nfatc14 | 1.1E-21 | 1.1750311 | 0.215 | 0.085 | 3.48E-17 | Lig/Ten | nfatc1 |
| ngfb6 | 1.1E-21 | 1.5060265 | 0.204 | 0.081 | 3.58E-17 | Lig/Ten | ngfb |
| skia8 | 1.2E-21 | 0.783309 | 0.655 | 0.448 | 4.06E-17 | Lig/Ten | skia |
| gstt1a4 | 2.6E-21 | 1.281226 | 0.195 | 0.075 | 8.55E-17 | Lig/Ten | gstt1a |
| bmp1a6 | 4.9E-21 | 0.961216 | 0.361 | 0.186 | 1.59E-16 | Lig/Ten | bmp1a |
| tmem119b7 | 5.9E-21 | 0.9489031 | 0.387 | 0.205 | 1.9E-16 | Lig/Ten | tmem119b |
| cpe5 | 6.1E-21 | 1.0668483 | 0.381 | 0.202 | 2E-16 | Lig/Ten | cpe |
| irf2bp2b5 | 7.6E-21 | 0.9522592 | 0.385 | 0.209 | 2.46E-16 | Lig/Ten | irf2bp2b |
| si:ch73-138n13.15 | 9.6E-21 | 0.5914885 | 0.604 | 0.376 | 3.13E-16 | Lig/Ten | si:ch73-138n13.1 |
| sost5 | 1.4E-20 | 1.5472227 | 0.195 | 0.078 | 4.6E-16 | Lig/Ten | sost |
| si:dkey-16l2.203 | 1.4E-20 | 1.4004065 | 0.17 | 0.063 | 4.65E-16 | Lig/Ten | si:dkey-16l2.20 |
| pros13 | 1.5E-20 | 0.9843512 | 0.261 | 0.116 | 4.72E-16 | Lig/Ten | pros1 |
| spry46 | 1.6E-20 | 0.8024561 | 0.354 | 0.178 | 5.12E-16 | Lig/Ten | spry4 |
| socs3a3 | 2.2E-20 | 0.5849211 | 0.909 | 0.794 | 7.28E-16 | Lig/Ten | socs3a |
| egr15 | 3.4E-20 | 0.5763955 | 0.845 | 0.627 | 1.12E-15 | Lig/Ten | egr1 |
| map2 | 1E-19 | 1.3234523 | 0.111 | 0.033 | 3.37E-15 | Lig/Ten | map2 |
| ifi464 | 1.1E-19 | 1.4907392 | 0.219 | 0.094 | 3.55E-15 | Lig/Ten | ifi46 |
| ctsa4 | 1.4E-19 | 1.1839643 | 0.254 | 0.117 | 4.68E-15 | Lig/Ten | ctsa |
| cldni3 | 1.8E-19 | 1.2938783 | 0.166 | 0.061 | 5.97E-15 | Lig/Ten | cldni |
| serpinh25 | 2.1E-19 | 1.044794 | 0.257 | 0.117 | 6.73E-15 | Lig/Ten | serpinh2 |
| clstn16 | 2.3E-19 | 1.2433081 | 0.281 | 0.135 | 7.61E-15 | Lig/Ten | clstn1 |
| ip6k2a6 | 2.5E-19 | 1.0116083 | 0.363 | 0.196 | 7.98E-15 | Lig/Ten | ip6k2a |
| b2m7 | 2.7E-19 | 0.4277995 | 0.843 | 0.685 | 8.74E-15 | Lig/Ten | b2m |
| ssr45 | 3.1E-19 | 0.6093278 | 0.777 | 0.607 | 1.01E-14 | Lig/Ten | ssr4 |
| fn1a5 | 4E-19 | 0.7108387 | 0.524 | 0.316 | 1.3E-14 | Lig/Ten | fn1a |
| ptprk4 | 4.2E-19 | 1.4729369 | 0.177 | 0.069 | 1.35E-14 | Lig/Ten | ptprk |
| acsl4b4 | 4.5E-19 | 1.252909 | 0.272 | 0.131 | 1.47E-14 | Lig/Ten | acsl4b |
| mrc25 | 7.7E-19 | 1.2696167 | 0.208 | 0.088 | 2.49E-14 | Lig/Ten | mrc2 |
| col6a17 | 1E-18 | 0.4134481 | 0.967 | 0.874 | 3.28E-14 | Lig/Ten | col6a1 |

| | | | | | | | |
|---|---|---|---|---|---|---|---|
| ctsh5 | 1.6E-18 | 1.2221168 | 0.239 | 0.11 | 5.05E-14 | Lig/Ten | ctsh |
| ctsz4 | 1.8E-18 | 1.0305189 | 0.314 | 0.162 | 5.89E-14 | Lig/Ten | ctsz |
| fbln54 | 1.9E-18 | 1.3076668 | 0.199 | 0.083 | 6.05E-14 | Lig/Ten | fbln5 |
| im:71523487 | 2.3E-18 | 0.7097332 | 0.416 | 0.235 | 7.53E-14 | Lig/Ten | im:7152348 |
| fkbp10b5 | 3E-18 | 0.8689802 | 0.429 | 0.254 | 9.85E-14 | Lig/Ten | fkbp10b |
| ctsba4 | 3.1E-18 | 0.8282376 | 0.575 | 0.383 | 1.01E-13 | Lig/Ten | ctsba |
| ano66 | 3.4E-18 | 1.1171491 | 0.261 | 0.124 | 1.11E-13 | Lig/Ten | ano6 |
| frzb7 | 3.7E-18 | 0.6853175 | 0.268 | 0.129 | 1.19E-13 | Lig/Ten | frzb |
| zgc:1102395 | 3.7E-18 | 0.8075506 | 0.403 | 0.231 | 1.2E-13 | Lig/Ten | zgc:110239 |
| col21a14 | 4.7E-18 | 1.1643647 | 0.192 | 0.08 | 1.53E-13 | Lig/Ten | col21a1 |
| map7d1b4 | 6.5E-18 | 0.9323969 | 0.35 | 0.189 | 2.11E-13 | Lig/Ten | map7d1b |
| skib8 | 6.8E-18 | 0.7244349 | 0.372 | 0.201 | 2.2E-13 | Lig/Ten | skib |
| b2ml8 | 1.7E-17 | 0.3656036 | 0.951 | 0.897 | 5.39E-13 | Lig/Ten | b2ml |
| sipa1l1 | 1.8E-17 | 1.7560709 | 0.106 | 0.033 | 5.7E-13 | Lig/Ten | sipa1l1 |
| slc39a134 | 1.9E-17 | 1.1770883 | 0.277 | 0.141 | 6.21E-13 | Lig/Ten | slc39a13 |
| si:dkey-7j14.64 | 2.1E-17 | 0.4564914 | 0.883 | 0.776 | 6.94E-13 | Lig/Ten | si:dkey-7j14.6 |
| rcan1a7 | 3E-17 | 0.9657458 | 0.394 | 0.229 | 9.63E-13 | Lig/Ten | rcan1a |
| srpx26 | 3.6E-17 | 0.8170057 | 0.429 | 0.253 | 1.16E-12 | Lig/Ten | srpx2 |
| lmod1a | 4.5E-17 | 1.6197512 | 0.106 | 0.034 | 1.47E-12 | Lig/Ten | lmod1a |
| nucb15 | 4.8E-17 | 0.9317454 | 0.365 | 0.207 | 1.57E-12 | Lig/Ten | nucb1 |
| bloc1s65 | 6E-17 | 0.8850667 | 0.33 | 0.179 | 1.96E-12 | Lig/Ten | bloc1s6 |
| si:ch211-195m9.32 | 7.5E-17 | 1.751848 | 0.126 | 0.044 | 2.45E-12 | Lig/Ten | si:ch211-195m9.3 |
| cd2765 | 1.3E-16 | 0.9234813 | 0.268 | 0.132 | 4.15E-12 | Lig/Ten | cd276 |
| mfap27 | 1.5E-16 | 0.7080424 | 0.56 | 0.383 | 4.73E-12 | Lig/Ten | mfap2 |
| txndc54 | 1.9E-16 | 0.8391625 | 0.361 | 0.206 | 6.08E-12 | Lig/Ten | txndc5 |
| arhgef1a4 | 1.9E-16 | 1.4750593 | 0.153 | 0.06 | 6.11E-12 | Lig/Ten | arhgef1a |
| serf25 | 2.1E-16 | 0.4321936 | 0.894 | 0.772 | 6.86E-12 | Lig/Ten | serf2 |
| cd635 | 2.3E-16 | 0.843077 | 0.912 | 0.827 | 7.61E-12 | Lig/Ten | cd63 |
| mmp14b6 | 2.4E-16 | 0.5747507 | 0.719 | 0.549 | 7.69E-12 | Lig/Ten | mmp14b |
| zmp:00000009911 | 2.5E-16 | 1.1742508 | 0.1 | 0.031 | 8.06E-12 | Lig/Ten | zmp:0000000991 |
| ppp1r1b | 2.5E-16 | 1.5542717 | 0.108 | 0.036 | 8.1E-12 | Lig/Ten | ppp1r1b |
| pfn25 | 3.3E-16 | 0.5398091 | 0.712 | 0.538 | 1.08E-11 | Lig/Ten | pfn2 |
| ENSDARG000000908 | 3.5E-16 | 0.9085714 | 0.204 | 0.09 | 1.15E-11 | Lig/Ten | ENSDARG00000090 |
| ppfibp1a3 | 4.2E-16 | 1.2598541 | 0.144 | 0.055 | 1.37E-11 | Lig/Ten | ppfibp1a |
| cxxc5a3 | 4.7E-16 | 1.0513616 | 0.228 | 0.107 | 1.52E-11 | Lig/Ten | cxxc5a |
| si:ch1073-291c23.28 | 5.2E-16 | 0.5809513 | 0.668 | 0.473 | 1.68E-11 | Lig/Ten | si:ch1073-291c23.2 |
| sgce3 | 6.2E-16 | 0.6674275 | 0.46 | 0.281 | 2.02E-11 | Lig/Ten | sgce |
| ZFP36L1.15 | 8.1E-16 | 0.4735179 | 0.945 | 0.873 | 2.62E-11 | Lig/Ten | ZFP36L1.1 |
| rabac15 | 9.5E-16 | 0.5832308 | 0.58 | 0.396 | 3.09E-11 | Lig/Ten | rabac1 |
| fosab5 | 1.1E-15 | 0.3048856 | 0.985 | 0.894 | 3.51E-11 | Lig/Ten | fosab |
| gtf2a13 | 1.2E-15 | 1.4883658 | 0.184 | 0.082 | 3.97E-11 | Lig/Ten | gtf2a1 |
| nxn2 | 1.4E-15 | 1.27805 | 0.162 | 0.066 | 4.49E-11 | Lig/Ten | nxn |
| mfsd13a2 | 1.4E-15 | 1.258073 | 0.146 | 0.057 | 4.61E-11 | Lig/Ten | mfsd13a |
| c46 | 1.5E-15 | 1.1568516 | 0.354 | 0.207 | 4.88E-11 | Lig/Ten | c4 |
| inhbaa6 | 1.7E-15 | 0.7915635 | 0.44 | 0.271 | 5.64E-11 | Lig/Ten | inhbaa |
| cd82b3 | 1.9E-15 | 0.5410121 | 0.56 | 0.374 | 6.29E-11 | Lig/Ten | cd82b |
| gulp1a5 | 2.2E-15 | 0.806757 | 0.336 | 0.187 | 7.21E-11 | Lig/Ten | gulp1a |
| tpm4b4 | 2.5E-15 | 1.1811887 | 0.186 | 0.084 | 8.11E-11 | Lig/Ten | tpm4b |

| | | | | | | | |
|---|---|---|---|---|---|---|---|
| lman15 | 2.8E-15 | 0.9273747 | 0.248 | 0.124 | 8.97E-11 | Lig/Ten | lman1 |
| prdx64 | 3.1E-15 | 1.141218 | 0.485 | 0.343 | 1E-10 | Lig/Ten | prdx6 |
| smco45 | 3.1E-15 | 0.8195636 | 0.274 | 0.141 | 1.02E-10 | Lig/Ten | smco4 |
| midn6 | 3.2E-15 | 0.5229648 | 0.624 | 0.433 | 1.03E-10 | Lig/Ten | midn |
| fam49a3 | 3.2E-15 | 1.2321595 | 0.173 | 0.075 | 1.03E-10 | Lig/Ten | fam49a |
| plpp2b4 | 4.9E-15 | 0.9569725 | 0.25 | 0.127 | 1.6E-10 | Lig/Ten | plpp2b |
| aldocb4 | 5.7E-15 | 1.0842558 | 0.175 | 0.075 | 1.86E-10 | Lig/Ten | aldocb |
| CABZ01064941.14 | 6.7E-15 | 1.1361638 | 0.188 | 0.085 | 2.17E-10 | Lig/Ten | CABZ01064941.1 |
| angptl47 | 7.7E-15 | 0.5258495 | 0.644 | 0.455 | 2.5E-10 | Lig/Ten | angptl4 |
| dpp75 | 9.8E-15 | 0.8243669 | 0.363 | 0.215 | 3.19E-10 | Lig/Ten | dpp7 |
| si:ch211-222l21.15 | 1E-14 | 0.354745 | 0.737 | 0.554 | 3.36E-10 | Lig/Ten | si:ch211-222l21.1 |
| lepr5 | 1.5E-14 | 1.1871877 | 0.153 | 0.063 | 4.91E-10 | Lig/Ten | lepr |
| ifi453 | 2E-14 | 1.3882054 | 0.376 | 0.228 | 6.43E-10 | Lig/Ten | ifi45 |
| si:ch73-380l3.26 | 2.1E-14 | 0.4901466 | 0.604 | 0.415 | 6.78E-10 | Lig/Ten | si:ch73-380l3.2 |
| nova26 | 2.1E-14 | 0.5294552 | 0.374 | 0.213 | 6.81E-10 | Lig/Ten | nova2 |
| bag35 | 2.1E-14 | 1.3183718 | 0.164 | 0.071 | 6.96E-10 | Lig/Ten | bag3 |
| yap13 | 2.7E-14 | 0.7055759 | 0.527 | 0.359 | 8.67E-10 | Lig/Ten | yap1 |
| smad3a3 | 3.6E-14 | 0.6409641 | 0.469 | 0.305 | 1.16E-09 | Lig/Ten | smad3a |
| thbs2b6 | 4.1E-14 | 0.4165856 | 0.425 | 0.25 | 1.33E-09 | Lig/Ten | thbs2b |
| ssr36 | 4.2E-14 | 0.5019675 | 0.752 | 0.577 | 1.38E-09 | Lig/Ten | ssr3 |
| aga | 4.3E-14 | 1.2468387 | 0.146 | 0.06 | 1.41E-09 | Lig/Ten | aga |
| ches15 | 4.5E-14 | 0.8380143 | 0.319 | 0.182 | 1.48E-09 | Lig/Ten | ches1 |
| nme2b.14 | 4.6E-14 | 0.2589896 | 0.954 | 0.918 | 1.49E-09 | Lig/Ten | nme2b.1 |
| hsd11b1la | 4.8E-14 | 1.1843175 | 0.126 | 0.048 | 1.56E-09 | Lig/Ten | hsd11b1la |
| eva1a3 | 4.8E-14 | 1.3628389 | 0.139 | 0.057 | 1.57E-09 | Lig/Ten | eva1a |
| arhgap42a4 | 5.5E-14 | 1.0200423 | 0.166 | 0.072 | 1.79E-09 | Lig/Ten | arhgap42a |
| C20H1orf1985 | 5.7E-14 | 0.7247922 | 0.325 | 0.184 | 1.87E-09 | Lig/Ten | C20H1orf198 |
| col27a1b6 | 6E-14 | 0.6298484 | 0.513 | 0.336 | 1.95E-09 | Lig/Ten | col27a1b |
| mxra5b4 | 6.4E-14 | 1.4039957 | 0.144 | 0.059 | 2.07E-09 | Lig/Ten | mxra5b |
| c1qtnf6a4 | 9.7E-14 | 1.2307358 | 0.146 | 0.061 | 3.14E-09 | Lig/Ten | c1qtnf6a |
| btg36 | 1E-13 | 0.8447333 | 0.246 | 0.126 | 3.29E-09 | Lig/Ten | btg3 |
| fthl275 | 1.6E-13 | 0.7034484 | 0.916 | 0.839 | 5.22E-09 | Lig/Ten | fthl27 |
| tsc22d35 | 2.1E-13 | 0.5969136 | 0.564 | 0.402 | 6.69E-09 | Lig/Ten | tsc22d3 |
| roraa4 | 2.2E-13 | 0.6867892 | 0.268 | 0.141 | 7.03E-09 | Lig/Ten | roraa |
| rassf2a4 | 2.2E-13 | 0.9611953 | 0.241 | 0.125 | 7.29E-09 | Lig/Ten | rassf2a |
| efnb2a4 | 2.3E-13 | 0.8302554 | 0.277 | 0.15 | 7.63E-09 | Lig/Ten | efnb2a |
| egln2 | 2.7E-13 | 1.5413416 | 0.104 | 0.038 | 8.62E-09 | Lig/Ten | egln2 |
| egr2a5 | 3.3E-13 | 0.7779904 | 0.197 | 0.094 | 1.09E-08 | Lig/Ten | egr2a |
| sh2d3cb | 4.4E-13 | 1.3244558 | 0.128 | 0.052 | 1.42E-08 | Lig/Ten | sh2d3cb |
| fstl1b5 | 4.5E-13 | 0.4476048 | 0.865 | 0.756 | 1.45E-08 | Lig/Ten | fstl1b |
| nr4a15 | 4.5E-13 | 1.1034284 | 0.206 | 0.102 | 1.46E-08 | Lig/Ten | nr4a1 |
| nedd4l4 | 4.8E-13 | 0.8789449 | 0.204 | 0.099 | 1.57E-08 | Lig/Ten | nedd4l |
| fmnl32 | 6.1E-13 | 0.9183792 | 0.259 | 0.138 | 1.99E-08 | Lig/Ten | fmnl3 |
| edil3a6 | 6.3E-13 | 0.8288053 | 0.575 | 0.435 | 2.04E-08 | Lig/Ten | edil3a |
| abhd17b1 | 6.8E-13 | 1.1796124 | 0.15 | 0.066 | 2.21E-08 | Lig/Ten | abhd17b |
| lepb1 | 7.4E-13 | 1.5509196 | 0.1 | 0.036 | 2.41E-08 | Lig/Ten | lepb |
| ostm12 | 7.7E-13 | 1.1069406 | 0.157 | 0.07 | 2.52E-08 | Lig/Ten | ostm1 |
| tapbp.11 | 8.4E-13 | 0.9477793 | 0.204 | 0.1 | 2.74E-08 | Lig/Ten | tapbp.1 |

| | | | | | | | |
|---|---|---|---|---|---|---|---|
| igflr12 | 9.5E-13 | 1.175166 | 0.111 | 0.042 | 3.1E-08 | Lig/Ten | igflr1 |
| gadd45ba4 | 1E-12 | 0.4520027 | 0.914 | 0.803 | 3.24E-08 | Lig/Ten | gadd45ba |
| entpd16 | 1.1E-12 | 0.8197001 | 0.177 | 0.081 | 3.63E-08 | Lig/Ten | entpd1 |
| ier2b4 | 2.2E-12 | 0.416986 | 0.816 | 0.648 | 7.11E-08 | Lig/Ten | ier2b |
| triqk4 | 2.4E-12 | 0.751289 | 0.285 | 0.162 | 7.86E-08 | Lig/Ten | triqk |
| c6.14 | 2.8E-12 | 0.2049618 | 0.35 | 0.205 | 9.07E-08 | Lig/Ten | c6.1 |
| mcl1b5 | 2.9E-12 | 0.4945449 | 0.653 | 0.482 | 9.47E-08 | Lig/Ten | mcl1b |
| tspan74 | 3.6E-12 | 0.6315593 | 0.356 | 0.215 | 1.17E-07 | Lig/Ten | tspan7 |
| dap6 | 5.4E-12 | 0.6925193 | 0.327 | 0.195 | 1.77E-07 | Lig/Ten | dap |
| add3a6 | 5.8E-12 | 0.5092425 | 0.615 | 0.447 | 1.9E-07 | Lig/Ten | add3a |
| angptl74 | 5.9E-12 | 0.5419685 | 0.334 | 0.203 | 1.91E-07 | Lig/Ten | angptl7 |
| itga11a5 | 6E-12 | 0.5901149 | 0.217 | 0.11 | 1.95E-07 | Lig/Ten | itga11a |
| qkia4 | 6E-12 | 0.6172046 | 0.615 | 0.467 | 1.96E-07 | Lig/Ten | qkia |
| tmem9b3 | 6.8E-12 | 0.8647332 | 0.204 | 0.102 | 2.2E-07 | Lig/Ten | tmem9b |
| olfml3b5 | 7.3E-12 | 0.4654024 | 0.666 | 0.513 | 2.36E-07 | Lig/Ten | olfml3b |
| pamr15 | 7.5E-12 | 0.7676241 | 0.215 | 0.109 | 2.45E-07 | Lig/Ten | pamr1 |
| ldlrad4a1 | 8E-12 | 1.4615879 | 0.1 | 0.038 | 2.61E-07 | Lig/Ten | ldlrad4a |
| slc7a10a6 | 1.2E-11 | 0.7679008 | 0.21 | 0.107 | 4E-07 | Lig/Ten | slc7a10a |
| ch25hl36 | 1.3E-11 | 0.6303411 | 0.21 | 0.106 | 4.16E-07 | Lig/Ten | ch25hl3 |
| tbx2a3 | 1.3E-11 | 1.0171228 | 0.139 | 0.061 | 4.28E-07 | Lig/Ten | tbx2a |
| irf1b7 | 1.5E-11 | 0.9455012 | 0.221 | 0.117 | 4.8E-07 | Lig/Ten | irf1b |
| si:dkeyp-23e4.38 | 1.5E-11 | 0.9647667 | 0.177 | 0.085 | 4.98E-07 | Lig/Ten | si:dkeyp-23e4.3 |
| smox4 | 1.6E-11 | 1.0605506 | 0.168 | 0.081 | 5.28E-07 | Lig/Ten | smox |
| creb3l3l3 | 2.1E-11 | 0.5853041 | 0.389 | 0.249 | 6.72E-07 | Lig/Ten | creb3l3l |
| klf12a5 | 2.2E-11 | 0.7817394 | 0.25 | 0.138 | 7.3E-07 | Lig/Ten | klf12a |
| tnfsf123 | 2.3E-11 | 0.9087725 | 0.257 | 0.146 | 7.34E-07 | Lig/Ten | tnfsf12 |
| prss166 | 2.8E-11 | 0.8749508 | 0.181 | 0.09 | 9.02E-07 | Lig/Ten | prss16 |
| slc2a102 | 3E-11 | 1.0984245 | 0.102 | 0.04 | 9.76E-07 | Lig/Ten | slc2a10 |
| tuba8l26 | 3E-11 | 0.8596329 | 0.341 | 0.217 | 9.84E-07 | Lig/Ten | tuba8l2 |
| si:ch73-269m14.23 | 3.1E-11 | 0.5893878 | 0.365 | 0.227 | 1.01E-06 | Lig/Ten | si:ch73-269m14.2 |
| scpep13 | 3.7E-11 | 0.7982141 | 0.239 | 0.131 | 1.22E-06 | Lig/Ten | scpep1 |
| nfatc3a4 | 3.8E-11 | 1.0298809 | 0.184 | 0.093 | 1.25E-06 | Lig/Ten | nfatc3a |
| loxl3b2 | 4.3E-11 | 0.6050621 | 0.272 | 0.154 | 1.39E-06 | Lig/Ten | loxl3b |
| arhgef45 | 4.4E-11 | 0.9935788 | 0.159 | 0.076 | 1.43E-06 | Lig/Ten | arhgef4 |
| rtn4b4 | 4.4E-11 | 0.8736245 | 0.184 | 0.092 | 1.43E-06 | Lig/Ten | rtn4b |
| si:rp71-36a1.53 | 4.5E-11 | 0.8710357 | 0.177 | 0.087 | 1.45E-06 | Lig/Ten | si:rp71-36a1.5 |
| pepd5 | 4.7E-11 | 0.5141205 | 0.35 | 0.212 | 1.54E-06 | Lig/Ten | pepd |
| fuca25 | 5.1E-11 | 0.7669554 | 0.257 | 0.146 | 1.65E-06 | Lig/Ten | fuca2 |
| cox4i23 | 5.4E-11 | 0.5021789 | 0.478 | 0.32 | 1.75E-06 | Lig/Ten | cox4i2 |
| eef1g4 | 5.7E-11 | 0.2379046 | 0.971 | 0.952 | 1.85E-06 | Lig/Ten | eef1g |
| rcn35 | 7.2E-11 | 0.5034106 | 0.412 | 0.262 | 2.33E-06 | Lig/Ten | rcn3 |
| fzd42 | 9.5E-11 | 1.0374109 | 0.122 | 0.052 | 3.08E-06 | Lig/Ten | fzd4 |
| btbd6b2 | 1E-10 | 1.213367 | 0.133 | 0.06 | 3.37E-06 | Lig/Ten | btbd6b |
| antxr1b2 | 1.1E-10 | 1.0235624 | 0.119 | 0.051 | 3.68E-06 | Lig/Ten | antxr1b |
| CABZ01092781.11 | 1.2E-10 | 1.0766607 | 0.117 | 0.05 | 3.75E-06 | Lig/Ten | CABZ01092781.1 |
| tmem473 | 2E-10 | 0.7990652 | 0.27 | 0.16 | 6.62E-06 | Lig/Ten | tmem47 |
| CTBP14 | 2E-10 | 0.6077453 | 0.285 | 0.168 | 6.67E-06 | Lig/Ten | CTBP1 |
| mxra8b5 | 2.1E-10 | 0.4387944 | 0.562 | 0.407 | 6.96E-06 | Lig/Ten | mxra8b |

| | | | | | | | |
|---|---|---|---|---|---|---|---|
| selenom4 | 2.4E-10 | 0.6769567 | 0.285 | 0.171 | 7.75E-06 | Lig/Ten | selenom |
| cremb2 | 2.5E-10 | 0.9467767 | 0.106 | 0.044 | 8.26E-06 | Lig/Ten | cremb |
| stt3b3 | 2.6E-10 | 0.6953543 | 0.204 | 0.108 | 8.58E-06 | Lig/Ten | stt3b |
| adam17a2 | 3E-10 | 0.7048908 | 0.19 | 0.098 | 9.65E-06 | Lig/Ten | adam17a |
| eef2b7 | 3E-10 | 0.2473295 | 0.956 | 0.909 | 9.8E-06 | Lig/Ten | eef2b |
| sdcbp24 | 3.5E-10 | 0.4021904 | 0.728 | 0.598 | 1.14E-05 | Lig/Ten | sdcbp2 |
| tmem39a4 | 3.6E-10 | 0.7592518 | 0.257 | 0.15 | 1.17E-05 | Lig/Ten | tmem39a |
| atp6ap1b3 | 4.1E-10 | 1.0788107 | 0.144 | 0.069 | 1.34E-05 | Lig/Ten | atp6ap1b |
| ctsc3 | 4.2E-10 | 0.8157824 | 0.144 | 0.067 | 1.38E-05 | Lig/Ten | ctsc |
| vkorc12 | 4.3E-10 | 1.0313639 | 0.115 | 0.05 | 1.41E-05 | Lig/Ten | vkorc1 |
| add15 | 4.7E-10 | 0.637617 | 0.243 | 0.138 | 1.54E-05 | Lig/Ten | add1 |
| itih53 | 5.3E-10 | 0.5158333 | 0.146 | 0.069 | 1.71E-05 | Lig/Ten | itih5 |
| ppib5 | 5.6E-10 | 0.2651388 | 0.954 | 0.917 | 1.83E-05 | Lig/Ten | ppib |
| kank47 | 5.8E-10 | 0.8123874 | 0.195 | 0.103 | 1.89E-05 | Lig/Ten | kank4 |
| nfkbiab6 | 5.8E-10 | 0.4679066 | 0.723 | 0.583 | 1.9E-05 | Lig/Ten | nfkbiab |
| boc7 | 6.3E-10 | 0.440397 | 0.52 | 0.372 | 2.06E-05 | Lig/Ten | boc |
| chp11 | 6.5E-10 | 0.9621876 | 0.122 | 0.054 | 2.12E-05 | Lig/Ten | chp1 |
| rps283 | 7.9E-10 | 0.2449002 | 0.993 | 0.978 | 2.57E-05 | Lig/Ten | rps28 |
| xbp14 | 8E-10 | 0.2915817 | 0.852 | 0.761 | 2.6E-05 | Lig/Ten | xbp1 |
| ddx215 | 8.7E-10 | 0.4722754 | 0.487 | 0.342 | 2.82E-05 | Lig/Ten | ddx21 |
| tppp33 | 9.8E-10 | 0.9145995 | 0.128 | 0.059 | 3.2E-05 | Lig/Ten | tppp3 |
| ckap44 | 1.3E-09 | 0.4456742 | 0.646 | 0.518 | 4.24E-05 | Lig/Ten | ckap4 |
| lpar15 | 1.4E-09 | 0.4729864 | 0.301 | 0.181 | 4.41E-05 | Lig/Ten | lpar1 |
| adh8a4 | 1.6E-09 | 0.5552295 | 0.33 | 0.211 | 5.26E-05 | Lig/Ten | adh8a |
| crtap3 | 1.8E-09 | 0.4871994 | 0.46 | 0.319 | 5.97E-05 | Lig/Ten | crtap |
| calua5 | 1.9E-09 | 0.5190788 | 0.456 | 0.326 | 6.3E-05 | Lig/Ten | calua |
| prdx44 | 2.1E-09 | 0.4768752 | 0.502 | 0.364 | 6.67E-05 | Lig/Ten | prdx4 |
| phldb1a3 | 2.2E-09 | 1.1391693 | 0.1 | 0.042 | 7.01E-05 | Lig/Ten | phldb1a |
| dok1b | 2.5E-09 | 1.2431652 | 0.1 | 0.042 | 8.06E-05 | Lig/Ten | dok1b |
| p4hb4 | 2.7E-09 | 0.322908 | 0.719 | 0.573 | 8.87E-05 | Lig/Ten | p4hb |
| serpine15 | 3.1E-09 | 0.4586792 | 0.834 | 0.762 | 9.95E-05 | Lig/Ten | serpine1 |
| ssr25 | 3.4E-09 | 0.3114101 | 0.81 | 0.69 | 0.000109 | Lig/Ten | ssr2 |
| sec61g4 | 3.4E-09 | 0.2191003 | 0.965 | 0.924 | 0.000111 | Lig/Ten | sec61g |
| psap5 | 3.9E-09 | 0.4710208 | 0.487 | 0.349 | 0.000126 | Lig/Ten | psap |
| zfp36l1b8 | 4.3E-09 | 0.3900988 | 0.688 | 0.546 | 0.000141 | Lig/Ten | zfp36l1b |
| cnpy43 | 4.7E-09 | 0.7673136 | 0.217 | 0.124 | 0.000152 | Lig/Ten | cnpy4 |
| olfml3a4 | 4.8E-09 | 0.667296 | 0.378 | 0.258 | 0.000157 | Lig/Ten | olfml3a |
| hbegfa6 | 5.1E-09 | 0.712747 | 0.52 | 0.402 | 0.000165 | Lig/Ten | hbegfa |
| mhc1zba6 | 5.6E-09 | 0.4045895 | 0.715 | 0.583 | 0.00018 | Lig/Ten | mhc1zba |
| barx16 | 5.7E-09 | 0.5771947 | 0.223 | 0.128 | 0.000186 | Lig/Ten | barx1 |
| jun5 | 6.1E-09 | 0.3109773 | 0.916 | 0.854 | 0.000197 | Lig/Ten | jun |
| srgap25 | 6.6E-09 | 0.7486701 | 0.162 | 0.084 | 0.000215 | Lig/Ten | srgap2 |
| cadm45 | 7.4E-09 | 0.7962407 | 0.15 | 0.076 | 0.000241 | Lig/Ten | cadm4 |
| rbms2b2 | 7.5E-09 | 0.6732405 | 0.131 | 0.062 | 0.000244 | Lig/Ten | rbms2b |
| egfl66 | 8.8E-09 | 1.0642552 | 0.117 | 0.055 | 0.000288 | Lig/Ten | egfl6 |
| fosl26 | 9.8E-09 | 0.3605854 | 0.573 | 0.428 | 0.00032 | Lig/Ten | fosl2 |
| kdelr2a4 | 9.9E-09 | 0.3693648 | 0.533 | 0.39 | 0.000323 | Lig/Ten | kdelr2a |
| chid15 | 1.1E-08 | 0.8568396 | 0.162 | 0.085 | 0.000342 | Lig/Ten | chid1 |

| | | | | | | | |
|---|---|---|---|---|---|---|---|
| zgc:1750885 | 1.1E-08 | 0.5443169 | 0.394 | 0.273 | 0.000361 | Lig/Ten | zgc:175088 |
| tagapa1 | 1.2E-08 | 0.8613124 | 0.148 | 0.076 | 0.000383 | Lig/Ten | tagapa |
| pgk15 | 1.2E-08 | 0.549297 | 0.319 | 0.207 | 0.000386 | Lig/Ten | pgk1 |
| slc25a56 | 1.3E-08 | 0.2350493 | 0.98 | 0.963 | 0.000408 | Lig/Ten | slc25a5 |
| pxdn1 | 1.3E-08 | 1.0048654 | 0.1 | 0.044 | 0.000417 | Lig/Ten | pxdn |
| golim4a3 | 1.5E-08 | 0.6480651 | 0.208 | 0.118 | 0.000473 | Lig/Ten | golim4a |
| C7H16orf874 | 1.6E-08 | 0.8470858 | 0.215 | 0.127 | 0.000513 | Lig/Ten | C7H16orf87 |
| hivep15 | 1.6E-08 | 0.4804273 | 0.254 | 0.153 | 0.000527 | Lig/Ten | hivep1 |
| rgl12 | 1.7E-08 | 0.4964977 | 0.217 | 0.125 | 0.000556 | Lig/Ten | rgl1 |
| svep14 | 1.7E-08 | 0.6058443 | 0.241 | 0.144 | 0.000567 | Lig/Ten | svep1 |
| myo1ea4 | 1.8E-08 | 0.9904217 | 0.155 | 0.082 | 0.000586 | Lig/Ten | myo1ea |
| lhfpl2a6 | 1.9E-08 | 0.6910786 | 0.192 | 0.107 | 0.000609 | Lig/Ten | lhfpl2a |
| ptmab6 | 1.9E-08 | 0.2491826 | 0.847 | 0.714 | 0.000626 | Lig/Ten | ptmab |
| nenf4 | 2E-08 | 0.5277353 | 0.179 | 0.097 | 0.000641 | Lig/Ten | nenf |
| selenof5 | 2E-08 | 0.3570593 | 0.6 | 0.467 | 0.000666 | Lig/Ten | selenof |
| b3glcta1 | 2.1E-08 | 1.1548175 | 0.108 | 0.05 | 0.000676 | Lig/Ten | b3glcta |
| net17 | 2.1E-08 | 0.3547447 | 0.511 | 0.372 | 0.000679 | Lig/Ten | net1 |
| npc24 | 2.1E-08 | 0.5135021 | 0.471 | 0.346 | 0.000693 | Lig/Ten | npc2 |
| pdzrn3b3 | 2.3E-08 | 0.9453179 | 0.15 | 0.079 | 0.000738 | Lig/Ten | pdzrn3b |
| pgam1a5 | 2.4E-08 | 0.447225 | 0.294 | 0.187 | 0.000776 | Lig/Ten | pgam1a |
| peli1a6 | 2.4E-08 | 0.7065947 | 0.219 | 0.129 | 0.000788 | Lig/Ten | peli1a |
| bmp65 | 2.5E-08 | 0.9219264 | 0.124 | 0.06 | 0.000804 | Lig/Ten | bmp6 |
| mgst3b2 | 2.8E-08 | 0.416051 | 0.518 | 0.384 | 0.000902 | Lig/Ten | mgst3b |
| rin26 | 2.9E-08 | 0.8149723 | 0.195 | 0.112 | 0.000938 | Lig/Ten | rin2 |
| zgc:1132084 | 3E-08 | 0.4540829 | 0.301 | 0.191 | 0.000984 | Lig/Ten | zgc:113208 |
| mych6 | 3.1E-08 | 0.330126 | 0.515 | 0.379 | 0.000993 | Lig/Ten | mych |
| postna10 | 3.2E-08 | 0.2227619 | 0.759 | 0.59 | 0.001045 | Lig/Ten | postna |
| arf52 | 3.2E-08 | 0.5375301 | 0.425 | 0.309 | 0.001053 | Lig/Ten | arf5 |
| tram15 | 3.4E-08 | 0.4929712 | 0.407 | 0.287 | 0.001099 | Lig/Ten | tram1 |
| acsl1a1 | 3.6E-08 | 1.1564731 | 0.142 | 0.075 | 0.001165 | Lig/Ten | acsl1a |
| pcdh18b4 | 3.7E-08 | 0.5541533 | 0.332 | 0.219 | 0.001198 | Lig/Ten | pcdh18b |
| lrp55 | 4.4E-08 | 0.6625594 | 0.197 | 0.112 | 0.001436 | Lig/Ten | lrp5 |
| gsk3ba3 | 4.8E-08 | 0.7304378 | 0.237 | 0.145 | 0.001563 | Lig/Ten | gsk3ba |
| lrp1ab6 | 5.3E-08 | 0.4753681 | 0.462 | 0.341 | 0.001722 | Lig/Ten | lrp1ab |
| rimkla3 | 5.7E-08 | 0.9547612 | 0.133 | 0.068 | 0.001838 | Lig/Ten | rimkla |
| calub2 | 6.5E-08 | 0.9122392 | 0.131 | 0.066 | 0.002126 | Lig/Ten | calub |
| si:ch1073-385f13.32 | 7.6E-08 | 0.7487093 | 0.124 | 0.061 | 0.002481 | Lig/Ten | si:ch1073-385f13.3 |
| dvl21 | 7.9E-08 | 0.6554899 | 0.217 | 0.129 | 0.002575 | Lig/Ten | dvl2 |
| tsku6 | 8.2E-08 | 0.2859403 | 0.465 | 0.318 | 0.002661 | Lig/Ten | tsku |
| sec61a16 | 8.2E-08 | 0.3684912 | 0.56 | 0.42 | 0.002669 | Lig/Ten | sec61a1 |
| sox4a6 | 8.4E-08 | 0.6592576 | 0.296 | 0.194 | 0.002722 | Lig/Ten | sox4a |
| maff8 | 8.9E-08 | 0.5802754 | 0.327 | 0.223 | 0.002885 | Lig/Ten | maff |
| cxcl18b7 | 9.4E-08 | 0.3803816 | 0.79 | 0.702 | 0.003045 | Lig/Ten | cxcl18b |
| sptan14 | 9.4E-08 | 0.3777253 | 0.365 | 0.244 | 0.003063 | Lig/Ten | sptan1 |
| cpne33 | 9.6E-08 | 0.6986657 | 0.188 | 0.108 | 0.003124 | Lig/Ten | cpne3 |
| serp16 | 1E-07 | 0.4277894 | 0.436 | 0.315 | 0.003336 | Lig/Ten | serp1 |
| sat1a.210 | 1E-07 | 0.370484 | 0.591 | 0.457 | 0.003414 | Lig/Ten | sat1a.2 |
| dap1b.15 | 1.2E-07 | 0.2005124 | 0.673 | 0.506 | 0.003968 | Lig/Ten | dap1b.1 |

| | | | | | | | |
|---|---|---|---|---|---|---|---|
| vat15 | 1.2E-07 | 0.3346338 | 0.438 | 0.315 | 0.00403 | Lig/Ten | vat1 |
| phc2b4 | 1.3E-07 | 0.9322102 | 0.142 | 0.075 | 0.004182 | Lig/Ten | phc2b |
| sulf2b7 | 1.3E-07 | 0.5239996 | 0.263 | 0.165 | 0.004359 | Lig/Ten | sulf2b |
| prelid3b6 | 1.4E-07 | 0.3556296 | 0.485 | 0.357 | 0.004434 | Lig/Ten | prelid3b |
| tpt14 | 1.4E-07 | 0.2204105 | 0.942 | 0.927 | 0.004531 | Lig/Ten | tpt1 |
| tnrc6a6 | 1.5E-07 | 0.6361983 | 0.376 | 0.272 | 0.004864 | Lig/Ten | tnrc6a |
| enosf11 | 1.5E-07 | 0.9254838 | 0.104 | 0.049 | 0.004968 | Lig/Ten | enosf1 |
| trib24 | 1.5E-07 | 0.6166304 | 0.162 | 0.088 | 0.005023 | Lig/Ten | trib2 |
| igf2b5 | 1.6E-07 | 0.5103211 | 0.551 | 0.436 | 0.005283 | Lig/Ten | igf2b |
| kazald36 | 1.7E-07 | 0.2162746 | 0.387 | 0.261 | 0.005633 | Lig/Ten | kazald3 |
| tmed24 | 1.7E-07 | 0.3859358 | 0.52 | 0.391 | 0.005662 | Lig/Ten | tmed2 |
| ifrd17 | 1.8E-07 | 0.4423877 | 0.396 | 0.282 | 0.005783 | Lig/Ten | ifrd1 |
| dbn13 | 1.9E-07 | 0.5413257 | 0.303 | 0.2 | 0.006129 | Lig/Ten | dbn1 |
| pttg1ipb2 | 1.9E-07 | 0.8019011 | 0.131 | 0.067 | 0.006265 | Lig/Ten | pttg1ipb |
| anapc163 | 1.9E-07 | 0.7042454 | 0.235 | 0.147 | 0.006267 | Lig/Ten | anapc16 |
| rgmb5 | 2E-07 | 0.6095093 | 0.201 | 0.119 | 0.006511 | Lig/Ten | rgmb |
| dad15 | 2.1E-07 | 0.4487588 | 0.513 | 0.399 | 0.00688 | Lig/Ten | dad1 |
| hmgb1b4 | 2.2E-07 | 0.4186415 | 0.365 | 0.252 | 0.007 | Lig/Ten | hmgb1b |
| zgc:77486.17 | 2.5E-07 | 0.486374 | 0.265 | 0.169 | 0.00813 | Lig/Ten | zgc:77486.1 |
| ralba3 | 2.5E-07 | 0.763854 | 0.162 | 0.09 | 0.008274 | Lig/Ten | ralba |
| nucb2b4 | 2.6E-07 | 0.7446515 | 0.197 | 0.118 | 0.008344 | Lig/Ten | nucb2b |
| mcfd23 | 2.6E-07 | 0.6129873 | 0.21 | 0.127 | 0.00854 | Lig/Ten | mcfd2 |
| plod26 | 2.7E-07 | 1.0584545 | 0.135 | 0.072 | 0.008686 | Lig/Ten | plod2 |
| pla2g151 | 2.9E-07 | 0.9372923 | 0.111 | 0.055 | 0.009469 | Lig/Ten | pla2g15 |
| os93 | 2.9E-07 | 0.6373758 | 0.314 | 0.215 | 0.009553 | Lig/Ten | os9 |
| itm2ba5 | 3E-07 | 0.2394237 | 0.847 | 0.741 | 0.009911 | Lig/Ten | itm2ba |
| si:dkey-42i9.63 | 3.2E-07 | 0.4529068 | 0.296 | 0.195 | 0.010436 | Lig/Ten | si:dkey-42i9.6 |
| tcf127 | 3.2E-07 | 0.217145 | 0.677 | 0.522 | 0.010452 | Lig/Ten | tcf12 |
| arhgef405 | 3.4E-07 | 0.5341168 | 0.29 | 0.191 | 0.010933 | Lig/Ten | arhgef40 |
| serinc56 | 3.8E-07 | 0.4436442 | 0.35 | 0.239 | 0.012287 | Lig/Ten | serinc5 |
| nr4a34 | 3.9E-07 | 0.6318015 | 0.104 | 0.051 | 0.012525 | Lig/Ten | nr4a3 |
| aldoaa5 | 3.9E-07 | 0.4231083 | 0.487 | 0.368 | 0.012779 | Lig/Ten | aldoaa |
| emp23 | 4E-07 | 0.4732554 | 0.241 | 0.152 | 0.013054 | Lig/Ten | emp2 |
| thbs2a8 | 4.7E-07 | 0.3807506 | 0.305 | 0.203 | 0.015445 | Lig/Ten | thbs2a |
| f11r.12 | 4.8E-07 | 0.642099 | 0.142 | 0.077 | 0.015589 | Lig/Ten | f11r.1 |
| klf6a4 | 5E-07 | 0.3755951 | 0.845 | 0.746 | 0.01618 | Lig/Ten | klf6a |
| specc1la4 | 5.2E-07 | 0.6132274 | 0.146 | 0.08 | 0.016785 | Lig/Ten | specc1la |
| rab2a4 | 5.4E-07 | 0.3260128 | 0.653 | 0.521 | 0.017517 | Lig/Ten | rab2a |
| skilb3 | 5.8E-07 | 0.6710881 | 0.144 | 0.079 | 0.018819 | Lig/Ten | skilb |
| mmp15a7 | 6.1E-07 | 0.7137693 | 0.15 | 0.083 | 0.019708 | Lig/Ten | mmp15a |
| etv5b6 | 6.3E-07 | 0.5940183 | 0.173 | 0.1 | 0.020434 | Lig/Ten | etv5b |
| dipk1c8 | 6.4E-07 | 0.5105666 | 0.192 | 0.115 | 0.020876 | Lig/Ten | dipk1c |
| ccdc3b4 | 6.6E-07 | 0.5067489 | 0.206 | 0.125 | 0.021533 | Lig/Ten | ccdc3b |
| si:ch1073-392o20.26 | 6.9E-07 | 0.3765852 | 0.35 | 0.242 | 0.022446 | Lig/Ten | si:ch1073-392o20.[illegible] |
| kdelr2b4 | 7.1E-07 | 0.3029779 | 0.44 | 0.32 | 0.023141 | Lig/Ten | kdelr2b |
| id18 | 7.4E-07 | 0.3921546 | 0.741 | 0.641 | 0.024124 | Lig/Ten | id1 |
| c1qtnf6b3 | 9.3E-07 | 0.5988576 | 0.139 | 0.076 | 0.030321 | Lig/Ten | c1qtnf6b |
| tmed94 | 9.9E-07 | 0.3647527 | 0.334 | 0.227 | 0.032337 | Lig/Ten | tmed9 |

| | | | | | | | |
|---|---|---|---|---|---|---|---|
| trim8b2 | 1E-06 | 0.5809085 | 0.217 | 0.136 | 0.033155 | Lig/Ten | trim8b |
| prss237 | 1.1E-06 | 0.3829171 | 0.303 | 0.2 | 0.035728 | Lig/Ten | prss23 |
| c65 | 1.1E-06 | 0.5547036 | 0.135 | 0.073 | 0.036017 | Lig/Ten | c6 |
| eif4ebp3l3 | 1.2E-06 | 0.5241811 | 0.239 | 0.154 | 0.038129 | Lig/Ten | eif4ebp3l |
| sox11b4 | 1.2E-06 | 0.4450405 | 0.31 | 0.21 | 0.039382 | Lig/Ten | sox11b |
| rab3il12 | 1.3E-06 | 0.6230796 | 0.124 | 0.065 | 0.041292 | Lig/Ten | rab3il1 |
| lgmn5 | 1.3E-06 | 0.6908676 | 0.261 | 0.171 | 0.042311 | Lig/Ten | lgmn |
| fnbp1l2 | 1.3E-06 | 0.687908 | 0.126 | 0.067 | 0.043675 | Lig/Ten | fnbp1l |
| tmem2585 | 1.4E-06 | 0.2970238 | 0.706 | 0.588 | 0.045432 | Lig/Ten | tmem258 |
| kdm6ba5 | 1.5E-06 | 0.4616825 | 0.321 | 0.221 | 0.048611 | Lig/Ten | kdm6ba |
| tmem176l.15 | 1.6E-06 | 0.2165142 | 0.852 | 0.741 | 0.051256 | Lig/Ten | tmem176l.1 |
| mtus1b1 | 1.6E-06 | 0.8177253 | 0.126 | 0.068 | 0.052223 | Lig/Ten | mtus1b |
| bahcc1a5 | 1.6E-06 | 0.8316127 | 0.153 | 0.087 | 0.053421 | Lig/Ten | bahcc1a |
| amotl2a3 | 1.7E-06 | 0.6943784 | 0.239 | 0.157 | 0.054985 | Lig/Ten | amotl2a |
| creg12 | 1.7E-06 | 0.6540164 | 0.133 | 0.072 | 0.055294 | Lig/Ten | creg1 |
| eml12 | 1.7E-06 | 0.8046815 | 0.108 | 0.056 | 0.055537 | Lig/Ten | eml1 |
| zeb1b5 | 1.8E-06 | 0.461695 | 0.155 | 0.088 | 0.059074 | Lig/Ten | zeb1b |
| pde4ba5 | 1.8E-06 | 0.5340546 | 0.192 | 0.117 | 0.06009 | Lig/Ten | pde4ba |
| glis2a5 | 1.9E-06 | 0.4666496 | 0.301 | 0.204 | 0.062529 | Lig/Ten | glis2a |
| nr1d15 | 2E-06 | 0.5326424 | 0.215 | 0.135 | 0.063851 | Lig/Ten | nr1d1 |
| dele11 | 2E-06 | 0.6784423 | 0.128 | 0.07 | 0.065643 | Lig/Ten | dele1 |
| slc25a25a6 | 2.3E-06 | 0.3521762 | 0.319 | 0.219 | 0.073564 | Lig/Ten | slc25a25a |
| igfbp2b4 | 2.3E-06 | 0.9574333 | 0.128 | 0.07 | 0.074806 | Lig/Ten | igfbp2b |
| atp6v0e11 | 2.5E-06 | 0.3779509 | 0.48 | 0.375 | 0.081761 | Lig/Ten | atp6v0e1 |
| frem1b5 | 2.6E-06 | 0.9530369 | 0.137 | 0.078 | 0.085122 | Lig/Ten | frem1b |
| il13ra15 | 2.8E-06 | 0.6502499 | 0.148 | 0.085 | 0.090213 | Lig/Ten | il13ra1 |
| si:dkey-188i13.114 | 2.9E-06 | 0.4256114 | 0.294 | 0.203 | 0.094498 | Lig/Ten | si:dkey-188i13.11 |
| mafba7 | 2.9E-06 | 0.4292258 | 0.498 | 0.39 | 0.095823 | Lig/Ten | mafba |
| si:cabz01069012.25 | 3.1E-06 | 0.4214193 | 0.175 | 0.103 | 0.101634 | Lig/Ten | si:cabz01069012.2 |
| hsp90b13 | 3.1E-06 | 0.3682591 | 0.454 | 0.345 | 0.101641 | Lig/Ten | hsp90b1 |
| si:ch73-44m9.3.14 | 3.2E-06 | 0.4408856 | 0.168 | 0.099 | 0.103524 | Lig/Ten | si:ch73-44m9.3.1 |
| si:dkey-11f4.206 | 3.3E-06 | 0.2602804 | 0.615 | 0.488 | 0.106315 | Lig/Ten | si:dkey-11f4.20 |
| svild4 | 3.3E-06 | 0.3514162 | 0.365 | 0.256 | 0.107144 | Lig/Ten | svild |
| tram24 | 3.5E-06 | 0.6197915 | 0.179 | 0.11 | 0.113557 | Lig/Ten | tram2 |
| ctsla5 | 3.7E-06 | 0.4361714 | 0.799 | 0.721 | 0.120607 | Lig/Ten | ctsla |
| amotl2b2 | 3.8E-06 | 0.8735314 | 0.139 | 0.08 | 0.122889 | Lig/Ten | amotl2b |
| ostc4 | 4.5E-06 | 0.3640757 | 0.398 | 0.291 | 0.146255 | Lig/Ten | ostc |
| dnajc3b3 | 5.1E-06 | 0.5225453 | 0.115 | 0.061 | 0.166672 | Lig/Ten | dnajc3b |
| serpinh1a5 | 5.5E-06 | 0.3711208 | 0.338 | 0.238 | 0.17916 | Lig/Ten | serpinh1a |
| golim4a.13 | 5.8E-06 | 0.6707562 | 0.162 | 0.097 | 0.187506 | Lig/Ten | golim4a.1 |
| cdc42ep54 | 6.3E-06 | 0.444538 | 0.405 | 0.3 | 0.204242 | Lig/Ten | cdc42ep5 |
| fam3c4 | 6.4E-06 | 0.5588379 | 0.168 | 0.102 | 0.20652 | Lig/Ten | fam3c |
| ckbb5 | 6.4E-06 | 0.4191472 | 0.285 | 0.199 | 0.209487 | Lig/Ten | ckbb |
| jpt1b4 | 6.7E-06 | 0.2673677 | 0.77 | 0.691 | 0.217708 | Lig/Ten | jpt1b |
| lamb1a1 | 7.3E-06 | 0.7536647 | 0.102 | 0.053 | 0.237456 | Lig/Ten | lamb1a |
| CABZ01058650.14 | 7.6E-06 | 0.698405 | 0.128 | 0.073 | 0.246936 | Lig/Ten | CABZ01058650.1 |
| CDPF14 | 7.7E-06 | 0.5742037 | 0.261 | 0.179 | 0.250016 | Lig/Ten | CDPF1 |
| ppp3r1b3 | 7.7E-06 | 0.524879 | 0.208 | 0.134 | 0.251288 | Lig/Ten | ppp3r1b |

| | | | | | | | |
|---|---|---|---|---|---|---|---|
| sh3pxd2ab2 | 7.7E-06 | 0.6634327 | 0.137 | 0.078 | 0.25177 | Lig/Ten | sh3pxd2ab |
| tmed36 | 8.1E-06 | 0.6811702 | 0.159 | 0.096 | 0.263719 | Lig/Ten | tmed3 |
| p3h23 | 8.3E-06 | 0.5549879 | 0.115 | 0.062 | 0.270456 | Lig/Ten | p3h2 |
| nfkbie3 | 8.9E-06 | 0.8492559 | 0.131 | 0.075 | 0.290524 | Lig/Ten | nfkbie |
| slc30a74 | 9E-06 | 0.4784611 | 0.181 | 0.112 | 0.292684 | Lig/Ten | slc30a7 |
| rer14 | 9.2E-06 | 0.2760957 | 0.301 | 0.206 | 0.298812 | Lig/Ten | rer1 |
| zgc:922425 | 9.4E-06 | 0.4272107 | 0.179 | 0.111 | 0.307159 | Lig/Ten | zgc:92242 |
| her66 | 1.1E-05 | 0.56168 | 0.52 | 0.433 | 0.358313 | Lig/Ten | her6 |
| camk1b3 | 1.1E-05 | 0.6162191 | 0.108 | 0.058 | 0.360109 | Lig/Ten | camk1b |
| pdia56 | 1.1E-05 | 0.3795336 | 0.248 | 0.166 | 0.367371 | Lig/Ten | pdia5 |
| sh3kbp12 | 1.2E-05 | 0.4655364 | 0.206 | 0.132 | 0.382176 | Lig/Ten | sh3kbp1 |
| fap3 | 1.3E-05 | 0.8941076 | 0.117 | 0.066 | 0.410379 | Lig/Ten | fap |
| pdia64 | 1.4E-05 | 0.436395 | 0.283 | 0.2 | 0.458379 | Lig/Ten | pdia6 |
| slc38a5b2 | 1.5E-05 | 0.3694833 | 0.456 | 0.353 | 0.479603 | Lig/Ten | slc38a5b |
| sec61b5 | 1.5E-05 | 0.2766658 | 0.706 | 0.583 | 0.482731 | Lig/Ten | sec61b |
| tomm20b5 | 1.5E-05 | 0.2886949 | 0.456 | 0.343 | 0.496273 | Lig/Ten | tomm20b |
| yif1a3 | 1.5E-05 | 0.5407072 | 0.195 | 0.124 | 0.50013 | Lig/Ten | yif1a |
| tfe3a4 | 1.5E-05 | 0.7066697 | 0.155 | 0.095 | 0.501882 | Lig/Ten | tfe3a |
| tiparp5 | 1.6E-05 | 0.5954146 | 0.195 | 0.125 | 0.50474 | Lig/Ten | tiparp |
| sec22ba4 | 1.7E-05 | 0.5771424 | 0.184 | 0.118 | 0.550694 | Lig/Ten | sec22ba |
| sorbs16 | 1.8E-05 | 0.437184 | 0.208 | 0.134 | 0.573994 | Lig/Ten | sorbs1 |
| bcl2a4 | 1.8E-05 | 0.3419439 | 0.235 | 0.155 | 0.598143 | Lig/Ten | bcl2a |
| cyp20a13 | 1.9E-05 | 0.4848681 | 0.179 | 0.113 | 0.62543 | Lig/Ten | cyp20a1 |
| ift203 | 1.9E-05 | 0.9025805 | 0.108 | 0.06 | 0.630199 | Lig/Ten | ift20 |
| bckdk3 | 2E-05 | 0.4726197 | 0.148 | 0.089 | 0.642313 | Lig/Ten | bckdk |
| bri32 | 2E-05 | 0.7696128 | 0.108 | 0.06 | 0.654403 | Lig/Ten | bri3 |
| fkbp10a5 | 2.2E-05 | 0.5133111 | 0.206 | 0.135 | 0.704847 | Lig/Ten | fkbp10a |
| cebpg2 | 2.2E-05 | 0.4642332 | 0.21 | 0.138 | 0.711987 | Lig/Ten | cebpg |
| adamts122 | 2.3E-05 | 0.9051545 | 0.1 | 0.054 | 0.738302 | Lig/Ten | adamts12 |
| mfge8a5 | 2.4E-05 | 0.5464344 | 0.186 | 0.12 | 0.767567 | Lig/Ten | mfge8a |
| cpvl5 | 2.4E-05 | 0.6549812 | 0.153 | 0.093 | 0.776045 | Lig/Ten | cpvl |
| slc35e13 | 2.4E-05 | 0.7817085 | 0.104 | 0.057 | 0.776365 | Lig/Ten | slc35e1 |
| si:dkey-238c7.164 | 2.8E-05 | 0.4008583 | 0.241 | 0.165 | 0.914512 | Lig/Ten | si:dkey-238c7.16 |
| npc13 | 3E-05 | 0.4392567 | 0.128 | 0.074 | 0.959495 | Lig/Ten | npc1 |
| si:ch211-11k18.42 | 3E-05 | 0.6680393 | 0.106 | 0.059 | 0.965362 | Lig/Ten | si:ch211-11k18.4 |
| atp6v1ba2 | 3E-05 | 0.5524993 | 0.104 | 0.057 | 0.967949 | Lig/Ten | atp6v1ba |
| COX7A23 | 3.2E-05 | 0.4598384 | 0.199 | 0.131 | 1 | Lig/Ten | COX7A2 |
| psat13 | 3.3E-05 | 0.6309947 | 0.119 | 0.069 | 1 | Lig/Ten | psat1 |
| clip24 | 3.4E-05 | 0.3621153 | 0.166 | 0.103 | 1 | Lig/Ten | clip2 |
| ppt15 | 3.7E-05 | 0.2903534 | 0.445 | 0.344 | 1 | Lig/Ten | ppt1 |
| ndfip14 | 3.8E-05 | 0.2474349 | 0.482 | 0.374 | 1 | Lig/Ten | ndfip1 |
| zgc:1527915 | 3.9E-05 | 0.8837872 | 0.166 | 0.106 | 1 | Lig/Ten | zgc:152791 |
| calr3b5 | 4.3E-05 | 0.3010538 | 0.365 | 0.27 | 1 | Lig/Ten | calr3b |
| csrnp1b6 | 4.3E-05 | 0.5343654 | 0.265 | 0.187 | 1 | Lig/Ten | csrnp1b |
| abi1a3 | 4.4E-05 | 0.6832937 | 0.117 | 0.068 | 1 | Lig/Ten | abi1a |
| itgb1a4 | 4.5E-05 | 0.2749223 | 0.358 | 0.264 | 1 | Lig/Ten | itgb1a |
| rab18b3 | 4.6E-05 | 0.4569483 | 0.179 | 0.115 | 1 | Lig/Ten | rab18b |
| ier5l4 | 4.7E-05 | 0.2615232 | 0.427 | 0.319 | 1 | Lig/Ten | ier5l |

| | | | | | | | |
|---|---|---|---|---|---|---|---|
| rab18a3 | 4.8E-05 | 0.3895093 | 0.228 | 0.154 | 1 | Lig/Ten | rab18a |
| tspan356 | 4.8E-05 | 0.4233921 | 0.19 | 0.124 | 1 | Lig/Ten | tspan35 |
| lamtor43 | 4.9E-05 | 0.580426 | 0.164 | 0.103 | 1 | Lig/Ten | lamtor4 |
| tuba8l33 | 5.1E-05 | 0.3064843 | 0.531 | 0.425 | 1 | Lig/Ten | tuba8l3 |
| cst14a.24 | 5.2E-05 | 0.4058031 | 0.451 | 0.364 | 1 | Lig/Ten | cst14a.2 |
| tril7 | 6E-05 | 0.2480931 | 0.17 | 0.107 | 1 | Lig/Ten | tril |
| nupr1a5 | 6.1E-05 | 0.3895955 | 0.345 | 0.257 | 1 | Lig/Ten | nupr1a |
| tmbim44 | 6.7E-05 | 0.4292065 | 0.179 | 0.116 | 1 | Lig/Ten | tmbim4 |
| cox7a2l1 | 7.1E-05 | 0.2743306 | 0.232 | 0.158 | 1 | Lig/Ten | cox7a2l |
| zgc:1536754 | 7.4E-05 | 0.3141729 | 0.35 | 0.262 | 1 | Lig/Ten | zgc:153675 |
| bnc25 | 7.7E-05 | 0.4525996 | 0.135 | 0.082 | 1 | Lig/Ten | bnc2 |
| ptgs2a6 | 7.7E-05 | 0.6969097 | 0.192 | 0.13 | 1 | Lig/Ten | ptgs2a |
| si:ch211-195b15.86 | 8.1E-05 | 1.0726745 | 0.144 | 0.091 | 1 | Lig/Ten | si:ch211-195b15.8 |
| dusp27 | 8.1E-05 | 0.6623538 | 0.296 | 0.223 | 1 | Lig/Ten | dusp2 |
| fam114a15 | 8.5E-05 | 0.4384228 | 0.17 | 0.11 | 1 | Lig/Ten | fam114a1 |
| odc15 | 8.5E-05 | 0.5821067 | 0.186 | 0.124 | 1 | Lig/Ten | odc1 |
| si:dkey-235d18.55 | 8.6E-05 | 0.4946047 | 0.155 | 0.098 | 1 | Lig/Ten | si:dkey-235d18.5 |
| rab22a4 | 8.6E-05 | 0.5634158 | 0.159 | 0.102 | 1 | Lig/Ten | rab22a |
| gosr23 | 8.7E-05 | 0.5870548 | 0.146 | 0.092 | 1 | Lig/Ten | gosr2 |
| kif1b4 | 9.3E-05 | 0.6229138 | 0.157 | 0.101 | 1 | Lig/Ten | kif1b |
| vasnb5 | 9.5E-05 | 0.2746594 | 0.215 | 0.144 | 1 | Lig/Ten | vasnb |
| ginm11 | 1E-04 | 0.2184019 | 0.438 | 0.334 | 1 | Lig/Ten | ginm1 |
| plod33 | 0.0001 | 0.3374495 | 0.219 | 0.151 | 1 | Lig/Ten | plod3 |
| sqstm16 | 0.0001 | 0.3829772 | 0.398 | 0.305 | 1 | Lig/Ten | sqstm1 |
| sec31a4 | 0.00011 | 0.4822725 | 0.168 | 0.109 | 1 | Lig/Ten | sec31a |
| eva1ba4 | 0.00011 | 0.4357382 | 0.166 | 0.108 | 1 | Lig/Ten | eva1ba |
| hint16 | 0.00011 | 0.2761339 | 0.489 | 0.386 | 1 | Lig/Ten | hint1 |
| gfra1a3 | 0.00012 | 0.2305973 | 0.1 | 0.056 | 1 | Lig/Ten | gfra1a |
| glud1b1 | 0.00012 | 0.4464554 | 0.166 | 0.107 | 1 | Lig/Ten | glud1b |
| kank35 | 0.00012 | 0.5900656 | 0.131 | 0.08 | 1 | Lig/Ten | kank3 |
| hist2h2l7 | 0.00013 | 0.2582943 | 0.473 | 0.374 | 1 | Lig/Ten | hist2h2l |
| tacc14 | 0.00013 | 0.4426756 | 0.294 | 0.216 | 1 | Lig/Ten | tacc1 |
| adamts15b.16 | 0.00013 | 0.4888304 | 0.148 | 0.093 | 1 | Lig/Ten | adamts15b.1 |
| suds32 | 0.00014 | 0.4541609 | 0.159 | 0.102 | 1 | Lig/Ten | suds3 |
| plxna35 | 0.00014 | 0.4976997 | 0.142 | 0.089 | 1 | Lig/Ten | plxna3 |
| cnpy13 | 0.00015 | 0.4705757 | 0.131 | 0.081 | 1 | Lig/Ten | cnpy1 |
| derl23 | 0.00015 | 0.2417345 | 0.148 | 0.093 | 1 | Lig/Ten | derl2 |
| zmiz1a6 | 0.00015 | 0.5360882 | 0.179 | 0.12 | 1 | Lig/Ten | zmiz1a |
| nabp1a5 | 0.00016 | 0.2610915 | 0.252 | 0.18 | 1 | Lig/Ten | nabp1a |
| dbi5 | 0.00016 | 0.3390369 | 0.549 | 0.442 | 1 | Lig/Ten | dbi |
| jcada4 | 0.00016 | 0.4377381 | 0.181 | 0.122 | 1 | Lig/Ten | jcada |
| si:dkey-25e12.34 | 0.00017 | 0.6824049 | 0.119 | 0.073 | 1 | Lig/Ten | si:dkey-25e12.3 |
| atp6v0ca4 | 0.00017 | 0.2384507 | 0.434 | 0.339 | 1 | Lig/Ten | atp6v0ca |
| klf91 | 0.00017 | 0.8587194 | 0.106 | 0.064 | 1 | Lig/Ten | klf9 |
| epn25 | 0.00017 | 0.5497118 | 0.166 | 0.109 | 1 | Lig/Ten | epn2 |
| rnaseka5 | 0.00018 | 0.4917418 | 0.162 | 0.106 | 1 | Lig/Ten | rnaseka |
| pdia34 | 0.00018 | 0.2478929 | 0.781 | 0.686 | 1 | Lig/Ten | pdia3 |
| lamp1b4 | 0.00019 | 0.4265772 | 0.142 | 0.089 | 1 | Lig/Ten | lamp1b |

| | | | | | | | |
|---|---|---|---|---|---|---|---|
| ap3b1a1 | 0.00019 | 0.4245516 | 0.135 | 0.084 | 1 | Lig/Ten | ap3b1a |
| cux1a5 | 0.0002 | 0.6799396 | 0.113 | 0.069 | 1 | Lig/Ten | cux1a |
| gnsa3 | 0.0002 | 0.280729 | 0.232 | 0.162 | 1 | Lig/Ten | gnsa |
| si:ch211-212k18.72 | 0.0002 | 0.4524465 | 0.108 | 0.064 | 1 | Lig/Ten | si:ch211-212k18.7 |
| ldha6 | 0.00022 | 0.3111723 | 0.31 | 0.23 | 1 | Lig/Ten | ldha |
| sgcg5 | 0.00023 | 0.3577076 | 0.144 | 0.092 | 1 | Lig/Ten | sgcg |
| rgs12b6 | 0.00023 | 0.4734902 | 0.139 | 0.089 | 1 | Lig/Ten | rgs12b |
| zgc:1716791 | 0.00025 | 0.8884609 | 0.111 | 0.068 | 1 | Lig/Ten | zgc:171679 |
| tuft1a5 | 0.00025 | 0.2445681 | 0.686 | 0.591 | 1 | Lig/Ten | tuft1a |
| selenon6 | 0.00026 | 0.237853 | 0.226 | 0.157 | 1 | Lig/Ten | selenon |
| cope5 | 0.00026 | 0.268325 | 0.336 | 0.257 | 1 | Lig/Ten | cope |
| adamtsl35 | 0.00027 | 0.3149754 | 0.115 | 0.07 | 1 | Lig/Ten | adamtsl3 |
| fam120a3 | 0.00027 | 0.2722555 | 0.254 | 0.181 | 1 | Lig/Ten | fam120a |
| CU638870.16 | 0.00027 | 0.2025208 | 0.42 | 0.329 | 1 | Lig/Ten | CU638870.1 |
| tmem594 | 0.00027 | 0.2303811 | 0.403 | 0.314 | 1 | Lig/Ten | tmem59 |
| hopx6 | 0.00028 | 0.2400852 | 0.358 | 0.27 | 1 | Lig/Ten | hopx |
| tgif17 | 0.00028 | 0.467206 | 0.277 | 0.206 | 1 | Lig/Ten | tgif1 |
| fkbp114 | 0.00028 | 0.2475914 | 0.226 | 0.16 | 1 | Lig/Ten | fkbp11 |
| dtnbp1a3 | 0.0003 | 0.4204002 | 0.168 | 0.112 | 1 | Lig/Ten | dtnbp1a |
| txnipa1 | 0.0003 | 0.2872252 | 0.321 | 0.245 | 1 | Lig/Ten | txnipa |
| nr1d2a4 | 0.00031 | 0.2908339 | 0.332 | 0.251 | 1 | Lig/Ten | nr1d2a |
| dnajb1b6 | 0.00031 | 0.2315668 | 0.553 | 0.464 | 1 | Lig/Ten | dnajb1b |
| znf5032 | 0.00031 | 0.3362936 | 0.17 | 0.113 | 1 | Lig/Ten | znf503 |
| anxa2a5 | 0.00031 | 0.2082083 | 0.927 | 0.891 | 1 | Lig/Ten | anxa2a |
| nocta5 | 0.00032 | 0.30453 | 0.17 | 0.114 | 1 | Lig/Ten | nocta |
| ctsd5 | 0.00032 | 0.2934173 | 0.219 | 0.154 | 1 | Lig/Ten | ctsd |
| stim1a | 0.00033 | 0.6792672 | 0.1 | 0.059 | 1 | Lig/Ten | stim1a |
| unc503 | 0.00035 | 0.434529 | 0.128 | 0.081 | 1 | Lig/Ten | unc50 |
| sdhdb3 | 0.00035 | 0.2994051 | 0.188 | 0.128 | 1 | Lig/Ten | sdhdb |
| rpn14 | 0.00036 | 0.4668322 | 0.281 | 0.213 | 1 | Lig/Ten | rpn1 |
| polr2d1 | 0.00037 | 0.3920017 | 0.21 | 0.149 | 1 | Lig/Ten | polr2d |
| ikbip4 | 0.00037 | 0.4422073 | 0.175 | 0.119 | 1 | Lig/Ten | ikbip |
| fhl1b5 | 0.00039 | 0.3867868 | 0.146 | 0.095 | 1 | Lig/Ten | fhl1b |
| tspan172 | 0.00039 | 0.4419431 | 0.124 | 0.077 | 1 | Lig/Ten | tspan17 |
| lsp1b8 | 0.00039 | 0.497959 | 0.33 | 0.258 | 1 | Lig/Ten | lsp1b |
| nfic4 | 0.00042 | 0.6657357 | 0.124 | 0.079 | 1 | Lig/Ten | nfic |
| yaf24 | 0.00043 | 0.5579966 | 0.162 | 0.109 | 1 | Lig/Ten | yaf2 |
| BX936337.14 | 0.00043 | 0.2367013 | 0.478 | 0.389 | 1 | Lig/Ten | BX936337.1 |
| adtrp12 | 0.00043 | 0.4684587 | 0.128 | 0.082 | 1 | Lig/Ten | adtrp1 |
| RAPH15 | 0.00043 | 0.6214829 | 0.139 | 0.091 | 1 | Lig/Ten | RAPH1 |
| bmp55 | 0.00045 | 0.4520659 | 0.137 | 0.088 | 1 | Lig/Ten | bmp5 |
| lrpap14 | 0.00045 | 0.3212933 | 0.25 | 0.181 | 1 | Lig/Ten | lrpap1 |
| sap30bp2 | 0.00046 | 0.307917 | 0.135 | 0.087 | 1 | Lig/Ten | sap30bp |
| arrdc3a5 | 0.00047 | 0.3467038 | 0.232 | 0.165 | 1 | Lig/Ten | arrdc3a |
| pthlha5 | 0.00047 | 0.2861761 | 0.117 | 0.072 | 1 | Lig/Ten | pthlha |
| nutf2l4 | 0.00048 | 0.3725893 | 0.153 | 0.102 | 1 | Lig/Ten | nutf2l |
| lman23 | 0.00048 | 0.2357092 | 0.316 | 0.236 | 1 | Lig/Ten | lman2 |
| pam162 | 0.00051 | 0.3843242 | 0.164 | 0.11 | 1 | Lig/Ten | pam16 |

| | | | | | | | |
|---|---|---|---|---|---|---|---|
| p4ha1a5 | 0.00051 | 0.5280855 | 0.146 | 0.097 | 1 | Lig/Ten | p4ha1a |
| rab134 | 0.00051 | 0.2903224 | 0.294 | 0.223 | 1 | Lig/Ten | rab13 |
| slc31a14 | 0.00051 | 0.2794734 | 0.268 | 0.198 | 1 | Lig/Ten | slc31a1 |
| zgc:1735875 | 0.00052 | 0.4259593 | 0.19 | 0.132 | 1 | Lig/Ten | zgc:173587 |
| sar1b4 | 0.00053 | 0.2036444 | 0.387 | 0.3 | 1 | Lig/Ten | sar1b |
| rspo34 | 0.00055 | 0.3939101 | 0.106 | 0.065 | 1 | Lig/Ten | rspo3 |
| six2a5 | 0.00055 | 0.4064325 | 0.153 | 0.102 | 1 | Lig/Ten | six2a |
| pear15 | 0.00056 | 0.2711509 | 0.19 | 0.131 | 1 | Lig/Ten | pear1 |
| tbc1d171 | 0.00057 | 0.460233 | 0.117 | 0.073 | 1 | Lig/Ten | tbc1d17 |
| tpd52l2b4 | 0.00058 | 0.2530608 | 0.292 | 0.22 | 1 | Lig/Ten | tpd52l2b |
| si:dkey-112e7.22 | 0.00058 | 0.2569077 | 0.303 | 0.226 | 1 | Lig/Ten | si:dkey-112e7.2 |
| nat93 | 0.00059 | 0.4551086 | 0.135 | 0.088 | 1 | Lig/Ten | nat9 |
| ap1g11 | 0.00061 | 0.2528522 | 0.139 | 0.091 | 1 | Lig/Ten | ap1g1 |
| selenot24 | 0.00062 | 0.3668093 | 0.155 | 0.104 | 1 | Lig/Ten | selenot2 |
| yipf33 | 0.00062 | 0.4430379 | 0.115 | 0.072 | 1 | Lig/Ten | yipf3 |
| sec11a3 | 0.00063 | 0.2203304 | 0.431 | 0.343 | 1 | Lig/Ten | sec11a |
| atp1a1a.16 | 0.00064 | 0.2389143 | 0.414 | 0.323 | 1 | Lig/Ten | atp1a1a.1 |
| arl5c5 | 0.00065 | 0.5672308 | 0.177 | 0.122 | 1 | Lig/Ten | arl5c |
| il15l6 | 0.00066 | 0.4661244 | 0.17 | 0.119 | 1 | Lig/Ten | il15l |
| sept9b3 | 0.00068 | 0.4806453 | 0.102 | 0.062 | 1 | Lig/Ten | sept9b |
| caprin1b3 | 0.00068 | 0.26303 | 0.378 | 0.297 | 1 | Lig/Ten | caprin1b |
| yipf43 | 0.0007 | 0.3371151 | 0.243 | 0.18 | 1 | Lig/Ten | yipf4 |
| six1a5 | 0.00074 | 0.2768842 | 0.188 | 0.131 | 1 | Lig/Ten | six1a |
| si:dkey-188i13.104 | 0.00074 | 0.3820021 | 0.321 | 0.248 | 1 | Lig/Ten | si:dkey-188i13.10 |
| mbnl25 | 0.00076 | 0.3058019 | 0.292 | 0.224 | 1 | Lig/Ten | mbnl2 |
| dnajc16 | 0.00078 | 0.2249324 | 0.363 | 0.283 | 1 | Lig/Ten | dnajc1 |
| cyc14 | 0.00078 | 0.2822909 | 0.235 | 0.17 | 1 | Lig/Ten | cyc1 |
| sertad44 | 0.00081 | 0.2701548 | 0.263 | 0.195 | 1 | Lig/Ten | sertad4 |
| sh3gl1b4 | 0.00084 | 0.2746199 | 0.321 | 0.244 | 1 | Lig/Ten | sh3gl1b |
| nfkbiaa6 | 0.00087 | 0.4466781 | 0.498 | 0.423 | 1 | Lig/Ten | nfkbiaa |
| nipa24 | 0.00087 | 0.4232148 | 0.119 | 0.076 | 1 | Lig/Ten | nipa2 |
| atp10a4 | 0.00089 | 0.5546863 | 0.144 | 0.097 | 1 | Lig/Ten | atp10a |
| cltca4 | 0.0009 | 0.2354875 | 0.268 | 0.197 | 1 | Lig/Ten | cltca |
| tmub12 | 0.00091 | 0.5234528 | 0.111 | 0.07 | 1 | Lig/Ten | tmub1 |
| si:dkey-17m8.14 | 0.00096 | 0.3450467 | 0.177 | 0.123 | 1 | Lig/Ten | si:dkey-17m8.1 |
| zgc:1949815 | 0.00102 | 0.3598587 | 0.126 | 0.082 | 1 | Lig/Ten | zgc:194981 |
| birc23 | 0.00103 | 0.2868046 | 0.192 | 0.136 | 1 | Lig/Ten | birc2 |
| gusb5 | 0.00104 | 0.4610219 | 0.184 | 0.13 | 1 | Lig/Ten | gusb |
| pim34 | 0.00105 | 0.5509421 | 0.155 | 0.107 | 1 | Lig/Ten | pim3 |
| tmem2631 | 0.00109 | 0.368665 | 0.212 | 0.155 | 1 | Lig/Ten | tmem263 |
| lysmd32 | 0.00109 | 0.3867634 | 0.155 | 0.105 | 1 | Lig/Ten | lysmd3 |
| wasla3 | 0.00113 | 0.2920628 | 0.241 | 0.178 | 1 | Lig/Ten | wasla |
| tex264a3 | 0.00114 | 0.3063965 | 0.175 | 0.122 | 1 | Lig/Ten | tex264a |
| pkd1a2 | 0.00114 | 0.6466277 | 0.113 | 0.073 | 1 | Lig/Ten | pkd1a |
| lhfpl2b | 0.00115 | 0.2764148 | 0.102 | 0.063 | 1 | Lig/Ten | lhfpl2b |
| papss2b3 | 0.00121 | 0.2780693 | 0.146 | 0.098 | 1 | Lig/Ten | papss2b |
| btbd6a5 | 0.00124 | 0.6446902 | 0.122 | 0.08 | 1 | Lig/Ten | btbd6a |
| arg28 | 0.00127 | 0.4400301 | 0.221 | 0.164 | 1 | Lig/Ten | arg2 |

| | | | | | | | |
|---|---|---|---|---|---|---|---|
| fam13b4 | 0.00129 | 0.5070952 | 0.17 | 0.12 | 1 | Lig/Ten | fam13b |
| chac15 | 0.0013 | 0.2735882 | 0.429 | 0.355 | 1 | Lig/Ten | chac1 |
| rorab6 | 0.00131 | 0.2954429 | 0.179 | 0.126 | 1 | Lig/Ten | rorab |
| CR848833.16 | 0.00139 | 0.3368775 | 0.113 | 0.073 | 1 | Lig/Ten | CR848833.1 |
| usp362 | 0.0014 | 0.2974744 | 0.265 | 0.201 | 1 | Lig/Ten | usp36 |
| MED94 | 0.0014 | 0.4104858 | 0.111 | 0.071 | 1 | Lig/Ten | MED9 |
| rab5c3 | 0.00141 | 0.3359804 | 0.296 | 0.231 | 1 | Lig/Ten | rab5c |
| zgc:560952 | 0.00144 | 0.299576 | 0.139 | 0.093 | 1 | Lig/Ten | zgc:56095 |
| col6a35 | 0.0015 | 0.2796032 | 0.308 | 0.238 | 1 | Lig/Ten | col6a3 |
| lamp24 | 0.00152 | 0.2130207 | 0.438 | 0.349 | 1 | Lig/Ten | lamp2 |
| zgc:1141742 | 0.00154 | 0.2760395 | 0.146 | 0.099 | 1 | Lig/Ten | zgc:114174 |
| fam3a4 | 0.00156 | 0.3429447 | 0.195 | 0.141 | 1 | Lig/Ten | fam3a |
| selenou1a2 | 0.00158 | 0.6000544 | 0.106 | 0.068 | 1 | Lig/Ten | selenou1a |
| si:ch211-198c19.34 | 0.00158 | 0.2604736 | 0.281 | 0.212 | 1 | Lig/Ten | si:ch211-198c19.3 |
| SGMS12 | 0.00167 | 0.2197071 | 0.1 | 0.062 | 1 | Lig/Ten | SGMS1 |
| copa4 | 0.0017 | 0.2466565 | 0.252 | 0.19 | 1 | Lig/Ten | copa |
| sypl2b2 | 0.00172 | 0.2818266 | 0.164 | 0.114 | 1 | Lig/Ten | sypl2b |
| manf4 | 0.00176 | 0.3388312 | 0.243 | 0.186 | 1 | Lig/Ten | manf |
| cmtm35 | 0.00186 | 0.39576 | 0.254 | 0.194 | 1 | Lig/Ten | cmtm3 |
| selenbp15 | 0.00189 | 0.3506398 | 0.119 | 0.078 | 1 | Lig/Ten | selenbp1 |
| myo1b2 | 0.00196 | 0.3333689 | 0.166 | 0.117 | 1 | Lig/Ten | myo1b |
| snx9b3 | 0.00204 | 0.2379739 | 0.243 | 0.181 | 1 | Lig/Ten | snx9b |
| g3bp15 | 0.00213 | 0.2336599 | 0.23 | 0.169 | 1 | Lig/Ten | g3bp1 |
| si:ch211-132g1.33 | 0.00215 | 0.358795 | 0.15 | 0.104 | 1 | Lig/Ten | si:ch211-132g1.3 |
| snai1b5 | 0.00216 | 0.4766587 | 0.159 | 0.113 | 1 | Lig/Ten | snai1b |
| rnf133 | 0.00216 | 0.2411058 | 0.181 | 0.129 | 1 | Lig/Ten | rnf13 |
| sc:d2174 | 0.00216 | 0.2581923 | 0.19 | 0.136 | 1 | Lig/Ten | sc:d217 |
| arf4a.14 | 0.00234 | 0.6182167 | 0.113 | 0.075 | 1 | Lig/Ten | arf4a.1 |
| gskip5 | 0.00249 | 0.204862 | 0.219 | 0.162 | 1 | Lig/Ten | gskip |
| praf23 | 0.00249 | 0.2320307 | 0.192 | 0.139 | 1 | Lig/Ten | praf2 |
| CU914776.15 | 0.00249 | 0.203964 | 0.126 | 0.084 | 1 | Lig/Ten | CU914776.1 |
| fkbp54 | 0.00265 | 0.2877967 | 0.257 | 0.197 | 1 | Lig/Ten | fkbp5 |
| zgc:1630801 | 0.00274 | 0.4085094 | 0.173 | 0.123 | 1 | Lig/Ten | zgc:163080 |
| tmem176l.3a3 | 0.00275 | 0.2333614 | 0.204 | 0.15 | 1 | Lig/Ten | tmem176l.3a |
| srprb4 | 0.00275 | 0.2383326 | 0.241 | 0.181 | 1 | Lig/Ten | srprb |
| blcap2 | 0.00286 | 0.3617012 | 0.195 | 0.144 | 1 | Lig/Ten | blcap |
| smoc26 | 0.00289 | 0.3143518 | 0.133 | 0.091 | 1 | Lig/Ten | smoc2 |
| baxa.14 | 0.00289 | 0.4243277 | 0.146 | 0.102 | 1 | Lig/Ten | baxa.1 |
| ets24 | 0.00293 | 0.2788026 | 0.215 | 0.16 | 1 | Lig/Ten | ets2 |
| bmpr2a4 | 0.00293 | 0.236882 | 0.237 | 0.178 | 1 | Lig/Ten | bmpr2a |
| si:ch211-66k16.271 | 0.00301 | 0.5463021 | 0.124 | 0.085 | 1 | Lig/Ten | si:ch211-66k16.27 |
| ndel1b4 | 0.00305 | 0.3187541 | 0.142 | 0.098 | 1 | Lig/Ten | ndel1b |
| arfgap25 | 0.00312 | 0.2877864 | 0.15 | 0.105 | 1 | Lig/Ten | arfgap2 |
| gorasp25 | 0.00313 | 0.4862845 | 0.155 | 0.111 | 1 | Lig/Ten | gorasp2 |
| tmem30ab4 | 0.00314 | 0.2669026 | 0.195 | 0.142 | 1 | Lig/Ten | tmem30ab |
| idh26 | 0.00318 | 0.2252194 | 0.423 | 0.344 | 1 | Lig/Ten | idh2 |
| higd2a4 | 0.00335 | 0.3311002 | 0.186 | 0.136 | 1 | Lig/Ten | higd2a |
| mbd3a2 | 0.00344 | 0.4127246 | 0.142 | 0.099 | 1 | Lig/Ten | mbd3a |

| | | | | | | | |
|---|---|---|---|---|---|---|---|
| gng13b1 | 0.00352 | 0.3671623 | 0.1 | 0.065 | 1 | Lig/Ten | gng13b |
| zgc:1103395 | 0.00357 | 0.3288804 | 0.277 | 0.216 | 1 | Lig/Ten | zgc:110339 |
| arl4ab6 | 0.00362 | 0.3284916 | 0.155 | 0.11 | 1 | Lig/Ten | arl4ab |
| ndufs33 | 0.00363 | 0.2324948 | 0.179 | 0.131 | 1 | Lig/Ten | ndufs3 |
| hipk3b1 | 0.00396 | 0.5064415 | 0.115 | 0.078 | 1 | Lig/Ten | hipk3b |
| ap2s13 | 0.00416 | 0.2577953 | 0.228 | 0.173 | 1 | Lig/Ten | ap2s1 |
| si:ch211-71n6.43 | 0.00435 | 0.304047 | 0.159 | 0.114 | 1 | Lig/Ten | si:ch211-71n6.4 |
| prrc13 | 0.00444 | 0.5042914 | 0.115 | 0.079 | 1 | Lig/Ten | prrc1 |
| uqcc33 | 0.00444 | 0.379097 | 0.122 | 0.083 | 1 | Lig/Ten | uqcc3 |
| socs3b6 | 0.00447 | 0.2464549 | 0.544 | 0.471 | 1 | Lig/Ten | socs3b |
| si:dkey-88p24.113 | 0.00451 | 0.3191704 | 0.119 | 0.082 | 1 | Lig/Ten | si:dkey-88p24.11 |
| arl14 | 0.00471 | 0.3568818 | 0.215 | 0.166 | 1 | Lig/Ten | arl1 |
| coro1ca6 | 0.00512 | 0.2690617 | 0.296 | 0.238 | 1 | Lig/Ten | coro1ca |
| rida3 | 0.00517 | 0.3562293 | 0.119 | 0.082 | 1 | Lig/Ten | rida |
| myo10l35 | 0.0052 | 0.3022328 | 0.142 | 0.1 | 1 | Lig/Ten | myo10l3 |
| psme22 | 0.00522 | 0.3991778 | 0.113 | 0.077 | 1 | Lig/Ten | psme2 |
| gas1a7 | 0.00526 | 0.2082617 | 0.212 | 0.16 | 1 | Lig/Ten | gas1a |
| gpr1081 | 0.00534 | 0.2524075 | 0.155 | 0.111 | 1 | Lig/Ten | gpr108 |
| fam114a25 | 0.00536 | 0.22534 | 0.159 | 0.115 | 1 | Lig/Ten | fam114a2 |
| si:dkey-23i12.53 | 0.00542 | 0.2368554 | 0.281 | 0.219 | 1 | Lig/Ten | si:dkey-23i12.5 |
| sav12 | 0.00568 | 0.2552357 | 0.17 | 0.125 | 1 | Lig/Ten | sav1 |
| tm9sf43 | 0.0057 | 0.2903029 | 0.111 | 0.075 | 1 | Lig/Ten | tm9sf4 |
| tspan365 | 0.00578 | 0.560172 | 0.208 | 0.162 | 1 | Lig/Ten | tspan36 |
| foxo3b5 | 0.0058 | 0.3910311 | 0.119 | 0.082 | 1 | Lig/Ten | foxo3b |
| SYDE11 | 0.00581 | 0.340083 | 0.1 | 0.066 | 1 | Lig/Ten | SYDE1 |
| pafah1b1b3 | 0.00595 | 0.2663212 | 0.265 | 0.208 | 1 | Lig/Ten | pafah1b1b |
| zmat53 | 0.00597 | 0.2626596 | 0.111 | 0.075 | 1 | Lig/Ten | zmat5 |
| fkbp25 | 0.00615 | 0.2656468 | 0.261 | 0.204 | 1 | Lig/Ten | fkbp2 |
| si:ch211-191j22.33 | 0.00618 | 0.3240012 | 0.164 | 0.12 | 1 | Lig/Ten | si:ch211-191j22.3 |
| dusp73 | 0.00627 | 0.5559744 | 0.108 | 0.074 | 1 | Lig/Ten | dusp7 |
| pdap1b5 | 0.00629 | 0.3911916 | 0.135 | 0.096 | 1 | Lig/Ten | pdap1b |
| twist1a5 | 0.00633 | 0.2577582 | 0.496 | 0.439 | 1 | Lig/Ten | twist1a |
| mia34 | 0.00634 | 0.222414 | 0.252 | 0.196 | 1 | Lig/Ten | mia3 |
| clasp23 | 0.0064 | 0.4774518 | 0.106 | 0.072 | 1 | Lig/Ten | clasp2 |
| chst141 | 0.00658 | 0.4799188 | 0.106 | 0.072 | 1 | Lig/Ten | chst14 |
| vdrb1 | 0.0066 | 0.2656938 | 0.197 | 0.148 | 1 | Lig/Ten | vdrb |
| dusp8a6 | 0.00667 | 0.4108808 | 0.128 | 0.09 | 1 | Lig/Ten | dusp8a |
| tbc1d14 | 0.00671 | 0.2195934 | 0.1 | 0.066 | 1 | Lig/Ten | tbc1d14 |
| spred2a2 | 0.00694 | 0.4280699 | 0.111 | 0.076 | 1 | Lig/Ten | spred2a |
| h2ax2 | 0.00704 | 0.3073355 | 0.104 | 0.071 | 1 | Lig/Ten | h2ax |
| zgc:1135318 | 0.00712 | 0.2582961 | 0.345 | 0.279 | 1 | Lig/Ten | zgc:113531 |
| ndufa83 | 0.00713 | 0.253415 | 0.29 | 0.233 | 1 | Lig/Ten | ndufa8 |
| il4r.16 | 0.00717 | 0.3080479 | 0.201 | 0.153 | 1 | Lig/Ten | il4r.1 |
| tspan9b3 | 0.00736 | 0.2598216 | 0.179 | 0.134 | 1 | Lig/Ten | tspan9b |
| atp6v1e1b5 | 0.00744 | 0.4307352 | 0.137 | 0.099 | 1 | Lig/Ten | atp6v1e1b |
| lamb1b4 | 0.00748 | 0.2437849 | 0.25 | 0.196 | 1 | Lig/Ten | lamb1b |
| map7d33 | 0.00751 | 0.5386503 | 0.115 | 0.08 | 1 | Lig/Ten | map7d3 |
| ugt2a75 | 0.00752 | 0.3171013 | 0.133 | 0.094 | 1 | Lig/Ten | ugt2a7 |

| | | | | | | | |
|---|---|---|---|---|---|---|---|
| ddah22 | 0.00761 | 0.3132909 | 0.173 | 0.128 | 1 | Lig/Ten | ddah2 |
| slc38a26 | 0.00769 | 0.3450928 | 0.115 | 0.08 | 1 | Lig/Ten | slc38a2 |
| tbcb4 | 0.00778 | 0.2819483 | 0.272 | 0.218 | 1 | Lig/Ten | tbcb |
| si:ch211-213a13.12 | 0.00781 | 0.3142961 | 0.173 | 0.129 | 1 | Lig/Ten | si:ch211-213a13.1 |
| tmco13 | 0.00783 | 0.3587193 | 0.117 | 0.082 | 1 | Lig/Ten | tmco1 |
| ddost4 | 0.00794 | 0.2741025 | 0.279 | 0.225 | 1 | Lig/Ten | ddost |
| mesd4 | 0.00801 | 0.2215438 | 0.197 | 0.151 | 1 | Lig/Ten | mesd |
| copg24 | 0.00817 | 0.3040063 | 0.195 | 0.15 | 1 | Lig/Ten | copg2 |
| afap14 | 0.00827 | 0.5867084 | 0.104 | 0.072 | 1 | Lig/Ten | afap1 |
| cdipt3 | 0.00832 | 0.25999 | 0.157 | 0.115 | 1 | Lig/Ten | cdipt |
| pfdn23 | 0.00843 | 0.2893843 | 0.104 | 0.071 | 1 | Lig/Ten | pfdn2 |
| lamc15 | 0.00846 | 0.3247224 | 0.173 | 0.129 | 1 | Lig/Ten | lamc1 |
| pdk2b4 | 0.00856 | 0.3741987 | 0.25 | 0.2 | 1 | Lig/Ten | pdk2b |
| ano5b5 | 0.0088 | 0.2241461 | 0.124 | 0.087 | 1 | Lig/Ten | ano5b |
| hspa56 | 0.009 | 0.2736827 | 0.558 | 0.492 | 1 | Lig/Ten | hspa5 |
| adam94 | 0.00912 | 0.3829803 | 0.1 | 0.067 | 1 | Lig/Ten | adam9 |
| stk353 | 0.00942 | 0.3775677 | 0.119 | 0.084 | 1 | Lig/Ten | stk35 |
| arcn1a3 | 0.00957 | 0.2531049 | 0.137 | 0.098 | 1 | Lig/Ten | arcn1a |
| anapc113 | 0.00965 | 0.2084377 | 0.173 | 0.129 | 1 | Lig/Ten | anapc11 |
| rasl122 | 0 | 6.9825458 | 0.919 | 0.052 | 0 | Perivascular | rasl12 |
| myh11a | 0 | 9.3373393 | 0.873 | 0.008 | 0 | Perivascular | myh11a |
| mcamb | 0 | 9.0192961 | 0.858 | 0.008 | 0 | Perivascular | mcamb |
| si:ch211-270g19.5 | 0 | 9.7735684 | 0.744 | 0.007 | 0 | Perivascular | si:ch211-270g19.5 |
| rbpms2a2 | 0 | 4.6280165 | 0.853 | 0.128 | 0 | Perivascular | rbpms2a |
| TCIM3 | 0 | 6.1752407 | 0.782 | 0.06 | 0 | Perivascular | TCIM |
| col4a26 | 0 | 4.1628915 | 0.906 | 0.213 | 0 | Perivascular | col4a2 |
| mylka4 | 0 | 4.5808494 | 0.777 | 0.102 | 0 | Perivascular | mylka |
| col4a15 | 0 | 4.0007911 | 0.893 | 0.222 | 0 | Perivascular | col4a1 |
| cxcl12b1 | 0 | 8.2869711 | 0.571 | 0.013 | 0 | Perivascular | cxcl12b |
| prx4 | 0 | 4.3992649 | 0.604 | 0.061 | 0 | Perivascular | prx |
| tpm4b5 | 0 | 4.6392116 | 0.609 | 0.08 | 0 | Perivascular | tpm4b |
| pln2 | 0 | 6.2319248 | 0.566 | 0.039 | 0 | Perivascular | pln |
| sh3d211 | 0 | 5.8900336 | 0.52 | 0.023 | 0 | Perivascular | sh3d21 |
| meox15 | 0 | 4.2548455 | 0.543 | 0.059 | 0 | Perivascular | meox1 |
| pcdh1b | 0 | 7.1560487 | 0.462 | 0.007 | 0 | Perivascular | pcdh1b |
| acta21 | 0 | 4.8875195 | 0.485 | 0.034 | 0 | Perivascular | acta2 |
| BX088707.3 | 0 | 5.148127 | 0.47 | 0.031 | 0 | Perivascular | BX088707.3 |
| itga12 | 0 | 5.1093661 | 0.447 | 0.028 | 0 | Perivascular | itga1 |
| atp1a1b | 0 | 5.6677305 | 0.419 | 0.019 | 0 | Perivascular | atp1a1b |
| rbpms2b | 0 | 10.241716 | 0.398 | 0.001 | 0 | Perivascular | rbpms2b |
| si:ch73-193i22.1 | 0 | 7.4461984 | 0.401 | 0.005 | 0 | Perivascular | si:ch73-193i22.1 |
| CR753876.12 | 0 | 4.2610302 | 0.426 | 0.034 | 0 | Perivascular | CR753876.1 |
| slc20a1a | 0 | 6.6578615 | 0.358 | 0.007 | 0 | Perivascular | slc20a1a |
| col18a1b | 0 | 7.5969589 | 0.35 | 0.004 | 0 | Perivascular | col18a1b |
| jam3a | 0 | 6.5132277 | 0.35 | 0.007 | 0 | Perivascular | jam3a |
| BX323087.11 | 0 | 5.1119465 | 0.358 | 0.019 | 0 | Perivascular | BX323087.1 |
| cntn3b | 0 | 5.6558137 | 0.302 | 0.013 | 0 | Perivascular | cntn3b |
| RAMP1 | 0 | 6.5381293 | 0.292 | 0.009 | 0 | Perivascular | RAMP1 |

| | | | | | | | |
|---|---|---|---|---|---|---|---|
| cables2b | 0 | 5.7601066 | 0.294 | 0.012 | 0 | Perivascular | cables2b |
| eef2l2 | 0 | 5.8578925 | 0.272 | 0.011 | 0 | Perivascular | eef2l2 |
| rassf4a | 0 | 9.4942268 | 0.259 | 0.001 | 0 | Perivascular | rassf4a |
| pdgfbb | 0 | 5.7013768 | 0.261 | 0.011 | 0 | Perivascular | pdgfbb |
| sorbs2b | 0 | 7.6053642 | 0.249 | 0.003 | 0 | Perivascular | sorbs2b |
| ptpreb | 0 | 6.518414 | 0.249 | 0.006 | 0 | Perivascular | ptpreb |
| ndufa4l2a | 0 | 9.350227 | 0.241 | 0.002 | 0 | Perivascular | ndufa4l2a |
| pde5ab1 | 0 | 6.2577611 | 0.216 | 0.006 | 0 | Perivascular | pde5ab |
| notch1b | 0 | 7.7440709 | 0.206 | 0.002 | 0 | Perivascular | notch1b |
| ppp1r14aa | 0 | 6.6336346 | 0.201 | 0.003 | 0 | Perivascular | ppp1r14aa |
| ano1 | 0 | 6.8842114 | 0.198 | 0.003 | 0 | Perivascular | ano1 |
| dachb1 | 0 | 5.7066417 | 0.195 | 0.006 | 0 | Perivascular | dachb |
| arr3b | 0 | 5.9387525 | 0.195 | 0.006 | 0 | Perivascular | arr3b |
| si:ch211-158d24.4 | 0 | 7.0236661 | 0.19 | 0.005 | 0 | Perivascular | si:ch211-158d24.4 |
| abcc9 | 0 | 8.576463 | 0.185 | 0.001 | 0 | Perivascular | abcc9 |
| spdya | 0 | 8.899586 | 0.168 | 0.001 | 0 | Perivascular | spdya |
| zgc:194629 | 0 | 8.1535502 | 0.157 | 0.001 | 0 | Perivascular | zgc:194629 |
| npr1b | 0 | 8.2009135 | 0.15 | 0.001 | 0 | Perivascular | npr1b |
| si:dkey-57k2.6 | 0 | 7.2084328 | 0.142 | 0.002 | 0 | Perivascular | si:dkey-57k2.6 |
| ampd3a | 0 | 7.256921 | 0.14 | 0.002 | 0 | Perivascular | ampd3a |
| atp2b2 | 0 | 6.1819388 | 0.129 | 0.003 | 0 | Perivascular | atp2b2 |
| aplnr2 | 0 | 7.6906781 | 0.122 | 0.001 | 0 | Perivascular | aplnr2 |
| CRACD | 0 | 7.0083308 | 0.119 | 0.002 | 0 | Perivascular | CRACD |
| atp2b1b | 0 | 7.3172338 | 0.119 | 0.002 | 0 | Perivascular | atp2b1b |
| rnd2 | 0 | 7.6211059 | 0.107 | 0.001 | 0 | Perivascular | rnd2 |
| cx39.4 | 0 | 6.8107072 | 0.107 | 0.001 | 0 | Perivascular | cx39.4 |
| arhgap29a | 0 | 6.4876412 | 0.102 | 0.002 | 0 | Perivascular | arhgap29a |
| col5a3a4 | 4E-306 | 3.5529014 | 0.528 | 0.064 | 1.2E-301 | Perivascular | col5a3a |
| plp1b | 5E-303 | 5.4417305 | 0.168 | 0.006 | 1.7E-298 | Perivascular | plp1b |
| BX901920.1 | 2E-301 | 5.7806121 | 0.249 | 0.014 | 7.6E-297 | Perivascular | BX901920.1 |
| hey2 | 2E-296 | 5.7744676 | 0.168 | 0.006 | 7E-292 | Perivascular | hey2 |
| adora2aa | 2E-294 | 5.9362808 | 0.165 | 0.006 | 4.9E-290 | Perivascular | adora2aa |
| tcap | 3E-294 | 6.6069848 | 0.221 | 0.011 | 8.2E-290 | Perivascular | tcap |
| ncaldb | 8E-287 | 5.8440673 | 0.15 | 0.005 | 2.6E-282 | Perivascular | ncaldb |
| sept154 | 2E-280 | 3.8667237 | 0.802 | 0.199 | 7E-276 | Perivascular | sept15 |
| oaz2a3 | 1E-271 | 4.3769434 | 0.528 | 0.074 | 3.7E-267 | Perivascular | oaz2a |
| kcne44 | 5E-261 | 4.4994678 | 0.464 | 0.058 | 1.6E-256 | Perivascular | kcne4 |
| notch33 | 1E-259 | 3.7114623 | 0.921 | 0.342 | 4E-255 | Perivascular | notch3 |
| rbm24a | 6E-259 | 5.0031772 | 0.284 | 0.022 | 2.1E-254 | Perivascular | rbm24a |
| lmod1b3 | 2E-258 | 3.7040712 | 0.581 | 0.093 | 7.6E-254 | Perivascular | lmod1b |
| eva1bb2 | 2E-255 | 4.3171257 | 0.371 | 0.037 | 4.9E-251 | Perivascular | eva1bb |
| epas1b1 | 2E-253 | 4.2752727 | 0.36 | 0.035 | 6.3E-249 | Perivascular | epas1b |
| prkg1b | 2E-253 | 5.3181869 | 0.173 | 0.008 | 7.2E-249 | Perivascular | prkg1b |
| smtnb | 4E-238 | 5.5249804 | 0.203 | 0.012 | 1.2E-233 | Perivascular | smtnb |
| fhl3b5 | 8E-232 | 3.8175597 | 0.586 | 0.108 | 2.7E-227 | Perivascular | fhl3b |
| epas1a3 | 3E-230 | 4.1357559 | 0.449 | 0.061 | 8.7E-226 | Perivascular | epas1a |
| cnnm2b1 | 2E-228 | 4.7147418 | 0.266 | 0.021 | 5.1E-224 | Perivascular | cnnm2b |
| tpm12 | 3E-225 | 3.3327981 | 0.896 | 0.332 | 1.1E-220 | Perivascular | tpm1 |

| | | | | | | | |
|---|---|---|---|---|---|---|---|
| stard13a | 2E-223 | 4.9498791 | 0.17 | 0.009 | 5E-219 | Perivascular | stard13a |
| OLFML2A | 7E-215 | 5.5011688 | 0.104 | 0.003 | 2.3E-210 | Perivascular | OLFML2A |
| adss1 | 1E-210 | 5.327936 | 0.137 | 0.006 | 3.1E-206 | Perivascular | adss1 |
| notch1a | 1E-210 | 4.833215 | 0.165 | 0.009 | 4.5E-206 | Perivascular | notch1a |
| fabp11a3 | 9E-210 | 3.9361804 | 0.467 | 0.07 | 2.8E-205 | Perivascular | fabp11a |
| akap12a | 6E-209 | 5.4232804 | 0.114 | 0.004 | 1.9E-204 | Perivascular | akap12a |
| si:ch211-191i18.22 | 1E-206 | 4.9588309 | 0.325 | 0.035 | 3.4E-202 | Perivascular | si:ch211-191i18.2 |
| lamb1b5 | 7E-203 | 3.0418248 | 0.726 | 0.192 | 2.4E-198 | Perivascular | lamb1b |
| hpca | 2E-199 | 4.1672439 | 0.307 | 0.032 | 5.7E-195 | Perivascular | hpca |
| daam2 | 4E-196 | 3.9770988 | 0.322 | 0.036 | 1.5E-191 | Perivascular | daam2 |
| cd248a7 | 2E-193 | 3.3460646 | 0.561 | 0.113 | 6.9E-189 | Perivascular | cd248a |
| tmsb4x4 | 2E-183 | 1.6914388 | 1 | 0.992 | 6.7E-179 | Perivascular | tmsb4x |
| lurap12 | 7E-180 | 3.7237393 | 0.36 | 0.048 | 2.1E-175 | Perivascular | lurap1 |
| si:ch1073-268j14.11 | 1E-179 | 3.6927456 | 0.332 | 0.041 | 3.8E-175 | Perivascular | si:ch1073-268j14.1 |
| fxyd6l4 | 2E-179 | 3.315696 | 0.751 | 0.25 | 5.2E-175 | Perivascular | fxyd6l |
| adra1aa | 2E-179 | 5.5737919 | 0.102 | 0.004 | 7.2E-175 | Perivascular | adra1aa |
| cald1a1 | 2E-174 | 2.5379967 | 0.949 | 0.62 | 5.4E-170 | Perivascular | cald1a |
| sh3bgrl21 | 8E-172 | 4.4290743 | 0.198 | 0.015 | 2.6E-167 | Perivascular | sh3bgrl2 |
| pkib | 1E-165 | 4.7311921 | 0.193 | 0.015 | 3.3E-161 | Perivascular | pkib |
| crip17 | 3E-165 | 2.7611949 | 0.929 | 0.459 | 1E-160 | Perivascular | crip1 |
| mical2b | 3E-162 | 4.5193832 | 0.183 | 0.014 | 1E-157 | Perivascular | mical2b |
| mustn1a5 | 4E-161 | 3.269192 | 0.442 | 0.079 | 1.3E-156 | Perivascular | mustn1a |
| myo1b3 | 3E-160 | 3.4063513 | 0.52 | 0.114 | 1.1E-155 | Perivascular | myo1b |
| cd281 | 7E-159 | 4.3707194 | 0.213 | 0.019 | 2.4E-154 | Perivascular | cd28 |
| pde4ba6 | 1E-154 | 3.3623122 | 0.513 | 0.114 | 3.1E-150 | Perivascular | pde4ba |
| rgs5a3 | 4E-153 | 4.5008154 | 0.589 | 0.149 | 1.3E-148 | Perivascular | rgs5a |
| mapkapk3 | 4E-151 | 4.0723157 | 0.256 | 0.029 | 1.4E-146 | Perivascular | mapkapk3 |
| rasgef1ba4 | 2E-150 | 3.2841399 | 0.589 | 0.157 | 7.9E-146 | Perivascular | rasgef1ba |
| rcan22 | 7E-149 | 3.4205584 | 0.358 | 0.056 | 2.2E-144 | Perivascular | rcan2 |
| BX322618.1 | 1E-146 | 5.6004144 | 0.109 | 0.005 | 4.4E-142 | Perivascular | BX322618.1 |
| atp1b41 | 5E-143 | 3.9209063 | 0.244 | 0.028 | 1.7E-138 | Perivascular | atp1b4 |
| add3b | 8E-143 | 4.1898158 | 0.198 | 0.018 | 2.7E-138 | Perivascular | add3b |
| calm2a3 | 9E-143 | 3.0609793 | 0.754 | 0.319 | 3E-138 | Perivascular | calm2a |
| pdgfrb6 | 6E-139 | 2.7092005 | 0.825 | 0.409 | 1.9E-134 | Perivascular | pdgfrb |
| kdrl | 7E-138 | 5.0554659 | 0.127 | 0.008 | 2.1E-133 | Perivascular | kdrl |
| tspan18a | 3E-136 | 4.3340025 | 0.183 | 0.017 | 1.1E-131 | Perivascular | tspan18a |
| itih54 | 5E-135 | 2.8780751 | 0.381 | 0.067 | 1.8E-130 | Perivascular | itih5 |
| si:ch211-195b13.17 | 1E-133 | 3.1166193 | 0.685 | 0.24 | 4.3E-129 | Perivascular | si:ch211-195b13.1 |
| atp2a33 | 3E-132 | 3.7912698 | 0.282 | 0.04 | 8.3E-128 | Perivascular | atp2a3 |
| tpm4a3 | 1E-131 | 2.1847674 | 0.891 | 0.551 | 3.5E-127 | Perivascular | tpm4a |
| stk17al3 | 2E-130 | 3.492015 | 0.386 | 0.074 | 6.5E-126 | Perivascular | stk17al |
| FP103004.1 | 2E-130 | 4.0695795 | 0.162 | 0.014 | 7.3E-126 | Perivascular | FP103004.1 |
| sept8a1 | 1E-129 | 3.1309993 | 0.505 | 0.129 | 3.4E-125 | Perivascular | sept8a |
| zak | 6E-126 | 4.0664949 | 0.231 | 0.028 | 2E-121 | Perivascular | zak |
| pcdh18a6 | 2E-124 | 3.1264293 | 0.401 | 0.08 | 5.1E-120 | Perivascular | pcdh18a |
| afap1l22 | 5E-118 | 3.7162422 | 0.254 | 0.036 | 1.6E-113 | Perivascular | afap1l2 |
| si:ch73-119p20.11 | 9E-118 | 3.9144478 | 0.236 | 0.031 | 2.9E-113 | Perivascular | si:ch73-119p20.1 |
| tagln6 | 2E-116 | 2.5754025 | 0.871 | 0.536 | 5.3E-112 | Perivascular | tagln |

| | | | | | | | |
|---|---|---|---|---|---|---|---|
| cobll1b4 | 5E-114 | 3.1594759 | 0.335 | 0.061 | 1.6E-109 | Perivascular | cobll1b |
| s1pr5a2 | 1E-109 | 3.293719 | 0.322 | 0.059 | 3.3E-105 | Perivascular | s1pr5a |
| im:71523488 | 9E-106 | 3.0502006 | 0.624 | 0.233 | 2.9E-101 | Perivascular | im:7152348 |
| BX908750.1 | 2E-104 | 4.5094484 | 0.109 | 0.008 | 5.7E-100 | Perivascular | BX908750.1 |
| arap3 | 1E-102 | 4.1617854 | 0.109 | 0.008 | 3.51E-98 | Perivascular | arap3 |
| tiam1a3 | 1E-102 | 3.1416888 | 0.315 | 0.06 | 3.91E-98 | Perivascular | tiam1a |
| zeb2a7 | 2E-102 | 2.3787508 | 0.718 | 0.339 | 7.77E-98 | Perivascular | zeb2a |
| akap12b4 | 4E-102 | 2.2085561 | 0.505 | 0.14 | 1.17E-97 | Perivascular | akap12b |
| nck2b | 4E-100 | 3.643296 | 0.261 | 0.043 | 1.26E-95 | Perivascular | nck2b |
| adgrl3.1 | 1E-99 | 3.4415443 | 0.239 | 0.036 | 3.45E-95 | Perivascular | adgrl3.1 |
| anks1b2 | 7E-99 | 3.2512164 | 0.305 | 0.058 | 2.25E-94 | Perivascular | anks1b |
| si:ch211-198c19.35 | 1.3E-98 | 1.9987094 | 0.619 | 0.209 | 4.25E-94 | Perivascular | si:ch211-198c19.3 |
| col18a1a5 | 9.6E-98 | 2.8522049 | 0.497 | 0.151 | 3.11E-93 | Perivascular | col18a1a |
| myl9b3 | 1.3E-96 | 2.1200063 | 0.766 | 0.423 | 4.07E-92 | Perivascular | myl9b |
| si:dkey-27i16.27 | 3.7E-95 | 2.574865 | 0.437 | 0.115 | 1.19E-90 | Perivascular | si:dkey-27i16.2 |
| CABZ01044746.11 | 6.3E-94 | 3.4563247 | 0.211 | 0.03 | 2.04E-89 | Perivascular | CABZ01044746.1 |
| ptmaa4 | 5.3E-93 | 1.4275445 | 0.954 | 0.835 | 1.73E-88 | Perivascular | ptmaa |
| mrvi1 | 1.6E-92 | 3.5759882 | 0.185 | 0.024 | 5.06E-88 | Perivascular | mrvi1 |
| nrip2 | 2.9E-90 | 3.9045593 | 0.165 | 0.02 | 9.41E-86 | Perivascular | nrip2 |
| mprip1 | 3.4E-90 | 2.9765109 | 0.335 | 0.074 | 1.1E-85 | Perivascular | mprip |
| CU914776.16 | 8E-90 | 2.3599263 | 0.358 | 0.082 | 2.61E-85 | Perivascular | CU914776.1 |
| rgs45 | 1.5E-88 | 3.3466416 | 0.317 | 0.068 | 4.83E-84 | Perivascular | rgs4 |
| marcksl1b8 | 6.6E-87 | 2.1004538 | 0.627 | 0.252 | 2.14E-82 | Perivascular | marcksl1b |
| arhgap363 | 2E-86 | 3.1251554 | 0.297 | 0.061 | 6.35E-82 | Perivascular | arhgap36 |
| CR735102.1 | 5.6E-86 | 4.0104932 | 0.112 | 0.01 | 1.82E-81 | Perivascular | CR735102.1 |
| BX284638.23 | 5.3E-84 | 3.0356751 | 0.228 | 0.038 | 1.73E-79 | Perivascular | BX284638.2 |
| prex25 | 3.7E-83 | 2.675091 | 0.353 | 0.086 | 1.2E-78 | Perivascular | prex2 |
| grk46 | 3.8E-77 | 2.481373 | 0.5 | 0.178 | 1.24E-72 | Perivascular | grk4 |
| neurl1b | 2E-76 | 2.9906041 | 0.239 | 0.045 | 6.65E-72 | Perivascular | neurl1b |
| calcrla | 8.7E-75 | 3.977116 | 0.129 | 0.015 | 2.84E-70 | Perivascular | calcrla |
| cyp2ad31 | 1.6E-74 | 3.2973138 | 0.155 | 0.021 | 5.19E-70 | Perivascular | cyp2ad3 |
| msna4 | 5.5E-74 | 1.4996887 | 0.904 | 0.75 | 1.8E-69 | Perivascular | msna |
| col15a1b3 | 1.9E-73 | 3.0236492 | 0.246 | 0.049 | 6.15E-69 | Perivascular | col15a1b |
| ppp1r14ab | 2.1E-72 | 4.1511189 | 0.107 | 0.01 | 6.72E-68 | Perivascular | ppp1r14ab |
| arhgap42a5 | 6E-72 | 2.6695209 | 0.299 | 0.071 | 1.94E-67 | Perivascular | arhgap42a |
| nrarpa3 | 7.3E-72 | 2.7905009 | 0.332 | 0.087 | 2.39E-67 | Perivascular | nrarpa |
| zgc:1538674 | 1.5E-71 | 1.4046505 | 0.881 | 0.752 | 4.85E-67 | Perivascular | zgc:153867 |
| itga4 | 3.7E-71 | 3.5738609 | 0.147 | 0.02 | 1.2E-66 | Perivascular | itga4 |
| tanc1b | 5.7E-71 | 3.4008773 | 0.195 | 0.033 | 1.85E-66 | Perivascular | tanc1b |
| BX248082.2 | 1.1E-70 | 3.5851815 | 0.137 | 0.017 | 3.58E-66 | Perivascular | BX248082.2 |
| ccdc3b5 | 1.8E-70 | 2.5088783 | 0.398 | 0.123 | 5.7E-66 | Perivascular | ccdc3b |
| flrt32 | 9.3E-68 | 3.1412514 | 0.254 | 0.056 | 3.02E-63 | Perivascular | flrt3 |
| ARHGEF17 | 4.6E-67 | 2.8975482 | 0.244 | 0.051 | 1.48E-62 | Perivascular | ARHGEF17 |
| CABZ01061591.11 | 1.5E-66 | 2.7417075 | 0.289 | 0.071 | 4.73E-62 | Perivascular | CABZ01061591.1 |
| prss356 | 2.5E-66 | 2.1892698 | 0.338 | 0.091 | 7.99E-62 | Perivascular | prss35 |
| pgfb | 2E-65 | 3.8104265 | 0.142 | 0.02 | 6.4E-61 | Perivascular | pgfb |
| ednraa7 | 3E-65 | 2.1447115 | 0.52 | 0.212 | 9.6E-61 | Perivascular | ednraa |
| pabpc41 | 1.9E-64 | 3.5854573 | 0.155 | 0.024 | 6.04E-60 | Perivascular | pabpc4 |

| | | | | | | | |
|---|---|---|---|---|---|---|---|
| cnnm2a | 3.6E-64 | 3.124906 | 0.208 | 0.04 | 1.17E-59 | Perivascular | cnnm2a |
| aatkb1 | 8.4E-64 | 3.3697583 | 0.152 | 0.023 | 2.72E-59 | Perivascular | aatkb |
| lims12 | 2.1E-62 | 2.0901803 | 0.579 | 0.293 | 6.73E-58 | Perivascular | lims1 |
| cfl13 | 8.1E-62 | 1.0617608 | 0.939 | 0.843 | 2.62E-57 | Perivascular | cfl1 |
| mef2cb2 | 2.5E-61 | 2.525595 | 0.277 | 0.069 | 8E-57 | Perivascular | mef2cb |
| ak13 | 2.6E-61 | 2.7973549 | 0.376 | 0.125 | 8.56E-57 | Perivascular | ak1 |
| pag1 | 1.1E-60 | 3.2677902 | 0.168 | 0.028 | 3.5E-56 | Perivascular | pag1 |
| lama45 | 1.1E-60 | 1.9929302 | 0.569 | 0.269 | 3.7E-56 | Perivascular | lama4 |
| sept21 | 7.6E-60 | 2.2326692 | 0.533 | 0.257 | 2.48E-55 | Perivascular | sept2 |
| actb25 | 1.6E-59 | 0.9451223 | 0.997 | 0.991 | 5.24E-55 | Perivascular | actb2 |
| parvab | 6.9E-59 | 2.8494772 | 0.195 | 0.038 | 2.26E-54 | Perivascular | parvab |
| f3a | 1.6E-57 | 3.1287559 | 0.137 | 0.02 | 5.36E-53 | Perivascular | f3a |
| pld1a4 | 7.6E-57 | 2.6500467 | 0.312 | 0.092 | 2.48E-52 | Perivascular | pld1a |
| adra1d | 1.7E-56 | 4.219077 | 0.102 | 0.012 | 5.54E-52 | Perivascular | adra1d |
| fzd43 | 5.1E-56 | 2.7275686 | 0.226 | 0.051 | 1.66E-51 | Perivascular | fzd4 |
| atp2a2b3 | 1.8E-55 | 2.6182065 | 0.325 | 0.101 | 5.86E-51 | Perivascular | atp2a2b |
| akt3a | 2.8E-55 | 3.0134363 | 0.155 | 0.026 | 8.96E-51 | Perivascular | akt3a |
| ppiab5 | 2.8E-55 | 0.9371189 | 0.944 | 0.896 | 9.16E-51 | Perivascular | ppiab |
| eva1a4 | 7.5E-55 | 2.9971093 | 0.234 | 0.056 | 2.44E-50 | Perivascular | eva1a |
| tagln25 | 9E-55 | 1.4089997 | 0.85 | 0.665 | 2.92E-50 | Perivascular | tagln2 |
| ralba4 | 9.3E-55 | 2.5676491 | 0.302 | 0.089 | 3.01E-50 | Perivascular | ralba |
| si:ch1073-303k11.22 | 8.3E-54 | 3.1013307 | 0.165 | 0.03 | 2.71E-49 | Perivascular | si:ch1073-303k11.2 |
| zgc:921407 | 5.1E-53 | 2.4683928 | 0.343 | 0.112 | 1.66E-48 | Perivascular | zgc:92140 |
| rgs5b8 | 1.5E-52 | 1.5821843 | 0.807 | 0.521 | 4.86E-48 | Perivascular | rgs5b |
| crip25 | 1.5E-52 | 1.4573889 | 0.282 | 0.076 | 5E-48 | Perivascular | crip2 |
| cttnbp2nlb | 1.9E-52 | 3.1047941 | 0.17 | 0.033 | 6.14E-48 | Perivascular | cttnbp2nlb |
| AL954322.21 | 3.9E-52 | 3.2874654 | 0.198 | 0.044 | 1.28E-47 | Perivascular | AL954322.2 |
| rftn22 | 1.1E-51 | 3.3614257 | 0.147 | 0.026 | 3.53E-47 | Perivascular | rftn2 |
| tns1b5 | 1.2E-51 | 1.950336 | 0.624 | 0.366 | 3.75E-47 | Perivascular | tns1b |
| si:dkey-172h23.25 | 3.3E-51 | 2.065833 | 0.393 | 0.145 | 1.07E-46 | Perivascular | si:dkey-172h23.2 |
| cdh26 | 2.8E-50 | 1.8046545 | 0.561 | 0.291 | 9.11E-46 | Perivascular | cdh2 |
| rtn1a3 | 6.6E-50 | 2.0929262 | 0.487 | 0.221 | 2.15E-45 | Perivascular | rtn1a |
| dysf1 | 9.5E-50 | 2.7739603 | 0.198 | 0.044 | 3.09E-45 | Perivascular | dysf |
| cadm3 | 7.3E-49 | 3.4993524 | 0.124 | 0.02 | 2.39E-44 | Perivascular | cadm3 |
| adam8a2 | 9.6E-49 | 2.1998979 | 0.368 | 0.134 | 3.14E-44 | Perivascular | adam8a |
| CR848833.17 | 3.2E-48 | 2.6045604 | 0.256 | 0.071 | 1.05E-43 | Perivascular | CR848833.1 |
| tbx2b4 | 3.4E-48 | 2.4653666 | 0.35 | 0.125 | 1.09E-43 | Perivascular | tbx2b |
| vcam1b | 1.3E-47 | 3.0101628 | 0.198 | 0.046 | 4.07E-43 | Perivascular | vcam1b |
| ENSDARG000001052( | 2.3E-47 | 2.6196502 | 0.231 | 0.06 | 7.62E-43 | Perivascular | ENSDARG00000105 |
| sptbn14 | 4.4E-47 | 1.7354621 | 0.513 | 0.248 | 1.44E-42 | Perivascular | sptbn1 |
| gapdhs4 | 7.6E-46 | 1.4077228 | 0.794 | 0.629 | 2.47E-41 | Perivascular | gapdhs |
| ENSDARG000000595( | 1.3E-45 | 2.9414519 | 0.15 | 0.029 | 4.11E-41 | Perivascular | ENSDARG00000059 |
| myl63 | 1.8E-45 | 2.0662546 | 0.459 | 0.214 | 5.97E-41 | Perivascular | myl6 |
| itpkb | 3E-45 | 2.7150739 | 0.193 | 0.046 | 9.84E-41 | Perivascular | itpkb |
| slc25a57 | 3.5E-45 | 0.7204038 | 0.985 | 0.963 | 1.13E-40 | Perivascular | slc25a5 |
| arhgef10la | 2E-44 | 3.0268265 | 0.135 | 0.024 | 6.64E-40 | Perivascular | arhgef10la |
| msx3 | 1.2E-43 | 3.3738478 | 0.135 | 0.025 | 3.94E-39 | Perivascular | msx3 |
| camk1b4 | 2.1E-43 | 2.4113618 | 0.218 | 0.057 | 6.82E-39 | Perivascular | camk1b |

| | | | | | | | |
|---|---|---|---|---|---|---|---|
| actb14 | 2E-42 | 0.9748211 | 0.944 | 0.866 | 6.54E-38 | Perivascular | actb1 |
| cnn3a5 | 1.5E-41 | 1.5245624 | 0.668 | 0.476 | 4.78E-37 | Perivascular | cnn3a |
| aldocb5 | 1.8E-41 | 2.4605208 | 0.249 | 0.075 | 5.87E-37 | Perivascular | aldocb |
| sept102 | 2.1E-41 | 1.7703526 | 0.528 | 0.301 | 6.95E-37 | Perivascular | sept10 |
| mef2ca1 | 3.7E-41 | 2.8287818 | 0.16 | 0.035 | 1.2E-36 | Perivascular | mef2ca |
| sorbs35 | 4.4E-41 | 2.0044287 | 0.419 | 0.189 | 1.42E-36 | Perivascular | sorbs3 |
| pmp22b4 | 5.3E-41 | 1.1578981 | 0.635 | 0.36 | 1.73E-36 | Perivascular | pmp22b |
| nr2f25 | 2.2E-40 | 2.1134944 | 0.317 | 0.114 | 7.12E-36 | Perivascular | nr2f2 |
| C7H4orf485 | 1.7E-39 | 1.6253935 | 0.482 | 0.242 | 5.55E-35 | Perivascular | C7H4orf48 |
| agtr24 | 3.8E-39 | 2.3705867 | 0.312 | 0.115 | 1.24E-34 | Perivascular | agtr2 |
| si:ch211-195m9.33 | 7E-39 | 2.8157376 | 0.178 | 0.044 | 2.28E-34 | Perivascular | si:ch211-195m9.3 |
| lamc16 | 7.2E-39 | 1.9237699 | 0.332 | 0.128 | 2.35E-34 | Perivascular | lamc1 |
| rem15 | 2E-38 | 2.4936354 | 0.218 | 0.063 | 6.52E-34 | Perivascular | rem1 |
| sik18 | 3E-38 | 1.888611 | 0.495 | 0.252 | 9.64E-34 | Perivascular | sik1 |
| rpz56 | 5.9E-38 | 1.3046666 | 0.566 | 0.306 | 1.91E-33 | Perivascular | rpz5 |
| kif1aa1 | 7.1E-38 | 2.6477938 | 0.122 | 0.023 | 2.32E-33 | Perivascular | kif1aa |
| ebf3a7 | 8.3E-38 | 1.9586618 | 0.442 | 0.215 | 2.71E-33 | Perivascular | ebf3a |
| myl9a4 | 9.7E-38 | 1.9633262 | 0.594 | 0.38 | 3.15E-33 | Perivascular | myl9a |
| si:ch211-127i16.2 | 1.8E-37 | 2.8837343 | 0.104 | 0.018 | 5.94E-33 | Perivascular | si:ch211-127i16.2 |
| dusp28 | 2.5E-37 | 1.861982 | 0.454 | 0.221 | 8.02E-33 | Perivascular | dusp2 |
| tln14 | 2.8E-37 | 1.1519419 | 0.749 | 0.61 | 9.16E-33 | Perivascular | tln1 |
| cfap451 | 3.3E-37 | 2.9901952 | 0.119 | 0.022 | 1.08E-32 | Perivascular | cfap45 |
| phc2b5 | 5.3E-37 | 2.3620328 | 0.239 | 0.075 | 1.72E-32 | Perivascular | phc2b |
| her67 | 3.2E-36 | 1.2499731 | 0.675 | 0.431 | 1.04E-31 | Perivascular | her6 |
| klf6a5 | 5.2E-36 | 1.0150525 | 0.873 | 0.746 | 1.68E-31 | Perivascular | klf6a |
| CABZ01058721.1 | 8.4E-36 | 2.5040701 | 0.168 | 0.042 | 2.73E-31 | Perivascular | CABZ01058721.1 |
| btbd6a6 | 3E-35 | 2.5773336 | 0.239 | 0.079 | 9.84E-31 | Perivascular | btbd6a |
| CABZ01068367.17 | 2.2E-34 | 1.8035432 | 0.596 | 0.392 | 7.01E-30 | Perivascular | CABZ01068367.1 |
| tes3 | 4.6E-34 | 2.2543952 | 0.231 | 0.074 | 1.51E-29 | Perivascular | tes |
| zgc:1748632 | 4.7E-34 | 1.5424217 | 0.442 | 0.229 | 1.52E-29 | Perivascular | zgc:174863 |
| abr | 1.2E-33 | 2.7232141 | 0.124 | 0.026 | 3.97E-29 | Perivascular | abr |
| lifrb4 | 1.3E-33 | 1.7444843 | 0.414 | 0.198 | 4.38E-29 | Perivascular | lifrb |
| niban2a4 | 1.5E-33 | 2.4921892 | 0.183 | 0.051 | 4.96E-29 | Perivascular | niban2a |
| cd28l | 2.2E-33 | 2.8239158 | 0.102 | 0.018 | 7.31E-29 | Perivascular | cd28l |
| map21 | 1.6E-32 | 2.4947926 | 0.14 | 0.032 | 5.06E-28 | Perivascular | map2 |
| neu3.54 | 2.4E-32 | 0.9310458 | 0.226 | 0.068 | 7.84E-28 | Perivascular | neu3.5 |
| itgb1b | 4.1E-32 | 2.2978169 | 0.173 | 0.047 | 1.32E-27 | Perivascular | itgb1b |
| gng13b2 | 6.1E-32 | 2.5959064 | 0.206 | 0.064 | 1.99E-27 | Perivascular | gng13b |
| rhogb4 | 1.2E-31 | 2.0129168 | 0.218 | 0.069 | 3.8E-27 | Perivascular | rhogb |
| kctd12.24 | 3.4E-31 | 1.8010375 | 0.431 | 0.225 | 1.12E-26 | Perivascular | kctd12.2 |
| mmrn2a | 1.3E-30 | 2.6143949 | 0.114 | 0.024 | 4.37E-26 | Perivascular | mmrn2a |
| palm2akap2.15 | 2.2E-30 | 2.1726268 | 0.239 | 0.084 | 7.05E-26 | Perivascular | palm2akap2.1 |
| pfn26 | 6.1E-30 | 1.1325095 | 0.68 | 0.538 | 1.99E-25 | Perivascular | pfn2 |
| ptenb5 | 1E-29 | 2.0743922 | 0.363 | 0.174 | 3.38E-25 | Perivascular | ptenb |
| pdgfab2 | 1.4E-29 | 2.7109197 | 0.145 | 0.037 | 4.39E-25 | Perivascular | pdgfab |
| dock72 | 1.4E-29 | 1.7750775 | 0.429 | 0.231 | 4.5E-25 | Perivascular | dock7 |
| hic14 | 1.6E-29 | 2.2368644 | 0.201 | 0.064 | 5.12E-25 | Perivascular | hic1 |
| utrn4 | 3.5E-29 | 1.7257787 | 0.383 | 0.187 | 1.14E-24 | Perivascular | utrn |

| | | | | | | | |
|---|---|---|---|---|---|---|---|
| arhgdig2 | 5.8E-29 | 2.2443908 | 0.226 | 0.079 | 1.89E-24 | Perivascular | arhgdig |
| col6a4a4 | 6.2E-29 | 2.5030354 | 0.183 | 0.056 | 2.03E-24 | Perivascular | col6a4a |
| zgc:664331 | 1.8E-28 | 2.1712301 | 0.277 | 0.112 | 5.77E-24 | Perivascular | zgc:66433 |
| iscub5 | 5.7E-28 | 2.0806994 | 0.381 | 0.192 | 1.84E-23 | Perivascular | iscub |
| fgd4a | 8.6E-28 | 2.6974102 | 0.117 | 0.027 | 2.78E-23 | Perivascular | fgd4a |
| tbx2a4 | 9E-28 | 2.2478437 | 0.19 | 0.061 | 2.92E-23 | Perivascular | tbx2a |
| flna6 | 1.1E-27 | 1.1218395 | 0.688 | 0.543 | 3.63E-23 | Perivascular | flna |
| mt-co23 | 2.3E-27 | 0.4791218 | 0.997 | 0.998 | 7.54E-23 | Perivascular | mt-co2 |
| rhoca3 | 3.7E-27 | 1.542864 | 0.442 | 0.257 | 1.22E-22 | Perivascular | rhoca |
| ppp1r9alb2 | 4.8E-27 | 2.4096607 | 0.15 | 0.042 | 1.55E-22 | Perivascular | ppp1r9alb |
| ywhabl4 | 5.7E-27 | 1.4290822 | 0.561 | 0.418 | 1.84E-22 | Perivascular | ywhabl |
| ilk3 | 6.8E-27 | 1.6339153 | 0.365 | 0.179 | 2.21E-22 | Perivascular | ilk |
| gask1b3 | 7.3E-27 | 2.5047901 | 0.142 | 0.039 | 2.39E-22 | Perivascular | gask1b |
| itm2ba6 | 4.4E-26 | 0.8435917 | 0.81 | 0.741 | 1.45E-21 | Perivascular | itm2ba |
| myo1cb3 | 1.7E-25 | 1.5966599 | 0.431 | 0.253 | 5.39E-21 | Perivascular | myo1cb |
| fam107b6 | 1.7E-25 | 1.3085049 | 0.404 | 0.208 | 5.62E-21 | Perivascular | fam107b |
| plekhg23 | 2E-25 | 2.0960875 | 0.206 | 0.073 | 6.6E-21 | Perivascular | plekhg2 |
| efhd21 | 4.2E-25 | 2.441023 | 0.15 | 0.044 | 1.38E-20 | Perivascular | efhd2 |
| prdx65 | 4.7E-25 | 1.7354579 | 0.495 | 0.344 | 1.52E-20 | Perivascular | prdx6 |
| susd61 | 5.8E-25 | 1.5359764 | 0.396 | 0.214 | 1.88E-20 | Perivascular | susd6 |
| rsu1 | 6.1E-25 | 2.2901557 | 0.155 | 0.046 | 1.99E-20 | Perivascular | rsu1 |
| mt-cyb4 | 7.7E-25 | 0.4620056 | 0.987 | 0.975 | 2.51E-20 | Perivascular | mt-cyb |
| gsna3 | 1.1E-24 | 1.2872151 | 0.525 | 0.357 | 3.47E-20 | Perivascular | gsna |
| capza1b4 | 1.1E-24 | 1.3148227 | 0.533 | 0.366 | 3.55E-20 | Perivascular | capza1b |
| pim17 | 1.3E-24 | 1.3179707 | 0.706 | 0.544 | 4.14E-20 | Perivascular | pim1 |
| tuba8l47 | 1.6E-24 | 0.8682015 | 0.792 | 0.734 | 5.12E-20 | Perivascular | tuba8l4 |
| pacsin3 | 1.9E-24 | 2.7233826 | 0.117 | 0.03 | 6.06E-20 | Perivascular | pacsin3 |
| mcl1b6 | 2.1E-24 | 1.2392471 | 0.622 | 0.482 | 6.73E-20 | Perivascular | mcl1b |
| clic44 | 3.1E-24 | 1.729226 | 0.368 | 0.198 | 1.02E-19 | Perivascular | clic4 |
| plk2b | 4.2E-24 | 2.9629222 | 0.114 | 0.029 | 1.38E-19 | Perivascular | plk2b |
| dnajb6b | 5.3E-24 | 2.5348682 | 0.114 | 0.029 | 1.73E-19 | Perivascular | dnajb6b |
| mob2a4 | 7.2E-24 | 1.8756461 | 0.294 | 0.137 | 2.36E-19 | Perivascular | mob2a |
| prkacaa | 9.1E-24 | 2.4708403 | 0.135 | 0.038 | 2.95E-19 | Perivascular | prkacaa |
| idh27 | 9.9E-24 | 1.2869582 | 0.51 | 0.343 | 3.23E-19 | Perivascular | idh2 |
| esyt2a3 | 1.5E-23 | 1.6582918 | 0.338 | 0.169 | 4.73E-19 | Perivascular | esyt2a |
| synm6 | 1.7E-23 | 1.3475076 | 0.414 | 0.23 | 5.45E-19 | Perivascular | synm |
| tuba8l34 | 1.8E-23 | 1.4931556 | 0.561 | 0.425 | 5.83E-19 | Perivascular | tuba8l3 |
| tbc1d1 | 1.9E-23 | 2.8755348 | 0.102 | 0.024 | 6.07E-19 | Perivascular | tbc1d1 |
| cirbpb4 | 3E-23 | 0.6625725 | 0.886 | 0.834 | 9.65E-19 | Perivascular | cirbpb |
| nr4a35 | 3.5E-23 | 2.7905479 | 0.157 | 0.05 | 1.15E-18 | Perivascular | nr4a3 |
| tspan18b5 | 3.6E-23 | 1.9509037 | 0.299 | 0.141 | 1.16E-18 | Perivascular | tspan18b |
| rap1b3 | 4.2E-23 | 1.2168709 | 0.53 | 0.365 | 1.37E-18 | Perivascular | rap1b |
| mt-co12 | 1E-22 | 0.4319011 | 1 | 0.995 | 3.25E-18 | Perivascular | mt-co1 |
| baz2ba7 | 1.3E-22 | 1.5277513 | 0.338 | 0.169 | 4.16E-18 | Perivascular | baz2ba |
| ppp1r12a2 | 1.4E-22 | 1.8188315 | 0.279 | 0.126 | 4.42E-18 | Perivascular | ppp1r12a |
| rac1a3 | 4.6E-22 | 1.7015174 | 0.348 | 0.187 | 1.5E-17 | Perivascular | rac1a |
| atp5pf4 | 5.1E-22 | 1.1425743 | 0.548 | 0.404 | 1.67E-17 | Perivascular | atp5pf |
| cyp26b14 | 6E-22 | 2.1109189 | 0.508 | 0.368 | 1.95E-17 | Perivascular | cyp26b1 |

| | | | | | | | |
|---|---|---|---|---|---|---|---|
| itpr1a | 8.2E-22 | 2.1205796 | 0.129 | 0.037 | 2.65E-17 | Perivascular | itpr1a |
| plpp2a1 | 1.3E-21 | 2.1512591 | 0.147 | 0.047 | 4.1E-17 | Perivascular | plpp2a |
| ubc6 | 1.3E-21 | 0.7188652 | 0.764 | 0.662 | 4.29E-17 | Perivascular | ubc |
| hsp90ab12 | 1.3E-21 | 0.3563615 | 0.995 | 0.993 | 4.34E-17 | Perivascular | hsp90ab1 |
| nt5c2b2 | 1.5E-21 | 2.1969544 | 0.173 | 0.061 | 4.72E-17 | Perivascular | nt5c2b |
| si:ch211-222k6.33 | 2E-21 | 1.8968289 | 0.251 | 0.11 | 6.42E-17 | Perivascular | si:ch211-222k6.3 |
| tns2a2 | 4.3E-21 | 2.0912249 | 0.17 | 0.06 | 1.39E-16 | Perivascular | tns2a |
| si:ch1073-429i10.33 | 6.8E-21 | 1.2538273 | 0.541 | 0.403 | 2.21E-16 | Perivascular | si:ch1073-429i10.3 |
| lipf3 | 7.5E-21 | 1.3704501 | 0.365 | 0.197 | 2.44E-16 | Perivascular | lipf |
| kdm2bb | 9E-21 | 2.2610042 | 0.162 | 0.056 | 2.94E-16 | Perivascular | kdm2bb |
| inka1a4 | 1E-20 | 1.5493929 | 0.332 | 0.174 | 3.28E-16 | Perivascular | inka1a |
| cxxc5b1 | 1.3E-20 | 2.8620518 | 0.107 | 0.029 | 4.38E-16 | Perivascular | cxxc5b |
| BX571880.1 | 1.5E-20 | 1.9626736 | 0.135 | 0.041 | 4.81E-16 | Perivascular | BX571880.1 |
| rgmb6 | 1.6E-20 | 1.8729672 | 0.259 | 0.119 | 5.27E-16 | Perivascular | rgmb |
| sema4ba5 | 1.8E-20 | 1.8482142 | 0.211 | 0.084 | 6.01E-16 | Perivascular | sema4ba |
| CR847566.13 | 2.7E-20 | 1.8035453 | 0.239 | 0.104 | 8.92E-16 | Perivascular | CR847566.1 |
| pbxip1b4 | 2.8E-20 | 1.5635189 | 0.398 | 0.24 | 8.94E-16 | Perivascular | pbxip1b |
| dgkza2 | 4E-20 | 2.1070735 | 0.162 | 0.057 | 1.3E-15 | Perivascular | dgkza |
| rhocb3 | 4.7E-20 | 1.8240067 | 0.292 | 0.147 | 1.53E-15 | Perivascular | rhocb |
| frzb8 | 4.9E-20 | 1.7468779 | 0.277 | 0.129 | 1.6E-15 | Perivascular | frzb |
| otud12 | 5.7E-20 | 1.9046026 | 0.165 | 0.058 | 1.84E-15 | Perivascular | otud1 |
| tuba1a4 | 6E-20 | 1.7289008 | 0.216 | 0.088 | 1.97E-15 | Perivascular | tuba1a |
| tgm2b2 | 6.4E-20 | 1.8436825 | 0.142 | 0.046 | 2.07E-15 | Perivascular | tgm2b |
| dab2ipb4 | 6.4E-20 | 1.7640225 | 0.249 | 0.113 | 2.09E-15 | Perivascular | dab2ipb |
| phlda27 | 7.5E-20 | 1.601579 | 0.383 | 0.219 | 2.45E-15 | Perivascular | phlda2 |
| CABZ01080702.12 | 9E-20 | 1.2486141 | 0.464 | 0.301 | 2.91E-15 | Perivascular | CABZ01080702.1 |
| ptmab7 | 1.2E-19 | 0.5174074 | 0.86 | 0.715 | 3.96E-15 | Perivascular | ptmab |
| rtn4rl2a1 | 1.6E-19 | 2.9679115 | 0.104 | 0.028 | 5.3E-15 | Perivascular | rtn4rl2a |
| actn16 | 1.8E-19 | 1.15513 | 0.475 | 0.317 | 5.89E-15 | Perivascular | actn1 |
| palm2akap2 | 1.8E-19 | 2.3310099 | 0.14 | 0.046 | 6.01E-15 | Perivascular | palm2akap2 |
| f3b8 | 2.3E-19 | 0.826106 | 0.566 | 0.379 | 7.39E-15 | Perivascular | f3b |
| gadd45ba5 | 2.7E-19 | 0.9874692 | 0.86 | 0.804 | 8.7E-15 | Perivascular | gadd45ba |
| st8sia6 | 2.8E-19 | 2.0968709 | 0.147 | 0.05 | 9.21E-15 | Perivascular | st8sia6 |
| eno1a4 | 3E-19 | 1.1658855 | 0.492 | 0.34 | 9.82E-15 | Perivascular | eno1a |
| socs3a4 | 4.7E-19 | 0.8378297 | 0.865 | 0.795 | 1.51E-14 | Perivascular | socs3a |
| zgc:1583435 | 4.8E-19 | 1.4774231 | 0.525 | 0.377 | 1.57E-14 | Perivascular | zgc:158343 |
| mt-co33 | 9.2E-19 | 0.3780552 | 0.997 | 0.997 | 2.99E-14 | Perivascular | mt-co3 |
| gna12a3 | 1.2E-18 | 1.9483327 | 0.188 | 0.075 | 4.02E-14 | Perivascular | gna12a |
| adam95 | 1.5E-18 | 1.8943742 | 0.175 | 0.067 | 4.87E-14 | Perivascular | adam9 |
| plcb3 | 1.9E-18 | 2.1282417 | 0.122 | 0.038 | 6.09E-14 | Perivascular | plcb3 |
| EPB41L23 | 2E-18 | 1.5383959 | 0.264 | 0.126 | 6.42E-14 | Perivascular | EPB41L2 |
| mrtfbb2 | 2.3E-18 | 1.8049621 | 0.183 | 0.072 | 7.59E-14 | Perivascular | mrtfbb |
| pdcd7 | 2.6E-18 | 2.2763664 | 0.137 | 0.046 | 8.52E-14 | Perivascular | pdcd7 |
| ppp1r3cb1 | 2.8E-18 | 2.237125 | 0.142 | 0.049 | 9.23E-14 | Perivascular | ppp1r3cb |
| aldoaa6 | 3.2E-18 | 1.1449431 | 0.508 | 0.368 | 1.05E-13 | Perivascular | aldoaa |
| sox4a7 | 3.7E-18 | 1.7352888 | 0.34 | 0.194 | 1.2E-13 | Perivascular | sox4a |
| pfn12 | 4.7E-18 | 1.1009417 | 0.462 | 0.308 | 1.53E-13 | Perivascular | pfn1 |
| CR855337.12 | 5.3E-18 | 1.987652 | 0.213 | 0.094 | 1.72E-13 | Perivascular | CR855337.1 |

| | | | | | | | |
|---|---|---|---|---|---|---|---|
| b2ml9 | 5.4E-18 | 0.6129811 | 0.909 | 0.898 | 1.77E-13 | Perivascular | b2ml |
| hsp90b14 | 9.6E-18 | 1.2275517 | 0.492 | 0.345 | 3.11E-13 | Perivascular | hsp90b1 |
| arhgef1b4 | 1.2E-17 | 1.2771708 | 0.416 | 0.26 | 3.92E-13 | Perivascular | arhgef1b |
| cnn24 | 2.2E-17 | 1.0013016 | 0.574 | 0.445 | 7.08E-13 | Perivascular | cnn2 |
| lrch4 | 2.6E-17 | 2.5290762 | 0.122 | 0.04 | 8.53E-13 | Perivascular | lrch4 |
| hmgb1b5 | 2.8E-17 | 1.2349148 | 0.404 | 0.252 | 9.21E-13 | Perivascular | hmgb1b |
| ehbp1l1a6 | 3.1E-17 | 1.5653162 | 0.213 | 0.094 | 1E-12 | Perivascular | ehbp1l1a |
| ppdpfb5 | 3.1E-17 | 0.5642722 | 0.83 | 0.778 | 1.02E-12 | Perivascular | ppdpfb |
| ddx511 | 3.2E-17 | 0.8180474 | 0.706 | 0.597 | 1.03E-12 | Perivascular | ddx5 |
| hmgn22 | 3.9E-17 | 1.4156328 | 0.662 | 0.566 | 1.28E-12 | Perivascular | hmgn2 |
| si:dkey-238c7.165 | 4.8E-17 | 1.5716082 | 0.302 | 0.165 | 1.55E-12 | Perivascular | si:dkey-238c7.16 |
| mef2d4 | 6.8E-17 | 1.4863082 | 0.332 | 0.187 | 2.21E-12 | Perivascular | mef2d |
| nr4a16 | 7.7E-17 | 1.5551559 | 0.223 | 0.101 | 2.52E-12 | Perivascular | nr4a1 |
| bahcc1b1 | 1.2E-16 | 1.7440019 | 0.246 | 0.121 | 3.99E-12 | Perivascular | bahcc1b |
| luzp14 | 1.5E-16 | 1.3789331 | 0.317 | 0.174 | 4.96E-12 | Perivascular | luzp1 |
| krt18a.16 | 1.6E-16 | 0.5130209 | 0.906 | 0.842 | 5.27E-12 | Perivascular | krt18a.1 |
| hspa84 | 1.9E-16 | 0.3633233 | 0.952 | 0.937 | 6.24E-12 | Perivascular | hspa8 |
| dapk31 | 2.8E-16 | 2.1235762 | 0.117 | 0.038 | 9.26E-12 | Perivascular | dapk3 |
| lpp6 | 3.2E-16 | 1.1774074 | 0.404 | 0.253 | 1.03E-11 | Perivascular | lpp |
| c7a6 | 3.6E-16 | 1.2670271 | 0.32 | 0.174 | 1.18E-11 | Perivascular | c7a |
| id2a8 | 3.7E-16 | 1.1616788 | 0.607 | 0.503 | 1.2E-11 | Perivascular | id2a |
| atp5mc3b4 | 5E-16 | 0.6307968 | 0.726 | 0.623 | 1.62E-11 | Perivascular | atp5mc3b |
| anxa133 | 6.7E-16 | 1.0841719 | 0.477 | 0.345 | 2.17E-11 | Perivascular | anxa13 |
| ppap2d5 | 1.1E-15 | 1.3418763 | 0.373 | 0.229 | 3.49E-11 | Perivascular | ppap2d |
| bcl2l105 | 1.1E-15 | 1.6458767 | 0.31 | 0.179 | 3.52E-11 | Perivascular | bcl2l10 |
| tpm25 | 3.6E-15 | 2.0002562 | 0.231 | 0.116 | 1.16E-10 | Perivascular | tpm2 |
| soga14 | 3.8E-15 | 1.5018488 | 0.249 | 0.126 | 1.24E-10 | Perivascular | soga1 |
| nfkbiab7 | 4.5E-15 | 0.9797419 | 0.68 | 0.583 | 1.47E-10 | Perivascular | nfkbiab |
| chmp5b3 | 5.1E-15 | 1.2482063 | 0.363 | 0.229 | 1.67E-10 | Perivascular | chmp5b |
| fstl1b6 | 5.3E-15 | 0.5462368 | 0.782 | 0.757 | 1.72E-10 | Perivascular | fstl1b |
| si:ch211-222l21.16 | 5.8E-15 | 0.4661378 | 0.713 | 0.554 | 1.9E-10 | Perivascular | si:ch211-222l21.1 |
| vclb | 5.9E-15 | 2.0956237 | 0.112 | 0.037 | 1.93E-10 | Perivascular | vclb |
| hmgn62 | 1E-14 | 0.7413834 | 0.665 | 0.603 | 3.32E-10 | Perivascular | hmgn6 |
| tmem176l.16 | 1.1E-14 | 0.6646774 | 0.784 | 0.742 | 3.45E-10 | Perivascular | tmem176l.1 |
| fhl1a1 | 1.2E-14 | 1.859531 | 0.107 | 0.035 | 3.85E-10 | Perivascular | fhl1a |
| pnrc25 | 1.2E-14 | 0.6680093 | 0.868 | 0.882 | 4.01E-10 | Perivascular | pnrc2 |
| chp12 | 1.7E-14 | 2.1589475 | 0.14 | 0.054 | 5.37E-10 | Perivascular | chp1 |
| zgc:922426 | 1.7E-14 | 1.7808245 | 0.221 | 0.111 | 5.65E-10 | Perivascular | zgc:92242 |
| kifap3a4 | 1.9E-14 | 1.5431155 | 0.201 | 0.092 | 6.14E-10 | Perivascular | kifap3a |
| sept7a4 | 2.1E-14 | 1.1717905 | 0.475 | 0.361 | 6.7E-10 | Perivascular | sept7a |
| banf13 | 2.1E-14 | 1.5962312 | 0.246 | 0.127 | 6.75E-10 | Perivascular | banf1 |
| dpysl35 | 2.4E-14 | 1.1014713 | 0.297 | 0.161 | 7.72E-10 | Perivascular | dpysl3 |
| chd4a2 | 4.6E-14 | 1.2416274 | 0.358 | 0.228 | 1.5E-09 | Perivascular | chd4a |
| palld2 | 4.9E-14 | 1.7770803 | 0.266 | 0.147 | 1.58E-09 | Perivascular | palld |
| zgc:1629446 | 6.4E-14 | 1.2210022 | 0.416 | 0.293 | 2.09E-09 | Perivascular | zgc:162944 |
| trak1a3 | 9.1E-14 | 1.8156254 | 0.147 | 0.06 | 2.95E-09 | Perivascular | trak1a |
| khdrbs1a2 | 9.3E-14 | 0.680231 | 0.703 | 0.653 | 3.04E-09 | Perivascular | khdrbs1a |
| pea15 | 9.7E-14 | 2.0682636 | 0.104 | 0.035 | 3.16E-09 | Perivascular | pea15 |

| | | | | | | | |
|---|---|---|---|---|---|---|---|
| cacnb2a1 | 1.3E-13 | 1.9981735 | 0.124 | 0.047 | 4.34E-09 | Perivascular | cacnb2a |
| cyp3c15 | 1.6E-13 | 1.564499 | 0.198 | 0.095 | 5.08E-09 | Perivascular | cyp3c1 |
| eif4ebp24 | 1.6E-13 | 1.4491777 | 0.338 | 0.215 | 5.16E-09 | Perivascular | eif4ebp2 |
| cox6a22 | 1.6E-13 | 1.8568813 | 0.127 | 0.048 | 5.35E-09 | Perivascular | cox6a2 |
| crfb42 | 2.1E-13 | 1.8498036 | 0.211 | 0.105 | 6.93E-09 | Perivascular | crfb4 |
| cyb5r14 | 3.5E-13 | 1.2980436 | 0.322 | 0.201 | 1.14E-08 | Perivascular | cyb5r1 |
| crema5 | 3.9E-13 | 1.7495114 | 0.213 | 0.109 | 1.28E-08 | Perivascular | crema |
| ywhaba3 | 4.9E-13 | 0.8573866 | 0.579 | 0.507 | 1.6E-08 | Perivascular | ywhaba |
| cav24 | 5.7E-13 | 1.7385242 | 0.287 | 0.172 | 1.86E-08 | Perivascular | cav2 |
| gadd45bb8 | 7E-13 | 1.5168306 | 0.345 | 0.22 | 2.27E-08 | Perivascular | gadd45bb |
| pde10a3 | 7.4E-13 | 1.9384651 | 0.147 | 0.062 | 2.4E-08 | Perivascular | pde10a |
| atp2b1a3 | 1.1E-12 | 1.2692322 | 0.365 | 0.246 | 3.59E-08 | Perivascular | atp2b1a |
| fermt23 | 1.3E-12 | 1.5351245 | 0.198 | 0.097 | 4.26E-08 | Perivascular | fermt2 |
| ywhaqa4 | 1.3E-12 | 1.180221 | 0.391 | 0.273 | 4.35E-08 | Perivascular | ywhaqa |
| pgam1a6 | 1.9E-12 | 1.2105732 | 0.307 | 0.187 | 6.24E-08 | Perivascular | pgam1a |
| myo18ab2 | 1.9E-12 | 1.2704264 | 0.305 | 0.187 | 6.26E-08 | Perivascular | myo18ab |
| atp5f1b5 | 2E-12 | 0.8090977 | 0.617 | 0.558 | 6.41E-08 | Perivascular | atp5f1b |
| sgip1a3 | 2E-12 | 1.9963522 | 0.129 | 0.052 | 6.65E-08 | Perivascular | sgip1a |
| txnipa2 | 2.1E-12 | 1.4414398 | 0.363 | 0.244 | 6.79E-08 | Perivascular | txnipa |
| glipr2l2 | 2.2E-12 | 1.618996 | 0.109 | 0.04 | 7.16E-08 | Perivascular | glipr2l |
| tmem35 | 2.6E-12 | 2.0542005 | 0.102 | 0.036 | 8.56E-08 | Perivascular | tmem35 |
| tril8 | 2.7E-12 | 1.4542638 | 0.211 | 0.106 | 8.86E-08 | Perivascular | tril |
| epn26 | 3.1E-12 | 1.5962583 | 0.211 | 0.109 | 1.02E-07 | Perivascular | epn2 |
| olfml2a2 | 3.3E-12 | 1.9040788 | 0.107 | 0.039 | 1.06E-07 | Perivascular | olfml2a |
| cap12 | 3.5E-12 | 1.8125311 | 0.124 | 0.049 | 1.13E-07 | Perivascular | cap1 |
| tent5ba4 | 5.8E-12 | 1.4083177 | 0.254 | 0.143 | 1.9E-07 | Perivascular | tent5ba |
| daam1a2 | 6.9E-12 | 1.356672 | 0.317 | 0.204 | 2.24E-07 | Perivascular | daam1a |
| stim1a1 | 7.8E-12 | 1.8677158 | 0.137 | 0.058 | 2.55E-07 | Perivascular | stim1a |
| psme14 | 7.9E-12 | 1.4133681 | 0.216 | 0.113 | 2.56E-07 | Perivascular | psme1 |
| ctsh6 | 8.3E-12 | 1.5396398 | 0.211 | 0.11 | 2.7E-07 | Perivascular | ctsh |
| ehbp1l1b7 | 8.8E-12 | 1.223695 | 0.236 | 0.127 | 2.88E-07 | Perivascular | ehbp1l1b |
| si:ch211-51c14.1 | 1E-11 | 1.8569494 | 0.109 | 0.041 | 3.29E-07 | Perivascular | si:ch211-51c14.1 |
| camk2g23 | 1.1E-11 | 1.7789113 | 0.226 | 0.126 | 3.53E-07 | Perivascular | camk2g2 |
| sipa1l21 | 1.4E-11 | 1.5929544 | 0.129 | 0.054 | 4.57E-07 | Perivascular | sipa1l2 |
| rab135 | 1.4E-11 | 1.1458409 | 0.34 | 0.222 | 4.65E-07 | Perivascular | rab13 |
| C25H11orf968 | 1.6E-11 | 0.9972772 | 0.5 | 0.383 | 5.06E-07 | Perivascular | C25H11orf96 |
| vaspb5 | 1.6E-11 | 1.6192478 | 0.195 | 0.1 | 5.13E-07 | Perivascular | vaspb |
| ier2b5 | 1.7E-11 | 0.9226701 | 0.723 | 0.65 | 5.41E-07 | Perivascular | ier2b |
| litaf1 | 1.9E-11 | 1.4996959 | 0.218 | 0.117 | 6.11E-07 | Perivascular | litaf |
| cox7c4 | 1.9E-11 | 1.079019 | 0.411 | 0.3 | 6.28E-07 | Perivascular | cox7c |
| lasp14 | 2.1E-11 | 0.92853 | 0.543 | 0.472 | 6.75E-07 | Perivascular | lasp1 |
| hirip31 | 2.1E-11 | 1.3077534 | 0.231 | 0.126 | 6.96E-07 | Perivascular | hirip3 |
| MYADM3 | 2.9E-11 | 1.373337 | 0.193 | 0.097 | 9.54E-07 | Perivascular | MYADM |
| h3f3b.1.12 | 3.1E-11 | 1.2581315 | 0.282 | 0.171 | 1.01E-06 | Perivascular | h3f3b.1.1 |
| mhc1zba7 | 3.3E-11 | 0.8004425 | 0.635 | 0.584 | 1.06E-06 | Perivascular | mhc1zba |
| foxc1b4 | 4E-11 | 1.8177845 | 0.142 | 0.063 | 1.29E-06 | Perivascular | foxc1b |
| ZFP36L1.16 | 4.2E-11 | 0.4089478 | 0.916 | 0.873 | 1.37E-06 | Perivascular | ZFP36L1.1 |
| wwc3 | 4.6E-11 | 1.9988017 | 0.114 | 0.046 | 1.5E-06 | Perivascular | wwc3 |

| | | | | | | | |
|---|---|---|---|---|---|---|---|
| sh3bgrl32 | 5E-11 | 1.4545196 | 0.236 | 0.134 | 1.61E-06 | Perivascular | sh3bgrl3 |
| gnb1b3 | 5E-11 | 0.6998793 | 0.622 | 0.568 | 1.62E-06 | Perivascular | gnb1b |
| eef1db6 | 5.9E-11 | 0.6084907 | 0.645 | 0.581 | 1.92E-06 | Perivascular | eef1db |
| anxa5b4 | 7.4E-11 | 0.8717317 | 0.492 | 0.403 | 2.4E-06 | Perivascular | anxa5b |
| ppm1la2 | 7.8E-11 | 1.5637575 | 0.155 | 0.072 | 2.52E-06 | Perivascular | ppm1la |
| plekha61 | 9.5E-11 | 1.8789376 | 0.132 | 0.058 | 3.09E-06 | Perivascular | plekha6 |
| COX5B.15 | 1.2E-10 | 0.6471518 | 0.602 | 0.542 | 3.82E-06 | Perivascular | COX5B.1 |
| sema6e | 1.3E-10 | 1.6429456 | 0.145 | 0.066 | 4.19E-06 | Perivascular | sema6e |
| rock2a2 | 1.7E-10 | 1.1896776 | 0.305 | 0.196 | 5.53E-06 | Perivascular | rock2a |
| pgk16 | 1.8E-10 | 1.3226619 | 0.312 | 0.207 | 5.73E-06 | Perivascular | pgk1 |
| ralgapa1 | 1.8E-10 | 1.7258239 | 0.155 | 0.074 | 5.93E-06 | Perivascular | ralgapa1 |
| ace3 | 1.9E-10 | 1.8781323 | 0.119 | 0.051 | 6.16E-06 | Perivascular | ace |
| hipk3b2 | 1.9E-10 | 1.603812 | 0.16 | 0.077 | 6.24E-06 | Perivascular | hipk3b |
| sept7b4 | 2.1E-10 | 1.1843388 | 0.307 | 0.198 | 6.74E-06 | Perivascular | sept7b |
| ywhaqb3 | 2.4E-10 | 0.6445599 | 0.591 | 0.538 | 7.95E-06 | Perivascular | ywhaqb |
| fkbp1aa4 | 2.5E-10 | 0.7456843 | 0.566 | 0.506 | 8.27E-06 | Perivascular | fkbp1aa |
| cox4i12 | 2.6E-10 | 0.4584493 | 0.789 | 0.778 | 8.57E-06 | Perivascular | cox4i1 |
| baz1b3 | 2.8E-10 | 1.1991791 | 0.282 | 0.174 | 9.06E-06 | Perivascular | baz1b |
| wu:fj16a037 | 3.2E-10 | 0.836691 | 0.315 | 0.192 | 1.05E-05 | Perivascular | wu:fj16a03 |
| osbpl5 | 3.5E-10 | 1.856672 | 0.117 | 0.05 | 1.15E-05 | Perivascular | osbpl5 |
| sdcbp25 | 4.3E-10 | 0.5743836 | 0.65 | 0.599 | 1.41E-05 | Perivascular | sdcbp2 |
| si:ch211-288d18.12 | 4.7E-10 | 1.7369924 | 0.162 | 0.082 | 1.53E-05 | Perivascular | si:ch211-288d18.1 |
| nutf2l5 | 4.8E-10 | 1.5770247 | 0.188 | 0.101 | 1.56E-05 | Perivascular | nutf2l |
| atrx4 | 4.8E-10 | 0.7636072 | 0.546 | 0.467 | 1.57E-05 | Perivascular | atrx |
| cox6a15 | 5.7E-10 | 0.7075162 | 0.563 | 0.492 | 1.85E-05 | Perivascular | cox6a1 |
| abracl4 | 6.1E-10 | 1.2599062 | 0.203 | 0.109 | 1.98E-05 | Perivascular | abracl |
| pfkfb36 | 6.1E-10 | 1.6457754 | 0.155 | 0.076 | 1.99E-05 | Perivascular | pfkfb3 |
| sorbs17 | 6.7E-10 | 1.4147243 | 0.228 | 0.134 | 2.18E-05 | Perivascular | sorbs1 |
| cremb3 | 6.8E-10 | 1.8015149 | 0.107 | 0.044 | 2.22E-05 | Perivascular | cremb |
| si:ch211-165e15.15 | 8.2E-10 | 1.5363819 | 0.142 | 0.067 | 2.68E-05 | Perivascular | si:ch211-165e15.1 |
| prkar1aa3 | 9.2E-10 | 1.3818571 | 0.241 | 0.144 | 3E-05 | Perivascular | prkar1aa |
| tpi1b4 | 1.1E-09 | 0.943889 | 0.449 | 0.358 | 3.47E-05 | Perivascular | tpi1b |
| irf2bpl4 | 1.2E-09 | 1.1962546 | 0.302 | 0.199 | 3.84E-05 | Perivascular | irf2bpl |
| magi1b | 1.7E-09 | 1.8478287 | 0.127 | 0.058 | 5.47E-05 | Perivascular | magi1b |
| hdac9b3 | 1.9E-09 | 1.4935104 | 0.145 | 0.07 | 6.03E-05 | Perivascular | hdac9b |
| slc25a3b5 | 1.9E-09 | 0.6875947 | 0.599 | 0.552 | 6.3E-05 | Perivascular | slc25a3b |
| dock10 | 1.9E-09 | 2.0408213 | 0.102 | 0.042 | 6.31E-05 | Perivascular | dock10 |
| angptl2b5 | 2.4E-09 | 1.0879965 | 0.239 | 0.141 | 7.77E-05 | Perivascular | angptl2b |
| nr2f1a4 | 2.4E-09 | 1.3220187 | 0.249 | 0.152 | 7.86E-05 | Perivascular | nr2f1a |
| mfap58 | 2.5E-09 | 0.515861 | 0.647 | 0.546 | 7.98E-05 | Perivascular | mfap5 |
| cfl23 | 2.5E-09 | 1.7395433 | 0.124 | 0.057 | 8.01E-05 | Perivascular | cfl2 |
| nid1a5 | 3E-09 | 1.0809672 | 0.338 | 0.235 | 9.6E-05 | Perivascular | nid1a |
| cirbpa1 | 3.6E-09 | 0.5765945 | 0.685 | 0.661 | 0.000116 | Perivascular | cirbpa |
| camk2n1a5 | 3.8E-09 | 1.4596287 | 0.162 | 0.083 | 0.000124 | Perivascular | camk2n1a |
| marcksb2 | 3.8E-09 | 1.2763122 | 0.185 | 0.101 | 0.000125 | Perivascular | marcksb |
| SYDE12 | 4.2E-09 | 1.6153623 | 0.137 | 0.066 | 0.000136 | Perivascular | SYDE1 |
| vamp34 | 4.3E-09 | 1.132518 | 0.343 | 0.25 | 0.000139 | Perivascular | vamp3 |
| ell4 | 4.4E-09 | 1.6223041 | 0.173 | 0.093 | 0.000144 | Perivascular | ell |

| | | | | | | | |
|---|---|---|---|---|---|---|---|
| bahcc1a6 | 4.8E-09 | 1.580041 | 0.165 | 0.087 | 0.000157 | Perivascular | bahcc1a |
| st6gal2a | 5.3E-09 | 2.2643779 | 0.102 | 0.043 | 0.000172 | Perivascular | st6gal2a |
| rtn35 | 5.4E-09 | 1.1450098 | 0.325 | 0.227 | 0.000176 | Perivascular | rtn3 |
| usp2a3 | 6E-09 | 1.7682827 | 0.142 | 0.071 | 0.000195 | Perivascular | usp2a |
| arpc4l3 | 7.4E-09 | 0.9836468 | 0.439 | 0.363 | 0.000242 | Perivascular | arpc4l |
| tubb53 | 7.6E-09 | 1.3793249 | 0.239 | 0.15 | 0.000246 | Perivascular | tubb5 |
| cycsb5 | 7.8E-09 | 0.8032922 | 0.467 | 0.374 | 0.000255 | Perivascular | cycsb |
| cmtm42 | 8.8E-09 | 1.4701458 | 0.201 | 0.116 | 0.000286 | Perivascular | cmtm4 |
| ppiaa3 | 9E-09 | 0.2728607 | 0.959 | 0.96 | 0.000291 | Perivascular | ppiaa |
| syt11a4 | 9.8E-09 | 1.0620494 | 0.261 | 0.165 | 0.000318 | Perivascular | syt11a |
| tagapa2 | 1E-08 | 1.4438527 | 0.15 | 0.076 | 0.000335 | Perivascular | tagapa |
| junba5 | 1.1E-08 | 0.7691985 | 0.86 | 0.846 | 0.000343 | Perivascular | junba |
| tgfb35 | 1.2E-08 | 1.539461 | 0.155 | 0.081 | 0.000392 | Perivascular | tgfb3 |
| ppm1ab2 | 1.3E-08 | 1.5559367 | 0.142 | 0.071 | 0.000408 | Perivascular | ppm1ab |
| iqgap14 | 1.3E-08 | 1.0505312 | 0.322 | 0.226 | 0.000422 | Perivascular | iqgap1 |
| prdx22 | 1.3E-08 | 0.9991872 | 0.409 | 0.33 | 0.000423 | Perivascular | prdx2 |
| niban2b2 | 1.4E-08 | 1.2441811 | 0.239 | 0.15 | 0.000467 | Perivascular | niban2b |
| grk5l | 1.4E-08 | 1.6312479 | 0.119 | 0.056 | 0.000469 | Perivascular | grk5l |
| cotl12 | 1.5E-08 | 1.2139086 | 0.185 | 0.102 | 0.000474 | Perivascular | cotl1 |
| sh2d3cb1 | 1.5E-08 | 1.6286822 | 0.114 | 0.052 | 0.000481 | Perivascular | sh2d3cb |
| mt-nd35 | 1.5E-08 | 0.7727329 | 0.442 | 0.361 | 0.000496 | Perivascular | mt-nd3 |
| sri4 | 1.5E-08 | 1.0877258 | 0.363 | 0.271 | 0.000496 | Perivascular | sri |
| clip25 | 1.6E-08 | 1.2576802 | 0.185 | 0.103 | 0.000524 | Perivascular | clip2 |
| ap1s12 | 1.7E-08 | 1.8818274 | 0.104 | 0.046 | 0.000547 | Perivascular | ap1s1 |
| icam36 | 1.8E-08 | 1.5298495 | 0.157 | 0.083 | 0.000572 | Perivascular | icam3 |
| cisd11 | 1.8E-08 | 1.5693123 | 0.132 | 0.064 | 0.00058 | Perivascular | cisd1 |
| arpc5a3 | 1.9E-08 | 1.3356009 | 0.244 | 0.155 | 0.000615 | Perivascular | arpc5a |
| lnpep2 | 2.1E-08 | 1.6067957 | 0.142 | 0.072 | 0.000668 | Perivascular | lnpep |
| ppp1r14bb4 | 2.2E-08 | 1.3563132 | 0.234 | 0.149 | 0.000701 | Perivascular | ppp1r14bb |
| slc38a27 | 2.5E-08 | 1.3348366 | 0.152 | 0.079 | 0.000801 | Perivascular | slc38a2 |
| hmgb2b3 | 2.7E-08 | 0.9309825 | 0.442 | 0.364 | 0.000881 | Perivascular | hmgb2b |
| raph1b6 | 2.9E-08 | 1.2036661 | 0.188 | 0.106 | 0.000952 | Perivascular | raph1b |
| mych7 | 2.9E-08 | 1.0588212 | 0.464 | 0.38 | 0.000955 | Perivascular | mych |
| prkar2aa3 | 3E-08 | 1.1499397 | 0.254 | 0.162 | 0.000986 | Perivascular | prkar2aa |
| nedd83 | 3.3E-08 | 1.393666 | 0.188 | 0.108 | 0.001079 | Perivascular | nedd8 |
| cox4i24 | 3.6E-08 | 0.9438697 | 0.404 | 0.321 | 0.001171 | Perivascular | cox4i2 |
| pfn2l3 | 3.7E-08 | 0.7692679 | 0.528 | 0.478 | 0.0012 | Perivascular | pfn2l |
| pkma4 | 4E-08 | 1.1954351 | 0.299 | 0.213 | 0.001286 | Perivascular | pkma |
| map3k31 | 4.2E-08 | 1.6574027 | 0.117 | 0.055 | 0.001372 | Perivascular | map3k3 |
| higd1a7 | 4.4E-08 | 0.6767173 | 0.533 | 0.458 | 0.001432 | Perivascular | higd1a |
| cdk62 | 5.1E-08 | 1.4006423 | 0.132 | 0.066 | 0.001642 | Perivascular | cdk6 |
| hsp70l5 | 5.2E-08 | 1.5588735 | 0.388 | 0.302 | 0.00168 | Perivascular | hsp70l |
| si:dkeyp-115e12.61 | 6.1E-08 | 1.8216848 | 0.107 | 0.049 | 0.00199 | Perivascular | si:dkeyp-115e12.6 |
| sat1a.211 | 6.3E-08 | 0.6413297 | 0.543 | 0.457 | 0.002057 | Perivascular | sat1a.2 |
| si:ch211-286b5.53 | 6.5E-08 | 0.9695864 | 0.345 | 0.258 | 0.002117 | Perivascular | si:ch211-286b5.5 |
| cspg43 | 6.5E-08 | 1.2691516 | 0.195 | 0.114 | 0.002123 | Perivascular | cspg4 |
| plce11 | 6.9E-08 | 1.5008474 | 0.104 | 0.047 | 0.002229 | Perivascular | plce1 |
| fosab6 | 7E-08 | 0.7289903 | 0.906 | 0.895 | 0.002286 | Perivascular | fosab |

| | | | | | | | |
|---|---|---|---|---|---|---|---|
| agap12 | 7E-08 | 1.4925354 | 0.145 | 0.076 | 0.002292 | Perivascular | agap1 |
| si:dkey-28b4.81 | 8.6E-08 | 2.0669749 | 0.107 | 0.05 | 0.002797 | Perivascular | si:dkey-28b4.8 |
| aoc27 | 8.8E-08 | 1.2938635 | 0.34 | 0.241 | 0.002866 | Perivascular | aoc2 |
| traf4a3 | 9.9E-08 | 1.6771757 | 0.142 | 0.075 | 0.003214 | Perivascular | traf4a |
| serpine16 | 1E-07 | 0.6862142 | 0.822 | 0.762 | 0.00339 | Perivascular | serpine1 |
| ube2v22 | 1.1E-07 | 0.6808908 | 0.561 | 0.531 | 0.003577 | Perivascular | ube2v2 |
| itga56 | 1.2E-07 | 1.0104137 | 0.348 | 0.257 | 0.003858 | Perivascular | itga5 |
| fkbp55 | 1.2E-07 | 1.2961193 | 0.287 | 0.197 | 0.004011 | Perivascular | fkbp5 |
| aopep5 | 1.2E-07 | 1.7002207 | 0.142 | 0.075 | 0.004032 | Perivascular | aopep |
| c6.15 | 1.3E-07 | 0.4773759 | 0.31 | 0.206 | 0.004166 | Perivascular | c6.1 |
| cdc42l4 | 1.3E-07 | 0.7470291 | 0.477 | 0.405 | 0.004203 | Perivascular | cdc42l |
| nckap13 | 1.5E-07 | 0.9935918 | 0.322 | 0.238 | 0.004742 | Perivascular | nckap1 |
| glud1b2 | 1.6E-07 | 1.3267495 | 0.183 | 0.107 | 0.005059 | Perivascular | glud1b |
| cdc42bpaa | 1.6E-07 | 1.663381 | 0.104 | 0.049 | 0.005098 | Perivascular | cdc42bpaa |
| arhgef46 | 1.7E-07 | 1.4013135 | 0.142 | 0.076 | 0.005688 | Perivascular | arhgef4 |
| specc1la5 | 1.8E-07 | 1.4989449 | 0.147 | 0.08 | 0.005823 | Perivascular | specc1la |
| cdipt4 | 1.8E-07 | 1.2782414 | 0.19 | 0.114 | 0.006006 | Perivascular | cdipt |
| stmn1b5 | 1.8E-07 | 1.1859937 | 0.203 | 0.124 | 0.006012 | Perivascular | stmn1b |
| atf63 | 2.2E-07 | 1.2946518 | 0.195 | 0.117 | 0.007172 | Perivascular | atf6 |
| pear16 | 2.2E-07 | 1.4532506 | 0.208 | 0.131 | 0.007238 | Perivascular | pear1 |
| gadd45ab7 | 2.3E-07 | 1.5895841 | 0.259 | 0.176 | 0.007452 | Perivascular | gadd45ab |
| rasal23 | 2.5E-07 | 1.4713775 | 0.137 | 0.072 | 0.008018 | Perivascular | rasal2 |
| phex4 | 2.6E-07 | 1.2557324 | 0.145 | 0.078 | 0.00841 | Perivascular | phex |
| jpt1b5 | 2.9E-07 | 0.5302939 | 0.695 | 0.692 | 0.009348 | Perivascular | jpt1b |
| rbms2b3 | 3E-07 | 1.6116125 | 0.122 | 0.063 | 0.009695 | Perivascular | rbms2b |
| ext1a3 | 3.1E-07 | 1.5474556 | 0.107 | 0.051 | 0.010232 | Perivascular | ext1a |
| nfia6 | 3.2E-07 | 0.9791801 | 0.256 | 0.17 | 0.010537 | Perivascular | nfia |
| anxa3b3 | 3.3E-07 | 1.0327737 | 0.33 | 0.248 | 0.01066 | Perivascular | anxa3b |
| dctn34 | 3.3E-07 | 1.2310399 | 0.231 | 0.151 | 0.010844 | Perivascular | dctn3 |
| txn7 | 3.5E-07 | 0.2596266 | 0.485 | 0.373 | 0.0113 | Perivascular | txn |
| si:ch1073-392o20.27 | 3.7E-07 | 1.0537318 | 0.325 | 0.243 | 0.012028 | Perivascular | si:ch1073-392o20.2 |
| cavin1b7 | 3.8E-07 | 0.3684858 | 0.706 | 0.646 | 0.012497 | Perivascular | cavin1b |
| atf36 | 4.5E-07 | 1.3131361 | 0.426 | 0.346 | 0.01454 | Perivascular | atf3 |
| plxnb1a1 | 6.1E-07 | 1.3408882 | 0.109 | 0.053 | 0.01976 | Perivascular | plxnb1a |
| rhoaa4 | 6.4E-07 | 0.6551073 | 0.482 | 0.421 | 0.020726 | Perivascular | rhoaa |
| cux1a6 | 7.1E-07 | 1.5478103 | 0.129 | 0.069 | 0.023236 | Perivascular | cux1a |
| cnbpb3 | 8.5E-07 | 0.6243574 | 0.604 | 0.577 | 0.027592 | Perivascular | cnbpb |
| igfbp1a4 | 8.7E-07 | 1.1734555 | 0.135 | 0.072 | 0.028182 | Perivascular | igfbp1a |
| rtca4 | 9.2E-07 | 0.9876571 | 0.272 | 0.188 | 0.029967 | Perivascular | rtca |
| hspa12a | 1.1E-06 | 1.6204253 | 0.117 | 0.061 | 0.036204 | Perivascular | hspa12a |
| tmem204 | 1.1E-06 | 1.5651082 | 0.102 | 0.049 | 0.036592 | Perivascular | tmem204 |
| setb3 | 1.1E-06 | 0.803247 | 0.444 | 0.376 | 0.037226 | Perivascular | setb |
| h3f3d6 | 1.2E-06 | 0.3394973 | 0.749 | 0.729 | 0.03888 | Perivascular | h3f3d |
| psme23 | 1.2E-06 | 1.3627348 | 0.14 | 0.077 | 0.040602 | Perivascular | psme2 |
| ano5b6 | 1.4E-06 | 1.2601815 | 0.152 | 0.087 | 0.045817 | Perivascular | ano5b |
| PKN21 | 1.5E-06 | 1.5974155 | 0.119 | 0.062 | 0.047473 | Perivascular | PKN2 |
| ppp1r71 | 1.5E-06 | 1.5407789 | 0.114 | 0.059 | 0.048061 | Perivascular | ppp1r7 |
| pard3ab1 | 1.5E-06 | 1.4950051 | 0.112 | 0.057 | 0.049451 | Perivascular | pard3ab |

| | | | | | | | |
|---|---|---|---|---|---|---|---|
| psmc33 | 1.6E-06 | 1.2212673 | 0.165 | 0.097 | 0.051595 | Perivascular | psmc3 |
| peli1a7 | 1.7E-06 | 0.9020641 | 0.206 | 0.129 | 0.05615 | Perivascular | peli1a |
| atp5mc15 | 1.8E-06 | 0.9230411 | 0.378 | 0.311 | 0.05888 | Perivascular | atp5mc1 |
| gng102 | 2.1E-06 | 1.5596722 | 0.119 | 0.064 | 0.068447 | Perivascular | gng10 |
| psmb53 | 2.3E-06 | 1.2307673 | 0.211 | 0.138 | 0.075366 | Perivascular | psmb5 |
| rras22 | 2.3E-06 | 1.6235515 | 0.124 | 0.068 | 0.075952 | Perivascular | rras2 |
| csnk1a12 | 2.9E-06 | 0.7269125 | 0.444 | 0.381 | 0.092847 | Perivascular | csnk1a1 |
| her97 | 2.9E-06 | 0.8438246 | 0.315 | 0.229 | 0.094242 | Perivascular | her9 |
| stom5 | 3.4E-06 | 0.9126386 | 0.345 | 0.27 | 0.109269 | Perivascular | stom |
| si:ch73-1a9.34 | 3.5E-06 | 0.5360552 | 0.711 | 0.72 | 0.113407 | Perivascular | si:ch73-1a9.3 |
| cct54 | 3.6E-06 | 0.8304303 | 0.434 | 0.382 | 0.117992 | Perivascular | cct5 |
| stard91 | 3.8E-06 | 1.2774302 | 0.102 | 0.051 | 0.124446 | Perivascular | stard9 |
| slc12a25 | 4.1E-06 | 1.5391978 | 0.216 | 0.144 | 0.131773 | Perivascular | slc12a2 |
| fosl27 | 4.1E-06 | 0.8346534 | 0.48 | 0.429 | 0.132504 | Perivascular | fosl2 |
| ptbp1b4 | 4.2E-06 | 1.0961006 | 0.261 | 0.19 | 0.136156 | Perivascular | ptbp1b |
| micu2 | 4.5E-06 | 1.2065516 | 0.112 | 0.058 | 0.145554 | Perivascular | micu2 |
| chaf1a2 | 4.5E-06 | 1.451098 | 0.104 | 0.053 | 0.147321 | Perivascular | chaf1a |
| rnf213a5 | 4.5E-06 | 1.0401177 | 0.343 | 0.269 | 0.147406 | Perivascular | rnf213a |
| sept9b4 | 4.8E-06 | 1.3570108 | 0.117 | 0.062 | 0.155467 | Perivascular | sept9b |
| atp5l2 | 4.8E-06 | 0.5236858 | 0.607 | 0.595 | 0.15641 | Perivascular | atp5l |
| tspan3a4 | 4.9E-06 | 1.1487521 | 0.234 | 0.163 | 0.158842 | Perivascular | tspan3a |
| egr2a6 | 4.9E-06 | 1.6447993 | 0.157 | 0.095 | 0.159006 | Perivascular | egr2a |
| amotl2b3 | 5.1E-06 | 1.1740588 | 0.14 | 0.08 | 0.165888 | Perivascular | amotl2b |
| phldb2a1 | 5.2E-06 | 1.213219 | 0.114 | 0.06 | 0.168327 | Perivascular | phldb2a |
| sptan15 | 5.3E-06 | 0.774701 | 0.322 | 0.245 | 0.17377 | Perivascular | sptan1 |
| psma14 | 5.4E-06 | 0.7870603 | 0.378 | 0.312 | 0.176096 | Perivascular | psma1 |
| got2b2 | 5.7E-06 | 1.5126268 | 0.147 | 0.088 | 0.18472 | Perivascular | got2b |
| RAPH16 | 6E-06 | 1.2815677 | 0.152 | 0.091 | 0.194326 | Perivascular | RAPH1 |
| arvcfb4 | 6.1E-06 | 1.2051612 | 0.168 | 0.103 | 0.197221 | Perivascular | arvcfb |
| znf106a1 | 6.2E-06 | 1.4369165 | 0.119 | 0.065 | 0.200216 | Perivascular | znf106a |
| fam171a2a3 | 6.6E-06 | 1.3985816 | 0.142 | 0.083 | 0.214578 | Perivascular | fam171a2a |
| ppp1caa4 | 6.6E-06 | 1.2369753 | 0.203 | 0.136 | 0.215878 | Perivascular | ppp1caa |
| psmc53 | 6.8E-06 | 1.0756889 | 0.246 | 0.174 | 0.221978 | Perivascular | psmc5 |
| tp534 | 7.8E-06 | 0.9319843 | 0.234 | 0.16 | 0.254001 | Perivascular | tp53 |
| lepr6 | 8.2E-06 | 1.5194322 | 0.117 | 0.064 | 0.265475 | Perivascular | lepr |
| marcksl1a7 | 9.1E-06 | 0.4368702 | 0.718 | 0.708 | 0.295175 | Perivascular | marcksl1a |
| dido12 | 9.9E-06 | 0.8016131 | 0.475 | 0.426 | 0.321686 | Perivascular | dido1 |
| fundc24 | 1E-05 | 0.8098806 | 0.368 | 0.302 | 0.334538 | Perivascular | fundc2 |
| en1b7 | 1.1E-05 | 1.4909327 | 0.173 | 0.11 | 0.346282 | Perivascular | en1b |
| arpc1a4 | 1.2E-05 | 0.8407427 | 0.376 | 0.318 | 0.406192 | Perivascular | arpc1a |
| dbn14 | 1.3E-05 | 0.8256111 | 0.277 | 0.201 | 0.41361 | Perivascular | dbn1 |
| smarca4a2 | 1.3E-05 | 1.2201501 | 0.157 | 0.096 | 0.4157 | Perivascular | smarca4a |
| cebpb6 | 1.4E-05 | 0.6265412 | 0.515 | 0.449 | 0.455832 | Perivascular | cebpb |
| spry25 | 1.4E-05 | 1.2597239 | 0.135 | 0.079 | 0.463687 | Perivascular | spry2 |
| chka5 | 1.4E-05 | 1.1135814 | 0.15 | 0.091 | 0.465221 | Perivascular | chka |
| nav33 | 1.5E-05 | 1.225499 | 0.14 | 0.083 | 0.483575 | Perivascular | nav3 |
| mrpl121 | 1.7E-05 | 1.2321738 | 0.185 | 0.124 | 0.561914 | Perivascular | mrpl12 |
| mt-nd24 | 1.8E-05 | 0.3771461 | 0.65 | 0.632 | 0.575743 | Perivascular | mt-nd2 |

| | | | | | | | |
|---|---|---|---|---|---|---|---|
| zgc:1034385 | 1.8E-05 | 1.2193979 | 0.147 | 0.089 | 0.584037 | Perivascular | zgc:103438 |
| zmp:00000007351 | 2E-05 | 1.0956687 | 0.109 | 0.059 | 0.656901 | Perivascular | zmp:0000000735 |
| cldn12 | 2E-05 | 1.424571 | 0.117 | 0.066 | 0.658308 | Perivascular | cldn12 |
| kmt2bb | 2E-05 | 0.7143958 | 0.421 | 0.36 | 0.659657 | Perivascular | kmt2bb |
| ssrp1a1 | 2.2E-05 | 1.3418857 | 0.107 | 0.058 | 0.700366 | Perivascular | ssrp1a |
| cox7b5 | 2.4E-05 | 0.6593364 | 0.444 | 0.397 | 0.795075 | Perivascular | cox7b |
| MYO1D4 | 2.4E-05 | 1.3049485 | 0.162 | 0.103 | 0.796064 | Perivascular | MYO1D |
| hsp70.25 | 2.5E-05 | 0.8391648 | 0.543 | 0.494 | 0.797844 | Perivascular | hsp70.2 |
| gstt1b5 | 2.6E-05 | 1.2065054 | 0.185 | 0.122 | 0.859101 | Perivascular | gstt1b |
| myo1ca6 | 2.7E-05 | 0.8781527 | 0.269 | 0.203 | 0.867647 | Perivascular | myo1ca |
| plxnb2a.14 | 2.7E-05 | 1.1690194 | 0.147 | 0.09 | 0.881652 | Perivascular | plxnb2a.1 |
| ddx612 | 2.8E-05 | 0.7617397 | 0.431 | 0.381 | 0.897272 | Perivascular | ddx61 |
| kif5ba3 | 2.8E-05 | 1.1226144 | 0.155 | 0.097 | 0.901406 | Perivascular | kif5ba |
| atp1b2a4 | 3E-05 | 1.1893512 | 0.137 | 0.081 | 0.962853 | Perivascular | atp1b2a |
| s1pr14 | 3E-05 | 1.2855613 | 0.15 | 0.093 | 0.967971 | Perivascular | s1pr1 |
| amotl2a4 | 3E-05 | 0.924905 | 0.226 | 0.157 | 0.976581 | Perivascular | amotl2a |
| anp32e3 | 3.1E-05 | 1.1773248 | 0.206 | 0.143 | 1 | Perivascular | anp32e |
| ano67 | 3.1E-05 | 1.1673864 | 0.188 | 0.125 | 1 | Perivascular | ano6 |
| maff9 | 3.4E-05 | 0.806209 | 0.297 | 0.224 | 1 | Perivascular | maff |
| gnai2a3 | 3.4E-05 | 0.9483422 | 0.343 | 0.283 | 1 | Perivascular | gnai2a |
| ralbb | 3.5E-05 | 1.2740956 | 0.109 | 0.061 | 1 | Perivascular | ralbb |
| ip6k2b6 | 3.5E-05 | 0.7339585 | 0.406 | 0.344 | 1 | Perivascular | ip6k2b |
| ssh2a4 | 3.7E-05 | 1.2008374 | 0.135 | 0.08 | 1 | Perivascular | ssh2a |
| tp53inp12 | 3.7E-05 | 1.7264452 | 0.107 | 0.06 | 1 | Perivascular | tp53inp1 |
| tegt3 | 3.8E-05 | 1.1727394 | 0.218 | 0.156 | 1 | Perivascular | tegt |
| dynll14 | 3.9E-05 | 0.6899131 | 0.48 | 0.427 | 1 | Perivascular | dynll1 |
| nocta6 | 4.1E-05 | 1.2294334 | 0.173 | 0.114 | 1 | Perivascular | nocta |
| arpc23 | 4.2E-05 | 0.5662061 | 0.48 | 0.445 | 1 | Perivascular | arpc2 |
| runx36 | 4.3E-05 | 0.8673949 | 0.16 | 0.274 | 1 | Perivascular | runx3 |
| nap1l12 | 4.4E-05 | 0.584058 | 0.543 | 0.527 | 1 | Perivascular | nap1l1 |
| angptl48 | 4.4E-05 | 0.7337787 | 0.49 | 0.457 | 1 | Perivascular | angptl4 |
| si:dkey-56m19.54 | 4.5E-05 | 1.1574794 | 0.14 | 0.086 | 1 | Perivascular | si:dkey-56m19.5 |
| elob.12 | 4.6E-05 | 1.0200048 | 0.287 | 0.229 | 1 | Perivascular | elob.1 |
| atp5fa14 | 4.7E-05 | 0.6037163 | 0.485 | 0.462 | 1 | Perivascular | atp5fa1 |
| ch25hl37 | 5.1E-05 | 1.4917888 | 0.162 | 0.107 | 1 | Perivascular | ch25hl3 |
| syncrip3 | 5.1E-05 | 0.5460033 | 0.515 | 0.496 | 1 | Perivascular | syncrip |
| upf3b3 | 5.2E-05 | 1.2058743 | 0.289 | 0.224 | 1 | Perivascular | upf3b |
| mbd25 | 5.3E-05 | 1.1330238 | 0.137 | 0.083 | 1 | Perivascular | mbd2 |
| pcbp23 | 5.5E-05 | 0.9815261 | 0.287 | 0.225 | 1 | Perivascular | pcbp2 |
| hsp70.35 | 6E-05 | 0.7116077 | 0.594 | 0.555 | 1 | Perivascular | hsp70.3 |
| PLEKHH34 | 6.1E-05 | 1.0524375 | 0.223 | 0.164 | 1 | Perivascular | PLEKHH3 |
| h3f3c3 | 6.4E-05 | 0.651466 | 0.495 | 0.466 | 1 | Perivascular | h3f3c |
| ppp2r2d3 | 6.4E-05 | 1.1352562 | 0.203 | 0.143 | 1 | Perivascular | ppp2r2d |
| ypel52 | 6.6E-05 | 1.4448648 | 0.112 | 0.064 | 1 | Perivascular | ypel5 |
| arhgef12b3 | 7.3E-05 | 0.7588262 | 0.266 | 0.198 | 1 | Perivascular | arhgef12b |
| atp5f1c4 | 7.3E-05 | 0.8622499 | 0.317 | 0.256 | 1 | Perivascular | atp5f1c |
| calr3b6 | 7.5E-05 | 0.8169854 | 0.33 | 0.271 | 1 | Perivascular | calr3b |
| si:dkey-23i12.74 | 8.2E-05 | 1.2580359 | 0.201 | 0.141 | 1 | Perivascular | si:dkey-23i12.7 |

| | | | | | | | |
|---|---|---|---|---|---|---|---|
| cav15 | 8.7E-05 | 0.331599 | 0.665 | 0.633 | 1 | Perivascular | cav1 |
| atf5b4 | 9.1E-05 | 1.0598136 | 0.312 | 0.253 | 1 | Perivascular | atf5b |
| hint17 | 9.3E-05 | 0.7062744 | 0.431 | 0.386 | 1 | Perivascular | hint1 |
| cnot6a2 | 9.5E-05 | 1.0160741 | 0.147 | 0.093 | 1 | Perivascular | cnot6a |
| gng53 | 9.7E-05 | 0.8738984 | 0.325 | 0.264 | 1 | Perivascular | gng5 |
| sh3bgrl5 | 0.0001 | 1.0351273 | 0.287 | 0.227 | 1 | Perivascular | sh3bgrl |
| ccng15 | 0.00011 | 0.4597741 | 0.533 | 0.506 | 1 | Perivascular | ccng1 |
| b2m8 | 0.00011 | 0.2902155 | 0.701 | 0.687 | 1 | Perivascular | b2m |
| tspo | 0.00011 | 1.4640656 | 0.117 | 0.07 | 1 | Perivascular | tspo |
| eno31 | 0.00012 | 0.7440652 | 0.353 | 0.302 | 1 | Perivascular | eno3 |
| atp5pd4 | 0.00014 | 0.6640589 | 0.411 | 0.369 | 1 | Perivascular | atp5pd |
| clic14 | 0.00014 | 1.0631706 | 0.16 | 0.105 | 1 | Perivascular | clic1 |
| ctnnd13 | 0.00014 | 0.8146878 | 0.231 | 0.166 | 1 | Perivascular | ctnnd1 |
| nid1b5 | 0.00014 | 1.1681813 | 0.129 | 0.08 | 1 | Perivascular | nid1b |
| serpinh1a6 | 0.00015 | 0.8303104 | 0.299 | 0.239 | 1 | Perivascular | serpinh1a |
| cox8a4 | 0.00016 | 0.4788379 | 0.563 | 0.564 | 1 | Perivascular | cox8a |
| cdv33 | 0.00016 | 0.7961859 | 0.297 | 0.238 | 1 | Perivascular | cdv3 |
| vdac25 | 0.00016 | 0.5321518 | 0.536 | 0.52 | 1 | Perivascular | vdac2 |
| march5l2 | 0.00018 | 1.3514358 | 0.109 | 0.064 | 1 | Perivascular | march5l |
| psmb33 | 0.00019 | 0.8043595 | 0.31 | 0.257 | 1 | Perivascular | psmb3 |
| map7d1b5 | 0.00019 | 0.9016968 | 0.251 | 0.19 | 1 | Perivascular | map7d1b |
| ppp2cb3 | 0.0002 | 0.7282256 | 0.348 | 0.292 | 1 | Perivascular | ppp2cb |
| selenow2a2 | 0.0002 | 0.9709078 | 0.221 | 0.164 | 1 | Perivascular | selenow2a |
| sqstm17 | 0.00021 | 0.6417931 | 0.36 | 0.306 | 1 | Perivascular | sqstm1 |
| acin1a3 | 0.00021 | 0.4547464 | 0.609 | 0.594 | 1 | Perivascular | acin1a |
| acadvl3 | 0.00021 | 1.1448785 | 0.15 | 0.099 | 1 | Perivascular | acadvl |
| hmgb2a1 | 0.00022 | 0.9379045 | 0.47 | 0.446 | 1 | Perivascular | hmgb2a |
| ankrd122 | 0.00023 | 1.0189061 | 0.302 | 0.25 | 1 | Perivascular | ankrd12 |
| abi1a4 | 0.00023 | 0.949056 | 0.114 | 0.068 | 1 | Perivascular | abi1a |
| sdhb4 | 0.00025 | 1.078016 | 0.231 | 0.176 | 1 | Perivascular | sdhb |
| dhx383 | 0.00025 | 1.1321525 | 0.15 | 0.099 | 1 | Perivascular | dhx38 |
| junbb5 | 0.00027 | 0.4868873 | 0.619 | 0.602 | 1 | Perivascular | junbb |
| ralbp13 | 0.00027 | 0.8138466 | 0.221 | 0.161 | 1 | Perivascular | ralbp1 |
| kdm6ba6 | 0.00028 | 0.9964157 | 0.279 | 0.222 | 1 | Perivascular | kdm6ba |
| ube2l3a3 | 0.00028 | 0.9871356 | 0.208 | 0.152 | 1 | Perivascular | ube2l3a |
| fhl3a4 | 0.0003 | 0.852866 | 0.114 | 0.069 | 1 | Perivascular | fhl3a |
| hmgn37 | 0.0003 | 0.8948564 | 0.221 | 0.163 | 1 | Perivascular | hmgn3 |
| tax1bp1b3 | 0.00033 | 0.8185575 | 0.332 | 0.282 | 1 | Perivascular | tax1bp1b |
| oaz1b8 | 0.00033 | 0.7378603 | 0.355 | 0.308 | 1 | Perivascular | oaz1b |
| pik3c2a3 | 0.00037 | 1.0965361 | 0.14 | 0.091 | 1 | Perivascular | pik3c2a |
| dab2ipa3 | 0.00037 | 0.9396351 | 0.165 | 0.113 | 1 | Perivascular | dab2ipa |
| cct46 | 0.00038 | 0.7423037 | 0.358 | 0.313 | 1 | Perivascular | cct4 |
| ubl3a6 | 0.00039 | 0.6275322 | 0.388 | 0.338 | 1 | Perivascular | ubl3a |
| rock13 | 0.00039 | 1.1165414 | 0.137 | 0.09 | 1 | Perivascular | rock1 |
| h1-104 | 0.0004 | 0.601467 | 0.266 | 0.205 | 1 | Perivascular | h1-10 |
| lrrfip1a3 | 0.0004 | 0.9447561 | 0.119 | 0.074 | 1 | Perivascular | lrrfip1a |
| napsa5 | 0.0004 | 0.7944693 | 0.228 | 0.17 | 1 | Perivascular | napsa |
| cltcb2 | 0.00041 | 0.8079716 | 0.162 | 0.109 | 1 | Perivascular | cltcb |

| | | | | | | | |
|---|---|---|---|---|---|---|---|
| zfhx36 | 0.00041 | 0.5225871 | 0.528 | 0.488 | 1 | Perivascular | zfhx3 |
| hdac13 | 0.00044 | 0.8437013 | 0.32 | 0.272 | 1 | Perivascular | hdac1 |
| si:ch1073-291c23.29 | 0.00044 | 0.547712 | 0.505 | 0.475 | 1 | Perivascular | si:ch1073-291c23.2 |
| rab11a3 | 0.00046 | 0.9207405 | 0.254 | 0.201 | 1 | Perivascular | rab11a |
| ywhabb3 | 0.00046 | 0.6325404 | 0.424 | 0.397 | 1 | Perivascular | ywhabb |
| ilf3b3 | 0.00047 | 0.8839872 | 0.241 | 0.188 | 1 | Perivascular | ilf3b |
| ndufa123 | 0.00049 | 0.915288 | 0.269 | 0.22 | 1 | Perivascular | ndufa12 |
| ndufb33 | 0.00049 | 0.7526642 | 0.33 | 0.279 | 1 | Perivascular | ndufb3 |
| ENSDARG000000356 | 0.00051 | 0.8999267 | 0.272 | 0.218 | 1 | Perivascular | ENSDARG00000035 |
| chd4b3 | 0.00052 | 0.8664049 | 0.31 | 0.26 | 1 | Perivascular | chd4b |
| gab1 | 0.00054 | 1.2178279 | 0.112 | 0.07 | 1 | Perivascular | gab1 |
| dlst3 | 0.00056 | 0.8959924 | 0.244 | 0.191 | 1 | Perivascular | dlst |
| ldha7 | 0.00057 | 0.8449452 | 0.279 | 0.231 | 1 | Perivascular | ldha |
| mtpn3 | 0.00058 | 0.7539588 | 0.348 | 0.313 | 1 | Perivascular | mtpn |
| hnrnpa0l.15 | 0.0006 | 0.3912978 | 0.657 | 0.668 | 1 | Perivascular | hnrnpa0l.1 |
| bag36 | 0.0006 | 1.2167117 | 0.114 | 0.072 | 1 | Perivascular | bag3 |
| mafba8 | 0.00061 | 0.9830964 | 0.411 | 0.391 | 1 | Perivascular | mafba |
| add3a7 | 0.00061 | 0.2978564 | 0.503 | 0.449 | 1 | Perivascular | add3a |
| kctd33 | 0.00062 | 0.940958 | 0.147 | 0.099 | 1 | Perivascular | kctd3 |
| higd2a5 | 0.00063 | 0.9878579 | 0.188 | 0.136 | 1 | Perivascular | higd2a |
| diaph25 | 0.00063 | 0.8849299 | 0.168 | 0.116 | 1 | Perivascular | diaph2 |
| snu13b3 | 0.00063 | 1.032512 | 0.231 | 0.182 | 1 | Perivascular | snu13b |
| il13ra16 | 0.00067 | 1.1758776 | 0.129 | 0.085 | 1 | Perivascular | il13ra1 |
| hip12 | 0.00069 | 1.1390878 | 0.107 | 0.066 | 1 | Perivascular | hip1 |
| romo14 | 0.00071 | 0.2945448 | 0.716 | 0.707 | 1 | Perivascular | romo1 |
| selenom5 | 0.00076 | 1.0034679 | 0.223 | 0.172 | 1 | Perivascular | selenom |
| rcan1a8 | 0.00082 | 0.8234041 | 0.282 | 0.231 | 1 | Perivascular | rcan1a |
| pxna3 | 0.00084 | 1.1953435 | 0.155 | 0.109 | 1 | Perivascular | pxna |
| cct2.14 | 0.00085 | 0.6879661 | 0.444 | 0.424 | 1 | Perivascular | cct2.1 |
| ndufa25 | 0.00086 | 1.1081335 | 0.14 | 0.094 | 1 | Perivascular | ndufa2 |
| capza1a5 | 0.00088 | 0.6501874 | 0.322 | 0.272 | 1 | Perivascular | capza1a |
| usp43 | 0.00091 | 0.9545034 | 0.142 | 0.096 | 1 | Perivascular | usp4 |
| aif1l4 | 0.00094 | 1.1289258 | 0.129 | 0.086 | 1 | Perivascular | aif1l |
| lin7c2 | 0.00094 | 0.8771046 | 0.147 | 0.1 | 1 | Perivascular | lin7c |
| eif3c1 | 0.00095 | 0.528877 | 0.51 | 0.508 | 1 | Perivascular | eif3c |
| pdlim14 | 0.00096 | 0.8671571 | 0.279 | 0.23 | 1 | Perivascular | pdlim1 |
| baz1a5 | 0.00096 | 1.2125339 | 0.16 | 0.114 | 1 | Perivascular | baz1a |
| vapb1 | 0.00101 | 0.7978119 | 0.292 | 0.242 | 1 | Perivascular | vapb |
| pdia76 | 0.00102 | 0.9036055 | 0.195 | 0.145 | 1 | Perivascular | pdia7 |
| smarcc1a2 | 0.00103 | 1.1325622 | 0.114 | 0.073 | 1 | Perivascular | smarcc1a |
| nfkbiaa7 | 0.00111 | 0.5593658 | 0.462 | 0.424 | 1 | Perivascular | nfkbiaa |
| knop13 | 0.00118 | 0.7745173 | 0.218 | 0.163 | 1 | Perivascular | knop1 |
| acadsb3 | 0.00118 | 1.3326164 | 0.102 | 0.064 | 1 | Perivascular | acadsb |
| si:dkey-276j7.15 | 0.00119 | 0.9411135 | 0.216 | 0.168 | 1 | Perivascular | si:dkey-276j7.1 |
| BX322530.12 | 0.00121 | 0.6431682 | 0.439 | 0.42 | 1 | Perivascular | BX322530.1 |
| myofl5 | 0.00125 | 0.7701396 | 0.213 | 0.16 | 1 | Perivascular | myofl |
| capns1a4 | 0.00129 | 0.6831862 | 0.409 | 0.401 | 1 | Perivascular | capns1a |
| zfr3 | 0.00131 | 0.8413173 | 0.16 | 0.113 | 1 | Perivascular | zfr |

| | | | | | | | |
|---|---|---|---|---|---|---|---|
| si:rp71-36a1.54 | 0.00133 | 0.8082441 | 0.132 | 0.087 | 1 | Perivascular | si:rp71-36a1.5 |
| sod12 | 0.00135 | 0.4660734 | 0.492 | 0.466 | 1 | Perivascular | sod1 |
| coro1ca7 | 0.00136 | 0.9212037 | 0.284 | 0.238 | 1 | Perivascular | coro1ca |
| dock9b6 | 0.00139 | 0.6964691 | 0.147 | 0.1 | 1 | Perivascular | dock9b |
| pomp2 | 0.00143 | 0.5774534 | 0.378 | 0.343 | 1 | Perivascular | pomp |
| tmem30aa3 | 0.00144 | 1.0917963 | 0.206 | 0.161 | 1 | Perivascular | tmem30aa |
| rbms1a3 | 0.00147 | 1.0281746 | 0.117 | 0.076 | 1 | Perivascular | rbms1a |
| ptp4a3a5 | 0.00147 | 1.0198286 | 0.168 | 0.12 | 1 | Perivascular | ptp4a3a |
| pcyt24 | 0.00148 | 1.0238516 | 0.14 | 0.096 | 1 | Perivascular | pcyt2 |
| ndufa84 | 0.0015 | 0.7479042 | 0.282 | 0.234 | 1 | Perivascular | ndufa8 |
| FP236812.14 | 0.00154 | 1.3306155 | 0.142 | 0.099 | 1 | Perivascular | FP236812.1 |
| rbm73 | 0.00155 | 1.0509123 | 0.129 | 0.087 | 1 | Perivascular | rbm7 |
| pacsin24 | 0.00158 | 0.9884449 | 0.102 | 0.064 | 1 | Perivascular | pacsin2 |
| taok1a2 | 0.00167 | 1.0489949 | 0.195 | 0.148 | 1 | Perivascular | taok1a |
| fhl2a4 | 0.00168 | 0.7654345 | 0.183 | 0.135 | 1 | Perivascular | fhl2a |
| jun6 | 0.00173 | 0.4490269 | 0.855 | 0.855 | 1 | Perivascular | jun |
| mycbp22 | 0.00176 | 0.8175425 | 0.279 | 0.231 | 1 | Perivascular | mycbp2 |
| vdac15 | 0.00181 | 0.7235216 | 0.358 | 0.324 | 1 | Perivascular | vdac1 |
| atp5meb2 | 0.00182 | 0.4244547 | 0.541 | 0.538 | 1 | Perivascular | atp5meb |
| cox5aa5 | 0.00187 | 0.7947063 | 0.327 | 0.287 | 1 | Perivascular | cox5aa |
| tmem119b8 | 0.00188 | 0.8387315 | 0.251 | 0.207 | 1 | Perivascular | tmem119b |
| hk14 | 0.00193 | 0.9794698 | 0.162 | 0.119 | 1 | Perivascular | hk1 |
| rad21a4 | 0.00196 | 0.7646923 | 0.251 | 0.201 | 1 | Perivascular | rad21a |
| pdap1b6 | 0.00197 | 1.0613652 | 0.137 | 0.096 | 1 | Perivascular | pdap1b |
| denr2 | 0.00221 | 0.8705122 | 0.266 | 0.224 | 1 | Perivascular | denr |
| hnrnpua3 | 0.00234 | 0.7508567 | 0.241 | 0.193 | 1 | Perivascular | hnrnpua |
| adka2 | 0.00236 | 0.7594351 | 0.185 | 0.138 | 1 | Perivascular | adka |
| tars14 | 0.00246 | 0.9559117 | 0.112 | 0.074 | 1 | Perivascular | tars1 |
| actr2a3 | 0.00273 | 0.7772695 | 0.251 | 0.206 | 1 | Perivascular | actr2a |
| ran1 | 0.00295 | 0.2921872 | 0.675 | 0.7 | 1 | Perivascular | ran |
| arid1ab | 0.00296 | 0.7137538 | 0.104 | 0.067 | 1 | Perivascular | arid1ab |
| prdx34 | 0.0031 | 0.7843807 | 0.185 | 0.141 | 1 | Perivascular | prdx3 |
| abhd17b2 | 0.00312 | 1.3066308 | 0.102 | 0.066 | 1 | Perivascular | abhd17b |
| cyc15 | 0.00325 | 0.8150545 | 0.216 | 0.171 | 1 | Perivascular | cyc1 |
| AL929435.14 | 0.00334 | 1.1662617 | 0.147 | 0.107 | 1 | Perivascular | AL929435.1 |
| lmnb22 | 0.00343 | 0.8306397 | 0.142 | 0.1 | 1 | Perivascular | lmnb2 |
| c1qtnf15 | 0.00345 | 1.0933243 | 0.112 | 0.075 | 1 | Perivascular | c1qtnf1 |
| ier2a6 | 0.00356 | 0.5518074 | 0.492 | 0.486 | 1 | Perivascular | ier2a |
| rab5c4 | 0.00364 | 0.6542515 | 0.277 | 0.231 | 1 | Perivascular | rab5c |
| aip2 | 0.00368 | 0.8699376 | 0.241 | 0.201 | 1 | Perivascular | aip |
| eva1ba5 | 0.0037 | 1.0878144 | 0.147 | 0.108 | 1 | Perivascular | eva1ba |
| gnas3 | 0.00374 | 0.8731032 | 0.198 | 0.156 | 1 | Perivascular | gnas |
| ube2v13 | 0.00379 | 0.818506 | 0.302 | 0.267 | 1 | Perivascular | ube2v1 |
| arhgap12 | 0.0038 | 1.2639461 | 0.117 | 0.081 | 1 | Perivascular | arhgap1 |
| midn7 | 0.00394 | 0.5148499 | 0.462 | 0.435 | 1 | Perivascular | midn |
| si:ch211-51e12.71 | 0.00406 | 0.8357307 | 0.183 | 0.139 | 1 | Perivascular | si:ch211-51e12.7 |
| cox7a2a5 | 0.00412 | 0.6964901 | 0.312 | 0.271 | 1 | Perivascular | cox7a2a |
| rnf213b2 | 0.00414 | 0.9539482 | 0.117 | 0.079 | 1 | Perivascular | rnf213b |

| | | | | | | | |
|---|---|---|---|---|---|---|---|
| marveld15 | 0.00416 | 0.8412343 | 0.251 | 0.21 | 1 | Perivascular | marveld1 |
| mdh23 | 0.00428 | 0.5255487 | 0.454 | 0.441 | 1 | Perivascular | mdh2 |
| gng74 | 0.00439 | 0.9377227 | 0.206 | 0.166 | 1 | Perivascular | gng7 |
| srp194 | 0.00459 | 0.9309395 | 0.183 | 0.141 | 1 | Perivascular | srp19 |
| ndrg45 | 0.00463 | 0.7773375 | 0.201 | 0.155 | 1 | Perivascular | ndrg4 |
| abhd2a3 | 0.00467 | 1.09422 | 0.124 | 0.087 | 1 | Perivascular | abhd2a |
| tcf3b3 | 0.0047 | 0.835375 | 0.259 | 0.221 | 1 | Perivascular | tcf3b |
| pdzrn3b4 | 0.0051 | 0.8861738 | 0.117 | 0.079 | 1 | Perivascular | pdzrn3b |
| ergic24 | 0.00517 | 0.803654 | 0.206 | 0.161 | 1 | Perivascular | ergic2 |
| psmd4b3 | 0.00523 | 0.797546 | 0.19 | 0.148 | 1 | Perivascular | psmd4b |
| si:ch211-132g1.34 | 0.00536 | 0.9611473 | 0.142 | 0.104 | 1 | Perivascular | si:ch211-132g1.3 |
| eif5a24 | 0.00536 | 0.426614 | 0.475 | 0.469 | 1 | Perivascular | eif5a2 |
| h3f3b.16 | 0.00557 | 0.4411017 | 0.508 | 0.496 | 1 | Perivascular | h3f3b.1 |
| gyg1a3 | 0.00563 | 0.8370318 | 0.223 | 0.183 | 1 | Perivascular | gyg1a |
| tet35 | 0.00569 | 0.9700897 | 0.137 | 0.099 | 1 | Perivascular | tet3 |
| atf4b8 | 0.00585 | 0.470302 | 0.622 | 0.635 | 1 | Perivascular | atf4b |
| hnrnpc1 | 0.00594 | 0.8949305 | 0.137 | 0.099 | 1 | Perivascular | hnrnpc |
| socs5b2 | 0.00602 | 0.9464937 | 0.129 | 0.092 | 1 | Perivascular | socs5b |
| mt-atp63 | 0.00613 | 0.2199369 | 0.957 | 0.965 | 1 | Perivascular | mt-atp6 |
| atp5f1d5 | 0.00622 | 0.5711852 | 0.365 | 0.335 | 1 | Perivascular | atp5f1d |
| CABZ01044053.13 | 0.00627 | 0.8510678 | 0.117 | 0.082 | 1 | Perivascular | CABZ01044053.1 |
| tubb4b5 | 0.00641 | 0.2894531 | 0.782 | 0.81 | 1 | Perivascular | tubb4b |
| prrg14 | 0.00648 | 0.6817443 | 0.251 | 0.207 | 1 | Perivascular | prrg1 |
| cacybp3 | 0.00671 | 0.8423941 | 0.195 | 0.153 | 1 | Perivascular | cacybp |
| phrf11 | 0.00685 | 0.8671657 | 0.183 | 0.142 | 1 | Perivascular | phrf1 |
| vgll4b3 | 0.00686 | 0.8656704 | 0.203 | 0.164 | 1 | Perivascular | vgll4b |
| esd2 | 0.00713 | 0.9215562 | 0.178 | 0.137 | 1 | Perivascular | esd |
| aco22 | 0.00731 | 0.8237009 | 0.117 | 0.082 | 1 | Perivascular | aco2 |
| cdc42se13 | 0.00741 | 0.8910932 | 0.112 | 0.077 | 1 | Perivascular | cdc42se1 |
| hnrnpabb3 | 0.00754 | 0.4917346 | 0.477 | 0.48 | 1 | Perivascular | hnrnpabb |
| cpda3 | 0.0078 | 0.9096694 | 0.114 | 0.08 | 1 | Perivascular | cpda |
| rnasekb4 | 0.0079 | 1.1136449 | 0.142 | 0.106 | 1 | Perivascular | rnasekb |
| smndc13 | 0.00793 | 0.7827146 | 0.244 | 0.208 | 1 | Perivascular | smndc1 |
| lyar1 | 0.00798 | 1.2710471 | 0.102 | 0.07 | 1 | Perivascular | lyar |
| ndufa53 | 0.00806 | 0.7319018 | 0.246 | 0.21 | 1 | Perivascular | ndufa5 |
| ube2al5 | 0.0081 | 0.7640158 | 0.239 | 0.2 | 1 | Perivascular | ube2al |
| adkb1 | 0.00822 | 1.137289 | 0.104 | 0.072 | 1 | Perivascular | adkb |
| ccdc93 | 0.00826 | 0.7827625 | 0.114 | 0.079 | 1 | Perivascular | ccdc9 |
| vcp3 | 0.00832 | 0.6165252 | 0.345 | 0.318 | 1 | Perivascular | vcp |
| myo1ea5 | 0.00838 | 0.9436853 | 0.117 | 0.082 | 1 | Perivascular | myo1ea |
| dhx162 | 0.00844 | 0.9993816 | 0.145 | 0.109 | 1 | Perivascular | dhx16 |
| dag14 | 0.00851 | 0.9811481 | 0.198 | 0.158 | 1 | Perivascular | dag1 |
| mphosph82 | 0.00851 | 0.7179087 | 0.312 | 0.277 | 1 | Perivascular | mphosph8 |
| C2H1orf354 | 0.00851 | 1.2137096 | 0.119 | 0.086 | 1 | Perivascular | C2H1orf35 |
| cct6a3 | 0.00886 | 0.5004397 | 0.472 | 0.481 | 1 | Perivascular | cct6a |
| pora5 | 0.00927 | 0.7289066 | 0.15 | 0.111 | 1 | Perivascular | pora |
| dnaja2b4 | 0.00959 | 1.0470746 | 0.147 | 0.112 | 1 | Perivascular | dnaja2b |
| mpp11 | 0.00984 | 1.0228946 | 0.109 | 0.077 | 1 | Perivascular | mpp1 |

| | | | | | | | |
|---|---|---|---|---|---|---|---|
| si:zfos-2326c3.23 | 0.00991 | 0.7873662 | 0.203 | 0.163 | 1 | Perivascular | si:zfos-2326c3.2 |
| tmem30ab5 | 0.00995 | 0.908768 | 0.178 | 0.142 | 1 | Perivascular | tmem30ab |
| itsn2b2 | 0.00999 | 0.7059364 | 0.183 | 0.142 | 1 | Perivascular | itsn2b |
| ntn1a7 | 0 | 3.4822801 | 0.785 | 0.116 | 0 | Stroma 5 | ntn1a |
| cfd2 | 0 | 5.8702763 | 0.729 | 0.095 | 0 | Stroma 5 | cfd |
| sostdc1a2 | 0 | 4.8532793 | 0.561 | 0.03 | 0 | Stroma 5 | sostdc1a |
| nr5a2 | 0 | 6.1660589 | 0.517 | 0.011 | 0 | Stroma 5 | nr5a2 |
| c4b3 | 0 | 4.1488154 | 0.531 | 0.051 | 0 | Stroma 5 | c4b |
| aldh1a25 | 0 | 3.6670137 | 0.506 | 0.051 | 0 | Stroma 5 | aldh1a2 |
| hp.1 | 0 | 6.1928376 | 0.461 | 0.017 | 0 | Stroma 5 | hp.1 |
| foxf2a | 0 | 5.9946061 | 0.293 | 0.005 | 0 | Stroma 5 | foxf2a |
| pcdh171 | 0 | 4.9913553 | 0.299 | 0.015 | 0 | Stroma 5 | pcdh17 |
| c3a.1 | 0 | 4.6570593 | 0.201 | 0.007 | 0 | Stroma 5 | c3a.1 |
| aoc1 | 0 | 8.0806553 | 0.19 | 0.002 | 0 | Stroma 5 | aoc1 |
| foxf2b | 0 | 5.9662848 | 0.184 | 0.004 | 0 | Stroma 5 | foxf2b |
| zgc:64106 | 0 | 5.4134984 | 0.176 | 0.005 | 0 | Stroma 5 | zgc:64106 |
| zmp:0000001268 | 0 | 6.4865806 | 0.106 | 0.001 | 0 | Stroma 5 | zmp:0000001268 |
| sema3fa4 | 1E-274 | 3.6162358 | 0.511 | 0.06 | 3.1E-270 | Stroma 5 | sema3fa |
| tmem178 | 7E-226 | 4.8471473 | 0.159 | 0.007 | 2.1E-221 | Stroma 5 | tmem178 |
| gsc5 | 3E-225 | 3.1347503 | 0.637 | 0.113 | 9.5E-221 | Stroma 5 | gsc |
| chrd | 3E-207 | 3.7076276 | 0.293 | 0.026 | 9.7E-203 | Stroma 5 | chrd |
| adcyap1r1a.1 | 1E-203 | 3.8987213 | 0.274 | 0.023 | 3.5E-199 | Stroma 5 | adcyap1r1a.1 |
| bnc26 | 3E-202 | 3.2907009 | 0.508 | 0.079 | 1E-197 | Stroma 5 | bnc2 |
| mmrn2a1 | 1E-199 | 4.0040965 | 0.271 | 0.023 | 4E-195 | Stroma 5 | mmrn2a |
| si:ch211-106h4.126 | 3E-172 | 2.9037576 | 0.986 | 0.455 | 8.3E-168 | Stroma 5 | si:ch211-106h4.12 |
| cxcl144 | 1E-171 | 2.8184632 | 0.788 | 0.245 | 3.7E-167 | Stroma 5 | cxcl14 |
| cxcl8b.11 | 4E-163 | 4.4181441 | 0.307 | 0.036 | 1.2E-158 | Stroma 5 | cxcl8b.1 |
| si:dkey-184p18.22 | 3E-160 | 3.5899232 | 0.383 | 0.056 | 1.1E-155 | Stroma 5 | si:dkey-184p18.2 |
| lhx62 | 5E-160 | 3.0857907 | 0.349 | 0.045 | 1.6E-155 | Stroma 5 | lhx6 |
| lmx1bb | 5E-143 | 4.1208561 | 0.123 | 0.006 | 1.5E-138 | Stroma 5 | lmx1bb |
| cxcl8b.3 | 4E-140 | 4.827567 | 0.115 | 0.006 | 1.4E-135 | Stroma 5 | cxcl8b.3 |
| ccl25b7 | 6E-140 | 1.8033683 | 1 | 0.662 | 1.8E-135 | Stroma 5 | ccl25b |
| rdh10a2 | 9E-137 | 3.6255234 | 0.293 | 0.038 | 2.8E-132 | Stroma 5 | rdh10a |
| asic1b | 7E-135 | 4.0165048 | 0.159 | 0.012 | 2.3E-130 | Stroma 5 | asic1b |
| si:ch211-37e10.21 | 2E-130 | 3.4744997 | 0.229 | 0.024 | 5.6E-126 | Stroma 5 | si:ch211-37e10.2 |
| plpp39 | 2E-126 | 2.1545561 | 0.902 | 0.452 | 7.6E-122 | Stroma 5 | plpp3 |
| slco2b17 | 2E-125 | 2.5926322 | 0.679 | 0.218 | 7.1E-121 | Stroma 5 | slco2b1 |
| cxcl12a9 | 2E-122 | 1.988963 | 0.964 | 0.681 | 6.3E-118 | Stroma 5 | cxcl12a |
| dll43 | 2E-119 | 3.1521842 | 0.288 | 0.041 | 5.3E-115 | Stroma 5 | dll4 |
| sfrp56 | 6E-115 | 1.9115502 | 0.684 | 0.207 | 2E-110 | Stroma 5 | sfrp5 |
| barx17 | 3E-112 | 2.3515168 | 0.514 | 0.126 | 8.7E-108 | Stroma 5 | barx1 |
| dcn7 | 2E-111 | 1.3739838 | 1 | 0.868 | 6.2E-107 | Stroma 5 | dcn |
| ifitm19 | 1E-105 | 1.4243169 | 1 | 0.789 | 4E-101 | Stroma 5 | ifitm1 |
| apoeb4 | 2E-104 | 2.7830678 | 0.684 | 0.245 | 5.6E-100 | Stroma 5 | apoeb |
| fgl15 | 7E-103 | 2.0800429 | 0.651 | 0.207 | 2.31E-98 | Stroma 5 | fgl1 |
| slc40a16 | 6E-102 | 2.3629637 | 0.612 | 0.199 | 1.85E-97 | Stroma 5 | slc40a1 |
| sema3e5 | 2E-101 | 2.5634312 | 0.419 | 0.093 | 5.43E-97 | Stroma 5 | sema3e |
| igfbp6b3 | 2E-100 | 2.9265974 | 0.324 | 0.059 | 5.01E-96 | Stroma 5 | igfbp6b |

| | | | | | | | | |
|---|---|---|---|---|---|---|---|---|
| vasna6 | 1.5E-98 | 2.3992238 | 0.461 | 0.113 | 5.03E-94 | Stroma 5 | | vasna |
| scara36 | 7.3E-91 | 2.0356937 | 0.492 | 0.132 | 2.36E-86 | Stroma 5 | | scara3 |
| neu3.55 | 2E-87 | 2.5752941 | 0.332 | 0.068 | 6.49E-83 | Stroma 5 | | neu3.5 |
| edil3a7 | 2E-87 | 1.9101126 | 0.818 | 0.433 | 6.63E-83 | Stroma 5 | | edil3a |
| rbp48 | 2.5E-87 | 1.268623 | 0.994 | 0.909 | 8.08E-83 | Stroma 5 | | rbp4 |
| zfpm2a | 5.7E-87 | 3.6859571 | 0.112 | 0.009 | 1.84E-82 | Stroma 5 | | zfpm2a |
| znf5361 | 3.8E-86 | 3.0884593 | 0.187 | 0.024 | 1.25E-81 | Stroma 5 | | znf536 |
| wu:fj16a038 | 3.2E-80 | 2.1426713 | 0.564 | 0.19 | 1.05E-75 | Stroma 5 | | wu:fj16a03 |
| zgc:162611 | 1.2E-79 | 3.2642576 | 0.19 | 0.026 | 3.83E-75 | Stroma 5 | | zgc:162611 |
| abi3bpb7 | 2.9E-79 | 1.183438 | 0.953 | 0.583 | 9.38E-75 | Stroma 5 | | abi3bpb |
| admb1 | 2.1E-78 | 3.123252 | 0.232 | 0.038 | 6.97E-74 | Stroma 5 | | admb |
| CR318588.3 | 1.7E-76 | 3.0643387 | 0.316 | 0.07 | 5.46E-72 | Stroma 5 | | CR318588.3 |
| si:dkey-285b23.41 | 3.6E-74 | 3.3019081 | 0.145 | 0.017 | 1.17E-69 | Stroma 5 | | si:dkey-285b23.4 |
| sfrp1b7 | 1.1E-73 | 1.878357 | 0.746 | 0.359 | 3.62E-69 | Stroma 5 | | sfrp1b |
| crabp1b1 | 4.7E-73 | 2.7054241 | 0.204 | 0.032 | 1.53E-68 | Stroma 5 | | crabp1b |
| sfrp1a5 | 2E-72 | 1.6551259 | 0.799 | 0.436 | 6.51E-68 | Stroma 5 | | sfrp1a |
| cdh6 | 1.2E-68 | 3.1544653 | 0.126 | 0.014 | 3.99E-64 | Stroma 5 | | cdh6 |
| rgs5a4 | 7.9E-68 | 0.9201557 | 0.486 | 0.15 | 2.58E-63 | Stroma 5 | | rgs5a |
| si:ch211-207e14.43 | 1.4E-65 | 2.0622541 | 0.299 | 0.068 | 4.47E-61 | Stroma 5 | | si:ch211-207e14.4 |
| adma6 | 1.5E-65 | 1.945492 | 0.592 | 0.232 | 4.74E-61 | Stroma 5 | | adma |
| clu7 | 1.6E-65 | 2.0959506 | 0.749 | 0.381 | 5.15E-61 | Stroma 5 | | clu |
| marcksl1a8 | 2.2E-63 | 1.1922627 | 0.944 | 0.706 | 7.29E-59 | Stroma 5 | | marcksl1a |
| CR318588.14 | 3.3E-60 | 3.0163568 | 0.603 | 0.294 | 1.06E-55 | Stroma 5 | | CR318588.1 |
| cxcl18b8 | 1.2E-59 | 1.344187 | 0.939 | 0.701 | 4.06E-55 | Stroma 5 | | cxcl18b |
| dkk22 | 2.8E-59 | 2.0399561 | 0.251 | 0.054 | 9.1E-55 | Stroma 5 | | dkk2 |
| fybb | 2.1E-58 | 3.1669736 | 0.128 | 0.017 | 6.76E-54 | Stroma 5 | | fybb |
| nr0b1 | 2.9E-57 | 3.375396 | 0.103 | 0.011 | 9.58E-53 | Stroma 5 | | nr0b1 |
| ca66 | 6.9E-57 | 1.7100307 | 0.715 | 0.345 | 2.25E-52 | Stroma 5 | | ca6 |
| osr17 | 2.5E-56 | 1.7911227 | 0.478 | 0.171 | 8.18E-52 | Stroma 5 | | osr1 |
| sult2st3 | 1.6E-55 | 3.3948381 | 0.131 | 0.018 | 5.14E-51 | Stroma 5 | | sult2st3 |
| apcdd1l7 | 3.3E-54 | 1.7528501 | 0.366 | 0.109 | 1.08E-49 | Stroma 5 | | apcdd1l |
| igf2a4 | 8.3E-54 | 2.3798391 | 0.321 | 0.091 | 2.71E-49 | Stroma 5 | | igf2a |
| dpt4 | 8.3E-53 | 1.8834816 | 0.436 | 0.153 | 2.69E-48 | Stroma 5 | | dpt |
| crhbp3 | 6.3E-52 | 2.6175937 | 0.196 | 0.039 | 2.05E-47 | Stroma 5 | | crhbp |
| pdgfrl7 | 7.2E-52 | 1.17419 | 0.86 | 0.583 | 2.33E-47 | Stroma 5 | | pdgfrl |
| socs3a5 | 2.3E-51 | 1.3145357 | 0.958 | 0.794 | 7.5E-47 | Stroma 5 | | socs3a |
| ebf3a.13 | 7.6E-51 | 2.226141 | 0.218 | 0.047 | 2.47E-46 | Stroma 5 | | ebf3a.1 |
| c6.16 | 2.6E-50 | 1.4834518 | 0.508 | 0.204 | 8.52E-46 | Stroma 5 | | c6.1 |
| CABZ01085139.13 | 8.3E-50 | 2.2523293 | 0.257 | 0.063 | 2.69E-45 | Stroma 5 | | CABZ01085139.1 |
| CR318588.44 | 4.7E-49 | 1.0453734 | 0.997 | 0.986 | 1.53E-44 | Stroma 5 | | CR318588.4 |
| pmp22a5 | 4.8E-49 | 1.1183442 | 0.888 | 0.688 | 1.56E-44 | Stroma 5 | | pmp22a |
| adh8a5 | 6.2E-49 | 1.4085037 | 0.514 | 0.21 | 2.02E-44 | Stroma 5 | | adh8a |
| CABZ01092746.18 | 5.7E-47 | 0.9298446 | 0.925 | 0.708 | 1.86E-42 | Stroma 5 | | CABZ01092746.1 |
| mdkb2 | 8.6E-46 | 2.2163902 | 0.193 | 0.041 | 2.78E-41 | Stroma 5 | | mdkb |
| twist1b6 | 1.1E-45 | 1.6534677 | 0.425 | 0.158 | 3.64E-41 | Stroma 5 | | twist1b |
| kera6 | 3.2E-45 | 1.6023506 | 0.444 | 0.17 | 1.04E-40 | Stroma 5 | | kera |
| rsrp16 | 7.8E-45 | 1.1493658 | 0.807 | 0.541 | 2.54E-40 | Stroma 5 | | rsrp1 |
| serpinf17 | 1.4E-44 | 1.0199934 | 0.821 | 0.553 | 4.71E-40 | Stroma 5 | | serpinf1 |

| | | | | | | | |
|---|---|---|---|---|---|---|---|
| phyhd11 | 2.5E-43 | 2.1580195 | 0.201 | 0.046 | 8.15E-39 | Stroma 5 | phyhd1 |
| cyp1d1 | 3.5E-42 | 2.7222716 | 0.142 | 0.026 | 1.13E-37 | Stroma 5 | cyp1d1 |
| id19 | 3.6E-42 | 1.2044053 | 0.869 | 0.64 | 1.16E-37 | Stroma 5 | id1 |
| zgc:92429 | 4E-42 | 2.7686098 | 0.112 | 0.017 | 1.31E-37 | Stroma 5 | zgc:92429 |
| fgl2a7 | 7.3E-42 | 1.6440388 | 0.413 | 0.157 | 2.38E-37 | Stroma 5 | fgl2a |
| hsd11b25 | 5.1E-41 | 1.7716721 | 0.299 | 0.093 | 1.67E-36 | Stroma 5 | hsd11b2 |
| si:dkey-192g7.33 | 6E-41 | 2.3604208 | 0.218 | 0.055 | 1.95E-36 | Stroma 5 | si:dkey-192g7.3 |
| cd282 | 6.1E-41 | 2.4832156 | 0.123 | 0.02 | 1.98E-36 | Stroma 5 | cd28 |
| fstl1a6 | 1E-40 | 1.452686 | 0.33 | 0.106 | 3.36E-36 | Stroma 5 | fstl1a |
| klf34 | 4.8E-40 | 1.8724851 | 0.307 | 0.1 | 1.56E-35 | Stroma 5 | klf3 |
| cxcl8a5 | 1.1E-39 | 1.7460256 | 0.553 | 0.264 | 3.61E-35 | Stroma 5 | cxcl8a |
| id2a9 | 1.8E-39 | 1.0795277 | 0.765 | 0.502 | 5.95E-35 | Stroma 5 | id2a |
| si:ch211-227m13.13 | 1.4E-38 | 1.8987403 | 0.209 | 0.053 | 4.45E-34 | Stroma 5 | si:ch211-227m13.1 |
| nrg2a | 1.5E-38 | 2.8313228 | 0.103 | 0.016 | 4.99E-34 | Stroma 5 | nrg2a |
| akap12b5 | 4.6E-38 | 1.4820096 | 0.38 | 0.142 | 1.48E-33 | Stroma 5 | akap12b |
| snx175 | 1.7E-37 | 1.5461414 | 0.503 | 0.241 | 5.44E-33 | Stroma 5 | snx17 |
| gsta.17 | 2E-37 | 1.2890357 | 0.634 | 0.356 | 6.37E-33 | Stroma 5 | gsta.1 |
| hsd17b31 | 2.3E-37 | 2.8999282 | 0.103 | 0.016 | 7.42E-33 | Stroma 5 | hsd17b3 |
| vmo1a6 | 5.4E-37 | 0.7707599 | 0.905 | 0.7 | 1.76E-32 | Stroma 5 | vmo1a |
| nfia7 | 1.1E-36 | 1.5024517 | 0.411 | 0.169 | 3.42E-32 | Stroma 5 | nfia |
| wu:fc23c091 | 1.1E-36 | 2.5572428 | 0.12 | 0.021 | 3.62E-32 | Stroma 5 | wu:fc23c09 |
| c7a7 | 1.2E-36 | 1.4681211 | 0.425 | 0.173 | 3.8E-32 | Stroma 5 | c7a |
| sema3d3 | 1.4E-36 | 2.2916802 | 0.182 | 0.044 | 4.43E-32 | Stroma 5 | sema3d |
| dnajb1b7 | 2.7E-36 | 1.3984866 | 0.715 | 0.463 | 8.94E-32 | Stroma 5 | dnajb1b |
| alx4b3 | 4.2E-36 | 2.0842576 | 0.212 | 0.057 | 1.37E-31 | Stroma 5 | alx4b |
| CR925728.14 | 5E-36 | 2.045567 | 0.282 | 0.093 | 1.64E-31 | Stroma 5 | CR925728.1 |
| sema3aa6 | 7.2E-35 | 1.5643743 | 0.31 | 0.108 | 2.34E-30 | Stroma 5 | sema3aa |
| si:ch211-212k18.73 | 1.4E-34 | 2.0314974 | 0.221 | 0.063 | 4.47E-30 | Stroma 5 | si:ch211-212k18.7 |
| stim1a2 | 1.5E-34 | 2.0820543 | 0.209 | 0.058 | 5.02E-30 | Stroma 5 | stim1a |
| rgs5b9 | 2.2E-34 | 0.8521832 | 0.818 | 0.522 | 7.3E-30 | Stroma 5 | rgs5b |
| zgc:1951736 | 6.1E-34 | 1.0978573 | 0.665 | 0.396 | 1.98E-29 | Stroma 5 | zgc:195173 |
| pcdh193 | 1.8E-33 | 2.0351845 | 0.196 | 0.052 | 5.97E-29 | Stroma 5 | pcdh19 |
| arhgef47 | 3.7E-33 | 2.1007488 | 0.24 | 0.075 | 1.21E-28 | Stroma 5 | arhgef4 |
| eda | 3.9E-33 | 2.3453635 | 0.103 | 0.018 | 1.26E-28 | Stroma 5 | eda |
| efna1b4 | 4.9E-33 | 1.9076342 | 0.349 | 0.142 | 1.58E-28 | Stroma 5 | efna1b |
| cebpb7 | 5.2E-33 | 1.6211443 | 0.684 | 0.448 | 1.71E-28 | Stroma 5 | cebpb |
| efna2a2 | 5.4E-33 | 2.1050507 | 0.176 | 0.044 | 1.75E-28 | Stroma 5 | efna2a |
| tent5c4 | 7.9E-33 | 1.9230363 | 0.254 | 0.083 | 2.57E-28 | Stroma 5 | tent5c |
| ntd57 | 7.9E-33 | 1.5111741 | 0.36 | 0.141 | 2.58E-28 | Stroma 5 | ntd5 |
| b2ml10 | 2.6E-32 | 0.670212 | 0.964 | 0.897 | 8.56E-28 | Stroma 5 | b2ml |
| antxr2a6 | 3.7E-32 | 1.3373534 | 0.587 | 0.334 | 1.19E-27 | Stroma 5 | antxr2a |
| tns2b2 | 4.3E-32 | 1.8979621 | 0.156 | 0.037 | 1.41E-27 | Stroma 5 | tns2b |
| slc43a3b4 | 1.2E-31 | 1.6615278 | 0.246 | 0.078 | 4.02E-27 | Stroma 5 | slc43a3b |
| il15ra | 1.5E-31 | 2.4265679 | 0.123 | 0.025 | 4.87E-27 | Stroma 5 | il15ra |
| abi3bpa6 | 1.7E-31 | 0.8476684 | 0.584 | 0.298 | 5.68E-27 | Stroma 5 | abi3bpa |
| ponzr16 | 2.9E-31 | 1.1533928 | 0.489 | 0.232 | 9.32E-27 | Stroma 5 | ponzr1 |
| junba6 | 6.9E-31 | 0.8968216 | 0.955 | 0.845 | 2.25E-26 | Stroma 5 | junba |
| CABZ01079281.11 | 9E-31 | 2.1090499 | 0.162 | 0.04 | 2.93E-26 | Stroma 5 | CABZ01079281.1 |

| | | | | | | | |
|---|---|---|---|---|---|---|---|
| nt5e5 | 1.4E-30 | 1.8861808 | 0.212 | 0.063 | 4.5E-26 | Stroma 5 | nt5e |
| fgf10a | 4.6E-30 | 2.5782519 | 0.117 | 0.024 | 1.49E-25 | Stroma 5 | fgf10a |
| tmem17610 | 1E-29 | 1.0141362 | 0.642 | 0.393 | 3.32E-25 | Stroma 5 | tmem176 |
| si:dkey-108k21.21 | 1.1E-29 | 2.5620956 | 0.101 | 0.018 | 3.7E-25 | Stroma 5 | si:dkey-108k21.21 |
| foxd13 | 4.4E-29 | 1.5104962 | 0.218 | 0.067 | 1.43E-24 | Stroma 5 | foxd1 |
| jun7 | 7E-29 | 0.9068377 | 0.972 | 0.854 | 2.26E-24 | Stroma 5 | jun |
| pim26 | 7.2E-29 | 1.3606765 | 0.455 | 0.225 | 2.33E-24 | Stroma 5 | pim2 |
| fosl1a5 | 1.5E-28 | 1.3884747 | 0.648 | 0.437 | 4.85E-24 | Stroma 5 | fosl1a |
| slc25a25a7 | 2.1E-28 | 1.4327237 | 0.436 | 0.218 | 6.67E-24 | Stroma 5 | slc25a25a |
| si:ch211-198c19.36 | 4.6E-28 | 0.9788889 | 0.455 | 0.21 | 1.51E-23 | Stroma 5 | si:ch211-198c19.3 |
| si:dkey-282h22.56 | 9.3E-28 | 1.0693551 | 0.221 | 0.07 | 3.02E-23 | Stroma 5 | si:dkey-282h22.5 |
| im:71523489 | 9.7E-28 | 1.8323593 | 0.455 | 0.235 | 3.15E-23 | Stroma 5 | im:7152348 |
| cwc275 | 2.3E-27 | 1.6892347 | 0.304 | 0.122 | 7.57E-23 | Stroma 5 | cwc27 |
| tnfsf10l8 | 2.5E-27 | 1.2708176 | 0.444 | 0.221 | 8.17E-23 | Stroma 5 | tnfsf10l |
| serpinb11 | 5.1E-27 | 1.9050342 | 0.207 | 0.066 | 1.67E-22 | Stroma 5 | serpinb1 |
| si:ch1073-291c23.21 | 7.1E-27 | 0.8859683 | 0.704 | 0.474 | 2.3E-22 | Stroma 5 | si:ch1073-291c23.2 |
| mknk2b6 | 7.3E-27 | 0.8004125 | 0.81 | 0.607 | 2.38E-22 | Stroma 5 | mknk2b |
| tenm4 | 7.4E-27 | 1.9107758 | 0.168 | 0.047 | 2.42E-22 | Stroma 5 | tenm4 |
| wnt116 | 9.2E-27 | 0.8434188 | 0.536 | 0.283 | 3E-22 | Stroma 5 | wnt11 |
| enc16 | 2.1E-26 | 1.4619591 | 0.307 | 0.123 | 6.74E-22 | Stroma 5 | enc1 |
| ccdc80l24 | 4.6E-26 | 2.7861797 | 0.285 | 0.114 | 1.51E-21 | Stroma 5 | ccdc80l2 |
| fzd24 | 2.8E-25 | 1.7390453 | 0.257 | 0.098 | 9.01E-21 | Stroma 5 | fzd2 |
| hhla2a.15 | 3E-25 | 1.7504027 | 0.229 | 0.081 | 9.67E-21 | Stroma 5 | hhla2a.1 |
| loxl18 | 7.2E-25 | 1.1025259 | 0.416 | 0.2 | 2.33E-20 | Stroma 5 | loxl1 |
| si:ch73-380l3.27 | 8E-25 | 0.9602153 | 0.64 | 0.415 | 2.61E-20 | Stroma 5 | si:ch73-380l3.2 |
| zgc:1140467 | 1.5E-24 | 1.5035829 | 0.26 | 0.099 | 4.73E-20 | Stroma 5 | zgc:114046 |
| slit34 | 2.3E-24 | 1.3759367 | 0.399 | 0.201 | 7.35E-20 | Stroma 5 | slit3 |
| btg26 | 2.8E-24 | 0.9273537 | 0.849 | 0.699 | 9.26E-20 | Stroma 5 | btg2 |
| id36 | 4.6E-24 | 0.9929627 | 0.718 | 0.511 | 1.5E-19 | Stroma 5 | id3 |
| smoc16 | 5E-24 | 1.4048668 | 0.408 | 0.207 | 1.64E-19 | Stroma 5 | smoc1 |
| hapln1a5 | 1.3E-23 | 0.2481898 | 0.768 | 0.477 | 4.34E-19 | Stroma 5 | hapln1a |
| cd74b | 2.5E-23 | 2.6165584 | 0.142 | 0.04 | 8.29E-19 | Stroma 5 | cd74b |
| si:ch211-250c4.43 | 2.6E-23 | 1.76024 | 0.154 | 0.044 | 8.6E-19 | Stroma 5 | si:ch211-250c4.4 |
| junbb6 | 3.3E-23 | 1.0254933 | 0.768 | 0.601 | 1.09E-18 | Stroma 5 | junbb |
| eif4ebp35 | 3.8E-23 | 1.2398351 | 0.363 | 0.173 | 1.25E-18 | Stroma 5 | eif4ebp3 |
| mcl1a7 | 4.1E-23 | 0.7611621 | 0.894 | 0.77 | 1.35E-18 | Stroma 5 | mcl1a |
| twist29 | 4.3E-23 | 0.870569 | 0.587 | 0.34 | 1.4E-18 | Stroma 5 | twist2 |
| c47 | 5.9E-23 | 0.9550037 | 0.427 | 0.207 | 1.93E-18 | Stroma 5 | c4 |
| add3a8 | 6.8E-23 | 0.9018446 | 0.665 | 0.447 | 2.22E-18 | Stroma 5 | add3a |
| tril9 | 1.1E-22 | 1.3723087 | 0.265 | 0.106 | 3.54E-18 | Stroma 5 | tril |
| vcam1b1 | 1.1E-22 | 1.8990369 | 0.156 | 0.046 | 3.61E-18 | Stroma 5 | vcam1b |
| fstl37 | 1.1E-22 | 1.2721739 | 0.52 | 0.321 | 3.69E-18 | Stroma 5 | fstl3 |
| tob1b7 | 1.2E-22 | 0.7593106 | 0.821 | 0.702 | 3.84E-18 | Stroma 5 | tob1b |
| zgc:1122345 | 1.3E-22 | 2.0190713 | 0.207 | 0.074 | 4.2E-18 | Stroma 5 | zgc:112234 |
| si:dkey-11f4.207 | 1.5E-22 | 0.778818 | 0.704 | 0.488 | 4.81E-18 | Stroma 5 | si:dkey-11f4.20 |
| mhc1zba8 | 2E-22 | 0.8043531 | 0.749 | 0.583 | 6.59E-18 | Stroma 5 | mhc1zba |
| tnfsf124 | 3.5E-22 | 1.459692 | 0.318 | 0.146 | 1.13E-17 | Stroma 5 | tnfsf12 |
| bhlhe406 | 4E-22 | 1.0218326 | 0.665 | 0.464 | 1.3E-17 | Stroma 5 | bhlhe40 |

| | | | | | | | |
|---|---|---|---|---|---|---|---|
| fosb7 | 7.2E-22 | 0.8799689 | 0.818 | 0.692 | 2.36E-17 | Stroma 5 | fosb |
| egr16 | 1.3E-21 | 0.943801 | 0.793 | 0.628 | 4.33E-17 | Stroma 5 | egr1 |
| dpysl36 | 2.4E-21 | 1.0677539 | 0.346 | 0.161 | 7.86E-17 | Stroma 5 | dpysl3 |
| zbtb16a5 | 5.1E-21 | 1.6568538 | 0.291 | 0.13 | 1.67E-16 | Stroma 5 | zbtb16a |
| tmsb15 | 1.1E-20 | 0.6474302 | 0.682 | 0.449 | 3.5E-16 | Stroma 5 | tmsb1 |
| anxa1a5 | 1.8E-20 | 0.4117994 | 0.983 | 0.904 | 5.71E-16 | Stroma 5 | anxa1a |
| tspan144 | 2.4E-20 | 1.5487841 | 0.274 | 0.121 | 7.64E-16 | Stroma 5 | tspan14 |
| nfil3-68 | 2.5E-20 | 1.0003829 | 0.514 | 0.305 | 8.12E-16 | Stroma 5 | nfil3-6 |
| fosab7 | 3.1E-20 | 0.6265711 | 0.983 | 0.895 | 1.02E-15 | Stroma 5 | fosab |
| zgc:1749044 | 3.3E-20 | 1.1249536 | 0.285 | 0.124 | 1.07E-15 | Stroma 5 | zgc:174904 |
| pfkfb37 | 3.9E-20 | 2.0127703 | 0.201 | 0.076 | 1.27E-15 | Stroma 5 | pfkfb3 |
| aldh9a1a.17 | 4.1E-20 | 0.7961245 | 0.665 | 0.481 | 1.33E-15 | Stroma 5 | aldh9a1a.1 |
| slc20a1b5 | 5.2E-20 | 1.1921866 | 0.486 | 0.294 | 1.69E-15 | Stroma 5 | slc20a1b |
| CU459056.15 | 9.4E-20 | 1.2972533 | 0.341 | 0.169 | 3.05E-15 | Stroma 5 | CU459056.1 |
| krt945 | 4.1E-19 | 0.747655 | 0.508 | 0.295 | 1.32E-14 | Stroma 5 | krt94 |
| lxn7 | 4.2E-19 | 0.4733263 | 0.891 | 0.701 | 1.38E-14 | Stroma 5 | lxn |
| col4a16 | 5.1E-19 | 0.3718884 | 0.43 | 0.226 | 1.67E-14 | Stroma 5 | col4a1 |
| ednraa8 | 8.3E-19 | 1.0389908 | 0.399 | 0.213 | 2.7E-14 | Stroma 5 | ednraa |
| mycb5 | 1E-18 | 0.8895604 | 0.791 | 0.663 | 3.3E-14 | Stroma 5 | mycb |
| arl4ab7 | 1.2E-18 | 1.2861639 | 0.254 | 0.11 | 3.82E-14 | Stroma 5 | arl4ab |
| adamts15a8 | 2.2E-18 | 1.2941524 | 0.304 | 0.143 | 7.19E-14 | Stroma 5 | adamts15a |
| rnd3a8 | 2.4E-18 | 0.9355445 | 0.478 | 0.285 | 7.93E-14 | Stroma 5 | rnd3a |
| btg16 | 2.5E-18 | 0.5950948 | 0.877 | 0.815 | 8.12E-14 | Stroma 5 | btg1 |
| BX663503.37 | 3.5E-18 | 0.6114809 | 0.64 | 0.402 | 1.12E-13 | Stroma 5 | BX663503.3 |
| si:ch211-212k18.85 | 4.1E-18 | 1.2650667 | 0.296 | 0.141 | 1.34E-13 | Stroma 5 | si:ch211-212k18.8 |
| opcml1 | 6.6E-18 | 1.5761766 | 0.173 | 0.062 | 2.14E-13 | Stroma 5 | opcml |
| six1a6 | 6.8E-18 | 1.1347523 | 0.282 | 0.13 | 2.21E-13 | Stroma 5 | six1a |
| tdh6 | 8.7E-18 | 1.1569871 | 0.416 | 0.24 | 2.82E-13 | Stroma 5 | tdh |
| zgc:1949896 | 1.1E-17 | 1.4921359 | 0.218 | 0.09 | 3.47E-13 | Stroma 5 | zgc:194989 |
| si:ch73-86n18.17 | 1.6E-17 | 1.1997023 | 0.316 | 0.153 | 5.05E-13 | Stroma 5 | si:ch73-86n18.1 |
| soul55 | 3.8E-17 | 0.7707999 | 0.469 | 0.276 | 1.24E-12 | Stroma 5 | soul5 |
| hmox1a7 | 5E-17 | 1.1147046 | 0.338 | 0.173 | 1.62E-12 | Stroma 5 | hmox1a |
| rflnb4 | 7.9E-17 | 1.5963212 | 0.246 | 0.111 | 2.56E-12 | Stroma 5 | rflnb |
| moxd14 | 8E-17 | 1.5547865 | 0.151 | 0.052 | 2.6E-12 | Stroma 5 | moxd1 |
| foxq1a | 1.1E-16 | 2.1112254 | 0.106 | 0.03 | 3.54E-12 | Stroma 5 | foxq1a |
| gpc43 | 1.3E-16 | 1.1154044 | 0.416 | 0.242 | 4.1E-12 | Stroma 5 | gpc4 |
| tob1a6 | 1.6E-16 | 0.6640232 | 0.76 | 0.609 | 5.26E-12 | Stroma 5 | tob1a |
| zfp36l25 | 1.6E-16 | 0.9764963 | 0.444 | 0.268 | 5.32E-12 | Stroma 5 | zfp36l2 |
| ctss2.1 | 2.7E-16 | 2.0891425 | 0.103 | 0.03 | 8.83E-12 | Stroma 5 | ctss2.1 |
| podn7 | 4.6E-16 | 1.0422533 | 0.332 | 0.172 | 1.5E-11 | Stroma 5 | podn |
| rbpjb3 | 1.4E-15 | 1.7011147 | 0.218 | 0.097 | 4.54E-11 | Stroma 5 | rbpjb |
| endouc4 | 1.5E-15 | 1.2132167 | 0.391 | 0.236 | 4.73E-11 | Stroma 5 | endouc |
| mmel11 | 1.9E-15 | 0.4557197 | 0.162 | 0.059 | 6.04E-11 | Stroma 5 | mmel1 |
| irf7 | 1.9E-15 | 1.7447576 | 0.134 | 0.046 | 6.24E-11 | Stroma 5 | irf7 |
| lpl6 | 2E-15 | 1.3153414 | 0.243 | 0.112 | 6.38E-11 | Stroma 5 | lpl |
| spon1b2 | 2.4E-15 | 1.4499279 | 0.154 | 0.056 | 7.9E-11 | Stroma 5 | spon1b |
| ccbe11 | 2.4E-15 | 1.5549898 | 0.14 | 0.049 | 7.96E-11 | Stroma 5 | ccbe1 |
| si:dkey-195m11.87 | 2.8E-15 | 1.3828064 | 0.263 | 0.127 | 8.97E-11 | Stroma 5 | si:dkey-195m11.8 |

| | | | | | | | |
|---|---|---|---|---|---|---|---|
| ubb7 | 2.8E-15 | 0.5588576 | 0.804 | 0.681 | 9.02E-11 | Stroma 5 | ubb |
| sult1st66 | 3.1E-15 | 1.2091311 | 0.249 | 0.116 | 1.01E-10 | Stroma 5 | sult1st6 |
| osr21 | 3.4E-15 | 1.8224958 | 0.103 | 0.031 | 1.11E-10 | Stroma 5 | osr2 |
| ier2b6 | 4.4E-15 | 0.7819231 | 0.779 | 0.649 | 1.44E-10 | Stroma 5 | ier2b |
| txnipa3 | 5.7E-15 | 0.9585314 | 0.413 | 0.244 | 1.85E-10 | Stroma 5 | txnipa |
| il6st6 | 1.8E-14 | 0.8742215 | 0.584 | 0.423 | 5.86E-10 | Stroma 5 | il6st |
| sox11a5 | 2.2E-14 | 1.0149456 | 0.511 | 0.343 | 7.02E-10 | Stroma 5 | sox11a |
| ddit37 | 2.3E-14 | 0.7380696 | 0.718 | 0.582 | 7.37E-10 | Stroma 5 | ddit3 |
| zeb1b6 | 3.3E-14 | 1.2138748 | 0.201 | 0.088 | 1.07E-09 | Stroma 5 | zeb1b |
| ITLN17 | 4.4E-14 | 0.7724791 | 0.374 | 0.21 | 1.45E-09 | Stroma 5 | ITLN1 |
| hist2h2l8 | 7.2E-14 | 0.7798037 | 0.539 | 0.374 | 2.36E-09 | Stroma 5 | hist2h2l |
| sat1a.212 | 7.4E-14 | 0.9014124 | 0.617 | 0.457 | 2.41E-09 | Stroma 5 | sat1a.2 |
| klf2a5 | 1E-13 | 0.7968455 | 0.461 | 0.285 | 3.4E-09 | Stroma 5 | klf2a |
| cgnl15 | 1.3E-13 | 1.2968169 | 0.173 | 0.072 | 4.08E-09 | Stroma 5 | cgnl1 |
| si:dkey-261h17.16 | 1.6E-13 | 0.4575922 | 0.478 | 0.29 | 5.2E-09 | Stroma 5 | si:dkey-261h17.1 |
| nabp1a6 | 1.8E-13 | 1.1745192 | 0.316 | 0.179 | 6.01E-09 | Stroma 5 | nabp1a |
| chac16 | 1.9E-13 | 0.9069592 | 0.5 | 0.354 | 6.18E-09 | Stroma 5 | chac1 |
| six2a6 | 2.3E-13 | 1.1787343 | 0.218 | 0.101 | 7.55E-09 | Stroma 5 | six2a |
| cebpd8 | 2.7E-13 | 0.5456148 | 0.737 | 0.577 | 8.7E-09 | Stroma 5 | cebpd |
| CU914487.13 | 2.9E-13 | 1.1247082 | 0.151 | 0.059 | 9.34E-09 | Stroma 5 | CU914487.1 |
| slc10a3 | 3E-13 | 1.7703835 | 0.126 | 0.045 | 9.72E-09 | Stroma 5 | slc10a3 |
| adam10a6 | 3.6E-13 | 0.7468035 | 0.536 | 0.381 | 1.18E-08 | Stroma 5 | adam10a |
| CU984600.16 | 3.7E-13 | 0.98925 | 0.313 | 0.17 | 1.2E-08 | Stroma 5 | CU984600.1 |
| klf12a6 | 5.5E-13 | 1.0862481 | 0.268 | 0.138 | 1.78E-08 | Stroma 5 | klf12a |
| ENSDARG00000908 | 8.1E-13 | 1.1810909 | 0.198 | 0.09 | 2.64E-08 | Stroma 5 | ENSDARG00000090 |
| pdgfra7 | 9.9E-13 | 0.5957878 | 0.631 | 0.46 | 3.24E-08 | Stroma 5 | pdgfra |
| cyp2aa72 | 1.1E-12 | 1.6448302 | 0.142 | 0.056 | 3.54E-08 | Stroma 5 | cyp2aa7 |
| tspan357 | 1.3E-12 | 1.1545598 | 0.243 | 0.123 | 4.38E-08 | Stroma 5 | tspan35 |
| b2m9 | 1.4E-12 | 0.6564882 | 0.779 | 0.686 | 4.47E-08 | Stroma 5 | b2m |
| dio16 | 1.6E-12 | 1.0346164 | 0.246 | 0.122 | 5.24E-08 | Stroma 5 | dio1 |
| ANPEP6 | 3.6E-12 | 1.0271558 | 0.346 | 0.21 | 1.17E-07 | Stroma 5 | ANPEP |
| tie11 | 3.9E-12 | 1.5702188 | 0.101 | 0.034 | 1.26E-07 | Stroma 5 | tie1 |
| col5a3a5 | 4E-12 | 0.692965 | 0.162 | 0.067 | 1.29E-07 | Stroma 5 | col5a3a |
| pappaa | 7.9E-12 | 1.7815214 | 0.106 | 0.037 | 2.58E-07 | Stroma 5 | pappaa |
| angptl49 | 1.2E-11 | 0.5711281 | 0.612 | 0.456 | 3.95E-07 | Stroma 5 | angptl4 |
| colec126 | 1.3E-11 | 0.5714447 | 0.598 | 0.438 | 4.21E-07 | Stroma 5 | colec12 |
| si:ch211-286o17.17 | 1.7E-11 | 1.0640896 | 0.349 | 0.216 | 5.42E-07 | Stroma 5 | si:ch211-286o17.1 |
| zgc:1735876 | 1.9E-11 | 1.0010613 | 0.249 | 0.131 | 6.25E-07 | Stroma 5 | zgc:173587 |
| gfra1a4 | 2.2E-11 | 1.4042018 | 0.137 | 0.055 | 7.15E-07 | Stroma 5 | gfra1a |
| gamt2 | 3E-11 | 1.3858008 | 0.162 | 0.072 | 9.63E-07 | Stroma 5 | gamt |
| igf2b6 | 3E-11 | 0.73757 | 0.595 | 0.436 | 9.71E-07 | Stroma 5 | igf2b |
| mfap59 | 3.6E-11 | 0.4275505 | 0.696 | 0.546 | 1.17E-06 | Stroma 5 | mfap5 |
| ecm1b7 | 3.7E-11 | 0.447511 | 0.642 | 0.471 | 1.21E-06 | Stroma 5 | ecm1b |
| aif1l5 | 4E-11 | 1.0910594 | 0.184 | 0.086 | 1.32E-06 | Stroma 5 | aif1l |
| dedd17 | 4.4E-11 | 0.5502277 | 0.687 | 0.552 | 1.43E-06 | Stroma 5 | dedd1 |
| napsa6 | 4.8E-11 | 1.1341408 | 0.293 | 0.169 | 1.55E-06 | Stroma 5 | napsa |
| lama46 | 5.5E-11 | 0.796248 | 0.411 | 0.27 | 1.79E-06 | Stroma 5 | lama4 |
| cilp24 | 6.1E-11 | 0.3829958 | 0.148 | 0.062 | 1.98E-06 | Stroma 5 | cilp2 |

| | | | | | | | |
|---|---|---|---|---|---|---|---|
| twist1a6 | 7.4E-11 | 0.6010866 | 0.595 | 0.438 | 2.4E-06 | Stroma 5 | twist1a |
| il11ra6 | 7.4E-11 | 0.8010263 | 0.38 | 0.24 | 2.41E-06 | Stroma 5 | il11ra |
| ccn2b4 | 8.5E-11 | 0.511517 | 0.344 | 0.206 | 2.78E-06 | Stroma 5 | ccn2b |
| rnd3b7 | 9.5E-11 | 0.6999761 | 0.478 | 0.331 | 3.08E-06 | Stroma 5 | rnd3b |
| clec3ba5 | 1.1E-10 | 0.6796324 | 0.243 | 0.126 | 3.71E-06 | Stroma 5 | clec3ba |
| jdp2b6 | 1.2E-10 | 0.7819586 | 0.707 | 0.571 | 3.92E-06 | Stroma 5 | jdp2b |
| ccdc1255 | 1.2E-10 | 1.2361983 | 0.165 | 0.075 | 4.01E-06 | Stroma 5 | ccdc125 |
| hdac46 | 1.4E-10 | 1.0593887 | 0.229 | 0.12 | 4.57E-06 | Stroma 5 | hdac4 |
| igfbp5b7 | 1.7E-10 | 0.4082641 | 0.712 | 0.571 | 5.61E-06 | Stroma 5 | igfbp5b |
| atf5b5 | 1.7E-10 | 0.7445461 | 0.394 | 0.252 | 5.68E-06 | Stroma 5 | atf5b |
| ablim23 | 1.9E-10 | 1.2834565 | 0.12 | 0.048 | 6.05E-06 | Stroma 5 | ablim2 |
| polr2a7 | 2E-10 | 0.8299741 | 0.494 | 0.363 | 6.48E-06 | Stroma 5 | polr2a |
| gpx4a6 | 2.3E-10 | 0.710584 | 0.447 | 0.303 | 7.41E-06 | Stroma 5 | gpx4a |
| anxa3b4 | 2.4E-10 | 0.8298105 | 0.383 | 0.248 | 7.78E-06 | Stroma 5 | anxa3b |
| ppap2d6 | 2.5E-10 | 0.6451382 | 0.372 | 0.229 | 8.09E-06 | Stroma 5 | ppap2d |
| cited4b6 | 2.6E-10 | 0.9599232 | 0.265 | 0.15 | 8.52E-06 | Stroma 5 | cited4b |
| si:dkey-85k7.77 | 2.7E-10 | 0.6871336 | 0.427 | 0.28 | 8.86E-06 | Stroma 5 | si:dkey-85k7.7 |
| ehbp1l1a7 | 2.9E-10 | 1.224311 | 0.19 | 0.094 | 9.38E-06 | Stroma 5 | ehbp1l1a |
| kirrel1a5 | 2.9E-10 | 0.7445208 | 0.469 | 0.326 | 9.53E-06 | Stroma 5 | kirrel1a |
| adgre105 | 3.7E-10 | 1.343123 | 0.162 | 0.076 | 1.19E-05 | Stroma 5 | adgre10 |
| si:ch211-212k18.97 | 4.3E-10 | 0.8613386 | 0.302 | 0.177 | 1.41E-05 | Stroma 5 | si:ch211-212k18.9 |
| si:cabz01069012.26 | 4.9E-10 | 1.1708968 | 0.201 | 0.103 | 1.58E-05 | Stroma 5 | si:cabz01069012.2 |
| ugt1b5 | 4.9E-10 | 1.6683409 | 0.109 | 0.043 | 1.61E-05 | Stroma 5 | ugt1b5 |
| mamdc2b3 | 5.1E-10 | 0.8612617 | 0.383 | 0.242 | 1.66E-05 | Stroma 5 | mamdc2b |
| pnrc26 | 5.3E-10 | 0.4086141 | 0.933 | 0.881 | 1.73E-05 | Stroma 5 | pnrc2 |
| rpl65 | 5.5E-10 | 0.3309732 | 0.975 | 0.959 | 1.79E-05 | Stroma 5 | rpl6 |
| creg26 | 6.4E-10 | 0.7981557 | 0.458 | 0.325 | 2.08E-05 | Stroma 5 | creg2 |
| tnrc187 | 6.4E-10 | 0.5820229 | 0.545 | 0.407 | 2.09E-05 | Stroma 5 | tnrc18 |
| fbln74 | 6.5E-10 | 0.6496991 | 0.201 | 0.1 | 2.11E-05 | Stroma 5 | fbln7 |
| limch1b7 | 1E-09 | 1.1513027 | 0.182 | 0.091 | 3.35E-05 | Stroma 5 | limch1b |
| zgc:1541641 | 1E-09 | 1.3057513 | 0.137 | 0.06 | 3.4E-05 | Stroma 5 | zgc:154164 |
| slc29a1a6 | 1.1E-09 | 0.9551796 | 0.318 | 0.198 | 3.65E-05 | Stroma 5 | slc29a1a |
| sorcs2 | 1.1E-09 | 1.3447191 | 0.103 | 0.04 | 3.69E-05 | Stroma 5 | sorcs2 |
| cox6a23 | 1.2E-09 | 1.3147065 | 0.117 | 0.048 | 3.99E-05 | Stroma 5 | cox6a2 |
| thbs1a5 | 1.2E-09 | 0.7371916 | 0.433 | 0.313 | 4.03E-05 | Stroma 5 | thbs1a |
| tgfbr34 | 1.3E-09 | 0.9929118 | 0.168 | 0.08 | 4.26E-05 | Stroma 5 | tgfbr3 |
| acsbg21 | 1.4E-09 | 1.3931269 | 0.115 | 0.047 | 4.66E-05 | Stroma 5 | acsbg2 |
| mlip4 | 1.5E-09 | 1.5697391 | 0.154 | 0.073 | 4.82E-05 | Stroma 5 | mlip |
| lrp1ab7 | 1.5E-09 | 0.7513127 | 0.466 | 0.341 | 4.97E-05 | Stroma 5 | lrp1ab |
| prelp8 | 1.6E-09 | 0.8141258 | 0.243 | 0.133 | 5.11E-05 | Stroma 5 | prelp |
| nt5c3a5 | 1.7E-09 | 1.0769611 | 0.279 | 0.17 | 5.61E-05 | Stroma 5 | nt5c3a |
| amotl2b4 | 1.8E-09 | 1.3795532 | 0.165 | 0.08 | 5.9E-05 | Stroma 5 | amotl2b |
| midn8 | 2E-09 | 1.0844583 | 0.525 | 0.434 | 6.58E-05 | Stroma 5 | midn |
| C9H3orf70 | 2.3E-09 | 1.1633518 | 0.128 | 0.056 | 7.45E-05 | Stroma 5 | C9H3orf70 |
| jund6 | 2.7E-09 | 0.6899521 | 0.679 | 0.57 | 8.77E-05 | Stroma 5 | jund |
| adh55 | 2.8E-09 | 0.5038495 | 0.642 | 0.517 | 8.97E-05 | Stroma 5 | adh5 |
| cyp3c16 | 3.1E-09 | 1.0720407 | 0.187 | 0.095 | 9.94E-05 | Stroma 5 | cyp3c1 |
| ablim1b8 | 3.4E-09 | 0.6125965 | 0.391 | 0.256 | 0.000111 | Stroma 5 | ablim1b |

| | | | | | | | |
|---|---|---|---|---|---|---|---|
| scara57 | 3.9E-09 | 0.821805 | 0.265 | 0.152 | 0.000127 | Stroma 5 | scara5 |
| fstb9 | 4.2E-09 | 0.4144425 | 0.52 | 0.361 | 0.000138 | Stroma 5 | fstb |
| hsp70l6 | 4.4E-09 | 0.6039441 | 0.43 | 0.302 | 0.000142 | Stroma 5 | hsp70l |
| igflr13 | 5.4E-09 | 1.6679991 | 0.103 | 0.042 | 0.000175 | Stroma 5 | igflr1 |
| pbx1a4 | 8.1E-09 | 0.7777341 | 0.352 | 0.232 | 0.000263 | Stroma 5 | pbx1a |
| htra1b6 | 8.6E-09 | 0.4086465 | 0.573 | 0.411 | 0.00028 | Stroma 5 | htra1b |
| aldh1a34 | 8.7E-09 | 0.8371695 | 0.207 | 0.111 | 0.000282 | Stroma 5 | aldh1a3 |
| zfp36l1b9 | 9.2E-09 | 0.6712536 | 0.662 | 0.546 | 0.000299 | Stroma 5 | zfp36l1b |
| il343 | 1.1E-08 | 1.1687673 | 0.12 | 0.052 | 0.000363 | Stroma 5 | il34 |
| eif15 | 1.3E-08 | 0.3877637 | 0.888 | 0.83 | 0.000434 | Stroma 5 | eif1 |
| dpp76 | 1.4E-08 | 0.8428689 | 0.327 | 0.215 | 0.000465 | Stroma 5 | dpp7 |
| furinb2 | 1.6E-08 | 1.3854061 | 0.123 | 0.055 | 0.00053 | Stroma 5 | furinb |
| rps27.23 | 2.2E-08 | 0.2355798 | 0.958 | 0.953 | 0.000719 | Stroma 5 | rps27.2 |
| bco2a4 | 2.2E-08 | 0.6619184 | 0.257 | 0.149 | 0.000724 | Stroma 5 | bco2a |
| nr1d2a5 | 2.5E-08 | 0.7375997 | 0.372 | 0.251 | 0.000819 | Stroma 5 | nr1d2a |
| ZFP36L1.17 | 2.6E-08 | 0.4378906 | 0.941 | 0.873 | 0.000856 | Stroma 5 | ZFP36L1.1 |
| atf37 | 2.7E-08 | 0.7601815 | 0.464 | 0.346 | 0.000869 | Stroma 5 | atf3 |
| spry47 | 2.8E-08 | 1.1694886 | 0.279 | 0.179 | 0.000907 | Stroma 5 | spry4 |
| stard92 | 2.8E-08 | 1.5446236 | 0.115 | 0.05 | 0.000913 | Stroma 5 | stard9 |
| socs1a7 | 2.8E-08 | 1.2488844 | 0.215 | 0.124 | 0.000914 | Stroma 5 | socs1a |
| hsp70.36 | 3E-08 | 0.5440066 | 0.648 | 0.554 | 0.000975 | Stroma 5 | hsp70.3 |
| FP236812.15 | 3.1E-08 | 0.8826293 | 0.187 | 0.099 | 0.000999 | Stroma 5 | FP236812.1 |
| adam19a5 | 3.1E-08 | 1.0101572 | 0.165 | 0.084 | 0.001018 | Stroma 5 | adam19a |
| icn4 | 3.2E-08 | 0.2011043 | 0.997 | 0.996 | 0.001052 | Stroma 5 | icn |
| si:rp71-79p20.24 | 3.9E-08 | 0.8656798 | 0.254 | 0.152 | 0.001272 | Stroma 5 | si:rp71-79p20.2 |
| bmp43 | 4.1E-08 | 1.2316878 | 0.137 | 0.066 | 0.001348 | Stroma 5 | bmp4 |
| adamtsl36 | 4.8E-08 | 1.1062377 | 0.142 | 0.069 | 0.001569 | Stroma 5 | adamtsl3 |
| znf2191 | 5E-08 | 1.3623887 | 0.12 | 0.055 | 0.001631 | Stroma 5 | znf219 |
| hspb16 | 5.1E-08 | 0.7420539 | 0.288 | 0.18 | 0.00166 | Stroma 5 | hspb1 |
| lamc17 | 5.4E-08 | 1.0001695 | 0.223 | 0.129 | 0.001765 | Stroma 5 | lamc1 |
| cd74a | 6.3E-08 | 2.6006651 | 0.101 | 0.043 | 0.002051 | Stroma 5 | cd74a |
| rdh12l | 6.7E-08 | 1.3632912 | 0.103 | 0.044 | 0.002187 | Stroma 5 | rdh12l |
| rasl11b7 | 7.4E-08 | 0.6912022 | 0.327 | 0.217 | 0.002393 | Stroma 5 | rasl11b |
| cmbl6 | 7.4E-08 | 0.8151458 | 0.279 | 0.175 | 0.002419 | Stroma 5 | cmbl |
| ppp1r107 | 8.1E-08 | 0.3649168 | 0.612 | 0.484 | 0.002625 | Stroma 5 | ppp1r10 |
| sept9b5 | 8.4E-08 | 1.077873 | 0.131 | 0.062 | 0.00272 | Stroma 5 | sept9b |
| foxo3b6 | 8.5E-08 | 1.4411556 | 0.156 | 0.082 | 0.002756 | Stroma 5 | foxo3b |
| foxc1b5 | 9E-08 | 1.2047148 | 0.131 | 0.063 | 0.002913 | Stroma 5 | foxc1b |
| mhc1zaa6 | 9.5E-08 | 0.8998382 | 0.176 | 0.093 | 0.003094 | Stroma 5 | mhc1zaa |
| lrp56 | 1.1E-07 | 0.8295388 | 0.201 | 0.112 | 0.003448 | Stroma 5 | lrp5 |
| map2k66 | 1.1E-07 | 0.7037987 | 0.296 | 0.19 | 0.003668 | Stroma 5 | map2k6 |
| ENSDARG0000007621 | 1.2E-07 | 0.4388011 | 0.707 | 0.624 | 0.003924 | Stroma 5 | ENSDARG00000076 |
| fbln55 | 1.2E-07 | 0.8883956 | 0.162 | 0.084 | 0.004055 | Stroma 5 | fbln5 |
| ccnl1a5 | 1.4E-07 | 0.6239179 | 0.38 | 0.262 | 0.004459 | Stroma 5 | ccnl1a |
| ch25hl38 | 1.4E-07 | 0.9861873 | 0.19 | 0.107 | 0.004646 | Stroma 5 | ch25hl3 |
| si:dkey-23i12.75 | 1.6E-07 | 1.1796085 | 0.232 | 0.141 | 0.005053 | Stroma 5 | si:dkey-23i12.7 |
| plxdc26 | 1.6E-07 | 0.9599266 | 0.251 | 0.158 | 0.005054 | Stroma 5 | plxdc2 |
| gadd45ba6 | 1.6E-07 | 0.5771766 | 0.86 | 0.804 | 0.005071 | Stroma 5 | gadd45ba |

| | | | | | | | |
|---|---|---|---|---|---|---|---|
| c1qtnf26 | 1.9E-07 | 0.9141849 | 0.173 | 0.092 | 0.006115 | Stroma 5 | c1qtnf2 |
| wls6 | 2.4E-07 | 0.5493032 | 0.511 | 0.395 | 0.007947 | Stroma 5 | wls |
| tnfaip2a4 | 2.5E-07 | 1.05752 | 0.165 | 0.089 | 0.008022 | Stroma 5 | tnfaip2a |
| si:ch211-63p21.14 | 2.7E-07 | 0.8354863 | 0.112 | 0.05 | 0.008648 | Stroma 5 | si:ch211-63p21.1 |
| ryr36 | 2.8E-07 | 0.8672697 | 0.168 | 0.09 | 0.009034 | Stroma 5 | ryr3 |
| celf29 | 3E-07 | 0.575771 | 0.31 | 0.198 | 0.009817 | Stroma 5 | celf2 |
| ms4a17a.98 | 3.5E-07 | 0.5380966 | 0.33 | 0.216 | 0.011321 | Stroma 5 | ms4a17a.9 |
| tsku7 | 3.7E-07 | 0.5604981 | 0.43 | 0.319 | 0.012118 | Stroma 5 | tsku |
| tmem119b9 | 4.4E-07 | 0.9321352 | 0.307 | 0.206 | 0.014459 | Stroma 5 | tmem119b |
| sulf2b8 | 4.6E-07 | 0.7597242 | 0.26 | 0.165 | 0.014953 | Stroma 5 | sulf2b |
| brd2a6 | 4.9E-07 | 0.5514149 | 0.545 | 0.447 | 0.015959 | Stroma 5 | brd2a |
| ism14 | 5.2E-07 | 0.9394581 | 0.209 | 0.125 | 0.016786 | Stroma 5 | ism1 |
| emp24 | 5.2E-07 | 0.8166534 | 0.243 | 0.153 | 0.016904 | Stroma 5 | emp2 |
| tcf7l1a7 | 6.4E-07 | 0.9617792 | 0.198 | 0.117 | 0.020681 | Stroma 5 | tcf7l1a |
| zgc:565859 | 6.9E-07 | 0.5087414 | 0.447 | 0.328 | 0.02256 | Stroma 5 | zgc:56585 |
| selenbp16 | 8E-07 | 0.9800875 | 0.148 | 0.078 | 0.026072 | Stroma 5 | selenbp1 |
| znf1134 | 8.4E-07 | 1.1770214 | 0.106 | 0.05 | 0.027297 | Stroma 5 | znf1134 |
| ace4 | 8.6E-07 | 0.8104236 | 0.109 | 0.051 | 0.028062 | Stroma 5 | ace |
| si:ch73-335l21.46 | 9.7E-07 | 0.9679215 | 0.615 | 0.571 | 0.031593 | Stroma 5 | si:ch73-335l21.4 |
| C25H11orf969 | 1E-06 | 0.746364 | 0.478 | 0.384 | 0.032412 | Stroma 5 | C25H11orf96 |
| il13ra17 | 1.1E-06 | 1.2294694 | 0.154 | 0.085 | 0.035711 | Stroma 5 | il13ra1 |
| sik2b5 | 1.2E-06 | 0.8610902 | 0.302 | 0.207 | 0.037563 | Stroma 5 | sik2b |
| nfixb8 | 1.3E-06 | 0.5033962 | 0.402 | 0.294 | 0.041966 | Stroma 5 | nfixb |
| SBNO26 | 1.4E-06 | 0.7278886 | 0.324 | 0.228 | 0.045838 | Stroma 5 | SBNO2 |
| rbm4.13 | 1.4E-06 | 0.9557999 | 0.176 | 0.101 | 0.046459 | Stroma 5 | rbm4.1 |
| h2ax17 | 1.5E-06 | 0.4226042 | 0.696 | 0.629 | 0.047203 | Stroma 5 | h2ax1 |
| hist1h4l.186 | 1.6E-06 | 0.7192862 | 0.207 | 0.121 | 0.050856 | Stroma 5 | hist1h4l.18 |
| ebf3a8 | 1.6E-06 | 0.6891835 | 0.318 | 0.216 | 0.053022 | Stroma 5 | ebf3a |
| hsp90aa1.25 | 1.8E-06 | 0.5074262 | 0.679 | 0.6 | 0.057384 | Stroma 5 | hsp90aa1.2 |
| zbtb45 | 1.8E-06 | 0.4194622 | 0.444 | 0.325 | 0.058789 | Stroma 5 | zbtb4 |
| si:dkey-7j14.65 | 1.9E-06 | 0.3024149 | 0.83 | 0.777 | 0.060463 | Stroma 5 | si:dkey-7j14.6 |
| fzd7a3 | 2.1E-06 | 1.2263079 | 0.145 | 0.08 | 0.067199 | Stroma 5 | fzd7a |
| kctd14 | 2.1E-06 | 1.1616586 | 0.145 | 0.079 | 0.068019 | Stroma 5 | kctd1 |
| rgma | 2.1E-06 | 1.2585105 | 0.106 | 0.051 | 0.068085 | Stroma 5 | rgma |
| dap1b.16 | 2.1E-06 | 0.3474995 | 0.628 | 0.507 | 0.069259 | Stroma 5 | dap1b.1 |
| ppp1r3cb2 | 2.3E-06 | 1.1341555 | 0.103 | 0.049 | 0.073408 | Stroma 5 | ppp1r3cb |
| nhsl25 | 2.4E-06 | 0.9507708 | 0.176 | 0.102 | 0.078417 | Stroma 5 | nhsl2 |
| si:ch211-160j14.36 | 2.6E-06 | 0.3437456 | 0.447 | 0.314 | 0.085975 | Stroma 5 | si:ch211-160j14.3 |
| ubc7 | 2.7E-06 | 0.5351004 | 0.735 | 0.663 | 0.088901 | Stroma 5 | ubc |
| etnk16 | 2.8E-06 | 0.680266 | 0.285 | 0.189 | 0.091383 | Stroma 5 | etnk1 |
| ms4a17a.114 | 3E-06 | 0.6182137 | 0.355 | 0.252 | 0.098269 | Stroma 5 | ms4a17a.11 |
| zcchc248 | 3.1E-06 | 0.5353952 | 0.363 | 0.256 | 0.100805 | Stroma 5 | zcchc24 |
| ttyh3b2 | 3.3E-06 | 1.1558286 | 0.109 | 0.054 | 0.106796 | Stroma 5 | ttyh3b |
| mvp7 | 3.5E-06 | 0.6894464 | 0.411 | 0.317 | 0.113872 | Stroma 5 | mvp |
| hsp70.15 | 3.6E-06 | 0.7821373 | 0.444 | 0.341 | 0.116467 | Stroma 5 | hsp70.1 |
| cnbpb4 | 3.6E-06 | 0.4595921 | 0.659 | 0.577 | 0.116898 | Stroma 5 | cnbpb |
| raraa7 | 4E-06 | 0.4639708 | 0.363 | 0.253 | 0.12935 | Stroma 5 | raraa |
| fosl28 | 4.1E-06 | 0.4252847 | 0.531 | 0.429 | 0.132243 | Stroma 5 | fosl2 |

| | | | | | | | |
|---|---|---|---|---|---|---|---|
| slc38a5b3 | 4.4E-06 | 0.6053463 | 0.444 | 0.353 | 0.144055 | Stroma 5 | slc38a5b |
| pmaip14 | 5E-06 | 0.8261046 | 0.165 | 0.095 | 0.161081 | Stroma 5 | pmaip1 |
| ehbp1l1b8 | 5.4E-06 | 0.7524724 | 0.207 | 0.127 | 0.176618 | Stroma 5 | ehbp1l1b |
| tagapa3 | 6.2E-06 | 0.7898787 | 0.14 | 0.076 | 0.200429 | Stroma 5 | tagapa |
| jupa2 | 6.2E-06 | 1.0108391 | 0.204 | 0.129 | 0.202075 | Stroma 5 | jupa |
| pfn13 | 6.4E-06 | 0.4470763 | 0.413 | 0.309 | 0.208301 | Stroma 5 | pfn1 |
| creb3l25 | 7.6E-06 | 0.6094149 | 0.517 | 0.427 | 0.24578 | Stroma 5 | creb3l2 |
| BX322530.13 | 8E-06 | 0.4520812 | 0.517 | 0.42 | 0.259623 | Stroma 5 | BX322530.1 |
| iscub6 | 9E-06 | 0.6716383 | 0.282 | 0.193 | 0.292552 | Stroma 5 | iscub |
| mfap28 | 9.3E-06 | 0.3327629 | 0.506 | 0.384 | 0.30322 | Stroma 5 | mfap2 |
| si:dkey-178e17.32 | 9.7E-06 | 0.9628016 | 0.101 | 0.049 | 0.31491 | Stroma 5 | si:dkey-178e17.3 |
| tiam1a4 | 9.9E-06 | 1.3937265 | 0.117 | 0.062 | 0.322427 | Stroma 5 | tiam1a |
| palm1b2 | 1.1E-05 | 0.8805561 | 0.148 | 0.084 | 0.362714 | Stroma 5 | palm1b |
| ptges5 | 1.3E-05 | 0.5556766 | 0.182 | 0.109 | 0.437179 | Stroma 5 | ptges |
| zgc:1527916 | 1.5E-05 | 0.6554979 | 0.176 | 0.106 | 0.487788 | Stroma 5 | zgc:152791 |
| agmo5 | 1.5E-05 | 0.8466237 | 0.145 | 0.082 | 0.498204 | Stroma 5 | agmo |
| fth1a3 | 1.6E-05 | 0.2631884 | 0.793 | 0.718 | 0.51426 | Stroma 5 | fth1a |
| adam8a3 | 1.9E-05 | 1.1841167 | 0.209 | 0.135 | 0.620149 | Stroma 5 | adam8a |
| phactr4b4 | 1.9E-05 | 0.9288516 | 0.131 | 0.073 | 0.62533 | Stroma 5 | phactr4b |
| parp12b | 2.1E-05 | 0.9481 | 0.128 | 0.071 | 0.670269 | Stroma 5 | parp12b |
| irf1b8 | 2.1E-05 | 1.0452362 | 0.187 | 0.118 | 0.677407 | Stroma 5 | irf1b |
| asap1b5 | 2.3E-05 | 0.8278067 | 0.226 | 0.152 | 0.744672 | Stroma 5 | asap1b |
| gadd45ga7 | 2.4E-05 | 0.4326158 | 0.372 | 0.277 | 0.79102 | Stroma 5 | gadd45ga |
| metrn7 | 2.5E-05 | 0.8826664 | 0.187 | 0.117 | 0.802594 | Stroma 5 | metrn |
| sdc45 | 2.7E-05 | 0.6483634 | 0.737 | 0.705 | 0.867561 | Stroma 5 | sdc4 |
| tlr5b4 | 2.8E-05 | 0.982282 | 0.131 | 0.074 | 0.926564 | Stroma 5 | tlr5b |
| slc20a23 | 2.9E-05 | 0.5811982 | 0.36 | 0.272 | 0.954438 | Stroma 5 | slc20a2 |
| sult6b16 | 3.2E-05 | 0.6281724 | 0.26 | 0.177 | 1 | Stroma 5 | sult6b1 |
| sntb26 | 3.3E-05 | 0.7100185 | 0.318 | 0.237 | 1 | Stroma 5 | sntb2 |
| dop1a5 | 3.4E-05 | 0.9342689 | 0.134 | 0.077 | 1 | Stroma 5 | dop1a |
| dhrs3a5 | 3.5E-05 | 1.0004204 | 0.123 | 0.068 | 1 | Stroma 5 | dhrs3a |
| myclb6 | 3.7E-05 | 0.8399291 | 0.215 | 0.141 | 1 | Stroma 5 | myclb |
| znf5033 | 4.1E-05 | 0.5216406 | 0.184 | 0.113 | 1 | Stroma 5 | znf503 |
| ier5l5 | 4.1E-05 | 0.4449141 | 0.411 | 0.32 | 1 | Stroma 5 | ier5l |
| serpina10a7 | 4.6E-05 | 0.3297741 | 0.483 | 0.378 | 1 | Stroma 5 | serpina10a |
| ak14 | 5.4E-05 | 0.705561 | 0.196 | 0.127 | 1 | Stroma 5 | ak1 |
| s100v24 | 5.5E-05 | 0.2856868 | 0.74 | 0.679 | 1 | Stroma 5 | s100v2 |
| ep300b3 | 5.7E-05 | 0.6764007 | 0.204 | 0.132 | 1 | Stroma 5 | ep300b |
| cpamd84 | 6.1E-05 | 0.620947 | 0.112 | 0.06 | 1 | Stroma 5 | cpamd8 |
| rcan23 | 6.1E-05 | 0.7701068 | 0.109 | 0.059 | 1 | Stroma 5 | rcan2 |
| ywhag19 | 6.3E-05 | 0.6111568 | 0.179 | 0.111 | 1 | Stroma 5 | ywhag1 |
| chka6 | 6.5E-05 | 0.8067922 | 0.151 | 0.091 | 1 | Stroma 5 | chka |
| si:ch211-246m6.53 | 7.5E-05 | 0.800158 | 0.103 | 0.055 | 1 | Stroma 5 | si:ch211-246m6.5 |
| itsn1 | 7.6E-05 | 0.9366053 | 0.106 | 0.057 | 1 | Stroma 5 | itsn1 |
| eef1db7 | 7.7E-05 | 0.3601412 | 0.656 | 0.581 | 1 | Stroma 5 | eef1db |
| si:ch73-46j18.57 | 7.9E-05 | 0.2917401 | 0.682 | 0.603 | 1 | Stroma 5 | si:ch73-46j18.5 |
| tasor22 | 7.9E-05 | 0.6799706 | 0.221 | 0.148 | 1 | Stroma 5 | tasor2 |
| ccni3 | 7.9E-05 | 0.3268802 | 0.662 | 0.594 | 1 | Stroma 5 | ccni |

| | | | | | | | |
|---|---|---|---|---|---|---|---|
| pdlim15 | 8.1E-05 | 0.648672 | 0.307 | 0.23 | 1 | Stroma 5 | pdlim1 |
| c1qtnf16 | 8.2E-05 | 0.7237951 | 0.131 | 0.075 | 1 | Stroma 5 | c1qtnf1 |
| rmdn15 | 9.2E-05 | 0.9106049 | 0.154 | 0.094 | 1 | Stroma 5 | rmdn1 |
| cbx7a6 | 9.4E-05 | 0.4269176 | 0.444 | 0.347 | 1 | Stroma 5 | cbx7a |
| ghra2 | 9.5E-05 | 0.5344377 | 0.31 | 0.228 | 1 | Stroma 5 | ghra |
| serpinb1l1 | 0.0001 | 1.0646365 | 0.112 | 0.063 | 1 | Stroma 5 | serpinb1l1 |
| sesn17 | 0.00011 | 0.4722834 | 0.299 | 0.218 | 1 | Stroma 5 | sesn1 |
| zyg11 | 0.00011 | 1.4886113 | 0.106 | 0.06 | 1 | Stroma 5 | zyg11 |
| ssh2a5 | 0.00012 | 0.9409908 | 0.134 | 0.08 | 1 | Stroma 5 | ssh2a |
| cipcb2 | 0.00012 | 0.8098778 | 0.134 | 0.079 | 1 | Stroma 5 | cipcb |
| pgrmc14 | 0.00013 | 0.4362519 | 0.447 | 0.364 | 1 | Stroma 5 | pgrmc1 |
| si:ch211-105c13.37 | 0.00013 | 0.2421963 | 0.665 | 0.569 | 1 | Stroma 5 | si:ch211-105c13.3 |
| nid1b6 | 0.00013 | 0.8912003 | 0.134 | 0.08 | 1 | Stroma 5 | nid1b |
| si:ch211-196l7.47 | 0.00013 | 0.3261532 | 0.561 | 0.476 | 1 | Stroma 5 | si:ch211-196l7.4 |
| zgc:66474 | 0.00014 | 0.8259568 | 0.142 | 0.087 | 1 | Stroma 5 | zgc:66474 |
| tle2a | 0.00014 | 0.8724867 | 0.109 | 0.061 | 1 | Stroma 5 | tle2a |
| si:dkey-164f24.28 | 0.00014 | 0.2031923 | 0.251 | 0.17 | 1 | Stroma 5 | si:dkey-164f24.2 |
| pcolcea8 | 0.00015 | 0.3692891 | 0.648 | 0.569 | 1 | Stroma 5 | pcolcea |
| psmg3 | 0.00016 | 0.765137 | 0.209 | 0.144 | 1 | Stroma 5 | psmg3 |
| eno32 | 0.00016 | 0.5728983 | 0.377 | 0.301 | 1 | Stroma 5 | eno3 |
| lmo4a4 | 0.00016 | 0.8782323 | 0.19 | 0.126 | 1 | Stroma 5 | lmo4a |
| itm2bb7 | 0.00016 | 0.6753622 | 0.148 | 0.09 | 1 | Stroma 5 | itm2bb |
| sorl12 | 0.00017 | 1.025468 | 0.123 | 0.072 | 1 | Stroma 5 | sorl1 |
| zmp:0000007116 | 0.00018 | 0.8228843 | 0.137 | 0.083 | 1 | Stroma 5 | zmp:0000000711 |
| si:ch211-15b10.63 | 0.00019 | 0.7249756 | 0.159 | 0.101 | 1 | Stroma 5 | si:ch211-15b10.6 |
| tcf128 | 0.0002 | 0.2656425 | 0.606 | 0.523 | 1 | Stroma 5 | tcf12 |
| tent5bb6 | 0.0002 | 0.7366993 | 0.209 | 0.143 | 1 | Stroma 5 | tent5bb |
| serpinh26 | 0.00021 | 0.8744591 | 0.179 | 0.118 | 1 | Stroma 5 | serpinh2 |
| UBB5 | 0.00022 | 0.4053352 | 0.634 | 0.563 | 1 | Stroma 5 | UBB |
| srgn6 | 0.00024 | 0.2215408 | 0.547 | 0.452 | 1 | Stroma 5 | srgn |
| si:dkey-51e6.16 | 0.00024 | 0.3796246 | 0.344 | 0.258 | 1 | Stroma 5 | si:dkey-51e6.1 |
| pdcd4b3 | 0.00025 | 0.7731005 | 0.168 | 0.108 | 1 | Stroma 5 | pdcd4b |
| proza4 | 0.00025 | 0.7184954 | 0.237 | 0.17 | 1 | Stroma 5 | proza |
| f3b9 | 0.00025 | 0.313909 | 0.472 | 0.38 | 1 | Stroma 5 | f3b |
| neu3.28 | 0.00025 | 0.577366 | 0.299 | 0.222 | 1 | Stroma 5 | neu3.2 |
| pttg1ipb3 | 0.00025 | 1.0457548 | 0.115 | 0.068 | 1 | Stroma 5 | pttg1ipb |
| si:ch211-281l24.34 | 0.00028 | 0.5496851 | 0.293 | 0.222 | 1 | Stroma 5 | si:ch211-281l24.3 |
| slc51a6 | 0.00028 | 0.2386705 | 0.235 | 0.159 | 1 | Stroma 5 | slc51a |
| prickle1b7 | 0.00028 | 0.6085199 | 0.251 | 0.178 | 1 | Stroma 5 | prickle1b |
| uxs11 | 0.00034 | 0.8200657 | 0.117 | 0.07 | 1 | Stroma 5 | uxs1 |
| nr1d2b6 | 0.00036 | 0.7595315 | 0.215 | 0.152 | 1 | Stroma 5 | nr1d2b |
| si:ch73-237c6.18 | 0.0004 | 0.2768634 | 0.464 | 0.366 | 1 | Stroma 5 | si:ch73-237c6.1 |
| emx27 | 0.00041 | 0.5195918 | 0.148 | 0.092 | 1 | Stroma 5 | emx2 |
| rhogb5 | 0.00044 | 0.9113064 | 0.117 | 0.07 | 1 | Stroma 5 | rhogb |
| sfxn5b7 | 0.00046 | 0.5962858 | 0.198 | 0.138 | 1 | Stroma 5 | sfxn5b |
| anxa45 | 0.00046 | 0.2550188 | 0.735 | 0.705 | 1 | Stroma 5 | anxa4 |
| waca3 | 0.00046 | 0.6813074 | 0.179 | 0.12 | 1 | Stroma 5 | waca |
| setbp15 | 0.00047 | 0.3492802 | 0.33 | 0.249 | 1 | Stroma 5 | setbp1 |

| | | | | | | | |
|---|---|---|---|---|---|---|---|
| egfra5 | 0.00048 | 0.5354682 | 0.332 | 0.259 | 1 | Stroma 5 | egfra |
| phldb1b5 | 0.00048 | 0.8060963 | 0.14 | 0.089 | 1 | Stroma 5 | phldb1b |
| tmem176l.17 | 0.00049 | 0.2034576 | 0.793 | 0.742 | 1 | Stroma 5 | tmem176l.1 |
| fbxl73 | 0.00049 | 0.7247993 | 0.126 | 0.077 | 1 | Stroma 5 | fbxl7 |
| apof4 | 0.00049 | 0.5037198 | 0.115 | 0.067 | 1 | Stroma 5 | apof |
| fbxo347 | 0.0005 | 0.9366546 | 0.142 | 0.09 | 1 | Stroma 5 | fbxo34 |
| mbd1a | 0.0005 | 0.825756 | 0.101 | 0.058 | 1 | Stroma 5 | mbd1a |
| zfyve212 | 0.00052 | 0.5184146 | 0.293 | 0.224 | 1 | Stroma 5 | zfyve21 |
| cpn17 | 0.00052 | 0.3758442 | 0.291 | 0.212 | 1 | Stroma 5 | cpn1 |
| mrpl233 | 0.00052 | 1.0979785 | 0.126 | 0.079 | 1 | Stroma 5 | mrpl23 |
| mgst1.15 | 0.00055 | 0.2976711 | 0.335 | 0.251 | 1 | Stroma 5 | mgst1.1 |
| ppp1r15a4 | 0.00056 | 0.6272011 | 0.279 | 0.21 | 1 | Stroma 5 | ppp1r15a |
| cd99l24 | 0.00056 | 0.8035751 | 0.271 | 0.208 | 1 | Stroma 5 | cd99l2 |
| rexo14 | 0.00057 | 0.7539167 | 0.176 | 0.119 | 1 | Stroma 5 | rexo1 |
| poldip2 | 0.00057 | 0.9965896 | 0.106 | 0.063 | 1 | Stroma 5 | poldip2 |
| clk4a5 | 0.0006 | 0.4957345 | 0.299 | 0.23 | 1 | Stroma 5 | clk4a |
| rcan1a9 | 0.00063 | 0.3291419 | 0.304 | 0.23 | 1 | Stroma 5 | rcan1a |
| peli1b6 | 0.00063 | 0.7327746 | 0.142 | 0.091 | 1 | Stroma 5 | peli1b |
| rbms2b4 | 0.00064 | 0.847265 | 0.106 | 0.063 | 1 | Stroma 5 | rbms2b |
| sgk16 | 0.00064 | 0.2700414 | 0.45 | 0.367 | 1 | Stroma 5 | sgk1 |
| mapk14b4 | 0.00067 | 0.7961585 | 0.137 | 0.087 | 1 | Stroma 5 | mapk14b |
| aff27 | 0.00068 | 0.485775 | 0.246 | 0.179 | 1 | Stroma 5 | aff2 |
| bahcc1b2 | 0.00071 | 0.6382906 | 0.179 | 0.121 | 1 | Stroma 5 | bahcc1b |
| mgst3b3 | 0.00072 | 0.3797567 | 0.458 | 0.385 | 1 | Stroma 5 | mgst3b |
| si:ch1073-185p12.24 | 0.00072 | 0.7042855 | 0.134 | 0.083 | 1 | Stroma 5 | si:ch1073-185p12.2 |
| wipf1b3 | 0.00074 | 0.7757676 | 0.159 | 0.105 | 1 | Stroma 5 | wipf1b |
| actb15 | 0.00078 | 0.2974499 | 0.902 | 0.867 | 1 | Stroma 5 | actb1 |
| emilin2b4 | 0.00079 | 0.6372103 | 0.115 | 0.069 | 1 | Stroma 5 | emilin2b |
| si:dkey-79d12.55 | 0.0008 | 0.6602991 | 0.187 | 0.13 | 1 | Stroma 5 | si:dkey-79d12.5 |
| sh3bp5a3 | 0.00082 | 0.5926159 | 0.19 | 0.131 | 1 | Stroma 5 | sh3bp5a |
| trib35 | 0.00082 | 0.847725 | 0.237 | 0.176 | 1 | Stroma 5 | trib3 |
| gli2a3 | 0.00087 | 0.9386575 | 0.115 | 0.071 | 1 | Stroma 5 | gli2a |
| arrdc3a6 | 0.00089 | 0.6698271 | 0.226 | 0.165 | 1 | Stroma 5 | arrdc3a |
| abtb2b5 | 0.00094 | 0.8490118 | 0.12 | 0.075 | 1 | Stroma 5 | abtb2b |
| smarca13 | 0.00095 | 0.8731561 | 0.128 | 0.082 | 1 | Stroma 5 | smarca1 |
| akap8l2 | 0.00096 | 0.6783977 | 0.257 | 0.196 | 1 | Stroma 5 | akap8l |
| acvr2aa | 0.00096 | 0.8456737 | 0.103 | 0.062 | 1 | Stroma 5 | acvr2aa |
| prkaa1 | 0.00099 | 0.640678 | 0.128 | 0.08 | 1 | Stroma 5 | prkaa1 |
| ralab3 | 0.00099 | 1.0207744 | 0.137 | 0.09 | 1 | Stroma 5 | ralab |
| acox14 | 0.00099 | 0.5408989 | 0.156 | 0.103 | 1 | Stroma 5 | acox1 |
| dock9b7 | 0.001 | 0.6509604 | 0.151 | 0.1 | 1 | Stroma 5 | dock9b |
| stom6 | 0.00111 | 0.4803307 | 0.341 | 0.27 | 1 | Stroma 5 | stom |
| spaca4l3 | 0.00111 | 0.3991853 | 0.131 | 0.083 | 1 | Stroma 5 | spaca4l |
| boc8 | 0.00113 | 0.4076981 | 0.441 | 0.373 | 1 | Stroma 5 | boc |
| scube34 | 0.00119 | 0.5720814 | 0.148 | 0.096 | 1 | Stroma 5 | scube3 |
| fam102ab1 | 0.00122 | 0.848818 | 0.109 | 0.067 | 1 | Stroma 5 | fam102ab |
| appb4 | 0.00122 | 0.2577378 | 0.545 | 0.469 | 1 | Stroma 5 | appb |
| si:dkey-188i13.105 | 0.00131 | 0.5211908 | 0.318 | 0.248 | 1 | Stroma 5 | si:dkey-188i13.10 |

| | | | | | | | |
|---|---|---|---|---|---|---|---|
| cmtm65 | 0.00133 | 0.6416258 | 0.196 | 0.141 | 1 | Stroma 5 | cmtm6 |
| mhc1uba3 | 0.0014 | 0.7087613 | 0.277 | 0.215 | 1 | Stroma 5 | mhc1uba |
| ramp28 | 0.00141 | 0.9450114 | 0.196 | 0.142 | 1 | Stroma 5 | ramp2 |
| fmnl2a4 | 0.00142 | 0.555393 | 0.162 | 0.11 | 1 | Stroma 5 | fmnl2a |
| cited4a4 | 0.00147 | 0.9308559 | 0.168 | 0.116 | 1 | Stroma 5 | cited4a |
| zgc:1530184 | 0.00148 | 0.4492113 | 0.299 | 0.233 | 1 | Stroma 5 | zgc:153018 |
| bzw1b5 | 0.00151 | 0.7118053 | 0.539 | 0.492 | 1 | Stroma 5 | bzw1b |
| si:dkey-262k9.22 | 0.00159 | 0.5310416 | 0.154 | 0.103 | 1 | Stroma 5 | si:dkey-262k9.2 |
| elmsan1b4 | 0.00161 | 0.4017182 | 0.419 | 0.357 | 1 | Stroma 5 | elmsan1b |
| dennd4a6 | 0.00174 | 0.5311752 | 0.134 | 0.087 | 1 | Stroma 5 | dennd4a |
| nck1a6 | 0.00177 | 0.8253333 | 0.123 | 0.08 | 1 | Stroma 5 | nck1a |
| gas2l12 | 0.00189 | 0.9441182 | 0.101 | 0.062 | 1 | Stroma 5 | gas2l1 |
| gas62 | 0.0019 | 0.9307136 | 0.106 | 0.067 | 1 | Stroma 5 | gas6 |
| wtap2 | 0.00198 | 0.6885425 | 0.179 | 0.127 | 1 | Stroma 5 | wtap |
| hspa12a1 | 0.002 | 0.6122081 | 0.101 | 0.061 | 1 | Stroma 5 | hspa12a |
| galnt13 | 0.00223 | 0.4274319 | 0.223 | 0.164 | 1 | Stroma 5 | galnt1 |
| si:dkey-81l17.62 | 0.00223 | 0.8003688 | 0.12 | 0.078 | 1 | Stroma 5 | si:dkey-81l17.6 |
| pepd6 | 0.0024 | 0.6869127 | 0.268 | 0.214 | 1 | Stroma 5 | pepd |
| pik3r3a4 | 0.00247 | 0.6160617 | 0.187 | 0.135 | 1 | Stroma 5 | pik3r3a |
| snai1b6 | 0.00256 | 0.6039208 | 0.162 | 0.113 | 1 | Stroma 5 | snai1b |
| nid2a3 | 0.00261 | 0.7118137 | 0.115 | 0.074 | 1 | Stroma 5 | nid2a |
| spred12 | 0.0027 | 0.5065776 | 0.182 | 0.128 | 1 | Stroma 5 | spred1 |
| anos1b3 | 0.00276 | 0.8229705 | 0.117 | 0.076 | 1 | Stroma 5 | anos1b |
| eif4ebp3l4 | 0.00277 | 0.707947 | 0.207 | 0.155 | 1 | Stroma 5 | eif4ebp3l |
| plekhj14 | 0.00279 | 0.5002397 | 0.173 | 0.122 | 1 | Stroma 5 | plekhj1 |
| ctdsp24 | 0.00282 | 0.5622698 | 0.246 | 0.191 | 1 | Stroma 5 | ctdsp2 |
| ubap2a3 | 0.00284 | 0.5194551 | 0.251 | 0.194 | 1 | Stroma 5 | ubap2a |
| pik3r3b5 | 0.00286 | 0.3952209 | 0.338 | 0.274 | 1 | Stroma 5 | pik3r3b |
| nfil3-54 | 0.00286 | 0.6994901 | 0.19 | 0.139 | 1 | Stroma 5 | nfil3-5 |
| nr2f1a5 | 0.00296 | 0.6127556 | 0.207 | 0.153 | 1 | Stroma 5 | nr2f1a |
| ssbp42 | 0.00297 | 0.6715705 | 0.117 | 0.076 | 1 | Stroma 5 | ssbp4 |
| bccip3 | 0.003 | 0.4336077 | 0.254 | 0.195 | 1 | Stroma 5 | bccip |
| per1b5 | 0.00301 | 0.5961759 | 0.26 | 0.203 | 1 | Stroma 5 | per1b |
| si:dkey-16l2.204 | 0.00311 | 0.8694528 | 0.101 | 0.063 | 1 | Stroma 5 | si:dkey-16l2.20 |
| slc38a28 | 0.00313 | 1.0005728 | 0.12 | 0.08 | 1 | Stroma 5 | slc38a2 |
| rbm4.25 | 0.00316 | 0.5683769 | 0.221 | 0.166 | 1 | Stroma 5 | rbm4.2 |
| srsf5a5 | 0.00331 | 0.2837906 | 0.628 | 0.58 | 1 | Stroma 5 | srsf5a |
| dcbld27 | 0.00333 | 0.3413063 | 0.179 | 0.125 | 1 | Stroma 5 | dcbld2 |
| ptpn32 | 0.00342 | 0.6828595 | 0.109 | 0.07 | 1 | Stroma 5 | ptpn3 |
| zgc:77486.18 | 0.00349 | 0.4471444 | 0.226 | 0.17 | 1 | Stroma 5 | zgc:77486.1 |
| phyh3 | 0.00359 | 0.7704727 | 0.134 | 0.092 | 1 | Stroma 5 | phyh |
| pdzrn3b5 | 0.00362 | 0.7543204 | 0.12 | 0.079 | 1 | Stroma 5 | pdzrn3b |
| si:dkey-7j14.54 | 0.00379 | 0.8417591 | 0.109 | 0.071 | 1 | Stroma 5 | si:dkey-7j14.5 |
| ece2b4 | 0.00383 | 0.774282 | 0.145 | 0.101 | 1 | Stroma 5 | ece2b |
| litaf2 | 0.00391 | 0.6081262 | 0.165 | 0.118 | 1 | Stroma 5 | litaf |
| dhrs96 | 0.00391 | 0.6465495 | 0.19 | 0.141 | 1 | Stroma 5 | dhrs9 |
| c66 | 0.00423 | 0.6796232 | 0.112 | 0.073 | 1 | Stroma 5 | c6 |
| tmem131l1 | 0.00428 | 0.7498382 | 0.182 | 0.132 | 1 | Stroma 5 | tmem131l |

| | | | | | | | |
|---|---|---|---|---|---|---|---|
| dlx2a2 | 0.00467 | 0.6564707 | 0.103 | 0.066 | 1 | Stroma 5 | dlx2a |
| ddx512 | 0.00494 | 0.3462549 | 0.631 | 0.598 | 1 | Stroma 5 | ddx5 |
| si:dkey-9i23.154 | 0.00499 | 0.4910181 | 0.204 | 0.152 | 1 | Stroma 5 | si:dkey-9i23.15 |
| ing3 | 0.00517 | 0.4049779 | 0.117 | 0.077 | 1 | Stroma 5 | ing3 |
| niban2b3 | 0.00537 | 0.4966351 | 0.201 | 0.15 | 1 | Stroma 5 | niban2b |
| slc25a36a4 | 0.00551 | 0.5840817 | 0.151 | 0.107 | 1 | Stroma 5 | slc25a36a |
| fam43a6 | 0.00591 | 0.5845573 | 0.106 | 0.069 | 1 | Stroma 5 | fam43a |
| hipk3b3 | 0.00593 | 0.5454752 | 0.117 | 0.078 | 1 | Stroma 5 | hipk3b |
| ezh15 | 0.00602 | 0.4914339 | 0.19 | 0.141 | 1 | Stroma 5 | ezh1 |
| st133 | 0.00605 | 0.4402697 | 0.268 | 0.213 | 1 | Stroma 5 | st13 |
| eif4a26 | 0.00616 | 0.5437277 | 0.243 | 0.193 | 1 | Stroma 5 | eif4a2 |
| romo15 | 0.00635 | 0.2292222 | 0.715 | 0.707 | 1 | Stroma 5 | romo1 |
| hic15 | 0.00637 | 0.4900329 | 0.101 | 0.065 | 1 | Stroma 5 | hic1 |
| rxrbb | 0.0066 | 0.6999581 | 0.112 | 0.075 | 1 | Stroma 5 | rxrbb |
| thrb4 | 0.00661 | 0.4145835 | 0.218 | 0.166 | 1 | Stroma 5 | thrb |
| prelid3b7 | 0.00669 | 0.3692953 | 0.413 | 0.358 | 1 | Stroma 5 | prelid3b |
| vamp54 | 0.00671 | 0.5892301 | 0.12 | 0.081 | 1 | Stroma 5 | vamp5 |
| rbm55 | 0.00674 | 0.4221639 | 0.24 | 0.186 | 1 | Stroma 5 | rbm5 |
| cdh27 | 0.00706 | 0.4421081 | 0.352 | 0.293 | 1 | Stroma 5 | cdh2 |
| gstk14 | 0.00717 | 0.5847544 | 0.209 | 0.162 | 1 | Stroma 5 | gstk1 |
| si:dkey-6n21.123 | 0.0078 | 0.6615112 | 0.173 | 0.13 | 1 | Stroma 5 | si:dkey-6n21.12 |
| zgc:1748887 | 0.00822 | 0.301722 | 0.274 | 0.218 | 1 | Stroma 5 | zgc:174888 |
| svila7 | 0.00823 | 0.4036434 | 0.198 | 0.149 | 1 | Stroma 5 | svila |
| adamts15b.17 | 0.00831 | 0.5093024 | 0.134 | 0.093 | 1 | Stroma 5 | adamts15b.1 |
| bag37 | 0.00881 | 1.1286225 | 0.106 | 0.072 | 1 | Stroma 5 | bag3 |
| SERF21 | 0.0089 | 0.4730737 | 0.14 | 0.098 | 1 | Stroma 5 | SERF2 |
| spink2.15 | 0.00891 | 0.4927994 | 0.176 | 0.129 | 1 | Stroma 5 | spink2.1 |
