## Supplemental Table 3 for "Dynamic cell fate plasticity and tissue integration drive functional synovial joint regeneration"

| Archetype | ID | Description | GeneRatio | BgRatio | p.adjust | Top Enriched Genes |
|---|---|---|---|---|---|---|
| 1 | GO:0030198 | extracellular matrix organization | 12/116 | 180/18569 | 4.89E-07 | col5a3b/paplnb/col7a1/ccdc80l2/postnb/si:ch211-196i2.1/adamtsl3/loxl3a/fbln1/lamb1a/ccdc80l1/col18a1a |
| 1 | GO:0043062 | extracellular structure organization | 12/116 | 180/18569 | 4.89E-07 | col5a3b/paplnb/col7a1/ccdc80l2/postnb/si:ch211-196i2.1/adamtsl3/loxl3a/fbln1/lamb1a/ccdc80l1/col18a1a |
| 1 | GO:0045229 | external encapsulating structure organization | 12/116 | 181/18569 | 4.89E-07 | col5a3b/paplnb/col7a1/ccdc80l2/postnb/si:ch211-196i2.1/adamtsl3/loxl3a/fbln1/lamb1a/ccdc80l1/col18a1a |
| 1 | GO:0048679 | regulation of axon regeneration | 4/116 | 12/18569 | 0.000138948 | col12a1a/clcf1/col12a1b/epha4b |
| 1 | GO:0070570 | regulation of neuron projection regeneration | 4/116 | 12/18569 | 0.000138948 | col12a1a/clcf1/col12a1b/epha4b |
| 1 | GO:0061564 | axon development | 14/116 | 488/18569 | 0.000365669 | col6a4a/coro1a/lamc3/wnt4/sema5a/col12a1a/musk/clcf1/col12a1b/epha4b/lamb1a/ccdc80l1/vwa1/sema3aa |
| 1 | GO:0051241 | negative regulation of multicellular organismal process | 9/116 | 204/18569 | 0.000778749 | fgl1/ism1/sema5a/hhla2a.1/loxl3a/musk/epha4b/axin2/sema3aa |
| 1 | GO:1990138 | neuron projection extension | 7/116 | 118/18569 | 0.001161501 | tnn/wnt4/sema5a/musk/nrn1a/epha4b/sema3aa |
| 1 | GO:0048812 | neuron projection morphogenesis | 13/116 | 500/18569 | 0.001650867 | tnn/col6a4a/coro1a/lamc3/wnt4/sema5a/musk/nrn1a/epha4b/lamb1a/ccdc80l1/vwa1/sema3aa |
| 1 | GO:0048588 | developmental cell growth | 7/116 | 135/18569 | 0.001931584 | tnn/wnt4/sema5a/musk/nrn1a/epha4b/sema3aa |
| 2 | GO:0040011 | locomotion | 13/111 | 409/18569 | 0.000934911 | sst6/sema7a/ccl19a.1/ackr3b/sema3d/ccl44/rhogb/sema3h/fgf18a/anxa5b/sulf1/sema3fb/ackr4b |
| 2 | GO:0006935 | chemotaxis | 9/111 | 214/18569 | 0.001709045 | sema7a/ccl19a.1/ackr3b/sema3d/ccl44/rhogb/sema3h/sema3fb/ackr4b |
| 2 | GO:0042330 | taxis | 9/111 | 214/18569 | 0.001709045 | sema7a/ccl19a.1/ackr3b/sema3d/ccl44/rhogb/sema3h/sema3fb/ackr4b |
| 2 | GO:0048843 | negative regulation of axon extension involved in axon g | 4/111 | 30/18569 | 0.006645711 | sema7a/sema3d/sema3h/sema3fb |
| 2 | GO:0050922 | negative regulation of chemotaxis | 4/111 | 35/18569 | 0.006645711 | sema7a/sema3d/sema3h/sema3fb |
| 2 | GO:0030517 | negative regulation of axon extension | 4/111 | 37/18569 | 0.006645711 | sema7a/sema3d/sema3h/sema3fb |
| 2 | GO:0010721 | negative regulation of cell development | 5/111 | 72/18569 | 0.006645711 | sema7a/sema3d/sema3h/qki2/sema3fb |
| 2 | GO:0050919 | negative chemotaxis | 4/111 | 38/18569 | 0.006645711 | sema7a/sema3d/sema3h/sema3fb |
| 2 | GO:0048841 | regulation of axon extension involved in axon guidance | 4/111 | 39/18569 | 0.006645711 | sema7a/sema3d/sema3h/sema3fb |
| 2 | GO:0050771 | negative regulation of axonogenesis | 4/111 | 39/18569 | 0.006645711 | sema7a/sema3d/sema3h/sema3fb |
| 3 | GO:0030198 | extracellular matrix organization | 14/117 | 180/18569 | 2.43E-09 | col10a1b/col10a1a/col5a2b/adamts18/loxl4/col22a1/si:dkey-32e6.6/emid1/col11a2/mibp2/col1a1a/col1a2/reck/col1a1b |
| 3 | GO:0043062 | extracellular structure organization | 14/117 | 180/18569 | 2.43E-09 | col10a1b/col10a1a/col5a2b/adamts18/loxl4/col22a1/si:dkey-32e6.6/emid1/col11a2/mibp2/col1a1a/col1a2/reck/col1a1b |
| 3 | GO:0045229 | external encapsulating structure organization | 14/117 | 181/18569 | 2.43E-09 | col10a1b/col10a1a/col5a2b/adamts18/loxl4/col22a1/si:dkey-32e6.6/emid1/col11a2/mibp2/col1a1a/col1a2/reck/col1a1b |
| 3 | GO:0001503 | ossification | 10/117 | 102/18569 | 1.97E-07 | bglap/panx3/tmem119a/bmp3/phospho1/sp7/entpd5a/spp1/mef2ca/col1a1a |
| 3 | GO:0001649 | osteoblast differentiation | 5/117 | 29/18569 | 0.000173185 | bglap/tmem119a/bmp3/sp7/mef2ca |
| 3 | GO:0001501 | skeletal system development | 13/117 | 407/18569 | 0.000243463 | bglap/bmp3/spns2/sp7/col11a2/spp1/mef2ca/fat3a/mkxa/sparc/col1a1a/col1a2/col1a1b |
| 3 | GO:0031214 | biomineral tissue development | 6/117 | 59/18569 | 0.000243463 | bglap/tmem119a/phospho1/entpd5a/fam20a/col1a1a |
| 3 | GO:0060348 | bone development | 5/117 | 69/18569 | 0.008380669 | bglap/sp7/spp1/mef2ca/col1a1a |
| 3 | GO:0030282 | bone mineralization | 4/117 | 48/18569 | 0.023595395 | tmem119a/phospho1/entpd5a/col1a1a |
| 3 | GO:0060027 | convergent extension involved in gastrulation | 5/117 | 90/18569 | 0.023604806 | ptk2ab/fyna/tnika/swap70b/csk |
| 4 | GO:0009074 | aromatic amino acid family catabolic process | 6/113 | 18/18569 | 3.59E-07 | pah/hgd/hpdb/gstz1/tat/qdpra |
| 4 | GO:0006558 | L-phenylalanine metabolic process | 5/113 | 10/18569 | 3.59E-07 | pah/hgd/gstz1/tat/qdpra |
| 4 | GO:0006559 | L-phenylalanine catabolic process | 5/113 | 10/18569 | 3.59E-07 | pah/hgd/gstz1/tat/qdpra |
| 4 | GO:1902221 | erythrose 4-phosphate/phosphoenolpyruvate family am | 5/113 | 10/18569 | 3.59E-07 | pah/hgd/gstz1/tat/qdpra |
| 4 | GO:1902222 | erythrose 4-phosphate/phosphoenolpyruvate family am | 5/113 | 10/18569 | 3.59E-07 | pah/hgd/gstz1/tat/qdpra |
| 4 | GO:0006570 | tyrosine metabolic process | 5/113 | 11/18569 | 5.45E-07 | pah/hgd/hpdb/gstz1/tat |
| 4 | GO:0009072 | aromatic amino acid metabolic process | 6/113 | 26/18569 | 1.26E-06 | pah/hgd/hpdb/gstz1/tat/qdpra |
| 4 | GO:0170039 | proteinogenic amino acid metabolic process | 8/113 | 123/18569 | 0.000100709 | pah/hgd/bhmt/zgc:153031/gstz1/tat/qdpra/cbsb |
| 4 | GO:1901605 | alpha-amino acid metabolic process | 9/113 | 173/18569 | 0.000118591 | pah/hgd/hpdb/bhmt/zgc:153031/gstz1/tat/qdpra/cbsb |
| 4 | GO:0170033 | L-amino acid metabolic process | 8/113 | 140/18569 | 0.000214434 | pah/hgd/bhmt/zgc:153031/gstz1/tat/qdpra/cbsb |
| 5 | GO:0030198 | extracellular matrix organization | 17/118 | 180/18569 | 4.08E-13 | loxa/postnb/col11a1b/col16a1/mmp2/loxl3a/col1a2/lum/abi3bpb/col8a2/col1a1a/col1a1b/col2a1b/col5a2a/adamtsl3/ndnf/mmp14a |
| 5 | GO:0043062 | extracellular structure organization | 17/118 | 180/18569 | 4.08E-13 | loxa/postnb/col11a1b/col16a1/mmp2/loxl3a/col1a2/lum/abi3bpb/col8a2/col1a1a/col1a1b/col2a1b/col5a2a/adamtsl3/ndnf/mmp14a |
| 5 | GO:0045229 | external encapsulating structure organization | 17/118 | 181/18569 | 4.08E-13 | loxa/postnb/col11a1b/col16a1/mmp2/loxl3a/col1a2/lum/abi3bpb/col8a2/col1a1a/col1a1b/col2a1b/col5a2a/adamtsl3/ndnf/mmp14a |
| 5 | GO:0001501 | skeletal system development | 19/118 | 407/18569 | 1.90E-09 | bcan/pitx2/ucmaa/ogna/nog3/loxa/meox1/sox6/nog2/col1a2/sparc/col1a1a/col1a1b/alcama/col2a1b/ndnf/runx2b/mmp14a/tgfb3 |
| 5 | GO:0061448 | connective tissue development | 12/118 | 205/18569 | 9.91E-07 | ogna/nog3/loxa/scxa/sox6/nog2/thbs4b/sparc/alcama/runx2b/mmp14a/tgfb3 |
| 5 | GO:0001503 | ossification | 9/118 | 102/18569 | 1.97E-06 | ucmaa/nog3/fap/nog2/col1a1a/runx2b/mmp14a/tgfb3/entpd5a |
| 5 | GO:0051216 | cartilage development | 10/118 | 192/18569 | 3.98E-05 | ogna/nog3/loxa/sox6/nog2/sparc/alcama/runx2b/mmp14a/tgfb3 |
| 5 | GO:0030199 | collagen fibril organization | 5/118 | 30/18569 | 0.000101663 | loxa/loxl3a/lum/col1a1a/col2a1b |
| 5 | GO:0051093 | negative regulation of developmental process | 8/118 | 174/18569 | 0.001157612 | tnmd/nog3/thbs2b/loxl3a/fap/nog2/s1pr1/serpinf1 |
| 5 | GO:0045595 | regulation of cell differentiation | 13/118 | 498/18569 | 0.001157612 | bcan/ucmaa/nog3/ccn4b/sox6/lgals2a/loxl3a/ccn5/nog2/six1a/alcama/runx2b/serpinf1 |
| 6 | GO:0030198 | extracellular matrix organization | 18/113 | 180/18569 | 1.10E-14 | mia/col2a1a/si:dkey-6n6.1/col9a2/loxl5a/col9a3/loxl5b/col9a1a/si:dkey-61l1.4/col11a1a/col11a2/col8a1b/col15a1b/col2a1b/adamts3/si:dkey-32e6.6/tgfbi/loxl3b |
| 6 | GO:0043062 | extracellular structure organization | 18/113 | 180/18569 | 1.10E-14 | mia/col2a1a/si:dkey-6n6.1/col9a2/loxl5a/col9a3/loxl5b/col9a1a/si:dkey-61l1.4/col11a1a/col11a2/col8a1b/col15a1b/col2a1b/adamts3/si:dkey-32e6.6/tgfbi/loxl3b |
| 6 | GO:0045229 | external encapsulating structure organization | 18/113 | 181/18569 | 1.10E-14 | mia/col2a1a/si:dkey-6n6.1/col9a2/loxl5a/col9a3/loxl5b/col9a1a/si:dkey-61l1.4/col11a1a/col11a2/col8a1b/col15a1b/col2a1b/adamts3/si:dkey-32e6.6/tgfbi/loxl3b |
| 6 | GO:0001501 | skeletal system development | 19/113 | 407/18569 | 1.03E-09 | matn1/ccn6/cnmd/col2a1a/acanb/snorc/epyc/ucmab/acana/col11a1a/col11a2/hapln1b/sox5/fgfrl1b/col2a1b/sox6/loxl3b/hmgcs1/alcama |
| 6 | GO:0030199 | collagen fibril organization | 6/113 | 30/18569 | 3.43E-06 | col2a1a/loxl5a/loxl5b/col11a2/col2a1b/loxl3b |
| 6 | GO:0051216 | cartilage development | 11/113 | 192/18569 | 3.43E-06 | matn1/ccn6/cnmd/snorc/epyc/col11a1a/col11a2/sox5/fgfrl1b/sox6/alcama |
| 6 | GO:0061448 | connective tissue development | 11/113 | 205/18569 | 5.76E-06 | matn1/ccn6/cnmd/snorc/epyc/col11a1a/col11a2/sox5/fgfrl1b/sox6/alcama |
| 6 | GO:0045597 | positive regulation of cell differentiation | 10/113 | 179/18569 | 1.45E-05 | ccn6/acanb/ccn1l2/si:ch211-106h11.3/acana/hapln1b/sox5/ccn1l1/sox6/alcama |
| 6 | GO:0051094 | positive regulation of developmental process | 10/113 | 269/18569 | 0.000522883 | ccn6/acanb/ccn1l2/si:ch211-106h11.3/acana/hapln1b/sox5/ccn1l1/sox6/alcama |
| 6 | GO:0045595 | regulation of cell differentiation | 13/113 | 498/18569 | 0.000864487 | ccn6/acanb/ccn1l2/ucmab/sox10/si:ch211-106h11.3/acana/hapln1b/sox5/ccn1l1/sox6/loxl3b/alcama |
